# SorCS2 modulates neurovascular coupling via glutamatergic and calcium signaling in astrocytes

**DOI:** 10.1101/2023.02.16.528727

**Authors:** Christian Staehr, Hande Login, Dmitry Postnov, Simin Berenji Ardestani, Stella Solveig Nolte, Hans Christian Beck, Anders Nykjaer, Vladimir V. Matchkov

**Affiliations:** Department of Biomedicine, Aarhus University, Aarhus, DK-8000, Denmark; Center of Functionally Integrative Neuroscience, Department of Clinical Medicine, Aarhus University Hospital Skejby, Aarhus, DK-8200, Denmark; Department for Clinical Biochemistry and Pharmacology, Odense University Hospital, Odense, DK-5000, Denmark; PROMEMO and DANDRITE, Aarhus University, Aarhus, DK-8000, Denmark; Department of Neurosurgery, Aarhus University Hospital Skejby, Aarhus, DK-8200, Denmark

**Keywords:** SorCS2, neurovascular coupling, astrocytes, calcium signalling, glutamate signalling

## Abstract

SorCS2 is involved in trafficking of membrane receptors and transporters. SorCS2 is implicated in brain disorders, but the mechanism remains uncertain. We hypothesized that SorCS2 expression is important for neurovascular coupling.

Brains from P8 and 2-month-old wild type mice were stained for SorCS2 and compared to SorCS2 knockouts (*Sorcs2^-/-^*). Changes in cerebral perfusion in response to sensory stimulation, i.e., neurovascular coupling, were compared *in vivo*. Neurovascular coupling was also assessed *ex vivo* in brain slices loaded with calcium-sensitive dye. Proteomics of astrocytes was analyzed for ingenuity pathways.

SorCS2 was strongly expressed in astrocytic endfeet of P8 mice but only in few astrocytes from 2-month-old brains. *Sorcs2^-/-^* mice demonstrated reduced neurovascular coupling. This was associated with reduced astrocytic calcium response to neuronal excitation in *Sorcs2^-/-^* mice. No difference in cerebral artery caliber nor in endothelial function was seen between wild type and *Sorcs2^-/-^* mice. Proteomics indicated reduced glutamatergic signaling and suppressed calcium signaling in *Sorcs2^-/-^* astrocytes.

We suggest that SorCS2 expression is important for neurovascular coupling due to modulation of glutamatergic and calcium signaling in astrocytes.

## Introduction

Sortilin Related VPS10 Domain Containing Receptor 2 (SorCS2) is a member of the VPS10P domain family of multifunctional receptors. It can participate in signaling from the plasma membrane and mediate trafficking of specific receptors and transporters between intracellular compartments and the plasma membrane to regulate their surface exposure and activitty.^1^ The relevance of SorCS2 to human brain disorders has been proposed,^2–5^ including an association of *SORCS2* with risk of ischemic stroke,^6^ bipolar disorder^7, 8^ and schizophrenia.^9^ However, the molecular mechanisms underlying the function of SorCS2 in these and other brain pathologies are still not fully resolved.

The expression of SorCS2 is controlled in a different temporal manner in different cell types. In particular SorCS2 is abundant at specific stages of development of the nervous system where it is indispensable for proper dopaminergic innervation.^10^ During adulthood, SorCS2 plays an important role in neuronal antioxidant response by contributing to the synthesis of glutathione, an important scavenger of reactive oxygen species (ROS).^5^ Hence, SorCS2 deficiency leads to oxidative brain damage, enhanced neuronal cell death and increased mortality during epilepsy.^5^ Whereas SorCS2 is prominent in neurons, its expression in resident astrocytes of adult mice seems to be low.^4, 5, 10, 11^ However, astrocytes upregulate SorCS2 after ischemic stroke, which has been suggested important for post-stroke angiogenesis.^4^

The survival of neuronal tissue during metabolic challenges depends to a large extent on the efficiency of neurovascular signaling.^12^ Neurovascular coupling is essential to match the metabolic demand of neuronal tissues to delivery of O_2_ and nutrients and for removal of waste products, which is managed by blood perfusion.^13^ When neurons are excited, this is sensed by the nearby astrocytes, which respond with intracellular Ca^2+^ waves spreading to the astrocytic endfeet.^12, 14^ As a consequence, the endfeet release a plethora of vasoactive substances, leading to relaxation of the adjacent arteriole and an increased blood flow in order to meet the metabolic demand of the neuronal tissue.^15^

Given that SorCS2 expression in astrocytes can be modified under metabolic challenges^4^ and that it can regulate the N-methyl-D-aspartate (NMDA) signalling,^2, 5, 16^ we speculated that SorCS2 may be important for regulation of neurovascular coupling. We tested our hypothesis by assessing neurovascular coupling in wild type and SorCS2 knockout (*Sorcs2^-/-^*) mice *in vivo* and *ex vivo* in brain slices. Cerebrovascular function was studied in isolated cerebral arteries. The mechanistic background underlying disturbances in astrocytic function was elucidated using ingenuity pathway analysis of proteomics of astrocytes. The results of this study provide deep insight into the important role of SorCS2 for the neurovascular unit.

## Results

### SorCS2 is detected in astrocytes, but its prevalence decreases with age

Immunohistochemical staining of brain slices from eight-day-old (P8) pups revealed strong staining in SorCS2 in GFAP-positive astrocytes surrounding arterioles of wildtype but not in *Sorcs2^-/-^* mice (Fig. 1A). Here, expression was enriched in a perinuclear compartment compatible with the trans-Golgi network in accordance with observations obtained in other cell types.^10^ However, higher magnification images revealed that SorCS2 was abundant in astrocytic aquaporin 4-positive endfeet that encase arterioles and play a critical role in vasoconstriction and dilation (Fig. 1B). SorCS2 was also expressed in astrocytes of 2-month-old mice (Fig. 2) although to a lesser extent compared to what was observed in P8 pups (Fig. 1). The neuronal SorCS2 expression was increased in adult mice compared to P8 pups (Fig. 1 and 2).

**Figure 1.**
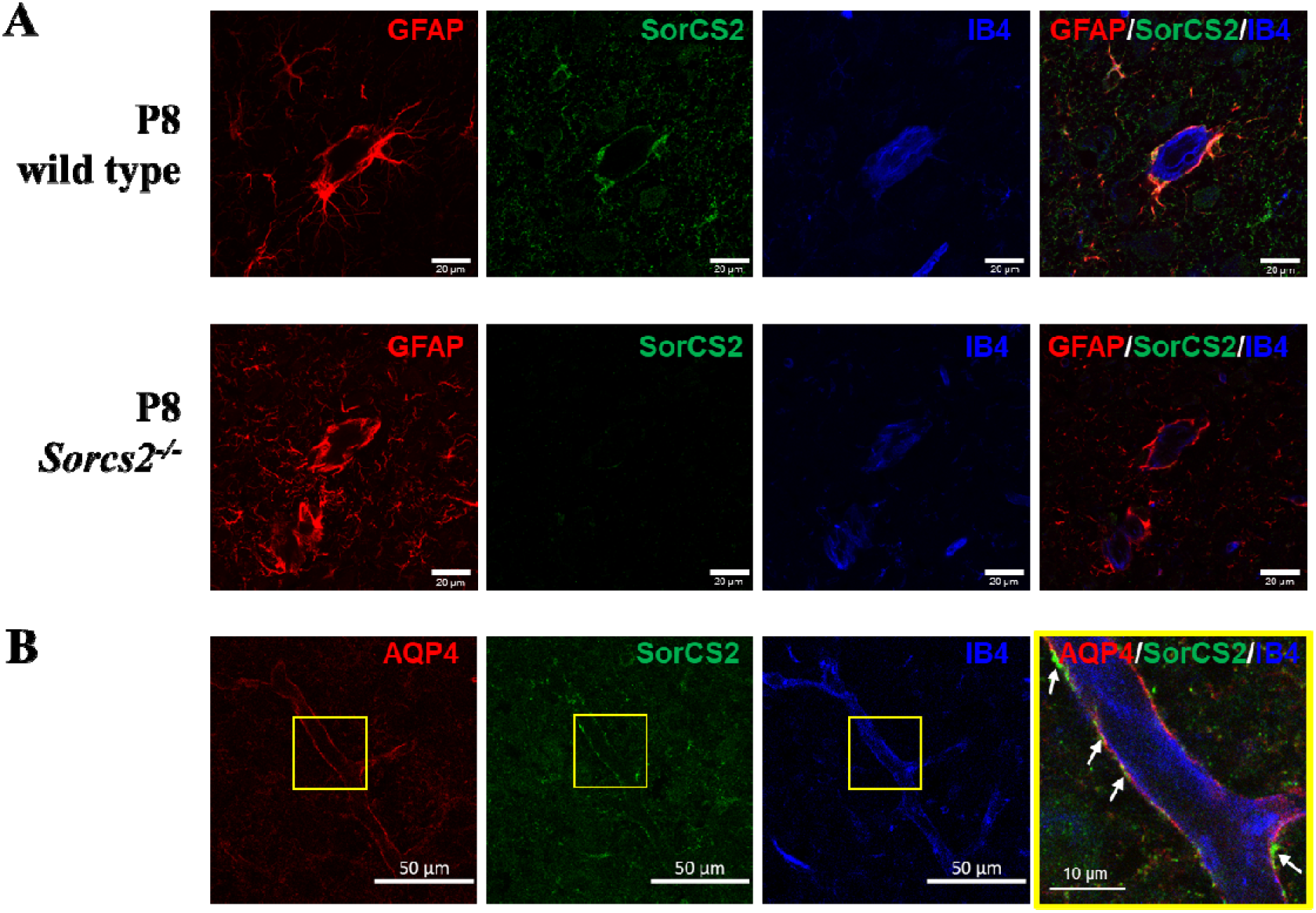
SorCS2 is intensively localizes to astrocytic endfeet in newborn mice. In the 8-days-old wild type mice (P8), SorCS2 staining colocalized with astrocytic marker, glial fibrillary acidic protein (GFAP), and surrounded vascular endothelial cells labeled with α-d-galactosyl-specific isolectin B4 (IB4), as indicated (A). *Sorcs2^-/-^* abolished SorCS2 specific staining in the brain of P8 mice, while GFAP and IB4 staining was not affected (bars correspond to 20 μm). An intensive aquaporin-4 (AQP4) and SorCS2 staining suggests astrocytic endfeet surrounding IB4 positive vascular endothelial cells in the brain of P8 wild type mice (B; bars correspond to 50 μm). Merged image suggests colocalization of AQP4 and SorCS2 in astrocytic endfeet surrounding blood vessel, as indicated by arrows in the magnified image (bar is 10 μm).

**Figure 2.**
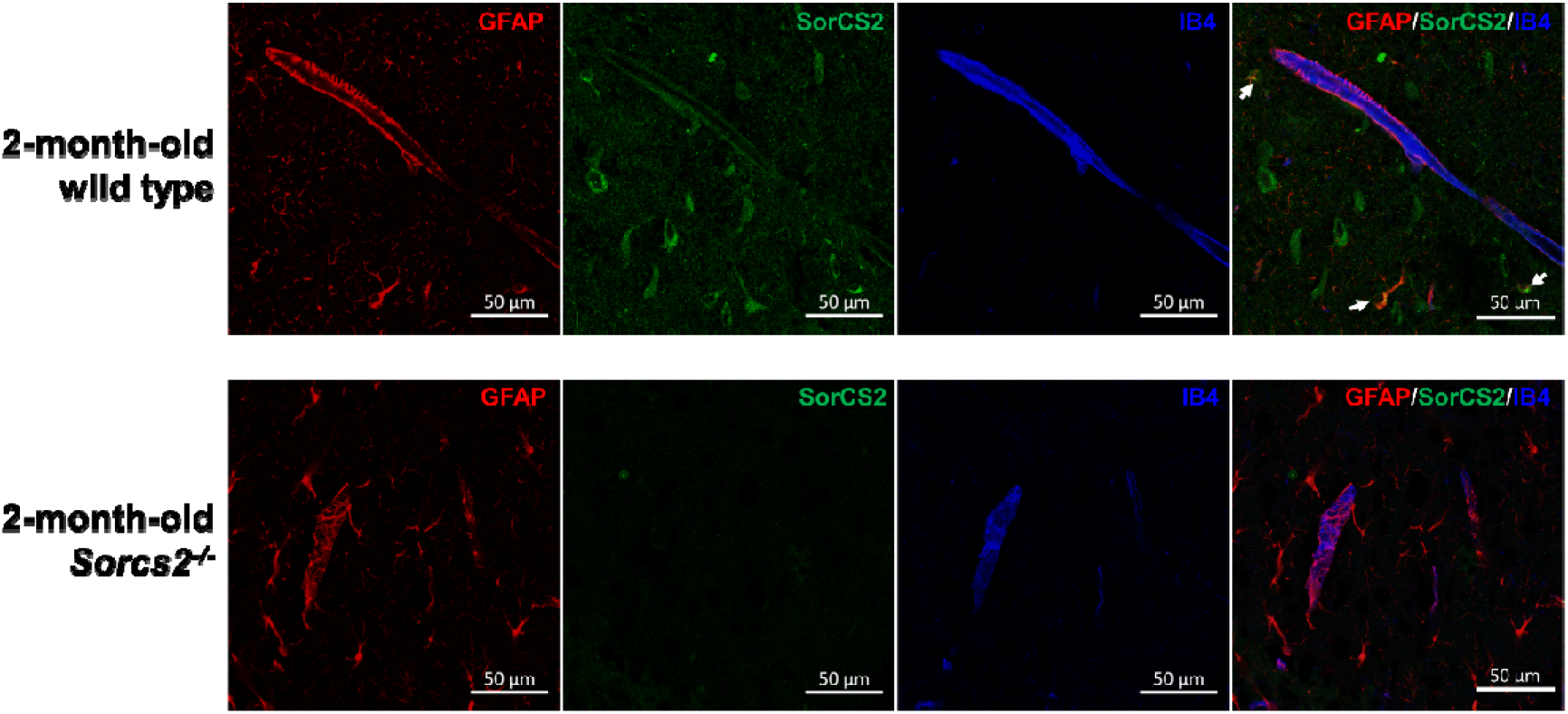
SorCS2 is expressed in some astrocytes of 2-month-old mice. Brain slices were stained with glial fibrillary acidic protein (GFAP) to stain astrocytes, SorCS2 and vascular endothelial cell label α-d-galactosyl-specific isolectin B4 (IB4), as indicated. In 2-month-old wild type mice, some GFAP staining was co-localized with SorCS2 staining as indicated by arrows. No SorCS2 staining was seen in brain slices from 2-month-old *Sorcs2^-/-^* mice. Bars correspond to 50 μm.

### Reduced neurovascular coupling in *Sorcs2^-/-^* mice

Neurovascular signaling was measured as relative changes in parenchymal perfusion (Fig. 3A and 3B) in the primary sensory cortex in response to whisker stimulation.^17^ The increase in parenchymal perfusion upon whisker stimulation was reduced in *Sorcs2^-/-^* mice compared to wild type mice (Fig. 3C). Laser speckle contrast imaging was also used to assess changes in arterial blood flow and arterial diameter in response to whisker stimulation (Fig. 4). Resting inner diameter of the supplying middle cerebral artery branch was similar in both genotypes (Fig. 3D). Whisker stimulation was associated with dilation of this artery, but the vasodilation observed *Sorcs2^-/-^* mice was smaller than that observed in wild type mice (Fig. 3E). Accordingly, arterial blood flow increase in response to neuronal excitation was reduced in *Sorcs2^-/-^* mice (Fig. 3F).

**Figure 3.**
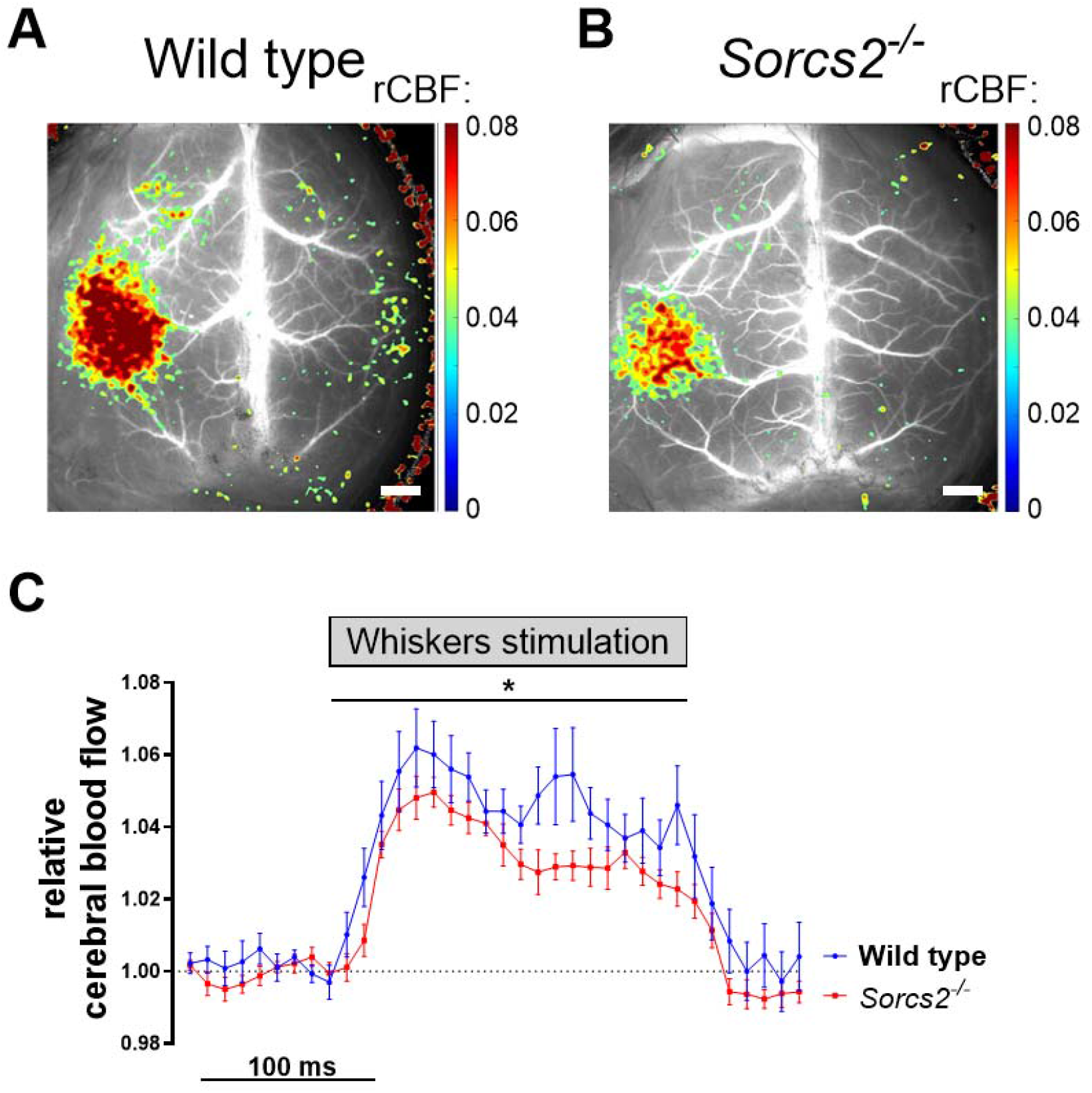
*Sorcs2^-/-^* mice exhibited reduced neurovascular responses *in vivo*. An averaged blood flow map (gray scale) with an overlaid relative cerebral blood flow response (rCBF; color) to whisker stimulation, which induced a local change in blood perfusion of the primary somatosensory cortex in wild type (A) and *Sorcs2^-/-^* mice (B). Bars correspond to 1000 μm. Increase in parenchymal perfusion of somatosensory cortex in response to contralateral whisker stimulation was reduced in *Sorcs2^-/-^* mice (*n* = 9) compared to wild type mice (*n* = 8; C). Responses in (C) were compared using two-way ANOVA. *, *P* < 0.05.

**Figure 4.**
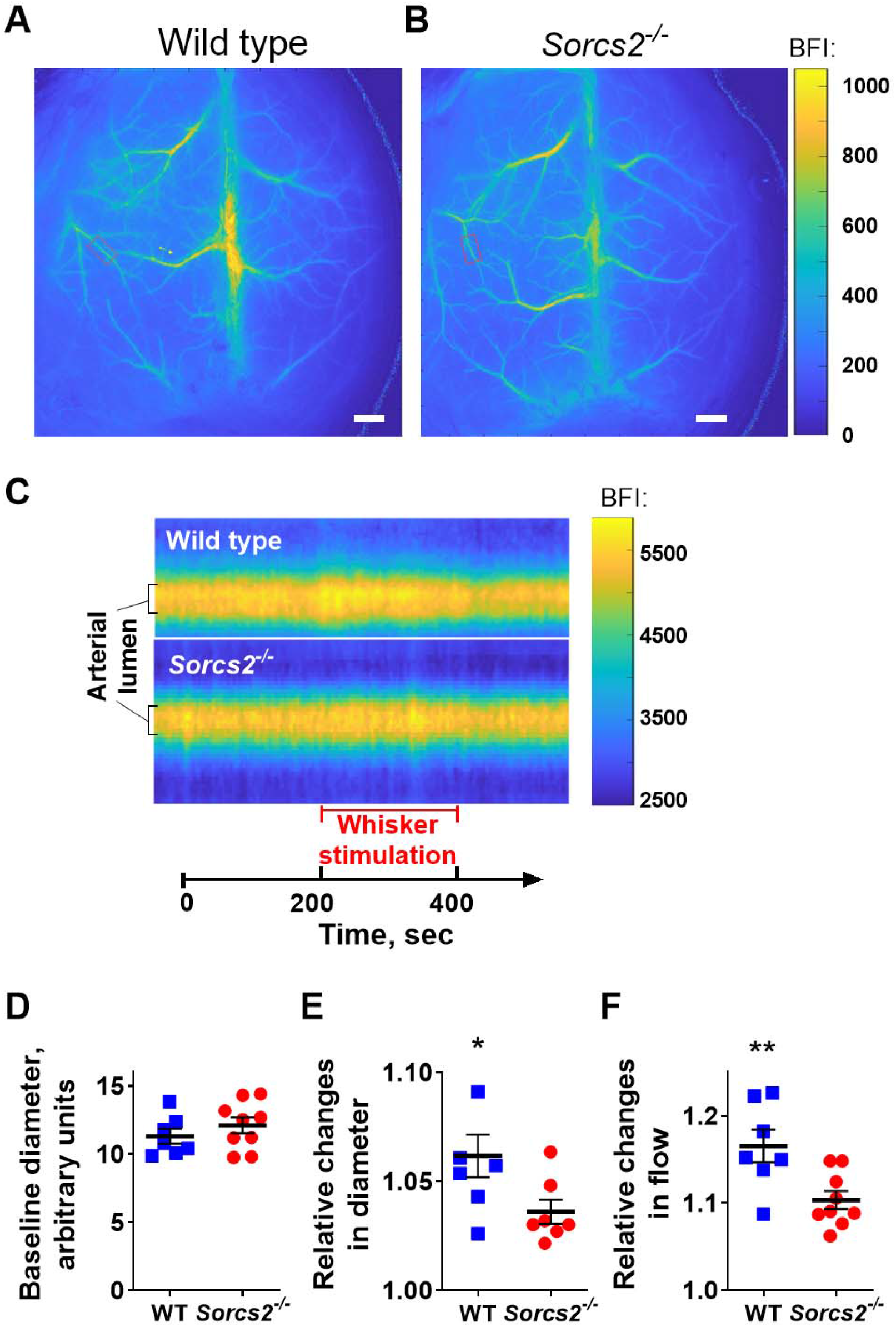
*Sorcs2^-/-^* mice showed reduced increase in vascular diameter and blood flow in response to sensory stimulation. Laser Speckle contrast imaging was used to assess changes in arterial diameter and arterial blood flow increase in response to whisker stimulation in wild type (WT) (A) and *Sorcs2^-/-^* mice (B). Color scale shows blood flow index (BFI). Bar correspond to 1000 μm. The red rectangle identifies the 3^rd^ branch of middle cerebral artery used for single vessel analysis. Representative single artery segmentation profiles show changes in diameter and blood flow in response to whisker stimulation, as indicated in X-axis (C). Color code corresponds to BFI. Y-axis indicates cross-section coordinate in respect to center of the vessel, i.e., inner diameter can be identified. There was no difference in baseline diameter of the 3^rd^ branch of middle cerebral artery between *Sorcs2^-/-^* and WT mice (D). Averaged peak responses to whisker stimulation revealed that *Sorcs2^-/-^* mice showed smaller increase in diameter (E) and blood flow (F) in the 3^rd^ branch of middle cerebral artery compared to WT. * and **, *P* < 0.05 and 0.01 (unpaired *t*-test). *n* = 7 - 9.

### Reduced intracellular Ca^2+^ signaling in astrocytes from *Sorcs2^-/-^* mice

Electric field stimulation of brain slices has previously been shown to excite neurons and lead to delayed elevation of intracellular Ca^2+^ in astrocytes.^17^ Electric field stimulation induced smaller elevation of intracellular Ca^2+^ in astrocytic endfeet in brain slices from *Sorcs2^-/-^* mice compared to wild type mice (Fig. 5). The slope of intracellular Ca^2+^ raise was also slower in astrocytes knocked out for SorCS2 than in wild type controls (0.047 ± 0.022 a.u./s vs. 0.212 ± 0.039 a.u./s, *n* = 4 - 5, *P* = 0.002).

**Figure 5.**
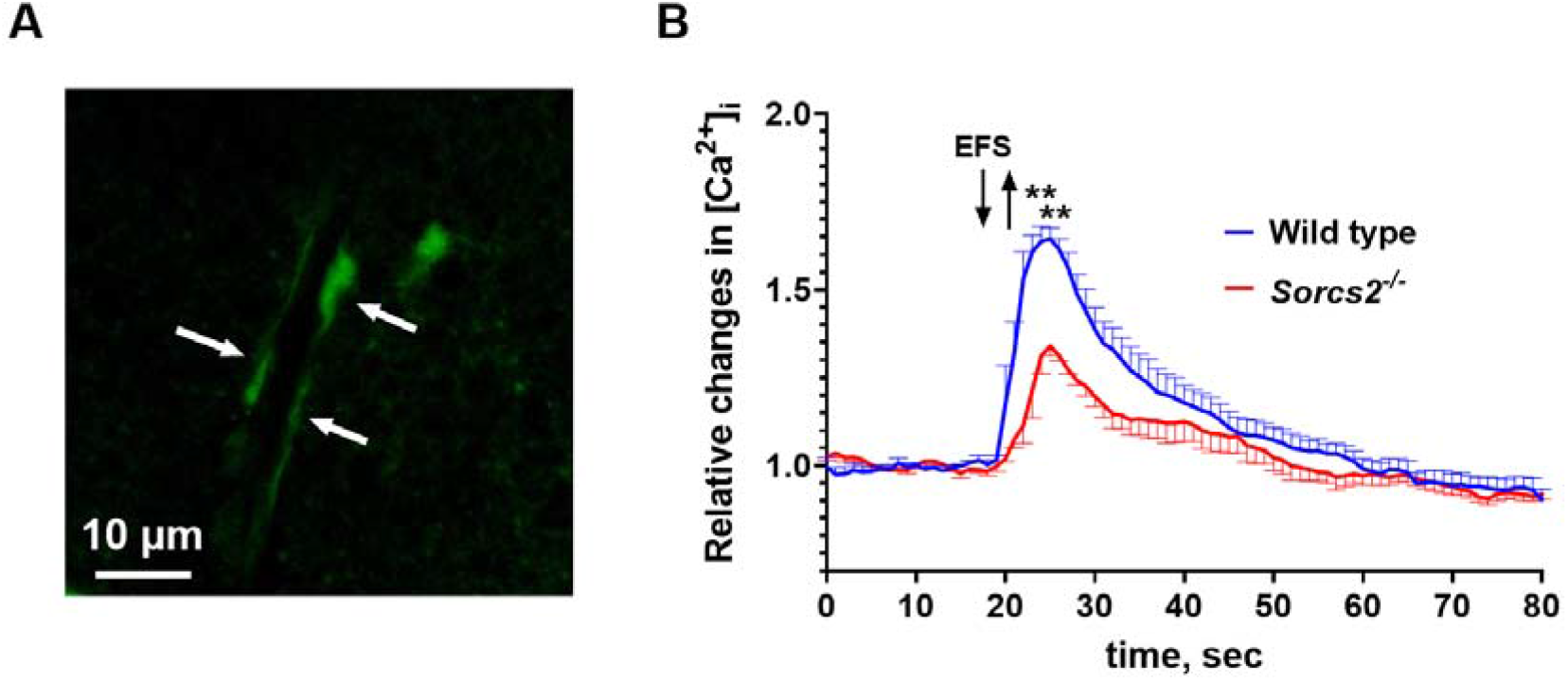
Astrocytes in brain slices from *Sorcs2^-/-^* mice showed reduced increase in intracellular Ca^2+^ in response to neuronal excitation. Brain slices were loaded with Ca^2+^ sensitive dye, Calcium Green-1/AM and that was preferentially allocated to astrocytes. White arrows indicate astrocytic endfeet surrounding a parenchymal arteriole (A). Electric field 2+ stimulation increased intracellular Ca^2+^ in astrocytes and this increase was larger in the brains from wild type mice (*n* = 5) than in brain slices from *Sorcs2^-/-^* mice (*n* = 4) (B). ** *P* < 0.01 (two-way ANOVA with Sidak’s multiple comparisons test).

### Similar endothelial function in cerebral arteries from *Sorcs2^-/-^* and wild type mice

Inner diameter of relaxed middle cerebral arteries *ex vivo* was similar between the groups (Fig. 6A). Thromboxane A2 analog, U46619, was used to pre-constrict arteries to 80% of their maximal tone. The level of preconstriction was the same for both wild type and *Sorcs2^-/-^* arteries (0.39 ± 0.08 mN/mm (*n* = 11) vs. 0.36 ± 0.03 mN/mm (*n* = 8), *P* = 0.69, for wild type and SorCS2 knockout mice, respectively). The acetylcholine-induced relaxation by 10^-5^ M carbachol was similar in both groups (Fig. 6B). This suggests that the endothelium-dependent vasorelaxation of cerebral arteries from SorCS2 knockout mice is unchanged in comparison with wild type mice.

**Figure 6.**
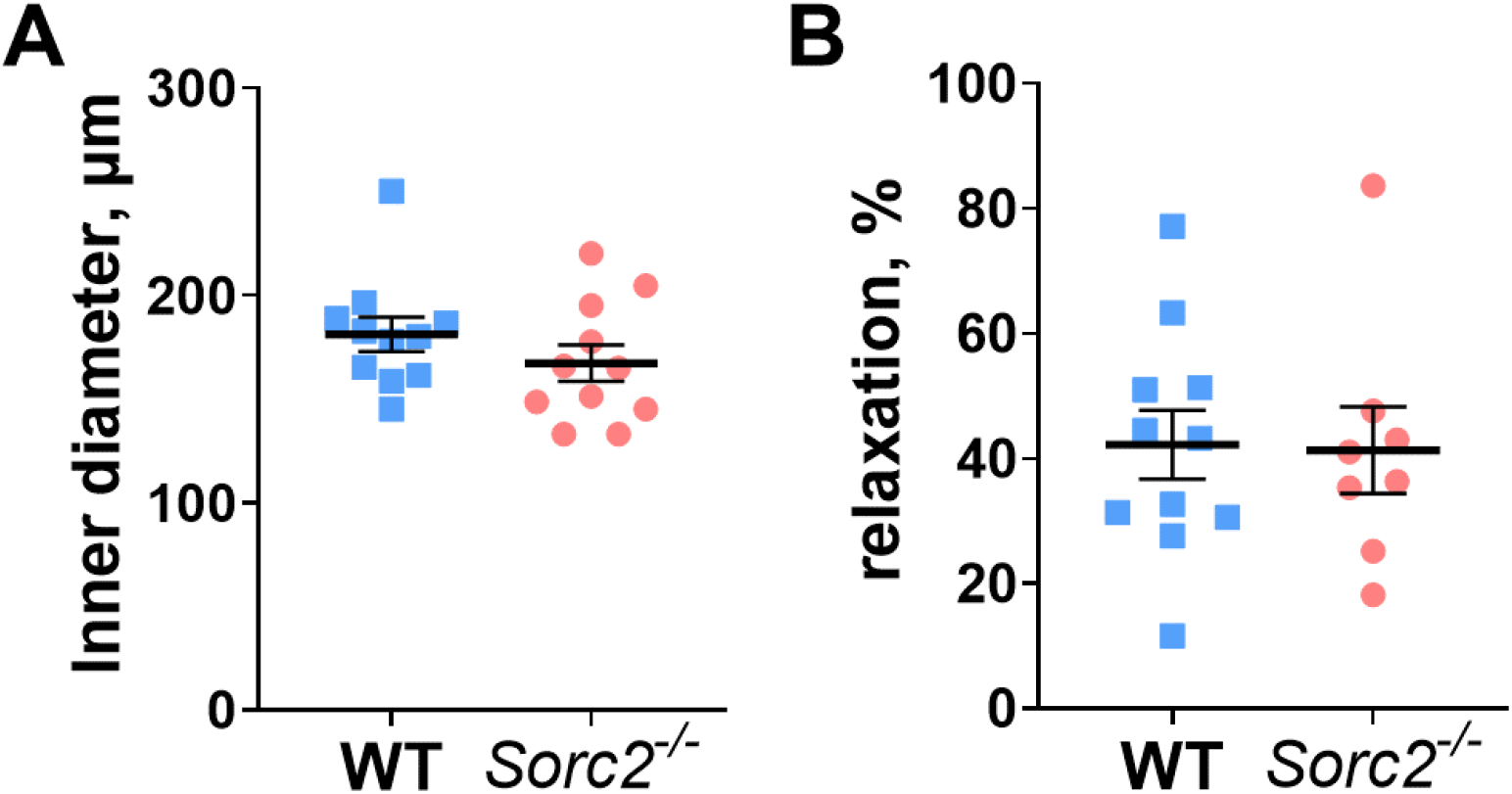
Endothelium-dependent vasorelaxation of the middle cerebral artery was unchanged in *Sorcs2^-/-^* mice. The diameter of isolated middle cerebral arteries from wild type (WT) and *Sorcs2^-/-^* mice was not different (A). There was no difference in the vasorelaxation in response to 10^-5^ M carbachol between wild type and *Sorcs2^-/-^* arteries. *n* = 8 – 11. *P* < 0.001.

### Reduced glutamatergic signaling linked to weakened astrocytic Ca^2+^ responses in *SorCS2^-/-^* mice

Proteomics analyses of isolated astrocytes from P8 mice identified overall 3210 proteins (Expanded View Table 1). Using a cut-off *P* < 0.05, 194 proteins were identified as differently expressed in wild type and *Sorcs2^-/-^* groups (*n* = 4); 84 were upregulated and 110 were downregulated in astrocytes from *Sorcs2^-/-^* mice (Fig. 7A). In astrocytes from 2-month-old mice, 3668 proteins were identified (Expanded View Table 2), among them 245 proteins were differently expressed (120 upregulated and 125 downregulated in *Sorcs2^-/-^)* in wild type and *Sorcs2^-/-^* mice (*n* = 4) (Fig. 7B).

**Figure 7.**
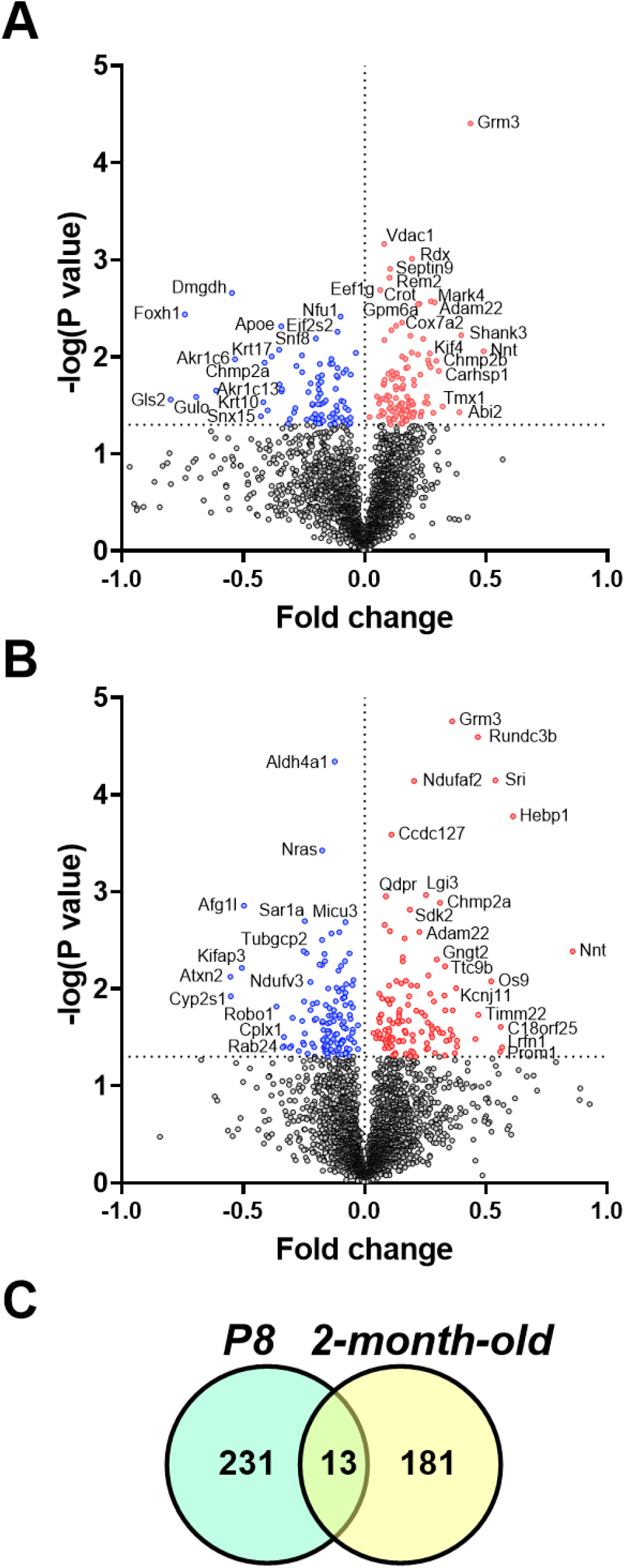
Volcano plot of differentially expressed proteins in astrocytes from wild type and *Sorcs2^-/-^* mice. This plot depicts fold changes in protein expression in astrocytes from *Sorcs2^-/-^* vs. wild type mice correlated to the probability that the protein is differentially expressed. The comparisons were done for astrocytes isolated from P8 mice (A) and 2-month-old mice (B). *P* < 0.05 was set as the significant threshold for differential expression and depicted on the graphs with horizontal dotted line. Dots in red denote significantly upregulated proteins, dots in blue denote significantly downregulated proteins. Gray dots indicate proteins that were not significantly different. Venn diagram (C) shows the number of overlapping differently expressed proteins, which expression was significantly changed in P8 and 2-month-old *Sorcs2^-/-^* mice vs. wild type mice. *n* = 4.

Importantly, metabotropic glutamate receptor 3 (Grm3) was shown significantly upregulated in astrocytes from SorCS2 knockout mice of both ages (Table 1; Expanded View Tables 1 and 2). This upregulation was the most prominently expression difference between the two genotypes in both age groups (Fig. 7A and 7B) and suggests modifications in glutamatergic signaling. Accordingly, both excitatory amino acid transporter 1 (EAAT1) and alanine serine cysteine transporter 1 (ASCT1) were upregulated in astrocytes from P8 *Sorcs2^-/-^* mice (Fig. 8A). The glutamatergic signaling underwent more significant remodeling in 2-month-old *Sorcs2^-/-^* astrocytes where both AMPA- (GluR2) and NMDA-glutamatergic (NMDAR1) receptors were downregulated (Fig. 8B). Moreover, mitochondrial pathways responsible for glutamate metabolization (i.e., glutamic-oxaloacetic transaminase 2 and glutamate dehydrogenase 1) were downregulated, as well as proteins that may contribute to glutamate secretion (i.e., syntaxin-1; synaptotagmin 1 and synaptojanin 1) (Expanded View Table 2). Altogether, this suggests that *Sorcs2^-/-^* astrocytes have suppressed glutamatergic signaling.

**Table 1.** Differentially expressed proteins, which expression was changed in the same direction in astrocytes from both P8 and 2-month-old *Sorcs2^-/-^* mice in comparison with wild type control.

| Protein | Abbreviation | SorCS2<sup>-/-</sup> vs. WT |  | Biological function |
| --- | --- | --- | --- | --- |
|  |  | P8 | 2-month-old |  |
| Disintegrin and metalloproteinase domain-containing protein 22 | Adam22 | 1.36 | 1.25 | Synapse formation |
| Heme-binding protein 1 | Hebp1 | 1.19 | 1.71 | Mitochondrial function |
| Matrin-3 | Matr3 | 0.83 | 0.77 | Cell differentiation |
| Metabotropic glutamate receptor 3 | Grm3 | 1.58 | 1.44 | Glutamate signaling |
| Mitochondrial NAD(P) transhydrogenase | Nnt | 1.63 | 2.27 | Mitochondrial function |
| Protein phosphatase 1 regulatory subunit 21 | Ppp1r21 | 1.02 | 1.10 | Endosomal sorting/maturation |
| Trafficking protein particle complex subunit 10 | Trappc10 | 0.75 | 0.87 | Intracellular trafficking |

**Figure 8.**
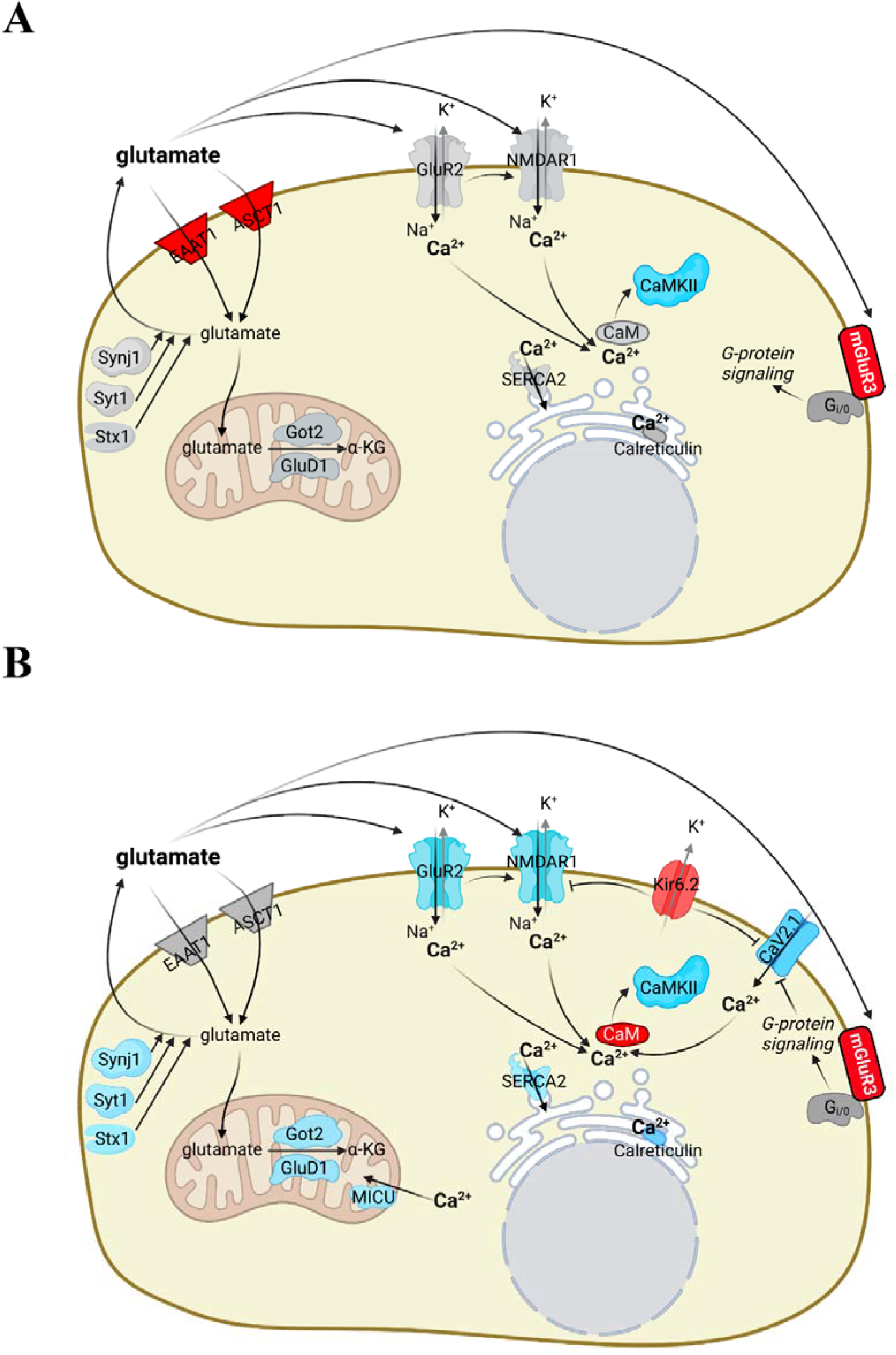
Ingenuity Pathway Analysis suggests suppression of glutamatergic signaling that linked to reduced intracellular Ca^2+^ signaling in astrocytes from *Sorcs2^-/-^* mice in comparison with wild type mice. Several differentially expressed proteins in astrocytes from P8 (A) and adult (B) mice are involved in glutamatergic signaling. These proteins were either downregulated (blue) or upregulated (red) or unchanged (grey) in Sorcs2^-/-^ mice in comparison with wild type. See also Supplementary excel files 1 and 2. EAAT1, excitatory amino acid transporter 1; ASCT1, alanine serine cysteine transporter 1; mGluR3, metabotropic glutamate receptor 3; GluR2, glutamate ionotropic receptor AMPA type subunit 2; NMDAR1, N-methyl-D-aspartate receptor (NMDAR) subunit 1; Kir6.2, ATP-sensitive K^+^ channel pore subunit 6.2; CaV2.1, P/Q voltage-dependent calcium channel; CaMKII, Calcium/calmodulin-dependent protein kinase II; CaM, calmodulin; SERCA2, sarco/endoplasmic reticulum Ca^2+^-ATPase 2; MICU3, mitochondrial calcium uptake protein 3; Stx1, syntaxin-1; Syt1, synaptotagmin 1; Synj1, synaptojanin 1; Got2, glutamic-oxaloacetic transaminase 2; and Glud1, glutamate dehydrogenase 1.

Suppressed glutamatergic signaling may lead to reduced intracellular Ca^2+^ signaling as seen in *Sorcs2^-/-^* mice (Fig. 3). In fact, reduced intracellular Ca^2+^ signaling in astrocytes from *Sorcs2^-/-^* mice was also suggested based on downregulation of Ca^2+^/calmodulin (CaM)-dependent protein kinase II (CaMKII) (Fig. 8). Moreover, reduction in the AMPA- and NMDA-receptors is expected to reduce Ca^2+^ influx (Fig. 8B). Upregulation of an ion channel subunit of the ATP-dependent K^+^ channels (Kir6.2) in the astrocytes from 2-month-old *Sorcs2^-/-^* mice may also indirectly reduce Ca^2+^ influx by hyperpolarizing membrane potential. Accordingly, the neuronal P/Q voltage-gated Ca^2+^ channel was also found downregulated in 2-month-old *Sorcs2^-/-^* (Fig. 8B). Downregulation of the sarco/endoplasmic reticulum Ca^2+^-ATPase and endoplasmic reticulum Ca^2+^ binding protein, calreticulin, further support the suggestion about suppressed intracellular Ca^2+^ signaling (Fig. 8B). Hence, proteomics suggested suppressed glutamatergic signaling associated with reduced intracellular Ca^2+^ signaling. Changes in these important signaling pathways may have important consequences for astrocytic phenotype and function.

Comparison of the results obtained from P8 and 2-month-old mice highlighted 13 proteins that were different between the genotypes in both age groups (Fig. 7C) and only 7 of them were changed in the same direction in both ages (Table 1). Notably, several of these differentially expressed proteins are known to modulate mitochondrial health and function (Table 1). Accordingly, oxidative phosphorylation was suggested to be diminished in 2-month-old *Sorcs2^-/-^* mice (Fig. 9; Expanded View table 3). The proteomics data proposed also changes in intracellular trafficking and protein turnover in astrocytes from *Sorcs2^-/-^* mice of both ages (Table 1; Expanded View Table 3). The overall number of canonical pathways, which were suggested to be significantly modified in *Sorcs2^-/-^* mice and are relevant to cerebral functions, was increased with age (Fig. 9; Expanded View Table 3). This further supports the suggestion that astrocytic phenotype in *Sorcs2^-/-^* mice changes with age.

**Figure 9.**
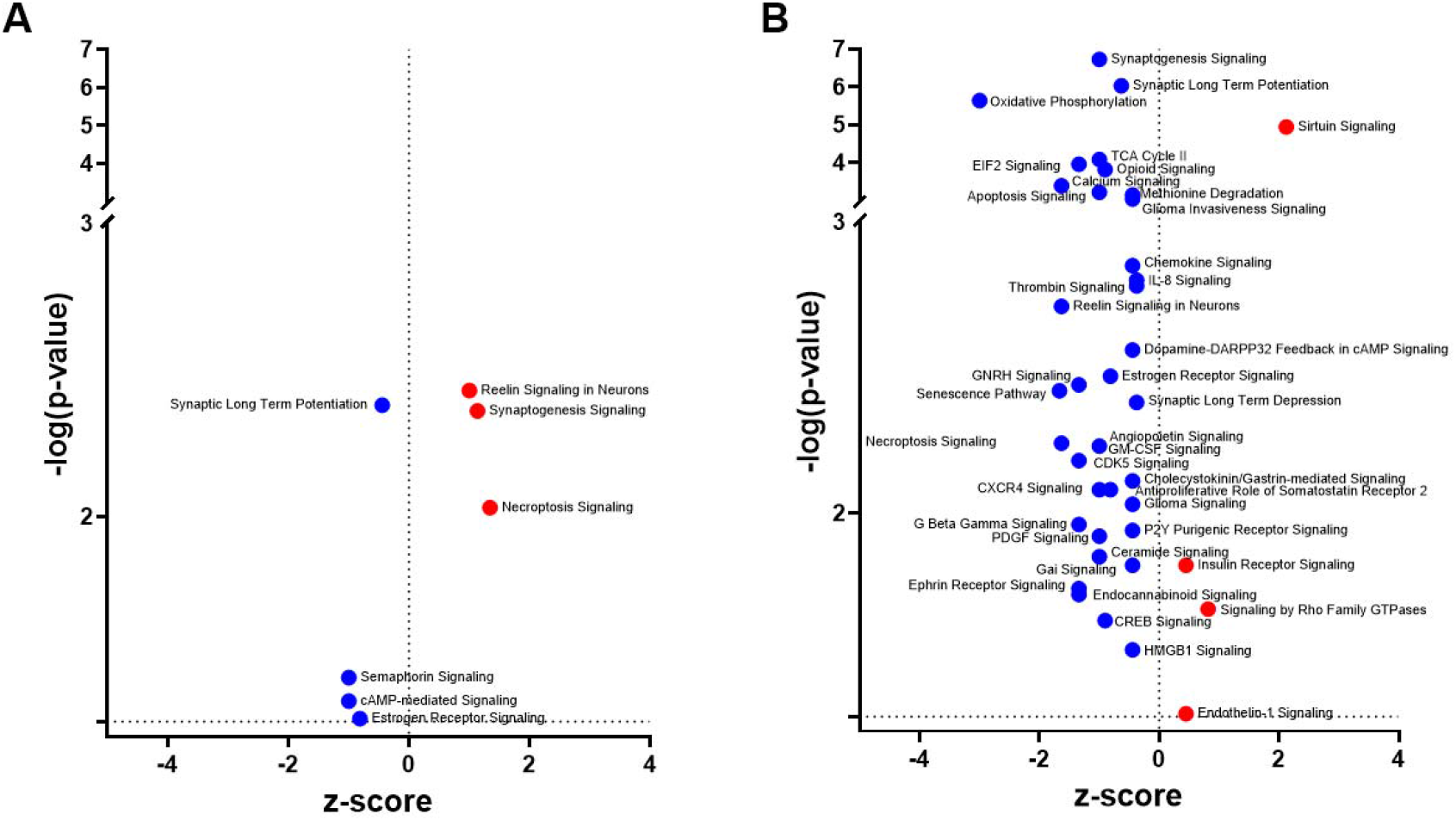
Ingenuity canonical pathways modified in astrocytes from P8 (A) and 2-month-old (B) *Sorcs2^-/-^* mice. Gene ontology pathways were analyzed for enrichment of differentially expressed proteins. A negative z-score (blue) indicates suppression of the pathway and a positive z-score (red) indicates enhancement in *Sorcs2*^-/-^ mice. The significance of changes is predicted by the Ingenuity Pathway Analysis (Qiagen). See also Expanded View Table 3. The horizontal dashed line indicates *P* = 0.05.

## Discussion

This study addressed the importance of astrocytic SorCS2 protein for integrity of neurovascular coupling. Using conventional SorCS2 knockout in mice, we found that the knockout of SorCS2 in the developing brain leads to diminished neurovascular signaling later in life. We identified that this was associated with reduced intracellular Ca^2+^ signaling in astrocytes from *Sorcs2^-/-^* mice upon neuronal excitation. Our data suggested that SorCS2 is important for sensing neuronal activity by astrocytes and, thus, explains why *Sorcs2^-/-^* is associated with disrupted cerebrovascular autoregulation.

### Importance of SorCS2 in neurovascular coupling

SorCS2 has originally been shown to be highly expressed in the mature central nervous system with primary localization to neurons.^3^ Previous papers have reported that SorCS2 is only sparsely expressed in astrocytes of healthy adult subjects.^4, 5, 10^ In the present study, we provided evidence that SorCS2 is expressed in some astrocytes in the brain of adult mice. However, the expression of SorCS2 in astrocytes was lower in the adult brain compared to P8 mice, where the expression was seen in almost all astrocytes. We found that astrocytic expression of SorCS2 was particularly dominant in astrocytic endfeet. The high expression of SorCS2 in astrocytic endfeet may suggest that SorCS2 is important in the development and function of the brain’s neurovascular unit.^12, 18^

Neuronal activity is sensed by astrocytes that express neurotransmitter receptors including glutamate receptors.^19, 20^ Glutamate receptors are localized in the astrocyte cell membrane in close proximity to neuronal synapses.^21^ Activation of the glutamate receptors upon neuronal excitation leads to increase of intracellular Ca^2+^ in astrocytes. ^14, 22^ This Ca^2+^ signal spreads to the astrocytic endfeet wrapping parenchymal arterioles.^12, 23^ Elevation of intracellular Ca^2+^ in astrocytic endfeet initiates a release of vasoactive substances relaxing arterioles and thus, increase blood flow to active neurons.^24^ In this study, we identified changes in astrocytes that may underlie the observed reduction of neurovascular signaling in *Sorcs2^-/-^* mice. We showed that knockout of SorCS2 was associated with reduced expression of AMPA and NMDA receptors in astrocytes. Interestingly, other studies have shown that SorCS2 regulated GluN2B expression in postsynaptic densities and ionotropic synaptic plasticity in the hippocampus, substantiating the importance of the receptor in controlling neurotransmission.^3, 16^ Such expression change may reduce the astrocytes’ sensitivity in sensing neuronal activity and thus, lead to reduced astrocytic Ca^2+^ responses.^25^ Accordingly, we observed significantly reduced astrocytic Ca^2+^ responses upon neuronal excitation. The glutamate-triggered Ca^2+^ response in astrocytes depends on a wide spectrum of proteins involved in Ca^2+^ homeostasis^12^ including Ca_V_2.1 channels and CaMKII^27^ that were both reduced in *Sorcs2^-/-^* mice. Furthermore, several Ca^2+^-dependent signaling pathways were predicted to be changed in astrocytes from *Sorcs2^-/-^* mice, including Ca^2+^ uptake by mitochondria^28^ and reduced capacity for storing Ca^2+^ in the sarcoplasmic reticulum.^29–30^ Hence, these changes may be a part of the explanation for the observed reduced Ca^2+^ responses in astrocytes from *Sorcs2^-/-^* mice. The reduced intracellular Ca^2+^ signaling can in turn reduce the exocytotic release of glutamate from astrocytes that can further reduce Ca^2+^ signaling leading to the negative feedback loop.^26^ Accordingly, mitochondrial pathways responsible for glutamate metabolization (Got2^31^ and Glud1^32^) and regulators of intracellular vesicle recycling and secretion that contribute to glutamate release and homeostasis (Synj1,^33^ Syt1^34^ and Stx1^35^) were found downregulated in astrocytes from 2-month-old *Sorcs2^-/-^* mice.

To summarize, our data suggest that SorCS2 is necessary for balanced expression of neurotransmitter receptors and key proteins in astrocytes involved in sensing neuronal activity. This is in line with the overall role of VPS10P domain receptor family proteins, including SorCS2, whose primary function is thought to be expressional regulation of a number of important proteins in the central nervous system.^36^

### SorCS2 signaling is not involved in endothelium-dependent relaxation of cerebral arteries

Recent studies have suggested that endothelium is an important component in the neurovascular unit.^17, 37^ We observed suppressed relaxation of the branch of the middle cerebral artery supplying the sensory cortex in response to whisker stimulation in *Sorcs2^-/-^* mice. We mounted therefore the isolated middle cerebral artery in the myograph to test whether changed endothelial function may underlie the reduced vasorelaxation. We found that endothelium-dependent vasorelaxation was similar in *Sorcs2^-/-^* and wild type mice. This observation seems to be in line with our immunohistochemical finding that SorCS2 is not expressed in the wall of brain arterioles. Thus, our data indicate that the importance of SorCS2 in the neurovascular unit is primarily attributed to astrocytes. Interestingly, previous characterization of SorCS2 expression in various organs suggested that SorCS2 is not expressed in large arteries of healthy subjects but prominently expressed in pulmonary arterioles.^38^ SorCS2 expression in the vasculature therefore seems to be organ specific.

### SorCS2 knockout leads to increased metabotropic glutamate receptor 3 expression

In both P8 and adult *Sorcs2^-/-^* mice, we observed increased expression of metabotropic glutamate receptor 3. Grm3 was previously demonstrated not to be involved in rapid astrocytic Ca^2+^ increases upon neuronal excitation.^39^ Interestingly, this is not the first time an increased expression of Grm3 has been reported in SorCS2 knockout mice.^5^ Under physiological conditions, Grm3 is expressed in astrocytes at all developmental stages.^39^ Since Grm3 activation can potentiate synthesis of glutathione, it was shown to be important in protection against oxidative stress.^40^ However, our proteomics analyses did not suggest any significant changes in oxidative stress response in astrocytes from neither P8 nor 2-month-old *Sorcs2^-/-^* mice. The reason and mechanism underlying increased Grm3 expression in *Sorcs2^-/-^* mice remains to be investigated.

### Neurovascular uncoupling may contribute to neurological abnormalities associated with changes in SorCS2 expression

A previous genome-wide association study suggested that SorCS2 variants increase predisposition to ischemic stroke.^6^ Accordingly, SorCS2 polymorphism was shown to be related to the prevalence of ischemic stroke in the Japanese population.^41^ There is evidence that the role of SorCS2 in stroke pathology is related to its role in oxidative stress response protecting the neuronal function.^5^ In spite of the minimal SorCS2 expression in healthy astrocytes, it markedly upregulates after ischemia and contributes to control the release of endostatin, a factor linked to post-stroke angiogenesis and neovascularization in the ischemic brain.^4^ In the present study, we suggest an additional mechanism that is modulated by SorCS2 and implicated in multiple neuropathological conditions. Disturbance in neurovascular coupling is involved in the pathology of stroke, Alzheimer’s disease, migraine and several other neurodegenerative diseases.^12^ Altogether, our results suggest that SorCS2-dependent regulation of glutamatergic and intracellular Ca^2+^ signaling in astrocytes is important for neurovascular coupling.

## Methods

All animal experiments conformed to guidelines from the European Communities Council Directive (86/609/EEC) for the Protection of Animals used for Experimental and other Scientific Purposes. The experiments were conducted with permission from the Animal Experiments Inspectorate of the Danish Ministry of Environment and Food (2016-15-0201-00982 and 2019-15-0201-00341) and reported in accordance with the ARRIVE (Animal Research: Reporting in vivo Experiments) guidelines.

### *Sorcs2^-/-^* mice

The *Sorcs2^-/-^* mice were generated as described previously.^10^ As no difference in experimental results between sexes was seen, equal number of male and female mice were included. Mice were housed at the Animal Facility at Aarhus University, Department of Biomedicine under a 12:12 light/dark cycle, and food and water was provided ad libitum.

### Assessment of neurovascular coupling using laser speckle contrast imaging

*In vivo* laser speckle contrast imaging was used to assess the changes in tissue perfusion and blood flow in response to neuronal excitation. Mice were anaesthetized with ketamine (80 mg/kg s.c., Ketaminol®vet, Intervet International, Boxmeer, The Netherlands) and xylazine (8 mg/kg s.c., Narcoxyl®vet, Intervet International). Maintenance of anesthesia was obtained by s.c. injection of 1/4 of the initial dose every 45 min. Preparation and experimental setup details were described previously.^17^ The mouse was placed on a heating pad and the head was fixed in a stereotaxic frame. The skin and periosteum were removed to expose the cranial surface. The recordings were made with an intact cranium to avoid changes in intracranial pressure. Temporal laser speckle contrast analysis was applied to diminish effects of static scattering from the skull.^42^ A coverslip was mounted in a 1 % agarose gel in 0.9% NaCl on top of the cranium to minimize optical reflections. At the end of experiment, anaesthetized mice were sacrificed by cervical dislocation.

Coherent near infrared light was delivered using a semiconductor laser diode (785nm, LDM785), controlled by the diode driver CLD1011LP (Thorlabs Inc., Newton, NJ). Backscattered light was recorded by a CMOS camera (acA2000-165umNIR; Basler AG, Ahrensburg, Germany) mounted on the video lens (VZM 200i, Edmund Optics, Barrington, NJ). Speckle images were then processed using temporal speckle contrast analysis^42^ and converted to arbitrary perfusion values as 1/K^2^, where K is the contrast calculated over 25 frames. Sensory cortex neurons were excited by mechanical whisker stimulation with 8 mm amplitude and 4 Hz frequency.

After baseline recording, 200 sec of whisker stimulation was applied on one side of the head. After 10 min rest, the experiment was repeated with whisker stimulation of the opposite side. The results of whisker stimulation were averaged for each mouse. The neurovascular coupling responses were assessed as an elevation in tissue perfusion at a region of interest in the second posterior bifurcation of the middle cerebral artery corresponding to primary somatosensory cortex.^43^ The changes in blood flow and arterial diameter were measured in 2^nd^ order middle cerebral artery that supply primary somatosensory cortex. Changes in perfusion were estimated in comparison to the baseline signal taken as 1. Data were analyzed using MATLAB software (ver. R2017b, MathWorks, Natick, MA). A segmentation algorithm was applied to extract blood flow and arterial diameter changes in middle cerebral artery.^44^

### Assessment of neurovascular coupling in brain slices

Mice were euthanized by cervical dislocation. The brain was transferred into ice-cold artificial cerebrospinal fluid (aCSF; in mM: NaCl 125, KCl 3, NaHCO_3_ 25.9, NaH_2_PO_4_ 1.25, MgCl_2_ 1, CaCl_2_ 2, L-ascorbic acid 0.4, glucose 4, gassed with 95 % O_2_ and 5 % CO_2_, and adjusted to pH 7.4) aerated with 95% O_2_ and 5% CO_2_. The prefrontal cortex located was sliced into 160 μm slices using vibratome (1200s, Leica Biosystems, Germany). Slices were incubated with Ca^2+^-sensitive fluorescent dye Calcium Green AM (5.33 mg/l; ThermoFisher Scientific, USA). For this purpose, 10 μg Calcium Green AM was diluted in 1.63 μl dimethylsulfoxide, DMSO; 0.33 μl Cremophor EL; 0.07 mg Pluronic acid, F127, and then, added to 1.88 ml pre-aerated a-CSF. During the incubation, the slices were heated at 37°C under constant gassing with 95% O_2_ and 5% CO_2_.

After 90 min incubation, the brain slices were washed in pre-heated and aerated aCSF and examined with confocal microscope (LSM 780, Zeiss, Germany). During the experiments, the slices were continuously superfused at 5 ml/min flow with aerated aCSF through the heating device (SH-27B / TC324B; Warner Instruments, Hamden, CT, USA). Neurons in the brain slices were stimulated with electric field stimulation (32 Hz, 14 V, 0.3 ms pulses, stimulation period 3 s) using two platinum electrodes placed 4 mm apart from each other. The electric field stimulation was repeated 5 times in each slice with 15 min interval between stimulations. The intracellular Ca^2+^ responses from the 5 stimulations in each brain slice were averaged for analysis. Fluorescence intensity was measured in astrocytic endfeet surrounding parenchymal arterioles located 30-50 μm from the cut surface in cortical tissue. An averaged fluorescence intensity over baseline period prior electric field stimulation (18 sec) was taken as 1 and used for calculations of relative changes in intracellular Ca^2+^.

### Isometric myograph

Mice were sacrificed with cervical dislocation. Middle cerebral arteries were dissected from the brain in ice-cold physiological salt solution (PSS; in mM: NaCl 115.8, KCl 2.82, KH_2_PO_4_ 1.18, MgSO_4_ 1.2, NaHCO_3_ 25, CaCl2 1.6, EDTA 0.03, glucose 5.5, gassed with 5 % CO_2_ in air and adjusted to pH 7.4). Two-millimeter arterial segments were mounted in a wire myograph (Danish Myo Technology A/S, Denmark) for isometric force measurements at 37°C.^45^ The artery diameter was set to 0.9 of the passive diameter at 100 mmHg to obtain maximal active force. Force (mN) was recorded with a PowerLab 4/25 – Chart8 acquisition system (ADInstruments Ltd., New Zealand) and converted to wall tension by dividing the force with double length of the arterial segment.

### Immunohistochemistry

Immediately after cervical dislocation, the mice were exsanguinated, and brain tissue was dissected. Tissues were fixed in 4% paraformaldehyde in Phosphate Buffered Saline (PBS; in mM: NaCl 137, KCl 2.7, Na_2_HPO_4_ 8.2, KH_2_PO_4_ 1.8, at pH 7.4) at 4 °C overnight, cryoprotected in 30% w/v sucrose in PBS at 4°C for 24 - 48 h, frozen in Tissue-Tek O.C.T (Sakura Finetek Europe, Alphen aan den Rijn, Netherlands) and cryosectioned at 12 μm. Antigen □ retrieval was performed by incubating the tissue for 2 h at 70 °C in Target Retrieval Solution (Dako, Santa Clara, CA, USA) using a 1:10 dilution in demineralized water. Sections were then cooled to room temperature, followed by a washing step in PBS for 10 minutes. The brain slices were incubated for 30 min in 1% bovine serum albumin (BSA) in PBS with 0.1% Triton X-100 (Tx), followed by overnight incubation at 4°C in 50 nM Tris-based buffer solution with 1% BSA containing anti-SorCS2 antibody (dilution 1:100; AF4237, R&D, Minneapolis, MN, USA), anti-Aquaporin4 antibody (1:2000; AQP-014, Alomone, Jerusalem, Isreal), Alexa Fluor® 647 galactosyl-specific isolectin (1:200; I32450, Thermo Fischer, Waltham, MA, USA), and anti-GFAP antibody (1:1000; Z0334, Dako). After washing out, specific fluorescent immunostainings were visualized using Alexa488-conjugated anti-sheep IgG and Alexa546-conjugated anti-rabbit IgG (1:300; Molecular Probes, Eugene, OR, USA).

### Flow cytometric isolation of astrocytes from mouse brain

#### Single Cell Suspension Preparation

Adult and P8 mice were perfused with ice-cold PBS, decapitated and the brains were collected. Dissected cortexes were collected in 24 well-plate and cut into small pieces to be digested in 1.5 ml of digestion buffer containing 1 mg/ml of collagenase/dispase cocktail (Roche) and 0.5 mg/ml of DNase I. Samples were incubated at 37°C for 40 min and the digestion was stopped by adding 10 mM EDTA pH 8. Cells were resuspended by gentle pipetting and cell suspensions were passed through a nylon mesh (pore size: 70 nm). 25 ml of 2% Fetal Bovine Serum (FBS) in Hank’s balanced salt solution (HBSS) were added to each sample. Cells were centrifuged 8 min at 350x g at 4°C and each pellet was resuspended with 10 ml of 37% density gradient medium Percoll (Cytiva). After a 20 min centrifugation at 2500x g at RT, the layer of myelin was discarded by using a pipette. 25 ml of HBSS-FBS were added and samples were centrifuged 8 min at 350x g at 4°C. Supernatants were removed and each pellet was dissolved in 300 μl of 0.5% Bovine Serum Albumin (BSA) in Dulbecco’s PBS (d-PBS).

#### Astrocyte labelling

100 μl aliquots of cell suspension were blocked with rat anti-mouse CD16/32 (1:50, BD, cat no: 553142) for 10 min at RT. Cells were afterwards incubated with phycoerythrin (PE) conjugated rat anti-mouse ASCA2 (1:100, Miltenyl Biotech, cat no: 130-123-284) and allophycocyanin (APC) conjugated rat anti-mouse CD45 (1:100, Biolegend, cat no: 103111) in blocking solution for 30 minutes on ice in the dark. 1 ml of 0.5% BSA in d-PBS was added to each sample and cells were centrifuged 10 min at 300x g at 4°C. The supernatant was discarded, and this washing step was repeated. 500 μl of 4% PFA in d-PBS was added to each sample and incubated for 15 min at RT. The cells were centrifuged 7 min at 700x g at 4°C. Each pellet was resuspended in 2.5 mM EDTA, 0.5% BSA in d-PBS.

#### Astrocyte Sorting

Cell sorting was performed using a 3-laser FACSAria^TM^III (BD Biosciences, San Jose, CA, USA), equipped with a 375 nm, 488 nm, and 633 nm laser. Instrument performance is checked on a daily basis by CS&T Quality Control Beads (BD Biosciences) and Accudrop Beads (BD Biosciences). No compensation was needed. Debris was removed in a cell gate, then single cells were selected in a forward scatter A versus forward scatter H plot and finally ASCA2 positive cells were selected and sorted from a CD45-APC versus ASCA2-PE plot (Expanded View figure 1). Cells were sorted into Eppendorf tubes containing 50 μl PBS. A 100 μm nozzle was used and the sort was kept at 4°C. The purity of the sort was determined by reanalyzing a small fraction of the sorted cells. The purity was above 97%. Data was acquired using BD FACSDiva Software version 8.0.2 (BD Biosciences).

### Proteome analysis

Freshly isolated astrocytes were immediately stored at −80°C prior to proteome analysis. Peptides for tandem mass tagging were prepared by incubating isolated astrocyte samples in 8M urea in the presence of 0.5 μg lys-C for four hours at 30°C initiated with 3 cycles (5 min each) in a water sonication bath followed by dilution to 1 M urea by tetraethylammonium bicarbonate (TEAB) and incubation overnight at 30° C after addition of 1 μg trypsin. The resulting peptide samples were labelled with 2 sets of tandem mass tags from a 10-plex Tandem Mass Tags (TMT) set using the mass tags 127N, 127C, 128N, 128C, 129N, 129C, 130N, 130C, and 131. A pool of all samples was labelled with mass tag 126 and served as reference channel (Expanded View Tables 1 and 2). Tagged peptide samples were mixed in two sets of tagged peptide mixture, which each were subsequently fractionated by high pH liquid chromatography, as previously described,^46^ and analysed by reversed phase nanoliquid chromatography tandem mass spectrometry (RP-nanoLC-MSMS), as previously described,^47^ with the exception that the LC gradient was 121 min from 10-14% A to 27-32% buffer A (0.1% Formic acid). All raw data files were processed using the Proteome Discoverer software (v. 2.4.0.305) and searched with the MSPepSearch and the Sequest HT search algorithm. The search parameters for the MSPepSearch were kept at default except for the precursor tolerance and the fragment tolerance were both set to 15 ppm. The TMT specific spectral library was prepared by Shen et al.^48^ and imported into Proteome Discoverer. Sequest HT search parameters were set to default except for MS accuracy of 8 ppm, MSMS accuracy of 0.05 Da for HCD data, with two missed cleavages allowed. Fixed modifications were set to carbamidomethylation at cysteine residues, TMT 6-plex N-terminal and TMT 6-plex on lysine residues. Variable modifications were set to methionine oxidation, deamidation of asparagine and glutamine and N-terminal acetylation. Sequest HT searches were performed against the Uniprot mouse database (25252 entries, downloaded 30^th^ September 2019).

### Ingenuity Pathway Analysis (IPA)

A list of differentially expressed proteins in astrocytes from SoCS2^-/-^ and wild type mice of P8 and 2 months of age was uploaded to IPA for functional interpretation. First, gene ontology pathways were analysed for enrichment of differentially expressed proteins. A negative z-score indicates suppression of that pathway, while a positive z-score indicates enhancement. The list of quantified proteins was uploaded into IPA software (Qiagen, Redwood City, CA, USA) for further analyses, where the proteins, which expression differed significantly between genotypes, were identified and their association with the changes in ingenuity canonical pathways was suggested.

### Statistics

Data are expressed as mean±SEM and *n* equals number of mice. Probability (*P)* values less than 0.05 were considered statistically significant. Data processing and statistical analyses were performed using Excel, Microsoft and with GraphPad Prism 8.4 and 9.2 software. The data were compared using Student’s unpaired *t*-test or two-way ANOVA where appropriate.

## Supporting information

Expanded View Table 1

Expanded View Table 2

Expanded View Table 3

Expanded View figure 1

## Author contributions

C.S, A.N., and V.V.M contributed to the conception and design of the study. C.S, H.L., S.B.A., S.S.N. and H.C.B. carried out the experiments. C.S, H.L., S.B.A., S.S.N., H.C.B., A.N. and V.V.M analyzed and interpreted the data. C.S., H.L., A.N. and V.V.M prepared the draft and finalized the manuscript. All authors provided critical feedback and approved the final version of the manuscript.

## Expanded View tables legends

**Expanded View Table 1.** Proteins mapped by proteomics of freshly isolated astrocytes from P8 *Sorcs2^-/-^* and wild type mice.

**Expanded View Table 2**. Proteins mapped by proteomics of freshly isolated astrocytes from 2-month-old *Sorcs2^-/-^* and wild type mice.

**Expanded View Table 3**. Ingenuity Canonical Pathways suggested by proteomics data analysis modified in P8 and 2-month-old *Sorcs2^-/-^* mice in comparison with matched wild type mice. See also Supplemental Excel File 1 and 2 for all detected proteins.

**Expanded View figure 1. Representative results of flow cytometric isolation of astrocytes from 2-month-old mouse brain.** Left panel shows ungated forward (FSC) and side (SSC) scatter plot of isolated cells. Middle panel identifies populations of ASCA2 positive and CD45 positive cells. Right panel reanalyzes the data to determine a fraction of sorted astrocytes.

## References

1. Malik AR and Willnow TE. VPS10P Domain Receptors: Sorting Out Brain Health and Disease. Trends Neurosci. 2020;43:870–885.

2. Ma Q, Yang J, Milner TA, Vonsattel JG, Palko ME, Tessarollo L and Hempstead BL. SorCS2-mediated NR2A trafficking regulates motor deficits in Huntington’s disease. JCI Insight. 2017;2.

3. Glerup S, Bolcho U, Molgaard S, Boggild S, Vaegter CB, Smith AH, Nieto-Gonzalez JL, Ovesen PL, Pedersen LF, Fjorback AN, Kjolby M, Login H, Holm MM, Andersen OM, Nyengaard JR, Willnow TE, Jensen K and Nykjaer A. SorCS2 is required for BDNF-dependent plasticity in the hippocampus. Mol Psychiatry. 2016;21:1740–1751.

4. Malik AR, Lips J, Gorniak-Walas M, Broekaart DWM, Asaro A, Kuffner MTC, Hoffmann CJ, Kikhia M, Dopatka M, Boehm-Sturm P, Mueller S, Dirnagl U, Aronica E, Harms C and Willnow TE. SorCS2 facilitates release of endostatin from astrocytes and controls post-stroke angiogenesis. Glia. 2020;68:1304–1316.

5. Malik AR, Szydlowska K, Nizinska K, Asaro A, van Vliet EA, Popp O, Dittmar G, Fritsche-Guenther R, Kirwan JA, Nykjaer A, Lukasiuk K, Aronica E and Willnow TE. SorCS2 Controls Functional Expression of Amino Acid Transporter EAAT3 and Protects Neurons from Oxidative Stress and Epilepsy-Induced Pathology. Cell Rep. 2019;26:2792–2804 e6.

6. Kaplan RC, Petersen AK, Chen MH, Teumer A, Glazer NL, Doring A, Lam CS, Friedrich N, Newman A, Muller M, Yang Q, Homuth G, Cappola A, Klopp N, Smith H, Ernst F, Psaty BM, Wichmann HE, Sawyer DB, Biffar R, Rotter JI, Gieger C, Sullivan LS, Volzke H, Rice K, Spyroglou A, Kroemer HK, Ida Chen YD, Manolopoulou J, Nauck M, Strickler HD, Goodarzi MO, Reincke M, Pollak MN, Bidlingmaier M, Vasan RS and Wallaschofski H. A genome-wide association study identifies novel loci associated with circulating IGF-I and IGFBP-3. Hum Mol Genet. 2011;20:1241–51.

7. Baum AE, Akula N, Cabanero M, Cardona I, Corona W, Klemens B, Schulze TG, Cichon S, Rietschel M, Nothen MM, Georgi A, Schumacher J, Schwarz M, Abou Jamra R, Hofels S, Propping P, Satagopan J, Detera-Wadleigh SD, Hardy J and McMahon FJ. A genome-wide association study implicates diacylglycerol kinase eta (DGKH) and several other genes in the etiology of bipolar disorder. Mol Psychiatry. 2008;13:197–207.

8. Ollila HM, Soronen P, Silander K, Palo OM, Kieseppa T, Kaunisto MA, Lonnqvist J, Peltonen L, Partonen T and Paunio T. Findings from bipolar disorder genome-wide association studies replicate in a Finnish bipolar family-cohort. Mol Psychiatry. 2009;14:351–3.

9. Christoforou A, McGhee KA, Morris SW, Thomson PA, Anderson S, McLean A, Torrance HS, Le Hellard S, Pickard BS, StClair D, Muir WJ, Blackwood DH, Porteous DJ and Evans KL. Convergence of linkage, association and GWAS findings for a candidate region for bipolar disorder and schizophrenia on chromosome 4p. Mol Psychiatry. 2011;16:240–2.

10. Glerup S, Olsen D, Vaegter CB, Gustafsen C, Sjoegaard SS, Hermey G, Kjolby M, Molgaard S, Ulrichsen M, Boggild S, Skeldal S, Fjorback AN, Nyengaard JR, Jacobsen J, Bender D, Bjarkam CR, Sorensen ES, Fuchtbauer EM, Eichele G, Madsen P, Willnow TE, Petersen CM and Nykjaer A. SorCS2 regulates dopaminergic wiring and is processed into an apoptotic two-chain receptor in peripheral glia. Neuron. 2014;82:1074–87.

11. Kozareva V, Martin C, Osorno T, Rudolph S, Guo C, Vanderburg C, Nadaf N, Regev A, Regehr WG and Macosko E. A transcriptomic atlas of mouse cerebellar cortex comprehensively defines cell types. Nature. 2021;598:214–219.

12. Nippert AR, Biesecker KR and Newman EA. Mechanisms Mediating Functional Hyperemia in the Brain. Neuroscientist. 2018;24:73–83.

13. Iadecola C. Neurovascular regulation in the normal brain and in Alzheimer’s disease. Nat Rev Neurosci. 2004;5:347–360.

14. Agulhon C, Petravicz J, McMullen AB, Sweger EJ, Minton SK, Taves SR, Casper KB, Fiacco TA and McCarthy KD. What is the role of astrocyte calcium in neurophysiology? Neuron. 2008;59:932–46.

15. Hosford PS and Gourine AV. What is the key mediator of the neurovascular coupling response? Neurosci Biobehav Rev. 2019;96:174–181.

16. Yang J, Ma Q, Dincheva I, Giza J, Jing D, Marinic T, Milner TA, Rajadhyaksha A, Lee FS and Hempstead BL. SorCS2 is required for social memory and trafficking of the NMDA receptor. Mol Psychiatry. 2021;26:927–940.

17. Staehr C, Rajanathan R, Postnov DD, Hangaard L, Bouzinova EV, Lykke-Hartmann K, Bach FW, Sandow SL, Aalkjaer C and Matchkov VV. Abnormal neurovascular coupling as a cause of excess cerebral vasodilation in familial migraine. Cardiovasc Res. 2020;116:2009–2020.

18. Iadecola C. The Neurovascular Unit Coming of Age: A Journey through Neurovascular Coupling in Health and Disease. Neuron. 2017;96:17–42.

19. Parri HR, Gould TM and Crunelli V. Spontaneous astrocytic Ca2+ oscillations in situ drive NMDAR-mediated neuronal excitation. Nature neuroscience. 2001;4:803–12.

20. Porter JT and McCarthy KD. Astrocytic neurotransmitter receptors in situ and in vivo. Progress in neurobiology. 1997;51:439–55.

21. van den Pol AN, Romano C and Ghosh P. Metabotropic glutamate receptor mGluR5 subcellular distribution and developmental expression in hypothalamus. The Journal of comparative neurology. 1995;362:134–50.

22. Petravicz J, Fiacco TA and McCarthy KD. Loss of IP3 receptor-dependent Ca2+ increases in hippocampal astrocytes does not affect baseline CA1 pyramidal neuron synaptic activity. The Journal of neuroscience: the official journal of the Society for Neuroscience. 2008;28:4967–73.

23. Straub SV, Bonev AD, Wilkerson MK and Nelson MT. Dynamic inositol trisphosphate-mediated calcium signals within astrocytic endfeet underlie vasodilation of cerebral arterioles. The Journal of general physiology. 2006;128:659–69.

24. Filosa JA and Blanco VM. Neurovascular coupling in the mammalian brain. Exp Physiol. 2007;92:641–6.

25. Lecrux C and Hamel E. Neuronal networks and mediators of cortical neurovascular coupling responses in normal and altered brain states. Philos Trans R Soc Lond B Biol Sci. 2016;371.

26. Parpura V, Grubisic V and Verkhratsky A. Ca(2+) sources for the exocytotic release of glutamate from astrocytes. Biochim Biophys Acta. 2011;1813:984–91.

27. Zhang X, Connelly J, Levitan ES, Sun D and Wang JQ. Calcium/Calmodulin-Dependent Protein Kinase II in Cerebrovascular Diseases. Transl Stroke Res. 2021;12:513–529.

28. Bantle CM, Hirst WD, Weihofen A and Shlevkov E. Mitochondrial Dysfunction in Astrocytes: A Role in Parkinson’s Disease? Front Cell Dev Biol. 2020;8:608026.

29. Simpson PB and Russell JT. Role of sarcoplasmic/endoplasmic-reticulum Ca2+-ATPases in mediating Ca2+ waves and local Ca2+-release microdomains in cultured glia. Biochem J. 1997;325(Pt 1):239–47.

30. Ni Fhlathartaigh M, McMahon J, Reynolds R, Connolly D, Higgins E, Counihan T and Fitzgerald U. Calreticulin and other components of endoplasmic reticulum stress in rat and human inflammatory demyelination. Acta Neuropathol Commun. 2013;1:37.

31. Rose J, Brian C, Pappa A, Panayiotidis MI and Franco R. Mitochondrial Metabolism in Astrocytes Regulates Brain Bioenergetics, Neurotransmission and Redox Balance. Front Neurosci. 2020;14:536682.

32. Frigerio F, Karaca M, De Roo M, Mlynarik V, Skytt DM, Carobbio S, Pajecka K, Waagepetersen HS, Gruetter R, Muller D and Maechler P. Deletion of glutamate dehydrogenase 1 (Glud1) in the central nervous system affects glutamate handling without altering synaptic transmission. J Neurochem. 2012;123:342–8.

33. Choudhry H, Aggarwal M and Pan PY. Mini-review: Synaptojanin 1 and its implications in membrane trafficking. Neurosci Lett. 2021;765:136288.

34. Mielnicka A and Michaluk P. Exocytosis in Astrocytes. Biomolecules. 2021;11.

35. Maienschein V, Marxen M, Volknandt W and Zimmermann H. A plethora of presynaptic proteins associated with ATP-storing organelles in cultured astrocytes. Glia. 1999;26:233–44.

36. Willnow TE, Petersen CM and Nykjaer A. VPS10P-domain receptors - regulators of neuronal viability and function. Nat Rev Neurosci. 2008;9:899–909.

37. Longden TA, Dabertrand F, Koide M, Gonzales AL, Tykocki NR, Brayden JE, Hill-Eubanks D and Nelson MT. Capillary K(+)-sensing initiates retrograde hyperpolarization to increase local cerebral blood flow. Nat Neurosci. 2017;20:717–726.

38. Boggild S, Molgaard S, Glerup S and Nyengaard JR. Spatiotemporal patterns of sortilin and SorCS2 localization during organ development. BMC Cell Biol. 2016;17:8–8.

39. Sun W, McConnell E, Pare JF, Xu Q, Chen M, Peng W, Lovatt D, Han X, Smith Y and Nedergaard M. Glutamate-dependent neuroglial calcium signaling differs between young and adult brain. Science. 2013;339:197–200.

40. Berent-Spillson A and Russell JW. Metabotropic glutamate receptor 3 protects neurons from glucose-induced oxidative injury by increasing intracellular glutathione concentration. J Neurochem. 2007;101:342–54.

41. Oguri M, Nagahiro t, Kamiya H, Ohno M, Kato K, Yokoi K, Yoshida T, Watanabe S, Metoki N, Yoshida H, Satoh K, Aoyagi Y, Nozawa Y and Yamada Y. Association of a polymorphism of ROR2 and ischemic stroke in Japanese individuals with chronic kidney disease. Experimental and therapeutic medicine. 2010;1:377–384.

42. Boas DA and Dunn AK. Laser speckle contrast imaging in biomedical optics. J Biomed Opt. 2010;15:011109.

43. Aronoff R and Petersen CC. Layer, column and cell-type specific genetic manipulation in mouse barrel cortex. Front Neurosci. 2008;2:64–71.

44. Postnov DD, Tuchin, V.V. and Sosnovtseva O.. Estimation of vessel diameter and blood flow dynamics from laser speckle images. Biomedical Optics Express. 2016;7:10.

45. Staehr C, Hangaard L, Bouzinova EV, Kim S, Rajanathan R, Boegh Jessen P, Luque N, Xie Z, Lykke-Hartmann K, Sandow SL, Aalkjaer C and Matchkov VV. Smooth muscle Ca(2+) sensitization causes hypercontractility of middle cerebral arteries in mice bearing the familial hemiplegic migraine type 2 associated mutation. J Cereb Blood Flow Metab. 2018;39:1570–1587.

46. Mulorz J, Spin JM, Beck HC, Tha Thi ML, Wagenhauser MU, Rasmussen LM, Lindholt JS, Tsao PSC and Steffensen LB. Hyperlipidemia does not affect development of elastase-induced abdominal aortic aneurysm in mice. Atherosclerosis. 2020;311:73–83.

47. Andersen LC, Palstrom NB, Diederichsen A, Lindholt JS, Rasmussen LM and Beck HC. Determining Plasma Protein Variation Parameters as a Prerequisite for Biomarker Studies-A TMT-Based LC-MSMS Proteome Investigation. Proteomes. 2021;9.

48. Shen J, Pagala VR, Breuer AM, Peng J, Bin M and Wang X. Spectral Library Search Improves Assignment of TMT Labeled MS/MS Spectra. J Proteome Res. 2018;17:3325–3331.

