## Supplementary material for "SorCS2 modulates neurovascular coupling via glutamatergic and calcium signaling in astrocytes": Expanded View Table 1

| | | | | | | | | | | | | **Sample name:** | **SorCS2 knockout mouse #1** | **SorCS2 knockout mouse #2** | **SorCS2 knockout mouse #3** | **SorCS2 knockout mouse #4** | **Wild type mouse #1** | **Wild type mouse #2** | **Wild type mouse #3** | **Wild type mouse #4** | | | |
|---|---|---|---|---|---|---|---|---|---|---|---|---|---|---|---|---|---|---|---|---|---|---|---|
| Accession | Description | Coverage [%] | # Peptides | # PSMs | # Unique Peptides | # AAs | MW [kDa] | calc. pI | Score MSPepSearch: MSPepSearch | Score Sequest HT: Sequest HT | # Peptides (by Search Engine): MSPepSearch | # Peptides (by Search Engine): Sequest HT | Abundance Ratio: (F1, 127N, Sample) / (F1, 126, Control) | Abundance Ratio: (F1, 128N, Sample) / (F1, 126, Control) | Abundance Ratio: (F1, 128C, Sample) / (F1, 126, Control) | Abundance Ratio: (F1, 129N, Sample) / (F1, 126, Control) | Abundance Ratio: (F1, 129C, Sample) / (F1, 126, Control) | Abundance Ratio: (F1, 130N, Sample) / (F1, 126, Control) | Abundance Ratio: (F1, 130C, Sample) / (F1, 126, Control) | Abundance Ratio: (F1, 131N, Sample) / (F1, 126, Control) | **Ratio SorCS2 knockout / wild type** | **Fold change in SorCS2 knockout vs. wild type** | ***t*-test, *P*-value** |
| Q9QYS2 | Metabotropic glutamate receptor 3 OS=Mus musculus OX=1009 | 17 | 14 | 28 | 13 | 879 | 99,1 | 7,75 | 13517 | 16,07 | 10 | 5 | 1,239 | 1,132 | 1,259 | 1,123 | 0,696 | 0,754 | 0,785 | 0,777 | 1,58 | 0,58 | 3,9E-05 |
| Q60932 | Voltage-dependent anion-selective channel protein 1 OS=Mus n | 49 | 12 | 28 | 12 | 296 | 32,3 | 8,43 | 8985 | 39,12 | 7 | 9 | 1,051 | 1,038 | 1,071 | 1,088 | 0,979 | 0,981 | 0,998 | 0,969 | 1,08 | 0,08 | 6,9E-04 |
| P26043 | Radixin OS=Mus musculus OX=10090 GN=Rdx PE=1 SV=3 | 12 | 7 | 11 | 2 | 583 | 68,5 | 6,2 | 4304 | 5,52 | 5 | 2 | 1,186 | 1,286 | 1,175 | 1,233 | 1,049 | 0,985 | 1,07 | 0,997 | 1,19 | 0,19 | 9,7E-04 |
| Q80UG5 | Septin-9 OS=Mus musculus OX=10090 GN=Septin9 PE=1 SV=1 | 20 | 11 | 13 | 11 | 583 | 65,5 | 8,9 | 6526 | 8,92 | 9 | 3 | 1,039 | 1,01 | 1,018 | 1,021 | 0,883 | 0,895 | 0,958 | 0,933 | 1,11 | 0,11 | 1,2E-03 |
| Q8VEL9 | GTP-binding protein REM 2 OS=Mus musculus OX=10090 GN=Re | 5 | 2 | 2 | 2 | 341 | 37,3 | 7,39 | 1076 | | 2 | | 1,039 | 1,073 | 1,076 | 1,081 | 0,923 | 0,96 | 0,996 | 0,982 | 1,11 | 0,11 | 1,5E-03 |
| Q9D8N0 | Elongation factor 1-gamma OS=Mus musculus OX=10090 GN=Ee | 26 | 12 | 16 | 12 | 437 | 50 | 6,74 | 4782 | 23,64 | 7 | 5 | 1,007 | 0,989 | 0,981 | 1,021 | 0,923 | 0,943 | 0,956 | 0,921 | 1,07 | 0,07 | 2,0E-03 |
| Q9DBT9 | Dimethylglycine dehydrogenase, mitochondrial OS=Mus muscul | 3 | 2 | 6 | 2 | 869 | 97,2 | 7,88 | 1747 | 8,26 | 1 | 1 | 1,421 | 1,555 | 1,436 | 1,636 | 2,309 | 2,069 | 1,998 | 1,86 | 0,73 | -0,27 | 2,2E-03 |
| Q8CIP4 | MAP/microtubule affinity-regulating kinase 4 OS=Mus musculus | 2 | 2 | 2 | 2 | 752 | 82,6 | 9,67 | | 4,05 | | 2 | 1,387 | 1,264 | 1,29 | 1,171 | 0,93 | 0,973 | 1,034 | 1,082 | 1,27 | 0,27 | 2,7E-03 |
| Q9R1V6 | Disintegrin and metalloproteinase domain-containing protein 22 | 14 | 10 | 13 | 10 | 904 | 99,7 | 7,83 | 2074 | 26,92 | 3 | 7 | 1,216 | 1,05 | 1,117 | 1 | 0,79 | 0,908 | 0,737 | 0,794 | 1,36 | 0,36 | 2,7E-03 |
| Q9DC50 | Peroxisomal carnitine O-octanoyltransferase OS=Mus musculus | 13 | 6 | 6 | 6 | 612 | 70,2 | 6,73 | 1361 | 4,68 | 2 | 4 | 1,213 | 1,279 | 1,391 | 1,271 | 1,074 | 1,014 | 1,029 | 1,137 | 1,21 | 0,21 | 2,8E-03 |
| P35802 | Neuronal membrane glycoprotein M6-a OS=Mus musculus OX=1 | 13 | 5 | 13 | 5 | 278 | 31,1 | 5,27 | 4805 | 15,44 | 2 | 4 | 1,102 | 1,081 | 1,087 | 1,212 | 0,913 | 0,966 | 0,919 | 0,81 | 1,24 | 0,24 | 2,9E-03 |
| O88621 | Forkhead box protein H1 OS=Mus musculus OX=10090 GN=Foxh | 3 | 1 | 1 | 1 | 401 | 44 | 8,9 | | 0 | | 1 | 0,897 | 0,762 | 0,773 | 0,83 | 1,324 | 1,277 | 1,681 | 1,945 | 0,52 | -0,48 | 3,7E-03 |
| Q9QZ23 | NFU1 iron-sulfur cluster scaffold homolog, mitochondrial OS=M | 13 | 3 | 5 | 3 | 255 | 28,5 | 5,03 | 2523 | | 3 | | 0,936 | 0,937 | 0,928 | 0,949 | 1,072 | 1,009 | 0,992 | 1,075 | 0,90 | -0,10 | 3,8E-03 |
| P48771 | Cytochrome c oxidase subunit 7A2, mitochondrial OS=Mus musc | 28 | 2 | 22 | 2 | 83 | 9,3 | 10,27 | 2969 | 53,34 | 1 | 2 | 1,068 | 1,064 | 1,019 | 1,116 | 0,846 | 0,94 | 0,976 | 0,889 | 1,17 | 0,17 | 4,4E-03 |
| Q922Q1 | Mitochondrial amidoxime reducing component 2 OS=Mus musc | 13 | 5 | 5 | 5 | 338 | 38,2 | 8,68 | 1843 | 5,72 | 3 | 2 | 1,063 | 1,036 | 1,122 | 1,064 | 0,876 | 0,986 | 0,944 | 0,961 | 1,14 | 0,14 | 4,8E-03 |
| P08226 | Apolipoprotein E OS=Mus musculus OX=10090 GN=Apoe PE=1 S | 23 | 7 | 10 | 7 | 311 | 35,8 | 5,68 | 3443 | 7,85 | 4 | 3 | 0,889 | 0,919 | 0,865 | 0,908 | 1,373 | 1,129 | 1,081 | 1,376 | 0,72 | -0,28 | 4,9E-03 |
| Q60598 | Src substrate cortactin OS=Mus musculus OX=10090 GN=Cttn PE | 31 | 17 | 25 | 17 | 546 | 61,2 | 5,4 | 7569 | 25,59 | 10 | 8 | 1,002 | 0,984 | 1,043 | 0,976 | 0,849 | 0,948 | 0,894 | 0,874 | 1,12 | 0,12 | 5,3E-03 |
| Q99L45 | Eukaryotic translation initiation factor 2 subunit 2 OS=Mus musc | 18 | 5 | 7 | 5 | 331 | 38,1 | 5,8 | 1055 | 4,79 | 2 | 3 | 0,885 | 0,92 | 0,943 | 0,891 | 1,065 | 0,967 | 1,053 | 1,002 | 0,89 | -0,11 | 5,5E-03 |
| Q4ACU6 | SH3 and multiple ankyrin repeat domains protein 3 OS=Mus mu | 1 | 1 | 1 | 1 | 1730 | 185,3 | 8,88 | 500 | | 1 | | 1,246 | 1,08 | 1,246 | 1,046 | 0,832 | 0,883 | 0,526 | 0,787 | 1,53 | 0,53 | 6,0E-03 |
| O55125 | Protein NipSnap homolog 1 OS=Mus musculus OX=10090 GN=N | 17 | 4 | 5 | 4 | 284 | 33,3 | 9,44 | 973 | 5,77 | 2 | 3 | 1,132 | 1,034 | 1,026 | 1,037 | 0,764 | 0,943 | 0,899 | 0,868 | 1,22 | 0,22 | 6,1E-03 |
| Q9CZ28 | Vacuolar-sorting protein SNF8 OS=Mus musculus OX=10090 GN | 10 | 2 | 2 | 2 | 258 | 28,9 | 6,65 | 757 | | 2 | | 0,959 | 0,994 | 0,919 | 1,111 | 1,248 | 1,132 | 1,233 | 1,175 | 0,83 | -0,17 | 6,5E-03 |
| Q3UZV7 | UPF0577 protein KIAA1324-like homolog OS=Mus musculus OX= | 2 | 2 | 2 | 2 | 1028 | 113,6 | 6,14 | | 2,46 | | 2 | 0,993 | 1,021 | 0,999 | 0,984 | 0,596 | 0,776 | 0,785 | 0,876 | 1,32 | 0,32 | 6,5E-03 |
| Q9D5V5 | Cullin-5 OS=Mus musculus OX=10090 GN=Cul5 PE=1 SV=3 | 7 | 6 | 8 | 6 | 780 | 90,9 | 7,81 | 3459 | 2,3 | 5 | 2 | 0,974 | 0,954 | 0,944 | 0,944 | 0,846 | 0,928 | 0,864 | 0,854 | 1,09 | 0,09 | 6,7E-03 |
| Q8C8R3-5 | Isoform 5 of Ankyrin-2 OS=Mus musculus OX=10090 GN=Ank2 | 38 | 117 | 235 | 1 | 3844 | 420,2 | 5,12 | 87436 | 253,25 | 77 | 51 | 0,608 | 0,473 | 0,669 | 0,69 | 0,873 | 1,192 | 0,919 | 0,866 | 0,63 | -0,37 | 8,4E-03 |
| Q61941 | NAD(P) transhydrogenase, mitochondrial OS=Mus musculus OX= | 9 | 8 | 10 | 8 | 1086 | 113,8 | 7,64 | 4031 | 2,99 | 5 | 3 | 1,285 | 1,156 | 1,204 | 1,403 | 1,095 | 0,566 | 0,648 | 0,779 | 1,63 | 0,63 | 8,7E-03 |
| Q8VBV7 | COP9 signalosome complex subunit 8 OS=Mus musculus OX=10 | 16 | 3 | 3 | 3 | 209 | 23,2 | 5,2 | 1314 | 2,33 | 2 | 1 | 0,91 | 0,941 | 0,825 | 1,029 | 0,654 | 0,789 | 0,754 | 0,735 | 1,26 | 0,26 | 9,0E-03 |
| Q9JME5 | AP-3 complex subunit beta-2 OS=Mus musculus OX=10090 GN= | 22 | 21 | 37 | 21 | 1082 | 119,1 | 5,63 | 14779 | 14,54 | 17 | 6 | 1,007 | 0,997 | 0,996 | 1,006 | 1,019 | 1,063 | 1,041 | 1,031 | 0,96 | -0,04 | 9,1E-03 |
| P33174 | Chromosome-associated kinesin KIF4 OS=Mus musculus OX=10 | 1 | 2 | 2 | 1 | 1231 | 139,4 | 6,68 | 521 | 2,59 | 1 | 1 | 1,03 | 1,074 | 1,101 | 1,11 | 0,686 | 0,993 | 0,837 | 0,726 | 1,33 | 0,33 | 9,2E-03 |
| Q6PDI5 | Proteasome adapter and scaffold protein ECM29 OS=Mus musc | 3 | 4 | 4 | 4 | 1840 | 203,6 | 7,06 | 958 | 2,7 | 2 | 2 | 1,169 | 0,948 | 1,138 | 1,102 | 0,828 | 0,818 | 0,877 | 0,957 | 1,25 | 0,25 | 9,5E-03 |
| Q76LS9 | Ubiquitin carboxyl-terminal hydrolase MINDY-1 OS=Mus muscul | 4 | 1 | 1 | 1 | 468 | 51,2 | 4,73 | 276 | | 1 | | 0,638 | 0,701 | 0,535 | 0,518 | 0,701 | 0,895 | 0,949 | 0,881 | 0,70 | -0,30 | 9,6E-03 |
| P70441 | Na(+)/H(+) exchange regulatory cofactor NHE-RF1 OS=Mus mus | 36 | 12 | 15 | 12 | 355 | 38,6 | 5,9 | 5570 | 16,1 | 6 | 6 | 1,02 | 1,097 | 1,106 | 1,068 | 0,836 | 0,907 | 1,014 | 0,919 | 1,17 | 0,17 | 9,9E-03 |
| Q9QWL7 | Keratin, type I cytoskeletal 17 OS=Mus musculus OX=10090 GN= | 24 | 13 | 29 | 6 | 433 | 48,1 | 5,06 | 10556 | 19,68 | 7 | 6 | 0,987 | 1,133 | 0,818 | 1,017 | 1,559 | 1,326 | 1,179 | 1,425 | 0,72 | -0,28 | 9,9E-03 |
| Q9DCB8 | Iron-sulfur cluster assembly 2 homolog, mitochondrial OS=Mus | 8 | 1 | 3 | 1 | 154 | 16,7 | 5,6 | 2017 | | 1 | | 1,163 | 1,116 | 1,095 | 1,124 | 0,902 | 1,023 | 0,863 | 1,038 | 1,18 | 0,18 | 1,1E-02 |
| Q8CDG3 | Deubiquitinating protein VCIP135 OS=Mus musculus OX=10090 | 8 | 7 | 9 | 7 | 1220 | 134,4 | 7,17 | 4631 | 2,15 | 7 | 1 | 0,963 | 0,948 | 0,887 | 0,92 | 1,147 | 1,094 | 1,029 | 0,997 | 0,87 | -0,13 | 1,1E-02 |
| P70694 | Estradiol 17 beta-dehydrogenase 5 OS=Mus musculus OX=1009 | 10 | 3 | 5 | 3 | 323 | 37 | 8,28 | 304 | 16,74 | 1 | 2 | 1,06 | 1,456 | 1,66 | 1,403 | 2,066 | 1,752 | 1,853 | 2,048 | 0,72 | -0,28 | 1,1E-02 |
| D3YZP9 | Coiled-coil domain-containing protein 6 OS=Mus musculus OX=1 | 2 | 1 | 1 | 1 | 469 | 52,9 | 7,34 | | 2,26 | | 1 | 1,137 | 1,171 | 1,05 | 1,197 | 0,883 | 0,724 | 1,038 | 0,865 | 1,30 | 0,30 | 1,1E-02 |
| Q8BJF9 | Charged multivesicular body protein 2b OS=Mus musculus OX=1 | 5 | 1 | 1 | 1 | 213 | 23,9 | 8,78 | 851 | | 1 | | 1,328 | 1,2 | 1,356 | 1,413 | 0,839 | 1,028 | 1,139 | 1,111 | 1,29 | 0,29 | 1,1E-02 |
| Q9DB34 | Charged multivesicular body protein 2a OS=Mus musculus OX=1 | 9 | 2 | 2 | 2 | 222 | 25,1 | 5,97 | 699 | 5,02 | 1 | 1 | 0,925 | 0,711 | 0,68 | 0,935 | 1,249 | 1,354 | 0,957 | 1,343 | 0,66 | -0,34 | 1,1E-02 |
| G5E8K5 | Ankyrin-3 OS=Mus musculus OX=10090 GN=Ank3 PE=1 SV=1 | 10 | 19 | 30 | 8 | 1961 | 213,9 | 6,7 | 13352 | 26,36 | 15 | 5 | 0,977 | 1,084 | 1,039 | 0,979 | 1,157 | 1,076 | 1,196 | 1,181 | 0,88 | -0,12 | 1,2E-02 |
| Q8CA95 | cAMP and cAMP-inhibited cGMP 3',5'-cyclic phosphodiesterase | 3 | 2 | 3 | 2 | 790 | 89,4 | 6,6 | 1311 | 2,97 | 2 | 1 | 0,974 | 1,015 | 1,004 | 0,924 | 1,132 | 1,042 | 1,207 | 1,104 | 0,87 | -0,13 | 1,2E-02 |
| Q9CZX0 | Elongator complex protein 3 OS=Mus musculus OX=10090 GN=E | 12 | 5 | 8 | 5 | 547 | 62,3 | 8,9 | 2457 | 1,77 | 4 | 1 | 0,984 | 0,938 | 0,827 | 0,987 | 1,186 | 1,165 | 1,1 | 1,026 | 0,83 | -0,17 | 1,2E-02 |
| Q9D061 | Acyl-CoA-binding domain-containing protein 6 OS=Mus musculu | 9 | 2 | 3 | 2 | 282 | 30,9 | 5,11 | 1519 | | 2 | | 1,237 | 1,017 | 1,021 | 1,154 | 0,771 | 0,934 | 0,928 | 0,739 | 1,31 | 0,31 | 1,2E-02 |
| Q9ERB5 | Solute carrier organic anion transporter family member 1C1 OS= | 1 | 1 | 1 | 1 | 715 | 78,3 | 8,81 | 417 | | 1 | | 0,896 | 0,759 | 0,882 | 0,763 | 1,088 | 1,097 | 0,949 | 1,293 | 0,75 | -0,25 | 1,2E-02 |
| Q62092 | Neuronal vesicle trafficking-associated protein 1 OS=Mus muscu | 4 | 1 | 1 | 1 | 185 | 20,9 | 6,21 | | 2,63 | | 1 | 1,166 | 1,123 | 0,996 | 0,999 | 0,941 | 0,827 | 0,878 | 0,929 | 1,20 | 0,20 | 1,3E-02 |
| Q9JME7 | Trafficking protein particle complex subunit 2-like protein OS=M | 6 | 1 | 1 | 1 | 139 | 16 | 6,77 | | 3,06 | | 1 | 1,067 | 0,974 | 1,054 | 1,054 | 0,562 | 0,787 | 0,792 | 0,921 | 1,35 | 0,35 | 1,3E-02 |
| P70232 | Neural cell adhesion molecule L1-like protein OS=Mus musculus | 6 | 8 | 12 | 8 | 1209 | 135 | 5,57 | 3266 | 20,95 | 4 | 4 | 1,065 | 0,935 | 1,122 | 1,054 | 0,958 | 0,865 | 0,83 | 0,842 | 1,19 | 0,19 | 1,3E-02 |
| Q9D0J8 | Parathymosin OS=Mus musculus OX=10090 GN=Ptms PE=1 SV=3 | 11 | 1 | 3 | 1 | 101 | 11,4 | 4,22 | 1645 | | 1 | | 0,998 | 1,094 | 1,014 | 1,037 | 1,189 | 1,086 | 1,132 | 1,185 | 0,90 | -0,10 | 1,3E-02 |
| Q8BKX1-2 | Isoform 2 of Brain-specific angiogenesis inhibitor 1-associated p | 12 | 6 | 8 | 6 | 522 | 57,6 | 8,88 | 4583 | 2,24 | 5 | 1 | 1,126 | 1,069 | 1,063 | 1,099 | 1,165 | 1,12 | 1,18 | 1,198 | 0,93 | -0,07 | 1,4E-02 |
| Q9CR86 | Calcium-regulated heat stable protein 1 OS=Mus musculus OX=1 | 5 | 1 | 1 | 1 | 148 | 16,1 | 8,21 | | 2,12 | | 1 | 0,964 | 1,141 | 1,174 | 0,965 | 0,594 | 0,931 | 0,736 | 0,761 | 1,40 | 0,40 | 1,4E-02 |
| Q8BGR6 | ADP-ribosylation factor-like protein 15 OS=Mus musculus OX=10 | 15 | 3 | 6 | 3 | 204 | 22,9 | 5,29 | 3271 | 3,19 | 2 | 1 | 0,986 | 0,922 | 1,002 | 0,961 | 0,779 | 0,896 | 0,897 | 0,851 | 1,13 | 0,13 | 1,4E-02 |
| Q9EQJ0 | Two pore calcium channel protein 1 OS=Mus musculus OX=1009 | 2 | 1 | 1 | 1 | 817 | 94,4 | 8,53 | 186 | | 1 | | 0,986 | 1,161 | 0,912 | 1,056 | 1,367 | 1,223 | 1,168 | 1,39 | 0,80 | -0,20 | 1,4E-02 |
| D3YXK2 | Scaffold attachment factor B1 OS=Mus musculus OX=10090 GN= | 2 | 2 | 4 | 2 | 937 | 105 | 5,35 | 2143 | | 2 | | 0,921 | 0,884 | 0,952 | 0,954 | 1,012 | 0,966 | 1,065 | 1,023 | 0,91 | -0,09 | 1,5E-02 |
| P12849 | cAMP-dependent protein kinase type I-beta regulatory subunit | 8 | 3 | 3 | 1 | 381 | 43,2 | 5,96 | 831 | 5,76 | 1 | 2 | 0,876 | 0,906 | 0,935 | 0,984 | 0,884 | 0,746 | 0,76 | 0,786 | 1,17 | 0,17 | 1,5E-02 |
| Q6I6G8 | E3 ubiquitin-protein ligase HECW2 OS=Mus musculus OX=10090 | 1 | 2 | 3 | 2 | 1578 | 176,1 | 5,41 | 1934 | | 2 | | 0,973 | 0,948 | 1,004 | 0,98 | 0,807 | 0,91 | 0,876 | 0,925 | 1,11 | 0,11 | 1,5E-02 |
| P62137 | Serine/threonine-protein phosphatase PP1-alpha catalytic subu | 34 | 9 | 14 | 3 | 330 | 37,5 | 6,33 | 5396 | 15,31 | 7 | 3 | 1,208 | 1,143 | 1,168 | 1,181 | 0,971 | 1,022 | 1,134 | 1,071 | 1,12 | 0,12 | 1,5E-02 |

| | | | | | | | | | | | | | | | | | | | | | | | |
|---|---|---|---|---|---|---|---|---|---|---|---|---|---|---|---|---|---|---|---|---|---|---|---|
| Q9CPW3 | 39S ribosomal protein L54, mitochondrial OS=Mus musculus OX | 7 | 1 | 1 | 1 | 135 | 15,4 | 10,04 | | 3,39 | | 1 | 0,968 | 0,901 | 0,906 | 1,145 | 0,742 | 0,826 | 0,784 | 0,651 | 1,31 | 0,31 | 1,5E-02 |
| Q9JIA1 | Leucine-rich glioma-inactivated protein 1 OS=Mus musculus OX= | 10 | 6 | 6 | 6 | 557 | 63,6 | 8,02 | 1688 | 5,05 | 3 | 3 | 1,03 | 1,098 | 0,973 | 1,044 | 0,852 | 0,916 | 0,964 | 0,94 | 1,13 | 0,13 | 1,5E-02 |
| P11798 | Calcium/calmodulin-dependent protein kinase type II subunit al | 23 | 11 | 23 | 7 | 478 | 54,1 | 7,08 | 11259 | 11,3 | 7 | 4 | 0,985 | 1,004 | 0,946 | 1,009 | 1,058 | 1,021 | 1,035 | 1,061 | 0,94 | -0,06 | 1,5E-02 |
| Q99NB8 | Ubiquilin-4 OS=Mus musculus OX=10090 GN=Ubqln4 PE=1 SV=1 | 4 | 1 | 1 | 1 | 596 | 63,5 | 5,03 | 352 | | 1 | | 0,758 | 0,649 | 0,694 | 0,692 | 0,929 | 0,801 | 0,788 | 0,982 | 0,80 | -0,20 | 1,5E-02 |
| P70208 | Plexin-A3 OS=Mus musculus OX=10090 GN=Plxna3 PE=1 SV=2 | 6 | 10 | 13 | 4 | 1872 | 207,8 | 7,09 | 4431 | 14,61 | 6 | 4 | 0,903 | 0,95 | 1,035 | 0,972 | 1,085 | 1,073 | 1,031 | 1,134 | 0,89 | -0,11 | 1,6E-02 |
| P25785 | Metalloproteinase inhibitor 2 OS=Mus musculus OX=10090 GN= | 3 | 1 | 1 | 1 | 220 | 24,3 | 7,49 | 338 | | 1 | | 1,091 | 0,983 | 1,092 | 1,081 | 0,849 | 1,001 | 0,934 | 0,922 | 1,15 | 0,15 | 1,6E-02 |
| O35680 | 28S ribosomal protein S12, mitochondrial OS=Mus musculus OX | 8 | 1 | 2 | 1 | 139 | 15,4 | 10,71 | | 2,15 | | 1 | 0,848 | 0,721 | 0,765 | 0,666 | 1,054 | 0,965 | 0,903 | 0,85 | 0,80 | -0,20 | 1,6E-02 |
| Q9Z0P5 | Twinfilin-2 OS=Mus musculus OX=10090 GN=Twf2 PE=1 SV=1 | 17 | 4 | 6 | 4 | 349 | 39,4 | 6,8 | 3223 | 2,31 | 3 | 1 | 0,861 | 0,941 | 0,921 | 0,884 | 0,932 | 1,039 | 1,062 | 1,019 | 0,89 | -0,11 | 1,6E-02 |
| Q60841 | Reelin OS=Mus musculus OX=10090 GN=Reln PE=1 SV=3 | 0 | 1 | 1 | 1 | 3461 | 387,2 | 5,73 | 259 | | 1 | | 1,357 | 1,161 | 1,261 | 1,295 | 1,104 | 1,049 | 1,158 | 1,14 | 1,14 | 0,14 | 1,7E-02 |
| Q8CI51 | PDZ and LIM domain protein 5 OS=Mus musculus OX=10090 GN | 5 | 2 | 3 | 2 | 591 | 63,3 | 8,25 | 730 | 2,13 | 1 | 1 | 1,139 | 1,255 | 1,197 | 1,153 | 1,06 | 1,019 | 1,101 | 1,117 | 1,10 | 0,10 | 1,7E-02 |
| P54728 | UV excision repair protein RAD23 homolog B OS=Mus musculus | 11 | 6 | 8 | 6 | 416 | 43,5 | 4,83 | 2134 | 12,96 | 3 | 4 | 0,942 | 0,914 | 0,885 | 0,955 | 0,834 | 0,867 | 0,877 | 0,789 | 1,10 | 0,10 | 1,7E-02 |
| Q3UHF7 | Transcription factor HIVEP2 OS=Mus musculus OX=10090 GN=H | 0 | 1 | 2 | 1 | 2430 | 266,5 | 6,96 | 652 | | 1 | | 1,04 | 0,831 | 0,91 | 0,852 | 0,6 | 0,79 | 0,742 | 0,692 | 1,29 | 0,29 | 1,7E-02 |
| Q6PFD5 | Disks large-associated protein 3 OS=Mus musculus OX=10090 GI | 4 | 3 | 3 | 3 | 977 | 105,8 | 8,76 | 1611 | | 3 | | 0,891 | 1,064 | 0,956 | 1,091 | 1,191 | 1,101 | 1,248 | 1,18 | 0,85 | -0,15 | 1,8E-02 |
| Q62418 | Drebrin-like protein OS=Mus musculus OX=10090 GN=Dbnl PE=1 | 23 | 8 | 12 | 8 | 436 | 48,7 | 4,92 | 3810 | 17,17 | 4 | 4 | 1,026 | 0,923 | 0,993 | 0,961 | 0,884 | 0,922 | 0,908 | 0,874 | 1,09 | 0,09 | 1,9E-02 |
| Q99LC9 | Peroxisome assembly factor 2 OS=Mus musculus OX=10090 GN= | 1 | 1 | 1 | 1 | 981 | 104,5 | 7,34 | 317 | | 1 | | 1,082 | 1,033 | 1,025 | 1,033 | 1,434 | 1,12 | 1,197 | 1,316 | 0,82 | -0,18 | 1,9E-02 |
| Q9QZQ8 | Core histone macro-H2A.1 OS=Mus musculus OX=10090 GN=H2 | 19 | 5 | 6 | 3 | 372 | 39,7 | 9,8 | 3878 | | 5 | | 0,855 | 0,898 | 0,976 | 1,074 | 1,172 | 1,036 | 1,19 | 1,164 | 0,83 | -0,17 | 1,9E-02 |
| Q5U405 | Transmembrane protease serine 13 OS=Mus musculus OX=1009 | 1 | 1 | 20 | 1 | 543 | 59,8 | 9,01 | 1387 | 46,12 | 1 | 1 | 1,168 | 0,955 | 0,883 | 0,962 | 1,581 | 1,278 | 1,368 | 1,144 | 0,74 | -0,26 | 1,9E-02 |
| Q3UHD6 | Sorting nexin-27 OS=Mus musculus OX=10090 GN=Snx27 PE=1 S | 15 | 8 | 9 | 8 | 539 | 61 | 6,4 | 2908 | 13,05 | 5 | 3 | 1,016 | 1,022 | 0,96 | 1,062 | 0,901 | 0,934 | 0,962 | 0,949 | 1,08 | 0,08 | 1,9E-02 |
| O55100 | Synaptogyrin-1 OS=Mus musculus OX=10090 GN=Syngr1 PE=1 S | 5 | 1 | 2 | 1 | 234 | 25,6 | 4,65 | 939 | | 1 | | 1,08 | 1,095 | 1,066 | 0,938 | 0,782 | 0,975 | 0,874 | 0,733 | 1,24 | 0,24 | 1,9E-02 |
| P61164 | Alpha-centractin OS=Mus musculus OX=10090 GN=Actr1a PE=1 | 38 | 11 | 18 | 6 | 376 | 42,6 | 6,64 | 10246 | 2,17 | 11 | 1 | 1,011 | 0,965 | 0,941 | 1,018 | 0,748 | 0,961 | 0,803 | 0,79 | 1,19 | 0,19 | 2,0E-02 |
| Q6PDY2 | 2-aminoethanethiol dioxygenase OS=Mus musculus OX=10090 ( | 16 | 4 | 5 | 4 | 256 | 28,4 | 5,97 | 2007 | 2,48 | 3 | 1 | 1,002 | 1,136 | 0,97 | 1,054 | 0,889 | 0,876 | 0,929 | 0,957 | 1,14 | 0,14 | 2,0E-02 |
| Q9Z2Z6 | Mitochondrial carnitine/acylcarnitine carrier protein OS=Mus m | 7 | 2 | 3 | 2 | 301 | 33 | 9,11 | 1311 | 2,6 | 1 | 1 | 0,981 | 0,876 | 1,054 | 1,04 | 0,787 | 0,86 | 0,796 | 0,901 | 1,18 | 0,18 | 2,1E-02 |
| P62482 | Voltage-gated potassium channel subunit beta-2 OS=Mus muscu | 4 | 2 | 2 | 2 | 367 | 41 | 9 | 1281 | | 2 | | 1,27 | 1,187 | 1,302 | 1,17 | 1,02 | 1,024 | 1,188 | 1,071 | 1,15 | 0,15 | 2,1E-02 |
| Q61137 | Astrotactin-1 OS=Mus musculus OX=10090 GN=Astn1 PE=1 SV=4 | 4 | 4 | 4 | 4 | 1302 | 144,8 | 5,21 | 1424 | 3,35 | 2 | 2 | 0,81 | 0,838 | 0,969 | 0,809 | 0,989 | 0,958 | 0,98 | 0,977 | 0,88 | -0,12 | 2,1E-02 |
| Q3UH53 | Protein sidekick-1 OS=Mus musculus OX=10090 GN=Sdk1 PE=1 S | 1 | 2 | 4 | 1 | 2193 | 240,2 | 6,77 | 1736 | | 2 | | 0,822 | 0,826 | 0,847 | 0,91 | 1,346 | 0,897 | 1,366 | 1,161 | 0,71 | -0,29 | 2,2E-02 |
| Q8BLN5 | Lanosterol synthase OS=Mus musculus OX=10090 GN=Lss PE=1 | 6 | 4 | 6 | 4 | 733 | 83,1 | 6,4 | 2391 | 0 | 3 | 1 | 1,184 | 1,093 | 1,225 | 1,069 | 0,969 | 1,065 | 1,016 | 1,004 | 1,13 | 0,13 | 2,2E-02 |
| Q9QXF8 | Glycine N-methyltransferase OS=Mus musculus OX=10090 GN=( | 13 | 2 | 7 | 2 | 293 | 32,7 | 7,44 | 4354 | | 2 | | 0,985 | 0,999 | 1,089 | 1,088 | 1,258 | 1,203 | 1,097 | 1,359 | 0,85 | -0,15 | 2,2E-02 |
| Q8VC28 | Aldo-keto reductase family 1 member C13 OS=Mus musculus OX | 5 | 2 | 3 | 2 | 323 | 37 | 7,08 | 2092 | | 2 | | 2,339 | 2,549 | 2,109 | 2,378 | 3,399 | 2,996 | 2,53 | 2,901 | 0,79 | -0,21 | 2,2E-02 |
| Q62283 | Tetraspanin-7 OS=Mus musculus OX=10090 GN=Tspan7 PE=1 SV | 6 | 1 | 2 | 1 | 249 | 27,5 | 7,2 | | 5,2 | | 1 | 0,74 | 0,871 | 0,79 | 0,768 | 0,869 | 1,047 | 1,287 | 1,336 | 0,70 | -0,30 | 2,3E-02 |
| O35083 | 1-acyl-sn-glycerol-3-phosphate acyltransferase alpha OS=Mus m | 5 | 1 | 1 | 1 | 285 | 31,7 | 9,14 | | 0 | | 1 | 0,92 | 0,881 | 0,993 | 0,776 | 1,013 | 1,023 | 1,226 | 1,239 | 0,79 | -0,21 | 2,3E-02 |
| Q9CQQ0 | Small integral membrane protein 8 OS=Mus musculus OX=1009( | 9 | 1 | 1 | 1 | 97 | 11,1 | 9,31 | | 0 | | 1 | 1,297 | 1,336 | 1,467 | 1,354 | 1,607 | 1,403 | 1,568 | 1,607 | 0,88 | -0,12 | 2,4E-02 |
| P27546 | Microtubule-associated protein 4 OS=Mus musculus OX=10090 | 30 | 27 | 55 | 19 | 1125 | 117,4 | 4,98 | 22242 | 42,21 | 17 | 13 | 1,043 | 1,023 | 1,037 | 1,099 | 0,882 | 0,98 | 1,004 | 0,964 | 1,10 | 0,10 | 2,5E-02 |
| P51863 | V-type proton ATPase subunit d 1 OS=Mus musculus OX=10090 | 20 | 9 | 13 | 9 | 351 | 40,3 | 5 | 5573 | 11,03 | 5 | 4 | 0,969 | 0,971 | 0,987 | 0,963 | 0,878 | 0,937 | 0,954 | 0,919 | 1,05 | 0,05 | 2,6E-02 |
| Q9CZD3 | Glycine--tRNA ligase OS=Mus musculus OX=10090 GN=Gars PE= | 22 | 18 | 24 | 18 | 729 | 81,8 | 6,65 | 10263 | 10,75 | 13 | 6 | 0,984 | 0,985 | 1,056 | 1,053 | 0,862 | 0,992 | 0,896 | 0,924 | 1,11 | 0,11 | 2,6E-02 |
| P51807 | Dynein light chain Tctex-type 1 OS=Mus musculus OX=10090 GN | 14 | 1 | 2 | 1 | 113 | 12,5 | 5,08 | 896 | 3,12 | 1 | 1 | 1,174 | 1,144 | 1,083 | 1,241 | 0,703 | 0,876 | 1,087 | 0,954 | 1,28 | 0,28 | 2,6E-02 |
| P26645 | Myristoylated alanine-rich C-kinase substrate OS=Mus musculus | 64 | 15 | 48 | 15 | 309 | 29,6 | 4,34 | 12431 | 124,37 | 8 | 9 | 1,084 | 1,118 | 0,97 | 1,116 | 0,783 | 1,014 | 0,918 | 0,873 | 1,20 | 0,20 | 2,6E-02 |
| Q571F8 | Glutaminase liver isoform, mitochondrial OS=Mus musculus OX= | 4 | 2 | 2 | 1 | 602 | 66,3 | 7,66 | 1057 | | 2 | | 0,81 | 1,848 | 1,439 | 1,63 | 2,163 | 2,325 | 2,033 | 1,986 | 0,67 | -0,33 | 2,6E-02 |
| Q8R081 | Heterogeneous nuclear ribonucleoprotein L OS=Mus musculus ( | 22 | 10 | 19 | 10 | 586 | 63,9 | 8,1 | 8189 | 17,22 | 8 | 5 | 0,97 | 0,872 | 0,963 | 1,093 | 1,263 | 1,035 | 1,196 | 1,179 | 0,83 | -0,17 | 2,6E-02 |
| O88809 | Neuronal migration protein doublecortin OS=Mus musculus OX= | 37 | 13 | 26 | 11 | 366 | 40,6 | 9,28 | 6779 | 41,38 | 4 | 10 | 1,009 | 0,973 | 1,058 | 0,993 | 0,768 | 0,933 | 0,879 | 0,941 | 1,15 | 0,15 | 2,7E-02 |
| Q80Y24 | Prickle-like protein 2 OS=Mus musculus OX=10090 GN=Prickle2 | 4 | 4 | 4 | 4 | 845 | 95,7 | 7,27 | 1665 | 3,07 | 3 | 1 | 1,049 | 1,013 | 1,11 | 1,07 | 0,957 | 1,026 | 0,995 | 0,967 | 1,08 | 0,08 | 2,7E-02 |
| P58710 | L-gulonolactone oxidase OS=Mus musculus OX=10090 GN=Gulo | 7 | 2 | 2 | 2 | 440 | 50,4 | 8,24 | 582 | 4,59 | 1 | 1 | 2,683 | 2,892 | 2,883 | 3,496 | 4,12 | 4,035 | 3,179 | 3,82 | 0,79 | -0,21 | 2,7E-02 |
| Q920I9 | WD repeat-containing protein 7 OS=Mus musculus OX=10090 G | 11 | 15 | 18 | 15 | 1489 | 163,3 | 6,9 | 5514 | 16,2 | 9 | 6 | 0,985 | 1,035 | 0,99 | 0,98 | 0,945 | 0,886 | 0,974 | 0,926 | 1,07 | 0,07 | 2,8E-02 |
| Q8K411 | Presequence protease, mitochondrial OS=Mus musculus OX=10( | 8 | 7 | 10 | 7 | 1036 | 117,3 | 7,2 | 6302 | | 7 | | 1,012 | 1,039 | 0,983 | 1,085 | 0,889 | 0,989 | 0,96 | 0,928 | 1,09 | 0,09 | 2,8E-02 |
| Q9CQF9 | Prenylcysteine oxidase OS=Mus musculus OX=10090 GN=Pcyox1 | 9 | 3 | 3 | 3 | 505 | 56,5 | 6,92 | 1531 | | 3 | | 0,967 | 1,105 | 0,957 | 1,012 | 1,248 | 1,138 | 1,236 | 1,06 | 0,86 | -0,14 | 2,8E-02 |
| Q61502 | Transcription factor E2F5 OS=Mus musculus OX=10090 GN=E2f5 | 3 | 1 | 1 | 1 | 335 | 36,5 | 5,1 | 416 | | 1 | | 0,988 | 1,105 | 0,845 | 1,024 | 0,736 | 0,828 | 0,744 | 0,895 | 1,24 | 0,24 | 2,8E-02 |
| Q9QZB7 | Actin-related protein 10 OS=Mus musculus OX=10090 GN=Actr1 | 8 | 3 | 6 | 3 | 417 | 46,2 | 7,61 | 1793 | 7,64 | 2 | 2 | 0,921 | 0,887 | 0,838 | 0,81 | 0,979 | 0,948 | 1,01 | 0,904 | 0,90 | -0,10 | 2,8E-02 |
| Q99LS3 | Phosphoserine phosphatase OS=Mus musculus OX=10090 GN=P | 10 | 2 | 2 | 2 | 225 | 25,1 | 6,14 | 1308 | | 2 | | 0,89 | 0,786 | 0,986 | 0,866 | 0,555 | 0,814 | 0,681 | 0,708 | 1,28 | 0,28 | 2,9E-02 |
| Q9EQH3 | Vacuolar protein sorting-associated protein 35 OS=Mus musculu | 16 | 12 | 22 | 12 | 796 | 91,7 | 5,44 | 8174 | 28,37 | 7 | 7 | 0,924 | 0,959 | 1,003 | 0,924 | 1,1 | 1,043 | 0,983 | 1,032 | 0,92 | -0,08 | 2,9E-02 |
| Q149F3 | Eukaryotic peptide chain release factor GTP-binding subunit ERF | 12 | 8 | 10 | 1 | 632 | 69,1 | 5,2 | 4331 | 5,26 | 6 | 2 | 1,079 | 0,999 | 1,076 | 1,089 | 0,743 | 1,035 | 0,887 | 0,849 | 1,21 | 0,21 | 2,9E-02 |
| P02535 | Keratin, type I cytoskeletal 10 OS=Mus musculus OX=10090 GN= | 18 | 11 | 41 | 7 | 570 | 57,7 | 5,11 | 13610 | 48,97 | 7 | 7 | 0,805 | 1,158 | 0,585 | 0,709 | 1,217 | 1,457 | 1,163 | 1,09 | 0,66 | -0,34 | 2,9E-02 |
| O35435 | Dihydroorotate dehydrogenase (quinone), mitochondrial OS=M | 14 | 3 | 4 | 3 | 395 | 42,7 | 9,55 | 1929 | | 3 | | 0,865 | 0,94 | 0,731 | 0,801 | 1,11 | 0,924 | 0,974 | 1,005 | 0,83 | -0,17 | 2,9E-02 |
| Q9CQJ6 | Density-regulated protein OS=Mus musculus OX=10090 GN=Der | 13 | 2 | 3 | 2 | 198 | 22,2 | 5,3 | 502 | 7,62 | 1 | 2 | 0,994 | 0,854 | 0,868 | 1,104 | 0,635 | 0,794 | 0,836 | 0,564 | 1,35 | 0,35 | 3,0E-02 |
| Q64433 | 10 kDa heat shock protein, mitochondrial OS=Mus musculus OX | 64 | 10 | 24 | 10 | 102 | 11 | 8,35 | 10669 | 15,12 | 6 | 5 | 1,017 | 1,014 | 0,954 | 1,069 | 0,719 | 0,989 | 0,856 | 0,77 | 1,22 | 0,22 | 3,0E-02 |
| Q61425 | Hydroxyacyl-coenzyme A dehydrogenase, mitochondrial OS=Mu | 12 | 3 | 5 | 3 | 314 | 34,4 | 8,65 | 1448 | 10,31 | 1 | 2 | 0,927 | 0,914 | 1,061 | 1,053 | 0,495 | 0,743 | 0,804 | 0,876 | 1,36 | 0,36 | 3,0E-02 |
| P97355 | Spermine synthase OS=Mus musculus OX=10090 GN=Sms PE=1 | 8 | 3 | 5 | 3 | 366 | 41,3 | 5,06 | 790 | 7,12 | 1 | 2 | 1,088 | 0,969 | 0,952 | 1,015 | 0,72 | 0,889 | 0,926 | 0,872 | 1,18 | 0,18 | 3,0E-02 |
| Q9WU78 | Programmed cell death 6-interacting protein OS=Mus musculus | 12 | 9 | 9 | 9 | 869 | 96 | 6,52 | 3906 | 4,54 | 6 | 3 | 1,221 | 1,083 | 1,215 | 1,243 | 0,945 | 1,105 | 1,101 | 1,014 | 1,14 | 0,14 | 3,0E-02 |
| Q8R1Q8 | Cytoplasmic dynein 1 light intermediate chain 1 OS=Mus muscul | 33 | 15 | 23 | 15 | 523 | 56,6 | 6,42 | 10139 | 17,29 | 10 | 6 | 1,007 | 1,025 | 1,016 | 1,037 | 0,917 | 0,999 | 0,943 | 0,992 | 1,06 | 0,06 | 3,0E-02 |
| Q8BHG1 | Nardilysin OS=Mus musculus OX=10090 GN=Nrdc PE=1 SV=1 | 3 | 3 | 3 | 3 | 1161 | 132,8 | 4,87 | 502 | 6,28 | 1 | 2 | 1,074 | 1,095 | 1,177 | 1,078 | 0,956 | 1,062 | 0,943 | 1,035 | 1,11 | 0,11 | 3,0E-02 |
| P51175 | Protoporphyrinogen oxidase OS=Mus musculus OX=10090 GN=F | 2 | 1 | 1 | 1 | 477 | 50,8 | 8,84 | 541 | | 1 | | 0,903 | 1,032 | 0,957 | 0,925 | 1,017 | 1,347 | 1,104 | 1,211 | 0,82 | -0,18 | 3,1E-02 |
| P00158 | Cytochrome b OS=Mus musculus OX=10090 GN=Mt-Cyb PE=1 SV | 2 | 1 | 1 | 1 | 381 | 43,2 | 7,97 | | 2,15 | | 1 | 0,952 | 0,875 | 1,158 | 0,943 | 0,815 | 0,867 | 0,737 | 0,715 | 1,25 | 0,25 | 3,1E-02 |
| Q8BL97 | Serine/arginine-rich splicing factor 7 OS=Mus musculus OX=1009 | 6 | 2 | 3 | 2 | 267 | 30,8 | 11,9 | 1690 | | 2 | | 1,011 | 0,996 | 1,118 | 1,116 | 1,31 | 1,133 | 1,39 | 1,174 | 0,85 | -0,15 | 3,1E-02 |
| P0C192 | Leucine-rich repeat-containing protein 4B OS=Mus musculus OX | 5 | 4 | 5 | 4 | 709 | 76,1 | 7,24 | 2782 | | 4 | | 0,88 | 1,075 | 0,818 | 0,927 | 1,019 | 1,109 | 1,112 | 1,189 | 0,84 | -0,16 | 3,1E-02 |
| O35874 | Neutral amino acid transporter A OS=Mus musculus OX=10090 ( | 12 | 4 | 8 | 4 | 532 | 56 | 5,87 | 1402 | 17,03 | 1 | 3 | 1,092 | 1,196 | 1,109 | 1,097 | 0,788 | 0,855 | 1,095 | 0,957 | 1,22 | 0,22 | 3,1E-02 |
| A2A699 | Protein FAM171A2 OS=Mus musculus OX=10090 GN=Fam171a2 | 10 | 7 | 8 | 7 | 822 | 87,4 | 8,05 | 3532 | 2,85 | 6 | 1 | 0,822 | 0,891 | 0,806 | 0,814 | 1,175 | 0,836 | 1,123 | 1,062 | 0,79 | -0,21 | 3,1E-02 |
| Q5SSL4 | Active breakpoint cluster region-related protein OS=Mus muscu | 8 | 8 | 9 | 7 | 859 | 97,6 | 6,58 | 4133 | 8,2 | 6 | 2 | 1,06 | 1,062 | 1,117 | 1,116 | 0,801 | 0,992 | 0,979 | 1,011 | 1,15 | 0,15 | 3,2E-02 |

| | | | | | | | | | | | | | | | | | | | | | | | |
|---|---|---|---|---|---|---|---|---|---|---|---|---|---|---|---|---|---|---|---|---|---|---|---|
| P62821 | Ras-related protein Rab-1A OS=Mus musculus OX=10090 GN=Ra | 47 | 9 | 22 | 4 | 205 | 22,7 | 6,21 | 10703 | 18,16 | 5 | 5 | 1,168 | 1,012 | 1,119 | 1,097 | 0,761 | 1,017 | 0,902 | 0,984 | 1,20 | 0,20 | 3,2E-02 |
| Q9R0S3 | Matrix metalloproteinase-17 OS=Mus musculus OX=10090 GN=I | 2 | 1 | 1 | 1 | 578 | 64,3 | 6,38 | 270 | | 1 | | 0,903 | 0,899 | 0,822 | 0,933 | 0,931 | 1,063 | 1,066 | 0,963 | 0,88 | -0,12 | 3,2E-02 |
| Q8VBT0 | Thioredoxin-related transmembrane protein 1 OS=Mus musculu | 4 | 1 | 1 | 1 | 278 | 31,4 | 5,29 | | 2,43 | | 1 | 1,193 | 1,237 | 1,451 | 1,165 | 0,714 | 1,18 | 0,964 | 0,904 | 1,34 | 0,34 | 3,2E-02 |
| Q3TDK6 | Protein rogdi homolog OS=Mus musculus OX=10090 GN=Rogdi I | 6 | 2 | 3 | 2 | 287 | 32,1 | 8,18 | 1916 | | 2 | | 1,049 | 1,032 | 0,913 | 1,134 | 0,953 | 0,865 | 0,836 | 0,9 | 1,16 | 0,16 | 3,3E-02 |
| Q80UK0 | SEC14 domain and spectrin repeat-containing protein 1 OS=Mus | 6 | 4 | 4 | 4 | 696 | 79,3 | 5,1 | 652 | 7,66 | 1 | 3 | 0,955 | 0,961 | 1,013 | 1,005 | 1,089 | 1,12 | 1,038 | 1,008 | 0,92 | -0,08 | 3,3E-02 |
| Q99LD9 | Translation initiation factor eIF-2B subunit beta OS=Mus muscul | 4 | 1 | 1 | 1 | 351 | 38,9 | 6,24 | 814 | | 1 | | 0,976 | 0,981 | 1,043 | 1,177 | 1,559 | 1,119 | 1,261 | 1,382 | 0,79 | -0,21 | 3,4E-02 |
| O08529 | Calpain-2 catalytic subunit OS=Mus musculus OX=10090 GN=Ca | 7 | 5 | 10 | 5 | 700 | 79,8 | 4,96 | 3209 | 11,28 | 4 | 2 | 0,994 | 0,943 | 1,015 | 0,99 | 0,746 | 0,937 | 0,916 | 0,841 | 1,15 | 0,15 | 3,4E-02 |
| Q6IRU2 | Tropomyosin alpha-4 chain OS=Mus musculus OX=10090 GN=Tp | 21 | 6 | 8 | 4 | 248 | 28,5 | 4,68 | 1258 | 14,53 | 2 | 4 | 0,995 | 1,008 | 0,999 | 1,067 | 0,985 | 0,833 | 0,949 | 0,896 | 1,11 | 0,11 | 3,4E-02 |
| P10637-3 | Isoform Tau-B of Microtubule-associated protein tau OS=Mus m | 68 | 28 | 107 | 1 | 364 | 38,2 | 9,33 | 44074 | 69,43 | 14 | 20 | 1,051 | 1,023 | 0,938 | 0,934 | 0,789 | 0,973 | 0,8 | 0,729 | 1,20 | 0,20 | 3,5E-02 |
| Q9QZ06 | Toll-interacting protein OS=Mus musculus OX=10090 GN=Tollip | 11 | 3 | 6 | 3 | 274 | 30,3 | 5,17 | 1322 | 7,4 | 1 | 2 | 1,104 | 1,117 | 1,035 | 1,178 | 1,011 | 1,022 | 0,774 | 0,93 | 1,19 | 0,19 | 3,5E-02 |
| P61924 | Coatomer subunit zeta-1 OS=Mus musculus OX=10090 GN=Cop | 8 | 1 | 1 | 1 | 177 | 20,2 | 4,81 | 632 | | 1 | | 0,933 | 0,932 | 0,922 | 1,05 | 0,762 | 0,749 | 0,931 | 0,836 | 1,17 | 0,17 | 3,5E-02 |
| Q7TMK9-2 | Isoform 2 of Heterogeneous nuclear ribonucleoprotein Q OS=M | 17 | 10 | 13 | 10 | 562 | 62,6 | 7,56 | 6035 | 7,3 | 8 | 2 | 0,942 | 0,919 | 1,013 | 1,002 | 1,033 | 1,014 | 1,036 | 1,061 | 0,94 | -0,06 | 3,5E-02 |
| Q8BW75 | Amine oxidase [flavin-containing] B OS=Mus musculus OX=1009 | 4 | 3 | 3 | 3 | 520 | 58,5 | 8,29 | 735 | 4,82 | 1 | 2 | 1,056 | 1,175 | 0,986 | 1,224 | 0,981 | 0,887 | 0,939 | 0,992 | 1,17 | 0,17 | 3,5E-02 |
| Q9CY57 | Chromatin target of PRMT1 protein OS=Mus musculus OX=1009 | 12 | 2 | 2 | 2 | 249 | 26,6 | 12,23 | 640 | 2,19 | 1 | 1 | 0,621 | 0,743 | 0,941 | 0,85 | 1,474 | 0,905 | 1,039 | 1,338 | 0,66 | -0,34 | 3,6E-02 |
| Q9R0Y5 | Adenylate kinase isoenzyme 1 OS=Mus musculus OX=10090 GN | 23 | 5 | 8 | 5 | 194 | 21,5 | 5,81 | 2298 | 8,02 | 2 | 3 | 0,855 | 0,922 | 0,927 | 0,895 | 1,122 | 0,987 | 1,01 | 0,937 | 0,89 | -0,11 | 3,6E-02 |
| Q8BWY9 | Protein CIP2A OS=Mus musculus OX=10090 GN=Cip2a PE=1 SV= | 1 | 1 | 1 | 1 | 907 | 102 | 6,28 | 402 | | 1 | | 1,15 | 0,876 | 0,978 | 0,981 | 0,741 | 0,875 | 0,859 | 0,82 | 1,21 | 0,21 | 3,6E-02 |
| P26638 | Serine--tRNA ligase, cytoplasmic OS=Mus musculus OX=10090 G | 25 | 11 | 15 | 11 | 512 | 58,4 | 6,3 | 5861 | 19,84 | 8 | 4 | 0,985 | 0,959 | 1,059 | 1,013 | 0,916 | 0,964 | 0,954 | 0,903 | 1,07 | 0,07 | 3,6E-02 |
| Q99LP6 | GrpE protein homolog 1, mitochondrial OS=Mus musculus OX=1 | 32 | 6 | 8 | 6 | 217 | 24,3 | 8,38 | 3919 | 5,81 | 5 | 1 | 1,151 | 1,011 | 1,126 | 1,021 | 1,01 | 0,989 | 0,948 | 0,908 | 1,12 | 0,12 | 3,6E-02 |
| Q9CWM4 | Prefoldin subunit 1 OS=Mus musculus OX=10090 GN=Pfdn1 PE= | 16 | 3 | 5 | 3 | 122 | 14,2 | 8,32 | 1489 | 7,63 | 1 | 2 | 1,128 | 1,054 | 1,088 | 1,142 | 0,65 | 1,022 | 0,984 | 0,821 | 1,27 | 0,27 | 3,7E-02 |
| Q9ERL9 | Guanylate cyclase soluble subunit alpha-1 OS=Mus musculus OX | 1 | 1 | 3 | 1 | 691 | 77,5 | 7,17 | 1668 | | 1 | | 1,166 | 0,944 | 1,19 | 1,201 | 0,902 | 1,012 | 0,971 | 0,801 | 1,22 | 0,22 | 3,7E-02 |
| P62484 | Abl interactor 2 OS=Mus musculus OX=10090 GN=Abi2 PE=1 SV | 13 | 6 | 9 | 3 | 446 | 49,4 | 6,01 | 2920 | 13,12 | 5 | 2 | 1,131 | 1 | 0,978 | 1,572 | 0,637 | 0,843 | 0,82 | 0,821 | 1,50 | 0,50 | 3,7E-02 |
| Q9Z1K5 | E3 ubiquitin-protein ligase ARIH1 OS=Mus musculus OX=10090 | 4 | 2 | 3 | 2 | 555 | 64 | 5,08 | 1523 | | 2 | | 0,787 | 0,826 | 0,791 | 0,652 | 1,341 | 0,868 | 1,055 | 0,958 | 0,72 | -0,28 | 3,7E-02 |
| Q9R257 | Heme-binding protein 1 OS=Mus musculus OX=10090 GN=Hebp | 38 | 6 | 6 | 6 | 190 | 21,1 | 5,26 | 3377 | 3,89 | 5 | 1 | 1,083 | 1,32 | 1,159 | 1,249 | 1,155 | 0,941 | 0,939 | 1,006 | 1,19 | 0,19 | 3,8E-02 |
| Q8R1N4 | NudC domain-containing protein 3 OS=Mus musculus OX=10090 | 6 | 2 | 2 | 2 | 363 | 40,9 | 5,26 | 1004 | | 2 | | 1,251 | 1,057 | 1,162 | 1,149 | 0,604 | 1,045 | 0,992 | 0,849 | 1,32 | 0,32 | 3,8E-02 |
| P47867 | Secretogranin-3 OS=Mus musculus OX=10090 GN=Scg3 PE=1 SV | 14 | 6 | 6 | 6 | 471 | 53,3 | 4,93 | 1751 | 2,54 | 3 | 3 | 0,918 | 0,938 | 1,028 | 0,968 | 0,842 | 0,866 | 0,92 | 0,906 | 1,09 | 0,09 | 3,8E-02 |
| Q9D0M3 | Cytochrome c1, heme protein, mitochondrial OS=Mus musculus | 24 | 6 | 16 | 6 | 325 | 35,3 | 9,16 | 8804 | 2,62 | 5 | 2 | 0,989 | 0,948 | 0,957 | 0,797 | 0,991 | 1,081 | 1,039 | 1,106 | 0,88 | -0,12 | 3,8E-02 |
| P63028 | Translationally-controlled tumor protein OS=Mus musculus OX= | 26 | 4 | 4 | 4 | 172 | 19,5 | 4,86 | | 9,87 | | 4 | 1,018 | 0,885 | 1,15 | 0,956 | 0,687 | 0,916 | 0,852 | 0,75 | 1,25 | 0,25 | 3,8E-02 |
| Q61147 | Ceruloplasmin OS=Mus musculus OX=10090 GN=Cp PE=1 SV=2 | 4 | 3 | 5 | 3 | 1061 | 121,1 | 5,85 | 1584 | 4,13 | 3 | 1 | 0,977 | 1,057 | 0,977 | 1,137 | 1,272 | 1,048 | 1,284 | 1,361 | 0,84 | -0,16 | 3,9E-02 |
| Q9WV98 | Mitochondrial import inner membrane translocase subunit Tim9 | 65 | 5 | 6 | 5 | 89 | 10,3 | 7,21 | 1039 | 10,76 | 1 | 4 | 1,076 | 1,092 | 1,141 | 1,095 | 0,806 | 0,958 | 1,079 | 0,954 | 1,16 | 0,16 | 3,9E-02 |
| Q8BRT1 | CLIP-associating protein 2 OS=Mus musculus OX=10090 GN=Cla | 18 | 20 | 30 | 20 | 1286 | 140,7 | 8,63 | 9953 | 24,84 | 13 | 10 | 1,018 | 0,954 | 1,018 | 0,951 | 0,855 | 0,915 | 0,959 | 0,913 | 1,08 | 0,08 | 3,9E-02 |
| O35678 | Monoglyceride lipase OS=Mus musculus OX=10090 GN=Mgll PE | 6 | 2 | 3 | 2 | 303 | 33,4 | 7,15 | 1946 | | 2 | | 0,839 | 0,941 | 1,087 | 1,104 | 1,24 | 1,179 | 1,129 | 1,135 | 0,85 | -0,15 | 4,0E-02 |
| Q8VIM0 | Hepatitis A virus cellular receptor 2 homolog OS=Mus musculus | 3 | 1 | 1 | 1 | 281 | 30,9 | 7,83 | 381 | | 1 | | 0,972 | 1,04 | 1,059 | 1,113 | 0,579 | 0,913 | 0,949 | 0,822 | 1,28 | 0,28 | 4,0E-02 |
| Q9DA73 | Coiled-coil domain-containing protein 89 OS=Mus musculus OX= | 5 | 1 | 1 | 1 | 370 | 43,2 | 5,62 | | 0 | | 1 | 1,082 | 0,707 | 1,04 | 1,707 | 2,193 | 1,726 | 3,637 | 1,964 | 0,48 | -0,52 | 4,0E-02 |
| Q9CQV1 | Mitochondrial import inner membrane translocase subunit TIM1 | 6 | 1 | 1 | 1 | 125 | 13,8 | 9,64 | 789 | | 1 | | 0,981 | 1,058 | 1,2 | 0,966 | 0,744 | 0,922 | 0,949 | 0,858 | 1,21 | 0,21 | 4,1E-02 |
| P60122 | RuvB-like 1 OS=Mus musculus OX=10090 GN=Ruvbl1 PE=1 SV=1 | 10 | 4 | 4 | 4 | 456 | 50,2 | 6,42 | 1675 | 7,1 | 2 | 2 | 1,154 | 0,945 | 1,088 | 1,024 | 0,889 | 0,982 | 0,887 | 0,933 | 1,14 | 0,14 | 4,1E-02 |
| Q91WE1 | Sorting nexin-15 OS=Mus musculus OX=10090 GN=Snx15 PE=1 S | 5 | 1 | 1 | 1 | 337 | 37,7 | 5,24 | 174 | | 1 | | 1,054 | 0,879 | 0,748 | 0,669 | 1,171 | 0,906 | 1,494 | 1,491 | 0,66 | -0,34 | 4,1E-02 |
| Q91VJ4 | Serine/threonine-protein kinase 38 OS=Mus musculus OX=1009 | 5 | 2 | 2 | 2 | 465 | 54,1 | 7,34 | 597 | | 2 | | 1,023 | 1,005 | 1,2 | 0,861 | 0,665 | 0,89 | 0,879 | 0,734 | 1,29 | 0,29 | 4,1E-02 |
| P54071 | Isocitrate dehydrogenase [NADP], mitochondrial OS=Mus muscu | 27 | 10 | 16 | 9 | 452 | 50,9 | 8,69 | 6119 | 17,53 | 7 | 4 | 1,03 | 0,959 | 1,026 | 1,06 | 0,969 | 0,961 | 0,972 | 0,938 | 1,06 | 0,06 | 4,1E-02 |
| Q3TDD9 | Protein phosphatase 1 regulatory subunit 21 OS=Mus musculus | 15 | 9 | 11 | 9 | 780 | 88,3 | 6,9 | 3307 | 14,83 | 4 | 5 | 0,973 | 0,982 | 0,97 | 0,971 | 0,96 | 0,947 | 0,971 | 0,938 | 1,02 | 0,02 | 4,2E-02 |
| P12960 | Contactin-1 OS=Mus musculus OX=10090 GN=Cntn1 PE=1 SV=1 | 38 | 34 | 79 | 34 | 1020 | 113,3 | 6,16 | 32209 | 63,7 | 22 | 17 | 1,06 | 1,189 | 1,017 | 1,004 | 0,748 | 0,772 | 0,988 | 0,972 | 1,23 | 0,23 | 4,2E-02 |
| Q9DBS5 | Kinesin light chain 4 OS=Mus musculus OX=10090 GN=Klc4 PE=1 | 6 | 2 | 2 | 1 | 619 | 68,6 | 6,09 | 1273 | | 2 | | 0,803 | 0,875 | 0,707 | 0,759 | 0,949 | 0,807 | 0,957 | 1,007 | 0,85 | -0,15 | 4,2E-02 |
| Q8BKX6 | Serine/threonine-protein kinase SMG1 OS=Mus musculus OX=10 | 0 | 1 | 1 | 1 | 3658 | 409,5 | 6,4 | 487 | | 1 | | 1,072 | 1,036 | 1,147 | 0,997 | 1,198 | 1,179 | 1,139 | 1,116 | 0,92 | -0,08 | 4,2E-02 |
| Q62351 | Transferrin receptor protein 1 OS=Mus musculus OX=10090 GN | 12 | 9 | 13 | 9 | 763 | 85,7 | 6,57 | 5248 | 11,83 | 6 | 3 | 0,926 | 1,003 | 0,957 | 0,993 | 1,024 | 1,053 | 0,989 | 1,049 | 0,94 | -0,06 | 4,2E-02 |
| Q9CR26 | Vacuolar protein sorting-associated protein VTA1 homolog OS=I | 13 | 3 | 4 | 3 | 309 | 33,9 | 6,13 | 2702 | | 3 | | 1,034 | 1,063 | 0,939 | 1,068 | 1,203 | 1,093 | 1,075 | 1,204 | 0,90 | -0,10 | 4,2E-02 |
| P56818 | Beta-secretase 1 OS=Mus musculus OX=10090 GN=Bace1 PE=1 S | 4 | 2 | 2 | 2 | 501 | 55,7 | 5,6 | 1039 | | 2 | | 0,986 | 1,109 | 1,042 | 1,102 | 0,76 | 1,019 | 0,925 | 0,91 | 1,17 | 0,17 | 4,2E-02 |
| Q8CHP5 | Partner of Y14 and mago OS=Mus musculus OX=10090 GN=Pym | 5 | 1 | 1 | 1 | 203 | 22,7 | 9,5 | | 3,07 | | 1 | 1,044 | 1,193 | 1,074 | 1,044 | 1,49 | 1,255 | 1,35 | 1,127 | 0,83 | -0,17 | 4,2E-02 |
| Q8K2Q9 | Shootin-1 OS=Mus musculus OX=10090 GN=Shtn1 PE=1 SV=1 | 22 | 13 | 18 | 13 | 631 | 71,3 | 5,44 | 7371 | 20,12 | 8 | 5 | 1,02 | 1,026 | 1,044 | 1,06 | 0,777 | 0,957 | 0,977 | 0,952 | 1,13 | 0,13 | 4,3E-02 |
| Q91WU0 | Carboxylesterase 1F OS=Mus musculus OX=10090 GN=Ces1f PE | 2 | 1 | 2 | 1 | 561 | 61,6 | 6,48 | 1008 | | 1 | | 0,977 | 0,904 | 0,915 | 0,872 | 1,314 | 0,942 | 1,069 | 1,197 | 0,81 | -0,19 | 4,3E-02 |
| Q3UNH4 | G protein-regulated inducer of neurite outgrowth 1 OS=Mus mu | 61 | 59 | 112 | 59 | 932 | 95,4 | 7,93 | 42272 | 106,14 | 34 | 30 | 1,027 | 0,966 | 1,107 | 0,999 | 0,862 | 0,981 | 0,936 | 0,922 | 1,11 | 0,11 | 4,3E-02 |
| Q8VD75 | Huntingtin-interacting protein 1 OS=Mus musculus OX=10090 G | 13 | 10 | 15 | 9 | 1029 | 115,1 | 5,43 | 7227 | 5,48 | 8 | 2 | 0,927 | 0,868 | 1,035 | 1,113 | 1,354 | 1,087 | 1,215 | 1,118 | 0,83 | -0,17 | 4,3E-02 |
| B9EJA2 | Cortactin-binding protein 2 OS=Mus musculus OX=10090 GN=Ct | 3 | 4 | 4 | 4 | 1648 | 178,7 | 8,19 | 1140 | 1,82 | 3 | 1 | 0,967 | 0,951 | 0,854 | 0,788 | 1,256 | 1,045 | 1,06 | 0,964 | 0,82 | -0,18 | 4,3E-02 |
| Q9QXB9 | Developmentally-regulated GTP-binding protein 2 OS=Mus mus | 6 | 3 | 5 | 3 | 364 | 40,7 | 8,88 | 1419 | 2,26 | 2 | 1 | 1,082 | 1,046 | 1,188 | 0,99 | 0,772 | 1,015 | 0,951 | 0,891 | 1,19 | 0,19 | 4,4E-02 |
| P07758 | Alpha-1-antitrypsin 1-1 OS=Mus musculus OX=10090 GN=Serpin | 2 | 1 | 1 | 1 | 413 | 46 | 5,72 | | 2,47 | | 1 | 1,231 | 1,444 | 1,446 | 1,363 | 1,965 | 1,478 | 1,737 | 1,538 | 0,82 | -0,18 | 4,4E-02 |
| Q9Z0Y1 | Dynactin subunit 3 OS=Mus musculus OX=10090 GN=Dctn3 PE= | 42 | 8 | 9 | 8 | 186 | 21 | 6,06 | 2596 | 11,5 | 4 | 4 | 1,001 | 0,959 | 1,062 | 0,967 | 0,752 | 0,919 | 0,933 | 0,898 | 1,14 | 0,14 | 4,4E-02 |
| Q6PHU5 | Sortilin OS=Mus musculus OX=10090 GN=Sort1 PE=1 SV=1 | 1 | 1 | 1 | 1 | 825 | 91,1 | 5,88 | | 2,3 | | 1 | 0,988 | 0,801 | 1,078 | 1,075 | 1,424 | 1,208 | 1,131 | 1,137 | 0,80 | -0,20 | 4,4E-02 |
| Q99J99 | 3-mercaptopyruvate sulfurtransferase OS=Mus musculus OX=10 | 40 | 10 | 16 | 10 | 297 | 33,1 | 6,47 | 8824 | 5,03 | 10 | 1 | 0,942 | 1,014 | 0,941 | 0,965 | 1,065 | 0,971 | 1,063 | 1,047 | 0,93 | -0,07 | 4,4E-02 |
| Q6Y5D8 | Rho GTPase-activating protein 10 OS=Mus musculus OX=10090 | 1 | 1 | 2 | 1 | 786 | 89,3 | 7,18 | 1111 | | 1 | | 1,026 | 0,917 | 0,932 | 0,936 | 0,594 | 0,814 | 0,876 | 0,839 | 1,22 | 0,22 | 4,5E-02 |
| Q8BKC5 | Importin-5 OS=Mus musculus OX=10090 GN=Ipo5 PE=1 SV=3 | 6 | 6 | 7 | 6 | 1097 | 123,5 | 4,93 | 3827 | 3,55 | 4 | 2 | 1,032 | 0,912 | 1,103 | 1,032 | 0,688 | 0,93 | 0,917 | 0,855 | 1,20 | 0,20 | 4,5E-02 |
| Q9ERT9 | Protein phosphatase 1 regulatory subunit 1A OS=Mus musculus | 43 | 5 | 9 | 5 | 171 | 18,7 | 5,25 | 2907 | 9,89 | 3 | 3 | 0,991 | 1,027 | 0,919 | 0,963 | 1,017 | 1,117 | 1,104 | 1,018 | 0,92 | -0,08 | 4,5E-02 |
| Q8C196 | Carbamoyl-phosphate synthase [ammonia], mitochondrial OS=M | 9 | 12 | 20 | 12 | 1500 | 164,5 | 6,92 | 8960 | 8,54 | 11 | 3 | 1,614 | 1,776 | 1,762 | 1,956 | 2,584 | 4,518 | 2,709 | 2,356 | 0,58 | -0,42 | 4,5E-02 |
| Q8BVI4 | Dihydropteridine reductase OS=Mus musculus OX=10090 GN=Q | 27 | 4 | 11 | 4 | 241 | 25,6 | 7,81 | 6544 | | 4 | | 1,165 | 1,059 | 1,042 | 1,033 | 1,418 | 1,17 | 1,152 | 1,243 | 0,86 | -0,14 | 4,6E-02 |
| Q8BMJ2 | Leucine--tRNA ligase, cytoplasmic OS=Mus musculus OX=10090 | 1 | 1 | 1 | 1 | 1178 | 134,1 | 7,05 | | 2,47 | | 1 | 1,073 | 0,885 | 0,991 | 1,074 | 0,702 | 0,93 | 0,878 | 0,847 | 1,20 | 0,20 | 4,6E-02 |
| Q9R1Q8 | Transgelin-3 OS=Mus musculus OX=10090 GN=Tagln3 PE=1 SV= | 42 | 10 | 25 | 9 | 199 | 22,5 | 7,33 | 11085 | 13,49 | 5 | 5 | 1,107 | 1,036 | 1,094 | 1,082 | 0,874 | 1,047 | 1,015 | 0,975 | 1,10 | 0,10 | 4,6E-02 |
| Q91WK2 | Eukaryotic translation initiation factor 3 subunit H OS=Mus mus | 9 | 3 | 3 | 3 | 352 | 39,8 | 6,67 | 985 | 2,3 | 2 | 1 | 0,917 | 0,941 | 1,06 | 0,963 | 1,007 | 1,066 | 1,2 | 1,122 | 0,88 | -0,12 | 4,7E-02 |
| P19157 | Glutathione S-transferase P 1 OS=Mus musculus OX=10090 GN= | 43 | 8 | 22 | 8 | 210 | 23,6 | 7,87 | 2674 | 45,45 | 4 | 5 | 1,042 | 1,066 | 0,955 | 1,029 | 1,081 | 1,086 | 1,085 | 1,078 | 0,95 | -0,05 | 4,8E-02 |

| | | | | | | | | | | | | | | | | | | | | | | | |
|---|---|---|---|---|---|---|---|---|---|---|---|---|---|---|---|---|---|---|---|---|---|---|---|
| Q3TJZ6 | Protein FAM98A OS=Mus musculus OX=10090 GN=Fam98a PE= | 3 | 2 | 2 | 2 | 515 | 55 | 8,95 | 722 | 2,12 | 1 | 1 | 0,888 | 0,949 | 0,865 | 1,129 | 1,196 | 1,024 | 1,291 | 1,131 | 0,83 | -0,17 | 4,8E-02 |
| P56564 | Excitatory amino acid transporter 1 OS=Mus musculus OX=1009 | 9 | 4 | 15 | 4 | 543 | 59,6 | 8,4 | 5683 | 17,14 | 2 | 2 | 1,006 | 1,031 | 1,01 | 0,964 | 0,695 | 0,803 | 0,964 | 0,927 | 1,18 | 0,18 | 4,8E-02 |
| Q3TLI0 | Trafficking protein particle complex subunit 10 OS=Mus musculu | 1 | 1 | 1 | 1 | 1259 | 141,5 | 5,96 | 502 | | 1 | | 0,684 | 1,133 | 1,092 | 0,981 | 1,188 | 1,122 | 1,405 | 1,439 | 0,75 | -0,25 | 4,9E-02 |
| Q8R344 | Coiled-coil domain-containing protein 12 OS=Mus musculus OX= | 4 | 1 | 1 | 1 | 166 | 18,9 | 7,21 | 465 | | 1 | | 1,221 | 1,065 | 1,011 | 1,033 | 0,85 | 1,015 | 0,905 | 0,971 | 1,16 | 0,16 | 4,9E-02 |
| Q8K310 | Matrin-3 OS=Mus musculus OX=10090 GN=Matr3 PE=1 SV=1 | 19 | 13 | 19 | 13 | 846 | 94,6 | 6,25 | 7153 | 13,5 | 9 | 4 | 0,846 | 0,822 | 1 | 0,95 | 1,089 | 0,917 | 1,213 | 1,155 | 0,83 | -0,17 | 4,9E-02 |
| O88712 | C-terminal-binding protein 1 OS=Mus musculus OX=10090 GN= | 18 | 6 | 10 | 6 | 441 | 47,7 | 6,77 | 2128 | 16,02 | 3 | 4 | 0,926 | 1,039 | 0,887 | 0,975 | 1,187 | 1,029 | 1,028 | 1,073 | 0,89 | -0,11 | 4,9E-02 |
| Q9CZ13 | Cytochrome b-c1 complex subunit 1, mitochondrial OS=Mus mu | 31 | 12 | 34 | 12 | 480 | 52,8 | 6,21 | 14645 | 40,97 | 8 | 8 | 1,004 | 0,957 | 0,971 | 0,982 | 0,989 | 1,027 | 1,054 | 1,108 | 0,94 | -0,06 | 5,0E-02 |
| Q60865 | Caprin-1 OS=Mus musculus OX=10090 GN=Caprin1 PE=1 SV=2 | 9 | 7 | 9 | 7 | 707 | 78,1 | 5,25 | 4764 | 2,03 | 6 | 1 | 1,102 | 0,999 | 1,064 | 1,108 | 0,861 | 0,998 | 0,962 | 1,024 | 1,11 | 0,11 | 5,0E-02 |
| Q99LC5 | Electron transfer flavoprotein subunit alpha, mitochondrial OS= | 38 | 10 | 21 | 10 | 333 | 35 | 8,38 | 11987 | 6,89 | 7 | 3 | 1,217 | 1,234 | 1,197 | 1,337 | 1,523 | 1,334 | 1,393 | 1,302 | 0,90 | -0,10 | 5,0E-02 |
| Q9QYH6 | Melanoma-associated antigen D1 OS=Mus musculus OX=10090 | 4 | 2 | 4 | 2 | 775 | 85,6 | 7,5 | 1811 | | 2 | | 1,082 | 1,061 | 0,867 | 1,132 | 0,573 | 1 | 0,823 | 0,674 | 1,35 | 0,35 | 5,0E-02 |
| Q60759 | Glutaryl-CoA dehydrogenase, mitochondrial OS=Mus musculus | 4 | 1 | 2 | 1 | 438 | 48,6 | 8,73 | 966 | | 1 | | 1,198 | 1,144 | 1,088 | 1,208 | 1,648 | 1,264 | 1,248 | 1,424 | 0,83 | -0,17 | 5,1E-02 |
| P35486 | Pyruvate dehydrogenase E1 component subunit alpha, somatic | 43 | 16 | 22 | 16 | 390 | 43,2 | 8,19 | 8315 | 20,35 | 11 | 6 | 1,035 | 1,063 | 1,001 | 0,981 | 0,875 | 0,972 | 0,971 | 0,969 | 1,08 | 0,08 | 5,1E-02 |
| Q62277 | Synaptophysin OS=Mus musculus OX=10090 GN=Syp PE=1 SV=2 | 10 | 3 | 13 | 3 | 314 | 34 | 4,94 | 7541 | 6,29 | 3 | 1 | 1,085 | 1,082 | 0,981 | 0,882 | 0,596 | 0,848 | 0,892 | 0,873 | 1,26 | 0,26 | 5,1E-02 |
| Q6KAR6 | Exocyst complex component 3 OS=Mus musculus OX=10090 GN | 8 | 5 | 6 | 5 | 755 | 86,4 | 6,2 | 3561 | | 5 | | 0,877 | 0,95 | 0,874 | 0,773 | 1,011 | 0,945 | 0,961 | 0,942 | 0,90 | -0,10 | 5,2E-02 |
| Q9JI44 | DNA methyltransferase 1-associated protein 1 OS=Mus musculu | 2 | 1 | 3 | 1 | 468 | 53,1 | 9,5 | 1297 | | 1 | | 0,92 | 0,934 | 1,014 | 1,002 | 0,898 | 0,896 | 0,888 | 0,936 | 1,07 | 0,07 | 5,2E-02 |
| Q8BX94 | Oxysterol-binding protein-related protein 2 OS=Mus musculus C | 5 | 2 | 2 | 1 | 484 | 55,3 | 6,2 | 526 | 2,85 | 1 | 1 | 0,796 | 0,872 | 0,775 | 0,873 | 1,276 | 1,05 | 0,919 | 0,921 | 0,80 | -0,20 | 5,2E-02 |
| P70677 | Caspase-3 OS=Mus musculus OX=10090 GN=Casp3 PE=1 SV=1 | 22 | 5 | 13 | 5 | 277 | 31,5 | 6,92 | 3891 | 15,14 | 3 | 3 | 1,023 | 1,044 | 1,127 | 1,121 | 0,665 | 0,99 | 0,974 | 0,913 | 1,22 | 0,22 | 5,2E-02 |
| Q9Z2Y8 | Pyridoxal phosphate homeostasis protein OS=Mus musculus OX | 3 | 1 | 2 | 1 | 274 | 30 | 8,27 | | 5,14 | | 1 | 1,135 | 1,052 | 1,055 | 1,172 | 1,731 | 1,265 | 1,233 | 1,336 | 0,79 | -0,21 | 5,2E-02 |
| Q9QYC0 | Alpha-adducin OS=Mus musculus OX=10090 GN=Add1 PE=1 SV= | 26 | 14 | 24 | 13 | 735 | 80,6 | 5,9 | 10495 | 19,91 | 11 | 5 | 1,031 | 1,002 | 1,009 | 0,98 | 0,862 | 0,97 | 0,969 | 0,953 | 1,07 | 0,07 | 5,3E-02 |
| Q8VEK0 | Cell cycle control protein 50A OS=Mus musculus OX=10090 GN= | 4 | 2 | 4 | 2 | 364 | 41 | 8,37 | 1854 | | 2 | | 1,319 | 0,966 | 1,343 | 1,095 | 0,787 | 0,862 | 1,102 | 0,884 | 1,30 | 0,30 | 5,3E-02 |
| Q91VM9 | Inorganic pyrophosphatase 2, mitochondrial OS=Mus musculus | 11 | 4 | 5 | 4 | 330 | 38,1 | 6,98 | 2765 | 2,59 | 3 | 1 | 0,896 | 0,859 | 0,966 | 0,952 | 1,016 | 1,067 | 0,939 | 0,999 | 0,91 | -0,09 | 5,3E-02 |
| Q9EQU5 | Protein SET OS=Mus musculus OX=10090 GN=Set PE=1 SV=1 | 10 | 3 | 3 | 3 | 289 | 33,4 | 4,32 | 1716 | | 3 | | 0,967 | 1,087 | 0,927 | 1,293 | 0,717 | 0,97 | 0,833 | 0,827 | 1,28 | 0,28 | 5,4E-02 |
| Q99J08 | SEC14-like protein 2 OS=Mus musculus OX=10090 GN=Sec14l2 P | 2 | 1 | 1 | 1 | 403 | 46,3 | 7,11 | 365 | | 1 | | 0,919 | 1,039 | 0,873 | 1,139 | 1,544 | 1,096 | 1,121 | 1,565 | 0,75 | -0,25 | 5,5E-02 |
| P46664 | Adenylosuccinate synthetase isozyme 2 OS=Mus musculus OX=1 | 13 | 7 | 9 | 7 | 456 | 50 | 6,38 | 1895 | 9,83 | 3 | 4 | 1,091 | 1,024 | 1,06 | 1,012 | 0,76 | 0,966 | 0,962 | 0,975 | 1,14 | 0,14 | 5,5E-02 |
| Q69ZQ1 | Myogenesis-regulating glycosidase OS=Mus musculus OX=1009 | 3 | 2 | 2 | 2 | 716 | 81,3 | 8,29 | 588 | 0 | 1 | 1 | 0,786 | 0,759 | 0,83 | 0,756 | 1,068 | 0,828 | 0,899 | 0,866 | 0,86 | -0,14 | 5,5E-02 |
| Q05D44 | Eukaryotic translation initiation factor 5B OS=Mus musculus OX= | 1 | 1 | 1 | 1 | 1216 | 137,5 | 5,59 | | 1,9 | | 1 | 0,998 | 0,829 | 0,899 | 0,955 | 0,546 | 0,834 | 0,84 | 0,719 | 1,25 | 0,25 | 5,5E-02 |
| Q64442 | Sorbitol dehydrogenase OS=Mus musculus OX=10090 GN=Sord | 27 | 9 | 23 | 9 | 357 | 38,2 | 7,02 | 9770 | 32,17 | 6 | 3 | 1,527 | 1,657 | 1,542 | 1,51 | 2,08 | 1,57 | 2,058 | 1,748 | 0,84 | -0,16 | 5,5E-02 |
| Q3UEB3 | Poly(U)-binding-splicing factor PUF60 OS=Mus musculus OX=10 | 4 | 3 | 3 | 3 | 564 | 60,2 | 5,29 | 606 | 5,19 | 1 | 2 | 1,286 | 1,727 | 1,071 | 1,334 | 1,12 | 1,034 | 0,932 | 0,976 | 1,33 | 0,33 | 5,5E-02 |
| Q9CQC9 | GTP-binding protein SAR1b OS=Mus musculus OX=10090 GN=Sa | 16 | 3 | 5 | 3 | 198 | 22,4 | 6,11 | 2870 | | 3 | | 0,902 | 0,93 | 0,843 | 0,871 | 1,046 | 0,916 | 0,944 | 1,118 | 0,88 | -0,12 | 5,5E-02 |
| Q9DBG6 | Dolichyl-diphosphooligosaccharide--protein glycosyltransferase | 17 | 8 | 14 | 8 | 631 | 69 | 5,81 | 6110 | 17,07 | 6 | 3 | 0,936 | 0,951 | 0,861 | 1,046 | 1,122 | 0,982 | 1,05 | 1,122 | 0,89 | -0,11 | 5,5E-02 |
| Q61207 | Prosaposin OS=Mus musculus OX=10090 GN=Psap PE=1 SV=2 | 16 | 8 | 13 | 8 | 557 | 61,4 | 5,19 | 4480 | 20,12 | 5 | 4 | 0,96 | 1,115 | 0,899 | 1,27 | 0,743 | 0,909 | 0,907 | 0,809 | 1,26 | 0,26 | 5,6E-02 |
| Q8K221 | Arfaptin-2 OS=Mus musculus OX=10090 GN=Arfip2 PE=1 SV=2 | 15 | 4 | 6 | 4 | 341 | 37,7 | 5,87 | 2250 | 2,45 | 4 | 1 | 0,845 | 0,805 | 0,721 | 0,738 | 0,882 | 0,823 | 0,836 | 0,873 | 0,91 | -0,09 | 5,6E-02 |
| Q921F2 | TAR DNA-binding protein 43 OS=Mus musculus OX=10090 GN=T | 6 | 2 | 3 | 2 | 414 | 44,5 | 6,7 | 1862 | | 2 | | 0,995 | 0,909 | 1,071 | 1,025 | 1,148 | 1,081 | 1,226 | 1,043 | 0,89 | -0,11 | 5,6E-02 |
| P63168 | Dynein light chain 1, cytoplasmic OS=Mus musculus OX=10090 | 39 | 5 | 15 | 3 | 89 | 10,4 | 7,4 | 8055 | 0 | 4 | 1 | 1,027 | 0,961 | 1,271 | 0,986 | 0,488 | 0,938 | 0,848 | 0,829 | 1,37 | 0,37 | 5,8E-02 |
| Q8JZZ7 | Adhesion G protein-coupled receptor L2 OS=Mus musculus OX= | 3 | 4 | 5 | 4 | 1487 | 166,5 | 6,71 | | 12,36 | | 4 | 0,866 | 0,877 | 0,894 | 0,826 | 0,965 | 0,983 | 0,95 | 0,858 | 0,92 | -0,08 | 5,8E-02 |
| Q9D1C8 | Vacuolar protein sorting-associated protein 28 homolog OS=Mu | 7 | 1 | 2 | 1 | 221 | 25,4 | 5,54 | | 6,95 | | 1 | 0,613 | 1,211 | 0,75 | 0,934 | 2,132 | 1,013 | 1,595 | 1,302 | 0,58 | -0,42 | 5,8E-02 |
| Q9Z0S1 | 3'(2'),5'-bisphosphate nucleotidase 1 OS=Mus musculus OX=100 | 4 | 1 | 2 | 1 | 308 | 33,2 | 5,82 | 1071 | | 1 | | 0,882 | 0,802 | 0,907 | 0,982 | 1,173 | 1,114 | 1,021 | 0,907 | 0,85 | -0,15 | 5,9E-02 |
| Q99K85 | Phosphoserine aminotransferase OS=Mus musculus OX=10090 | 23 | 9 | 13 | 9 | 370 | 40,4 | 8,03 | 6757 | 8,07 | 7 | 3 | 0,988 | 1,062 | 0,995 | 1,055 | 0,893 | 1,026 | 0,92 | 0,938 | 1,09 | 0,09 | 5,9E-02 |
| O55017 | Voltage-dependent N-type calcium channel subunit alpha-1B OS | 1 | 1 | 1 | 1 | 2327 | 261,3 | 8,59 | | 0 | | 1 | 1,014 | 0,873 | 1,209 | 0,973 | 0,784 | 0,745 | 0,954 | 0,805 | 1,24 | 0,24 | 5,9E-02 |
| Q8BK63 | Casein kinase I isoform alpha OS=Mus musculus OX=10090 GN= | 3 | 1 | 2 | 1 | 337 | 38,9 | 9,57 | 1039 | | 1 | | 0,933 | 0,824 | 0,89 | 0,924 | 0,664 | 0,753 | 0,721 | 0,904 | 1,17 | 0,17 | 5,9E-02 |
| Q8CI94 | Glycogen phosphorylase, brain form OS=Mus musculus OX=100 | 19 | 16 | 31 | 12 | 843 | 96,7 | 6,73 | 10835 | 24,48 | 12 | 6 | 1,005 | 1,001 | 0,991 | 1,051 | 0,845 | 0,942 | 0,974 | 0,976 | 1,08 | 0,08 | 5,9E-02 |
| Q9JHS4 | ATP-dependent Clp protease ATP-binding subunit clpX-like, mit | 4 | 2 | 2 | 2 | 634 | 69,2 | 7,71 | 914 | | 2 | | 0,906 | 1,019 | 0,999 | 0,997 | 1,334 | 1,055 | 1,032 | 1,185 | 0,85 | -0,15 | 6,0E-02 |
| Q8K0V2 | DCN1-like protein 3 OS=Mus musculus OX=10090 GN=Dcun1d3 | 4 | 1 | 1 | 1 | 304 | 34,4 | 5,12 | 423 | | 1 | | 0,943 | 1,014 | 0,937 | 1,011 | 0,992 | 1,074 | 1,326 | 1,254 | 0,84 | -0,16 | 6,0E-02 |
| Q9Z130 | Heterogeneous nuclear ribonucleoprotein D-like OS=Mus muscu | 13 | 3 | 4 | 3 | 301 | 33,5 | 7,31 | 2266 | | 3 | | 0,908 | 0,982 | 1,016 | 1,055 | 1,128 | 1,009 | 1,268 | 1,123 | 0,87 | -0,13 | 6,1E-02 |
| Q8BW66 | Solute carrier family 25 member 48 OS=Mus musculus OX=1009 | 3 | 1 | 1 | 1 | 306 | 33,4 | 9,03 | 448 | | 1 | | 1,097 | 1,034 | 1,064 | 1,008 | 0,508 | 0,943 | 0,951 | 0,83 | 1,30 | 0,30 | 6,1E-02 |
| Q920Q8 | Influenza virus NS1A-binding protein homolog OS=Mus musculu | 2 | 1 | 2 | 1 | 642 | 71,5 | 5,47 | 947 | | 1 | | 0,906 | 0,739 | 1,031 | 1,052 | 1,247 | 1,032 | 1,13 | 1,099 | 0,83 | -0,17 | 6,1E-02 |
| Q9CZW4 | Long-chain-fatty-acid--CoA ligase 3 OS=Mus musculus OX=1009 | 4 | 3 | 4 | 2 | 720 | 80,4 | 8,54 | 785 | 6,23 | 2 | 1 | 1,12 | 1,174 | 1,136 | 1,07 | 0,782 | 0,967 | 1,145 | 0,862 | 1,20 | 0,20 | 6,2E-02 |
| P62962 | Profilin-1 OS=Mus musculus OX=10090 GN=Pfn1 PE=1 SV=2 | 55 | 8 | 20 | 8 | 140 | 14,9 | 8,28 | 10561 | 6,69 | 6 | 2 | 0,991 | 0,99 | 0,984 | 1,019 | 0,867 | 0,996 | 0,898 | 0,952 | 1,07 | 0,07 | 6,2E-02 |
| Q80UU9 | Membrane-associated progesterone receptor component 2 OS= | 6 | 2 | 4 | 1 | 217 | 23,3 | 5,15 | 2432 | | 2 | | 1,229 | 1,325 | 1,268 | 1,234 | 1,12 | 0,94 | 1,278 | 1,028 | 1,16 | 0,16 | 6,2E-02 |
| Q9QYK7 | RING finger protein 11 OS=Mus musculus OX=10090 GN=Rnf11 | 14 | 2 | 3 | 2 | 154 | 17,4 | 4,78 | 775 | 5,23 | 1 | 1 | 0,991 | 1,02 | 0,998 | 0,926 | 1,108 | 1,01 | 1,027 | 1,06 | 0,94 | -0,06 | 6,3E-02 |
| Q8C854 | Myelin expression factor 2 OS=Mus musculus OX=10090 GN=My | 9 | 6 | 6 | 6 | 591 | 63,3 | 8,87 | 1294 | 7,92 | 2 | 4 | 0,842 | 0,781 | 0,982 | 1,008 | 0,944 | 1,04 | 1,259 | 1,187 | 0,82 | -0,18 | 6,3E-02 |
| Q8C5W3 | Tubulin-specific chaperone cofactor E-like protein OS=Mus mus | 2 | 1 | 1 | 1 | 424 | 48 | 5,53 | 662 | | 1 | | 0,834 | 0,885 | 0,906 | 0,932 | 0,676 | 0,895 | 0,668 | 0,793 | 1,17 | 0,17 | 6,3E-02 |
| Q78JW9 | Ubiquitin domain-containing protein UBFD1 OS=Mus musculus | 14 | 3 | 3 | 3 | 368 | 40,1 | 8,85 | 1279 | 3,58 | 2 | 1 | 1,049 | 0,97 | 0,969 | 0,967 | 0,8 | 0,977 | 0,915 | 0,881 | 1,11 | 0,11 | 6,3E-02 |
| P51880 | Fatty acid-binding protein, brain OS=Mus musculus OX=10090 G | 52 | 6 | 20 | 6 | 132 | 14,9 | 5,63 | 5458 | 34,65 | 2 | 6 | 0,989 | 1,048 | 1,19 | 0,985 | 0,918 | 0,963 | 0,908 | 0,967 | 1,12 | 0,12 | 6,4E-02 |
| Q8CDD9 | Ligand-dependent nuclear receptor-interacting factor 1 OS=Mus | 1 | 1 | 1 | 1 | 755 | 82,9 | 9,5 | 398 | | 1 | | 1,077 | 1,142 | 1,343 | 1,051 | 0,928 | 1,088 | 0,961 | 0,954 | 1,17 | 0,17 | 6,4E-02 |
| O35459 | Delta(3,5)-Delta(2,4)-dienoyl-CoA isomerase, mitochondrial OS= | 23 | 5 | 5 | 5 | 327 | 36,1 | 7,71 | 2635 | 0 | 4 | 1 | 1,018 | 0,961 | 1,17 | 1,188 | 0,938 | 0,925 | 0,933 | 1,006 | 1,14 | 0,14 | 6,4E-02 |
| Q9CZG3 | COMM domain-containing protein 8 OS=Mus musculus OX=100 | 13 | 2 | 2 | 2 | 183 | 20,8 | 5,59 | 775 | | 2 | | 1,148 | 1,069 | 1,018 | 1,185 | 1,324 | 1,091 | 1,567 | 1,391 | 0,82 | -0,18 | 6,5E-02 |
| Q99LF4 | tRNA-splicing ligase RtcB homolog OS=Mus musculus OX=10090 | 18 | 7 | 10 | 7 | 505 | 55,2 | 7,23 | 4604 | 4,57 | 5 | 2 | 0,998 | 1,014 | 0,918 | 0,999 | 1,042 | 1,075 | 1,012 | 1,03 | 0,94 | -0,06 | 6,5E-02 |
| P62627 | Dynein light chain roadblock-type 1 OS=Mus musculus OX=1009 | 74 | 5 | 14 | 5 | 96 | 11 | 7,25 | 2780 | 22 | 3 | 4 | 1,012 | 1,108 | 0,937 | 1,192 | 0,964 | 0,932 | 0,95 | 0,855 | 1,15 | 0,15 | 6,5E-02 |
| Q9D3D9 | ATP synthase subunit delta, mitochondrial OS=Mus musculus O | 17 | 3 | 6 | 3 | 168 | 17,6 | 5,08 | 1855 | 8,88 | 2 | 2 | 1,025 | 1,158 | 0,944 | 1,198 | 0,779 | 0,918 | 0,864 | 1,04 | 1,20 | 0,20 | 6,5E-02 |
| Q62393-3 | Isoform 3 of Tumor protein D52 OS=Mus musculus OX=10090 G | 17 | 3 | 4 | 3 | 199 | 21,6 | 5,03 | 1497 | 2,16 | 2 | 1 | 1,052 | 1,043 | 1,059 | 0,859 | 1,58 | 1,065 | 1,264 | 1,186 | 0,79 | -0,21 | 6,5E-02 |
| P56391 | Cytochrome c oxidase subunit 6B1 OS=Mus musculus OX=10090 | 42 | 6 | 27 | 6 | 86 | 10,1 | 8,72 | 8686 | 29,08 | 5 | 3 | 1,04 | 1,029 | 1,003 | 1,101 | 0,672 | 1,019 | 0,922 | 0,876 | 1,20 | 0,20 | 6,5E-02 |
| Q04207 | Transcription factor p65 OS=Mus musculus OX=10090 GN=Rela | 2 | 1 | 1 | 1 | 549 | 60,2 | 5,76 | 170 | | 1 | | 0,925 | 1,139 | 1,001 | 1,237 | 1,204 | 1,216 | 1,257 | 1,267 | 0,87 | -0,13 | 6,6E-02 |
| Q8BYL4 | Tyrosine--tRNA ligase, mitochondrial OS=Mus musculus OX=100 | 5 | 2 | 3 | 2 | 472 | 52,6 | 9,16 | 1610 | | 2 | | 1,069 | 0,897 | 1,023 | 1,108 | 0,81 | 0,907 | 0,959 | 0,92 | 1,14 | 0,14 | 6,6E-02 |
| Q8R550 | SH3 domain-containing kinase-binding protein 1 OS=Mus muscu | 11 | 7 | 7 | 7 | 709 | 78,1 | 7,55 | 2194 | 8,96 | 4 | 3 | 1,083 | 1,11 | 1,171 | 1,13 | 1,077 | 1,042 | 1,101 | 1,075 | 1,05 | 0,05 | 6,6E-02 |

| | | | | | | | | | | | | | | | | | | | | | | | |
|---|---|---|---|---|---|---|---|---|---|---|---|---|---|---|---|---|---|---|---|---|---|---|---|
| Q80YF9 | Rho GTPase-activating protein 33 OS=Mus musculus OX=10090 | 2 | 2 | 2 | 2 | 1305 | 139,7 | 9,19 | 472 | 2,66 | 1 | 1 | 0,987 | 0,946 | 0,957 | 0,879 | 0,801 | 0,82 | 0,767 | 0,956 | 1,13 | 0,13 | 6,6E-02 |
| Q69ZR9 | Protein TASOR OS=Mus musculus OX=10090 GN=Tasor PE=1 SV | 1 | 1 | 1 | 1 | 1610 | 181,3 | 5,67 | 175 | | 1 | | 0,888 | 0,958 | 1,086 | 1,092 | 0,824 | 0,757 | 0,888 | 0,96 | 1,17 | 0,17 | 6,6E-02 |
| O88448 | Kinesin light chain 2 OS=Mus musculus OX=10090 GN=Klc2 PE=1 | 21 | 11 | 17 | 8 | 599 | 66,6 | 7,21 | 5458 | 23,52 | 6 | 6 | 1,077 | 1,029 | 1,066 | 1,203 | 0,742 | 1,034 | 0,952 | 0,98 | 1,18 | 0,18 | 6,6E-02 |
| O70251 | Elongation factor 1-beta OS=Mus musculus OX=10090 GN=Eef1l | 15 | 3 | 4 | 2 | 225 | 24,7 | 4,69 | | 11,39 | | 3 | 1,157 | 1,003 | 1,18 | 1,349 | 0,934 | 1,008 | 0,969 | 1,082 | 1,17 | 0,17 | 6,6E-02 |
| Q9Z247 | Peptidyl-prolyl cis-trans isomerase FKBP9 OS=Mus musculus OX | 5 | 3 | 3 | 3 | 570 | 63 | 5,21 | 1414 | | 3 | | 0,848 | 0,971 | 0,799 | 0,95 | 1,225 | 1,21 | 0,875 | 1,555 | 0,73 | -0,27 | 6,6E-02 |
| P68037 | Ubiquitin-conjugating enzyme E2 L3 OS=Mus musculus OX=1009 | 16 | 3 | 5 | 3 | 154 | 17,9 | 8,51 | 3211 | | 3 | | 1,13 | 1,155 | 1,123 | 1,199 | 0,792 | 1,103 | 1,056 | 1,019 | 1,16 | 0,16 | 6,6E-02 |
| Q6PD19 | Armadillo-like helical domain-containing protein 3 OS=Mus mus | 2 | 1 | 1 | 1 | 689 | 78,6 | 6,6 | 512 | | 1 | | 0,931 | 1,108 | 1,054 | 1,191 | 0,967 | 0,788 | 1,017 | 0,763 | 1,21 | 0,21 | 6,7E-02 |
| Q3UHD1 | Adhesion G protein-coupled receptor B1 OS=Mus musculus OX= | 1 | 1 | 2 | 1 | 1582 | 173,2 | 7,56 | 395 | 2,65 | 1 | 1 | 1,041 | 0,892 | 0,839 | 0,793 | 1,155 | 0,99 | 1,047 | 0,976 | 0,86 | -0,14 | 6,7E-02 |
| Q8VDP4 | Cell cycle and apoptosis regulator protein 2 OS=Mus musculus C | 6 | 3 | 3 | 3 | 922 | 102,9 | 5,25 | 621 | 6,23 | 1 | 2 | 0,617 | 0,488 | 0,641 | 0,707 | 0,821 | 0,617 | 0,932 | 0,795 | 0,78 | -0,22 | 6,7E-02 |
| Q80TS3 | Adhesion G protein-coupled receptor L3 OS=Mus musculus OX= | 4 | 5 | 7 | 5 | 1537 | 171 | 6,71 | 2018 | 4,18 | 3 | 3 | 0,971 | 1,025 | 0,888 | 1,008 | 0,925 | 0,81 | 0,934 | 0,827 | 1,11 | 0,11 | 6,7E-02 |
| P56375 | Acylphosphatase-2 OS=Mus musculus OX=10090 GN=Acyp2 PE= | 12 | 1 | 2 | 1 | 106 | 11,9 | 9,26 | 1512 | | 1 | | 0,977 | 0,975 | 1,047 | 1,169 | 0,569 | 0,999 | 0,727 | 0,92 | 1,30 | 0,30 | 6,8E-02 |
| Q99K51 | Plastin-3 OS=Mus musculus OX=10090 GN=Pls3 PE=1 SV=3 | 3 | 2 | 2 | 2 | 630 | 70,7 | 5,62 | 398 | 0 | 1 | 1 | 1,105 | 1,199 | 0,993 | 1,047 | 1,809 | 1,263 | 1,173 | 1,41 | 0,77 | -0,23 | 6,8E-02 |
| P67778 | Prohibitin OS=Mus musculus OX=10090 GN=Phb PE=1 SV=1 | 40 | 12 | 27 | 12 | 272 | 29,8 | 5,76 | 12363 | 15,81 | 8 | 4 | 1,005 | 1,044 | 0,965 | 1,024 | 0,833 | 0,918 | 0,969 | 0,982 | 1,09 | 0,09 | 6,8E-02 |
| Q8BNY6 | Neuronal calcium sensor 1 OS=Mus musculus OX=10090 GN=Nc | 17 | 4 | 4 | 4 | 190 | 21,9 | 4,83 | 1409 | 4,68 | 2 | 2 | 1,083 | 0,991 | 1,125 | 1,065 | 0,735 | 1,028 | 0,912 | 0,972 | 1,17 | 0,17 | 6,8E-02 |
| Q8BHK2 | Sodium channel subunit beta-3 OS=Mus musculus OX=10090 GN | 10 | 2 | 2 | 2 | 215 | 24,8 | 4,74 | 1145 | | 2 | | 1,033 | 1,436 | 1,261 | 1,338 | 0,74 | 1,156 | 1,101 | 0,944 | 1,29 | 0,29 | 6,8E-02 |
| Q80UM3 | N-alpha-acetyltransferase 15, NatA auxiliary subunit OS=Mus m | 2 | 2 | 2 | 2 | 865 | 100,9 | 7,62 | 491 | 2,5 | 1 | 1 | 1,194 | 1,069 | 1,103 | 1,241 | 0,863 | 1,086 | 1,095 | 0,955 | 1,15 | 0,15 | 6,8E-02 |
| Q8VCW8 | Medium-chain acyl-CoA ligase ACSF2, mitochondrial OS=Mus m | 12 | 6 | 7 | 6 | 615 | 67,9 | 8,18 | 2725 | 2,25 | 5 | 1 | 1,039 | 1,104 | 0,947 | 1,315 | 0,681 | 0,934 | 0,861 | 1,002 | 1,27 | 0,27 | 6,8E-02 |
| Q60597 | 2-oxoglutarate dehydrogenase, mitochondrial OS=Mus musculu | 22 | 21 | 33 | 21 | 1023 | 116,4 | 6,83 | 15506 | 10,3 | 16 | 5 | 0,892 | 0,9 | 0,899 | 0,827 | 1,067 | 0,953 | 0,882 | 0,993 | 0,90 | -0,10 | 6,8E-02 |
| P19096 | Fatty acid synthase OS=Mus musculus OX=10090 GN=Fasn PE=1 | 33 | 72 | 133 | 72 | 2504 | 272,3 | 6,58 | 64730 | 68,41 | 58 | 25 | 1,001 | 1,001 | 1,094 | 0,997 | 0,878 | 0,978 | 0,977 | 0,962 | 1,08 | 0,08 | 6,8E-02 |
| Q9Z2U1 | Proteasome subunit alpha type-5 OS=Mus musculus OX=10090 | 38 | 7 | 16 | 7 | 241 | 26,4 | 4,79 | 7174 | 12,2 | 6 | 2 | 1,028 | 0,995 | 1,006 | 1,064 | 0,913 | 1,034 | 0,936 | 0,935 | 1,07 | 0,07 | 6,9E-02 |
| O09061 | Proteasome subunit beta type-1 OS=Mus musculus OX=10090 G | 30 | 6 | 10 | 6 | 240 | 26,4 | 7,81 | 3848 | 7,07 | 5 | 2 | 0,981 | 1,143 | 0,914 | 1,128 | 1,356 | 1,136 | 1,175 | 1,165 | 0,86 | -0,14 | 6,9E-02 |
| Q91V61 | Sideroflexin-3 OS=Mus musculus OX=10090 GN=Sfxn3 PE=1 SV= | 29 | 8 | 16 | 7 | 321 | 35,4 | 9,51 | 5589 | 24,25 | 5 | 4 | 0,92 | 0,918 | 0,906 | 0,917 | 1,11 | 0,921 | 0,963 | 1,038 | 0,91 | -0,09 | 6,9E-02 |
| P16045 | Galectin-1 OS=Mus musculus OX=10090 GN=Lgals1 PE=1 SV=3 | 9 | 2 | 2 | 2 | 135 | 14,9 | 5,49 | 1351 | | 2 | | 0,927 | 1,036 | 0,931 | 1,167 | 0,835 | 0,841 | 0,932 | 0,908 | 1,16 | 0,16 | 6,9E-02 |
| O88291 | DBIRD complex subunit ZNF326 OS=Mus musculus OX=10090 GI | 3 | 1 | 1 | 1 | 580 | 65,2 | 5,19 | | 2,4 | | 1 | 0,825 | 0,972 | 1,05 | 0,81 | 1,326 | 1,263 | 1,238 | 0,873 | 0,78 | -0,22 | 6,9E-02 |
| O35490 | Betaine--homocysteine S-methyltransferase 1 OS=Mus musculu | 14 | 4 | 9 | 4 | 407 | 45 | 7,9 | 3887 | 6,37 | 3 | 1 | 1,63 | 2,371 | 1,549 | 2,04 | 3,044 | 2,2 | 2,612 | 2,196 | 0,76 | -0,24 | 6,9E-02 |
| Q71M36 | Chondroitin sulfate proteoglycan 5 OS=Mus musculus OX=1009( | 12 | 4 | 4 | 1 | 566 | 60,4 | 4,54 | 2511 | | 4 | | 1,211 | 1,011 | 0,948 | 1,051 | 2,293 | 1,439 | 1,117 | 1,624 | 0,65 | -0,35 | 6,9E-02 |
| Q9D1E6 | Tubulin-folding cofactor B OS=Mus musculus OX=10090 GN=Tbc | 18 | 4 | 8 | 4 | 244 | 27,4 | 5,24 | 2626 | 4,18 | 3 | 2 | 1,182 | 1,163 | 1,541 | 1,112 | 1,427 | 1,431 | 1,5 | 1,547 | 0,85 | -0,15 | 6,9E-02 |
| P61804 | Dolichyl-diphosphooligosaccharide--protein glycosyltransferase | 9 | 1 | 1 | 1 | 113 | 12,5 | 7,08 | 417 | | 1 | | 0,882 | 0,827 | 0,956 | 1,084 | 0,555 | 0,849 | 0,852 | 0,697 | 1,27 | 0,27 | 6,9E-02 |
| P10126 | Elongation factor 1-alpha 1 OS=Mus musculus OX=10090 GN=Ee | 38 | 18 | 50 | 12 | 462 | 50,1 | 9,01 | 19517 | 45,54 | 10 | 11 | 1,023 | 0,922 | 1,05 | 0,95 | 0,734 | 0,948 | 0,904 | 0,873 | 1,14 | 0,14 | 7,0E-02 |
| Q91ZA3 | Propionyl-CoA carboxylase alpha chain, mitochondrial OS=Mus r | 14 | 8 | 9 | 8 | 724 | 79,9 | 7,25 | 3596 | 6,08 | 6 | 2 | 1,017 | 1,126 | 1,011 | 1,144 | 1,546 | 1,111 | 1,213 | 1,302 | 0,83 | -0,17 | 7,0E-02 |
| Q99M51 | Cytoplasmic protein NCK1 OS=Mus musculus OX=10090 GN=Ncl | 7 | 2 | 2 | 2 | 377 | 42,9 | 6,47 | 257 | 3,56 | 1 | 1 | 1,039 | 1,203 | 1,214 | 1,031 | 0,708 | 0,972 | 0,928 | 1,072 | 1,22 | 0,22 | 7,0E-02 |
| P02088 | Hemoglobin subunit beta-1 OS=Mus musculus OX=10090 GN=H | 54 | 8 | 30 | 8 | 147 | 15,8 | 7,65 | 14074 | 24,15 | 7 | 4 | 0,902 | 1,149 | 0,827 | 0,98 | 1,548 | 1,027 | 1,17 | 1,254 | 0,77 | -0,23 | 7,0E-02 |
| Q922F4 | Tubulin beta-6 chain OS=Mus musculus OX=10090 GN=Tubb6 PI | 28 | 13 | 257 | 1 | 447 | 50,1 | 4,89 | 106004 | 191,39 | 10 | 11 | 1,077 | 0,963 | 0,975 | 1,147 | 1,245 | 1,645 | 1,11 | 2,032 | 0,69 | -0,31 | 7,0E-02 |
| Q9CR84 | ATP synthase F(0) complex subunit C1, mitochondrial OS=Mus n | 5 | 1 | 1 | 1 | 136 | 14,2 | 10,01 | | 2,7 | | 1 | 1,255 | 0,99 | 0,924 | 0,875 | 0,77 | 0,921 | 0,807 | 0,703 | 1,26 | 0,26 | 7,1E-02 |
| Q9Z1S8 | GRB2-associated-binding protein 2 OS=Mus musculus OX=1009( | 1 | 1 | 1 | 1 | 665 | 73,2 | 8,31 | 422 | | 1 | | 1,116 | 1,055 | 1,247 | 1,069 | 1,143 | 0,88 | 0,883 | 0,904 | 1,18 | 0,18 | 7,1E-02 |
| Q9JI91 | Alpha-actinin-2 OS=Mus musculus OX=10090 GN=Actn2 PE=1 SV | 8 | 7 | 9 | 2 | 894 | 103,8 | 5,45 | 4420 | 2,18 | 6 | 1 | 1,187 | 1,279 | 1,018 | 1,205 | 1,091 | 0,959 | 1,062 | 1,035 | 1,13 | 0,13 | 7,1E-02 |
| Q9Z280 | Phospholipase D1 OS=Mus musculus OX=10090 GN=Pld1 PE=1 S | 1 | 1 | 3 | 1 | 1074 | 123,9 | 8,72 | 1187 | | 1 | | 0,732 | 0,737 | 0,584 | 0,62 | 1,7 | 0,423 | 2,092 | 2,746 | 0,38 | -0,62 | 7,2E-02 |
| Q8BTX9 | Inactive hydroxysteroid dehydrogenase-like protein 1 OS=Mus n | 11 | 4 | 6 | 4 | 330 | 36,8 | 8,48 | 3497 | 2,44 | 3 | 1 | 1,017 | 0,884 | 0,929 | 0,963 | 0,85 | 0,868 | 0,933 | 0,808 | 1,10 | 0,10 | 7,2E-02 |
| Q9CR95 | Adaptin ear-binding coat-associated protein 1 OS=Mus musculu | 9 | 2 | 3 | 2 | 275 | 29,6 | 6,38 | 1214 | 3,18 | 1 | 1 | 1,078 | 1,089 | 1,085 | 0,991 | 0,835 | 1,038 | 0,833 | 1,012 | 1,14 | 0,14 | 7,2E-02 |
| Q5RKR3 | Immunoglobulin superfamily containing leucine-rich repeat prot | 3 | 2 | 2 | 2 | 745 | 79,7 | 5,49 | 915 | | 2 | | 0,956 | 1,263 | 1,22 | 1,038 | 0,766 | 0,802 | 0,988 | 1,046 | 1,24 | 0,24 | 7,2E-02 |
| P59672 | Ankyrin repeat and SAM domain-containing protein 1A OS=Mus | 1 | 1 | 1 | 1 | 1150 | 125,2 | 6,05 | 558 | | 1 | | 0,863 | 0,933 | 1,212 | 1,3 | 1,89 | 1,455 | 1,202 | 1,347 | 0,73 | -0,27 | 7,2E-02 |
| Q8BH04 | Phosphoenolpyruvate carboxykinase [GTP], mitochondrial OS=N | 7 | 3 | 3 | 3 | 640 | 70,5 | 7,28 | 1268 | | 3 | | 0,75 | 0,976 | 0,766 | 1,06 | 1,29 | 0,988 | 1,042 | 1,114 | 0,80 | -0,20 | 7,2E-02 |
| P61961 | Ubiquitin-fold modifier 1 OS=Mus musculus OX=10090 GN=Ufm | 18 | 2 | 4 | 2 | 85 | 9,1 | 9,31 | 1557 | 2,17 | 1 | 1 | 1,006 | 0,952 | 0,891 | 0,968 | 0,779 | 0,965 | 0,855 | 0,801 | 1,12 | 0,12 | 7,3E-02 |
| Q61187 | Tumor susceptibility gene 101 protein OS=Mus musculus OX=10 | 14 | 5 | 5 | 5 | 391 | 44,1 | 6,71 | 2952 | 0 | 4 | 1 | 0,94 | 0,882 | 0,789 | 0,911 | 0,782 | 0,812 | 0,819 | 0,815 | 1,09 | 0,09 | 7,3E-02 |
| Q9D394-4 | Isoform 4 of Protein RUFY3 OS=Mus musculus OX=10090 GN=Ru | 36 | 20 | 35 | 20 | 519 | 57,4 | 5,85 | 9060 | 48,69 | 8 | 13 | 0,994 | 0,922 | 0,992 | 0,902 | 0,907 | 0,84 | 0,931 | 0,853 | 1,08 | 0,08 | 7,3E-02 |
| Q66JS6 | Eukaryotic translation initiation factor 3 subunit J-B OS=Mus mu | 15 | 5 | 10 | 5 | 263 | 29,5 | 4,81 | 3435 | 13,49 | 3 | 2 | 1,013 | 1,021 | 0,911 | 1,088 | 0,821 | 0,945 | 0,952 | 0,905 | 1,11 | 0,11 | 7,3E-02 |
| P60824 | Cold-inducible RNA-binding protein OS=Mus musculus OX=1009 | 6 | 1 | 1 | 1 | 172 | 18,6 | 9,61 | 810 | | 1 | | 1,048 | 0,923 | 0,915 | 1,184 | 0,744 | 0,759 | 1,024 | 0,715 | 1,26 | 0,26 | 7,4E-02 |
| Q6ZQ29-2 | Isoform 2 of Serine/threonine-protein kinase TAO2 OS=Mus mu | 13 | 11 | 15 | 10 | 1055 | 119,9 | 7,28 | 5257 | 17,47 | 8 | 3 | 1,002 | 0,942 | 0,937 | 0,954 | 0,773 | 0,944 | 0,883 | 0,897 | 1,10 | 0,10 | 7,4E-02 |
| Q64516-1 | Isoform 1 of Glycerol kinase OS=Mus musculus OX=10090 GN=G | 7 | 3 | 3 | 3 | 524 | 57,4 | 5,64 | 1004 | 1,92 | 2 | 1 | 1,077 | 0,998 | 1,392 | 1,38 | 0,847 | 0,802 | 1,017 | 1,107 | 1,28 | 0,28 | 7,4E-02 |
| O88485 | Cytoplasmic dynein 1 intermediate chain 1 OS=Mus musculus O | 11 | 7 | 10 | 6 | 628 | 70,7 | 5,12 | 2306 | 15,95 | 3 | 4 | 1,123 | 1,01 | 1,071 | 1,046 | 1,047 | 0,939 | 0,962 | 1,01 | 1,07 | 0,07 | 7,5E-02 |
| Q61655 | ATP-dependent RNA helicase DDX19A OS=Mus musculus OX=10 | 4 | 2 | 3 | 2 | 478 | 53,9 | 6,67 | 1549 | | 2 | | 0,986 | 0,887 | 1,022 | 0,912 | 0,654 | 0,949 | 0,641 | 0,855 | 1,23 | 0,23 | 7,5E-02 |
| Q9CQ80 | Vacuolar protein-sorting-associated protein 25 OS=Mus musculu | 11 | 2 | 4 | 2 | 176 | 20,7 | 6,34 | 2531 | | 2 | | 1,005 | 0,998 | 0,894 | 0,959 | 0,806 | 0,937 | 0,909 | 0,877 | 1,09 | 0,09 | 7,5E-02 |
| Q99KC8 | von Willebrand factor A domain-containing protein 5A OS=Mus | 3 | 2 | 2 | 2 | 793 | 87,1 | 6,58 | 567 | 2,68 | 1 | 1 | 1,075 | 1,146 | 1,011 | 1,428 | 0,986 | 0,942 | 0,981 | 0,957 | 1,21 | 0,21 | 7,5E-02 |
| Q9WVD5 | Mitochondrial ornithine transporter 1 OS=Mus musculus OX=10 | 4 | 1 | 2 | 1 | 301 | 32,8 | 8,85 | 1000 | | 1 | | 1,247 | 1,229 | 1,085 | 1,208 | 1,88 | 1,439 | 1,292 | 1,347 | 0,80 | -0,20 | 7,6E-02 |
| Q61411 | GTPase HRas OS=Mus musculus OX=10090 GN=Hras PE=1 SV=2 | 42 | 7 | 15 | 1 | 189 | 21,3 | 5,31 | 7853 | 6,08 | 7 | 1 | 1,023 | 0,869 | 1,018 | 1,039 | 0,923 | 0,914 | 0,609 | 0,789 | 1,22 | 0,22 | 7,6E-02 |
| Q9Z2U0 | Proteasome subunit alpha type-7 OS=Mus musculus OX=10090 | 33 | 8 | 14 | 8 | 248 | 27,8 | 8,46 | 6835 | 5,77 | 5 | 3 | 0,954 | 1,002 | 0,877 | 1,05 | 0,837 | 0,874 | 0,922 | 0,898 | 1,10 | 0,10 | 7,6E-02 |
| Q99J39 | Malonyl-CoA decarboxylase, mitochondrial OS=Mus musculus O | 5 | 1 | 1 | 1 | 492 | 54,7 | 8,94 | 317 | | 1 | | 0,871 | 0,99 | 0,972 | 1,371 | 1,311 | 1,482 | 1,123 | 1,704 | 0,75 | -0,25 | 7,6E-02 |
| Q9CX00 | IST1 homolog OS=Mus musculus OX=10090 GN=Ist1 PE=1 SV=1 | 2 | 1 | 3 | 1 | 362 | 39,4 | 5,44 | | 8,92 | | 1 | 1,189 | 1,163 | 1,253 | 1,161 | 0,744 | 1,175 | 1,025 | 1,03 | 1,20 | 0,20 | 7,7E-02 |
| Q8BGT8 | Phytanoyl-CoA hydroxylase-interacting protein-like OS=Mus mu | 26 | 10 | 12 | 10 | 375 | 42,3 | 6,35 | 5500 | 7,46 | 7 | 3 | 1,008 | 1,091 | 0,97 | 0,977 | 1,069 | 1,1 | 1,056 | 1,071 | 0,94 | -0,06 | 7,7E-02 |
| Q6PGE7 | Sodium-dependent proline transporter OS=Mus musculus OX=1 | 1 | 1 | 1 | 1 | 637 | 71 | 6,61 | | 0 | | 1 | 0,93 | 0,871 | 1,134 | 0,944 | 0,567 | 0,817 | 0,754 | 0,93 | 1,26 | 0,26 | 7,7E-02 |
| Q9CZ42 | ATP-dependent (S)-NAD(P)H-hydrate dehydratase OS=Mus mus | 23 | 7 | 12 | 7 | 343 | 36,7 | 7,77 | 5824 | 5,59 | 6 | 1 | 0,982 | 0,912 | 0,968 | 1,01 | 0,77 | 0,903 | 0,892 | 0,943 | 1,10 | 0,10 | 7,7E-02 |
| P46935 | E3 ubiquitin-protein ligase NEDD4 OS=Mus musculus OX=10090 | 10 | 8 | 10 | 8 | 887 | 102,6 | 5,26 | 3668 | 2,97 | 6 | 2 | 1,079 | 0,968 | 1,086 | 1,073 | 0,891 | 0,96 | 1,009 | 1,009 | 1,09 | 0,09 | 7,7E-02 |
| P56565 | Protein S100-A1 OS=Mus musculus OX=10090 GN=S100a1 PE=1 | 14 | 1 | 1 | 1 | 94 | 10,5 | 4,5 | 420 | | 1 | | 0,722 | 0,724 | 0,767 | 0,901 | 0,778 | 1,127 | 0,952 | 0,963 | 0,82 | -0,18 | 7,7E-02 |
| Q5SV85-2 | Isoform 2 of Synergin gamma OS=Mus musculus OX=10090 GN= | 8 | 7 | 7 | 7 | 1138 | 122 | 5 | 2333 | 2,51 | 6 | 1 | 1,048 | 1,01 | 1,001 | 0,995 | 0,934 | 0,81 | 1 | 0,951 | 1,10 | 0,10 | 7,7E-02 |
| B5X0G2 | Major urinary protein 17 OS=Mus musculus OX=10090 GN=Mup | 20 | 3 | 3 | 2 | 180 | 20,6 | 5,02 | 1028 | | 3 | | 0,752 | 0,834 | 0,781 | 0,871 | 1,49 | 0,804 | 1,024 | 1,163 | 0,72 | -0,28 | 7,8E-02 |

| | | | | | | | | | | | | | | | | | | | | | | | |
|---|---|---|---|---|---|---|---|---|---|---|---|---|---|---|---|---|---|---|---|---|---|---|---|
| Q91Y86 | Mitogen-activated protein kinase 8 OS=Mus musculus OX=1009 | 19 | 7 | 10 | 4 | 384 | 44,2 | 7,69 | 2518 | 17,93 | 4 | 3 | 0,929 | 0,995 | 0,911 | 1,005 | 1,004 | 1,02 | 1,152 | 1,019 | 0,92 | -0,08 | 7,8E-02 |
| Q920P3 | BMP/retinoic acid-inducible neural-specific protein 1 OS=Mus m | 6 | 4 | 5 | 4 | 760 | 88,6 | 8,97 | 1937 | 4,7 | 2 | 2 | 0,941 | 1,049 | 0,975 | 1,023 | 0,954 | 0,894 | 0,908 | 0,973 | 1,07 | 0,07 | 7,8E-02 |
| P25688 | Uricase OS=Mus musculus OX=10090 GN=Uox PE=1 SV=2 | 19 | 5 | 7 | 5 | 303 | 35 | 8,35 | 2667 | 2,29 | 4 | 1 | 1,549 | 1,801 | 1,7 | 1,964 | 2,684 | 1,775 | 2,19 | 2,118 | 0,80 | -0,20 | 7,8E-02 |
| Q6P5U7 | NACHT and WD repeat domain-containing protein 2 OS=Mus mu | 3 | 3 | 4 | 3 | 1742 | 197,3 | 6,15 | 1726 | 2,22 | 2 | 1 | 0,839 | 0,966 | 0,858 | 0,943 | 1,272 | 0,855 | 1,089 | 1,303 | 0,80 | -0,20 | 7,8E-02 |
| P61264 | Syntaxin-1B OS=Mus musculus OX=10090 GN=Stx1b PE=1 SV=1 | 48 | 16 | 41 | 14 | 288 | 33,2 | 5,38 | 15675 | 43,52 | 10 | 6 | 1,02 | 0,967 | 1,127 | 0,998 | 0,831 | 0,972 | 0,987 | 0,898 | 1,11 | 0,11 | 7,9E-02 |
| P49615 | Cyclin-dependent-like kinase 5 OS=Mus musculus OX=10090 GN | 18 | 5 | 9 | 5 | 292 | 33,3 | 7,66 | 2830 | 3,81 | 3 | 2 | 0,967 | 0,981 | 1,057 | 1,084 | 0,875 | 0,983 | 0,963 | 0,954 | 1,08 | 0,08 | 7,9E-02 |
| Q61166 | Microtubule-associated protein RP/EB family member 1 OS=Mu | 48 | 14 | 36 | 11 | 268 | 30 | 5,22 | 8922 | 72,15 | 5 | 10 | 0,976 | 0,947 | 1,092 | 1,003 | 0,822 | 0,998 | 0,911 | 0,878 | 1,11 | 0,11 | 7,9E-02 |
| Q9DCS3 | Enoyl-[acyl-carrier-protein] reductase, mitochondrial OS=Mus m | 9 | 3 | 5 | 3 | 373 | 40,3 | 9,07 | 2199 | 2,25 | 3 | 1 | 1,141 | 1,086 | 1,131 | 0,983 | 0,929 | 1,018 | 1,023 | 1,012 | 1,09 | 0,09 | 7,9E-02 |
| Q9CQM0 | Nicolin-1 OS=Mus musculus OX=10090 GN=Nicn1 PE=1 SV=1 | 8 | 2 | 2 | 2 | 213 | 24,3 | 7,08 | 1137 | | 2 | | 1,031 | 1,047 | 1,111 | 0,982 | 1,393 | 1,027 | 1,248 | 1,182 | 0,86 | -0,14 | 7,9E-02 |
| Q3V3R1 | Monofunctional C1-tetrahydrofolate synthase, mitochondrial OS | 2 | 2 | 3 | 2 | 977 | 105,7 | 7,02 | 905 | 2,46 | 1 | 1 | 0,684 | 0,78 | 0,669 | 0,561 | 1,091 | 0,704 | 1,013 | 0,764 | 0,75 | -0,25 | 8,0E-02 |
| P61458 | Pterin-4-alpha-carbinolamine dehydratase OS=Mus musculus O) | 23 | 2 | 2 | 2 | 104 | 12 | 6,8 | 567 | 2,83 | 1 | 1 | 1,037 | 1,356 | 1,305 | 1,446 | 1,82 | 1,494 | 1,514 | 1,391 | 0,83 | -0,17 | 8,0E-02 |
| Q8R0J7 | Vacuolar protein sorting-associated protein 37B OS=Mus muscu | 14 | 3 | 6 | 3 | 285 | 31 | 7,05 | 2321 | 3,48 | 3 | 1 | 0,978 | 1,011 | 1,002 | 1,164 | 1,176 | 1,031 | 1,388 | 1,457 | 0,82 | -0,18 | 8,0E-02 |
| Q8VEH3 | ADP-ribosylation factor-like protein 8A OS=Mus musculus OX=1( | 23 | 5 | 6 | 2 | 186 | 21,4 | 7,77 | 1718 | 12,08 | 1 | 4 | 0,892 | 0,896 | 1,108 | 0,876 | 0,814 | 0,804 | 0,872 | 0,795 | 1,15 | 0,15 | 8,0E-02 |
| Q9D0B6 | Protein PBDC1 OS=Mus musculus OX=10090 GN=Pbdc1 PE=1 SV | 17 | 2 | 2 | 2 | 198 | 22,2 | 4,55 | 220 | 2,63 | 1 | 1 | 1,238 | 0,953 | 1,235 | 1,05 | 0,723 | 1,043 | 0,767 | 1,022 | 1,26 | 0,26 | 8,0E-02 |
| Q7TNR6 | Immunoglobulin superfamily member 21 OS=Mus musculus OX= | 12 | 5 | 6 | 5 | 468 | 51,9 | 6,95 | 1583 | 8,23 | 2 | 3 | 1,022 | 0,918 | 0,99 | 0,996 | 0,646 | 0,97 | 0,883 | 0,823 | 1,18 | 0,18 | 8,1E-02 |
| P46096 | Synaptotagmin-1 OS=Mus musculus OX=10090 GN=Syt1 PE=1 S\ | 40 | 20 | 42 | 20 | 421 | 47,4 | 8,53 | 7924 | 81,44 | 8 | 12 | 1,018 | 1,039 | 1,032 | 1,011 | 0,781 | 0,949 | 0,995 | 0,967 | 1,11 | 0,11 | 8,1E-02 |
| Q80U40 | RIMS-binding protein 2 OS=Mus musculus OX=10090 GN=Rimbp | 1 | 1 | 1 | 1 | 1072 | 118,3 | 5,35 | 659 | | 1 | | 0,705 | 0,847 | 0,766 | 0,795 | 0,945 | 0,775 | 1,401 | 1,551 | 0,67 | -0,33 | 8,1E-02 |
| P62301 | 40S ribosomal protein S13 OS=Mus musculus OX=10090 GN=Rp: | 35 | 6 | 8 | 6 | 151 | 17,2 | 10,54 | 1545 | 15,69 | 2 | 4 | 1,051 | 0,895 | 1,161 | 1 | 1,306 | 1,184 | 1,038 | 1,246 | 0,86 | -0,14 | 8,2E-02 |
| P48428 | Tubulin-specific chaperone A OS=Mus musculus OX=10090 GN= | 45 | 8 | 9 | 8 | 108 | 12,8 | 5,27 | 3015 | 10,59 | 4 | 4 | 1,042 | 1,178 | 1,042 | 1,317 | 0,749 | 1,047 | 1,015 | 0,98 | 1,21 | 0,21 | 8,2E-02 |
| E9Q401 | Ryanodine receptor 2 OS=Mus musculus OX=10090 GN=Ryr2 PE | 0 | 2 | 2 | 1 | 4966 | 564,5 | 6,09 | 568 | | 2 | | 0,819 | 0,965 | 0,839 | 0,782 | 1,264 | 0,889 | 0,917 | 1,232 | 0,79 | -0,21 | 8,2E-02 |
| Q60870 | Receptor expression-enhancing protein 5 OS=Mus musculus OX | 11 | 3 | 4 | 3 | 185 | 21 | 8,38 | 1051 | 5,92 | 1 | 2 | 1,101 | 1,011 | 1,155 | 0,953 | 0,748 | 1,034 | 0,915 | 0,907 | 1,17 | 0,17 | 8,3E-02 |
| B1AWN6 | Sodium channel protein type 2 subunit alpha OS=Mus musculus | 5 | 10 | 15 | 10 | 2006 | 227,8 | 5,67 | 4425 | 18,62 | 6 | 4 | 1,025 | 0,961 | 1,137 | 1,012 | 1,004 | 0,906 | 0,956 | 0,902 | 1,10 | 0,10 | 8,3E-02 |
| P62812 | Gamma-aminobutyric acid receptor subunit alpha-1 OS=Mus mu | 7 | 3 | 3 | 2 | 455 | 51,7 | 9,31 | 1707 | | 3 | | 1,211 | 0,915 | 1,022 | 0,953 | 0,718 | 0,917 | 0,93 | 0,858 | 1,20 | 0,20 | 8,3E-02 |
| P29758 | Ornithine aminotransferase, mitochondrial OS=Mus musculus O | 24 | 8 | 10 | 8 | 439 | 48,3 | 6,62 | 2593 | 17,78 | 3 | 5 | 0,955 | 0,947 | 1,043 | 1,125 | 0,864 | 1 | 0,826 | 0,913 | 1,13 | 0,13 | 8,3E-02 |
| Q7TMB8 | Cytoplasmic FMR1-interacting protein 1 OS=Mus musculus OX= | 12 | 15 | 25 | 1 | 1253 | 145,1 | 6,9 | 9965 | 7,69 | 11 | 4 | 0,946 | 1,005 | 0,889 | 0,984 | 1,333 | 1,015 | 1,019 | 1,11 | 0,85 | -0,15 | 8,4E-02 |
| O88587 | Catechol O-methyltransferase OS=Mus musculus OX=10090 GN= | 8 | 3 | 5 | 3 | 265 | 29,5 | 5,83 | 1547 | 0 | 2 | 1 | 1,015 | 1,121 | 0,867 | 1,012 | 1,286 | 1,006 | 1,159 | 1,435 | 0,82 | -0,18 | 8,4E-02 |
| Q80VC9-5 | Isoform 5 of Calmodulin-regulated spectrin-associated protein 3 | 14 | 13 | 16 | 1 | 1279 | 138,2 | 8,54 | 6635 | 9,17 | 9 | 4 | 1,069 | 0,951 | 1,052 | 0,978 | 0,969 | 0,981 | 0,844 | 0,762 | 1,14 | 0,14 | 8,4E-02 |
| Q9CQX2 | Cytochrome b5 type B OS=Mus musculus OX=10090 GN=Cyb5b | 20 | 3 | 4 | 2 | 146 | 16,3 | 4,89 | 2185 | 0 | 2 | 1 | 1,289 | 1,366 | 1,186 | 1,23 | 0,838 | 1,066 | 1,289 | 1,052 | 1,19 | 0,19 | 8,5E-02 |
| Q8CAE9 | Podocalyxin-like protein 2 OS=Mus musculus OX=10090 GN=Poc | 5 | 2 | 2 | 2 | 603 | 64,9 | 4,36 | 327 | 0 | 1 | 1 | 0,889 | 0,781 | 0,856 | 1,075 | 1,188 | 1,093 | 0,968 | 1,007 | 0,85 | -0,15 | 8,5E-02 |
| Q3UUJ4 | STE20-related kinase adapter protein alpha OS=Mus musculus C | 4 | 2 | 2 | 2 | 431 | 48,2 | 6,43 | 1116 | | 2 | | 1,238 | 1,184 | 1,036 | 1,365 | 0,823 | 1,139 | 1,055 | 1,019 | 1,19 | 0,19 | 8,5E-02 |
| P16546 | Spectrin alpha chain, non-erythrocytic 1 OS=Mus musculus OX= | 50 | 134 | 292 | 134 | 2472 | 284,4 | 5,33 | 116964 | 304,56 | 85 | 65 | 1,022 | 0,982 | 0,986 | 0,994 | 0,864 | 0,969 | 0,977 | 0,948 | 1,06 | 0,06 | 8,5E-02 |
| Q9QXZ0 | Microtubule-actin cross-linking factor 1 OS=Mus musculus OX=1 | 14 | 92 | 127 | 91 | 7354 | 831,4 | 5,43 | 45333 | 132,44 | 55 | 43 | 1,005 | 0,982 | 0,969 | 0,97 | 0,883 | 0,953 | 0,972 | 0,945 | 1,05 | 0,05 | 8,5E-02 |
| Q8VBW6 | NEDD8-activating enzyme E1 regulatory subunit OS=Mus muscu | 3 | 1 | 4 | 1 | 534 | 60,2 | 5,52 | 2428 | | 1 | | 0,894 | 0,999 | 0,899 | 0,984 | 1,218 | 0,9 | 1,14 | 1,125 | 0,86 | -0,14 | 8,5E-02 |
| P47934 | Carnitine O-acetyltransferase OS=Mus musculus OX=10090 GN= | 3 | 2 | 2 | 2 | 626 | 70,8 | 8,44 | 470 | 2,36 | 1 | 1 | 1,103 | 1,08 | 1,09 | 1,055 | 1 | 0,958 | 1,093 | 1,029 | 1,06 | 0,06 | 8,5E-02 |
| Q64331 | Unconventional myosin-VI OS=Mus musculus OX=10090 GN=My | 5 | 4 | 7 | 4 | 1265 | 146,3 | 8,85 | 2163 | 7,63 | 2 | 2 | 1,003 | 0,939 | 1,186 | 0,955 | 0,816 | 0,833 | 0,949 | 0,938 | 1,15 | 0,15 | 8,5E-02 |
| Q9WVA4 | Transgelin-2 OS=Mus musculus OX=10090 GN=Tagln2 PE=1 SV= | 11 | 2 | 2 | 1 | 199 | 22,4 | 8,24 | 694 | 2,36 | 1 | 1 | 1,07 | 0,977 | 1,132 | 1,158 | 1,007 | 0,869 | 1,06 | 0,782 | 1,17 | 0,17 | 8,5E-02 |
| Q9R1J0 | Sterol-4-alpha-carboxylate 3-dehydrogenase, decarboxylating O | 16 | 5 | 8 | 5 | 362 | 40,7 | 7,85 | 4199 | | 5 | | 1,162 | 1,091 | 1,153 | 1,08 | 0,751 | 1,121 | 0,816 | 1,046 | 1,20 | 0,20 | 8,5E-02 |
| Q80TL4 | PHD finger protein 24 OS=Mus musculus OX=10090 GN=Phf24 P | 20 | 9 | 13 | 9 | 400 | 45,2 | 5,77 | 7013 | 2,54 | 9 | 1 | 0,948 | 1,052 | 1,017 | 1,032 | 0,955 | 0,951 | 0,997 | 0,906 | 1,06 | 0,06 | 8,6E-02 |
| Q7TN79 | A-kinase anchor protein 7 isoform gamma OS=Mus musculus O) | 7 | 4 | 9 | 4 | 314 | 35,5 | 7,87 | 2967 | 10,72 | 2 | 2 | 0,992 | 1,039 | 1,1 | 0,938 | 0,848 | 0,988 | 0,959 | 0,89 | 1,10 | 0,10 | 8,7E-02 |
| Q9QXD6 | Fructose-1,6-bisphosphatase 1 OS=Mus musculus OX=10090 GN | 14 | 3 | 4 | 3 | 338 | 36,9 | 6,57 | 1954 | 2,06 | 2 | 1 | 1,241 | 1,485 | 1,284 | 1,394 | 2,613 | 1,552 | 1,538 | 1,814 | 0,72 | -0,28 | 8,7E-02 |
| Q8BIK4 | Dedicator of cytokinesis protein 9 OS=Mus musculus OX=10090 | 0 | 1 | 1 | 1 | 2055 | 235,2 | 7,25 | 559 | | 1 | | 0,963 | 0,721 | 0,902 | 0,897 | 0,822 | 0,785 | 0,689 | 0,65 | 1,18 | 0,18 | 8,8E-02 |
| Q8R1B5 | Complexin-3 OS=Mus musculus OX=10090 GN=Cplx3 PE=1 SV=1 | 6 | 1 | 2 | 1 | 158 | 17,6 | 4,89 | 1565 | | 1 | | 1,142 | 0,879 | 0,951 | 0,925 | 0,956 | 1,286 | 1,404 | 1,879 | 0,71 | -0,29 | 8,8E-02 |
| Q8JZN5 | Complex I assembly factor ACAD9, mitochondrial OS=Mus musc | 7 | 4 | 4 | 4 | 625 | 68,7 | 7,46 | 2737 | | 4 | | 0,951 | 1,03 | 1,125 | 1,071 | 1,043 | 1,213 | 1,257 | 1,147 | 0,90 | -0,10 | 8,8E-02 |
| E9Q3L2 | Phosphatidylinositol 4-kinase alpha OS=Mus musculus OX=1009 | 3 | 6 | 6 | 6 | 2105 | 236,9 | 7,06 | 2986 | 2,25 | 4 | 2 | 1,05 | 1,077 | 1,193 | 1,038 | 1,283 | 1,148 | 1,133 | 1,193 | 0,92 | -0,08 | 8,8E-02 |
| Q9DBS1 | Transmembrane protein 43 OS=Mus musculus OX=10090 GN=Tr | 6 | 2 | 4 | 2 | 400 | 44,8 | 7,36 | 2334 | | 2 | | 0,95 | 0,991 | 0,874 | 0,92 | 1,127 | 1,064 | 0,94 | 0,993 | 0,91 | -0,09 | 8,8E-02 |
| Q8BG40 | Katanin p80 WD40 repeat-containing subunit B1 OS=Mus muscu | 1 | 1 | 1 | 1 | 658 | 72,6 | 7,27 | 600 | | 1 | | 0,915 | 0,76 | 0,913 | 0,809 | 0,591 | 0,758 | 0,632 | 0,846 | 1,20 | 0,20 | 8,8E-02 |
| O88935 | Synapsin-1 OS=Mus musculus OX=10090 GN=Syn1 PE=1 SV=2 | 56 | 25 | 68 | 21 | 706 | 74,1 | 9,8 | 31671 | 42,14 | 18 | 11 | 0,975 | 1,016 | 0,977 | 0,981 | 1,126 | 1,013 | 1,006 | 1,041 | 0,94 | -0,06 | 8,9E-02 |
| P48722 | Heat shock 70 kDa protein 4L OS=Mus musculus OX=10090 GN= | 21 | 16 | 19 | 15 | 838 | 94,3 | 5,74 | 7372 | 9,33 | 11 | 5 | 0,985 | 0,938 | 1,051 | 0,95 | 1,111 | 1,034 | 1,023 | 1,024 | 0,94 | -0,06 | 8,9E-02 |
| Q01730 | Ras suppressor protein 1 OS=Mus musculus OX=10090 GN=Rsu1 | 9 | 2 | 2 | 2 | 277 | 31,5 | 8,88 | 1140 | | 2 | | 1,058 | 1,371 | 1,029 | 1,035 | 0,943 | 0,898 | 1,047 | 0,855 | 1,20 | 0,20 | 8,9E-02 |
| Q9ES97 | Reticulon-3 OS=Mus musculus OX=10090 GN=Rtn3 PE=1 SV=2 | 10 | 11 | 18 | 11 | 964 | 103,8 | 4,92 | 9826 | 7,58 | 10 | 1 | 1,1 | 1,055 | 0,92 | 1,118 | 0,783 | 0,878 | 1,011 | 0,972 | 1,15 | 0,15 | 9,0E-02 |
| Q9D824 | Pre-mRNA 3'-end-processing factor FIP1 OS=Mus musculus OX= | 3 | 1 | 1 | 1 | 581 | 64,9 | 5,77 | 328 | | 1 | | 0,932 | 0,628 | 1,105 | 0,693 | 1,508 | 1,025 | 1,655 | 0,898 | 0,66 | -0,34 | 9,0E-02 |
| Q9D8Y0 | EF-hand domain-containing protein D2 OS=Mus musculus OX=1( | 15 | 4 | 5 | 4 | 240 | 26,8 | 5,06 | 198 | 10,4 | 1 | 3 | 0,935 | 0,894 | 1,02 | 0,952 | 0,819 | 0,975 | 0,804 | 0,807 | 1,12 | 0,12 | 9,0E-02 |
| P32883 | GTPase KRas OS=Mus musculus OX=10090 GN=Kras PE=1 SV=1 | 47 | 8 | 11 | 3 | 189 | 21,6 | 6,77 | 5329 | 2,94 | 7 | 1 | 0,796 | 0,909 | 0,978 | 0,734 | 1,195 | 0,936 | 1,026 | 0,928 | 0,84 | -0,16 | 9,0E-02 |
| Q6ZPJ3 | (E3-independent) E2 ubiquitin-conjugating enzyme UBE2O OS=N | 15 | 15 | 21 | 14 | 1288 | 140,7 | 5,06 | 9155 | 12,89 | 11 | 5 | 0,98 | 0,941 | 0,978 | 1,005 | 0,997 | 1,012 | 1,016 | 0,994 | 0,97 | -0,03 | 9,0E-02 |
| Q6ZQA6 | Immunoglobulin superfamily member 3 OS=Mus musculus OX= | 13 | 13 | 19 | 13 | 1194 | 134,6 | 6,05 | 6899 | 17,17 | 8 | 6 | 0,943 | 0,966 | 0,932 | 0,979 | 1,223 | 0,965 | 1,011 | 1,083 | 0,89 | -0,11 | 9,1E-02 |
| Q9DBC3 | Cap-specific mRNA (nucleoside-2'-O-)-methyltransferase 1 OS=N | 1 | 1 | 1 | 1 | 837 | 95,6 | 7,27 | 220 | | 1 | | 0,937 | 0,991 | 0,869 | 1,033 | 1,612 | 1,253 | 0,978 | 1,118 | 0,77 | -0,23 | 9,1E-02 |
| Q9ET01 | Glycogen phosphorylase, liver form OS=Mus musculus OX=1009 | 5 | 4 | 10 | 1 | 850 | 97,4 | 7,09 | 3095 | 11,37 | 2 | 3 | 1,427 | 1,496 | 1,364 | 1,123 | 1,83 | 1,616 | 1,768 | 1,32 | 0,83 | -0,17 | 9,1E-02 |
| Q4VAA2 | Protein CDV3 OS=Mus musculus OX=10090 GN=Cdv3 PE=1 SV=2 | 21 | 3 | 3 | 3 | 281 | 29,7 | 6,1 | 1112 | 2,73 | 2 | 1 | 0,87 | 1,179 | 0,93 | 0,956 | 0,6 | 0,889 | 0,928 | 0,678 | 1,27 | 0,27 | 9,1E-02 |
| P49722 | Proteasome subunit alpha type-2 OS=Mus musculus OX=10090 | 21 | 7 | 12 | 7 | 234 | 25,9 | 7,43 | 5681 | 6,65 | 7 | 1 | 1,051 | 0,931 | 0,938 | 0,97 | 0,759 | 0,994 | 0,873 | 0,781 | 1,14 | 0,14 | 9,1E-02 |
| E9Q4S1 | High affinity cAMP-specific and IBMX-insensitive 3',5'-cyclic pho | 1 | 1 | 1 | 1 | 865 | 96,7 | 6,86 | 383 | | 1 | | 1,119 | 0,928 | 1,05 | 1,437 | 1,81 | 1,092 | 1,457 | 1,747 | 0,74 | -0,26 | 9,3E-02 |
| P47740 | Aldehyde dehydrogenase family 3 member A2 OS=Mus musculu | 7 | 3 | 4 | 3 | 484 | 53,9 | 8,35 | 665 | 9,22 | 1 | 3 | 1,066 | 1,053 | 1,005 | 1,182 | 1,348 | 1,151 | 1,083 | 1,383 | 0,87 | -0,13 | 9,3E-02 |
| Q9JMD3 | START domain-containing protein 10 OS=Mus musculus OX=100 | 11 | 2 | 4 | 2 | 291 | 32,9 | 7,12 | 839 | 6,25 | 1 | 2 | 5,151 | 6,19 | 4,799 | 6,234 | 12,709 | 6,816 | 7,492 | 6,983 | 0,66 | -0,34 | 9,3E-02 |
| Q9Z0P4 | Paralemmin-1 OS=Mus musculus OX=10090 GN=Palm PE=1 SV= | 47 | 17 | 33 | 16 | 383 | 41,6 | 4,84 | 5587 | 53,3 | 6 | 12 | 1,039 | 0,956 | 1,003 | 1,042 | 0,682 | 1,012 | 0,917 | 0,852 | 1,17 | 0,17 | 9,3E-02 |
| Q922U2 | Keratin, type II cytoskeletal 5 OS=Mus musculus OX=10090 GN= | 21 | 20 | 35 | 9 | 580 | 61,7 | 7,75 | 5872 | 44,17 | 5 | 16 | 0,956 | 1,045 | 0,725 | 0,751 | 1,38 | 1,422 | 1,011 | 0,89 | 0,74 | -0,26 | 9,4E-02 |
| Q8C7X2 | ER membrane protein complex subunit 1 OS=Mus musculus OX= | 7 | 5 | 5 | 5 | 997 | 111,5 | 7,43 | 3110 | | 5 | | 1 | 0,908 | 0,901 | 0,933 | 0,942 | 1,003 | 1,054 | 0,999 | 0,94 | -0,06 | 9,4E-02 |

| | | | | | | | | | | | | | | | | | | | | | | | |
|---|---|---|---|---|---|---|---|---|---|---|---|---|---|---|---|---|---|---|---|---|---|---|---|
| P15209 | BDNF/NT-3 growth factors receptor OS=Mus musculus OX=1009 | 9 | 7 | 10 | 2 | 821 | 92,1 | 6,55 | 3829 | 5,32 | 4 | 3 | 0,976 | 0,952 | 1,111 | 0,993 | 1,105 | 1,154 | 1,135 | 1,013 | 0,91 | -0,09 | 9,4E-02 |
| P62911 | 60S ribosomal protein L32 OS=Mus musculus OX=10090 GN=Rpl | 21 | 3 | 3 | 3 | 135 | 15,9 | 11,33 | 1232 | 2 | 2 | 1 | 0,901 | 1,018 | 1,145 | 0,959 | 1,179 | 0,988 | 1,231 | 1,331 | 0,85 | -0,15 | 9,4E-02 |
| O70228 | Probable phospholipid-transporting ATPase IIA OS=Mus muscul | 3 | 2 | 5 | 2 | 1047 | 118,5 | 7,69 | | 16,62 | | 2 | 0,96 | 0,967 | 0,915 | 0,86 | 0,928 | 1,08 | 1,01 | 0,999 | 0,92 | -0,08 | 9,5E-02 |
| Q9CQZ1 | Heat shock factor-binding protein 1 OS=Mus musculus OX=1009 | 24 | 2 | 2 | 2 | 76 | 8,6 | 4,31 | 1484 | | 2 | | 1,008 | 1,212 | 1,096 | 1,034 | 1,031 | 1,03 | 0,947 | 0,901 | 1,11 | 0,11 | 9,5E-02 |
| Q6P5F6 | Zinc transporter ZIP10 OS=Mus musculus OX=10090 GN=Slc39a1 | 1 | 1 | 2 | 1 | 833 | 94,3 | 6,71 | 299 | 1,95 | 1 | 1 | 1,145 | 1,529 | 0,948 | 1,031 | 0,666 | 1,007 | 0,988 | 0,784 | 1,35 | 0,35 | 9,5E-02 |
| Q9D6Y7 | Mitochondrial peptide methionine sulfoxide reductase OS=Mus | 11 | 2 | 3 | 2 | 233 | 26 | 8,41 | 1992 | | 2 | | 1,395 | 1,275 | 1,612 | 1,24 | 1,681 | 1,792 | 1,347 | 1,734 | 0,84 | -0,16 | 9,5E-02 |
| Q78JT3 | 3-hydroxyanthranilate 3,4-dioxygenase OS=Mus musculus OX=1 | 5 | 1 | 4 | 1 | 286 | 32,8 | 6,54 | | 8,53 | | 1 | 0,896 | 1,068 | 0,894 | 0,994 | 1,646 | 1,014 | 1,301 | 1,072 | 0,77 | -0,23 | 9,5E-02 |
| Q3U182 | CREB-regulated transcription coactivator 2 OS=Mus musculus O | 1 | 1 | 1 | 1 | 692 | 73,2 | 7,01 | 510 | | 1 | | 0,651 | 0,694 | 0,515 | 1,206 | 1,378 | 3,586 | 0,691 | 2,666 | 0,37 | -0,63 | 9,6E-02 |
| O70479 | BTB/POZ domain-containing adapter for CUL3-mediated RhoA d | 2 | 1 | 1 | 1 | 316 | 36,1 | 7,84 | 409 | | 1 | | 0,902 | 0,941 | 0,896 | 1,072 | 0,795 | 0,538 | 0,963 | 0,752 | 1,25 | 0,25 | 9,6E-02 |
| Q6IRU5-2 | Isoform 2 of Clathrin light chain B OS=Mus musculus OX=10090 | 31 | 9 | 15 | 9 | 211 | 23,2 | 4,68 | 2123 | 30,84 | 3 | 6 | 1,112 | 1,005 | 1,093 | 0,983 | 0,71 | 1,046 | 0,92 | 0,913 | 1,17 | 0,17 | 9,6E-02 |
| Q8C729 | Protein FAM126B OS=Mus musculus OX=10090 GN=Fam126b PE | 1 | 1 | 1 | 1 | 530 | 58,6 | 7,72 | 380 | | 1 | | 1,086 | 1,154 | 1,005 | 1,129 | 0,839 | 1,1 | 0,89 | 1,012 | 1,14 | 0,14 | 9,6E-02 |
| Q9JI19 | Acidic fibroblast growth factor intracellular-binding protein OS= | 5 | 2 | 2 | 2 | 357 | 41,2 | 6,76 | 326 | 2,35 | 1 | 1 | 0,631 | 0,888 | 0,586 | 0,696 | 1,187 | 0,735 | 0,871 | 0,922 | 0,75 | -0,25 | 9,6E-02 |
| P70388 | DNA repair protein RAD50 OS=Mus musculus OX=10090 GN=Ra | 1 | 1 | 1 | 1 | 1312 | 153,4 | 6,95 | 361 | | 1 | | 0,942 | 0,877 | 1,238 | 0,911 | 0,512 | 0,89 | 0,818 | 0,815 | 1,31 | 0,31 | 9,6E-02 |
| Q8VHH5 | Arf-GAP with GTPase, ANK repeat and PH domain-containing pr | 11 | 9 | 12 | 6 | 910 | 97,9 | 7,75 | 5259 | 11,08 | 7 | 3 | 1,175 | 0,907 | 1,047 | 0,925 | 0,595 | 1,009 | 0,835 | 0,783 | 1,26 | 0,26 | 9,6E-02 |
| O35954 | Membrane-associated phosphatidylinositol transfer protein 1 O | 6 | 6 | 6 | 6 | 1243 | 134,9 | 6,06 | 1080 | 6,92 | 3 | 3 | 0,996 | 0,98 | 1,077 | 0,954 | 1,189 | 0,985 | 1,083 | 1,195 | 0,90 | -0,10 | 9,6E-02 |
| Q91ZT5 | FYVE, RhoGEF and PH domain-containing protein 4 OS=Mus mu | 5 | 3 | 6 | 3 | 766 | 86,5 | 5,95 | 2972 | | 3 | | 1,012 | 1,1 | 1,087 | 1,203 | 0,967 | 1,102 | 0,942 | 0,971 | 1,11 | 0,11 | 9,7E-02 |
| Q8CHW4 | Translation initiation factor eIF-2B subunit epsilon OS=Mus mus | 1 | 1 | 1 | 1 | 717 | 80 | 5,07 | 572 | | 1 | | 1,035 | 0,924 | 1,028 | 1,104 | 0,731 | 1,063 | 0,877 | 0,754 | 1,19 | 0,19 | 9,7E-02 |
| Q9CPQ8 | ATP synthase subunit g, mitochondrial OS=Mus musculus OX=10 | 25 | 2 | 3 | 2 | 103 | 11,4 | 9,74 | 1060 | 3,27 | 1 | 1 | 1,132 | 1,119 | 1,086 | 1,169 | 0,766 | 1,066 | 1,084 | 0,998 | 1,15 | 0,15 | 9,7E-02 |
| Q9ER00 | Syntaxin-12 OS=Mus musculus OX=10090 GN=Stx12 PE=1 SV=1 | 33 | 7 | 10 | 7 | 274 | 31,2 | 5,44 | 2235 | 21,26 | 3 | 5 | 1,104 | 1,004 | 1,046 | 1,1 | 0,925 | 0,985 | 1,032 | 1,036 | 1,07 | 0,07 | 9,8E-02 |
| P62838 | Ubiquitin-conjugating enzyme E2 D2 OS=Mus musculus OX=100 | 24 | 3 | 3 | 2 | 147 | 16,7 | 7,83 | 1093 | 3,8 | 2 | 1 | 0,936 | 0,961 | 1,003 | 1,008 | 0,51 | 0,997 | 0,871 | 0,681 | 1,28 | 0,28 | 9,8E-02 |
| P34914 | Bifunctional epoxide hydrolase 2 OS=Mus musculus OX=10090 G | 6 | 2 | 2 | 2 | 554 | 62,5 | 6,19 | 1331 | | 2 | | 1,394 | 1,241 | 1,492 | 1,267 | 1,953 | 1,338 | 1,553 | 1,65 | 0,83 | -0,17 | 9,8E-02 |
| Q7SIG6 | Arf-GAP with SH3 domain, ANK repeat and PH domain-containir | 4 | 3 | 4 | 3 | 958 | 106,7 | 6,65 | 1367 | 2,94 | 3 | 1 | 0,877 | 1,243 | 0,72 | 0,739 | 0,971 | 1,462 | 1,129 | 1,27 | 0,74 | -0,26 | 9,8E-02 |
| Q8BP92 | Reticulocalbin-2 OS=Mus musculus OX=10090 GN=Rcn2 PE=1 SV | 6 | 2 | 2 | 2 | 320 | 37,2 | 4,42 | 430 | 2 | 1 | 1 | 0,729 | 0,477 | 1,201 | 0,378 | 2,543 | 1,095 | 1,473 | 0,892 | 0,46 | -0,54 | 9,8E-02 |
| O70133 | ATP-dependent RNA helicase A OS=Mus musculus OX=10090 GN | 10 | 14 | 14 | 14 | 1380 | 149,4 | 6,83 | 6826 | 8,07 | 11 | 3 | 0,854 | 0,804 | 0,978 | 1,041 | 1,213 | 0,934 | 1,112 | 1,045 | 0,85 | -0,15 | 9,8E-02 |
| Q6IME9 | Keratin, type II cytoskeletal 72 OS=Mus musculus OX=10090 GN | 10 | 7 | 10 | 1 | 520 | 56,7 | 7,4 | 2519 | 8,08 | 4 | 4 | 0,672 | 1,028 | 0,501 | 0,625 | 1,187 | 1,068 | 0,889 | 0,799 | 0,72 | -0,28 | 9,8E-02 |
| P63141 | Potassium voltage-gated channel subfamily A member 2 OS=Mu | 3 | 1 | 1 | 1 | 499 | 56,7 | 4,86 | 312 | | 1 | | 1,084 | 1,22 | 0,94 | 1,326 | 1,408 | 1,384 | 1,214 | 1,303 | 0,86 | -0,14 | 9,8E-02 |
| Q99JP7 | Glutathione hydrolase 7 OS=Mus musculus OX=10090 GN=Ggt7 | 7 | 4 | 7 | 4 | 662 | 70,2 | 5,06 | 4814 | | 4 | | 1,093 | 1,052 | 0,909 | 0,93 | 1,393 | 0,955 | 1,209 | 1,214 | 0,84 | -0,16 | 9,9E-02 |
| Q8CBW3 | Abl interactor 1 OS=Mus musculus OX=10090 GN=Abi1 PE=1 SV= | 15 | 7 | 10 | 4 | 481 | 52,3 | 7,64 | 3101 | 18,07 | 5 | 3 | 1,024 | 1,003 | 0,96 | 1,025 | 0,813 | 0,961 | 0,99 | 0,923 | 1,09 | 0,09 | 9,9E-02 |
| Q8C145 | Zinc transporter ZIP6 OS=Mus musculus OX=10090 GN=Slc39a6 | 3 | 2 | 3 | 2 | 765 | 86,3 | 6,84 | 2225 | | 2 | | 1,095 | 1,277 | 1,165 | 1,195 | 0,969 | 0,941 | 1,091 | 1,192 | 1,13 | 0,13 | 9,9E-02 |
| Q811D0 | Disks large homolog 1 OS=Mus musculus OX=10090 GN=Dlg1 PE | 14 | 11 | 13 | 5 | 905 | 100,1 | 5,8 | 4082 | 4,77 | 8 | 3 | 0,885 | 0,846 | 0,827 | 0,896 | 0,63 | 0,884 | 0,793 | 0,695 | 1,15 | 0,15 | 9,9E-02 |
| Q6PIE5 | Sodium/potassium-transporting ATPase subunit alpha-2 OS=Mu | 38 | 43 | 181 | 24 | 1020 | 112,1 | 5,55 | 84854 | 127,88 | 29 | 21 | 1,034 | 0,958 | 1,032 | 0,976 | 0,781 | 0,966 | 0,933 | 0,953 | 1,10 | 0,10 | 1,0E-01 |
| P53986 | Monocarboxylate transporter 1 OS=Mus musculus OX=10090 GI | 7 | 2 | 3 | 2 | 493 | 53,2 | 7,47 | 1674 | | 2 | | 0,913 | 1,124 | 0,978 | 0,959 | 0,617 | 0,727 | 1,028 | 0,834 | 1,24 | 0,24 | 1,0E-01 |
| Q9CRA4 | Methylsterol monooxygenase 1 OS=Mus musculus OX=10090 GI | 5 | 1 | 1 | 1 | 293 | 34,8 | 7,66 | 397 | | 1 | | 0,769 | 0,979 | 0,817 | 0,831 | 1,099 | 0,794 | 1,062 | 1,159 | 0,83 | -0,17 | 1,0E-01 |
| Q61176 | Arginase-1 OS=Mus musculus OX=10090 GN=Arg1 PE=1 SV=1 | 28 | 6 | 10 | 6 | 323 | 34,8 | 7,01 | 5845 | | 6 | | 0,971 | 1,372 | 1,06 | 0,99 | 1,593 | 1,202 | 1,359 | 1,235 | 0,82 | -0,18 | 1,0E-01 |
| Q8C0C7 | Phenylalanine--tRNA ligase alpha subunit OS=Mus musculus OX= | 9 | 5 | 7 | 5 | 508 | 57,6 | 8,28 | 3148 | 2,69 | 4 | 1 | 0,964 | 0,954 | 1,049 | 1,117 | 1,137 | 1,13 | 1,047 | 1,107 | 0,92 | -0,08 | 1,0E-01 |
| Q8R2X8 | Golgin-45 OS=Mus musculus OX=10090 GN=Blzf1 PE=1 SV=2 | 3 | 1 | 3 | 1 | 403 | 45,5 | 7,59 | 663 | | 1 | | 1,035 | 1,036 | 0,868 | 0,883 | 1,835 | 1,101 | 1,094 | 1,208 | 0,73 | -0,27 | 1,0E-01 |
| Q8K0L9 | Zinc finger and BTB domain-containing protein 20 OS=Mus musc | 1 | 1 | 1 | 1 | 741 | 81 | 6,55 | 475 | | 1 | | 0,922 | 0,912 | 1,065 | 0,976 | 0,93 | 0,879 | 0,898 | 0,728 | 1,13 | 0,13 | 1,0E-01 |
| P46638 | Ras-related protein Rab-11B OS=Mus musculus OX=10090 GN=F | 40 | 10 | 18 | 10 | 218 | 24,5 | 5,94 | 5774 | 18,93 | 7 | 3 | 1,092 | 0,969 | 0,988 | 1,07 | 0,872 | 0,99 | 0,958 | 0,985 | 1,08 | 0,08 | 1,0E-01 |
| Q8JZL3 | Thiamine-triphosphatase OS=Mus musculus OX=10090 GN=Thtp | 17 | 4 | 4 | 4 | 224 | 24,2 | 4,72 | 2148 | | 4 | | 0,828 | 1,092 | 0,951 | 1 | 1,158 | 0,988 | 1,535 | 1,172 | 0,80 | -0,20 | 1,0E-01 |
| Q9CQU5 | ZW10 interactor OS=Mus musculus OX=10090 GN=Zwint PE=1 S | 3 | 1 | 1 | 1 | 252 | 28,7 | 8,43 | | 2,29 | | 1 | 0,955 | 1,125 | 0,978 | 0,93 | 0,674 | 0,618 | 0,89 | 1,016 | 1,25 | 0,25 | 1,0E-01 |
| P00688 | Pancreatic alpha-amylase OS=Mus musculus OX=10090 GN=Am | 2 | 1 | 2 | 1 | 508 | 57,3 | 7,28 | 1003 | | 1 | | 0,598 | 0,6 | 0,536 | 0,531 | 0,648 | 0,836 | 0,535 | 0,854 | 0,79 | -0,21 | 1,0E-01 |
| Q61171 | Peroxiredoxin-2 OS=Mus musculus OX=10090 GN=Prdx2 PE=1 S | 35 | 7 | 31 | 6 | 198 | 21,8 | 5,41 | 9171 | 55,3 | 6 | 4 | 1,05 | 1,013 | 1,041 | 1,097 | 0,766 | 1,094 | 0,908 | 0,898 | 1,15 | 0,15 | 1,0E-01 |
| Q6PHN9 | Ras-related protein Rab-35 OS=Mus musculus OX=10090 GN=Ra | 40 | 10 | 15 | 9 | 201 | 23 | 8,29 | 6986 | 6,01 | 9 | 2 | 1,031 | 0,937 | 1,177 | 0,988 | 0,875 | 0,995 | 0,912 | 0,909 | 1,12 | 0,12 | 1,0E-01 |
| Q80W47 | WD repeat domain phosphoinositide-interacting protein 2 OS=M | 4 | 2 | 2 | 2 | 445 | 48,4 | 5,86 | 667 | 0 | 1 | 1 | 1,012 | 0,949 | 1,224 | 1,092 | 0,698 | 1,075 | 0,935 | 0,786 | 1,22 | 0,22 | 1,0E-01 |
| Q8CAY6 | Acetyl-CoA acetyltransferase, cytosolic OS=Mus musculus OX=1 | 35 | 13 | 19 | 13 | 397 | 41,3 | 7,5 | 4875 | 13,05 | 6 | 7 | 1,087 | 1,021 | 1,054 | 1,077 | 0,722 | 1,026 | 0,982 | 0,97 | 1,15 | 0,15 | 1,0E-01 |
| Q60931 | Voltage-dependent anion-selective channel protein 3 OS=Mus n | 44 | 11 | 19 | 11 | 283 | 30,7 | 8,79 | 5621 | 28,26 | 7 | 5 | 1,154 | 1,042 | 1,025 | 1,082 | 0,879 | 1,03 | 0,956 | 1,059 | 1,10 | 0,10 | 1,0E-01 |
| A2ASS6 | Titin OS=Mus musculus OX=10090 GN=Ttn PE=1 SV=1 | 0 | 4 | 7 | 3 | 35213 | 3904,1 | 6,2 | 2473 | | 4 | | 1,021 | 0,881 | 0,946 | 0,914 | 1,142 | 0,979 | 0,99 | 1,018 | 0,91 | -0,09 | 1,0E-01 |
| Q9CQI6 | Coactosin-like protein OS=Mus musculus OX=10090 GN=Cotl1 P | 36 | 8 | 10 | 8 | 142 | 15,9 | 5,4 | 3243 | 11,99 | 3 | 5 | 0,897 | 1,048 | 1,095 | 0,992 | 0,658 | 1,007 | 0,902 | 0,813 | 1,19 | 0,19 | 1,0E-01 |
| Q8R050 | Eukaryotic peptide chain release factor GTP-binding subunit ERF | 15 | 10 | 12 | 3 | 636 | 68,6 | 5,21 | 5451 | 11,18 | 7 | 3 | 1,051 | 0,941 | 1,04 | 0,982 | 0,791 | 1,004 | 0,847 | 0,953 | 1,12 | 0,12 | 1,1E-01 |
| Q64010 | Adapter molecule crk OS=Mus musculus OX=10090 GN=Crk PE= | 13 | 3 | 3 | 3 | 304 | 33,8 | 5,55 | 2250 | | 3 | | 0,878 | 1,246 | 0,841 | 1,157 | 2,101 | 0,756 | 1,705 | 2,072 | 0,62 | -0,38 | 1,1E-01 |
| Q07113 | Cation-independent mannose-6-phosphate receptor OS=Mus m | 1 | 2 | 2 | 2 | 2483 | 273,6 | 5,71 | 1038 | | 2 | | 1,033 | 0,985 | 0,908 | 1,139 | 1,25 | 1,198 | 1,01 | 1,145 | 0,88 | -0,12 | 1,1E-01 |
| Q921M3 | Splicing factor 3B subunit 3 OS=Mus musculus OX=10090 GN=Sf | 4 | 3 | 3 | 3 | 1217 | 135,5 | 5,26 | 1248 | 1,93 | 2 | 1 | 1,04 | 0,907 | 1,136 | 1,151 | 1,412 | 1,065 | 1,369 | 1,154 | 0,85 | -0,15 | 1,1E-01 |
| P63037 | DnaJ homolog subfamily A member 1 OS=Mus musculus OX=100 | 19 | 7 | 11 | 7 | 397 | 44,8 | 7,08 | 3081 | 13,29 | 4 | 4 | 1,084 | 0,998 | 1,108 | 1,004 | 0,665 | 0,988 | 0,944 | 0,977 | 1,17 | 0,17 | 1,1E-01 |
| Q9D6E4 | Tetratricopeptide repeat protein 9B OS=Mus musculus OX=1009 | 5 | 1 | 2 | 1 | 239 | 25,9 | 9,48 | 538 | | 1 | | 0,889 | 1 | 1,017 | 1,044 | 1,147 | 1,083 | 1,095 | 0,987 | 0,92 | -0,08 | 1,1E-01 |
| Q91W52 | Transmembrane protein 19 OS=Mus musculus OX=10090 GN=Tr | 4 | 1 | 2 | 1 | 336 | 36,3 | 7,72 | 964 | | 1 | | 1,007 | 0,879 | 1,113 | 1,015 | 0,9 | 0,811 | 0,97 | 0,893 | 1,12 | 0,12 | 1,1E-01 |
| P17427 | AP-2 complex subunit alpha-2 OS=Mus musculus OX=10090 GN= | 24 | 24 | 47 | 17 | 938 | 104 | 6,93 | 21778 | 24,08 | 16 | 8 | 1,008 | 0,987 | 0,986 | 0,992 | 1,06 | 0,975 | 1,039 | 1,048 | 0,96 | -0,04 | 1,1E-01 |
| P61089 | Ubiquitin-conjugating enzyme E2 N OS=Mus musculus OX=1009 | 37 | 5 | 8 | 5 | 152 | 17,1 | 6,57 | 2397 | 15,37 | 4 | 1 | 1,026 | 1,015 | 0,878 | 0,996 | 0,83 | 0,979 | 0,906 | 0,806 | 1,11 | 0,11 | 1,1E-01 |
| Q6WVG3 | BTB/POZ domain-containing protein KCTD12 OS=Mus musculus | 21 | 4 | 9 | 4 | 327 | 35,9 | 5,81 | 3430 | 5,55 | 4 | 2 | 0,886 | 0,83 | 0,945 | 0,7 | 1,926 | 0,741 | 1,084 | 1,797 | 0,61 | -0,39 | 1,1E-01 |
| Q3UHB1 | 5'-nucleotidase domain-containing protein 3 OS=Mus musculus | 9 | 5 | 5 | 5 | 546 | 63,1 | 8,56 | 2197 | 5,21 | 3 | 2 | 1,01 | 0,961 | 0,97 | 0,949 | 0,944 | 0,87 | 0,964 | 0,929 | 1,05 | 0,05 | 1,1E-01 |
| P53657 | Pyruvate kinase PKLR OS=Mus musculus OX=10090 GN=Pklr PE= | 7 | 3 | 7 | 2 | 574 | 62,3 | 7,09 | 3951 | 4,87 | 3 | 1 | 1,425 | 1,333 | 1,207 | 1,419 | 2,096 | 1,405 | 1,395 | 1,856 | 0,80 | -0,20 | 1,1E-01 |
| Q6PGB8 | Probable global transcription activator SNF2L1 OS=Mus muscul | 1 | 1 | 1 | 1 | 1046 | 121,6 | 8,27 | 216 | | 1 | | 0,836 | 1 | 0,816 | 0,885 | 1,519 | 0,768 | 1,213 | 1,255 | 0,74 | -0,26 | 1,1E-01 |
| Q9D023 | Mitochondrial pyruvate carrier 2 OS=Mus musculus OX=10090 G | 10 | 2 | 2 | 2 | 127 | 14,3 | 10,61 | 840 | 0 | 1 | 1 | 0,976 | 1,299 | 1,169 | 1,417 | 2,179 | 1,245 | 1,496 | 1,595 | 0,75 | -0,25 | 1,1E-01 |
| Q922J3 | CAP-Gly domain-containing linker protein 1 OS=Mus musculus C | 19 | 25 | 38 | 22 | 1391 | 155,7 | 5,24 | 14197 | 37,02 | 16 | 9 | 0,954 | 0,932 | 0,948 | 0,951 | 0,852 | 0,964 | 0,908 | 0,872 | 1,05 | 0,05 | 1,1E-01 |
| O70311 | Glycylpeptide N-tetradecanoyltransferase 2 OS=Mus musculus C | 4 | 2 | 2 | 2 | 529 | 60,4 | 7,8 | 1568 | | 2 | | 1,039 | 0,906 | 0,994 | 0,973 | 0,882 | 0,921 | 0,86 | 0,971 | 1,08 | 0,08 | 1,1E-01 |
| Q9QZX7 | Serine racemase OS=Mus musculus OX=10090 GN=Srr PE=1 SV= | 14 | 3 | 3 | 3 | 339 | 36,3 | 6,02 | 1160 | 3,28 | 2 | 1 | 0,911 | 0,939 | 1,037 | 0,88 | 1,108 | 1,036 | 0,98 | 0,984 | 0,92 | -0,08 | 1,1E-01 |

| | | | | | | | | | | | | | | | | | | | | | | | |
|---|---|---|---|---|---|---|---|---|---|---|---|---|---|---|---|---|---|---|---|---|---|---|---|
| Q501J6 | Probable ATP-dependent RNA helicase DDX17 OS=Mus musculu | 15 | 10 | 13 | 4 | 650 | 72,4 | 8,59 | 6089 | 2,26 | 8 | 2 | 1,283 | 1,124 | 1,365 | 0,934 | 1,34 | 1,369 | 1,523 | 1,286 | 0,85 | -0,15 | 1,1E-01 |
| Q8R1G2 | Carboxymethylenebutenolidase homolog OS=Mus musculus OX | 4 | 1 | 2 | 1 | 245 | 27,9 | 7,18 | | 5,22 | | 1 | 1,663 | 1,499 | 1,836 | 1,47 | 2,505 | 1,689 | 1,971 | 1,803 | 0,81 | -0,19 | 1,1E-01 |
| P47199 | Quinone oxidoreductase OS=Mus musculus OX=10090 GN=Cryz | 17 | 3 | 4 | 3 | 331 | 35,2 | 8,07 | 2475 | | 3 | | 1,167 | 1,102 | 1,069 | 1,001 | 1,274 | 1,192 | 1,226 | 1,068 | 0,91 | -0,09 | 1,1E-01 |
| Q9R060 | Cytosolic Fe-S cluster assembly factor NUBP1 OS=Mus musculus | 3 | 1 | 1 | 1 | 320 | 34,1 | 5,9 | 695 | | 1 | | 0,886 | 0,897 | 0,962 | 1,05 | 0,817 | 0,958 | 0,838 | 0,752 | 1,13 | 0,13 | 1,1E-01 |
| O35326 | Serine/arginine-rich splicing factor 5 OS=Mus musculus OX=100 | 8 | 2 | 3 | 2 | 269 | 30,9 | 11,56 | 1212 | | 2 | | 0,915 | 0,923 | 1,164 | 1,291 | 0,97 | 0,826 | 0,88 | 0,885 | 1,21 | 0,21 | 1,1E-01 |
| P11627 | Neural cell adhesion molecule L1 OS=Mus musculus OX=10090 | 19 | 24 | 46 | 23 | 1260 | 140,9 | 5,97 | 16018 | 56,38 | 14 | 10 | 1,073 | 1,065 | 1,025 | 1,099 | 0,79 | 1,068 | 0,975 | 0,976 | 1,12 | 0,12 | 1,1E-01 |
| Q3THG9 | Alanyl-tRNA editing protein Aarsd1 OS=Mus musculus OX=1009 | 13 | 5 | 8 | 5 | 412 | 44,9 | 6,42 | 3708 | 2,65 | 4 | 1 | 0,946 | 1,024 | 0,931 | 0,919 | 0,725 | 0,98 | 0,81 | 0,866 | 1,13 | 0,13 | 1,1E-01 |
| Q8BHC9 | Alpha-(1,3)-fucosyltransferase 11 OS=Mus musculus OX=10090 | 2 | 1 | 1 | 1 | 489 | 55,5 | 6,23 | 633 | | 1 | | 0,898 | 0,808 | 1,055 | 0,878 | 0,746 | 0,723 | 0,906 | 0,766 | 1,16 | 0,16 | 1,1E-01 |
| Q9CR09 | Ubiquitin-fold modifier-conjugating enzyme 1 OS=Mus musculu | 11 | 2 | 3 | 2 | 167 | 19,5 | 7,4 | 1448 | | 2 | | 1,067 | 1,002 | 1,115 | 1,018 | 0,846 | 1,11 | 0,854 | 0,886 | 1,14 | 0,14 | 1,1E-01 |
| P84086 | Complexin-2 OS=Mus musculus OX=10090 GN=Cplx2 PE=1 SV=1 | 43 | 4 | 6 | 4 | 134 | 15,4 | 5,08 | 3574 | | 4 | | 0,985 | 0,864 | 1,027 | 0,891 | 0,581 | 0,946 | 0,839 | 0,757 | 1,21 | 0,21 | 1,1E-01 |
| P52825 | Carnitine O-palmitoyltransferase 2, mitochondrial OS=Mus mus | 6 | 3 | 3 | 3 | 658 | 73,9 | 8,37 | 1209 | 2,84 | 2 | 1 | 1,087 | 1,234 | 1,259 | 1,242 | 1,114 | 1,198 | 0,991 | 1,082 | 1,10 | 0,10 | 1,1E-01 |
| P51150 | Ras-related protein Rab-7a OS=Mus musculus OX=10090 GN=Ra | 47 | 10 | 18 | 10 | 207 | 23,5 | 6,7 | 9742 | 2,72 | 9 | 2 | 1,059 | 0,981 | 1,082 | 0,983 | 0,981 | 1,012 | 0,945 | 0,882 | 1,07 | 0,07 | 1,1E-01 |
| Q8BKZ9 | Pyruvate dehydrogenase protein X component, mitochondrial O | 30 | 10 | 12 | 10 | 501 | 54 | 7,75 | 5336 | 0 | 8 | 2 | 0,977 | 0,859 | 1,019 | 0,967 | 1,053 | 1,02 | 0,977 | 1,067 | 0,93 | -0,07 | 1,1E-01 |
| Q9QYG0 | Protein NDRG2 OS=Mus musculus OX=10090 GN=Ndrg2 PE=1 S | 29 | 9 | 14 | 9 | 371 | 40,8 | 5,4 | 6744 | 7,69 | 7 | 3 | 1,018 | 0,928 | 1,271 | 0,978 | 0,71 | 0,938 | 0,977 | 0,85 | 1,21 | 0,21 | 1,1E-01 |
| Q3UHL1 | CaM kinase-like vesicle-associated protein OS=Mus musculus O | 29 | 11 | 20 | 11 | 512 | 54,8 | 5,54 | 8176 | 17,24 | 8 | 6 | 0,892 | 1,006 | 0,93 | 0,916 | 0,95 | 1,001 | 0,983 | 1,045 | 0,94 | -0,06 | 1,1E-01 |
| Q8WTY4 | Anamorsin OS=Mus musculus OX=10090 GN=Ciapin1 PE=1 SV=1 | 19 | 4 | 6 | 4 | 309 | 33,4 | 5,2 | 1069 | 9,43 | 1 | 3 | 1,008 | 0,959 | 0,974 | 1,166 | 0,715 | 0,949 | 0,907 | 0,974 | 1,16 | 0,16 | 1,1E-01 |
| Q9QUG9 | RAS guanyl-releasing protein 2 OS=Mus musculus OX=10090 GN | 12 | 6 | 7 | 6 | 608 | 69,4 | 7,66 | 1436 | 12,48 | 2 | 5 | 0,926 | 0,934 | 1,052 | 0,955 | 0,736 | 0,948 | 0,913 | 0,863 | 1,12 | 0,12 | 1,1E-01 |
| Q91VS7 | Microsomal glutathione S-transferase 1 OS=Mus musculus OX=1 | 8 | 1 | 1 | 1 | 155 | 17,5 | 9,66 | | 2,22 | | 1 | 1,433 | 1,495 | 1,541 | 1,649 | 2,011 | 1,74 | 1,823 | 3,33 | 0,69 | -0,31 | 1,1E-01 |
| Q8BHJ5 | F-box-like/WD repeat-containing protein TBL1XR1 OS=Mus mus | 4 | 2 | 2 | 2 | 514 | 55,6 | 5,63 | 737 | | 2 | | 0,939 | 1,048 | 0,907 | 0,925 | 1,122 | 0,943 | 1,323 | 1,065 | 0,86 | -0,14 | 1,1E-01 |
| Q3TEA8 | Heterochromatin protein 1-binding protein 3 OS=Mus musculus | 13 | 8 | 10 | 8 | 554 | 60,8 | 9,7 | 2828 | 12,48 | 3 | 5 | 0,836 | 0,795 | 0,963 | 0,999 | 1,115 | 0,869 | 1,171 | 1,046 | 0,86 | -0,14 | 1,1E-01 |
| O70589 | Peripheral plasma membrane protein CASK OS=Mus musculus C | 23 | 19 | 24 | 19 | 926 | 105 | 6,43 | 9555 | 19,64 | 14 | 5 | 1,042 | 1,039 | 1,014 | 0,996 | 0,97 | 0,991 | 1,002 | 1,018 | 1,03 | 0,03 | 1,1E-01 |
| P12710 | Fatty acid-binding protein, liver OS=Mus musculus OX=10090 G | 35 | 5 | 9 | 5 | 127 | 14,2 | 8,56 | 5458 | | 5 | | 1,604 | 1,993 | 1,682 | 2,164 | 3,164 | 1,796 | 2,469 | 2,319 | 0,76 | -0,24 | 1,1E-01 |
| P61202 | COP9 signalosome complex subunit 2 OS=Mus musculus OX=10 | 15 | 6 | 8 | 6 | 443 | 51,6 | 5,53 | 3270 | 2,07 | 4 | 2 | 1,073 | 0,959 | 1,031 | 1,028 | 0,913 | 0,987 | 0,989 | 0,981 | 1,06 | 0,06 | 1,1E-01 |
| P23819-3 | Isoform 3 of Glutamate receptor 2 OS=Mus musculus OX=10090 | 12 | 12 | 18 | 9 | 939 | 104,5 | 7,87 | 6115 | 13,22 | 7 | 5 | 1,074 | 1,021 | 1,039 | 1,012 | 0,777 | 0,991 | 0,989 | 0,985 | 1,11 | 0,11 | 1,1E-01 |
| Q8K2M0 | 39S ribosomal protein L38, mitochondrial OS=Mus musculus OX | 6 | 2 | 2 | 2 | 380 | 45 | 8,1 | 266 | 2,75 | 1 | 1 | 0,994 | 1,136 | 1,016 | 1,052 | 1,309 | 1 | 1,181 | 1,261 | 0,88 | -0,12 | 1,1E-01 |
| O35638 | Cohesin subunit SA-2 OS=Mus musculus OX=10090 GN=Stag2 P | 1 | 1 | 2 | 1 | 1231 | 141,2 | 5,43 | 778 | | 1 | | 1,009 | 0,802 | 2,152 | 1,451 | 0,599 | 0,751 | 0,831 | 0,957 | 1,73 | 0,73 | 1,1E-01 |
| Q8JZQ9 | Eukaryotic translation initiation factor 3 subunit B OS=Mus mus | 15 | 10 | 10 | 10 | 803 | 91,3 | 5,02 | 3720 | 4,24 | 7 | 3 | 0,952 | 0,943 | 0,95 | 0,972 | 1,106 | 0,953 | 1,016 | 0,988 | 0,94 | -0,06 | 1,1E-01 |
| Q99N96 | 39S ribosomal protein L1, mitochondrial OS=Mus musculus OX= | 10 | 3 | 3 | 3 | 336 | 37,6 | 8,72 | 1129 | 3,91 | 2 | 1 | 1,052 | 1,104 | 0,755 | 0,846 | 2,319 | 1,111 | 1,305 | 1,191 | 0,63 | -0,37 | 1,1E-01 |
| P32037 | Solute carrier family 2, facilitated glucose transporter member 3 | 9 | 4 | 5 | 4 | 493 | 53,4 | 4,98 | 1688 | 3,41 | 3 | 1 | 1,048 | 1,019 | 1,117 | 0,881 | 0,699 | 1,001 | 0,894 | 0,882 | 1,17 | 0,17 | 1,1E-01 |
| P21300 | Aldo-keto reductase family 1 member B7 OS=Mus musculus OX= | 3 | 1 | 1 | 1 | 316 | 36 | 7,25 | 669 | | 1 | | 0,887 | 0,759 | 0,918 | 0,959 | 0,555 | 0,908 | 0,622 | 0,774 | 1,23 | 0,23 | 1,1E-01 |
| Q8VCI0 | Phospholipase B-like 1 OS=Mus musculus OX=10090 GN=Plbd1 | 2 | 1 | 1 | 1 | 550 | 63 | 8,87 | 233 | | 1 | | 0,894 | 1,085 | 0,719 | 1,253 | 1,66 | 1,014 | 1,138 | 1,642 | 0,72 | -0,28 | 1,2E-01 |
| Q9JIF0 | Protein arginine N-methyltransferase 1 OS=Mus musculus OX=1 | 21 | 7 | 9 | 5 | 371 | 42,4 | 5,43 | 4397 | 8,39 | 6 | 1 | 0,865 | 0,949 | 0,911 | 0,945 | 1,043 | 0,91 | 0,959 | 1,026 | 0,93 | -0,07 | 1,2E-01 |
| Q5RJH6 | Protein SMG7 OS=Mus musculus OX=10090 GN=Smg7 PE=2 SV= | 1 | 1 | 1 | 1 | 1138 | 126,8 | 7,99 | 281 | | 1 | | 0,94 | 0,87 | 0,935 | 1,024 | 1,114 | 1,09 | 1 | 0,941 | 0,91 | -0,09 | 1,2E-01 |
| Q9QYF9 | Protein NDRG3 OS=Mus musculus OX=10090 GN=Ndrg3 PE=1 S | 24 | 6 | 10 | 6 | 375 | 41,5 | 5,25 | 4249 | 4,32 | 6 | 2 | 0,918 | 0,948 | 0,981 | 0,914 | 1,041 | 0,91 | 1,064 | 1,022 | 0,93 | -0,07 | 1,2E-01 |
| P11352 | Glutathione peroxidase 1 OS=Mus musculus OX=10090 GN=Gpx | 6 | 2 | 2 | 2 | 201 | 22,3 | 7,21 | 1009 | | 2 | | 0,839 | 1,225 | 1,039 | 1,099 | 1,599 | 1,28 | 1,068 | 1,256 | 0,81 | -0,19 | 1,2E-01 |
| E9PUL5 | Proline-rich transmembrane protein 2 OS=Mus musculus OX=10 | 15 | 3 | 9 | 3 | 346 | 35,9 | 4,63 | 2181 | 15,46 | 2 | 2 | 0,99 | 0,855 | 1,176 | 0,872 | 0,516 | 0,966 | 0,779 | 0,764 | 1,29 | 0,29 | 1,2E-01 |
| Q8K4F5 | Protein ABHD11 OS=Mus musculus OX=10090 GN=Abhd11 PE=1 | 4 | 1 | 1 | 1 | 307 | 33,5 | 9,61 | 775 | | 1 | | 1,076 | 1,071 | 1,195 | 1,184 | 0,791 | 0,816 | 1,135 | 1,085 | 1,18 | 0,18 | 1,2E-01 |
| Q8BRV5 | Uncharacterized protein KIAA1671 OS=Mus musculus OX=1009 | 6 | 2 | 7 | 2 | 308 | 34,6 | 6,09 | 3394 | | 2 | | 0,962 | 0,98 | 0,929 | 0,884 | 0,705 | 0,926 | 0,871 | 0,87 | 1,11 | 0,11 | 1,2E-01 |
| Q9JIX8 | Apoptotic chromatin condensation inducer in the nucleus OS=M | 1 | 1 | 1 | 1 | 1338 | 150,6 | 5,91 | | 2,24 | | 1 | 1,074 | 0,943 | 1,291 | 1,075 | 1,241 | 1,247 | 1,296 | 1,163 | 0,89 | -0,11 | 1,2E-01 |
| Q63932 | Dual specificity mitogen-activated protein kinase kinase 2 OS=M | 15 | 6 | 8 | 3 | 401 | 44,4 | 7,05 | 3333 | 6,54 | 5 | 2 | 0,981 | 1,043 | 1,248 | 1,141 | 0,927 | 1,093 | 0,947 | 0,93 | 1,13 | 0,13 | 1,2E-01 |
| Q69ZR2 | E3 ubiquitin-protein ligase HECTD1 OS=Mus musculus OX=1009 | 1 | 3 | 4 | 3 | 2618 | 289,9 | 5,41 | 2195 | | 3 | | 0,718 | 0,817 | 0,812 | 0,825 | 0,92 | 0,879 | 0,768 | 0,987 | 0,89 | -0,11 | 1,2E-01 |
| Q9QXV0 | ProSAAS OS=Mus musculus OX=10090 GN=Pcsk1n PE=1 SV=2 | 17 | 3 | 3 | 3 | 258 | 27,3 | 5,85 | 1391 | 3,96 | 2 | 1 | 0,867 | 1,115 | 0,876 | 1,083 | 0,774 | 0,954 | 0,862 | 0,779 | 1,17 | 0,17 | 1,2E-01 |
| Q61543 | Golgi apparatus protein 1 OS=Mus musculus OX=10090 GN=Glg | 5 | 5 | 6 | 5 | 1175 | 133,6 | 6,84 | 1805 | 5,55 | 4 | 1 | 0,797 | 0,824 | 1,02 | 1,024 | 1,147 | 0,921 | 1,048 | 1,134 | 0,86 | -0,14 | 1,2E-01 |
| Q9CQM9 | Glutaredoxin-3 OS=Mus musculus OX=10090 GN=Glrx3 PE=1 SV | 16 | 5 | 9 | 5 | 337 | 37,8 | 5,59 | 4443 | 7,53 | 3 | 2 | 1,048 | 1,151 | 1,075 | 1,23 | 0,933 | 1,079 | 1,086 | 1,011 | 1,10 | 0,10 | 1,2E-01 |
| Q8BHL3 | TBC1 domain family member 10B OS=Mus musculus OX=10090 | 4 | 3 | 3 | 3 | 798 | 87,2 | 8,9 | 1264 | 2,1 | 2 | 1 | 1,075 | 1,002 | 1,152 | 0,742 | 0,7 | 0,863 | 0,81 | 0,885 | 1,22 | 0,22 | 1,2E-01 |
| O08585 | Clathrin light chain A OS=Mus musculus OX=10090 GN=Clta PE= | 31 | 10 | 17 | 10 | 235 | 25,6 | 4,58 | 3300 | 31,72 | 3 | 7 | 0,99 | 1,07 | 0,979 | 1,038 | 0,599 | 1,058 | 0,826 | 0,888 | 1,21 | 0,21 | 1,2E-01 |
| Q9JHU9 | Inositol-3-phosphate synthase 1 OS=Mus musculus OX=10090 G | 5 | 3 | 3 | 3 | 557 | 60,9 | 6,42 | 1335 | 2,59 | 2 | 1 | 1,028 | 0,92 | 0,933 | 1,092 | 0,746 | 0,978 | 0,913 | 0,874 | 1,13 | 0,13 | 1,2E-01 |
| Q6DFW0 | Guanine nucleotide exchange C9orf72 homolog OS=Mus muscu | 2 | 1 | 1 | 1 | 481 | 54,2 | 6,09 | 333 | | 1 | | 1,155 | 1,029 | 0,956 | 1,181 | 1,286 | 1,254 | 1,083 | 1,201 | 0,90 | -0,10 | 1,2E-01 |
| Q5KU39 | Vacuolar protein sorting-associated protein 41 homolog OS=Mu | 1 | 1 | 1 | 1 | 853 | 98,5 | 5,81 | | 2,55 | | 1 | 0,898 | 0,82 | 0,897 | 0,855 | 0,852 | 0,978 | 0,917 | 1,004 | 0,93 | -0,07 | 1,2E-01 |
| Q99JT1 | Glutamyl-tRNA(Gln) amidotransferase subunit B, mitochondrial | 3 | 1 | 1 | 1 | 557 | 62,1 | 8,54 | 577 | | 1 | | 0,765 | 1,701 | 0,907 | 1,086 | 1,734 | 1,217 | 2,76 | 1,573 | 0,61 | -0,39 | 1,2E-01 |
| Q9D0F9 | Phosphoglucomutase-1 OS=Mus musculus OX=10090 GN=Pgm1 | 24 | 12 | 17 | 12 | 562 | 61,4 | 6,57 | 4665 | 20,81 | 7 | 5 | 0,977 | 1,089 | 1,132 | 1,136 | 1,388 | 1,199 | 1,161 | 1,105 | 0,89 | -0,11 | 1,2E-01 |
| Q8K296 | Myotubularin-related protein 3 OS=Mus musculus OX=10090 G | 1 | 1 | 1 | 1 | 1196 | 133,8 | 5,83 | 266 | | 1 | | 0,856 | 0,786 | 0,82 | 0,941 | 0,876 | 0,676 | 0,731 | 0,726 | 1,13 | 0,13 | 1,2E-01 |
| Q80UM7 | Mannosyl-oligosaccharide glucosidase OS=Mus musculus OX=1 | 5 | 3 | 4 | 3 | 834 | 91,8 | 9 | 1027 | 5,79 | 2 | 1 | 1,014 | 0,975 | 1,029 | 1,214 | 1,322 | 1,063 | 1,14 | 1,457 | 0,85 | -0,15 | 1,2E-01 |
| Q9EPK2 | Protein XRP2 OS=Mus musculus OX=10090 GN=Rp2 PE=1 SV=3 | 2 | 1 | 2 | 1 | 347 | 39,4 | 5,24 | 1199 | | 1 | | 0,897 | 0,945 | 0,949 | 0,921 | 1,171 | 1,01 | 1,024 | 0,908 | 0,90 | -0,10 | 1,2E-01 |
| Q9CQS4 | Solute carrier family 25 member 46 OS=Mus musculus OX=1009 | 5 | 2 | 2 | 2 | 418 | 46,2 | 7,64 | 951 | | 2 | | 1,054 | 0,885 | 0,919 | 0,955 | 1,436 | 0,951 | 1,13 | 1,083 | 0,83 | -0,17 | 1,2E-01 |
| Q9DB41 | Mitochondrial glutamate carrier 2 OS=Mus musculus OX=10090 | 9 | 3 | 4 | 1 | 320 | 34,1 | 9,19 | 1100 | 5,64 | 2 | 1 | 1,58 | 1,415 | 1,391 | 1,529 | 1,108 | 1,419 | 1,052 | 1,485 | 1,17 | 0,17 | 1,2E-01 |
| P63276 | 40S ribosomal protein S17 OS=Mus musculus OX=10090 GN=Rp | 16 | 4 | 7 | 4 | 135 | 15,5 | 9,85 | 3101 | | 4 | | 0,965 | 0,874 | 0,987 | 1,034 | 0,94 | 0,909 | 0,815 | 0,887 | 1,09 | 0,09 | 1,2E-01 |
| Q9Z1F9 | SUMO-activating enzyme subunit 2 OS=Mus musculus OX=1009 | 1 | 1 | 1 | 1 | 638 | 70,5 | 5,24 | 435 | | 1 | | 1,075 | 0,95 | 0,93 | 1,151 | 0,998 | 0,872 | 0,944 | 0,743 | 1,15 | 0,15 | 1,2E-01 |
| Q8BHL5 | Engulfment and cell motility protein 2 OS=Mus musculus OX=10 | 8 | 5 | 6 | 5 | 732 | 83,8 | 5,95 | 2240 | 4,44 | 4 | 2 | 0,942 | 0,887 | 1,114 | 1,005 | 1,262 | 1,011 | 1,185 | 1,043 | 0,88 | -0,12 | 1,2E-01 |
| Q8BTG7-3 | Isoform 3 of Protein NDRG4 OS=Mus musculus OX=10090 GN=N | 8 | 3 | 4 | 3 | 404 | 44,4 | 6,23 | 761 | 7,94 | 1 | 2 | 1,109 | 1,004 | 1,231 | 1,022 | 0,705 | 1,079 | 0,983 | 0,916 | 1,19 | 0,19 | 1,2E-01 |
| O70252 | Heme oxygenase 2 OS=Mus musculus OX=10090 GN=Hmox2 PE | 6 | 2 | 2 | 2 | 315 | 35,7 | 5,87 | 719 | 2,93 | 1 | 1 | 0,985 | 1,001 | 1,108 | 1,085 | 0,556 | 0,972 | 0,922 | 0,972 | 1,22 | 0,22 | 1,2E-01 |
| O88986 | 2-amino-3-ketobutyrate coenzyme A ligase, mitochondrial OS=M | 7 | 2 | 3 | 2 | 416 | 44,9 | 7,33 | 1443 | | 2 | | 0,929 | 0,936 | 1,072 | 1,147 | 1,409 | 1,095 | 1,014 | 1,385 | 0,83 | -0,17 | 1,2E-01 |
| Q8BH95 | Enoyl-CoA hydratase, mitochondrial OS=Mus musculus OX=1009 | 27 | 7 | 10 | 7 | 290 | 31,5 | 8,48 | 932 | 22,46 | 2 | 5 | 0,983 | 0,97 | 0,977 | 1,046 | 1,037 | 1,033 | 1,025 | 1,185 | 0,93 | -0,07 | 1,2E-01 |
| Q8R5H6 | Wiskott-Aldrich syndrome protein family member 1 OS=Mus mu | 12 | 6 | 8 | 5 | 559 | 61,5 | 6,37 | 1320 | 6,75 | 3 | 3 | 0,953 | 0,819 | 0,999 | 0,762 | 1,044 | 0,928 | 1,015 | 0,984 | 0,89 | -0,11 | 1,2E-01 |
| Q8BML9 | Glutamine--tRNA ligase OS=Mus musculus OX=10090 GN=Qars | 3 | 2 | 2 | 2 | 775 | 87,6 | 7,31 | 583 | | 2 | | 0,806 | 0,923 | 0,943 | 1,027 | 1,13 | 0,923 | 1,012 | 1,107 | 0,89 | -0,11 | 1,2E-01 |

| | | | | | | | | | | | | | | | | | | | | | | | |
|---|---|---|---|---|---|---|---|---|---|---|---|---|---|---|---|---|---|---|---|---|---|---|---|
| Q9CQ54 | NADH dehydrogenase [ubiquinone] 1 subunit C2 OS=Mus muscu | 33 | 6 | 8 | 6 | 120 | 14,2 | 9,2 | 3208 | 6,54 | 4 | 2 | 1,051 | 0,924 | 1,149 | 1,025 | 0,977 | 0,998 | 0,911 | 0,864 | 1,11 | 0,11 | 1,2E-01 |
| P27601 | Guanine nucleotide-binding protein subunit alpha-13 OS=Mus m | 18 | 7 | 13 | 6 | 377 | 44 | 8,21 | 5893 | 10,52 | 4 | 3 | 1,033 | 0,962 | 0,984 | 1 | 0,952 | 0,879 | 0,948 | 0,996 | 1,05 | 0,05 | 1,2E-01 |
| P47964 | 60S ribosomal protein L36 OS=Mus musculus OX=10090 GN=Rpl | 18 | 2 | 6 | 2 | 105 | 12,2 | 11,34 | 1236 | 10,02 | 1 | 2 | 0,967 | 0,871 | 1,051 | 0,977 | 1,017 | 1,032 | 1,091 | 1,019 | 0,93 | -0,07 | 1,2E-01 |
| Q99P72 | Reticulon-4 OS=Mus musculus OX=10090 GN=Rtn4 PE=1 SV=2 | 31 | 33 | 65 | 33 | 1162 | 126,5 | 4,54 | 21862 | 94,87 | 20 | 17 | 1,066 | 0,999 | 1,07 | 0,991 | 0,808 | 1,02 | 0,955 | 0,979 | 1,10 | 0,10 | 1,2E-01 |
| Q5FW60 | Major urinary protein 20 OS=Mus musculus OX=10090 GN=Mup | 15 | 2 | 3 | 1 | 181 | 20,9 | 4,89 | 1994 | | 2 | | 2,496 | 2,447 | 2,249 | 2,188 | 4,479 | 2,678 | 2,539 | 2,936 | 0,74 | -0,26 | 1,2E-01 |
| O70401 | Tetraspanin-6 OS=Mus musculus OX=10090 GN=Tspan6 PE=1 SV | 10 | 2 | 2 | 2 | 245 | 27,3 | 7,74 | | 6 | | 2 | 1,203 | 1,191 | 1,231 | 1,232 | 0,822 | 1,188 | 1,201 | 1,001 | 1,15 | 0,15 | 1,2E-01 |
| P37040 | NADPH--cytochrome P450 reductase OS=Mus musculus OX=100 | 17 | 10 | 19 | 10 | 678 | 77 | 5,53 | 6754 | 12,67 | 5 | 5 | 1,019 | 0,955 | 1,009 | 0,948 | 1,313 | 1,026 | 1,043 | 1,053 | 0,89 | -0,11 | 1,2E-01 |
| Q61656 | Probable ATP-dependent RNA helicase DDX5 OS=Mus musculus | 20 | 13 | 18 | 7 | 614 | 69,2 | 8,92 | 8821 | 7,8 | 9 | 4 | 0,877 | 0,963 | 0,88 | 1,029 | 1,127 | 0,885 | 1,126 | 1,104 | 0,88 | -0,12 | 1,3E-01 |
| Q61387 | Cytochrome c oxidase subunit 7A-related protein, mitochondria | 12 | 1 | 1 | 1 | 111 | 12,4 | 9,6 | 251 | | 1 | | 0,838 | 0,844 | 0,697 | 0,565 | 1,672 | 0,646 | 1,1 | 1,097 | 0,65 | -0,35 | 1,3E-01 |
| Q8R0A7 | Uncharacterized protein KIAA0513 OS=Mus musculus OX=1009 | 3 | 1 | 1 | 1 | 407 | 46,3 | 5,02 | | 3,54 | | 1 | 0,694 | 0,566 | 0,939 | 0,784 | 1,259 | 0,815 | 0,791 | 1,577 | 0,67 | -0,33 | 1,3E-01 |
| O35598 | Disintegrin and metalloproteinase domain-containing protein 1 | 4 | 2 | 3 | 2 | 749 | 83,9 | 7,94 | 1300 | | 2 | | 0,892 | 0,884 | 0,878 | 0,77 | 1,238 | 1,098 | 0,932 | 0,83 | 0,84 | -0,16 | 1,3E-01 |
| Q9CZM2 | 60S ribosomal protein L15 OS=Mus musculus OX=10090 GN=Rpl | 29 | 5 | 8 | 5 | 204 | 24,1 | 11,62 | 2671 | 8,99 | 3 | 3 | 1,035 | 0,942 | 1,013 | 1,068 | 1,17 | 1,021 | 1,123 | 1,051 | 0,93 | -0,07 | 1,3E-01 |
| P63330 | Serine/threonine-protein phosphatase 2A catalytic subunit alph | 40 | 11 | 14 | 11 | 309 | 35,6 | 5,54 | 4750 | 9,62 | 7 | 4 | 0,953 | 0,912 | 0,934 | 0,944 | 1,047 | 1,007 | 0,953 | 0,932 | 0,95 | -0,05 | 1,3E-01 |
| O35685 | Nuclear migration protein nudC OS=Mus musculus OX=10090 G | 42 | 12 | 22 | 12 | 332 | 38,3 | 5,26 | 10755 | 5,43 | 10 | 2 | 0,971 | 0,998 | 0,992 | 1,018 | 0,864 | 0,999 | 0,968 | 0,93 | 1,06 | 0,06 | 1,3E-01 |
| Q8BGH4 | Receptor expression-enhancing protein 1 OS=Mus musculus OX | 20 | 4 | 4 | 4 | 201 | 22,3 | 9,5 | 1498 | 2,52 | 2 | 2 | 0,894 | 0,959 | 0,837 | 0,847 | 1,084 | 0,858 | 1,069 | 1,496 | 0,78 | -0,22 | 1,3E-01 |
| P38060 | Hydroxymethylglutaryl-CoA lyase, mitochondrial OS=Mus muscu | 11 | 3 | 7 | 3 | 325 | 34,2 | 8,41 | 2330 | 13,58 | 3 | 1 | 1,005 | 1,033 | 1,066 | 1,033 | 1,36 | 1,038 | 1,096 | 1,15 | 0,89 | -0,11 | 1,3E-01 |
| Q8BH55 | Threonine synthase-like 1 OS=Mus musculus OX=10090 GN=Thr | 5 | 3 | 3 | 3 | 747 | 83 | 7,2 | 1100 | 2,99 | 2 | 1 | 1,153 | 1,117 | 1,176 | 1,207 | 1,131 | 1,381 | 1,222 | 1,364 | 0,91 | -0,09 | 1,3E-01 |
| Q9WTL7 | Acyl-protein thioesterase 2 OS=Mus musculus OX=10090 GN=Ly | 10 | 3 | 7 | 3 | 231 | 24,8 | 7,23 | 3590 | 4,61 | 2 | 1 | 1,02 | 0,898 | 1,05 | 0,925 | 0,816 | 0,982 | 0,89 | 0,833 | 1,11 | 0,11 | 1,3E-01 |
| Q3UQ44 | Ras GTPase-activating-like protein IQGAP2 OS=Mus musculus O | 1 | 1 | 1 | 1 | 1575 | 180,4 | 5,64 | 461 | | 1 | | 1,032 | 1,095 | 1,031 | 1,211 | 1,874 | 1,007 | 1,367 | 1,42 | 0,77 | -0,23 | 1,3E-01 |
| A2ACJ2 | Fanconi anemia core complex-associated protein 100 OS=Mus n | 4 | 2 | 2 | 2 | 879 | 94,2 | 5,36 | 493 | 3,03 | 1 | 1 | 0,688 | 0,899 | 0,61 | 0,602 | 1,683 | 1,014 | 1,385 | 4,637 | 0,32 | -0,68 | 1,3E-01 |
| Q6PCZ4 | Melanoma-associated antigen E1 OS=Mus musculus OX=10090 | 1 | 1 | 2 | 1 | 918 | 101,6 | 6,67 | 679 | | 1 | | 1,09 | 0,934 | 1,116 | 1,088 | 0,773 | 1,062 | 0,94 | 0,939 | 1,14 | 0,14 | 1,3E-01 |
| O35887 | Calumenin OS=Mus musculus OX=10090 GN=Calu PE=1 SV=1 | 18 | 6 | 7 | 6 | 315 | 37 | 4,67 | 3751 | | 6 | | 1,019 | 1,063 | 1,137 | 1,153 | 0,989 | 0,913 | 1,067 | 1,062 | 1,08 | 0,08 | 1,3E-01 |
| P83882 | 60S ribosomal protein L36a OS=Mus musculus OX=10090 GN=R | 8 | 2 | 2 | 2 | 106 | 12,4 | 10,58 | 1317 | | 2 | | 0,93 | 0,845 | 0,919 | 0,975 | 0,619 | 0,881 | 0,753 | 0,908 | 1,16 | 0,16 | 1,3E-01 |
| P60761 | Neurogranin OS=Mus musculus OX=10090 GN=Nrgn PE=1 SV=1 | 32 | 3 | 5 | 3 | 78 | 7,5 | 7,05 | 1187 | 10,59 | 2 | 2 | 0,699 | 0,875 | 1,489 | 0,986 | 0,477 | 0,935 | 0,62 | 0,636 | 1,52 | 0,52 | 1,3E-01 |
| Q8VDM4 | 26S proteasome non-ATPase regulatory subunit 2 OS=Mus musc | 18 | 13 | 17 | 13 | 908 | 100,1 | 5,17 | 8206 | 5,43 | 12 | 2 | 1 | 1,015 | 0,964 | 1,032 | 1,071 | 1,004 | 1,05 | 1,199 | 0,93 | -0,07 | 1,3E-01 |
| Q99104 | Unconventional myosin-Va OS=Mus musculus OX=10090 GN=M | 27 | 47 | 64 | 47 | 1853 | 215,4 | 8,63 | 24533 | 53,92 | 28 | 20 | 1,07 | 0,955 | 0,999 | 0,975 | 0,789 | 0,942 | 0,961 | 0,962 | 1,09 | 0,09 | 1,3E-01 |
| Q925N0 | Sideroflexin-5 OS=Mus musculus OX=10090 GN=Sfxn5 PE=1 SV= | 13 | 5 | 9 | 5 | 342 | 37,3 | 9,44 | 4126 | 7,36 | 2 | 3 | 1,046 | 1,015 | 1,004 | 1,007 | 0,822 | 0,989 | 0,981 | 0,984 | 1,08 | 0,08 | 1,3E-01 |
| Q64471 | Glutathione S-transferase theta-1 OS=Mus musculus OX=10090 | 3 | 1 | 1 | 1 | 240 | 27,4 | 7,27 | 807 | | 1 | | 1,228 | 1,093 | 1,242 | 1,647 | 1,664 | 1,643 | 1,396 | 1,471 | 0,84 | -0,16 | 1,3E-01 |
| P55096 | ATP-binding cassette sub-family D member 3 OS=Mus musculus | 5 | 4 | 7 | 4 | 659 | 75,4 | 9,26 | 1938 | 12,11 | 2 | 2 | 1,035 | 0,959 | 0,992 | 0,919 | 0,795 | 0,83 | 0,94 | 0,982 | 1,10 | 0,10 | 1,3E-01 |
| P59648 | FXYD domain-containing ion transport regulator 7 OS=Mus mus | 16 | 1 | 3 | 1 | 80 | 8,5 | 7,84 | 2344 | | 1 | | 0,964 | 1,166 | 0,765 | 1,25 | 0,819 | 0,917 | 0,841 | 0,775 | 1,24 | 0,24 | 1,3E-01 |
| Q8R0F8 | Acylpyruvase FAHD1, mitochondrial OS=Mus musculus OX=1009 | 12 | 2 | 2 | 2 | 227 | 25,2 | 7,69 | 814 | 0 | 1 | 1 | 0,934 | 1,047 | 1,042 | 0,984 | 1,218 | 1,011 | 1,072 | 1,069 | 0,92 | -0,08 | 1,3E-01 |
| Q8BIE6 | FERM domain-containing protein 4A OS=Mus musculus OX=100 | 14 | 11 | 12 | 11 | 1020 | 113,8 | 8,92 | 4671 | 5,88 | 8 | 3 | 1,056 | 1,019 | 1,02 | 1,044 | 1,043 | 0,878 | 1,014 | 0,93 | 1,07 | 0,07 | 1,3E-01 |
| O88696 | ATP-dependent Clp protease proteolytic subunit, mitochondrial | 19 | 4 | 6 | 4 | 272 | 29,8 | 7,47 | 3471 | 2,35 | 3 | 1 | 0,907 | 1,328 | 0,854 | 0,831 | 1,114 | 1,305 | 1,123 | 1,269 | 0,81 | -0,19 | 1,3E-01 |
| P63147 | Ubiquitin-conjugating enzyme E2 B OS=Mus musculus OX=1009 | 11 | 1 | 1 | 1 | 152 | 17,3 | 5,01 | | 2,05 | | 1 | 0,959 | 1,243 | 0,938 | 1,314 | 1,21 | 1,423 | 1,29 | 1,276 | 0,86 | -0,14 | 1,3E-01 |
| Q91W90 | Thioredoxin domain-containing protein 5 OS=Mus musculus OX | 6 | 3 | 3 | 2 | 417 | 46,4 | 5,78 | 2434 | | 3 | | 1,299 | 1,215 | 0,75 | 1,304 | 0,555 | 0,968 | 0,846 | 1,02 | 1,35 | 0,35 | 1,3E-01 |
| Q9DC53 | Copine-8 OS=Mus musculus OX=10090 GN=Cpne8 PE=2 SV=3 | 10 | 5 | 5 | 3 | 577 | 64,6 | 5,97 | 1382 | 8,32 | 2 | 3 | 0,88 | 1,06 | 1,022 | 0,98 | 1,03 | 1,089 | 1,083 | 1,033 | 0,93 | -0,07 | 1,3E-01 |
| P24549 | Retinal dehydrogenase 1 OS=Mus musculus OX=10090 GN=Aldh | 11 | 6 | 7 | 3 | 501 | 54,4 | 7,8 | 1796 | 11,29 | 3 | 3 | 1,982 | 1,936 | 1,913 | 2,219 | 2,572 | 2,222 | 2,033 | 2,166 | 0,90 | -0,10 | 1,3E-01 |
| O88844 | Isocitrate dehydrogenase [NADP] cytoplasmic OS=Mus musculus | 44 | 22 | 48 | 21 | 414 | 46,6 | 7,17 | 16930 | 55,48 | 12 | 13 | 1,018 | 0,939 | 1,074 | 0,997 | 0,729 | 1,005 | 0,925 | 0,916 | 1,13 | 0,13 | 1,3E-01 |
| Q61584 | Fragile X mental retardation syndrome-related protein 1 OS=Mu | 2 | 2 | 3 | 1 | 677 | 76,2 | 6,98 | 1462 | | 2 | | 0,974 | 0,793 | 0,792 | 0,93 | 1,32 | 0,885 | 1,044 | 0,974 | 0,83 | -0,17 | 1,3E-01 |
| Q99K48 | Non-POU domain-containing octamer-binding protein OS=Mus | 20 | 11 | 18 | 8 | 473 | 54,5 | 8,95 | 9487 | 2,57 | 10 | 1 | 0,815 | 0,819 | 1,137 | 1,016 | 1,348 | 0,96 | 1,203 | 1,078 | 0,83 | -0,17 | 1,3E-01 |
| P23506 | Protein-L-isoaspartate(D-aspartate) O-methyltransferase OS=M | 22 | 4 | 6 | 4 | 227 | 24,6 | 7,65 | 3197 | 2,05 | 4 | 1 | 0,952 | 0,958 | 1,021 | 1,031 | 0,853 | 0,979 | 0,972 | 0,904 | 1,07 | 0,07 | 1,3E-01 |
| Q8VHY0 | Chondroitin sulfate proteoglycan 4 OS=Mus musculus OX=1009 | 4 | 6 | 7 | 6 | 2327 | 252,2 | 5,44 | 1232 | 9,4 | 3 | 3 | 1,009 | 1,019 | 1,049 | 1,03 | 1,141 | 1,002 | 1,092 | 1,082 | 0,95 | -0,05 | 1,3E-01 |
| Q8BVA4 | Leiomodin-1 OS=Mus musculus OX=10090 GN=Lmod1 PE=1 SV= | 1 | 1 | 1 | 1 | 595 | 66,3 | 9,19 | 481 | | 1 | | 1,004 | 0,95 | 1,019 | 1,097 | 0,968 | 0,978 | 0,891 | 0,976 | 1,07 | 0,07 | 1,3E-01 |
| Q64176 | Carboxylesterase 1E OS=Mus musculus OX=10090 GN=Ces1e PE | 3 | 1 | 2 | 1 | 562 | 61,5 | 6,15 | 1128 | | 1 | | 1,725 | 1,763 | 1,798 | 1,652 | 2,974 | 1,561 | 2,338 | 2,115 | 0,77 | -0,23 | 1,3E-01 |
| Q9CR00 | 26S proteasome non-ATPase regulatory subunit 9 OS=Mus musc | 30 | 6 | 6 | 6 | 222 | 24,7 | 6,43 | 1941 | 5,87 | 3 | 3 | 1 | 1,044 | 0,966 | 1,003 | 0,793 | 0,945 | 0,996 | 0,951 | 1,09 | 0,09 | 1,3E-01 |
| Q62210 | Baculoviral IAP repeat-containing protein 2 OS=Mus musculus O | 3 | 1 | 1 | 1 | 612 | 69,6 | 6,54 | 148 | | 1 | | 1,205 | 1,03 | 1,086 | 1,303 | 2,53 | 1,181 | 1,312 | 1,767 | 0,68 | -0,32 | 1,3E-01 |
| Q8VE22 | 28S ribosomal protein S23, mitochondrial OS=Mus musculus OX | 30 | 5 | 5 | 5 | 177 | 20,3 | 8,59 | 997 | 4,48 | 3 | 2 | 1,03 | 1,062 | 1,093 | 1,049 | 1,025 | 1,276 | 1,115 | 1,379 | 0,88 | -0,12 | 1,3E-01 |
| G3X9C2 | F-box only protein 50 OS=Mus musculus OX=10090 GN=Nccrp1 | 4 | 1 | 1 | 1 | 266 | 30,4 | 7,12 | | 0 | | 1 | 0,825 | 0,727 | 0,733 | 0,764 | 1,824 | 0,702 | 0,837 | 1,731 | 0,60 | -0,40 | 1,3E-01 |
| Q8CHC4 | Synaptojanin-1 OS=Mus musculus OX=10090 GN=Synj1 PE=1 SV | 18 | 22 | 30 | 22 | 1574 | 172,5 | 6,89 | 11012 | 23,84 | 16 | 8 | 1,094 | 1,145 | 1,035 | 0,935 | 0,947 | 0,983 | 0,983 | 0,977 | 1,08 | 0,08 | 1,3E-01 |
| Q80WG5 | Volume-regulated anion channel subunit LRRC8A OS=Mus musc | 2 | 1 | 1 | 1 | 810 | 94,1 | 7,94 | | 0 | | 1 | 0,881 | 0,823 | 0,654 | 1,08 | 1,051 | 0,999 | 0,921 | 1,286 | 0,81 | -0,19 | 1,3E-01 |
| Q9ESP1 | Stromal cell-derived factor 2-like protein 1 OS=Mus musculus O | 6 | 1 | 1 | 1 | 221 | 23,6 | 7,42 | | 3,58 | | 1 | 2,362 | 2,711 | 1,944 | 2,07 | 4,419 | 2,51 | 2,553 | 2,936 | 0,73 | -0,27 | 1,3E-01 |
| Q9Z0J0 | NPC intracellular cholesterol transporter 2 OS=Mus musculus O | 25 | 3 | 5 | 3 | 149 | 16,4 | 7,68 | 702 | 10,54 | 1 | 3 | 0,976 | 0,941 | 1,176 | 1,455 | 0,901 | 1,004 | 0,989 | 0,712 | 1,26 | 0,26 | 1,3E-01 |
| Q68FE6 | Rho family-interacting cell polarization regulator 1 OS=Mus mus | 1 | 1 | 1 | 1 | 1223 | 132,3 | 5,88 | | 4,21 | | 1 | 1,044 | 0,831 | 1,033 | 1,098 | 1,454 | 1,036 | 1,101 | 1,172 | 0,84 | -0,16 | 1,3E-01 |
| Q8C0E2 | Vacuolar protein sorting-associated protein 26B OS=Mus muscu | 15 | 4 | 8 | 4 | 336 | 39,1 | 7,37 | 2412 | 12,09 | 2 | 3 | 0,799 | 0,845 | 0,811 | 0,788 | 1,056 | 0,834 | 0,833 | 0,894 | 0,90 | -0,10 | 1,3E-01 |
| Q8C7M3 | E3 ubiquitin-protein ligase TRIM9 OS=Mus musculus OX=10090 | 4 | 2 | 2 | 2 | 817 | 90,8 | 7,24 | 435 | 2,66 | 1 | 1 | 0,95 | 0,803 | 0,87 | 0,866 | 1,091 | 0,796 | 1,08 | 1,049 | 0,87 | -0,13 | 1,3E-01 |
| Q99JF5 | Diphosphomevalonate decarboxylase OS=Mus musculus OX=10 | 4 | 2 | 2 | 2 | 401 | 44 | 6,3 | 416 | 1,97 | 1 | 1 | 1,01 | 0,928 | 0,898 | 1,063 | 0,9 | 1,006 | 0,691 | 0,726 | 1,17 | 0,17 | 1,3E-01 |
| Q4LDG0 | Bile acyl-CoA synthetase OS=Mus musculus OX=10090 GN=Slc27 | 1 | 1 | 1 | 1 | 689 | 76,2 | 8,38 | 503 | | 1 | | 0,944 | 0,855 | 0,898 | 0,755 | 0,847 | 0,832 | 0,66 | 0,65 | 1,15 | 0,15 | 1,3E-01 |
| P58389 | Serine/threonine-protein phosphatase 2A activator OS=Mus mu | 7 | 2 | 2 | 2 | 323 | 36,7 | 6,39 | 1281 | | 2 | | 1,056 | 1,024 | 0,779 | 0,935 | 0,989 | 1,143 | 1,136 | 1,03 | 0,88 | -0,12 | 1,4E-01 |
| P06909 | Complement factor H OS=Mus musculus OX=10090 GN=Cfh PE= | 1 | 1 | 1 | 1 | 1234 | 139 | 6,99 | 472 | | 1 | | 1,948 | 1,426 | 1,696 | 1,827 | 4,234 | 1,936 | 1,861 | 2,74 | 0,64 | -0,36 | 1,4E-01 |
| Q6IFX2 | Keratin, type I cytoskeletal 42 OS=Mus musculus OX=10090 GN= | 25 | 11 | 26 | 5 | 452 | 50,1 | 5,16 | 11038 | 10,37 | 7 | 4 | 0,885 | 1,142 | 0,823 | 0,836 | 1,166 | 1,216 | 0,954 | 1,017 | 0,85 | -0,15 | 1,4E-01 |
| O35381 | Acidic leucine-rich nuclear phosphoprotein 32 family member A | 15 | 4 | 7 | 4 | 247 | 28,5 | 4,07 | 3076 | 2,65 | 2 | 2 | 1,067 | 1,098 | 1,038 | 1,223 | 0,879 | 0,992 | 1,097 | 1,035 | 1,11 | 0,11 | 1,4E-01 |
| Q3TTY5 | Keratin, type II cytoskeletal 2 epidermal OS=Mus musculus OX= | 12 | 13 | 28 | 8 | 707 | 70,9 | 8,06 | 10781 | 31,07 | 6 | 7 | 0,987 | 1,29 | 0,683 | 0,838 | 1,199 | 1,168 | 1,376 | 1,055 | 0,79 | -0,21 | 1,4E-01 |
| P07724 | Serum albumin OS=Mus musculus OX=10090 GN=Alb PE=1 SV=3 | 43 | 27 | 117 | 27 | 608 | 68,6 | 6,07 | 46422 | 83,37 | 22 | 9 | 0,963 | 1,134 | 0,93 | 1,051 | 1,443 | 0,972 | 1,232 | 1,169 | 0,85 | -0,15 | 1,4E-01 |
| Q9DCT1 | 1,5-anhydro-D-fructose reductase OS=Mus musculus OX=10090 | 3 | 1 | 1 | 1 | 301 | 34,4 | 7,33 | 699 | | 1 | | 0,829 | 0,722 | 0,917 | 0,622 | 0,932 | 0,763 | 0,967 | 0,995 | 0,84 | -0,16 | 1,4E-01 |
| P52480-2 | Isoform M1 of Pyruvate kinase PKM OS=Mus musculus OX=100 | 41 | 22 | 50 | 21 | 531 | 57,9 | 7,14 | 15306 | 77,9 | 11 | 13 | 0,999 | 0,973 | 0,985 | 1,008 | 0,995 | 1,006 | 1,007 | 1,028 | 0,98 | -0,02 | 1,4E-01 |

| | | | | | | | | | | | | | | | | | | | | | | | |
|---|---|---|---|---|---|---|---|---|---|---|---|---|---|---|---|---|---|---|---|---|---|---|---|
| Q9WV92-8 | Isoform 8 of Band 4.1-like protein 3 OS=Mus musculus OX=1009 | 18 | 17 | 26 | 1 | 1052 | 116,7 | 5,22 | 13244 | 8,77 | 13 | 4 | 0,822 | 1,045 | 1,097 | 1,09 | 0,992 | 1,385 | 1,106 | 1,399 | 0,83 | -0,17 | 1,4E-01 |
| P14873 | Microtubule-associated protein 1B OS=Mus musculus OX=10090 | 45 | 110 | 269 | 109 | 2464 | 270,1 | 4,83 | 103901 | 277,28 | 61 | 61 | 1,077 | 0,937 | 1,063 | 0,976 | 0,763 | 1,002 | 0,931 | 0,937 | 1,12 | 0,12 | 1,4E-01 |
| Q64519 | Syndecan-3 OS=Mus musculus OX=10090 GN=Sdc3 PE=1 SV=2 | 4 | 1 | 3 | 1 | 442 | 46 | 4,59 | 728 | 0 | 1 | 1 | 1,197 | 1,033 | 0,995 | 1,209 | 0,931 | 1,027 | 1,09 | 0,912 | 1,12 | 0,12 | 1,4E-01 |
| Q8BG67 | Protein EFR3 homolog A OS=Mus musculus OX=10090 GN=Efr3a | 1 | 1 | 1 | 1 | 819 | 92,6 | 6,83 | 402 | | 1 | | 1,21 | 1,035 | 1,156 | 1,218 | 0,942 | 1,031 | 1,141 | 1,093 | 1,10 | 0,10 | 1,4E-01 |
| Q9JMH6 | Thioredoxin reductase 1, cytoplasmic OS=Mus musculus OX=10( | 9 | 5 | 8 | 5 | 613 | 67 | 7,44 | 3225 | 8,48 | 2 | 3 | 1,055 | 1,014 | 0,952 | 1,025 | 0,733 | 1,032 | 0,88 | 0,945 | 1,13 | 0,13 | 1,4E-01 |
| P61226 | Ras-related protein Rap-2b OS=Mus musculus OX=10090 GN=Ra | 19 | 4 | 9 | 3 | 183 | 20,5 | 4,81 | 4706 | 2,36 | 3 | 1 | 1,195 | 1,226 | 1,093 | 1,152 | 0,916 | 1,134 | 1,175 | 0,969 | 1,11 | 0,11 | 1,4E-01 |
| Q8BI84 | Transport and Golgi organization protein 1 homolog OS=Mus mu | 0 | 1 | 1 | 1 | 1930 | 213,5 | 4,75 | 506 | | 1 | | 1,054 | 1,073 | 1,093 | 1,001 | 1,202 | 1,057 | 1,223 | 1,067 | 0,93 | -0,07 | 1,4E-01 |
| P61027 | Ras-related protein Rab-10 OS=Mus musculus OX=10090 GN=Ra | 25 | 6 | 19 | 5 | 200 | 22,5 | 8,38 | 2657 | 35,4 | 1 | 5 | 1,113 | 1,002 | 1,047 | 1,087 | 0,704 | 1,018 | 1,025 | 0,959 | 1,15 | 0,15 | 1,4E-01 |
| P39688 | Tyrosine-protein kinase Fyn OS=Mus musculus OX=10090 GN=Fy | 13 | 7 | 11 | 6 | 537 | 60,6 | 6,67 | 5759 | 6,02 | 5 | 2 | 1,033 | 0,964 | 1,187 | 0,97 | 0,879 | 1,012 | 0,918 | 0,942 | 1,11 | 0,11 | 1,4E-01 |
| Q80U63 | Mitofusin-2 OS=Mus musculus OX=10090 GN=Mfn2 PE=1 SV=3 | 6 | 4 | 4 | 4 | 757 | 86,1 | 6,77 | 1674 | 2,3 | 3 | 1 | 1,248 | 1,22 | 1,161 | 1,356 | 1,424 | 1,197 | 1,388 | 1,569 | 0,89 | -0,11 | 1,4E-01 |
| P06837 | Neuromodulin OS=Mus musculus OX=10090 GN=Gap43 PE=1 SV | 89 | 33 | 110 | 33 | 227 | 23,6 | 4,73 | 32762 | 150,6 | 13 | 25 | 0,944 | 0,938 | 1,217 | 1,01 | 0,665 | 1,112 | 0,739 | 0,784 | 1,25 | 0,25 | 1,4E-01 |
| Q99JH7 | Calsyntenin-3 OS=Mus musculus OX=10090 GN=Clstn3 PE=1 SV= | 1 | 1 | 1 | 1 | 956 | 105,8 | 5,34 | 185 | | 1 | | 0,931 | 0,976 | 0,965 | 1,021 | 0,989 | 1,06 | 1,053 | 0,981 | 0,95 | -0,05 | 1,4E-01 |
| O35295 | Transcriptional activator protein Pur-beta OS=Mus musculus OX | 23 | 6 | 10 | 6 | 324 | 33,9 | 5,43 | 5869 | 0 | 5 | 1 | 0,971 | 0,988 | 0,982 | 0,98 | 1,161 | 1,021 | 0,98 | 1,028 | 0,94 | -0,06 | 1,4E-01 |
| P61028 | Ras-related protein Rab-8B OS=Mus musculus OX=10090 GN=Ra | 13 | 3 | 10 | 1 | 207 | 23,6 | 9,07 | 5124 | 4,98 | 2 | 1 | 1,068 | 0,926 | 0,881 | 0,989 | 0,987 | 0,863 | 0,816 | 0,795 | 1,12 | 0,12 | 1,4E-01 |
| A6H6A9 | Rab GTPase-activating protein 1-like OS=Mus musculus OX=100 | 2 | 2 | 2 | 1 | 815 | 92,3 | 5,31 | | 3,95 | | 2 | 1,314 | 1,176 | 1,22 | 1,166 | 0,779 | 1,022 | 1,198 | 1,179 | 1,17 | 0,17 | 1,4E-01 |
| Q3TLH4 | Protein PRRC2C OS=Mus musculus OX=10090 GN=Prrc2c PE=1 S | 1 | 3 | 3 | 3 | 2846 | 310,7 | 9,1 | 526 | 0 | 2 | 1 | 0,966 | 0,957 | 0,961 | 0,86 | 0,844 | 1,242 | 1,086 | 1,419 | 0,82 | -0,18 | 1,4E-01 |
| P32020 | Non-specific lipid-transfer protein OS=Mus musculus OX=10090 | 17 | 10 | 29 | 10 | 547 | 59,1 | 7,44 | 12063 | 13,45 | 8 | 5 | 1,025 | 1,13 | 1,065 | 1,122 | 1,299 | 1,077 | 1,128 | 1,208 | 0,92 | -0,08 | 1,4E-01 |
| P30115 | Glutathione S-transferase A3 OS=Mus musculus OX=10090 GN= | 9 | 2 | 2 | 2 | 221 | 25,3 | 8,73 | 734 | 2,77 | 1 | 1 | 2,095 | 3,356 | 2,625 | 2,595 | 4,11 | 2,514 | 3,259 | 3,7 | 0,79 | -0,21 | 1,4E-01 |
| Q80TE7 | Leucine-rich repeat-containing protein 7 OS=Mus musculus OX= | 2 | 3 | 5 | 3 | 1490 | 166,8 | 6,61 | 2063 | | 3 | | 0,913 | 1,04 | 0,943 | 1,088 | 0,883 | 0,903 | 0,898 | 0,983 | 1,09 | 0,09 | 1,4E-01 |
| B0V2N1-3 | Isoform 3 of Receptor-type tyrosine-protein phosphatase S OS= | 10 | 14 | 17 | 10 | 1501 | 168,2 | 6,81 | 4940 | 20,66 | 7 | 7 | 0,982 | 0,941 | 0,907 | 0,944 | 1,113 | 0,958 | 1,039 | 0,947 | 0,93 | -0,07 | 1,4E-01 |
| P61205 | ADP-ribosylation factor 3 OS=Mus musculus OX=10090 GN=Arf3 | 47 | 8 | 17 | 1 | 181 | 20,6 | 7,43 | 8294 | 6,44 | 6 | 3 | 0,912 | 0,918 | 0,828 | 1,053 | 0,864 | 0,717 | 0,776 | 0,917 | 1,13 | 0,13 | 1,4E-01 |
| Q9WV54 | Acid ceramidase OS=Mus musculus OX=10090 GN=Asah1 PE=1 S | 5 | 2 | 3 | 2 | 394 | 44,6 | 8,46 | 565 | 2,26 | 1 | 1 | 0,779 | 0,969 | 0,712 | 0,856 | 1,694 | 0,838 | 1,173 | 0,951 | 0,71 | -0,29 | 1,4E-01 |
| P70336 | Rho-associated protein kinase 2 OS=Mus musculus OX=10090 G | 6 | 7 | 7 | 7 | 1388 | 160,5 | 5,99 | 2553 | 8,5 | 4 | 3 | 1,065 | 1,072 | 1,046 | 0,983 | 0,882 | 0,997 | 1,039 | 0,984 | 1,07 | 0,07 | 1,4E-01 |
| P29788 | Vitronectin OS=Mus musculus OX=10090 GN=Vtn PE=1 SV=2 | 3 | 1 | 1 | 1 | 478 | 54,8 | 5,88 | 627 | | 1 | | 1,243 | 1,224 | 1,472 | 1,72 | 2,919 | 1,331 | 1,839 | 1,95 | 0,70 | -0,30 | 1,4E-01 |
| A2ALU4 | Protein Shroom2 OS=Mus musculus OX=10090 GN=Shroom2 PE | 7 | 8 | 10 | 8 | 1481 | 164,6 | 6,7 | 6079 | 2,94 | 7 | 1 | 1,107 | 1,072 | 1,026 | 1,193 | 0,941 | 1,078 | 0,96 | 1,075 | 1,08 | 0,08 | 1,4E-01 |
| Q9ERB0 | Synaptosomal-associated protein 29 OS=Mus musculus OX=100 | 24 | 6 | 6 | 6 | 260 | 29,6 | 5,38 | 2455 | 3,55 | 4 | 2 | 0,923 | 0,953 | 0,98 | 0,94 | 0,749 | 0,899 | 0,918 | 0,932 | 1,09 | 0,09 | 1,4E-01 |
| Q9QXX4 | Calcium-binding mitochondrial carrier protein Aralar2 OS=Mus r | 7 | 4 | 4 | 2 | 676 | 74,4 | 8,6 | 1460 | 2,63 | 3 | 1 | 1,01 | 0,618 | 1,127 | 0,82 | 1,202 | 1,095 | 1,079 | 1,003 | 0,82 | -0,18 | 1,4E-01 |
| Q61792 | LIM and SH3 domain protein 1 OS=Mus musculus OX=10090 GN | 48 | 14 | 21 | 13 | 263 | 30 | 7,05 | 7128 | 23,28 | 7 | 7 | 1,023 | 0,962 | 1,124 | 1,008 | 0,763 | 0,988 | 1,007 | 0,919 | 1,12 | 0,12 | 1,4E-01 |
| Q9D1B9 | 39S ribosomal protein L28, mitochondrial OS=Mus musculus OX | 5 | 1 | 1 | 1 | 257 | 30,2 | 9,29 | 592 | | 1 | | 1,12 | 1,014 | 1,007 | 1,054 | 1,693 | 0,862 | 1,433 | 1,399 | 0,78 | -0,22 | 1,4E-01 |
| Q60575 | Kinesin-like protein KIF1B OS=Mus musculus OX=10090 GN=Kif1 | 5 | 8 | 9 | 2 | 1816 | 204 | 5,62 | 4394 | 2,43 | 7 | 1 | 0,788 | 0,815 | 1,082 | 0,999 | 1,223 | 1,013 | 1,047 | 0,997 | 0,86 | -0,14 | 1,4E-01 |
| Q9JL56 | Glycerophosphodiester phosphodiesterase 1 OS=Mus musculus | 9 | 3 | 5 | 3 | 331 | 37,6 | 6,9 | 3275 | | 3 | | 0,907 | 0,966 | 0,941 | 1,005 | 1 | 0,946 | 1,028 | 1,057 | 0,95 | -0,05 | 1,4E-01 |
| Q91VK4 | Integral membrane protein 2C OS=Mus musculus OX=10090 GN | 4 | 1 | 1 | 1 | 269 | 30,5 | 8,59 | 682 | | 1 | | 1,226 | 1,235 | 1,196 | 1,11 | 0,75 | 1,071 | 1,11 | 1,172 | 1,16 | 0,16 | 1,4E-01 |
| Q9WUM4 | Coronin-1C OS=Mus musculus OX=10090 GN=Coro1c PE=1 SV=2 | 30 | 18 | 24 | 16 | 474 | 53,1 | 7,08 | 5607 | 42,15 | 8 | 10 | 1,006 | 0,95 | 1,035 | 0,928 | 0,887 | 0,988 | 0,825 | 0,933 | 1,08 | 0,08 | 1,4E-01 |
| Q8CI75 | DIS3-like exonuclease 2 OS=Mus musculus OX=10090 GN=Dis3l2 | 3 | 2 | 3 | 2 | 870 | 97,7 | 5,9 | 722 | 5,21 | 1 | 1 | 0,971 | 1,061 | 1,08 | 1,045 | 0,82 | 0,94 | 1,084 | 0,911 | 1,11 | 0,11 | 1,4E-01 |
| Q8R0N6 | Hydroxyacid-oxoacid transhydrogenase, mitochondrial OS=Mus | 3 | 1 | 1 | 1 | 465 | 49,9 | 7,58 | 409 | | 1 | | 0,867 | 0,855 | 0,848 | 1,084 | 1,22 | 0,791 | 1,189 | 1,377 | 0,80 | -0,20 | 1,4E-01 |
| Q91WQ3 | Tyrosine--tRNA ligase, cytoplasmic OS=Mus musculus OX=10090 | 40 | 20 | 25 | 20 | 528 | 59,1 | 7,01 | 10532 | 16,2 | 15 | 7 | 1,044 | 0,995 | 0,969 | 1,019 | 0,859 | 0,984 | 0,938 | 1,004 | 1,06 | 0,06 | 1,4E-01 |
| Q61016 | Guanine nucleotide-binding protein G(I)/G(S)/G(O) subunit gam | 31 | 2 | 5 | 2 | 68 | 7,5 | 8,51 | 2590 | 3,2 | 1 | 1 | 1,098 | 0,825 | 1,137 | 0,903 | 0,789 | 0,979 | 0,783 | 0,818 | 1,18 | 0,18 | 1,4E-01 |
| Q3TKT4 | Transcription activator BRG1 OS=Mus musculus OX=10090 GN= | 1 | 1 | 1 | 1 | 1613 | 181,3 | 8 | 228 | | 1 | | 0,732 | 1,074 | 0,843 | 1,048 | 1,41 | 1,149 | 1,102 | 0,916 | 0,81 | -0,19 | 1,4E-01 |
| Q9CR62 | Mitochondrial 2-oxoglutarate/malate carrier protein OS=Mus m | 24 | 6 | 7 | 6 | 314 | 34,1 | 9,94 | 2755 | 5,15 | 3 | 3 | 0,983 | 0,992 | 0,993 | 1,097 | 0,976 | 0,949 | 0,893 | 1,005 | 1,06 | 0,06 | 1,4E-01 |
| O88569 | Heterogeneous nuclear ribonucleoproteins A2/B1 OS=Mus musc | 65 | 19 | 35 | 18 | 353 | 37,4 | 8,95 | 12609 | 44,93 | 12 | 9 | 0,85 | 0,832 | 1,019 | 1,076 | 1,095 | 0,952 | 1,167 | 1,07 | 0,88 | -0,12 | 1,4E-01 |
| Q9DCU9 | 4-hydroxy-2-oxoglutarate aldolase, mitochondrial OS=Mus musc | 5 | 1 | 1 | 1 | 321 | 34,6 | 7,72 | 796 | | 1 | | 1,323 | 1,368 | 1,018 | 1,317 | 2,317 | 1,071 | 1,646 | 1,795 | 0,74 | -0,26 | 1,4E-01 |
| P00520 | Tyrosine-protein kinase ABL1 OS=Mus musculus OX=10090 GN= | 1 | 1 | 1 | 1 | 1123 | 122,6 | 8,65 | 388 | | 1 | | 1,07 | 1,109 | 0,956 | 1,076 | 1,519 | 0,924 | 1,249 | 1,427 | 0,82 | -0,18 | 1,4E-01 |
| Q9QZ25 | Vascular non-inflammatory molecule 3 OS=Mus musculus OX=1( | 2 | 1 | 3 | 1 | 500 | 56,3 | 6,29 | | 8,7 | | 1 | 0,9 | 1,026 | 0,868 | 0,968 | 1,332 | 0,821 | 1,156 | 1,234 | 0,83 | -0,17 | 1,4E-01 |
| Q9JHI5 | Isovaleryl-CoA dehydrogenase, mitochondrial OS=Mus musculus | 19 | 8 | 13 | 8 | 424 | 46,3 | 8,29 | 5506 | 8 | 5 | 4 | 0,911 | 0,862 | 1,416 | 0,927 | 0,671 | 0,871 | 0,85 | 0,803 | 1,29 | 0,29 | 1,4E-01 |
| O88343 | Electrogenic sodium bicarbonate cotransporter 1 OS=Mus musc | 1 | 2 | 2 | 2 | 1079 | 121,4 | 6,84 | | 4,45 | | 2 | 1,046 | 0,873 | 1,101 | 1,001 | 0,74 | 0,844 | 1,061 | 0,808 | 1,16 | 0,16 | 1,4E-01 |
| F8VPN2 | Testis-expressed protein 15 OS=Mus musculus OX=10090 GN=T | 0 | 1 | 1 | 1 | 2785 | 311,1 | 6,39 | | 0 | | 1 | 0,941 | 1,236 | 1,175 | 1,01 | 1,001 | 0,972 | 0,871 | 1,009 | 1,13 | 0,13 | 1,4E-01 |
| Q91WD5 | NADH dehydrogenase [ubiquinone] iron-sulfur protein 2, mitoch | 22 | 9 | 15 | 9 | 463 | 52,6 | 6,99 | 5196 | 0 | 7 | 2 | 0,982 | 1,053 | 0,893 | 0,963 | 1,251 | 0,974 | 0,985 | 1,324 | 0,86 | -0,14 | 1,5E-01 |
| Q01097 | Glutamate receptor ionotropic, NMDA 2B OS=Mus musculus OX | 1 | 2 | 2 | 2 | 1482 | 165,9 | 6,87 | 884 | | 2 | | 0,812 | 0,885 | 0,688 | 0,82 | 1,333 | 0,812 | 0,966 | 0,906 | 0,80 | -0,20 | 1,5E-01 |
| P24270 | Catalase OS=Mus musculus OX=10090 GN=Cat PE=1 SV=4 | 48 | 23 | 43 | 23 | 527 | 59,8 | 7,88 | 20455 | 12,03 | 20 | 4 | 1,103 | 1,139 | 0,996 | 1,131 | 1,367 | 1,132 | 1,152 | 1,151 | 0,91 | -0,09 | 1,5E-01 |
| Q60764 | Probable E3 ubiquitin-protein ligase makorin-3 OS=Mus musculu | 1 | 1 | 1 | 1 | 544 | 59,4 | 6,64 | | 2,25 | | 1 | 1,022 | 1,026 | 1,028 | 1,174 | 0,862 | 1,012 | 1,049 | 0,959 | 1,09 | 0,09 | 1,5E-01 |
| O09114 | Prostaglandin-H2 D-isomerase OS=Mus musculus OX=10090 GN | 10 | 3 | 4 | 3 | 189 | 21,1 | 8,25 | 661 | 0 | 1 | 2 | 1,038 | 0,972 | 0,831 | 1,641 | 0,898 | 0,773 | 0,859 | 0,73 | 1,37 | 0,37 | 1,5E-01 |
| P06797 | Cathepsin L1 OS=Mus musculus OX=10090 GN=Ctsl PE=1 SV=2 | 10 | 2 | 3 | 2 | 334 | 37,5 | 6,83 | 795 | 3,15 | 2 | 1 | 0,953 | 1,106 | 0,999 | 0,991 | 0,919 | 0,861 | 1,036 | 0,899 | 1,09 | 0,09 | 1,5E-01 |
| Q8K341 | Alpha-tubulin N-acetyltransferase 1 OS=Mus musculus OX=1009 | 33 | 13 | 30 | 13 | 421 | 47,1 | 9,98 | 16841 | 2,2 | 11 | 2 | 1,005 | 1,043 | 1,131 | 1,034 | 1,001 | 1,045 | 0,985 | 0,972 | 1,05 | 0,05 | 1,5E-01 |
| O89086 | RNA-binding protein 3 OS=Mus musculus OX=10090 GN=Rbm3 I | 27 | 3 | 7 | 3 | 153 | 16,6 | 7,5 | 2841 | 5,2 | 2 | 2 | 1,032 | 1,088 | 0,949 | 1,095 | 0,92 | 1,054 | 0,966 | 0,91 | 1,08 | 0,08 | 1,5E-01 |
| P70296 | Phosphatidylethanolamine-binding protein 1 OS=Mus musculus | 45 | 6 | 22 | 6 | 187 | 20,8 | 5,4 | 8927 | 28,66 | 6 | 3 | 0,981 | 1,001 | 0,932 | 1,027 | 0,776 | 1,017 | 0,907 | 0,886 | 1,10 | 0,10 | 1,5E-01 |
| Q9WVJ3 | Carboxypeptidase Q OS=Mus musculus OX=10090 GN=Cpq PE=1 | 4 | 2 | 2 | 2 | 470 | 51,8 | 6,21 | 1374 | | 2 | | 0,76 | 0,822 | 0,788 | 0,874 | 0,83 | 0,802 | 1,069 | 1,012 | 0,87 | -0,13 | 1,5E-01 |
| Q8BRJ3 | RELT-like protein 2 OS=Mus musculus OX=10090 GN=Rell2 PE=1 | 7 | 1 | 1 | 1 | 303 | 32,3 | 6,18 | 196 | | 1 | | 0,875 | 0,986 | 0,899 | 1,068 | 1,171 | 1,305 | 0,882 | 1,126 | 0,85 | -0,15 | 1,5E-01 |
| P16054 | Protein kinase C epsilon type OS=Mus musculus OX=10090 GN= | 19 | 10 | 15 | 10 | 737 | 83,5 | 7,03 | 3911 | 17,87 | 6 | 5 | 0,948 | 0,96 | 1,065 | 0,999 | 1,175 | 0,947 | 1,175 | 1,075 | 0,91 | -0,09 | 1,5E-01 |
| Q8BYN5 | FSD1-like protein OS=Mus musculus OX=10090 GN=Fsd1l PE=1 S | 21 | 9 | 12 | 9 | 507 | 56,9 | 6,7 | 1421 | 21,16 | 3 | 6 | 0,998 | 0,95 | 1,024 | 0,969 | 0,722 | 0,999 | 0,906 | 0,912 | 1,11 | 0,11 | 1,5E-01 |
| Q99L13 | 3-hydroxyisobutyrate dehydrogenase, mitochondrial OS=Mus m | 5 | 2 | 3 | 2 | 335 | 35,4 | 8,13 | 1529 | 3,41 | 1 | 1 | 1,004 | 1,058 | 1,023 | 1,077 | 0,935 | 0,926 | 0,966 | 1,078 | 1,07 | 0,07 | 1,5E-01 |
| P47757-4 | Isoform 3 of F-actin-capping protein subunit beta OS=Mus musc | 44 | 14 | 24 | 4 | 301 | 33,7 | 6,43 | 6942 | 28,67 | 10 | 7 | 1,223 | 1,096 | 1,064 | 1,123 | 0,713 | 1,083 | 1,115 | 0,947 | 1,17 | 0,17 | 1,5E-01 |
| Q91ZR1 | Ras-related protein Rab-4B OS=Mus musculus OX=10090 GN=Ra | 12 | 2 | 2 | 2 | 213 | 23,6 | 6,06 | 787 | | 2 | | 0,782 | 0,833 | 0,777 | 0,741 | 1,13 | 0,683 | 0,91 | 1,127 | 0,81 | -0,19 | 1,5E-01 |
| P83510 | Traf2 and NCK-interacting protein kinase OS=Mus musculus OX= | 14 | 13 | 16 | 9 | 1323 | 150,3 | 7,27 | 7198 | 8,37 | 10 | 3 | 0,816 | 0,871 | 0,844 | 0,942 | 1,035 | 0,833 | 0,957 | 0,984 | 0,91 | -0,09 | 1,5E-01 |
| P63080 | Gamma-aminobutyric acid receptor subunit beta-3 OS=Mus mu | 12 | 4 | 4 | 4 | 473 | 54,1 | 9,1 | 1793 | | 4 | | 0,952 | 0,856 | 0,925 | 1,039 | 1,708 | 0,978 | 1,05 | 1,161 | 0,77 | -0,23 | 1,5E-01 |
| Q9CXZ1 | NADH dehydrogenase [ubiquinone] iron-sulfur protein 4, mitoch | 29 | 5 | 5 | 5 | 175 | 19,8 | 9,99 | 2877 | 2,01 | 4 | 1 | 0,975 | 0,946 | 0,915 | 1,033 | 0,784 | 0,911 | 0,941 | 0,937 | 1,08 | 0,08 | 1,5E-01 |

| | | | | | | | | | | | | | | | | | | | | | | | |
|---|---|---|---|---|---|---|---|---|---|---|---|---|---|---|---|---|---|---|---|---|---|---|---|
| P28184 | Metallothionein-3 OS=Mus musculus OX=10090 GN=Mt3 PE=1 S | 34 | 2 | 2 | 2 | 68 | 7 | 7,47 | | 4,93 | | 2 | 1,212 | 1,001 | 1,249 | 0,846 | 0,521 | 1,218 | 0,77 | 0,586 | 1,39 | 0,39 | 1,5E-01 |
| Q9D0K2 | Succinyl-CoA:3-ketoacid coenzyme A transferase 1, mitochondri | 34 | 15 | 54 | 15 | 520 | 56 | 8,53 | 26839 | 18,68 | 11 | 6 | 0,935 | 0,941 | 0,945 | 0,889 | 1,154 | 0,973 | 0,928 | 0,992 | 0,92 | -0,08 | 1,5E-01 |
| P14094 | Sodium/potassium-transporting ATPase subunit beta-1 OS=Mus | 39 | 15 | 40 | 15 | 304 | 35,2 | 8,65 | 17166 | 32,1 | 10 | 8 | 1,013 | 1,008 | 1,124 | 0,946 | 0,744 | 0,966 | 1,01 | 0,915 | 1,13 | 0,13 | 1,5E-01 |
| Q80ZX8 | Sperm-associated antigen 1 OS=Mus musculus OX=10090 GN=Sp | 1 | 1 | 1 | 1 | 901 | 100,6 | 5,62 | 264 | | | 1 | 1,077 | 1,099 | 0,923 | 1,089 | 0,473 | 0,973 | 1,033 | 0,836 | 1,26 | 0,26 | 1,5E-01 |
| Q91Y97 | Fructose-bisphosphate aldolase B OS=Mus musculus OX=10090 | 38 | 10 | 23 | 9 | 364 | 39,5 | 8,27 | 6418 | 28,41 | 7 | 4 | 1,568 | 1,511 | 1,429 | 1,754 | 2,472 | 1,554 | 1,797 | 1,815 | 0,82 | -0,18 | 1,5E-01 |
| P49443 | Protein phosphatase 1A OS=Mus musculus OX=10090 GN=Ppm1 | 13 | 6 | 6 | 5 | 382 | 42,4 | 5,36 | 1929 | 5,31 | 3 | 3 | 0,926 | 0,908 | 1,007 | 0,917 | 1,007 | 1,014 | 0,923 | 1,036 | 0,94 | -0,06 | 1,5E-01 |
| E9Q6B2 | Coiled-coil domain-containing protein 85C OS=Mus musculus OX | 5 | 1 | 1 | 1 | 420 | 45,3 | 6,96 | 455 | | | 1 | 1,495 | 1,5 | 1,567 | 1,975 | 1,285 | 1,432 | 1,564 | 1,411 | 1,15 | 0,15 | 1,5E-01 |
| P63328-2 | Isoform 2 of Serine/threonine-protein phosphatase 2B catalytic | 33 | 15 | 51 | 10 | 511 | 57,6 | 6,3 | 25041 | 31,68 | 8 | 9 | 1,029 | 1,013 | 0,983 | 0,954 | 0,748 | 0,96 | 0,952 | 0,958 | 1,10 | 0,10 | 1,5E-01 |
| P58252 | Elongation factor 2 OS=Mus musculus OX=10090 GN=Eef2 PE=1 | 31 | 27 | 69 | 26 | 858 | 95,3 | 6,83 | 35166 | 18,96 | 20 | 10 | 1,003 | 0,99 | 0,981 | 1,017 | 0,777 | 0,979 | 0,953 | 0,965 | 1,09 | 0,09 | 1,5E-01 |
| Q99JP4 | Anaphase-promoting complex subunit CDC26 OS=Mus musculus | 26 | 1 | 2 | 1 | 85 | 9,8 | 6,81 | 343 | | | 1 | 0,749 | 1,011 | 0,756 | 0,712 | 0,997 | 0,785 | 0,978 | 1,078 | 0,84 | -0,16 | 1,5E-01 |
| Q3UQ84 | Threonine--tRNA ligase, mitochondrial OS=Mus musculus OX=10 | 5 | 4 | 4 | 4 | 723 | 81,6 | 7,87 | 1302 | 2,28 | 3 | 1 | 0,995 | 1,117 | 1,104 | 1,122 | 1,109 | 1,095 | 1,201 | 1,284 | 0,93 | -0,07 | 1,5E-01 |
| P05063 | Fructose-bisphosphate aldolase C OS=Mus musculus OX=10090 | 50 | 23 | 36 | 18 | 363 | 39,4 | 7,12 | 11661 | 44,87 | 15 | 12 | 0,959 | 1,018 | 0,929 | 1,042 | 0,964 | 1,114 | 1,044 | 1,21 | 0,91 | -0,09 | 1,5E-01 |
| P51660 | Peroxisomal multifunctional enzyme type 2 OS=Mus musculus C | 33 | 19 | 28 | 19 | 735 | 79,4 | 8,57 | 11228 | 21,92 | 16 | 5 | 1,141 | 1,184 | 1,133 | 1,216 | 1,129 | 1,157 | 1,069 | 1,14 | 1,04 | 0,04 | 1,5E-01 |
| O88487 | Cytoplasmic dynein 1 intermediate chain 2 OS=Mus musculus O) | 16 | 8 | 9 | 7 | 612 | 68,4 | 5,29 | 4829 | 7,68 | 5 | 3 | 1,121 | 1,015 | 1,086 | 1,097 | 0,724 | 1,027 | 1,089 | 0,934 | 1,14 | 0,14 | 1,5E-01 |
| A2A7S8 | Uncharacterized protein KIAA1522 OS=Mus musculus OX=10090 | 2 | 1 | 1 | 1 | 1013 | 104,8 | 9,69 | 122 | | | 1 | 0,889 | 0,722 | 0,687 | 0,629 | 1,03 | 0,781 | 0,739 | 0,964 | 0,83 | -0,17 | 1,5E-01 |
| Q8CJ19 | [F-actin]-monooxygenase MICAL3 OS=Mus musculus OX=10090 | 7 | 12 | 13 | 12 | 1993 | 223,6 | 5,47 | 6465 | 6,79 | 9 | 3 | 1,035 | 0,986 | 1,068 | 1,063 | 0,925 | 1,018 | 0,997 | 1,021 | 1,05 | 0,05 | 1,5E-01 |
| P14824 | Annexin A6 OS=Mus musculus OX=10090 GN=Anxa6 PE=1 SV=3 | 28 | 18 | 25 | 18 | 673 | 75,8 | 5,5 | 10734 | 18,24 | 13 | 7 | 0,995 | 1,005 | 0,981 | 1,042 | 0,912 | 0,976 | 1,017 | 0,951 | 1,04 | 0,04 | 1,5E-01 |
| P22315 | Ferrochelatase, mitochondrial OS=Mus musculus OX=10090 GN= | 12 | 5 | 8 | 5 | 420 | 47,1 | 8,91 | 3478 | 6,05 | 3 | 2 | 0,98 | 1,094 | 0,955 | 0,995 | 1,149 | 1,07 | 0,981 | 1,267 | 0,90 | -0,10 | 1,5E-01 |
| Q9JLJ2 | 4-trimethylaminobutyraldehyde dehydrogenase OS=Mus muscu | 24 | 9 | 19 | 9 | 494 | 53,5 | 6,98 | 5027 | 32,48 | 6 | 3 | 0,932 | 1,075 | 1,131 | 1,151 | 1,188 | 1,134 | 1,151 | 1,147 | 0,93 | -0,07 | 1,5E-01 |
| Q9CZ30 | Obg-like ATPase 1 OS=Mus musculus OX=10090 GN=Ola1 PE=1 S | 20 | 8 | 12 | 8 | 396 | 44,7 | 7,81 | 4288 | 11,16 | 5 | 3 | 1,063 | 0,997 | 1,06 | 1,041 | 0,753 | 1,043 | 0,937 | 1 | 1,11 | 0,11 | 1,5E-01 |
| Q9JKD3 | Secretory carrier-associated membrane protein 5 OS=Mus musc | 9 | 2 | 10 | 2 | 235 | 26,1 | 8,54 | 5278 | 0 | 2 | 1 | 1,016 | 1,028 | 1,024 | 0,931 | 0,892 | 0,951 | 0,907 | 1,015 | 1,06 | 0,06 | 1,5E-01 |
| Q9CYH2 | Peroxiredoxin-like 2A OS=Mus musculus OX=10090 GN=Prxl2a P | 17 | 4 | 5 | 4 | 218 | 24,4 | 9,17 | 1019 | 5,47 | 2 | 2 | 0,941 | 0,949 | 0,96 | 0,956 | 0,81 | 0,924 | 0,955 | 0,911 | 1,06 | 0,06 | 1,5E-01 |
| Q8R1F1 | Protein Niban 2 OS=Mus musculus OX=10090 GN=Niban2 PE=1 S | 1 | 1 | 2 | 1 | 749 | 84,8 | 5,94 | 1059 | | | 1 | 1,14 | 1,223 | 1,056 | 1,211 | 1,082 | 1,321 | 1,415 | 1,761 | 0,83 | -0,17 | 1,6E-01 |
| Q8K2C9 | Very-long-chain (3R)-3-hydroxyacyl-CoA dehydratase 3 OS=Mus | 5 | 2 | 2 | 2 | 362 | 43,1 | 9,13 | 819 | 0 | 1 | 1 | 1,271 | 1,265 | 1,435 | 1,249 | 1,302 | 1,237 | 1,185 | 1,084 | 1,09 | 0,09 | 1,6E-01 |
| Q920R0 | Alsin OS=Mus musculus OX=10090 GN=Als2 PE=1 SV=3 | 3 | 3 | 3 | 3 | 1651 | 182,5 | 6,3 | 445 | 2,84 | 2 | 1 | 1,034 | 1,051 | 1,185 | 1,059 | 1,718 | 1,136 | 1,166 | 1,226 | 0,83 | -0,17 | 1,6E-01 |
| Q99P47 | Contactin-associated protein-like 4 OS=Mus musculus OX=10090 | 1 | 1 | 1 | 1 | 1310 | 144,6 | 7,06 | | 2,78 | | 1 | 0,765 | 0,696 | 0,636 | 0,92 | 0,797 | 0,947 | 0,883 | 0,839 | 0,87 | -0,13 | 1,6E-01 |
| P11725 | Ornithine carbamoyltransferase, mitochondrial OS=Mus muscul | 7 | 2 | 2 | 2 | 354 | 39,7 | 8,73 | | 2,98 | | 2 | 0,995 | 1,475 | 1,239 | 1,346 | 2,325 | 1,312 | 1,783 | 1,325 | 0,75 | -0,25 | 1,6E-01 |
| O09173 | Homogentisate 1,2-dioxygenase OS=Mus musculus OX=10090 G | 2 | 1 | 1 | 1 | 445 | 49,9 | 7,24 | | 0 | | 1 | 1,007 | 0,865 | 0,936 | 0,798 | 1,859 | 1,024 | 0,892 | 1,252 | 0,72 | -0,28 | 1,6E-01 |
| Q9DCR2 | AP-3 complex subunit sigma-1 OS=Mus musculus OX=10090 GN | 10 | 2 | 2 | 2 | 193 | 21,7 | 5,39 | 843 | 3,21 | 1 | 1 | 1,052 | 0,972 | 1,148 | 0,94 | 0,422 | 1,048 | 0,868 | 0,859 | 1,29 | 0,29 | 1,6E-01 |
| Q6NS60 | F-box only protein 41 OS=Mus musculus OX=10090 GN=Fbxo41 | 9 | 6 | 7 | 5 | 873 | 94,3 | 8,24 | 2658 | 7,22 | 5 | 2 | 0,945 | 0,986 | 1,1 | 1,097 | 1,442 | 1,023 | 1,047 | 1,349 | 0,85 | -0,15 | 1,6E-01 |
| Q60864 | Stress-induced-phosphoprotein 1 OS=Mus musculus OX=10090 | 29 | 21 | 28 | 21 | 543 | 62,5 | 6,8 | 11254 | 24,6 | 12 | 10 | 1,053 | 0,924 | 1,106 | 1,001 | 0,78 | 1,011 | 0,946 | 0,941 | 1,11 | 0,11 | 1,6E-01 |
| Q9D0L7 | Armadillo repeat-containing protein 10 OS=Mus musculus OX=1 | 19 | 5 | 8 | 5 | 306 | 33,3 | 7,99 | 4038 | 6,71 | 3 | 2 | 1,056 | 1,049 | 1,073 | 0,986 | 0,936 | 1,041 | 0,974 | 1,018 | 1,05 | 0,05 | 1,6E-01 |
| Q8K4L3 | Supervillin OS=Mus musculus OX=10090 GN=Svil PE=1 SV=1 | 0 | 1 | 1 | 1 | 2170 | 243 | 6,87 | 421 | | | 1 | 0,948 | 0,971 | 0,866 | 0,901 | 1,368 | 0,921 | 0,903 | 1,314 | 0,82 | -0,18 | 1,6E-01 |
| P40142 | Transketolase OS=Mus musculus OX=10090 GN=Tkt PE=1 SV=1 | 41 | 26 | 59 | 25 | 623 | 67,6 | 7,5 | 21696 | 65,46 | 14 | 14 | 1,053 | 1,014 | 0,961 | 0,993 | 0,8 | 0,97 | 0,998 | 0,942 | 1,08 | 0,08 | 1,6E-01 |
| P11404 | Fatty acid-binding protein, heart OS=Mus musculus OX=10090 G | 43 | 6 | 10 | 6 | 133 | 14,8 | 6,57 | 2048 | 19,14 | 1 | 5 | 0,932 | 0,968 | 0,923 | 1,016 | 0,632 | 1,038 | 0,841 | 0,761 | 1,17 | 0,17 | 1,6E-01 |
| P0C027 | Diphosphoinositol polyphosphate phosphohydrolase 3-alpha OS | 25 | 4 | 6 | 3 | 164 | 18,6 | 5,69 | 2733 | 2,05 | 3 | 1 | 1,039 | 0,937 | 0,855 | 1,055 | 0,495 | 1,046 | 0,8 | 0,755 | 1,26 | 0,26 | 1,6E-01 |
| P51791 | H(+)/Cl(-) exchange transporter 3 OS=Mus musculus OX=10090 ( | 1 | 1 | 2 | 1 | 818 | 90,8 | 6,28 | 803 | | | 1 | 0,983 | 1,004 | 1,015 | 0,885 | 1,196 | 0,999 | 1,077 | 0,983 | 0,91 | -0,09 | 1,6E-01 |
| O08997 | Copper transport protein ATOX1 OS=Mus musculus OX=10090 G | 28 | 1 | 1 | 1 | 68 | 7,3 | 6,51 | | 3,92 | | 1 | 0,726 | 0,746 | 0,782 | 0,866 | 0,844 | 1,086 | 0,935 | 0,754 | 0,86 | -0,14 | 1,6E-01 |
| Q8C0L0 | Thioredoxin-related transmembrane protein 4 OS=Mus musculu | 9 | 2 | 2 | 2 | 335 | 37,1 | 4,37 | 782 | 2,36 | 1 | 1 | 0,826 | 0,979 | 0,879 | 1,033 | 0,644 | 0,828 | 0,939 | 0,81 | 1,15 | 0,15 | 1,6E-01 |
| P00329 | Alcohol dehydrogenase 1 OS=Mus musculus OX=10090 GN=Adh | 7 | 2 | 4 | 2 | 375 | 39,7 | 8,1 | 3065 | | | 2 | 1,813 | 1,988 | 1,623 | 1,765 | 3,481 | 1,864 | 1,912 | 2,403 | 0,74 | -0,26 | 1,6E-01 |
| Q91YM2 | Rho GTPase-activating protein 35 OS=Mus musculus OX=10090 ( | 8 | 12 | 17 | 12 | 1499 | 170,3 | 6,61 | 5528 | 7,57 | 6 | 7 | 1,007 | 0,874 | 0,978 | 0,892 | 0,883 | 0,923 | 0,861 | 0,852 | 1,07 | 0,07 | 1,6E-01 |
| P16460 | Argininosuccinate synthase OS=Mus musculus OX=10090 GN=As | 17 | 7 | 8 | 7 | 412 | 46,6 | 8,22 | 2516 | 5,8 | 5 | 2 | 1,097 | 1,081 | 1,118 | 1,203 | 1,388 | 1,114 | 1,231 | 1,18 | 0,92 | -0,08 | 1,6E-01 |
| Q8BJ71 | Nuclear pore complex protein Nup93 OS=Mus musculus OX=100 | 2 | 2 | 2 | 2 | 819 | 93,2 | 5,72 | 1430 | | | 2 | 0,953 | 0,834 | 0,988 | 1,097 | 0,61 | 0,872 | 0,961 | 0,836 | 1,18 | 0,18 | 1,6E-01 |
| Q8VC12 | Urocanate hydratase OS=Mus musculus OX=10090 GN=Uroc1 PI | 4 | 2 | 2 | 2 | 676 | 74,5 | 7,53 | 541 | 4,44 | 1 | 1 | 1,309 | 1,308 | 1,353 | 1,554 | 2,31 | 1,428 | 1,43 | 1,743 | 0,80 | -0,20 | 1,6E-01 |
| Q80XI4 | Phosphatidylinositol 5-phosphate 4-kinase type-2 beta OS=Mus | 24 | 10 | 19 | 10 | 416 | 47,3 | 7,33 | 10542 | 0 | 10 | 1 | 1,049 | 1,087 | 0,934 | 1,108 | 0,785 | 0,959 | 0,983 | 1,028 | 1,11 | 0,11 | 1,6E-01 |
| O35943 | Frataxin, mitochondrial OS=Mus musculus OX=10090 GN=Fxn PI | 3 | 1 | 1 | 1 | 207 | 22,9 | 8,02 | 495 | | | 1 | 1,144 | 1,045 | 0,734 | 1,427 | 0,223 | 0,815 | 1,124 | 0,675 | 1,53 | 0,53 | 1,6E-01 |
| Q9ERR1 | Nuclear distribution protein nudE-like 1 OS=Mus musculus OX=1 | 13 | 4 | 5 | 4 | 345 | 38,3 | 5,24 | 2326 | 2,31 | 3 | 1 | 0,968 | 1,08 | 0,873 | 1,084 | 1,007 | 0,821 | 0,905 | 0,857 | 1,12 | 0,12 | 1,6E-01 |
| P62071 | Ras-related protein R-Ras2 OS=Mus musculus OX=10090 GN=Rr | 36 | 7 | 15 | 6 | 204 | 23,4 | 6,01 | 3864 | 26,23 | 5 | 4 | 0,978 | 0,951 | 1,022 | 0,922 | 0,642 | 0,933 | 0,994 | 0,778 | 1,16 | 0,16 | 1,6E-01 |
| P31230 | Aminoacyl tRNA synthase complex-interacting multifunctional p | 22 | 5 | 5 | 5 | 310 | 34 | 8,35 | 1316 | 2,48 | 3 | 2 | 0,979 | 0,817 | 0,984 | 0,847 | 0,784 | 0,891 | 0,827 | 0,81 | 1,10 | 0,10 | 1,6E-01 |
| Q9DB73 | NADH-cytochrome b5 reductase 1 OS=Mus musculus OX=10090 | 6 | 2 | 2 | 2 | 305 | 34,1 | 8,87 | 361 | 1,66 | 1 | 1 | 1,077 | 1,085 | 0,974 | 1,269 | 0,966 | 0,968 | 0,915 | 1,094 | 1,12 | 0,12 | 1,6E-01 |
| P18572 | Basigin OS=Mus musculus OX=10090 GN=Bsg PE=1 SV=2 | 19 | 7 | 14 | 6 | 389 | 42,4 | 5,85 | 7441 | 0 | 6 | 1 | 1,008 | 0,951 | 0,956 | 0,945 | 1,014 | 1,005 | 0,98 | 0,973 | 0,97 | -0,03 | 1,6E-01 |
| P08249 | Malate dehydrogenase, mitochondrial OS=Mus musculus OX=10 | 55 | 20 | 75 | 20 | 338 | 35,6 | 8,68 | 36254 | 43,84 | 15 | 9 | 1,042 | 0,998 | 1,012 | 1,029 | 0,839 | 1,055 | 0,916 | 0,973 | 1,08 | 0,08 | 1,6E-01 |
| Q9DB05 | Alpha-soluble NSF attachment protein OS=Mus musculus OX=10 | 39 | 10 | 16 | 8 | 295 | 33,2 | 5,45 | 7265 | 16,57 | 7 | 5 | 1,017 | 1,041 | 1,016 | 1,001 | 1,025 | 1,091 | 1,012 | 1,108 | 0,96 | -0,04 | 1,6E-01 |
| P41105 | 60S ribosomal protein L28 OS=Mus musculus OX=10090 GN=Rpl | 39 | 7 | 9 | 7 | 137 | 15,7 | 12,02 | 2100 | 11,63 | 2 | 5 | 1,181 | 0,953 | 0,987 | 1,15 | 0,912 | 1,001 | 0,975 | 0,995 | 1,10 | 0,10 | 1,6E-01 |
| Q9WV27 | Sodium/potassium-transporting ATPase subunit alpha-4 OS=Mu | 9 | 10 | 64 | 1 | 1032 | 114,8 | 5,71 | 29660 | 29,21 | 6 | 5 | 1,028 | 0,975 | 0,742 | 1,055 | 1,308 | 0,919 | 1,053 | 1,672 | 0,77 | -0,23 | 1,6E-01 |
| Q62073 | Mitogen-activated protein kinase kinase kinase 7 OS=Mus musc | 4 | 2 | 2 | 2 | 579 | 64,2 | 6,67 | 224 | 3,07 | 1 | 1 | 0,949 | 0,786 | 0,965 | 0,963 | 1,201 | 0,867 | 1,06 | 1,053 | 0,88 | -0,12 | 1,6E-01 |
| P14148 | 60S ribosomal protein L7 OS=Mus musculus OX=10090 GN=Rpl7 | 36 | 16 | 24 | 16 | 270 | 31,4 | 10,89 | 5670 | 46,98 | 6 | 10 | 1,022 | 0,924 | 1,032 | 1,043 | 0,908 | 0,952 | 0,954 | 0,997 | 1,06 | 0,06 | 1,6E-01 |
| Q3UV17 | Keratin, type II cytoskeletal 2 oral OS=Mus musculus OX=10090 | 9 | 7 | 15 | 2 | 594 | 62,8 | 8,43 | 4947 | 11,17 | 3 | 5 | 0,885 | 1,08 | 0,677 | 0,785 | 2,17 | 1,316 | 1,052 | 0,832 | 0,64 | -0,36 | 1,6E-01 |
| Q9DBT5 | AMP deaminase 2 OS=Mus musculus OX=10090 GN=Ampd2 PE= | 5 | 4 | 5 | 4 | 798 | 92 | 6,23 | 971 | 8,57 | 2 | 2 | 0,979 | 0,935 | 0,989 | 0,895 | 0,994 | 0,977 | 1,007 | 0,968 | 0,96 | -0,04 | 1,6E-01 |
| Q5PR69 | Cancer-related regulator of actin dynamics OS=Mus musculus O | 5 | 4 | 4 | 4 | 1207 | 132,2 | 5,4 | 1708 | 2,26 | 3 | 1 | 1,081 | 0,98 | 1,026 | 1,035 | 1,586 | 1,033 | 1,222 | 1,084 | 0,84 | -0,16 | 1,6E-01 |
| Q9Z1D1 | Eukaryotic translation initiation factor 3 subunit G OS=Mus mus | 12 | 3 | 3 | 3 | 320 | 35,6 | 5,9 | 561 | 5,83 | 1 | 2 | 1,271 | 1,251 | 0,868 | 1,284 | 1,045 | 1,023 | 0,911 | 1,029 | 1,17 | 0,17 | 1,6E-01 |
| Q91ZJ5 | UTP--glucose-1-phosphate uridylyltransferase OS=Mus musculu | 22 | 13 | 21 | 13 | 508 | 56,9 | 7,61 | 9356 | 14,52 | 10 | 4 | 1,086 | 1,128 | 1,124 | 1,237 | 1,097 | 1,086 | 1,113 | 1,062 | 1,05 | 0,05 | 1,6E-01 |
| Q8R5C5 | Beta-centractin OS=Mus musculus OX=10090 GN=Actr1b PE=1 S | 29 | 8 | 9 | 3 | 376 | 42,3 | 6,4 | 4935 | 2,85 | 7 | 1 | 0,878 | 0,853 | 0,977 | 0,839 | 1,499 | 0,914 | 1,01 | 0,992 | 0,80 | -0,20 | 1,6E-01 |
| O89023 | Tripeptidyl-peptidase 1 OS=Mus musculus OX=10090 GN=Tpp1 I | 2 | 1 | 3 | 1 | 562 | 61,3 | 6,57 | 1941 | | | 1 | 1,434 | 1,537 | 1,364 | 1,435 | 1,905 | 1,381 | 1,484 | 1,889 | 0,87 | -0,13 | 1,6E-01 |
| Q9CQV8 | 14-3-3 protein beta/alpha OS=Mus musculus OX=10090 GN=Yw | 56 | 16 | 86 | 6 | 246 | 28,1 | 4,83 | 33374 | 73,53 | 11 | 11 | 0,893 | 1,022 | 0,907 | 1,056 | 0,863 | 0,951 | 0,901 | 0,877 | 1,08 | 0,08 | 1,6E-01 |

| | | | | | | | | | | | | | | | | | | | | | | | |
|---|---|---|---|---|---|---|---|---|---|---|---|---|---|---|---|---|---|---|---|---|---|---|---|
| Q8VE98 | CD276 antigen OS=Mus musculus OX=10090 GN=Cd276 PE=1 SV | 6 | 2 | 2 | 2 | 316 | 34 | 4,82 | 1126 | | 2 | | 1,262 | 1,259 | 1,321 | 1,05 | 1,206 | 1,166 | 0,99 | 1,031 | 1,11 | 0,11 | 1,7E-01 |
| Q99KE1 | NAD-dependent malic enzyme, mitochondrial OS=Mus musculu | 6 | 3 | 5 | 2 | 589 | 65,8 | 7,61 | 2991 | | 3 | | 0,896 | 0,894 | 0,703 | 0,736 | 1,397 | 0,72 | 0,919 | 1,186 | 0,76 | -0,24 | 1,7E-01 |
| Q9WVJ2 | 26S proteasome non-ATPase regulatory subunit 13 OS=Mus mu | 18 | 7 | 11 | 7 | 376 | 42,8 | 5,71 | 5007 | 4,79 | 4 | 3 | 1,04 | 1,034 | 1,046 | 1,08 | 0,778 | 1,067 | 0,963 | 0,997 | 1,10 | 0,10 | 1,7E-01 |
| Q9EPE9 | Manganese-transporting ATPase 13A1 OS=Mus musculus OX=10 | 4 | 3 | 3 | 3 | 1200 | 132,3 | 8,03 | 967 | 0 | 2 | 1 | 1,051 | 1,276 | 1,193 | 1,299 | 1,24 | 1,554 | 1,217 | 1,959 | 0,81 | -0,19 | 1,7E-01 |
| O88746 | Target of Myb protein 1 OS=Mus musculus OX=10090 GN=Tom1 | 5 | 2 | 3 | 2 | 492 | 54,3 | 4,94 | 797 | 2,47 | 1 | 1 | 0,78 | 0,805 | 0,726 | 0,808 | 0,996 | 0,742 | 1,072 | 0,811 | 0,86 | -0,14 | 1,7E-01 |
| Q91VR8 | Protein BRICK1 OS=Mus musculus OX=10090 GN=Brk1 PE=1 SV= | 20 | 2 | 4 | 2 | 75 | 8,8 | 5,45 | 1980 | 1,83 | 1 | 1 | 0,964 | 0,949 | 0,924 | 0,801 | 0,631 | 0,869 | 0,919 | 0,75 | 1,15 | 0,15 | 1,7E-01 |
| P24369 | Peptidyl-prolyl cis-trans isomerase B OS=Mus musculus OX=100 | 42 | 10 | 20 | 10 | 216 | 23,7 | 9,55 | 5968 | 33,85 | 5 | 5 | 1,09 | 1,163 | 0,982 | 1,453 | 0,869 | 1,005 | 1,097 | 1,015 | 1,18 | 0,18 | 1,7E-01 |
| Q9Z2M7 | Phosphomannomutase 2 OS=Mus musculus OX=10090 GN=Pmr | 13 | 3 | 4 | 2 | 242 | 27,6 | 6,42 | 1370 | 0 | 2 | 1 | 0,969 | 1,079 | 1,176 | 1,159 | 1,361 | 1,088 | 1,174 | 1,228 | 0,90 | -0,10 | 1,7E-01 |
| Q9QXS6 | Drebrin OS=Mus musculus OX=10090 GN=Dbn1 PE=1 SV=4 | 38 | 24 | 44 | 24 | 706 | 77,2 | 4,49 | 13265 | 61,36 | 14 | 13 | 1,045 | 0,982 | 1,008 | 0,965 | 0,874 | 1,001 | 0,919 | 0,988 | 1,06 | 0,06 | 1,7E-01 |
| Q9WUC3 | Lymphocyte antigen 6H OS=Mus musculus OX=10090 GN=Ly6h | 22 | 3 | 8 | 3 | 139 | 14,7 | 7,53 | 834 | 17,12 | 1 | 2 | 1,16 | 1,259 | 0,831 | 1,112 | 0,572 | 1,127 | 0,944 | 0,774 | 1,28 | 0,28 | 1,7E-01 |
| Q6ZQ06 | Centrosomal protein of 162 kDa OS=Mus musculus OX=10090 G | 1 | 2 | 8 | 2 | 1403 | 160,8 | 5,53 | 482 | 18,84 | 1 | 1 | 0,902 | 0,962 | 0,86 | 0,868 | 1,35 | 0,781 | 1,119 | 1,092 | 0,83 | -0,17 | 1,7E-01 |
| Q9JKF7 | 39S ribosomal protein L39, mitochondrial OS=Mus musculus OX | 3 | 1 | 1 | 1 | 336 | 38,5 | 7,94 | 553 | | 1 | | 0,881 | 0,779 | 1,007 | 0,921 | 0,664 | 0,81 | 0,921 | 0,738 | 1,15 | 0,15 | 1,7E-01 |
| Q9CR59 | Growth arrest and DNA damage-inducible proteins-interacting p | 4 | 1 | 1 | 1 | 222 | 25,8 | 10,27 | | 2,73 | | 1 | 0,927 | 0,944 | 0,825 | 0,813 | 1,016 | 1,137 | 0,87 | 0,915 | 0,89 | -0,11 | 1,7E-01 |
| O88447 | Kinesin light chain 1 OS=Mus musculus OX=10090 GN=Klc1 PE=1 | 26 | 12 | 17 | 9 | 541 | 61,4 | 5,68 | 6137 | 13,96 | 7 | 5 | 0,978 | 1,077 | 1,079 | 1,002 | 0,968 | 0,932 | 1,011 | 1,019 | 1,05 | 0,05 | 1,7E-01 |
| P80313 | T-complex protein 1 subunit eta OS=Mus musculus OX=10090 G | 41 | 22 | 34 | 22 | 544 | 59,6 | 7,84 | 18308 | 18,96 | 18 | 4 | 1,057 | 0,975 | 1,116 | 0,986 | 0,868 | 1,027 | 0,978 | 0,967 | 1,08 | 0,08 | 1,7E-01 |
| Q8BJZ4 | 28S ribosomal protein S35, mitochondrial OS=Mus musculus OX | 4 | 1 | 1 | 1 | 320 | 36 | 8,59 | | 2,42 | | 1 | 0,933 | 0,793 | 0,944 | 0,919 | 1,484 | 0,921 | 1,05 | 0,97 | 0,81 | -0,19 | 1,7E-01 |
| Q8CI32 | BAG family molecular chaperone regulator 5 OS=Mus musculus | 2 | 1 | 1 | 1 | 447 | 50,9 | 6,05 | | 1,72 | | 1 | 0,814 | 1,074 | 1,088 | 0,885 | 1,047 | 1,1 | 1,009 | 1,359 | 0,86 | -0,14 | 1,7E-01 |
| Q9CZU6 | Citrate synthase, mitochondrial OS=Mus musculus OX=10090 GI | 28 | 13 | 35 | 13 | 464 | 51,7 | 8,57 | 14728 | 20,82 | 10 | 4 | 1,007 | 0,988 | 0,975 | 0,955 | 0,925 | 0,989 | 0,951 | 0,953 | 1,03 | 0,03 | 1,7E-01 |
| P0DP27 | Calmodulin-2 OS=Mus musculus OX=10090 GN=Calm2 PE=1 SV= | 52 | 11 | 111 | 11 | 149 | 16,8 | 4,22 | 46810 | 88,86 | 9 | 5 | 0,991 | 1,132 | 0,896 | 1,146 | 0,744 | 1,052 | 0,927 | 0,898 | 1,15 | 0,15 | 1,7E-01 |
| P58064 | 28S ribosomal protein S6, mitochondrial OS=Mus musculus OX= | 33 | 4 | 5 | 4 | 125 | 14,3 | 9,5 | 689 | 9,35 | 1 | 3 | 0,999 | 0,867 | 0,866 | 0,895 | 0,889 | 0,849 | 0,865 | 0,769 | 1,08 | 0,08 | 1,7E-01 |
| Q921W4 | Quinone oxidoreductase-like protein 1 OS=Mus musculus OX=10 | 8 | 3 | 3 | 3 | 348 | 38,7 | 5,97 | 1176 | 2,36 | 2 | 1 | 1,14 | 1,093 | 1,024 | 1,219 | 0,796 | 1,036 | 1,036 | 1,11 | 1,13 | 0,13 | 1,7E-01 |
| P62196 | 26S proteasome regulatory subunit 8 OS=Mus musculus OX=100 | 21 | 9 | 12 | 9 | 406 | 45,6 | 7,55 | 3847 | 9,84 | 5 | 4 | 1,108 | 1,029 | 0,973 | 1,055 | 0,876 | 0,964 | 1,028 | 1,018 | 1,07 | 0,07 | 1,7E-01 |
| P17156 | Heat shock-related 70 kDa protein 2 OS=Mus musculus OX=100 | 26 | 18 | 41 | 4 | 633 | 69,6 | 5,67 | 16368 | 40,46 | 10 | 10 | 0,91 | 1,067 | 1,042 | 0,934 | 1,23 | 0,907 | 1,241 | 1,116 | 0,88 | -0,12 | 1,7E-01 |
| Q68EF0 | Rab-3A-interacting protein OS=Mus musculus OX=10090 GN=Ra | 2 | 1 | 1 | 1 | 428 | 47,1 | 6,71 | 391 | | 1 | | 0,913 | 1 | 0,982 | 0,968 | 0,796 | 0,867 | 0,987 | 0,931 | 1,08 | 0,08 | 1,7E-01 |
| P41216 | Long-chain-fatty-acid--CoA ligase 1 OS=Mus musculus OX=1009 | 5 | 3 | 3 | 3 | 699 | 77,9 | 7,15 | 730 | 2,53 | 2 | 1 | 1,05 | 1,022 | 1,097 | 1,077 | 1,137 | 1,046 | 1,09 | 1,169 | 0,96 | -0,04 | 1,7E-01 |
| P01864 | Ig gamma-2A chain C region secreted form OS=Mus musculus O | 7 | 2 | 2 | 2 | 335 | 36,6 | 8,22 | 1508 | | 2 | | 0,913 | 0,817 | 0,797 | 0,718 | 0,767 | 0,969 | 0,958 | 1,347 | 0,80 | -0,20 | 1,7E-01 |
| Q8CHT1 | Ephexin-1 OS=Mus musculus OX=10090 GN=Ngef PE=1 SV=1 | 7 | 4 | 4 | 4 | 710 | 82,1 | 5,99 | 1606 | 0 | 3 | 1 | 1,009 | 0,996 | 1,211 | 1,012 | 1,538 | 1,405 | 0,971 | 1,162 | 0,83 | -0,17 | 1,7E-01 |
| P67871 | Casein kinase II subunit beta OS=Mus musculus OX=10090 GN=C | 14 | 3 | 10 | 3 | 215 | 24,9 | 5,55 | 3434 | 8,25 | 2 | 2 | 0,959 | 0,934 | 0,99 | 0,938 | 1,594 | 1,279 | 0,906 | 1,005 | 0,80 | -0,20 | 1,7E-01 |
| Q8VD04 | GRIP1-associated protein 1 OS=Mus musculus OX=10090 GN=Gr | 6 | 4 | 4 | 4 | 806 | 92,7 | 5,25 | 1827 | | 4 | | 0,896 | 0,825 | 1,242 | 0,988 | 0,9 | 0,777 | 0,854 | 0,832 | 1,17 | 0,17 | 1,7E-01 |
| P31650 | Sodium- and chloride-dependent GABA transporter 3 OS=Mus m | 6 | 5 | 5 | 5 | 627 | 69,9 | 6,98 | 2207 | 6,37 | 3 | 2 | 1,019 | 1,02 | 0,935 | 0,897 | 0,529 | 1,002 | 0,852 | 0,842 | 1,20 | 0,20 | 1,7E-01 |
| P50446 | Keratin, type II cytoskeletal 6A OS=Mus musculus OX=10090 GN | 21 | 14 | 17 | 4 | 553 | 59,3 | 7,94 | 4359 | 15,2 | 7 | 7 | 0,863 | 0,955 | 0,819 | 1,109 | 1,577 | 1,173 | 0,917 | 1,049 | 0,79 | -0,21 | 1,7E-01 |
| P50172 | Corticosteroid 11-beta-dehydrogenase isozyme 1 OS=Mus musc | 8 | 2 | 4 | 2 | 292 | 32,3 | 8,46 | 1347 | 2,49 | 1 | 1 | 1,038 | 1,052 | 1,103 | 1,231 | 1,18 | 1,229 | 1,185 | 1,13 | 0,94 | -0,06 | 1,7E-01 |
| P22723 | Gamma-aminobutyric acid receptor subunit gamma-2 OS=Mus r | 3 | 1 | 1 | 1 | 474 | 55,1 | 8,66 | | 0 | | 1 | 1,321 | 1,548 | 1,575 | 1,336 | 1,494 | 1,148 | 1,077 | 1,341 | 1,14 | 0,14 | 1,7E-01 |
| Q01853 | Transitional endoplasmic reticulum ATPase OS=Mus musculus O | 42 | 31 | 65 | 31 | 806 | 89,3 | 5,26 | 26081 | 62,8 | 22 | 11 | 1,005 | 0,972 | 1,004 | 0,994 | 0,781 | 1,005 | 0,969 | 0,912 | 1,08 | 0,08 | 1,7E-01 |
| Q3UM45 | Protein phosphatase 1 regulatory subunit 7 OS=Mus musculus | 18 | 6 | 8 | 6 | 361 | 41,3 | 4,92 | 2734 | 8,32 | 3 | 3 | 0,956 | 0,983 | 0,95 | 0,98 | 0,999 | 0,985 | 1,082 | 0,968 | 0,96 | -0,04 | 1,7E-01 |
| Q3UX10 | Tubulin alpha chain-like 3 OS=Mus musculus OX=10090 GN=Tub | 10 | 5 | 33 | 1 | 446 | 50 | 5,58 | 15136 | 14,09 | 3 | 2 | 0,935 | 0,748 | 0,84 | 0,905 | 1,196 | 1,092 | 0,9 | 0,826 | 0,85 | -0,15 | 1,7E-01 |
| Q9CQI3 | Glia maturation factor beta OS=Mus musculus OX=10090 GN=Gi | 18 | 3 | 5 | 3 | 142 | 16,7 | 5,16 | 2835 | | 3 | | 0,883 | 1,074 | 1,171 | 1,11 | 0,877 | 0,97 | 0,995 | 0,978 | 1,11 | 0,11 | 1,7E-01 |
| Q61696 | Heat shock 70 kDa protein 1A OS=Mus musculus OX=10090 GN= | 13 | 8 | 21 | 2 | 641 | 70 | 5,72 | 8626 | 21,29 | 6 | 3 | 0,835 | 0,805 | 0,969 | 0,914 | 1,066 | 0,976 | 0,879 | 0,937 | 0,91 | -0,09 | 1,7E-01 |
| Q8BYJ6 | TBC1 domain family member 4 OS=Mus musculus OX=10090 GN | 1 | 1 | 1 | 1 | 1307 | 147,4 | 7,33 | | 2,75 | | 1 | 0,832 | 1,138 | 0,886 | 1,009 | 1,621 | 1,032 | 0,933 | 1,328 | 0,79 | -0,21 | 1,7E-01 |
| Q68ED2 | Metabotropic glutamate receptor 7 OS=Mus musculus OX=1009 | 11 | 8 | 11 | 7 | 915 | 102,2 | 7,88 | 2569 | 18,31 | 4 | 5 | 1,062 | 0,993 | 0,93 | 1,025 | 0,916 | 0,976 | 0,951 | 0,974 | 1,05 | 0,05 | 1,7E-01 |
| Q62465 | Synaptic vesicle membrane protein VAT-1 homolog OS=Mus mu | 39 | 13 | 34 | 13 | 406 | 43,1 | 6,37 | 14661 | 25,07 | 7 | 8 | 1,019 | 1,01 | 0,976 | 1,019 | 0,896 | 1,007 | 0,989 | 0,969 | 1,04 | 0,04 | 1,7E-01 |
| P97379 | Ras GTPase-activating protein-binding protein 2 OS=Mus muscu | 4 | 2 | 2 | 2 | 482 | 54,1 | 5,62 | 1480 | | 2 | | 1,055 | 0,782 | 1,004 | 0,97 | 0,801 | 0,883 | 0,792 | 0,92 | 1,12 | 0,12 | 1,7E-01 |
| P99027 | 60S acidic ribosomal protein P2 OS=Mus musculus OX=10090 GI | 61 | 5 | 14 | 5 | 115 | 11,6 | 4,54 | 4848 | 14,24 | 3 | 2 | 1,049 | 1,12 | 1,044 | 1,358 | 1,011 | 1,05 | 1,031 | 1,021 | 1,11 | 0,11 | 1,7E-01 |
| P35564 | Calnexin OS=Mus musculus OX=10090 GN=Canx PE=1 SV=1 | 22 | 14 | 23 | 14 | 591 | 67,2 | 4,64 | 6302 | 21,22 | 7 | 7 | 1,05 | 1,054 | 1,071 | 1,085 | 0,736 | 1,011 | 1,033 | 1,031 | 1,12 | 0,12 | 1,7E-01 |
| Q9D826 | Peroxisomal sarcosine oxidase OS=Mus musculus OX=10090 GN | 4 | 1 | 1 | 1 | 390 | 43,8 | 7,55 | 717 | | 1 | | 1,613 | 1,786 | 1,326 | 1,661 | 2,832 | 1,728 | 1,694 | 1,891 | 0,78 | -0,22 | 1,7E-01 |
| P46061 | Ran GTPase-activating protein 1 OS=Mus musculus OX=10090 G | 6 | 3 | 4 | 2 | 589 | 63,5 | 4,68 | 1782 | | 3 | | 1,066 | 1,061 | 1,142 | 1,218 | 1,142 | 1,212 | 1,169 | 1,217 | 0,95 | -0,05 | 1,8E-01 |
| Q61036 | Serine/threonine-protein kinase PAK 3 OS=Mus musculus OX=10 | 18 | 10 | 15 | 4 | 559 | 62,4 | 5,4 | 8174 | 3,84 | 9 | 2 | 0,977 | 1,023 | 0,873 | 1,088 | 1,014 | 1,057 | 1,078 | 1,326 | 0,89 | -0,11 | 1,8E-01 |
| Q9EQI8 | 39S ribosomal protein L46, mitochondrial OS=Mus musculus OX | 11 | 3 | 3 | 3 | 283 | 32,1 | 7,4 | 2283 | | 3 | | 0,914 | 0,886 | 0,863 | 0,835 | 0,966 | 1,041 | 0,911 | 0,85 | 0,93 | -0,07 | 1,8E-01 |
| Q60854 | Serpin B6 OS=Mus musculus OX=10090 GN=Serpinb6 PE=1 SV=1 | 9 | 3 | 3 | 3 | 378 | 42,6 | 5,74 | 1929 | | 3 | | 0,98 | 0,996 | 0,976 | 0,962 | 0,939 | 1,121 | 1,015 | 1,097 | 0,94 | -0,06 | 1,8E-01 |
| P62830 | 60S ribosomal protein L23 OS=Mus musculus OX=10090 GN=Rpl | 34 | 4 | 9 | 4 | 140 | 14,9 | 10,51 | 3522 | 7,36 | 3 | 3 | 1,099 | 1,033 | 1,026 | 1,077 | 1,39 | 1,052 | 1,202 | 1,077 | 0,90 | -0,10 | 1,8E-01 |
| Q99MK8 | Beta-adrenergic receptor kinase 1 OS=Mus musculus OX=10090 | 11 | 7 | 9 | 7 | 689 | 79,6 | 7,28 | 3284 | 9,49 | 4 | 4 | 0,997 | 0,894 | 1,008 | 0,918 | 0,937 | 1,16 | 0,99 | 1,078 | 0,92 | -0,08 | 1,8E-01 |
| O70166 | Stathmin-3 OS=Mus musculus OX=10090 GN=Stmn3 PE=1 SV=1 | 37 | 8 | 12 | 8 | 180 | 20,9 | 7,49 | 836 | 27,16 | 1 | 7 | 0,98 | 0,992 | 1,008 | 0,954 | 0,875 | 0,917 | 1,021 | 0,918 | 1,05 | 0,05 | 1,8E-01 |
| Q8R4E6 | Purine-rich element-binding protein gamma OS=Mus musculus | 3 | 1 | 1 | 1 | 350 | 39,9 | 9,51 | 513 | | 1 | | 1,033 | 0,822 | 0,898 | 0,658 | 0,957 | 0,899 | 1,031 | 1,048 | 0,87 | -0,13 | 1,8E-01 |
| Q61818 | Retinoic acid-induced protein 1 OS=Mus musculus OX=10090 GI | 1 | 1 | 1 | 1 | 1889 | 201,4 | 8,75 | 208 | | 1 | | 0,552 | 1,034 | 1,031 | 0,794 | 2,022 | 0,908 | 1,142 | 1,04 | 0,67 | -0,33 | 1,8E-01 |
| P56399 | Ubiquitin carboxyl-terminal hydrolase 5 OS=Mus musculus OX=1 | 24 | 15 | 23 | 15 | 858 | 95,8 | 5,01 | 8007 | 25,32 | 9 | 7 | 0,976 | 1 | 1,02 | 0,974 | 0,741 | 1,022 | 0,929 | 0,913 | 1,10 | 0,10 | 1,8E-01 |
| P13707 | Glycerol-3-phosphate dehydrogenase [NAD(+)], cytoplasmic OS= | 5 | 2 | 6 | 2 | 349 | 37,5 | 7,17 | | 27,6 | | 2 | 2,585 | 3,258 | 2,285 | 3,449 | 4,339 | 2,664 | 3,552 | 3,73 | 0,81 | -0,19 | 1,8E-01 |
| Q9JIN6 | Calcium-activated potassium channel subunit beta-4 OS=Mus m | 7 | 1 | 1 | 1 | 210 | 23,8 | 7,25 | 126 | | 1 | | 1,029 | 0,812 | 0,881 | 0,936 | 0,971 | 0,991 | 0,971 | 1,012 | 0,93 | -0,07 | 1,8E-01 |
| P30276 | G2/mitotic-specific cyclin-B2 OS=Mus musculus OX=10090 GN=C | 3 | 1 | 1 | 1 | 398 | 45,4 | 8,92 | | 0 | | 1 | 0,993 | 0,956 | 1,022 | 0,896 | 0,744 | 0,95 | 0,993 | 0,639 | 1,16 | 0,16 | 1,8E-01 |
| Q01768 | Nucleoside diphosphate kinase B OS=Mus musculus OX=10090 C | 62 | 10 | 30 | 5 | 152 | 17,4 | 7,5 | 6745 | 54,12 | 6 | 6 | 0,978 | 0,972 | 1,097 | 0,972 | 0,463 | 1,031 | 0,916 | 0,834 | 1,24 | 0,24 | 1,8E-01 |
| Q61838 | Pregnancy zone protein OS=Mus musculus OX=10090 GN=Pzp P | 4 | 5 | 11 | 5 | 1495 | 165,7 | 6,68 | 6692 | 2,48 | 4 | 1 | 1,745 | 1,595 | 2,073 | 1,507 | 3,812 | 1,75 | 1,936 | 2,376 | 0,70 | -0,30 | 1,8E-01 |
| Q9JHU4 | Cytoplasmic dynein 1 heavy chain 1 OS=Mus musculus OX=1009 | 29 | 144 | 266 | 144 | 4644 | 531,7 | 6,42 | 110204 | 221,73 | 95 | 54 | 1,018 | 0,957 | 1,017 | 0,957 | 0,863 | 1,003 | 0,941 | 0,937 | 1,05 | 0,05 | 1,8E-01 |
| P83940 | Elongin-C OS=Mus musculus OX=10090 GN=Eloc PE=1 SV=1 | 59 | 6 | 7 | 6 | 112 | 12,5 | 4,78 | 4180 | 3,18 | 5 | 1 | 0,973 | 1,02 | 0,974 | 0,912 | 0,683 | 0,973 | 0,939 | 0,866 | 1,12 | 0,12 | 1,8E-01 |
| P00493 | Hypoxanthine-guanine phosphoribosyltransferase OS=Mus mus | 26 | 4 | 7 | 4 | 218 | 24,6 | 6,68 | 3558 | 2,13 | 4 | 1 | 1,015 | 1,134 | 0,976 | 1,067 | 1,297 | 1,073 | 1,053 | 1,168 | 0,91 | -0,09 | 1,8E-01 |
| P35803 | Neuronal membrane glycoprotein M6-b OS=Mus musculus OX=1 | 13 | 4 | 10 | 4 | 328 | 36,2 | 6,14 | 5817 | | 4 | | 0,957 | 0,973 | 0,962 | 0,934 | 0,614 | 0,847 | 0,991 | 0,886 | 1,15 | 0,15 | 1,8E-01 |
| Q3UGY8 | Brefeldin A-inhibited guanine nucleotide-exchange protein 3 OS | 1 | 2 | 2 | 2 | 2170 | 239,9 | 5,88 | 1221 | | 2 | | 0,855 | 1,002 | 1,19 | 0,828 | 0,888 | 0,846 | 0,852 | 0,753 | 1,16 | 0,16 | 1,8E-01 |

| | | | | | | | | | | | | | | | | | | | | | | | |
|---|---|---|---|---|---|---|---|---|---|---|---|---|---|---|---|---|---|---|---|---|---|---|---|
| Q99PT1 | Rho GDP-dissociation inhibitor 1 OS=Mus musculus OX=10090 G | 40 | 8 | 30 | 8 | 204 | 23,4 | 5,2 | 13690 | 18,13 | 5 | 4 | 0,958 | 1,037 | 1,071 | 1,073 | 0,585 | 1 | 1,004 | 0,923 | 1,18 | 0,18 | 1,8E-01 |
| Q6ZWV3 | 60S ribosomal protein L10 OS=Mus musculus OX=10090 GN=Rpl | 28 | 7 | 10 | 7 | 214 | 24,6 | 10,08 | 3819 | 6,74 | 4 | 3 | 0,89 | 0,749 | 0,88 | 0,75 | 1,084 | 0,896 | 0,861 | 0,841 | 0,89 | -0,11 | 1,8E-01 |
| O70443 | Guanine nucleotide-binding protein G(z) subunit alpha OS=Mus | 37 | 11 | 16 | 9 | 355 | 40,8 | 7,61 | 6157 | 10,48 | 9 | 2 | 1,021 | 1,027 | 1,024 | 0,993 | 1,02 | 1,028 | 1,038 | 1,102 | 0,97 | -0,03 | 1,8E-01 |
| Q9CQF8 | Ribosomal protein 63, mitochondrial OS=Mus musculus OX=100 | 22 | 3 | 5 | 3 | 102 | 11,9 | 10,15 | 2406 | | 3 | | 0,959 | 0,936 | 1,055 | 0,95 | 1,147 | 1,059 | 0,974 | 1,003 | 0,93 | -0,07 | 1,8E-01 |
| Q9QXK3 | Coatomer subunit gamma-2 OS=Mus musculus OX=10090 GN=C | 2 | 2 | 7 | 1 | 871 | 97,6 | 5,8 | 3108 | | 2 | | 0,967 | 0,839 | 0,993 | 0,805 | 0,595 | 0,92 | 0,738 | 0,842 | 1,16 | 0,16 | 1,8E-01 |
| Q569Z6 | Thyroid hormone receptor-associated protein 3 OS=Mus muscu | 1 | 1 | 2 | 1 | 951 | 108,1 | 10,17 | 979 | | 1 | | 0,859 | 1,162 | 0,883 | 0,968 | 1,232 | 0,889 | 1,16 | 1,238 | 0,86 | -0,14 | 1,8E-01 |
| Q9CWZ7 | Gamma-soluble NSF attachment protein OS=Mus musculus OX= | 30 | 10 | 17 | 10 | 312 | 34,7 | 5,41 | 10768 | 5,44 | 9 | 2 | 1,061 | 0,962 | 0,984 | 1,007 | 0,756 | 1,007 | 0,923 | 0,97 | 1,10 | 0,10 | 1,8E-01 |
| A2APV2 | Formin-like protein 2 OS=Mus musculus OX=10090 GN=Fmnl2 P | 1 | 1 | 1 | 1 | 1086 | 123 | 7,53 | 557 | | 1 | | 0,756 | 0,972 | 0,768 | 0,859 | 0,978 | 0,765 | 1,038 | 1,275 | 0,83 | -0,17 | 1,8E-01 |
| Q99JI4 | 26S proteasome non-ATPase regulatory subunit 6 OS=Mus musc | 17 | 7 | 11 | 7 | 389 | 45,5 | 5,52 | 4400 | 11,92 | 5 | 3 | 0,974 | 1,011 | 1,11 | 1,102 | 0,967 | 0,985 | 1,04 | 0,98 | 1,06 | 0,06 | 1,8E-01 |
| Q8VCL2 | Protein SCO2 homolog, mitochondrial OS=Mus musculus OX=10 | 3 | 1 | 1 | 1 | 255 | 28,9 | 8,29 | 697 | | 1 | | 1,188 | 0,895 | 1,034 | 1,107 | 1,677 | 1,27 | 1,164 | 1,032 | 0,82 | -0,18 | 1,8E-01 |
| Q8BHE3 | Caytaxin OS=Mus musculus OX=10090 GN=Atcay PE=1 SV=1 | 13 | 3 | 4 | 3 | 372 | 42,2 | 4,68 | 2321 | 2,71 | 2 | 1 | 0,882 | 0,838 | 0,993 | 0,853 | 1,126 | 0,816 | 0,993 | 1,112 | 0,88 | -0,12 | 1,8E-01 |
| P62500 | TSC22 domain family protein 1 OS=Mus musculus OX=10090 GN | 2 | 2 | 3 | 2 | 1077 | 109,7 | 5,83 | 1623 | 2,27 | 2 | 1 | 1,006 | 1,086 | 0,852 | 1,105 | 0,738 | 0,937 | 0,983 | 0,917 | 1,13 | 0,13 | 1,8E-01 |
| Q3UUQ7 | GPI inositol-deacylase OS=Mus musculus OX=10090 GN=Pgap1 I | 1 | 1 | 1 | 1 | 922 | 104,5 | 8,91 | 514 | | 1 | | 0,769 | 0,878 | 1,272 | 0,896 | 0,794 | 0,797 | 0,812 | 0,746 | 1,21 | 0,21 | 1,8E-01 |
| P13595-2 | Isoform 2 of Neural cell adhesion molecule 1 OS=Mus musculus | 47 | 42 | 200 | 3 | 848 | 93,5 | 4,91 | 72895 | 220,67 | 30 | 25 | 1,095 | 1,243 | 0,743 | 1,221 | 0,71 | 0,884 | 0,9 | 1,016 | 1,23 | 0,23 | 1,8E-01 |
| O08807 | Peroxiredoxin-4 OS=Mus musculus OX=10090 GN=Prdx4 PE=1 S | 16 | 4 | 9 | 3 | 274 | 31 | 7,15 | 5556 | 3,49 | 4 | 1 | 0,966 | 1,082 | 0,854 | 0,942 | 0,725 | 0,802 | 0,921 | 0,963 | 1,13 | 0,13 | 1,8E-01 |
| Q61206 | Platelet-activating factor acetylhydrolase IB subunit beta OS=Mu | 12 | 2 | 6 | 2 | 229 | 25,6 | 5,92 | 3699 | | 2 | | 0,882 | 0,807 | 1,095 | 0,922 | 0,398 | 0,979 | 0,823 | 0,678 | 1,29 | 0,29 | 1,8E-01 |
| Q60648 | Ganglioside GM2 activator OS=Mus musculus OX=10090 GN=Gn | 4 | 1 | 1 | 1 | 193 | 20,8 | 5,9 | 480 | | 1 | | 1,124 | 1,06 | 0,848 | 1,219 | 0,642 | 0,991 | 0,933 | 0,994 | 1,19 | 0,19 | 1,8E-01 |
| Q8BFP9 | [Pyruvate dehydrogenase (acetyl-transferring)] kinase isozyme 1 | 3 | 1 | 1 | 1 | 434 | 49 | 8,19 | | 2,09 | | 1 | 0,82 | 1,101 | 1,566 | 1,098 | 2,313 | 1,375 | 1,274 | 1,363 | 0,72 | -0,28 | 1,8E-01 |
| P42932 | T-complex protein 1 subunit theta OS=Mus musculus OX=10090 | 47 | 29 | 55 | 29 | 548 | 59,5 | 5,62 | 28339 | 28,72 | 22 | 7 | 0,992 | 0,977 | 0,959 | 0,966 | 0,827 | 0,933 | 0,969 | 0,963 | 1,05 | 0,05 | 1,8E-01 |
| Q80Z24 | Neuronal growth regulator 1 OS=Mus musculus OX=10090 GN=I | 11 | 3 | 5 | 3 | 348 | 37,9 | 6,52 | 2275 | 2,84 | 3 | 1 | 0,968 | 1,046 | 0,865 | 1,156 | 0,925 | 0,871 | 0,92 | 0,94 | 1,10 | 0,10 | 1,9E-01 |
| Q811U4 | Mitofusin-1 OS=Mus musculus OX=10090 GN=Mfn1 PE=1 SV=3 | 1 | 1 | 1 | 1 | 741 | 83,7 | 6,51 | | 0 | | 1 | 0,822 | 0,765 | 0,954 | 0,901 | 1,342 | 0,733 | 0,97 | 1,284 | 0,80 | -0,20 | 1,9E-01 |
| Q99L04 | Dehydrogenase/reductase SDR family member 1 OS=Mus muscu | 22 | 7 | 8 | 7 | 313 | 34 | 8,35 | 2272 | 9,63 | 4 | 3 | 0,99 | 0,976 | 1,041 | 1,006 | 1,058 | 1,077 | 0,971 | 1,084 | 0,96 | -0,04 | 1,9E-01 |
| O09133 | Submaxillary gland androgen-regulated protein 2, isoform alpha | 4 | 1 | 1 | 1 | 150 | 16,6 | 9,85 | 590 | | 1 | | 0,979 | 1,153 | 0,727 | 0,978 | 1,299 | 0,788 | 1,351 | 1,392 | 0,79 | -0,21 | 1,9E-01 |
| E9Q7G0 | Nuclear mitotic apparatus protein 1 OS=Mus musculus OX=1009 | 1 | 2 | 3 | 2 | 2094 | 235,5 | 5,87 | 1083 | | 2 | | 0,82 | 0,886 | 0,925 | 0,909 | 1,13 | 1,016 | 0,815 | 0,992 | 0,90 | -0,10 | 1,9E-01 |
| P11881 | Inositol 1,4,5-trisphosphate receptor type 1 OS=Mus musculus C | 0 | 1 | 1 | 1 | 2749 | 313 | 6,04 | 471 | | 1 | | 0,867 | 0,973 | 0,904 | 0,968 | 1,161 | 0,92 | 1,001 | 0,975 | 0,91 | -0,09 | 1,9E-01 |
| Q8VDM6 | Heterogeneous nuclear ribonucleoprotein U-like protein 1 OS=N | 3 | 2 | 4 | 2 | 859 | 95,9 | 6,58 | 1576 | | 2 | | 0,962 | 0,988 | 1,066 | 1,152 | 1,325 | 0,976 | 1,119 | 1,301 | 0,88 | -0,12 | 1,9E-01 |
| P97820 | Mitogen-activated protein kinase kinase kinase kinase 4 OS=Mu | 10 | 10 | 11 | 6 | 1233 | 140,5 | 7,47 | 4542 | 11,36 | 6 | 4 | 0,955 | 1,044 | 1,162 | 1,104 | 0,702 | 1,042 | 1,064 | 0,893 | 1,15 | 0,15 | 1,9E-01 |
| Q3UHC7 | Disabled homolog 2-interacting protein OS=Mus musculus OX=1 | 3 | 4 | 5 | 3 | 1189 | 131,6 | 8,72 | 878 | 7,45 | 2 | 2 | 1,026 | 1,105 | 0,909 | 1,097 | 1,14 | 1,058 | 1,068 | 1,236 | 0,92 | -0,08 | 1,9E-01 |
| P62862 | 40S ribosomal protein S30 OS=Mus musculus OX=10090 GN=Fau | 31 | 2 | 3 | 2 | 59 | 6,6 | 12,15 | 1194 | 2,85 | 1 | 1 | 1,17 | 0,976 | 0,802 | 0,985 | 0,731 | 0,984 | 0,862 | 0,807 | 1,16 | 0,16 | 1,9E-01 |
| Q9CQ40 | 39S ribosomal protein L49, mitochondrial OS=Mus musculus OX | 11 | 2 | 2 | 2 | 166 | 19,1 | 9,5 | 740 | 2,04 | 1 | 1 | 0,939 | 0,989 | 1,063 | 0,912 | 1,14 | 1,021 | 1,134 | 0,948 | 0,92 | -0,08 | 1,9E-01 |
| Q9Z1J3 | Cysteine desulfurase, mitochondrial OS=Mus musculus OX=1009 | 9 | 4 | 4 | 4 | 459 | 50,5 | 8,16 | 1414 | 4,5 | 2 | 2 | 0,991 | 0,9 | 0,948 | 0,977 | 0,934 | 0,881 | 0,805 | 0,96 | 1,07 | 0,07 | 1,9E-01 |
| P62889 | 60S ribosomal protein L30 OS=Mus musculus OX=10090 GN=Rpl | 34 | 3 | 6 | 3 | 115 | 12,8 | 9,63 | 775 | 8,65 | 1 | 2 | 1,053 | 0,961 | 1,038 | 1,097 | 0,791 | 0,942 | 1,014 | 1,034 | 1,10 | 0,10 | 1,9E-01 |
| P21661 | Neuroendocrine convertase 2 OS=Mus musculus OX=10090 GN= | 3 | 2 | 2 | 2 | 637 | 70,7 | 6,42 | 1212 | | 2 | | 0,974 | 1,046 | 1,12 | 1,19 | 1,02 | 0,96 | 1,037 | 1,019 | 1,07 | 0,07 | 1,9E-01 |
| P60867 | 40S ribosomal protein S20 OS=Mus musculus OX=10090 GN=Rp | 24 | 4 | 6 | 4 | 119 | 13,4 | 9,94 | 4026 | | 4 | | 0,938 | 0,962 | 1,021 | 1,1 | 0,856 | 0,997 | 0,98 | 0,895 | 1,08 | 0,08 | 1,9E-01 |
| Q61598 | Rab GDP dissociation inhibitor beta OS=Mus musculus OX=1009 | 46 | 20 | 36 | 15 | 445 | 50,5 | 6,25 | 9520 | 46,82 | 13 | 10 | 0,976 | 0,938 | 0,956 | 0,916 | 0,812 | 0,985 | 0,901 | 0,857 | 1,06 | 0,06 | 1,9E-01 |
| P60840 | Alpha-endosulfine OS=Mus musculus OX=10090 GN=Ensa PE=1 | 24 | 2 | 2 | 2 | 121 | 13,3 | 7,24 | 659 | 2,94 | 1 | 1 | 0,94 | 0,864 | 1,049 | 0,814 | 1,033 | 1,069 | 0,902 | 1,036 | 0,91 | -0,09 | 1,9E-01 |
| Q6PD10 | Inositol hexakisphosphate kinase 1 OS=Mus musculus OX=1009 | 3 | 1 | 2 | 1 | 433 | 49,3 | 7,23 | | 5,13 | | 1 | 0,882 | 1,123 | 1,239 | 0,999 | 1,384 | 0,968 | 1,254 | 1,365 | 0,85 | -0,15 | 1,9E-01 |
| Q02257 | Junction plakoglobin OS=Mus musculus OX=10090 GN=Jup PE=1 | 14 | 8 | 10 | 7 | 745 | 81,7 | 6,14 | 3346 | 8,3 | 5 | 3 | 0,963 | 1,181 | 0,967 | 0,977 | 1,318 | 1,08 | 1,148 | 1,029 | 0,89 | -0,11 | 1,9E-01 |
| Q6P6M7 | O-phosphoseryl-tRNA(Sec) selenium transferase OS=Mus muscu | 1 | 1 | 2 | 1 | 504 | 55,3 | 8,06 | | 4,57 | | 1 | 1,194 | 1,141 | 0,906 | 1,186 | 0,901 | 1,125 | 0,927 | 0,976 | 1,13 | 0,13 | 1,9E-01 |
| Q9CPV4 | Glyoxalase domain-containing protein 4 OS=Mus musculus OX= | 24 | 6 | 9 | 6 | 298 | 33,3 | 5,47 | 5151 | 3,02 | 5 | 1 | 0,988 | 0,891 | 0,995 | 1,028 | 1,203 | 1,147 | 0,868 | 1,173 | 0,89 | -0,11 | 1,9E-01 |
| Q8CGC7 | Bifunctional glutamate/proline--tRNA ligase OS=Mus musculus C | 7 | 9 | 11 | 9 | 1512 | 170 | 7,66 | 2978 | 10,65 | 4 | 5 | 0,97 | 0,949 | 0,994 | 0,978 | 0,954 | 1,233 | 0,971 | 1,131 | 0,91 | -0,09 | 1,9E-01 |
| O08582 | GTP-binding protein 1 OS=Mus musculus OX=10090 GN=Gtpbp1 | 4 | 2 | 4 | 2 | 668 | 72,3 | 8,29 | 739 | 6,23 | 1 | 1 | 1,091 | 1,148 | 1,09 | 1,213 | 1,242 | 1,048 | 1,332 | 1,424 | 0,90 | -0,10 | 1,9E-01 |
| P20029 | Endoplasmic reticulum chaperone BiP OS=Mus musculus OX=10 | 45 | 30 | 77 | 28 | 655 | 72,4 | 5,16 | 26941 | 94,95 | 21 | 17 | 1,007 | 0,98 | 1,011 | 1,097 | 0,964 | 0,953 | 1,023 | 0,98 | 1,04 | 0,04 | 1,9E-01 |
| P19253 | 60S ribosomal protein L13a OS=Mus musculus OX=10090 GN=Rp | 13 | 3 | 3 | 3 | 203 | 23,4 | 11,02 | 1738 | 2,5 | 2 | 1 | 1,134 | 0,879 | 1,103 | 1,187 | 0,845 | 0,958 | 0,99 | 1,041 | 1,12 | 0,12 | 1,9E-01 |
| Q9DCJ5 | NADH dehydrogenase [ubiquinone] 1 alpha subcomplex subunit | 48 | 8 | 15 | 8 | 172 | 20 | 8,46 | 5099 | 21,82 | 4 | 6 | 1,013 | 0,912 | 1,005 | 0,976 | 0,953 | 0,992 | 0,871 | 0,853 | 1,06 | 0,06 | 1,9E-01 |
| A2RT62 | F-box/LRR-repeat protein 16 OS=Mus musculus OX=10090 GN=F | 2 | 1 | 1 | 1 | 479 | 51,8 | 6,6 | | 3,29 | | 1 | 0,881 | 0,971 | 0,776 | 1,014 | 1,314 | 1,058 | 0,866 | 1,031 | 0,85 | -0,15 | 1,9E-01 |
| P62774 | Myotrophin OS=Mus musculus OX=10090 GN=Mtpn PE=1 SV=2 | 41 | 6 | 10 | 6 | 118 | 12,9 | 5,52 | 3892 | 10,87 | 4 | 4 | 1,011 | 0,872 | 0,941 | 0,978 | 0,787 | 0,976 | 0,84 | 0,902 | 1,08 | 0,08 | 1,9E-01 |
| O70435 | Proteasome subunit alpha type-3 OS=Mus musculus OX=10090 | 36 | 12 | 22 | 12 | 255 | 28,4 | 5,44 | 9856 | 18,12 | 8 | 5 | 0,975 | 0,93 | 1,049 | 1,077 | 0,735 | 1,078 | 0,887 | 0,871 | 1,13 | 0,13 | 1,9E-01 |
| P97300-3 | Isoform 3 of Neuroplastin OS=Mus musculus OX=10090 GN=Npt | 31 | 8 | 21 | 1 | 277 | 30,8 | 7,49 | 3204 | 42,59 | 3 | 5 | 0,827 | 1,015 | 0,992 | 0,811 | 0,752 | 1,263 | 1,67 | 1,12 | 0,76 | -0,24 | 1,9E-01 |
| Q7TNC6 | Kinesin-like protein KIF26B OS=Mus musculus OX=10090 GN=Kif | 0 | 1 | 1 | 1 | 2112 | 225,4 | 8,5 | 388 | | 1 | | 1 | 0,856 | 0,899 | 0,844 | 0,714 | 0,885 | 0,892 | 0,78 | 1,10 | 0,10 | 1,9E-01 |
| Q6RHR9 | Membrane-associated guanylate kinase, WW and PDZ domain-c | 3 | 4 | 4 | 4 | 1471 | 161,9 | 7,36 | 2072 | | 4 | | 0,913 | 0,925 | 1,346 | 1,065 | 0,921 | 0,819 | 0,99 | 0,892 | 1,17 | 0,17 | 1,9E-01 |
| Q9CS84 | Neurexin-1 OS=Mus musculus OX=10090 GN=Nrxn1 PE=1 SV=3 | 11 | 14 | 28 | 11 | 1514 | 166,1 | 6,09 | 9194 | 25,91 | 9 | 7 | 0,979 | 0,998 | 0,975 | 1,123 | 1,052 | 1,223 | 1,285 | 0,981 | 0,90 | -0,10 | 1,9E-01 |
| Q8CAA7 | Glucose 1,6-bisphosphate synthase OS=Mus musculus OX=1009 | 13 | 11 | 18 | 11 | 621 | 70,2 | 6,49 | 8507 | 8,68 | 7 | 4 | 0,964 | 0,948 | 0,979 | 0,999 | 0,857 | 0,997 | 0,942 | 0,911 | 1,05 | 0,05 | 1,9E-01 |
| Q9D3R6 | Katanin p60 ATPase-containing subunit A-like 2 OS=Mus muscul | 2 | 1 | 1 | 1 | 539 | 61,1 | 7,52 | 913 | | 1 | | 1,044 | 0,845 | 1,201 | 0,998 | 0,359 | 1,034 | 0,97 | 0,732 | 1,32 | 0,32 | 1,9E-01 |
| Q9WVK4 | EH domain-containing protein 1 OS=Mus musculus OX=10090 G | 19 | 10 | 15 | 9 | 534 | 60,6 | 6,83 | 6692 | 6,21 | 7 | 3 | 0,996 | 1,012 | 1,081 | 0,963 | 1,034 | 1,042 | 1,044 | 1,114 | 0,96 | -0,04 | 1,9E-01 |
| P61294 | Ras-related protein Rab-6B OS=Mus musculus OX=10090 GN=Ra | 46 | 13 | 21 | 7 | 208 | 23,4 | 5,53 | 12543 | 8,75 | 10 | 3 | 0,958 | 1,014 | 1,07 | 0,926 | 0,815 | 0,966 | 0,904 | 0,989 | 1,08 | 0,08 | 1,9E-01 |
| Q62167 | ATP-dependent RNA helicase DDX3X OS=Mus musculus OX=100 | 16 | 11 | 15 | 10 | 662 | 73,1 | 7,18 | 7825 | 2,62 | 10 | 1 | 0,968 | 0,742 | 0,886 | 0,862 | 1,173 | 0,8 | 0,943 | 1,101 | 0,86 | -0,14 | 1,9E-01 |
| P13541 | Myosin-3 OS=Mus musculus OX=10090 GN=Myh3 PE=2 SV=2 | 4 | 7 | 7 | 1 | 1940 | 223,7 | 5,81 | 1571 | 8,31 | 4 | 3 | 0,712 | 0,676 | 0,713 | 0,719 | 3,342 | 0,87 | 0,952 | 1,123 | 0,45 | -0,55 | 1,9E-01 |
| Q8CG46 | Structural maintenance of chromosomes protein 5 OS=Mus mus | 1 | 1 | 1 | 1 | 1101 | 128,7 | 8,56 | 299 | | 1 | | 1,2 | 0,955 | 1,049 | 1,122 | 0,622 | 0,853 | 1,144 | 0,991 | 1,20 | 0,20 | 1,9E-01 |
| Q07456 | Protein AMBP OS=Mus musculus OX=10090 GN=Ambp PE=1 SV= | 3 | 1 | 1 | 1 | 349 | 39 | 6,32 | 741 | | 1 | | 0,857 | 0,932 | 0,894 | 0,83 | 1,508 | 0,766 | 1,038 | 1,106 | 0,80 | -0,20 | 1,9E-01 |
| Q6PA06 | Atlastin-2 OS=Mus musculus OX=10090 GN=Atl2 PE=1 SV=1 | 9 | 4 | 6 | 3 | 583 | 66,2 | 5,43 | 3063 | 2,13 | 3 | 1 | 0,821 | 0,971 | 0,834 | 0,861 | 1,415 | 0,871 | 0,906 | 1,048 | 0,82 | -0,18 | 1,9E-01 |
| P62242 | 40S ribosomal protein S8 OS=Mus musculus OX=10090 GN=Rps | 41 | 9 | 13 | 9 | 208 | 24,2 | 10,32 | 4272 | 11,61 | 5 | 4 | 1,01 | 0,928 | 1,045 | 1,047 | 0,793 | 0,997 | 0,96 | 0,966 | 1,08 | 0,08 | 1,9E-01 |
| Q9JJK2 | LanC-like protein 2 OS=Mus musculus OX=10090 GN=Lancl2 PE= | 2 | 1 | 2 | 1 | 450 | 50,7 | 7,28 | 1765 | | 1 | | 1,169 | 0,936 | 0,959 | 0,928 | 0,551 | 0,923 | 1,032 | 0,796 | 1,21 | 0,21 | 1,9E-01 |
| P46660 | Alpha-internexin OS=Mus musculus OX=10090 GN=Ina PE=1 SV= | 28 | 12 | 14 | 9 | 501 | 55,3 | 5,4 | 4765 | 18,99 | 6 | 6 | 1,024 | 0,97 | 1,162 | 0,962 | 0,687 | 0,935 | 1,047 | 0,931 | 1,14 | 0,14 | 2,0E-01 |
| Q3UMU9 | Hepatoma-derived growth factor-related protein 2 OS=Mus mus | 2 | 2 | 3 | 1 | 669 | 74,2 | 8,66 | 1456 | | 2 | | 0,664 | 0,551 | 0,537 | 0,381 | 1,971 | 0,562 | 0,863 | 0,659 | 0,53 | -0,47 | 2,0E-01 |

| | | | | | | | | | | | | | | | | | | | | | | | |
|---|---|---|---|---|---|---|---|---|---|---|---|---|---|---|---|---|---|---|---|---|---|---|---|
| Q3U2A8 | Valine--tRNA ligase, mitochondrial OS=Mus musculus OX=10090 | 2 | 2 | 2 | 2 | 1060 | 118,4 | 7,14 | 637 | | 2 | | 1,195 | 1,025 | 1,185 | 1,233 | 1,085 | 1,072 | 1,065 | 1,134 | 1,06 | 0,06 | 2,0E-01 |
| P34152 | Focal adhesion kinase 1 OS=Mus musculus OX=10090 GN=Ptk2 F | 1 | 2 | 2 | 2 | 1052 | 119,2 | 6,62 | 558 | 2,14 | 1 | 1 | 1,026 | 0,96 | 0,877 | 0,976 | 1,081 | 0,959 | 1,033 | 1 | 0,94 | -0,06 | 2,0E-01 |
| Q3UH93 | Plexin-D1 OS=Mus musculus OX=10090 GN=Plxnd1 PE=1 SV=1 | 4 | 6 | 6 | 6 | 1925 | 211,5 | 7,11 | 1917 | 6,89 | 3 | 3 | 0,954 | 1,07 | 0,906 | 0,969 | 1,062 | 0,951 | 1,045 | 1,158 | 0,92 | -0,08 | 2,0E-01 |
| Q8BGD5 | Carnitine O-palmitoyltransferase 1, brain isoform OS=Mus musc | 2 | 1 | 1 | 1 | 798 | 90 | 7,49 | 352 | | 1 | | 1,174 | 1,784 | 0,947 | 1,657 | 1,224 | 1,063 | 1,132 | 0,947 | 1,27 | 0,27 | 2,0E-01 |
| Q9D906 | Ubiquitin-like modifier-activating enzyme ATG7 OS=Mus muscul | 1 | 1 | 1 | 1 | 698 | 77,5 | 6,4 | | 2,25 | | 1 | 1,258 | 1,341 | 1,232 | 1,344 | 1,276 | 1,58 | 1,27 | 1,664 | 0,89 | -0,11 | 2,0E-01 |
| Q9ERD7 | Tubulin beta-3 chain OS=Mus musculus OX=10090 GN=Tubb3 PI | 56 | 22 | 393 | 6 | 450 | 50,4 | 4,93 | 167952 | 313,79 | 18 | 17 | 1,013 | 0,919 | 0,949 | 0,921 | 0,765 | 0,867 | 0,997 | 0,87 | 1,09 | 0,09 | 2,0E-01 |
| Q99PV0 | Pre-mRNA-processing-splicing factor 8 OS=Mus musculus OX=10 | 2 | 4 | 5 | 4 | 2335 | 273,4 | 8,84 | 2331 | | 4 | | 0,866 | 1,015 | 1,065 | 1,007 | 1,179 | 0,995 | 1,08 | 1,036 | 0,92 | -0,08 | 2,0E-01 |
| Q61937 | Nucleophosmin OS=Mus musculus OX=10090 GN=Npm1 PE=1 S' | 13 | 4 | 5 | 4 | 292 | 32,5 | 4,77 | 916 | 4,49 | 1 | 3 | 0,997 | 0,826 | 1,017 | 1,068 | 0,866 | 0,933 | 0,924 | 0,863 | 1,09 | 0,09 | 2,0E-01 |
| Q9CRD0 | OCIA domain-containing protein 1 OS=Mus musculus OX=10090 | 12 | 3 | 3 | 3 | 247 | 27,6 | 7,81 | 1229 | 3,56 | 2 | 1 | 0,98 | 1,011 | 0,936 | 0,989 | 0,905 | 0,968 | 0,978 | 0,932 | 1,04 | 0,04 | 2,0E-01 |
| Q9QYA2 | Mitochondrial import receptor subunit TOM40 homolog OS=Mu | 3 | 1 | 1 | 1 | 361 | 37,9 | 7,74 | 580 | | 1 | | 1,519 | 1,294 | 1,051 | 1,012 | 0,929 | 0,925 | 1,169 | 1,088 | 1,19 | 0,19 | 2,0E-01 |
| Q62425 | Cytochrome c oxidase subunit NDUFA4 OS=Mus musculus OX=1 | 56 | 5 | 11 | 5 | 82 | 9,3 | 9,52 | 4351 | 10,51 | 3 | 2 | 1,029 | 0,886 | 1,024 | 0,913 | 0,813 | 0,941 | 0,934 | 0,891 | 1,08 | 0,08 | 2,0E-01 |
| Q923X4 | Glutaredoxin-2, mitochondrial OS=Mus musculus OX=10090 GN | 6 | 1 | 1 | 1 | 156 | 17,3 | 9,95 | 706 | | 1 | | 0,861 | 0,606 | 1,229 | 1,22 | 1,6 | 1,102 | 1,132 | 1,182 | 0,78 | -0,22 | 2,0E-01 |
| Q7TPB0 | Phospholipid phosphatase-related protein type 3 OS=Mus musc | 15 | 9 | 14 | 9 | 716 | 76,6 | 6,01 | 2970 | 28,75 | 4 | 6 | 1,025 | 1,056 | 0,987 | 0,985 | 0,928 | 0,987 | 1,008 | 0,99 | 1,04 | 0,04 | 2,0E-01 |
| Q9EPW0 | Inositol polyphosphate-4-phosphatase type I A OS=Mus musculu | 4 | 4 | 5 | 4 | 939 | 105,5 | 7,05 | 1950 | 7,2 | 2 | 2 | 1,051 | 1,103 | 1,094 | 1,092 | 1,29 | 1,191 | 1,06 | 1,101 | 0,93 | -0,07 | 2,0E-01 |
| Q9D924 | Iron-sulfur cluster assembly 1 homolog, mitochondrial OS=Mus | 12 | 1 | 1 | 1 | 129 | 14,2 | 9,07 | 539 | | 1 | | 0,91 | 1,056 | 1,187 | 1,13 | 0,58 | 0,915 | 1,078 | 0,994 | 1,20 | 0,20 | 2,0E-01 |
| P70372 | ELAV-like protein 1 OS=Mus musculus OX=10090 GN=Elavl1 PE= | 17 | 5 | 6 | 5 | 326 | 36,1 | 9,04 | 2145 | 5,39 | 4 | 1 | 0,878 | 0,826 | 0,826 | 1,014 | 1,119 | 0,86 | 0,993 | 0,97 | 0,90 | -0,10 | 2,0E-01 |
| O35114 | Lysosome membrane protein 2 OS=Mus musculus OX=10090 GN | 9 | 3 | 4 | 3 | 478 | 54 | 5,1 | 478 | 10,05 | 1 | 2 | 1,134 | 0,959 | 1,25 | 1,196 | 0,978 | 1,087 | 1,036 | 1,052 | 1,09 | 0,09 | 2,0E-01 |
| Q6P5E4 | UDP-glucose:glycoprotein glucosyltransferase 1 OS=Mus muscul | 5 | 8 | 9 | 8 | 1551 | 176,3 | 5,62 | 2109 | 8,97 | 4 | 5 | 0,925 | 0,982 | 0,916 | 0,981 | 1,262 | 0,903 | 1,059 | 1,021 | 0,90 | -0,10 | 2,0E-01 |
| D3YVF0 | A-kinase anchor protein 5 OS=Mus musculus OX=10090 GN=Aka | 30 | 10 | 13 | 10 | 745 | 79,4 | 4,75 | 4685 | 13,86 | 7 | 4 | 1,111 | 1,228 | 0,911 | 1,016 | 1,029 | 0,912 | 0,939 | 0,972 | 1,11 | 0,11 | 2,0E-01 |
| Q9CQN1 | Heat shock protein 75 kDa, mitochondrial OS=Mus musculus OX | 15 | 9 | 12 | 8 | 706 | 80,2 | 6,68 | 5437 | 4,67 | 8 | 2 | 0,775 | 0,746 | 1,039 | 0,763 | 1,228 | 0,815 | 1,038 | 0,897 | 0,84 | -0,16 | 2,0E-01 |
| Q9D7X3 | Dual specificity protein phosphatase 3 OS=Mus musculus OX=10 | 23 | 3 | 5 | 3 | 185 | 20,5 | 6,54 | 2716 | 3,65 | 3 | 1 | 1,153 | 1,164 | 0,975 | 1,204 | 1,498 | 1,133 | 1,134 | 1,308 | 0,89 | -0,11 | 2,0E-01 |
| Q3TWL2 | Type 1 phosphatidylinositol 4,5-bisphosphate 4-phosphatase OS | 4 | 1 | 1 | 1 | 284 | 30 | 8,82 | | 0 | | 1 | 1,031 | 0,745 | 0,972 | 1,206 | 1,471 | 1,09 | 1,335 | 0,936 | 0,82 | -0,18 | 2,0E-01 |
| Q9CQL7 | MORF4 family-associated protein 1 OS=Mus musculus OX=1009 | 6 | 1 | 2 | 1 | 125 | 14,2 | 4,73 | 719 | | 1 | | 1,019 | 0,85 | 1,159 | 0,879 | 0,725 | 0,84 | 0,98 | 0,857 | 1,15 | 0,15 | 2,0E-01 |
| P11983 | T-complex protein 1 subunit alpha OS=Mus musculus OX=10090 | 24 | 14 | 27 | 14 | 556 | 60,4 | 6,16 | 14875 | 8,13 | 13 | 3 | 1,034 | 0,978 | 1,105 | 0,945 | 0,845 | 1,038 | 0,946 | 0,931 | 1,08 | 0,08 | 2,0E-01 |
| Q9QZB6 | Nuclear receptor subfamily 4 group A member 3 OS=Mus muscu | 2 | 1 | 3 | 1 | 627 | 68,4 | 7,88 | 1172 | | 1 | | 1,223 | 0,926 | 0,688 | 0,775 | 1,383 | 0,883 | 1,099 | 1,138 | 0,80 | -0,20 | 2,0E-01 |
| Q6ZWN5 | 40S ribosomal protein S9 OS=Mus musculus OX=10090 GN=Rps9 | 30 | 7 | 10 | 7 | 194 | 22,6 | 10,65 | 5216 | 5,66 | 5 | 2 | 0,913 | 0,88 | 0,971 | 1,002 | 1,157 | 1,001 | 0,935 | 0,988 | 0,92 | -0,08 | 2,0E-01 |
| Q9Z1R2 | Large proline-rich protein BAG6 OS=Mus musculus OX=10090 GI | 5 | 3 | 4 | 3 | 1154 | 121 | 5,71 | 1926 | | 3 | | 1,116 | 1,05 | 1,161 | 1,016 | 0,873 | 1,17 | 0,991 | 0,856 | 1,12 | 0,12 | 2,0E-01 |
| P18654 | Ribosomal protein S6 kinase alpha-3 OS=Mus musculus OX=1009 | 5 | 3 | 3 | 3 | 740 | 83,6 | 6,89 | 1692 | | 3 | | 0,958 | 0,919 | 0,988 | 1,072 | 0,744 | 0,984 | 0,894 | 0,957 | 1,10 | 0,10 | 2,0E-01 |
| Q8BXK8 | Arf-GAP with GTPase, ANK repeat and PH domain-containing pro | 9 | 7 | 9 | 4 | 857 | 94,4 | 7,94 | 3292 | 8,95 | 5 | 2 | 0,969 | 1,077 | 0,989 | 0,945 | 1,097 | 1,033 | 1,069 | 0,991 | 0,95 | -0,05 | 2,0E-01 |
| P08414 | Calcium/calmodulin-dependent protein kinase type IV OS=Mus r | 17 | 9 | 12 | 9 | 469 | 52,6 | 4,93 | 3420 | 13,42 | 4 | 6 | 1,016 | 0,975 | 1,126 | 0,952 | 0,686 | 1,01 | 0,973 | 0,929 | 1,13 | 0,13 | 2,0E-01 |
| Q9WV69 | Dematin OS=Mus musculus OX=10090 GN=Dmtn PE=1 SV=1 | 17 | 6 | 8 | 6 | 405 | 45,4 | 8,41 | 3214 | 4,9 | 4 | 2 | 1,025 | 0,941 | 1,118 | 0,929 | 1,106 | 1,099 | 1,005 | 1,086 | 0,93 | -0,07 | 2,0E-01 |
| Q920A5 | Retinoid-inducible serine carboxypeptidase OS=Mus musculus C | 2 | 1 | 1 | 1 | 452 | 50,9 | 5,66 | 573 | | 1 | | 1,299 | 0,941 | 1,186 | 1,313 | 0,928 | 1,247 | 1 | 0,898 | 1,16 | 0,16 | 2,0E-01 |
| P16015 | Carbonic anhydrase 3 OS=Mus musculus OX=10090 GN=Ca3 PE= | 13 | 3 | 5 | 3 | 260 | 29,3 | 7,37 | 3581 | | 3 | | 1,217 | 1,523 | 1,39 | 1,455 | 3,846 | 1,5 | 1,887 | 1,546 | 0,64 | -0,36 | 2,0E-01 |
| Q3V038 | Tetratricopeptide repeat protein 9A OS=Mus musculus OX=1009 | 12 | 2 | 2 | 2 | 219 | 24,3 | 8,85 | 1124 | | 2 | | 0,876 | 1,25 | 1,105 | 1,14 | 1,171 | 0,863 | 0,906 | 0,764 | 1,18 | 0,18 | 2,0E-01 |
| P28667 | MARCKS-related protein OS=Mus musculus OX=10090 GN=Marc | 70 | 9 | 27 | 9 | 200 | 20,2 | 4,61 | 9498 | 47,67 | 2 | 8 | 1,045 | 0,893 | 1,058 | 1,006 | 0,628 | 1,138 | 0,846 | 0,725 | 1,20 | 0,20 | 2,0E-01 |
| Q9D020 | Cytosolic 5'-nucleotidase 3A OS=Mus musculus OX=10090 GN=N | 7 | 2 | 3 | 2 | 331 | 37,2 | 6,65 | 1648 | | 2 | | 0,888 | 0,947 | 1,114 | 0,884 | 1,278 | 1,064 | 0,859 | 1,287 | 0,85 | -0,15 | 2,0E-01 |
| P31324 | cAMP-dependent protein kinase type II-beta regulatory subunit | 50 | 18 | 31 | 14 | 416 | 46,1 | 4,98 | 11123 | 33,39 | 10 | 8 | 1,038 | 0,97 | 1,042 | 0,993 | 0,762 | 0,985 | 0,949 | 1,011 | 1,09 | 0,09 | 2,0E-01 |
| Q59J78 | NADH dehydrogenase [ubiquinone] 1 alpha subcomplex assemb | 14 | 2 | 4 | 2 | 168 | 19,6 | 8,25 | 2798 | | 2 | | 0,931 | 0,998 | 1,04 | 1,279 | 0,917 | 0,91 | 0,991 | 0,983 | 1,12 | 0,12 | 2,1E-01 |
| Q9D1K7 | UPF0687 protein C20orf27 homolog OS=Mus musculus OX=1009 | 6 | 1 | 1 | 1 | 174 | 19,5 | 6,68 | 678 | | 1 | | 0,77 | 1,025 | 1,002 | 1,137 | 0,589 | 0,896 | 0,999 | 0,787 | 1,20 | 0,20 | 2,1E-01 |
| O35188 | Fractalkine OS=Mus musculus OX=10090 GN=Cx3cl1 PE=1 SV=3 | 3 | 1 | 1 | 1 | 395 | 42,1 | 5,54 | 275 | | 1 | | 0,924 | 0,997 | 0,978 | 0,766 | 0,725 | 0,944 | 0,651 | 0,865 | 1,15 | 0,15 | 2,1E-01 |
| Q04690 | Neurofibromin OS=Mus musculus OX=10090 GN=Nf1 PE=1 SV=1 | 3 | 7 | 8 | 7 | 2841 | 319,4 | 7,39 | 2585 | 2,38 | 6 | 1 | 0,992 | 1,003 | 0,913 | 1,068 | 1,033 | 1,003 | 1,08 | 1,065 | 0,95 | -0,05 | 2,1E-01 |
| Q3TW96 | UDP-N-acetylhexosamine pyrophosphorylase-like protein 1 OS= | 2 | 1 | 1 | 1 | 507 | 56,6 | 5,43 | 672 | | 1 | | 1,165 | 1,554 | 1,294 | 1,388 | 2,229 | 1,283 | 1,617 | 1,509 | 0,81 | -0,19 | 2,1E-01 |
| P06151 | L-lactate dehydrogenase A chain OS=Mus musculus OX=10090 G | 38 | 16 | 27 | 15 | 332 | 36,5 | 7,74 | 12138 | 21,5 | 10 | 6 | 0,973 | 1,012 | 1,025 | 1,027 | 0,915 | 1,061 | 0,921 | 0,93 | 1,05 | 0,05 | 2,1E-01 |
| Q91VF2 | Histamine N-methyltransferase OS=Mus musculus OX=10090 GI | 8 | 3 | 3 | 3 | 295 | 33,6 | 5,06 | 1465 | | 3 | | 0,945 | 0,78 | 0,778 | 0,928 | 0,86 | 0,856 | 1,086 | 1,399 | 0,82 | -0,18 | 2,1E-01 |
| P20918 | Plasminogen OS=Mus musculus OX=10090 GN=Plg PE=1 SV=3 | 5 | 2 | 3 | 2 | 812 | 90,7 | 6,6 | 74 | 5,24 | 1 | 1 | 0,798 | 0,795 | 0,812 | 0,824 | 2,308 | 0,912 | 0,98 | 0,955 | 0,63 | -0,37 | 2,1E-01 |
| Q61771 | Kinesin-like protein KIF3B OS=Mus musculus OX=10090 GN=Kif3 | 11 | 8 | 12 | 6 | 747 | 85,2 | 7,69 | 6486 | 5,71 | 7 | 2 | 0,95 | 0,829 | 0,986 | 0,891 | 0,866 | 0,899 | 0,871 | 0,75 | 1,08 | 0,08 | 2,1E-01 |
| O55033 | Cytoplasmic protein NCK2 OS=Mus musculus OX=10090 GN=Ncl | 8 | 4 | 5 | 4 | 380 | 42,9 | 6,95 | 2424 | 2,66 | 3 | 1 | 0,994 | 0,95 | 1,073 | 1,034 | 0,799 | 1,043 | 0,964 | 0,92 | 1,09 | 0,09 | 2,1E-01 |
| Q99LD4 | COP9 signalosome complex subunit 1 OS=Mus musculus OX=10( | 9 | 3 | 4 | 3 | 471 | 53,4 | 6,84 | 1953 | 2,74 | 2 | 1 | 0,858 | 1,087 | 0,981 | 1,039 | 1,396 | 0,882 | 1,252 | 1,114 | 0,85 | -0,15 | 2,1E-01 |
| G3X9K3 | Brefeldin A-inhibited guanine nucleotide-exchange protein 1 OS | 1 | 2 | 2 | 2 | 1846 | 208,4 | 5,86 | 272 | 2,02 | 1 | 1 | 0,834 | 0,752 | 0,831 | 0,764 | 1,107 | 0,789 | 0,844 | 0,859 | 0,88 | -0,12 | 2,1E-01 |
| Q9Z0H8 | CAP-Gly domain-containing linker protein 2 OS=Mus musculus C | 38 | 33 | 45 | 1 | 1047 | 115,8 | 6,48 | 19856 | 30,84 | 23 | 10 | 0,917 | 0,951 | 0,848 | 0,828 | 1,105 | 0,811 | 1,043 | 0,978 | 0,90 | -0,10 | 2,1E-01 |
| Q9D0V7 | Receptor-binding cancer antigen expressed on SiSo cells OS=Mu | 11 | 1 | 1 | 1 | 213 | 24,3 | 6,29 | 97 | | 1 | | 1,19 | 1,374 | 0,853 | 1,62 | 0,71 | 0,895 | 1,211 | 1,111 | 1,28 | 0,28 | 2,1E-01 |
| Q6PCN3 | Tau-tubulin kinase 1 OS=Mus musculus OX=10090 GN=Ttbk1 PE | 3 | 3 | 3 | 3 | 1308 | 141,5 | 5,62 | | 1,92 | | 3 | 0,834 | 0,946 | 0,969 | 0,829 | 1,104 | 0,906 | 0,947 | 0,944 | 0,92 | -0,08 | 2,1E-01 |
| P53811 | Phosphatidylinositol transfer protein beta isoform OS=Mus mus | 17 | 4 | 4 | 3 | 271 | 31,5 | 6,95 | 1853 | 2,57 | 3 | 1 | 0,86 | 1,481 | 0,815 | 1,169 | 1,643 | 1,035 | 1,633 | 1,234 | 0,78 | -0,22 | 2,1E-01 |
| Q8VC30 | Triokinase/FMN cyclase OS=Mus musculus OX=10090 GN=Tkfc F | 12 | 5 | 7 | 5 | 578 | 59,7 | 6,92 | 3924 | 3,31 | 5 | 1 | 1,334 | 1,395 | 1,218 | 1,444 | 2,103 | 1,36 | 1,444 | 1,484 | 0,84 | -0,16 | 2,1E-01 |
| Q9R0Q7 | Prostaglandin E synthase 3 OS=Mus musculus OX=10090 GN=Ptg | 29 | 4 | 7 | 4 | 160 | 18,7 | 4,55 | 2069 | 9,22 | 2 | 3 | 0,873 | 1,004 | 0,932 | 0,902 | 0,985 | 0,97 | 0,971 | 0,949 | 0,96 | -0,04 | 2,1E-01 |
| Q99KG3 | RNA-binding protein 10 OS=Mus musculus OX=10090 GN=Rbm1 | 2 | 1 | 1 | 1 | 930 | 103,4 | 5,92 | 104 | | 1 | | 1,548 | 2,422 | 0,617 | 0,856 | 1,343 | 2,5 | 1,909 | 5,919 | 0,47 | -0,53 | 2,1E-01 |
| P50427 | Steryl-sulfatase OS=Mus musculus OX=10090 GN=Sts PE=1 SV=1 | 3 | 1 | 1 | 1 | 624 | 66,5 | 8,5 | 228 | | 1 | | 0,965 | 1,052 | 0,702 | 0,891 | 1,742 | 0,64 | 1,186 | 1,414 | 0,72 | -0,28 | 2,1E-01 |
| Q9CQT1 | Methylthioribose-1-phosphate isomerase OS=Mus musculus OX | 11 | 3 | 3 | 3 | 369 | 39,4 | 5,91 | 1178 | | 3 | | 0,752 | 0,687 | 1,463 | 0,931 | 0,797 | 0,695 | 0,447 | 0,799 | 1,40 | 0,40 | 2,1E-01 |
| Q8JZR6 | Electroneutral sodium bicarbonate exchanger 1 OS=Mus muscul | 2 | 1 | 1 | 1 | 1089 | 122,3 | 6,6 | 495 | | 1 | | 0,846 | 1,252 | 0,77 | 0,974 | 1,4 | 0,886 | 1,542 | 1,054 | 0,79 | -0,21 | 2,1E-01 |
| Q810B6 | Rabankyrin-5 OS=Mus musculus OX=10090 GN=Ankfy1 PE=1 SV= | 3 | 3 | 3 | 3 | 1169 | 128,6 | 5,91 | 1485 | | 3 | | 1,313 | 1,483 | 1,058 | 1,342 | 1,432 | 1,384 | 1,366 | 1,586 | 0,90 | -0,10 | 2,1E-01 |
| Q99PJ0 | Neurotrimin OS=Mus musculus OX=10090 GN=Ntm PE=1 SV=2 | 28 | 8 | 17 | 8 | 344 | 38 | 7,81 | 7140 | 15,35 | 6 | 4 | 0,964 | 1,044 | 0,943 | 0,961 | 0,773 | 0,961 | 0,875 | 0,996 | 1,09 | 0,09 | 2,1E-01 |
| Q5SWU9 | Acetyl-CoA carboxylase 1 OS=Mus musculus OX=10090 GN=Acac | 8 | 18 | 21 | 18 | 2345 | 265,1 | 6,39 | 8476 | 6,43 | 15 | 3 | 1,067 | 1,082 | 0,97 | 1,05 | 2,767 | 1,587 | 1,012 | 1,086 | 0,65 | -0,35 | 2,1E-01 |
| Q61316 | Heat shock 70 kDa protein 4 OS=Mus musculus OX=10090 GN=H | 38 | 30 | 53 | 27 | 841 | 94,1 | 5,24 | 18133 | 61,64 | 15 | 18 | 1,024 | 0,892 | 1,05 | 0,997 | 0,769 | 1,02 | 0,897 | 0,927 | 1,10 | 0,10 | 2,1E-01 |
| E9PVA8 | eIF-2-alpha kinase activator GCN1 OS=Mus musculus OX=10090 | 2 | 5 | 6 | 5 | 2671 | 292,8 | 7,36 | 2309 | 1,85 | 4 | 1 | 1,098 | 1,12 | 1,099 | 1,088 | 0,984 | 1,077 | 1,066 | 1,117 | 1,04 | 0,04 | 2,1E-01 |
| Q9CQ21 | Malignant T-cell-amplified sequence 2 OS=Mus musculus OX=10 | 7 | 1 | 3 | 1 | 181 | 20,4 | 9,11 | 1044 | 2,09 | 1 | 1 | 0,896 | 0,945 | 0,916 | 1,122 | 1,147 | 1,164 | 0,956 | 1,017 | 0,91 | -0,09 | 2,1E-01 |

| | | | | | | | | | | | | | | | | | | | | | | | |
|---|---|---|---|---|---|---|---|---|---|---|---|---|---|---|---|---|---|---|---|---|---|---|---|
| Q8C7H1 | Methylmalonic aciduria type A homolog, mitochondrial OS=Mus | 3 | 1 | 1 | 1 | 415 | 45,9 | 9,32 | | 2,94 | | 1 | 0,959 | 1,005 | 1,173 | 1,02 | 0,601 | 0,778 | 0,952 | 1,134 | 1,20 | 0,20 | 2,1E-01 |
| Q8R3S6 | Exocyst complex component 1 OS=Mus musculus OX=10090 GN | 5 | 5 | 5 | 5 | 894 | 101,8 | 6,52 | 2406 | 2,64 | 4 | 1 | 1,043 | 1,058 | 0,983 | 1,062 | 0,725 | 1,033 | 0,939 | 1,031 | 1,11 | 0,11 | 2,1E-01 |
| P97370 | Sodium/potassium-transporting ATPase subunit beta-3 OS=Mus | 10 | 3 | 8 | 3 | 278 | 31,8 | 8,51 | 5312 | 2,23 | 3 | 1 | 0,977 | 0,949 | 1,085 | 1,008 | 0,794 | 1,034 | 0,925 | 0,944 | 1,09 | 0,09 | 2,1E-01 |
| Q9WUK2 | Eukaryotic translation initiation factor 4H OS=Mus musculus OX= | 22 | 6 | 10 | 6 | 248 | 27,3 | 7,23 | 4182 | 4,5 | 5 | 2 | 1,047 | 1,156 | 1,025 | 1,133 | 0,896 | 1,1 | 1,038 | 1,028 | 1,07 | 0,07 | 2,1E-01 |
| Q64514 | Tripeptidyl-peptidase 2 OS=Mus musculus OX=10090 GN=Tpp2 I | 15 | 15 | 22 | 15 | 1262 | 139,8 | 6,58 | 8849 | 17,81 | 11 | 6 | 1,039 | 0,98 | 1,018 | 1,021 | 1,037 | 1,092 | 0,987 | 1,145 | 0,95 | -0,05 | 2,1E-01 |
| Q9CR68 | Cytochrome b-c1 complex subunit Rieske, mitochondrial OS=Mu | 36 | 11 | 19 | 11 | 274 | 29,3 | 8,7 | 9684 | 10,53 | 9 | 3 | 1,065 | 0,994 | 1,09 | 1,063 | 0,868 | 0,977 | 1,058 | 1,042 | 1,07 | 0,07 | 2,1E-01 |
| P01942 | Hemoglobin subunit alpha OS=Mus musculus OX=10090 GN=Hb | 43 | 6 | 19 | 6 | 142 | 15,1 | 8,22 | 10440 | 4,48 | 6 | 1 | 0,905 | 1,18 | 0,875 | 0,868 | 2,003 | 0,881 | 1,111 | 1,251 | 0,73 | -0,27 | 2,1E-01 |
| Q8R3Z5 | Voltage-dependent L-type calcium channel subunit beta-1 OS=M | 7 | 5 | 5 | 3 | 597 | 65,4 | 6,62 | 1855 | 4,16 | 3 | 2 | 0,852 | 0,948 | 1,057 | 0,959 | 0,771 | 0,979 | 0,856 | 0,876 | 1,10 | 0,10 | 2,1E-01 |
| O09044 | Synaptosomal-associated protein 23 OS=Mus musculus OX=100 | 11 | 2 | 2 | 1 | 210 | 23,2 | 4,98 | 933 | | 2 | | 0,883 | 0,915 | 0,971 | 0,752 | 1,572 | 0,799 | 1,089 | 1,009 | 0,79 | -0,21 | 2,1E-01 |
| Q6PF93 | Phosphatidylinositol 3-kinase catalytic subunit type 3 OS=Mus m | 1 | 1 | 2 | 1 | 887 | 101,4 | 6,73 | 1061 | | 1 | | 1,132 | 0,96 | 1,06 | 0,856 | 1,251 | 1,409 | 1,038 | 0,966 | 0,86 | -0,14 | 2,1E-01 |
| O35526 | Syntaxin-1A OS=Mus musculus OX=10090 GN=Stx1a PE=1 SV=3 | 45 | 14 | 21 | 12 | 288 | 33 | 5,24 | 7175 | 15,12 | 8 | 6 | 1,021 | 1 | 1,284 | 0,964 | 0,832 | 0,904 | 1,087 | 0,94 | 1,13 | 0,13 | 2,1E-01 |
| F8VPU2 | FERM, ARHGEF and pleckstrin domain-containing protein 1 OS=I | 16 | 14 | 20 | 14 | 1048 | 118,8 | 7,88 | 8584 | 16,23 | 10 | 4 | 0,974 | 1,038 | 1,026 | 0,94 | 0,88 | 0,948 | 0,995 | 0,968 | 1,05 | 0,05 | 2,1E-01 |
| P26883 | Peptidyl-prolyl cis-trans isomerase FKBP1A OS=Mus musculus O | 29 | 2 | 6 | 2 | 108 | 11,9 | 8,16 | 2565 | | 2 | | 0,827 | 0,825 | 0,809 | 0,84 | 1,594 | 0,827 | 1,091 | 0,809 | 0,76 | -0,24 | 2,1E-01 |
| Q99J09 | Methylosome protein 50 OS=Mus musculus OX=10090 GN=Wdr | 5 | 1 | 3 | 1 | 342 | 36,9 | 5,27 | 1891 | | 1 | | 0,941 | 1,054 | 0,98 | 0,328 | 0,776 | 1,612 | 1,559 | 0,89 | 0,68 | -0,32 | 2,1E-01 |
| O08663 | Methionine aminopeptidase 2 OS=Mus musculus OX=10090 GN | 8 | 3 | 4 | 3 | 478 | 52,9 | 5,82 | 663 | 7,88 | 1 | 3 | 1,266 | 1,586 | 1,369 | 1,46 | 2,261 | 1,502 | 1,471 | 1,561 | 0,84 | -0,16 | 2,1E-01 |
| P14231 | Sodium/potassium-transporting ATPase subunit beta-2 OS=Mus | 27 | 6 | 20 | 6 | 290 | 33,3 | 8,31 | 5314 | 29,7 | 3 | 4 | 1,022 | 1,017 | 0,912 | 1,073 | 0,805 | 0,948 | 0,942 | 1,018 | 1,08 | 0,08 | 2,1E-01 |
| Q8BX70 | Vacuolar protein sorting-associated protein 13C OS=Mus muscu | 1 | 2 | 2 | 2 | 3748 | 419,8 | 6,81 | 715 | | 2 | | 1,057 | 0,818 | 0,96 | 0,879 | 1,283 | 1,32 | 0,825 | 1,002 | 0,84 | -0,16 | 2,1E-01 |
| Q9Z2K1 | Keratin, type I cytoskeletal 16 OS=Mus musculus OX=10090 GN= | 12 | 7 | 20 | 3 | 469 | 51,6 | 5,2 | 7298 | 14,91 | 4 | 4 | 1,066 | 1,259 | 0,918 | 0,89 | 1,227 | 1,366 | 1,019 | 1,141 | 0,87 | -0,13 | 2,1E-01 |
| Q9QXJ1 | Amyloid-beta A4 precursor protein-binding family B member 1 ( | 4 | 2 | 3 | 2 | 710 | 77,3 | 5 | 608 | 0 | 1 | 1 | 0,822 | 1,08 | 1,204 | 0,864 | 1,116 | 0,931 | 1,376 | 1,287 | 0,84 | -0,16 | 2,2E-01 |
| Q922H2 | [Pyruvate dehydrogenase (acetyl-transferring)] kinase isozyme 3 | 2 | 1 | 2 | 1 | 415 | 47,9 | 8,82 | | 4,64 | | 1 | 0,82 | 0,705 | 1,184 | 0,852 | 0,54 | 0,806 | 0,774 | 0,776 | 1,23 | 0,23 | 2,2E-01 |
| P63325 | 40S ribosomal protein S10 OS=Mus musculus OX=10090 GN=Rp: | 15 | 3 | 12 | 3 | 165 | 18,9 | 10,15 | 4898 | 19,32 | 2 | 2 | 1,413 | 1,319 | 1,151 | 1,533 | 1,733 | 1,277 | 1,496 | 1,608 | 0,89 | -0,11 | 2,2E-01 |
| Q9DCX2 | ATP synthase subunit d, mitochondrial OS=Mus musculus OX=10 | 68 | 12 | 26 | 12 | 161 | 18,7 | 5,69 | 4046 | 72,55 | 4 | 8 | 1,023 | 0,982 | 0,955 | 1,039 | 0,793 | 1,043 | 0,91 | 0,948 | 1,08 | 0,08 | 2,2E-01 |
| Q8VEA4 | Mitochondrial intermembrane space import and assembly prote | 6 | 1 | 2 | 1 | 139 | 15,5 | 4,32 | 1172 | | 1 | | 0,908 | 1,149 | 0,768 | 1,107 | 0,666 | 0,823 | 1,003 | 0,818 | 1,19 | 0,19 | 2,2E-01 |
| O88544 | COP9 signalosome complex subunit 4 OS=Mus musculus OX=10 | 33 | 11 | 14 | 11 | 406 | 46,3 | 5,83 | 5561 | 6,18 | 8 | 4 | 1,031 | 0,981 | 0,896 | 0,95 | 1,025 | 0,982 | 1,049 | 0,983 | 0,96 | -0,04 | 2,2E-01 |
| Q99MD9 | Nuclear autoantigenic sperm protein OS=Mus musculus OX=100 | 1 | 1 | 1 | 1 | 773 | 83,9 | 4,37 | | 1,97 | | 1 | 1,017 | 1,147 | 0,904 | 1,1 | 1,3 | 1,052 | 1,02 | 1,338 | 0,88 | -0,12 | 2,2E-01 |
| Q8CI08 | SLAIN motif-containing protein 2 OS=Mus musculus OX=10090 ( | 3 | 1 | 1 | 1 | 581 | 62,3 | 9,47 | 576 | | 1 | | 0,98 | 1,148 | 1,024 | 0,976 | 1,527 | 0,922 | 1,096 | 1,359 | 0,84 | -0,16 | 2,2E-01 |
| Q80YA7 | Dipeptidyl peptidase 8 OS=Mus musculus OX=10090 GN=Dpp8 F | 5 | 5 | 5 | 5 | 892 | 102,1 | 5,81 | 2148 | 0 | 4 | 1 | 0,974 | 0,928 | 0,853 | 0,943 | 1,11 | 0,917 | 0,931 | 1,026 | 0,93 | -0,07 | 2,2E-01 |
| Q8N9S3 | Activator of 90 kDa heat shock protein ATPase homolog 2 OS=M | 3 | 1 | 1 | 1 | 331 | 37,6 | 6,44 | 623 | | 1 | | 1,283 | 1,127 | 1,333 | 1,381 | 1,85 | 1,518 | 1,33 | 1,234 | 0,86 | -0,14 | 2,2E-01 |
| Q9D964 | Glycine amidinotransferase, mitochondrial OS=Mus musculus O | 1 | 1 | 1 | 1 | 423 | 48,3 | 7,88 | 548 | | 1 | | 1,001 | 0,816 | 0,944 | 1,099 | 0,926 | 0,879 | 0,78 | 0,904 | 1,11 | 0,11 | 2,2E-01 |
| Q62189 | U1 small nuclear ribonucleoprotein A OS=Mus musculus OX=10 | 7 | 2 | 2 | 2 | 287 | 31,8 | 9,8 | 749 | 2,67 | 1 | 1 | 0,893 | 0,835 | 0,965 | 1,054 | 0,749 | 0,852 | 0,933 | 0,877 | 1,10 | 0,10 | 2,2E-01 |
| Q6P4T2 | U5 small nuclear ribonucleoprotein 200 kDa helicase OS=Mus m | 2 | 4 | 4 | 4 | 2136 | 244,4 | 6,06 | 773 | 3,92 | 2 | 2 | 0,955 | 1,006 | 1,087 | 1,066 | 1,204 | 1,096 | 1,053 | 1,03 | 0,94 | -0,06 | 2,2E-01 |
| Q9WTI7 | Unconventional myosin-Ic OS=Mus musculus OX=10090 GN=My | 2 | 2 | 2 | 2 | 1063 | 121,9 | 9,35 | | 0 | | 2 | 1,079 | 1,005 | 1,006 | 1,338 | 2,875 | 1,492 | 1,113 | 1,231 | 0,66 | -0,34 | 2,2E-01 |
| Q5XG73 | Acyl-CoA-binding domain-containing protein 5 OS=Mus musculu | 6 | 2 | 3 | 2 | 508 | 56,6 | 5,41 | 1294 | | 2 | | 0,851 | 0,721 | 0,893 | 0,985 | 1,316 | 0,7 | 1,21 | 1,026 | 0,81 | -0,19 | 2,2E-01 |
| Q8BXR1 | Probable cationic amino acid transporter OS=Mus musculus OX= | 1 | 1 | 1 | 1 | 771 | 83,9 | 5,35 | 294 | | 1 | | 1,047 | 0,98 | 1,003 | 0,907 | 1,253 | 1,064 | 0,881 | 1,261 | 0,88 | -0,12 | 2,2E-01 |
| Q9CQ92 | Mitochondrial fission 1 protein OS=Mus musculus OX=10090 GN | 16 | 2 | 5 | 2 | 152 | 17 | 8,53 | 3057 | | 2 | | 1,023 | 1,156 | 0,994 | 1,27 | 0,911 | 1,087 | 1,021 | 1,02 | 1,10 | 0,10 | 2,2E-01 |
| P97315 | Cysteine and glycine-rich protein 1 OS=Mus musculus OX=1009( | 34 | 4 | 4 | 4 | 193 | 20,6 | 8,57 | 1935 | 3,19 | 3 | 1 | 1,051 | 1,045 | 1,043 | 0,994 | 0,702 | 0,961 | 1,123 | 0,854 | 1,14 | 0,14 | 2,2E-01 |
| P56135 | ATP synthase subunit f, mitochondrial OS=Mus musculus OX=10 | 20 | 2 | 4 | 2 | 88 | 10,3 | 9,95 | 2148 | 2,11 | 1 | 1 | 1,102 | 0,935 | 0,984 | 1,044 | 0,818 | 0,948 | 0,985 | 1,008 | 1,08 | 0,08 | 2,2E-01 |
| Q6PHZ2 | Calcium/calmodulin-dependent protein kinase type II subunit de | 18 | 10 | 24 | 4 | 499 | 56,3 | 7,25 | 8136 | 15,26 | 5 | 6 | 1,025 | 1,168 | 0,935 | 0,96 | 0,966 | 0,948 | 0,947 | 0,939 | 1,08 | 0,08 | 2,2E-01 |
| Q9JIS5 | Synaptic vesicle glycoprotein 2A OS=Mus musculus OX=10090 G | 17 | 11 | 12 | 10 | 742 | 82,6 | 5,57 | 5411 | 6,13 | 8 | 3 | 1,009 | 0,986 | 0,982 | 0,871 | 0,62 | 0,901 | 0,982 | 0,883 | 1,14 | 0,14 | 2,2E-01 |
| Q7TQH0 | Ataxin-2-like protein OS=Mus musculus OX=10090 GN=Atxn2l PI | 2 | 2 | 2 | 2 | 1049 | 110,6 | 8,85 | 1519 | | 2 | | 1,112 | 1,088 | 1,051 | 1,095 | 0,72 | 1,028 | 0,91 | 1,166 | 1,14 | 0,14 | 2,2E-01 |
| Q9WV91 | Prostaglandin F2 receptor negative regulator OS=Mus musculus | 7 | 6 | 6 | 6 | 879 | 98,7 | 6,61 | 2445 | 6,82 | 4 | 2 | 0,903 | 0,938 | 1,053 | 0,961 | 0,962 | 1,081 | 1,118 | 0,971 | 0,93 | -0,07 | 2,2E-01 |
| Q9JI11 | Serine/threonine-protein kinase 4 OS=Mus musculus OX=10090 | 3 | 1 | 1 | 1 | 487 | 55,5 | 5,19 | 880 | | 1 | | 1,148 | 0,883 | 1,077 | 1,056 | 0,886 | 0,699 | 1,022 | 1,032 | 1,14 | 0,14 | 2,2E-01 |
| P62878 | E3 ubiquitin-protein ligase RBX1 OS=Mus musculus OX=10090 G | 7 | 1 | 1 | 1 | 108 | 12,3 | 6,96 | 797 | | 1 | | 1,109 | 1,1 | 1,527 | 1,031 | 0,664 | 1,157 | 1,161 | 0,883 | 1,23 | 0,23 | 2,2E-01 |
| Q9DB60 | Prostamide/prostaglandin F synthase OS=Mus musculus OX=10 | 13 | 3 | 4 | 3 | 201 | 21,7 | 6,74 | 1995 | 2,31 | 2 | 1 | 1,001 | 1,033 | 1,077 | 0,944 | 0,956 | 1,011 | 0,909 | 0,985 | 1,05 | 0,05 | 2,2E-01 |
| Q921G7 | Electron transfer flavoprotein-ubiquinone oxidoreductase, mito | 17 | 8 | 9 | 8 | 616 | 68 | 7,58 | 4806 | | 8 | | 0,978 | 0,941 | 1,143 | 1,016 | 1,196 | 0,974 | 1,132 | 1,128 | 0,92 | -0,08 | 2,2E-01 |
| Q9DBL7 | Bifunctional coenzyme A synthase OS=Mus musculus OX=10090 | 4 | 2 | 2 | 2 | 563 | 62 | 7,11 | 696 | 2,67 | 1 | 1 | 1,001 | 0,82 | 1,134 | 0,928 | 0,636 | 0,793 | 0,922 | 0,981 | 1,17 | 0,17 | 2,2E-01 |
| P09671 | Superoxide dismutase [Mn], mitochondrial OS=Mus musculus O | 27 | 7 | 15 | 7 | 222 | 24,6 | 8,62 | 9039 | 2,92 | 5 | 2 | 0,966 | 1,037 | 0,864 | 1,106 | 0,784 | 0,985 | 0,931 | 0,908 | 1,10 | 0,10 | 2,2E-01 |
| Q99JY9 | Actin-related protein 3 OS=Mus musculus OX=10090 GN=Actr3 I | 32 | 12 | 27 | 10 | 418 | 47,3 | 5,88 | 12255 | 18,16 | 8 | 6 | 0,999 | 1,087 | 1,113 | 1,018 | 0,784 | 1,078 | 1,034 | 0,936 | 1,10 | 0,10 | 2,2E-01 |
| Q6ZWU9 | 40S ribosomal protein S27 OS=Mus musculus OX=10090 GN=Rp: | 38 | 3 | 4 | 3 | 84 | 9,5 | 9,45 | 1628 | 2,48 | 2 | 1 | 0,916 | 0,905 | 0,951 | 1,11 | 0,945 | 1,075 | 1,085 | 1,102 | 0,92 | -0,08 | 2,2E-01 |
| Q9ER88 | 28S ribosomal protein S29, mitochondrial OS=Mus musculus OX | 5 | 2 | 2 | 2 | 391 | 44,7 | 8,94 | 873 | | 2 | | 1,207 | 0,984 | 0,966 | 1,078 | 1,488 | 1,068 | 1,117 | 1,161 | 0,88 | -0,12 | 2,2E-01 |
| Q9JM14 | 5'(3')-deoxyribonucleotidase, cytosolic type OS=Mus musculus ( | 16 | 3 | 4 | 3 | 200 | 23,1 | 5,49 | 1357 | 2,74 | 2 | 1 | 0,803 | 0,984 | 0,829 | 1,12 | 1,077 | 1,195 | 0,996 | 0,96 | 0,88 | -0,12 | 2,2E-01 |
| Q9QUM9 | Proteasome subunit alpha type-6 OS=Mus musculus OX=10090 ( | 33 | 8 | 9 | 8 | 246 | 27,4 | 6,76 | 2151 | 13,8 | 4 | 4 | 1,021 | 0,947 | 1,182 | 1,036 | 0,827 | 1,108 | 0,923 | 0,908 | 1,11 | 0,11 | 2,2E-01 |
| Q91YT0 | NADH dehydrogenase [ubiquinone] flavoprotein 1, mitochondri | 32 | 14 | 31 | 14 | 464 | 50,8 | 8,21 | 8853 | 40,79 | 8 | 6 | 0,946 | 0,909 | 0,934 | 0,899 | 1,276 | 1,001 | 0,89 | 0,979 | 0,89 | -0,11 | 2,2E-01 |
| Q99N28 | Cell adhesion molecule 3 OS=Mus musculus OX=10090 GN=Cadr | 32 | 9 | 22 | 9 | 396 | 42,9 | 5,8 | 9048 | 24,1 | 6 | 4 | 0,98 | 1,024 | 1,112 | 0,986 | 0,88 | 1,028 | 0,993 | 0,962 | 1,06 | 0,06 | 2,2E-01 |
| P35980 | 60S ribosomal protein L18 OS=Mus musculus OX=10090 GN=Rpl | 37 | 6 | 11 | 6 | 188 | 21,6 | 11,78 | 6101 | 2,22 | 5 | 1 | 0,936 | 0,895 | 0,913 | 0,869 | 1,336 | 0,879 | 0,863 | 1,166 | 0,85 | -0,15 | 2,2E-01 |
| Q99JY8 | Phospholipid phosphatase 3 OS=Mus musculus OX=10090 GN=P | 2 | 1 | 2 | 1 | 312 | 35,2 | 9,07 | 1215 | | 1 | | 1,118 | 0,677 | 1,021 | 0,812 | 0,425 | 0,859 | 0,752 | 0,827 | 1,27 | 0,27 | 2,2E-01 |
| Q6P8X1 | Sorting nexin-6 OS=Mus musculus OX=10090 GN=Snx6 PE=1 SV= | 15 | 6 | 7 | 6 | 406 | 46,6 | 6,16 | 3567 | 0 | 5 | 1 | 0,942 | 1,067 | 0,947 | 1,011 | 0,857 | 0,948 | 0,995 | 0,943 | 1,06 | 0,06 | 2,2E-01 |
| Q9WUQ2 | Prolactin regulatory element-binding protein OS=Mus musculus | 3 | 1 | 2 | 1 | 417 | 45,4 | 8,76 | 1218 | | 1 | | 1,121 | 1,426 | 0,975 | 1,17 | 0,906 | 0,94 | 1,113 | 1,133 | 1,15 | 0,15 | 2,2E-01 |
| P60710 | Actin, cytoplasmic 1 OS=Mus musculus OX=10090 GN=Actb PE=1 | 80 | 39 | 490 | 18 | 375 | 41,7 | 5,48 | 200729 | 422,73 | 30 | 23 | 0,983 | 0,984 | 0,967 | 1,037 | 1,08 | 1,034 | 0,962 | 1,056 | 0,96 | -0,04 | 2,2E-01 |
| P28738 | Kinesin heavy chain isoform 5C OS=Mus musculus OX=10090 GN | 40 | 37 | 61 | 27 | 956 | 109,2 | 6,19 | 24205 | 63,12 | 26 | 14 | 1,045 | 1,007 | 1,019 | 0,989 | 0,797 | 0,986 | 1,039 | 0,949 | 1,08 | 0,08 | 2,2E-01 |
| Q3UH60 | Disco-interacting protein 2 homolog B OS=Mus musculus OX=10 | 5 | 7 | 8 | 7 | 1574 | 171 | 8,09 | 4120 | 2,32 | 6 | 1 | 1,074 | 1,079 | 0,958 | 1,061 | 0,952 | 1,076 | 0,916 | 0,986 | 1,06 | 0,06 | 2,2E-01 |
| Q9ESK9 | RB1-inducible coiled-coil protein 1 OS=Mus musculus OX=10090 | 2 | 3 | 3 | 3 | 1588 | 182,2 | 5,49 | 913 | 2,78 | 2 | 1 | 0,953 | 0,95 | 0,964 | 1,182 | 0,801 | 0,99 | 1,013 | 0,811 | 1,12 | 0,12 | 2,2E-01 |
| P70699 | Lysosomal alpha-glucosidase OS=Mus musculus OX=10090 GN=( | 3 | 2 | 2 | 2 | 953 | 106,2 | 5,83 | 1334 | | 2 | | 0,954 | 1,128 | 1,036 | 1,097 | 1,165 | 1,049 | 1,085 | 1,191 | 0,94 | -0,06 | 2,2E-01 |
| O08547 | Vesicle-trafficking protein SEC22b OS=Mus musculus OX=10090 | 32 | 6 | 7 | 6 | 215 | 24,7 | 8,51 | 2899 | 4,71 | 5 | 1 | 1,008 | 0,983 | 0,928 | 1,032 | 0,805 | 0,858 | 1,083 | 0,844 | 1,10 | 0,10 | 2,2E-01 |
| Q8R3T5 | Syntaxin-binding protein 6 OS=Mus musculus OX=10090 GN=Stx | 4 | 1 | 1 | 1 | 210 | 23,7 | 9,19 | 634 | | 1 | | 1,003 | 1,147 | 0,783 | 1,194 | 1,15 | 1,31 | 1,183 | 1,058 | 0,88 | -0,12 | 2,2E-01 |

| | | | | | | | | | | | | | | | | | | | | | | | |
|---|---|---|---|---|---|---|---|---|---|---|---|---|---|---|---|---|---|---|---|---|---|---|---|
| Q3ULJ0 | Glycerol-3-phosphate dehydrogenase 1-like protein OS=Mus mu | 20 | 8 | 13 | 8 | 351 | 38,2 | 6,77 | 1868 | 31,02 | 2 | 6 | 1,04 | 1,02 | 1,035 | 1 | 0,63 | 1,101 | 0,924 | 0,911 | 1,15 | 0,15 | 2,2E-01 |
| Q921H8 | 3-ketoacyl-CoA thiolase A, peroxisomal OS=Mus musculus OX=1 | 27 | 8 | 21 | 3 | 424 | 43,9 | 8,44 | 8030 | 29,4 | 7 | 2 | 0,903 | 0,915 | 0,876 | 0,917 | 0,987 | 0,936 | 0,861 | 1,068 | 0,94 | -0,06 | 2,2E-01 |
| P62761 | Visinin-like protein 1 OS=Mus musculus OX=10090 GN=Vsnl1 PE | 37 | 8 | 15 | 5 | 191 | 22,1 | 5,15 | 6006 | 14,44 | 5 | 4 | 0,939 | 0,888 | 0,928 | 0,886 | 0,908 | 0,961 | 0,969 | 0,915 | 0,97 | -0,03 | 2,2E-01 |
| P52019 | Squalene monooxygenase OS=Mus musculus OX=10090 GN=Sql | 3 | 1 | 1 | 1 | 572 | 63,7 | 8,47 | 533 | | 1 | | 1,186 | 1,362 | 1,278 | 1,153 | 1,953 | 1,495 | 1,385 | 1,12 | 0,84 | -0,16 | 2,2E-01 |
| P42125 | Enoyl-CoA delta isomerase 1, mitochondrial OS=Mus musculus ( | 12 | 3 | 8 | 3 | 289 | 32,2 | 8,98 | 4924 | | 3 | | 0,889 | 0,913 | 0,798 | 0,955 | 1,461 | 0,877 | 0,944 | 1,01 | 0,83 | -0,17 | 2,3E-01 |
| Q8VEH5 | EPM2A-interacting protein 1 OS=Mus musculus OX=10090 GN=E | 4 | 2 | 2 | 2 | 606 | 70,1 | 5,87 | 1451 | | 2 | | 1,308 | 0,996 | 1,188 | 1,134 | 0,823 | 1,136 | 1,015 | 1,126 | 1,13 | 0,13 | 2,3E-01 |
| P62835 | Ras-related protein Rap-1A OS=Mus musculus OX=10090 GN=Ra | 37 | 6 | 12 | 2 | 184 | 21 | 6,67 | 5912 | 8,31 | 4 | 2 | 0,998 | 0,955 | 1,038 | 0,918 | 1,052 | 1,173 | 0,946 | 1,028 | 0,93 | -0,07 | 2,3E-01 |
| Q8BSQ9 | Protein polybromo-1 OS=Mus musculus OX=10090 GN=Pbrm1 P | 0 | 1 | 5 | 1 | 1634 | 187,1 | 6,9 | 1788 | | 1 | | 0,857 | 0,887 | 0,634 | 0,842 | 0,988 | 0,714 | 1,044 | 0,981 | 0,86 | -0,14 | 2,3E-01 |
| P97352 | Protein S100-A13 OS=Mus musculus OX=10090 GN=S100a13 PE | 33 | 3 | 3 | 3 | 98 | 11,2 | 6,13 | 1413 | 3,47 | 2 | 1 | 0,92 | 1,058 | 0,85 | 1,018 | 0,656 | 0,876 | 0,935 | 0,938 | 1,13 | 0,13 | 2,3E-01 |
| Q63844 | Mitogen-activated protein kinase 3 OS=Mus musculus OX=1009( | 29 | 12 | 24 | 8 | 380 | 43 | 6,61 | 11719 | 9,35 | 9 | 4 | 0,975 | 0,877 | 1,039 | 0,98 | 0,795 | 0,999 | 0,899 | 0,889 | 1,08 | 0,08 | 2,3E-01 |
| Q99MX0 | Transketolase-like protein 1 OS=Mus musculus OX=10090 GN=T | 2 | 2 | 2 | 1 | 595 | 65,2 | 6,02 | 1390 | | 2 | | 1,02 | 1,069 | 0,932 | 1,178 | 0,287 | 0,85 | 1,022 | 1,046 | 1,31 | 0,31 | 2,3E-01 |
| P12023 | Amyloid-beta A4 protein OS=Mus musculus OX=10090 GN=App | 8 | 5 | 9 | 5 | 770 | 86,7 | 4,79 | 4316 | 2,8 | 4 | 2 | 0,923 | 0,86 | 0,951 | 0,929 | 1,42 | 0,885 | 1,062 | 0,949 | 0,85 | -0,15 | 2,3E-01 |
| Q9Z0U1 | Tight junction protein ZO-2 OS=Mus musculus OX=10090 GN=Tj | 3 | 4 | 4 | 4 | 1167 | 131,2 | 6,79 | 2197 | | 4 | | 0,959 | 0,929 | 0,961 | 1,077 | 0,991 | 0,922 | 0,887 | 0,913 | 1,06 | 0,06 | 2,3E-01 |
| Q8BMF3 | NADP-dependent malic enzyme, mitochondrial OS=Mus muscul | 3 | 2 | 3 | 1 | 604 | 67,1 | 7,83 | 1363 | 2,43 | 1 | 1 | 0,768 | 0,751 | 1,081 | 0,715 | 0,67 | 0,719 | 0,761 | 0,696 | 1,16 | 0,16 | 2,3E-01 |
| P21107-2 | Isoform 2 of Tropomyosin alpha-3 chain OS=Mus musculus OX= | 60 | 26 | 46 | 10 | 248 | 29 | 4,78 | 16698 | 54,02 | 15 | 13 | 1,069 | 1,163 | 0,851 | 1,135 | 0,741 | 1,015 | 0,935 | 1,012 | 1,14 | 0,14 | 2,3E-01 |
| P56812 | Programmed cell death protein 5 OS=Mus musculus OX=10090 ( | 14 | 2 | 5 | 2 | 126 | 14,3 | 5,68 | | 10,63 | | 2 | 1,021 | 0,995 | 0,995 | 1,208 | 0,991 | 0,985 | 0,926 | 1,02 | 1,08 | 0,08 | 2,3E-01 |
| Q5DQR4 | Syntaxin-binding protein 5-like OS=Mus musculus OX=10090 GN | 4 | 3 | 4 | 3 | 1185 | 131,8 | 6,86 | 1930 | 2,57 | 3 | 1 | 0,843 | 0,86 | 0,954 | 0,871 | 0,88 | 1,019 | 0,881 | 0,98 | 0,94 | -0,06 | 2,3E-01 |
| Q3THW5 | Histone H2A.V OS=Mus musculus OX=10090 GN=H2afv PE=1 SV | 26 | 3 | 8 | 2 | 128 | 13,5 | 10,58 | 2349 | 13,22 | 2 | 1 | 0,925 | 0,939 | 1,428 | 1,659 | 0,938 | 0,782 | 1,234 | 0,882 | 1,29 | 0,29 | 2,3E-01 |
| Q6ZQK5 | Arf-GAP with coiled-coil, ANK repeat and PH domain-containing | 4 | 2 | 2 | 2 | 770 | 87,2 | 6,68 | 1115 | | 2 | | 0,798 | 1,044 | 0,821 | 1,078 | 1,15 | 0,932 | 1,149 | 1,001 | 0,88 | -0,12 | 2,3E-01 |
| Q9EQ80 | NIF3-like protein 1 OS=Mus musculus OX=10090 GN=Nif3l1 PE= | 7 | 2 | 4 | 2 | 376 | 41,7 | 6,76 | 2116 | | 2 | | 0,942 | 1,114 | 0,949 | 0,988 | 1,137 | 0,992 | 1,104 | 1,037 | 0,94 | -0,06 | 2,3E-01 |
| O35206 | Collagen alpha-1(XV) chain OS=Mus musculus OX=10090 GN=Co | 1 | 1 | 1 | 1 | 1367 | 140,4 | 4,89 | 336 | | 1 | | 0,93 | 0,921 | 0,853 | 1,095 | 1,667 | 0,828 | 1,091 | 1,194 | 0,79 | -0,21 | 2,3E-01 |
| Q91WG5 | 5'-AMP-activated protein kinase subunit gamma-2 OS=Mus mus | 1 | 1 | 1 | 1 | 566 | 62,9 | 9,36 | | 1,93 | | 1 | 0,865 | 0,815 | 1,172 | 0,965 | 0,891 | 0,897 | 0,812 | 0,755 | 1,14 | 0,14 | 2,3E-01 |
| P12032 | Metalloproteinase inhibitor 1 OS=Mus musculus OX=10090 GN= | 4 | 1 | 3 | 1 | 205 | 22,6 | 8,82 | 1189 | | 1 | | 0,965 | 0,932 | 1,434 | 0,874 | 0,709 | 0,955 | 0,954 | 0,825 | 1,22 | 0,22 | 2,3E-01 |
| Q91YQ3 | Cold shock domain-containing protein C2 OS=Mus musculus OX | 12 | 3 | 5 | 3 | 154 | 16,8 | 7,55 | 604 | 10,43 | 1 | 2 | 0,967 | 0,902 | 0,974 | 0,921 | 0,569 | 0,906 | 0,94 | 0,879 | 1,14 | 0,14 | 2,3E-01 |
| Q64012 | RNA-binding protein Raly OS=Mus musculus OX=10090 GN=Raly | 13 | 3 | 3 | 3 | 312 | 33,2 | 8,84 | | 7,15 | | 3 | 0,916 | 0,704 | 1,247 | 1,048 | 1,649 | 1,012 | 1,23 | 1,027 | 0,80 | -0,20 | 2,3E-01 |
| Q8QZT1 | Acetyl-CoA acetyltransferase, mitochondrial OS=Mus musculus ( | 48 | 19 | 45 | 19 | 424 | 44,8 | 8,51 | 18558 | 47,5 | 11 | 11 | 1,051 | 0,94 | 0,963 | 0,967 | 0,76 | 1,035 | 0,905 | 0,893 | 1,09 | 0,09 | 2,3E-01 |
| Q8BNE1 | TRPM8 channel-associated factor 1 OS=Mus musculus OX=1009 | 1 | 1 | 1 | 1 | 924 | 102,7 | 6,96 | | 2,32 | | 1 | 0,938 | 0,873 | 1,107 | 1,077 | 2,066 | 1,103 | 1,048 | 1,12 | 0,75 | -0,25 | 2,3E-01 |
| Q91XV3 | Brain acid soluble protein 1 OS=Mus musculus OX=10090 GN=B | 96 | 32 | 190 | 32 | 226 | 22,1 | 4,51 | 58403 | 319,62 | 12 | 23 | 1,029 | 0,927 | 1,153 | 0,958 | 0,661 | 1,209 | 0,736 | 0,737 | 1,22 | 0,22 | 2,3E-01 |
| Q9R0X4 | Acyl-coenzyme A thioesterase 9, mitochondrial OS=Mus muscul | 11 | 5 | 6 | 5 | 439 | 50,5 | 8,59 | 2285 | 5,56 | 3 | 2 | 0,987 | 0,845 | 0,874 | 0,832 | 1,04 | 0,892 | 0,899 | 0,972 | 0,93 | -0,07 | 2,3E-01 |
| Q9R0Q6 | Actin-related protein 2/3 complex subunit 1A OS=Mus musculus | 27 | 9 | 23 | 9 | 370 | 41,6 | 8,18 | 5456 | 38,86 | 5 | 5 | 0,968 | 0,927 | 1,013 | 0,932 | 0,716 | 1,01 | 0,856 | 0,913 | 1,10 | 0,10 | 2,3E-01 |
| Q80TT8 | Cullin-9 OS=Mus musculus OX=10090 GN=Cul9 PE=1 SV=2 | 1 | 1 | 1 | 1 | 1865 | 209 | 5,63 | 457 | | 1 | | 0,999 | 1 | 0,779 | 0,897 | 0,982 | 1,05 | 1,091 | 0,906 | 0,91 | -0,09 | 2,3E-01 |
| Q99KP3 | Lambda-crystallin homolog OS=Mus musculus OX=10090 GN=Cr | 3 | 1 | 1 | 1 | 319 | 35,2 | 5,86 | 257 | | 1 | | 0,825 | 1,278 | 0,693 | 0,899 | 1,607 | 0,673 | 1,325 | 1,345 | 0,75 | -0,25 | 2,3E-01 |
| P50518 | V-type proton ATPase subunit E 1 OS=Mus musculus OX=10090 | 38 | 9 | 19 | 9 | 226 | 26,1 | 8,43 | 7288 | 17,02 | 4 | 5 | 0,935 | 0,909 | 0,834 | 0,869 | 0,975 | 0,909 | 0,869 | 0,979 | 0,95 | -0,05 | 2,3E-01 |
| Q9QZC2 | Plexin-C1 OS=Mus musculus OX=10090 GN=Plxnc1 PE=1 SV=1 | 2 | 3 | 4 | 3 | 1574 | 176,4 | 7,75 | 1633 | 2,28 | 2 | 1 | 0,786 | 0,729 | 0,782 | 0,889 | 0,907 | 0,883 | 0,759 | 0,891 | 0,93 | -0,07 | 2,3E-01 |
| Q9CYZ2 | Tumor protein D54 OS=Mus musculus OX=10090 GN=Tpd52l2 P | 14 | 2 | 2 | 2 | 220 | 24 | 6,15 | 746 | | 2 | | 0,718 | 0,746 | 0,568 | 0,603 | 1,177 | 0,563 | 1,008 | 0,679 | 0,77 | -0,23 | 2,3E-01 |
| Q9EQ20 | Methylmalonate-semialdehyde dehydrogenase [acylating], mito | 21 | 9 | 13 | 9 | 535 | 57,9 | 8,07 | 6559 | 6,15 | 8 | 2 | 1,008 | 1,092 | 1,043 | 1,127 | 1,003 | 0,962 | 0,97 | 1,108 | 1,06 | 0,06 | 2,3E-01 |
| Q9JM76 | Actin-related protein 2/3 complex subunit 3 OS=Mus musculus ( | 14 | 3 | 7 | 3 | 178 | 20,5 | 8,59 | 1572 | 12,14 | 1 | 2 | 1,082 | 1,067 | 1,026 | 0,997 | 0,656 | 1,063 | 1,036 | 0,912 | 1,14 | 0,14 | 2,3E-01 |
| P35438 | Glutamate receptor ionotropic, NMDA 1 OS=Mus musculus OX= | 4 | 5 | 7 | 5 | 938 | 105,4 | 8,84 | 2810 | 2,32 | 4 | 1 | 0,922 | 0,987 | 0,838 | 0,848 | 1,047 | 0,853 | 0,941 | 1,104 | 0,91 | -0,09 | 2,3E-01 |
| Q8VD37 | SH3-containing GRB2-like protein 3-interacting protein 1 OS=Mu | 15 | 10 | 12 | 10 | 806 | 86 | 7,87 | 4902 | 11,5 | 7 | 4 | 0,974 | 0,95 | 0,952 | 0,932 | 1,066 | 0,917 | 0,989 | 1,005 | 0,96 | -0,04 | 2,3E-01 |
| P14211 | Calreticulin OS=Mus musculus OX=10090 GN=Calr PE=1 SV=1 | 42 | 20 | 32 | 20 | 416 | 48 | 4,49 | 10642 | 37,87 | 11 | 10 | 0,956 | 0,905 | 0,97 | 1,14 | 0,713 | 1,036 | 0,907 | 0,87 | 1,13 | 0,13 | 2,3E-01 |
| Q9CQW1 | Synaptobrevin homolog YKT6 OS=Mus musculus OX=10090 GN= | 37 | 7 | 9 | 7 | 198 | 22,3 | 6,35 | 3597 | 8,16 | 4 | 3 | 1,027 | 0,914 | 1,005 | 0,945 | 0,651 | 0,958 | 0,929 | 0,941 | 1,12 | 0,12 | 2,3E-01 |
| Q91YW3 | DnaJ homolog subfamily C member 3 OS=Mus musculus OX=10( | 2 | 1 | 1 | 1 | 504 | 57,4 | 5,85 | 611 | | 1 | | 1,045 | 0,804 | 1,036 | 1,15 | 0,714 | 0,939 | 0,998 | 0,879 | 1,14 | 0,14 | 2,3E-01 |
| Q61361 | Brevican core protein OS=Mus musculus OX=10090 GN=Bcan PE | 7 | 3 | 3 | 3 | 883 | 95,8 | 4,93 | 1056 | | 3 | | 0,882 | 0,665 | 0,714 | 0,846 | 0,697 | 0,647 | 0,79 | 0,644 | 1,12 | 0,12 | 2,3E-01 |
| Q8BGG7 | Ubiquitin-associated and SH3 domain-containing protein B OS=I | 2 | 1 | 1 | 1 | 638 | 71,4 | 6,81 | 251 | | 1 | | 1,366 | 1,223 | 1,223 | 1,685 | 1,673 | 1,547 | 1,443 | 1,472 | 0,90 | -0,10 | 2,3E-01 |
| Q3UTJ2 | Sorbin and SH3 domain-containing protein 2 OS=Mus musculus | 2 | 2 | 2 | 2 | 1180 | 132,3 | 8,53 | 1382 | | 2 | | 0,843 | 0,691 | 0,771 | 1,011 | 0,756 | 1,115 | 0,949 | 1,044 | 0,86 | -0,14 | 2,3E-01 |
| Q00493 | Carboxypeptidase E OS=Mus musculus OX=10090 GN=Cpe PE=1 | 29 | 12 | 18 | 12 | 476 | 53,2 | 5,19 | 8944 | 10,51 | 10 | 4 | 1,015 | 0,926 | 0,956 | 0,998 | 0,925 | 0,976 | 0,915 | 0,95 | 1,03 | 0,03 | 2,3E-01 |
| Q922B2 | Aspartate--tRNA ligase, cytoplasmic OS=Mus musculus OX=1009 | 25 | 11 | 16 | 11 | 501 | 57,1 | 6,49 | 7457 | 8,76 | 10 | 1 | 1,05 | 1,067 | 1,042 | 1,149 | 1,183 | 1,041 | 1,107 | 1,253 | 0,94 | -0,06 | 2,3E-01 |
| Q9WUM3 | Coronin-1B OS=Mus musculus OX=10090 GN=Coro1b PE=1 SV=1 | 11 | 6 | 6 | 5 | 484 | 53,9 | 5,78 | 2277 | 5,28 | 4 | 2 | 0,834 | 0,961 | 1,004 | 0,911 | 1,063 | 0,882 | 1,008 | 1,048 | 0,93 | -0,07 | 2,3E-01 |
| Q8K2Z2 | Pre-mRNA-processing factor 39 OS=Mus musculus OX=10090 GI | 1 | 1 | 1 | 1 | 665 | 77,9 | 5,53 | 409 | | 1 | | 0,984 | 1,341 | 0,59 | 1,381 | 0,696 | 0,777 | 0,861 | 0,946 | 1,31 | 0,31 | 2,3E-01 |
| Q01405 | Protein transport protein Sec23A OS=Mus musculus OX=10090 ( | 7 | 4 | 4 | 4 | 765 | 86,1 | 7,08 | 2565 | | 4 | | 0,96 | 1,204 | 1,109 | 0,942 | 1,137 | 1,15 | 1,063 | 1,258 | 0,91 | -0,09 | 2,3E-01 |
| Q8BWA8 | Rho guanine nucleotide exchange factor 19 OS=Mus musculus C | 1 | 1 | 2 | 1 | 802 | 88,8 | 8,34 | | 4,56 | | 1 | 3,232 | 7,135 | 2,635 | 1,153 | 1,29 | 0,991 | 3,363 | 1,159 | 2,08 | 1,08 | 2,3E-01 |
| P12961 | Neuroendocrine protein 7B2 OS=Mus musculus OX=10090 GN=S | 19 | 4 | 6 | 4 | 212 | 23,9 | 5,81 | 1610 | 8,12 | 3 | 2 | 0,9 | 0,97 | 0,912 | 0,985 | 1,421 | 1,027 | 1,017 | 0,907 | 0,86 | -0,14 | 2,4E-01 |
| Q6R891 | Neurabin-2 OS=Mus musculus OX=10090 GN=Ppp1r9b PE=1 SV= | 13 | 8 | 9 | 8 | 817 | 89,5 | 4,92 | 3390 | 5,67 | 6 | 2 | 1,015 | 1,059 | 0,998 | 1,003 | 1,042 | 1,304 | 1,081 | 1,008 | 0,92 | -0,08 | 2,4E-01 |
| Q99LB7 | Sarcosine dehydrogenase, mitochondrial OS=Mus musculus OX= | 3 | 3 | 3 | 3 | 919 | 101,6 | 6,74 | 1704 | | 3 | | 1,217 | 1,216 | 1,168 | 1,367 | 1,547 | 1,374 | 1,425 | 1,133 | 0,91 | -0,09 | 2,4E-01 |
| Q9QYX7 | Protein piccolo OS=Mus musculus OX=10090 GN=Pclo PE=1 SV= | 9 | 38 | 60 | 37 | 5068 | 550,5 | 6,51 | 25532 | 39,63 | 25 | 15 | 1,098 | 0,988 | 1,07 | 0,994 | 0,95 | 1,006 | 0,993 | 1,031 | 1,04 | 0,04 | 2,4E-01 |
| Q6A065 | Centrosomal protein of 170 kDa OS=Mus musculus OX=10090 G | 10 | 17 | 21 | 16 | 1588 | 174,9 | 7,17 | 8505 | 9,8 | 11 | 6 | 0,906 | 0,758 | 1,385 | 0,988 | 0,814 | 0,822 | 0,848 | 0,847 | 1,21 | 0,21 | 2,4E-01 |
| Q3TBN1 | Transmembrane protein 248 OS=Mus musculus OX=10090 GN= | 3 | 1 | 1 | 1 | 314 | 35 | 6,37 | 306 | | 1 | | 0,993 | 1,158 | 0,905 | 0,947 | 1,649 | 0,888 | 1,417 | 1,018 | 0,81 | -0,19 | 2,4E-01 |
| Q9WTP6 | Adenylate kinase 2, mitochondrial OS=Mus musculus OX=10090 | 18 | 3 | 4 | 3 | 239 | 26,5 | 7,39 | 1956 | | 3 | | 0,915 | 0,997 | 0,943 | 1,043 | 1,495 | 1,08 | 0,845 | 1,204 | 0,84 | -0,16 | 2,4E-01 |
| Q05920 | Pyruvate carboxylase, mitochondrial OS=Mus musculus OX=100 | 30 | 30 | 48 | 30 | 1178 | 129,6 | 6,71 | 18111 | 38,78 | 20 | 11 | 1,024 | 0,914 | 1,075 | 1,076 | 1,226 | 0,961 | 1,11 | 1,145 | 0,92 | -0,08 | 2,4E-01 |
| Q3U0M1 | Trafficking protein particle complex subunit 9 OS=Mus musculus | 3 | 4 | 4 | 4 | 1148 | 128,2 | 6,47 | 1253 | 4,37 | 2 | 2 | 1,08 | 1,001 | 0,998 | 0,966 | 0,971 | 1,031 | 0,944 | 0,921 | 1,05 | 0,05 | 2,4E-01 |
| Q9EST4 | Proteasome assembly chaperone 2 OS=Mus musculus OX=1009( | 3 | 1 | 2 | 1 | 264 | 29,5 | 6,05 | 1123 | | 1 | | 1,056 | 1,237 | 1,286 | 1,042 | 0,683 | 1,131 | 1,102 | 1,065 | 1,16 | 0,16 | 2,4E-01 |
| P20060 | Beta-hexosaminidase subunit beta OS=Mus musculus OX=1009( | 9 | 4 | 4 | 4 | 536 | 61,1 | 8,12 | 1473 | 2,22 | 3 | 1 | 1,12 | 1,188 | 1,166 | 1,212 | 1,598 | 1,24 | 1,239 | 1,144 | 0,90 | -0,10 | 2,4E-01 |
| Q60875 | Rho guanine nucleotide exchange factor 2 OS=Mus musculus O) | 6 | 5 | 5 | 5 | 985 | 111,9 | 7,25 | 2853 | | 5 | | 1,219 | 1,105 | 1,063 | 1 | 1,275 | 1,301 | 1,018 | 1,206 | 0,91 | -0,09 | 2,4E-01 |
| Q9EQQ9 | Protein O-GlcNAcase OS=Mus musculus OX=10090 GN=Oga PE= | 7 | 7 | 9 | 7 | 916 | 103,1 | 4,92 | 4129 | 5,06 | 5 | 2 | 0,979 | 0,989 | 1,013 | 1,042 | 0,798 | 0,977 | 0,935 | 1,037 | 1,07 | 0,07 | 2,4E-01 |
| P62881 | Guanine nucleotide-binding protein subunit beta-5 OS=Mus mu: | 19 | 5 | 8 | 5 | 395 | 43,5 | 6,46 | 2871 | 12,94 | 3 | 2 | 1,064 | 1,175 | 1,104 | 1,034 | 1,179 | 1,04 | 1,183 | 1,275 | 0,94 | -0,06 | 2,4E-01 |

| | | | | | | | | | | | | | | | | | | | | | | | |
|---|---|---|---|---|---|---|---|---|---|---|---|---|---|---|---|---|---|---|---|---|---|---|---|
| O70439 | Syntaxin-7 OS=Mus musculus OX=10090 GN=Stx7 PE=1 SV=3 | 13 | 3 | 5 | 3 | 261 | 29,8 | 5,78 | 2264 | 2,75 | 2 | 1 | 1,041 | 1,032 | 1,208 | 1,151 | 0,922 | 1,101 | 1,073 | 1,031 | 1,07 | 0,07 | 2,4E-01 |
| Q9CQC6 | Basic leucine zipper and W2 domain-containing protein 1 OS=M | 13 | 6 | 8 | 6 | 419 | 48 | 5,92 | 1395 | 14,63 | 2 | 4 | 1,083 | 1,044 | 1,088 | 1,083 | 0,905 | 0,992 | 1,085 | 1,084 | 1,06 | 0,06 | 2,4E-01 |
| Q8K1A6 | Coiled-coil and C2 domain-containing protein 1A OS=Mus muscu | 3 | 3 | 4 | 3 | 943 | 103,6 | 7,84 | 1297 | 5,25 | 2 | 1 | 1,018 | 0,917 | 0,959 | 0,94 | 0,785 | 0,806 | 1,039 | 0,882 | 1,09 | 0,09 | 2,4E-01 |
| Q9D8B3 | Charged multivesicular body protein 4b OS=Mus musculus OX=1 | 27 | 6 | 11 | 6 | 224 | 24,9 | 4,82 | 2422 | 14,81 | 2 | 4 | 0,889 | 0,915 | 0,898 | 0,905 | 1,001 | 0,802 | 1,076 | 1,514 | 0,82 | -0,18 | 2,4E-01 |
| Q6PAR5 | GTPase-activating protein and VPS9 domain-containing protein | 7 | 7 | 9 | 7 | 1458 | 162,3 | 5,19 | 3691 | 4,71 | 6 | 2 | 0,897 | 0,928 | 1,015 | 0,9 | 1,136 | 0,897 | 1,001 | 1 | 0,93 | -0,07 | 2,4E-01 |
| P97447 | Four and a half LIM domains protein 1 OS=Mus musculus OX=10 | 9 | 2 | 2 | 2 | 280 | 31,9 | 8,37 | 317 | 0 | 1 | 1 | 0,773 | 0,975 | 0,831 | 0,841 | 0,787 | 0,738 | 0,864 | 0,767 | 1,08 | 0,08 | 2,4E-01 |
| Q6ZWX6 | Eukaryotic translation initiation factor 2 subunit 1 OS=Mus musc | 25 | 8 | 9 | 8 | 315 | 36,1 | 5,08 | 1829 | 12,82 | 3 | 5 | 1,023 | 0,994 | 1,092 | 0,992 | 0,798 | 0,975 | 1,027 | 1,003 | 1,08 | 0,08 | 2,4E-01 |
| A2AGL3 | Ryanodine receptor 3 OS=Mus musculus OX=10090 GN=Ryr3 PE | 1 | 2 | 5 | 1 | 4863 | 551 | 5,78 | 1667 | | 2 | | 0,948 | 0,999 | 0,784 | 0,78 | 1,602 | 0,898 | 0,959 | 0,967 | 0,79 | -0,21 | 2,4E-01 |
| Q8VE37 | Regulator of chromosome condensation OS=Mus musculus OX= | 5 | 1 | 2 | 1 | 421 | 44,9 | 8,1 | | 7,09 | | 1 | 1,273 | 0,955 | 1,148 | 1,103 | 2,289 | 1,339 | 1,215 | 1,098 | 0,75 | -0,25 | 2,4E-01 |
| Q8R086 | Sulfite oxidase, mitochondrial OS=Mus musculus OX=10090 GN= | 3 | 2 | 4 | 1 | 546 | 60,7 | 6,54 | 1701 | 2,12 | 1 | 1 | 0,718 | 0,921 | 1,205 | 0,969 | 1,17 | 1,071 | 0,969 | 1,187 | 0,87 | -0,13 | 2,4E-01 |
| Q8C4Q6 | Axin interactor, dorsalization-associated protein OS=Mus muscu | 2 | 1 | 1 | 1 | 305 | 34,9 | 6,74 | | 2,1 | | 1 | 0,849 | 0,766 | 1,009 | 0,774 | 0,751 | 1,055 | 0,986 | 1,115 | 0,87 | -0,13 | 2,4E-01 |
| P47791 | Glutathione reductase, mitochondrial OS=Mus musculus OX=10 | 15 | 5 | 6 | 5 | 500 | 53,6 | 7,99 | 1356 | 5,42 | 3 | 3 | 0,926 | 0,941 | 0,878 | 1,293 | 1,078 | 1,024 | 1,6 | 1,179 | 0,83 | -0,17 | 2,4E-01 |
| O35607 | Bone morphogenetic protein receptor type-2 OS=Mus musculus | 5 | 4 | 7 | 4 | 1038 | 114,9 | 6,23 | 2517 | 2,51 | 3 | 1 | 1,004 | 0,998 | 0,936 | 0,837 | 1,578 | 0,945 | 1,035 | 1,007 | 0,83 | -0,17 | 2,4E-01 |
| Q62421 | Endophilin-A3 OS=Mus musculus OX=10090 GN=Sh3gl3 PE=1 SV | 2 | 1 | 1 | 1 | 347 | 38,9 | 5,12 | | 2,46 | | 1 | 1,187 | 0,968 | 1,289 | 1,115 | 0,639 | 0,979 | 1,02 | 1,21 | 1,18 | 0,18 | 2,4E-01 |
| Q6KAU4 | Multivesicular body subunit 12B OS=Mus musculus OX=10090 G | 4 | 1 | 1 | 1 | 317 | 35,4 | 7,93 | 467 | | 1 | | 1,065 | 1,104 | 0,974 | 0,993 | 0,996 | 1,437 | 1,089 | 1,135 | 0,89 | -0,11 | 2,4E-01 |
| P62746 | Rho-related GTP-binding protein RhoB OS=Mus musculus OX=10 | 33 | 7 | 10 | 7 | 196 | 22,1 | 5,24 | 4305 | 12,21 | 4 | 3 | 0,978 | 0,994 | 1,125 | 0,993 | 0,574 | 1,046 | 1,012 | 0,871 | 1,17 | 0,17 | 2,4E-01 |
| O54781 | SRSF protein kinase 2 OS=Mus musculus OX=10090 GN=Srpk2 P | 9 | 5 | 6 | 5 | 681 | 76,7 | 4,91 | 1480 | 6,65 | 2 | 3 | 0,96 | 1,08 | 0,946 | 1,077 | 0,864 | 1,018 | 0,938 | 0,986 | 1,07 | 0,07 | 2,4E-01 |
| Q9WUB4 | Dynactin subunit 6 OS=Mus musculus OX=10090 GN=Dctn6 PE= | 7 | 2 | 3 | 2 | 190 | 20,7 | 6,67 | 1918 | | 2 | | 0,986 | 0,968 | 0,974 | 1,124 | 0,864 | 1,077 | 0,895 | 0,899 | 1,08 | 0,08 | 2,4E-01 |
| Q8BHN3 | Neutral alpha-glucosidase AB OS=Mus musculus OX=10090 GN= | 20 | 19 | 23 | 19 | 944 | 106,8 | 6,06 | 9478 | 9,15 | 14 | 5 | 0,993 | 1,077 | 1,055 | 1,092 | 1,258 | 1,034 | 1,061 | 1,15 | 0,94 | -0,06 | 2,4E-01 |
| Q8R3R8 | Gamma-aminobutyric acid receptor-associated protein-like 1 OS | 16 | 3 | 3 | 1 | 117 | 14 | 8,73 | 540 | 8,83 | 1 | 2 | 1,052 | 1,093 | 1,267 | 1,174 | 0,795 | 1,169 | 1,033 | 1,1 | 1,12 | 0,12 | 2,4E-01 |
| Q9JK48 | Endophilin-B1 OS=Mus musculus OX=10090 GN=Sh3glb1 PE=1 S | 13 | 6 | 7 | 6 | 365 | 40,8 | 6,04 | 1832 | 12,55 | 3 | 3 | 0,969 | 0,922 | 0,966 | 0,988 | 0,788 | 0,9 | 0,957 | 0,972 | 1,06 | 0,06 | 2,4E-01 |
| P54823 | Probable ATP-dependent RNA helicase DDX6 OS=Mus musculus | 17 | 6 | 6 | 6 | 483 | 54,2 | 8,66 | 2442 | 4,77 | 4 | 2 | 0,867 | 0,855 | 0,918 | 0,873 | 0,914 | 0,839 | 0,947 | 1,021 | 0,94 | -0,06 | 2,4E-01 |
| P47857 | ATP-dependent 6-phosphofructokinase, muscle type OS=Mus m | 25 | 16 | 26 | 13 | 780 | 85,2 | 8 | 8405 | 33,83 | 8 | 8 | 1,07 | 1,037 | 1,149 | 1,058 | 0,948 | 1,054 | 1,06 | 1,061 | 1,05 | 0,05 | 2,4E-01 |
| Q6PD03 | Serine/threonine-protein phosphatase 2A 56 kDa regulatory sub | 5 | 2 | 3 | 2 | 486 | 56,3 | 7,03 | 1149 | 2,45 | 1 | 1 | 1,056 | 0,988 | 1,138 | 0,962 | 0,775 | 1,133 | 0,865 | 0,928 | 1,12 | 0,12 | 2,4E-01 |
| O35226-2 | Isoform Rpn10B of 26S proteasome non-ATPase regulatory subu | 26 | 8 | 12 | 1 | 379 | 41 | 4,79 | 6010 | 4,94 | 6 | 2 | 1,053 | 1,018 | 1,222 | 1,1 | 1,109 | 0,84 | 1,101 | 0,931 | 1,10 | 0,10 | 2,4E-01 |
| O35454 | Chloride transport protein 6 OS=Mus musculus OX=10090 GN=C | 5 | 3 | 4 | 3 | 870 | 96,9 | 7,02 | 1636 | | 3 | | 0,516 | 0,775 | 0,567 | 0,664 | 0,858 | 0,506 | 1,022 | 0,767 | 0,80 | -0,20 | 2,4E-01 |
| Q05793 | Basement membrane-specific heparan sulfate proteoglycan core | 1 | 2 | 2 | 2 | 3707 | 398 | 6,32 | 375 | 3,3 | 1 | 1 | 0,798 | 1,562 | 0,621 | 0,713 | 1,071 | 1,446 | 1,44 | 1,007 | 0,74 | -0,26 | 2,4E-01 |
| Q8CCJ4 | APC membrane recruitment protein 2 OS=Mus musculus OX=10 | 4 | 3 | 5 | 3 | 672 | 69,9 | 6,64 | 2286 | 1,76 | 1 | 2 | 0,892 | 0,785 | 0,739 | 0,828 | 0,849 | 0,939 | 0,822 | 0,848 | 0,94 | -0,06 | 2,4E-01 |
| Q61576 | Peptidyl-prolyl cis-trans isomerase FKBP10 OS=Mus musculus O | 3 | 2 | 2 | 2 | 581 | 64,7 | 5,64 | 349 | 2,4 | 1 | 1 | 0,955 | 1,145 | 0,789 | 0,918 | 0,891 | 0,828 | 0,822 | 0,876 | 1,11 | 0,11 | 2,4E-01 |
| Q6PDL0 | Cytoplasmic dynein 1 light intermediate chain 2 OS=Mus muscul | 18 | 6 | 7 | 6 | 492 | 54,2 | 6,28 | 2451 | 7,7 | 4 | 2 | 0,931 | 1,087 | 1,11 | 1,309 | 0,627 | 0,968 | 1,127 | 1,027 | 1,18 | 0,18 | 2,4E-01 |
| O35609 | Secretory carrier-associated membrane protein 3 OS=Mus musc | 11 | 3 | 3 | 3 | 349 | 38,4 | 7,64 | 2119 | | 3 | | 0,929 | 1,085 | 0,899 | 1,011 | 1,079 | 0,933 | 1,061 | 1,258 | 0,91 | -0,09 | 2,4E-01 |
| Q99LX0 | Protein/nucleic acid deglycase DJ-1 OS=Mus musculus OX=1009 | 34 | 5 | 16 | 5 | 189 | 20 | 6,77 | 7842 | 5,28 | 4 | 2 | 0,978 | 0,992 | 1,194 | 0,997 | 0,853 | 1,063 | 0,939 | 0,96 | 1,09 | 0,09 | 2,4E-01 |
| P62827 | GTP-binding nuclear protein Ran OS=Mus musculus OX=10090 G | 30 | 9 | 15 | 9 | 216 | 24,4 | 7,49 | 7913 | 10,57 | 7 | 3 | 1,044 | 0,963 | 1,044 | 1,031 | 0,837 | 1,023 | 1,01 | 0,971 | 1,06 | 0,06 | 2,5E-01 |
| P31648 | Sodium- and chloride-dependent GABA transporter 1 OS=Mus m | 5 | 2 | 6 | 2 | 599 | 67 | 7,96 | 3194 | | 2 | | 0,983 | 0,926 | 0,995 | 0,929 | 0,734 | 0,892 | 1,03 | 0,849 | 1,09 | 0,09 | 2,5E-01 |
| Q9CWK8 | Sorting nexin-2 OS=Mus musculus OX=10090 GN=Snx2 PE=1 SV= | 16 | 7 | 10 | 7 | 519 | 58,4 | 5,12 | 3148 | 9,33 | 5 | 3 | 0,999 | 0,923 | 0,967 | 0,943 | 1,211 | 0,953 | 0,979 | 1,004 | 0,92 | -0,08 | 2,5E-01 |
| Q60829 | Protein phosphatase 1 regulatory subunit 1B OS=Mus musculus | 29 | 3 | 3 | 3 | 194 | 21,8 | 4,65 | 690 | 0 | 2 | 1 | 1,104 | 1,651 | 1,657 | 1,412 | 2,026 | 1,754 | 1,212 | 2,024 | 0,83 | -0,17 | 2,5E-01 |
| Q9CQ85 | Mitochondrial import inner membrane translocase subunit Tim2 | 8 | 1 | 1 | 1 | 194 | 20,1 | 8,22 | 656 | | 1 | | 1,537 | 1,315 | 1,197 | 1,469 | 1,172 | 1,034 | 1,305 | 1,423 | 1,12 | 0,12 | 2,5E-01 |
| Q9CY64 | Biliverdin reductase A OS=Mus musculus OX=10090 GN=Blvra P | 7 | 2 | 3 | 2 | 295 | 33,5 | 7,02 | 831 | 2,29 | 1 | 1 | 0,856 | 1,033 | 1,078 | 0,939 | 1,197 | 1,025 | 1,023 | 1,005 | 0,92 | -0,08 | 2,5E-01 |
| P35235 | Tyrosine-protein phosphatase non-receptor type 11 OS=Mus mu | 5 | 3 | 3 | 3 | 597 | 68,4 | 7,3 | 746 | 5,76 | 1 | 2 | 0,904 | 0,755 | 0,99 | 0,912 | 0,633 | 0,936 | 0,834 | 0,741 | 1,13 | 0,13 | 2,5E-01 |
| Q9R1C0 | Transcription initiation factor TFIID subunit 7 OS=Mus musculus | 2 | 1 | 1 | 1 | 341 | 39,1 | 5,33 | 307 | | 1 | | 0,932 | 0,849 | 1,181 | 0,937 | 0,622 | 0,876 | 0,929 | 0,941 | 1,16 | 0,16 | 2,5E-01 |
| Q60631 | Growth factor receptor-bound protein 2 OS=Mus musculus OX= | 41 | 8 | 10 | 8 | 217 | 25,2 | 6,32 | 3435 | 8,86 | 4 | 5 | 0,933 | 0,861 | 1,346 | 0,883 | 0,827 | 0,905 | 0,875 | 0,82 | 1,17 | 0,17 | 2,5E-01 |
| Q99P88 | Nuclear pore complex protein Nup155 OS=Mus musculus OX=10 | 1 | 1 | 1 | 1 | 1391 | 155 | 6,15 | 499 | | 1 | | 1,253 | 1,321 | 1,092 | 1,184 | 1,693 | 1,242 | 1,09 | 1,605 | 0,86 | -0,14 | 2,5E-01 |
| Q6KCD5 | Nipped-B-like protein OS=Mus musculus OX=10090 GN=Nipbl P | 0 | 1 | 1 | 1 | 2798 | 315,3 | 7,91 | 320 | | 1 | | 1,012 | 1,043 | 1,351 | 0,976 | 0,565 | 1,083 | 0,976 | 1,013 | 1,20 | 0,20 | 2,5E-01 |
| O09111 | NADH dehydrogenase [ubiquinone] 1 beta subcomplex subunit | 7 | 1 | 1 | 1 | 151 | 17,4 | 5,22 | 903 | | 1 | | 1,002 | 0,888 | 1,201 | 0,911 | 0,769 | 0,898 | 0,955 | 0,952 | 1,12 | 0,12 | 2,5E-01 |
| Q6DFV7 | Nuclear receptor coactivator 7 OS=Mus musculus OX=10090 GN | 1 | 1 | 1 | 1 | 943 | 106,3 | 5,43 | 423 | | 1 | | 0,87 | 0,677 | 0,833 | 0,78 | 1,124 | 0,725 | 0,893 | 0,89 | 0,87 | -0,13 | 2,5E-01 |
| Q9QZD9 | Eukaryotic translation initiation factor 3 subunit I OS=Mus musc | 15 | 4 | 5 | 4 | 325 | 36,4 | 5,64 | 2523 | 2,74 | 3 | 1 | 0,951 | 0,925 | 1,015 | 1,06 | 0,937 | 0,946 | 0,824 | 0,999 | 1,07 | 0,07 | 2,5E-01 |
| O35166 | Golgi SNAP receptor complex member 2 OS=Mus musculus OX= | 8 | 2 | 4 | 2 | 212 | 24,7 | 8,21 | 1956 | | 2 | | 0,828 | 0,789 | 0,629 | 0,61 | 4,516 | 1,231 | 0,985 | 0,7 | 0,38 | -0,62 | 2,5E-01 |
| Q9CWS0 | N(G),N(G)-dimethylarginine dimethylaminohydrolase 1 OS=Mus | 50 | 13 | 21 | 12 | 285 | 31,4 | 5,97 | 11194 | 13,29 | 12 | 3 | 1,017 | 0,987 | 0,924 | 1,055 | 0,833 | 1,017 | 0,897 | 0,981 | 1,07 | 0,07 | 2,5E-01 |
| Q8BGZ4 | Cell division cycle protein 23 homolog OS=Mus musculus OX=10 | 1 | 1 | 1 | 1 | 597 | 68,5 | 7,18 | 462 | | 1 | | 0,812 | 0,941 | 0,746 | 0,955 | 1,007 | 0,934 | 1,247 | 0,806 | 0,86 | -0,14 | 2,5E-01 |
| Q6ZWR6 | Nesprin-1 OS=Mus musculus OX=10090 GN=Syne1 PE=1 SV=2 | 0 | 1 | 1 | 1 | 8799 | 1009,3 | 5,59 | 379 | | 1 | | 0,872 | 0,793 | 0,901 | 0,742 | 1,048 | 0,985 | 0,76 | 0,886 | 0,90 | -0,10 | 2,5E-01 |
| Q80TV8 | CLIP-associating protein 1 OS=Mus musculus OX=10090 GN=Cla | 5 | 7 | 7 | 7 | 1535 | 169,1 | 9,03 | 3029 | 1,96 | 6 | 1 | 1,038 | 0,96 | 0,897 | 0,803 | 0,542 | 0,844 | 0,924 | 0,878 | 1,16 | 0,16 | 2,5E-01 |
| Q8BMI3 | ADP-ribosylation factor-binding protein GGA3 OS=Mus musculu | 2 | 1 | 1 | 1 | 718 | 77,9 | 5,66 | 381 | | 1 | | 1,169 | 1,122 | 1,269 | 1,261 | 1,687 | 1,349 | 1,109 | 1,314 | 0,88 | -0,12 | 2,5E-01 |
| Q8K0C9 | GDP-mannose 4,6 dehydratase OS=Mus musculus OX=10090 GN | 9 | 3 | 5 | 3 | 372 | 42 | 7,03 | 2514 | | 3 | | 1,039 | 0,97 | 0,921 | 1,035 | 1,086 | 1,155 | 0,913 | 1,121 | 0,93 | -0,07 | 2,5E-01 |
| Q9CQW2 | ADP-ribosylation factor-like protein 8B OS=Mus musculus OX=10 | 20 | 4 | 4 | 1 | 186 | 21,5 | 8,43 | 740 | 8,02 | 1 | 3 | 1,061 | 1,038 | 0,992 | 1,214 | 0,44 | 0,857 | 1,161 | 1,016 | 1,24 | 0,24 | 2,5E-01 |
| Q8K2T1 | NmrA-like family domain-containing protein 1 OS=Mus musculu | 6 | 2 | 3 | 2 | 309 | 34,4 | 6,86 | 1075 | | 2 | | 0,827 | 0,785 | 1,065 | 0,959 | 0,972 | 0,882 | 1,182 | 1,061 | 0,89 | -0,11 | 2,5E-01 |
| P61358 | 60S ribosomal protein L27 OS=Mus musculus OX=10090 GN=Rpl | 43 | 6 | 10 | 6 | 136 | 15,8 | 10,56 | 3895 | 5,91 | 3 | 3 | 0,923 | 1,22 | 0,931 | 1,041 | 0,966 | 0,911 | 0,923 | 0,956 | 1,10 | 0,10 | 2,5E-01 |
| O54901 | OX-2 membrane glycoprotein OS=Mus musculus OX=10090 GN= | 17 | 6 | 20 | 6 | 278 | 31,2 | 8,79 | 11694 | 7,23 | 4 | 2 | 1,082 | 1,034 | 0,888 | 1,105 | 0,706 | 1,005 | 0,956 | 1,002 | 1,12 | 0,12 | 2,5E-01 |
| Q569Z5 | Probable ATP-dependent RNA helicase DDX46 OS=Mus musculu | 1 | 1 | 1 | 1 | 1032 | 117,4 | 9,26 | 474 | | 1 | | 1,075 | 1,045 | 1,073 | 1,143 | 0,681 | 1,059 | 1,106 | 0,991 | 1,13 | 0,13 | 2,5E-01 |
| Q8BG16 | Sodium-dependent neutral amino acid transporter B(0)AT2 OS= | 3 | 1 | 1 | 1 | 729 | 81,7 | 5,19 | 516 | | 1 | | 1,283 | 1,225 | 1,165 | 1,142 | 1,431 | 1,086 | 1,352 | 1,366 | 0,92 | -0,08 | 2,5E-01 |
| Q91V64 | Isochorismatase domain-containing protein 1 OS=Mus musculus | 6 | 2 | 2 | 2 | 297 | 32 | 7,39 | 472 | 2,41 | 1 | 1 | 0,909 | 1,012 | 1,034 | 0,994 | 0,684 | 1,088 | 0,907 | 0,817 | 1,13 | 0,13 | 2,5E-01 |
| Q8BT60 | Copine-3 OS=Mus musculus OX=10090 GN=Cpne3 PE=1 SV=2 | 6 | 3 | 3 | 2 | 533 | 59,5 | 5,78 | 840 | 2,85 | 1 | 2 | 1,174 | 0,98 | 1,028 | 0,902 | 0,988 | 0,99 | 0,871 | 0,907 | 1,09 | 0,09 | 2,5E-01 |
| Q8K021 | Secretory carrier-associated membrane protein 1 OS=Mus musc | 19 | 5 | 18 | 5 | 338 | 38 | 7,71 | 10277 | 5,39 | 5 | 1 | 0,996 | 1,013 | 1,052 | 1,047 | 1,122 | 1,041 | 1,049 | 1,025 | 0,97 | -0,03 | 2,5E-01 |
| P62245 | 40S ribosomal protein S15a OS=Mus musculus OX=10090 GN=R | 43 | 6 | 8 | 6 | 130 | 14,8 | 10,13 | 1273 | 10,61 | 2 | 5 | 0,996 | 1,051 | 1,147 | 0,982 | 1,08 | 0,94 | 0,935 | 0,964 | 1,07 | 0,07 | 2,5E-01 |
| Q8BGT5 | Alanine aminotransferase 2 OS=Mus musculus OX=10090 GN=G | 4 | 1 | 2 | 1 | 522 | 57,9 | 8 | | 9,83 | | 1 | 1,636 | 1,535 | 1,382 | 1,704 | 2,619 | 1,345 | 1,658 | 2,077 | 0,81 | -0,19 | 2,5E-01 |
| Q11136 | Xaa-Pro dipeptidase OS=Mus musculus OX=10090 GN=Pepd PE= | 7 | 3 | 4 | 3 | 493 | 55 | 5,78 | 1216 | 6,9 | 2 | 2 | 1,059 | 0,994 | 1,044 | 1,093 | 0,831 | 1,144 | 0,942 | 0,923 | 1,09 | 0,09 | 2,5E-01 |

| | | | | | | | | | | | | | | | | | | | | | | | |
|---|---|---|---|---|---|---|---|---|---|---|---|---|---|---|---|---|---|---|---|---|---|---|---|
| Q5U5M8 | Biogenesis of lysosome-related organelles complex 1 subunit 3 ( | 8 | 1 | 1 | 1 | 195 | 20,4 | 4,92 | 448 | | 1 | | 0,62 | 0,691 | 1,199 | 0,604 | 2,222 | 0,969 | 0,745 | 1,013 | 0,63 | -0,37 | 2,5E-01 |
| A2BDX3 | Adenylyltransferase and sulfurtransferase MOCS3 OS=Mus mus | 2 | 1 | 1 | 1 | 460 | 49,3 | 7,49 | 587 | | 1 | | 0,984 | 1,153 | 0,733 | 0,655 | 0,788 | 0,544 | 0,623 | 0,875 | 1,25 | 0,25 | 2,5E-01 |
| P50285 | Dimethylaniline monooxygenase [N-oxide-forming] 1 OS=Mus m | 5 | 2 | 4 | 2 | 532 | 59,9 | 8,47 | 2493 | | 2 | | 1,785 | 1,972 | 1,765 | 2,051 | 2,559 | 1,758 | 2,089 | 2,075 | 0,89 | -0,11 | 2,5E-01 |
| Q9JLZ3 | Methylglutaconyl-CoA hydratase, mitochondrial OS=Mus muscu | 17 | 4 | 9 | 4 | 314 | 33,4 | 9,51 | 2098 | 14,31 | 3 | 2 | 1,005 | 1,025 | 1,069 | 1,07 | 0,86 | 1,096 | 0,963 | 0,991 | 1,07 | 0,07 | 2,5E-01 |
| B1AY13 | Ubiquitin carboxyl-terminal hydrolase 24 OS=Mus musculus OX= | 1 | 2 | 2 | 2 | 2617 | 293,8 | 6,19 | 936 | | 2 | | 1,015 | 1,125 | 1,137 | 1,093 | 1,393 | 1,521 | 1,173 | 0,942 | 0,87 | -0,13 | 2,5E-01 |
| O88543 | COP9 signalosome complex subunit 3 OS=Mus musculus OX=10 | 6 | 2 | 4 | 2 | 423 | 47,8 | 6,65 | 1394 | 5,78 | 1 | 1 | 0,926 | 0,974 | 0,96 | 0,924 | 1,01 | 0,97 | 0,928 | 0,983 | 0,97 | -0,03 | 2,5E-01 |
| P38647 | Stress-70 protein, mitochondrial OS=Mus musculus OX=10090 G | 37 | 24 | 47 | 23 | 679 | 73,4 | 6,07 | 21394 | 37,53 | 18 | 10 | 1,049 | 0,96 | 1,061 | 0,996 | 0,913 | 0,991 | 0,929 | 1,042 | 1,05 | 0,05 | 2,5E-01 |
| P00405 | Cytochrome c oxidase subunit 2 OS=Mus musculus OX=10090 G | 11 | 4 | 6 | 4 | 227 | 26 | 4,73 | 4237 | | 4 | | 0,96 | 0,881 | 0,886 | 0,913 | 0,98 | 1,067 | 0,873 | 0,943 | 0,94 | -0,06 | 2,5E-01 |
| A2AAE1 | Transmembrane protein KIAA1109 OS=Mus musculus OX=1009 | 0 | 1 | 1 | 1 | 5005 | 555 | 6,61 | 158 | | 1 | | 1,299 | 1,69 | 1,326 | 1,43 | 1,474 | 1,279 | 1,215 | 1,239 | 1,10 | 0,10 | 2,5E-01 |
| P62320 | Small nuclear ribonucleoprotein Sm D3 OS=Mus musculus OX=1 | 8 | 1 | 1 | 1 | 126 | 13,9 | 10,32 | | 2,44 | | 1 | 0,843 | 0,954 | 0,9 | 0,943 | 1,509 | 0,801 | 0,984 | 1,112 | 0,83 | -0,17 | 2,5E-01 |
| P61963 | DDB1- and CUL4-associated factor 7 OS=Mus musculus OX=100 | 6 | 2 | 2 | 2 | 342 | 38,9 | 5,52 | 1233 | | 2 | | 1,3 | 1,206 | 1,28 | 1,114 | 1,21 | 1,238 | 1,041 | 1,094 | 1,07 | 0,07 | 2,6E-01 |
| Q3UQN2 | F-BAR domain only protein 2 OS=Mus musculus OX=10090 GN=I | 1 | 1 | 1 | 1 | 809 | 88,7 | 6,89 | | 0 | | 1 | 1,169 | 0,856 | 0,927 | 1,072 | 0,987 | 1,226 | 1,08 | 1,174 | 0,90 | -0,10 | 2,6E-01 |
| Q9QUI0 | Transforming protein RhoA OS=Mus musculus OX=10090 GN=Rh | 30 | 5 | 12 | 5 | 193 | 21,8 | 6,1 | 4627 | 9,86 | 3 | 2 | 1,054 | 0,957 | 1,212 | 1,023 | 0,678 | 1,128 | 0,976 | 0,921 | 1,15 | 0,15 | 2,6E-01 |
| Q8R0Y6 | Cytosolic 10-formyltetrahydrofolate dehydrogenase OS=Mus mu | 33 | 27 | 36 | 24 | 902 | 98,6 | 5,91 | 17079 | 38,54 | 20 | 9 | 1,223 | 1,306 | 1,156 | 1,323 | 1,414 | 1,214 | 1,363 | 1,307 | 0,95 | -0,05 | 2,6E-01 |
| Q8BP67 | 60S ribosomal protein L24 OS=Mus musculus OX=10090 GN=Rpl | 23 | 4 | 6 | 4 | 157 | 17,8 | 11,25 | 2717 | 5,03 | 2 | 2 | 1,068 | 0,896 | 1,01 | 0,945 | 0,953 | 0,899 | 0,962 | 0,898 | 1,06 | 0,06 | 2,6E-01 |
| P82349 | Beta-sarcoglycan OS=Mus musculus OX=10090 GN=Sgcb PE=1 S | 4 | 1 | 1 | 1 | 320 | 34,9 | 8,6 | 219 | | 1 | | 1,304 | 1,313 | 1,147 | 1,257 | 1,141 | 1,333 | 1,159 | 0,973 | 1,09 | 0,09 | 2,6E-01 |
| Q91YT7 | YTH domain-containing family protein 2 OS=Mus musculus OX= | 2 | 1 | 1 | 1 | 579 | 62,2 | 8,79 | | 2,17 | | 1 | 1,064 | 0,979 | 1,03 | 0,865 | 0,719 | 0,838 | 0,986 | 0,998 | 1,11 | 0,11 | 2,6E-01 |
| Q9DCD0 | 6-phosphogluconate dehydrogenase, decarboxylating OS=Mus r | 20 | 10 | 11 | 10 | 483 | 53,2 | 7,23 | 5090 | 2,58 | 8 | 2 | 0,956 | 0,996 | 1,033 | 0,959 | 0,994 | 0,947 | 0,965 | 0,916 | 1,03 | 0,03 | 2,6E-01 |
| Q68FL6 | Methionine--tRNA ligase, cytoplasmic OS=Mus musculus OX=10 | 3 | 2 | 3 | 2 | 902 | 101,4 | 7,14 | 1501 | | 2 | | 1,149 | 0,809 | 1,431 | 0,931 | 2,288 | 1,154 | 1,302 | 1,119 | 0,74 | -0,26 | 2,6E-01 |
| Q9D1Q6 | Endoplasmic reticulum resident protein 44 OS=Mus musculus O | 19 | 7 | 11 | 7 | 406 | 46,8 | 5,27 | 5175 | 3,53 | 7 | 1 | 0,937 | 0,98 | 0,908 | 1,028 | 1,261 | 0,887 | 1,204 | 0,97 | 0,89 | -0,11 | 2,6E-01 |
| Q8R1I1 | Cytochrome b-c1 complex subunit 9 OS=Mus musculus OX=100 | 27 | 1 | 2 | 1 | 64 | 7,4 | 9,19 | 1495 | | 1 | | 1,068 | 0,974 | 0,897 | 1,225 | 0,899 | 0,979 | 0,94 | 0,981 | 1,10 | 0,10 | 2,6E-01 |
| Q61749 | Translation initiation factor eIF-2B subunit delta OS=Mus muscu | 8 | 3 | 3 | 3 | 524 | 57,6 | 9,25 | 1954 | | 3 | | 1,009 | 0,995 | 0,989 | 1,077 | 1,094 | 0,815 | 0,936 | 0,921 | 1,08 | 0,08 | 2,6E-01 |
| Q6NZJ6 | Eukaryotic translation initiation factor 4 gamma 1 OS=Mus musc | 5 | 6 | 6 | 6 | 1600 | 176 | 5,4 | 2207 | 5,08 | 4 | 2 | 1,022 | 0,924 | 1,176 | 1 | 0,876 | 1,029 | 0,978 | 0,929 | 1,08 | 0,08 | 2,6E-01 |
| P68368 | Tubulin alpha-4A chain OS=Mus musculus OX=10090 GN=Tuba4 | 48 | 24 | 272 | 3 | 448 | 49,9 | 5,06 | 105048 | 339,26 | 22 | 12 | 1,076 | 1,195 | 1,118 | 1,112 | 0,769 | 1,097 | 1,157 | 1,034 | 1,11 | 0,11 | 2,6E-01 |
| O55013 | Trafficking protein particle complex subunit 3 OS=Mus musculus | 6 | 1 | 1 | 1 | 180 | 20,3 | 4,96 | 313 | | 1 | | 0,916 | 1,046 | 0,913 | 1,058 | 1,074 | 0,895 | 1,198 | 1,153 | 0,91 | -0,09 | 2,6E-01 |
| Q0VE82 | Copine-7 OS=Mus musculus OX=10090 GN=Cpne7 PE=1 SV=1 | 12 | 6 | 6 | 5 | 557 | 61,9 | 5,4 | 2794 | 5,51 | 4 | 2 | 0,903 | 0,902 | 0,921 | 0,862 | 0,889 | 0,926 | 0,915 | 0,94 | 0,98 | -0,02 | 2,6E-01 |
| Q9WUP7 | Ubiquitin carboxyl-terminal hydrolase isozyme L5 OS=Mus musc | 13 | 3 | 7 | 3 | 329 | 37,6 | 5,33 | 2145 | 3,98 | 2 | 2 | 1,064 | 1,129 | 1,15 | 1,343 | 1,334 | 1,154 | 1,349 | 1,226 | 0,93 | -0,07 | 2,6E-01 |
| Q9R0D8 | WD repeat-containing protein 54 OS=Mus musculus OX=10090 | 3 | 1 | 2 | 1 | 334 | 35,6 | 6,21 | | 4,47 | | 1 | 0,832 | 0,961 | 1,122 | 1,072 | 1,674 | 1,03 | 0,951 | 1,199 | 0,82 | -0,18 | 2,6E-01 |
| Q9QYJ0 | DnaJ homolog subfamily A member 2 OS=Mus musculus OX=10 | 17 | 7 | 14 | 7 | 412 | 45,7 | 6,48 | 5761 | 9,01 | 6 | 2 | 0,972 | 0,943 | 1,052 | 1,027 | 0,934 | 0,967 | 1,013 | 0,918 | 1,04 | 0,04 | 2,6E-01 |
| P16330 | 2',3'-cyclic-nucleotide 3'-phosphodiesterase OS=Mus musculus | 17 | 9 | 13 | 9 | 420 | 47,1 | 8,97 | 4715 | 15,39 | 5 | 5 | 1,159 | 1,042 | 1,077 | 1,05 | 0,958 | 0,958 | 1,081 | 1,099 | 1,06 | 0,06 | 2,6E-01 |
| Q61699 | Heat shock protein 105 kDa OS=Mus musculus OX=10090 GN=H | 33 | 26 | 47 | 24 | 858 | 96,3 | 5,53 | 16117 | 44,21 | 17 | 13 | 0,961 | 0,912 | 1,032 | 1,008 | 0,784 | 0,94 | 0,969 | 0,965 | 1,07 | 0,07 | 2,6E-01 |
| O08539 | Myc box-dependent-interacting protein 1 OS=Mus musculus OX | 23 | 13 | 25 | 11 | 588 | 64,4 | 5,03 | 10681 | 29,21 | 10 | 3 | 1,001 | 0,862 | 0,975 | 0,93 | 0,863 | 0,929 | 0,894 | 0,915 | 1,05 | 0,05 | 2,6E-01 |
| Q9QZM0 | Ubiquilin-2 OS=Mus musculus OX=10090 GN=Ubqln2 PE=1 SV=2 | 14 | 6 | 10 | 4 | 638 | 67,3 | 5,22 | 5017 | 5,17 | 6 | 1 | 0,889 | 1,144 | 0,957 | 1,15 | 0,898 | 0,875 | 0,995 | 1,005 | 1,10 | 0,10 | 2,6E-01 |
| Q6P9N1 | Hyccin OS=Mus musculus OX=10090 GN=Fam126a PE=1 SV=3 | 7 | 3 | 3 | 3 | 521 | 57,3 | 7,88 | 1256 | | 3 | | 0,834 | 0,859 | 0,793 | 0,799 | 1,437 | 1,092 | 0,872 | 0,687 | 0,80 | -0,20 | 2,6E-01 |
| Q9D164 | FXYD domain-containing ion transport regulator 6 OS=Mus mus | 23 | 1 | 1 | 1 | 94 | 10,4 | 5,11 | 721 | | 1 | | 0,954 | 1,567 | 1,05 | 1,541 | 1,056 | 1,131 | 1,109 | 1,008 | 1,19 | 0,19 | 2,6E-01 |
| P63001 | Ras-related C3 botulinum toxin substrate 1 OS=Mus musculus O | 40 | 8 | 23 | 4 | 192 | 21,4 | 8,5 | 9006 | 32,93 | 6 | 4 | 1,059 | 0,977 | 1,036 | 0,955 | 0,648 | 1,142 | 0,857 | 0,864 | 1,15 | 0,15 | 2,6E-01 |
| Q3UJP5 | Protein C8orf37 homolog OS=Mus musculus OX=10090 PE=1 SV | 11 | 2 | 2 | 2 | 209 | 23,8 | 6,86 | 315 | 0 | 1 | 1 | 1,235 | 1,086 | 1,303 | 0,985 | 0,987 | 1,064 | 1,217 | 0,609 | 1,19 | 0,19 | 2,6E-01 |
| Q8BWS5 | G protein-regulated inducer of neurite outgrowth 3 OS=Mus mu | 2 | 1 | 1 | 1 | 763 | 80,4 | 7,02 | | 2,23 | | 1 | 1,252 | 1,275 | 1,318 | 1,679 | 1,884 | 1,602 | 1,532 | 1,289 | 0,88 | -0,12 | 2,6E-01 |
| Q60739 | BAG family molecular chaperone regulator 1 OS=Mus musculus | 9 | 3 | 5 | 3 | 355 | 39,7 | 8,46 | 3071 | | 3 | | 0,943 | 1,002 | 1,008 | 1,016 | 1,085 | 0,988 | 0,975 | 1,103 | 0,96 | -0,04 | 2,6E-01 |
| Q62172 | RalA-binding protein 1 OS=Mus musculus OX=10090 GN=Ralbp1 | 1 | 1 | 1 | 1 | 648 | 75 | 5,92 | 357 | | 1 | | 1,026 | 1,076 | 1,03 | 1,122 | 0,802 | 0,981 | 1,059 | 1,08 | 1,08 | 0,08 | 2,6E-01 |
| Q62348 | Translin OS=Mus musculus OX=10090 GN=Tsn PE=1 SV=1 | 14 | 4 | 5 | 3 | 228 | 26,2 | 6,44 | 1707 | 4,06 | 3 | 1 | 1,174 | 0,982 | 1,176 | 1,33 | 0,995 | 1,197 | 1,102 | 0,879 | 1,12 | 0,12 | 2,6E-01 |
| Q921H9 | Cytochrome c oxidase assembly factor 7 OS=Mus musculus OX= | 8 | 2 | 3 | 2 | 231 | 25,6 | 6,29 | 1560 | | 2 | | 0,847 | 1,208 | 1,182 | 1,006 | 1,155 | 1,07 | 1,307 | 1,193 | 0,90 | -0,10 | 2,6E-01 |
| Q9D8P4 | 39S ribosomal protein L17, mitochondrial OS=Mus musculus OX | 7 | 1 | 1 | 1 | 176 | 20,2 | 9,82 | 625 | | 1 | | 0,85 | 0,718 | 0,862 | 1,126 | 1,365 | 1,017 | 0,871 | 0,979 | 0,84 | -0,16 | 2,6E-01 |
| P39053-3 | Isoform 3 of Dynamin-1 OS=Mus musculus OX=10090 GN=Dnm1 | 34 | 34 | 70 | 1 | 851 | 95,9 | 6,76 | 25010 | 71,02 | 20 | 19 | 0,858 | 0,797 | 0,853 | 0,747 | 0,637 | 0,879 | 0,782 | 0,637 | 1,11 | 0,11 | 2,6E-01 |
| Q8VI75 | Importin-4 OS=Mus musculus OX=10090 GN=Ipo4 PE=1 SV=1 | 4 | 4 | 5 | 4 | 1082 | 119,2 | 5,03 | 2773 | | 4 | | 1,078 | 1,076 | 1,03 | 1,099 | 2,233 | 1,134 | 1,068 | 1,21 | 0,76 | -0,24 | 2,6E-01 |
| Q8JZK9 | Hydroxymethylglutaryl-CoA synthase, cytoplasmic OS=Mus mus | 16 | 8 | 15 | 7 | 520 | 57,5 | 5,99 | 3598 | 25,93 | 4 | 4 | 0,995 | 0,991 | 0,931 | 1,025 | 0,773 | 1,002 | 0,879 | 1 | 1,08 | 0,08 | 2,6E-01 |
| Q8K4Q0 | Regulatory-associated protein of mTOR OS=Mus musculus OX=1 | 1 | 1 | 1 | 1 | 1335 | 149,4 | 6,87 | 568 | | 1 | | 1,354 | 1,054 | 0,99 | 1,224 | 1,668 | 1,098 | 1,22 | 1,366 | 0,86 | -0,14 | 2,6E-01 |
| P34884 | Macrophage migration inhibitory factor OS=Mus musculus OX=1 | 20 | 3 | 5 | 3 | 115 | 12,5 | 7,34 | | 11,54 | | 3 | 0,891 | 0,9 | 0,922 | 0,828 | 1,508 | 0,937 | 0,929 | 0,898 | 0,83 | -0,17 | 2,6E-01 |
| P18760 | Cofilin-1 OS=Mus musculus OX=10090 GN=Cfl1 PE=1 SV=3 | 75 | 18 | 44 | 17 | 166 | 18,5 | 8,09 | 13277 | 68,45 | 9 | 11 | 0,987 | 1,132 | 0,918 | 1,123 | 0,709 | 1,134 | 0,911 | 0,906 | 1,14 | 0,14 | 2,6E-01 |
| O09131 | Glutathione S-transferase omega-1 OS=Mus musculus OX=1009 | 28 | 11 | 14 | 11 | 240 | 27,5 | 7,36 | 4071 | 17,21 | 6 | 5 | 0,974 | 0,955 | 1,054 | 1,079 | 0,581 | 1,146 | 0,886 | 0,861 | 1,17 | 0,17 | 2,6E-01 |
| Q3UPL0 | Protein transport protein Sec31A OS=Mus musculus OX=10090 | 10 | 9 | 16 | 9 | 1230 | 133,5 | 6,76 | 4502 | 19,42 | 7 | 3 | 1,131 | 1,028 | 1,078 | 0,948 | 1,143 | 0,954 | 1,191 | 1,302 | 0,91 | -0,09 | 2,6E-01 |
| Q9JKY5 | Huntingtin-interacting protein 1-related protein OS=Mus muscu | 7 | 6 | 7 | 5 | 1068 | 119,4 | 6,52 | 3436 | 3,05 | 5 | 1 | 1,005 | 0,945 | 1,104 | 1,062 | 0,658 | 0,915 | 1,117 | 0,932 | 1,14 | 0,14 | 2,6E-01 |
| P52760 | 2-iminobutanoate/2-iminopropanoate deaminase OS=Mus mus | 7 | 1 | 1 | 1 | 135 | 14,2 | 8,68 | | 3,26 | | 1 | 1,308 | 1,202 | 1,379 | 1,37 | 2,931 | 1,422 | 1,465 | 1,331 | 0,74 | -0,26 | 2,7E-01 |
| P54227 | Stathmin OS=Mus musculus OX=10090 GN=Stmn1 PE=1 SV=2 | 83 | 22 | 70 | 17 | 149 | 17,3 | 5,97 | 18903 | 103,58 | 8 | 17 | 0,94 | 0,964 | 1,019 | 0,943 | 0,879 | 1,009 | 0,884 | 0,921 | 1,05 | 0,05 | 2,7E-01 |
| P62192 | 26S proteasome regulatory subunit 4 OS=Mus musculus OX=10 | 23 | 10 | 13 | 10 | 440 | 49,2 | 6,21 | 5121 | 11,06 | 6 | 4 | 0,99 | 0,961 | 1,01 | 0,916 | 0,799 | 0,97 | 0,939 | 0,952 | 1,06 | 0,06 | 2,7E-01 |
| P21279 | Guanine nucleotide-binding protein G(q) subunit alpha OS=Mus | 43 | 15 | 24 | 9 | 359 | 42,1 | 5,68 | 11068 | 21,06 | 12 | 5 | 1,123 | 1,028 | 0,974 | 1,11 | 0,79 | 1,068 | 1,027 | 1 | 1,09 | 0,09 | 2,7E-01 |
| P20108 | Thioredoxin-dependent peroxide reductase, mitochondrial OS=I | 38 | 8 | 19 | 8 | 257 | 28,1 | 7,58 | 5020 | 30,72 | 5 | 5 | 0,975 | 1,084 | 0,871 | 1,054 | 0,814 | 0,991 | 0,973 | 0,9 | 1,08 | 0,08 | 2,7E-01 |
| O09012 | Peroxisomal targeting signal 1 receptor OS=Mus musculus OX=1 | 3 | 2 | 2 | 2 | 639 | 70,7 | 4,55 | 958 | | 2 | | 0,921 | 0,765 | 1,027 | 0,859 | 1,733 | 0,828 | 1,031 | 0,998 | 0,78 | -0,22 | 2,7E-01 |
| Q99KW9 | T-cell immunomodulatory protein OS=Mus musculus OX=10090 | 1 | 1 | 1 | 1 | 610 | 67,4 | 5,73 | 835 | | 1 | | 0,883 | 0,781 | 1,113 | 0,785 | 0,703 | 0,895 | 0,721 | 0,806 | 1,14 | 0,14 | 2,7E-01 |
| Q9D6K8 | FUN14 domain-containing protein 2 OS=Mus musculus OX=100 | 12 | 2 | 3 | 2 | 151 | 16,6 | 9,7 | 1559 | | 2 | | 1,04 | 0,979 | 1,015 | 1,006 | 0,944 | 1,231 | 1,004 | 1,239 | 0,91 | -0,09 | 2,7E-01 |
| O88456 | Calpain small subunit 1 OS=Mus musculus OX=10090 GN=Capns | 9 | 2 | 2 | 2 | 269 | 28,4 | 5,63 | 1211 | | 2 | | 0,791 | 0,956 | 0,842 | 0,924 | 1,045 | 0,916 | 0,972 | 0,852 | 0,93 | -0,07 | 2,7E-01 |
| Q924Z4 | Ceramide synthase 2 OS=Mus musculus OX=10090 GN=Cers2 PE | 4 | 1 | 1 | 1 | 380 | 45 | 8,75 | 534 | | 1 | | 0,929 | 1,024 | 1,033 | 0,89 | 1,22 | 1,024 | 0,885 | 1,143 | 0,91 | -0,09 | 2,7E-01 |
| P11276 | Fibronectin OS=Mus musculus OX=10090 GN=Fn1 PE=1 SV=4 | 1 | 1 | 1 | 1 | 2477 | 272,4 | 5,59 | 459 | | 1 | | 0,771 | 0,787 | 0,608 | 0,52 | 1,128 | 0,859 | 0,798 | 0,555 | 0,80 | -0,20 | 2,7E-01 |
| Q8K0G5 | EARP and GARP complex-interacting protein 1 OS=Mus musculu | 9 | 3 | 4 | 3 | 386 | 43,1 | 5,14 | 988 | 2,7 | 2 | 1 | 1,021 | 0,983 | 0,985 | 1,071 | 0,789 | 1,026 | 0,901 | 1,04 | 1,08 | 0,08 | 2,7E-01 |
| P08752 | Guanine nucleotide-binding protein G(i) subunit alpha-2 OS=Mu | 41 | 13 | 26 | 7 | 355 | 40,5 | 5,45 | 9072 | 21,6 | 8 | 5 | 0,929 | 0,977 | 0,849 | 0,975 | 1,127 | 0,872 | 1,071 | 0,97 | 0,92 | -0,08 | 2,7E-01 |

| | | | | | | | | | | | | | | | | | | | | | | |
|---|---|---|---|---|---|---|---|---|---|---|---|---|---|---|---|---|---|---|---|---|---|---|
| P55821 | Stathmin-2 OS=Mus musculus OX=10090 GN=Stmn2 PE=1 SV=1 | 51 | 11 | 32 | 5 | 179 | 20,8 | 8,32 | 6711 | 63,31 | 4 | 8 | 0,882 | 1 | 0,994 | 1,072 | 0,936 | 0,83 | 0,975 | 0,959 | 1,07 | 0,07 | 2,7E-01 |
| Q8VHR0 | Protocadherin 18 OS=Mus musculus OX=10090 GN=Pcdh18 PE= | 1 | 1 | 1 | 1 | 1134 | 125,3 | 5,25 | | 2,44 | | 1 | 1,066 | 1 | 0,965 | 0,903 | 0,934 | 1,057 | 1,15 | 1,065 | 0,94 | -0,06 | 2,7E-01 |
| P42208 | Septin-2 OS=Mus musculus OX=10090 GN=Septin2 PE=1 SV=2 | 16 | 6 | 9 | 5 | 361 | 41,5 | 6,55 | 4642 | 2,83 | 5 | 1 | 1,09 | 0,938 | 1,246 | 0,936 | 0,86 | 0,979 | 1,047 | 0,915 | 1,11 | 0,11 | 2,7E-01 |
| P97784 | Cryptochrome-1 OS=Mus musculus OX=10090 GN=Cry1 PE=1 SV | 1 | 1 | 1 | 1 | 606 | 68 | 7,97 | | 2,06 | | 1 | 1,206 | 0,921 | 0,977 | 0,914 | 0,757 | 1,042 | 0,892 | 0,89 | 1,12 | 0,12 | 2,7E-01 |
| Q9EQF5 | Dihydropyrimidinase OS=Mus musculus OX=10090 GN=Dpys PE= | 6 | 2 | 3 | 1 | 519 | 56,7 | 7,2 | 1753 | 2,04 | 1 | 1 | 1,063 | 1,209 | 0,982 | 1,75 | 1,578 | 1,152 | 1,595 | 1,703 | 0,83 | -0,17 | 2,7E-01 |
| Q9D1G1 | Ras-related protein Rab-1B OS=Mus musculus OX=10090 GN=Ra | 56 | 9 | 16 | 4 | 201 | 22,2 | 5,73 | 7859 | 11,09 | 8 | 4 | 1,079 | 1,039 | 0,961 | 1,084 | 0,878 | 1,03 | 0,99 | 1,039 | 1,06 | 0,06 | 2,7E-01 |
| Q8BGB7 | Enolase-phosphatase E1 OS=Mus musculus OX=10090 GN=Enop | 8 | 2 | 2 | 2 | 257 | 28,6 | 4,92 | 940 | | 2 | | 1,362 | 1,372 | 1,316 | 1,278 | 1,294 | 1,633 | 1,253 | 1,789 | 0,89 | -0,11 | 2,7E-01 |
| Q9QYS9 | Protein quaking OS=Mus musculus OX=10090 GN=Qki PE=1 SV= | 7 | 2 | 2 | 2 | 341 | 37,6 | 8,56 | 733 | | 2 | | 1,398 | 1,548 | 1,455 | 1,533 | 1,176 | 1,111 | 1,665 | 1,358 | 1,12 | 0,12 | 2,7E-01 |
| Q60692 | Proteasome subunit beta type-6 OS=Mus musculus OX=10090 G | 17 | 4 | 5 | 4 | 238 | 25,4 | 5,11 | 1963 | 6,85 | 2 | 2 | 1,014 | 0,934 | 0,914 | 0,988 | 0,931 | 0,971 | 0,885 | 0,922 | 1,04 | 0,04 | 2,7E-01 |
| Q9QZ73 | DCN1-like protein 1 OS=Mus musculus OX=10090 GN=Dcun1d1 | 3 | 1 | 1 | 1 | 259 | 30,1 | 5,34 | | 2,33 | | 1 | 0,872 | 1,019 | 0,914 | 0,775 | 0,452 | 0,935 | 0,816 | 0,815 | 1,19 | 0,19 | 2,7E-01 |
| Q8C031 | Leucine-rich repeat-containing protein 4C OS=Mus musculus OX | 2 | 1 | 1 | 1 | 640 | 71,9 | 7,15 | | 0 | | 1 | 1,001 | 1,321 | 0,996 | 1,335 | 1,76 | 1,3 | 0,941 | 1,667 | 0,82 | -0,18 | 2,7E-01 |
| Q8BYA0 | Tubulin-specific chaperone D OS=Mus musculus OX=10090 GN= | 3 | 3 | 3 | 3 | 1196 | 133,2 | 6,51 | 1992 | | 3 | | 1,106 | 0,753 | 0,872 | 0,923 | 1,578 | 0,819 | 1,106 | 1,007 | 0,81 | -0,19 | 2,7E-01 |
| P52734 | FYVE, RhoGEF and PH domain-containing protein 1 OS=Mus mu: | 3 | 2 | 2 | 2 | 960 | 106,3 | 6,58 | 914 | | 2 | | 0,919 | 1,109 | 0,921 | 0,909 | 1,25 | 0,88 | 1,06 | 1,103 | 0,90 | -0,10 | 2,7E-01 |
| Q6Q477 | Plasma membrane calcium-transporting ATPase 4 OS=Mus musc | 19 | 21 | 37 | 9 | 1205 | 133 | 6,13 | 12755 | 46,9 | 14 | 9 | 0,828 | 0,859 | 0,701 | 0,819 | 1,094 | 0,702 | 0,975 | 0,87 | 0,88 | -0,12 | 2,7E-01 |
| Q8VHW2 | Voltage-dependent calcium channel gamma-8 subunit OS=Mus | 2 | 1 | 1 | 1 | 423 | 43,4 | 9,2 | | 0 | | 1 | 1,414 | 1,041 | 1,626 | 1,264 | 1,04 | 1,206 | 1,37 | 1,014 | 1,15 | 0,15 | 2,7E-01 |
| Q6NZL0 | Protein SOGA3 OS=Mus musculus OX=10090 GN=Soga3 PE=1 SV | 13 | 11 | 12 | 11 | 945 | 103,4 | 6,14 | 3010 | 19,02 | 5 | 6 | 0,983 | 0,991 | 0,937 | 0,994 | 0,919 | 0,851 | 1,025 | 0,926 | 1,05 | 0,05 | 2,7E-01 |
| Q8C1E7 | Transmembrane protein 120A OS=Mus musculus OX=10090 GN | 2 | 1 | 1 | 1 | 343 | 40,7 | 8,88 | | 2,43 | | 1 | 1,028 | 0,897 | 1,311 | 1,039 | 0,576 | 1,08 | 0,933 | 1,005 | 1,19 | 0,19 | 2,7E-01 |
| P02798 | Metallothionein-2 OS=Mus musculus OX=10090 GN=Mt2 PE=1 S | 20 | 1 | 1 | 1 | 61 | 6,1 | 7,83 | | 2,11 | | 1 | 0,888 | 1,365 | 1,066 | 1,294 | 0,867 | 1,108 | 1,066 | 0,989 | 1,14 | 0,14 | 2,7E-01 |
| Q99J27 | Acetyl-coenzyme A transporter 1 OS=Mus musculus OX=10090 ( | 4 | 1 | 1 | 1 | 550 | 61 | 8,16 | 97 | | 1 | | 1,374 | 1,248 | 0,906 | 0,905 | 2,017 | 1,175 | 1,191 | 1,201 | 0,79 | -0,21 | 2,7E-01 |
| S4R1M9 | Oxysterol-binding protein-related protein 10 OS=Mus musculus | 1 | 1 | 1 | 1 | 766 | 83,8 | 8,28 | 528 | | 1 | | 1,14 | 1,136 | 0,914 | 0,973 | 1,025 | 1,327 | 1,004 | 1,314 | 0,89 | -0,11 | 2,7E-01 |
| Q9R0A0 | Peroxisomal membrane protein PEX14 OS=Mus musculus OX=1( | 3 | 1 | 1 | 1 | 376 | 41,2 | 5,11 | 525 | | 1 | | 0,769 | 1,065 | 1,174 | 0,991 | 1,601 | 1,208 | 1,052 | 0,943 | 0,83 | -0,17 | 2,7E-01 |
| P28656 | Nucleosome assembly protein 1-like 1 OS=Mus musculus OX=10 | 10 | 3 | 6 | 2 | 391 | 45,3 | 4,46 | 3929 | | 3 | | 0,852 | 1,01 | 0,663 | 0,913 | 0,988 | 0,928 | 0,966 | 0,916 | 0,91 | -0,09 | 2,8E-01 |
| Q61545 | RNA-binding protein EWS OS=Mus musculus OX=10090 GN=Ews | 2 | 1 | 3 | 1 | 655 | 68,4 | 9,33 | 1602 | | 1 | | 0,932 | 1,273 | 0,994 | 1,066 | 0,918 | 0,921 | 1,095 | 0,917 | 1,11 | 0,11 | 2,8E-01 |
| Q60930 | Voltage-dependent anion-selective channel protein 2 OS=Mus n | 34 | 9 | 14 | 9 | 295 | 31,7 | 7,49 | 5159 | 8,68 | 6 | 3 | 1,034 | 0,997 | 1,01 | 1,014 | 1,332 | 1,02 | 1,007 | 1,064 | 0,92 | -0,08 | 2,8E-01 |
| Q6A0D4 | Raftlin OS=Mus musculus OX=10090 GN=Rftn1 PE=1 SV=4 | 5 | 2 | 2 | 2 | 554 | 61,5 | 7,68 | 504 | | 2 | | 1,07 | 1,153 | 1,25 | 1,826 | 1,485 | 1,526 | 1,537 | 1,575 | 0,87 | -0,13 | 2,8E-01 |
| Q9WV80 | Sorting nexin-1 OS=Mus musculus OX=10090 GN=Snx1 PE=1 SV= | 8 | 3 | 3 | 3 | 522 | 58,9 | 5,22 | 568 | 3,92 | 2 | 1 | 0,873 | 1,066 | 0,711 | 1,011 | 0,959 | 0,881 | 1,229 | 1,127 | 0,87 | -0,13 | 2,8E-01 |
| O08795 | Glucosidase 2 subunit beta OS=Mus musculus OX=10090 GN=Pr | 13 | 6 | 9 | 6 | 521 | 58,8 | 4,46 | 1689 | 14,69 | 1 | 5 | 1,013 | 1,031 | 1,19 | 1,171 | 0,877 | 1,034 | 1,092 | 1,079 | 1,08 | 0,08 | 2,8E-01 |
| P17182 | Alpha-enolase OS=Mus musculus OX=10090 GN=Eno1 PE=1 SV= | 68 | 30 | 92 | 24 | 434 | 47,1 | 6,8 | 34477 | 112,44 | 20 | 13 | 0,991 | 0,981 | 0,984 | 0,954 | 1,175 | 1,03 | 0,961 | 0,979 | 0,94 | -0,06 | 2,8E-01 |
| Q8BG05 | Heterogeneous nuclear ribonucleoprotein A3 OS=Mus musculus | 46 | 17 | 25 | 15 | 379 | 39,6 | 9,01 | 10758 | 19,65 | 12 | 5 | 0,92 | 0,886 | 0,92 | 1,109 | 0,976 | 0,957 | 1,157 | 1,073 | 0,92 | -0,08 | 2,8E-01 |
| Q5SW19 | Clustered mitochondria protein homolog OS=Mus musculus OX= | 2 | 2 | 2 | 2 | 1315 | 148 | 6,02 | 616 | | 2 | | 0,859 | 0,768 | 1,1 | 0,944 | 1,068 | 1,147 | 1,034 | 0,863 | 0,89 | -0,11 | 2,8E-01 |
| Q9JLN9 | Serine/threonine-protein kinase mTOR OS=Mus musculus OX=1( | 2 | 4 | 5 | 4 | 2549 | 288,6 | 7,17 | 951 | 6,28 | 2 | 2 | 0,841 | 0,952 | 1,132 | 1,335 | 0,982 | 0,915 | 0,754 | 1,022 | 1,16 | 0,16 | 2,8E-01 |
| P11031 | Activated RNA polymerase II transcriptional coactivator p15 OS= | 9 | 1 | 1 | 1 | 127 | 14,4 | 9,6 | 657 | | 1 | | 1,035 | 0,896 | 0,826 | 1,078 | 0,873 | 0,819 | 0,954 | 0,879 | 1,09 | 0,09 | 2,8E-01 |
| P57776 | Elongation factor 1-delta OS=Mus musculus OX=10090 GN=Eef1 | 36 | 8 | 10 | 7 | 281 | 31,3 | 5,02 | 4283 | 7,82 | 6 | 2 | 0,994 | 0,949 | 0,956 | 1,044 | 1,154 | 0,952 | 0,996 | 1,078 | 0,94 | -0,06 | 2,8E-01 |
| Q80ZJ1 | Ras-related protein Rap-2a OS=Mus musculus OX=10090 GN=Ra | 19 | 4 | 7 | 3 | 183 | 20,6 | 4,82 | 3962 | 3,53 | 3 | 1 | 0,948 | 1,109 | 1,039 | 1,17 | 0,767 | 1,12 | 0,971 | 0,993 | 1,11 | 0,11 | 2,8E-01 |
| Q8R0X7 | Sphingosine-1-phosphate lyase 1 OS=Mus musculus OX=10090 ( | 4 | 2 | 3 | 2 | 568 | 63,6 | 9,1 | 1350 | | 2 | | 1,048 | 0,696 | 0,826 | 0,878 | 0,928 | 1,002 | 0,952 | 0,923 | 0,91 | -0,09 | 2,8E-01 |
| Q8BIJ6 | Isoleucine--tRNA ligase, mitochondrial OS=Mus musculus OX=1( | 8 | 6 | 6 | 6 | 1012 | 112,7 | 6,81 | 2168 | 6,78 | 4 | 2 | 1,055 | 1,016 | 1,015 | 0,991 | 1,184 | 1 | 1,022 | 1,076 | 0,95 | -0,05 | 2,8E-01 |
| Q99JY0 | Trifunctional enzyme subunit beta, mitochondrial OS=Mus musc | 16 | 8 | 10 | 8 | 475 | 51,4 | 9,38 | 1611 | 18,04 | 3 | 5 | 0,97 | 1,162 | 1,016 | 1,237 | 1,328 | 1,021 | 1,184 | 1,293 | 0,91 | -0,09 | 2,8E-01 |
| O55222 | Integrin-linked protein kinase OS=Mus musculus OX=10090 GN= | 3 | 1 | 1 | 1 | 452 | 51,3 | 8,07 | 340 | | 1 | | 0,928 | 0,88 | 0,97 | 1,312 | 1,25 | 0,53 | 0,879 | 0,438 | 1,32 | 0,32 | 2,8E-01 |
| P57780 | Alpha-actinin-4 OS=Mus musculus OX=10090 GN=Actn4 PE=1 SV | 36 | 29 | 42 | 21 | 912 | 104,9 | 5,41 | 19051 | 33,85 | 23 | 8 | 0,952 | 1,036 | 0,884 | 0,973 | 1,212 | 0,893 | 1,075 | 1,013 | 0,92 | -0,08 | 2,8E-01 |
| P97765 | WW domain-binding protein 2 OS=Mus musculus OX=10090 GN | 18 | 4 | 5 | 4 | 261 | 28 | 6,33 | 872 | 7,04 | 2 | 2 | 1,121 | 1,069 | 1,174 | 1,13 | 1,024 | 1,174 | 1,01 | 1,082 | 1,05 | 0,05 | 2,8E-01 |
| P24472 | Glutathione S-transferase A4 OS=Mus musculus OX=10090 GN= | 9 | 2 | 3 | 2 | 222 | 25,5 | 7,39 | | 7,78 | | 2 | 0,997 | 1,22 | 0,993 | 1,126 | 1,364 | 1,14 | 1,102 | 1,12 | 0,92 | -0,08 | 2,8E-01 |
| Q8BMK4 | Cytoskeleton-associated protein 4 OS=Mus musculus OX=10090 | 20 | 11 | 17 | 11 | 575 | 63,7 | 5,64 | 5540 | 13,85 | 7 | 5 | 0,854 | 0,823 | 0,875 | 0,949 | 1,166 | 0,757 | 1,05 | 0,957 | 0,89 | -0,11 | 2,8E-01 |
| Q9QYJ3 | DnaJ homolog subfamily B member 1 OS=Mus musculus OX=10( | 11 | 4 | 5 | 4 | 340 | 38,1 | 8,63 | 1276 | 2,68 | 2 | 2 | 0,977 | 0,838 | 1,026 | 0,914 | 0,826 | 0,915 | 0,945 | 0,829 | 1,07 | 0,07 | 2,8E-01 |
| Q91V41 | Ras-related protein Rab-14 OS=Mus musculus OX=10090 GN=Ra | 72 | 12 | 22 | 11 | 215 | 23,9 | 6,21 | 9702 | 6,92 | 11 | 2 | 1,049 | 0,914 | 1,145 | 1,063 | 0,817 | 1,095 | 1,005 | 0,883 | 1,10 | 0,10 | 2,8E-01 |
| Q8BGN8 | Synaptoporin OS=Mus musculus OX=10090 GN=Synpr PE=1 SV= | 5 | 1 | 1 | 1 | 265 | 29,2 | 6,3 | 654 | | 1 | | 0,866 | 0,948 | 0,792 | 0,854 | 1,056 | 0,807 | 0,861 | 1,135 | 0,90 | -0,10 | 2,8E-01 |
| Q922S4 | cGMP-dependent 3',5'-cyclic phosphodiesterase OS=Mus muscu | 13 | 8 | 9 | 8 | 939 | 105,6 | 5,41 | 3517 | 2,49 | 6 | 2 | 1,061 | 0,976 | 0,896 | 0,827 | 1,069 | 0,963 | 0,965 | 1,032 | 0,93 | -0,07 | 2,8E-01 |
| P56959 | RNA-binding protein FUS OS=Mus musculus OX=10090 GN=Fus I | 6 | 2 | 3 | 2 | 518 | 52,6 | 9,36 | 1625 | 1,98 | 1 | 1 | 0,792 | 0,833 | 0,835 | 0,912 | 0,741 | 0,676 | 0,868 | 0,844 | 1,08 | 0,08 | 2,8E-01 |
| A2RSQ0 | DENN domain-containing protein 5B OS=Mus musculus OX=10( | 1 | 1 | 1 | 1 | 1274 | 144,5 | 6,68 | 245 | | 1 | | 0,798 | 0,799 | 1,294 | 0,76 | 4,463 | 0,889 | 0,842 | 1,55 | 0,47 | -0,53 | 2,8E-01 |
| P60904 | DnaJ homolog subfamily C member 5 OS=Mus musculus OX=10( | 42 | 6 | 17 | 6 | 198 | 22,1 | 5,07 | 5091 | 34,61 | 5 | 3 | 1,069 | 1,099 | 0,984 | 1,112 | 0,897 | 0,931 | 1,049 | 1,113 | 1,07 | 0,07 | 2,8E-01 |
| Q8CIQ7 | Dedicator of cytokinesis protein 3 OS=Mus musculus OX=10090 | 3 | 4 | 5 | 4 | 2027 | 232,8 | 6,99 | 1715 | 6,51 | 3 | 1 | 0,784 | 0,937 | 0,946 | 1,032 | 1,354 | 0,859 | 0,998 | 1,038 | 0,87 | -0,13 | 2,8E-01 |
| Q9WV68 | Peroxisomal 2,4-dienoyl-CoA reductase OS=Mus musculus OX=1 | 5 | 1 | 1 | 1 | 292 | 31,3 | 8,81 | 735 | | 1 | | 1,623 | 1,769 | 2,033 | 1,599 | 3,598 | 1,498 | 1,747 | 2,449 | 0,76 | -0,24 | 2,8E-01 |
| P50153 | Guanine nucleotide-binding protein G(I)/G(S)/G(O) subunit gam | 52 | 3 | 5 | 3 | 75 | 8,4 | 7,08 | 1239 | 8,25 | 1 | 2 | 1,129 | 0,937 | 1,38 | 0,998 | 0,936 | 1,238 | 0,84 | 0,747 | 1,18 | 0,18 | 2,8E-01 |
| Q8BGC4 | Prostaglandin reductase-3 OS=Mus musculus OX=10090 GN=Zac | 3 | 1 | 1 | 1 | 377 | 40,5 | 7,42 | 360 | | 1 | | 1,413 | 1,361 | 1,572 | 1,78 | 0,817 | 1,246 | 1,588 | 1,533 | 1,18 | 0,18 | 2,8E-01 |
| P49817 | Caveolin-1 OS=Mus musculus OX=10090 GN=Cav1 PE=1 SV=1 | 6 | 1 | 1 | 1 | 178 | 20,5 | 6,02 | 709 | | 1 | | 0,881 | 0,862 | 0,807 | 1,083 | 1,176 | 0,807 | 1,821 | 0,93 | 0,77 | -0,23 | 2,8E-01 |
| Q8BWM0 | Prostaglandin E synthase 2 OS=Mus musculus OX=10090 GN=Pt | 9 | 3 | 4 | 3 | 384 | 43,3 | 9 | 1496 | 2,65 | 2 | 1 | 0,912 | 0,919 | 0,888 | 0,824 | 0,842 | 0,97 | 0,95 | 0,952 | 0,95 | -0,05 | 2,8E-01 |
| Q9CXP8 | Guanine nucleotide-binding protein G(I)/G(S)/G(O) subunit gam | 37 | 2 | 3 | 2 | 68 | 7,2 | 7,85 | 670 | 4,96 | 1 | 2 | 1,024 | 1,079 | 1,067 | 0,942 | 0,898 | 1,005 | 0,963 | 1,041 | 1,05 | 0,05 | 2,8E-01 |
| Q8BWT1 | 3-ketoacyl-CoA thiolase, mitochondrial OS=Mus musculus OX=1 | 37 | 14 | 18 | 14 | 397 | 41,8 | 8,09 | 5109 | 20,99 | 8 | 8 | 1,056 | 0,961 | 1,295 | 1,022 | 1,76 | 1,112 | 1,216 | 1,071 | 0,84 | -0,16 | 2,8E-01 |
| Q8BHG2 | CXXC motif containing zinc binding protein OS=Mus musculus O | 11 | 1 | 2 | 1 | 160 | 18 | 5,12 | 526 | | 1 | | 0,987 | 1,145 | 0,911 | 0,993 | 1,107 | 1,069 | 1,073 | 1,029 | 0,94 | -0,06 | 2,9E-01 |
| P97817 | Cerebellar degeneration-related protein 2 OS=Mus musculus OX | 4 | 1 | 1 | 1 | 455 | 52,1 | 5 | 234 | | 1 | | 0,761 | 0,843 | 0,645 | 0,807 | 1,279 | 0,583 | 0,797 | 1,196 | 0,79 | -0,21 | 2,9E-01 |
| Q9JMB8 | Contactin-6 OS=Mus musculus OX=10090 GN=Cntn6 PE=1 SV=2 | 13 | 9 | 12 | 9 | 1028 | 113,7 | 6,64 | 3209 | 11,38 | 6 | 4 | 1,035 | 1,024 | 0,917 | 1,23 | 1,155 | 1,129 | 1,021 | 1,328 | 0,91 | -0,09 | 2,9E-01 |
| Q9D883 | Splicing factor U2AF 35 kDa subunit OS=Mus musculus OX=1009 | 3 | 1 | 1 | 1 | 239 | 27,8 | 8,81 | | 2,19 | | 1 | 0,871 | 0,965 | 0,908 | 0,91 | 1,277 | 0,774 | 1,107 | 0,998 | 0,88 | -0,12 | 2,9E-01 |
| P01837 | Immunoglobulin kappa constant OS=Mus musculus OX=10090 G | 18 | 2 | 10 | 2 | 107 | 11,9 | 5,9 | 5129 | | 2 | | 1,715 | 1,692 | 1,56 | 1,731 | 2,412 | 1,62 | 1,558 | 2,067 | 0,87 | -0,13 | 2,9E-01 |
| Q9R1P1 | Proteasome subunit beta type-3 OS=Mus musculus OX=10090 G | 20 | 4 | 8 | 4 | 205 | 22,9 | 6,55 | 3843 | 4,98 | 3 | 1 | 1,066 | 1,111 | 0,983 | 1,124 | 1,193 | 1,106 | 1,041 | 1,159 | 0,95 | -0,05 | 2,9E-01 |
| Q3TKY6 | Spliceosome-associated protein CWC27 homolog OS=Mus musc | 1 | 1 | 1 | 1 | 469 | 53,5 | 5,53 | 239 | | 1 | | 0,871 | 0,949 | 1,17 | 1,333 | 1,249 | 1,063 | 1,145 | 1,638 | 0,85 | -0,15 | 2,9E-01 |
| Q9D8T7 | SRA stem-loop-interacting RNA-binding protein, mitochondrial ( | 40 | 4 | 5 | 4 | 112 | 12,6 | 9,82 | 2247 | 3,9 | 3 | 1 | 1,08 | 1,06 | 1,079 | 1,169 | 1,284 | 1,094 | 1,095 | 1,153 | 0,95 | -0,05 | 2,9E-01 |

| | | | | | | | | | | | | | | | | | | | | | | | |
|---|---|---|---|---|---|---|---|---|---|---|---|---|---|---|---|---|---|---|---|---|---|---|---|
| Q62165 | Dystroglycan OS=Mus musculus OX=10090 GN=Dag1 PE=1 SV=4 | 11 | 8 | 10 | 8 | 893 | 96,8 | 8,44 | 3083 | 7,45 | 5 | 4 | 1,003 | 1,013 | 0,887 | 1,128 | 0,96 | 0,978 | 0,918 | 0,937 | 1,06 | 0,06 | 2,9E-01 |
| Q9JKK7 | Tropomodulin-2 OS=Mus musculus OX=10090 GN=Tmod2 PE=1 | 28 | 9 | 13 | 9 | 351 | 39,5 | 5,35 | 8202 | 2,66 | 9 | 1 | 0,999 | 1,05 | 1,057 | 1,054 | 0,655 | 1,116 | 0,981 | 0,95 | 1,12 | 0,12 | 2,9E-01 |
| Q810U4-5 | Isoform 5 of Neuronal cell adhesion molecule OS=Mus musculus | 28 | 6 | 14 | 1 | 260 | 28,9 | 8,43 | 2562 | 23,94 | 4 | 4 | 1,299 | 1,173 | 1,193 | 1,381 | 2,958 | 1,255 | 1,203 | 1,565 | 0,72 | -0,28 | 2,9E-01 |
| Q7TT37 | Elongator complex protein 1 OS=Mus musculus OX=10090 GN=E | 4 | 5 | 6 | 5 | 1333 | 149,5 | 6 | 3119 | 3,25 | 4 | 1 | 0,994 | 0,864 | 0,994 | 0,945 | 0,94 | 1,093 | 0,955 | 1,027 | 0,95 | -0,05 | 2,9E-01 |
| Q9R1P0 | Proteasome subunit alpha type-4 OS=Mus musculus OX=10090 | 21 | 6 | 19 | 6 | 261 | 29,5 | 7,72 | 7459 | 16,86 | 3 | 4 | 1,013 | 0,975 | 0,985 | 1,036 | 0,896 | 1,04 | 0,956 | 0,965 | 1,04 | 0,04 | 2,9E-01 |
| Q91VU0 | Protein FAM3C OS=Mus musculus OX=10090 GN=Fam3c PE=1 S | 13 | 3 | 4 | 3 | 227 | 24,7 | 8,29 | 1342 | 4,61 | 1 | 2 | 0,87 | 0,827 | 0,727 | 1,017 | 1,42 | 0,822 | 0,992 | 0,892 | 0,83 | -0,17 | 2,9E-01 |
| Q8BPM0 | Disheveled-associated activator of morphogenesis 1 OS=Mus mu | 6 | 5 | 5 | 5 | 1077 | 123,3 | 7,46 | 1746 | 5,4 | 3 | 2 | 0,934 | 0,974 | 0,968 | 0,976 | 0,859 | 0,9 | 1,006 | 0,935 | 1,04 | 0,04 | 2,9E-01 |
| Q7TNF0 | Double C2-like domain-containing protein alpha OS=Mus muscu | 4 | 1 | 2 | 1 | 405 | 44,6 | 7,12 | 1022 | | 1 | | 0,896 | 1,08 | 0,816 | 0,937 | 1,265 | 0,928 | 1,065 | 0,924 | 0,89 | -0,11 | 2,9E-01 |
| P56212 | cAMP-regulated phosphoprotein 19 OS=Mus musculus OX=1009 | 44 | 3 | 3 | 3 | 112 | 12,3 | 9,09 | 1330 | 3,29 | 2 | 1 | 1,401 | 1,188 | 1,175 | 1,381 | 0,868 | 1,446 | 1,136 | 1,071 | 1,14 | 0,14 | 2,9E-01 |
| P60843 | Eukaryotic initiation factor 4A-I OS=Mus musculus OX=10090 GN | 22 | 8 | 16 | 4 | 406 | 46,1 | 5,48 | 7923 | 5,67 | 7 | 2 | 1,044 | 0,91 | 1,136 | 1,072 | 0,839 | 1,093 | 0,935 | 0,966 | 1,09 | 0,09 | 2,9E-01 |
| Q80XA6 | RalBP1-associated Eps domain-containing protein 2 OS=Mus mu | 4 | 2 | 2 | 1 | 521 | 57,7 | 7,53 | 1023 | | 2 | | 0,943 | 1,097 | 0,945 | 1,177 | 0,998 | 1,084 | 0,764 | 0,898 | 1,11 | 0,11 | 2,9E-01 |
| Q91W43 | Glycine dehydrogenase (decarboxylating), mitochondrial OS=Mu | 4 | 3 | 3 | 3 | 1025 | 113,2 | 7,56 | 1146 | | 3 | | 1,153 | 1,12 | 0,993 | 1,086 | 1,227 | 1,092 | 1,038 | 1,471 | 0,90 | -0,10 | 2,9E-01 |
| Q9D8X2 | Coiled-coil domain-containing protein 124 OS=Mus musculus OX | 10 | 2 | 2 | 2 | 217 | 25,3 | 9,64 | 1357 | | 2 | | 0,731 | 0,841 | 0,924 | 0,839 | 0,749 | 0,906 | 0,727 | 0,631 | 1,11 | 0,11 | 2,9E-01 |
| Q9DBS2 | Tumor protein p63-regulated gene 1-like protein OS=Mus muscu | 6 | 2 | 2 | 2 | 266 | 29,8 | 7,37 | 1115 | | 2 | | 1,038 | 0,87 | 1,225 | 1,006 | 0,777 | 1,626 | 1,11 | 1,693 | 0,80 | -0,20 | 2,9E-01 |
| Q9Z1Z2 | Serine-threonine kinase receptor-associated protein OS=Mus m | 29 | 8 | 16 | 8 | 350 | 38,4 | 5,12 | 4839 | 14,18 | 6 | 5 | 1,007 | 0,983 | 0,911 | 0,958 | 0,931 | 1,035 | 0,988 | 1,084 | 0,96 | -0,04 | 2,9E-01 |
| Q9D4C9 | Clavesin-1 OS=Mus musculus OX=10090 GN=Clvs1 PE=1 SV=1 | 6 | 2 | 3 | 1 | 354 | 40,6 | 6,73 | 1419 | | 2 | | 1,418 | 1,377 | 1,206 | 1,457 | 1,296 | 1,2 | 1,386 | 1,263 | 1,06 | 0,06 | 2,9E-01 |
| Q6ZPE2 | Myotubularin-related protein 5 OS=Mus musculus OX=10090 GN | 9 | 12 | 16 | 12 | 1867 | 208,6 | 7,12 | 6777 | 5,1 | 10 | 3 | 1,046 | 1,055 | 1,052 | 0,98 | 1,146 | 0,997 | 1,089 | 1,066 | 0,96 | -0,04 | 2,9E-01 |
| Q68FD5 | Clathrin heavy chain 1 OS=Mus musculus OX=10090 GN=Cltc PE | 37 | 55 | 124 | 55 | 1675 | 191,4 | 5,69 | 50387 | 122,57 | 42 | 24 | 0,959 | 0,975 | 0,984 | 0,94 | 0,968 | 1,033 | 0,988 | 0,958 | 0,98 | -0,02 | 2,9E-01 |
| Q8VEK3 | Heterogeneous nuclear ribonucleoprotein U OS=Mus musculus | 21 | 18 | 21 | 18 | 800 | 87,9 | 6,24 | 6909 | 19,97 | 9 | 9 | 0,864 | 0,775 | 1,043 | 0,986 | 0,956 | 0,952 | 1,053 | 1,007 | 0,92 | -0,08 | 2,9E-01 |
| Q9Z2Q5 | 39S ribosomal protein L40, mitochondrial OS=Mus musculus OX | 14 | 2 | 2 | 2 | 206 | 24,3 | 9,47 | | 2,44 | | 2 | 0,626 | 0,696 | 0,765 | 0,824 | 0,792 | 0,65 | 0,893 | 0,923 | 0,89 | -0,11 | 2,9E-01 |
| P10107 | Annexin A1 OS=Mus musculus OX=10090 GN=Anxa1 PE=1 SV=2 | 3 | 1 | 1 | 1 | 346 | 38,7 | 7,37 | 340 | | 1 | | 1,005 | 0,84 | 0,924 | 0,55 | 2,073 | 0,784 | 1,01 | 0,901 | 0,70 | -0,30 | 2,9E-01 |
| P56593 | Cytochrome P450 2A12 OS=Mus musculus OX=10090 GN=Cyp2a | 2 | 1 | 3 | 1 | 492 | 56,1 | 9,19 | 1588 | | 1 | | 1,587 | 1,459 | 1,545 | 1,579 | 1,991 | 1,316 | 1,79 | 1,743 | 0,90 | -0,10 | 2,9E-01 |
| A2BH40 | AT-rich interactive domain-containing protein 1A OS=Mus musc | 1 | 1 | 1 | 1 | 2283 | 241,9 | 6,68 | 59 | | 1 | | 0,636 | 0,61 | 1,092 | 0,587 | 1,965 | 0,697 | 0,863 | 0,86 | 0,67 | -0,33 | 2,9E-01 |
| Q32M21 | Gasdermin-A2 OS=Mus musculus OX=10090 GN=Gsdma2 PE=2 S | 3 | 1 | 1 | 1 | 443 | 49,8 | 5,9 | | 2,66 | | 1 | 0,866 | 0,959 | 1,089 | 1,013 | 1,613 | 0,895 | 1,026 | 1,146 | 0,84 | -0,16 | 2,9E-01 |
| P04925 | Major prion protein OS=Mus musculus OX=10090 GN=Prnp PE= | 12 | 3 | 4 | 3 | 254 | 28 | 9,33 | 2894 | | 3 | | 1,098 | 1,03 | 1,006 | 1,039 | 0,954 | 1,083 | 1,022 | 0,932 | 1,05 | 0,05 | 2,9E-01 |
| Q7TNG5 | Echinoderm microtubule-associated protein-like 2 OS=Mus mus | 6 | 3 | 6 | 3 | 649 | 70,7 | 6,28 | 1094 | 2,57 | 2 | 1 | 1,063 | 1,063 | 0,981 | 1,106 | 1,408 | 1,032 | 1,184 | 1,022 | 0,91 | -0,09 | 2,9E-01 |
| Q8CC88 | von Willebrand factor A domain-containing protein 8 OS=Mus m | 1 | 2 | 2 | 1 | 1905 | 213,3 | 6,6 | 1368 | | 2 | | 0,909 | 1,001 | 0,874 | 1,065 | 1,298 | 1,072 | 0,921 | 0,987 | 0,90 | -0,10 | 2,9E-01 |
| Q64511 | DNA topoisomerase 2-beta OS=Mus musculus OX=10090 GN=To | 3 | 6 | 6 | 6 | 1612 | 181,8 | 8,29 | 2529 | 4,73 | 4 | 2 | 1,092 | 0,951 | 1,048 | 1,19 | 1,194 | 0,979 | 1,258 | 1,217 | 0,92 | -0,08 | 2,9E-01 |
| Q80TL7 | Protein MON2 homolog OS=Mus musculus OX=10090 GN=Mon2 | 1 | 1 | 1 | 1 | 1715 | 189 | 6,13 | 462 | | 1 | | 1,127 | 1,089 | 1,207 | 1,156 | 1,54 | 1,026 | 1,302 | 1,215 | 0,90 | -0,10 | 2,9E-01 |
| P99029 | Peroxiredoxin-5, mitochondrial OS=Mus musculus OX=10090 GN | 41 | 10 | 20 | 10 | 210 | 21,9 | 8,85 | 7490 | 21,25 | 4 | 7 | 0,971 | 0,933 | 1,094 | 1,033 | 1,158 | 1,11 | 0,995 | 1,011 | 0,94 | -0,06 | 2,9E-01 |
| Q924T2 | 28S ribosomal protein S2, mitochondrial OS=Mus musculus OX= | 2 | 1 | 1 | 1 | 291 | 32,3 | 9,14 | 351 | | 1 | | 0,667 | 0,908 | 0,85 | 0,64 | 0,867 | 0,782 | 0,799 | 0,995 | 0,89 | -0,11 | 2,9E-01 |
| Q922J9 | Fatty acyl-CoA reductase 1 OS=Mus musculus OX=10090 GN=Fa | 2 | 1 | 1 | 1 | 515 | 59,4 | 9,19 | 433 | | 1 | | 0,882 | 1,483 | 0,963 | 1,412 | 1,896 | 1,201 | 1,232 | 1,431 | 0,82 | -0,18 | 2,9E-01 |
| Q3UVX5 | Metabotropic glutamate receptor 5 OS=Mus musculus OX=1009 | 11 | 11 | 14 | 11 | 1203 | 131,8 | 7,84 | 6557 | 8,3 | 7 | 4 | 1,046 | 0,904 | 1,089 | 0,995 | 1,01 | 0,952 | 0,965 | 0,893 | 1,06 | 0,06 | 2,9E-01 |
| O55142 | 60S ribosomal protein L35a OS=Mus musculus OX=10090 GN=R | 21 | 3 | 5 | 3 | 110 | 12,5 | 10,89 | 2332 | 2,36 | 2 | 1 | 0,934 | 0,996 | 0,979 | 0,995 | 1,123 | 0,967 | 1,093 | 0,94 | 0,95 | -0,05 | 2,9E-01 |
| Q9D8E6 | 60S ribosomal protein L4 OS=Mus musculus OX=10090 GN=Rpl4 | 27 | 12 | 19 | 12 | 419 | 47,1 | 11 | 4559 | 32,23 | 4 | 8 | 0,982 | 0,889 | 0,988 | 0,965 | 1,002 | 1,013 | 0,966 | 0,963 | 0,97 | -0,03 | 2,9E-01 |
| Q99KI0 | Aconitate hydratase, mitochondrial OS=Mus musculus OX=1009 | 39 | 29 | 67 | 29 | 780 | 85,4 | 7,93 | 23151 | 78,48 | 20 | 14 | 0,976 | 1 | 0,974 | 0,975 | 0,975 | 0,979 | 0,93 | 0,979 | 1,02 | 0,02 | 2,9E-01 |
| P57746 | V-type proton ATPase subunit D OS=Mus musculus OX=10090 G | 25 | 7 | 13 | 7 | 247 | 28,4 | 9,45 | 7143 | 2,73 | 6 | 1 | 1,082 | 1,015 | 1,035 | 0,991 | 0,896 | 1,055 | 0,935 | 1,038 | 1,05 | 0,05 | 2,9E-01 |
| Q9CQJ8 | NADH dehydrogenase [ubiquinone] 1 beta subcomplex subunit | 21 | 3 | 4 | 3 | 179 | 22 | 7,8 | 2218 | | 3 | | 0,998 | 0,719 | 1,063 | 0,896 | 1,667 | 1,039 | 0,89 | 0,967 | 0,81 | -0,19 | 2,9E-01 |
| P61759 | Prefoldin subunit 3 OS=Mus musculus OX=10090 GN=Vbp1 PE= | 17 | 3 | 3 | 3 | 196 | 22,4 | 6,28 | | 8,94 | | 3 | 0,877 | 1,01 | 0,993 | 1,036 | 0,554 | 1,015 | 0,967 | 0,877 | 1,15 | 0,15 | 2,9E-01 |
| Q2NL51 | Glycogen synthase kinase-3 alpha OS=Mus musculus OX=10090 | 26 | 8 | 11 | 4 | 490 | 51,6 | 8,81 | 2889 | 10,57 | 5 | 3 | 0,855 | 0,709 | 1,025 | 0,752 | 1,092 | 0,923 | 0,861 | 0,87 | 0,89 | -0,11 | 3,0E-01 |
| Q9Z218 | Dipeptidyl aminopeptidase-like protein 6 OS=Mus musculus OX= | 21 | 14 | 22 | 13 | 804 | 91,2 | 6,68 | 8730 | 19,87 | 10 | 5 | 0,993 | 0,988 | 1,061 | 0,974 | 1,238 | 1,067 | 1,009 | 0,98 | 0,94 | -0,06 | 3,0E-01 |
| O35405 | Phospholipase D3 OS=Mus musculus OX=10090 GN=Pld3 PE=1 S | 10 | 5 | 8 | 5 | 488 | 54,4 | 6,52 | 3559 | 8,01 | 3 | 2 | 0,916 | 0,988 | 1,06 | 1,17 | 0,923 | 1,022 | 0,864 | 1,021 | 1,08 | 0,08 | 3,0E-01 |
| Q99KK9 | Probable histidine--tRNA ligase, mitochondrial OS=Mus musculu | 10 | 6 | 8 | 4 | 505 | 56,9 | 8,32 | 2718 | 6,61 | 3 | 3 | 1,008 | 1,133 | 1,021 | 1,044 | 1,164 | 0,996 | 1,076 | 1,256 | 0,94 | -0,06 | 3,0E-01 |
| Q64332 | Synapsin-2 OS=Mus musculus OX=10090 GN=Syn2 PE=1 SV=2 | 28 | 12 | 17 | 8 | 586 | 63,3 | 8,43 | 8361 | 12,2 | 10 | 3 | 0,957 | 0,951 | 1,029 | 1,015 | 1,434 | 0,927 | 0,995 | 1,116 | 0,88 | -0,12 | 3,0E-01 |
| P57716 | Nicastrin OS=Mus musculus OX=10090 GN=Ncstn PE=1 SV=3 | 2 | 1 | 1 | 1 | 708 | 78,4 | 6,09 | | 2,55 | | 1 | 0,958 | 0,897 | 0,803 | 1,11 | 1,198 | 0,807 | 1,09 | 1,178 | 0,88 | -0,12 | 3,0E-01 |
| Q9JLB0-2 | Isoform Alpha of MAGUK p55 subfamily member 6 OS=Mus mus | 22 | 11 | 12 | 10 | 539 | 60,9 | 6,18 | 5301 | 7,5 | 7 | 4 | 1,06 | 1,069 | 1,04 | 1,041 | 1,005 | 0,992 | 1,018 | 1,09 | 1,03 | 0,03 | 3,0E-01 |
| Q56A07 | Sodium channel subunit beta-2 OS=Mus musculus OX=10090 GN | 10 | 2 | 2 | 2 | 215 | 24,2 | 6,54 | 842 | | 2 | | 0,754 | 0,813 | 1,364 | 1,077 | 0,989 | 0,853 | 0,701 | 0,769 | 1,21 | 0,21 | 3,0E-01 |
| P22005 | Proenkephalin-A OS=Mus musculus OX=10090 GN=Penk PE=1 S | 6 | 1 | 1 | 1 | 268 | 31 | 5,68 | 679 | | 1 | | 0,827 | 0,942 | 0,713 | 0,597 | 0,64 | 0,638 | 0,787 | 0,636 | 1,14 | 0,14 | 3,0E-01 |
| Q9Z2H2 | Regulator of G-protein signaling 6 OS=Mus musculus OX=10090 | 11 | 5 | 7 | 3 | 472 | 54,5 | 7,42 | 1634 | 7,47 | 3 | 3 | 0,876 | 0,88 | 0,863 | 0,879 | 0,824 | 1 | 0,942 | 0,901 | 0,95 | -0,05 | 3,0E-01 |
| P47962 | 60S ribosomal protein L5 OS=Mus musculus OX=10090 GN=Rpl5 | 44 | 13 | 17 | 13 | 297 | 34,4 | 9,77 | 6322 | 16,94 | 9 | 4 | 1,047 | 0,961 | 1,02 | 1,135 | 1,005 | 1,033 | 0,924 | 1,005 | 1,05 | 0,05 | 3,0E-01 |
| Q61687 | Transcriptional regulator ATRX OS=Mus musculus OX=10090 GN | 1 | 1 | 1 | 1 | 2476 | 278,4 | 6,68 | 164 | | 1 | | 0,759 | 0,645 | 0,819 | 0,929 | 0,658 | 0,949 | 1,104 | 0,941 | 0,86 | -0,14 | 3,0E-01 |
| P30416 | Peptidyl-prolyl cis-trans isomerase FKBP4 OS=Mus musculus OX | 20 | 7 | 9 | 7 | 458 | 51,5 | 5,72 | 2736 | 10,47 | 4 | 3 | 0,951 | 0,829 | 0,859 | 0,999 | 0,681 | 0,897 | 0,882 | 0,883 | 1,09 | 0,09 | 3,0E-01 |
| P05784 | Keratin, type I cytoskeletal 18 OS=Mus musculus OX=10090 GN= | 22 | 7 | 30 | 6 | 423 | 47,5 | 5,33 | 16019 | 6,89 | 6 | 1 | 2,735 | 2,426 | 2,031 | 2,211 | 4,482 | 2,077 | 2,17 | 3,328 | 0,78 | -0,22 | 3,0E-01 |
| Q9JJY3 | Sphingomyelin phosphodiesterase 3 OS=Mus musculus OX=1009 | 10 | 6 | 6 | 6 | 655 | 71,2 | 5,88 | 1616 | 5,12 | 3 | 3 | 1,168 | 0,984 | 1,132 | 1,057 | 1,094 | 1,032 | 1,016 | 0,988 | 1,05 | 0,05 | 3,0E-01 |
| Q8VHJ5 | Serine/threonine-protein kinase MARK1 OS=Mus musculus OX= | 9 | 7 | 9 | 3 | 795 | 88,3 | 9,39 | 4712 | 2,68 | 6 | 1 | 1,194 | 1,085 | 0,964 | 1,105 | 1,419 | 1,232 | 0,904 | 1,353 | 0,89 | -0,11 | 3,0E-01 |
| P70398 | Probable ubiquitin carboxyl-terminal hydrolase FAF-X OS=Mus n | 6 | 15 | 19 | 14 | 2559 | 290,5 | 5,87 | 9138 | 5,92 | 11 | 4 | 1,025 | 0,972 | 1,033 | 0,958 | 0,863 | 1,053 | 0,965 | 0,901 | 1,05 | 0,05 | 3,0E-01 |
| P45591 | Cofilin-2 OS=Mus musculus OX=10090 GN=Cfl2 PE=1 SV=1 | 36 | 8 | 14 | 7 | 166 | 18,7 | 7,88 | 6670 | 2,34 | 6 | 2 | 1,024 | 0,972 | 0,883 | 1,148 | 0,744 | 1,052 | 0,922 | 0,929 | 1,10 | 0,10 | 3,0E-01 |
| O88983 | Syntaxin-8 OS=Mus musculus OX=10090 GN=Stx8 PE=1 SV=1 | 4 | 1 | 1 | 1 | 236 | 26,9 | 5,01 | 445 | | 1 | | 1,176 | 1,257 | 1,278 | 1,568 | 1,888 | 1,35 | 1,402 | 1,346 | 0,88 | -0,12 | 3,0E-01 |
| Q3UMB5 | Guanine nucleotide exchange protein SMCR8 OS=Mus musculus | 1 | 1 | 1 | 1 | 935 | 104,9 | 5,4 | | 2,25 | | 1 | 1,337 | 1,013 | 1,298 | 1,21 | 1,29 | 1,223 | 1,246 | 1,754 | 0,88 | -0,12 | 3,0E-01 |
| Q5FWI3 | Cell surface hyaluronidase OS=Mus musculus OX=10090 GN=Cel | 1 | 1 | 1 | 1 | 1383 | 153,7 | 7,78 | 414 | | 1 | | 1,003 | 1,288 | 0,86 | 1,21 | 1,005 | 1,057 | 1,786 | 1,448 | 0,82 | -0,18 | 3,0E-01 |
| O35551 | Rab GTPase-binding effector protein 1 OS=Mus musculus OX=10 | 15 | 13 | 13 | 13 | 862 | 99,5 | 5,01 | 4829 | 7,93 | 9 | 4 | 1,018 | 1,009 | 0,961 | 0,982 | 1,121 | 0,912 | 1,046 | 1,12 | 0,95 | -0,05 | 3,0E-01 |
| Q62261 | Spectrin beta chain, non-erythrocytic 1 OS=Mus musculus OX=1 | 47 | 112 | 206 | 102 | 2363 | 274,1 | 5,58 | 82036 | 187,17 | 73 | 48 | 0,988 | 0,964 | 1,006 | 0,962 | 0,895 | 0,951 | 1 | 0,965 | 1,03 | 0,03 | 3,0E-01 |
| Q3UHD3 | Microtubule-associated tumor suppressor candidate 2 homolog | 1 | 1 | 1 | 1 | 1353 | 147,3 | 8,24 | | 2,49 | | 1 | 1,086 | 1,169 | 1,028 | 1,155 | 1,349 | 0,871 | 0,7 | 0,874 | 1,17 | 0,17 | 3,0E-01 |
| P70207 | Plexin-A2 OS=Mus musculus OX=10090 GN=Plxna2 PE=1 SV=2 | 6 | 13 | 18 | 5 | 1894 | 211,4 | 6,54 | 6179 | 13,78 | 9 | 5 | 0,877 | 1,018 | 0,637 | 0,74 | 1,808 | 0,619 | 1,063 | 0,966 | 0,73 | -0,27 | 3,0E-01 |
| Q7TQD2 | Tubulin polymerization-promoting protein OS=Mus musculus O | 18 | 4 | 5 | 4 | 218 | 23,6 | 9,42 | 1667 | 1,95 | 2 | 2 | 1,093 | 0,992 | 0,989 | 0,796 | 0,656 | 0,907 | 0,944 | 0,945 | 1,12 | 0,12 | 3,1E-01 |

| | | | | | | | | | | | | | | | | | | | | | | | |
|---|---|---|---|---|---|---|---|---|---|---|---|---|---|---|---|---|---|---|---|---|---|---|---|
| Q9WTX5 | S-phase kinase-associated protein 1 OS=Mus musculus OX=1009 | 21 | 4 | 4 | 4 | 163 | 18,7 | 4,54 | 1373 | 2,61 | 3 | 1 | 0,951 | 0,895 | 0,946 | 0,859 | 0,679 | 0,945 | 0,812 | 0,924 | 1,09 | 0,09 | 3,1E-01 |
| O55143 | Sarcoplasmic/endoplasmic reticulum calcium ATPase 2 OS=Mus | 27 | 26 | 45 | 21 | 1044 | 114,8 | 5,34 | 16141 | 41,38 | 17 | 11 | 1,001 | 1,015 | 1,076 | 0,981 | 0,987 | 1,061 | 1,11 | 1,061 | 0,97 | -0,03 | 3,1E-01 |
| P28740-1 | Isoform 1 of Kinesin-like protein KIF2A OS=Mus musculus OX=10 | 24 | 16 | 23 | 16 | 716 | 80,8 | 6,16 | 9333 | 15,31 | 11 | 6 | 1,072 | 0,958 | 1,185 | 0,937 | 0,724 | 1,072 | 0,993 | 0,942 | 1,11 | 0,11 | 3,1E-01 |
| Q9WTL4 | Insulin receptor-related protein OS=Mus musculus OX=10090 GI | 1 | 1 | 1 | 1 | 1300 | 144,8 | 6,64 | 702 | | 1 | | 0,959 | 0,842 | 0,822 | 0,955 | 1,421 | 0,913 | 0,828 | 1,025 | 0,85 | -0,15 | 3,1E-01 |
| Q3UIJ9 | Myocardial zonula adherens protein OS=Mus musculus OX=1009 | 2 | 1 | 1 | 1 | 466 | 53,9 | 6,51 | 315 | | 1 | | 1,058 | 0,915 | 0,99 | 0,787 | 0,936 | 0,727 | 0,958 | 0,756 | 1,11 | 0,11 | 3,1E-01 |
| Q01341 | Adenylate cyclase type 6 OS=Mus musculus OX=10090 GN=Adcy | 1 | 1 | 1 | 1 | 1165 | 130,2 | 7,87 | 331 | | 1 | | 0,928 | 1,233 | 1,182 | 0,775 | 1,8 | 1,136 | 1,043 | 1,077 | 0,81 | -0,19 | 3,1E-01 |
| P46471 | 26S proteasome regulatory subunit 7 OS=Mus musculus OX=100 | 29 | 12 | 18 | 12 | 433 | 48,6 | 5,95 | 7836 | 14,92 | 7 | 5 | 0,968 | 0,992 | 0,975 | 0,938 | 0,983 | 0,947 | 1,026 | 1,009 | 0,98 | -0,02 | 3,1E-01 |
| P63280 | SUMO-conjugating enzyme UBC9 OS=Mus musculus OX=10090 | 8 | 1 | 2 | 1 | 158 | 18 | 8,66 | 1203 | | 1 | | 1,14 | 1,003 | 0,9 | 1,336 | 0,982 | 0,898 | 1,012 | 1,043 | 1,11 | 0,11 | 3,1E-01 |
| P14115 | 60S ribosomal protein L27a OS=Mus musculus OX=10090 GN=R | 28 | 5 | 9 | 5 | 148 | 16,6 | 11,12 | 5156 | 5,09 | 3 | 2 | 1,078 | 1,063 | 0,929 | 1,191 | 0,957 | 1,237 | 1,179 | 1,298 | 0,91 | -0,09 | 3,1E-01 |
| O55126 | Protein NipSnap homolog 2 OS=Mus musculus OX=10090 GN=N | 21 | 5 | 5 | 5 | 281 | 32,9 | 9,26 | 2162 | 6,57 | 3 | 2 | 1,108 | 0,916 | 1,332 | 1,109 | 0,91 | 1,145 | 0,994 | 0,976 | 1,11 | 0,11 | 3,1E-01 |
| Q3UKC1 | Tax1-binding protein 1 homolog OS=Mus musculus OX=10090 G | 1 | 1 | 1 | 1 | 814 | 93,6 | 5,33 | 523 | | 1 | | 0,871 | 1,042 | 0,905 | 0,859 | 0,999 | 0,787 | 1,153 | 1,186 | 0,89 | -0,11 | 3,1E-01 |
| P06801 | NADP-dependent malic enzyme OS=Mus musculus OX=10090 G | 19 | 10 | 16 | 10 | 572 | 63,9 | 7,44 | 6293 | 9,28 | 8 | 3 | 0,999 | 0,954 | 0,895 | 0,917 | 0,678 | 0,932 | 0,919 | 0,937 | 1,09 | 0,09 | 3,1E-01 |
| P12367 | cAMP-dependent protein kinase type II-alpha regulatory subuni | 19 | 8 | 10 | 4 | 401 | 45,4 | 4,89 | 4292 | 9,3 | 5 | 3 | 1,037 | 0,97 | 1,086 | 1,047 | 0,695 | 1,069 | 1,048 | 0,93 | 1,11 | 0,11 | 3,1E-01 |
| P07310 | Creatine kinase M-type OS=Mus musculus OX=10090 GN=Ckm F | 10 | 4 | 8 | 1 | 381 | 43 | 7,06 | 3212 | 2,41 | 3 | 1 | 0,916 | 0,783 | 0,743 | 0,798 | 6,412 | 1,075 | 0,984 | 0,844 | 0,35 | -0,65 | 3,1E-01 |
| Q8C8N2 | Protein SCAI OS=Mus musculus OX=10090 GN=Scai PE=1 SV=2 | 2 | 1 | 1 | 1 | 606 | 70,2 | 8,6 | 815 | | 1 | | 0,942 | 1,054 | 1,005 | 0,808 | 0,728 | 0,895 | 0,974 | 0,882 | 1,09 | 0,09 | 3,1E-01 |
| P80560 | Receptor-type tyrosine-protein phosphatase N2 OS=Mus muscu | 2 | 2 | 2 | 2 | 1001 | 111,4 | 5,95 | 985 | | 2 | | 0,968 | 0,808 | 0,91 | 0,97 | 0,956 | 0,947 | 0,911 | 1,071 | 0,94 | -0,06 | 3,1E-01 |
| Q9ES56 | Trafficking protein particle complex subunit 4 OS=Mus musculus | 4 | 1 | 1 | 1 | 219 | 24,4 | 6,21 | 588 | | 1 | | 1,129 | 0,889 | 0,978 | 1,107 | 1,064 | 0,951 | 0,823 | 0,931 | 1,09 | 0,09 | 3,1E-01 |
| Q9JIP4 | Pannexin-1 OS=Mus musculus OX=10090 GN=Panx1 PE=1 SV=3 | 2 | 1 | 1 | 1 | 426 | 48,1 | 7,03 | 732 | | 1 | | 1,149 | 1,076 | 1,061 | 1,14 | 1,056 | 0,94 | 1,118 | 1,106 | 1,05 | 0,05 | 3,1E-01 |
| P11499 | Heat shock protein HSP 90-beta OS=Mus musculus OX=10090 G | 49 | 41 | 118 | 23 | 724 | 83,2 | 5,03 | 48759 | 97,89 | 29 | 18 | 1,007 | 0,934 | 1,029 | 1,08 | 0,816 | 0,986 | 1,006 | 0,999 | 1,06 | 0,06 | 3,1E-01 |
| P63042 | Stathmin-4 OS=Mus musculus OX=10090 GN=Stmn4 PE=2 SV=1 | 10 | 3 | 4 | 1 | 189 | 22,1 | 5,99 | | 8,18 | | 3 | 1,149 | 1,015 | 1,425 | 0,939 | 0,921 | 1,194 | 0,909 | 0,941 | 1,14 | 0,14 | 3,1E-01 |
| Q8C863 | E3 ubiquitin-protein ligase Itchy OS=Mus musculus OX=10090 G | 1 | 1 | 1 | 1 | 864 | 98,9 | 6,28 | 353 | | 1 | | 1,225 | 0,954 | 0,828 | 0,759 | 0,613 | 0,797 | 0,84 | 0,96 | 1,17 | 0,17 | 3,1E-01 |
| Q9Z188 | Dual specificity tyrosine-phosphorylation-regulated kinase 1B O | 2 | 1 | 1 | 1 | 629 | 69,1 | 9,09 | | 2,87 | | 1 | 1,125 | 0,906 | 1,075 | 1,094 | 1,056 | 0,815 | 0,689 | 1,133 | 1,14 | 0,14 | 3,1E-01 |
| Q01149 | Collagen alpha-2(I) chain OS=Mus musculus OX=10090 GN=Col1 | 1 | 1 | 1 | 1 | 1372 | 129,5 | 9,19 | 505 | | 1 | | 0,831 | 0,988 | 0,868 | 0,558 | 1,95 | 0,925 | 0,687 | 0,981 | 0,71 | -0,29 | 3,1E-01 |
| Q80XR2 | Calcium-transporting ATPase type 2C member 1 OS=Mus muscu | 3 | 2 | 3 | 2 | 918 | 100,2 | 6,83 | | 8,58 | | 2 | 1,078 | 1,051 | 0,974 | 0,875 | 1,241 | 1,087 | 0,867 | 1,208 | 0,90 | -0,10 | 3,1E-01 |
| O88545 | COP9 signalosome complex subunit 6 OS=Mus musculus OX=10 | 8 | 2 | 3 | 2 | 324 | 35,9 | 5,73 | 830 | 5,04 | 1 | 1 | 0,905 | 0,792 | 0,857 | 0,767 | 1,161 | 0,775 | 0,898 | 0,876 | 0,90 | -0,10 | 3,1E-01 |
| P47941 | Crk-like protein OS=Mus musculus OX=10090 GN=Crkl PE=1 SV= | 8 | 2 | 4 | 2 | 303 | 33,8 | 6,74 | 1899 | | 2 | | 1,07 | 0,984 | 0,996 | 1,003 | 1,218 | 1,169 | 0,969 | 0,988 | 0,93 | -0,07 | 3,1E-01 |
| P53612 | Geranylgeranyl transferase type-2 subunit beta OS=Mus muscul | 6 | 2 | 2 | 2 | 339 | 37,8 | 5,16 | 1430 | | 2 | | 0,935 | 0,966 | 0,939 | 1,017 | 0,975 | 0,916 | 1,192 | 1,05 | 0,93 | -0,07 | 3,1E-01 |
| Q9Z1X4 | Interleukin enhancer-binding factor 3 OS=Mus musculus OX=10 | 8 | 7 | 9 | 7 | 898 | 96 | 8,76 | 4085 | 3,4 | 6 | 1 | 0,859 | 0,91 | 0,916 | 1,022 | 0,983 | 0,872 | 1,104 | 1,007 | 0,93 | -0,07 | 3,1E-01 |
| P47743 | Metabotropic glutamate receptor 8 OS=Mus musculus OX=1009 | 2 | 2 | 2 | 1 | 908 | 101,8 | 8,15 | 586 | 2,21 | 1 | 1 | 1,005 | 1,332 | 1,058 | 1,438 | 1,527 | 1,248 | 1,428 | 1,202 | 0,89 | -0,11 | 3,1E-01 |
| P31001 | Desmin OS=Mus musculus OX=10090 GN=Des PE=1 SV=3 | 8 | 4 | 6 | 1 | 469 | 53,5 | 5,27 | 2612 | 2,66 | 3 | 1 | 0,685 | 0,572 | 0,821 | 0,603 | 5,945 | 0,943 | 0,967 | 0,512 | 0,32 | -0,68 | 3,1E-01 |
| Q91VM5 | RNA binding motif protein, X-linked-like-1 OS=Mus musculus OX | 19 | 8 | 10 | 8 | 388 | 42,1 | 9,99 | 3305 | 4,95 | 5 | 3 | 0,887 | 0,817 | 0,892 | 1,001 | 0,998 | 0,817 | 1,063 | 1,008 | 0,93 | -0,07 | 3,1E-01 |
| Q8R464 | Cell adhesion molecule 4 OS=Mus musculus OX=10090 GN=Cadr | 24 | 6 | 16 | 6 | 388 | 42,7 | 6,3 | 3285 | 31,31 | 3 | 5 | 0,934 | 1,052 | 0,997 | 1,023 | 1,183 | 0,933 | 1,055 | 1,088 | 0,94 | -0,06 | 3,1E-01 |
| Q9DC63 | F-box only protein 3 OS=Mus musculus OX=10090 GN=Fbxo3 PE | 8 | 2 | 2 | 2 | 480 | 55,2 | 5,02 | 1001 | | 2 | | 0,865 | 1,051 | 0,691 | 0,859 | 1,215 | 0,785 | 1,038 | 0,94 | 0,87 | -0,13 | 3,1E-01 |
| Q8VEM8 | Phosphate carrier protein, mitochondrial OS=Mus musculus OX= | 23 | 9 | 15 | 9 | 357 | 39,6 | 9,26 | 3650 | 25,09 | 4 | 5 | 1,054 | 1,037 | 1,031 | 1,061 | 0,845 | 0,977 | 0,997 | 1,117 | 1,06 | 0,06 | 3,1E-01 |
| Q9WU60 | Attractin OS=Mus musculus OX=10090 GN=Atrn PE=1 SV=3 | 1 | 1 | 2 | 1 | 1428 | 158 | 7,27 | 968 | | 1 | | 1,072 | 1,132 | 1,006 | 0,794 | 1,213 | 0,926 | 1,234 | 1,081 | 0,90 | -0,10 | 3,1E-01 |
| Q6RT24 | Centromere-associated protein E OS=Mus musculus OX=10090 | 0 | 1 | 1 | 1 | 2474 | 286,3 | 5,31 | 403 | | 1 | | 1,189 | 1,462 | 0,965 | 1,548 | 0,923 | 1,245 | 1,1 | 1,227 | 1,15 | 0,15 | 3,1E-01 |
| O08749 | Dihydrolipoyl dehydrogenase, mitochondrial OS=Mus musculus | 28 | 15 | 32 | 15 | 509 | 54,2 | 7,9 | 11749 | 32,11 | 8 | 7 | 1,001 | 0,945 | 0,946 | 0,937 | 0,87 | 0,998 | 0,906 | 0,92 | 1,04 | 0,04 | 3,1E-01 |
| Q8K2K6 | Arf-GAP domain and FG repeat-containing protein 1 OS=Mus mu | 11 | 5 | 9 | 5 | 561 | 58 | 8,63 | 4022 | 6,96 | 3 | 2 | 1,025 | 0,883 | 0,987 | 1,083 | 0,669 | 0,924 | 1,013 | 0,982 | 1,11 | 0,11 | 3,1E-01 |
| Q9EP89 | Serine beta-lactamase-like protein LACTB, mitochondrial OS=Mu | 2 | 1 | 1 | 1 | 551 | 60,7 | 8,9 | 333 | | 1 | | 0,976 | 1,064 | 1,183 | 1,045 | 1,164 | 1,076 | 1,04 | 1,32 | 0,93 | -0,07 | 3,2E-01 |
| Q9CQF0 | 39S ribosomal protein L11, mitochondrial OS=Mus musculus OX | 34 | 5 | 6 | 5 | 192 | 20,7 | 9,73 | 2956 | | 5 | | 0,886 | 0,991 | 1,17 | 0,879 | 1,493 | 1,073 | 0,97 | 1 | 0,87 | -0,13 | 3,2E-01 |
| Q9Z1S5 | Neuronal-specific septin-3 OS=Mus musculus OX=10090 GN=Se | 25 | 10 | 17 | 9 | 350 | 40 | 6,81 | 8246 | 14,32 | 6 | 4 | 1,06 | 0,878 | 1,174 | 0,945 | 0,896 | 1,036 | 0,899 | 0,904 | 1,09 | 0,09 | 3,2E-01 |
| Q8K0S0 | Phytanoyl-CoA hydroxylase-interacting protein OS=Mus muscul | 4 | 1 | 2 | 1 | 330 | 37,5 | 7,01 | 1105 | | 1 | | 1,096 | 0,798 | 1,288 | 0,925 | 0,675 | 0,974 | 0,989 | 0,906 | 1,16 | 0,16 | 3,2E-01 |
| O55023 | Inositol monophosphatase 1 OS=Mus musculus OX=10090 GN=I | 27 | 8 | 9 | 8 | 277 | 30,4 | 5,19 | 3891 | 9,49 | 5 | 3 | 1,037 | 1,019 | 0,985 | 1,158 | 0,852 | 1,106 | 0,986 | 0,975 | 1,07 | 0,07 | 3,2E-01 |
| Q5HZI2 | C2 calcium-dependent domain-containing protein 4C OS=Mus m | 3 | 1 | 1 | 1 | 419 | 44,6 | 9,73 | | 2,03 | | 1 | 0,913 | 0,796 | 0,794 | 0,895 | 1,024 | 0,76 | 0,997 | 0,911 | 0,92 | -0,08 | 3,2E-01 |
| P97461 | 40S ribosomal protein S5 OS=Mus musculus OX=10090 GN=Rps | 19 | 5 | 5 | 5 | 204 | 22,9 | 9,72 | 1942 | 4,9 | 3 | 2 | 0,91 | 0,834 | 1,005 | 1,004 | 1,047 | 0,921 | 1,072 | 0,953 | 0,94 | -0,06 | 3,2E-01 |
| P60766 | Cell division control protein 42 homolog OS=Mus musculus OX= | 36 | 7 | 15 | 1 | 191 | 21,2 | 6,55 | 5688 | 10,31 | 5 | 4 | 0,912 | 0,897 | 1,107 | 0,676 | 1,763 | 1,215 | 0,766 | 0,892 | 0,77 | -0,23 | 3,2E-01 |
| Q9DCJ1 | Target of rapamycin complex subunit LST8 OS=Mus musculus O | 9 | 2 | 2 | 2 | 326 | 35,8 | 5,86 | 1373 | | 2 | | 0,921 | 1,047 | 1,167 | 0,93 | 1,067 | 1,005 | 1,063 | 1,359 | 0,90 | -0,10 | 3,2E-01 |
| Q7TMY8 | E3 ubiquitin-protein ligase HUWE1 OS=Mus musculus OX=10090 | 2 | 8 | 8 | 8 | 4377 | 482,3 | 5,22 | 3451 | 5,5 | 6 | 2 | 0,918 | 1,005 | 1,003 | 1,048 | 1,208 | 1,001 | 0,942 | 1,104 | 0,93 | -0,07 | 3,2E-01 |
| Q6P2B1 | Transportin-3 OS=Mus musculus OX=10090 GN=Tnpo3 PE=1 SV= | 3 | 2 | 3 | 2 | 923 | 104,1 | 5,57 | 1539 | 2,47 | 1 | 1 | 1,021 | 0,91 | 1,164 | 0,891 | 0,636 | 1,036 | 0,989 | 0,849 | 1,14 | 0,14 | 3,2E-01 |
| Q64674 | Spermidine synthase OS=Mus musculus OX=10090 GN=Srm PE= | 15 | 4 | 4 | 4 | 302 | 34 | 5,5 | 2063 | 2,09 | 3 | 1 | 1,109 | 1,092 | 1,049 | 1,131 | 0,797 | 1,137 | 1,084 | 1,029 | 1,08 | 0,08 | 3,2E-01 |
| P99028 | Cytochrome b-c1 complex subunit 6, mitochondrial OS=Mus mu | 47 | 5 | 6 | 5 | 89 | 10,4 | 4,87 | 3896 | 2,64 | 4 | 1 | 0,982 | 0,998 | 0,927 | 0,969 | 0,791 | 1,039 | 0,875 | 0,935 | 1,06 | 0,06 | 3,2E-01 |
| P97450 | ATP synthase-coupling factor 6, mitochondrial OS=Mus musculu | 51 | 6 | 11 | 6 | 108 | 12,5 | 9,36 | 3665 | 14,49 | 5 | 3 | 0,869 | 0,94 | 1,032 | 1,004 | 0,855 | 0,878 | 0,956 | 0,96 | 1,05 | 0,05 | 3,2E-01 |
| P59325 | Eukaryotic translation initiation factor 5 OS=Mus musculus OX= | 15 | 5 | 6 | 5 | 429 | 48,9 | 5,52 | 2392 | 6,1 | 4 | 2 | 1,039 | 1,238 | 0,984 | 1,095 | 1,092 | 1,152 | 1,132 | 1,268 | 0,94 | -0,06 | 3,2E-01 |
| P55302 | Alpha-2-macroglobulin receptor-associated protein OS=Mus mu | 8 | 4 | 4 | 4 | 360 | 42,2 | 7,87 | 1574 | 4,59 | 2 | 2 | 0,992 | 1,035 | 1,072 | 1,2 | 0,814 | 1,036 | 1,117 | 0,993 | 1,09 | 0,09 | 3,2E-01 |
| Q3TZX8 | Polynucleotide 5'-hydroxyl-kinase NOL9 OS=Mus musculus OX= | 1 | 1 | 2 | 1 | 714 | 80,8 | 9,45 | 615 | | 1 | | 0,724 | 1,024 | 0,817 | 0,88 | 5,38 | 1,273 | 0,799 | 0,813 | 0,42 | -0,58 | 3,2E-01 |
| Q9QYY8 | Spastin OS=Mus musculus OX=10090 GN=Spast PE=1 SV=3 | 1 | 1 | 1 | 1 | 614 | 66,4 | 9,69 | | 2,85 | | 1 | 0,948 | 1,028 | 1,068 | 1,013 | 1,098 | 0,778 | 0,932 | 0,946 | 1,08 | 0,08 | 3,2E-01 |
| Q9D7G0 | Ribose-phosphate pyrophosphokinase 1 OS=Mus musculus OX= | 14 | 4 | 6 | 4 | 318 | 34,8 | 6,98 | 1733 | 5,95 | 2 | 3 | 1,01 | 1,055 | 1,096 | 1,033 | 1,47 | 1,036 | 1,014 | 1,137 | 0,90 | -0,10 | 3,2E-01 |
| Q61301 | Catenin alpha-2 OS=Mus musculus OX=10090 GN=Ctnna2 PE=1 | 32 | 27 | 42 | 21 | 953 | 105,2 | 5,71 | 19261 | 25,75 | 20 | 8 | 0,986 | 1,044 | 0,995 | 0,984 | 0,951 | 0,904 | 1,022 | 1,002 | 1,03 | 0,03 | 3,2E-01 |
| P80316 | T-complex protein 1 subunit epsilon OS=Mus musculus OX=1009 | 26 | 16 | 25 | 16 | 541 | 59,6 | 6,02 | 9703 | 20 | 10 | 8 | 1,061 | 0,852 | 1,159 | 0,933 | 0,87 | 1,015 | 0,885 | 0,909 | 1,09 | 0,09 | 3,2E-01 |
| Q60780 | Growth arrest-specific protein 7 OS=Mus musculus OX=10090 G | 7 | 4 | 4 | 3 | 421 | 48,1 | 7,83 | 852 | 8,69 | 1 | 3 | 1,046 | 0,999 | 1,215 | 1,027 | 0,626 | 1,114 | 1,022 | 1,012 | 1,14 | 0,14 | 3,2E-01 |
| Q8VCR2-2 | Isoform 2 of 17-beta-hydroxysteroid dehydrogenase 13 OS=Mu | 10 | 2 | 3 | 2 | 300 | 33,4 | 9,31 | 1433 | | 2 | | 0,955 | 0,802 | 1,236 | 0,966 | 1,356 | 0,898 | 1,124 | 1,142 | 0,88 | -0,12 | 3,2E-01 |
| P51881 | ADP/ATP translocase 2 OS=Mus musculus OX=10090 GN=Slc25a | 34 | 12 | 22 | 5 | 298 | 32,9 | 9,73 | 4419 | 37,52 | 5 | 7 | 0,911 | 0,91 | 1,052 | 0,893 | 0,993 | 0,998 | 0,946 | 0,998 | 0,96 | -0,04 | 3,2E-01 |
| Q76MZ3 | Serine/threonine-protein phosphatase 2A 65 kDa regulatory sub | 35 | 18 | 49 | 18 | 589 | 65,3 | 5,11 | 24366 | 15,18 | 16 | 7 | 1,002 | 0,926 | 1,035 | 0,987 | 0,794 | 1,047 | 0,934 | 0,931 | 1,07 | 0,07 | 3,2E-01 |
| P35279 | Ras-related protein Rab-6A OS=Mus musculus OX=10090 GN=Ra | 39 | 10 | 17 | 4 | 208 | 23,6 | 5,54 | 11483 | 3,3 | 9 | 1 | 0,994 | 1,145 | 1,201 | 1,113 | 1,168 | 1,028 | 1,028 | 0,965 | 1,06 | 0,06 | 3,2E-01 |
| Q8K010 | 5-oxoprolinase OS=Mus musculus OX=10090 GN=Oplah PE=1 SV | 5 | 6 | 6 | 6 | 1288 | 137,5 | 6,28 | 2156 | 0 | 5 | 1 | 0,892 | 1,003 | 0,887 | 0,884 | 1,434 | 0,881 | 0,957 | 0,955 | 0,87 | -0,13 | 3,2E-01 |

| | | | | | | | | | | | | | | | | | | | | | | | |
|---|---|---|---|---|---|---|---|---|---|---|---|---|---|---|---|---|---|---|---|---|---|---|---|
| O54988 | STE20-like serine/threonine-protein kinase OS=Mus musculus O | 5 | 4 | 4 | 4 | 1233 | 141,4 | 5,14 | 536 | 2,22 | 2 | 2 | 0,744 | 0,675 | 0,713 | 0,516 | 0,675 | 0,82 | 0,827 | 0,632 | 0,90 | -0,10 | 3,2E-01 |
| Q9QZ88 | Vacuolar protein sorting-associated protein 29 OS=Mus musculu | 33 | 6 | 10 | 6 | 182 | 20,5 | 6,79 | 4567 | 5,6 | 4 | 2 | 1,038 | 1,034 | 0,947 | 1,034 | 0,803 | 0,973 | 1,022 | 1,015 | 1,06 | 0,06 | 3,2E-01 |
| Q9QXL8 | Nucleoside diphosphate kinase 7 OS=Mus musculus OX=10090 ( | 2 | 1 | 1 | 1 | 395 | 44,4 | 7,12 | 460 | | 1 | | 0,859 | 1,082 | 1,079 | 1,06 | 1,283 | 0,964 | 1,04 | 1,176 | 0,91 | -0,09 | 3,2E-01 |
| P60229 | Eukaryotic translation initiation factor 3 subunit E OS=Mus musc | 4 | 2 | 3 | 2 | 445 | 52,2 | 6,04 | 1670 | | 2 | | 1,365 | 1,504 | 1,347 | 1,242 | 1,793 | 1,404 | 1,324 | 1,439 | 0,92 | -0,08 | 3,2E-01 |
| Q99MN9 | Propionyl-CoA carboxylase beta chain, mitochondrial OS=Mus m | 15 | 6 | 9 | 6 | 541 | 58,4 | 7,66 | 4406 | | 6 | | 0,984 | 1,146 | 0,96 | 1,204 | 0,957 | 0,935 | 1,067 | 1,043 | 1,07 | 0,07 | 3,2E-01 |
| Q62523 | Zyxin OS=Mus musculus OX=10090 GN=Zyx PE=1 SV=2 | 3 | 1 | 2 | 1 | 564 | 60,5 | 6,4 | 1351 | | 1 | | 1,21 | 1,124 | 1,162 | 1,204 | 1,675 | 1,064 | 1,189 | 1,345 | 0,89 | -0,11 | 3,2E-01 |
| O35129 | Prohibitin-2 OS=Mus musculus OX=10090 GN=Phb2 PE=1 SV=1 | 36 | 11 | 14 | 11 | 299 | 33,3 | 9,83 | 3460 | 23,52 | 4 | 7 | 1,064 | 0,964 | 0,964 | 0,981 | 0,815 | 0,975 | 0,945 | 1,022 | 1,06 | 0,06 | 3,3E-01 |
| Q8C996 | Transmembrane protein 163 OS=Mus musculus OX=10090 GN=1 | 18 | 3 | 4 | 3 | 288 | 31,2 | 7,93 | 1087 | 4,66 | 3 | 1 | 1,395 | 1,256 | 1,145 | 1,214 | 1,026 | 1,306 | 1,287 | 0,926 | 1,10 | 0,10 | 3,3E-01 |
| P42337 | Phosphatidylinositol 4,5-bisphosphate 3-kinase catalytic subunit | 1 | 1 | 1 | 1 | 1068 | 124,3 | 7,15 | | 0 | | 1 | 0,886 | 0,579 | 0,77 | 0,919 | 0,567 | 0,681 | 0,776 | 0,746 | 1,14 | 0,14 | 3,3E-01 |
| Q8K0T0 | Reticulon-1 OS=Mus musculus OX=10090 GN=Rtn1 PE=1 SV=1 | 20 | 12 | 25 | 12 | 780 | 83,5 | 4,58 | 10940 | 24,34 | 8 | 6 | 1,083 | 1,021 | 0,978 | 0,982 | 0,785 | 0,991 | 1,021 | 1,007 | 1,07 | 0,07 | 3,3E-01 |
| Q9DC51 | Guanine nucleotide-binding protein G(i) subunit alpha OS=Mus | 31 | 11 | 22 | 4 | 354 | 40,5 | 5,69 | 8409 | 15,79 | 7 | 4 | 0,947 | 0,884 | 0,937 | 0,96 | 0,838 | 0,987 | 0,876 | 0,871 | 1,04 | 0,04 | 3,3E-01 |
| Q9CT10 | Ran-binding protein 3 OS=Mus musculus OX=10090 GN=Ranbp3 | 4 | 2 | 2 | 2 | 491 | 52,5 | 5,12 | 970 | | 2 | | 0,895 | 0,845 | 0,844 | 0,836 | 1,11 | 0,882 | 0,807 | 0,904 | 0,92 | -0,08 | 3,3E-01 |
| O08989 | Ras-related protein M-Ras OS=Mus musculus OX=10090 GN=Mr | 18 | 3 | 4 | 3 | 208 | 23,9 | 8,78 | 739 | 8,72 | 1 | 2 | 1,045 | 0,957 | 0,988 | 1,036 | 0,979 | 1,141 | 1,034 | 1,041 | 0,96 | -0,04 | 3,3E-01 |
| Q9Z126 | Platelet factor 4 OS=Mus musculus OX=10090 GN=Pf4 PE=1 SV= | 9 | 1 | 1 | 1 | 105 | 11,2 | 9,29 | 788 | | 1 | | 0,854 | 0,631 | 0,646 | 1,099 | 4,409 | 0,836 | 0,859 | 0,927 | 0,46 | -0,54 | 3,3E-01 |
| P24638 | Lysosomal acid phosphatase OS=Mus musculus OX=10090 GN=A | 2 | 1 | 1 | 1 | 423 | 48,5 | 7,02 | 602 | | 1 | | 1,075 | 0,962 | 0,963 | 1,08 | 2,039 | 1,082 | 0,876 | 1,184 | 0,79 | -0,21 | 3,3E-01 |
| Q923T9 | Calcium/calmodulin-dependent protein kinase type II subunit ga | 16 | 9 | 30 | 2 | 529 | 59,6 | 7,58 | 12510 | 10,47 | 5 | 5 | 1,036 | 1,011 | 1,328 | 0,837 | 0,894 | 0,976 | 1,04 | 0,824 | 1,13 | 0,13 | 3,3E-01 |
| Q3TAS6 | ER membrane protein complex subunit 10 OS=Mus musculus OX | 4 | 1 | 1 | 1 | 258 | 27 | 5,91 | | 2,47 | | 1 | 0,85 | 0,817 | 0,882 | 0,903 | 0,786 | 0,878 | 0,872 | 0,774 | 1,04 | 0,04 | 3,3E-01 |
| P62843 | 40S ribosomal protein S15 OS=Mus musculus OX=10090 GN=Rp: | 9 | 2 | 2 | 2 | 145 | 17 | 10,39 | | 6,16 | | 2 | 0,964 | 0,863 | 1,039 | 0,882 | 0,711 | 0,888 | 0,934 | 0,932 | 1,08 | 0,08 | 3,3E-01 |
| Q9D898 | Actin-related protein 2/3 complex subunit 5-like protein OS=Mu | 38 | 5 | 9 | 5 | 153 | 17 | 6,8 | 2679 | 8,45 | 3 | 2 | 1,017 | 1,082 | 0,973 | 1,111 | 0,881 | 1,057 | 1,068 | 0,943 | 1,06 | 0,06 | 3,3E-01 |
| P46460 | Vesicle-fusing ATPase OS=Mus musculus OX=10090 GN=Nsf PE= | 42 | 33 | 72 | 33 | 744 | 82,6 | 6,95 | 23354 | 97,58 | 19 | 18 | 1,004 | 1,036 | 0,942 | 0,962 | 0,953 | 1,053 | 1,029 | 1,04 | 0,97 | -0,03 | 3,3E-01 |
| Q60605 | Myosin light polypeptide 6 OS=Mus musculus OX=10090 GN=My | 52 | 8 | 14 | 7 | 151 | 16,9 | 4,65 | 4928 | 17,09 | 5 | 4 | 0,904 | 0,848 | 0,899 | 0,971 | 2,228 | 0,864 | 0,902 | 1,02 | 0,72 | -0,28 | 3,3E-01 |
| A2AAJ9 | Obscurin OS=Mus musculus OX=10090 GN=Obscn PE=1 SV=3 | 1 | 2 | 2 | 2 | 8886 | 966 | 5,9 | 678 | | 2 | | 1,192 | 0,949 | 1,087 | 1,185 | 0,902 | 0,978 | 1,069 | 1,138 | 1,08 | 0,08 | 3,3E-01 |
| Q5SX39 | Myosin-4 OS=Mus musculus OX=10090 GN=Myh4 PE=2 SV=1 | 11 | 22 | 26 | 4 | 1939 | 222,7 | 5,74 | 8383 | 28,08 | 13 | 9 | 0,742 | 0,709 | 0,494 | 0,687 | 4,585 | 0,615 | 0,719 | 0,828 | 0,39 | -0,61 | 3,3E-01 |
| P63242 | Eukaryotic translation initiation factor 5A-1 OS=Mus musculus C | 30 | 5 | 11 | 5 | 154 | 16,8 | 5,24 | 3162 | 20,51 | 2 | 3 | 0,866 | 0,958 | 0,941 | 0,978 | 0,587 | 0,938 | 0,893 | 0,947 | 1,11 | 0,11 | 3,3E-01 |
| Q64105 | Sepiapterin reductase OS=Mus musculus OX=10090 GN=Spr PE= | 36 | 8 | 13 | 8 | 261 | 27,9 | 5,74 | 6287 | 4,07 | 6 | 2 | 1,021 | 1,061 | 0,928 | 0,965 | 1,116 | 0,967 | 1,083 | 1,002 | 0,95 | -0,05 | 3,3E-01 |
| Q8BYH7 | TBC1 domain family member 17 OS=Mus musculus OX=10090 G | 2 | 2 | 2 | 2 | 645 | 72,8 | 5,25 | | 2,47 | | 2 | 1,15 | 1,222 | 0,997 | 1,009 | 1,071 | 0,859 | 0,881 | 1,171 | 1,10 | 0,10 | 3,3E-01 |
| Q8C3W1 | Uncharacterized protein C1orf198 homolog OS=Mus musculus C | 10 | 2 | 2 | 2 | 322 | 35,3 | 5,24 | 1190 | | 2 | | 1,225 | 0,995 | 1,061 | 1,274 | 1,078 | 1,124 | 1,083 | 0,944 | 1,08 | 0,08 | 3,3E-01 |
| P35922 | Synaptic functional regulator FMR1 OS=Mus musculus OX=1009 | 9 | 5 | 7 | 3 | 614 | 68,9 | 7,62 | 3085 | 2,33 | 4 | 1 | 1,101 | 1,002 | 1,124 | 1,216 | 1,089 | 1,036 | 1,066 | 1,06 | 1,05 | 0,05 | 3,3E-01 |
| Q9CQ60 | 6-phosphogluconolactonase OS=Mus musculus OX=10090 GN=F | 25 | 5 | 5 | 5 | 257 | 27,2 | 5,85 | 1784 | 5,37 | 3 | 2 | 0,982 | 0,963 | 1,019 | 0,955 | 0,864 | 1,011 | 0,947 | 0,955 | 1,04 | 0,04 | 3,3E-01 |
| P84091 | AP-2 complex subunit mu OS=Mus musculus OX=10090 GN=Ap2 | 35 | 15 | 27 | 15 | 435 | 49,6 | 9,54 | 11215 | 20,64 | 10 | 6 | 0,967 | 0,92 | 1,047 | 0,912 | 0,804 | 0,95 | 0,969 | 0,919 | 1,06 | 0,06 | 3,3E-01 |
| Q07079 | Insulin-like growth factor-binding protein 5 OS=Mus musculus O | 4 | 1 | 1 | 1 | 271 | 30,4 | 8,12 | 667 | | 1 | | 0,909 | 0,953 | 1,065 | 1,19 | 2,21 | 1,164 | 0,826 | 1,205 | 0,76 | -0,24 | 3,3E-01 |
| P21550 | Beta-enolase OS=Mus musculus OX=10090 GN=Eno3 PE=1 SV=3 | 24 | 10 | 17 | 4 | 434 | 47 | 7,18 | 6354 | 24,93 | 5 | 5 | 0,555 | 0,679 | 0,75 | 0,729 | 10,24 | 0,866 | 0,868 | 0,691 | 0,21 | -0,79 | 3,3E-01 |
| Q400C8 | Archaemetzincin-2 OS=Mus musculus OX=10090 GN=Amz2 PE=2 | 3 | 1 | 1 | 1 | 359 | 41,3 | 7,66 | 570 | | 1 | | 1,043 | 0,989 | 0,835 | 1,255 | 1,269 | 1,209 | 1,157 | 0,953 | 0,90 | -0,10 | 3,3E-01 |
| Q9CQA3 | Succinate dehydrogenase [ubiquinone] iron-sulfur subunit, mito | 28 | 7 | 9 | 7 | 282 | 31,8 | 8,68 | 4591 | 3,84 | 6 | 1 | 1,029 | 0,949 | 0,996 | 1,029 | 0,853 | 1,028 | 0,922 | 1,011 | 1,05 | 0,05 | 3,3E-01 |
| Q8K1M6 | Dynamin-1-like protein OS=Mus musculus OX=10090 GN=Dnm1 | 23 | 18 | 27 | 18 | 742 | 82,6 | 7,05 | 12821 | 27,63 | 13 | 5 | 1,004 | 0,964 | 1,045 | 0,96 | 0,904 | 1,002 | 0,949 | 0,992 | 1,03 | 0,03 | 3,3E-01 |
| Q9Z1Z0 | General vesicular transport factor p115 OS=Mus musculus OX=1 | 8 | 7 | 8 | 7 | 959 | 106,9 | 4,93 | 4357 | 2,01 | 6 | 1 | 1,005 | 1,001 | 0,914 | 1,035 | 0,903 | 0,892 | 0,912 | 1,054 | 1,05 | 0,05 | 3,3E-01 |
| Q9ERU3 | Zinc finger protein 22 OS=Mus musculus OX=10090 GN=Znf22 P | 5 | 1 | 1 | 1 | 237 | 27,3 | 10,29 | 385 | | 1 | | 0,837 | 1,105 | 0,899 | 1,046 | 1,174 | 0,836 | 1,154 | 1,153 | 0,90 | -0,10 | 3,3E-01 |
| Q8BR63 | Protein FAM177A1 OS=Mus musculus OX=10090 GN=Fam177a1 | 15 | 2 | 3 | 2 | 207 | 23,6 | 4,59 | 594 | 4,37 | 1 | 1 | 0,929 | 1,196 | 0,884 | 1,086 | 1,479 | 0,969 | 1,08 | 1,12 | 0,88 | -0,12 | 3,3E-01 |
| P51410 | 60S ribosomal protein L9 OS=Mus musculus OX=10090 GN=Rpl9 | 22 | 5 | 8 | 5 | 192 | 21,9 | 9,95 | 2299 | 7,27 | 4 | 3 | 1,099 | 1,003 | 1,144 | 1,03 | 1,052 | 1,187 | 1,106 | 1,109 | 0,96 | -0,04 | 3,4E-01 |
| Q60634 | Flotillin-2 OS=Mus musculus OX=10090 GN=Flot2 PE=1 SV=2 | 35 | 15 | 21 | 15 | 428 | 47 | 5,2 | 10166 | 11,71 | 12 | 3 | 0,948 | 1,002 | 0,927 | 0,94 | 1,137 | 0,908 | 0,991 | 0,992 | 0,95 | -0,05 | 3,4E-01 |
| Q8CCB4 | Vacuolar protein sorting-associated protein 53 homolog OS=Mu | 5 | 4 | 4 | 4 | 832 | 94,4 | 6,61 | 1721 | | 4 | | 0,763 | 0,841 | 0,924 | 0,894 | 2,074 | 0,707 | 0,971 | 0,954 | 0,73 | -0,27 | 3,4E-01 |
| O70250 | Phosphoglycerate mutase 2 OS=Mus musculus OX=10090 GN=P. | 21 | 7 | 17 | 2 | 253 | 28,8 | 8,5 | 5969 | 21,21 | 4 | 4 | 0,712 | 0,774 | 0,572 | 0,669 | 4,914 | 0,682 | 0,647 | 0,857 | 0,38 | -0,62 | 3,4E-01 |
| Q8BV13 | COP9 signalosome complex subunit 7b OS=Mus musculus OX=1( | 11 | 2 | 2 | 2 | 264 | 29,7 | 6,32 | | 7,39 | | 2 | 0,954 | 0,875 | 0,874 | 1,005 | 0,871 | 0,948 | 0,915 | 0,747 | 1,07 | 0,07 | 3,4E-01 |
| P63321 | Ras-related protein Ral-A OS=Mus musculus OX=10090 GN=Rala | 31 | 8 | 22 | 7 | 206 | 23,5 | 7,11 | 10527 | 16,83 | 6 | 4 | 1,028 | 0,826 | 1,195 | 0,92 | 0,71 | 1,094 | 0,846 | 0,85 | 1,13 | 0,13 | 3,4E-01 |
| Q9Z0L1 | Sphingosine 1-phosphate receptor 4 OS=Mus musculus OX=100 | 2 | 1 | 1 | 1 | 386 | 42,2 | 9,77 | 453 | | 1 | | 0,371 | 0,292 | 0,199 | 0,274 | 12,263 | 0,532 | 0,076 | 0,623 | 0,08 | -0,92 | 3,4E-01 |
| Q01065 | Calcium/calmodulin-dependent 3',5'-cyclic nucleotide phosphoc | 12 | 5 | 5 | 5 | 535 | 61,2 | 5,72 | 1913 | 4,63 | 3 | 2 | 1,097 | 1,054 | 1,019 | 0,989 | 1,036 | 1,072 | 1,088 | 1,07 | 0,97 | -0,03 | 3,4E-01 |
| P54731 | FAS-associated factor 1 OS=Mus musculus OX=10090 GN=Faf1 P | 6 | 3 | 4 | 3 | 649 | 73,8 | 4,86 | 1402 | | 3 | | 0,945 | 0,964 | 0,839 | 0,938 | 1,121 | 0,863 | 1,119 | 0,896 | 0,92 | -0,08 | 3,4E-01 |
| Q9JLM8 | Serine/threonine-protein kinase DCLK1 OS=Mus musculus OX=1 | 37 | 25 | 48 | 10 | 756 | 84,1 | 8,87 | 17332 | 55,72 | 15 | 14 | 0,97 | 0,903 | 0,999 | 0,935 | 0,639 | 0,947 | 0,936 | 0,954 | 1,10 | 0,10 | 3,4E-01 |
| Q06890 | Clusterin OS=Mus musculus OX=10090 GN=Clu PE=1 SV=1 | 2 | 1 | 1 | 1 | 448 | 51,6 | 5,67 | 715 | | 1 | | 1,068 | 0,909 | 1,046 | 0,868 | 1,05 | 0,873 | 0,825 | 0,846 | 1,08 | 0,08 | 3,4E-01 |
| P05132 | cAMP-dependent protein kinase catalytic subunit alpha OS=Mus | 27 | 11 | 21 | 4 | 351 | 40,5 | 8,79 | 4298 | 41,33 | 5 | 6 | 0,832 | 0,726 | 1,194 | 0,869 | 0,712 | 0,869 | 0,75 | 0,843 | 1,14 | 0,14 | 3,4E-01 |
| Q9EP69 | Phosphatidylinositide phosphatase SAC1 OS=Mus musculus OX= | 8 | 5 | 5 | 5 | 587 | 66,9 | 7,3 | 2124 | 2,61 | 4 | 1 | 0,985 | 1,112 | 1,084 | 1,042 | 1,054 | 1,084 | 0,92 | 0,971 | 1,05 | 0,05 | 3,4E-01 |
| Q80W54 | CAAX prenyl protease 1 homolog OS=Mus musculus OX=10090 ( | 4 | 2 | 3 | 2 | 475 | 54,7 | 6,95 | | 7,61 | | 2 | 1,17 | 1,122 | 1,027 | 1,139 | 1,127 | 1,037 | 1,096 | 1,041 | 1,04 | 0,04 | 3,4E-01 |
| Q9D819 | Inorganic pyrophosphatase OS=Mus musculus OX=10090 GN=Pp | 48 | 17 | 30 | 17 | 289 | 32,6 | 5,6 | 14195 | 13,28 | 15 | 5 | 1,008 | 1,068 | 0,927 | 1,14 | 0,87 | 1,004 | 1,041 | 0,986 | 1,06 | 0,06 | 3,4E-01 |
| P68404-2 | Isoform Beta-II of Protein kinase C beta type OS=Mus musculus | 14 | 12 | 13 | 7 | 673 | 76,8 | 7,02 | 4515 | 11,76 | 7 | 5 | 0,948 | 0,973 | 1,343 | 0,976 | 0,835 | 1,052 | 1,012 | 0,897 | 1,12 | 0,12 | 3,4E-01 |
| Q9CQK7 | RWD domain-containing protein 1 OS=Mus musculus OX=10090 | 5 | 1 | 1 | 1 | 243 | 27,8 | 4,26 | 542 | | 1 | | 0,998 | 0,958 | 1,193 | 1,169 | 1,012 | 1,036 | 0,975 | 1,041 | 1,06 | 0,06 | 3,4E-01 |
| Q99NB9 | Splicing factor 3B subunit 1 OS=Mus musculus OX=10090 GN=Sf | 1 | 1 | 1 | 1 | 1304 | 145,7 | 7,09 | 340 | | 1 | | 0,901 | 0,966 | 1,331 | 0,903 | 0,759 | 0,829 | 0,89 | 1,099 | 1,15 | 0,15 | 3,4E-01 |
| P28474 | Alcohol dehydrogenase class-3 OS=Mus musculus OX=10090 GN | 27 | 9 | 18 | 9 | 374 | 39,5 | 7,25 | 8773 | 10,68 | 7 | 2 | 0,97 | 1,01 | 0,95 | 1,102 | 0,701 | 1,057 | 0,966 | 0,961 | 1,09 | 0,09 | 3,4E-01 |
| P17225 | Polypyrimidine tract-binding protein 1 OS=Mus musculus OX=1( | 3 | 1 | 1 | 1 | 527 | 56,4 | 8,34 | | 3,96 | | 1 | 0,854 | 0,918 | 1,055 | 1,101 | 1,367 | 0,789 | 1,128 | 1,2 | 0,88 | -0,12 | 3,4E-01 |
| Q5PR73 | GTP-binding protein Di-Ras2 OS=Mus musculus OX=10090 GN=D | 27 | 4 | 5 | 4 | 199 | 22,5 | 8,76 | 3075 | | 4 | | 0,967 | 0,999 | 1,042 | 0,961 | 0,965 | 1,051 | 1,058 | 1,012 | 0,97 | -0,03 | 3,4E-01 |
| Q99JI6 | Ras-related protein Rap-1b OS=Mus musculus OX=10090 GN=Ra | 41 | 8 | 15 | 4 | 184 | 20,8 | 5,78 | 6665 | 10,88 | 5 | 3 | 1,051 | 1,053 | 1,076 | 1,038 | 0,9 | 1,046 | 0,824 | 1,147 | 1,08 | 0,08 | 3,4E-01 |
| Q62318 | Transcription intermediary factor 1-beta OS=Mus musculus OX= | 2 | 1 | 2 | 1 | 834 | 88,8 | 5,77 | 648 | 2,24 | 1 | 1 | 0,961 | 0,87 | 0,974 | 1,172 | 1,186 | 1,008 | 1,073 | 1,021 | 0,93 | -0,07 | 3,4E-01 |
| P97872 | Dimethylaniline monooxygenase [N-oxide-forming] 5 OS=Mus m | 5 | 2 | 3 | 2 | 533 | 60 | 8,73 | 2562 | | 2 | | 2,408 | 2,654 | 2,074 | 2,371 | 3,794 | 2,156 | 2,428 | 2,689 | 0,86 | -0,14 | 3,4E-01 |
| Q9Z0K8 | Pantetheinase OS=Mus musculus OX=10090 GN=Vnn1 PE=1 SV= | 3 | 1 | 1 | 1 | 512 | 57,1 | 5,94 | 802 | | 1 | | 1,085 | 1,103 | 0,878 | 0,828 | 1,615 | 0,75 | 1,148 | 1,165 | 0,83 | -0,17 | 3,4E-01 |
| Q5SX40 | Myosin-1 OS=Mus musculus OX=10090 GN=Myh1 PE=1 SV=1 | 9 | 17 | 18 | 3 | 1942 | 223,2 | 5,76 | 4969 | 20,43 | 10 | 7 | 0,845 | 1,014 | 0,62 | 1,005 | 6,494 | 0,874 | 0,874 | 1 | 0,38 | -0,62 | 3,4E-01 |
| Q6P5H2 | Nestin OS=Mus musculus OX=10090 GN=Nes PE=1 SV=1 | 2 | 3 | 4 | 3 | 1864 | 207 | 4,34 | 726 | 4,67 | 2 | 1 | 0,954 | 1,244 | 0,933 | 1,279 | 1,134 | 1,018 | 1,467 | 1,358 | 0,89 | -0,11 | 3,4E-01 |

| | | | | | | | | | | | | | | | | | | | | | | | |
|---|---|---|---|---|---|---|---|---|---|---|---|---|---|---|---|---|---|---|---|---|---|---|---|
| Q91WP6 | Serine protease inhibitor A3N OS=Mus musculus OX=10090 GN= | 5 | 1 | 1 | 1 | 418 | 46,7 | 5,82 | 338 | | 1 | | 0,941 | 1,397 | 0,778 | 0,849 | 2,1 | 0,732 | 1,169 | 1,271 | 0,75 | -0,25 | 3,4E-01 |
| E9PXF8 | Myotubularin-related protein 13 OS=Mus musculus OX=10090 G | 1 | 1 | 1 | 1 | 1872 | 210,3 | 7,28 | 557 | | 1 | | 1,64 | 2,014 | 1,476 | 2,135 | 1,379 | 1,41 | 2,101 | 1,414 | 1,15 | 0,15 | 3,4E-01 |
| Q9CQZ6 | NADH dehydrogenase [ubiquinone] 1 beta subcomplex subunit | 18 | 3 | 4 | 3 | 104 | 11,7 | 9,04 | 1051 | 4,84 | 1 | 2 | 1,016 | 0,767 | 1,033 | 0,967 | 0,748 | 0,781 | 1,003 | 0,903 | 1,10 | 0,10 | 3,4E-01 |
| Q60714 | Long-chain fatty acid transport protein 1 OS=Mus musculus OX= | 5 | 3 | 6 | 3 | 646 | 71,2 | 8,38 | 3514 | | 3 | | 1,16 | 0,983 | 1,04 | 1,052 | 1,143 | 0,922 | 0,949 | 0,965 | 1,06 | 0,06 | 3,4E-01 |
| P05977 | Myosin light chain 1/3, skeletal muscle isoform OS=Mus muscul | 14 | 3 | 4 | 3 | 188 | 20,6 | 5,03 | 621 | 5,27 | 1 | 2 | 0,466 | 0,677 | 0,562 | 0,618 | 10,789 | 0,579 | 0,496 | 0,872 | 0,18 | -0,82 | 3,4E-01 |
| P33175 | Kinesin heavy chain isoform 5A OS=Mus musculus OX=10090 GN | 9 | 11 | 14 | 4 | 1027 | 116,9 | 5,94 | 5007 | 14,43 | 7 | 4 | 0,987 | 0,966 | 0,937 | 1,446 | 1,777 | 0,915 | 1,426 | 1,132 | 0,83 | -0,17 | 3,4E-01 |
| Q9QYB8 | Beta-adducin OS=Mus musculus OX=10090 GN=Add2 PE=1 SV=4 | 15 | 11 | 23 | 10 | 725 | 80,6 | 6,21 | 10140 | 14,22 | 7 | 5 | 1,065 | 1,001 | 1,003 | 1,03 | 0,801 | 1,023 | 1,021 | 1,02 | 1,06 | 0,06 | 3,4E-01 |
| O08599 | Syntaxin-binding protein 1 OS=Mus musculus OX=10090 GN=Stx | 49 | 33 | 81 | 1 | 594 | 67,5 | 6,96 | 34847 | 64,91 | 19 | 14 | 1,192 | 0,757 | 0,976 | 0,903 | 0,818 | 1,645 | 0,886 | 1,371 | 0,81 | -0,19 | 3,5E-01 |
| P08556 | GTPase NRas OS=Mus musculus OX=10090 GN=Nras PE=1 SV=1 | 42 | 8 | 14 | 2 | 189 | 21,2 | 5,17 | 6681 | 7,89 | 7 | 2 | 1,453 | 0,965 | 1,5 | 1,203 | 1,118 | 1,352 | 1,029 | 1,026 | 1,13 | 0,13 | 3,5E-01 |
| Q8R4H2 | Rho guanine nucleotide exchange factor 12 OS=Mus musculus O | 2 | 2 | 2 | 2 | 1543 | 172,2 | 5,74 | 1179 | | 2 | | 1,078 | 1,396 | 1,089 | 1,021 | 1,256 | 1,207 | 1,302 | 1,182 | 0,93 | -0,07 | 3,5E-01 |
| P07146 | Anionic trypsin-2 OS=Mus musculus OX=10090 GN=Prss2 PE=1 S | 12 | 2 | 9 | 2 | 246 | 26,2 | 4,63 | 1455 | 17,19 | 1 | 1 | 1,002 | 1,203 | 0,787 | 0,945 | 1,68 | 0,926 | 1,207 | 0,93 | 0,83 | -0,17 | 3,5E-01 |
| Q62048 | Astrocytic phosphoprotein PEA-15 OS=Mus musculus OX=10090 | 46 | 6 | 8 | 5 | 130 | 15 | 5,02 | 4063 | 6,29 | 5 | 1 | 1,002 | 1,043 | 0,855 | 1,18 | 0,883 | 0,937 | 1,018 | 0,946 | 1,08 | 0,08 | 3,5E-01 |
| A6X935 | Inter alpha-trypsin inhibitor, heavy chain 4 OS=Mus musculus O | 2 | 2 | 2 | 2 | 942 | 104,6 | 6,4 | 740 | 0 | 1 | 1 | 0,995 | 1,017 | 1,003 | 0,953 | 6,214 | 1,025 | 0,861 | 1,179 | 0,43 | -0,57 | 3,5E-01 |
| Q7TSH3 | Zinc finger protein 516 OS=Mus musculus OX=10090 GN=Znf516 | 1 | 1 | 2 | 1 | 1157 | 124,7 | 8,51 | 523 | | 1 | | 0,494 | 0,5 | 0,478 | 0,418 | 7,292 | 0,51 | 0,402 | 0,616 | 0,21 | -0,79 | 3,5E-01 |
| P60879 | Synaptosomal-associated protein 25 OS=Mus musculus OX=100 | 70 | 17 | 52 | 16 | 206 | 23,3 | 4,77 | 18230 | 43,59 | 13 | 6 | 0,977 | 1,038 | 0,928 | 1,019 | 0,909 | 0,962 | 1,005 | 0,958 | 1,03 | 0,03 | 3,5E-01 |
| Q8BGB5 | LIM domain-containing protein 2 OS=Mus musculus OX=10090 | 5 | 1 | 1 | 1 | 128 | 14,2 | 9,04 | 521 | | 1 | | 0,852 | 0,774 | 1,443 | 0,935 | 1,025 | 0,608 | 0,735 | 0,913 | 1,22 | 0,22 | 3,5E-01 |
| Q61151 | Serine/threonine-protein phosphatase 2A 56 kDa regulatory sub | 15 | 6 | 8 | 4 | 467 | 54,7 | 6,95 | 3078 | 7,36 | 4 | 2 | 1,02 | 0,968 | 1,128 | 0,879 | 0,74 | 1,061 | 0,933 | 0,918 | 1,09 | 0,09 | 3,5E-01 |
| P62869 | Elongin-B OS=Mus musculus OX=10090 GN=Elob PE=1 SV=1 | 70 | 5 | 10 | 5 | 118 | 13,2 | 5,01 | 4508 | 5,71 | 4 | 2 | 0,979 | 1,016 | 1,04 | 1,168 | 0,87 | 1,12 | 1,014 | 0,918 | 1,07 | 0,07 | 3,5E-01 |
| P56480 | ATP synthase subunit beta, mitochondrial OS=Mus musculus OX | 53 | 23 | 130 | 23 | 529 | 56,3 | 5,34 | 52916 | 106,78 | 15 | 16 | 0,958 | 0,936 | 1,1 | 0,955 | 1,298 | 1 | 0,971 | 1,025 | 0,92 | -0,08 | 3,5E-01 |
| Q924N4 | Solute carrier family 12 member 6 OS=Mus musculus OX=10090 | 1 | 2 | 2 | 1 | 1150 | 127,4 | 7,03 | 1098 | | 2 | | 0,74 | 0,942 | 0,897 | 0,673 | 0,69 | 1,16 | 0,773 | 1,255 | 0,84 | -0,16 | 3,5E-01 |
| Q9R062 | Glycogenin-1 OS=Mus musculus OX=10090 GN=Gyg1 PE=1 SV=3 | 7 | 2 | 2 | 2 | 333 | 37,4 | 5,29 | 829 | | 2 | | 1,268 | 1,165 | 1,094 | 1,263 | 1,652 | 1,204 | 1,215 | 1,203 | 0,91 | -0,09 | 3,5E-01 |
| Q3TDX8 | Cytochrome b5 reductase 4 OS=Mus musculus OX=10090 GN=C | 1 | 1 | 1 | 1 | 528 | 59,7 | 6,35 | | 2,45 | | 1 | 1,14 | 1,097 | 0,922 | 1,004 | 0,711 | 1,07 | 0,861 | 1,099 | 1,11 | 0,11 | 3,5E-01 |
| Q9JHQ5 | Leucine zipper transcription factor-like protein 1 OS=Mus muscu | 11 | 3 | 4 | 3 | 299 | 34,8 | 5,17 | 803 | 4,36 | 2 | 1 | 1,202 | 1,051 | 1,29 | 1,142 | 1,088 | 1,25 | 1,031 | 1,018 | 1,07 | 0,07 | 3,5E-01 |
| P26516 | 26S proteasome non-ATPase regulatory subunit 7 OS=Mus musc | 7 | 2 | 2 | 2 | 321 | 36,5 | 6,77 | 1215 | | 2 | | 1,061 | 0,944 | 1,34 | 1,09 | 1,189 | 1,327 | 1,039 | 1,315 | 0,91 | -0,09 | 3,5E-01 |
| P47911 | 60S ribosomal protein L6 OS=Mus musculus OX=10090 GN=Rpl6 | 36 | 13 | 21 | 13 | 296 | 33,5 | 10,7 | 7266 | 19,45 | 8 | 5 | 1,078 | 0,895 | 0,959 | 1,054 | 0,848 | 0,988 | 0,962 | 0,972 | 1,06 | 0,06 | 3,5E-01 |
| Q91Z69 | SLIT-ROBO Rho GTPase-activating protein 1 OS=Mus musculus C | 3 | 3 | 7 | 2 | 1062 | 121,4 | 6,74 | 3186 | 7,39 | 2 | 2 | 0,921 | 0,928 | 1,131 | 0,86 | 0,721 | 0,95 | 0,886 | 0,956 | 1,09 | 0,09 | 3,5E-01 |
| P13412 | Troponin I, fast skeletal muscle OS=Mus musculus OX=10090 GN | 6 | 1 | 1 | 1 | 182 | 21,3 | 8,5 | | 0 | | 1 | 0,611 | 0,895 | 0,668 | 0,831 | 12,013 | 0,686 | 0,542 | 1,166 | 0,21 | -0,79 | 3,5E-01 |
| Q8C170 | Unconventional myosin-IXa OS=Mus musculus OX=10090 GN=M | 1 | 1 | 1 | 1 | 2542 | 291,9 | 8,97 | | 0 | | 1 | 0,71 | 0,875 | 0,943 | 1,048 | 1,22 | 0,895 | 0,918 | 0,962 | 0,90 | -0,10 | 3,5E-01 |
| P12970 | 60S ribosomal protein L7a OS=Mus musculus OX=10090 GN=Rpl | 29 | 8 | 13 | 8 | 266 | 30 | 10,56 | 5715 | 17,1 | 6 | 2 | 1,088 | 0,913 | 1,038 | 1,033 | 0,942 | 0,96 | 1,046 | 0,943 | 1,05 | 0,05 | 3,5E-01 |
| P08730 | Keratin, type I cytoskeletal 13 OS=Mus musculus OX=10090 GN= | 13 | 9 | 30 | 1 | 437 | 47,7 | 4,86 | 10329 | 28,76 | 5 | 7 | 0,64 | 0,697 | 0,801 | 0,68 | 6,885 | 0,814 | 0,595 | 0,771 | 0,31 | -0,69 | 3,5E-01 |
| Q9DCN2 | NADH-cytochrome b5 reductase 3 OS=Mus musculus OX=10090 | 45 | 11 | 21 | 11 | 301 | 34,1 | 8,38 | 7988 | 13,51 | 10 | 3 | 1,047 | 1,058 | 0,918 | 1,09 | 0,887 | 0,926 | 1,06 | 1,017 | 1,06 | 0,06 | 3,5E-01 |
| Q9D4H1 | Exocyst complex component 2 OS=Mus musculus OX=10090 GN | 8 | 7 | 9 | 7 | 924 | 103,9 | 7,18 | 2428 | 11,54 | 4 | 4 | 0,978 | 1,002 | 0,82 | 0,923 | 1,066 | 0,877 | 1,047 | 0,972 | 0,94 | -0,06 | 3,5E-01 |
| Q8K386 | Ras-related protein Rab-15 OS=Mus musculus OX=10090 GN=Ra | 11 | 2 | 5 | 1 | 212 | 24,3 | 5,71 | 2657 | 2,19 | 1 | 1 | 0,901 | 1,027 | 0,938 | 1,495 | 1,256 | 1,106 | 1,234 | 1,361 | 0,88 | -0,12 | 3,5E-01 |
| Q9D1M4 | Eukaryotic translation elongation factor 1 epsilon-1 OS=Mus mu | 6 | 1 | 1 | 1 | 174 | 19,8 | 8,59 | 261 | | 1 | | 0,92 | 0,824 | 0,81 | 1,031 | 0,691 | 0,916 | 0,651 | 0,953 | 1,12 | 0,12 | 3,5E-01 |
| Q99L85 | Elongator complex protein 5 OS=Mus musculus OX=10090 GN=E | 3 | 1 | 4 | 1 | 300 | 33,5 | 5,49 | 2621 | | 1 | | 1,081 | 0,903 | 1,079 | 1,07 | 1,007 | 1,051 | 0,93 | 0,932 | 1,05 | 0,05 | 3,5E-01 |
| P05366 | Serum amyloid A-1 protein OS=Mus musculus OX=10090 GN=Sa | 7 | 1 | 2 | 1 | 122 | 13,8 | 7,03 | 977 | | 1 | | 0,647 | 0,602 | 0,742 | 0,567 | 7,556 | 0,808 | 0,542 | 0,628 | 0,27 | -0,73 | 3,5E-01 |
| Q9WUA2 | Phenylalanine--tRNA ligase beta subunit OS=Mus musculus OX= | 5 | 3 | 4 | 3 | 589 | 65,7 | 7,12 | 1483 | 5,03 | 2 | 1 | 0,89 | 0,949 | 1,024 | 0,95 | 0,918 | 0,936 | 0,914 | 0,932 | 1,03 | 0,03 | 3,5E-01 |
| P39447 | Tight junction protein ZO-1 OS=Mus musculus OX=10090 GN=Tj | 4 | 4 | 4 | 4 | 1745 | 194,6 | 6,64 | 1526 | | 4 | | 0,83 | 0,931 | 0,975 | 1,011 | 1,124 | 0,829 | 1,025 | 1,074 | 0,92 | -0,08 | 3,5E-01 |
| P58044 | Isopentenyl-diphosphate Delta-isomerase 1 OS=Mus musculus C | 10 | 3 | 3 | 3 | 227 | 26,3 | 6,16 | 1311 | 3,01 | 2 | 1 | 1,021 | 0,994 | 1,167 | 1,113 | 0,832 | 1,202 | 1,041 | 0,813 | 1,10 | 0,10 | 3,5E-01 |
| P34022 | Ran-specific GTPase-activating protein OS=Mus musculus OX=10 | 25 | 4 | 6 | 4 | 203 | 23,6 | 5,22 | 787 | 8,94 | 1 | 3 | 0,95 | 0,984 | 1,051 | 0,984 | 1,144 | 1,102 | 1,011 | 0,926 | 0,95 | -0,05 | 3,5E-01 |
| Q61646 | Haptoglobin OS=Mus musculus OX=10090 GN=Hp PE=1 SV=1 | 3 | 1 | 1 | 1 | 347 | 38,7 | 6,29 | | 2,3 | | 1 | 0,754 | 0,768 | 0,796 | 0,819 | 4,405 | 0,765 | 0,731 | 0,883 | 0,46 | -0,54 | 3,5E-01 |
| Q99MD6 | Thioredoxin reductase 3 OS=Mus musculus OX=10090 GN=Txnr | 4 | 2 | 2 | 2 | 652 | 71,3 | 8,09 | 228 | 2,08 | 1 | 1 | 0,874 | 1,017 | 1,118 | 1,034 | 2,574 | 1,029 | 0,867 | 1,171 | 0,72 | -0,28 | 3,5E-01 |
| Q60700 | Mitogen-activated protein kinase kinase kinase 12 OS=Mus mus | 4 | 3 | 3 | 3 | 888 | 96 | 6,24 | 1655 | | 3 | | 0,834 | 0,701 | 1,073 | 0,994 | 1,618 | 0,826 | 1,001 | 0,946 | 0,82 | -0,18 | 3,5E-01 |
| Q99KQ4 | Nicotinamide phosphoribosyltransferase OS=Mus musculus OX= | 4 | 1 | 1 | 1 | 491 | 55,4 | 7,15 | | 3,27 | | 1 | 1,015 | 1,393 | 1,04 | 1,261 | 1,84 | 1,077 | 1,037 | 1,649 | 0,84 | -0,16 | 3,5E-01 |
| Q62448 | Eukaryotic translation initiation factor 4 gamma 2 OS=Mus musc | 4 | 3 | 3 | 3 | 906 | 102 | 7,14 | 1114 | 0 | 2 | 1 | 0,885 | 0,807 | 1,098 | 0,95 | 1,08 | 1,037 | 0,926 | 0,98 | 0,93 | -0,07 | 3,5E-01 |
| Q9WUB3 | Glycogen phosphorylase, muscle form OS=Mus musculus OX=10 | 4 | 4 | 10 | 1 | 842 | 97,2 | 7,11 | 3340 | 8,28 | 3 | 2 | 0,894 | 0,708 | 0,757 | 0,794 | 4,138 | 0,642 | 0,749 | 1,004 | 0,48 | -0,52 | 3,5E-01 |
| Q9EPV8 | Ubiquitin-like protein 5 OS=Mus musculus OX=10090 GN=Ubl5 P | 12 | 1 | 1 | 1 | 73 | 8,5 | 8,44 | | 1,75 | | 1 | 0,817 | 1,059 | 0,933 | 0,93 | 1 | 0,86 | 1 | 1,313 | 0,90 | -0,10 | 3,5E-01 |
| P26339 | Chromogranin-A OS=Mus musculus OX=10090 GN=Chga PE=1 S | 4 | 2 | 2 | 2 | 463 | 51,8 | 4,72 | 578 | 0 | 1 | 1 | 1,128 | 1,02 | 1,097 | 1,219 | 1,126 | 1,015 | 1,12 | 0,981 | 1,05 | 0,05 | 3,5E-01 |
| P08553 | Neurofilament medium polypeptide OS=Mus musculus OX=100 | 25 | 20 | 25 | 17 | 848 | 95,9 | 4,77 | 8110 | 26,34 | 10 | 11 | 1,049 | 1,002 | 1,049 | 1,014 | 0,832 | 1,048 | 1,032 | 0,997 | 1,05 | 0,05 | 3,5E-01 |
| P43274 | Histone H1.4 OS=Mus musculus OX=10090 GN=Hist1h1e PE=1 S | 40 | 13 | 23 | 3 | 219 | 22 | 11,11 | 2190 | 73,93 | 2 | 11 | 0,92 | 0,843 | 0,97 | 1,094 | 1,063 | 0,891 | 1,158 | 1,022 | 0,93 | -0,07 | 3,5E-01 |
| P28665 | Murinoglobulin-1 OS=Mus musculus OX=10090 GN=Mug1 PE=1 | 1 | 2 | 2 | 2 | 1476 | 165,2 | 6,42 | 720 | 2,51 | 1 | 1 | 1,1 | 0,877 | 1,065 | 0,846 | 1,254 | 1,189 | 0,874 | 1,005 | 0,90 | -0,10 | 3,5E-01 |
| B9EKR1 | Receptor-type tyrosine-protein phosphatase zeta OS=Mus musc | 1 | 2 | 2 | 2 | 2312 | 254,2 | 4,88 | 1184 | | 2 | | 1,129 | 0,922 | 1,061 | 1,094 | 0,681 | 1,074 | 1,138 | 0,855 | 1,12 | 0,12 | 3,5E-01 |
| O35215 | D-dopachrome decarboxylase OS=Mus musculus OX=10090 GN= | 33 | 3 | 3 | 3 | 118 | 13,1 | 6,54 | 1489 | 2,27 | 2 | 1 | 0,946 | 1,015 | 0,887 | 1,102 | 0,987 | 1,157 | 1,067 | 0,986 | 0,94 | -0,06 | 3,5E-01 |
| B9EHT4 | CAP-Gly domain-containing linker protein 3 OS=Mus musculus C | 7 | 3 | 4 | 3 | 547 | 59,5 | 7,94 | 2036 | 0 | 2 | 1 | 0,944 | 0,978 | 0,834 | 0,897 | 1,31 | 0,832 | 0,954 | 0,985 | 0,90 | -0,10 | 3,5E-01 |
| Q8BH60 | Golgi-associated PDZ and coiled-coil motif-containing protein O | 4 | 2 | 2 | 2 | 463 | 50,6 | 6,25 | 338 | 1,63 | 1 | 1 | 1,134 | 1,049 | 1,065 | 1,17 | 1,21 | 1,026 | 1,263 | 1,153 | 0,95 | -0,05 | 3,5E-01 |
| P14869 | 60S acidic ribosomal protein P0 OS=Mus musculus OX=10090 G | 40 | 11 | 21 | 11 | 317 | 34,2 | 6,25 | 10079 | 11,59 | 9 | 4 | 0,968 | 0,959 | 1,059 | 0,995 | 0,93 | 1,14 | 1,014 | 1,108 | 0,95 | -0,05 | 3,6E-01 |
| Q3USJ8 | F-BAR and double SH3 domains protein 2 OS=Mus musculus OX | 2 | 1 | 1 | 1 | 740 | 84,2 | 5,66 | 380 | | 1 | | 0,589 | 0,788 | 0,815 | 0,81 | 1,044 | 0,801 | 0,769 | 0,741 | 0,89 | -0,11 | 3,6E-01 |
| Q91XD6 | Vacuolar protein-sorting-associated protein 36 OS=Mus muscul | 3 | 1 | 1 | 1 | 386 | 43,7 | 7,15 | | 3,57 | | 1 | 1,088 | 0,963 | 1,064 | 0,871 | 0,627 | 0,951 | 1,056 | 0,932 | 1,12 | 0,12 | 3,6E-01 |
| Q8BIG7 | Catechol O-methyltransferase domain-containing protein 1 OS= | 8 | 2 | 3 | 2 | 262 | 28,9 | 8,32 | | 6,86 | | 2 | 0,768 | 0,738 | 1,115 | 0,67 | 0,816 | 1,037 | 0,949 | 0,927 | 0,88 | -0,12 | 3,6E-01 |
| Q9CPU0 | Lactoylglutathione lyase OS=Mus musculus OX=10090 GN=Glo1 | 8 | 3 | 7 | 3 | 184 | 20,8 | 5,47 | 4200 | 2,66 | 2 | 1 | 1,055 | 1,171 | 1,13 | 1,288 | 1,042 | 1,233 | 1,02 | 1,075 | 1,06 | 0,06 | 3,6E-01 |
| Q9QZE5 | Coatomer subunit gamma-1 OS=Mus musculus OX=10090 GN=C | 14 | 10 | 12 | 9 | 874 | 97,5 | 5,35 | 4973 | 9,44 | 6 | 4 | 1,044 | 0,959 | 1,061 | 1,059 | 0,895 | 0,991 | 1,02 | 1,051 | 1,04 | 0,04 | 3,6E-01 |
| Q7TN29 | Stromal membrane-associated protein 2 OS=Mus musculus OX= | 3 | 1 | 1 | 1 | 428 | 46,5 | 8,87 | 696 | | 1 | | 0,677 | 0,83 | 0,813 | 0,79 | 0,739 | 0,63 | 0,439 | 0,895 | 1,15 | 0,15 | 3,6E-01 |
| Q9Z2I0 | Mitochondrial proton/calcium exchanger protein OS=Mus musc | 10 | 7 | 9 | 7 | 738 | 82,9 | 6,52 | 4759 | 3,19 | 5 | 2 | 0,941 | 1,01 | 1,176 | 1,068 | 0,747 | 1,085 | 1,054 | 0,945 | 1,10 | 0,10 | 3,6E-01 |
| P61082 | NEDD8-conjugating enzyme Ubc12 OS=Mus musculus OX=1009 | 39 | 8 | 15 | 8 | 183 | 20,9 | 7,69 | 4579 | 16,91 | 4 | 5 | 0,961 | 0,965 | 1,12 | 0,955 | 0,724 | 1,113 | 0,867 | 0,937 | 1,10 | 0,10 | 3,6E-01 |
| Q6PFE7 | Tomoregulin-1 OS=Mus musculus OX=10090 GN=Tmeff1 PE=2 S | 5 | 2 | 2 | 2 | 372 | 40,1 | 6,81 | 1008 | | 2 | | 0,859 | 0,978 | 0,987 | 0,835 | 2,381 | 0,853 | 0,872 | 1,03 | 0,71 | -0,29 | 3,6E-01 |

| | | | | | | | | | | | | | | | | | | | | | | | |
|---|---|---|---|---|---|---|---|---|---|---|---|---|---|---|---|---|---|---|---|---|---|---|---|
| Q8K012 | Formin-binding protein 1-like OS=Mus musculus OX=10090 GN= | 5 | 4 | 4 | 4 | 605 | 69,8 | 6,64 | 1938 | 2,11 | 3 | 1 | 1,032 | 1,057 | 0,97 | 0,917 | 0,799 | 1,072 | 1,012 | 0,761 | 1,09 | 0,09 | 3,6E-01 |
| P97471 | Mothers against decapentaplegic homolog 4 OS=Mus musculus | 1 | 1 | 1 | 1 | 551 | 60,3 | 6,99 | 457 | | 1 | | 1,045 | 0,832 | 1,156 | 1,141 | 0,556 | 1,13 | 1,018 | 0,892 | 1,16 | 0,16 | 3,6E-01 |
| P35288 | Ras-related protein Rab-23 OS=Mus musculus OX=10090 GN=Ra | 15 | 3 | 3 | 3 | 237 | 26,7 | 6,79 | 714 | 4,72 | 1 | 2 | 0,779 | 0,842 | 0,996 | 0,87 | 0,926 | 0,906 | 0,905 | 0,934 | 0,95 | -0,05 | 3,6E-01 |
| Q9CZ04 | COP9 signalosome complex subunit 7a OS=Mus musculus OX=1( | 13 | 3 | 4 | 3 | 275 | 30,2 | 7,87 | 2088 | | 3 | | 0,975 | 0,971 | 0,971 | 1,124 | 0,891 | 0,94 | 0,986 | 1,031 | 1,05 | 0,05 | 3,6E-01 |
| Q9D6Y9 | 1,4-alpha-glucan-branching enzyme OS=Mus musculus OX=1009 | 2 | 1 | 2 | 1 | 702 | 80,3 | 6,43 | 1318 | | 1 | | 0,523 | 0,648 | 0,805 | 0,692 | 1,094 | 0,618 | 0,641 | 0,81 | 0,84 | -0,16 | 3,6E-01 |
| P30681 | High mobility group protein B2 OS=Mus musculus OX=10090 GN | 18 | 5 | 8 | 5 | 210 | 24,1 | 7,31 | 2178 | 8,96 | 3 | 2 | 0,922 | 0,955 | 0,994 | 1,029 | 1,844 | 0,977 | 0,991 | 0,958 | 0,82 | -0,18 | 3,6E-01 |
| E9Q5C9 | Nucleolar and coiled-body phosphoprotein 1 OS=Mus musculus | 1 | 1 | 3 | 1 | 702 | 73,7 | 9,36 | 1371 | | 1 | | 0,446 | 0,415 | 0,42 | 0,412 | 9,038 | 0,433 | 0,309 | 0,496 | 0,16 | -0,84 | 3,6E-01 |
| P68510 | 14-3-3 protein eta OS=Mus musculus OX=10090 GN=Ywhah PE= | 57 | 17 | 61 | 10 | 246 | 28,2 | 4,89 | 21135 | 71,21 | 9 | 12 | 1,013 | 0,994 | 0,948 | 1,019 | 0,835 | 1,084 | 0,956 | 0,869 | 1,06 | 0,06 | 3,6E-01 |
| Q6NXK7 | Inactive dipeptidyl peptidase 10 OS=Mus musculus OX=10090 G | 15 | 12 | 15 | 11 | 797 | 90,8 | 6,48 | 5695 | 13,33 | 8 | 4 | 0,947 | 1,028 | 0,959 | 1,003 | 1,028 | 0,938 | 0,906 | 0,936 | 1,03 | 0,03 | 3,6E-01 |
| Q921I1 | Serotransferrin OS=Mus musculus OX=10090 GN=Tf PE=1 SV=1 | 21 | 13 | 17 | 13 | 697 | 76,7 | 7,18 | 6515 | 16,21 | 9 | 4 | 1,005 | 0,97 | 1,016 | 0,996 | 1,699 | 0,959 | 1,001 | 1,029 | 0,85 | -0,15 | 3,6E-01 |
| Q9CR16 | Peptidyl-prolyl cis-trans isomerase D OS=Mus musculus OX=100 | 30 | 10 | 13 | 10 | 370 | 40,7 | 7,43 | 4475 | 15,87 | 4 | 6 | 1,042 | 0,955 | 1,04 | 0,994 | 0,747 | 1,08 | 0,954 | 0,963 | 1,08 | 0,08 | 3,6E-01 |
| Q8BK30 | NADH dehydrogenase [ubiquinone] flavoprotein 3, mitochondri | 16 | 2 | 2 | 2 | 104 | 11,8 | 9,35 | 1358 | | 2 | | 1,233 | 0,944 | 0,81 | 1,266 | 0,72 | 0,845 | 1,161 | 0,951 | 1,16 | 0,16 | 3,6E-01 |
| Q9DCQ7 | Alpha-1-antitrypsin 1-6 OS=Mus musculus OX=10090 GN=Serpin | 2 | 1 | 1 | 1 | 411 | 46,3 | 8,59 | | 3,1 | | 1 | 0,274 | 0,324 | 0,154 | 0,198 | 16,031 | 0,275 | 0,01 | 0,318 | 0,06 | -0,94 | 3,6E-01 |
| Q8BTY2-2 | Isoform 2 of Sodium bicarbonate cotransporter 3 OS=Mus musc | 2 | 2 | 2 | 1 | 936 | 104,6 | 6,38 | 634 | 2,34 | 1 | 1 | 1,05 | 0,917 | 0,808 | 0,859 | 4,67 | 1,156 | 0,735 | 0,825 | 0,49 | -0,51 | 3,6E-01 |
| Q8VCT4 | Carboxylesterase 1D OS=Mus musculus OX=10090 GN=Ces1d PE | 5 | 3 | 3 | 3 | 565 | 61,7 | 6,61 | 1255 | | 3 | | 0,951 | 1,204 | 1,128 | 1,398 | 1,96 | 0,923 | 1,432 | 1,291 | 0,83 | -0,17 | 3,6E-01 |
| O55091 | Protein IMPACT OS=Mus musculus OX=10090 GN=Impact PE=1 S | 3 | 1 | 1 | 1 | 318 | 36,3 | 5,05 | 722 | | 1 | | 0,89 | 1,111 | 1,033 | 1,195 | 0,974 | 1,031 | 0,991 | 0,971 | 1,07 | 0,07 | 3,6E-01 |
| Q3URD3 | Sarcolemmal membrane-associated protein OS=Mus musculus O | 2 | 2 | 3 | 2 | 845 | 96,9 | 5,33 | 1284 | | 2 | | 0,477 | 0,48 | 0,386 | 0,386 | 14,128 | 0,39 | 0,205 | 0,575 | 0,11 | -0,89 | 3,6E-01 |
| O54983 | Ketimine reductase mu-crystallin OS=Mus musculus OX=10090 ( | 28 | 9 | 12 | 9 | 313 | 33,5 | 5,67 | 6427 | | 9 | | 1,009 | 1,124 | 0,835 | 1,098 | 0,945 | 0,874 | 0,994 | 0,975 | 1,07 | 0,07 | 3,6E-01 |
| P54923 | [Protein ADP-ribosylarginine] hydrolase OS=Mus musculus OX=1 | 16 | 4 | 5 | 4 | 362 | 40 | 5,76 | 2279 | 2,14 | 3 | 1 | 0,903 | 0,913 | 0,931 | 0,986 | 1,19 | 0,908 | 1,005 | 0,904 | 0,93 | -0,07 | 3,6E-01 |
| Q505D1 | Serine/threonine-protein phosphatase 6 regulatory ankyrin repe | 3 | 2 | 2 | 2 | 1053 | 112,8 | 6,27 | 1248 | | 2 | | 0,712 | 0,78 | 0,866 | 0,956 | 1,009 | 0,957 | 0,706 | 0,99 | 0,90 | -0,10 | 3,6E-01 |
| Q80TL0 | Protein phosphatase 1E OS=Mus musculus OX=10090 GN=Ppm1 | 5 | 3 | 4 | 3 | 749 | 83,4 | 4,97 | 821 | 3,45 | 2 | 1 | 0,982 | 0,928 | 0,868 | 0,936 | 1,055 | 0,94 | 1,095 | 0,857 | 0,94 | -0,06 | 3,6E-01 |
| P50516 | V-type proton ATPase catalytic subunit A OS=Mus musculus OX= | 41 | 26 | 66 | 26 | 617 | 68,3 | 5,58 | 25856 | 63,95 | 17 | 12 | 1,081 | 1,008 | 0,983 | 0,966 | 0,866 | 1,099 | 0,922 | 0,929 | 1,06 | 0,06 | 3,6E-01 |
| Q05BC3 | Echinoderm microtubule-associated protein-like 1 OS=Mus mus | 7 | 5 | 6 | 5 | 814 | 89,6 | 7,02 | 385 | 13,12 | 1 | 4 | 0,928 | 0,892 | 0,909 | 0,918 | 0,836 | 0,973 | 1,064 | 0,961 | 0,95 | -0,05 | 3,6E-01 |
| Q9JJZ2 | Tubulin alpha-8 chain OS=Mus musculus OX=10090 GN=Tuba8 F | 33 | 16 | 172 | 1 | 449 | 50 | 5,1 | 67315 | 215,35 | 14 | 9 | 1,033 | 1,07 | 0,936 | 1,2 | 0,96 | 1,189 | 1,152 | 1,279 | 0,93 | -0,07 | 3,6E-01 |
| Q9DCC4 | Pyrroline-5-carboxylate reductase 3 OS=Mus musculus OX=1009 | 21 | 5 | 9 | 5 | 274 | 28,7 | 7,27 | 1218 | 12,97 | 3 | 3 | 0,99 | 0,968 | 1,045 | 0,93 | 0,706 | 1,037 | 0,954 | 0,941 | 1,08 | 0,08 | 3,6E-01 |
| O70325 | Phospholipid hydroperoxide glutathione peroxidase OS=Mus mu | 28 | 6 | 9 | 6 | 197 | 22,2 | 8,46 | 4421 | 5,35 | 6 | 2 | 0,98 | 0,999 | 0,987 | 1,023 | 1,184 | 0,934 | 1,065 | 1,015 | 0,95 | -0,05 | 3,6E-01 |
| Q8CIV8 | Tubulin-specific chaperone E OS=Mus musculus OX=10090 GN=T | 2 | 1 | 1 | 1 | 524 | 59 | 6,29 | 530 | | 1 | | 0,957 | 1,071 | 1,119 | 1,009 | 1,088 | 1,017 | 1,01 | 1,944 | 0,82 | -0,18 | 3,6E-01 |
| Q9JII6 | Aldo-keto reductase family 1 member A1 OS=Mus musculus OX= | 35 | 13 | 24 | 13 | 325 | 36,6 | 7,39 | 7332 | 35,27 | 8 | 6 | 0,976 | 0,948 | 1,137 | 1,072 | 0,997 | 1,031 | 0,954 | 0,967 | 1,05 | 0,05 | 3,6E-01 |
| P58771 | Tropomyosin alpha-1 chain OS=Mus musculus OX=10090 GN=Tp | 50 | 21 | 29 | 12 | 284 | 32,7 | 4,74 | 6276 | 40,58 | 7 | 14 | 0,86 | 0,835 | 1,031 | 0,902 | 2,825 | 0,931 | 0,894 | 0,876 | 0,66 | -0,34 | 3,6E-01 |
| Q0VGU4 | Neurosecretory protein VGF OS=Mus musculus OX=10090 GN=V | 21 | 10 | 17 | 10 | 617 | 68,2 | 4,7 | 5682 | 15,75 | 6 | 4 | 0,994 | 0,949 | 0,906 | 0,948 | 1,066 | 0,963 | 0,915 | 0,996 | 0,96 | -0,04 | 3,6E-01 |
| Q9CZU3 | Exosome RNA helicase MTR4 OS=Mus musculus OX=10090 GN= | 2 | 2 | 2 | 1 | 1040 | 117,6 | 6,4 | 807 | | 2 | | 0,429 | 0,463 | 0,369 | 0,397 | 13,059 | 0,427 | 0,18 | 0,447 | 0,12 | -0,88 | 3,6E-01 |
| P46978 | Dolichyl-diphosphooligosaccharide--protein glycosyltransferase | 1 | 1 | 1 | 1 | 705 | 80,5 | 8,1 | | 2,66 | | 1 | 1,23 | 0,896 | 1,357 | 1,024 | 1,469 | 0,923 | 1,26 | 1,523 | 0,87 | -0,13 | 3,7E-01 |
| Q91XQ0 | Dynein heavy chain 8, axonemal OS=Mus musculus OX=10090 G | 0 | 1 | 1 | 1 | 4731 | 540,9 | 6,18 | | 2,28 | | 1 | 0,913 | 0,815 | 0,96 | 0,634 | 0,402 | 1,064 | 0,634 | 0,609 | 1,23 | 0,23 | 3,7E-01 |
| Q8K4M5 | COMM domain-containing protein 1 OS=Mus musculus OX=100! | 14 | 2 | 2 | 2 | 188 | 21 | 7,59 | | 4,97 | | 2 | 0,98 | 1,079 | 0,951 | 1,072 | 1,407 | 0,966 | 1,09 | 1,024 | 0,91 | -0,09 | 3,7E-01 |
| P55258 | Ras-related protein Rab-8A OS=Mus musculus OX=10090 GN=Ra | 12 | 3 | 9 | 1 | 207 | 23,7 | 9,07 | 5540 | | 3 | | 0,991 | 1,104 | 1,021 | 1,036 | 0,925 | 0,92 | 1,092 | 1,027 | 1,05 | 0,05 | 3,7E-01 |
| Q9WUR9 | Adenylate kinase 4, mitochondrial OS=Mus musculus OX=10090 | 10 | 2 | 2 | 2 | 223 | 25 | 7,53 | 1751 | | 2 | | 0,966 | 1,189 | 1,12 | 0,856 | 0,768 | 0,777 | 0,735 | 1,271 | 1,16 | 0,16 | 3,7E-01 |
| Q8K0D0 | Cyclin-dependent kinase 17 OS=Mus musculus OX=10090 GN=C | 1 | 1 | 1 | 1 | 523 | 59,5 | 9,03 | | 2,09 | | 1 | 0,985 | 0,789 | 0,983 | 1,05 | 0,935 | 0,849 | 0,892 | 0,9 | 1,06 | 0,06 | 3,7E-01 |
| Q6P5F9 | Exportin-1 OS=Mus musculus OX=10090 GN=Xpo1 PE=1 SV=1 | 6 | 6 | 9 | 6 | 1071 | 123 | 6,07 | 1180 | 18,09 | 2 | 4 | 1,006 | 0,988 | 0,971 | 0,974 | 1,038 | 0,947 | 1,067 | 0,994 | 0,97 | -0,03 | 3,7E-01 |
| Q9DC16 | Endoplasmic reticulum-Golgi intermediate compartment protein | 13 | 2 | 3 | 2 | 290 | 32,5 | 7,06 | 1248 | 2,63 | 2 | 1 | 0,888 | 0,983 | 0,935 | 0,94 | 1,017 | 1,01 | 1,004 | 0,871 | 0,96 | -0,04 | 3,7E-01 |
| P13542 | Myosin-8 OS=Mus musculus OX=10090 GN=Myh8 PE=2 SV=2 | 7 | 15 | 18 | 1 | 1937 | 222,6 | 5,83 | 6165 | 18,6 | 8 | 7 | 0,815 | 0,96 | 0,731 | 0,894 | 6,25 | 0,769 | 0,754 | 0,932 | 0,39 | -0,61 | 3,7E-01 |
| Q9Z1L5 | Voltage-dependent calcium channel subunit alpha-2/delta-3 OS | 4 | 4 | 6 | 4 | 1091 | 122,7 | 5,73 | 1815 | 8,13 | 2 | 2 | 1,056 | 0,958 | 0,891 | 0,969 | 0,717 | 1,106 | 0,849 | 0,859 | 1,10 | 0,10 | 3,7E-01 |
| Q920E5 | Farnesyl pyrophosphate synthase OS=Mus musculus OX=10090 | 20 | 8 | 8 | 8 | 353 | 40,6 | 5,66 | 2981 | 5,27 | 5 | 3 | 0,969 | 0,929 | 1,127 | 0,959 | 0,789 | 1,071 | 0,923 | 0,917 | 1,08 | 0,08 | 3,7E-01 |
| P50396 | Rab GDP dissociation inhibitor alpha OS=Mus musculus OX=100! | 54 | 26 | 74 | 21 | 447 | 50,5 | 5,08 | 23033 | 89,52 | 17 | 16 | 1,034 | 1,093 | 0,81 | 1,217 | 0,737 | 1,019 | 0,946 | 1,028 | 1,11 | 0,11 | 3,7E-01 |
| Q3TNA1 | Xylulose kinase OS=Mus musculus OX=10090 GN=Xylb PE=1 SV= | 7 | 2 | 3 | 2 | 551 | 59,5 | 6,9 | 1832 | | 2 | | 1,212 | 1,24 | 0,938 | 0,928 | 1,955 | 1,044 | 1,084 | 1,143 | 0,83 | -0,17 | 3,7E-01 |
| Q64336 | T-box brain protein 1 OS=Mus musculus OX=10090 GN=Tbr1 PE= | 1 | 1 | 1 | 1 | 681 | 73,9 | 7,33 | 377 | | 1 | | 1,2 | 0,966 | 0,9 | 0,893 | 1,129 | 0,888 | 0,813 | 0,651 | 1,14 | 0,14 | 3,7E-01 |
| Q61315 | Adenomatous polyposis coli protein OS=Mus musculus OX=1009 | 3 | 7 | 7 | 7 | 2845 | 310,9 | 7,58 | 1635 | 8,39 | 3 | 4 | 1,009 | 1,003 | 0,946 | 0,932 | 1,178 | 0,892 | 1,121 | 0,968 | 0,94 | -0,06 | 3,7E-01 |
| Q9D1R9 | 60S ribosomal protein L34 OS=Mus musculus OX=10090 GN=Rpl | 15 | 2 | 6 | 2 | 117 | 13,3 | 11,47 | 2878 | | 2 | | 0,856 | 0,913 | 0,992 | 1,048 | 0,855 | 0,931 | 0,977 | 0,838 | 1,06 | 0,06 | 3,7E-01 |
| Q9JKC6 | Cell cycle exit and neuronal differentiation protein 1 OS=Mus m | 37 | 8 | 15 | 8 | 149 | 15 | 8,97 | 6299 | 18,89 | 5 | 5 | 1,04 | 0,768 | 1,153 | 0,767 | 0,73 | 0,974 | 0,82 | 0,771 | 1,13 | 0,13 | 3,7E-01 |
| P08551 | Neurofilament light polypeptide OS=Mus musculus OX=10090 G | 12 | 7 | 9 | 5 | 543 | 61,5 | 4,64 | 4360 | 4,86 | 5 | 2 | 0,965 | 0,924 | 1,068 | 0,917 | 0,752 | 0,699 | 1,152 | 0,855 | 1,12 | 0,12 | 3,7E-01 |
| Q8VDK1 | Deaminated glutathione amidase OS=Mus musculus OX=10090 ( | 5 | 1 | 3 | 1 | 323 | 35,7 | 7,94 | | 7,03 | | 1 | 1,155 | 1,245 | 1,055 | 1,031 | 1,596 | 0,871 | 1,375 | 1,264 | 0,88 | -0,12 | 3,7E-01 |
| P97371 | Proteasome activator complex subunit 1 OS=Mus musculus OX= | 37 | 8 | 10 | 8 | 249 | 28,7 | 5,97 | 5398 | 6,55 | 6 | 2 | 1,097 | 1,187 | 0,911 | 1,274 | 0,77 | 1,175 | 1,076 | 0,997 | 1,11 | 0,11 | 3,7E-01 |
| Q9WV85 | Nucleoside diphosphate kinase 3 OS=Mus musculus OX=10090 ( | 10 | 1 | 1 | 1 | 169 | 19,1 | 6,68 | 701 | | 1 | | 0,908 | 0,777 | 1,296 | 0,989 | 1,173 | 0,799 | 0,638 | 0,736 | 1,19 | 0,19 | 3,7E-01 |
| Q8VCV1 | Alpha/beta hydrolase domain-containing protein 17C OS=Mus n | 5 | 2 | 2 | 2 | 320 | 35,1 | 5,64 | 1224 | | 2 | | 0,926 | 0,983 | 0,863 | 1,066 | 1,165 | 1,079 | 0,991 | 0,888 | 0,93 | -0,07 | 3,7E-01 |
| Q5XJV6 | Serine/threonine-protein kinase LMTK3 OS=Mus musculus OX=1 | 1 | 1 | 1 | 1 | 1424 | 150,8 | 4,88 | 709 | | 1 | | 0,854 | 1,217 | 1,162 | 0,95 | 0,718 | 1,043 | 1,033 | 0,945 | 1,12 | 0,12 | 3,7E-01 |
| Q69ZK0 | Phosphatidylinositol 3,4,5-trisphosphate-dependent Rac exchan | 1 | 2 | 2 | 2 | 1650 | 184,8 | 6,29 | 410 | 3,52 | 1 | 1 | 1,197 | 0,952 | 1,023 | 1,117 | 1,283 | 1,082 | 1,19 | 1,034 | 0,93 | -0,07 | 3,7E-01 |
| Q9CYT6 | Adenylyl cyclase-associated protein 2 OS=Mus musculus OX=10( | 25 | 9 | 13 | 8 | 476 | 52,8 | 6,43 | 4233 | 4,81 | 7 | 2 | 1,044 | 1,04 | 1,076 | 1,129 | 1,284 | 1,066 | 1,023 | 1,153 | 0,95 | -0,05 | 3,7E-01 |
| Q8R2R9 | AP-3 complex subunit mu-2 OS=Mus musculus OX=10090 GN=A | 15 | 5 | 6 | 4 | 418 | 46,9 | 7,56 | 2750 | 3,1 | 5 | 1 | 0,892 | 1,004 | 0,917 | 0,93 | 1,196 | 0,965 | 0,892 | 0,963 | 0,93 | -0,07 | 3,7E-01 |
| Q02566 | Myosin-6 OS=Mus musculus OX=10090 GN=Myh6 PE=1 SV=2 | 5 | 12 | 15 | 1 | 1938 | 223,4 | 5,73 | 5927 | 18,98 | 7 | 5 | 0,619 | 0,658 | 0,331 | 0,5 | 16,274 | 0,556 | 0,185 | 0,46 | 0,12 | -0,88 | 3,7E-01 |
| Q14BI2 | Metabotropic glutamate receptor 2 OS=Mus musculus OX=1009 | 7 | 6 | 6 | 5 | 872 | 95,8 | 8,06 | 1717 | 4,64 | 3 | 3 | 1,025 | 0,893 | 0,881 | 0,921 | 0,805 | 0,964 | 0,938 | 0,806 | 1,06 | 0,06 | 3,7E-01 |
| Q9DCC8 | Mitochondrial import receptor subunit TOM20 homolog OS=Mu | 9 | 1 | 2 | 1 | 145 | 16,3 | 8,6 | 1359 | | 1 | | 0,881 | 0,874 | 0,67 | 0,796 | 1,076 | 0,588 | 0,994 | 1,041 | 0,87 | -0,13 | 3,7E-01 |
| O88910 | MAGUK p55 subfamily member 3 OS=Mus musculus OX=10090 | 5 | 3 | 3 | 3 | 568 | 64,4 | 6,34 | 1104 | 2,35 | 2 | 1 | 1,174 | 0,679 | 0,854 | 0,973 | 1,039 | 0,861 | 0,7 | 0,454 | 1,20 | 0,20 | 3,7E-01 |
| Q9EP72 | ER membrane protein complex subunit 7 OS=Mus musculus OX= | 10 | 2 | 2 | 2 | 241 | 26,3 | 9,23 | 1243 | | 2 | | 0,938 | 0,918 | 0,868 | 0,906 | 0,86 | 0,921 | 0,962 | 1,055 | 0,96 | -0,04 | 3,7E-01 |
| Q9QUR6 | Prolyl endopeptidase OS=Mus musculus OX=10090 GN=Prep PE | 27 | 19 | 35 | 19 | 710 | 80,7 | 5,73 | 16376 | 18,72 | 14 | 7 | 0,992 | 0,979 | 0,96 | 1,06 | 1,069 | 1,076 | 1,015 | 0,963 | 0,97 | -0,03 | 3,7E-01 |
| Q9WU28 | Prefoldin subunit 5 OS=Mus musculus OX=10090 GN=Pfdn5 PE= | 30 | 4 | 6 | 4 | 154 | 17,3 | 6,33 | 2086 | 5,81 | 2 | 2 | 1,011 | 1,084 | 1,053 | 1,044 | 0,838 | 1,144 | 0,889 | 1,044 | 1,07 | 0,07 | 3,7E-01 |
| Q8CIT9 | Suprabasin OS=Mus musculus OX=10090 GN=Sbsn PE=2 SV=1 | 2 | 2 | 2 | 2 | 700 | 72,3 | 7,31 | | 2,41 | | 2 | 0,901 | 1,059 | 0,858 | 0,743 | 1,137 | 0,676 | 0,826 | 3,155 | 0,61 | -0,39 | 3,7E-01 |

| | | | | | | | | | | | | | | | | | | | | | | | |
|---|---|---|---|---|---|---|---|---|---|---|---|---|---|---|---|---|---|---|---|---|---|---|---|
| Q9QYB1 | Chloride intracellular channel protein 4 OS=Mus musculus OX=1 | 6 | 1 | 2 | 1 | 253 | 28,7 | 5,59 | 1127 | | 1 | | 1,005 | 0,976 | 1,066 | 0,863 | 1,343 | 0,9 | 1,029 | 1,035 | 0,91 | -0,09 | 3,7E-01 |
| Q32NY4 | Metal transporter CNNM3 OS=Mus musculus OX=10090 GN=Cn | 2 | 1 | 2 | 1 | 713 | 76,2 | 5,58 | 1173 | | 1 | | 0,88 | 0,924 | 0,985 | 1,002 | 0,563 | 0,911 | 1,002 | 0,924 | 1,12 | 0,12 | 3,7E-01 |
| Q9D7A8 | Armadillo repeat-containing protein 1 OS=Mus musculus OX=10 | 4 | 2 | 2 | 2 | 282 | 31,2 | 5,57 | 1360 | | 2 | | 0,938 | 0,938 | 0,852 | 0,885 | 0,658 | 0,905 | 0,93 | 0,868 | 1,07 | 0,07 | 3,7E-01 |
| Q03157 | Amyloid-like protein 1 OS=Mus musculus OX=10090 GN=Aplp1 I | 3 | 2 | 3 | 2 | 653 | 72,7 | 5,67 | 1284 | 0 | 1 | 1 | 0,969 | 1,033 | 0,943 | 1,031 | 1,065 | 1,173 | 0,893 | 1,086 | 0,94 | -0,06 | 3,7E-01 |
| O70310 | Glycylpeptide N-tetradecanoyltransferase 1 OS=Mus musculus ( | 2 | 1 | 1 | 1 | 496 | 56,9 | 8 | 398 | | 1 | | 1,093 | 0,856 | 1,061 | 1,016 | 1,361 | 1,142 | 0,904 | 1,04 | 0,91 | -0,09 | 3,7E-01 |
| P18872 | Guanine nucleotide-binding protein G(o) subunit alpha OS=Mus | 49 | 18 | 69 | 17 | 354 | 40,1 | 5,53 | 23474 | 73,4 | 10 | 12 | 1,031 | 0,929 | 1,066 | 0,927 | 0,862 | 1,032 | 0,904 | 0,959 | 1,05 | 0,05 | 3,7E-01 |
| P14246 | Solute carrier family 2, facilitated glucose transporter member 2 | 3 | 1 | 1 | 1 | 523 | 57,1 | 6,73 | | 2,5 | | 1 | 1,248 | 1,507 | 1,475 | 1,482 | 2,24 | 1,594 | 1,199 | 1,543 | 0,87 | -0,13 | 3,7E-01 |
| Q9JKV1 | Proteasomal ubiquitin receptor ADRM1 OS=Mus musculus OX=1 | 9 | 4 | 4 | 4 | 407 | 42 | 5,07 | 1207 | 2,79 | 3 | 1 | 1,103 | 1,073 | 0,84 | 1,018 | 0,597 | 0,99 | 1,003 | 0,999 | 1,12 | 0,12 | 3,8E-01 |
| Q8K596 | Sodium/calcium exchanger 2 OS=Mus musculus OX=10090 GN=! | 15 | 13 | 28 | 13 | 921 | 100,6 | 5,12 | 8852 | 25,04 | 10 | 5 | 0,96 | 0,974 | 1,025 | 0,913 | 1,054 | 1,005 | 0,943 | 0,994 | 0,97 | -0,03 | 3,8E-01 |
| Q9D1K2 | V-type proton ATPase subunit F OS=Mus musculus OX=10090 GI | 30 | 4 | 7 | 4 | 119 | 13,4 | 5,82 | 2117 | 9,57 | 2 | 2 | 1,048 | 0,925 | 1,069 | 0,965 | 0,968 | 0,959 | 0,976 | 0,973 | 1,03 | 0,03 | 3,8E-01 |
| P68372 | Tubulin beta-4B chain OS=Mus musculus OX=10090 GN=Tubb4b | 62 | 27 | 441 | 2 | 445 | 49,8 | 4,89 | 188477 | 332,07 | 22 | 19 | 1,006 | 0,956 | 1,143 | 0,973 | 0,963 | 1,001 | 0,954 | 0,992 | 1,04 | 0,04 | 3,8E-01 |
| P48318 | Glutamate decarboxylase 1 OS=Mus musculus OX=10090 GN=Ga | 3 | 2 | 2 | 1 | 593 | 66,6 | 7,17 | 916 | | 2 | | 0,866 | 0,751 | 0,962 | 0,819 | 0,544 | 0,946 | 0,813 | 0,732 | 1,12 | 0,12 | 3,8E-01 |
| Q6PAM0 | 5'-AMP-activated protein kinase subunit beta-2 OS=Mus muscul | 4 | 1 | 1 | 1 | 271 | 30,2 | 6,46 | | 2,56 | | 1 | 0,997 | 0,97 | 0,943 | 0,992 | 1,209 | 0,866 | 1,006 | 1,103 | 0,93 | -0,07 | 3,8E-01 |
| Q9CRC8 | Leucine-rich repeat-containing protein 40 OS=Mus musculus OX | 8 | 5 | 5 | 5 | 602 | 68 | 6,89 | 1133 | 4,55 | 3 | 2 | 1,037 | 0,859 | 0,842 | 1,038 | 1,207 | 0,852 | 1,013 | 1,05 | 0,92 | -0,08 | 3,8E-01 |
| Q9D1T0 | Leucine-rich repeat and immunoglobulin-like domain-containing | 4 | 2 | 2 | 2 | 614 | 69,1 | 8,31 | 695 | 2,82 | 1 | 1 | 0,971 | 0,95 | 0,923 | 0,924 | 0,887 | 0,88 | 0,841 | 1,012 | 1,04 | 0,04 | 3,8E-01 |
| Q8CFI0 | E3 ubiquitin-protein ligase NEDD4-like OS=Mus musculus OX=10 | 6 | 4 | 4 | 4 | 1004 | 115,3 | 6,15 | 1179 | 0 | 3 | 1 | 1,024 | 1,048 | 0,944 | 1,257 | 0,99 | 0,842 | 1,058 | 1,062 | 1,08 | 0,08 | 3,8E-01 |
| Q3TJD7 | PDZ and LIM domain protein 7 OS=Mus musculus OX=10090 GN | 3 | 1 | 1 | 1 | 457 | 50,1 | 8,47 | | 2,37 | | 1 | 0,886 | 0,74 | 0,757 | 0,857 | 4,704 | 0,777 | 0,808 | 0,708 | 0,46 | -0,54 | 3,8E-01 |
| P50429 | Arylsulfatase B OS=Mus musculus OX=10090 GN=Arsb PE=1 SV= | 4 | 2 | 2 | 2 | 534 | 59,6 | 7,24 | 1018 | | 2 | | 1,282 | 1,104 | 0,863 | 1,007 | 1,373 | 1,076 | 1,048 | 1,196 | 0,91 | -0,09 | 3,8E-01 |
| Q9D428 | Golgin subfamily A member 7B OS=Mus musculus OX=10090 GN | 13 | 2 | 3 | 2 | 167 | 18,3 | 6,01 | 664 | 5,87 | 1 | 1 | 0,898 | 0,945 | 1,077 | 1,069 | 2,164 | 1,06 | 0,905 | 0,998 | 0,78 | -0,22 | 3,8E-01 |
| O35633 | Vesicular inhibitory amino acid transporter OS=Mus musculus O | 7 | 3 | 3 | 3 | 525 | 57,3 | 6,64 | 1804 | | 3 | | 0,99 | 0,958 | 0,948 | 0,867 | 1,276 | 0,999 | 0,905 | 0,924 | 0,92 | -0,08 | 3,8E-01 |
| Q5U458 | DnaJ homolog subfamily C member 11 OS=Mus musculus OX=10 | 10 | 5 | 6 | 5 | 559 | 63,2 | 8,32 | 625 | 9,73 | 2 | 3 | 1,048 | 1,193 | 0,954 | 0,982 | 1,206 | 0,956 | 1,161 | 1,146 | 0,93 | -0,07 | 3,8E-01 |
| Q3UVL4 | Vacuolar protein sorting-associated protein 51 homolog OS=Mu | 3 | 2 | 2 | 2 | 782 | 86,1 | 6,24 | 699 | 1,63 | 1 | 1 | 0,904 | 0,951 | 1,006 | 1,298 | 2,252 | 1,009 | 1,098 | 1,005 | 0,78 | -0,22 | 3,8E-01 |
| Q8BWF0 | Succinate-semialdehyde dehydrogenase, mitochondrial OS=Mus | 27 | 12 | 17 | 12 | 523 | 55,9 | 8,25 | 4511 | 23,36 | 8 | 4 | 0,973 | 0,995 | 0,981 | 0,995 | 0,973 | 0,891 | 0,979 | 1,005 | 1,02 | 0,02 | 3,8E-01 |
| P14576 | Signal recognition particle 54 kDa protein OS=Mus musculus OX | 7 | 4 | 4 | 4 | 504 | 55,7 | 8,75 | 1922 | 2,68 | 3 | 1 | 1,004 | 0,878 | 1,014 | 1,052 | 1,164 | 1,021 | 1,004 | 0,974 | 0,95 | -0,05 | 3,8E-01 |
| P16858 | Glyceraldehyde-3-phosphate dehydrogenase OS=Mus musculus | 69 | 27 | 133 | 27 | 333 | 35,8 | 8,25 | 44991 | 150,07 | 15 | 19 | 1,021 | 1,068 | 1,124 | 0,978 | 0,925 | 1,128 | 0,975 | 0,954 | 1,05 | 0,05 | 3,8E-01 |
| Q8R5J9 | PRA1 family protein 3 OS=Mus musculus OX=10090 GN=Arl6ip5 | 7 | 2 | 3 | 2 | 188 | 21,5 | 9,61 | 1170 | 0 | 1 | 1 | 0,965 | 0,973 | 0,998 | 0,942 | 0,861 | 0,915 | 1,024 | 0,942 | 1,04 | 0,04 | 3,8E-01 |
| P21107 | Tropomyosin alpha-3 chain OS=Mus musculus OX=10090 GN=Tp | 42 | 19 | 33 | 1 | 285 | 33 | 4,72 | 9534 | 45,8 | 8 | 13 | 1,131 | 0,956 | 1,093 | 1,249 | 1,449 | 1,322 | 0,973 | 1,141 | 0,91 | -0,09 | 3,8E-01 |
| P13020 | Gelsolin OS=Mus musculus OX=10090 GN=Gsn PE=1 SV=3 | 9 | 7 | 9 | 7 | 780 | 85,9 | 6,18 | 1319 | 15,18 | 2 | 5 | 0,941 | 1,127 | 0,911 | 0,886 | 1,951 | 0,844 | 0,948 | 1,1 | 0,80 | -0,20 | 3,8E-01 |
| Q91Z67 | SLIT-ROBO Rho GTPase-activating protein 2 OS=Mus musculus C | 22 | 18 | 32 | 17 | 1071 | 120,7 | 6,64 | 12774 | 25,59 | 14 | 7 | 0,921 | 0,967 | 1,018 | 0,853 | 1,19 | 0,903 | 0,968 | 0,97 | 0,93 | -0,07 | 3,8E-01 |
| Q5M8N4 | Epimerase family protein SDR39U1 OS=Mus musculus OX=1009( | 3 | 1 | 1 | 1 | 293 | 31,4 | 9,31 | 748 | | 1 | | 1,186 | 1,059 | 0,998 | 0,968 | 1,4 | 1,094 | 1,036 | 1,053 | 0,92 | -0,08 | 3,8E-01 |
| Q8BVA5 | Lipid droplet-associated hydrolase OS=Mus musculus OX=10090 | 4 | 1 | 1 | 1 | 326 | 37,3 | 8,28 | | 0 | | 1 | 0,675 | 1,112 | 0,892 | 0,636 | 2,217 | 0,722 | 0,891 | 0,87 | 0,71 | -0,29 | 3,8E-01 |
| P23591 | GDP-L-fucose synthase OS=Mus musculus OX=10090 GN=Tsta3 I | 11 | 3 | 4 | 3 | 321 | 35,9 | 6,74 | 1059 | 5,68 | 2 | 2 | 0,912 | 1,265 | 0,959 | 1,077 | 1,377 | 0,952 | 1,111 | 1,224 | 0,90 | -0,10 | 3,8E-01 |
| Q8BVU5 | ADP-ribose pyrophosphatase, mitochondrial OS=Mus musculus | 9 | 4 | 4 | 4 | 350 | 38,6 | 6,76 | 1758 | 2,12 | 3 | 1 | 0,902 | 0,897 | 1,071 | 1,026 | 1,802 | 0,967 | 0,854 | 1,095 | 0,83 | -0,17 | 3,8E-01 |
| P06728 | Apolipoprotein A-IV OS=Mus musculus OX=10090 GN=Apoa4 PE | 2 | 1 | 1 | 1 | 395 | 45 | 5,47 | | 2,31 | | 1 | 0,601 | 0,595 | 0,476 | 0,706 | 10,295 | 0,436 | 0,352 | 0,582 | 0,20 | -0,80 | 3,8E-01 |
| Q922E4 | Ethanolamine-phosphate cytidylyltransferase OS=Mus musculus | 17 | 5 | 8 | 5 | 404 | 45,2 | 6,58 | 2946 | 2,16 | 3 | 2 | 1,064 | 1,072 | 0,928 | 1,193 | 1,267 | 1,041 | 1,083 | 1,142 | 0,94 | -0,06 | 3,8E-01 |
| Q9D2G2 | Dihydrolipoyllysine-residue succinyltransferase component of 2- | 23 | 10 | 15 | 10 | 454 | 49 | 8,95 | 4081 | 17,9 | 4 | 6 | 0,986 | 0,965 | 1,016 | 0,951 | 0,909 | 1,021 | 0,948 | 0,935 | 1,03 | 0,03 | 3,8E-01 |
| Q9ERK4 | Exportin-2 OS=Mus musculus OX=10090 GN=Cse1l PE=1 SV=1 | 8 | 7 | 7 | 7 | 971 | 110,4 | 5,77 | 2195 | 10,63 | 3 | 4 | 1,061 | 1,016 | 1,147 | 1,038 | 1,053 | 1,128 | 1,033 | 1,305 | 0,94 | -0,06 | 3,8E-01 |
| P05201 | Aspartate aminotransferase, cytoplasmic OS=Mus musculus OX= | 33 | 13 | 22 | 13 | 413 | 46,2 | 7,14 | 7717 | 28,46 | 7 | 7 | 1,044 | 1,09 | 0,936 | 1,107 | 1,015 | 1,096 | 0,953 | 0,896 | 1,05 | 0,05 | 3,8E-01 |
| Q8BHY3 | Anoctamin-1 OS=Mus musculus OX=10090 GN=Ano1 PE=1 SV=2 | 1 | 1 | 1 | 1 | 960 | 110,8 | 7,83 | 285 | | 1 | | 0,958 | 0,805 | 0,853 | 0,963 | 1,442 | 0,884 | 0,857 | 0,937 | 0,87 | -0,13 | 3,8E-01 |
| Q8BVQ5 | Protein phosphatase methylesterase 1 OS=Mus musculus OX=10 | 10 | 3 | 4 | 3 | 386 | 42,2 | 5,97 | 744 | 4,76 | 1 | 2 | 0,806 | 0,837 | 0,875 | 0,956 | 0,575 | 0,875 | 0,949 | 0,743 | 1,11 | 0,11 | 3,8E-01 |
| Q62433 | Protein NDRG1 OS=Mus musculus OX=10090 GN=Ndrg1 PE=1 SV | 12 | 3 | 7 | 3 | 394 | 43 | 6,1 | 2372 | 13,25 | 3 | 1 | 1,055 | 1,04 | 0,972 | 1,144 | 0,81 | 1,111 | 1,018 | 1 | 1,07 | 0,07 | 3,8E-01 |
| Q9EPK7 | Exportin-7 OS=Mus musculus OX=10090 GN=Xpo7 PE=1 SV=3 | 2 | 2 | 2 | 2 | 1087 | 123,7 | 6,38 | 378 | 3,2 | 1 | 1 | 1,114 | 1,095 | 1,009 | 1,043 | 1,119 | 1,033 | 1,016 | 0,905 | 1,05 | 0,05 | 3,8E-01 |
| Q9QY76 | Vesicle-associated membrane protein-associated protein B OS=I | 26 | 7 | 10 | 6 | 243 | 26,9 | 7,78 | 5572 | 5,38 | 5 | 2 | 0,967 | 1,061 | 1,047 | 1,19 | 1,027 | 1,112 | 1,357 | 1,091 | 0,93 | -0,07 | 3,8E-01 |
| Q60676 | Serine/threonine-protein phosphatase 5 OS=Mus musculus OX= | 13 | 6 | 6 | 6 | 499 | 56,8 | 6,2 | 1437 | 8,84 | 2 | 4 | 1,039 | 0,816 | 0,945 | 1,011 | 0,707 | 0,977 | 0,99 | 0,823 | 1,09 | 0,09 | 3,8E-01 |
| P58404 | Striatin-4 OS=Mus musculus OX=10090 GN=Strn4 PE=1 SV=2 | 11 | 6 | 10 | 5 | 760 | 81,6 | 5,38 | 5185 | | 6 | | 1,076 | 0,843 | 1,206 | 0,954 | 1,042 | 1,065 | 0,771 | 0,781 | 1,11 | 0,11 | 3,8E-01 |
| Q8BFZ9 | Erlin-2 OS=Mus musculus OX=10090 GN=Erlin2 PE=1 SV=1 | 17 | 6 | 9 | 5 | 340 | 37,8 | 5,5 | 3152 | 7,55 | 3 | 3 | 1,049 | 1,011 | 1,024 | 1,141 | 1,063 | 1,042 | 1,117 | 1,144 | 0,97 | -0,03 | 3,9E-01 |
| Q8JZU2 | Tricarboxylate transport protein, mitochondrial OS=Mus muscul | 13 | 4 | 7 | 4 | 311 | 33,9 | 9,89 | 3801 | | 4 | | 1,001 | 0,942 | 0,955 | 0,917 | 1,224 | 0,905 | 0,948 | 1,011 | 0,93 | -0,07 | 3,9E-01 |
| B1AZP2 | Disks large-associated protein 4 OS=Mus musculus OX=10090 GI | 9 | 6 | 11 | 6 | 992 | 108 | 7,08 | 4587 | 8,29 | 5 | 2 | 0,844 | 0,823 | 1,382 | 0,839 | 0,853 | 0,8 | 0,877 | 0,843 | 1,15 | 0,15 | 3,9E-01 |
| Q99MI1 | ELKS/Rab6-interacting/CAST family member 1 OS=Mus musculu | 20 | 22 | 34 | 18 | 1120 | 128,3 | 5,87 | 17829 | 9,3 | 17 | 6 | 1,016 | 0,961 | 1,002 | 0,987 | 0,759 | 0,962 | 1,058 | 0,949 | 1,06 | 0,06 | 3,9E-01 |
| Q9JKB1 | Ubiquitin carboxyl-terminal hydrolase isozyme L3 OS=Mus musc | 26 | 6 | 7 | 6 | 230 | 26,1 | 5,05 | 3015 | 2,72 | 5 | 1 | 1,021 | 1,156 | 1,097 | 1,22 | 1,164 | 1,133 | 1,184 | 1,177 | 0,96 | -0,04 | 3,9E-01 |
| O08788 | Dynactin subunit 1 OS=Mus musculus OX=10090 GN=Dctn1 PE= | 28 | 32 | 54 | 30 | 1281 | 141,6 | 5,9 | 25233 | 35,12 | 21 | 13 | 1,012 | 1,014 | 0,997 | 1,012 | 0,859 | 0,957 | 1,054 | 1,008 | 1,04 | 0,04 | 3,9E-01 |
| Q91YR1 | Twinfilin-1 OS=Mus musculus OX=10090 GN=Twf1 PE=1 SV=2 | 13 | 5 | 7 | 5 | 350 | 40,1 | 6,67 | 2193 | 8,45 | 3 | 2 | 1,026 | 1,073 | 0,98 | 1,069 | 0,906 | 0,961 | 1,097 | 1,012 | 1,04 | 0,04 | 3,9E-01 |
| Q6P5F7 | Protein tweety homolog 3 OS=Mus musculus OX=10090 GN=Tty | 6 | 3 | 4 | 3 | 524 | 57,7 | 5,85 | | 9,29 | | 3 | 0,97 | 0,922 | 1,025 | 0,813 | 0,715 | 0,957 | 0,869 | 0,927 | 1,08 | 0,08 | 3,9E-01 |
| Q9CWG8 | Protein arginine methyltransferase NDUFAF7, mitochondrial OS | 5 | 2 | 2 | 2 | 436 | 48,4 | 6,95 | 1276 | | 2 | | 1,193 | 1,009 | 1,05 | 1,174 | 1,237 | 1,011 | 1,174 | 1,283 | 0,94 | -0,06 | 3,9E-01 |
| Q9D1A2 | Cytosolic non-specific dipeptidase OS=Mus musculus OX=10090 | 22 | 10 | 15 | 10 | 475 | 52,7 | 5,66 | 7582 | 9,9 | 9 | 2 | 0,988 | 0,989 | 1,065 | 1,093 | 0,998 | 1,021 | 0,95 | 1,042 | 1,03 | 0,03 | 3,9E-01 |
| Q91YJ5 | Translation initiation factor IF-2, mitochondrial OS=Mus muscul | 4 | 2 | 2 | 2 | 727 | 81,2 | 7,24 | 665 | 3,13 | 1 | 1 | 0,798 | 0,86 | 1,113 | 0,862 | 0,853 | 0,962 | 0,727 | 0,766 | 1,10 | 0,10 | 3,9E-01 |
| O54829 | Regulator of G-protein signaling 7 OS=Mus musculus OX=10090 | 7 | 3 | 4 | 1 | 469 | 54,8 | 8,13 | 1131 | 3,09 | 2 | 2 | 0,717 | 0,715 | 0,957 | 0,533 | 2,218 | 0,589 | 0,954 | 0,624 | 0,67 | -0,33 | 3,9E-01 |
| Q8BNU0 | Armadillo repeat-containing protein 6 OS=Mus musculus OX=10 | 18 | 8 | 11 | 8 | 468 | 50,7 | 6 | 3168 | 14,15 | 5 | 4 | 1 | 0,894 | 1,261 | 0,872 | 0,718 | 1,11 | 0,924 | 0,819 | 1,13 | 0,13 | 3,9E-01 |
| Q60967 | Bifunctional 3'-phosphoadenosine 5'-phosphosulfate synthase 1 | 3 | 2 | 2 | 2 | 624 | 70,7 | 6,77 | 1107 | | 2 | | 0,969 | 0,931 | 0,858 | 0,999 | 1,001 | 0,603 | 1,003 | 0,771 | 1,11 | 0,11 | 3,9E-01 |
| Q9EQ51 | Vomeronasal type-1 receptor 47 OS=Mus musculus OX=10090 G | 4 | 1 | 1 | 1 | 310 | 35,4 | 9,29 | | 0 | | 1 | 1,155 | 1,158 | 1,06 | 1,169 | 1,244 | 0,95 | 0,964 | 1,11 | 1,06 | 0,06 | 3,9E-01 |
| Q9CYH6 | Ribosome biogenesis regulatory protein homolog OS=Mus musc | 2 | 1 | 1 | 1 | 365 | 41,5 | 10,77 | 531 | | 1 | | 0,91 | 0,975 | 1,006 | 0,88 | 0,912 | 0,931 | 0,836 | 0,95 | 1,04 | 0,04 | 3,9E-01 |
| P62254 | Ubiquitin-conjugating enzyme E2 G1 OS=Mus musculus OX=100 | 4 | 1 | 1 | 1 | 170 | 19,5 | 5,3 | 715 | | 1 | | 0,865 | 0,933 | 1,16 | 1,214 | 1,004 | 1,163 | 1,177 | 1,181 | 0,92 | -0,08 | 3,9E-01 |
| Q14CH7 | Alanine--tRNA ligase, mitochondrial OS=Mus musculus OX=1009 | 3 | 2 | 2 | 2 | 980 | 106,7 | 6,18 | 979 | | 2 | | 1,017 | 1,641 | 1,212 | 1,315 | 1,509 | 1,449 | 1,349 | 1,38 | 0,91 | -0,09 | 3,9E-01 |
| Q9DC07 | LIM zinc-binding domain-containing Nebulette OS=Mus musculu | 10 | 3 | 3 | 2 | 270 | 31,1 | 8,31 | 915 | 0 | 2 | 1 | 1,074 | 1,208 | 1,188 | 1,078 | 1,448 | 1,241 | 0,992 | 1,237 | 0,92 | -0,08 | 3,9E-01 |
| Q8R555 | Cartilage acidic protein 1 OS=Mus musculus OX=10090 GN=Crta | 1 | 1 | 1 | 1 | 646 | 70,3 | 5,14 | 380 | | 1 | | 0,871 | 0,965 | 0,984 | 1,033 | 1,471 | 0,836 | 1,114 | 0,958 | 0,88 | -0,12 | 3,9E-01 |

| | | | | | | | | | | | | | | | | | | | | | | | |
|---|---|---|---|---|---|---|---|---|---|---|---|---|---|---|---|---|---|---|---|---|---|---|---|
| Q8BZ36 | RAD50-interacting protein 1 OS=Mus musculus OX=10090 GN=R | 1 | 1 | 1 | 1 | 792 | 90 | 5,14 | 311 | | 1 | | 0,805 | 0,791 | 0,894 | 0,955 | 1,032 | 0,918 | 0,818 | 0,895 | 0,94 | -0,06 | 3,9E-01 |
| Q99LB2 | Dehydrogenase/reductase SDR family member 4 OS=Mus muscu | 23 | 5 | 6 | 5 | 279 | 29,9 | 9,38 | 3600 | | 5 | | 1,096 | 1,06 | 1,157 | 1,334 | 1,104 | 1,153 | 1,071 | 1,084 | 1,05 | 0,05 | 3,9E-01 |
| Q91XM9 | Disks large homolog 2 OS=Mus musculus OX=10090 GN=Dlg2 PE | 17 | 13 | 19 | 9 | 852 | 94,8 | 6,24 | 8591 | 6,94 | 10 | 3 | 0,995 | 0,973 | 0,981 | 0,951 | 1,506 | 0,964 | 0,96 | 0,972 | 0,89 | -0,11 | 3,9E-01 |
| Q9DBR7 | Protein phosphatase 1 regulatory subunit 12A OS=Mus musculu | 4 | 4 | 5 | 4 | 1029 | 114,9 | 5,49 | 1126 | 5,65 | 2 | 2 | 1,074 | 1,026 | 1,086 | 1,134 | 1,155 | 0,808 | 0,964 | 1,096 | 1,07 | 0,07 | 3,9E-01 |
| Q80V91 | Probable E3 ubiquitin-protein ligase DTX3 OS=Mus musculus OX | 10 | 2 | 2 | 2 | 347 | 38 | 8,73 | 783 | 2,46 | 1 | 1 | 0,982 | 1,142 | 0,789 | 0,972 | 0,992 | 1,03 | 1,178 | 0,997 | 0,93 | -0,07 | 3,9E-01 |
| P06745 | Glucose-6-phosphate isomerase OS=Mus musculus OX=10090 G | 32 | 17 | 27 | 16 | 558 | 62,7 | 8,13 | 9303 | 32,63 | 9 | 10 | 0,98 | 0,945 | 1,097 | 0,952 | 0,755 | 1,098 | 0,967 | 0,851 | 1,08 | 0,08 | 3,9E-01 |
| P67984 | 60S ribosomal protein L22 OS=Mus musculus OX=10090 GN=Rpl | 37 | 5 | 6 | 5 | 128 | 14,8 | 9,19 | 2352 | 4,09 | 5 | 1 | 0,964 | 1,051 | 0,82 | 1,001 | 1,116 | 0,824 | 1,229 | 1,032 | 0,91 | -0,09 | 3,9E-01 |
| Q80VP1 | Epsin-1 OS=Mus musculus OX=10090 GN=Epn1 PE=1 SV=3 | 2 | 1 | 1 | 1 | 575 | 60,2 | 4,81 | 780 | | 1 | | 0,79 | 0,866 | 0,708 | 0,815 | 0,758 | 0,866 | 0,82 | 0,902 | 0,95 | -0,05 | 3,9E-01 |
| Q80TF4 | Kelch-like protein 13 OS=Mus musculus OX=10090 GN=Klhl13 PE | 2 | 1 | 1 | 1 | 654 | 73,7 | 6,9 | 505 | | 1 | | 0,831 | 0,896 | 1,274 | 1,02 | 1,713 | 1,026 | 0,92 | 1,109 | 0,84 | -0,16 | 3,9E-01 |
| Q8CHI8 | E1A-binding protein p400 OS=Mus musculus OX=10090 GN=Ep4 | 0 | 1 | 1 | 1 | 3072 | 337 | 9,14 | 374 | | 1 | | 0,898 | 0,715 | 1,305 | 0,839 | 1,324 | 1,189 | 0,961 | 0,883 | 0,86 | -0,14 | 3,9E-01 |
| O54833 | Casein kinase II subunit alpha' OS=Mus musculus OX=10090 GN= | 9 | 3 | 3 | 3 | 350 | 41,2 | 8,56 | 886 | 4,79 | 1 | 2 | 0,956 | 0,919 | 0,93 | 0,903 | 0,978 | 0,869 | 0,913 | 0,815 | 1,04 | 0,04 | 3,9E-01 |
| Q3U1F9 | Phosphoprotein associated with glycosphingolipid-enriched mic | 4 | 1 | 1 | 1 | 429 | 46,5 | 4,81 | 742 | | 1 | | 0,795 | 1,04 | 0,705 | 1,019 | 0,891 | 0,955 | 0,969 | 1,08 | 0,91 | -0,09 | 3,9E-01 |
| Q8K4P8 | E3 ubiquitin-protein ligase HECW1 OS=Mus musculus OX=10090 | 2 | 4 | 4 | 4 | 1604 | 179,4 | 5,39 | 1165 | 4,93 | 2 | 2 | 1,052 | 0,923 | 1,027 | 0,973 | 0,968 | 1,041 | 0,975 | 1,273 | 0,93 | -0,07 | 4,0E-01 |
| Q80VJ3 | 2'-deoxynucleoside 5'-phosphate N-hydrolase 1 OS=Mus muscul | 8 | 1 | 1 | 1 | 173 | 19 | 5,38 | 347 | | 1 | | 0,73 | 0,916 | 0,747 | 1,047 | 1,195 | 0,773 | 0,914 | 0,981 | 0,89 | -0,11 | 4,0E-01 |
| Q5SNZ0 | Girdin OS=Mus musculus OX=10090 GN=Ccdc88a PE=1 SV=2 | 11 | 20 | 22 | 20 | 1873 | 215,8 | 6,24 | 7580 | 28,67 | 12 | 8 | 0,975 | 0,937 | 0,968 | 0,962 | 0,916 | 0,909 | 0,97 | 0,976 | 1,02 | 0,02 | 4,0E-01 |
| P48774 | Glutathione S-transferase Mu 5 OS=Mus musculus OX=10090 GI | 13 | 3 | 3 | 2 | 224 | 26,6 | 7,21 | 524 | 5,81 | 1 | 2 | 0,932 | 0,891 | 0,99 | 0,83 | 0,609 | 0,932 | 0,888 | 0,911 | 1,09 | 0,09 | 4,0E-01 |
| Q9Z2H5 | Band 4.1-like protein 1 OS=Mus musculus OX=10090 GN=Epb41 | 29 | 23 | 68 | 22 | 879 | 98,3 | 5,62 | 33966 | 28,54 | 18 | 7 | 1,029 | 0,944 | 0,999 | 0,917 | 0,859 | 0,969 | 0,953 | 0,973 | 1,04 | 0,04 | 4,0E-01 |
| Q9EQF6 | Dihydropyrimidinase-related protein 5 OS=Mus musculus OX=1( | 67 | 32 | 113 | 31 | 564 | 61,5 | 7,09 | 51733 | 98,94 | 24 | 13 | 1,03 | 0,978 | 0,998 | 0,967 | 0,796 | 1,06 | 0,951 | 0,961 | 1,05 | 0,05 | 4,0E-01 |
| P49442 | Inositol polyphosphate 1-phosphatase OS=Mus musculus OX=1( | 7 | 2 | 2 | 2 | 396 | 43,3 | 5,01 | 782 | 2,33 | 1 | 1 | 0,83 | 1,03 | 0,953 | 1,023 | 0,75 | 1,035 | 0,952 | 0,807 | 1,08 | 0,08 | 4,0E-01 |
| Q9CXJ4 | ATP-binding cassette sub-family B member 8, mitochondrial OS= | 2 | 1 | 1 | 1 | 717 | 78 | 9,07 | 441 | | 1 | | 0,667 | 0,88 | 0,725 | 1,153 | 1,266 | 0,836 | 0,809 | 1,071 | 0,86 | -0,14 | 4,0E-01 |
| P59017 | Bcl-2-like protein 13 OS=Mus musculus OX=10090 GN=Bcl2l13 P | 11 | 4 | 4 | 4 | 434 | 46,7 | 4,59 | 580 | 8,53 | 1 | 3 | 1,073 | 1,028 | 0,962 | 0,991 | 1,038 | 1,035 | 0,877 | 0,936 | 1,04 | 0,04 | 4,0E-01 |
| Q8R395 | COMM domain-containing protein 5 OS=Mus musculus OX=100 | 4 | 1 | 1 | 1 | 224 | 24,5 | 8,06 | | 0 | | 1 | 1,22 | 0,678 | 0,88 | 0,889 | 1,835 | 0,842 | 0,836 | 1,102 | 0,79 | -0,21 | 4,0E-01 |
| P62996 | Transformer-2 protein homolog beta OS=Mus musculus OX=100 | 6 | 1 | 1 | 1 | 288 | 33,6 | 11,25 | 516 | | 1 | | 0,79 | 1,029 | 0,694 | 0,913 | 1,004 | 0,818 | 0,82 | 1,221 | 0,89 | -0,11 | 4,0E-01 |
| Q9CQF3 | Cleavage and polyadenylation specificity factor subunit 5 OS=M | 7 | 1 | 2 | 1 | 227 | 26,2 | 8,82 | 804 | | 1 | | 0,688 | 0,815 | 1,351 | 0,643 | 1,741 | 0,859 | 1,291 | 0,661 | 0,77 | -0,23 | 4,0E-01 |
| Q64462 | Cytochrome P450 4B1 OS=Mus musculus OX=10090 GN=Cyp4b1 | 2 | 1 | 1 | 1 | 511 | 58,9 | 8,44 | 418 | | 1 | | 1,102 | 1,33 | 1,212 | 1,124 | 1,026 | 0,926 | 1,247 | 1,228 | 1,08 | 0,08 | 4,0E-01 |
| Q8R0S2 | IQ motif and SEC7 domain-containing protein 1 OS=Mus muscul | 10 | 9 | 11 | 8 | 961 | 107,9 | 7,14 | 3883 | 4,79 | 6 | 3 | 1,01 | 0,983 | 0,968 | 0,938 | 0,823 | 0,833 | 1,036 | 0,999 | 1,06 | 0,06 | 4,0E-01 |
| Q8BH59 | Calcium-binding mitochondrial carrier protein Aralar1 OS=Mus r | 14 | 8 | 16 | 6 | 677 | 74,5 | 8,25 | 6779 | 7,86 | 6 | 3 | 0,992 | 0,994 | 0,934 | 0,916 | 0,945 | 0,895 | 0,964 | 0,942 | 1,02 | 0,02 | 4,0E-01 |
| P14206 | 40S ribosomal protein SA OS=Mus musculus OX=10090 GN=Rps | 18 | 4 | 13 | 4 | 295 | 32,8 | 4,87 | 5957 | 6,31 | 3 | 1 | 1,004 | 1,009 | 0,987 | 1,02 | 1,541 | 0,921 | 1,054 | 1,01 | 0,89 | -0,11 | 4,0E-01 |
| P63216 | Guanine nucleotide-binding protein G(I)/G(S)/G(O) subunit gam | 23 | 1 | 5 | 1 | 75 | 8,3 | 7,78 | 1477 | 8,19 | 1 | 1 | 1,156 | 1,04 | 1,023 | 0,833 | 1,028 | 1,2 | 1,065 | 1,041 | 0,93 | -0,07 | 4,0E-01 |
| Q9DCZ4 | MICOS complex subunit Mic26 OS=Mus musculus OX=10090 GN | 13 | 2 | 4 | 2 | 198 | 22,6 | 9,25 | 904 | 8,43 | 1 | 2 | 1,051 | 1,134 | 0,918 | 1,014 | 1,301 | 1,016 | 1,162 | 0,96 | 0,93 | -0,07 | 4,0E-01 |
| Q3U1J4 | DNA damage-binding protein 1 OS=Mus musculus OX=10090 GN | 12 | 12 | 12 | 12 | 1140 | 126,8 | 5,26 | 4200 | 12,27 | 7 | 5 | 0,926 | 0,893 | 0,94 | 1,007 | 0,804 | 0,968 | 0,929 | 0,911 | 1,04 | 0,04 | 4,0E-01 |
| P04117 | Fatty acid-binding protein, adipocyte OS=Mus musculus OX=100 | 11 | 1 | 1 | 1 | 132 | 14,6 | 8,4 | 901 | | 1 | | 4,789 | 4,948 | 4,503 | 3,961 | 5,711 | 3,413 | 5,056 | 6,557 | 0,88 | -0,12 | 4,0E-01 |
| Q80X90 | Filamin-B OS=Mus musculus OX=10090 GN=Flnb PE=1 SV=3 | 1 | 2 | 2 | 1 | 2602 | 277,7 | 5,71 | 675 | 1,6 | 1 | 1 | 1,083 | 1,001 | 0,981 | 1,073 | 1,003 | 1,31 | 1,105 | 0,999 | 0,94 | -0,06 | 4,0E-01 |
| P62908 | 40S ribosomal protein S3 OS=Mus musculus OX=10090 GN=Rps | 54 | 12 | 23 | 12 | 243 | 26,7 | 9,66 | 6990 | 34,14 | 5 | 8 | 0,991 | 0,948 | 1,047 | 1,045 | 0,861 | 0,997 | 1,005 | 1,012 | 1,04 | 0,04 | 4,0E-01 |
| Q61644 | Protein kinase C and casein kinase substrate in neurons protein | 24 | 9 | 15 | 8 | 441 | 50,5 | 5,24 | 4632 | 20,68 | 7 | 2 | 1,051 | 0,968 | 1,02 | 0,967 | 0,872 | 1,002 | 1,064 | 0,884 | 1,05 | 0,05 | 4,0E-01 |
| D3YZU1 | SH3 and multiple ankyrin repeat domains protein 1 OS=Mus mu | 4 | 7 | 9 | 6 | 2167 | 226,2 | 8,34 | 2997 | 2,05 | 6 | 1 | 0,966 | 0,998 | 1,067 | 1,043 | 1,143 | 0,979 | 1,181 | 0,98 | 0,95 | -0,05 | 4,0E-01 |
| P40240 | CD9 antigen OS=Mus musculus OX=10090 GN=Cd9 PE=1 SV=2 | 3 | 1 | 1 | 1 | 226 | 25,2 | 7,23 | | 2,89 | | 1 | 0,843 | 1,202 | 0,606 | 1,107 | 0,643 | 0,615 | 0,984 | 0,923 | 1,19 | 0,19 | 4,0E-01 |
| Q8BLK3 | Limbic system-associated membrane protein OS=Mus musculus | 24 | 7 | 32 | 7 | 341 | 38,1 | 6,65 | 14983 | 16,86 | 5 | 3 | 1,021 | 1,085 | 0,891 | 1,027 | 1,035 | 0,989 | 0,885 | 0,925 | 1,05 | 0,05 | 4,0E-01 |
| P32921 | Tryptophan--tRNA ligase, cytoplasmic OS=Mus musculus OX=10 | 15 | 6 | 7 | 6 | 481 | 54,3 | 6,89 | 2791 | 5,77 | 4 | 2 | 0,863 | 0,986 | 0,918 | 0,867 | 0,834 | 0,962 | 1,04 | 0,986 | 0,95 | -0,05 | 4,0E-01 |
| Q5SSH7 | Zinc finger ZZ-type and EF-hand domain-containing protein 1 OS | 1 | 1 | 1 | 1 | 2924 | 328,1 | 6,11 | 420 | | 1 | | 1,068 | 1,637 | 1,089 | 0,988 | 1,085 | 1,49 | 1,275 | 1,612 | 0,88 | -0,12 | 4,0E-01 |
| Q9QXS1 | Plectin OS=Mus musculus OX=10090 GN=Plec PE=1 SV=3 | 10 | 44 | 49 | 44 | 4691 | 533,9 | 5,96 | 21699 | 21,63 | 34 | 10 | 1,049 | 1,003 | 1,055 | 1 | 1,25 | 0,994 | 1,021 | 1,057 | 0,95 | -0,05 | 4,0E-01 |
| P84084 | ADP-ribosylation factor 5 OS=Mus musculus OX=10090 GN=Arf5 | 31 | 6 | 8 | 3 | 180 | 20,5 | 6,79 | 3213 | 2,06 | 4 | 2 | 1,107 | 0,96 | 1,177 | 1,027 | 1,02 | 0,932 | 1,115 | 0,985 | 1,05 | 0,05 | 4,0E-01 |
| O54724 | Caveolae-associated protein 1 OS=Mus musculus OX=10090 GN | 3 | 1 | 2 | 1 | 392 | 43,9 | 5,52 | | 5,5 | | 1 | 0,914 | 1,113 | 0,773 | 1,067 | 0,516 | 1,004 | 0,892 | 0,966 | 1,14 | 0,14 | 4,0E-01 |
| Q8R2Y2 | Cell surface glycoprotein MUC18 OS=Mus musculus OX=10090 ( | 6 | 2 | 2 | 2 | 648 | 71,5 | 5,83 | 1269 | | 2 | | 0,721 | 0,613 | 0,769 | 0,775 | 0,96 | 0,595 | 0,788 | 0,839 | 0,90 | -0,10 | 4,0E-01 |
| Q61733 | 28S ribosomal protein S31, mitochondrial OS=Mus musculus OX | 7 | 2 | 3 | 2 | 384 | 43,9 | 8,51 | | 8,42 | | 2 | 1,071 | 0,783 | 1,141 | 1,142 | 1,05 | 1,128 | 1,043 | 1,281 | 0,92 | -0,08 | 4,0E-01 |
| Q8R164 | Valacyclovir hydrolase OS=Mus musculus OX=10090 GN=Bphl PE | 12 | 3 | 5 | 3 | 291 | 32,8 | 8,94 | 1601 | 4,06 | 2 | 1 | 0,947 | 1,062 | 0,931 | 0,959 | 1,328 | 0,816 | 1,061 | 1,084 | 0,91 | -0,09 | 4,0E-01 |
| Q8BRJ4 | Protein phosphatase 1 regulatory subunit 3E OS=Mus musculus | 5 | 1 | 1 | 1 | 279 | 30,6 | 11,15 | 434 | | 1 | | 0,746 | 0,67 | 0,776 | 1,122 | 1,192 | 1,147 | 0,699 | 0,836 | 0,86 | -0,14 | 4,0E-01 |
| P97792 | Coxsackievirus and adenovirus receptor homolog OS=Mus musc | 42 | 14 | 52 | 14 | 365 | 39,9 | 6,96 | 18035 | 59,58 | 9 | 11 | 1,069 | 0,975 | 0,918 | 0,963 | 0,988 | 0,999 | 0,868 | 0,91 | 1,04 | 0,04 | 4,0E-01 |
| P63011 | Ras-related protein Rab-3A OS=Mus musculus OX=10090 GN=Ra | 37 | 7 | 20 | 4 | 220 | 25 | 5,03 | 11997 | 1,71 | 6 | 1 | 0,984 | 1,065 | 0,973 | 0,952 | 0,913 | 0,997 | 0,996 | 0,954 | 1,03 | 0,03 | 4,1E-01 |
| Q9D554 | Splicing factor 3A subunit 3 OS=Mus musculus OX=10090 GN=Sf | 2 | 1 | 1 | 1 | 501 | 58,8 | 5,34 | 430 | | 1 | | 1,074 | 1,063 | 0,946 | 1,215 | 0,877 | 0,91 | 1,159 | 1,047 | 1,08 | 0,08 | 4,1E-01 |
| P70206 | Plexin-A1 OS=Mus musculus OX=10090 GN=Plxna1 PE=1 SV=1 | 18 | 30 | 47 | 23 | 1894 | 211 | 6,9 | 20159 | 28,43 | 22 | 9 | 0,979 | 0,933 | 1,013 | 0,893 | 1,192 | 0,974 | 0,954 | 0,933 | 0,94 | -0,06 | 4,1E-01 |
| Q8K212 | Phosphofurin acidic cluster sorting protein 1 OS=Mus musculus | 13 | 8 | 11 | 8 | 961 | 104,8 | 7,74 | 6110 | 3,8 | 7 | 2 | 0,988 | 0,921 | 0,958 | 0,865 | 0,99 | 0,91 | 0,91 | 1,131 | 0,95 | -0,05 | 4,1E-01 |
| P97390 | Vacuolar protein sorting-associated protein 45 OS=Mus musculu | 3 | 2 | 3 | 2 | 570 | 65 | 8,25 | 1329 | | 2 | | 0,963 | 0,695 | 1,167 | 0,703 | 1,508 | 0,97 | 0,959 | 0,783 | 0,84 | -0,16 | 4,1E-01 |
| Q9CZW5 | Mitochondrial import receptor subunit TOM70 OS=Mus musculu | 29 | 14 | 14 | 14 | 611 | 67,5 | 7,53 | 5693 | 18,51 | 8 | 6 | 1,023 | 0,951 | 0,992 | 0,924 | 0,782 | 1,023 | 0,959 | 0,928 | 1,05 | 0,05 | 4,1E-01 |
| Q91ZP9 | N-terminal EF-hand calcium-binding protein 2 OS=Mus musculus | 14 | 3 | 3 | 3 | 389 | 43,4 | 5,3 | 1520 | | 3 | | 0,792 | 0,912 | 0,799 | 1,259 | 1,384 | 0,912 | 0,894 | 1,138 | 0,87 | -0,13 | 4,1E-01 |
| G3X982 | Aldehyde oxidase 3 OS=Mus musculus OX=10090 GN=Aox3 PE= | 3 | 2 | 2 | 2 | 1335 | 146,8 | 6,46 | 1275 | | 2 | | 1,433 | 2,508 | 1,367 | 1,302 | 1,336 | 2,048 | 2,029 | 2,546 | 0,83 | -0,17 | 4,1E-01 |
| P12787 | Cytochrome c oxidase subunit 5A, mitochondrial OS=Mus muscu | 28 | 6 | 22 | 6 | 146 | 16,1 | 6,54 | 9214 | 21,12 | 3 | 4 | 0,992 | 1,022 | 0,924 | 1,103 | 0,98 | 1,001 | 0,96 | 0,964 | 1,03 | 0,03 | 4,1E-01 |
| Q62193 | Replication protein A 32 kDa subunit OS=Mus musculus OX=100 | 3 | 1 | 1 | 1 | 270 | 29,7 | 6,15 | | 2,34 | | 1 | 1,198 | 1,417 | 1,001 | 1,383 | 1,24 | 1,012 | 0,966 | 1,322 | 1,10 | 0,10 | 4,1E-01 |
| P62751 | 60S ribosomal protein L23a OS=Mus musculus OX=10090 GN=R | 42 | 10 | 19 | 10 | 156 | 17,7 | 10,45 | 8056 | 17,41 | 6 | 5 | 1,034 | 0,984 | 1,009 | 1,136 | 1,506 | 1,07 | 1,018 | 1,009 | 0,90 | -0,10 | 4,1E-01 |
| Q8CI43 | Myosin light chain 6B OS=Mus musculus OX=10090 GN=Myl6b F | 12 | 2 | 7 | 1 | 207 | 22,7 | 5,53 | 2272 | 12,2 | 1 | 2 | 0,855 | 1,214 | 1,114 | 0,892 | 1,777 | 0,774 | 1,151 | 1,172 | 0,84 | -0,16 | 4,1E-01 |
| P48193 | Protein 4.1 OS=Mus musculus OX=10090 GN=Epb41 PE=1 SV=2 | 8 | 6 | 6 | 5 | 858 | 95,9 | 5,6 | 3419 | | 6 | | 1,013 | 1,11 | 0,997 | 1,203 | 1,045 | 1,032 | 1,039 | 1,037 | 1,04 | 0,04 | 4,1E-01 |
| P45376 | Aldo-keto reductase family 1 member B1 OS=Mus musculus OX= | 24 | 9 | 15 | 9 | 316 | 35,7 | 7,18 | 8799 | 2,06 | 8 | 1 | 0,996 | 0,941 | 1,012 | 1,012 | 1,082 | 1,024 | 1,042 | 0,937 | 0,97 | -0,03 | 4,1E-01 |
| P52196 | Thiosulfate sulfurtransferase OS=Mus musculus OX=10090 GN= | 26 | 7 | 11 | 7 | 297 | 33,4 | 7,85 | 4396 | 8,51 | 6 | 2 | 0,971 | 0,954 | 0,968 | 0,989 | 1,001 | 0,842 | 0,993 | 0,91 | 1,04 | 0,04 | 4,1E-01 |
| P29416 | Beta-hexosaminidase subunit alpha OS=Mus musculus OX=1009 | 4 | 2 | 2 | 2 | 528 | 60,6 | 6,54 | 1051 | | 2 | | 1,188 | 1,141 | 1,294 | 0,985 | 1,403 | 1,293 | 1,155 | 1,093 | 0,93 | -0,07 | 4,1E-01 |
| Q9DBG1 | Sterol 26-hydroxylase, mitochondrial OS=Mus musculus OX=100 | 2 | 1 | 1 | 1 | 533 | 60,7 | 9,14 | 634 | | 1 | | 0,965 | 0,73 | 1,223 | 1,185 | 2,072 | 0,817 | 1,351 | 0,943 | 0,79 | -0,21 | 4,1E-01 |

| | | | | | | | | | | | | | | | | | | | | | | | |
|---|---|---|---|---|---|---|---|---|---|---|---|---|---|---|---|---|---|---|---|---|---|---|---|
| O88444 | Adenylate cyclase type 1 OS=Mus musculus OX=10090 GN=Adcy | 2 | 2 | 2 | 2 | 1118 | 123,3 | 8,47 | 634 | 0 | 1 | 1 | 1,126 | 1,22 | 1,203 | 0,891 | 0,584 | 1,064 | 1,217 | 1,024 | 1,14 | 0,14 | 4,1E-01 |
| P24547 | Inosine-5'-monophosphate dehydrogenase 2 OS=Mus musculus | 21 | 7 | 9 | 6 | 514 | 55,8 | 7,28 | 3744 | 6,53 | 6 | 2 | 1,035 | 1,061 | 1,082 | 1,232 | 1,37 | 1,116 | 1,111 | 1,094 | 0,94 | -0,06 | 4,1E-01 |
| P48758 | Carbonyl reductase [NADPH] 1 OS=Mus musculus OX=10090 GN | 22 | 6 | 9 | 5 | 277 | 30,6 | 8,31 | 2977 | 7,44 | 3 | 3 | 1,088 | 1,082 | 1,563 | 1,318 | 1,08 | 1,264 | 1,129 | 1,151 | 1,09 | 0,09 | 4,1E-01 |
| Q9D379 | Epoxide hydrolase 1 OS=Mus musculus OX=10090 GN=Ephx1 PE | 3 | 1 | 1 | 1 | 455 | 52,5 | 8,35 | 265 | | 1 | | 0,565 | 0,702 | 0,647 | 0,682 | 0,774 | 0,672 | 0,636 | 0,665 | 0,95 | -0,05 | 4,1E-01 |
| Q3TC72 | Fumarylacetoacetate hydrolase domain-containing protein 2A C | 12 | 3 | 3 | 3 | 313 | 34,7 | 8,16 | 1061 | 3,24 | 2 | 1 | 1,131 | 1,023 | 0,983 | 0,964 | 1,012 | 0,986 | 0,834 | 1,054 | 1,06 | 0,06 | 4,1E-01 |
| Q99NE5-7 | Isoform 7 of Regulating synaptic membrane exocytosis protein 1 | 9 | 8 | 11 | 2 | 1184 | 132,9 | 9,33 | 7519 | | 8 | | 1,145 | 0,874 | 1,303 | 1,056 | 0,691 | 0,975 | 1,163 | 1,069 | 1,12 | 0,12 | 4,1E-01 |
| Q8CGK3 | Lon protease homolog, mitochondrial OS=Mus musculus OX=10 | 6 | 5 | 8 | 5 | 949 | 105,8 | 6,57 | 4444 | | 5 | | 0,968 | 0,983 | 1,048 | 0,982 | 0,995 | 1,19 | 0,988 | 0,994 | 0,96 | -0,04 | 4,1E-01 |
| P42669 | Transcriptional activator protein Pur-alpha OS=Mus musculus O | 11 | 4 | 5 | 4 | 321 | 34,9 | 6,44 | 1181 | 4,85 | 2 | 2 | 0,89 | 0,973 | 1,023 | 0,997 | 0,724 | 1,017 | 0,873 | 1,006 | 1,07 | 0,07 | 4,1E-01 |
| P10639 | Thioredoxin OS=Mus musculus OX=10090 GN=Txn PE=1 SV=3 | 31 | 4 | 8 | 4 | 105 | 11,7 | 4,92 | 3488 | 7,58 | 3 | 1 | 0,98 | 1,072 | 0,968 | 1,213 | 0,873 | 1,021 | 1,093 | 0,989 | 1,06 | 0,06 | 4,1E-01 |
| Q80WQ2 | Protein VAC14 homolog OS=Mus musculus OX=10090 GN=Vac14 | 7 | 5 | 5 | 5 | 782 | 88 | 6,13 | 1806 | 4,8 | 3 | 2 | 1,111 | 1,101 | 1,141 | 1,089 | 1,427 | 0,98 | 1,165 | 1,196 | 0,93 | -0,07 | 4,1E-01 |
| P27773 | Protein disulfide-isomerase A3 OS=Mus musculus OX=10090 GN | 48 | 28 | 58 | 28 | 505 | 56,6 | 6,21 | 25855 | 44,82 | 19 | 11 | 1,012 | 0,948 | 1,038 | 1,097 | 0,987 | 0,98 | 1,011 | 1,005 | 1,03 | 0,03 | 4,1E-01 |
| P63085 | Mitogen-activated protein kinase 1 OS=Mus musculus OX=1009 | 33 | 14 | 24 | 10 | 358 | 41,2 | 6,98 | 12320 | 11,85 | 12 | 3 | 1,051 | 0,949 | 1,001 | 1,049 | 0,699 | 1,101 | 0,971 | 0,969 | 1,08 | 0,08 | 4,1E-01 |
| Q9DB75 | Cell death-inducing p53-target protein 1 OS=Mus musculus OX= | 4 | 1 | 2 | 1 | 208 | 21,8 | 5,66 | | 4,98 | | 1 | 1,09 | 1,008 | 0,912 | 0,839 | 1,04 | 0,985 | 1,001 | 1,021 | 0,95 | -0,05 | 4,1E-01 |
| Q80WT0 | Junctophilin-4 OS=Mus musculus OX=10090 GN=Jph4 PE=2 SV=1 | 1 | 1 | 1 | 1 | 628 | 66 | 6,71 | 370 | | 1 | | 1,009 | 1,088 | 1,196 | 0,994 | 1,093 | 1,368 | 1,285 | 0,926 | 0,92 | -0,08 | 4,1E-01 |
| Q9DBE0 | Cysteine sulfinic acid decarboxylase OS=Mus musculus OX=1009 | 3 | 1 | 1 | 1 | 493 | 55,1 | 6,61 | | 1,75 | | 1 | 1,163 | 1,472 | 1,178 | 1,161 | 1,361 | 1,11 | 1,379 | 1,531 | 0,92 | -0,08 | 4,1E-01 |
| Q9Z2Q6 | Septin-5 OS=Mus musculus OX=10090 GN=Septin5 PE=1 SV=2 | 43 | 19 | 31 | 18 | 369 | 42,7 | 6,67 | 12392 | 24,97 | 14 | 7 | 1,002 | 0,961 | 0,953 | 0,959 | 0,95 | 0,999 | 0,891 | 0,948 | 1,02 | 0,02 | 4,1E-01 |
| P40336 | Vacuolar protein sorting-associated protein 26A OS=Mus muscu | 13 | 4 | 5 | 4 | 327 | 38,1 | 6,57 | 2946 | | 4 | | 0,913 | 0,97 | 0,952 | 0,956 | 1,74 | 0,805 | 0,931 | 1,05 | 0,84 | -0,16 | 4,1E-01 |
| A2AKG8 | Focadhesin OS=Mus musculus OX=10090 GN=Focad PE=1 SV=1 | 0 | 1 | 1 | 1 | 1798 | 198,8 | 6,74 | 406 | | 1 | | 1,111 | 0,849 | 1,118 | 0,974 | 0,59 | 1,082 | 0,987 | 0,953 | 1,12 | 0,12 | 4,1E-01 |
| Q69Z98 | Serine/threonine-protein kinase BRSK2 OS=Mus musculus OX=1 | 11 | 8 | 10 | 6 | 735 | 81,7 | 8,79 | 3015 | 6,3 | 4 | 4 | 0,926 | 0,965 | 1,05 | 0,929 | 0,857 | 0,948 | 0,888 | 1,017 | 1,04 | 0,04 | 4,1E-01 |
| Q6ZQ82 | Rho GTPase-activating protein 26 OS=Mus musculus OX=10090 | 3 | 2 | 3 | 2 | 814 | 92 | 6,68 | 446 | 1,66 | 1 | 1 | 0,991 | 0,972 | 0,75 | 0,98 | 0,93 | 1,156 | 1,097 | 0,838 | 0,92 | -0,08 | 4,1E-01 |
| Q8R366 | Immunoglobulin superfamily member 8 OS=Mus musculus OX=1 | 10 | 5 | 6 | 5 | 611 | 65 | 7,99 | 3489 | 3,19 | 4 | 1 | 0,95 | 0,923 | 0,999 | 0,941 | 1,492 | 0,91 | 0,897 | 1,01 | 0,88 | -0,12 | 4,1E-01 |
| Q8VCH0 | 3-ketoacyl-CoA thiolase B, peroxisomal OS=Mus musculus OX=1 | 19 | 6 | 15 | 1 | 424 | 44 | 8,51 | 4108 | 29,4 | 5 | 2 | 0,89 | 1,154 | 0,95 | 0,991 | 2,28 | 0,636 | 0,857 | 1,525 | 0,75 | -0,25 | 4,1E-01 |
| P02468 | Laminin subunit gamma-1 OS=Mus musculus OX=10090 GN=Lan | 1 | 1 | 1 | 1 | 1607 | 177,2 | 5,21 | 739 | | 1 | | 0,892 | 0,888 | 0,686 | 0,995 | 1,203 | 0,553 | 1,207 | 1,089 | 0,85 | -0,15 | 4,1E-01 |
| Q61490 | CD166 antigen OS=Mus musculus OX=10090 GN=Alcam PE=1 SV | 11 | 6 | 8 | 6 | 583 | 65,1 | 6,15 | 975 | 14,94 | 2 | 5 | 0,976 | 0,897 | 1,076 | 0,926 | 0,991 | 0,905 | 0,901 | 0,922 | 1,04 | 0,04 | 4,1E-01 |
| O35857 | Mitochondrial import inner membrane translocase subunit TIM4 | 7 | 3 | 4 | 3 | 452 | 51,1 | 8,13 | 974 | 5,55 | 1 | 2 | 0,906 | 0,881 | 0,834 | 0,961 | 0,926 | 1,008 | 0,864 | 0,923 | 0,96 | -0,04 | 4,1E-01 |
| P56395 | Cytochrome b5 OS=Mus musculus OX=10090 GN=Cyb5a PE=1 SV | 30 | 4 | 8 | 4 | 134 | 15,2 | 5,07 | 5603 | | 4 | | 0,988 | 1,082 | 1,112 | 1,047 | 1,262 | 0,959 | 1,121 | 1,123 | 0,95 | -0,05 | 4,2E-01 |
| Q9R0Q3 | Transmembrane emp24 domain-containing protein 2 OS=Mus m | 10 | 2 | 2 | 2 | 201 | 22,7 | 5,17 | 1094 | | 2 | | 0,992 | 1,024 | 1,217 | 1,019 | 1,035 | 1,125 | 0,768 | 1,002 | 1,08 | 0,08 | 4,2E-01 |
| P09103 | Protein disulfide-isomerase OS=Mus musculus OX=10090 GN=P4 | 32 | 18 | 35 | 18 | 509 | 57 | 4,88 | 13709 | 33,53 | 13 | 5 | 0,958 | 0,968 | 0,965 | 1,064 | 1,161 | 0,97 | 0,97 | 1,034 | 0,96 | -0,04 | 4,2E-01 |
| P59913 | Protein-L-isoaspartate O-methyltransferase domain-containing | 2 | 1 | 1 | 1 | 357 | 40,7 | 5,66 | 350 | | 1 | | 1,025 | 0,962 | 0,959 | 0,799 | 1,48 | 0,912 | 1,211 | 0,738 | 0,86 | -0,14 | 4,2E-01 |
| Q9Z1P6 | NADH dehydrogenase [ubiquinone] 1 alpha subcomplex subunit | 55 | 9 | 19 | 9 | 113 | 12,6 | 10,17 | 8560 | 9,52 | 7 | 3 | 1,034 | 0,951 | 0,969 | 1,006 | 1,137 | 1,024 | 0,96 | 0,989 | 0,96 | -0,04 | 4,2E-01 |
| Q8BJI1 | Sodium-dependent neutral amino acid transporter SLC6A17 OS= | 6 | 3 | 7 | 3 | 727 | 81 | 6,23 | 1479 | 7,21 | 1 | 3 | 1,201 | 0,917 | 0,964 | 1,092 | 0,892 | 1,007 | 0,984 | 1,041 | 1,06 | 0,06 | 4,2E-01 |
| P35123 | Ubiquitin carboxyl-terminal hydrolase 4 OS=Mus musculus OX=1 | 2 | 2 | 2 | 1 | 962 | 108,3 | 5,64 | | 5,34 | | 2 | 0,966 | 0,685 | 1,038 | 0,884 | 1,054 | 0,747 | 1,022 | 1,153 | 0,90 | -0,10 | 4,2E-01 |
| Q91YS8 | Calcium/calmodulin-dependent protein kinase type 1 OS=Mus m | 12 | 4 | 5 | 3 | 374 | 41,6 | 5,35 | 1705 | 5,52 | 3 | 2 | 0,94 | 1,075 | 0,995 | 1,202 | 1,175 | 1,1 | 1,263 | 0,968 | 0,93 | -0,07 | 4,2E-01 |
| Q69ZX6 | ATPase MORC2A OS=Mus musculus OX=10090 GN=Morc2a PE=1 | 1 | 1 | 2 | 1 | 1030 | 117,3 | 8,34 | 386 | | 1 | | 1,042 | 1,19 | 0,964 | 1,267 | 1,436 | 0,803 | 1,268 | 1,6 | 0,87 | -0,13 | 4,2E-01 |
| Q8BX10 | Serine/threonine-protein phosphatase PGAM5, mitochondrial O | 26 | 7 | 9 | 7 | 288 | 32 | 9,04 | 4198 | 5,67 | 6 | 1 | 1,019 | 1,021 | 0,999 | 1,029 | 0,921 | 1,05 | 1,008 | 0,993 | 1,02 | 0,02 | 4,2E-01 |
| Q9D3A9 | Protein tweety homolog 1 OS=Mus musculus OX=10090 GN=Tty | 3 | 1 | 3 | 1 | 450 | 49 | 5,03 | 1722 | | 1 | | 1,168 | 1,173 | 0,993 | 0,77 | 2,928 | 0,732 | 0,881 | 1,342 | 0,70 | -0,30 | 4,2E-01 |
| Q8CGY8 | UDP-N-acetylglucosamine--peptide N-acetylglucosaminyltransfe | 11 | 10 | 11 | 10 | 1046 | 116,9 | 6,7 | 3621 | 14,78 | 6 | 5 | 1,021 | 1,078 | 1,061 | 1,054 | 0,993 | 1,075 | 1,091 | 1,234 | 0,96 | -0,04 | 4,2E-01 |
| Q3UHJ0 | AP2-associated protein kinase 1 OS=Mus musculus OX=10090 G | 16 | 10 | 19 | 10 | 959 | 103,3 | 6,7 | 8400 | 10,67 | 9 | 3 | 0,904 | 1,091 | 1,04 | 0,992 | 1,456 | 0,937 | 1,07 | 0,994 | 0,90 | -0,10 | 4,2E-01 |
| Q9EPU0 | Regulator of nonsense transcripts 1 OS=Mus musculus OX=1009 | 9 | 7 | 10 | 7 | 1124 | 123,9 | 6,61 | 4245 | 4,42 | 6 | 1 | 1,026 | 1,017 | 1,057 | 1,102 | 1,03 | 1,066 | 0,996 | 1,027 | 1,02 | 0,02 | 4,2E-01 |
| Q8CHX7 | Raftlin-2 OS=Mus musculus OX=10090 GN=Rftn2 PE=1 SV=3 | 5 | 2 | 3 | 2 | 500 | 54,9 | 5,5 | 1699 | | 2 | | 0,927 | 0,924 | 0,822 | 0,756 | 1,191 | 0,684 | 0,991 | 0,952 | 0,90 | -0,10 | 4,2E-01 |
| Q9ERG2 | Striatin-3 OS=Mus musculus OX=10090 GN=Strn3 PE=1 SV=1 | 13 | 7 | 11 | 6 | 796 | 87,1 | 5,29 | 5643 | 3,9 | 6 | 1 | 0,986 | 0,936 | 0,854 | 0,964 | 0,965 | 0,971 | 0,962 | 0,944 | 0,97 | -0,03 | 4,2E-01 |
| Q99M87 | DnaJ homolog subfamily A member 3, mitochondrial OS=Mus m | 7 | 3 | 3 | 3 | 480 | 52,4 | 9,22 | 525 | 4,42 | 1 | 2 | 0,989 | 1,017 | 1,004 | 1,149 | 0,956 | 0,929 | 1,138 | 0,916 | 1,06 | 0,06 | 4,2E-01 |
| Q811J3 | Iron-responsive element-binding protein 2 OS=Mus musculus O | 1 | 1 | 1 | 1 | 963 | 104,9 | 6,93 | 325 | | 1 | | 0,951 | 1 | 1,012 | 0,886 | 1,333 | 1,043 | 0,772 | 1,113 | 0,90 | -0,10 | 4,2E-01 |
| Q8C3Y4 | Kinetochore-associated protein 1 OS=Mus musculus OX=10090 | 0 | 1 | 1 | 1 | 2207 | 250,2 | 6 | 309 | | 1 | | 1,17 | 1,303 | 0,87 | 1,191 | 1,335 | 1,024 | 1,289 | 1,289 | 0,92 | -0,08 | 4,2E-01 |
| P63073 | Eukaryotic translation initiation factor 4E OS=Mus musculus OX= | 6 | 1 | 2 | 1 | 217 | 25 | 6,15 | 1141 | | 1 | | 1,127 | 1,1 | 1,111 | 1,181 | 1,575 | 1,245 | 1,034 | 1,089 | 0,91 | -0,09 | 4,2E-01 |
| Q9Z0H8-2 | Isoform 2 of CAP-Gly domain-containing linker protein 2 OS=Mu | 39 | 33 | 45 | 1 | 1012 | 111,8 | 6,55 | 19724 | 30,84 | 23 | 10 | 0,893 | 1,415 | 1,145 | 1,132 | 0,444 | 1,167 | 0,995 | 1,253 | 1,19 | 0,19 | 4,2E-01 |
| Q8BXA7 | PH domain leucine-rich repeat-containing protein phosphatase 1 | 1 | 1 | 1 | 1 | 1320 | 145,9 | 5,95 | 320 | | 1 | | 1,182 | 0,909 | 1,133 | 1,34 | 1,092 | 0,976 | 0,787 | 1,253 | 1,11 | 0,11 | 4,2E-01 |
| Q91VW3 | SH3 domain-binding glutamic acid-rich-like protein 3 OS=Mus m | 52 | 5 | 8 | 5 | 93 | 10,5 | 5,14 | 959 | 12,43 | 2 | 4 | 0,901 | 1,073 | 0,95 | 0,927 | 1,025 | 1,154 | 0,875 | 1,033 | 0,94 | -0,06 | 4,2E-01 |
| P10649 | Glutathione S-transferase Mu 1 OS=Mus musculus OX=10090 GI | 39 | 10 | 17 | 9 | 218 | 26 | 7,94 | 7298 | 10,9 | 6 | 4 | 1,093 | 1,377 | 1,288 | 1,302 | 1,647 | 1,2 | 1,338 | 1,272 | 0,93 | -0,07 | 4,2E-01 |
| Q91X91 | Nicotinate-nucleotide pyrophosphorylase [carboxylating] OS=M | 4 | 1 | 1 | 1 | 299 | 31,5 | 6,71 | 616 | | 1 | | 0,895 | 0,907 | 1,42 | 1,14 | 1,465 | 1,289 | 0,894 | 1,309 | 0,88 | -0,12 | 4,2E-01 |
| Q9CRB9 | MICOS complex subunit Mic19 OS=Mus musculus OX=10090 GN | 11 | 3 | 4 | 3 | 227 | 26,3 | 8,37 | 3326 | | 3 | | 1,008 | 0,979 | 1,128 | 0,967 | 1,043 | 1,106 | 1,02 | 1,054 | 0,97 | -0,03 | 4,2E-01 |
| Q8K400 | Syntaxin-binding protein 5 OS=Mus musculus OX=10090 GN=Stx | 2 | 2 | 2 | 2 | 1152 | 127,6 | 7,21 | 987 | | 2 | | 1,095 | 1,413 | 1,289 | 1,253 | 2,018 | 1,146 | 1,2 | 1,407 | 0,88 | -0,12 | 4,2E-01 |
| Q6P9Q6 | FK506-binding protein 15 OS=Mus musculus OX=10090 GN=Fkb | 1 | 1 | 1 | 1 | 1216 | 132,9 | 5,07 | 210 | | 1 | | 1,294 | 0,81 | 0,666 | 1,069 | 1,856 | 0,84 | 0,822 | 1,28 | 0,80 | -0,20 | 4,2E-01 |
| Q64096 | Guanine nucleotide exchange factor DBS OS=Mus musculus OX= | 2 | 2 | 2 | 2 | 1149 | 129 | 5,9 | 720 | | 2 | | 1,111 | 1,083 | 1,15 | 1,117 | 0,764 | 1,169 | 1,175 | 1,021 | 1,08 | 0,08 | 4,3E-01 |
| Q9D0W5 | Peptidyl-prolyl cis-trans isomerase-like 1 OS=Mus musculus OX= | 11 | 2 | 3 | 2 | 166 | 18,2 | 7,99 | 1102 | 2,16 | 1 | 1 | 0,9 | 1,06 | 1,077 | 1,022 | 1,098 | 1,1 | 1,109 | 0,944 | 0,95 | -0,05 | 4,3E-01 |
| Q8BP48 | Methionine aminopeptidase 1 OS=Mus musculus OX=10090 GN | 11 | 3 | 4 | 3 | 386 | 43,2 | 7,17 | 980 | 2,73 | 2 | 1 | 1,11 | 1,105 | 1,328 | 1,219 | 0,843 | 0,953 | 1,164 | 1,367 | 1,10 | 0,10 | 4,3E-01 |
| Q8BHC4 | Dephospho-CoA kinase domain-containing protein OS=Mus mus | 10 | 2 | 4 | 2 | 231 | 26,5 | 9,58 | 525 | 2,17 | 1 | 1 | 1,16 | 1,276 | 1,232 | 1,289 | 1,555 | 1,312 | 1,25 | 1,15 | 0,94 | -0,06 | 4,3E-01 |
| Q9D6Z1 | Nucleolar protein 56 OS=Mus musculus OX=10090 GN=Nop56 P | 5 | 3 | 4 | 3 | 580 | 64,4 | 9,14 | 1826 | 2,53 | 2 | 1 | 0,853 | 1,024 | 1,014 | 1,071 | 1,159 | 0,873 | 1,063 | 1,143 | 0,93 | -0,07 | 4,3E-01 |
| O54734 | Dolichyl-diphosphooligosaccharide--protein glycosyltransferase | 13 | 5 | 8 | 5 | 441 | 49 | 5,83 | 3868 | 5,77 | 4 | 1 | 0,979 | 1,052 | 0,907 | 1,015 | 1,251 | 0,988 | 1,047 | 0,928 | 0,94 | -0,06 | 4,3E-01 |
| P45952 | Medium-chain specific acyl-CoA dehydrogenase, mitochondrial | 21 | 7 | 10 | 7 | 421 | 46,5 | 8,37 | 6821 | | 7 | | 1,042 | 1,05 | 1,174 | 1,074 | 1,452 | 1,068 | 1,009 | 1,161 | 0,93 | -0,07 | 4,3E-01 |
| Q9Z268 | RasGAP-activating-like protein 1 OS=Mus musculus OX=10090 G | 5 | 4 | 4 | 4 | 799 | 89,3 | 6,37 | 996 | 4,95 | 2 | 2 | 1,02 | 0,916 | 0,906 | 0,941 | 0,956 | 0,977 | 0,847 | 0,861 | 1,04 | 0,04 | 4,3E-01 |
| P47915 | 60S ribosomal protein L29 OS=Mus musculus OX=10090 GN=Rpl | 24 | 4 | 5 | 4 | 160 | 17,6 | 11,84 | 1594 | 1,75 | 2 | 2 | 0,976 | 0,68 | 1,394 | 0,859 | 0,742 | 1,004 | 0,808 | 0,805 | 1,16 | 0,16 | 4,3E-01 |
| Q9D6K5 | Synaptojanin-2-binding protein OS=Mus musculus OX=10090 GN | 18 | 2 | 2 | 2 | 145 | 15,8 | 6,3 | 1027 | | 2 | | 1,004 | 1,028 | 0,972 | 0,874 | 1,038 | 1,041 | 0,921 | 1,029 | 0,96 | -0,04 | 4,3E-01 |
| Q8BHA3 | D-aminoacyl-tRNA deacylase 2 OS=Mus musculus OX=10090 GN | 10 | 1 | 1 | 1 | 168 | 18,2 | 7,91 | 559 | | 1 | | 1,398 | 1,66 | 1,862 | 1,773 | 2,364 | 1,457 | 1,406 | 2,519 | 0,86 | -0,14 | 4,3E-01 |
| Q810U3 | Neurofascin OS=Mus musculus OX=10090 GN=Nfasc PE=1 SV=1 | 5 | 4 | 5 | 4 | 1240 | 137,9 | 6,19 | 2737 | | 4 | | 1,004 | 0,973 | 1,061 | 1,199 | 1,454 | 0,909 | 1,087 | 1,211 | 0,91 | -0,09 | 4,3E-01 |

| | | | | | | | | | | | | | | | | | | | | | | | |
|---|---|---|---|---|---|---|---|---|---|---|---|---|---|---|---|---|---|---|---|---|---|---|---|
| Q91VI7 | Ribonuclease inhibitor OS=Mus musculus OX=10090 GN=Rnh1 P | 11 | 3 | 5 | 3 | 456 | 49,8 | 4,78 | 2811 | | 3 | | 0,986 | 0,942 | 1,057 | 0,914 | 1,101 | 0,879 | 1,008 | 1,13 | 0,95 | -0,05 | 4,3E-01 |
| Q8K019 | Bcl-2-associated transcription factor 1 OS=Mus musculus OX=10 | 1 | 1 | 1 | 1 | 919 | 105,9 | 9,99 | 237 | | 1 | | 0,764 | 0,98 | 0,771 | 0,767 | 0,889 | 0,903 | 0,844 | 0,835 | 0,95 | -0,05 | 4,3E-01 |
| Q3TIV5 | Zinc finger CCCH domain-containing protein 15 OS=Mus muscul | 2 | 1 | 1 | 1 | 426 | 48,3 | 5,3 | 724 | | 1 | | 0,901 | 0,912 | 1,115 | 1,019 | 0,778 | 1,085 | 0,937 | 0,869 | 1,08 | 0,08 | 4,3E-01 |
| P97798 | Neogenin OS=Mus musculus OX=10090 GN=Neo1 PE=1 SV=1 | 8 | 8 | 15 | 8 | 1493 | 163,1 | 6,47 | 7367 | 2,01 | 7 | 2 | 0,949 | 1,113 | 0,846 | 0,973 | 1,054 | 0,933 | 1,037 | 1,071 | 0,95 | -0,05 | 4,3E-01 |
| Q61599 | Rho GDP-dissociation inhibitor 2 OS=Mus musculus OX=10090 G | 19 | 3 | 4 | 3 | 200 | 22,8 | 5,11 | 1594 | 2,47 | 2 | 1 | 0,986 | 1,281 | 0,919 | 1,477 | 0,817 | 1,228 | 1,084 | 1,006 | 1,13 | 0,13 | 4,3E-01 |
| Q8R001 | Microtubule-associated protein RP/EB family member 2 OS=Mu | 22 | 10 | 16 | 8 | 326 | 36,9 | 5,38 | 7422 | 14,06 | 7 | 3 | 1,045 | 0,997 | 1,032 | 1,096 | 0,959 | 1,007 | 1,083 | 1,01 | 1,03 | 0,03 | 4,3E-01 |
| Q60899 | ELAV-like protein 2 OS=Mus musculus OX=10090 GN=Elavl2 PE= | 9 | 3 | 3 | 1 | 360 | 39,6 | 9,13 | 718 | 5,58 | 1 | 2 | 1,027 | 0,89 | 1,106 | 1,009 | 1,217 | 0,954 | 0,941 | 1,228 | 0,93 | -0,07 | 4,3E-01 |
| C0HK80 | Adipocyte-related X-chromosome expressed sequence 2 OS=Mu | 11 | 2 | 2 | 2 | 180 | 20,1 | 9,6 | 366 | 2,67 | 1 | 1 | 1,07 | 0,874 | 1,129 | 0,915 | 0,843 | 0,94 | 1,027 | 0,936 | 1,06 | 0,06 | 4,3E-01 |
| Q7TQF7 | Amphiphysin OS=Mus musculus OX=10090 GN=Amph PE=1 SV=1 | 27 | 18 | 39 | 16 | 686 | 75 | 4,63 | 17142 | 32,84 | 12 | 8 | 1,057 | 1,01 | 1,003 | 0,963 | 0,856 | 1,059 | 0,967 | 0,994 | 1,04 | 0,04 | 4,3E-01 |
| Q8QZS1 | 3-hydroxyisobutyryl-CoA hydrolase, mitochondrial OS=Mus mus | 12 | 4 | 4 | 4 | 385 | 43 | 8,06 | 1492 | 2,77 | 2 | 2 | 1,146 | 1,067 | 1,185 | 1,115 | 1,265 | 1,15 | 1,133 | 1,109 | 0,97 | -0,03 | 4,3E-01 |
| P19536 | Cytochrome c oxidase subunit 5B, mitochondrial OS=Mus muscu | 43 | 8 | 44 | 8 | 128 | 13,8 | 8,38 | 23338 | 8,46 | 6 | 2 | 1,052 | 1,022 | 0,943 | 1,026 | 0,938 | 1,038 | 0,948 | 1,006 | 1,03 | 0,03 | 4,3E-01 |
| Q8R317 | Ubiquilin-1 OS=Mus musculus OX=10090 GN=Ubqln1 PE=1 SV=1 | 9 | 4 | 7 | 2 | 582 | 61,9 | 4,94 | 2571 | 5,17 | 4 | 1 | 0,796 | 0,835 | 0,758 | 0,999 | 1,354 | 0,899 | 0,816 | 0,796 | 0,88 | -0,12 | 4,3E-01 |
| Q5F2E8 | Serine/threonine-protein kinase TAO1 OS=Mus musculus OX=10 | 5 | 5 | 5 | 4 | 1001 | 116 | 7,55 | 1551 | 0 | 4 | 1 | 0,872 | 1,454 | 1,213 | 0,974 | 2,571 | 0,924 | 0,942 | 1,449 | 0,77 | -0,23 | 4,3E-01 |
| P62141 | Serine/threonine-protein phosphatase PP1-beta catalytic subun | 34 | 9 | 15 | 3 | 327 | 37,2 | 6,19 | 5027 | 15,34 | 7 | 3 | 0,84 | 0,854 | 1,184 | 0,922 | 2,563 | 0,896 | 0,923 | 0,853 | 0,73 | -0,27 | 4,3E-01 |
| Q61037-6 | Isoform E of Tuberin OS=Mus musculus OX=10090 GN=Tsc2 | 3 | 3 | 5 | 3 | 1786 | 198,8 | 7,01 | 2134 | | 3 | | 1,036 | 1,074 | 0,939 | 0,914 | 1,149 | 0,891 | 1,064 | 1,085 | 0,95 | -0,05 | 4,3E-01 |
| Q8CGZ0 | Calcium homeostasis endoplasmic reticulum protein OS=Mus m | 1 | 1 | 1 | 1 | 936 | 106,1 | 9,14 | 717 | | 1 | | 1,02 | 1,13 | 1,185 | 1,149 | 3,081 | 0,953 | 0,805 | 1,408 | 0,72 | -0,28 | 4,3E-01 |
| A3KMP2 | Tetratricopeptide repeat protein 38 OS=Mus musculus OX=1009 | 3 | 1 | 1 | 1 | 465 | 52,2 | 6,3 | 428 | | 1 | | 0,624 | 0,842 | 0,96 | 0,639 | 1,602 | 0,747 | 0,841 | 0,655 | 0,80 | -0,20 | 4,3E-01 |
| Q78IK4 | MICOS complex subunit Mic27 OS=Mus musculus OX=10090 GN | 17 | 4 | 6 | 4 | 265 | 29,2 | 9,31 | 2336 | 2,23 | 3 | 1 | 0,827 | 0,805 | 0,767 | 0,848 | 0,933 | 0,741 | 0,82 | 0,913 | 0,95 | -0,05 | 4,3E-01 |
| O54916 | RalBP1-associated Eps domain-containing protein 1 OS=Mus mu | 3 | 3 | 4 | 1 | 795 | 86,5 | 5,58 | 2728 | | 3 | | 0,915 | 0,741 | 1,366 | 0,98 | 1,214 | 0,878 | 0,713 | 0,544 | 1,19 | 0,19 | 4,3E-01 |
| Q499X9 | Methionine--tRNA ligase, mitochondrial OS=Mus musculus OX=1 | 2 | 1 | 1 | 1 | 586 | 65,8 | 8,09 | 422 | | 1 | | 0,798 | 1,133 | 1,217 | 0,879 | 1,219 | 1,159 | 1,032 | 0,996 | 0,91 | -0,09 | 4,3E-01 |
| Q9CQ75 | NADH dehydrogenase [ubiquinone] 1 alpha subcomplex subunit | 56 | 6 | 10 | 6 | 99 | 10,9 | 9,99 | 5162 | 5,09 | 4 | 3 | 0,906 | 0,855 | 0,775 | 0,902 | 0,765 | 0,936 | 0,778 | 0,791 | 1,05 | 0,05 | 4,3E-01 |
| Q3ULB5 | Serine/threonine-protein kinase PAK 6 OS=Mus musculus OX=10 | 2 | 1 | 2 | 1 | 682 | 74,8 | 9,44 | 1162 | | 1 | | 0,999 | 1,141 | 0,882 | 1,415 | 0,602 | 1,094 | 1,083 | 1,097 | 1,14 | 0,14 | 4,4E-01 |
| P17183 | Gamma-enolase OS=Mus musculus OX=10090 GN=Eno2 PE=1 SV | 26 | 9 | 17 | 6 | 434 | 47,3 | 5,11 | 7171 | 20,7 | 7 | 3 | 0,94 | 0,869 | 0,991 | 0,952 | 0,988 | 1,018 | 0,895 | 0,973 | 0,97 | -0,03 | 4,4E-01 |
| P14152 | Malate dehydrogenase, cytoplasmic OS=Mus musculus OX=1009 | 38 | 13 | 30 | 13 | 334 | 36,5 | 6,58 | 11730 | 35,84 | 7 | 7 | 1,042 | 1,004 | 0,919 | 1,039 | 0,711 | 1,142 | 0,916 | 0,926 | 1,08 | 0,08 | 4,4E-01 |
| Q9R0H0 | Peroxisomal acyl-coenzyme A oxidase 1 OS=Mus musculus OX=1 | 24 | 12 | 15 | 12 | 661 | 74,6 | 8,48 | 7524 | 10,49 | 9 | 4 | 1,051 | 1,105 | 1,153 | 1,107 | 1,197 | 1,061 | 1,117 | 1,161 | 0,97 | -0,03 | 4,4E-01 |
| Q99KH8 | Serine/threonine-protein kinase 24 OS=Mus musculus OX=1009 | 5 | 2 | 2 | 2 | 431 | 47,9 | 5,43 | 641 | 2,57 | 1 | 1 | 0,921 | 0,94 | 0,869 | 0,897 | 0,556 | 0,958 | 0,895 | 0,907 | 1,09 | 0,09 | 4,4E-01 |
| Q9D0G0 | 28S ribosomal protein S30, mitochondrial OS=Mus musculus OX | 8 | 3 | 3 | 3 | 442 | 49,9 | 9,38 | 1720 | | 3 | | 1,045 | 0,88 | 0,973 | 0,945 | 0,951 | 0,916 | 1,005 | 1,238 | 0,94 | -0,06 | 4,4E-01 |
| Q64518 | Sarcoplasmic/endoplasmic reticulum calcium ATPase 3 OS=Mus | 7 | 6 | 8 | 2 | 1038 | 113,6 | 5,85 | 2721 | 7,42 | 3 | 3 | 0,847 | 0,782 | 0,862 | 1,032 | 1,645 | 0,811 | 0,992 | 0,772 | 0,83 | -0,17 | 4,4E-01 |
| O55042 | Alpha-synuclein OS=Mus musculus OX=10090 GN=Snca PE=1 SV | 61 | 11 | 38 | 8 | 140 | 14,5 | 4,77 | 22515 | 21,21 | 7 | 4 | 1,035 | 1,025 | 0,85 | 1,057 | 0,62 | 1,213 | 0,847 | 0,849 | 1,12 | 0,12 | 4,4E-01 |
| Q9JIY5 | Serine protease HTRA2, mitochondrial OS=Mus musculus OX=10 | 5 | 2 | 2 | 2 | 458 | 49,3 | 9,6 | 926 | | 2 | | 0,869 | 1,025 | 0,875 | 0,959 | 0,993 | 0,859 | 1,015 | 1,045 | 0,95 | -0,05 | 4,4E-01 |
| Q8QZY1 | Eukaryotic translation initiation factor 3 subunit L OS=Mus musc | 8 | 4 | 5 | 4 | 564 | 66,6 | 6,44 | 1167 | 7,72 | 2 | 3 | 0,731 | 0,896 | 0,844 | 0,887 | 1,024 | 0,77 | 0,928 | 0,855 | 0,94 | -0,06 | 4,4E-01 |
| Q9QYB5 | Gamma-adducin OS=Mus musculus OX=10090 GN=Add3 PE=1 S | 14 | 8 | 13 | 8 | 706 | 78,7 | 5,95 | 5455 | 0 | 7 | 1 | 0,958 | 1,052 | 0,87 | 1,009 | 0,988 | 0,893 | 0,961 | 0,895 | 1,04 | 0,04 | 4,4E-01 |
| Q6P8M1 | Putative deoxyribonuclease TATDN1 OS=Mus musculus OX=100 | 3 | 1 | 1 | 1 | 295 | 33,3 | 6,18 | 248 | | 1 | | 1,138 | 1,079 | 1,12 | 1,22 | 1,416 | 1,347 | 0,937 | 1,221 | 0,93 | -0,07 | 4,4E-01 |
| Q3ULD5 | Methylcrotonoyl-CoA carboxylase beta chain, mitochondrial OS= | 6 | 3 | 4 | 3 | 563 | 61,3 | 8 | 1689 | 3,49 | 2 | 1 | 1,11 | 0,898 | 1,057 | 0,905 | 2,096 | 0,956 | 0,789 | 1,116 | 0,80 | -0,20 | 4,4E-01 |
| P62900 | 60S ribosomal protein L31 OS=Mus musculus OX=10090 GN=Rpl | 22 | 4 | 6 | 4 | 125 | 14,5 | 10,54 | 2591 | 0 | 3 | 1 | 1,017 | 0,905 | 1,057 | 0,932 | 0,527 | 1,251 | 0,881 | 0,733 | 1,15 | 0,15 | 4,4E-01 |
| Q8K003 | Translation machinery-associated protein 7 OS=Mus musculus C | 14 | 1 | 1 | 1 | 64 | 7,1 | 9,99 | | 2,47 | | 1 | 1,381 | 1,235 | 1,345 | 1,164 | 0,938 | 1,416 | 1,095 | 1,29 | 1,08 | 0,08 | 4,4E-01 |
| Q9CY34 | NEDD8-conjugating enzyme UBE2F OS=Mus musculus OX=10090 | 4 | 1 | 1 | 1 | 185 | 21,1 | 6,54 | 531 | | 1 | | 0,956 | 0,946 | 1,096 | 1,076 | 0,799 | 1,107 | 0,976 | 0,946 | 1,06 | 0,06 | 4,4E-01 |
| Q9CYR0 | Single-stranded DNA-binding protein, mitochondrial OS=Mus m | 5 | 1 | 1 | 1 | 152 | 17,3 | 9,92 | 775 | | 1 | | 0,832 | 0,965 | 1,434 | 1,166 | 0,68 | 1,056 | 1,065 | 1,063 | 1,14 | 0,14 | 4,4E-01 |
| Q9Z2Y3 | Homer protein homolog 1 OS=Mus musculus OX=10090 GN=Ho | 25 | 8 | 14 | 8 | 366 | 41,4 | 5,53 | 5337 | 11,09 | 6 | 2 | 1,133 | 0,96 | 1,012 | 1,028 | 1,161 | 1,063 | 1,004 | 1,065 | 0,96 | -0,04 | 4,4E-01 |
| O08553 | Dihydropyrimidinase-related protein 2 OS=Mus musculus OX=10 | 68 | 33 | 321 | 29 | 572 | 62,2 | 6,38 | 121957 | 362,75 | 24 | 23 | 0,97 | 0,909 | 1,03 | 0,969 | 0,908 | 0,995 | 0,941 | 0,933 | 1,03 | 0,03 | 4,4E-01 |
| Q66L44 | Voltage-dependent calcium channel beta subunit-associated reg | 3 | 2 | 2 | 2 | 698 | 74,1 | 6,33 | 802 | | 2 | | 0,861 | 0,766 | 1,756 | 1,024 | 1,042 | 0,705 | 0,949 | 0,937 | 1,21 | 0,21 | 4,4E-01 |
| P63101 | 14-3-3 protein zeta/delta OS=Mus musculus OX=10090 GN=Ywh | 62 | 22 | 108 | 15 | 245 | 27,8 | 4,79 | 39088 | 113,44 | 15 | 14 | 0,945 | 0,914 | 1,001 | 0,995 | 0,752 | 1,095 | 0,9 | 0,864 | 1,07 | 0,07 | 4,4E-01 |
| Q62083 | PRKCA-binding protein OS=Mus musculus OX=10090 GN=Pick1 F | 5 | 2 | 2 | 2 | 416 | 46,6 | 5,52 | 623 | 2,52 | 1 | 1 | 1,076 | 1,191 | 1,364 | 1,35 | 0,913 | 1,269 | 0,904 | 1,416 | 1,11 | 0,11 | 4,4E-01 |
| Q810U4 | Neuronal cell adhesion molecule OS=Mus musculus OX=10090 G | 12 | 12 | 23 | 7 | 1256 | 138,4 | 5,91 | 6259 | 26,87 | 7 | 6 | 1,084 | 1,077 | 1,125 | 1,136 | 1,37 | 1,127 | 1,011 | 1,164 | 0,95 | -0,05 | 4,4E-01 |
| Q8BW96 | Calcium/calmodulin-dependent protein kinase type 1D OS=Mus | 14 | 6 | 9 | 5 | 385 | 42,9 | 7,17 | 4112 | 5,55 | 4 | 2 | 0,854 | 0,864 | 0,944 | 0,922 | 0,863 | 0,846 | 0,866 | 0,92 | 1,03 | 0,03 | 4,4E-01 |
| P99024 | Tubulin beta-5 chain OS=Mus musculus OX=10090 GN=Tubb5 PI | 65 | 27 | 457 | 4 | 444 | 49,6 | 4,89 | 191622 | 362,15 | 23 | 19 | 1,09 | 0,982 | 1,031 | 1,017 | 0,839 | 1,091 | 0,994 | 1,009 | 1,05 | 0,05 | 4,5E-01 |
| Q91X97 | Neurocalcin-delta OS=Mus musculus OX=10090 GN=Ncald PE=1 | 33 | 7 | 13 | 4 | 193 | 22,2 | 5,35 | 4795 | 7,44 | 5 | 3 | 0,898 | 0,956 | 1,138 | 0,924 | 1,243 | 0,902 | 1,014 | 1,049 | 0,93 | -0,07 | 4,5E-01 |
| Q9Z2W9 | Glutamate receptor 3 OS=Mus musculus OX=10090 GN=Gria3 PI | 4 | 4 | 5 | 1 | 888 | 100,5 | 8,38 | 2559 | 1,83 | 3 | 1 | 1,103 | 0,931 | 1,085 | 1,1 | 0,912 | 0,872 | 1,199 | 0,961 | 1,07 | 0,07 | 4,5E-01 |
| P47708 | Rabphilin-3A OS=Mus musculus OX=10090 GN=Rph3a PE=1 SV= | 19 | 9 | 13 | 9 | 681 | 75,4 | 8,27 | 5954 | 3,6 | 8 | 1 | 1,062 | 1,19 | 1,301 | 1,044 | 1,2 | 1,18 | 1,279 | 1,153 | 0,96 | -0,04 | 4,5E-01 |
| P70349 | Histidine triad nucleotide-binding protein 1 OS=Mus musculus C | 40 | 6 | 11 | 6 | 126 | 13,8 | 6,87 | 4326 | 9,33 | 4 | 2 | 1,032 | 1,006 | 1,259 | 0,928 | 0,593 | 1,255 | 0,928 | 0,951 | 1,13 | 0,13 | 4,5E-01 |
| Q9CRW3 | UPF0538 protein C2orf76 homolog OS=Mus musculus OX=10090 | 6 | 1 | 1 | 1 | 126 | 14,6 | 7,55 | 455 | | 1 | | 0,901 | 1,085 | 0,656 | 0,907 | 1,361 | 0,732 | 1,127 | 0,867 | 0,87 | -0,13 | 4,5E-01 |
| Q8C6E0 | Cilia- and flagella-associated protein 36 OS=Mus musculus OX=1 | 6 | 1 | 1 | 1 | 343 | 39,6 | 4,93 | | 0 | | 1 | 0,742 | 0,743 | 0,525 | 0,717 | 0,531 | 0,795 | 0,956 | 0,778 | 0,89 | -0,11 | 4,5E-01 |
| Q9ESW4 | Acylglycerol kinase, mitochondrial OS=Mus musculus OX=10090 | 2 | 1 | 1 | 1 | 421 | 46,9 | 8,4 | 357 | | 1 | | 0,969 | 0,928 | 0,91 | 1,117 | 0,795 | 0,958 | 0,903 | 1,041 | 1,06 | 0,06 | 4,5E-01 |
| Q9QY30 | Bile salt export pump OS=Mus musculus OX=10090 GN=Abcb11 | 1 | 1 | 1 | 1 | 1321 | 146,7 | 7,66 | 304 | | 1 | | 1,094 | 0,837 | 0,797 | 0,962 | 0,967 | 0,676 | 1,421 | 1,186 | 0,87 | -0,13 | 4,5E-01 |
| Q8BLR5 | PH and SEC7 domain-containing protein 4 OS=Mus musculus OX | 1 | 1 | 1 | 1 | 1005 | 112,7 | 5,24 | 588 | | 1 | | 1,406 | 1,366 | 1,153 | 1,15 | 2,257 | 1,307 | 0,974 | 1,449 | 0,85 | -0,15 | 4,5E-01 |
| Q8K4G5 | Actin-binding LIM protein 1 OS=Mus musculus OX=10090 GN=Al | 17 | 12 | 17 | 12 | 861 | 96,7 | 8,63 | 7102 | 9,01 | 9 | 3 | 0,937 | 0,88 | 1,009 | 0,96 | 1,061 | 1,004 | 0,931 | 0,926 | 0,97 | -0,03 | 4,5E-01 |
| P62311 | U6 snRNA-associated Sm-like protein LSm3 OS=Mus musculus O | 6 | 1 | 1 | 1 | 102 | 11,8 | 4,7 | 574 | | 1 | | 1,048 | 1,015 | 0,863 | 1,341 | 1,039 | 0,954 | 0,969 | 0,975 | 1,08 | 0,08 | 4,5E-01 |
| Q9WVR4 | Fragile X mental retardation syndrome-related protein 2 OS=Mu | 7 | 3 | 5 | 1 | 673 | 73,7 | 6,23 | 1621 | 2,01 | 2 | 1 | 1,227 | 1,422 | 0,918 | 1,342 | 1,701 | 0,966 | 1,368 | 1,491 | 0,89 | -0,11 | 4,5E-01 |
| P70195 | Proteasome subunit beta type-7 OS=Mus musculus OX=10090 G | 14 | 4 | 14 | 4 | 277 | 29,9 | 7,99 | 8173 | 3,35 | 4 | 1 | 1,104 | 1,149 | 0,925 | 1,095 | 0,99 | 1,107 | 1,043 | 0,933 | 1,05 | 0,05 | 4,5E-01 |
| Q9WV18 | Gamma-aminobutyric acid type B receptor subunit 1 OS=Mus m | 3 | 3 | 5 | 3 | 960 | 108,1 | 8,21 | 1806 | 2,53 | 3 | 1 | 0,765 | 0,882 | 0,886 | 0,928 | 0,963 | 0,905 | 1,059 | 0,764 | 0,94 | -0,06 | 4,5E-01 |
| P27005 | Protein S100-A8 OS=Mus musculus OX=10090 GN=S100a8 PE=1 | 8 | 1 | 1 | 1 | 89 | 10,3 | 5,68 | | 0 | | 1 | 0,785 | 0,557 | 2,705 | 0,709 | 1,063 | 0,592 | 0,888 | 0,519 | 1,55 | 0,55 | 4,5E-01 |
| P16125 | L-lactate dehydrogenase B chain OS=Mus musculus OX=10090 G | 28 | 10 | 30 | 9 | 334 | 36,5 | 6,05 | 10056 | 31,75 | 5 | 6 | 1,036 | 1,048 | 1,051 | 1,016 | 0,875 | 1,136 | 0,959 | 1,003 | 1,04 | 0,04 | 4,5E-01 |
| Q64737 | Trifunctional purine biosynthetic protein adenosine-3 OS=Mus r | 8 | 7 | 7 | 7 | 1010 | 107,4 | 6,68 | 4359 | 2,23 | 6 | 1 | 0,97 | 0,944 | 1,034 | 1,031 | 0,903 | 0,979 | 1,011 | 0,982 | 1,03 | 0,03 | 4,5E-01 |
| Q8BG18 | N-terminal EF-hand calcium-binding protein 1 OS=Mus musculu | 14 | 3 | 4 | 3 | 352 | 40,9 | 4,89 | 2528 | | 3 | | 0,966 | 0,834 | 0,753 | 0,903 | 0,645 | 0,831 | 0,96 | 0,761 | 1,08 | 0,08 | 4,5E-01 |
| P18242 | Cathepsin D OS=Mus musculus OX=10090 GN=Ctsd PE=1 SV=1 | 28 | 10 | 15 | 10 | 410 | 44,9 | 7,15 | 7928 | 2,11 | 9 | 1 | 1,031 | 1,133 | 1,037 | 1,171 | 1,057 | 1,06 | 1,138 | 0,948 | 1,04 | 0,04 | 4,5E-01 |

| | | | | | | | | | | | | | | | | | | | | | | | |
|---|---|---|---|---|---|---|---|---|---|---|---|---|---|---|---|---|---|---|---|---|---|---|---|
| Q02788 | Collagen alpha-2(VI) chain OS=Mus musculus OX=10090 GN=Col | 2 | 1 | 2 | 1 | 1034 | 110,3 | 6,42 | 892 | | 1 | | 1,138 | 1,349 | 1,053 | 0,875 | 1,44 | 1,014 | 0,94 | 1,66 | 0,87 | -0,13 | 4,5E-01 |
| Q0KL02 | Triple functional domain protein OS=Mus musculus OX=10090 G | 5 | 14 | 17 | 12 | 3102 | 347,6 | 6,35 | 6070 | 13,3 | 8 | 6 | 0,927 | 1,002 | 0,967 | 1,086 | 0,913 | 0,94 | 0,969 | 1,026 | 1,03 | 0,03 | 4,5E-01 |
| Q9CQR2 | 40S ribosomal protein S21 OS=Mus musculus OX=10090 GN=Rp: | 39 | 4 | 6 | 4 | 83 | 9,1 | 8,51 | 2386 | 2,16 | 3 | 1 | 0,924 | 1,045 | 0,83 | 0,933 | 1,015 | 0,745 | 0,911 | 0,827 | 1,07 | 0,07 | 4,5E-01 |
| Q91V92 | ATP-citrate synthase OS=Mus musculus OX=10090 GN=Acly PE= | 23 | 23 | 29 | 23 | 1091 | 119,7 | 7,44 | 11913 | 15,34 | 16 | 7 | 1,044 | 0,95 | 1,028 | 0,938 | 0,96 | 1,003 | 0,942 | 0,959 | 1,02 | 0,02 | 4,5E-01 |
| Q9Z0J4 | Nitric oxide synthase, brain OS=Mus musculus OX=10090 GN=N | 1 | 2 | 2 | 2 | 1429 | 160,4 | 7,15 | 613 | 2,62 | 1 | 1 | 0,826 | 0,905 | 0,958 | 0,9 | 1,101 | 0,875 | 0,864 | 0,945 | 0,95 | -0,05 | 4,5E-01 |
| Q9D4H2 | GRIP and coiled-coil domain-containing protein 1 OS=Mus musc | 1 | 1 | 1 | 1 | 778 | 87,6 | 5,54 | 367 | | 1 | | 1,324 | 1,003 | 1,047 | 0,97 | 0,727 | 1,053 | 1,194 | 0,962 | 1,10 | 0,10 | 4,5E-01 |
| Q9CQ91 | NADH dehydrogenase [ubiquinone] 1 alpha subcomplex subunit | 8 | 1 | 1 | 1 | 84 | 9,3 | 8,47 | | 2,79 | | 1 | 1,186 | 1,028 | 0,901 | 0,941 | 0,779 | 1,02 | 1,04 | 0,939 | 1,07 | 0,07 | 4,5E-01 |
| Q640R3 | Hepatocyte cell adhesion molecule OS=Mus musculus OX=1009( | 6 | 2 | 2 | 2 | 418 | 46,3 | 9,42 | 1065 | | 2 | | 1,249 | 0,808 | 1,143 | 1,247 | 0,882 | 0,814 | 1,151 | 1,159 | 1,11 | 0,11 | 4,5E-01 |
| Q9D823 | 60S ribosomal protein L37 OS=Mus musculus OX=10090 GN=Rpl | 14 | 2 | 2 | 2 | 97 | 11,1 | 11,74 | 861 | 0 | 1 | 1 | 0,943 | 0,757 | 0,896 | 0,886 | 0,561 | 0,904 | 0,846 | 0,884 | 1,09 | 0,09 | 4,5E-01 |
| Q07417 | Short-chain specific acyl-CoA dehydrogenase, mitochondrial OS= | 8 | 3 | 3 | 3 | 412 | 44,9 | 8,47 | 1533 | | 3 | | 1,366 | 1,61 | 1,202 | 1,223 | 1,946 | 1,442 | 1,127 | 1,505 | 0,90 | -0,10 | 4,5E-01 |
| Q9WTU6 | Mitogen-activated protein kinase 9 OS=Mus musculus OX=1009( | 12 | 5 | 12 | 1 | 423 | 48,2 | 5,97 | 4713 | 15,18 | 3 | 2 | 1,006 | 1,16 | 0,995 | 0,821 | 0,593 | 1,107 | 1,002 | 0,86 | 1,12 | 0,12 | 4,5E-01 |
| O70591 | Prefoldin subunit 2 OS=Mus musculus OX=10090 GN=Pfdn2 PE= | 33 | 5 | 12 | 5 | 154 | 16,5 | 6,58 | 6608 | | 5 | | 1,01 | 0,895 | 0,876 | 0,997 | 0,934 | 0,927 | 0,964 | 0,783 | 1,05 | 0,05 | 4,5E-01 |
| P35276 | Ras-related protein Rab-3D OS=Mus musculus OX=10090 GN=Ra | 19 | 4 | 11 | 1 | 219 | 24,4 | 4,93 | 5708 | 2,09 | 4 | 1 | 1,06 | 1,131 | 1,036 | 1,158 | 1,054 | 1,228 | 0,754 | 1,023 | 1,08 | 0,08 | 4,6E-01 |
| Q8VDL4 | ADP-dependent glucokinase OS=Mus musculus OX=10090 GN=A | 3 | 1 | 1 | 1 | 496 | 53,9 | 5,62 | 191 | | 1 | | 1,05 | 0,965 | 0,734 | 0,842 | 1,425 | 0,73 | 0,805 | 1,201 | 0,86 | -0,14 | 4,6E-01 |
| P08775 | DNA-directed RNA polymerase II subunit RPB1 OS=Mus musculu | 0 | 1 | 1 | 1 | 1970 | 217 | 7,37 | 513 | | 1 | | 0,891 | 0,887 | 0,864 | 1,024 | 0,847 | 1,002 | 1,427 | 0,843 | 0,89 | -0,11 | 4,6E-01 |
| P05064 | Fructose-bisphosphate aldolase A OS=Mus musculus OX=10090 | 65 | 30 | 78 | 25 | 364 | 39,3 | 8,09 | 21040 | 129,14 | 14 | 22 | 0,964 | 0,977 | 0,974 | 0,977 | 1,694 | 0,956 | 0,952 | 0,896 | 0,87 | -0,13 | 4,6E-01 |
| Q80U49 | Centrosomal protein of 170 kDa protein B OS=Mus musculus OX | 6 | 7 | 7 | 6 | 1574 | 170,7 | 6,87 | 4739 | | 7 | | 0,764 | 0,868 | 0,901 | 0,911 | 1,767 | 0,673 | 0,878 | 0,905 | 0,82 | -0,18 | 4,6E-01 |
| P62748 | Hippocalcin-like protein 1 OS=Mus musculus OX=10090 GN=Hpc | 32 | 7 | 8 | 1 | 193 | 22,3 | 5,5 | 4211 | 2,22 | 6 | 1 | 0,684 | 1,097 | 1,078 | 1,093 | 0,695 | 1,02 | 1,015 | 0,813 | 1,12 | 0,12 | 4,6E-01 |
| Q6P1I6 | PH and SEC7 domain-containing protein 2 OS=Mus musculus OX | 5 | 4 | 4 | 4 | 770 | 84,2 | 5,21 | 1198 | 5,16 | 2 | 2 | 0,922 | 0,929 | 0,935 | 0,93 | 0,885 | 0,883 | 0,894 | 0,98 | 1,02 | 0,02 | 4,6E-01 |
| Q9CZL5 | Pterin-4-alpha-carbinolamine dehydratase 2 OS=Mus musculus ( | 11 | 1 | 1 | 1 | 136 | 14,8 | 9,16 | 644 | | 1 | | 1,245 | 1,115 | 0,924 | 1,34 | 1,715 | 0,873 | 1,169 | 1,521 | 0,88 | -0,12 | 4,6E-01 |
| Q63918 | Caveolae-associated protein 2 OS=Mus musculus OX=10090 GN: | 2 | 1 | 1 | 1 | 418 | 46,7 | 5,21 | 359 | | 1 | | 0,825 | 0,792 | 0,817 | 2,497 | 0,749 | 0,553 | 1,116 | 1,116 | 1,40 | 0,40 | 4,6E-01 |
| P62814 | V-type proton ATPase subunit B, brain isoform OS=Mus musculu | 46 | 23 | 61 | 23 | 511 | 56,5 | 5,81 | 23815 | 61,93 | 16 | 11 | 0,997 | 0,962 | 0,932 | 0,941 | 1,161 | 0,993 | 0,902 | 0,957 | 0,95 | -0,05 | 4,6E-01 |
| B2RY56 | RNA-binding protein 25 OS=Mus musculus OX=10090 GN=Rbm2 | 2 | 1 | 1 | 1 | 838 | 99,5 | 6,32 | 609 | | 1 | | 0,727 | 0,968 | 0,751 | 1,003 | 1,389 | 0,733 | 0,998 | 0,834 | 0,87 | -0,13 | 4,6E-01 |
| Q6ZPQ6 | Membrane-associated phosphatidylinositol transfer protein 2 O | 2 | 2 | 3 | 2 | 1335 | 147,9 | 7,93 | 1697 | | 2 | | 1,235 | 1,207 | 1,105 | 1,22 | 1,125 | 1,299 | 1,067 | 1,086 | 1,04 | 0,04 | 4,6E-01 |
| P28271 | Cytoplasmic aconitate hydratase OS=Mus musculus OX=10090 ( | 13 | 10 | 14 | 10 | 889 | 98,1 | 7,5 | 3568 | 12,01 | 4 | 7 | 1,034 | 0,995 | 1,053 | 1,054 | 0,903 | 1,108 | 0,983 | 1,003 | 1,03 | 0,03 | 4,6E-01 |
| P0DN34 | NADH dehydrogenase [ubiquinone] 1 beta subcomplex subunit | 19 | 1 | 2 | 1 | 57 | 7 | 8,21 | | 7,91 | | 1 | 0,941 | 0,953 | 0,825 | 1,02 | 0,72 | 0,964 | 0,868 | 0,966 | 1,06 | 0,06 | 4,6E-01 |
| Q9CWJ9 | Bifunctional purine biosynthesis protein PURH OS=Mus musculu | 21 | 10 | 14 | 10 | 592 | 64,2 | 6,76 | 8488 | 3,2 | 9 | 1 | 0,957 | 0,956 | 0,907 | 0,916 | 0,872 | 0,954 | 0,923 | 0,92 | 1,02 | 0,02 | 4,6E-01 |
| Q9CX34 | Protein SGT1 homolog OS=Mus musculus OX=10090 GN=Sugt1 I | 4 | 1 | 1 | 1 | 336 | 38,1 | 5,45 | | 2,32 | | 1 | 1,054 | 1,453 | 0,838 | 1,097 | 1,273 | 0,693 | 0,976 | 0,956 | 1,14 | 0,14 | 4,6E-01 |
| Q9D0M1 | Phosphoribosyl pyrophosphate synthase-associated protein 1 O | 10 | 3 | 5 | 1 | 356 | 39,4 | 7,2 | 1415 | 3,5 | 2 | 1 | 1,236 | 1,175 | 1,097 | 1,086 | 1,257 | 1,085 | 1,104 | 1,408 | 0,95 | -0,05 | 4,6E-01 |
| P47753 | F-actin-capping protein subunit alpha-1 OS=Mus musculus OX=1 | 30 | 6 | 14 | 3 | 286 | 32,9 | 5,55 | 3541 | 25,99 | 4 | 4 | 0,752 | 0,861 | 0,715 | 0,772 | 0,94 | 0,658 | 0,907 | 0,813 | 0,93 | -0,07 | 4,7E-01 |
| Q91VZ6 | Stromal membrane-associated protein 1 OS=Mus musculus OX= | 5 | 2 | 6 | 2 | 440 | 47,6 | 8,51 | 2132 | 6,53 | 1 | 1 | 0,891 | 0,902 | 1,05 | 0,846 | 0,804 | 1,01 | 0,738 | 0,907 | 1,07 | 0,07 | 4,7E-01 |
| Q8VBT9 | Tether containing UBX domain for GLUT4 OS=Mus musculus OX: | 16 | 6 | 7 | 6 | 550 | 59,8 | 6,96 | 2122 | 10,94 | 3 | 4 | 1,041 | 0,979 | 1,014 | 1,029 | 1,023 | 0,938 | 0,999 | 1,027 | 1,02 | 0,02 | 4,7E-01 |
| Q8K2Q7 | BRO1 domain-containing protein BROX OS=Mus musculus OX=1 | 10 | 4 | 4 | 4 | 411 | 46,2 | 7,69 | 1336 | 4,53 | 2 | 2 | 1,012 | 0,994 | 1,048 | 1,07 | 0,924 | 1,012 | 1,064 | 1,019 | 1,03 | 0,03 | 4,7E-01 |
| Q9DCW4 | Electron transfer flavoprotein subunit beta OS=Mus musculus O | 46 | 11 | 17 | 11 | 255 | 27,6 | 8,1 | 9568 | 2,91 | 11 | 1 | 0,926 | 0,916 | 0,954 | 1,044 | 0,986 | 0,951 | 0,981 | 1,023 | 0,97 | -0,03 | 4,7E-01 |
| Q8C0M9 | Isoaspartyl peptidase/L-asparaginase OS=Mus musculus OX=10( | 25 | 8 | 11 | 8 | 326 | 33,9 | 7,65 | 5473 | 4,34 | 6 | 2 | 0,962 | 1,042 | 1,017 | 0,99 | 0,92 | 0,986 | 0,967 | 1,043 | 1,02 | 0,02 | 4,7E-01 |
| Q5FWK3 | Rho GTPase-activating protein 1 OS=Mus musculus OX=10090 G | 9 | 4 | 11 | 4 | 439 | 50,4 | 6,44 | 5160 | 2,32 | 2 | 2 | 0,925 | 0,988 | 0,955 | 1,008 | 0,948 | 0,892 | 1,001 | 0,946 | 1,02 | 0,02 | 4,7E-01 |
| A2AHC3 | Calmodulin-regulated spectrin-associated protein 1 OS=Mus mu | 5 | 5 | 7 | 5 | 1581 | 175,8 | 6,95 | 3106 | 1,77 | 4 | 1 | 0,976 | 1,062 | 1,072 | 1,047 | 1,404 | 0,974 | 1,039 | 1,048 | 0,93 | -0,07 | 4,7E-01 |
| Q80VD1 | Protein FAM98B OS=Mus musculus OX=10090 GN=Fam98b PE=: | 3 | 1 | 1 | 1 | 429 | 45,3 | 8,5 | 233 | | 1 | | 1,029 | 1,346 | 1,289 | 1,062 | 1,259 | 1,128 | 1,309 | 1,309 | 0,94 | -0,06 | 4,7E-01 |
| Q9D892 | Inosine triphosphate pyrophosphatase OS=Mus musculus OX=1( | 10 | 2 | 4 | 2 | 198 | 21,9 | 5,87 | | 11,79 | | 2 | 1,065 | 0,891 | 1,018 | 0,997 | 0,665 | 1,079 | 0,903 | 1,02 | 1,08 | 0,08 | 4,7E-01 |
| P61161 | Actin-related protein 2 OS=Mus musculus OX=10090 GN=Actr2 I | 34 | 13 | 38 | 13 | 394 | 44,7 | 6,74 | 12186 | 37,51 | 8 | 7 | 0,976 | 1,046 | 1,06 | 0,956 | 0,916 | 1,006 | 1,001 | 1,01 | 1,03 | 0,03 | 4,7E-01 |
| Q9QWI6 | SRC kinase signaling inhibitor 1 OS=Mus musculus OX=10090 GN | 30 | 30 | 44 | 30 | 1250 | 134,8 | 9,32 | 15168 | 33,97 | 19 | 11 | 0,992 | 1,056 | 1,022 | 0,995 | 0,982 | 1,002 | 1,02 | 1,009 | 1,01 | 0,01 | 4,7E-01 |
| Q9WUL7 | ADP-ribosylation factor-like protein 3 OS=Mus musculus OX=10( | 4 | 1 | 1 | 1 | 182 | 20,5 | 7,24 | | 2,94 | | 1 | 1,471 | 1,16 | 1,276 | 1,034 | 0,729 | 1,395 | 1,292 | 0,977 | 1,12 | 0,12 | 4,7E-01 |
| A2AJI0 | MAP7 domain-containing protein 1 OS=Mus musculus OX=1009 | 4 | 3 | 3 | 3 | 846 | 93,2 | 10,15 | 1010 | 2,13 | 2 | 1 | 0,782 | 0,835 | 0,894 | 0,739 | 1,128 | 0,72 | 1,02 | 0,72 | 0,91 | -0,09 | 4,7E-01 |
| P10518 | Delta-aminolevulinic acid dehydratase OS=Mus musculus OX=1C | 4 | 1 | 1 | 1 | 330 | 36 | 6,79 | 122 | | 1 | | 2,283 | 2,681 | 1,111 | 0,7 | 1,593 | 1,326 | 1,301 | 1,073 | 1,28 | 0,28 | 4,7E-01 |
| Q3UHE1 | Membrane-associated phosphatidylinositol transfer protein 3 O | 1 | 1 | 1 | 1 | 974 | 106,4 | 7,12 | 427 | | 1 | | 1,153 | 1,06 | 1,05 | 1,125 | 1,204 | 0,692 | 1,115 | 1,025 | 1,09 | 0,09 | 4,7E-01 |
| P60521 | Gamma-aminobutyric acid receptor-associated protein-like 2 OS | 38 | 5 | 7 | 4 | 117 | 13,7 | 8,1 | 3238 | 5,74 | 3 | 2 | 0,977 | 1,007 | 1,105 | 1,092 | 1,073 | 1,029 | 0,972 | 0,988 | 1,03 | 0,03 | 4,7E-01 |
| P63005 | Platelet-activating factor acetylhydrolase IB subunit alpha OS=M | 37 | 12 | 19 | 12 | 410 | 46,6 | 7,37 | 4603 | 31,61 | 4 | 9 | 0,976 | 0,913 | 1,084 | 0,94 | 0,777 | 0,994 | 0,963 | 0,984 | 1,05 | 0,05 | 4,7E-01 |
| P62264 | 40S ribosomal protein S14 OS=Mus musculus OX=10090 GN=Rp: | 40 | 6 | 8 | 6 | 151 | 16,3 | 10,05 | 4960 | 1,92 | 5 | 1 | 1 | 1,099 | 1,092 | 1,097 | 1,153 | 1,055 | 1,087 | 1,09 | 0,98 | -0,02 | 4,7E-01 |
| Q8BLR2 | Copine-4 OS=Mus musculus OX=10090 GN=Cpne4 PE=1 SV=1 | 12 | 6 | 21 | 5 | 557 | 62,4 | 6,33 | 3847 | 42,88 | 5 | 3 | 0,847 | 0,637 | 1,591 | 0,575 | 2,676 | 0,847 | 1,008 | 0,702 | 0,70 | -0,30 | 4,7E-01 |
| P36993 | Protein phosphatase 1B OS=Mus musculus OX=10090 GN=Ppm1 | 4 | 2 | 2 | 1 | 390 | 42,8 | 5,19 | 671 | 2,1 | 1 | 1 | 0,963 | 1,236 | 1,304 | 1,28 | 0,696 | 1,386 | 1,188 | 1,003 | 1,12 | 0,12 | 4,7E-01 |
| Q6ZQ58 | La-related protein 1 OS=Mus musculus OX=10090 GN=Larp1 PE= | 6 | 4 | 4 | 4 | 1072 | 121,1 | 8,79 | 1966 | 2,83 | 3 | 1 | 0,881 | 0,803 | 0,897 | 0,807 | 0,818 | 0,743 | 0,817 | 0,891 | 1,04 | 0,04 | 4,7E-01 |
| Q8BG02 | Serine/threonine-protein phosphatase 2A 55 kDa regulatory sub | 6 | 3 | 7 | 1 | 447 | 51,4 | 6,37 | 3017 | 5,21 | 2 | 1 | 1,134 | 0,975 | 0,943 | 0,997 | 0,891 | 1,077 | 0,97 | 0,934 | 1,05 | 0,05 | 4,7E-01 |
| Q9DCS9 | NADH dehydrogenase [ubiquinone] 1 beta subcomplex subunit | 26 | 4 | 7 | 4 | 176 | 21 | 8,03 | 3524 | 0 | 3 | 1 | 0,96 | 1,12 | 0,851 | 1,305 | 0,688 | 1,082 | 0,997 | 1,058 | 1,11 | 0,11 | 4,7E-01 |
| Q9CPR4 | 60S ribosomal protein L17 OS=Mus musculus OX=10090 GN=Rpl | 17 | 3 | 4 | 3 | 184 | 21,4 | 10,18 | | 7,96 | | 3 | 1,01 | 0,955 | 0,913 | 1,02 | 0,713 | 0,982 | 0,984 | 0,996 | 1,06 | 0,06 | 4,7E-01 |
| Q8BHC1 | Ras-related protein Rab-39B OS=Mus musculus OX=10090 GN=F | 11 | 4 | 4 | 4 | 213 | 24,6 | 7,83 | 841 | 3,73 | 2 | 2 | 1,243 | 1,141 | 1,149 | 1,007 | 1,028 | 1,125 | 1,053 | 1,159 | 1,04 | 0,04 | 4,7E-01 |
| Q8C078 | Calcium/calmodulin-dependent protein kinase kinase 2 OS=Mus | 14 | 8 | 9 | 8 | 588 | 64,6 | 5,9 | 5430 | 0 | 7 | 1 | 1,071 | 0,983 | 1,076 | 1,018 | 0,906 | 1,086 | 1,026 | 0,997 | 1,03 | 0,03 | 4,7E-01 |
| Q6NXK8 | Acid-sensing ion channel 1 OS=Mus musculus OX=10090 GN=Asi | 2 | 1 | 1 | 1 | 526 | 59,6 | 5,58 | 408 | | 1 | | 1,129 | 1,376 | 1,535 | 1,527 | 1,112 | 0,918 | 1,258 | 1,697 | 1,12 | 0,12 | 4,7E-01 |
| Q922X9 | Protein arginine N-methyltransferase 7 OS=Mus musculus OX=1 | 2 | 1 | 1 | 1 | 692 | 78,3 | 5,74 | 219 | | 1 | | 0,698 | 0,904 | 0,654 | 0,799 | 2,287 | 0,582 | 0,616 | 0,822 | 0,71 | -0,29 | 4,7E-01 |
| Q99NE5 | Regulating synaptic membrane exocytosis protein 1 OS=Mus mu | 6 | 8 | 9 | 2 | 1463 | 163,1 | 9,42 | 5910 | 0 | 7 | 1 | 0,941 | 0,941 | 0,843 | 0,72 | 0,71 | 0,822 | 0,977 | 0,674 | 1,08 | 0,08 | 4,8E-01 |
| Q9JI46 | Diphosphoinositol polyphosphate phosphohydrolase 1 OS=Mus | 27 | 4 | 5 | 3 | 168 | 19 | 6,34 | 2373 | 2,05 | 3 | 1 | 0,693 | 0,785 | 0,947 | 0,956 | 0,634 | 0,99 | 0,691 | 0,758 | 1,10 | 0,10 | 4,8E-01 |
| Q8VED9 | Galectin-related protein OS=Mus musculus OX=10090 GN=Lgals | 14 | 3 | 4 | 3 | 172 | 18,9 | 5,35 | 1471 | 2,16 | 2 | 1 | 0,794 | 0,919 | 1,039 | 0,901 | 0,678 | 1,036 | 0,813 | 0,854 | 1,08 | 0,08 | 4,8E-01 |
| P09405 | Nucleolin OS=Mus musculus OX=10090 GN=Ncl PE=1 SV=2 | 21 | 16 | 19 | 16 | 707 | 76,7 | 4,75 | 7948 | 15,98 | 10 | 6 | 0,948 | 1,033 | 0,973 | 1,161 | 1,01 | 0,887 | 1,013 | 1,029 | 1,04 | 0,04 | 4,8E-01 |
| P35979 | 60S ribosomal protein L12 OS=Mus musculus OX=10090 GN=Rpl | 32 | 4 | 7 | 4 | 165 | 17,8 | 9,42 | 747 | 21,27 | 1 | 4 | 0,972 | 0,888 | 0,942 | 0,973 | 0,838 | 0,886 | 0,991 | 0,942 | 1,03 | 0,03 | 4,8E-01 |
| Q9Z1B3 | 1-phosphatidylinositol 4,5-bisphosphate phosphodiesterase bet | 18 | 20 | 33 | 20 | 1216 | 138,3 | 6,13 | 10372 | 35,54 | 12 | 10 | 1,026 | 0,977 | 0,923 | 0,966 | 0,86 | 1,003 | 0,969 | 0,947 | 1,03 | 0,03 | 4,8E-01 |
| P60041 | Somatostatin OS=Mus musculus OX=10090 GN=Sst PE=1 SV=1 | 7 | 1 | 1 | 1 | 116 | 12,7 | 5,52 | | 2,6 | | 1 | 0,708 | 0,985 | 0,989 | 0,851 | 0,598 | 1,017 | 0,775 | 0,812 | 1,10 | 0,10 | 4,8E-01 |

| | | | | | | | | | | | | | | | | | | | | | | | |
|---|---|---|---|---|---|---|---|---|---|---|---|---|---|---|---|---|---|---|---|---|---|---|---|
| Q812A2 | SLIT-ROBO Rho GTPase-activating protein 3 OS=Mus musculus C | 15 | 15 | 19 | 15 | 1099 | 124,3 | 6,67 | 5216 | 23,56 | 7 | 8 | 1,024 | 1,028 | 1,021 | 0,995 | 1,018 | 0,937 | 1,033 | 1,011 | 1,02 | 0,02 | 4,8E-01 |
| Q07813 | Apoptosis regulator BAX OS=Mus musculus OX=10090 GN=Bax F | 10 | 1 | 2 | 1 | 192 | 21,4 | 4,98 | 463 | 3,58 | 1 | 1 | 0,954 | 0,977 | 0,501 | 0,805 | 1,128 | 0,478 | 0,895 | 1,433 | 0,82 | -0,18 | 4,8E-01 |
| Q9CR51 | V-type proton ATPase subunit G 1 OS=Mus musculus OX=10090 | 9 | 1 | 1 | 1 | 118 | 13,7 | 7,97 | 890 | | 1 | | 0,839 | 0,939 | 0,836 | 1,108 | 0,583 | 0,927 | 1,004 | 0,868 | 1,10 | 0,10 | 4,8E-01 |
| Q8CH77 | Neuron navigator 1 OS=Mus musculus OX=10090 GN=Nav1 PE=: | 14 | 20 | 29 | 19 | 1875 | 202,2 | 8,06 | 9191 | 23,99 | 11 | 9 | 1,072 | 0,984 | 1,077 | 1,005 | 0,976 | 0,958 | 1,047 | 1,054 | 1,03 | 0,03 | 4,8E-01 |
| Q3B7Z2 | Oxysterol-binding protein 1 OS=Mus musculus OX=10090 GN=O | 9 | 4 | 6 | 4 | 805 | 88,7 | 7,2 | 3041 | | 4 | | 0,986 | 0,941 | 0,991 | 1,106 | 1,043 | 0,985 | 0,981 | 0,853 | 1,04 | 0,04 | 4,8E-01 |
| Q9DBC7 | cAMP-dependent protein kinase type I-alpha regulatory subunit | 13 | 4 | 4 | 2 | 381 | 43,2 | 5,35 | 1804 | 3,42 | 3 | 1 | 0,883 | 0,907 | 0,955 | 0,871 | 0,961 | 0,907 | 0,991 | 0,861 | 0,97 | -0,03 | 4,8E-01 |
| Q9DCL9 | Multifunctional protein ADE2 OS=Mus musculus OX=10090 GN= | 17 | 8 | 15 | 8 | 425 | 47 | 7,23 | 4966 | 9,6 | 5 | 3 | 0,97 | 1,068 | 1,21 | 0,984 | 0,992 | 1,037 | 1,03 | 1,003 | 1,04 | 0,04 | 4,8E-01 |
| Q69ZX8 | Actin-binding LIM protein 3 OS=Mus musculus OX=10090 GN=Al | 3 | 2 | 2 | 1 | 682 | 77,6 | 8,54 | 1056 | | 2 | | 0,845 | 1,065 | 1,069 | 0,922 | 1,368 | 0,925 | 0,962 | 0,999 | 0,92 | -0,08 | 4,8E-01 |
| P29391 | Ferritin light chain 1 OS=Mus musculus OX=10090 GN=Ftl1 PE=1 | 26 | 3 | 6 | 3 | 183 | 20,8 | 6 | 1178 | 5,42 | 2 | 2 | 1,014 | 1,038 | 0,829 | 1,04 | 1,211 | 0,912 | 0,886 | 1,235 | 0,92 | -0,08 | 4,8E-01 |
| Q56A10 | Zinc finger protein 608 OS=Mus musculus OX=10090 GN=Znf608 | 1 | 1 | 1 | 1 | 1511 | 162,2 | 8,81 | 448 | | 1 | | 0,513 | 0,406 | 2,373 | 0,597 | 0,045 | 1,044 | 0,722 | 0,525 | 1,66 | 0,66 | 4,8E-01 |
| O35350 | Calpain-1 catalytic subunit OS=Mus musculus OX=10090 GN=Ca| | 3 | 2 | 4 | 2 | 713 | 82,1 | 5,87 | 2103 | | 2 | | 1 | 1,047 | 0,877 | 0,935 | 0,904 | 0,772 | 1,006 | 0,982 | 1,05 | 0,05 | 4,8E-01 |
| Q91YI0 | Argininosuccinate lyase OS=Mus musculus OX=10090 GN=Asl PE | 2 | 1 | 2 | 1 | 464 | 51,7 | 6,96 | 1269 | | 1 | | 0,932 | 1,143 | 1,062 | 1,131 | 1,132 | 1,326 | 1,014 | 1,052 | 0,94 | -0,06 | 4,8E-01 |
| Q8VE33 | Ganglioside-induced differentiation-associated protein 1-like 1 C | 14 | 4 | 9 | 4 | 370 | 42,3 | 6,6 | 4682 | 2,5 | 3 | 1 | 0,989 | 1,037 | 0,933 | 1,046 | 1,018 | 1,131 | 0,966 | 1,021 | 0,97 | -0,03 | 4,8E-01 |
| A6H6A9-3 | Isoform 3 of Rab GTPase-activating protein 1-like OS=Mus musc | 14 | 5 | 10 | 3 | 370 | 42,7 | 7,33 | 3496 | 7,24 | 4 | 1 | 1,088 | 1,074 | 1,147 | 1,083 | 0,987 | 1,106 | 1,069 | 1,125 | 1,02 | 0,02 | 4,8E-01 |
| Q9D1H8 | 39S ribosomal protein L53, mitochondrial OS=Mus musculus OX | 8 | 1 | 1 | 1 | 118 | 12,7 | 9,52 | 698 | | 1 | | 0,904 | 1,017 | 0,706 | 1,051 | 1,012 | 0,892 | 1,128 | 0,929 | 0,93 | -0,07 | 4,8E-01 |
| Q60625 | Intercellular adhesion molecule 5 OS=Mus musculus OX=10090 | 3 | 4 | 4 | 4 | 917 | 96,9 | 6,32 | 1383 | 2,56 | 3 | 1 | 1,033 | 1,184 | 1 | 0,9 | 0,869 | 1,065 | 1,006 | 0,96 | 1,06 | 0,06 | 4,8E-01 |
| P61255 | 60S ribosomal protein L26 OS=Mus musculus OX=10090 GN=Rpl | 44 | 8 | 12 | 8 | 145 | 17,2 | 10,55 | 1878 | 20,77 | 2 | 6 | 1,164 | 1,039 | 0,992 | 1,081 | 1,029 | 1,039 | 1,012 | 1,078 | 1,03 | 0,03 | 4,8E-01 |
| Q8BH57 | WD repeat-containing protein 48 OS=Mus musculus OX=10090 | 12 | 7 | 8 | 7 | 676 | 76 | 7,17 | 2357 | 9,06 | 4 | 3 | 0,997 | 1,074 | 0,919 | 0,925 | 0,993 | 1,018 | 1,027 | 0,99 | 0,97 | -0,03 | 4,8E-01 |
| Q9Z0E0 | Neurochondrin OS=Mus musculus OX=10090 GN=Ncdn PE=1 SV | 10 | 7 | 13 | 7 | 729 | 78,8 | 5,54 | 6347 | 3,25 | 5 | 2 | 0,983 | 0,931 | 1,111 | 0,927 | 0,589 | 1,132 | 0,948 | 0,919 | 1,10 | 0,10 | 4,8E-01 |
| P61222 | ATP-binding cassette sub-family E member 1 OS=Mus musculus | 12 | 6 | 9 | 6 | 599 | 67,3 | 8,34 | 4063 | 0 | 6 | 1 | 0,998 | 1,024 | 1,071 | 1,053 | 1,085 | 1,067 | 1,139 | 0,971 | 0,97 | -0,03 | 4,8E-01 |
| Q69ZT1 | Fanconi-associated nuclease 1 OS=Mus musculus OX=10090 GN | 1 | 1 | 2 | 1 | 1020 | 112,9 | 7,44 | 1243 | | 1 | | 0,943 | 1,02 | 1,017 | 0,915 | 0,773 | 0,992 | 1,022 | 0,923 | 1,05 | 0,05 | 4,8E-01 |
| Q2PFD7 | PH and SEC7 domain-containing protein 3 OS=Mus musculus OX | 9 | 11 | 14 | 11 | 1037 | 114,7 | 6,24 | 5448 | 7,99 | 6 | 5 | 1,015 | 1,024 | 0,9 | 0,98 | 0,82 | 0,893 | 1,024 | 1,012 | 1,05 | 0,05 | 4,8E-01 |
| P53702 | Cytochrome c-type heme lyase OS=Mus musculus OX=10090 GN | 3 | 1 | 1 | 1 | 272 | 31 | 7,12 | 391 | | 1 | | 1,387 | 1,242 | 1,074 | 1,468 | 1,346 | 1,112 | 1,248 | 1,163 | 1,06 | 0,06 | 4,8E-01 |
| Q80Z38 | SH3 and multiple ankyrin repeat domains protein 2 OS=Mus mu | 9 | 12 | 12 | 11 | 1476 | 158,9 | 7,18 | 3904 | 13,47 | 7 | 5 | 1,122 | 1,042 | 1,101 | 1,079 | 1,017 | 1,086 | 1,098 | 1,069 | 1,02 | 0,02 | 4,8E-01 |
| Q8CHG7 | Rap guanine nucleotide exchange factor 2 OS=Mus musculus OX | 4 | 4 | 4 | 4 | 1496 | 166,3 | 6,61 | 304 | 4,58 | 1 | 3 | 0,844 | 0,815 | 1,025 | 1,002 | 0,793 | 0,932 | 0,813 | 0,948 | 1,06 | 0,06 | 4,8E-01 |
| Q8BTJ4 | Bis(5'-adenosyl)-triphosphatase enpp4 OS=Mus musculus OX=1( | 2 | 1 | 1 | 1 | 456 | 51,6 | 6,73 | 421 | | 1 | | 0,895 | 0,98 | 1,075 | 1,242 | 1,059 | 1,146 | 1,19 | 1,043 | 0,94 | -0,06 | 4,8E-01 |
| P61971 | Nuclear transport factor 2 OS=Mus musculus OX=10090 GN=Nu | 28 | 2 | 2 | 2 | 127 | 14,5 | 5,38 | 269 | 2,65 | 1 | 1 | 0,67 | 0,882 | 0,758 | 0,619 | 0,521 | 0,907 | 0,813 | 1,075 | 0,88 | -0,12 | 4,8E-01 |
| Q8JZW4 | Copine-5 OS=Mus musculus OX=10090 GN=Cpne5 PE=1 SV=1 | 8 | 4 | 5 | 2 | 593 | 65,6 | 5,77 | 3325 | | 4 | | 1,082 | 1,029 | 1,551 | 1,103 | 0,98 | 1,238 | 1,084 | 1,068 | 1,09 | 0,09 | 4,8E-01 |
| Q8CHT0 | Delta-1-pyrroline-5-carboxylate dehydrogenase, mitochondrial ( | 17 | 9 | 11 | 9 | 562 | 61,8 | 8,24 | 4269 | 6,36 | 6 | 3 | 1,047 | 0,988 | 0,999 | 1,066 | 1,43 | 0,971 | 0,908 | 1,144 | 0,92 | -0,08 | 4,8E-01 |
| Q9D2V7 | Coronin-7 OS=Mus musculus OX=10090 GN=Coro7 PE=1 SV=2 | 5 | 3 | 3 | 3 | 922 | 100,7 | 5,77 | 867 | 2,75 | 2 | 1 | 0,983 | 1,055 | 1,009 | 1,028 | 0,917 | 0,984 | 0,97 | 1,088 | 1,03 | 0,03 | 4,8E-01 |
| Q9JMG1 | Endothelial differentiation-related factor 1 OS=Mus musculus O | 10 | 1 | 1 | 1 | 148 | 16,4 | 9,99 | 351 | | 1 | | 0,97 | 1,026 | 1,384 | 1,141 | 1,166 | 1,066 | 1,18 | 1,5 | 0,92 | -0,08 | 4,9E-01 |
| P56376 | Acylphosphatase-1 OS=Mus musculus OX=10090 GN=Acyp1 PE= | 27 | 2 | 2 | 2 | 99 | 11,2 | 9,04 | 593 | | 2 | | 1,128 | 0,912 | 1,025 | 0,957 | 1,202 | 1,045 | 0,893 | 1,122 | 0,94 | -0,06 | 4,9E-01 |
| Q80VP0 | Tectonin beta-propeller repeat-containing protein 1 OS=Mus mu | 2 | 2 | 2 | 2 | 1166 | 130,2 | 6,48 | 1272 | | 2 | | 1,228 | 0,994 | 0,812 | 0,919 | 0,986 | 0,989 | 0,844 | 0,845 | 1,08 | 0,08 | 4,9E-01 |
| Q60715 | Prolyl 4-hydroxylase subunit alpha-1 OS=Mus musculus OX=100 | 2 | 1 | 1 | 1 | 534 | 60,9 | 5,9 | | 2,44 | | 1 | 0,958 | 0,767 | 0,892 | 0,982 | 1,031 | 0,839 | 0,972 | 0,943 | 0,95 | -0,05 | 4,9E-01 |
| Q9JKF6 | Nectin-1 OS=Mus musculus OX=10090 GN=Nectin1 PE=1 SV=3 | 13 | 6 | 7 | 6 | 515 | 57 | 6,35 | 2018 | 8,16 | 2 | 4 | 1,043 | 0,82 | 0,907 | 0,924 | 0,988 | 1,086 | 0,845 | 0,975 | 0,95 | -0,05 | 4,9E-01 |
| Q3UGR5 | Haloacid dehalogenase-like hydrolase domain-containing protei | 7 | 2 | 2 | 2 | 259 | 28,7 | 6,05 | 1099 | | 2 | | 0,797 | 0,829 | 1,111 | 1,056 | 1,013 | 0,941 | 0,968 | 1,138 | 0,93 | -0,07 | 4,9E-01 |
| Q64374 | Regucalcin OS=Mus musculus OX=10090 GN=Rgn PE=1 SV=1 | 12 | 3 | 4 | 3 | 299 | 33,4 | 5,34 | 959 | 3 | 2 | 1 | 1,176 | 0,895 | 0,964 | 1,003 | 1,042 | 0,73 | 1,134 | 1,859 | 0,85 | -0,15 | 4,9E-01 |
| Q8BMJ3 | Eukaryotic translation initiation factor 1A, X-chromosomal OS=N | 18 | 2 | 5 | 2 | 144 | 16,5 | 5,24 | 2203 | 2,45 | 2 | 1 | 1,179 | 1,034 | 0,95 | 1,094 | 1,121 | 0,995 | 1,202 | 1,13 | 0,96 | -0,04 | 4,9E-01 |
| Q9D6R2 | Isocitrate dehydrogenase [NAD] subunit alpha, mitochondrial O | 25 | 10 | 19 | 10 | 366 | 39,6 | 6,73 | 7633 | 20,14 | 7 | 4 | 1,02 | 0,909 | 1,063 | 0,939 | 1,002 | 0,985 | 0,866 | 0,94 | 1,04 | 0,04 | 4,9E-01 |
| Q99JW2 | Aminoacylase-1 OS=Mus musculus OX=10090 GN=Acy1 PE=1 SV | 5 | 2 | 2 | 2 | 408 | 45,8 | 6,32 | 1264 | | 2 | | 0,969 | 0,902 | 0,688 | 1,08 | 2,127 | 1,109 | 0,618 | 0,806 | 0,78 | -0,22 | 4,9E-01 |
| P22892 | AP-1 complex subunit gamma-1 OS=Mus musculus OX=10090 G | 6 | 6 | 7 | 6 | 822 | 91,3 | 6,8 | 3117 | 5,32 | 4 | 2 | 1,026 | 0,997 | 1,124 | 0,983 | 0,761 | 1,168 | 0,968 | 0,971 | 1,07 | 0,07 | 4,9E-01 |
| O89079 | Coatomer subunit epsilon OS=Mus musculus OX=10090 GN=Cop | 14 | 4 | 5 | 4 | 308 | 34,5 | 5,06 | 1829 | 4,64 | 3 | 1 | 1,024 | 0,94 | 1,05 | 0,966 | 0,981 | 0,791 | 0,857 | 1,123 | 1,06 | 0,06 | 4,9E-01 |
| P35283 | Ras-related protein Rab-12 OS=Mus musculus OX=10090 GN=Ra | 13 | 2 | 3 | 2 | 243 | 27,3 | 8,41 | 1444 | 2,8 | 1 | 1 | 0,937 | 0,937 | 1,057 | 0,822 | 1,047 | 0,951 | 0,929 | 0,986 | 0,96 | -0,04 | 4,9E-01 |
| Q9R0N7-3 | Isoform 3 of Synaptotagmin-7 OS=Mus musculus OX=10090 GN= | 10 | 3 | 3 | 3 | 447 | 50,6 | 9,11 | 1803 | | 3 | | 0,838 | 0,799 | 0,875 | 0,821 | 0,832 | 0,927 | 0,801 | 0,865 | 0,97 | -0,03 | 4,9E-01 |
| O08917 | Flotillin-1 OS=Mus musculus OX=10090 GN=Flot1 PE=1 SV=1 | 29 | 11 | 16 | 11 | 428 | 47,5 | 7,15 | 5242 | 19,03 | 6 | 5 | 0,914 | 0,897 | 1,246 | 0,969 | 1,134 | 0,881 | 0,837 | 0,861 | 1,08 | 0,08 | 4,9E-01 |
| P29595 | NEDD8 OS=Mus musculus OX=10090 GN=Nedd8 PE=1 SV=2 | 17 | 2 | 6 | 2 | 81 | 9 | 7,25 | 3164 | | 2 | | 0,789 | 0,927 | 0,681 | 0,91 | 0,907 | 0,846 | 1,054 | 0,751 | 0,93 | -0,07 | 4,9E-01 |
| Q62108 | Disks large homolog 4 OS=Mus musculus OX=10090 GN=Dlg4 PE | 25 | 18 | 24 | 15 | 724 | 80,4 | 5,87 | 10587 | 3,88 | 15 | 3 | 0,904 | 1,051 | 0,907 | 0,913 | 0,78 | 0,873 | 0,995 | 0,953 | 1,05 | 0,05 | 4,9E-01 |
| Q8BGD9 | Eukaryotic translation initiation factor 4B OS=Mus musculus OX= | 13 | 6 | 6 | 6 | 611 | 68,8 | 5,67 | 2016 | 0 | 5 | 1 | 0,921 | 0,966 | 1,087 | 1,09 | 0,881 | 1,073 | 0,938 | 0,998 | 1,04 | 0,04 | 4,9E-01 |
| Q9WTX6 | Cullin-1 OS=Mus musculus OX=10090 GN=Cul1 PE=1 SV=1 | 6 | 4 | 6 | 4 | 776 | 89,6 | 8 | 2465 | 2,47 | 2 | 2 | 1,081 | 0,917 | 1,113 | 0,882 | 0,953 | 1,018 | 0,96 | 0,871 | 1,05 | 0,05 | 4,9E-01 |
| Q9DB20 | ATP synthase subunit O, mitochondrial OS=Mus musculus OX=1( | 52 | 11 | 21 | 11 | 213 | 23,3 | 9,99 | 9633 | 14,36 | 8 | 5 | 1,053 | 0,931 | 1,058 | 0,992 | 0,798 | 1,077 | 0,982 | 0,985 | 1,05 | 0,05 | 4,9E-01 |
| Q4KMM3 | Oxidation resistance protein 1 OS=Mus musculus OX=10090 GN | 11 | 7 | 10 | 7 | 866 | 95,9 | 5,33 | 3484 | 3,27 | 7 | 1 | 1,088 | 1,081 | 0,899 | 0,991 | 1,056 | 1,001 | 1,011 | 1,158 | 0,96 | -0,04 | 4,9E-01 |
| P29341 | Polyadenylate-binding protein 1 OS=Mus musculus OX=10090 G | 25 | 17 | 24 | 17 | 636 | 70,6 | 9,5 | 8445 | 28,56 | 9 | 8 | 0,929 | 0,95 | 1,013 | 0,995 | 1,052 | 0,927 | 0,952 | 1,078 | 0,97 | -0,03 | 4,9E-01 |
| P17665 | Cytochrome c oxidase subunit 7C, mitochondrial OS=Mus muscu | 29 | 3 | 4 | 3 | 63 | 7,3 | 11 | 987 | 9,88 | 2 | 1 | 1,014 | 0,896 | 1,233 | 1,201 | 1 | 1,236 | 1,075 | 1,367 | 0,93 | -0,07 | 4,9E-01 |
| Q9CWX2 | Complex I intermediate-associated protein 30, mitochondrial OS | 14 | 3 | 3 | 3 | 328 | 37,8 | 8,6 | 1062 | | 3 | | 1,036 | 1,03 | 1,164 | 1,319 | 1,282 | 1,043 | 1,126 | 1,406 | 0,94 | -0,06 | 4,9E-01 |
| P15864 | Histone H1.2 OS=Mus musculus OX=10090 GN=Hist1h1c PE=1 S\ | 47 | 16 | 25 | 8 | 212 | 21,3 | 11 | 5220 | 64,65 | 3 | 13 | 0,851 | 0,678 | 1,021 | 1,054 | 0,923 | 0,83 | 1,165 | 1,013 | 0,92 | -0,08 | 4,9E-01 |
| Q9D1G5 | Leucine-rich repeat-containing protein 57 OS=Mus musculus OX | 17 | 4 | 4 | 4 | 239 | 26,7 | 8,35 | 2753 | | 4 | | 0,908 | 0,959 | 1,05 | 0,914 | 1,009 | 0,961 | 0,932 | 0,72 | 1,06 | 0,06 | 4,9E-01 |
| E9Q7X7 | Neurexin-2 OS=Mus musculus OX=10090 GN=Nrxn2 PE=1 SV=1 | 3 | 5 | 10 | 3 | 1710 | 184,8 | 5,9 | 303 | 17,84 | 1 | 5 | 1,182 | 1,156 | 1,174 | 1,192 | 0,995 | 1,285 | 0,988 | 1,214 | 1,05 | 0,05 | 4,9E-01 |
| Q922P8 | Transmembrane protein 132A OS=Mus musculus OX=10090 GN | 3 | 2 | 3 | 2 | 1018 | 110,2 | 5,63 | 1638 | | 2 | | 1,012 | 1,018 | 0,946 | 1,11 | 1,447 | 0,854 | 1,1 | 1,056 | 0,92 | -0,08 | 4,9E-01 |
| P26443 | Glutamate dehydrogenase 1, mitochondrial OS=Mus musculus C | 44 | 24 | 80 | 24 | 558 | 61,3 | 8 | 26195 | 113,36 | 16 | 17 | 0,985 | 1,017 | 1,059 | 1,003 | 0,974 | 0,953 | 1,02 | 1,042 | 1,02 | 0,02 | 5,0E-01 |
| Q2WF71 | Leucine-rich repeat and fibronectin type III domain-containing p | 1 | 1 | 1 | 1 | 766 | 81,9 | 7,59 | 759 | | 1 | | 1,078 | 1,053 | 0,99 | 0,928 | 1,05 | 1,129 | 1,194 | 0,89 | 0,95 | -0,05 | 5,0E-01 |
| Q9WTK3 | Glycosylphosphatidylinositol anchor attachment 1 protein OS=N | 2 | 1 | 2 | 1 | 621 | 67,9 | 8,4 | | 2,96 | | 1 | 0,985 | 0,706 | 1,176 | 1,004 | 0,741 | 0,658 | 1,126 | 0,932 | 1,12 | 0,12 | 5,0E-01 |
| P08003 | Protein disulfide-isomerase A4 OS=Mus musculus OX=10090 GN | 32 | 21 | 31 | 21 | 638 | 71,9 | 5,31 | 16190 | 13,73 | 16 | 5 | 1,012 | 0,952 | 1,006 | 1,13 | 1,015 | 0,956 | 1,022 | 0,99 | 1,03 | 0,03 | 5,0E-01 |
| Q7TPH6 | E3 ubiquitin-protein ligase MYCBP2 OS=Mus musculus OX=1009 | 0 | 1 | 1 | 1 | 4749 | 520,9 | 7,09 | 384 | | 1 | | 1,042 | 0,864 | 1,047 | 0,907 | 0,599 | 0,913 | 1,084 | 0,939 | 1,09 | 0,09 | 5,0E-01 |
| Q791V5 | Mitochondrial carrier homolog 2 OS=Mus musculus OX=10090 G | 7 | 2 | 2 | 2 | 303 | 33,5 | 8,25 | 829 | 2,47 | 1 | 1 | 1,057 | 1,072 | 1,001 | 1,001 | 0,775 | 1,104 | 1,034 | 1,005 | 1,05 | 0,05 | 5,0E-01 |
| A6H5Z3 | Exocyst complex component 6B OS=Mus musculus OX=10090 G | 3 | 2 | 2 | 2 | 810 | 94,1 | 6,39 | 1046 | | 2 | | 1,394 | 1,014 | 1,528 | 1,158 | 1,222 | 1,116 | 1,155 | 1,255 | 1,07 | 0,07 | 5,0E-01 |

| | | | | | | | | | | | | | | | | | | | | | | | |
|---|---|---|---|---|---|---|---|---|---|---|---|---|---|---|---|---|---|---|---|---|---|---|---|
| P03893 | NADH-ubiquinone oxidoreductase chain 2 OS=Mus musculus OX | 2 | 1 | 1 | 1 | 345 | 38,7 | 9,92 | 550 | | 1 | | 1,251 | 0,834 | 0,829 | 0,944 | 0,943 | 0,965 | 0,972 | 1,414 | 0,90 | -0,10 | 5,0E-01 |
| Q9R111 | Guanine deaminase OS=Mus musculus OX=10090 GN=Gda PE=1 | 33 | 15 | 22 | 15 | 454 | 51 | 5,53 | 8398 | 19,18 | 9 | 6 | 1,038 | 1,107 | 0,885 | 1,048 | 0,831 | 1,133 | 0,941 | 0,946 | 1,06 | 0,06 | 5,0E-01 |
| Q61879 | Myosin-10 OS=Mus musculus OX=10090 GN=Myh10 PE=1 SV=2 | 36 | 71 | 111 | 57 | 1976 | 228,9 | 5,54 | 41939 | 121,46 | 43 | 31 | 1,011 | 0,977 | 0,951 | 1,028 | 0,901 | 0,93 | 1,036 | 0,998 | 1,03 | 0,03 | 5,0E-01 |
| Q8BRK8 | 5'-AMP-activated protein kinase catalytic subunit alpha-2 OS=M | 7 | 4 | 4 | 4 | 552 | 62 | 7,87 | 1678 | 2,08 | 3 | 1 | 0,922 | 1,032 | 1,184 | 1,01 | 0,919 | 1,024 | 1,044 | 0,985 | 1,04 | 0,04 | 5,0E-01 |
| Q8CI71 | Syndetin OS=Mus musculus OX=10090 GN=Vps50 PE=1 SV=2 | 3 | 2 | 2 | 2 | 964 | 111,1 | 6,07 | 733 | 2,41 | 1 | 1 | 1,065 | 1,192 | 0,779 | 1,373 | 0,458 | 1,142 | 1,035 | 1,173 | 1,16 | 0,16 | 5,0E-01 |
| P56382 | ATP synthase subunit epsilon, mitochondrial OS=Mus musculus | 27 | 2 | 2 | 2 | 52 | 5,8 | 10,01 | | 4,64 | | 2 | 0,827 | 0,784 | 1,106 | 0,896 | 0,892 | 1,315 | 0,942 | 0,837 | 0,91 | -0,09 | 5,0E-01 |
| Q9CZ44 | NSFL1 cofactor p47 OS=Mus musculus OX=10090 GN=Nsfl1c PE= | 44 | 14 | 21 | 14 | 370 | 40,7 | 5,15 | 8397 | 15,85 | 10 | 6 | 0,948 | 0,955 | 0,876 | 1,06 | 0,969 | 0,977 | 0,985 | 1,022 | 0,97 | -0,03 | 5,0E-01 |
| Q8VCA8 | Secernin-2 OS=Mus musculus OX=10090 GN=Scrn2 PE=1 SV=1 | 4 | 1 | 1 | 1 | 425 | 46,6 | 5,71 | 271 | | 1 | | 0,638 | 0,778 | 0,598 | 0,94 | 0,887 | 0,67 | 0,925 | 0,752 | 0,91 | -0,09 | 5,0E-01 |
| P51174 | Long-chain specific acyl-CoA dehydrogenase, mitochondrial OS= | 29 | 10 | 12 | 10 | 430 | 47,9 | 8,31 | 6275 | 6,88 | 9 | 3 | 1,001 | 0,992 | 0,956 | 1,077 | 1,25 | 0,93 | 1,047 | 1,008 | 0,95 | -0,05 | 5,0E-01 |
| Q60790 | Ras GTPase-activating protein 3 OS=Mus musculus OX=10090 GI | 2 | 2 | 2 | 2 | 834 | 95,9 | 7,34 | 1050 | | 2 | | 1,045 | 1,03 | 1,366 | 1,196 | 1,855 | 0,991 | 1,341 | 1,057 | 0,88 | -0,12 | 5,0E-01 |
| Q9QYR9 | Acyl-coenzyme A thioesterase 2, mitochondrial OS=Mus muscul | 9 | 4 | 6 | 1 | 453 | 49,6 | 7,36 | 1665 | 9,39 | 2 | 2 | 1,058 | 1,465 | 0,824 | 1,191 | 1,25 | 0,923 | 0,996 | 0,926 | 1,11 | 0,11 | 5,0E-01 |
| Q91VE6 | MKI67 FHA domain-interacting nucleolar phosphoprotein OS=M | 2 | 1 | 3 | 1 | 317 | 36,2 | 10,24 | 1610 | | 1 | | 0,985 | 1,073 | 0,962 | 1,083 | 0,934 | 0,969 | 1,146 | 1,329 | 0,94 | -0,06 | 5,0E-01 |
| P29533 | Vascular cell adhesion protein 1 OS=Mus musculus OX=10090 G | 2 | 1 | 1 | 1 | 739 | 81,3 | 5,3 | | 2,39 | | 1 | 0,785 | 0,757 | 0,592 | 0,865 | 0,958 | 0,885 | 0,717 | 0,688 | 0,92 | -0,08 | 5,0E-01 |
| Q60668 | Heterogeneous nuclear ribonucleoprotein D0 OS=Mus musculus | 11 | 5 | 7 | 5 | 355 | 38,3 | 7,81 | 2555 | 9,13 | 3 | 2 | 0,87 | 1,09 | 0,867 | 1,031 | 1,002 | 0,887 | 1,001 | 1,231 | 0,94 | -0,06 | 5,0E-01 |
| P59281 | Rho GTPase-activating protein 39 OS=Mus musculus OX=10090 | 2 | 2 | 2 | 2 | 1107 | 125,1 | 7,97 | 970 | | 2 | | 1,222 | 1,102 | 1,05 | 1,13 | 1,028 | 1,094 | 1,215 | 0,993 | 1,04 | 0,04 | 5,0E-01 |
| Q9R0P5 | Destrin OS=Mus musculus OX=10090 GN=Dstn PE=1 SV=3 | 41 | 8 | 15 | 8 | 165 | 18,5 | 7,97 | 4806 | 19,95 | 5 | 4 | 1,093 | 1,034 | 1,118 | 0,981 | 1,349 | 1,192 | 0,966 | 0,992 | 0,94 | -0,06 | 5,0E-01 |
| Q99020 | Heterogeneous nuclear ribonucleoprotein A/B OS=Mus musculu | 5 | 2 | 4 | 2 | 285 | 30,8 | 7,91 | 702 | 10,24 | 1 | 1 | 0,908 | 0,895 | 1,041 | 1,106 | 1,089 | 0,882 | 1,166 | 1,039 | 0,95 | -0,05 | 5,0E-01 |
| P97351 | 40S ribosomal protein S3a OS=Mus musculus OX=10090 GN=Rp: | 45 | 14 | 21 | 14 | 264 | 29,9 | 9,73 | 8304 | 14,42 | 10 | 5 | 1,061 | 1,073 | 1,164 | 1,08 | 1,161 | 1,006 | 1,009 | 1,078 | 1,03 | 0,03 | 5,0E-01 |
| Q9JKC8 | AP-3 complex subunit mu-1 OS=Mus musculus OX=10090 GN=A | 11 | 3 | 3 | 2 | 418 | 46,9 | 6,93 | 1584 | | 3 | | 0,968 | 1,052 | 1,225 | 1,076 | 0,813 | 1,404 | 1,145 | 1,379 | 0,91 | -0,09 | 5,0E-01 |
| Q61290 | Voltage-dependent R-type calcium channel subunit alpha-1E OS | 1 | 2 | 2 | 2 | 2272 | 257,1 | 8,18 | 794 | | 2 | | 0,872 | 0,888 | 1,313 | 1,038 | 1,723 | 1,164 | 0,868 | 0,972 | 0,87 | -0,13 | 5,0E-01 |
| Q9D0E1 | Heterogeneous nuclear ribonucleoprotein M OS=Mus musculus | 27 | 16 | 19 | 16 | 729 | 77,6 | 8,63 | 5330 | 20,22 | 6 | 11 | 0,823 | 0,849 | 1,065 | 1,009 | 0,9 | 0,94 | 1,107 | 1,012 | 0,95 | -0,05 | 5,0E-01 |
| P60487 | Pyridoxal phosphate phosphatase OS=Mus musculus OX=10090 | 25 | 6 | 8 | 6 | 292 | 31,5 | 5,74 | 3193 | 7,39 | 4 | 2 | 0,926 | 1,026 | 0,989 | 1,037 | 1,033 | 0,95 | 0,961 | 0,941 | 1,02 | 0,02 | 5,0E-01 |
| O70293 | G protein-coupled receptor kinase 6 OS=Mus musculus OX=100! | 4 | 2 | 3 | 2 | 576 | 65,9 | 8,1 | 1545 | | 2 | | 0,787 | 0,809 | 0,79 | 0,822 | 0,779 | 0,806 | 0,79 | 0,947 | 0,97 | -0,03 | 5,0E-01 |
| P54822 | Adenylosuccinate lyase OS=Mus musculus OX=10090 GN=Adsl P | 10 | 3 | 3 | 3 | 484 | 54,8 | 7,27 | 1276 | | 3 | | 0,9 | 1,285 | 0,988 | 1,003 | 1,287 | 0,961 | 1,084 | 1,149 | 0,93 | -0,07 | 5,1E-01 |
| O55234 | Proteasome subunit beta type-5 OS=Mus musculus OX=10090 G | 13 | 3 | 4 | 3 | 264 | 28,5 | 7,02 | 1735 | 2,63 | 2 | 1 | 1,025 | 1,091 | 0,942 | 1,092 | 1,019 | 0,841 | 0,92 | 1,154 | 1,05 | 0,05 | 5,1E-01 |
| Q9DC69 | NADH dehydrogenase [ubiquinone] 1 alpha subcomplex subunit | 23 | 8 | 12 | 8 | 377 | 42,5 | 9,74 | 5353 | 9,73 | 5 | 3 | 1,016 | 0,905 | 1,11 | 0,994 | 0,961 | 1,015 | 0,96 | 0,964 | 1,03 | 0,03 | 5,1E-01 |
| P24668 | Cation-dependent mannose-6-phosphate receptor OS=Mus mus | 3 | 1 | 1 | 1 | 278 | 31,2 | 5,39 | 486 | | 1 | | 1,26 | 1,165 | 0,832 | 1,214 | 0,999 | 0,725 | 1,117 | 1,221 | 1,10 | 0,10 | 5,1E-01 |
| Q6P9R2 | Serine/threonine-protein kinase OSR1 OS=Mus musculus OX=10 | 8 | 4 | 7 | 4 | 527 | 58,2 | 6,43 | 2921 | | 4 | | 0,695 | 0,78 | 0,646 | 0,622 | 0,578 | 0,668 | 0,894 | 0,831 | 0,92 | -0,08 | 5,1E-01 |
| Q69ZH9 | Rho GTPase-activating protein 23 OS=Mus musculus OX=10090 | 5 | 5 | 6 | 5 | 1483 | 161,7 | 8,98 | 2763 | 2,83 | 4 | 1 | 1,024 | 1,011 | 0,98 | 1,044 | 1,316 | 0,939 | 1,067 | 0,979 | 0,94 | -0,06 | 5,1E-01 |
| P97372 | Proteasome activator complex subunit 2 OS=Mus musculus OX= | 11 | 3 | 3 | 3 | 239 | 27 | 5,76 | 1242 | 3,79 | 2 | 1 | 1,121 | 0,909 | 1,009 | 1,076 | 0,703 | 1,008 | 1,11 | 1,013 | 1,07 | 0,07 | 5,1E-01 |
| Q8BK72 | 28S ribosomal protein S27, mitochondrial OS=Mus musculus OX | 5 | 2 | 2 | 2 | 415 | 47,7 | 5,5 | 1495 | | 2 | | 1,081 | 0,969 | 0,96 | 0,844 | 0,938 | 1,005 | 0,899 | 0,846 | 1,05 | 0,05 | 5,1E-01 |
| Q99M28 | RNA-binding protein with serine-rich domain 1 OS=Mus musculu | 5 | 1 | 1 | 1 | 305 | 34,2 | 11,84 | 681 | | 1 | | 0,62 | 0,709 | 0,759 | 0,784 | 0,797 | 0,533 | 1,193 | 0,75 | 0,88 | -0,12 | 5,1E-01 |
| Q05186 | Reticulocalbin-1 OS=Mus musculus OX=10090 GN=Rcn1 PE=1 SV | 14 | 5 | 6 | 5 | 325 | 38,1 | 4,84 | 3056 | | 5 | | 0,959 | 1,019 | 0,933 | 0,991 | 1,204 | 0,831 | 0,995 | 1,103 | 0,94 | -0,06 | 5,1E-01 |
| Q6VNS1 | NT-3 growth factor receptor OS=Mus musculus OX=10090 GN=N | 9 | 7 | 10 | 2 | 825 | 92,7 | 6,67 | 3587 | 5,31 | 4 | 3 | 1,027 | 0,908 | 0,901 | 0,998 | 1,086 | 0,967 | 0,882 | 1,057 | 0,96 | -0,04 | 5,1E-01 |
| Q71RI9 | Kynurenine--oxoglutarate transaminase 3 OS=Mus musculus OX | 2 | 1 | 1 | 1 | 455 | 51,1 | 8,37 | 541 | | 1 | | 0,814 | 1,213 | 0,861 | 0,974 | 1,113 | 0,793 | 1,221 | 1,095 | 0,91 | -0,09 | 5,1E-01 |
| P48036 | Annexin A5 OS=Mus musculus OX=10090 GN=Anxa5 PE=1 SV=1 | 20 | 7 | 11 | 7 | 319 | 35,7 | 4,96 | 3575 | 3,11 | 4 | 3 | 1,07 | 1,147 | 1,086 | 1,357 | 1,214 | 1,088 | 1,091 | 1,057 | 1,05 | 0,05 | 5,1E-01 |
| Q3UHH2 | Solute carrier family 22 member 23 OS=Mus musculus OX=1009 | 8 | 4 | 5 | 4 | 689 | 74,3 | 7,83 | 1381 | 5,42 | 2 | 2 | 1,236 | 1,244 | 1,013 | 1,071 | 1,143 | 0,993 | 1,124 | 1,114 | 1,04 | 0,04 | 5,1E-01 |
| Q9ES28 | Rho guanine nucleotide exchange factor 7 OS=Mus musculus OX | 12 | 11 | 12 | 11 | 862 | 97 | 6,8 | 2968 | 15,97 | 5 | 6 | 1,028 | 0,973 | 1,109 | 0,993 | 1,027 | 0,944 | 1,021 | 1,011 | 1,02 | 0,02 | 5,1E-01 |
| P60766-1 | Isoform 1 of Cell division control protein 42 homolog OS=Mus m | 36 | 7 | 16 | 1 | 191 | 21,3 | 6,04 | 5688 | 13,13 | 5 | 4 | 1,205 | 0,716 | 1,337 | 1,015 | 1,199 | 1,054 | 1,194 | 1,219 | 0,92 | -0,08 | 5,1E-01 |
| P07901 | Heat shock protein HSP 90-alpha OS=Mus musculus OX=10090 ( | 44 | 36 | 95 | 21 | 733 | 84,7 | 5,01 | 43112 | 76,9 | 25 | 16 | 0,959 | 0,966 | 0,989 | 0,98 | 0,897 | 0,992 | 0,966 | 0,977 | 1,02 | 0,02 | 5,1E-01 |
| Q62077 | 1-phosphatidylinositol 4,5-bisphosphate phosphodiesterase gan | 1 | 2 | 3 | 2 | 1302 | 149,6 | 5,8 | 1745 | | 2 | | 0,895 | 0,971 | 0,924 | 1,036 | 1,246 | 0,908 | 1,025 | 0,892 | 0,94 | -0,06 | 5,1E-01 |
| Q8R570 | Synaptosomal-associated protein 47 OS=Mus musculus OX=100! | 24 | 7 | 7 | 7 | 413 | 46,5 | 5,76 | 3964 | | 7 | | 1,236 | 1,048 | 1,132 | 1,286 | 1,301 | 1,065 | 1,28 | 1,271 | 0,96 | -0,04 | 5,1E-01 |
| Q99KF1 | Transmembrane emp24 domain-containing protein 9 OS=Mus m | 9 | 2 | 2 | 2 | 235 | 27,1 | 8,41 | 1310 | | 2 | | 1,193 | 0,802 | 1,374 | 1,099 | 1,154 | 1,165 | 1,31 | 1,189 | 0,93 | -0,07 | 5,1E-01 |
| Q99KV1 | DnaJ homolog subfamily B member 11 OS=Mus musculus OX=1( | 18 | 4 | 5 | 4 | 358 | 40,5 | 6,32 | 2079 | | 4 | | 1,002 | 1,033 | 0,799 | 0,998 | 0,944 | 0,951 | 1,049 | 1,062 | 0,96 | -0,04 | 5,1E-01 |
| O35969 | Guanidinoacetate N-methyltransferase OS=Mus musculus OX=1 | 10 | 1 | 1 | 1 | 236 | 26,3 | 5,71 | 233 | | 1 | | 0,882 | 1,008 | 0,893 | 1,231 | 0,959 | 0,813 | 0,935 | 1,044 | 1,07 | 0,07 | 5,1E-01 |
| Q8VCT3 | Aminopeptidase B OS=Mus musculus OX=10090 GN=Rnpep PE= | 8 | 4 | 4 | 4 | 650 | 72,4 | 5,35 | 483 | 7,31 | 1 | 3 | 1,085 | 1,11 | 0,817 | 0,964 | 1,054 | 0,993 | 0,952 | 1,232 | 0,94 | -0,06 | 5,1E-01 |
| Q9Z2X1 | Heterogeneous nuclear ribonucleoprotein F OS=Mus musculus ( | 17 | 5 | 5 | 4 | 415 | 45,7 | 5,49 | 3193 | | 5 | | 0,912 | 1,019 | 1,098 | 1,116 | 0,619 | 1,081 | 1,017 | 1,089 | 1,09 | 0,09 | 5,1E-01 |
| Q80U28 | MAP kinase-activating death domain protein OS=Mus musculus | 3 | 4 | 6 | 4 | 1577 | 175,1 | 6,04 | 2222 | 4,8 | 3 | 1 | 1,004 | 0,951 | 1,066 | 0,965 | 1,13 | 1,014 | 0,966 | 0,999 | 0,97 | -0,03 | 5,1E-01 |
| P45878 | Peptidyl-prolyl cis-trans isomerase FKBP2 OS=Mus musculus OX: | 25 | 4 | 4 | 4 | 140 | 15,3 | 8,88 | 644 | 6,23 | 1 | 3 | 0,864 | 1,015 | 0,971 | 1,1 | 0,876 | 0,932 | 0,984 | 1 | 1,04 | 0,04 | 5,1E-01 |
| Q8BJH1 | Zinc finger C2HC domain-containing protein 1A OS=Mus muscul | 45 | 13 | 24 | 13 | 324 | 35,1 | 9,88 | 6559 | 24,41 | 6 | 7 | 1,102 | 0,911 | 1,186 | 1,074 | 1,039 | 1,037 | 0,975 | 1,054 | 1,04 | 0,04 | 5,1E-01 |
| Q641K1 | Cytosolic carboxypeptidase 1 OS=Mus musculus OX=10090 GN= | 2 | 2 | 3 | 2 | 1218 | 137,1 | 6,52 | 1162 | | 2 | | 1,29 | 1,221 | 1,51 | 1,444 | 1,45 | 1,17 | 1,339 | 1,256 | 1,05 | 0,05 | 5,1E-01 |
| P56393 | Cytochrome c oxidase subunit 7B, mitochondrial OS=Mus muscu | 9 | 1 | 2 | 1 | 80 | 9 | 10,27 | 1209 | | 1 | | 0,804 | 0,734 | 1,18 | 0,616 | 0,639 | 1,034 | 0,745 | 0,427 | 1,17 | 0,17 | 5,1E-01 |
| P35710 | Transcription factor SOX-5 OS=Mus musculus OX=10090 GN=So: | 2 | 1 | 2 | 1 | 763 | 84 | 6,6 | 428 | | 1 | | 1,073 | 1,03 | 1,055 | 0,943 | 1,082 | 1,211 | 1,136 | 0,883 | 0,95 | -0,05 | 5,1E-01 |
| P55264 | Adenosine kinase OS=Mus musculus OX=10090 GN=Adk PE=1 S\ | 14 | 4 | 11 | 4 | 361 | 40,1 | 6,21 | 4289 | 13,21 | 2 | 3 | 1,396 | 1,294 | 1,163 | 1,374 | 1,626 | 1,41 | 1,193 | 1,295 | 0,95 | -0,05 | 5,1E-01 |
| P35278 | Ras-related protein Rab-5C OS=Mus musculus OX=10090 GN=Ra | 16 | 3 | 9 | 1 | 216 | 23,4 | 8,41 | 3469 | 4,68 | 3 | 1 | 0,841 | 1,108 | 0,938 | 1,229 | 0,966 | 1,143 | 1,102 | 1,178 | 0,94 | -0,06 | 5,1E-01 |
| Q78RX3 | Small integral membrane protein 12 OS=Mus musculus OX=100! | 9 | 1 | 1 | 1 | 92 | 10,8 | 9,04 | | 2,29 | | 1 | 0,999 | 0,837 | 0,905 | 0,83 | 0,618 | 0,978 | 0,937 | 0,786 | 1,08 | 0,08 | 5,1E-01 |
| Q8VCM8 | Nicalin OS=Mus musculus OX=10090 GN=Ncln PE=1 SV=2 | 2 | 1 | 1 | 1 | 563 | 62,9 | 6,49 | | 2,19 | | 1 | 0,708 | 0,851 | 0,732 | 0,775 | 0,909 | 0,874 | 0,693 | 0,755 | 0,95 | -0,05 | 5,1E-01 |
| Q9JIF7 | Coatomer subunit beta OS=Mus musculus OX=10090 GN=Copb1 | 7 | 4 | 4 | 4 | 953 | 107 | 6 | 1429 | 3,36 | 3 | 1 | 0,868 | 0,856 | 0,967 | 0,964 | 1,004 | 0,969 | 0,907 | 0,887 | 0,97 | -0,03 | 5,1E-01 |
| Q80YN3 | Breast carcinoma-amplified sequence 1 homolog OS=Mus musc | 2 | 1 | 2 | 1 | 633 | 67,3 | 6,21 | | 5,19 | | 1 | 1,033 | 0,838 | 1,025 | 0,889 | 0,663 | 0,87 | 1,012 | 0,983 | 1,07 | 0,07 | 5,2E-01 |
| O88741 | Ganglioside-induced differentiation-associated protein 1 OS=Mu | 16 | 6 | 9 | 6 | 358 | 41,3 | 8,37 | 2015 | 20,71 | 3 | 4 | 1,091 | 1,06 | 1,127 | 1,035 | 0,858 | 1,091 | 1,113 | 1,078 | 1,04 | 0,04 | 5,2E-01 |
| Q8VE62 | Polyadenylate-binding protein-interacting protein 1 OS=Mus mu | 3 | 1 | 2 | 1 | 400 | 45,7 | 4,55 | 719 | | 1 | | 0,976 | 1,167 | 1,205 | 1,025 | 0,957 | 1,045 | 1,166 | 1,01 | 1,05 | 0,05 | 5,2E-01 |
| P35293 | Ras-related protein Rab-18 OS=Mus musculus OX=10090 GN=Ra | 37 | 7 | 9 | 6 | 206 | 23 | 5,36 | 4482 | 2,66 | 6 | 1 | 0,927 | 1,102 | 1,013 | 1,131 | 1,144 | 1,002 | 1,093 | 1,085 | 0,97 | -0,03 | 5,2E-01 |
| Q8CG76 | Aflatoxin B1 aldehyde reductase member 2 OS=Mus musculus C | 12 | 3 | 3 | 3 | 367 | 40,6 | 8,12 | 994 | 2,88 | 2 | 1 | 1,073 | 0,896 | 0,941 | 0,934 | 1,407 | 0,873 | 0,895 | 1,026 | 0,92 | -0,08 | 5,2E-01 |
| P47738 | Aldehyde dehydrogenase, mitochondrial OS=Mus musculus OX= | 16 | 9 | 16 | 7 | 519 | 56,5 | 7,62 | 5464 | 16,99 | 5 | 5 | 1,337 | 1,314 | 1,122 | 1,544 | 1,439 | 1,035 | 1,3 | 1,21 | 1,07 | 0,07 | 5,2E-01 |
| Q9R1V7 | Disintegrin and metalloproteinase domain-containing protein 23 | 3 | 2 | 3 | 2 | 829 | 91,5 | 7,68 | 235 | 4,61 | 1 | 1 | 0,984 | 1,038 | 0,969 | 0,908 | 0,753 | 1,074 | 0,963 | 0,911 | 1,05 | 0,05 | 5,2E-01 |

| | | | | | | | | | | | | | | | | | | | | | | | |
|---|---|---|---|---|---|---|---|---|---|---|---|---|---|---|---|---|---|---|---|---|---|---|---|
| Q00612 | Glucose-6-phosphate 1-dehydrogenase X OS=Mus musculus OX= | 16 | 8 | 10 | 8 | 515 | 59,2 | 6,49 | 3414 | 6,69 | 5 | 3 | 0,867 | 1,003 | 0,937 | 1,004 | 1,06 | 0,917 | 1 | 0,957 | 0,97 | -0,03 | 5,2E-01 |
| Q9JHW2 | Omega-amidase NIT2 OS=Mus musculus OX=10090 GN=Nit2 PE= | 13 | 3 | 3 | 3 | 276 | 30,5 | 6,9 | 1600 | | 3 | | 1,122 | 1,023 | 0,848 | 1,059 | 1,314 | 0,89 | 1,043 | 1,095 | 0,93 | -0,07 | 5,2E-01 |
| Q8BG92 | Clavesin-2 OS=Mus musculus OX=10090 GN=Clvs2 PE=1 SV=1 | 10 | 3 | 3 | 2 | 327 | 37,9 | 6,2 | 1736 | | 3 | | 0,68 | 0,313 | 2,086 | 0,591 | 0,489 | 0,835 | 0,702 | 0,531 | 1,44 | 0,44 | 5,2E-01 |
| O09159 | Lysosomal alpha-mannosidase OS=Mus musculus OX=10090 GN | 1 | 1 | 1 | 1 | 1013 | 114,6 | 8,13 | 624 | | 1 | | 1,304 | 1,381 | 0,808 | 1,521 | 0,973 | 1,285 | 0,996 | 1,274 | 1,11 | 0,11 | 5,2E-01 |
| Q9D4H8 | Cullin-2 OS=Mus musculus OX=10090 GN=Cul2 PE=1 SV=2 | 5 | 4 | 4 | 4 | 745 | 86,8 | 7,01 | 1980 | 2,85 | 3 | 1 | 1,029 | 0,969 | 1,226 | 0,941 | 0,994 | 1,095 | 1,076 | 1,217 | 0,95 | -0,05 | 5,2E-01 |
| Q9WUR2 | Enoyl-CoA delta isomerase 2, mitochondrial OS=Mus musculus ( | 4 | 2 | 2 | 2 | 391 | 43,2 | 8,92 | 732 | 3,73 | 1 | 1 | 0,802 | 1,031 | 1,145 | 1,029 | 0,958 | 0,939 | 0,968 | 0,944 | 1,05 | 0,05 | 5,2E-01 |
| Q8K3G9 | DCC-interacting protein 13-beta OS=Mus musculus OX=10090 G | 1 | 1 | 1 | 1 | 662 | 73,8 | 5,03 | | 2,26 | | 1 | 0,857 | 0,708 | 0,815 | 0,838 | 0,96 | 0,74 | 0,793 | 0,887 | 0,95 | -0,05 | 5,2E-01 |
| Q9WUA3 | ATP-dependent 6-phosphofructokinase, platelet type OS=Mus n | 10 | 8 | 15 | 5 | 784 | 85,4 | 7,11 | 6421 | 6,43 | 7 | 2 | 0,999 | 1,102 | 1,025 | 1,02 | 0,896 | 1,087 | 0,997 | 1,039 | 1,03 | 0,03 | 5,2E-01 |
| P70168 | Importin subunit beta-1 OS=Mus musculus OX=10090 GN=Kpnb | 10 | 8 | 14 | 8 | 876 | 97,1 | 4,78 | 2891 | 29,77 | 3 | 6 | 1,023 | 0,945 | 1,039 | 1,065 | 1,115 | 1,018 | 1,039 | 0,999 | 0,98 | -0,02 | 5,2E-01 |
| Q4QQM4 | Tumor protein p53-inducible protein 11 OS=Mus musculus OX= | 8 | 1 | 1 | 1 | 189 | 20,9 | 9,41 | 384 | | 1 | | 0,876 | 0,817 | 0,762 | 0,776 | 1,536 | 0,582 | 1,209 | 0,563 | 0,83 | -0,17 | 5,2E-01 |
| Q8R066 | Complement C1q tumor necrosis factor-related protein 4 OS=M | 8 | 2 | 3 | 2 | 326 | 35 | 8,98 | 2031 | | 2 | | 1,459 | 1,219 | 1,209 | 1,37 | 1,677 | 1,15 | 0,921 | 1,022 | 1,10 | 0,10 | 5,2E-01 |
| Q6IFZ6 | Keratin, type II cytoskeletal 1b OS=Mus musculus OX=10090 GN | 8 | 8 | 21 | 1 | 572 | 61,3 | 8,02 | 5631 | 26,33 | 4 | 6 | 1,176 | 1,004 | 1,005 | 1,022 | 2,123 | 0,967 | 1,058 | 0,863 | 0,84 | -0,16 | 5,2E-01 |
| Q8R146 | Acylamino-acid-releasing enzyme OS=Mus musculus OX=10090 | 8 | 6 | 6 | 6 | 732 | 81,5 | 5,59 | 1862 | 6,2 | 4 | 2 | 1,12 | 1,128 | 1,041 | 1,211 | 0,947 | 1,259 | 1,029 | 1,062 | 1,05 | 0,05 | 5,2E-01 |
| Q6PB44 | Tyrosine-protein phosphatase non-receptor type 23 OS=Mus mu | 4 | 5 | 6 | 5 | 1692 | 185,1 | 6,8 | 1827 | 5,51 | 3 | 2 | 1,167 | 1,138 | 1,183 | 1,23 | 1,606 | 1,098 | 1,233 | 1,108 | 0,94 | -0,06 | 5,2E-01 |
| Q80ZI6 | E3 ubiquitin-protein ligase LRSAM1 OS=Mus musculus OX=1009 | 6 | 4 | 5 | 4 | 727 | 83,9 | 6,14 | 1880 | 2,38 | 3 | 1 | 1,048 | 1,003 | 1,026 | 1,027 | 1,247 | 0,927 | 1,121 | 1,001 | 0,96 | -0,04 | 5,2E-01 |
| Q8C0I1 | Alkyldihydroxyacetonephosphate synthase, peroxisomal OS=Mu | 3 | 2 | 2 | 2 | 645 | 71,6 | 7,5 | 1277 | | 2 | | 1,011 | 0,902 | 0,889 | 1,04 | 0,823 | 1,109 | 1,038 | 1,075 | 0,95 | -0,05 | 5,2E-01 |
| Q91WG4 | Elongator complex protein 2 OS=Mus musculus OX=10090 GN=E | 4 | 3 | 3 | 3 | 831 | 93 | 5,82 | 584 | 2,7 | 1 | 2 | 1,014 | 1,139 | 1,055 | 1,143 | 0,807 | 1,054 | 1,252 | 0,973 | 1,06 | 0,06 | 5,2E-01 |
| Q9D074 | E3 ubiquitin-protein ligase MGRN1 OS=Mus musculus OX=1009( | 2 | 1 | 1 | 1 | 532 | 58,4 | 4,88 | 513 | | 1 | | 0,935 | 0,865 | 1,053 | 1,045 | 0,845 | 1,151 | 1,001 | 1,127 | 0,95 | -0,05 | 5,2E-01 |
| Q9CVB6 | Actin-related protein 2/3 complex subunit 2 OS=Mus musculus ( | 30 | 10 | 36 | 10 | 300 | 34,3 | 7,36 | 18812 | 5,82 | 8 | 5 | 0,945 | 0,92 | 1,005 | 0,913 | 0,837 | 0,992 | 0,944 | 0,905 | 1,03 | 0,03 | 5,2E-01 |
| O35927 | Catenin delta-2 OS=Mus musculus OX=10090 GN=Ctnnd2 PE=1 S | 16 | 16 | 21 | 14 | 1247 | 134,9 | 7,65 | 8113 | 13,83 | 11 | 6 | 0,947 | 1,028 | 1,097 | 0,921 | 0,998 | 0,876 | 0,98 | 1,004 | 1,03 | 0,03 | 5,2E-01 |
| Q9R099 | Transducin beta-like protein 2 OS=Mus musculus OX=10090 GN= | 3 | 1 | 1 | 1 | 442 | 49,6 | 9,04 | 631 | | 1 | | 0,849 | 0,92 | 0,953 | 0,765 | 1,093 | 0,748 | 0,808 | 1,12 | 0,93 | -0,07 | 5,2E-01 |
| Q80U58 | Pumilio homolog 2 OS=Mus musculus OX=10090 GN=Pum2 PE= | 1 | 1 | 1 | 1 | 1066 | 114,2 | 7,08 | 669 | | 1 | | 0,956 | 0,798 | 0,88 | 0,857 | 0,503 | 0,888 | 0,899 | 0,918 | 1,09 | 0,09 | 5,3E-01 |
| Q9CQC7 | NADH dehydrogenase [ubiquinone] 1 beta subcomplex subunit | 54 | 7 | 10 | 7 | 129 | 15,1 | 9,89 | 5152 | 5,62 | 7 | 1 | 1,037 | 0,894 | 1,006 | 0,993 | 0,95 | 1,044 | 1,04 | 0,998 | 0,97 | -0,03 | 5,3E-01 |
| Q7TPD3 | Roundabout homolog 2 OS=Mus musculus OX=10090 GN=Robo | 11 | 11 | 12 | 11 | 1470 | 161,1 | 6,33 | 4058 | 8,21 | 7 | 4 | 1,019 | 0,97 | 0,96 | 0,988 | 0,955 | 1,024 | 1,082 | 0,963 | 0,98 | -0,02 | 5,3E-01 |
| P59759 | Myocardin-related transcription factor B OS=Mus musculus OX= | 2 | 2 | 3 | 2 | 1080 | 117,5 | 6,16 | 1066 | | 2 | | 1,027 | 0,878 | 0,991 | 1,061 | 0,838 | 1,023 | 1 | 0,942 | 1,04 | 0,04 | 5,3E-01 |
| Q68ED7 | CREB-regulated transcription coactivator 1 OS=Mus musculus O | 3 | 2 | 3 | 2 | 630 | 66,9 | 6,05 | 1784 | | 2 | | 0,997 | 0,949 | 1,001 | 0,831 | 0,809 | 0,895 | 1,21 | 1,144 | 0,93 | -0,07 | 5,3E-01 |
| Q8K3H0 | DCC-interacting protein 13-alpha OS=Mus musculus OX=10090 ( | 7 | 4 | 4 | 4 | 707 | 79,3 | 5,41 | 1688 | 2,55 | 3 | 1 | 1,022 | 0,995 | 0,902 | 1,007 | 0,883 | 1,069 | 0,909 | 0,932 | 1,04 | 0,04 | 5,3E-01 |
| Q61390 | T-complex protein 1 subunit zeta-2 OS=Mus musculus OX=1009( | 13 | 6 | 12 | 1 | 531 | 58,1 | 7,36 | 5289 | 10,34 | 5 | 2 | 1,148 | 0,919 | 0,912 | 0,908 | 0,835 | 0,862 | 1,003 | 0,991 | 1,05 | 0,05 | 5,3E-01 |
| Q9CZS1 | Aldehyde dehydrogenase X, mitochondrial OS=Mus musculus O | 32 | 15 | 21 | 13 | 519 | 57,5 | 7,02 | 6215 | 28,36 | 7 | 8 | 1,029 | 1,034 | 1,02 | 0,998 | 0,925 | 1,06 | 1,006 | 1,012 | 1,02 | 0,02 | 5,3E-01 |
| O55029 | Coatomer subunit beta' OS=Mus musculus OX=10090 GN=Copb | 7 | 7 | 7 | 7 | 905 | 102,4 | 5,3 | 2017 | 7,87 | 4 | 3 | 1,056 | 1,049 | 1,032 | 1,137 | 1,089 | 0,954 | 0,993 | 1,117 | 1,03 | 0,03 | 5,3E-01 |
| Q8K354 | Carbonyl reductase [NADPH] 3 OS=Mus musculus OX=10090 GN | 6 | 2 | 3 | 1 | 277 | 30,9 | 6,57 | 1261 | | 2 | | 1,133 | 0,866 | 0,727 | 1,19 | 1,077 | 1,089 | 1,069 | 0,982 | 0,93 | -0,07 | 5,3E-01 |
| Q91VT4 | Carbonyl reductase family member 4 OS=Mus musculus OX=100 | 11 | 2 | 3 | 2 | 236 | 25,4 | 9,69 | 1984 | | 2 | | 0,827 | 0,765 | 0,797 | 0,926 | 0,708 | 0,794 | 0,779 | 0,895 | 1,04 | 0,04 | 5,3E-01 |
| Q9Z1Q5 | Chloride intracellular channel protein 1 OS=Mus musculus OX=1 | 15 | 3 | 3 | 3 | 241 | 27 | 5,17 | 711 | 2,38 | 2 | 1 | 0,927 | 0,972 | 1,103 | 1,061 | 1,188 | 0,931 | 1,069 | 1,052 | 0,96 | -0,04 | 5,3E-01 |
| Q9EPL2 | Calsyntenin-1 OS=Mus musculus OX=10090 GN=Clstn1 PE=1 SV= | 2 | 3 | 3 | 3 | 979 | 108,8 | 4,92 | 937 | 2,26 | 2 | 1 | 0,869 | 0,989 | 0,868 | 0,912 | 0,941 | 0,796 | 1,089 | 0,992 | 0,95 | -0,05 | 5,3E-01 |
| P60469 | Liprin-alpha-3 OS=Mus musculus OX=10090 GN=Ppfia3 PE=1 SV= | 21 | 22 | 31 | 19 | 1194 | 133,3 | 5,64 | 11154 | 27,16 | 14 | 9 | 0,981 | 0,922 | 0,961 | 0,936 | 0,977 | 0,917 | 0,972 | 0,989 | 0,99 | -0,01 | 5,3E-01 |
| P55088 | Aquaporin-4 OS=Mus musculus OX=10090 GN=Aqp4 PE=1 SV=2 | 6 | 2 | 2 | 2 | 323 | 34,4 | 7,42 | 1551 | | 2 | | 1,05 | 1,064 | 0,929 | 1,009 | 0,77 | 1,04 | 1,208 | 0,716 | 1,09 | 0,09 | 5,3E-01 |
| P08113 | Endoplasmin OS=Mus musculus OX=10090 GN=Hsp90b1 PE=1 S | 31 | 26 | 45 | 24 | 802 | 92,4 | 4,82 | 15177 | 44,32 | 16 | 15 | 0,968 | 0,912 | 0,983 | 1,103 | 0,937 | 0,939 | 0,998 | 0,978 | 1,03 | 0,03 | 5,3E-01 |
| Q9DC70 | NADH dehydrogenase [ubiquinone] iron-sulfur protein 7, mitoch | 22 | 5 | 8 | 5 | 224 | 24,7 | 9,92 | 3571 | 4,68 | 4 | 1 | 1,095 | 0,941 | 0,986 | 1,001 | 1,024 | 0,99 | 0,968 | 0,944 | 1,02 | 0,02 | 5,3E-01 |
| Q8BZ98 | Dynamin-3 OS=Mus musculus OX=10090 GN=Dnm3 PE=1 SV=1 | 20 | 19 | 37 | 7 | 863 | 97,1 | 8,35 | 15512 | 33,83 | 11 | 9 | 0,962 | 0,926 | 0,947 | 0,922 | 1,054 | 0,939 | 0,968 | 0,89 | 0,98 | -0,02 | 5,3E-01 |
| Q9Z0R4 | Intersectin-1 OS=Mus musculus OX=10090 GN=Itsn1 PE=1 SV=2 | 13 | 17 | 20 | 15 | 1714 | 194,2 | 7,91 | 7554 | 9,37 | 11 | 6 | 0,943 | 1,065 | 0,918 | 0,921 | 0,862 | 0,89 | 0,977 | 0,993 | 1,03 | 0,03 | 5,3E-01 |
| Q99PU8 | ATP-dependent RNA helicase DHX30 OS=Mus musculus OX=100 | 1 | 1 | 1 | 1 | 1217 | 136,6 | 8,75 | 610 | | 1 | | 1,152 | 1,245 | 1,003 | 0,99 | 1,221 | 1,007 | 1,241 | 1,135 | 0,95 | -0,05 | 5,3E-01 |
| Q8BWG8 | Beta-arrestin-1 OS=Mus musculus OX=10090 GN=Arrb1 PE=1 SV | 13 | 5 | 9 | 5 | 418 | 46,9 | 6,28 | 1173 | 19,22 | 2 | 3 | 1,024 | 0,912 | 1,145 | 0,956 | 0,774 | 1,068 | 1,001 | 0,98 | 1,06 | 0,06 | 5,3E-01 |
| Q8BGX2 | Mitochondrial import inner membrane translocase subunit Tim2 | 8 | 2 | 2 | 2 | 266 | 29,4 | 6,73 | 764 | 2,27 | 1 | 1 | 0,924 | 0,614 | 0,961 | 1,277 | 1,748 | 1,186 | 0,798 | 0,749 | 0,84 | -0,16 | 5,3E-01 |
| Q64521 | Glycerol-3-phosphate dehydrogenase, mitochondrial OS=Mus m | 23 | 16 | 24 | 16 | 727 | 80,9 | 6,61 | 10985 | 4,47 | 13 | 4 | 0,98 | 1,03 | 0,966 | 1,008 | 0,936 | 1,004 | 1,056 | 1,08 | 0,98 | -0,02 | 5,3E-01 |
| Q8BJY1 | 26S proteasome non-ATPase regulatory subunit 5 OS=Mus musc | 22 | 8 | 11 | 8 | 504 | 55,9 | 5,21 | 5268 | 2,91 | 7 | 1 | 0,998 | 1,021 | 0,932 | 0,989 | 1,064 | 0,962 | 0,955 | 1,049 | 0,98 | -0,02 | 5,3E-01 |
| Q68FF6 | ARF GTPase-activating protein GIT1 OS=Mus musculus OX=1009 | 21 | 15 | 18 | 14 | 770 | 85,2 | 6,93 | 5670 | 18,65 | 8 | 7 | 0,988 | 0,879 | 1,114 | 0,888 | 0,885 | 0,953 | 0,952 | 0,929 | 1,04 | 0,04 | 5,4E-01 |
| P97393 | Rho GTPase-activating protein 5 OS=Mus musculus OX=10090 G | 3 | 4 | 4 | 4 | 1501 | 172 | 6,34 | 903 | 2,25 | 3 | 1 | 0,966 | 1,11 | 1,087 | 0,952 | 0,979 | 0,976 | 1,004 | 1,042 | 1,03 | 0,03 | 5,4E-01 |
| Q8BU88 | 39S ribosomal protein L22, mitochondrial OS=Mus musculus OX | 12 | 2 | 2 | 2 | 206 | 23,8 | 9,86 | 1320 | | 2 | | 0,904 | 0,855 | 0,615 | 0,785 | 0,986 | 0,834 | 0,717 | 0,842 | 0,93 | -0,07 | 5,4E-01 |
| Q9QZR0 | E3 ubiquitin-protein ligase RNF25 OS=Mus musculus OX=10090 | 2 | 1 | 1 | 1 | 456 | 51,2 | 6,29 | 430 | | 1 | | 0,817 | 0,912 | 0,862 | 0,961 | 0,976 | 0,668 | 0,813 | 0,903 | 1,06 | 0,06 | 5,4E-01 |
| Q8CDC7 | Zinc finger and BTB domain-containing protein 9 OS=Mus muscu | 2 | 1 | 2 | 1 | 459 | 49,5 | 7,03 | 626 | | 1 | | 1,095 | 0,923 | 1,012 | 1,137 | 0,993 | 1,086 | 1,186 | 1,064 | 0,96 | -0,04 | 5,4E-01 |
| Q8C8R3-2 | Isoform 2 of Ankyrin-2 OS=Mus musculus OX=10090 GN=Ank2 | 38 | 120 | 244 | 3 | 3926 | 428,5 | 5,16 | 91623 | 255,45 | 81 | 52 | 0,943 | 1,096 | 0,969 | 1,049 | 1,247 | 0,881 | 1,005 | 1,155 | 0,95 | -0,05 | 5,4E-01 |
| Q91Z53 | Glyoxylate reductase/hydroxypyruvate reductase OS=Mus musc | 11 | 4 | 4 | 3 | 328 | 35,3 | 7,65 | 1835 | 2,17 | 3 | 1 | 0,877 | 1,082 | 1,117 | 0,999 | 0,876 | 1,016 | 1,011 | 1,007 | 1,04 | 0,04 | 5,4E-01 |
| Q9D6M3 | Mitochondrial glutamate carrier 1 OS=Mus musculus OX=10090 | 27 | 7 | 13 | 5 | 323 | 34,6 | 9,09 | 5221 | 14,76 | 5 | 4 | 0,995 | 1,032 | 1,041 | 1,058 | 0,902 | 1,097 | 0,992 | 1,024 | 1,03 | 0,03 | 5,4E-01 |
| Q9Z2I8 | Succinate--CoA ligase [GDP-forming] subunit beta, mitochondria | 11 | 5 | 5 | 5 | 433 | 46,8 | 7,02 | 1842 | 4,18 | 3 | 2 | 0,971 | 1,037 | 1,058 | 1,209 | 1,268 | 1,04 | 1,001 | 1,173 | 0,95 | -0,05 | 5,4E-01 |
| Q921F4 | Heterogeneous nuclear ribonucleoprotein L-like OS=Mus muscu | 4 | 2 | 2 | 2 | 591 | 64,1 | 5,85 | 1053 | | 2 | | 0,799 | 1,237 | 1,022 | 1,152 | 1,299 | 1,008 | 1,061 | 1,14 | 0,93 | -0,07 | 5,4E-01 |
| Q3THE2 | Myosin regulatory light chain 12B OS=Mus musculus OX=10090 | 51 | 7 | 12 | 7 | 172 | 19,8 | 4,84 | 6959 | 0 | 7 | 1 | 1,014 | 1,01 | 1,037 | 1,12 | 1,123 | 0,932 | 1,043 | 0,95 | 1,03 | 0,03 | 5,4E-01 |
| P17710 | Hexokinase-1 OS=Mus musculus OX=10090 GN=Hk1 PE=1 SV=3 | 31 | 32 | 58 | 32 | 974 | 108,2 | 6,8 | 24821 | 61,12 | 24 | 12 | 1,012 | 0,966 | 0,959 | 0,965 | 0,989 | 0,981 | 0,951 | 1,037 | 0,99 | -0,01 | 5,4E-01 |
| Q9D967 | Magnesium-dependent phosphatase 1 OS=Mus musculus OX=1( | 4 | 1 | 2 | 1 | 164 | 18,6 | 6,8 | 966 | | 1 | | 1,094 | 0,997 | 1,021 | 0,942 | 0,916 | 1,104 | 0,901 | 0,988 | 1,04 | 0,04 | 5,4E-01 |
| Q8CHP8 | Glycerol-3-phosphate phosphatase OS=Mus musculus OX=1009( | 13 | 4 | 5 | 4 | 321 | 34,5 | 5,35 | 2702 | 2,09 | 3 | 1 | 0,978 | 0,929 | 0,99 | 0,942 | 0,773 | 1,015 | 0,928 | 0,98 | 1,04 | 0,04 | 5,4E-01 |
| Q8BIF9 | Zinc finger protein 787 OS=Mus musculus OX=10090 GN=Znf787 | 3 | 1 | 1 | 1 | 381 | 40,5 | 8,24 | 148 | | 1 | | 1,245 | 1,275 | 1,199 | 1,353 | 1,176 | 1,252 | 1,164 | 1,344 | 1,03 | 0,03 | 5,4E-01 |
| Q80TM9 | Nischarin OS=Mus musculus OX=10090 GN=Nisch PE=1 SV=2 | 1 | 1 | 1 | 1 | 1593 | 174,9 | 5,16 | 549 | | 1 | | 0,896 | 0,839 | 1,014 | 1,018 | 1,135 | 0,864 | 1,141 | 0,862 | 0,94 | -0,06 | 5,4E-01 |
| Q80ZK0 | 28S ribosomal protein S10, mitochondrial OS=Mus musculus OX | 11 | 1 | 1 | 1 | 160 | 18,7 | 8,84 | 369 | | 1 | | 1,098 | 0,89 | 0,931 | 1,126 | 0,888 | 1,133 | 1,248 | 1,026 | 0,94 | -0,06 | 5,4E-01 |
| Q8BFR5 | Elongation factor Tu, mitochondrial OS=Mus musculus OX=1009 | 41 | 15 | 40 | 15 | 452 | 49,5 | 7,56 | 18042 | 31,22 | 11 | 5 | 0,989 | 0,973 | 0,994 | 1,013 | 0,827 | 0,993 | 1,018 | 1,012 | 1,03 | 0,03 | 5,4E-01 |
| Q9D7I5 | Phospholysine phosphohistidine inorganic pyrophosphate phos | 13 | 3 | 3 | 3 | 270 | 29,1 | 5,1 | 727 | 6,12 | 1 | 2 | 0,897 | 1,086 | 0,969 | 0,948 | 0,719 | 1,214 | 0,847 | 0,824 | 1,08 | 0,08 | 5,4E-01 |
| Q8VEB2 | Protein salvador homolog 1 OS=Mus musculus OX=10090 GN=Sa | 2 | 1 | 3 | 1 | 386 | 44,8 | 9,06 | 1352 | | 1 | | 0,874 | 1,285 | 0,731 | 1,245 | 0,939 | 0,781 | 1,109 | 0,914 | 1,10 | 0,10 | 5,4E-01 |

| | | | | | | | | | | | | | | | | | | | | | | | |
|---|---|---|---|---|---|---|---|---|---|---|---|---|---|---|---|---|---|---|---|---|---|---|---|
| Q91VR5 | ATP-dependent RNA helicase DDX1 OS=Mus musculus OX=1009 | 6 | 5 | 7 | 5 | 740 | 82,4 | 7,21 | 3280 | 4,95 | 4 | 2 | 0,942 | 0,998 | 0,942 | 0,905 | 0,854 | 0,977 | 0,955 | 0,916 | 1,02 | 0,02 | 5,4E-01 |
| Q9CX86 | Heterogeneous nuclear ribonucleoprotein A0 OS=Mus musculus | 17 | 4 | 7 | 4 | 305 | 30,5 | 9,31 | 2875 | 3,31 | 4 | 1 | 0,916 | 0,881 | 1,362 | 1,321 | 1,135 | 1,274 | 1,344 | 1,09 | 0,93 | -0,07 | 5,4E-01 |
| Q8VCY8 | Phospholipid phosphatase-related protein type 2 OS=Mus musc | 6 | 1 | 1 | 1 | 343 | 36,9 | 9,5 | | 3,65 | | 1 | 1,78 | 1,452 | 1,364 | 1,476 | 2,305 | 1,589 | 1,155 | 1,674 | 0,90 | -0,10 | 5,5E-01 |
| Q8JZR0 | Long-chain-fatty-acid--CoA ligase 5 OS=Mus musculus OX=10090 | 6 | 4 | 4 | 4 | 683 | 76,2 | 7,09 | 1886 | | 4 | | 1,038 | 1,062 | 1,076 | 1,077 | 1,49 | 0,975 | 1,019 | 1,074 | 0,93 | -0,07 | 5,5E-01 |
| Q9R0P9 | Ubiquitin carboxyl-terminal hydrolase isozyme L1 OS=Mus musc | 56 | 14 | 35 | 14 | 223 | 24,8 | 5,24 | 7581 | 51,51 | 6 | 11 | 0,965 | 0,977 | 1,043 | 0,922 | 0,757 | 1,076 | 0,998 | 0,887 | 1,05 | 0,05 | 5,5E-01 |
| Q9DBJ1 | Phosphoglycerate mutase 1 OS=Mus musculus OX=10090 GN=P | 57 | 14 | 37 | 9 | 254 | 28,8 | 7,18 | 17526 | 15,08 | 11 | 4 | 0,909 | 1,027 | 0,846 | 1,185 | 1,153 | 1,103 | 0,973 | 0,964 | 0,95 | -0,05 | 5,5E-01 |
| Q9WUM5 | Succinate--CoA ligase [ADP/GDP-forming] subunit alpha, mitoch | 18 | 5 | 10 | 5 | 346 | 36,1 | 9,39 | 4224 | 13,99 | 3 | 3 | 1,095 | 1,114 | 1,105 | 1,081 | 1,153 | 1,147 | 1,053 | 1,104 | 0,99 | -0,01 | 5,5E-01 |
| P52792 | Hexokinase-4 OS=Mus musculus OX=10090 GN=Gck PE=1 SV=1 | 5 | 2 | 4 | 2 | 465 | 52,1 | 5,29 | | 10,72 | | 2 | 1,445 | 1,24 | 1,184 | 1,452 | 1,595 | 1,137 | 1,474 | 1,42 | 0,95 | -0,05 | 5,5E-01 |
| P97300 | Neuroplastin OS=Mus musculus OX=10090 GN=Nptn PE=1 SV=3 | 30 | 11 | 23 | 3 | 397 | 44,3 | 7,74 | 5456 | 32,41 | 6 | 5 | 1,011 | 0,966 | 0,962 | 0,97 | 0,947 | 1,136 | 0,992 | 0,951 | 0,97 | -0,03 | 5,5E-01 |
| Q91VK1 | Basic leucine zipper and W2 domain-containing protein 2 OS=M | 5 | 3 | 3 | 3 | 419 | 48 | 6,68 | 824 | 2,81 | 2 | 1 | 0,984 | 0,98 | 0,932 | 1,016 | 0,707 | 1,062 | 0,983 | 0,959 | 1,05 | 0,05 | 5,5E-01 |
| Q3TIR3 | Synembryn-A OS=Mus musculus OX=10090 GN=Ric8a PE=1 SV=: | 4 | 2 | 2 | 2 | 530 | 59,8 | 5,68 | 468 | 1,97 | 1 | 1 | 0,965 | 1,137 | 0,979 | 0,965 | 0,778 | 1,128 | 1,059 | 0,841 | 1,06 | 0,06 | 5,5E-01 |
| O35704 | Serine palmitoyltransferase 1 OS=Mus musculus OX=10090 GN= | 3 | 2 | 2 | 2 | 473 | 52,5 | 6,4 | 368 | 2,61 | 1 | 1 | 1,031 | 0,959 | 0,958 | 0,983 | 0,79 | 1,017 | 1,022 | 0,957 | 1,04 | 0,04 | 5,5E-01 |
| O35658 | Complement component 1 Q subcomponent-binding protein, m | 5 | 1 | 4 | 1 | 278 | 31 | 4,92 | 471 | 4,72 | 1 | 1 | 0,749 | 1,055 | 0,826 | 1,084 | 1,207 | 0,776 | 1,051 | 0,991 | 0,92 | -0,08 | 5,5E-01 |
| Q9D517 | 1-acyl-sn-glycerol-3-phosphate acyltransferase gamma OS=Mus | 5 | 2 | 2 | 2 | 376 | 43,3 | 8,51 | 885 | | 2 | | 0,969 | 1,104 | 0,827 | 0,823 | 0,8 | 0,816 | 0,848 | 1,038 | 1,06 | 0,06 | 5,5E-01 |
| P53810 | Phosphatidylinositol transfer protein alpha isoform OS=Mus mu | 28 | 7 | 11 | 6 | 271 | 31,9 | 6,37 | 6694 | 2,57 | 5 | 2 | 1,003 | 0,992 | 1,09 | 1,077 | 1,064 | 0,989 | 1 | 1,032 | 1,02 | 0,02 | 5,5E-01 |
| Q91YE6 | Importin-9 OS=Mus musculus OX=10090 GN=Ipo9 PE=1 SV=3 | 4 | 4 | 5 | 4 | 1041 | 116 | 4,81 | 1523 | 2,23 | 3 | 1 | 0,976 | 1,027 | 0,985 | 1,087 | 0,969 | 1,026 | 1,096 | 1,082 | 0,98 | -0,02 | 5,5E-01 |
| P63166 | Small ubiquitin-related modifier 1 OS=Mus musculus OX=10090 | 12 | 1 | 1 | 1 | 101 | 11,6 | 5,52 | 546 | | 1 | | 0,933 | 1,148 | 0,926 | 1,007 | 1,329 | 0,864 | 0,84 | 0,551 | 1,12 | 0,12 | 5,5E-01 |
| Q61024 | Asparagine synthetase [glutamine-hydrolyzing] OS=Mus muscul | 3 | 2 | 4 | 2 | 561 | 64,2 | 6,58 | 2791 | | 2 | | 0,981 | 0,967 | 1,014 | 1,057 | 1,06 | 0,999 | 0,891 | 0,966 | 1,03 | 0,03 | 5,5E-01 |
| Q9WTR5 | Cadherin-13 OS=Mus musculus OX=10090 GN=Cdh13 PE=1 SV=2 | 9 | 5 | 11 | 5 | 714 | 78,1 | 5,12 | 4198 | 11,78 | 4 | 3 | 0,977 | 0,83 | 0,815 | 1,212 | 0,6 | 1,367 | 0,715 | 0,637 | 1,16 | 0,16 | 5,5E-01 |
| Q01279 | Epidermal growth factor receptor OS=Mus musculus OX=10090 | 3 | 2 | 2 | 2 | 1210 | 134,8 | 6,86 | 782 | | 2 | | 0,716 | 0,781 | 0,929 | 0,991 | 1,952 | 0,822 | 0,697 | 0,729 | 0,81 | -0,19 | 5,5E-01 |
| Q61239 | Protein farnesyltransferase/geranylgeranyltransferase type-1 su | 3 | 1 | 1 | 1 | 377 | 44 | 4,93 | 506 | | 1 | | 1,171 | 0,735 | 1,139 | 0,78 | 0,869 | 0,951 | 0,835 | 0,871 | 1,08 | 0,08 | 5,5E-01 |
| Q921S7 | 39S ribosomal protein L37, mitochondrial OS=Mus musculus OX | 4 | 1 | 1 | 1 | 423 | 48,3 | 8,84 | 469 | | 1 | | 0,995 | 0,799 | 0,681 | 0,825 | 0,881 | 0,803 | 0,983 | 0,826 | 0,94 | -0,06 | 5,5E-01 |
| P70392 | Ras-specific guanine nucleotide-releasing factor 2 OS=Mus musc | 1 | 1 | 3 | 1 | 1189 | 135,6 | 7,85 | 1671 | | 1 | | 1,158 | 1,151 | 1,15 | 0,909 | 1,174 | 1,037 | 1,038 | 1,372 | 0,95 | -0,05 | 5,5E-01 |
| Q08460 | Calcium-activated potassium channel subunit alpha-1 OS=Mus n | 3 | 3 | 3 | 3 | 1209 | 134,3 | 7,15 | 1028 | 2,58 | 2 | 1 | 1,128 | 1,018 | 1,082 | 1,092 | 1,153 | 1,051 | 1,045 | 0,952 | 1,03 | 0,03 | 5,5E-01 |
| Q3TYD6 | Serine/threonine-protein kinase LMTK2 OS=Mus musculus OX=1 | 0 | 1 | 1 | 1 | 1471 | 160,4 | 4,51 | | 2,18 | | 1 | 1,021 | 1,113 | 0,91 | 0,938 | 1,005 | 0,99 | 1,121 | 1,005 | 0,97 | -0,03 | 5,5E-01 |
| P62702 | 40S ribosomal protein S4, X isoform OS=Mus musculus OX=1009 | 31 | 11 | 23 | 11 | 263 | 29,6 | 10,15 | 9700 | 13,01 | 8 | 3 | 1,09 | 0,91 | 1,015 | 1,075 | 0,981 | 1,034 | 0,959 | 1,005 | 1,03 | 0,03 | 5,5E-01 |
| P31938 | Dual specificity mitogen-activated protein kinase kinase 1 OS=M | 24 | 7 | 9 | 4 | 393 | 43,4 | 6,7 | 2821 | 12,16 | 4 | 4 | 1,128 | 1,215 | 0,986 | 1,142 | 0,929 | 1,135 | 1,16 | 1,069 | 1,04 | 0,04 | 5,5E-01 |
| Q9JMA1 | Ubiquitin carboxyl-terminal hydrolase 14 OS=Mus musculus OX= | 9 | 4 | 4 | 3 | 493 | 56 | 5,24 | 1744 | 2,89 | 3 | 1 | 0,935 | 0,881 | 0,954 | 0,824 | 0,872 | 1,065 | 0,839 | 0,966 | 0,96 | -0,04 | 5,5E-01 |
| Q8VIM9 | Immunity-related GTPase family Q protein OS=Mus musculus O) | 9 | 5 | 7 | 5 | 583 | 59,3 | 4,83 | 2658 | 6,04 | 4 | 1 | 0,893 | 0,932 | 0,976 | 0,996 | 0,685 | 0,95 | 0,87 | 1,077 | 1,06 | 0,06 | 5,5E-01 |
| Q8R5M8 | Cell adhesion molecule 1 OS=Mus musculus OX=10090 GN=Cadr | 12 | 5 | 6 | 5 | 456 | 49,8 | 5,03 | 1994 | 5,67 | 4 | 2 | 1,061 | 0,825 | 1,02 | 0,84 | 1,016 | 1,116 | 0,914 | 0,899 | 0,95 | -0,05 | 5,5E-01 |
| P61021 | Ras-related protein Rab-5B OS=Mus musculus OX=10090 GN=Ra | 22 | 4 | 9 | 2 | 215 | 23,7 | 8,13 | 3117 | 7,84 | 3 | 2 | 0,953 | 0,92 | 0,926 | 0,874 | 1,206 | 0,897 | 0,881 | 0,892 | 0,95 | -0,05 | 5,5E-01 |
| P61087 | Ubiquitin-conjugating enzyme E2 K OS=Mus musculus OX=1009( | 19 | 3 | 5 | 3 | 200 | 22,4 | 5,44 | 1863 | 5,72 | 2 | 1 | 0,967 | 1,022 | 1,087 | 0,99 | 1,064 | 1,148 | 0,955 | 1,02 | 0,97 | -0,03 | 5,5E-01 |
| P97427 | Dihydropyrimidinase-related protein 1 OS=Mus musculus OX=1( | 58 | 32 | 167 | 24 | 572 | 62,1 | 7,12 | 67445 | 138,09 | 19 | 21 | 1,035 | 0,99 | 1,044 | 1,022 | 0,97 | 1,024 | 0,989 | 1,053 | 1,01 | 0,01 | 5,5E-01 |
| Q9CQZ5 | NADH dehydrogenase [ubiquinone] 1 alpha subcomplex subunit | 26 | 3 | 6 | 3 | 131 | 15,3 | 10,11 | 3533 | | 3 | | 1,034 | 1,082 | 1,072 | 1,028 | 1,077 | 0,996 | 1,086 | 1,138 | 0,98 | -0,02 | 5,5E-01 |
| B2RSH2 | Guanine nucleotide-binding protein G(i) subunit alpha-1 OS=Mu | 25 | 10 | 24 | 3 | 354 | 40,3 | 5,97 | 8201 | 19,12 | 6 | 5 | 0,965 | 1,048 | 1,003 | 0,864 | 1,04 | 0,965 | 0,785 | 0,924 | 1,04 | 0,04 | 5,5E-01 |
| Q8CJG0 | Protein argonaute-2 OS=Mus musculus OX=10090 GN=Ago2 PE= | 4 | 3 | 3 | 3 | 860 | 97,2 | 9,19 | 661 | | 3 | | 0,985 | 0,711 | 1,48 | 0,881 | 2,205 | 1,018 | 1,024 | 0,728 | 0,82 | -0,18 | 5,5E-01 |
| Q69ZK9 | Neuroligin-2 OS=Mus musculus OX=10090 GN=Nlgn2 PE=1 SV=2 | 4 | 3 | 3 | 3 | 836 | 90,9 | 6,18 | 1074 | 2,23 | 2 | 1 | 1,09 | 1,054 | 1,075 | 1,161 | 0,799 | 1,183 | 1,037 | 1,139 | 1,05 | 0,05 | 5,6E-01 |
| P61514 | 60S ribosomal protein L37a OS=Mus musculus OX=10090 GN=R | 64 | 6 | 8 | 6 | 92 | 10,3 | 10,43 | 2075 | 8,25 | 4 | 4 | 1,139 | 1,054 | 1,178 | 1,079 | 1,372 | 1,117 | 1,112 | 1,042 | 0,96 | -0,04 | 5,6E-01 |
| Q9DAU1 | Protein canopy homolog 3 OS=Mus musculus OX=10090 GN=Cn | 10 | 2 | 3 | 2 | 276 | 30,5 | 5,62 | 1218 | 2,65 | 1 | 1 | 1,085 | 1,087 | 0,89 | 1,149 | 1,086 | 1,118 | 0,986 | 0,791 | 1,06 | 0,06 | 5,6E-01 |
| Q9CU62 | Structural maintenance of chromosomes protein 1A OS=Mus m | 1 | 1 | 1 | 1 | 1233 | 143,1 | 7,64 | 702 | | 1 | | 0,904 | 0,888 | 0,759 | 0,982 | 1,145 | 0,554 | 1,185 | 1,027 | 0,90 | -0,10 | 5,6E-01 |
| P53026 | 60S ribosomal protein L10a OS=Mus musculus OX=10090 GN=R | 28 | 7 | 11 | 7 | 217 | 24,9 | 9,98 | 5750 | 7,63 | 5 | 2 | 0,996 | 0,9 | 1,006 | 1,052 | 0,946 | 0,974 | 0,933 | 1,011 | 1,02 | 0,02 | 5,6E-01 |
| Q8CIN6 | CUGBP Elav-like family member 3 OS=Mus musculus OX=10090 | 3 | 1 | 1 | 1 | 465 | 50,5 | 8,68 | 583 | | 1 | | 1,343 | 1,603 | 1,42 | 0,956 | 1,189 | 1,335 | 1,093 | 1,337 | 1,07 | 0,07 | 5,6E-01 |
| Q5U4F6 | WD repeat-containing protein 34 OS=Mus musculus OX=10090 ( | 2 | 1 | 1 | 1 | 535 | 57,9 | 6,52 | 310 | | 1 | | 1,061 | 1,015 | 1,209 | 1,207 | 0,814 | 1,306 | 0,857 | 1,19 | 1,08 | 0,08 | 5,6E-01 |
| O89051 | Integral membrane protein 2B OS=Mus musculus OX=10090 GN | 4 | 1 | 1 | 1 | 266 | 30,2 | 5,3 | 491 | | 1 | | 1,079 | 1,373 | 0,925 | 1,204 | 0,954 | 1,056 | 1,221 | 1,079 | 1,06 | 0,06 | 5,6E-01 |
| P08207 | Protein S100-A10 OS=Mus musculus OX=10090 GN=S100a10 PE | 10 | 1 | 1 | 1 | 97 | 11,2 | 6,77 | 428 | | 1 | | 0,931 | 0,653 | 1,021 | 1,011 | 1,126 | 0,996 | 0,708 | 1,104 | 0,92 | -0,08 | 5,6E-01 |
| Q05816 | Fatty acid-binding protein 5 OS=Mus musculus OX=10090 GN=F | 71 | 9 | 36 | 9 | 135 | 15,1 | 6,54 | 8518 | 43,04 | 5 | 5 | 1,022 | 1,017 | 1,018 | 1,022 | 0,989 | 1,07 | 0,923 | 1,021 | 1,02 | 0,02 | 5,6E-01 |
| Q60823 | RAC-beta serine/threonine-protein kinase OS=Mus musculus OX | 7 | 3 | 3 | 2 | 481 | 55,7 | 6,37 | 767 | 2,59 | 2 | 1 | 1,246 | 1,072 | 1,367 | 1,136 | 2,045 | 1,082 | 1,093 | 1,194 | 0,89 | -0,11 | 5,6E-01 |
| Q8CJH3 | Plexin-B1 OS=Mus musculus OX=10090 GN=Plxnb1 PE=1 SV=2 | 2 | 4 | 5 | 4 | 2119 | 231,2 | 5,55 | 2594 | 2,27 | 3 | 1 | 1,087 | 1,014 | 1,194 | 1,12 | 1,026 | 1,104 | 0,997 | 1,159 | 1,03 | 0,03 | 5,6E-01 |
| Q8C0D5 | Elongation factor-like GTPase 1 OS=Mus musculus OX=10090 GN | 3 | 3 | 3 | 3 | 1127 | 125,7 | 6,16 | 1661 | | 3 | | 0,922 | 0,993 | 0,879 | 0,996 | 0,911 | 0,946 | 1,013 | 1,014 | 0,98 | -0,02 | 5,6E-01 |
| Q61990 | Poly(rC)-binding protein 2 OS=Mus musculus OX=10090 GN=Pcb | 28 | 7 | 12 | 4 | 362 | 38,2 | 6,79 | 4260 | 14,34 | 4 | 4 | 0,931 | 1,081 | 1,102 | 1,141 | 1,234 | 1,157 | 1,021 | 1,016 | 0,96 | -0,04 | 5,6E-01 |
| P70698 | CTP synthase 1 OS=Mus musculus OX=10090 GN=Ctps1 PE=1 SV | 19 | 9 | 12 | 8 | 591 | 66,6 | 6,58 | 7042 | | 9 | | 0,926 | 0,94 | 1,077 | 0,968 | 1,022 | 1,053 | 0,96 | 0,975 | 0,98 | -0,02 | 5,6E-01 |
| Q8JZP2 | Synapsin-3 OS=Mus musculus OX=10090 GN=Syn3 PE=1 SV=2 | 19 | 8 | 13 | 7 | 579 | 63,3 | 9,38 | 5763 | 10,51 | 6 | 3 | 0,967 | 1 | 1,054 | 0,987 | 0,935 | 0,968 | 1,024 | 1,013 | 1,02 | 0,02 | 5,6E-01 |
| Q63810 | Calcineurin subunit B type 1 OS=Mus musculus OX=10090 GN=P | 35 | 7 | 9 | 7 | 170 | 19,3 | 4,81 | 2233 | 14,36 | 3 | 5 | 1,112 | 0,93 | 0,932 | 0,971 | 0,925 | 1,138 | 1,017 | 1,015 | 0,96 | -0,04 | 5,6E-01 |
| Q3TWI9 | CSC1-like protein 2 OS=Mus musculus OX=10090 GN=Tmem63b | 2 | 1 | 1 | 1 | 832 | 94,7 | 7,49 | | 0 | | 1 | 1,069 | 0,783 | 0,911 | 0,986 | 0,796 | 0,818 | 0,989 | 0,956 | 1,05 | 0,05 | 5,6E-01 |
| Q148V7 | RAB11-binding protein RELCH OS=Mus musculus OX=10090 GN= | 8 | 9 | 13 | 1 | 1216 | 134,5 | 5,34 | 6522 | 6,32 | 8 | 1 | 0,911 | 0,961 | 1,001 | 1,194 | 0,993 | 1,011 | 1,021 | 1,257 | 0,95 | -0,05 | 5,6E-01 |
| Q8K3J1 | NADH dehydrogenase [ubiquinone] iron-sulfur protein 8, mitoch | 25 | 5 | 10 | 5 | 212 | 24 | 6,21 | 6471 | | 5 | | 0,985 | 0,894 | 1,021 | 1,077 | 1,084 | 0,897 | 1,028 | 1,12 | 0,96 | -0,04 | 5,6E-01 |
| P35821 | Tyrosine-protein phosphatase non-receptor type 1 OS=Mus mus | 4 | 2 | 3 | 2 | 432 | 49,6 | 6,16 | 1131 | | 2 | | 1,055 | 0,995 | 1,231 | 0,981 | 1,023 | 0,976 | 1,05 | 1,065 | 1,04 | 0,04 | 5,6E-01 |
| P62073 | Mitochondrial import inner membrane translocase subunit Tim1 | 30 | 3 | 3 | 3 | 90 | 10,3 | 6,29 | 1033 | 2 | 2 | 1 | 0,97 | 0,935 | 1,273 | 0,909 | 0,922 | 1,016 | 1,111 | 0,764 | 1,07 | 0,07 | 5,6E-01 |
| Q91WS0 | CDGSH iron-sulfur domain-containing protein 1 OS=Mus muscul | 36 | 4 | 10 | 4 | 108 | 12,1 | 9,06 | 5690 | 3,05 | 3 | 1 | 0,921 | 1,074 | 1,009 | 0,964 | 0,94 | 1,082 | 1,006 | 1,049 | 0,97 | -0,03 | 5,7E-01 |
| A8Y5H7 | SEC14-like protein 1 OS=Mus musculus OX=10090 GN=Sec14l1 F | 1 | 1 | 1 | 1 | 715 | 81,2 | 6,34 | 442 | | 1 | | 0,942 | 0,781 | 1,001 | 0,938 | 0,627 | 0,996 | 0,911 | 0,902 | 1,07 | 0,07 | 5,7E-01 |
| Q71M36-2 | Isoform 2 of Chondroitin sulfate proteoglycan 5 OS=Mus muscul | 13 | 4 | 6 | 1 | 539 | 57,3 | 4,44 | 1793 | 12,35 | 3 | 1 | 0,944 | 0,984 | 0,975 | 0,859 | 1,187 | 0,899 | 0,979 | 0,881 | 0,95 | -0,05 | 5,7E-01 |
| Q5SQX6 | Cytoplasmic FMR1-interacting protein 2 OS=Mus musculus OX=: | 16 | 20 | 32 | 6 | 1253 | 145,6 | 7,05 | 13104 | 12,44 | 14 | 6 | 1,053 | 0,992 | 1,203 | 0,998 | 1,08 | 1,141 | 1,141 | 1,022 | 0,97 | -0,03 | 5,7E-01 |
| Q9JIA7 | Sphingosine kinase 2 OS=Mus musculus OX=10090 GN=Sphk2 PI | 3 | 1 | 1 | 1 | 617 | 65,6 | 6,57 | 347 | | 1 | | 1,027 | 1,16 | 0,78 | 1,242 | 1,797 | 0,676 | 1,197 | 1,148 | 0,87 | -0,13 | 5,7E-01 |
| Q3TVI8 | Pre-B-cell leukemia transcription factor-interacting protein 1 OS | 3 | 2 | 2 | 2 | 727 | 81,1 | 5,36 | 991 | | 2 | | 0,949 | 1,057 | 0,75 | 1,129 | 1,148 | 0,779 | 1,196 | 1,064 | 0,93 | -0,07 | 5,7E-01 |
| Q9CPR5 | 39S ribosomal protein L15, mitochondrial OS=Mus musculus OX | 11 | 4 | 4 | 4 | 295 | 33,5 | 10,07 | 900 | 3,74 | 2 | 2 | 1,056 | 0,853 | 1,007 | 0,951 | 0,819 | 0,961 | 0,995 | 0,951 | 1,04 | 0,04 | 5,7E-01 |

| | | | | | | | | | | | | | | | | | | | | | | | |
|---|---|---|---|---|---|---|---|---|---|---|---|---|---|---|---|---|---|---|---|---|---|---|---|
| Q9DCD6 | Gamma-aminobutyric acid receptor-associated protein OS=Mus | 20 | 3 | 3 | 1 | 117 | 13,9 | 8,79 | 751 | 8,83 | 1 | 2 | 0,892 | 0,884 | 0,955 | 0,919 | 0,777 | 0,925 | 0,796 | 1,012 | 1,04 | 0,04 | 5,7E-01 |
| Q922Q4 | Pyrroline-5-carboxylate reductase 2 OS=Mus musculus OX=1009 | 13 | 4 | 4 | 3 | 320 | 33,6 | 7,77 | 2276 | | 4 | | 1,025 | 1,158 | 0,944 | 0,99 | 0,771 | 1,002 | 0,898 | 1,2 | 1,06 | 0,06 | 5,7E-01 |
| P62874 | Guanine nucleotide-binding protein G(I)/G(S)/G(T) subunit beta- | 44 | 11 | 43 | 7 | 340 | 37,4 | 6 | 13903 | 65,53 | 5 | 7 | 0,951 | 0,996 | 1,127 | 0,979 | 0,978 | 1,103 | 0,975 | 1,135 | 0,97 | -0,03 | 5,7E-01 |
| P70202 | Latexin OS=Mus musculus OX=10090 GN=Lxn PE=1 SV=2 | 8 | 1 | 2 | 1 | 222 | 25,5 | 5,74 | 831 | 4,54 | 1 | 1 | 0,937 | 1,013 | 0,927 | 0,927 | 0,816 | 1,132 | 0,959 | 1,072 | 0,96 | -0,04 | 5,7E-01 |
| Q6P9K8 | Caskin-1 OS=Mus musculus OX=10090 GN=Caskin1 PE=1 SV=2 | 17 | 16 | 20 | 16 | 1431 | 150,4 | 9,17 | 10301 | 7,02 | 13 | 3 | 1,009 | 0,981 | 1,126 | 0,995 | 1,134 | 1,001 | 1,026 | 1,056 | 0,97 | -0,03 | 5,7E-01 |
| P59999 | Actin-related protein 2/3 complex subunit 4 OS=Mus musculus ( | 23 | 4 | 8 | 4 | 168 | 19,7 | 8,43 | 3096 | 5,91 | 2 | 2 | 0,99 | 1,048 | 1,047 | 1,016 | 0,81 | 1,11 | 1 | 1,025 | 1,04 | 0,04 | 5,7E-01 |
| Q8C1B7 | Septin-11 OS=Mus musculus OX=10090 GN=Septin11 PE=1 SV=4 | 35 | 14 | 26 | 7 | 431 | 49,7 | 6,68 | 15981 | 3,31 | 14 | 1 | 0,904 | 0,88 | 1,042 | 0,917 | 1,021 | 0,987 | 0,886 | 0,96 | 0,97 | -0,03 | 5,7E-01 |
| P62754 | 40S ribosomal protein S6 OS=Mus musculus OX=10090 GN=Rps6 | 27 | 9 | 15 | 9 | 249 | 28,7 | 10,84 | 6563 | 2,12 | 6 | 3 | 1,003 | 1,002 | 1,007 | 1,116 | 0,951 | 0,975 | 1,003 | 1,097 | 1,03 | 0,03 | 5,7E-01 |
| Q9D0T1 | NHP2-like protein 1 OS=Mus musculus OX=10090 GN=Snu13 PE= | 9 | 1 | 1 | 1 | 128 | 14,2 | 8,46 | 767 | | 1 | | 0,891 | 0,945 | 1,009 | 1,141 | 1,227 | 0,85 | 0,964 | 1,2 | 0,94 | -0,06 | 5,7E-01 |
| Q9QXY6 | EH domain-containing protein 3 OS=Mus musculus OX=10090 G | 15 | 7 | 9 | 6 | 535 | 60,8 | 6,46 | 2038 | 16,46 | 2 | 5 | 0,897 | 1,021 | 1,123 | 0,923 | 0,985 | 1,021 | 1,083 | 1,008 | 0,97 | -0,03 | 5,7E-01 |
| Q9QXG2 | Rab proteins geranylgeranyltransferase component A 1 OS=Mus | 3 | 2 | 4 | 2 | 665 | 73,9 | 4,68 | 1794 | 2,61 | 1 | 1 | 0,955 | 0,887 | 1,218 | 0,969 | 1,432 | 0,997 | 0,955 | 0,97 | 0,93 | -0,07 | 5,7E-01 |
| Q9Z2I9 | Succinate--CoA ligase [ADP-forming] subunit beta, mitochondria | 22 | 11 | 19 | 11 | 463 | 50,1 | 7,01 | 8213 | 16,11 | 6 | 5 | 1,052 | 0,982 | 0,975 | 0,923 | 0,889 | 1,005 | 0,955 | 0,994 | 1,02 | 0,02 | 5,7E-01 |
| Q02053 | Ubiquitin-like modifier-activating enzyme 1 OS=Mus musculus O | 31 | 30 | 62 | 30 | 1058 | 117,7 | 5,66 | 28697 | 52,72 | 21 | 10 | 0,948 | 0,934 | 1,006 | 1,029 | 0,862 | 1,074 | 0,911 | 0,949 | 1,03 | 0,03 | 5,7E-01 |
| Q9QUH0 | Glutaredoxin-1 OS=Mus musculus OX=10090 GN=Glrx PE=1 SV=3 | 17 | 2 | 2 | 2 | 107 | 11,9 | 8,37 | 1240 | | 2 | | 0,675 | 0,928 | 0,689 | 0,812 | 0,979 | 0,756 | 0,845 | 0,721 | 0,94 | -0,06 | 5,7E-01 |
| Q80TR1 | Adhesion G protein-coupled receptor L1 OS=Mus musculus OX= | 3 | 3 | 3 | 3 | 1466 | 161,6 | 6,48 | 927 | 3,3 | 2 | 1 | 0,984 | 0,669 | 0,816 | 0,959 | 1,024 | 0,967 | 0,853 | 0,797 | 0,94 | -0,06 | 5,7E-01 |
| P20152 | Vimentin OS=Mus musculus OX=10090 GN=Vim PE=1 SV=3 | 40 | 15 | 21 | 12 | 466 | 53,7 | 5,12 | 8635 | 9,4 | 11 | 4 | 0,955 | 0,952 | 1,073 | 1,092 | 1,089 | 0,796 | 1,05 | 0,958 | 1,05 | 0,05 | 5,7E-01 |
| P43006 | Excitatory amino acid transporter 2 OS=Mus musculus OX=1009 | 11 | 7 | 13 | 7 | 572 | 62 | 6,68 | 3774 | 14,53 | 6 | 1 | 1,061 | 0,994 | 1,091 | 0,856 | 0,692 | 1,064 | 1,025 | 0,984 | 1,06 | 0,06 | 5,7E-01 |
| O35417 | Proenkephalin-B OS=Mus musculus OX=10090 GN=Pdyn PE=2 SV | 4 | 1 | 1 | 1 | 248 | 28 | 6,06 | 551 | | 1 | | 1,019 | 0,867 | 1,07 | 1,132 | 1,083 | 1,192 | 0,863 | 1,175 | 0,95 | -0,05 | 5,7E-01 |
| Q62086 | Serum paraoxonase/arylesterase 2 OS=Mus musculus OX=10090 | 2 | 1 | 1 | 1 | 354 | 39,6 | 5,83 | | 2,71 | | 1 | 0,868 | 1,212 | 0,747 | 0,985 | 0,658 | 0,787 | 0,882 | 1,145 | 1,10 | 0,10 | 5,7E-01 |
| P49312 | Heterogeneous nuclear ribonucleoprotein A1 OS=Mus musculus | 41 | 14 | 20 | 11 | 320 | 34,2 | 9,23 | 5904 | 27,92 | 8 | 7 | 0,866 | 0,785 | 1,069 | 1,028 | 0,887 | 0,919 | 1,138 | 1,012 | 0,95 | -0,05 | 5,7E-01 |
| Q9CQ45 | Neudesin OS=Mus musculus OX=10090 GN=Nenf PE=1 SV=1 | 20 | 3 | 5 | 3 | 171 | 18,9 | 5,27 | 535 | 12,4 | 1 | 2 | 1 | 1,125 | 0,887 | 1,104 | 0,932 | 0,945 | 1,089 | 0,995 | 1,04 | 0,04 | 5,7E-01 |
| P08121 | Collagen alpha-1(III) chain OS=Mus musculus OX=10090 GN=Col | 1 | 1 | 1 | 1 | 1464 | 138,9 | 6,52 | 590 | | 1 | | 0,86 | 0,849 | 0,928 | 1,054 | 1,213 | 0,618 | 0,974 | 0,47 | 1,13 | 0,13 | 5,7E-01 |
| P68134 | Actin, alpha skeletal muscle OS=Mus musculus OX=10090 GN=A | 47 | 24 | 334 | 1 | 377 | 42 | 5,39 | 136641 | 278,07 | 16 | 17 | 1,095 | 0,994 | 1,271 | 1,072 | 1,012 | 1,265 | 1,024 | 1,393 | 0,94 | -0,06 | 5,7E-01 |
| Q8C079 | Striatin-interacting protein 1 OS=Mus musculus OX=10090 GN=S | 4 | 3 | 3 | 3 | 837 | 95,5 | 6,25 | 1203 | 0 | 2 | 1 | 1,245 | 1 | 1,03 | 0,996 | 0,926 | 1,18 | 1,292 | 1,104 | 0,95 | -0,05 | 5,7E-01 |
| Q9CQH3 | NADH dehydrogenase [ubiquinone] 1 beta subcomplex subunit | 19 | 4 | 4 | 4 | 189 | 21,7 | 9,41 | 2174 | 3,94 | 3 | 1 | 0,984 | 1,035 | 0,804 | 0,961 | 0,944 | 0,938 | 0,988 | 1,046 | 0,97 | -0,03 | 5,8E-01 |
| Q9Z1G4 | V-type proton ATPase 116 kDa subunit a isoform 1 OS=Mus mus | 24 | 23 | 45 | 23 | 839 | 96,4 | 6,76 | 21571 | 31,1 | 16 | 8 | 1,045 | 1,029 | 0,997 | 0,988 | 0,925 | 1,041 | 1,035 | 0,987 | 1,02 | 0,02 | 5,8E-01 |
| Q80Y17 | Lethal(2) giant larvae protein homolog 1 OS=Mus musculus OX= | 2 | 2 | 2 | 2 | 1036 | 112,5 | 6,48 | 704 | | 2 | | 1,054 | 1,093 | 1,172 | 1,36 | 0,665 | 1,252 | 1,082 | 1,302 | 1,09 | 0,09 | 5,8E-01 |
| O35136 | Neural cell adhesion molecule 2 OS=Mus musculus OX=10090 G | 26 | 21 | 29 | 21 | 837 | 93,1 | 6,07 | 9061 | 39,49 | 10 | 11 | 0,96 | 0,992 | 0,963 | 0,917 | 0,968 | 0,928 | 0,939 | 0,955 | 1,01 | 0,01 | 5,8E-01 |
| P48962 | ADP/ATP translocase 1 OS=Mus musculus OX=10090 GN=Slc25a | 44 | 15 | 30 | 8 | 298 | 32,9 | 9,72 | 5712 | 59,69 | 6 | 10 | 0,958 | 1,006 | 1,041 | 0,998 | 0,832 | 0,926 | 1,031 | 1,078 | 1,04 | 0,04 | 5,8E-01 |
| P35550 | rRNA 2'-O-methyltransferase fibrillarin OS=Mus musculus OX=10 | 15 | 5 | 7 | 5 | 327 | 34,3 | 10,24 | 2280 | 7,16 | 4 | 1 | 1,106 | 0,929 | 1,147 | 0,956 | 1,311 | 0,908 | 1,133 | 1,025 | 0,95 | -0,05 | 5,8E-01 |
| Q8BHZ0 | Protein FAM49A OS=Mus musculus OX=10090 GN=Fam49a PE=1 | 15 | 4 | 8 | 3 | 323 | 37,3 | 6,01 | 4407 | | 4 | | 0,975 | 1,092 | 1,123 | 0,884 | 0,643 | 1,064 | 0,944 | 1,136 | 1,08 | 0,08 | 5,8E-01 |
| P39054 | Dynamin-2 OS=Mus musculus OX=10090 GN=Dnm2 PE=1 SV=2 | 18 | 18 | 32 | 4 | 870 | 98,1 | 7,43 | 12504 | 24,8 | 12 | 7 | 0,9 | 0,979 | 0,962 | 0,989 | 0,95 | 0,914 | 1,046 | 1,003 | 0,98 | -0,02 | 5,8E-01 |
| Q5SXY1 | Cytospin-B OS=Mus musculus OX=10090 GN=Specc1 PE=1 SV=2 | 1 | 2 | 2 | 2 | 1067 | 118 | 6,64 | 316 | 2,18 | 1 | 1 | 0,994 | 0,986 | 0,846 | 0,782 | 0,687 | 1,009 | 0,944 | 0,749 | 1,06 | 0,06 | 5,8E-01 |
| Q9Z204 | Heterogeneous nuclear ribonucleoproteins C1/C2 OS=Mus musc | 31 | 11 | 21 | 11 | 313 | 34,4 | 5,05 | 9982 | 8,69 | 8 | 3 | 0,896 | 0,867 | 1,109 | 1,093 | 0,868 | 1,006 | 1,191 | 1,124 | 0,95 | -0,05 | 5,8E-01 |
| P58871 | 182 kDa tankyrase-1-binding protein OS=Mus musculus OX=100 | 3 | 3 | 3 | 3 | 1720 | 181,7 | 4,88 | 1761 | | 3 | | 1,392 | 0,757 | 1,221 | 1,077 | 1,166 | 0,954 | 1,039 | 0,951 | 1,08 | 0,08 | 5,8E-01 |
| Q8R035 | Peptidyl-tRNA hydrolase ICT1, mitochondrial OS=Mus musculus | 9 | 2 | 2 | 2 | 206 | 23,5 | 10,18 | 641 | 0 | 1 | 1 | 1,296 | 0,808 | 1,229 | 0,99 | 1,284 | 0,932 | 0,8 | 0,95 | 1,09 | 0,09 | 5,8E-01 |
| O35345 | Importin subunit alpha-7 OS=Mus musculus OX=10090 GN=Kpna | 5 | 2 | 6 | 2 | 536 | 59,9 | 4,94 | 1861 | | 2 | | 0,942 | 0,892 | 1,003 | 0,892 | 0,684 | 1,344 | 0,874 | 1,18 | 0,91 | -0,09 | 5,8E-01 |
| P40124 | Adenylyl cyclase-associated protein 1 OS=Mus musculus OX=100 | 39 | 18 | 32 | 17 | 474 | 51,5 | 7,52 | 9219 | 38,19 | 11 | 10 | 1,048 | 0,969 | 1,076 | 1,07 | 0,894 | 1,11 | 0,989 | 1,048 | 1,03 | 0,03 | 5,8E-01 |
| O70492 | Sorting nexin-3 OS=Mus musculus OX=10090 GN=Snx3 PE=1 SV= | 17 | 3 | 3 | 2 | 162 | 18,7 | 8,66 | 2172 | | 3 | | 0,963 | 1,073 | 1,038 | 0,934 | 1,163 | 0,811 | 1,058 | 1,192 | 0,95 | -0,05 | 5,8E-01 |
| P28650 | Adenylosuccinate synthetase isozyme 1 OS=Mus musculus OX=1 | 4 | 1 | 3 | 1 | 457 | 50,2 | 8,43 | 323 | 4,88 | 1 | 1 | 1,48 | 1,451 | 1,11 | 1,266 | 1,248 | 1,36 | 1,387 | 1,038 | 1,05 | 0,05 | 5,8E-01 |
| B2RUJ5 | Amyloid-beta A4 precursor protein-binding family A member 1 ( | 9 | 6 | 7 | 5 | 842 | 92,9 | 4,88 | 2373 | 11,29 | 4 | 3 | 1,143 | 1,033 | 1,075 | 1,16 | 1,169 | 1,031 | 1,216 | 1,111 | 0,97 | -0,03 | 5,8E-01 |
| P70362 | Ubiquitin recognition factor in ER-associated degradation protei | 5 | 2 | 3 | 2 | 307 | 34,5 | 6,7 | 1100 | 2,11 | 1 | 1 | 0,998 | 1,024 | 1,056 | 1,068 | 1,087 | 1,034 | 0,998 | 0,95 | 1,02 | 0,02 | 5,8E-01 |
| P58854 | Gamma-tubulin complex component 3 OS=Mus musculus OX=10 | 1 | 1 | 1 | 1 | 905 | 103,4 | 8,32 | | 1,95 | | 1 | 1,207 | 1,066 | 1,097 | 0,822 | 1,117 | 1,042 | 0,937 | 0,868 | 1,06 | 0,06 | 5,8E-01 |
| O35099 | Mitogen-activated protein kinase kinase kinase 5 OS=Mus musc | 1 | 1 | 1 | 1 | 1380 | 154,4 | 5,78 | | 0 | | 1 | 0,992 | 0,834 | 1,226 | 1,164 | 1,144 | 0,672 | 0,805 | 1,225 | 1,10 | 0,10 | 5,8E-01 |
| Q9QUR8 | Semaphorin-7A OS=Mus musculus OX=10090 GN=Sema7a PE=1 | 11 | 5 | 5 | 5 | 664 | 74,9 | 7,74 | 795 | 0 | 3 | 2 | 0,941 | 1,551 | 1,166 | 1,471 | 1,341 | 1,169 | 1,412 | 1,591 | 0,93 | -0,07 | 5,8E-01 |
| Q9CR57 | 60S ribosomal protein L14 OS=Mus musculus OX=10090 GN=Rpl | 19 | 6 | 9 | 5 | 217 | 23,5 | 11,02 | 3086 | 6,71 | 5 | 1 | 0,936 | 0,906 | 1,22 | 0,894 | 0,994 | 1,115 | 0,99 | 1,049 | 0,95 | -0,05 | 5,8E-01 |
| P48453 | Serine/threonine-protein phosphatase 2B catalytic subunit beta | 36 | 15 | 42 | 10 | 525 | 59,1 | 5,91 | 23015 | 14,22 | 11 | 4 | 0,967 | 0,99 | 0,993 | 1,041 | 0,998 | 0,968 | 1,054 | 1,027 | 0,99 | -0,01 | 5,8E-01 |
| Q8CIG8 | Protein arginine N-methyltransferase 5 OS=Mus musculus OX=1 | 6 | 4 | 5 | 4 | 637 | 72,6 | 6,42 | 1764 | 1,85 | 3 | 1 | 0,861 | 0,899 | 0,83 | 0,926 | 0,833 | 0,951 | 0,812 | 0,832 | 1,03 | 0,03 | 5,8E-01 |
| P62880 | Guanine nucleotide-binding protein G(I)/G(S)/G(T) subunit beta- | 26 | 7 | 27 | 2 | 340 | 37,3 | 6 | 12873 | 16,39 | 5 | 3 | 0,921 | 0,966 | 1,153 | 0,919 | 1,16 | 1,215 | 0,89 | 0,923 | 0,95 | -0,05 | 5,8E-01 |
| O35250 | Exocyst complex component 7 OS=Mus musculus OX=10090 GN | 13 | 7 | 9 | 7 | 697 | 79,9 | 6,98 | 4003 | 2,62 | 7 | 1 | 1,089 | 1,144 | 1,133 | 1,071 | 1,12 | 1,12 | 1,073 | 1,073 | 1,01 | 0,01 | 5,8E-01 |
| Q9JHG6 | Calcipressin-1 OS=Mus musculus OX=10090 GN=Rcan1 PE=1 SV= | 4 | 1 | 1 | 1 | 251 | 28,1 | 5,41 | 587 | | 1 | | 0,772 | 0,887 | 0,78 | 0,828 | 0,945 | 0,632 | 0,864 | 0,63 | 1,06 | 0,06 | 5,8E-01 |
| P24527 | Leukotriene A-4 hydrolase OS=Mus musculus OX=10090 GN=Lta | 15 | 8 | 13 | 8 | 611 | 69 | 6,42 | 7218 | 3,12 | 7 | 1 | 0,903 | 0,964 | 1,239 | 0,954 | 0,867 | 1,155 | 0,914 | 0,891 | 1,06 | 0,06 | 5,9E-01 |
| Q9CY27 | Very-long-chain enoyl-CoA reductase OS=Mus musculus OX=100 | 9 | 3 | 3 | 3 | 308 | 36,1 | 9,55 | 1334 | 2,76 | 2 | 1 | 0,82 | 0,973 | 0,826 | 0,946 | 0,924 | 0,731 | 0,927 | 0,843 | 1,04 | 0,04 | 5,9E-01 |
| Q9Z1W8 | Potassium-transporting ATPase alpha chain 2 OS=Mus musculus | 6 | 7 | 46 | 1 | 1035 | 114,7 | 6,33 | 20661 | 24,23 | 3 | 4 | 1,186 | 1,068 | 0,875 | 1,046 | 0,851 | 1,101 | 0,953 | 1,072 | 1,05 | 0,05 | 5,9E-01 |
| P62984 | Ubiquitin-60S ribosomal protein L40 OS=Mus musculus OX=1009 | 52 | 12 | 75 | 11 | 128 | 14,7 | 9,83 | 37854 | 26,22 | 10 | 5 | 1 | 0,991 | 1,125 | 0,992 | 0,994 | 1,106 | 0,937 | 0,956 | 1,03 | 0,03 | 5,9E-01 |
| Q91YP2 | Neurolysin, mitochondrial OS=Mus musculus OX=10090 GN=Nln | 19 | 13 | 18 | 13 | 704 | 80,4 | 6,44 | 8148 | 4,63 | 10 | 4 | 0,961 | 0,985 | 0,98 | 0,972 | 0,931 | 1,025 | 0,994 | 0,995 | 0,99 | -0,01 | 5,9E-01 |
| Q9Z1N5 | Spliceosome RNA helicase Ddx39b OS=Mus musculus OX=10090 | 11 | 5 | 7 | 5 | 428 | 49 | 5,67 | 3532 | | 5 | | 0,853 | 0,789 | 0,76 | 0,859 | 1,103 | 0,718 | 0,856 | 0,785 | 0,94 | -0,06 | 5,9E-01 |
| Q8BL66 | Early endosome antigen 1 OS=Mus musculus OX=10090 GN=Eea | 2 | 3 | 3 | 3 | 1411 | 160,8 | 5,77 | 1659 | | 3 | | 1,026 | 0,811 | 1,168 | 0,858 | 0,889 | 0,868 | 0,887 | 1,016 | 1,06 | 0,06 | 5,9E-01 |
| O89116 | Vesicle transport through interaction with t-SNAREs homolog 1A | 4 | 1 | 1 | 1 | 217 | 25 | 6,4 | 819 | | 1 | | 0,935 | 1,007 | 0,884 | 0,77 | 1,116 | 0,673 | 1,279 | 0,857 | 0,92 | -0,08 | 5,9E-01 |
| P70175 | Disks large homolog 3 OS=Mus musculus OX=10090 GN=Dlg3 PE | 21 | 16 | 25 | 15 | 849 | 93,4 | 6,79 | 10630 | 13,35 | 13 | 6 | 1,039 | 0,983 | 1,057 | 0,964 | 0,997 | 0,987 | 0,996 | 1,011 | 1,01 | 0,01 | 5,9E-01 |
| P63054 | Calmodulin regulator protein PCP4 OS=Mus musculus OX=10090 | 44 | 3 | 10 | 3 | 62 | 6,8 | 6,71 | 1129 | 27,12 | 1 | 2 | 0,884 | 0,971 | 0,938 | 0,981 | 0,7 | 1,107 | 0,953 | 0,805 | 1,06 | 0,06 | 5,9E-01 |
| Q3UMY5 | Echinoderm microtubule-associated protein-like 4 OS=Mus mus | 6 | 5 | 7 | 5 | 988 | 110 | 6,54 | 2324 | 5,72 | 3 | 2 | 0,734 | 0,735 | 0,961 | 0,73 | 0,335 | 0,983 | 0,705 | 0,799 | 1,12 | 0,12 | 5,9E-01 |
| Q8BFQ8 | Glutamine amidotransferase-like class 1 domain-containing prot | 12 | 3 | 4 | 3 | 220 | 23,3 | 7,05 | 2207 | | 3 | | 0,956 | 0,953 | 1,139 | 0,94 | 1,098 | 0,928 | 0,881 | 0,927 | 1,04 | 0,04 | 5,9E-01 |
| Q9DBX6 | Cytochrome P450 2S1 OS=Mus musculus OX=10090 GN=Cyp2s1 | 3 | 1 | 1 | 1 | 501 | 55,6 | 8,78 | 388 | | 1 | | 0,755 | 0,805 | 1,234 | 0,834 | 1,619 | 0,643 | 1,005 | 0,897 | 0,87 | -0,13 | 5,9E-01 |
| P56379 | ATP synthase subunit ATP5MPL, mitochondrial OS=Mus musculu | 14 | 1 | 1 | 1 | 58 | 6,7 | 9,99 | | 2,44 | | 1 | 0,947 | 0,953 | 0,888 | 0,897 | 0,682 | 0,893 | 0,977 | 0,971 | 1,05 | 0,05 | 5,9E-01 |

| | | | | | | | | | | | | | | | | | | | | | | | |
|---|---|---|---|---|---|---|---|---|---|---|---|---|---|---|---|---|---|---|---|---|---|---|---|
| B2RXS4 | Plexin-B2 OS=Mus musculus OX=10090 GN=Plxnb2 PE=1 SV=1 | 11 | 14 | 19 | 14 | 1842 | 206,1 | 5,87 | 7003 | 21,04 | 11 | 5 | 1,004 | 0,987 | 0,953 | 1,044 | 1,008 | 1,023 | 0,923 | 0,967 | 1,02 | 0,02 | 5,9E-01 |
| Q9D6S7 | Ribosome-recycling factor, mitochondrial OS=Mus musculus OX | 6 | 1 | 1 | 1 | 262 | 29 | 9,85 | 682 | | 1 | | 0,664 | 0,828 | 0,881 | 0,575 | 0,564 | 0,712 | 0,64 | 0,826 | 1,08 | 0,08 | 5,9E-01 |
| P19324 | Serpin H1 OS=Mus musculus OX=10090 GN=Serpinh1 PE=1 SV=3 | 9 | 3 | 5 | 3 | 417 | 46,5 | 8,82 | 3807 | | 3 | | 1,289 | 1,015 | 1,054 | 1,368 | 1,335 | 1,027 | 0,906 | 1,169 | 1,07 | 0,07 | 5,9E-01 |
| P62259 | 14-3-3 protein epsilon OS=Mus musculus OX=10090 GN=Ywhae | 60 | 22 | 135 | 19 | 255 | 29,2 | 4,74 | 50665 | 163,53 | 10 | 18 | 0,998 | 0,997 | 0,938 | 1,113 | 0,776 | 1,105 | 0,984 | 1,003 | 1,05 | 0,05 | 5,9E-01 |
| Q8C460 | ERI1 exoribonuclease 3 OS=Mus musculus OX=10090 GN=Eri3 P | 8 | 2 | 3 | 2 | 337 | 37,2 | 8,07 | 1370 | 2,45 | 2 | 1 | 1,149 | 1,57 | 1,295 | 1,186 | 0,814 | 1,567 | 1,2 | 1,209 | 1,09 | 0,09 | 5,9E-01 |
| Q6P1B1 | Xaa-Pro aminopeptidase 1 OS=Mus musculus OX=10090 GN=Xp | 15 | 9 | 11 | 9 | 623 | 69,5 | 5,54 | 5432 | 4,61 | 7 | 2 | 0,982 | 1,062 | 0,97 | 1,028 | 0,946 | 1,146 | 1,033 | 1,022 | 0,97 | -0,03 | 5,9E-01 |
| Q8R307 | Vacuolar protein sorting-associated protein 18 homolog OS=Mu | 2 | 2 | 2 | 2 | 973 | 110,1 | 6,09 | 1119 | | 2 | | 1,04 | 0,833 | 0,78 | 0,896 | 0,772 | 0,717 | 1,043 | 0,811 | 1,06 | 0,06 | 5,9E-01 |
| P28798 | Progranulin OS=Mus musculus OX=10090 GN=Grn PE=1 SV=2 | 7 | 3 | 5 | 3 | 589 | 63,4 | 6,8 | 2165 | 0 | 3 | 1 | 1,159 | 1,089 | 0,911 | 1,312 | 1,174 | 1,203 | 0,907 | 0,929 | 1,06 | 0,06 | 5,9E-01 |
| P17742 | Peptidyl-prolyl cis-trans isomerase A OS=Mus musculus OX=100 | 65 | 12 | 55 | 12 | 164 | 18 | 7,9 | 10340 | 111,2 | 8 | 9 | 0,996 | 0,986 | 1,012 | 0,998 | 0,858 | 1,085 | 0,944 | 0,997 | 1,03 | 0,03 | 5,9E-01 |
| Q8R1V4 | Transmembrane emp24 domain-containing protein 4 OS=Mus m | 4 | 1 | 1 | 1 | 227 | 26 | 8,18 | 572 | | 1 | | 0,926 | 0,875 | 0,939 | 1,051 | 0,907 | 0,708 | 1,019 | 0,98 | 1,05 | 0,05 | 5,9E-01 |
| Q9CWF2 | Tubulin beta-2B chain OS=Mus musculus OX=10090 GN=Tubb2b | 69 | 28 | 549 | 2 | 445 | 49,9 | 4,89 | 229533 | 416,49 | 23 | 21 | 1,038 | 0,978 | 1,042 | 1,05 | 0,95 | 1,028 | 0,98 | 1,077 | 1,02 | 0,02 | 5,9E-01 |
| P55012 | Solute carrier family 12 member 2 OS=Mus musculus OX=10090 | 3 | 3 | 3 | 3 | 1205 | 131 | 7,33 | 1985 | | 3 | | 0,974 | 0,88 | 1,11 | 0,869 | 0,954 | 1,063 | 1,06 | 0,909 | 0,96 | -0,04 | 5,9E-01 |
| Q8BG51 | Mitochondrial Rho GTPase 1 OS=Mus musculus OX=10090 GN=F | 5 | 2 | 3 | 2 | 631 | 72,2 | 6,49 | 2132 | | 2 | | 1,035 | 1,182 | 1,105 | 1,158 | 1,492 | 1,002 | 1,122 | 1,115 | 0,95 | -0,05 | 5,9E-01 |
| Q9QZQ1 | Afadin OS=Mus musculus OX=10090 GN=Afdn PE=1 SV=3 | 9 | 15 | 19 | 15 | 1820 | 206,4 | 6,32 | 8765 | 11,18 | 12 | 3 | 0,989 | 0,924 | 1,02 | 0,979 | 0,946 | 1,016 | 0,994 | 1,014 | 0,99 | -0,01 | 5,9E-01 |
| Q8C7K6 | Prenylcysteine oxidase-like OS=Mus musculus OX=10090 GN=Pc | 3 | 1 | 1 | 1 | 495 | 54,8 | 7,65 | 722 | | 1 | | 0,976 | 1,031 | 0,699 | 1,157 | 0,942 | 1,113 | 1,123 | 0,932 | 0,94 | -0,06 | 5,9E-01 |
| Q9Z110 | Delta-1-pyrroline-5-carboxylate synthase OS=Mus musculus OX= | 8 | 6 | 8 | 6 | 795 | 87,2 | 7,55 | 3012 | 5,57 | 4 | 2 | 1,025 | 1,085 | 0,977 | 1,022 | 0,84 | 1,106 | 1,045 | 1,367 | 0,94 | -0,06 | 5,9E-01 |
| Q8BIF0 | CD99 antigen-like protein 2 OS=Mus musculus OX=10090 GN=C | 8 | 1 | 1 | 1 | 237 | 25,4 | 4,86 | 414 | | 1 | | 0,89 | 0,813 | 0,607 | 0,832 | 1,199 | 0,475 | 0,999 | 0,84 | 0,89 | -0,11 | 5,9E-01 |
| Q9CQ65 | S-methyl-5'-thioadenosine phosphorylase OS=Mus musculus OX | 9 | 2 | 2 | 2 | 283 | 31 | 7,14 | 783 | | 2 | | 1,045 | 1,201 | 1,054 | 1,233 | 0,613 | 1,252 | 1,22 | 1,098 | 1,08 | 0,08 | 5,9E-01 |
| Q80TJ1 | Calcium-dependent secretion activator 1 OS=Mus musculus OX= | 17 | 15 | 17 | 15 | 1355 | 153 | 5,74 | 6559 | 11,56 | 12 | 3 | 1,02 | 1,012 | 1,031 | 1,002 | 1,029 | 0,947 | 1,057 | 0,974 | 1,01 | 0,01 | 6,0E-01 |
| Q9CQV6 | Microtubule-associated proteins 1A/1B light chain 3B OS=Mus n | 37 | 6 | 15 | 3 | 125 | 14,6 | 8,43 | 7058 | 9,74 | 4 | 4 | 0,878 | 0,82 | 1,104 | 0,954 | 0,921 | 0,91 | 0,859 | 0,924 | 1,04 | 0,04 | 6,0E-01 |
| P16332 | Methylmalonyl-CoA mutase, mitochondrial OS=Mus musculus O | 3 | 2 | 2 | 2 | 748 | 82,8 | 6,89 | 1209 | | 2 | | 1,437 | 1,004 | 1,258 | 1,251 | 1,746 | 1,176 | 1,27 | 1,132 | 0,93 | -0,07 | 6,0E-01 |
| Q9D710 | Thioredoxin-related transmembrane protein 2 OS=Mus musculu | 12 | 2 | 3 | 2 | 295 | 33,9 | 8,75 | 696 | 5,68 | 1 | 1 | 0,963 | 1,151 | 0,981 | 0,989 | 1,024 | 0,894 | 1,251 | 1,109 | 0,95 | -0,05 | 6,0E-01 |
| Q8K0C4 | Lanosterol 14-alpha demethylase OS=Mus musculus OX=10090 | 8 | 4 | 5 | 4 | 503 | 56,7 | 8,41 | 1715 | 5,46 | 2 | 2 | 1,036 | 0,976 | 1,007 | 1,149 | 0,928 | 0,993 | 1,047 | 1,086 | 1,03 | 0,03 | 6,0E-01 |
| Q9QYI3 | DnaJ homolog subfamily C member 7 OS=Mus musculus OX=100 | 2 | 1 | 3 | 1 | 494 | 56,4 | 6,49 | 1114 | | 1 | | 0,833 | 0,806 | 1,147 | 1,109 | 0,946 | 0,985 | 0,882 | 0,873 | 1,06 | 0,06 | 6,0E-01 |
| P62305 | Small nuclear ribonucleoprotein E OS=Mus musculus OX=10090 | 13 | 1 | 3 | 1 | 92 | 10,8 | 9,44 | 2136 | | 1 | | 0,777 | 1,046 | 0,892 | 1,058 | 1,149 | 0,745 | 1,146 | 0,992 | 0,94 | -0,06 | 6,0E-01 |
| Q9D8W5 | 26S proteasome non-ATPase regulatory subunit 12 OS=Mus mu | 10 | 5 | 5 | 5 | 456 | 52,9 | 7,06 | 1658 | 4,77 | 3 | 2 | 1,047 | 1 | 1,039 | 1,04 | 0,831 | 1,133 | 1,053 | 0,963 | 1,04 | 0,04 | 6,0E-01 |
| Q9R1P4 | Proteasome subunit alpha type-1 OS=Mus musculus OX=10090 | 43 | 12 | 18 | 12 | 263 | 29,5 | 6,46 | 9676 | 6,39 | 10 | 3 | 1,07 | 1,107 | 0,92 | 1,104 | 0,841 | 1,003 | 1,127 | 1,062 | 1,04 | 0,04 | 6,0E-01 |
| Q7TSJ2 | Microtubule-associated protein 6 OS=Mus musculus OX=10090 | 49 | 33 | 51 | 33 | 906 | 96,4 | 9,5 | 26084 | 25,12 | 27 | 8 | 1,075 | 1,017 | 0,968 | 1,011 | 0,942 | 1,029 | 1,035 | 0,997 | 1,02 | 0,02 | 6,0E-01 |
| A2RSJ4 | UHRF1-binding protein 1-like OS=Mus musculus OX=10090 GN= | 1 | 1 | 1 | 1 | 1457 | 161,8 | 6,61 | 545 | | 1 | | 0,814 | 0,925 | 0,754 | 0,844 | 0,734 | 0,883 | 0,85 | 0,758 | 1,03 | 0,03 | 6,0E-01 |
| Q9Z2G6 | Protein sel-1 homolog 1 OS=Mus musculus OX=10090 GN=Sel1l | 3 | 2 | 3 | 2 | 790 | 88,3 | 5,57 | 1225 | 2,2 | 2 | 1 | 1,451 | 0,982 | 0,91 | 1,418 | 1,077 | 0,992 | 1,048 | 1,296 | 1,08 | 0,08 | 6,0E-01 |
| P54763 | Ephrin type-B receptor 2 OS=Mus musculus OX=10090 GN=Ephb | 8 | 7 | 8 | 5 | 986 | 109,8 | 5,71 | 2669 | 10,2 | 4 | 3 | 1,026 | 0,965 | 0,869 | 1,061 | 1,12 | 1,037 | 0,91 | 0,989 | 0,97 | -0,03 | 6,0E-01 |
| O89053 | Coronin-1A OS=Mus musculus OX=10090 GN=Coro1a PE=1 SV=5 | 27 | 16 | 28 | 14 | 461 | 51 | 6,48 | 11703 | 21,56 | 11 | 6 | 0,97 | 0,871 | 0,995 | 0,887 | 0,83 | 0,985 | 0,892 | 0,918 | 1,03 | 0,03 | 6,0E-01 |
| Q9WUD1 | STIP1 homology and U box-containing protein 1 OS=Mus muscu | 14 | 4 | 8 | 4 | 304 | 34,9 | 6,01 | 3804 | 6,34 | 3 | 2 | 1,065 | 1,011 | 1,123 | 1,13 | 1,061 | 1,173 | 1,052 | 1,132 | 0,98 | -0,02 | 6,0E-01 |
| Q61081 | Hsp90 co-chaperone Cdc37 OS=Mus musculus OX=10090 GN=C | 17 | 6 | 7 | 6 | 379 | 44,6 | 5,34 | 3488 | 2,16 | 5 | 1 | 1,149 | 0,884 | 0,882 | 1,063 | 0,91 | 0,967 | 0,997 | 0,951 | 1,04 | 0,04 | 6,0E-01 |
| P28741 | Kinesin-like protein KIF3A OS=Mus musculus OX=10090 GN=Kif3 | 3 | 2 | 2 | 2 | 701 | 80,1 | 6,54 | 663 | 2,38 | 1 | 1 | 1,07 | 0,811 | 0,933 | 0,702 | 0,853 | 0,939 | 1,072 | 0,86 | 0,94 | -0,06 | 6,0E-01 |
| Q8R0H9 | ADP-ribosylation factor-binding protein GGA1 OS=Mus musculu | 6 | 4 | 4 | 4 | 635 | 69,9 | 5,27 | 2102 | | 4 | | 0,859 | 0,879 | 0,964 | 0,992 | 0,651 | 0,835 | 1,062 | 0,941 | 1,06 | 0,06 | 6,0E-01 |
| P70236 | Dual specificity mitogen-activated protein kinase kinase 6 OS=M | 9 | 3 | 4 | 3 | 334 | 37,4 | 7,39 | 2167 | | 3 | | 0,946 | 0,753 | 0,998 | 1,028 | 0,647 | 0,869 | 1,061 | 0,915 | 1,07 | 0,07 | 6,0E-01 |
| Q3UYG8 | ADP-ribose glycohydrolase MACROD2 OS=Mus musculus OX=10 | 2 | 1 | 1 | 1 | 475 | 52,1 | 4,75 | 700 | | 1 | | 0,928 | 1,098 | 1,147 | 1,128 | 0,984 | 1,275 | 0,92 | 0,901 | 1,05 | 0,05 | 6,0E-01 |
| P50136 | 2-oxoisovalerate dehydrogenase subunit alpha, mitochondrial C | 2 | 1 | 1 | 1 | 442 | 50,3 | 8,06 | 720 | | 1 | | 0,719 | 0,737 | 0,735 | 0,93 | 0,821 | 0,864 | 0,832 | 0,731 | 0,96 | -0,04 | 6,0E-01 |
| Q9WUU7 | Cathepsin Z OS=Mus musculus OX=10090 GN=Ctsz PE=1 SV=1 | 8 | 2 | 3 | 2 | 306 | 34 | 6,6 | 1078 | 2,3 | 1 | 1 | 1,075 | 1,061 | 0,941 | 1,26 | 1,336 | 1,22 | 0,976 | 1,038 | 0,95 | -0,05 | 6,0E-01 |
| P63017 | Heat shock cognate 71 kDa protein OS=Mus musculus OX=1009 | 61 | 43 | 176 | 29 | 646 | 70,8 | 5,52 | 73471 | 153,15 | 33 | 22 | 1,02 | 0,958 | 1,009 | 0,939 | 0,9 | 1,022 | 0,946 | 0,986 | 1,02 | 0,02 | 6,0E-01 |
| Q921J2 | GTP-binding protein Rheb OS=Mus musculus OX=10090 GN=Rhe | 23 | 5 | 5 | 5 | 184 | 20,4 | 5,92 | 1064 | 6,14 | 2 | 3 | 0,928 | 0,92 | 1,171 | 1,037 | 1,302 | 0,954 | 1,043 | 0,975 | 0,95 | -0,05 | 6,0E-01 |
| Q9CPP6 | NADH dehydrogenase [ubiquinone] 1 alpha subcomplex subunit | 23 | 2 | 3 | 2 | 116 | 13,4 | 8,1 | | 10,33 | | 2 | 0,998 | 1,07 | 0,901 | 0,855 | 2,019 | 0,872 | 0,79 | 0,808 | 0,85 | -0,15 | 6,0E-01 |
| P58281 | Dynamin-like 120 kDa protein, mitochondrial OS=Mus musculus | 21 | 20 | 29 | 20 | 960 | 111,3 | 7,55 | 9526 | 40,18 | 11 | 10 | 1,02 | 1,047 | 1,029 | 1,009 | 0,994 | 1,029 | 1,014 | 1,04 | 1,01 | 0,01 | 6,0E-01 |
| P07356 | Annexin A2 OS=Mus musculus OX=10090 GN=Anxa2 PE=1 SV=2 | 20 | 7 | 7 | 7 | 339 | 38,7 | 7,69 | 4061 | 2,39 | 6 | 1 | 1,048 | 1,221 | 0,834 | 1,073 | 1,184 | 1,202 | 1,107 | 0,91 | 0,95 | -0,05 | 6,1E-01 |
| Q9CXS4 | Centromere protein V OS=Mus musculus OX=10090 GN=Cenpv | 14 | 2 | 4 | 2 | 252 | 27,5 | 9,79 | 525 | 2,72 | 1 | 1 | 0,976 | 0,989 | 1,085 | 0,969 | 0,933 | 0,738 | 1,145 | 1,009 | 1,05 | 0,05 | 6,1E-01 |
| Q6PB90 | Protocadherin beta-14 OS=Mus musculus OX=10090 GN=Pcdhb | 2 | 2 | 2 | 2 | 796 | 87 | 4,89 | 555 | 2,3 | 1 | 1 | 0,997 | 0,952 | 1,214 | 0,904 | 1,015 | 1,109 | 0,956 | 1,168 | 0,96 | -0,04 | 6,1E-01 |
| Q91WJ8 | Far upstream element-binding protein 1 OS=Mus musculus OX= | 7 | 4 | 4 | 3 | 651 | 68,5 | 7,93 | 899 | 4,99 | 2 | 2 | 0,7 | 0,83 | 1,237 | 0,912 | 0,777 | 0,829 | 0,804 | 0,998 | 1,08 | 0,08 | 6,1E-01 |
| P20444 | Protein kinase C alpha type OS=Mus musculus OX=10090 GN=Pr | 10 | 8 | 10 | 4 | 672 | 76,8 | 7,14 | 1908 | 16,54 | 3 | 5 | 0,963 | 0,971 | 1,146 | 0,963 | 0,903 | 1,095 | 0,921 | 0,988 | 1,03 | 0,03 | 6,1E-01 |
| Q61330 | Contactin-2 OS=Mus musculus OX=10090 GN=Cntn2 PE=1 SV=2 | 1 | 1 | 1 | 1 | 1040 | 113,1 | 7,62 | 538 | | 1 | | 1,362 | 1,206 | 1,054 | 1,026 | 1,5 | 1,018 | 1,158 | 1,249 | 0,94 | -0,06 | 6,1E-01 |
| Q8BLQ9 | Cell adhesion molecule 2 OS=Mus musculus OX=10090 GN=Cadr | 30 | 12 | 23 | 12 | 435 | 47,5 | 5,41 | 9572 | 15,85 | 9 | 3 | 1,013 | 0,925 | 1,098 | 0,927 | 0,985 | 1,003 | 0,954 | 0,924 | 1,03 | 0,03 | 6,1E-01 |
| Q6NVF9 | Cleavage and polyadenylation specificity factor subunit 6 OS=M | 5 | 2 | 3 | 2 | 551 | 59,1 | 7,15 | 1825 | | 2 | | 1,078 | 0,902 | 1,094 | 1,131 | 1,048 | 0,927 | 1,2 | 1,214 | 0,96 | -0,04 | 6,1E-01 |
| Q62093 | Serine/arginine-rich splicing factor 2 OS=Mus musculus OX=100 | 7 | 2 | 2 | 2 | 221 | 25,5 | 11,85 | 1127 | | 2 | | 0,807 | 0,945 | 0,877 | 0,894 | 1,147 | 0,861 | 0,829 | 0,858 | 0,95 | -0,05 | 6,1E-01 |
| Q8R3Q2 | Serine/threonine-protein phosphatase 6 regulatory subunit 2 O | 4 | 3 | 3 | 3 | 923 | 100,4 | 4,82 | 953 | 1,76 | 2 | 1 | 0,844 | 0,752 | 0,744 | 0,897 | 0,629 | 0,99 | 0,939 | 0,868 | 0,94 | -0,06 | 6,1E-01 |
| P63038 | 60 kDa heat shock protein, mitochondrial OS=Mus musculus OX | 46 | 29 | 83 | 29 | 573 | 60,9 | 6,18 | 36641 | 69,73 | 21 | 13 | 1,026 | 0,992 | 1,049 | 0,99 | 1,046 | 1,045 | 0,993 | 1,014 | 0,99 | -0,01 | 6,1E-01 |
| P53395 | Lipoamide acyltransferase component of branched-chain alpha- | 2 | 1 | 1 | 1 | 482 | 53,2 | 8,6 | | 2,54 | | 1 | 0,978 | 1,187 | 1,143 | 0,912 | 1,365 | 1,069 | 1,047 | 0,971 | 0,95 | -0,05 | 6,1E-01 |
| Q63829 | COMM domain-containing protein 3 OS=Mus musculus OX=100 | 4 | 1 | 1 | 1 | 195 | 22 | 5,59 | 512 | | 1 | | 0,958 | 1,005 | 1,06 | 1,213 | 1,184 | 1,038 | 1,024 | 1,134 | 0,97 | -0,03 | 6,1E-01 |
| Q9D219 | B-cell CLL/lymphoma 9 protein OS=Mus musculus OX=10090 GN | 1 | 1 | 1 | 1 | 1425 | 148,9 | 8,91 | 198 | | 1 | | 1,116 | 1,539 | 0,963 | 1,438 | 1,242 | 1,182 | 1,164 | 1,972 | 0,91 | -0,09 | 6,1E-01 |
| P32507 | Nectin-2 OS=Mus musculus OX=10090 GN=Nectin2 PE=1 SV=2 | 8 | 3 | 3 | 3 | 530 | 57,3 | 4,86 | 1526 | 2,26 | 2 | 1 | 1,11 | 1,169 | 1,016 | 1,168 | 1,208 | 1,183 | 1,092 | 1,082 | 0,98 | -0,02 | 6,1E-01 |
| Q6PER3 | Microtubule-associated protein RP/EB family member 3 OS=Mu | 17 | 7 | 14 | 4 | 281 | 31,9 | 5,54 | 3947 | 27,65 | 4 | 3 | 1,009 | 1,06 | 0,951 | 1,046 | 0,889 | 1,003 | 1,083 | 0,992 | 1,02 | 0,02 | 6,1E-01 |
| Q9ESX5 | H/ACA ribonucleoprotein complex subunit DKC1 OS=Mus muscu | 2 | 1 | 1 | 1 | 509 | 57,4 | 9,28 | 261 | | 1 | | 0,915 | 0,808 | 0,79 | 1,098 | 1,103 | 0,866 | 0,958 | 0,874 | 0,95 | -0,05 | 6,1E-01 |
| P24288 | Branched-chain-amino-acid aminotransferase, cytosolic OS=Mu | 6 | 2 | 2 | 2 | 386 | 42,8 | 5,39 | 709 | 0 | 1 | 1 | 1,068 | 0,921 | 0,98 | 0,868 | 1,41 | 0,964 | 0,838 | 0,913 | 0,93 | -0,07 | 6,2E-01 |
| Q9JHW4 | Selenocysteine-specific elongation factor OS=Mus musculus OX= | 5 | 2 | 3 | 2 | 583 | 63,5 | 8,29 | 1545 | | 2 | | 1,204 | 1,025 | 0,914 | 0,937 | 1,062 | 0,86 | 1,176 | 1,197 | 0,95 | -0,05 | 6,2E-01 |
| Q3UHX2 | 28 kDa heat- and acid-stable phosphoprotein OS=Mus musculus | 19 | 4 | 6 | 4 | 181 | 20,6 | 7,39 | 1332 | 13,03 | 2 | 3 | 1,078 | 1,016 | 1,006 | 0,993 | 0,932 | 0,907 | 1,108 | 1,039 | 1,03 | 0,03 | 6,2E-01 |
| Q91VR2 | ATP synthase subunit gamma, mitochondrial OS=Mus musculus | 34 | 11 | 21 | 11 | 298 | 32,9 | 9,01 | 10585 | 11,16 | 8 | 4 | 1,01 | 0,91 | 0,999 | 0,956 | 0,994 | 1,061 | 0,908 | 0,994 | 0,98 | -0,02 | 6,2E-01 |

| | | | | | | | | | | | | | | | | | | | | | | | |
|---|---|---|---|---|---|---|---|---|---|---|---|---|---|---|---|---|---|---|---|---|---|---|---|
| P28652 | Calcium/calmodulin-dependent protein kinase type II subunit be | 31 | 16 | 60 | 11 | 542 | 60,4 | 7,28 | 26917 | 39,85 | 10 | 9 | 1,029 | 0,978 | 1,028 | 0,941 | 0,956 | 0,993 | 0,974 | 1,003 | 1,01 | 0,01 | 6,2E-01 |
| P70670 | Nascent polypeptide-associated complex subunit alpha, muscle- | 2 | 3 | 3 | 3 | 2187 | 220,4 | 9,35 | 950 | 5,05 | 2 | 1 | 0,998 | 1,194 | 1,207 | 0,927 | 1,031 | 1,065 | 1,194 | 0,813 | 1,05 | 0,05 | 6,2E-01 |
| Q922W5 | Pyrroline-5-carboxylate reductase 1, mitochondrial OS=Mus mu | 10 | 2 | 2 | 1 | 309 | 32,4 | 6,86 | 561 | | 2 | | 0,822 | 0,944 | 0,992 | 0,932 | 1,121 | 0,882 | 0,938 | 0,889 | 0,96 | -0,04 | 6,2E-01 |
| Q91VH6 | Protein MEMO1 OS=Mus musculus OX=10090 GN=Memo1 PE=1 | 12 | 3 | 4 | 3 | 297 | 33,7 | 7,14 | 2328 | | 3 | | 1,112 | 1,132 | 1,365 | 1,089 | 1,318 | 1,103 | 1,007 | 1,075 | 1,04 | 0,04 | 6,2E-01 |
| Q8R1B4 | Eukaryotic translation initiation factor 3 subunit C OS=Mus musc | 3 | 2 | 2 | 2 | 911 | 105,5 | 5,78 | 1076 | | 2 | | 0,84 | 1,212 | 0,83 | 0,921 | 1,165 | 0,77 | 0,975 | 1,166 | 0,93 | -0,07 | 6,2E-01 |
| Q8VDQ8 | NAD-dependent protein deacetylase sirtuin-2 OS=Mus musculus | 11 | 3 | 3 | 3 | 389 | 43,2 | 5,35 | 942 | 0 | 2 | 1 | 0,851 | 0,836 | 0,813 | 0,837 | 0,725 | 0,866 | 0,872 | 0,8 | 1,02 | 0,02 | 6,2E-01 |
| Q8CBY8 | Dynactin subunit 4 OS=Mus musculus OX=10090 GN=Dctn4 PE= | 6 | 2 | 2 | 2 | 467 | 53 | 7,72 | 1198 | | 2 | | 1,359 | 1,649 | 1,173 | 1,298 | 1,714 | 1,421 | 1,165 | 1,497 | 0,95 | -0,05 | 6,2E-01 |
| O35639 | Annexin A3 OS=Mus musculus OX=10090 GN=Anxa3 PE=1 SV=4 | 3 | 1 | 1 | 1 | 323 | 36,4 | 5,76 | 374 | | 1 | | 1,049 | 0,999 | 0,851 | 0,992 | 1,261 | 1,09 | 0,942 | 0,817 | 0,95 | -0,05 | 6,2E-01 |
| Q6PEB6 | MOB-like protein phocein OS=Mus musculus OX=10090 GN=Mo | 17 | 3 | 4 | 3 | 225 | 26 | 5,78 | 3040 | | 3 | | 0,898 | 1,057 | 0,924 | 0,941 | 0,851 | 1,004 | 0,981 | 1,121 | 0,97 | -0,03 | 6,2E-01 |
| P54285 | Voltage-dependent L-type calcium channel subunit beta-3 OS=M | 8 | 4 | 6 | 2 | 484 | 54,5 | 6,27 | 3729 | | 4 | | 0,973 | 0,946 | 0,912 | 0,773 | 1,089 | 0,862 | 0,934 | 0,864 | 0,96 | -0,04 | 6,2E-01 |
| Q91XF0 | Pyridoxine-5'-phosphate oxidase OS=Mus musculus OX=10090 G | 8 | 2 | 2 | 2 | 261 | 30,1 | 8,22 | 491 | 1,9 | 1 | 1 | 0,605 | 0,92 | 0,849 | 0,889 | 1,139 | 0,854 | 0,791 | 0,722 | 0,93 | -0,07 | 6,2E-01 |
| Q03265 | ATP synthase subunit alpha, mitochondrial OS=Mus musculus O | 49 | 33 | 108 | 33 | 553 | 59,7 | 9,19 | 50509 | 76,19 | 20 | 19 | 0,99 | 0,936 | 1,03 | 0,94 | 0,959 | 1,003 | 0,923 | 0,953 | 1,02 | 0,02 | 6,2E-01 |
| P84104 | Serine/arginine-rich splicing factor 3 OS=Mus musculus OX=1009 | 16 | 3 | 3 | 3 | 164 | 19,3 | 11,65 | | 4,31 | | 3 | 0,866 | 0,899 | 1,05 | 1,112 | 0,935 | 0,982 | 1,058 | 1,097 | 0,96 | -0,04 | 6,2E-01 |
| Q9WV34 | MAGUK p55 subfamily member 2 OS=Mus musculus OX=10090 | 18 | 9 | 10 | 8 | 552 | 61,5 | 6,44 | 3998 | 5,44 | 7 | 3 | 0,97 | 0,975 | 0,94 | 1,093 | 1,03 | 0,957 | 0,911 | 0,992 | 1,02 | 0,02 | 6,2E-01 |
| Q8CCS6 | Polyadenylate-binding protein 2 OS=Mus musculus OX=10090 G | 5 | 2 | 3 | 2 | 302 | 32,3 | 5,17 | | 6,97 | | 2 | 0,896 | 0,871 | 1,085 | 1,215 | 1,05 | 1,06 | 1,064 | 1,063 | 0,96 | -0,04 | 6,2E-01 |
| Q64487 | Receptor-type tyrosine-protein phosphatase delta OS=Mus mus | 13 | 19 | 23 | 15 | 1912 | 214,3 | 6,55 | 6312 | 30,19 | 7 | 12 | 0,996 | 1,029 | 1,08 | 0,95 | 1,156 | 1,009 | 0,975 | 1,016 | 0,98 | -0,02 | 6,2E-01 |
| P09055 | Integrin beta-1 OS=Mus musculus OX=10090 GN=Itgb1 PE=1 SV= | 11 | 7 | 13 | 7 | 798 | 88,2 | 5,94 | 7286 | 2,65 | 6 | 1 | 1,042 | 1,004 | 0,985 | 1,056 | 1,124 | 0,955 | 1,035 | 1,053 | 0,98 | -0,02 | 6,2E-01 |
| O08756 | 3-hydroxyacyl-CoA dehydrogenase type-2 OS=Mus musculus OX | 38 | 8 | 10 | 8 | 261 | 27,4 | 8,41 | 3917 | 10,17 | 6 | 2 | 1,119 | 1,225 | 1,061 | 1,26 | 1,15 | 1,081 | 1,155 | 1,174 | 1,02 | 0,02 | 6,2E-01 |
| P63087 | Serine/threonine-protein phosphatase PP1-gamma catalytic sub | 33 | 9 | 15 | 2 | 323 | 37 | 6,54 | 4836 | 19,37 | 6 | 4 | 1,022 | 0,865 | 0,941 | 0,892 | 1,092 | 0,771 | 1,15 | 0,902 | 0,95 | -0,05 | 6,2E-01 |
| Q9D7N9 | Adipocyte plasma membrane-associated protein OS=Mus muscu | 2 | 1 | 1 | 1 | 415 | 46,4 | 6,32 | 666 | | 1 | | 0,922 | 1,08 | 1,073 | 1,052 | 0,819 | 1,116 | 1,137 | 0,867 | 1,05 | 0,05 | 6,2E-01 |
| A2AFR3 | FERM and PDZ domain-containing protein 4 OS=Mus musculus O | 1 | 1 | 1 | 1 | 1320 | 144,9 | 5,25 | 353 | | 1 | | 1,102 | 0,994 | 0,989 | 1,15 | 0,939 | 0,76 | 1,451 | 0,728 | 1,09 | 0,09 | 6,2E-01 |
| P10637-5 | Isoform Tau-D of Microtubule-associated protein tau OS=Mus m | 72 | 30 | 110 | 3 | 372 | 38,9 | 9,5 | 44857 | 76,59 | 15 | 21 | 0,983 | 0,804 | 0,893 | 0,853 | 0,618 | 0,978 | 0,926 | 0,828 | 1,05 | 0,05 | 6,2E-01 |
| P27659 | 60S ribosomal protein L3 OS=Mus musculus OX=10090 GN=Rpl3 | 29 | 13 | 16 | 13 | 403 | 46,1 | 10,21 | 5432 | 16,03 | 7 | 6 | 0,983 | 0,906 | 0,936 | 1,081 | 0,912 | 0,943 | 0,976 | 0,988 | 1,02 | 0,02 | 6,2E-01 |
| Q99L43 | Phosphatidate cytidylyltransferase 2 OS=Mus musculus OX=100 | 7 | 2 | 2 | 2 | 444 | 51,3 | 7,05 | 1609 | | 2 | | 0,943 | 0,946 | 0,689 | 1,257 | 1,347 | 0,77 | 1,127 | 0,943 | 0,92 | -0,08 | 6,2E-01 |
| E9Q6P5 | Tetratricopeptide repeat protein 7B OS=Mus musculus OX=1009 | 3 | 3 | 3 | 3 | 843 | 94,1 | 6,9 | 634 | 4,61 | 1 | 2 | 0,904 | 1,017 | 0,959 | 0,933 | 0,934 | 1,046 | 1,035 | 0,891 | 0,98 | -0,02 | 6,2E-01 |
| Q3UU96 | Serine/threonine-protein kinase MRCK alpha OS=Mus musculus | 3 | 4 | 4 | 2 | 1719 | 195,4 | 6,51 | 1134 | 5,87 | 2 | 2 | 0,86 | 0,907 | 1,205 | 1,256 | 1,067 | 1,152 | 1,028 | 1,206 | 0,95 | -0,05 | 6,2E-01 |
| O35681 | Synaptotagmin-3 OS=Mus musculus OX=10090 GN=Syt3 PE=1 S\ | 3 | 2 | 2 | 2 | 587 | 63,2 | 6,65 | 222 | 2,31 | 1 | 1 | 1,084 | 1,174 | 1,093 | 1,073 | 0,969 | 1,021 | 1,204 | 1,113 | 1,03 | 0,03 | 6,2E-01 |
| Q8JZQ2 | AFG3-like protein 2 OS=Mus musculus OX=10090 GN=Afg3l2 PE= | 6 | 5 | 5 | 5 | 802 | 89,5 | 8,6 | 1763 | 4,19 | 3 | 2 | 0,877 | 1,025 | 0,999 | 0,986 | 0,989 | 0,861 | 0,936 | 1,006 | 1,03 | 0,03 | 6,2E-01 |
| Q04887 | Transcription factor SOX-9 OS=Mus musculus OX=10090 GN=Sox | 2 | 1 | 1 | 1 | 507 | 56 | 6,81 | 351 | | 1 | | 0,958 | 0,858 | 0,883 | 1,056 | 0,951 | 1,041 | 0,898 | 0,975 | 0,97 | -0,03 | 6,3E-01 |
| Q91XY4 | Protocadherin gamma-A4 OS=Mus musculus OX=10090 GN=Pcd | 3 | 2 | 3 | 2 | 930 | 100,3 | 4,87 | 1278 | 2,68 | 2 | 1 | 1,184 | 1,008 | 1,086 | 1,067 | 1,25 | 1,082 | 1,087 | 1,046 | 0,97 | -0,03 | 6,3E-01 |
| Q3UDE2 | Tubulin--tyrosine ligase-like protein 12 OS=Mus musculus OX=1 | 9 | 5 | 7 | 5 | 639 | 74 | 5,63 | 3126 | 2,28 | 4 | 1 | 0,865 | 0,959 | 0,976 | 0,997 | 0,654 | 1,023 | 0,975 | 0,962 | 1,05 | 0,05 | 6,3E-01 |
| P62918 | 60S ribosomal protein L8 OS=Mus musculus OX=10090 GN=Rpl8 | 32 | 9 | 17 | 9 | 257 | 28 | 11,03 | 5211 | 20,5 | 4 | 7 | 1,133 | 0,901 | 1,014 | 0,997 | 1,222 | 0,977 | 0,974 | 1,027 | 0,96 | -0,04 | 6,3E-01 |
| P61620 | Protein transport protein Sec61 subunit alpha isoform 1 OS=Mu | 2 | 1 | 1 | 1 | 476 | 52,2 | 8,06 | | 0 | | 1 | 1,151 | 1,125 | 0,757 | 0,987 | 0,808 | 1,137 | 0,729 | 1,071 | 1,07 | 0,07 | 6,3E-01 |
| Q8K0U4 | Heat shock 70 kDa protein 12A OS=Mus musculus OX=10090 GN | 26 | 14 | 19 | 14 | 675 | 74,8 | 6,77 | 6000 | 26,34 | 8 | 6 | 1,031 | 0,969 | 1,034 | 1,029 | 0,872 | 1,026 | 1,07 | 1,002 | 1,02 | 0,02 | 6,3E-01 |
| P62855 | 40S ribosomal protein S26 OS=Mus musculus OX=10090 GN=Rp: | 8 | 1 | 4 | 1 | 115 | 13 | 11 | 2598 | | 1 | | 1,127 | 0,9 | 0,91 | 0,885 | 1,419 | 0,827 | 0,925 | 0,947 | 0,93 | -0,07 | 6,3E-01 |
| Q8R5H1 | Ubiquitin carboxyl-terminal hydrolase 15 OS=Mus musculus OX= | 4 | 4 | 4 | 3 | 981 | 112,3 | 5,17 | 1568 | 2,49 | 3 | 1 | 0,952 | 1,056 | 0,953 | 1,143 | 0,968 | 1,034 | 1,016 | 0,987 | 1,02 | 0,02 | 6,3E-01 |
| P28663 | Beta-soluble NSF attachment protein OS=Mus musculus OX=100 | 57 | 17 | 30 | 15 | 298 | 33,5 | 5,47 | 10664 | 31,18 | 12 | 7 | 1,003 | 0,966 | 1,032 | 0,983 | 0,821 | 1,033 | 1,039 | 0,983 | 1,03 | 0,03 | 6,3E-01 |
| Q8BU30 | Isoleucine--tRNA ligase, cytoplasmic OS=Mus musculus OX=1009 | 2 | 2 | 5 | 2 | 1262 | 144,2 | 6,55 | 1506 | | 2 | | 1,162 | 1,375 | 1,147 | 0,693 | 1,059 | 1,441 | 1,237 | 0,997 | 0,92 | -0,08 | 6,3E-01 |
| Q9CQX8 | 28S ribosomal protein S36, mitochondrial OS=Mus musculus OX | 57 | 3 | 7 | 3 | 102 | 11,1 | 9,99 | 2030 | 11,92 | 2 | 3 | 0,781 | 1,025 | 0,829 | 0,899 | 0,938 | 0,745 | 0,895 | 0,817 | 1,04 | 0,04 | 6,3E-01 |
| O88531 | Palmitoyl-protein thioesterase 1 OS=Mus musculus OX=10090 G | 9 | 2 | 2 | 2 | 306 | 34,5 | 8 | 802 | 3,05 | 1 | 1 | 1,074 | 1,161 | 1,069 | 1,374 | 0,733 | 1,204 | 1,215 | 1,238 | 1,07 | 0,07 | 6,3E-01 |
| Q9ERS2 | NADH dehydrogenase [ubiquinone] 1 alpha subcomplex subunit | 38 | 5 | 17 | 5 | 144 | 16,8 | 9,48 | 10498 | | 5 | | 1,074 | 0,983 | 0,973 | 1,015 | 0,995 | 1,077 | 1,031 | 1,002 | 0,99 | -0,01 | 6,3E-01 |
| P70377 | Fibroblast growth factor 13 OS=Mus musculus OX=10090 GN=Fg | 4 | 1 | 1 | 1 | 245 | 27,6 | 9,91 | 543 | | 1 | | 0,782 | 0,949 | 1 | 0,879 | 1,391 | 0,739 | 1,01 | 0,788 | 0,92 | -0,08 | 6,3E-01 |
| Q9Z1G3 | V-type proton ATPase subunit C 1 OS=Mus musculus OX=10090 | 30 | 15 | 23 | 15 | 382 | 43,9 | 7,46 | 10329 | 10,04 | 12 | 3 | 1,017 | 0,926 | 0,864 | 0,945 | 1,015 | 0,946 | 0,924 | 0,943 | 0,98 | -0,02 | 6,3E-01 |
| O55128 | Histone deacetylase complex subunit SAP18 OS=Mus musculus | 5 | 1 | 1 | 1 | 153 | 17,6 | 9,35 | 534 | | 1 | | 0,906 | 0,658 | 0,877 | 1,054 | 0,847 | 0,864 | 0,952 | 1,017 | 0,95 | -0,05 | 6,3E-01 |
| O08848 | 60 kDa SS-A/Ro ribonucleoprotein OS=Mus musculus OX=10090 | 3 | 2 | 3 | 1 | 538 | 60,1 | 7,9 | 816 | 3,11 | 1 | 1 | 0,871 | 0,99 | 0,906 | 1,027 | 1,003 | 0,994 | 0,87 | 0,804 | 1,03 | 0,03 | 6,3E-01 |
| P11531 | Dystrophin OS=Mus musculus OX=10090 GN=Dmd PE=1 SV=3 | 0 | 2 | 2 | 2 | 3678 | 425,6 | 5,94 | 337 | 2,07 | 1 | 1 | 0,904 | 0,665 | 1,127 | 0,816 | 0,462 | 1,073 | 0,735 | 0,912 | 1,10 | 0,10 | 6,3E-01 |
| Q99NB1 | Acetyl-coenzyme A synthetase 2-like, mitochondrial OS=Mus mu | 5 | 3 | 3 | 3 | 682 | 74,6 | 6,98 | 902 | 2,19 | 2 | 1 | 0,879 | 0,923 | 0,916 | 1,02 | 0,989 | 0,996 | 0,901 | 0,929 | 0,98 | -0,02 | 6,3E-01 |
| P62281 | 40S ribosomal protein S11 OS=Mus musculus OX=10090 GN=Rp: | 52 | 10 | 19 | 10 | 158 | 18,4 | 10,3 | 6184 | 23,8 | 4 | 6 | 0,925 | 0,917 | 1,028 | 0,994 | 1,155 | 0,959 | 0,938 | 0,933 | 0,97 | -0,03 | 6,3E-01 |
| Q6ZPK7 | Lateral signaling target protein 2 homolog OS=Mus musculus OX | 2 | 1 | 2 | 1 | 905 | 99,7 | 5,1 | 307 | | 1 | | 0,858 | 1,287 | 0,657 | 0,973 | 1,153 | 0,727 | 1,059 | 1,175 | 0,92 | -0,08 | 6,3E-01 |
| Q8BMF4 | Dihydrolipoyllysine-residue acetyltransferase component of pyr | 18 | 12 | 19 | 12 | 642 | 67,9 | 8,57 | 10796 | 5,96 | 9 | 3 | 1,086 | 1,053 | 1,466 | 1,006 | 1,025 | 1,227 | 1,067 | 1,06 | 1,05 | 0,05 | 6,3E-01 |
| P97411 | Islet cell autoantigen 1 OS=Mus musculus OX=10090 GN=Ica1 PE | 6 | 3 | 5 | 2 | 478 | 54,3 | 5,8 | 2621 | | 3 | | 0,85 | 0,72 | 0,861 | 1,023 | 0,671 | 0,76 | 1,077 | 0,723 | 1,07 | 0,07 | 6,3E-01 |
| Q9WV55 | Vesicle-associated membrane protein-associated protein A OS=I | 33 | 8 | 16 | 7 | 249 | 27,8 | 8,4 | 8511 | 0 | 8 | 1 | 0,958 | 0,969 | 0,977 | 0,926 | 0,844 | 0,873 | 1,136 | 1,142 | 0,96 | -0,04 | 6,3E-01 |
| Q4VAE3 | Transmembrane protein 65 OS=Mus musculus OX=10090 GN=Tr | 11 | 2 | 5 | 2 | 234 | 24,9 | 7,78 | 1776 | 0 | 1 | 1 | 1,139 | 1,131 | 1,143 | 0,945 | 1,454 | 1,06 | 0,93 | 1,158 | 0,95 | -0,05 | 6,3E-01 |
| Q8BYW1 | Rho GTPase-activating protein 25 OS=Mus musculus OX=10090 | 2 | 1 | 1 | 1 | 648 | 73,3 | 6,38 | 282 | | 1 | | 0,665 | 1,096 | 1,005 | 1,074 | 1,125 | 0,752 | 1,508 | 0,85 | 0,91 | -0,09 | 6,3E-01 |
| Q5XJY5 | Coatomer subunit delta OS=Mus musculus OX=10090 GN=Arcn1 | 14 | 8 | 10 | 8 | 511 | 57,2 | 6,21 | 4938 | 0 | 7 | 1 | 1,121 | 1,026 | 0,97 | 1,001 | 1,08 | 1,003 | 1,058 | 1,05 | 0,98 | -0,02 | 6,3E-01 |
| Q6ZWV7 | 60S ribosomal protein L35 OS=Mus musculus OX=10090 GN=Rpl | 15 | 2 | 2 | 2 | 123 | 14,5 | 11,05 | 1527 | | 2 | | 1,084 | 1,019 | 0,984 | 1,057 | 0,891 | 1,148 | 0,874 | 1,086 | 1,04 | 0,04 | 6,3E-01 |
| Q6A0A9 | Constitutive coactivator of PPAR-gamma-like protein 1 OS=Mus | 7 | 7 | 7 | 7 | 1112 | 121,6 | 8,92 | 3048 | 3,5 | 6 | 1 | 0,871 | 0,771 | 0,9 | 0,93 | 1,005 | 0,773 | 0,964 | 0,856 | 0,96 | -0,04 | 6,3E-01 |
| Q8C015 | Serine/threonine-protein kinase PAK 5 OS=Mus musculus OX=1 | 3 | 2 | 3 | 2 | 719 | 80,9 | 8,21 | 1161 | 2,93 | 1 | 1 | 0,876 | 0,995 | 0,949 | 1,001 | 0,94 | 0,965 | 0,92 | 1,093 | 0,98 | -0,02 | 6,3E-01 |
| Q91V14 | Solute carrier family 12 member 5 OS=Mus musculus OX=10090 | 8 | 9 | 13 | 8 | 1138 | 126,2 | 6,74 | 6568 | 10,41 | 7 | 2 | 0,998 | 0,973 | 0,952 | 0,951 | 0,96 | 0,972 | 0,975 | 0,993 | 0,99 | -0,01 | 6,3E-01 |
| P37172 | Activin receptor type-1 OS=Mus musculus OX=10090 GN=Acvr1 | 2 | 1 | 1 | 1 | 509 | 57,2 | 7,33 | | 2,05 | | 1 | 0,861 | 0,944 | 0,896 | 0,855 | 1,103 | 0,775 | 0,873 | 0,586 | 1,07 | 0,07 | 6,3E-01 |
| Q69ZP3-2 | Isoform 2 of Probable hydrolase PNKD OS=Mus musculus OX=1 | 12 | 1 | 1 | 1 | 142 | 15,6 | 10,18 | 523 | | 1 | | 0,895 | 1,181 | 0,925 | 1,053 | 1,146 | 1,013 | 0,926 | 1,137 | 0,96 | -0,04 | 6,4E-01 |
| Q6PAJ1 | Breakpoint cluster region protein OS=Mus musculus OX=10090 | 3 | 4 | 5 | 3 | 1270 | 143 | 6,92 | 2873 | | 4 | | 1,19 | 0,893 | 1,226 | 1,069 | 0,868 | 1,22 | 0,985 | 1,093 | 1,05 | 0,05 | 6,4E-01 |
| P10922 | Histone H1.0 OS=Mus musculus OX=10090 GN=H1f0 PE=2 SV=4 | 14 | 3 | 4 | 3 | 194 | 20,8 | 10,9 | 618 | 8,6 | 1 | 2 | 0,939 | 0,884 | 1,095 | 1,234 | 0,939 | 0,778 | 1,132 | 1,08 | 1,06 | 0,06 | 6,4E-01 |
| Q9CQ62 | 2,4-dienoyl-CoA reductase, mitochondrial OS=Mus musculus OX | 17 | 4 | 6 | 4 | 335 | 36,2 | 8,95 | 2019 | 9,33 | 2 | 2 | 0,954 | 1,022 | 1,048 | 1,293 | 1,191 | 0,998 | 1,134 | 1,164 | 0,96 | -0,04 | 6,4E-01 |
| E9Q5G3 | Kinesin-like protein KIF23 OS=Mus musculus OX=10090 GN=Kif2 | 1 | 1 | 1 | 1 | 953 | 108,7 | 8,66 | 680 | | 1 | | 0,869 | 0,947 | 1,036 | 0,897 | 0,657 | 0,973 | 0,945 | 1 | 1,05 | 0,05 | 6,4E-01 |

| | | | | | | | | | | | | | | | | | | | | | | | |
|---|---|---|---|---|---|---|---|---|---|---|---|---|---|---|---|---|---|---|---|---|---|---|---|
| P59016 | Vacuolar protein sorting-associated protein 33B OS=Mus muscu | 5 | 2 | 2 | 2 | 617 | 70,5 | 6,86 | 654 | 2,83 | 1 | 1 | 1,382 | 1,164 | 1,05 | 1,2 | 1,082 | 1,296 | 1,16 | 1,09 | 1,04 | 0,04 | 6,4E-01 |
| Q8BKG3 | Inactive tyrosine-protein kinase 7 OS=Mus musculus OX=10090 | 1 | 1 | 1 | 1 | 1062 | 117,5 | 6,84 | | 2,45 | | 1 | 1,106 | 0,992 | 0,806 | 1,036 | 0,902 | 1,049 | 0,906 | 0,939 | 1,04 | 0,04 | 6,4E-01 |
| Q5DTY9 | BTB/POZ domain-containing protein KCTD16 OS=Mus musculus | 4 | 2 | 2 | 2 | 427 | 48,9 | 8,19 | 328 | 2,95 | 1 | 1 | 1,263 | 1,324 | 1,171 | 1,363 | 0,929 | 1,17 | 1,444 | 1,341 | 1,05 | 0,05 | 6,4E-01 |
| Q6URW6 | Myosin-14 OS=Mus musculus OX=10090 GN=Myh14 PE=1 SV=1 | 4 | 8 | 17 | 1 | 2000 | 228,4 | 5,55 | 4640 | 23,66 | 4 | 5 | 1,546 | 2,077 | 1,768 | 1,378 | 1,856 | 1,164 | 1,509 | 1,807 | 1,07 | 0,07 | 6,4E-01 |
| Q68FM6 | Protein phosphatase 1 regulatory subunit 29 OS=Mus musculus | 3 | 2 | 3 | 2 | 823 | 90 | 7,52 | 2031 | | 2 | | 0,879 | 0,812 | 1,153 | 0,911 | 0,95 | 1,071 | 0,978 | 0,916 | 0,96 | -0,04 | 6,4E-01 |
| Q8C166 | Copine-1 OS=Mus musculus OX=10090 GN=Cpne1 PE=1 SV=1 | 6 | 3 | 3 | 3 | 536 | 58,8 | 5,66 | 740 | 1,79 | 2 | 1 | 1,148 | 1,253 | 1,178 | 1,08 | 1,247 | 1,236 | 1,039 | 1,262 | 0,97 | -0,03 | 6,4E-01 |
| P19221 | Prothrombin OS=Mus musculus OX=10090 GN=F2 PE=1 SV=1 | 1 | 1 | 1 | 1 | 618 | 70,2 | 6,43 | 220 | | 1 | | 0,821 | 0,766 | 0,793 | 0,911 | 0,733 | 0,701 | 0,966 | 0,758 | 1,04 | 0,04 | 6,4E-01 |
| Q61735 | Leukocyte surface antigen CD47 OS=Mus musculus OX=10090 G | 8 | 2 | 5 | 2 | 303 | 33,1 | 8,63 | 3258 | | 2 | | 1,033 | 1,13 | 0,958 | 1,043 | 1,17 | 0,971 | 0,972 | 0,923 | 1,03 | 0,03 | 6,4E-01 |
| Q9DAW9 | Calponin-3 OS=Mus musculus OX=10090 GN=Cnn3 PE=1 SV=1 | 14 | 3 | 5 | 3 | 330 | 36,4 | 5,72 | 1793 | 5,08 | 2 | 2 | 1,099 | 1,112 | 1,223 | 1,154 | 1,165 | 0,982 | 0,923 | 1,329 | 1,04 | 0,04 | 6,4E-01 |
| Q61120 | SHC-transforming protein 3 OS=Mus musculus OX=10090 GN=Sh | 3 | 1 | 1 | 1 | 474 | 52,1 | 7,3 | 640 | | 1 | | 0,834 | 1,076 | 0,881 | 1,263 | 1,147 | 1,083 | 0,969 | 1,061 | 0,95 | -0,05 | 6,4E-01 |
| Q99L47 | Hsc70-interacting protein OS=Mus musculus OX=10090 GN=St13 | 23 | 9 | 23 | 9 | 371 | 41,6 | 5,26 | 8579 | 18,12 | 8 | 4 | 0,924 | 0,873 | 0,903 | 0,941 | 0,901 | 0,938 | 0,948 | 0,893 | 0,99 | -0,01 | 6,4E-01 |
| Q07076 | Annexin A7 OS=Mus musculus OX=10090 GN=Anxa7 PE=1 SV=2 | 13 | 6 | 6 | 6 | 463 | 49,9 | 6,18 | 2394 | 4,36 | 4 | 2 | 1,052 | 0,987 | 0,977 | 0,976 | 0,971 | 1,051 | 0,947 | 1,101 | 0,98 | -0,02 | 6,4E-01 |
| Q03137 | Ephrin type-A receptor 4 OS=Mus musculus OX=10090 GN=Epha | 9 | 8 | 10 | 6 | 986 | 109,7 | 6,51 | 3493 | 9,45 | 4 | 4 | 0,98 | 0,993 | 0,986 | 1,698 | 1,048 | 0,958 | 1,048 | 1,238 | 1,09 | 0,09 | 6,4E-01 |
| P05532 | Mast/stem cell growth factor receptor Kit OS=Mus musculus OX | 6 | 5 | 7 | 5 | 979 | 109,3 | 7,21 | 4079 | 2,49 | 4 | 1 | 0,973 | 0,981 | 0,894 | 0,943 | 0,988 | 0,901 | 1,006 | 0,955 | 0,98 | -0,02 | 6,4E-01 |
| Q925I1 | ATPase family AAA domain-containing protein 3 OS=Mus muscu | 10 | 7 | 8 | 7 | 591 | 66,7 | 9,29 | 3608 | 2,9 | 5 | 2 | 1,029 | 0,969 | 0,953 | 1,137 | 0,726 | 1,042 | 1,073 | 1,065 | 1,05 | 0,05 | 6,4E-01 |
| Q922H4 | Mannose-1-phosphate guanyltransferase alpha OS=Mus muscul | 8 | 3 | 3 | 3 | 420 | 46,2 | 7,62 | 1317 | 2,19 | 2 | 1 | 0,857 | 1,183 | 1,011 | 0,938 | 1,404 | 0,777 | 1,044 | 1,048 | 0,93 | -0,07 | 6,4E-01 |
| Q505D9 | Tripartite motif-containing protein 67 OS=Mus musculus OX=10 | 8 | 4 | 4 | 4 | 768 | 82,7 | 7,08 | | 12,8 | | 4 | 0,853 | 1,03 | 0,955 | 1,027 | 1,143 | 0,899 | 0,909 | 1,054 | 0,97 | -0,03 | 6,5E-01 |
| Q5GH67 | XK-related protein 4 OS=Mus musculus OX=10090 GN=Xkr4 PE= | 7 | 3 | 5 | 3 | 647 | 71,5 | 8,03 | 2001 | 2,49 | 2 | 1 | 0,891 | 0,982 | 1,135 | 0,992 | 0,742 | 1,034 | 0,947 | 1,098 | 1,05 | 0,05 | 6,5E-01 |
| P28658 | Ataxin-10 OS=Mus musculus OX=10090 GN=Atxn10 PE=1 SV=2 | 10 | 4 | 4 | 4 | 475 | 53,7 | 5,25 | 2634 | | 4 | | 0,842 | 0,85 | 0,897 | 0,836 | 0,823 | 0,666 | 1,023 | 0,765 | 1,05 | 0,05 | 6,5E-01 |
| P35436 | Glutamate receptor ionotropic, NMDA 2A OS=Mus musculus OX | 1 | 1 | 1 | 1 | 1464 | 165,3 | 7,01 | 392 | | 1 | | 1,177 | 0,894 | 0,891 | 0,893 | 1,836 | 0,778 | 0,931 | 0,814 | 0,88 | -0,12 | 6,5E-01 |
| Q9D7B6 | Isobutyryl-CoA dehydrogenase, mitochondrial OS=Mus musculu | 8 | 3 | 3 | 3 | 413 | 45 | 8,13 | | 6,19 | | 3 | 1,081 | 1,027 | 1,193 | 1,312 | 1,056 | 1,006 | 1,207 | 1,189 | 1,03 | 0,03 | 6,5E-01 |
| Q924C1 | Exportin-5 OS=Mus musculus OX=10090 GN=Xpo5 PE=1 SV=1 | 1 | 1 | 1 | 1 | 1204 | 136,9 | 5,82 | 341 | | 1 | | 0,856 | 0,817 | 0,786 | 0,77 | 0,493 | 0,785 | 0,978 | 0,777 | 1,06 | 0,06 | 6,5E-01 |
| O54865 | Guanylate cyclase soluble subunit beta-1 OS=Mus musculus OX= | 5 | 2 | 4 | 2 | 620 | 70,6 | 5,31 | 1936 | | 2 | | 0,962 | 0,927 | 1,115 | 1,019 | 1,04 | 0,934 | 1 | 0,958 | 1,02 | 0,02 | 6,5E-01 |
| Q9D0F3 | Protein ERGIC-53 OS=Mus musculus OX=10090 GN=Lman1 PE=1 | 3 | 2 | 2 | 2 | 517 | 57,8 | 6,34 | 1421 | | 2 | | 1,042 | 0,935 | 1,122 | 1,145 | 1,252 | 1 | 1,092 | 1,04 | 0,97 | -0,03 | 6,5E-01 |
| Q8R1T1 | Charged multivesicular body protein 7 OS=Mus musculus OX=10 | 3 | 1 | 2 | 1 | 451 | 50,6 | 5,22 | 1305 | | 1 | | 1,007 | 0,871 | 0,784 | 1,196 | 0,846 | 0,889 | 0,919 | 1,018 | 1,05 | 0,05 | 6,5E-01 |
| O35465 | Peptidyl-prolyl cis-trans isomerase FKBP8 OS=Mus musculus OX= | 16 | 5 | 7 | 5 | 402 | 43,5 | 5,16 | 3570 | 3,08 | 4 | 1 | 0,94 | 0,808 | 1,032 | 0,804 | 0,85 | 0,991 | 0,955 | 0,909 | 0,97 | -0,03 | 6,5E-01 |
| Q6P4S6 | Serine/threonine-protein kinase SIK3 OS=Mus musculus OX=100 | 1 | 1 | 1 | 1 | 1311 | 145,7 | 6,92 | | 0 | | 1 | 0,752 | 0,679 | 1,089 | 0,557 | 0,503 | 1,056 | 0,653 | 0,538 | 1,12 | 0,12 | 6,5E-01 |
| Q9D9V3 | Ethylmalonyl-CoA decarboxylase OS=Mus musculus OX=10090 G | 2 | 1 | 1 | 1 | 322 | 35,4 | 7,01 | 662 | | 1 | | 0,992 | 0,969 | 1,088 | 1,022 | 0,953 | 0,984 | 1,125 | 0,906 | 1,03 | 0,03 | 6,5E-01 |
| P06684 | Complement C5 OS=Mus musculus OX=10090 GN=C5 PE=1 SV=2 | 1 | 1 | 1 | 1 | 1680 | 188,8 | 6,81 | 228 | | 1 | | 0,933 | 1,308 | 0,51 | 1,244 | 0,763 | 0,783 | 0,865 | 1,187 | 1,11 | 0,11 | 6,5E-01 |
| P60670 | Nuclear protein localization protein 4 homolog OS=Mus musculu | 6 | 3 | 3 | 3 | 608 | 68 | 6,46 | 1013 | 2,68 | 2 | 1 | 1,002 | 1,129 | 0,882 | 1,024 | 1,163 | 0,832 | 1,177 | 1,046 | 0,96 | -0,04 | 6,5E-01 |
| Q9ESN6 | Tripartite motif-containing protein 2 OS=Mus musculus OX=100 | 19 | 11 | 18 | 8 | 744 | 81,4 | 6,96 | 6865 | 17,68 | 9 | 5 | 0,99 | 1,021 | 1,001 | 0,967 | 0,949 | 0,979 | 0,966 | 1,041 | 1,01 | 0,01 | 6,5E-01 |
| Q91W50 | Cold shock domain-containing protein E1 OS=Mus musculus OX= | 14 | 11 | 14 | 11 | 798 | 88,7 | 6,37 | 5623 | 10,27 | 8 | 4 | 1,042 | 0,954 | 0,988 | 0,967 | 0,926 | 1,029 | 1,063 | 1 | 0,98 | -0,02 | 6,5E-01 |
| O88643 | Serine/threonine-protein kinase PAK 1 OS=Mus musculus OX=10 | 30 | 15 | 29 | 10 | 545 | 60,7 | 5,74 | 16171 | 9,13 | 13 | 5 | 1,013 | 0,944 | 1,155 | 1,191 | 0,78 | 1,218 | 1,062 | 1,037 | 1,05 | 0,05 | 6,5E-01 |
| Q8VE73 | Cullin-7 OS=Mus musculus OX=10090 GN=Cul7 PE=1 SV=2 | 1 | 1 | 1 | 1 | 1689 | 192,2 | 6,29 | 448 | | 1 | | 0,893 | 0,911 | 1,195 | 0,735 | 1,374 | 0,747 | 0,957 | 0,966 | 0,92 | -0,08 | 6,5E-01 |
| P62317 | Small nuclear ribonucleoprotein Sm D2 OS=Mus musculus OX=1 | 22 | 3 | 4 | 3 | 118 | 13,5 | 9,91 | 1507 | 2,34 | 2 | 1 | 0,845 | 0,958 | 0,901 | 0,991 | 1,117 | 0,787 | 0,979 | 0,955 | 0,96 | -0,04 | 6,5E-01 |
| Q62188 | Dihydropyrimidinase-related protein 3 OS=Mus musculus OX=10 | 68 | 31 | 271 | 26 | 570 | 61,9 | 6,49 | 109258 | 220,98 | 17 | 24 | 1 | 0,941 | 1,038 | 0,957 | 0,841 | 1,05 | 0,981 | 0,971 | 1,02 | 0,02 | 6,5E-01 |
| Q5XF89 | Probable cation-transporting ATPase 13A3 OS=Mus musculus O | 1 | 1 | 1 | 1 | 1219 | 137,4 | 6,67 | 295 | | 1 | | 1,684 | 1,697 | 1,675 | 1,707 | 1,487 | 2,006 | 1,76 | 1,713 | 0,97 | -0,03 | 6,5E-01 |
| Q8BH44 | Coronin-2B OS=Mus musculus OX=10090 GN=Coro2b PE=1 SV=2 | 8 | 3 | 3 | 3 | 480 | 54,9 | 8,27 | 2304 | | 3 | | 0,891 | 1,124 | 1,037 | 0,933 | 1,253 | 0,922 | 0,95 | 1,034 | 0,96 | -0,04 | 6,5E-01 |
| Q9D051 | Pyruvate dehydrogenase E1 component subunit beta, mitochon | 40 | 10 | 26 | 10 | 359 | 38,9 | 6,87 | 9115 | 32,46 | 7 | 4 | 1,076 | 1,028 | 1,051 | 1,112 | 0,946 | 1,039 | 1,099 | 1,105 | 1,02 | 0,02 | 6,5E-01 |
| Q9Z2W0 | Aspartyl aminopeptidase OS=Mus musculus OX=10090 GN=Dnp | 15 | 6 | 9 | 6 | 473 | 52,2 | 7,25 | 2444 | 2,04 | 4 | 3 | 1,096 | 1,157 | 1,141 | 1,25 | 1,371 | 1,26 | 1,156 | 1,014 | 0,97 | -0,03 | 6,5E-01 |
| Q11011 | Puromycin-sensitive aminopeptidase OS=Mus musculus OX=100 | 27 | 22 | 32 | 22 | 920 | 103,3 | 5,9 | 15116 | 19,26 | 16 | 7 | 1,061 | 1,016 | 1,011 | 1,07 | 0,979 | 1,124 | 1,009 | 1,123 | 0,98 | -0,02 | 6,5E-01 |
| Q8BG39 | Synaptic vesicle glycoprotein 2B OS=Mus musculus OX=10090 G | 5 | 3 | 4 | 2 | 683 | 77,4 | 5,57 | 2350 | 0 | 2 | 1 | 0,941 | 1,001 | 0,814 | 0,969 | 0,975 | 0,922 | 0,923 | 0,801 | 1,03 | 0,03 | 6,5E-01 |
| Q9CXW2 | 28S ribosomal protein S22, mitochondrial OS=Mus musculus OX | 6 | 2 | 3 | 2 | 359 | 41,2 | 8,56 | 641 | 0 | 1 | 1 | 0,972 | 1,017 | 0,956 | 0,93 | 1,024 | 0,974 | 0,934 | 0,874 | 1,02 | 0,02 | 6,5E-01 |
| Q6PB66 | Leucine-rich PPR motif-containing protein, mitochondrial OS=M | 18 | 26 | 29 | 26 | 1392 | 156,5 | 6,83 | 11191 | 27,62 | 16 | 10 | 0,987 | 0,934 | 1,039 | 0,994 | 0,889 | 1,046 | 0,965 | 0,981 | 1,02 | 0,02 | 6,5E-01 |
| Q80XI3 | Eukaryotic translation initiation factor 4 gamma 3 OS=Mus musc | 2 | 3 | 3 | 3 | 1579 | 174,8 | 5,53 | 1867 | | 3 | | 0,989 | 1,031 | 0,978 | 1,028 | 1,003 | 0,944 | 0,887 | 1,102 | 1,02 | 0,02 | 6,5E-01 |
| P52503 | NADH dehydrogenase [ubiquinone] iron-sulfur protein 6, mitoch | 62 | 6 | 9 | 6 | 116 | 13 | 8,65 | 2843 | 10,4 | 3 | 3 | 0,991 | 0,967 | 0,929 | 0,884 | 0,905 | 1,06 | 0,866 | 1,041 | 0,97 | -0,03 | 6,6E-01 |
| Q5M8N0 | CB1 cannabinoid receptor-interacting protein 1 OS=Mus muscul | 21 | 3 | 9 | 3 | 164 | 18,6 | 7,96 | 4898 | | 3 | | 0,834 | 0,877 | 1,137 | 0,938 | 0,595 | 1,13 | 0,933 | 0,886 | 1,07 | 0,07 | 6,6E-01 |
| O35643 | AP-1 complex subunit beta-1 OS=Mus musculus OX=10090 GN=A | 16 | 18 | 31 | 5 | 943 | 103,9 | 5,17 | 12473 | 30,59 | 11 | 9 | 1,009 | 0,901 | 1,056 | 0,913 | 0,918 | 1,112 | 1,01 | 0,946 | 0,97 | -0,03 | 6,6E-01 |
| Q8VEF1 | Protein Aster-A OS=Mus musculus OX=10090 GN=Gramd1a PE= | 1 | 1 | 1 | 1 | 722 | 80,6 | 6,86 | 410 | | 1 | | 0,721 | 0,701 | 1,696 | 0,962 | 1,187 | 0,636 | 0,831 | 0,94 | 1,14 | 0,14 | 6,6E-01 |
| Q6PGF7 | Exocyst complex component 8 OS=Mus musculus OX=10090 GN | 11 | 8 | 8 | 8 | 716 | 81 | 5,4 | 2916 | 4,91 | 5 | 3 | 0,958 | 0,97 | 1,041 | 0,914 | 0,888 | 0,992 | 0,932 | 1,001 | 1,02 | 0,02 | 6,6E-01 |
| Q3TQB2 | FAD-dependent oxidoreductase domain-containing protein 1 OS | 5 | 2 | 2 | 2 | 487 | 54,1 | 7,53 | 1027 | | 2 | | 1,343 | 1,409 | 0,964 | 1,304 | 1,323 | 1,139 | 1,35 | 1,425 | 0,96 | -0,04 | 6,6E-01 |
| P03995 | Glial fibrillary acidic protein OS=Mus musculus OX=10090 GN=G | 16 | 7 | 8 | 5 | 430 | 49,9 | 5,34 | 3522 | 7,43 | 4 | 3 | 1,071 | 0,983 | 1,323 | 1,147 | 1,315 | 0,727 | 1,022 | 1,188 | 1,06 | 0,06 | 6,6E-01 |
| Q8R2Y0 | Monoacylglycerol lipase ABHD6 OS=Mus musculus OX=10090 G | 7 | 2 | 2 | 2 | 336 | 38,2 | 8,47 | 1671 | | 2 | | 0,932 | 0,996 | 0,878 | 0,963 | 0,729 | 0,869 | 0,876 | 1,132 | 1,05 | 0,05 | 6,6E-01 |
| Q6P069 | Sorcin OS=Mus musculus OX=10090 GN=Sri PE=1 SV=1 | 20 | 4 | 7 | 4 | 198 | 21,6 | 5,59 | 3478 | 2,4 | 3 | 1 | 1,023 | 1,076 | 1,319 | 1,036 | 1,515 | 0,808 | 0,96 | 0,839 | 1,08 | 0,08 | 6,6E-01 |
| Q04447 | Creatine kinase B-type OS=Mus musculus OX=10090 GN=Ckb PE | 58 | 22 | 94 | 18 | 381 | 42,7 | 5,67 | 49964 | 48,66 | 18 | 10 | 0,961 | 0,978 | 0,935 | 0,976 | 0,991 | 0,91 | 0,987 | 1,006 | 0,99 | -0,01 | 6,6E-01 |
| Q8BG94 | COMM domain-containing protein 7 OS=Mus musculus OX=100 | 6 | 1 | 1 | 1 | 200 | 22,6 | 5,94 | 665 | | 1 | | 1,177 | 1,318 | 1,242 | 1,153 | 0,933 | 1,148 | 1,481 | 1,105 | 1,05 | 0,05 | 6,6E-01 |
| Q61189 | Methylosome subunit pICln OS=Mus musculus OX=10090 GN=C | 6 | 1 | 2 | 1 | 236 | 26 | 4,12 | | 5,98 | | 1 | 0,976 | 1,066 | 0,861 | 1,049 | 0,752 | 1,016 | 1,006 | 1,028 | 1,04 | 0,04 | 6,6E-01 |
| Q99JB2 | Stomatin-like protein 2, mitochondrial OS=Mus musculus OX=10 | 15 | 4 | 4 | 4 | 353 | 38,4 | 8,87 | 2465 | | 4 | | 0,77 | 0,944 | 0,741 | 0,817 | 0,627 | 0,699 | 0,746 | 1,021 | 1,06 | 0,06 | 6,6E-01 |
| Q6NVE8 | WD repeat-containing protein 44 OS=Mus musculus OX=10090 | 3 | 2 | 2 | 2 | 915 | 101,5 | 5,24 | 1154 | | 2 | | 0,964 | 1,023 | 0,779 | 1,018 | 0,819 | 0,888 | 1,023 | 0,923 | 1,04 | 0,04 | 6,6E-01 |
| Q99MN1 | Lysine--tRNA ligase OS=Mus musculus OX=10090 GN=Kars PE=1 | 8 | 4 | 4 | 4 | 595 | 67,8 | 5,94 | 2446 | | 4 | | 0,965 | 0,987 | 1,122 | 1,125 | 1,202 | 1,06 | 1,012 | 1,036 | 0,97 | -0,03 | 6,6E-01 |
| P62858 | 40S ribosomal protein S28 OS=Mus musculus OX=10090 GN=Rp | 13 | 1 | 3 | 1 | 69 | 7,8 | 10,7 | 1922 | | 1 | | 0,872 | 0,857 | 0,974 | 0,956 | 0,895 | 0,894 | 0,879 | 0,933 | 1,02 | 0,02 | 6,6E-01 |
| Q9QUR7 | Peptidyl-prolyl cis-trans isomerase NIMA-interacting 1 OS=Mus | 21 | 3 | 4 | 3 | 165 | 18,4 | 8,79 | 976 | 0 | 2 | 1 | 1,026 | 1,031 | 0,984 | 1,036 | 0,986 | 1,083 | 0,972 | 1,1 | 0,98 | -0,02 | 6,6E-01 |
| P63213 | Guanine nucleotide-binding protein G(I)/G(S)/G(O) subunit gam | 65 | 5 | 14 | 5 | 71 | 7,8 | 7,99 | 7520 | 5,46 | 3 | 2 | 0,991 | 1,015 | 1,018 | 1,074 | 0,753 | 1,108 | 1,085 | 1 | 1,04 | 0,04 | 6,6E-01 |
| Q62426 | Cystatin-B OS=Mus musculus OX=10090 GN=Cstb PE=1 SV=1 | 40 | 3 | 4 | 3 | 98 | 11 | 7,39 | 1909 | 3,15 | 3 | 1 | 1,015 | 1,093 | 1,094 | 1,039 | 1,044 | 1,085 | 0,989 | 1,22 | 0,98 | -0,02 | 6,6E-01 |
| Q8BH74 | Nuclear pore complex protein Nup107 OS=Mus musculus OX=10 | 1 | 1 | 1 | 1 | 926 | 106,7 | 5,43 | 367 | | 1 | | 1,005 | 1,096 | 1,356 | 0,916 | 1,82 | 0,917 | 1,161 | 0,904 | 0,91 | -0,09 | 6,6E-01 |

| | | | | | | | | | | | | | | | | | | | | | | | |
|---|---|---|---|---|---|---|---|---|---|---|---|---|---|---|---|---|---|---|---|---|---|---|---|
| P97807 | Fumarate hydratase, mitochondrial OS=Mus musculus OX=1009 | 28 | 15 | 29 | 15 | 507 | 54,3 | 9,04 | 10303 | 14,73 | 7 | 8 | 1,011 | 0,963 | 1,108 | 0,956 | 1,122 | 1,029 | 1 | 0,974 | 0,98 | -0,02 | 6,6E-01 |
| Q8VCD6 | Receptor expression-enhancing protein 2 OS=Mus musculus OX | 17 | 5 | 6 | 5 | 254 | 28,4 | 9,41 | 2642 | 0 | 4 | 1 | 1,091 | 1,125 | 1,04 | 1,117 | 1,017 | 1,121 | 1,105 | 1,21 | 0,98 | -0,02 | 6,6E-01 |
| Q3TWN3 | Metal transporter CNNM2 OS=Mus musculus OX=10090 GN=Cn | 1 | 1 | 1 | 1 | 875 | 96,6 | 6,21 | 643 | | 1 | | 1,634 | 1,305 | 1,154 | 1,021 | 1,839 | 1,655 | 1,058 | 1,012 | 0,92 | -0,08 | 6,7E-01 |
| Q91XL9 | Oxysterol-binding protein-related protein 1 OS=Mus musculus C | 6 | 5 | 10 | 4 | 950 | 107,7 | 6,44 | 3957 | 7,43 | 4 | 2 | 0,987 | 0,952 | 0,878 | 1,052 | 0,846 | 0,951 | 1,025 | 0,953 | 1,02 | 0,02 | 6,7E-01 |
| P08228 | Superoxide dismutase [Cu-Zn] OS=Mus musculus OX=10090 GN | 63 | 13 | 44 | 13 | 154 | 15,9 | 6,51 | 20646 | 27,61 | 9 | 7 | 0,985 | 1,071 | 1,207 | 1,071 | 1,044 | 1,514 | 1,006 | 1,01 | 0,95 | -0,05 | 6,7E-01 |
| P62806 | Histone H4 OS=Mus musculus OX=10090 GN=Hist1h4a PE=1 SV | 32 | 4 | 12 | 4 | 103 | 11,4 | 11,36 | 6431 | 2,92 | 3 | 1 | 0,952 | 0,866 | 1,192 | 1,462 | 1,126 | 0,886 | 1,192 | 0,995 | 1,07 | 0,07 | 6,7E-01 |
| Q9DBE8 | Alpha-1,3/1,6-mannosyltransferase ALG2 OS=Mus musculus OX | 7 | 2 | 2 | 2 | 415 | 47,4 | 7,97 | 683 | 2,5 | 1 | 1 | 1,376 | 0,974 | 1,356 | 0,837 | 0,895 | 1,313 | 1,09 | 0,945 | 1,07 | 0,07 | 6,7E-01 |
| P15208 | Insulin receptor OS=Mus musculus OX=10090 GN=Insr PE=1 SV= | 3 | 4 | 5 | 4 | 1372 | 155,5 | 5,95 | 2251 | 0 | 3 | 1 | 0,952 | 0,927 | 0,931 | 1,053 | 0,813 | 1,072 | 1,19 | 0,945 | 0,96 | -0,04 | 6,7E-01 |
| Q3U0V1 | Far upstream element-binding protein 2 OS=Mus musculus OX= | 16 | 8 | 11 | 7 | 748 | 76,7 | 7,33 | 4266 | 2,98 | 7 | 2 | 1,03 | 0,931 | 0,969 | 1,081 | 1,474 | 0,902 | 0,924 | 0,966 | 0,94 | -0,06 | 6,7E-01 |
| Q9D5T0 | ATPase family AAA domain-containing protein 1 OS=Mus muscu | 12 | 4 | 4 | 4 | 361 | 40,7 | 6,9 | 1950 | 2,74 | 3 | 1 | 1,227 | 1,007 | 1,142 | 1,159 | 1,126 | 1,131 | 1,168 | 1,199 | 0,98 | -0,02 | 6,7E-01 |
| P58802 | TBC1 domain family member 10A OS=Mus musculus OX=10090 | 6 | 2 | 2 | 2 | 500 | 56,2 | 7,85 | 467 | 2,97 | 1 | 1 | 0,841 | 0,967 | 0,943 | 0,947 | 0,557 | 0,817 | 0,816 | 1,245 | 1,08 | 0,08 | 6,7E-01 |
| Q99PU5 | Long-chain-fatty-acid--CoA ligase ACSBG1 OS=Mus musculus OX | 13 | 9 | 13 | 9 | 721 | 80,4 | 5,94 | 6472 | 6,9 | 9 | 1 | 0,95 | 0,988 | 0,987 | 1,008 | 1,021 | 0,94 | 0,984 | 0,948 | 1,01 | 0,01 | 6,7E-01 |
| Q2TPA8 | Hydroxysteroid dehydrogenase-like protein 2 OS=Mus musculus | 6 | 3 | 3 | 3 | 490 | 54,2 | 6,74 | 1193 | 1,96 | 2 | 1 | 0,842 | 0,867 | 0,937 | 0,813 | 1,02 | 0,922 | 0,781 | 0,841 | 0,97 | -0,03 | 6,7E-01 |
| O08532-5 | Isoform 2E of Voltage-dependent calcium channel subunit alpha | 22 | 22 | 33 | 21 | 1091 | 123,2 | 5,27 | 12756 | 20,18 | 14 | 9 | 0,981 | 0,915 | 0,92 | 0,923 | 0,929 | 0,927 | 0,984 | 0,936 | 0,99 | -0,01 | 6,7E-01 |
| Q6GQT9 | Nodal modulator 1 OS=Mus musculus OX=10090 GN=Nomo1 PE | 3 | 3 | 3 | 3 | 1214 | 133,3 | 6,09 | 1126 | 2,18 | 2 | 1 | 0,903 | 0,977 | 1,013 | 1,019 | 1,078 | 0,868 | 0,943 | 0,93 | 1,02 | 0,02 | 6,7E-01 |
| O35344 | Importin subunit alpha-4 OS=Mus musculus OX=10090 GN=Kpn | 11 | 5 | 6 | 5 | 521 | 57,7 | 4,94 | 1795 | 5,05 | 3 | 2 | 1,077 | 1,088 | 0,928 | 1,189 | 1,104 | 1,014 | 0,961 | 1,089 | 1,03 | 0,03 | 6,7E-01 |
| Q6PHS9 | Voltage-dependent calcium channel subunit alpha-2/delta-2 OS | 5 | 8 | 9 | 7 | 1154 | 130,3 | 5,66 | 2982 | 8,94 | 4 | 4 | 0,949 | 0,967 | 0,94 | 1,032 | 1,011 | 0,986 | 0,872 | 0,953 | 1,02 | 0,02 | 6,7E-01 |
| Q9EQJ9 | Membrane-associated guanylate kinase, WW and PDZ domain-c | 1 | 1 | 1 | 1 | 1476 | 161,6 | 8,05 | 378 | | 1 | | 0,984 | 1,328 | 0,752 | 1,165 | 0,785 | 1,099 | 0,899 | 1,172 | 1,07 | 0,07 | 6,7E-01 |
| Q9CYG7 | Mitochondrial import receptor subunit TOM34 OS=Mus muscul | 4 | 1 | 1 | 1 | 309 | 34,3 | 9,14 | | 2,34 | | 1 | 1,272 | 0,973 | 1,202 | 1,06 | 1,113 | 1,197 | 1,313 | 1,044 | 0,97 | -0,03 | 6,7E-01 |
| Q9CQ76 | Nephrocan OS=Mus musculus OX=10090 GN=Nepn PE=1 SV=1 | 3 | 1 | 1 | 1 | 512 | 57,7 | 8,24 | | 0 | | 1 | 0,918 | 0,757 | 0,968 | 1,08 | 0,98 | 0,882 | 0,894 | 0,835 | 1,04 | 0,04 | 6,7E-01 |
| Q8VDJ3 | Vigilin OS=Mus musculus OX=10090 GN=Hdlbp PE=1 SV=1 | 5 | 7 | 7 | 7 | 1268 | 141,7 | 6,87 | 3127 | 4,18 | 5 | 2 | 1,272 | 1,352 | 1,27 | 1,266 | 1,605 | 1,233 | 1,119 | 1,395 | 0,96 | -0,04 | 6,7E-01 |
| P70288 | Histone deacetylase 2 OS=Mus musculus OX=10090 GN=Hdac2 | 2 | 1 | 1 | 1 | 488 | 55,3 | 5,91 | 335 | | 1 | | 0,813 | 0,636 | 0,882 | 0,796 | 1,07 | 0,818 | 0,704 | 0,713 | 0,95 | -0,05 | 6,7E-01 |
| Q3TPX4 | Exocyst complex component 5 OS=Mus musculus OX=10090 GN | 5 | 3 | 5 | 3 | 708 | 81,7 | 6,71 | 1688 | 2,47 | 3 | 1 | 0,968 | 1,131 | 0,983 | 1,119 | 1,158 | 0,956 | 1,142 | 1,058 | 0,97 | -0,03 | 6,7E-01 |
| P61022 | Calcineurin B homologous protein 1 OS=Mus musculus OX=100 | 12 | 2 | 2 | 2 | 195 | 22,4 | 5,1 | 572 | 2,89 | 1 | 1 | 0,881 | 0,944 | 0,887 | 0,962 | 0,804 | 0,888 | 1,225 | 0,924 | 0,96 | -0,04 | 6,7E-01 |
| Q8K070 | Sterile alpha motif domain-containing protein 14 OS=Mus musc | 6 | 2 | 3 | 2 | 417 | 45,1 | 9,47 | 1164 | | 2 | | 1,004 | 1,094 | 1,005 | 0,991 | 1,434 | 0,865 | 0,868 | 1,174 | 0,94 | -0,06 | 6,7E-01 |
| P46735 | Unconventional myosin-Ib OS=Mus musculus OX=10090 GN=My | 6 | 6 | 8 | 6 | 1107 | 128,5 | 9,26 | 3886 | 2,57 | 5 | 1 | 0,923 | 0,847 | 0,816 | 0,966 | 1,091 | 0,845 | 0,834 | 0,904 | 0,97 | -0,03 | 6,7E-01 |
| Q8CCK0 | Core histone macro-H2A.2 OS=Mus musculus OX=10090 GN=H2 | 17 | 5 | 5 | 3 | 372 | 40,1 | 9,69 | 2513 | 2,66 | 4 | 1 | 0,818 | 0,889 | 1,296 | 0,996 | 0,987 | 0,794 | 1,379 | 1,126 | 0,93 | -0,07 | 6,7E-01 |
| P49586 | Choline-phosphate cytidylyltransferase A OS=Mus musculus OX | 5 | 2 | 2 | 1 | 367 | 41,6 | 7,03 | 1086 | | 2 | | 0,84 | 0,757 | 0,821 | 0,886 | 0,946 | 0,705 | 0,97 | 0,803 | 0,96 | -0,04 | 6,7E-01 |
| P26231 | Catenin alpha-1 OS=Mus musculus OX=10090 GN=Ctnna1 PE=1 | 9 | 8 | 11 | 2 | 906 | 100 | 6,23 | 6083 | 2,97 | 7 | 1 | 0,983 | 1,089 | 0,816 | 0,921 | 0,725 | 0,928 | 1,07 | 0,925 | 1,04 | 0,04 | 6,7E-01 |
| Q8BFQ4 | WD repeat-containing protein 82 OS=Mus musculus OX=10090 | 5 | 2 | 2 | 2 | 313 | 35,1 | 7,69 | 977 | | 2 | | 0,799 | 0,708 | 0,903 | 1,078 | 0,749 | 0,917 | 1,074 | 0,931 | 0,95 | -0,05 | 6,7E-01 |
| P21460 | Cystatin-C OS=Mus musculus OX=10090 GN=Cst3 PE=1 SV=2 | 25 | 4 | 7 | 4 | 140 | 15,5 | 9 | 1725 | 7,5 | 2 | 3 | 0,873 | 0,908 | 0,978 | 0,947 | 0,85 | 0,956 | 0,958 | 0,879 | 1,02 | 0,02 | 6,7E-01 |
| Q9EQ06 | Estradiol 17-beta-dehydrogenase 11 OS=Mus musculus OX=100 | 6 | 1 | 2 | 1 | 298 | 32,9 | 8,66 | 1262 | | 1 | | 1,171 | 1,101 | 1,148 | 1,156 | 1,21 | 1,004 | 1,069 | 1,2 | 1,02 | 0,02 | 6,7E-01 |
| Q3TCN2 | Putative phospholipase B-like 2 OS=Mus musculus OX=10090 G | 5 | 3 | 4 | 3 | 594 | 66,2 | 6,13 | 1362 | | 3 | | 1,013 | 1,097 | 1,095 | 1,221 | 1,335 | 1,017 | 0,968 | 0,925 | 1,04 | 0,04 | 6,7E-01 |
| P48678 | Prelamin-A/C OS=Mus musculus OX=10090 GN=Lmna PE=1 SV= | 18 | 13 | 14 | 13 | 665 | 74,2 | 6,98 | 3103 | 18,07 | 6 | 7 | 0,79 | 1,259 | 0,854 | 0,761 | 1,013 | 0,767 | 0,848 | 1,336 | 0,92 | -0,08 | 6,8E-01 |
| Q9WV92 | Band 4.1-like protein 3 OS=Mus musculus OX=10090 GN=Epb41 | 22 | 18 | 26 | 1 | 929 | 103,3 | 5,31 | 13025 | 13,21 | 13 | 5 | 1,321 | 1,084 | 1,081 | 1,21 | 1,02 | 1,332 | 1,059 | 1,127 | 1,03 | 0,03 | 6,8E-01 |
| Q80SW1 | S-adenosylhomocysteine hydrolase-like protein 1 OS=Mus musc | 26 | 12 | 16 | 12 | 530 | 58,9 | 6,89 | 5269 | 13,12 | 8 | 4 | 1,018 | 0,989 | 1,087 | 1,037 | 0,794 | 1,058 | 1,044 | 1,107 | 1,03 | 0,03 | 6,8E-01 |
| Q9CRY7 | Lysophospholipase D GDPD1 OS=Mus musculus OX=10090 GN= | 12 | 4 | 4 | 4 | 314 | 35,8 | 8,31 | 1737 | | 4 | | 1,107 | 1,102 | 1,227 | 1,022 | 1,169 | 1,031 | 1,088 | 1,081 | 1,02 | 0,02 | 6,8E-01 |
| P20357 | Microtubule-associated protein 2 OS=Mus musculus OX=10090 | 38 | 65 | 131 | 65 | 1828 | 199 | 4,91 | 53621 | 131,59 | 38 | 32 | 1,062 | 0,966 | 0,95 | 0,965 | 0,938 | 1,033 | 0,953 | 0,961 | 1,01 | 0,01 | 6,8E-01 |
| Q9D6Z0 | Alpha-ketoglutarate-dependent dioxygenase alkB homolog 7, m | 10 | 1 | 1 | 1 | 221 | 25 | 6,54 | | 0 | | 1 | 1,105 | 1,159 | 1,066 | 1,19 | 1,068 | 0,91 | 1,638 | 1,181 | 0,94 | -0,06 | 6,8E-01 |
| Q99KB8 | Hydroxyacylglutathione hydrolase, mitochondrial OS=Mus musc | 9 | 2 | 3 | 2 | 309 | 34,1 | 7,75 | 1443 | 3,8 | 2 | 1 | 1,07 | 1,143 | 1,038 | 1,025 | 1,101 | 1,092 | 0,962 | 1,049 | 1,02 | 0,02 | 6,8E-01 |
| Q60960 | Importin subunit alpha-5 OS=Mus musculus OX=10090 GN=Kpn | 3 | 1 | 1 | 1 | 538 | 60,1 | 5,01 | 175 | | 1 | | 0,543 | 1,191 | 0,273 | 0,63 | 0,255 | 0,473 | 1,383 | 1,094 | 0,82 | -0,18 | 6,8E-01 |
| Q99KP6 | Pre-mRNA-processing factor 19 OS=Mus musculus OX=10090 G | 10 | 5 | 8 | 5 | 504 | 55,2 | 6,61 | 2856 | 4,54 | 3 | 2 | 1,037 | 1,003 | 1,077 | 1,039 | 1,218 | 0,943 | 1,067 | 1,031 | 0,98 | -0,02 | 6,8E-01 |
| Q9WTX8 | Mitotic spindle assembly checkpoint protein MAD1 OS=Mus mu | 1 | 1 | 2 | 1 | 717 | 83,5 | 5,66 | 844 | | 1 | | 1,572 | 1,054 | 0,96 | 1,362 | 2,199 | 0,647 | 1,822 | 0,955 | 0,88 | -0,12 | 6,8E-01 |
| Q8BKS9 | Pumilio homolog 3 OS=Mus musculus OX=10090 GN=Pum3 PE= | 1 | 1 | 1 | 1 | 647 | 72,8 | 9,66 | 693 | | 1 | | 0,94 | 0,834 | 1,015 | 1,059 | 0,826 | 1,015 | 1,043 | 1,097 | 0,97 | -0,03 | 6,8E-01 |
| P70404 | Isocitrate dehydrogenase [NAD] subunit gamma 1, mitochondri | 18 | 5 | 11 | 5 | 393 | 42,8 | 9,01 | 6560 | | 5 | | 1,131 | 1,135 | 1,171 | 1,172 | 1,358 | 1,076 | 1,065 | 1,232 | 0,97 | -0,03 | 6,8E-01 |
| Q61245 | Collagen alpha-1(XI) chain OS=Mus musculus OX=10090 GN=Col | 0 | 1 | 1 | 1 | 1804 | 180,9 | 5,21 | 462 | | 1 | | 1,023 | 0,893 | 1,054 | 0,892 | 1,244 | 1,349 | 0,693 | 0,855 | 0,93 | -0,07 | 6,8E-01 |
| Q9CZX8 | 40S ribosomal protein S19 OS=Mus musculus OX=10090 GN=Rp | 40 | 7 | 11 | 7 | 145 | 16,1 | 10,4 | 3115 | 10,63 | 5 | 3 | 1,075 | 0,821 | 0,941 | 1,048 | 1,469 | 0,902 | 0,89 | 0,891 | 0,94 | -0,06 | 6,8E-01 |
| P54869 | Hydroxymethylglutaryl-CoA synthase, mitochondrial OS=Mus m | 12 | 6 | 8 | 5 | 508 | 56,8 | 8,41 | 3281 | 5,41 | 4 | 2 | 1,11 | 1,29 | 1,286 | 1,639 | 1,708 | 1,145 | 1,434 | 1,317 | 0,95 | -0,05 | 6,8E-01 |
| P63323 | 40S ribosomal protein S12 OS=Mus musculus OX=10090 GN=Rp | 8 | 1 | 5 | 1 | 132 | 14,5 | 7,24 | 2896 | | 1 | | 1,016 | 0,842 | 0,896 | 1,033 | 0,866 | 0,892 | 1,052 | 1,106 | 0,97 | -0,03 | 6,8E-01 |
| Q8VDP3 | [F-actin]-monooxygenase MICAL1 OS=Mus musculus OX=10090 | 1 | 1 | 1 | 1 | 1048 | 116,7 | 6,05 | 429 | | 1 | | 1,035 | 1,334 | 0,996 | 1,573 | 1,582 | 1,026 | 1,271 | 1,365 | 0,94 | -0,06 | 6,8E-01 |
| Q9Z0R6 | Intersectin-2 OS=Mus musculus OX=10090 GN=Itsn2 PE=1 SV=2 | 2 | 3 | 3 | 1 | 1659 | 188,8 | 8,13 | 1331 | 3,34 | 2 | 1 | 1,215 | 1,142 | 1,069 | 1,018 | 0,58 | 1,414 | 1,082 | 1,064 | 1,07 | 0,07 | 6,8E-01 |
| O54774 | AP-3 complex subunit delta-1 OS=Mus musculus OX=10090 GN= | 12 | 14 | 17 | 14 | 1199 | 135 | 7,37 | 6094 | 12,31 | 10 | 4 | 1,033 | 1,035 | 1,065 | 1,06 | 0,922 | 1,065 | 1,092 | 1,048 | 1,02 | 0,02 | 6,8E-01 |
| Q99M80 | Receptor-type tyrosine-protein phosphatase T OS=Mus musculu | 1 | 1 | 1 | 1 | 1454 | 162,9 | 6,84 | 474 | | 1 | | 1,101 | 1,171 | 1,002 | 0,954 | 1,175 | 1,204 | 1,02 | 0,96 | 0,97 | -0,03 | 6,8E-01 |
| Q9WTS4 | Teneurin-1 OS=Mus musculus OX=10090 GN=Tenm1 PE=1 SV=1 | 0 | 1 | 1 | 1 | 2731 | 305,6 | 6,37 | 430 | | 1 | | 0,95 | 0,716 | 0,973 | 0,784 | 0,536 | 0,86 | 0,984 | 0,848 | 1,06 | 0,06 | 6,8E-01 |
| Q8CI95 | Oxysterol-binding protein-related protein 11 OS=Mus musculus | 2 | 1 | 1 | 1 | 751 | 83,6 | 7,01 | 370 | | 1 | | 0,946 | 1,031 | 1,14 | 1,085 | 1,046 | 1,211 | 1,069 | 0,984 | 0,97 | -0,03 | 6,8E-01 |
| O55022 | Membrane-associated progesterone receptor component 1 OS= | 24 | 7 | 16 | 6 | 195 | 21,7 | 4,7 | 7956 | 2,16 | 5 | 2 | 0,982 | 1,119 | 0,925 | 1,038 | 1 | 0,926 | 1,084 | 0,963 | 1,02 | 0,02 | 6,9E-01 |
| Q9CQQ7 | ATP synthase F(0) complex subunit B1, mitochondrial OS=Mus m | 30 | 10 | 20 | 10 | 256 | 28,9 | 9,06 | 7259 | 24,43 | 6 | 5 | 1,011 | 0,922 | 1,086 | 0,925 | 0,932 | 0,994 | 0,907 | 1,029 | 1,02 | 0,02 | 6,9E-01 |
| P12382 | ATP-dependent 6-phosphofructokinase, liver type OS=Mus mus | 17 | 10 | 19 | 5 | 780 | 85,3 | 7,17 | 6683 | 10,74 | 7 | 4 | 1,01 | 1,071 | 0,946 | 1,126 | 0,947 | 1,035 | 1,068 | 1,024 | 1,02 | 0,02 | 6,9E-01 |
| Q62417 | Sorbin and SH3 domain-containing protein 1 OS=Mus musculus | 1 | 1 | 1 | 1 | 1290 | 143 | 8,25 | | 2,21 | | 1 | 1,044 | 1,341 | 1,147 | 1,42 | 0,928 | 1,639 | 0,804 | 1,233 | 1,08 | 0,08 | 6,9E-01 |
| P08030 | Adenine phosphoribosyltransferase OS=Mus musculus OX=1009 | 22 | 3 | 3 | 3 | 180 | 19,7 | 6,79 | 1609 | | 3 | | 1,095 | 1,113 | 1,171 | 1,163 | 1,291 | 1,054 | 1,015 | 1,072 | 1,02 | 0,02 | 6,9E-01 |
| Q9CR70 | EKC/KEOPS complex subunit Lage3 OS=Mus musculus OX=1009 | 8 | 1 | 1 | 1 | 148 | 15,8 | 8,13 | 539 | | 1 | | 1,027 | 0,926 | 0,946 | 1,24 | 0,98 | 1,111 | 1,152 | 1,034 | 0,97 | -0,03 | 6,9E-01 |
| P61922 | 4-aminobutyrate aminotransferase, mitochondrial OS=Mus mus | 27 | 12 | 22 | 12 | 500 | 56,4 | 8,09 | 9229 | 20,67 | 9 | 5 | 1,009 | 0,883 | 0,957 | 0,887 | 0,968 | 0,982 | 0,903 | 0,942 | 0,98 | -0,02 | 6,9E-01 |
| P68040 | Receptor of activated protein C kinase 1 OS=Mus musculus OX= | 19 | 5 | 6 | 5 | 317 | 35,1 | 7,69 | 3832 | | 5 | | 0,862 | 0,88 | 0,952 | 1,163 | 1,005 | 0,78 | 1,031 | 0,889 | 1,04 | 0,04 | 6,9E-01 |
| Q91V36 | Nuclear receptor-binding protein 2 OS=Mus musculus OX=1009 | 7 | 4 | 6 | 4 | 499 | 57,3 | 6,35 | 1519 | 6,83 | 2 | 2 | 1,034 | 0,978 | 1,354 | 0,963 | 1,053 | 1,061 | 1,016 | 1,043 | 1,04 | 0,04 | 6,9E-01 |
| Q923D2 | Flavin reductase (NADPH) OS=Mus musculus OX=10090 GN=Blv | 13 | 2 | 2 | 2 | 206 | 22,2 | 7,01 | 1530 | | 2 | | 1,214 | 0,903 | 0,992 | 1,228 | 1,384 | 1,057 | 0,989 | 1,107 | 0,96 | -0,04 | 6,9E-01 |

| | | | | | | | | | | | | | | | | | | | | | | | |
|---|---|---|---|---|---|---|---|---|---|---|---|---|---|---|---|---|---|---|---|---|---|---|---|
| P62823 | Ras-related protein Rab-3C OS=Mus musculus OX=10090 GN=Ra | 34 | 7 | 18 | 4 | 227 | 25,9 | 5,24 | 10188 | 1,71 | 6 | 1 | 0,851 | 1,136 | 0,969 | 0,939 | 0,833 | 0,974 | 0,992 | 0,978 | 1,03 | 0,03 | 6,9E-01 |
| Q7M6Y3 | Phosphatidylinositol-binding clathrin assembly protein OS=Mus | 8 | 5 | 5 | 4 | 660 | 71,5 | 7,91 | 675 | 6,55 | 1 | 4 | 1,014 | 0,922 | 1,038 | 0,86 | 0,464 | 1,109 | 0,972 | 1,034 | 1,07 | 0,07 | 6,9E-01 |
| O70161 | Phosphatidylinositol 4-phosphate 5-kinase type-1 gamma OS=M | 15 | 7 | 8 | 2 | 661 | 72,4 | 5,59 | 2458 | 16,8 | 4 | 3 | 1,072 | 1,39 | 1,337 | 1,353 | 1,537 | 1,013 | 1,055 | 1,309 | 1,05 | 0,05 | 6,9E-01 |
| Q3UHD9 | Arf-GAP with GTPase, ANK repeat and PH domain-containing pr | 12 | 8 | 11 | 8 | 1186 | 124,4 | 9,89 | 4221 | 5,71 | 6 | 3 | 0,916 | 0,911 | 1,159 | 1,036 | 1,178 | 0,893 | 1,076 | 1,015 | 0,97 | -0,03 | 6,9E-01 |
| Q8BIW1 | Exopolyphosphatase PRUNE1 OS=Mus musculus OX=10090 GN= | 17 | 7 | 9 | 7 | 454 | 50,2 | 5,11 | 2560 | 6,77 | 4 | 3 | 0,998 | 0,892 | 1,057 | 0,897 | 0,933 | 0,996 | 0,965 | 1,025 | 0,98 | -0,02 | 6,9E-01 |
| P11438 | Lysosome-associated membrane glycoprotein 1 OS=Mus muscu | 5 | 2 | 4 | 2 | 406 | 43,8 | 8,4 | 1151 | 2,83 | 2 | 1 | 1,014 | 0,989 | 0,876 | 0,895 | 0,934 | 1,039 | 0,983 | 0,895 | 0,98 | -0,02 | 6,9E-01 |
| Q03958 | Prefoldin subunit 6 OS=Mus musculus OX=10090 GN=Pfdn6 PE= | 24 | 3 | 4 | 3 | 127 | 14,4 | 8,88 | 1910 | 0 | 2 | 1 | 0,84 | 0,938 | 0,909 | 0,851 | 0,92 | 0,84 | 0,901 | 0,928 | 0,99 | -0,01 | 6,9E-01 |
| Q8K2C6 | NAD-dependent protein deacylase sirtuin-5, mitochondrial OS=I | 15 | 3 | 3 | 3 | 310 | 34,1 | 8,53 | 1815 | | 3 | | 1,052 | 1,091 | 0,876 | 1,011 | 1,302 | 0,824 | 1,088 | 0,998 | 0,96 | -0,04 | 6,9E-01 |
| Q8BGH7 | CDC42 small effector protein 2 OS=Mus musculus OX=10090 GN | 23 | 1 | 1 | 1 | 84 | 9,2 | 8,35 | 563 | | 1 | | 1,276 | 1,391 | 1,147 | 1,271 | 1,337 | 1,197 | 1,329 | 1,096 | 1,03 | 0,03 | 6,9E-01 |
| O35453 | Serine protease hepsin OS=Mus musculus OX=10090 GN=Hpn P | 3 | 1 | 1 | 1 | 436 | 46,8 | 7,43 | 273 | | 1 | | 0,873 | 1,147 | 1,129 | 1,003 | 0,962 | 0,862 | 1,034 | 1,149 | 1,04 | 0,04 | 6,9E-01 |
| Q8VDI9 | Alpha-1,2-mannosyltransferase ALG9 OS=Mus musculus OX=100 | 2 | 1 | 1 | 1 | 611 | 69,5 | 8,94 | 396 | | 1 | | 1,265 | 0,941 | 1,06 | 1,387 | 1,29 | 1,301 | 1,191 | 1,059 | 0,96 | -0,04 | 7,0E-01 |
| Q6PAK3 | Protein arginine N-methyltransferase 8 OS=Mus musculus OX=1 | 12 | 4 | 4 | 2 | 394 | 45,2 | 6,93 | 1873 | 2,27 | 3 | 1 | 1,112 | 1,067 | 0,898 | 1,436 | 1,065 | 0,98 | 1,348 | 1,364 | 0,95 | -0,05 | 7,0E-01 |
| Q99K01 | Pyridoxal-dependent decarboxylase domain-containing protein | 5 | 3 | 3 | 3 | 787 | 87,3 | 5,48 | 704 | 0 | 2 | 1 | 1,443 | 1,179 | 1,285 | 1,187 | 0,761 | 1,25 | 1,512 | 1,293 | 1,06 | 0,06 | 7,0E-01 |
| Q8C3I8 | Protein HGH1 homolog OS=Mus musculus OX=10090 GN=Hgh1 | 9 | 2 | 2 | 2 | 393 | 42,9 | 4,74 | 1427 | | 2 | | 1,035 | 1,059 | 0,909 | 0,982 | 1,454 | 1,05 | 0,763 | 0,961 | 0,94 | -0,06 | 7,0E-01 |
| Q505F5 | Leucine-rich repeat-containing protein 47 OS=Mus musculus OX | 13 | 6 | 6 | 6 | 581 | 63,6 | 8,1 | 2628 | 4,86 | 4 | 2 | 0,971 | 0,992 | 1,004 | 1,001 | 1,101 | 0,981 | 0,982 | 0,958 | 0,99 | -0,01 | 7,0E-01 |
| Q9DBZ5 | Eukaryotic translation initiation factor 3 subunit K OS=Mus musc | 13 | 2 | 2 | 2 | 218 | 25,1 | 4,93 | 758 | | 2 | | 0,913 | 0,898 | 0,911 | 0,822 | 0,873 | 0,709 | 1,127 | 0,66 | 1,05 | 0,05 | 7,0E-01 |
| Q8BGQ7 | Alanine--tRNA ligase, cytoplasmic OS=Mus musculus OX=10090 | 21 | 20 | 35 | 20 | 968 | 106,8 | 5,67 | 12588 | 39,23 | 13 | 7 | 0,954 | 0,919 | 0,953 | 0,973 | 0,873 | 0,983 | 0,92 | 0,977 | 1,01 | 0,01 | 7,0E-01 |
| Q8BMQ8 | Vacuolar fusion protein MON1 homolog B OS=Mus musculus OX | 2 | 1 | 1 | 1 | 553 | 60 | 5,57 | 315 | | 1 | | 0,908 | 0,849 | 0,948 | 0,958 | 0,932 | 0,82 | 0,724 | 1,062 | 1,04 | 0,04 | 7,0E-01 |
| Q9D0R2 | Threonine--tRNA ligase, cytoplasmic OS=Mus musculus OX=100 | 6 | 4 | 4 | 4 | 722 | 83,3 | 7,36 | 1539 | 2,24 | 3 | 1 | 0,799 | 0,739 | 1,115 | 0,716 | 1,66 | 0,9 | 0,631 | 0,605 | 0,89 | -0,11 | 7,0E-01 |
| Q9DCT8 | Cysteine-rich protein 2 OS=Mus musculus OX=10090 GN=Crip2 | 40 | 5 | 10 | 5 | 208 | 22,7 | 8,63 | 2597 | 12,77 | 3 | 3 | 0,923 | 0,968 | 0,998 | 0,966 | 1,093 | 0,909 | 0,924 | 1,002 | 0,98 | -0,02 | 7,0E-01 |
| Q02248 | Catenin beta-1 OS=Mus musculus OX=10090 GN=Ctnnb1 PE=1 S | 28 | 17 | 31 | 16 | 781 | 85,4 | 5,86 | 15446 | 19,93 | 15 | 5 | 0,976 | 0,968 | 1,001 | 0,934 | 1,172 | 0,92 | 0,943 | 0,943 | 0,98 | -0,02 | 7,0E-01 |
| Q6DFV3 | Rho GTPase-activating protein 21 OS=Mus musculus OX=10090 | 1 | 2 | 2 | 2 | 1944 | 215,6 | 7,64 | 754 | | 2 | | 1,345 | 0,979 | 0,975 | 1,278 | 2,132 | 0,939 | 1,005 | 0,995 | 0,90 | -0,10 | 7,0E-01 |
| Q3TXX4 | Vesicular glutamate transporter 1 OS=Mus musculus OX=10090 | 5 | 4 | 6 | 4 | 560 | 61,6 | 7,34 | 1126 | 12,26 | 2 | 2 | 0,968 | 1,019 | 1,015 | 0,957 | 0,801 | 0,979 | 0,999 | 1,081 | 1,03 | 0,03 | 7,0E-01 |
| Q9ERE7 | LRP chaperone MESD OS=Mus musculus OX=10090 GN=Mesd P | 9 | 2 | 3 | 2 | 224 | 25,2 | 6,34 | 1450 | | 2 | | 1,24 | 1,008 | 0,945 | 1,073 | 1,13 | 1,153 | 1,037 | 1,058 | 0,97 | -0,03 | 7,0E-01 |
| Q8K0D5 | Elongation factor G, mitochondrial OS=Mus musculus OX=1009 | 1 | 1 | 1 | 1 | 751 | 83,5 | 6,92 | 537 | | 1 | | 1,355 | 1,562 | 1,116 | 0,941 | 1,149 | 1,082 | 1,425 | 1,06 | 1,05 | 0,05 | 7,0E-01 |
| P84096 | Rho-related GTP-binding protein RhoG OS=Mus musculus OX=1 | 10 | 2 | 3 | 1 | 191 | 21,3 | 8,12 | 1213 | 3,47 | 2 | 1 | 0,732 | 1,033 | 0,81 | 0,855 | 0,953 | 0,627 | 1,157 | 0,897 | 0,94 | -0,06 | 7,0E-01 |
| Q91W61 | F-box/LRR-repeat protein 15 OS=Mus musculus OX=10090 GN=F | 4 | 1 | 1 | 1 | 300 | 33,1 | 7,43 | | 2,05 | | 1 | 0,966 | 0,916 | 1,244 | 0,929 | 1,346 | 0,886 | 0,66 | 0,9 | 1,07 | 0,07 | 7,0E-01 |
| P10630 | Eukaryotic initiation factor 4A-II OS=Mus musculus OX=10090 G | 15 | 5 | 8 | 1 | 407 | 46,4 | 5,48 | 4440 | 2,14 | 5 | 1 | 1,06 | 0,929 | 1,152 | 1,094 | 1,243 | 0,878 | 0,668 | 1,211 | 1,06 | 0,06 | 7,0E-01 |
| P70303 | CTP synthase 2 OS=Mus musculus OX=10090 GN=Ctps2 PE=1 SV | 4 | 2 | 3 | 1 | 586 | 65,5 | 6,49 | 2115 | | 2 | | 1,061 | 0,978 | 1,009 | 0,854 | 1,333 | 0,721 | 1,002 | 1,06 | 0,95 | -0,05 | 7,0E-01 |
| P84075 | Neuron-specific calcium-binding protein hippocalcin OS=Mus mu | 39 | 8 | 9 | 2 | 193 | 22,4 | 4,97 | 4428 | 5,58 | 6 | 2 | 1,063 | 1,271 | 1,176 | 1,006 | 1,02 | 1,312 | 1,026 | 1,006 | 1,03 | 0,03 | 7,0E-01 |
| Q9ET26 | E3 ubiquitin-protein ligase RNF114 OS=Mus musculus OX=1009 | 12 | 2 | 2 | 2 | 229 | 25,7 | 7,03 | 1186 | | 2 | | 0,815 | 0,626 | 0,851 | 0,961 | 0,617 | 0,624 | 1,044 | 0,773 | 1,06 | 0,06 | 7,0E-01 |
| P62270 | 40S ribosomal protein S18 OS=Mus musculus OX=10090 GN=Rp | 38 | 7 | 13 | 7 | 152 | 17,7 | 10,99 | 5105 | 10,51 | 4 | 3 | 0,974 | 0,934 | 0,961 | 1,033 | 1,176 | 0,938 | 0,911 | 0,978 | 0,97 | -0,03 | 7,0E-01 |
| Q91YX5 | Acyl-CoA:lysophosphatidylglycerol acyltransferase 1 OS=Mus mu | 2 | 1 | 1 | 1 | 370 | 43,1 | 8,79 | 351 | | 1 | | 1,167 | 1,107 | 0,987 | 1,223 | 1,198 | 1,387 | 0,894 | 0,767 | 1,06 | 0,06 | 7,1E-01 |
| Q8VHK9 | ATP-dependent DNA/RNA helicase DHX36 OS=Mus musculus OX | 1 | 1 | 1 | 1 | 1001 | 113,8 | 8,29 | 784 | | 1 | | 1,047 | 1,073 | 1,333 | 1,284 | 1,001 | 1,345 | 1,182 | 1,04 | 1,04 | 0,04 | 7,1E-01 |
| P21956 | Lactadherin OS=Mus musculus OX=10090 GN=Mfge8 PE=1 SV=3 | 5 | 2 | 2 | 2 | 463 | 51,2 | 6,52 | 319 | 2,37 | 1 | 1 | 0,99 | 1,175 | 1,039 | 0,993 | 1,316 | 0,957 | 0,892 | 0,85 | 1,05 | 0,05 | 7,1E-01 |
| P54775 | 26S proteasome regulatory subunit 6B OS=Mus musculus OX=1 | 23 | 9 | 11 | 9 | 418 | 47,4 | 5,21 | 3991 | 7,45 | 6 | 3 | 0,958 | 1,112 | 1,539 | 0,976 | 2,145 | 0,954 | 1,009 | 0,985 | 0,90 | -0,10 | 7,1E-01 |
| Q06335 | Amyloid-like protein 2 OS=Mus musculus OX=10090 GN=Aplp2 | 4 | 3 | 3 | 2 | 707 | 80,4 | 4,7 | 1345 | 2,95 | 2 | 1 | 0,79 | 1,201 | 1,015 | 0,872 | 0,766 | 0,916 | 0,873 | 1,135 | 1,05 | 0,05 | 7,1E-01 |
| O35864 | COP9 signalosome complex subunit 5 OS=Mus musculus OX=10 | 8 | 2 | 2 | 2 | 334 | 37,5 | 6,54 | 1199 | | 2 | | 0,926 | 0,987 | 1,109 | 1,184 | 0,628 | 1,145 | 1,19 | 1,021 | 1,06 | 0,06 | 7,1E-01 |
| Q9DD18 | D-aminoacyl-tRNA deacylase 1 OS=Mus musculus OX=10090 GN | 4 | 1 | 1 | 1 | 209 | 23,4 | 7,87 | 255 | | 1 | | 0,656 | 0,513 | 1,012 | 0,595 | 0,786 | 0,635 | 0,643 | 0,518 | 1,08 | 0,08 | 7,1E-01 |
| Q09PK2 | Retroviral-like aspartic protease 1 OS=Mus musculus OX=10090 | 2 | 1 | 2 | 1 | 339 | 37,2 | 6,1 | | 3,86 | | 1 | 1,252 | 2,605 | 0,677 | 0,53 | 1,244 | 0,954 | 0,667 | 1,41 | 1,18 | 0,18 | 7,1E-01 |
| Q61739 | Integrin alpha-6 OS=Mus musculus OX=10090 GN=Itga6 PE=1 SV | 1 | 1 | 1 | 1 | 1091 | 122,1 | 7,03 | 555 | | 1 | | 0,969 | 0,595 | 1,021 | 1,176 | 1,004 | 1,067 | 1,037 | 0,86 | 0,95 | -0,05 | 7,1E-01 |
| Q9DCM0 | Persulfide dioxygenase ETHE1, mitochondrial OS=Mus musculus | 12 | 2 | 2 | 2 | 254 | 27,7 | 7,23 | 1304 | | 2 | | 0,962 | 1,155 | 0,951 | 0,819 | 1,04 | 0,753 | 1,084 | 0,845 | 1,04 | 0,04 | 7,1E-01 |
| Q6ZQ38 | Cullin-associated NEDD8-dissociated protein 1 OS=Mus musculu | 17 | 19 | 24 | 19 | 1230 | 136,2 | 5,78 | 7803 | 29,48 | 10 | 10 | 0,997 | 0,907 | 0,963 | 0,921 | 0,802 | 1,017 | 0,941 | 0,95 | 1,02 | 0,02 | 7,1E-01 |
| O08810 | 116 kDa U5 small nuclear ribonucleoprotein component OS=Mu | 3 | 3 | 3 | 2 | 971 | 109,3 | 5 | 1338 | 2,09 | 2 | 1 | 1,101 | 0,956 | 0,942 | 1,151 | 1,072 | 0,911 | 1,147 | 1,139 | 0,97 | -0,03 | 7,1E-01 |
| Q7TT50 | Serine/threonine-protein kinase MRCK beta OS=Mus musculus | 8 | 12 | 14 | 10 | 1713 | 194,6 | 6,46 | 5829 | 7,81 | 9 | 3 | 0,964 | 0,913 | 1,016 | 0,959 | 1,045 | 0,904 | 0,948 | 0,891 | 1,02 | 0,02 | 7,1E-01 |
| Q8BLF1 | Neutral cholesterol ester hydrolase 1 OS=Mus musculus OX=10 | 14 | 5 | 8 | 5 | 408 | 45,7 | 7,05 | 3853 | | 5 | | 0,998 | 0,902 | 0,955 | 1,006 | 0,981 | 0,883 | 1,007 | 0,933 | 1,01 | 0,01 | 7,1E-01 |
| Q9Z0X1 | Apoptosis-inducing factor 1, mitochondrial OS=Mus musculus O | 17 | 8 | 14 | 8 | 612 | 66,7 | 9,17 | 6803 | 7,02 | 6 | 2 | 1,083 | 1,139 | 1,052 | 1,064 | 1,213 | 1,06 | 1,002 | 1,141 | 0,98 | -0,02 | 7,1E-01 |
| O35593 | 26S proteasome non-ATPase regulatory subunit 14 OS=Mus mu | 9 | 2 | 3 | 2 | 310 | 34,6 | 6,52 | 1100 | 0 | 2 | 1 | 1,101 | 1,025 | 1,13 | 0,953 | 1,037 | 1,037 | 1,139 | 0,898 | 1,02 | 0,02 | 7,1E-01 |
| Q9D2N9 | Vacuolar protein sorting-associated protein 33A OS=Mus muscu | 4 | 3 | 3 | 3 | 598 | 67,5 | 7,08 | 2079 | | 3 | | 1,068 | 1,008 | 1,114 | 1,152 | 0,942 | 1,073 | 1,063 | 1,176 | 1,02 | 0,02 | 7,1E-01 |
| Q9CPY7 | Cytosol aminopeptidase OS=Mus musculus OX=10090 GN=Lap3 | 23 | 10 | 16 | 10 | 519 | 56,1 | 7,72 | 7772 | 11,67 | 8 | 3 | 1,063 | 1,001 | 1,049 | 1,001 | 0,901 | 1,095 | 1,044 | 1,006 | 1,02 | 0,02 | 7,1E-01 |
| A2ALS5 | Rap1 GTPase-activating protein 1 OS=Mus musculus OX=10090 | 9 | 4 | 5 | 4 | 663 | 73,4 | 5,87 | 1440 | 7,83 | 2 | 2 | 1,019 | 0,948 | 1,007 | 1,1 | 0,868 | 1,253 | 1,092 | 0,995 | 0,97 | -0,03 | 7,1E-01 |
| Q9Z1K7 | Adenomatous polyposis coli protein 2 OS=Mus musculus OX=10 | 1 | 2 | 2 | 2 | 2274 | 243 | 8,88 | | 2,15 | | 2 | 0,981 | 0,901 | 1,053 | 1,04 | 0,923 | 0,923 | 1,205 | 0,776 | 1,04 | 0,04 | 7,1E-01 |
| Q9D7N3 | 28S ribosomal protein S9, mitochondrial OS=Mus musculus OX= | 7 | 3 | 3 | 3 | 390 | 44,9 | 8,81 | 624 | 2,33 | 1 | 2 | 0,981 | 0,938 | 0,934 | 0,906 | 0,916 | 0,932 | 1,039 | 0,79 | 1,02 | 0,02 | 7,1E-01 |
| Q9Z2C9 | Myotubularin-related protein 7 OS=Mus musculus OX=10090 G | 2 | 1 | 1 | 1 | 660 | 75,6 | 6,43 | | 2,31 | | 1 | 0,813 | 0,67 | 0,78 | 0,96 | 0,879 | 0,666 | 0,753 | 0,81 | 1,04 | 0,04 | 7,1E-01 |
| Q8CGA4 | Maturin OS=Mus musculus OX=10090 GN=Mturn PE=1 SV=1 | 6 | 1 | 1 | 1 | 131 | 14,8 | 4,17 | 668 | | 1 | | 0,965 | 0,872 | 0,878 | 1,326 | 1,103 | 1,191 | 0,96 | 0,972 | 0,96 | -0,04 | 7,1E-01 |
| P63318 | Protein kinase C gamma type OS=Mus musculus OX=10090 GN= | 14 | 9 | 15 | 8 | 697 | 78,3 | 7,46 | 4829 | 9,15 | 5 | 4 | 0,858 | 0,96 | 0,991 | 0,879 | 0,91 | 0,941 | 0,965 | 0,924 | 0,99 | -0,01 | 7,1E-01 |
| P27661 | Histone H2AX OS=Mus musculus OX=10090 GN=H2afx PE=1 SV= | 17 | 4 | 8 | 2 | 143 | 15,1 | 10,74 | | 22,1 | | 4 | 0,787 | 0,723 | 1,079 | 0,898 | 0,892 | 0,67 | 1,296 | 0,863 | 0,94 | -0,06 | 7,2E-01 |
| O70493 | Sorting nexin-12 OS=Mus musculus OX=10090 GN=Snx12 PE=1 S | 17 | 3 | 4 | 2 | 165 | 19,1 | 7,34 | 2585 | | 3 | | 0,891 | 1,009 | 1,101 | 1,072 | 0,837 | 0,994 | 1,303 | 1,105 | 0,96 | -0,04 | 7,2E-01 |
| Q9ERR7 | Selenoprotein F OS=Mus musculus OX=10090 GN=Selenof PE=1 | 4 | 1 | 1 | 1 | 162 | 17,8 | 5,35 | 735 | | 1 | | 0,953 | 0,717 | 1,027 | 1,046 | 0,831 | 0,819 | 0,93 | 1,026 | 1,04 | 0,04 | 7,2E-01 |
| Q9Z2A5 | Arginyl-tRNA--protein transferase 1 OS=Mus musculus OX=1009 | 4 | 2 | 3 | 2 | 516 | 59,1 | 8,19 | 1942 | | 2 | | 1,097 | 1,144 | 1 | 1,259 | 1,14 | 1,276 | 1,001 | 1,204 | 0,97 | -0,03 | 7,2E-01 |
| Q6PB93 | Polypeptide N-acetylgalactosaminyltransferase 2 OS=Mus musc | 2 | 1 | 1 | 1 | 570 | 64,5 | 8,53 | 461 | | 1 | | 0,519 | 0,819 | 0,886 | 0,771 | 0,719 | 0,864 | 0,859 | 0,693 | 0,96 | -0,04 | 7,2E-01 |
| Q8CFA2 | Aminomethyltransferase, mitochondrial OS=Mus musculus OX= | 3 | 1 | 1 | 1 | 403 | 44 | 8,7 | | 2,9 | | 1 | 0,916 | 0,919 | 0,949 | 0,942 | 0,917 | 0,697 | 0,86 | 1,119 | 1,04 | 0,04 | 7,2E-01 |
| Q8K3C3 | Protein LZIC OS=Mus musculus OX=10090 GN=Lzic PE=1 SV=1 | 5 | 1 | 1 | 1 | 190 | 21,5 | 4,96 | | 2,68 | | 1 | 1,066 | 1,338 | 0,776 | 1,172 | 0,936 | 1,323 | 1,028 | 1,297 | 0,95 | -0,05 | 7,2E-01 |
| Q03173 | Protein enabled homolog OS=Mus musculus OX=10090 GN=Ena | 7 | 4 | 5 | 4 | 802 | 85,8 | 7,72 | 1688 | 2,66 | 3 | 2 | 1,006 | 0,94 | 1,041 | 0,9 | 0,885 | 0,933 | 0,981 | 1,022 | 1,02 | 0,02 | 7,2E-01 |
| Q02819 | Nucleobindin-1 OS=Mus musculus OX=10090 GN=Nucb1 PE=1 S | 10 | 4 | 5 | 4 | 459 | 53,4 | 5,07 | 2130 | 3,15 | 3 | 1 | 0,896 | 1,048 | 0,883 | 1,085 | 1,057 | 1,034 | 0,955 | 0,954 | 0,98 | -0,02 | 7,2E-01 |

| | | | | | | | | | | | | | | | | | | | | | | | |
|---|---|---|---|---|---|---|---|---|---|---|---|---|---|---|---|---|---|---|---|---|---|---|---|
| Q9D0L4 | Uncharacterized aarF domain-containing protein kinase 1 OS=M | 5 | 2 | 3 | 2 | 525 | 59,7 | 8,05 | 393 | 4,5 | 1 | 2 | 0,885 | 0,902 | 0,85 | 1,081 | 0,888 | 0,974 | 0,901 | 1,051 | 0,97 | -0,03 | 7,2E-01 |
| Q9CQ22 | Ragulator complex protein LAMTOR1 OS=Mus musculus OX=100 | 10 | 1 | 4 | 1 | 161 | 17,7 | 5,15 | 236 | 7,55 | 1 | 1 | 1,166 | 1,134 | 1,423 | 1,176 | 1,212 | 1,073 | 1,379 | 1,089 | 1,03 | 0,03 | 7,2E-01 |
| P14131 | 40S ribosomal protein S16 OS=Mus musculus OX=10090 GN=Rp: | 34 | 6 | 10 | 6 | 146 | 16,4 | 10,21 | 2798 | 12,92 | 3 | 3 | 1,017 | 0,905 | 0,976 | 0,984 | 0,948 | 0,945 | 1,014 | 1,022 | 0,99 | -0,01 | 7,2E-01 |
| Q91ZU6 | Dystonin OS=Mus musculus OX=10090 GN=Dst PE=1 SV=2 | 2 | 12 | 15 | 11 | 7393 | 833,7 | 5,31 | 5688 | 2,1 | 11 | 1 | 1,036 | 0,967 | 0,919 | 0,983 | 1,048 | 0,911 | 1,008 | 0,994 | 0,99 | -0,01 | 7,2E-01 |
| Q9Z140 | Copine-6 OS=Mus musculus OX=10090 GN=Cpne6 PE=1 SV=1 | 12 | 6 | 8 | 5 | 557 | 61,7 | 5,59 | 4226 | 2,46 | 5 | 1 | 0,825 | 0,798 | 1,151 | 0,818 | 1,335 | 0,837 | 0,881 | 0,768 | 0,94 | -0,06 | 7,2E-01 |
| P98203 | Armadillo repeat protein deleted in velo-cardio-facial syndrome | 6 | 5 | 5 | 5 | 962 | 105 | 6,64 | 2322 | 2,42 | 4 | 1 | 1,001 | 0,979 | 1,034 | 0,972 | 1,14 | 0,936 | 1,015 | 0,965 | 0,98 | -0,02 | 7,2E-01 |
| Q8BYW9 | EGF domain-specific O-linked N-acetylglucosamine transferase ( | 2 | 1 | 1 | 1 | 527 | 61,4 | 6,81 | 530 | | 1 | | 1,16 | 1,076 | 0,847 | 0,833 | 0,999 | 1,14 | 0,7 | 1,302 | 0,95 | -0,05 | 7,2E-01 |
| Q9D486 | C-Maf-inducing protein OS=Mus musculus OX=10090 GN=Cmip | 3 | 2 | 3 | 2 | 773 | 86,2 | 6,81 | 1335 | | 2 | | 0,989 | 1,129 | 0,874 | 1,183 | 0,982 | 0,998 | 1,04 | 1,049 | 1,03 | 0,03 | 7,2E-01 |
| Q91X78 | Erlin-1 OS=Mus musculus OX=10090 GN=Erlin1 PE=1 SV=2 | 9 | 3 | 3 | 2 | 348 | 39,2 | 7,21 | 684 | 2,27 | 2 | 1 | 0,991 | 1,132 | 0,999 | 1,022 | 1,305 | 0,775 | 0,942 | 1,33 | 0,95 | -0,05 | 7,2E-01 |
| P38585 | Tubulin--tyrosine ligase OS=Mus musculus OX=10090 GN=Ttl PE | 8 | 2 | 2 | 2 | 377 | 43,1 | 6,46 | 552 | | 2 | | 0,754 | 0,896 | 1,996 | 0,889 | 2,799 | 0,827 | 0,902 | 0,842 | 0,84 | -0,16 | 7,2E-01 |
| Q8BFU3 | RING finger protein 214 OS=Mus musculus OX=10090 GN=Rnf21 | 17 | 10 | 10 | 10 | 668 | 73,6 | 6,23 | 2745 | 6,96 | 5 | 5 | 1,12 | 1,099 | 1,065 | 1,039 | 1,075 | 1,032 | 1,123 | 1,054 | 1,01 | 0,01 | 7,3E-01 |
| P62897 | Cytochrome c, somatic OS=Mus musculus OX=10090 GN=Cycs P | 45 | 7 | 15 | 7 | 105 | 11,6 | 9,58 | 3315 | 35,38 | 4 | 5 | 1,02 | 1,025 | 0,974 | 0,826 | 0,865 | 1,303 | 0,883 | 0,959 | 0,96 | -0,04 | 7,3E-01 |
| Q06185 | ATP synthase subunit e, mitochondrial OS=Mus musculus OX=1( | 49 | 5 | 8 | 5 | 71 | 8,2 | 9,35 | 4779 | | 5 | | 0,962 | 0,867 | 1,003 | 0,917 | 1,103 | 0,844 | 0,975 | 0,918 | 0,98 | -0,02 | 7,3E-01 |
| Q64523 | Histone H2A type 2-C OS=Mus musculus OX=10090 GN=Hist2h2 | 16 | 3 | 7 | 1 | 129 | 14 | 10,9 | | 18,76 | | 3 | 0,905 | 0,769 | 1,632 | 0,906 | 1,462 | 0,723 | 0,961 | 0,676 | 1,10 | 0,10 | 7,3E-01 |
| P41241 | Tyrosine-protein kinase CSK OS=Mus musculus OX=10090 GN=C | 9 | 4 | 4 | 4 | 450 | 50,7 | 7,06 | 1764 | 0 | 3 | 1 | 0,935 | 1,009 | 1,15 | 0,936 | 0,967 | 0,844 | 1,033 | 1,081 | 1,03 | 0,03 | 7,3E-01 |
| Q91VD9 | NADH-ubiquinone oxidoreductase 75 kDa subunit, mitochondria | 40 | 23 | 35 | 23 | 727 | 79,7 | 5,72 | 12929 | 43,54 | 14 | 10 | 0,978 | 0,916 | 0,989 | 0,952 | 0,892 | 0,99 | 0,906 | 1,13 | 0,98 | -0,02 | 7,3E-01 |
| P63158 | High mobility group protein B1 OS=Mus musculus OX=10090 GN | 22 | 6 | 9 | 6 | 215 | 24,9 | 5,74 | 1308 | 16,04 | 3 | 3 | 0,943 | 0,904 | 0,892 | 1,026 | 0,853 | 1,015 | 0,998 | 0,968 | 0,98 | -0,02 | 7,3E-01 |
| P70697 | Uroporphyrinogen decarboxylase OS=Mus musculus OX=10090 | 7 | 2 | 2 | 2 | 367 | 40,7 | 6,65 | 703 | 2,87 | 1 | 1 | 0,821 | 0,966 | 0,877 | 0,994 | 0,979 | 0,713 | 0,977 | 1,127 | 0,96 | -0,04 | 7,3E-01 |
| Q9JIW9 | Ras-related protein Ral-B OS=Mus musculus OX=10090 GN=Ralb | 29 | 6 | 13 | 5 | 206 | 23,3 | 6,62 | 6455 | 4,88 | 6 | 2 | 0,909 | 0,952 | 0,884 | 0,993 | 1,031 | 0,852 | 0,917 | 0,87 | 1,02 | 0,02 | 7,3E-01 |
| Q9D2R0 | Acetoacetyl-CoA synthetase OS=Mus musculus OX=10090 GN=A | 4 | 3 | 4 | 3 | 672 | 75,2 | 6,71 | 1010 | 4,23 | 2 | 1 | 0,822 | 0,996 | 0,864 | 0,94 | 1,006 | 0,783 | 0,983 | 0,942 | 0,98 | -0,02 | 7,3E-01 |
| Q8C8T8 | Pre-rRNA-processing protein TSR2 homolog OS=Mus musculus ( | 8 | 1 | 2 | 1 | 191 | 20,9 | 4,23 | 413 | | 1 | | 0,665 | 1,359 | 1,687 | 0,987 | 1,43 | 0,862 | 1,168 | 0,86 | 1,09 | 0,09 | 7,3E-01 |
| Q0P5W1 | Vacuolar protein sorting-associated protein 8 homolog OS=Mus | 1 | 1 | 1 | 1 | 1427 | 161 | 5,58 | 535 | | 1 | | 0,844 | 0,681 | 0,923 | 0,983 | 0,997 | 0,716 | 0,887 | 0,962 | 0,96 | -0,04 | 7,3E-01 |
| Q8C167 | Prolyl endopeptidase-like OS=Mus musculus OX=10090 GN=Pre| | 5 | 3 | 6 | 3 | 725 | 83,1 | 6,65 | 1515 | 8,17 | 2 | 2 | 1,094 | 1,068 | 1,07 | 1,098 | 1,006 | 1,141 | 1,058 | 1,184 | 0,99 | -0,01 | 7,3E-01 |
| Q3UFY7 | 7-methylguanosine phosphate-specific 5'-nucleotidase OS=Mus | 10 | 3 | 3 | 3 | 300 | 34,4 | 6,24 | 335 | 4,28 | 1 | 2 | 0,997 | 0,947 | 1,156 | 0,902 | 1,111 | 0,893 | 1,022 | 1,082 | 0,97 | -0,03 | 7,3E-01 |
| E9Q557 | Desmoplakin OS=Mus musculus OX=10090 GN=Dsp PE=1 SV=1 | 4 | 16 | 21 | 16 | 2883 | 332,7 | 6,8 | 6975 | 15,65 | 8 | 8 | 0,831 | 1,459 | 0,779 | 0,635 | 0,933 | 0,79 | 0,907 | 1,398 | 0,92 | -0,08 | 7,3E-01 |
| Q9D172 | Glutamine amidotransferase-like class 1 domain-containing prot | 23 | 5 | 7 | 5 | 266 | 28,1 | 8,78 | 1830 | 7,33 | 3 | 2 | 1,031 | 1,062 | 1,1 | 1,111 | 0,814 | 1,235 | 1,029 | 1,098 | 1,03 | 0,03 | 7,3E-01 |
| P97950 | Ras-related protein Rab-33A OS=Mus musculus OX=10090 GN=F | 13 | 3 | 3 | 2 | 237 | 26,5 | 7,88 | 2013 | | 3 | | 0,977 | 1,088 | 1,056 | 1,014 | 1,211 | 0,987 | 0,987 | 0,834 | 1,03 | 0,03 | 7,3E-01 |
| Q9R1L5 | Microtubule-associated serine/threonine-protein kinase 1 OS=N | 4 | 5 | 5 | 5 | 1570 | 170,9 | 8,44 | 2325 | 0 | 4 | 1 | 1,173 | 1,241 | 0,997 | 0,959 | 1,258 | 1,037 | 1,343 | 0,904 | 0,96 | -0,04 | 7,4E-01 |
| Q61335 | B-cell receptor-associated protein 31 OS=Mus musculus OX=100 | 18 | 4 | 6 | 4 | 245 | 27,9 | 8,7 | 3222 | 2,9 | 3 | 1 | 0,974 | 0,91 | 0,931 | 1,063 | 0,649 | 0,942 | 1,604 | 0,971 | 0,93 | -0,07 | 7,4E-01 |
| Q9D6F9 | Tubulin beta-4A chain OS=Mus musculus OX=10090 GN=Tubb4a | 58 | 24 | 365 | 4 | 444 | 49,6 | 4,88 | 156873 | 275,58 | 19 | 18 | 0,999 | 0,983 | 0,977 | 0,984 | 0,882 | 0,971 | 1,082 | 0,949 | 1,02 | 0,02 | 7,4E-01 |
| Q9WTR1 | Transient receptor potential cation channel subfamily V membe | 8 | 4 | 4 | 4 | 756 | 85,9 | 6,99 | 1039 | | 4 | | 1,027 | 1,238 | 0,887 | 1,113 | 1,174 | 0,861 | 1,191 | 1,194 | 0,96 | -0,04 | 7,4E-01 |
| Q3TXS7 | 26S proteasome non-ATPase regulatory subunit 1 OS=Mus musc | 16 | 13 | 20 | 12 | 953 | 105,7 | 5,39 | 8008 | 15,19 | 10 | 4 | 1,048 | 0,986 | 1,059 | 1,053 | 1,192 | 1,042 | 0,961 | 1,024 | 0,98 | -0,02 | 7,4E-01 |
| Q8R313 | Exocyst complex component 6 OS=Mus musculus OX=10090 GN | 4 | 3 | 3 | 3 | 802 | 93 | 6,15 | 1854 | | 3 | | 1,154 | 1,034 | 1,14 | 0,978 | 1,267 | 0,955 | 0,971 | 0,993 | 1,03 | 0,03 | 7,4E-01 |
| Q8CJ40 | Rootletin OS=Mus musculus OX=10090 GN=Crocc PE=1 SV=2 | 1 | 2 | 2 | 2 | 2009 | 226,8 | 5,55 | 1188 | | 2 | | 0,668 | 2,08 | 0,735 | 0,472 | 1,377 | 0,564 | 0,65 | 2,083 | 0,85 | -0,15 | 7,4E-01 |
| Q8K2F8 | Protein LSM14 homolog A OS=Mus musculus OX=10090 GN=Lsn | 2 | 1 | 1 | 1 | 462 | 50,5 | 9,52 | 290 | | 1 | | 1,154 | 0,798 | 0,945 | 0,866 | 1,048 | 0,943 | 0,796 | 0,842 | 1,04 | 0,04 | 7,4E-01 |
| Q8R1Y2 | bMERB domain-containing protein 1 OS=Mus musculus OX=100 | 5 | 1 | 1 | 1 | 203 | 23,5 | 5,63 | 477 | | 1 | | 0,683 | 1,17 | 0,828 | 0,833 | 0,667 | 0,784 | 1,342 | 0,974 | 0,93 | -0,07 | 7,4E-01 |
| D3Z7P3 | Glutaminase kidney isoform, mitochondrial OS=Mus musculus C | 19 | 9 | 12 | 9 | 674 | 73,9 | 7,99 | 5672 | 9,86 | 7 | 3 | 1,001 | 0,851 | 1,051 | 0,858 | 0,837 | 1,128 | 0,949 | 0,957 | 0,97 | -0,03 | 7,4E-01 |
| D3YZI9 | PiggyBac transposable element-derived protein 5 OS=Mus musc | 2 | 1 | 1 | 1 | 523 | 58,3 | 8,85 | 378 | | 1 | | 0,928 | 0,665 | 0,748 | 0,746 | 0,952 | 0,703 | 0,724 | 0,573 | 1,05 | 0,05 | 7,4E-01 |
| Q80W04 | Transmembrane and coiled-coil domains protein 2 OS=Mus mus | 2 | 1 | 1 | 1 | 706 | 77 | 6,84 | 541 | | 1 | | 0,902 | 0,857 | 0,785 | 0,987 | 1,291 | 0,865 | 0,644 | 0,928 | 0,95 | -0,05 | 7,4E-01 |
| P62743 | AP-2 complex subunit sigma OS=Mus musculus OX=10090 GN=A | 6 | 1 | 1 | 1 | 142 | 17 | 6,18 | 551 | | 1 | | 0,574 | 0,697 | 0,921 | 0,587 | 0,867 | 0,653 | 0,86 | 0,555 | 0,95 | -0,05 | 7,4E-01 |
| Q9R0M6 | Ras-related protein Rab-9A OS=Mus musculus OX=10090 GN=Ra | 13 | 2 | 2 | 2 | 201 | 22,9 | 5,66 | 486 | 2,49 | 1 | 1 | 1,415 | 0,886 | 1,039 | 1,129 | 1,176 | 1,025 | 1,113 | 1,335 | 0,96 | -0,04 | 7,4E-01 |
| Q3TY65 | Islet cell autoantigen 1-like protein OS=Mus musculus OX=1009( | 7 | 3 | 5 | 2 | 431 | 48,1 | 4,91 | 1931 | 2,46 | 2 | 1 | 0,965 | 0,946 | 1,069 | 0,913 | 0,838 | 1,115 | 1,114 | 0,933 | 0,97 | -0,03 | 7,4E-01 |
| Q9CPT4 | Myeloid-derived growth factor OS=Mus musculus OX=10090 GN | 23 | 3 | 5 | 3 | 166 | 18 | 6,79 | 1798 | 0 | 2 | 1 | 1,097 | 1,065 | 1,043 | 1,182 | 1,24 | 1,054 | 1,137 | 1,034 | 0,98 | -0,02 | 7,4E-01 |
| Q60749 | KH domain-containing, RNA-binding, signal transduction-associa | 7 | 3 | 3 | 3 | 443 | 48,3 | 8,72 | | 8,76 | | 3 | 0,878 | 0,922 | 1,138 | 1,143 | 0,756 | 0,903 | 1,177 | 1,082 | 1,04 | 0,04 | 7,4E-01 |
| F6SEU4 | Ras/Rap GTPase-activating protein SynGAP OS=Mus musculus O | 10 | 10 | 12 | 9 | 1340 | 148,1 | 8,98 | 5223 | 5,79 | 9 | 1 | 1,021 | 0,996 | 1,131 | 1,007 | 1,037 | 1,031 | 1,167 | 0,989 | 0,98 | -0,02 | 7,4E-01 |
| Q00PI9 | Heterogeneous nuclear ribonucleoprotein U-like protein 2 OS=N | 10 | 7 | 8 | 7 | 745 | 84,9 | 4,89 | 2295 | 5,85 | 4 | 3 | 0,852 | 0,822 | 1,188 | 0,996 | 1,088 | 1,003 | 1,126 | 0,796 | 0,96 | -0,04 | 7,4E-01 |
| A2AP18 | 1-phosphatidylinositol 4,5-bisphosphate phosphodiesterase eta | 1 | 2 | 2 | 2 | 1501 | 164,2 | 7,49 | 575 | 2,26 | 1 | 1 | 1,093 | 0,952 | 0,944 | 1,079 | 0,951 | 0,891 | 1,166 | 0,96 | 1,03 | 0,03 | 7,4E-01 |
| P60335 | Poly(rC)-binding protein 1 OS=Mus musculus OX=10090 GN=Pcb | 35 | 8 | 16 | 5 | 356 | 37,5 | 7,09 | 4246 | 26,12 | 4 | 6 | 1,095 | 1,077 | 1,131 | 1,128 | 1,036 | 1,071 | 1,135 | 1,148 | 1,01 | 0,01 | 7,4E-01 |
| O35963 | Ras-related protein Rab-33B OS=Mus musculus OX=10090 GN=F | 10 | 2 | 2 | 1 | 229 | 25,8 | 7,69 | 1449 | | 2 | | 0,856 | 0,848 | 1,102 | 0,796 | 1,226 | 1,057 | 0,839 | 0,673 | 0,95 | -0,05 | 7,4E-01 |
| Q2PZL6 | Protocadherin Fat 4 OS=Mus musculus OX=10090 GN=Fat4 PE=1 | 0 | 2 | 2 | 2 | 4981 | 540 | 4,91 | | 2,35 | | 2 | 0,976 | 0,923 | 1,05 | 1,021 | 1,08 | 1,052 | 0,839 | 0,911 | 1,02 | 0,02 | 7,4E-01 |
| Q9D338 | 39S ribosomal protein L19, mitochondrial OS=Mus musculus OX | 3 | 1 | 2 | 1 | 292 | 33,6 | 9,44 | 1423 | | 1 | | 1,086 | 1,043 | 1,008 | 0,995 | 0,906 | 1,083 | 0,835 | 1,192 | 1,03 | 0,03 | 7,4E-01 |
| P17047 | Lysosome-associated membrane glycoprotein 2 OS=Mus muscu | 2 | 1 | 1 | 1 | 415 | 45,7 | 7,39 | | 2,22 | | 1 | 1,051 | 0,978 | 0,821 | 1,144 | 1,08 | 0,954 | 1,017 | 0,822 | 1,03 | 0,03 | 7,4E-01 |
| Q99PL5 | Ribosome-binding protein 1 OS=Mus musculus OX=10090 GN=R | 8 | 11 | 12 | 11 | 1605 | 172,8 | 9,33 | 5803 | 8,13 | 8 | 3 | 1,015 | 1,003 | 1,002 | 1,098 | 1,141 | 0,959 | 1,023 | 1,056 | 0,99 | -0,01 | 7,4E-01 |
| Q8BL86 | Metallo-beta-lactamase domain-containing protein 2 OS=Mus n | 16 | 3 | 4 | 3 | 279 | 31,2 | 6,86 | 955 | 4,81 | 2 | 1 | 1,626 | 1,14 | 1,182 | 1,575 | 1,605 | 1,299 | 1,313 | 1,509 | 0,96 | -0,04 | 7,4E-01 |
| Q3UUI3 | Acyl-coenzyme A thioesterase THEM4 OS=Mus musculus OX=10 | 3 | 1 | 1 | 1 | 230 | 26 | 9,64 | 384 | | 1 | | 0,915 | 0,949 | 0,943 | 1,106 | 0,968 | 1,025 | 0,968 | 0,88 | 1,02 | 0,02 | 7,4E-01 |
| Q9JKR6 | Hypoxia up-regulated protein 1 OS=Mus musculus OX=10090 GI | 10 | 9 | 9 | 9 | 999 | 111,1 | 5,19 | 2822 | 7,53 | 6 | 3 | 1,021 | 0,978 | 0,967 | 1,069 | 0,933 | 0,987 | 1,098 | 1,079 | 0,98 | -0,02 | 7,4E-01 |
| Q8C1B1 | Calmodulin-regulated spectrin-associated protein 2 OS=Mus mu | 4 | 5 | 6 | 5 | 1461 | 164,2 | 6,86 | 2636 | 2,17 | 4 | 1 | 1,09 | 0,924 | 1,056 | 0,937 | 0,969 | 1,016 | 1,035 | 0,92 | 1,02 | 0,02 | 7,4E-01 |
| Q9WV60 | Glycogen synthase kinase-3 beta OS=Mus musculus OX=10090 ( | 28 | 11 | 17 | 7 | 420 | 46,7 | 8,78 | 3401 | 34,15 | 3 | 8 | 1,035 | 1,044 | 1,111 | 1,008 | 0,991 | 1,145 | 0,936 | 1,058 | 1,02 | 0,02 | 7,5E-01 |
| Q80Y98 | Phospholipase DDHD2 OS=Mus musculus OX=10090 GN=Ddhd2 | 1 | 1 | 1 | 1 | 699 | 79,5 | 5,31 | 728 | | 1 | | 0,956 | 1 | 0,954 | 0,696 | 0,701 | 1,098 | 0,854 | 0,805 | 1,04 | 0,04 | 7,5E-01 |
| Q8CIB5 | Fermitin family homolog 2 OS=Mus musculus OX=10090 GN=Fer | 5 | 3 | 3 | 3 | 680 | 77,8 | 6,7 | 2069 | | 3 | | 0,998 | 1,038 | 0,908 | 1,082 | 1,035 | 0,972 | 0,961 | 1,131 | 0,98 | -0,02 | 7,5E-01 |
| Q8BR92 | Paralemmin-2 OS=Mus musculus OX=10090 GN=Palm2 PE=1 SV | 20 | 7 | 12 | 6 | 376 | 42,1 | 5,15 | 4121 | 12,89 | 3 | 4 | 0,992 | 1,106 | 0,882 | 1,162 | 1,159 | 0,97 | 1,056 | 1,056 | 0,98 | -0,02 | 7,5E-01 |
| Q9D8W7 | OCIA domain-containing protein 2 OS=Mus musculus OX=10090 | 12 | 2 | 2 | 2 | 154 | 16,9 | 9,41 | 1238 | | 2 | | 0,933 | 0,865 | 1,164 | 0,867 | 0,861 | 1,043 | 0,959 | 1,082 | 0,97 | -0,03 | 7,5E-01 |
| O88737 | Protein bassoon OS=Mus musculus OX=10090 GN=Bsn PE=1 SV= | 8 | 22 | 24 | 21 | 3942 | 418,6 | 7,71 | 11971 | 2,66 | 20 | 2 | 1,014 | 1,029 | 0,918 | 0,928 | 1,019 | 0,914 | 0,989 | 1,018 | 0,99 | -0,01 | 7,5E-01 |
| Q8R0P4 | Mth938 domain-containing protein OS=Mus musculus OX=1009 | 20 | 2 | 2 | 2 | 122 | 13,2 | 7,99 | | 4,9 | | 2 | 1,028 | 1,266 | 1,17 | 0,92 | 1,138 | 0,882 | 1,14 | 1,092 | 1,03 | 0,03 | 7,5E-01 |
| P43276 | Histone H1.5 OS=Mus musculus OX=10090 GN=Hist1h1b PE=1 S | 32 | 11 | 25 | 8 | 223 | 22,6 | 10,92 | 4975 | 57,02 | 2 | 9 | 0,85 | 0,798 | 0,951 | 1,138 | 0,895 | 0,826 | 1,152 | 1,003 | 0,96 | -0,04 | 7,5E-01 |

| | | | | | | | | | | | | | | | | | | | | | | | |
|---|---|---|---|---|---|---|---|---|---|---|---|---|---|---|---|---|---|---|---|---|---|---|---|
| Q8K4Z3 | NAD(P)H-hydrate epimerase OS=Mus musculus OX=10090 GN=N | 25 | 5 | 9 | 5 | 282 | 31 | 7,69 | 4386 | 0 | 4 | 2 | 1,025 | 0,913 | 0,967 | 1,039 | 0,915 | 1,024 | 0,951 | 1,003 | 1,01 | 0,01 | 7,5E-01 |
| Q80TK0 | AP2-interacting clathrin-endocytosis protein OS=Mus musculus | 3 | 3 | 3 | 3 | 1239 | 134 | 6,61 | 868 | 2,56 | 2 | 1 | 0,791 | 0,992 | 0,906 | 1,014 | 0,732 | 0,764 | 0,956 | 1,114 | 1,04 | 0,04 | 7,5E-01 |
| O88455 | 7-dehydrocholesterol reductase OS=Mus musculus OX=10090 G | 2 | 1 | 1 | 1 | 471 | 53,9 | 8,35 | 777 | | 1 | | 0,869 | 0,84 | 0,923 | 0,945 | 0,928 | 0,858 | 0,716 | 0,99 | 1,02 | 0,02 | 7,5E-01 |
| Q9D708 | Protein S100-A16 OS=Mus musculus OX=10090 GN=S100a16 PE | 10 | 1 | 1 | 1 | 124 | 14,3 | 5,86 | 663 | | 1 | | 0,917 | 1,065 | 0,893 | 0,896 | 1,447 | 0,978 | 0,89 | 0,679 | 0,94 | -0,06 | 7,5E-01 |
| Q7TMR0 | Lysosomal Pro-X carboxypeptidase OS=Mus musculus OX=10090 | 3 | 1 | 1 | 1 | 491 | 55 | 6,79 | 248 | | 1 | | 1,162 | 0,888 | 1,016 | 1,258 | 1,618 | 0,941 | 1,12 | 0,891 | 0,95 | -0,05 | 7,5E-01 |
| Q6P9J9 | Anoctamin-6 OS=Mus musculus OX=10090 GN=Ano6 PE=1 SV=1 | 1 | 1 | 1 | 1 | 911 | 106,2 | 6,76 | 517 | | 1 | | 0,939 | 0,779 | 1,058 | 0,836 | 0,81 | 0,811 | 0,987 | 0,905 | 1,03 | 0,03 | 7,5E-01 |
| Q9CQB5 | CDGSH iron-sulfur domain-containing protein 2 OS=Mus muscul | 11 | 1 | 1 | 1 | 135 | 15,2 | 9,51 | 410 | | 1 | | 0,84 | 1,124 | 0,904 | 1,037 | 0,968 | 0,952 | 1,154 | 0,939 | 0,97 | -0,03 | 7,5E-01 |
| Q9DBF1 | Alpha-aminoadipic semialdehyde dehydrogenase OS=Mus musc | 14 | 7 | 9 | 7 | 539 | 58,8 | 7,47 | 3304 | 7,4 | 4 | 3 | 1,095 | 0,998 | 1,106 | 1,06 | 1,19 | 1,011 | 1,096 | 1,025 | 0,99 | -0,01 | 7,5E-01 |
| Q8BPM2 | Mitogen-activated protein kinase kinase kinase 5 OS=Mu | 3 | 2 | 2 | 2 | 847 | 95 | 7,83 | 858 | | 2 | | 0,881 | 0,905 | 1,01 | 1,06 | 0,963 | 0,661 | 1,08 | 1,017 | 1,04 | 0,04 | 7,5E-01 |
| P50544 | Very long-chain specific acyl-CoA dehydrogenase, mitochondrial | 20 | 11 | 14 | 11 | 656 | 70,8 | 8,75 | 6631 | 9,79 | 8 | 3 | 0,962 | 1,056 | 1,017 | 1,114 | 1,025 | 0,972 | 1,078 | 1,137 | 0,99 | -0,01 | 7,5E-01 |
| Q80VC9-14 | Isoform 14 of Calmodulin-regulated spectrin-associated protein | 14 | 13 | 16 | 1 | 1280 | 138,3 | 8,54 | 6611 | 9,17 | 9 | 4 | 1,002 | 0,813 | 0,712 | 0,927 | 0,691 | 0,798 | 0,986 | 0,862 | 1,04 | 0,04 | 7,5E-01 |
| Q9CZY3 | Ubiquitin-conjugating enzyme E2 variant 1 OS=Mus musculus O | 27 | 4 | 8 | 3 | 147 | 16,3 | 7,96 | 1882 | 9,47 | 2 | 2 | 0,875 | 0,944 | 0,879 | 1,005 | 0,971 | 0,909 | 0,847 | 1,045 | 0,98 | -0,02 | 7,5E-01 |
| P62313 | U6 snRNA-associated Sm-like protein LSm6 OS=Mus musculus O | 46 | 4 | 5 | 4 | 80 | 9,1 | 9,58 | 1854 | 0 | 3 | 1 | 0,931 | 0,945 | 0,918 | 1,034 | 0,889 | 0,871 | 1,014 | 0,995 | 1,02 | 0,02 | 7,5E-01 |
| Q6NZC7 | SEC23-interacting protein OS=Mus musculus OX=10090 GN=Sec | 1 | 1 | 1 | 1 | 998 | 110,7 | 5,94 | 479 | | 1 | | 0,864 | 1,041 | 1,709 | 1,037 | 1,041 | 1,247 | 1,275 | 1,348 | 0,95 | -0,05 | 7,5E-01 |
| Q9DCN1 | Peroxisomal NADH pyrophosphatase NUDT12 OS=Mus musculus | 2 | 1 | 1 | 1 | 462 | 51,5 | 7,12 | 676 | | 1 | | 0,826 | 0,905 | 0,959 | 1,003 | 0,826 | 1,112 | 0,818 | 1,047 | 0,97 | -0,03 | 7,6E-01 |
| Q6ZQI3 | Malectin OS=Mus musculus OX=10090 GN=Mlec PE=1 SV=2 | 8 | 3 | 5 | 3 | 291 | 32,3 | 6,05 | 1934 | 0 | 2 | 1 | 0,834 | 0,841 | 1,082 | 0,911 | 0,926 | 0,844 | 0,936 | 1,056 | 0,98 | -0,02 | 7,6E-01 |
| Q99LT0 | Protein dpy-30 homolog OS=Mus musculus OX=10090 GN=Dpy3 | 9 | 1 | 1 | 1 | 99 | 11,2 | 4,88 | 679 | | 1 | | 1,167 | 1,155 | 1,204 | 1,294 | 1,096 | 1,131 | 1,126 | 1,642 | 0,96 | -0,04 | 7,6E-01 |
| Q91VN4 | MICOS complex subunit Mic25 OS=Mus musculus OX=10090 GN | 14 | 4 | 4 | 4 | 273 | 29,8 | 8,41 | 1309 | 4,35 | 2 | 2 | 1,072 | 0,929 | 1,144 | 0,895 | 0,94 | 1,122 | 0,913 | 0,968 | 1,02 | 0,02 | 7,6E-01 |
| Q9DCT5 | Stromal cell-derived factor 2 OS=Mus musculus OX=10090 GN=S | 7 | 1 | 1 | 1 | 211 | 23,1 | 7,33 | 225 | | 1 | | 1,023 | 1,102 | 1,01 | 1,027 | 1,739 | 0,69 | 1,182 | 0,851 | 0,93 | -0,07 | 7,6E-01 |
| P60764 | Ras-related C3 botulinum toxin substrate 3 OS=Mus musculus O | 40 | 7 | 17 | 3 | 192 | 21,4 | 8,15 | 4689 | 25,39 | 6 | 5 | 0,904 | 0,94 | 1,146 | 0,905 | 0,631 | 1,057 | 1,029 | 1,027 | 1,04 | 0,04 | 7,6E-01 |
| Q8BRF7 | Sec1 family domain-containing protein 1 OS=Mus musculus OX= | 4 | 2 | 3 | 2 | 639 | 72,3 | 6,38 | | 9,7 | | 2 | 1,087 | 0,947 | 1,079 | 1,118 | 0,969 | 1,122 | 0,984 | 1,255 | 0,98 | -0,02 | 7,6E-01 |
| P70429 | Ena/VASP-like protein OS=Mus musculus OX=10090 GN=Evl PE= | 12 | 3 | 3 | 3 | 414 | 44,3 | 8,85 | 937 | 2,54 | 2 | 1 | 1,038 | 1,211 | 1,445 | 1,001 | 1,375 | 1,302 | 1,266 | 0,933 | 0,96 | -0,04 | 7,6E-01 |
| Q8BG32 | 26S proteasome non-ATPase regulatory subunit 11 OS=Mus mus | 23 | 11 | 16 | 11 | 422 | 47,4 | 6,48 | 6278 | 13,26 | 7 | 5 | 1,058 | 0,977 | 0,972 | 1,023 | 0,914 | 1,061 | 1,023 | 0,984 | 1,01 | 0,01 | 7,6E-01 |
| Q3V0G7 | GTPase-activating Rap/Ran-GAP domain-like protein 3 OS=Mus | 2 | 2 | 2 | 2 | 1038 | 115,4 | 7,17 | 710 | | 2 | | 0,717 | 0,679 | 0,71 | 0,731 | 0,638 | 0,641 | 0,781 | 0,73 | 1,02 | 0,02 | 7,6E-01 |
| B8ZXI1 | Queuine tRNA-ribosyltransferase accessory subunit 2 OS=Mus m | 3 | 1 | 1 | 1 | 415 | 46,3 | 6,16 | 196 | | 1 | | 0,747 | 0,925 | 1,536 | 1,004 | 1,069 | 1,046 | 0,899 | 0,976 | 1,06 | 0,06 | 7,6E-01 |
| O08915 | AH receptor-interacting protein OS=Mus musculus OX=10090 G | 7 | 2 | 2 | 2 | 330 | 37,6 | 6,4 | 1124 | | 2 | | 1,142 | 1,298 | 1,091 | 1,114 | 1,229 | 1,008 | 1,319 | 0,965 | 1,03 | 0,03 | 7,6E-01 |
| P62267 | 40S ribosomal protein S23 OS=Mus musculus OX=10090 GN=Rp | 27 | 6 | 13 | 6 | 143 | 15,8 | 10,49 | 7466 | 4,82 | 3 | 3 | 1,142 | 0,949 | 1,053 | 1,168 | 1,123 | 0,99 | 1,059 | 1,068 | 1,02 | 0,02 | 7,6E-01 |
| Q9CQN7 | 39S ribosomal protein L41, mitochondrial OS=Mus musculus OX | 17 | 2 | 2 | 2 | 135 | 15,3 | 9,82 | 1445 | | 2 | | 0,973 | 1,116 | 1,039 | 1,265 | 0,83 | 0,899 | 1,213 | 1,287 | 1,04 | 0,04 | 7,6E-01 |
| P61982 | 14-3-3 protein gamma OS=Mus musculus OX=10090 GN=Ywhag | 54 | 18 | 87 | 9 | 247 | 28,3 | 4,89 | 36234 | 58,06 | 11 | 11 | 0,926 | 0,852 | 1,051 | 0,88 | 0,782 | 1,109 | 0,819 | 0,891 | 1,03 | 0,03 | 7,6E-01 |
| Q80UP8 | Sodium-dependent phosphate transporter 2 OS=Mus musculus | 2 | 1 | 1 | 1 | 656 | 70,8 | 6,48 | | 2,18 | | 1 | 0,996 | 0,84 | 0,964 | 1,134 | 1,099 | 0,662 | 0,985 | 1,043 | 1,04 | 0,04 | 7,6E-01 |
| P04104 | Keratin, type II cytoskeletal 1 OS=Mus musculus OX=10090 GN= | 6 | 8 | 34 | 4 | 637 | 65,6 | 8,15 | 6857 | 67,29 | 5 | 6 | 0,727 | 1,778 | 0,653 | 0,8 | 1,478 | 1,169 | 0,859 | 0,836 | 0,91 | -0,09 | 7,6E-01 |
| P19783 | Cytochrome c oxidase subunit 4 isoform 1, mitochondrial OS=M | 41 | 7 | 18 | 7 | 169 | 19,5 | 9,23 | 8497 | 12,19 | 6 | 3 | 0,997 | 0,869 | 1,048 | 0,93 | 1,003 | 0,915 | 0,959 | 0,911 | 1,01 | 0,01 | 7,6E-01 |
| P70663 | SPARC-like protein 1 OS=Mus musculus OX=10090 GN=Sparcl1 F | 13 | 6 | 6 | 6 | 650 | 72,2 | 4,6 | 2332 | 2,2 | 5 | 1 | 0,967 | 1,044 | 1,016 | 1,088 | 0,823 | 0,995 | 1,037 | 1,166 | 1,02 | 0,02 | 7,7E-01 |
| P61080 | Ubiquitin-conjugating enzyme E2 D1 OS=Mus musculus OX=100 | 20 | 2 | 2 | 1 | 147 | 16,6 | 7,42 | 368 | 3,8 | 1 | 1 | 1,016 | 0,835 | 1,323 | 1,34 | 0,459 | 1,53 | 1,129 | 1,08 | 1,08 | 0,08 | 7,7E-01 |
| Q3UMR5 | Calcium uniporter protein, mitochondrial OS=Mus musculus OX= | 15 | 5 | 7 | 5 | 350 | 39,7 | 8,56 | 3849 | 0 | 5 | 1 | 1,03 | 0,995 | 0,862 | 0,931 | 0,782 | 0,895 | 1,065 | 0,987 | 1,02 | 0,02 | 7,7E-01 |
| P63046 | Sulfotransferase 4A1 OS=Mus musculus OX=10090 GN=Sult4a1 | 7 | 2 | 2 | 2 | 284 | 33 | 5,53 | | 5,1 | | 2 | 1,051 | 0,921 | 1,038 | 0,983 | 0,878 | 1,185 | 0,905 | 0,929 | 1,02 | 0,02 | 7,7E-01 |
| Q921L3 | Calcium load-activated calcium channel OS=Mus musculus OX=1 | 7 | 1 | 1 | 1 | 188 | 21,2 | 9,74 | 281 | | 1 | | 0,893 | 0,921 | 1,108 | 1,062 | 0,961 | 0,968 | 1,078 | 1,052 | 0,98 | -0,02 | 7,7E-01 |
| Q9JJU8 | SH3 domain-binding glutamic acid-rich-like protein OS=Mus mus | 11 | 1 | 3 | 1 | 114 | 12,8 | 4,92 | | 11,11 | | 1 | 0,858 | 1,246 | 1,028 | 1,101 | 0,849 | 1,309 | 0,919 | 0,995 | 1,04 | 0,04 | 7,7E-01 |
| Q91X72 | Hemopexin OS=Mus musculus OX=10090 GN=Hpx PE=1 SV=2 | 3 | 1 | 2 | 1 | 460 | 51,3 | 7,8 | 1168 | | 1 | | 0,978 | 0,933 | 1,027 | 1,181 | 1,305 | 0,823 | 1,163 | 0,975 | 0,97 | -0,03 | 7,7E-01 |
| Q9CY97 | RNA polymerase II subunit A C-terminal domain phosphatase SS | 7 | 1 | 1 | 1 | 194 | 22,5 | 5,21 | 205 | | 1 | | 0,961 | 0,945 | 1,334 | 1,154 | 1,064 | 0,82 | 1,469 | 1,246 | 0,96 | -0,04 | 7,7E-01 |
| P01831 | Thy-1 membrane glycoprotein OS=Mus musculus OX=10090 GN | 36 | 6 | 27 | 6 | 162 | 18,1 | 8,97 | 13275 | 17,86 | 6 | 2 | 1,035 | 1,066 | 0,895 | 1,051 | 0,995 | 1,096 | 0,969 | 1,047 | 0,99 | -0,01 | 7,7E-01 |
| Q9CQ69 | Cytochrome b-c1 complex subunit 8 OS=Mus musculus OX=1009 | 32 | 4 | 8 | 4 | 82 | 9,8 | 10,26 | 1259 | 10,65 | 1 | 3 | 1,12 | 0,861 | 1,052 | 0,924 | 0,879 | 1,015 | 1,014 | 0,966 | 1,02 | 0,02 | 7,7E-01 |
| Q05512 | Serine/threonine-protein kinase MARK2 OS=Mus musculus OX= | 7 | 4 | 5 | 3 | 776 | 86,3 | 9,67 | 2970 | | 4 | | 0,96 | 0,979 | 0,982 | 0,984 | 0,961 | 0,962 | 0,972 | 1,032 | 0,99 | -0,01 | 7,7E-01 |
| O89112 | Glutathione S-transferase LANCL1 OS=Mus musculus OX=10090 | 5 | 2 | 3 | 2 | 399 | 45,3 | 7,77 | 2073 | | 2 | | 1,066 | 1,066 | 1,012 | 0,896 | 1,056 | 1,028 | 0,974 | 1,036 | 0,99 | -0,01 | 7,7E-01 |
| O88685 | 26S proteasome regulatory subunit 6A OS=Mus musculus OX=1 | 26 | 11 | 19 | 11 | 442 | 49,5 | 5,19 | 5980 | 17,71 | 7 | 5 | 1,103 | 1,069 | 1,035 | 1,09 | 0,926 | 1,098 | 1,085 | 1,129 | 1,01 | 0,01 | 7,7E-01 |
| Q3UV70 | [Pyruvate dehydrogenase [acetyl-transferring]]-phosphatase 1, | 2 | 1 | 1 | 1 | 538 | 61,1 | 6,67 | 495 | | 1 | | 0,943 | 1,078 | 1,117 | 0,853 | 1,132 | 1,186 | 0,721 | 1,103 | 0,96 | -0,04 | 7,7E-01 |
| P43275 | Histone H1.1 OS=Mus musculus OX=10090 GN=Hist1h1a PE=1 S | 17 | 6 | 8 | 3 | 213 | 21,8 | 10,93 | 884 | 20,34 | 1 | 5 | 0,859 | 0,621 | 0,897 | 1,125 | 0,679 | 0,754 | 1,301 | 0,982 | 0,94 | -0,06 | 7,7E-01 |
| Q93092 | Transaldolase OS=Mus musculus OX=10090 GN=Taldo1 PE=1 SV | 26 | 8 | 10 | 8 | 337 | 37,4 | 7,03 | 3131 | 9,45 | 5 | 3 | 1,032 | 0,977 | 1,026 | 1,042 | 1,074 | 1,069 | 0,988 | 0,89 | 1,01 | 0,01 | 7,7E-01 |
| E9Q555 | E3 ubiquitin-protein ligase RNF213 OS=Mus musculus OX=1009 | 0 | 2 | 6 | 2 | 5152 | 584,4 | 6,8 | 1303 | | 2 | | 0,893 | 0,917 | 1,046 | 1,171 | 1,178 | 1,101 | 0,993 | 0,869 | 0,97 | -0,03 | 7,7E-01 |
| Q9JLQ2 | ARF GTPase-activating protein GIT2 OS=Mus musculus OX=1009 | 2 | 2 | 2 | 1 | 708 | 78,7 | 7,72 | 566 | 2,64 | 1 | 1 | 1,215 | 1,06 | 1,299 | 1,384 | 1,167 | 1,426 | 0,996 | 1,231 | 1,03 | 0,03 | 7,7E-01 |
| P80315 | T-complex protein 1 subunit delta OS=Mus musculus OX=10090 | 30 | 17 | 27 | 17 | 539 | 58 | 8,02 | 6878 | 34,5 | 7 | 11 | 1,016 | 0,866 | 0,913 | 0,95 | 0,94 | 0,969 | 0,873 | 0,917 | 1,01 | 0,01 | 7,7E-01 |
| Q3TFD2 | Lysophosphatidylcholine acyltransferase 1 OS=Mus musculus O | 2 | 1 | 1 | 1 | 534 | 59,7 | 6,34 | | 2,94 | | 1 | 1,099 | 1,016 | 0,971 | 0,978 | 1,042 | 0,953 | 0,97 | 1,169 | 0,98 | -0,02 | 7,7E-01 |
| Q99LD8 | N(G),N(G)-dimethylarginine dimethylaminohydrolase 2 OS=Mus | 28 | 8 | 14 | 7 | 285 | 29,6 | 6,01 | 5660 | 12,83 | 4 | 4 | 0,971 | 0,914 | 1,018 | 0,916 | 0,894 | 1,096 | 0,842 | 0,913 | 1,02 | 0,02 | 7,7E-01 |
| P68254 | 14-3-3 protein theta OS=Mus musculus OX=10090 GN=Ywhaq P | 51 | 19 | 83 | 12 | 245 | 27,8 | 4,78 | 31220 | 98,3 | 10 | 14 | 0,838 | 0,923 | 0,949 | 0,876 | 0,736 | 0,993 | 0,924 | 0,86 | 1,02 | 0,02 | 7,7E-01 |
| Q99M71 | Mammalian ependymin-related protein 1 OS=Mus musculus OX | 10 | 2 | 2 | 2 | 224 | 25,5 | 7,58 | 1297 | | 2 | | 0,83 | 0,851 | 0,963 | 0,889 | 0,993 | 0,917 | 0,953 | 0,745 | 0,98 | -0,02 | 7,7E-01 |
| P15379 | CD44 antigen OS=Mus musculus OX=10090 GN=Cd44 PE=1 SV=3 | 1 | 1 | 1 | 1 | 778 | 85,6 | 4,96 | 470 | | 1 | | 1,098 | 1,154 | 1,116 | 1,318 | 2,255 | 0,736 | 0,795 | 1,33 | 0,92 | -0,08 | 7,7E-01 |
| Q5SUR0 | Phosphoribosylformylglycinamidine synthase OS=Mus musculus | 1 | 1 | 1 | 1 | 1337 | 144,5 | 5,67 | 625 | | 1 | | 1,199 | 1,384 | 1,091 | 1,234 | 1,035 | 0,995 | 1,324 | 1,41 | 1,03 | 0,03 | 7,7E-01 |
| Q80T41 | Gamma-aminobutyric acid type B receptor subunit 2 OS=Mus m | 1 | 1 | 1 | 1 | 940 | 105,6 | 8,72 | 684 | | 1 | | 0,926 | 0,807 | 0,865 | 0,907 | 1,239 | 0,879 | 0,702 | 0,827 | 0,96 | -0,04 | 7,7E-01 |
| Q91V12 | Cytosolic acyl coenzyme A thioester hydrolase OS=Mus musculu | 20 | 8 | 13 | 8 | 381 | 42,5 | 8,68 | 3903 | 16,84 | 5 | 5 | 0,94 | 0,908 | 1,131 | 0,872 | 0,867 | 0,969 | 0,961 | 0,978 | 1,02 | 0,02 | 7,7E-01 |
| P62309 | Small nuclear ribonucleoprotein G OS=Mus musculus OX=10090 | 9 | 1 | 3 | 1 | 76 | 8,5 | 8,88 | 1816 | | 1 | | 1,018 | 0,922 | 1,038 | 1,147 | 0,827 | 1,026 | 1,229 | 0,926 | 1,03 | 0,03 | 7,7E-01 |
| P12815 | Programmed cell death protein 6 OS=Mus musculus OX=10090 | 6 | 1 | 1 | 1 | 191 | 21,9 | 5,4 | 676 | | 1 | | 1,087 | 0,732 | 1,225 | 0,966 | 0,959 | 0,938 | 1,291 | 0,982 | 0,96 | -0,04 | 7,7E-01 |
| Q9QYK9 | Calcium/calmodulin-dependent protein kinase type 1B OS=Mus | 3 | 1 | 1 | 1 | 343 | 38,5 | 6,47 | 589 | | 1 | | 0,878 | 1,034 | 0,904 | 0,791 | 0,867 | 0,869 | 1,368 | 0,688 | 0,95 | -0,05 | 7,8E-01 |
| Q8BZA9 | Fructose-2,6-bisphosphatase TIGAR OS=Mus musculus OX=1009 | 10 | 2 | 2 | 2 | 269 | 29,2 | 8,18 | 1068 | | 2 | | 0,81 | 1,062 | 0,927 | 0,949 | 0,916 | 0,764 | 1,139 | 1,044 | 0,97 | -0,03 | 7,8E-01 |
| Q8BFR4 | N-acetylglucosamine-6-sulfatase OS=Mus musculus OX=10090 G | 4 | 2 | 3 | 2 | 544 | 61,1 | 8,24 | 1206 | 2,65 | 1 | 1 | 0,931 | 0,993 | 1,016 | 1,124 | 0,829 | 1,172 | 0,935 | 1,029 | 1,02 | 0,02 | 7,8E-01 |
| Q8K1J6 | CCA tRNA nucleotidyltransferase 1, mitochondrial OS=Mus musc | 4 | 2 | 2 | 2 | 434 | 49,9 | 8,56 | 796 | 2,83 | 1 | 1 | 1,015 | 0,811 | 0,9 | 1,118 | 0,648 | 1,13 | 1,033 | 0,885 | 1,04 | 0,04 | 7,8E-01 |

| | | | | | | | | | | | | | | | | | | | | | | | |
|---|---|---|---|---|---|---|---|---|---|---|---|---|---|---|---|---|---|---|---|---|---|---|---|
| Q8BGZ1 | Hippocalcin-like protein 4 OS=Mus musculus OX=10090 GN=Hpc | 20 | 5 | 6 | 2 | 191 | 22,2 | 4,89 | 1147 | 10,35 | 2 | 3 | 0,902 | 1,062 | 0,599 | 1,016 | 0,783 | 0,944 | 0,925 | 1,069 | 0,96 | -0,04 | 7,8E-01 |
| Q91YQ5 | Dolichyl-diphosphooligosaccharide--protein glycosyltransferase | 22 | 14 | 21 | 14 | 608 | 68,5 | 6,46 | 10054 | 7,16 | 11 | 3 | 0,911 | 1,005 | 0,884 | 1,071 | 0,945 | 0,836 | 1,056 | 0,96 | 1,02 | 0,02 | 7,8E-01 |
| Q811Q9 | Choline-phosphate cytidylyltransferase B OS=Mus musculus OX= | 6 | 2 | 3 | 1 | 369 | 41,9 | 6,4 | 1872 | | 2 | | 1,068 | 0,971 | 0,992 | 0,974 | 0,9 | 1,352 | 0,987 | 0,897 | 0,97 | -0,03 | 7,8E-01 |
| Q99KK7 | Dipeptidyl peptidase 3 OS=Mus musculus OX=10090 GN=Dpp3 F | 12 | 8 | 9 | 8 | 738 | 82,8 | 5,38 | 2662 | 7,03 | 5 | 3 | 0,974 | 0,964 | 0,91 | 1,084 | 0,874 | 1,007 | 1,006 | 1,118 | 0,98 | -0,02 | 7,8E-01 |
| P40630 | Transcription factor A, mitochondrial OS=Mus musculus OX=100 | 10 | 3 | 4 | 3 | 243 | 28 | 9,69 | 1085 | 4,51 | 1 | 2 | 1,028 | 1,054 | 1,135 | 1,087 | 1,05 | 1,165 | 1,156 | 0,99 | 0,99 | -0,01 | 7,8E-01 |
| O35382 | Exocyst complex component 4 OS=Mus musculus OX=10090 GN | 4 | 3 | 4 | 3 | 975 | 110,5 | 6,49 | 2677 | | 3 | | 0,806 | 1,03 | 0,852 | 0,844 | 0,748 | 0,786 | 1,016 | 0,89 | 1,03 | 0,03 | 7,8E-01 |
| Q9Z2W8 | Glutamate receptor 4 OS=Mus musculus OX=10090 GN=Gria4 PI | 4 | 4 | 5 | 1 | 902 | 100,8 | 8,07 | 2266 | 1,83 | 3 | 1 | 0,954 | 1,016 | 1,065 | 1,096 | 1,305 | 0,775 | 1,049 | 0,859 | 1,04 | 0,04 | 7,8E-01 |
| Q91VR7 | Microtubule-associated proteins 1A/1B light chain 3A OS=Mus n | 36 | 6 | 16 | 3 | 121 | 14,3 | 8,68 | 7980 | 4,68 | 3 | 3 | 1,055 | 0,89 | 1,367 | 0,989 | 0,861 | 1,155 | 1,088 | 1,056 | 1,03 | 0,03 | 7,8E-01 |
| P62960 | Nuclease-sensitive element-binding protein 1 OS=Mus musculus | 28 | 6 | 15 | 2 | 322 | 35,7 | 9,88 | 8039 | 6,96 | 5 | 2 | 0,933 | 0,957 | 0,699 | 0,949 | 0,845 | 0,904 | 0,894 | 0,819 | 1,02 | 0,02 | 7,8E-01 |
| Q9ESU6 | Bromodomain-containing protein 4 OS=Mus musculus OX=1009 | 1 | 1 | 1 | 1 | 1400 | 155,8 | 9,19 | 455 | | 1 | | 0,803 | 0,433 | 1,239 | 0,502 | 0,821 | 1,174 | 0,612 | 0,632 | 0,92 | -0,08 | 7,8E-01 |
| Q9EQC5 | N-terminal kinase-like protein OS=Mus musculus OX=10090 GN= | 1 | 1 | 1 | 1 | 806 | 89,1 | 6,44 | 367 | | 1 | | 1,071 | 0,893 | 1,035 | 0,81 | 0,954 | 0,846 | 1,055 | 0,864 | 1,02 | 0,02 | 7,8E-01 |
| O55242 | Sigma non-opioid intracellular receptor 1 OS=Mus musculus OX= | 6 | 1 | 1 | 1 | 223 | 25,2 | 5,91 | 208 | | 1 | | 0,546 | 0,779 | 0,838 | 0,662 | 0,828 | 0,76 | 0,703 | 0,624 | 0,97 | -0,03 | 7,8E-01 |
| Q3UYC0 | Protein phosphatase 1H OS=Mus musculus OX=10090 GN=Ppm1 | 15 | 5 | 6 | 5 | 513 | 56,3 | 6,73 | 1930 | 5,08 | 4 | 2 | 0,955 | 0,816 | 0,937 | 0,88 | 0,807 | 1,018 | 0,848 | 0,85 | 1,02 | 0,02 | 7,8E-01 |
| Q9CPX6 | Ubiquitin-like-conjugating enzyme ATG3 OS=Mus musculus OX= | 14 | 5 | 7 | 5 | 314 | 35,8 | 4,72 | 4157 | | 5 | | 1,108 | 0,935 | 0,921 | 0,996 | 0,856 | 0,966 | 1,035 | 1,034 | 1,02 | 0,02 | 7,8E-01 |
| Q62245 | Son of sevenless homolog 1 OS=Mus musculus OX=10090 GN=S | 2 | 3 | 3 | 3 | 1319 | 150,8 | 6,89 | 1841 | | 3 | | 0,999 | 1,029 | 1,017 | 1,005 | 1,029 | 0,939 | 0,97 | 1,076 | 1,01 | 0,01 | 7,8E-01 |
| Q8BKE6 | Cytochrome P450 20A1 OS=Mus musculus OX=10090 GN=Cyp20 | 4 | 1 | 1 | 1 | 462 | 52,1 | 6,95 | 459 | | 1 | | 1,515 | 1,634 | 1,418 | 2,146 | 1,683 | 1,533 | 1,629 | 1,678 | 1,03 | 0,03 | 7,8E-01 |
| Q8CFV4 | Neuritin OS=Mus musculus OX=10090 GN=Nrn1 PE=1 SV=1 | 11 | 1 | 1 | 1 | 142 | 15,3 | 6,99 | 217 | | 1 | | 0,989 | 1,095 | 0,796 | 1,28 | 1,146 | 1,045 | 1,003 | 1,087 | 0,97 | -0,03 | 7,8E-01 |
| Q922D8 | C-1-tetrahydrofolate synthase, cytoplasmic OS=Mus musculus O | 16 | 14 | 17 | 14 | 935 | 101,1 | 7,14 | 7626 | 7,95 | 12 | 2 | 1,073 | 1,331 | 1,073 | 1,055 | 1,145 | 1,012 | 1,179 | 1,11 | 1,02 | 0,02 | 7,8E-01 |
| Q8BMA6 | Signal recognition particle subunit SRP68 OS=Mus musculus OX= | 3 | 2 | 2 | 2 | 625 | 70,5 | 8,57 | | 5,16 | | 2 | 1,051 | 1,141 | 1,115 | 1,029 | 1,04 | 0,965 | 1,076 | 1,193 | 1,01 | 0,01 | 7,8E-01 |
| P09602 | Non-histone chromosomal protein HMG-17 OS=Mus musculus O | 16 | 1 | 1 | 1 | 90 | 9,4 | 9,99 | | 2,05 | | 1 | 0,711 | 0,58 | 1,114 | 0,758 | 0,924 | 1,164 | 0,764 | 0,514 | 0,94 | -0,06 | 7,9E-01 |
| P70408 | Cadherin-10 OS=Mus musculus OX=10090 GN=Cdh10 PE=1 SV=3 | 2 | 1 | 1 | 1 | 788 | 88,3 | 4,94 | 725 | | 1 | | 0,928 | 1,285 | 1,086 | 1,18 | 1,176 | 1,166 | 1,066 | 1,162 | 0,98 | -0,02 | 7,9E-01 |
| Q91YN9 | BAG family molecular chaperone regulator 2 OS=Mus musculus | 5 | 1 | 1 | 1 | 210 | 23,5 | 6,42 | 756 | | 1 | | 1,062 | 1,157 | 1,143 | 1,26 | 0,542 | 1,307 | 1,453 | 1,089 | 1,05 | 0,05 | 7,9E-01 |
| P14733 | Lamin-B1 OS=Mus musculus OX=10090 GN=Lmnb1 PE=1 SV=3 | 28 | 16 | 25 | 16 | 588 | 66,7 | 5,16 | 9128 | 33,28 | 11 | 7 | 0,883 | 0,904 | 1,085 | 1,073 | 0,888 | 0,808 | 1,323 | 1,069 | 0,97 | -0,03 | 7,9E-01 |
| Q9ER72 | Cysteine--tRNA ligase, cytoplasmic OS=Mus musculus OX=10090 | 6 | 5 | 6 | 5 | 831 | 94,8 | 6,76 | 1800 | 5,52 | 2 | 3 | 1,042 | 0,986 | 0,993 | 1,126 | 0,804 | 0,906 | 1,09 | 1,233 | 1,03 | 0,03 | 7,9E-01 |
| O88952 | Protein lin-7 homolog C OS=Mus musculus OX=10090 GN=Lin7c | 50 | 7 | 14 | 7 | 197 | 21,8 | 8,43 | 3058 | 30,51 | 4 | 5 | 1,037 | 0,97 | 0,965 | 1 | 0,994 | 0,982 | 0,974 | 1,002 | 1,01 | 0,01 | 7,9E-01 |
| Q3UGP9 | Leucine-rich repeat-containing protein 58 OS=Mus musculus OX | 2 | 1 | 1 | 1 | 366 | 40,1 | 6,62 | 480 | | 1 | | 0,865 | 0,912 | 0,748 | 0,82 | 0,94 | 0,718 | 0,955 | 0,633 | 1,03 | 0,03 | 7,9E-01 |
| P61957 | Small ubiquitin-related modifier 2 OS=Mus musculus OX=10090 | 27 | 3 | 5 | 3 | 95 | 10,9 | 5,5 | 3001 | | 3 | | 0,941 | 0,993 | 1,041 | 1,063 | 0,968 | 0,989 | 1,06 | 1,062 | 0,99 | -0,01 | 7,9E-01 |
| Q8VIJ6 | Splicing factor, proline- and glutamine-rich OS=Mus musculus O | 18 | 11 | 18 | 9 | 699 | 75,4 | 9,44 | 8981 | 4,72 | 9 | 2 | 0,885 | 0,946 | 1,069 | 1,117 | 0,96 | 0,846 | 1,228 | 1,094 | 0,97 | -0,03 | 7,9E-01 |
| O35143 | ATPase inhibitor, mitochondrial OS=Mus musculus OX=10090 GI | 43 | 6 | 7 | 6 | 106 | 12,2 | 9,64 | 1262 | 7,56 | 2 | 4 | 1,075 | 0,938 | 0,949 | 0,926 | 0,631 | 1,075 | 1,186 | 0,852 | 1,04 | 0,04 | 7,9E-01 |
| O70456 | 14-3-3 protein sigma OS=Mus musculus OX=10090 GN=Sfn PE=1 | 19 | 6 | 21 | 1 | 248 | 27,7 | 4,78 | 7639 | 23,47 | 3 | 5 | 0,877 | 2,263 | 0,708 | 0,682 | 1,469 | 0,776 | 0,655 | 2,217 | 0,89 | -0,11 | 7,9E-01 |
| Q9Z0L0 | Trophoblast glycoprotein OS=Mus musculus OX=10090 GN=Tpb | 2 | 1 | 1 | 1 | 426 | 46,4 | 6,83 | | 2,4 | | 1 | 1,108 | 0,938 | 1,075 | 0,749 | 1 | 1,187 | 0,797 | 1,014 | 0,97 | -0,03 | 7,9E-01 |
| P62852 | 40S ribosomal protein S25 OS=Mus musculus OX=10090 GN=Rp | 30 | 4 | 5 | 4 | 125 | 13,7 | 10,11 | 1217 | 7,21 | 2 | 2 | 0,956 | 0,89 | 1,102 | 0,902 | 0,942 | 0,992 | 0,887 | 0,969 | 1,02 | 0,02 | 7,9E-01 |
| Q8BYI9 | Tenascin-R OS=Mus musculus OX=10090 GN=Tnr PE=1 SV=2 | 1 | 1 | 1 | 1 | 1358 | 149,5 | 4,94 | | 1,87 | | 1 | 1,201 | 0,967 | 1,293 | 1,287 | 1,338 | 1,107 | 1,094 | 1,322 | 0,98 | -0,02 | 7,9E-01 |
| Q8BMS1 | Trifunctional enzyme subunit alpha, mitochondrial OS=Mus mus | 23 | 14 | 21 | 14 | 763 | 82,6 | 9,14 | 9350 | 10,67 | 10 | 4 | 1,082 | 1,106 | 1,049 | 1,081 | 1,212 | 1,011 | 0,988 | 1,049 | 1,01 | 0,01 | 7,9E-01 |
| O54879 | High mobility group protein B3 OS=Mus musculus OX=10090 GN | 14 | 2 | 4 | 2 | 200 | 23 | 8,37 | 709 | 10,91 | 1 | 1 | 0,924 | 0,902 | 1,118 | 1,095 | 1,174 | 0,925 | 0,958 | 1,071 | 0,98 | -0,02 | 7,9E-01 |
| Q9CZC8 | Secernin-1 OS=Mus musculus OX=10090 GN=Scrn1 PE=1 SV=1 | 32 | 13 | 21 | 13 | 414 | 46,3 | 4,79 | 8701 | 20,04 | 9 | 4 | 0,986 | 1,001 | 1,046 | 1 | 1,019 | 1,064 | 0,954 | 0,964 | 1,01 | 0,01 | 7,9E-01 |
| Q6DFW4 | Nucleolar protein 58 OS=Mus musculus OX=10090 GN=Nop58 P | 5 | 2 | 2 | 2 | 536 | 60,3 | 8,34 | 1114 | | 2 | | 0,766 | 0,944 | 0,945 | 0,921 | 0,9 | 0,82 | 1,154 | 0,804 | 0,97 | -0,03 | 7,9E-01 |
| Q9CPW4 | Actin-related protein 2/3 complex subunit 5 OS=Mus musculus ( | 56 | 8 | 10 | 8 | 151 | 16,3 | 5,67 | 2160 | 19,85 | 3 | 6 | 1,003 | 1,031 | 0,929 | 0,992 | 0,924 | 1,004 | 0,978 | 1,016 | 1,01 | 0,01 | 7,9E-01 |
| P85094 | Isochorismatase domain-containing protein 2A OS=Mus muscul | 7 | 1 | 1 | 1 | 206 | 22,4 | 8,02 | 429 | | 1 | | 0,943 | 0,861 | 1,098 | 1,078 | 1,28 | 1,099 | 0,875 | 0,854 | 0,97 | -0,03 | 7,9E-01 |
| P35282 | Ras-related protein Rab-21 OS=Mus musculus OX=10090 GN=Ra | 22 | 4 | 4 | 3 | 222 | 24,1 | 7,94 | 1486 | 6,38 | 2 | 2 | 0,807 | 1,142 | 0,895 | 0,81 | 0,856 | 0,799 | 0,97 | 0,932 | 1,03 | 0,03 | 7,9E-01 |
| Q99J77 | Sialic acid synthase OS=Mus musculus OX=10090 GN=Nans PE=1 | 16 | 4 | 5 | 4 | 359 | 40 | 7,06 | 2368 | | 4 | | 0,798 | 0,7 | 0,712 | 0,707 | 0,862 | 0,66 | 0,741 | 0,708 | 0,98 | -0,02 | 7,9E-01 |
| Q9D3P8 | Plasminogen receptor (KT) OS=Mus musculus OX=10090 GN=Plg | 22 | 3 | 4 | 3 | 147 | 17,3 | 9,5 | 2445 | | 3 | | 0,897 | 1,214 | 1,064 | 1,143 | 0,819 | 0,993 | 1,381 | 0,973 | 1,04 | 0,04 | 7,9E-01 |
| Q61140 | Breast cancer anti-estrogen resistance protein 1 OS=Mus muscu | 7 | 4 | 5 | 4 | 874 | 94,2 | 5,68 | 2225 | | 4 | | 0,934 | 1,313 | 1,126 | 0,951 | 1,302 | 0,961 | 1 | 1,193 | 0,97 | -0,03 | 7,9E-01 |
| Q7TMF5 | Serpin A12 OS=Mus musculus OX=10090 GN=Serpina12 PE=1 SV | 2 | 1 | 1 | 1 | 413 | 47,6 | 9,2 | | 0 | | 1 | 0,984 | 1,171 | 0,934 | 1,085 | 1,25 | 0,754 | 1,214 | 1,093 | 0,97 | -0,03 | 7,9E-01 |
| Q9WTK7 | Serine/threonine-protein kinase STK11 OS=Mus musculus OX=1 | 6 | 2 | 2 | 2 | 436 | 49,2 | 6,84 | 350 | 2,65 | 1 | 1 | 0,882 | 0,94 | 1,085 | 0,96 | 0,936 | 0,943 | 0,916 | 1,019 | 1,01 | 0,01 | 7,9E-01 |
| Q8BNW9 | Kelch repeat and BTB domain-containing protein 11 OS=Mus mu | 4 | 3 | 3 | 3 | 633 | 67,9 | 5,39 | 359 | 2,23 | 2 | 1 | 0,834 | 1,023 | 1,002 | 0,98 | 0,772 | 1,104 | 1,189 | 0,889 | 0,97 | -0,03 | 7,9E-01 |
| Q9D358 | Low molecular weight phosphotyrosine protein phosphatase OS | 25 | 4 | 5 | 4 | 158 | 18,2 | 6,74 | 2268 | 2,15 | 3 | 1 | 1,024 | 0,939 | 1,062 | 0,958 | 0,988 | 1,074 | 0,99 | 0,971 | 0,99 | -0,01 | 7,9E-01 |
| O89026 | Roundabout homolog 1 OS=Mus musculus OX=10090 GN=Robo | 2 | 2 | 2 | 2 | 1612 | 176,3 | 6,18 | 471 | 0 | 1 | 1 | 0,948 | 1,043 | 0,964 | 1,214 | 1,153 | 0,864 | 1,032 | 1,028 | 1,02 | 0,02 | 8,0E-01 |
| Q9QXL1 | Kinesin-like protein KIF21B OS=Mus musculus OX=10090 GN=Kif | 14 | 22 | 27 | 20 | 1668 | 186,1 | 7,08 | 7288 | 33,11 | 9 | 13 | 0,97 | 0,956 | 1,048 | 0,858 | 0,884 | 0,988 | 0,991 | 0,918 | 1,01 | 0,01 | 8,0E-01 |
| Q9DCV4 | Regulator of microtubule dynamics protein 1 OS=Mus musculus | 3 | 1 | 2 | 1 | 305 | 35 | 8,7 | 869 | | 1 | | 1,11 | 0,938 | 1,007 | 1,324 | 1,067 | 1,087 | 1,138 | 1,182 | 0,98 | -0,02 | 8,0E-01 |
| Q91YJ2 | Sorting nexin-4 OS=Mus musculus OX=10090 GN=Snx4 PE=1 SV= | 14 | 5 | 7 | 5 | 450 | 51,7 | 5,8 | 780 | 11,5 | 2 | 3 | 1,075 | 0,975 | 1,287 | 1,126 | 0,934 | 1,063 | 1,21 | 1,16 | 1,02 | 0,02 | 8,0E-01 |
| Q8C2Q3 | RNA-binding protein 14 OS=Mus musculus OX=10090 GN=Rbm1 | 4 | 2 | 2 | 2 | 669 | 69,4 | 9,67 | 656 | 2,71 | 1 | 1 | 1,247 | 1,267 | 1,318 | 1,46 | 1,486 | 1,135 | 1,318 | 1,259 | 1,02 | 0,02 | 8,0E-01 |
| Q3U487 | E3 ubiquitin-protein ligase HECTD3 OS=Mus musculus OX=1009 | 1 | 1 | 1 | 1 | 861 | 97,3 | 5,47 | 396 | | 1 | | 1,182 | 1,121 | 0,984 | 1,132 | 1,03 | 0,999 | 1,238 | 1,079 | 1,02 | 0,02 | 8,0E-01 |
| P27612 | Phospholipase A-2-activating protein OS=Mus musculus OX=100 | 7 | 5 | 6 | 5 | 794 | 87,2 | 6,14 | 2004 | 5,52 | 4 | 1 | 0,941 | 0,859 | 0,981 | 0,941 | 0,839 | 1,009 | 0,948 | 0,877 | 1,01 | 0,01 | 8,0E-01 |
| Q9D6J5 | NADH dehydrogenase [ubiquinone] 1 beta subcomplex subunit | 21 | 2 | 6 | 2 | 186 | 21,9 | 6,64 | 2310 | | 2 | | 1,018 | 1,044 | 1,03 | 0,928 | 0,949 | 1,002 | 1,282 | 0,885 | 0,98 | -0,02 | 8,0E-01 |
| Q9DAS9 | Guanine nucleotide-binding protein G(I)/G(S)/G(O) subunit gam | 35 | 2 | 5 | 2 | 72 | 8 | 8,97 | 1250 | 5,49 | 1 | 1 | 1,079 | 0,888 | 0,831 | 1,066 | 1,145 | 0,928 | 0,977 | 0,903 | 0,98 | -0,02 | 8,0E-01 |
| P58468 | Protein FAM207A OS=Mus musculus OX=10090 GN=Fam207a PI | 4 | 1 | 1 | 1 | 219 | 24,8 | 11 | 309 | | 1 | | 1,106 | 0,74 | 1,102 | 1,282 | 1,141 | 0,984 | 1,062 | 1,173 | 0,97 | -0,03 | 8,0E-01 |
| Q9QXL2 | Kinesin-like protein KIF21A OS=Mus musculus OX=10090 GN=Kif | 5 | 8 | 10 | 6 | 1672 | 186,4 | 6,27 | 5229 | 2,36 | 7 | 1 | 0,913 | 1,028 | 1,075 | 1,06 | 1,109 | 0,931 | 1,073 | 0,895 | 1,02 | 0,02 | 8,0E-01 |
| Q8C419 | Probable G-protein coupled receptor 158 OS=Mus musculus OX= | 1 | 1 | 1 | 1 | 1200 | 134,3 | 8,09 | 358 | | 1 | | 1,057 | 0,926 | 1,375 | 0,924 | 1,013 | 1,368 | 1,003 | 1,044 | 0,97 | -0,03 | 8,0E-01 |
| Q9WUB0 | RanBP-type and C3HC4-type zinc finger-containing protein 1 OS | 2 | 1 | 1 | 1 | 508 | 57,5 | 6,19 | 276 | | 1 | | 0,894 | 1,001 | 1,256 | 1,055 | 0,866 | 1,094 | 1,035 | 1,111 | 1,02 | 0,02 | 8,0E-01 |
| Q8BFY9 | Transportin-1 OS=Mus musculus OX=10090 GN=Tnpo1 PE=1 SV= | 2 | 2 | 2 | 2 | 898 | 102,3 | 4,98 | 987 | | 2 | | 0,76 | 0,525 | 1,105 | 0,738 | 1,067 | 0,616 | 0,794 | 0,464 | 1,06 | 0,06 | 8,0E-01 |
| Q63880 | Carboxylesterase 3A OS=Mus musculus OX=10090 GN=Ces3a PE | 2 | 1 | 1 | 1 | 571 | 63,3 | 6,14 | | 2,65 | | 1 | 1,335 | 1,368 | 1,324 | 1,639 | 1,593 | 1,743 | 1,348 | 1,143 | 0,97 | -0,03 | 8,0E-01 |
| P23818 | Glutamate receptor 1 OS=Mus musculus OX=10090 GN=Gria1 PI | 9 | 7 | 10 | 5 | 907 | 101,5 | 7,69 | 3369 | 6,46 | 4 | 3 | 1,056 | 0,922 | 0,97 | 0,955 | 1,062 | 0,949 | 0,852 | 1,108 | 0,98 | -0,02 | 8,0E-01 |
| Q78PY7 | Staphylococcal nuclease domain-containing protein 1 OS=Mus n | 11 | 9 | 9 | 9 | 910 | 102 | 7,43 | 4162 | 2,88 | 8 | 1 | 1,035 | 1,045 | 0,963 | 1,104 | 1,093 | 0,965 | 0,963 | 1,078 | 1,01 | 0,01 | 8,0E-01 |
| Q9D7S9 | Charged multivesicular body protein 5 OS=Mus musculus OX=10 | 7 | 1 | 1 | 1 | 219 | 24,6 | 4,79 | 501 | | 1 | | 0,756 | 1,118 | 0,974 | 1,161 | 1,199 | 0,857 | 0,95 | 0,876 | 1,03 | 0,03 | 8,0E-01 |

| | | | | | | | | | | | | | | | | | | | | | | | |
|---|---|---|---|---|---|---|---|---|---|---|---|---|---|---|---|---|---|---|---|---|---|---|---|
| O55057 | Retinal rod rhodopsin-sensitive cGMP 3',5'-cyclic phosphodieste | 8 | 1 | 1 | 1 | 150 | 17,3 | 5,67 | 247 | | 1 | | 0,756 | 1,346 | 0,845 | 1,187 | 1,018 | 0,687 | 1,312 | 0,917 | 1,05 | 0,05 | 8,0E-01 |
| Q71LX4 | Talin-2 OS=Mus musculus OX=10090 GN=Tln2 PE=1 SV=3 | 14 | 25 | 29 | 22 | 2375 | 253,5 | 5,8 | 10525 | 16,38 | 17 | 8 | 1,002 | 0,958 | 0,982 | 0,974 | 0,954 | 0,972 | 1,006 | 0,969 | 1,00 | 0,00 | 8,0E-01 |
| P35585 | AP-1 complex subunit mu-1 OS=Mus musculus OX=10090 GN=A | 8 | 4 | 5 | 4 | 423 | 48,5 | 7,3 | 578 | 6,79 | 1 | 3 | 0,971 | 0,993 | 1,087 | 1,011 | 0,752 | 1,088 | 1,089 | 1,044 | 1,02 | 0,02 | 8,0E-01 |
| Q9D3S3 | Sorting nexin-29 OS=Mus musculus OX=10090 GN=Snx29 PE=1 S | 1 | 1 | 3 | 1 | 818 | 91,3 | 6,13 | 1716 | | 1 | | 1,037 | 0,976 | 0,99 | 1,129 | 1,077 | 1,094 | 1 | 0,903 | 1,01 | 0,01 | 8,0E-01 |
| Q99LR1 | Lysophosphatidylserine lipase ABHD12 OS=Mus musculus OX=1( | 12 | 4 | 5 | 4 | 398 | 45,2 | 8,72 | 2559 | | 4 | | 1,063 | 1,058 | 0,992 | 0,955 | 0,789 | 1,107 | 1,191 | 1,076 | 0,98 | -0,02 | 8,0E-01 |
| O35737 | Heterogeneous nuclear ribonucleoprotein H OS=Mus musculus ( | 10 | 4 | 4 | 3 | 449 | 49,2 | 6,3 | 1702 | 4,69 | 2 | 2 | 0,765 | 0,896 | 1,059 | 0,988 | 0,829 | 0,967 | 1,016 | 0,974 | 0,98 | -0,02 | 8,0E-01 |
| Q8BTM8 | Filamin-A OS=Mus musculus OX=10090 GN=Flna PE=1 SV=5 | 11 | 22 | 24 | 21 | 2647 | 281 | 6,04 | 8316 | 16,04 | 14 | 8 | 0,938 | 0,872 | 1,05 | 1,186 | 1,423 | 0,79 | 1,113 | 0,882 | 0,96 | -0,04 | 8,1E-01 |
| A2AN08 | E3 ubiquitin-protein ligase UBR4 OS=Mus musculus OX=10090 G | 3 | 13 | 15 | 13 | 5180 | 571,9 | 6,06 | 5567 | 9,71 | 8 | 5 | 0,996 | 1,036 | 0,979 | 0,998 | 0,985 | 0,907 | 1,052 | 1,112 | 0,99 | -0,01 | 8,1E-01 |
| Q99LI2 | Chloride channel CLIC-like protein 1 OS=Mus musculus OX=1009 | 4 | 2 | 2 | 2 | 539 | 60,6 | 5,68 | 513 | | 2 | | 0,978 | 0,905 | 1,136 | 1,14 | 0,672 | 1,018 | 0,993 | 1,326 | 1,04 | 0,04 | 8,1E-01 |
| Q8R3D1 | TBC1 domain family member 13 OS=Mus musculus OX=10090 G | 2 | 1 | 1 | 1 | 400 | 46,4 | 5,36 | 600 | | 1 | | 0,84 | 0,95 | 0,854 | 1,035 | 0,575 | 1,103 | 0,994 | 0,881 | 1,04 | 0,04 | 8,1E-01 |
| Q8BP47 | Asparagine--tRNA ligase, cytoplasmic OS=Mus musculus OX=100 | 7 | 3 | 5 | 3 | 559 | 64,2 | 5,86 | 1330 | 5,63 | 2 | 1 | 0,85 | 0,99 | 0,758 | 1,018 | 0,961 | 0,804 | 0,979 | 0,947 | 0,98 | -0,02 | 8,1E-01 |
| P39053-4 | Isoform 4 of Dynamin-1 OS=Mus musculus OX=10090 GN=Dnm1 | 36 | 35 | 71 | 2 | 864 | 97,4 | 7,17 | 25572 | 70,94 | 21 | 19 | 0,529 | 0,928 | 0,798 | 0,834 | 1,22 | 0,654 | 0,835 | 0,554 | 0,95 | -0,05 | 8,1E-01 |
| Q91Z38 | Tetratricopeptide repeat protein 1 OS=Mus musculus OX=10090 | 3 | 1 | 1 | 1 | 292 | 33,2 | 5,01 | | 2,68 | | 1 | 0,88 | 1,001 | 0,755 | 0,836 | 0,69 | 0,864 | 0,846 | 0,99 | 1,02 | 0,02 | 8,1E-01 |
| Q8VDC0 | Probable leucine--tRNA ligase, mitochondrial OS=Mus musculus | 1 | 1 | 2 | 1 | 902 | 101,4 | 8,19 | 1204 | | 1 | | 0,877 | 0,834 | 1,128 | 0,986 | 1,016 | 1,014 | 0,972 | 0,896 | 0,98 | -0,02 | 8,1E-01 |
| P59108 | Copine-2 OS=Mus musculus OX=10090 GN=Cpne2 PE=1 SV=1 | 13 | 6 | 7 | 5 | 548 | 61 | 5,96 | 3630 | 2,76 | 5 | 1 | 0,996 | 1,093 | 1,061 | 0,93 | 0,813 | 1,025 | 1,088 | 1,079 | 1,02 | 0,02 | 8,1E-01 |
| P97823 | Acyl-protein thioesterase 1 OS=Mus musculus OX=10090 GN=Ly | 6 | 1 | 1 | 1 | 230 | 24,7 | 6,62 | 569 | | 1 | | 0,892 | 0,794 | 0,94 | 1,001 | 0,802 | 0,895 | 0,964 | 1,033 | 0,98 | -0,02 | 8,1E-01 |
| Q8VCN5 | Cystathionine gamma-lyase OS=Mus musculus OX=10090 GN=C | 4 | 1 | 1 | 1 | 398 | 43,5 | 7,65 | | 2,74 | | 1 | 1,199 | 1,547 | 1,395 | 1,445 | 1,674 | 1,398 | 1,038 | 1,324 | 1,03 | 0,03 | 8,1E-01 |
| O89001 | Carboxypeptidase D OS=Mus musculus OX=10090 GN=Cpd PE=1 | 1 | 1 | 1 | 1 | 1377 | 152,3 | 6,18 | 327 | | 1 | | 1,174 | 0,892 | 0,899 | 0,814 | 0,741 | 1,21 | 0,908 | 1,049 | 0,97 | -0,03 | 8,1E-01 |
| P83741 | Serine/threonine-protein kinase WNK1 OS=Mus musculus OX=1 | 2 | 5 | 6 | 2 | 2377 | 250,8 | 6,43 | 1794 | 6,94 | 2 | 3 | 1,106 | 0,828 | 0,932 | 1,002 | 0,677 | 1,073 | 1,167 | 1,076 | 0,97 | -0,03 | 8,1E-01 |
| P52623 | Uridine-cytidine kinase 1 OS=Mus musculus OX=10090 GN=Uck1 | 4 | 1 | 1 | 1 | 277 | 31 | 7,88 | 284 | | 1 | | 0,689 | 1,027 | 0,783 | 0,936 | 0,847 | 0,606 | 1,138 | 0,979 | 0,96 | -0,04 | 8,1E-01 |
| Q9EPR5 | VPS10 domain-containing receptor SorCS2 OS=Mus musculus O | 1 | 1 | 1 | 1 | 1159 | 128,8 | 7,34 | 533 | | 1 | | 0,933 | 0,661 | 1,917 | 1,113 | 1,236 | 1,415 | 0,992 | 1,263 | 0,94 | -0,06 | 8,1E-01 |
| P50580 | Proliferation-associated protein 2G4 OS=Mus musculus OX=100 | 28 | 10 | 13 | 10 | 394 | 43,7 | 6,86 | 6549 | 5,09 | 9 | 2 | 1,039 | 1,125 | 1,05 | 1,078 | 1,029 | 1,133 | 1,008 | 1,088 | 1,01 | 0,01 | 8,1E-01 |
| Q8K183 | Pyridoxal kinase OS=Mus musculus OX=10090 GN=Pdxk PE=1 SV | 16 | 4 | 7 | 4 | 312 | 35 | 6,29 | 2914 | 2,57 | 3 | 1 | 0,947 | 0,829 | 0,828 | 0,898 | 0,901 | 0,944 | 0,818 | 0,794 | 1,01 | 0,01 | 8,1E-01 |
| P05202 | Aspartate aminotransferase, mitochondrial OS=Mus musculus O | 44 | 19 | 41 | 19 | 430 | 47,4 | 9 | 15607 | 48,74 | 12 | 9 | 1,015 | 1,017 | 1,057 | 0,946 | 0,984 | 1,071 | 0,95 | 0,996 | 1,01 | 0,01 | 8,1E-01 |
| Q3UUG6 | TBC1 domain family member 24 OS=Mus musculus OX=10090 G | 12 | 6 | 6 | 6 | 561 | 63,2 | 7,24 | 3018 | 3,05 | 5 | 1 | 0,925 | 1,036 | 1,027 | 1,006 | 0,997 | 0,899 | 1,063 | 1,083 | 0,99 | -0,01 | 8,1E-01 |
| Q5SSM3 | Rho GTPase-activating protein 44 OS=Mus musculus OX=10090 | 2 | 1 | 2 | 1 | 814 | 88,9 | 6,6 | 925 | | 1 | | 1,101 | 1,109 | 1,097 | 1,008 | 1,327 | 1,252 | 0,973 | 0,873 | 0,98 | -0,02 | 8,1E-01 |
| Q9CY58 | Plasminogen activator inhibitor 1 RNA-binding protein OS=Mus | 18 | 6 | 7 | 6 | 407 | 44,7 | 8,54 | 2283 | 7,29 | 4 | 3 | 1,001 | 1,011 | 0,977 | 0,971 | 0,964 | 0,995 | 1,028 | 0,954 | 1,00 | 0,00 | 8,1E-01 |
| O70475 | UDP-glucose 6-dehydrogenase OS=Mus musculus OX=10090 GN | 3 | 1 | 1 | 1 | 493 | 54,8 | 7,56 | 768 | | 1 | | 0,824 | 1,152 | 1,093 | 1,101 | 1,405 | 1,12 | 0,888 | 0,897 | 0,97 | -0,03 | 8,1E-01 |
| Q9R0N0 | Galactokinase OS=Mus musculus OX=10090 GN=Galk1 PE=1 SV= | 6 | 2 | 3 | 2 | 392 | 42,3 | 5,26 | 1657 | | 2 | | 1,176 | 1,111 | 1,346 | 1,365 | 1,056 | 1,565 | 1,305 | 1,194 | 0,98 | -0,02 | 8,1E-01 |
| Q9CR98 | Protein FAM136A OS=Mus musculus OX=10090 GN=Fam136a PE | 13 | 2 | 2 | 2 | 138 | 15,7 | 7,61 | 833 | 3,06 | 1 | 1 | 1,122 | 0,831 | 0,886 | 0,958 | 1,06 | 0,827 | 0,987 | 0,84 | 1,02 | 0,02 | 8,2E-01 |
| Q6A4J8 | Ubiquitin carboxyl-terminal hydrolase 7 OS=Mus musculus OX=1 | 1 | 1 | 1 | 1 | 1103 | 128,4 | 5,5 | 443 | | 1 | | 0,935 | 0,933 | 0,985 | 0,755 | 0,611 | 1,07 | 1,018 | 0,795 | 1,03 | 0,03 | 8,2E-01 |
| Q9WUZ9 | Ectonucleoside triphosphate diphosphohydrolase 5 OS=Mus mu | 3 | 1 | 1 | 1 | 427 | 47,1 | 5,29 | 203 | | 1 | | 1,491 | 1,625 | 1,378 | 1,549 | 1,074 | 1,416 | 1,704 | 1,696 | 1,03 | 0,03 | 8,2E-01 |
| P53996 | Cellular nucleic acid-binding protein OS=Mus musculus OX=1009 | 20 | 3 | 3 | 3 | 178 | 19,6 | 7,58 | 1041 | 2,39 | 2 | 1 | 0,824 | 0,763 | 0,852 | 0,959 | 0,834 | 0,976 | 0,861 | 0,783 | 0,98 | -0,02 | 8,2E-01 |
| P56380 | Bis(5'-nucleosyl)-tetraphosphatase [asymmetrical] OS=Mus mus | 15 | 2 | 2 | 2 | 147 | 17 | 6,28 | 1709 | | 2 | | 0,9 | 0,928 | 0,931 | 1,106 | 0,785 | 1,039 | 1,001 | 0,969 | 1,02 | 0,02 | 8,2E-01 |
| Q9JJW6 | Aly/REF export factor 2 OS=Mus musculus OX=10090 GN=Alyref | 13 | 2 | 3 | 2 | 218 | 23,7 | 10,04 | 2029 | | 2 | | 0,826 | 0,897 | 0,959 | 0,984 | 0,728 | 1,091 | 0,857 | 1,082 | 0,98 | -0,02 | 8,2E-01 |
| Q3UHA3 | Spatacsin OS=Mus musculus OX=10090 GN=Spg11 PE=1 SV=3 | 0 | 1 | 1 | 1 | 2430 | 273,8 | 5,94 | 309 | | 1 | | 1,225 | 1,205 | 0,774 | 1,247 | 1,19 | 0,886 | 1,26 | 1,253 | 0,97 | -0,03 | 8,2E-01 |
| B9EJ80 | PDZ domain-containing protein 8 OS=Mus musculus OX=10090 ( | 2 | 2 | 2 | 1 | 1147 | 127,7 | 6,04 | 889 | | 2 | | 0,786 | 0,88 | 1,212 | 0,681 | 0,803 | 1,118 | 0,897 | 0,869 | 0,97 | -0,03 | 8,2E-01 |
| P10605 | Cathepsin B OS=Mus musculus OX=10090 GN=Ctsb PE=1 SV=2 | 21 | 6 | 10 | 6 | 339 | 37,3 | 5,91 | 2518 | 17,74 | 3 | 3 | 1,014 | 1,056 | 0,946 | 1,242 | 1,021 | 1,208 | 1,013 | 0,934 | 1,02 | 0,02 | 8,2E-01 |
| O55137 | Acyl-coenzyme A thioesterase 1 OS=Mus musculus OX=10090 G | 14 | 5 | 8 | 2 | 419 | 46,1 | 6,58 | 2205 | 11,72 | 3 | 3 | 0,992 | 1,297 | 0,856 | 1,054 | 1,084 | 0,87 | 1,184 | 1,173 | 0,97 | -0,03 | 8,2E-01 |
| Q99N95 | 39S ribosomal protein L3, mitochondrial OS=Mus musculus OX= | 12 | 4 | 6 | 4 | 348 | 39,1 | 9,55 | 1879 | 4,92 | 2 | 2 | 1,129 | 0,948 | 1,108 | 1,168 | 0,991 | 1,046 | 1,165 | 1,219 | 0,98 | -0,02 | 8,2E-01 |
| Q9Z0H4 | CUGBP Elav-like family member 2 OS=Mus musculus OX=10090 | 4 | 2 | 5 | 2 | 508 | 54,2 | 8,76 | 2201 | 2,69 | 1 | 2 | 1,2 | 0,995 | 0,985 | 1,076 | 1,076 | 0,964 | 1,047 | 1,113 | 1,01 | 0,01 | 8,2E-01 |
| P15116 | Cadherin-2 OS=Mus musculus OX=10090 GN=Cdh2 PE=1 SV=2 | 6 | 4 | 7 | 4 | 906 | 99,7 | 4,78 | 2795 | 5,71 | 3 | 1 | 1,04 | 1,185 | 0,94 | 1,219 | 1,141 | 0,986 | 1,221 | 1,113 | 0,98 | -0,02 | 8,2E-01 |
| Q9Z127 | Large neutral amino acids transporter small subunit 1 OS=Mus n | 6 | 3 | 3 | 3 | 512 | 55,8 | 7,9 | 1833 | | 3 | | 1,244 | 1,081 | 1,114 | 1,246 | 1,535 | 1,032 | 0,956 | 1,029 | 1,03 | 0,03 | 8,2E-01 |
| Q9CZT8 | Ras-related protein Rab-3B OS=Mus musculus OX=10090 GN=Ra | 27 | 5 | 10 | 3 | 219 | 24,7 | 5,11 | 5113 | 2,07 | 5 | 1 | 1 | 1,165 | 1,028 | 0,972 | 1,02 | 1,111 | 0,96 | 1,024 | 1,01 | 0,01 | 8,2E-01 |
| Q9DBG3 | AP-2 complex subunit beta OS=Mus musculus OX=10090 GN=Ap | 21 | 21 | 44 | 8 | 937 | 104,5 | 5,38 | 17216 | 38,86 | 14 | 10 | 1,029 | 1,018 | 1,01 | 0,972 | 0,928 | 1,047 | 0,95 | 1,069 | 1,01 | 0,01 | 8,2E-01 |
| Q9JHK4 | Geranylgeranyl transferase type-2 subunit alpha OS=Mus muscu | 2 | 1 | 1 | 1 | 567 | 64,9 | 5,8 | 665 | | 1 | | 0,862 | 0,761 | 0,986 | 1,053 | 0,728 | 1,049 | 1,035 | 0,943 | 0,98 | -0,02 | 8,2E-01 |
| Q3UEG6 | Alanine--glyoxylate aminotransferase 2, mitochondrial OS=Mus | 2 | 1 | 1 | 1 | 513 | 57,1 | 7,93 | 338 | | 1 | | 1,278 | 0,901 | 1,12 | 0,939 | 1,386 | 1,136 | 0,929 | 0,919 | 0,97 | -0,03 | 8,2E-01 |
| O08739 | AMP deaminase 3 OS=Mus musculus OX=10090 GN=Ampd3 PE= | 1 | 1 | 1 | 1 | 766 | 88,6 | 7,33 | | 2,56 | | 1 | 1,248 | 1,478 | 1,081 | 1,176 | 1,042 | 1,203 | 1,227 | 1,405 | 1,02 | 0,02 | 8,2E-01 |
| P22682 | E3 ubiquitin-protein ligase CBL OS=Mus musculus OX=10090 GN | 2 | 1 | 1 | 1 | 913 | 100,5 | 6,67 | 749 | | 1 | | 1,104 | 1,021 | 0,805 | 1,025 | 1,038 | 0,889 | 0,949 | 1,011 | 1,02 | 0,02 | 8,2E-01 |
| O70318 | Band 4.1-like protein 2 OS=Mus musculus OX=10090 GN=Epb41 | 8 | 6 | 7 | 5 | 988 | 109,9 | 5,43 | 4852 | | 6 | | 0,931 | 0,931 | 0,935 | 0,881 | 0,828 | 0,785 | 1,038 | 0,97 | 1,02 | 0,02 | 8,2E-01 |
| Q61033 | Lamina-associated polypeptide 2, isoforms alpha/zeta OS=Mus r | 12 | 5 | 7 | 1 | 693 | 75,1 | 8,05 | 2922 | 2,95 | 4 | 1 | 0,882 | 0,678 | 0,801 | 0,975 | 0,392 | 0,74 | 1,195 | 0,843 | 1,05 | 0,05 | 8,2E-01 |
| Q80YS6 | Actin filament-associated protein 1 OS=Mus musculus OX=1009( | 5 | 3 | 4 | 3 | 731 | 80,6 | 8,68 | 644 | 8,48 | 1 | 2 | 1,032 | 1,131 | 0,853 | 1,235 | 1,01 | 1,018 | 1,238 | 1,076 | 0,98 | -0,02 | 8,2E-01 |
| Q99LC3 | NADH dehydrogenase [ubiquinone] 1 alpha subcomplex subunit | 15 | 6 | 9 | 6 | 355 | 40,6 | 7,78 | 2141 | 17,06 | 2 | 4 | 1,135 | 0,948 | 1,218 | 1,073 | 1,015 | 1,257 | 1,068 | 1,106 | 0,98 | -0,02 | 8,2E-01 |
| Q7TMS5 | ATP-binding cassette sub-family G member 2 OS=Mus musculus | 2 | 1 | 1 | 1 | 657 | 72,9 | 8,54 | 732 | | 1 | | 0,951 | 1,149 | 0,92 | 0,868 | 0,951 | 0,84 | 0,938 | 1,085 | 1,02 | 0,02 | 8,2E-01 |
| G5E829 | Plasma membrane calcium-transporting ATPase 1 OS=Mus musc | 30 | 36 | 79 | 19 | 1220 | 134,7 | 5,91 | 28209 | 71,8 | 25 | 15 | 0,981 | 0,949 | 0,982 | 0,855 | 0,891 | 0,97 | 0,953 | 0,987 | 0,99 | -0,01 | 8,2E-01 |
| P26369 | Splicing factor U2AF 65 kDa subunit OS=Mus musculus OX=1009 | 6 | 2 | 2 | 2 | 475 | 53,5 | 9,09 | 977 | | 2 | | 0,911 | 1,09 | 1,038 | 1,084 | 0,888 | 0,98 | 1,093 | 1,101 | 1,02 | 0,02 | 8,2E-01 |
| Q08331 | Calretinin OS=Mus musculus OX=10090 GN=Calb2 PE=1 SV=3 | 22 | 6 | 9 | 6 | 271 | 31,4 | 5,02 | 2486 | 11,2 | 2 | 5 | 0,912 | 1,014 | 1,02 | 1,08 | 1,062 | 1,023 | 0,986 | 0,911 | 1,01 | 0,01 | 8,2E-01 |
| Q6PGN3 | Serine/threonine-protein kinase DCLK2 OS=Mus musculus OX=1 | 10 | 7 | 7 | 6 | 756 | 82,9 | 8,44 | 3681 | 2,3 | 6 | 1 | 0,955 | 0,928 | 1,228 | 1,144 | 1,387 | 0,905 | 1,044 | 1,036 | 0,97 | -0,03 | 8,2E-01 |
| Q9CXT8 | Mitochondrial-processing peptidase subunit beta OS=Mus musc | 7 | 3 | 4 | 3 | 489 | 54,6 | 6,99 | 1570 | 2,28 | 2 | 1 | 0,982 | 1,166 | 0,983 | 1,123 | 1,19 | 0,701 | 1,165 | 1,084 | 1,03 | 0,03 | 8,2E-01 |
| Q3UH66-5 | Isoform 5 of Serine/threonine-protein kinase WNK2 OS=Mus mu | 9 | 14 | 16 | 12 | 2213 | 233,9 | 6,1 | 6532 | 12,32 | 8 | 6 | 0,992 | 0,976 | 1,025 | 1,003 | 0,965 | 1,039 | 0,935 | 1,091 | 0,99 | -0,01 | 8,3E-01 |
| Q8BK64 | Activator of 90 kDa heat shock protein ATPase homolog 1 OS=M | 11 | 4 | 6 | 4 | 338 | 38,1 | 5,53 | 2067 | 2,61 | 2 | 2 | 0,954 | 0,862 | 1,071 | 0,983 | 0,854 | 1,051 | 0,991 | 0,918 | 1,01 | 0,01 | 8,3E-01 |
| Q8BH24 | Transmembrane 9 superfamily member 4 OS=Mus musculus OX | 1 | 1 | 1 | 1 | 643 | 74,6 | 7,23 | | 1,94 | | 1 | 0,96 | 1,228 | 1,441 | 1,061 | 1,578 | 0,94 | 1,033 | 0,97 | 1,04 | 0,04 | 8,3E-01 |
| A2AGT5 | Cytoskeleton-associated protein 5 OS=Mus musculus OX=10090 | 8 | 16 | 20 | 16 | 2032 | 225,5 | 7,96 | 5116 | 17,94 | 8 | 8 | 1,084 | 0,884 | 1,127 | 0,972 | 0,966 | 1,058 | 0,976 | 1,013 | 1,01 | 0,01 | 8,3E-01 |
| P62717 | 60S ribosomal protein L18a OS=Mus musculus OX=10090 GN=Rp | 22 | 5 | 8 | 5 | 176 | 20,7 | 10,71 | 2579 | 9,4 | 3 | 3 | 0,909 | 0,839 | 0,947 | 1,015 | 1,011 | 0,92 | 0,871 | 0,951 | 0,99 | -0,01 | 8,3E-01 |
| P28651 | Carbonic anhydrase-related protein OS=Mus musculus OX=1009 | 15 | 3 | 3 | 3 | 291 | 33,1 | 4,78 | 1374 | | 3 | | 0,85 | 1,047 | 0,747 | 1,026 | 0,938 | 0,803 | 0,837 | 1,196 | 0,97 | -0,03 | 8,3E-01 |

| | | | | | | | | | | | | | | | | | | | | | | | |
|---|---|---|---|---|---|---|---|---|---|---|---|---|---|---|---|---|---|---|---|---|---|---|---|
| P12658 | Calbindin OS=Mus musculus OX=10090 GN=Calb1 PE=1 SV=2 | 10 | 2 | 4 | 2 | 261 | 30 | 4,83 | | 12,77 | | 2 | 0,959 | 1,198 | 1,1 | 0,95 | 1,108 | 0,936 | 1,051 | 1,184 | 0,98 | -0,02 | 8,3E-01 |
| P70414 | Sodium/calcium exchanger 1 OS=Mus musculus OX=10090 GN=S | 10 | 9 | 16 | 9 | 970 | 108 | 5 | 5289 | 11,17 | 5 | 4 | 1,029 | 0,999 | 0,921 | 0,972 | 1,037 | 0,934 | 0,923 | 0,995 | 1,01 | 0,01 | 8,3E-01 |
| Q8BWT5 | Disco-interacting protein 2 homolog A OS=Mus musculus OX=10 | 3 | 5 | 5 | 5 | 1523 | 165,1 | 7,61 | 1531 | 5,01 | 3 | 2 | 1,05 | 0,889 | 1,215 | 1,047 | 1,279 | 1,011 | 0,993 | 1,005 | 0,98 | -0,02 | 8,3E-01 |
| Q9Z0N1 | Eukaryotic translation initiation factor 2 subunit 3, X-linked OS=I | 22 | 8 | 12 | 8 | 472 | 51 | 8,4 | 5436 | 8,42 | 5 | 3 | 1,071 | 1,08 | 1,024 | 1,052 | 0,985 | 0,994 | 1,034 | 1,173 | 1,01 | 0,01 | 8,3E-01 |
| Q8CH72 | E3 ubiquitin-protein ligase TRIM32 OS=Mus musculus OX=10090 | 8 | 4 | 4 | 4 | 655 | 72 | 6,9 | 1544 | 0 | 3 | 1 | 1,073 | 1,019 | 1,598 | 1,208 | 1,005 | 1,408 | 1,502 | 1,141 | 0,97 | -0,03 | 8,3E-01 |
| Q9WTQ5 | A-kinase anchor protein 12 OS=Mus musculus OX=10090 GN=Ak | 22 | 25 | 41 | 25 | 1684 | 180,6 | 4,44 | 18907 | 40,47 | 17 | 10 | 1,064 | 1,084 | 0,921 | 1,073 | 0,914 | 1,068 | 1,051 | 1,061 | 1,01 | 0,01 | 8,3E-01 |
| Q8VHG2 | Angiomotin OS=Mus musculus OX=10090 GN=Amot PE=1 SV=3 | 1 | 2 | 2 | 1 | 1126 | 120,8 | 7,5 | 432 | 2,26 | 1 | 1 | 1,03 | 1,001 | 1,343 | 1,277 | 1,303 | 1,063 | 1,2 | 0,984 | 1,02 | 0,02 | 8,3E-01 |
| Q64337 | Sequestosome-1 OS=Mus musculus OX=10090 GN=Sqstm1 PE=1 | 7 | 2 | 2 | 2 | 442 | 48,1 | 5,21 | 1019 | | 2 | | 0,901 | 0,813 | 0,807 | 0,814 | 0,887 | 0,759 | 0,91 | 0,734 | 1,01 | 0,01 | 8,3E-01 |
| P50247 | Adenosylhomocysteinase OS=Mus musculus OX=10090 GN=Ahc | 22 | 9 | 10 | 9 | 432 | 47,7 | 6,54 | 4576 | 4,86 | 7 | 2 | 0,969 | 1,012 | 1,033 | 1,057 | 0,909 | 1,047 | 1,083 | 0,994 | 1,01 | 0,01 | 8,3E-01 |
| P97291 | Cadherin-8 OS=Mus musculus OX=10090 GN=Cdh8 PE=1 SV=2 | 2 | 1 | 1 | 1 | 799 | 88,2 | 4,74 | | 3,79 | | 1 | 1,23 | 1,135 | 1,084 | 1,076 | 1,137 | 1,086 | 0,983 | 1,406 | 0,98 | -0,02 | 8,3E-01 |
| Q9CY50 | Translocon-associated protein subunit alpha OS=Mus musculus | 4 | 1 | 2 | 1 | 286 | 32 | 4,45 | 1209 | | 1 | | 0,851 | 0,715 | 0,885 | 0,915 | 0,915 | 0,817 | 0,886 | 0,795 | 0,99 | -0,01 | 8,3E-01 |
| Q6PIC6 | Sodium/potassium-transporting ATPase subunit alpha-3 OS=Mu | 49 | 53 | 280 | 36 | 1013 | 111,6 | 5,41 | 120845 | 240,56 | 36 | 28 | 1,006 | 0,956 | 1,045 | 0,912 | 0,944 | 1,058 | 0,972 | 0,979 | 0,99 | -0,01 | 8,3E-01 |
| Q8C6G8 | WD repeat-containing protein 26 OS=Mus musculus OX=10090 ( | 3 | 2 | 3 | 2 | 641 | 70,5 | 6,16 | 1495 | | 2 | | 0,8 | 1,079 | 0,622 | 1,271 | 0,593 | 0,83 | 0,861 | 1,303 | 1,05 | 0,05 | 8,3E-01 |
| Q3TZM9 | GDP-Man:Man(3)GlcNAc(2)-PP-Dol alpha-1,2-mannosyltransfera | 2 | 1 | 1 | 1 | 492 | 55,2 | 8,29 | 295 | | 1 | | 1,059 | 1,361 | 1,325 | 1,347 | 1,495 | 1,157 | 1,105 | 1,235 | 1,02 | 0,02 | 8,3E-01 |
| Q9D8U8 | Sorting nexin-5 OS=Mus musculus OX=10090 GN=Snx5 PE=1 SV= | 6 | 2 | 2 | 2 | 404 | 46,8 | 6,62 | 542 | 0 | 1 | 1 | 0,938 | 1,349 | 1,042 | 1,488 | 1,571 | 1,01 | 1,406 | 1,001 | 0,97 | -0,03 | 8,3E-01 |
| Q9Z2V5 | Histone deacetylase 6 OS=Mus musculus OX=10090 GN=Hdac6 I | 4 | 4 | 5 | 4 | 1149 | 125,7 | 5,78 | 2050 | 2,11 | 3 | 1 | 0,919 | 0,97 | 0,922 | 0,897 | 0,994 | 0,889 | 0,933 | 0,916 | 0,99 | -0,01 | 8,3E-01 |
| O88492 | Perilipin-4 OS=Mus musculus OX=10090 GN=Plin4 PE=1 SV=2 | 14 | 3 | 3 | 3 | 1403 | 139,3 | 8,59 | 1666 | | 3 | | 2,35 | 1,574 | 1,418 | 1,398 | 2,758 | 1,034 | 1,246 | 2,108 | 0,94 | -0,06 | 8,3E-01 |
| Q8C052 | Microtubule-associated protein 1S OS=Mus musculus OX=10090 | 9 | 8 | 9 | 8 | 973 | 102,9 | 7,02 | 3824 | 8,34 | 5 | 3 | 0,869 | 0,862 | 0,87 | 0,91 | 0,849 | 0,931 | 0,898 | 0,807 | 1,01 | 0,01 | 8,3E-01 |
| Q91WT9 | Cystathionine beta-synthase OS=Mus musculus OX=10090 GN=( | 9 | 3 | 4 | 3 | 561 | 61,5 | 6,48 | 1930 | 2,61 | 2 | 1 | 1,259 | 1,208 | 1,321 | 1,314 | 1,227 | 1,308 | 1,155 | 1,365 | 1,01 | 0,01 | 8,3E-01 |
| Q9DCT2 | NADH dehydrogenase [ubiquinone] iron-sulfur protein 3, mitoch | 25 | 6 | 9 | 6 | 263 | 30,1 | 7,17 | 5399 | | 6 | | 0,985 | 1,06 | 1,101 | 1,052 | 1,038 | 0,962 | 1,129 | 1,032 | 1,01 | 0,01 | 8,3E-01 |
| A2AG50 | MAP7 domain-containing protein 2 OS=Mus musculus OX=1009 | 2 | 2 | 2 | 1 | 781 | 86 | 8,81 | | 1,97 | | 2 | 0,92 | 1,106 | 0,907 | 1,08 | 0,622 | 0,957 | 1,208 | 1,105 | 1,03 | 0,03 | 8,3E-01 |
| P35700 | Peroxiredoxin-1 OS=Mus musculus OX=10090 GN=Prdx1 PE=1 S\ | 74 | 16 | 46 | 14 | 199 | 22,2 | 8,12 | 15947 | 55,16 | 11 | 8 | 0,924 | 0,978 | 1,007 | 0,966 | 0,959 | 1,007 | 0,95 | 0,939 | 1,01 | 0,01 | 8,3E-01 |
| O88342 | WD repeat-containing protein 1 OS=Mus musculus OX=10090 G | 47 | 22 | 43 | 22 | 606 | 66,4 | 6,6 | 14349 | 39,92 | 16 | 11 | 1,05 | 0,976 | 1,034 | 1,027 | 0,951 | 1,105 | 0,982 | 1,085 | 0,99 | -0,01 | 8,3E-01 |
| Q9EPN1 | Neurobeachin OS=Mus musculus OX=10090 GN=Nbea PE=1 SV=1 | 10 | 25 | 34 | 25 | 2936 | 326,5 | 6,2 | 10829 | 42,58 | 15 | 12 | 1,017 | 1,019 | 1,033 | 1,018 | 1,064 | 1,014 | 0,996 | 1,026 | 1,00 | 0,00 | 8,3E-01 |
| Q91VC3 | Eukaryotic initiation factor 4A-III OS=Mus musculus OX=10090 G | 21 | 8 | 10 | 7 | 411 | 46,8 | 6,73 | 2938 | 5,55 | 5 | 4 | 0,866 | 1,048 | 1,021 | 1,151 | 1,023 | 0,958 | 1,046 | 1,005 | 1,01 | 0,01 | 8,3E-01 |
| P48725 | Pericentrin OS=Mus musculus OX=10090 GN=Pcnt PE=1 SV=2 | 0 | 1 | 1 | 1 | 2898 | 329,3 | 5,36 | 376 | | 1 | | 0,895 | 1,228 | 1,168 | 1,019 | 0,996 | 1,096 | 1,039 | 1,11 | 1,02 | 0,02 | 8,4E-01 |
| B1AS29 | Glutamate receptor ionotropic, kainate 3 OS=Mus musculus OX= | 3 | 2 | 2 | 2 | 919 | 104 | 7,52 | 417 | 1,78 | 1 | 1 | 1,024 | 1,196 | 0,976 | 1,027 | 1,235 | 0,85 | 1,036 | 1,188 | 0,98 | -0,02 | 8,4E-01 |
| Q9JKB3 | Y-box-binding protein 3 OS=Mus musculus OX=10090 GN=Ybx3 | 20 | 6 | 11 | 2 | 361 | 38,8 | 9,69 | 5858 | 2,54 | 5 | 1 | 0,923 | 1,106 | 0,953 | 1,03 | 0,908 | 1,025 | 1,225 | 0,926 | 0,98 | -0,02 | 8,4E-01 |
| Q80X85 | 28S ribosomal protein S7, mitochondrial OS=Mus musculus OX= | 8 | 2 | 2 | 2 | 242 | 28 | 9,94 | 627 | 3,31 | 1 | 1 | 1,117 | 0,843 | 1,165 | 1,302 | 1,137 | 1,182 | 1,068 | 1,125 | 0,98 | -0,02 | 8,4E-01 |
| Q8K009 | Mitochondrial 10-formyltetrahydrofolate dehydrogenase OS=M | 7 | 6 | 6 | 3 | 923 | 101,5 | 6,29 | 2464 | 5,11 | 4 | 2 | 0,897 | 0,997 | 0,957 | 0,932 | 1,104 | 0,979 | 0,919 | 0,833 | 0,99 | -0,01 | 8,4E-01 |
| P36916 | Guanine nucleotide-binding protein-like 1 OS=Mus musculus OX | 10 | 5 | 6 | 5 | 607 | 68,7 | 5,68 | 1163 | 7,52 | 2 | 3 | 0,993 | 0,947 | 1,075 | 1,16 | 1,274 | 0,969 | 0,902 | 0,947 | 1,02 | 0,02 | 8,4E-01 |
| Q9QZE7 | Translin-associated protein X OS=Mus musculus OX=10090 GN=1 | 10 | 3 | 4 | 3 | 290 | 32,9 | 6,55 | 2226 | | 3 | | 1,039 | 1,369 | 0,829 | 1,165 | 1,027 | 1,179 | 1,177 | 0,907 | 1,03 | 0,03 | 8,4E-01 |
| Q61738 | Integrin alpha-7 OS=Mus musculus OX=10090 GN=Itga7 PE=1 SV | 1 | 1 | 1 | 1 | 1179 | 129,2 | 5,71 | 575 | | 1 | | 1,315 | 1,298 | 1,314 | 1,45 | 1,367 | 1,341 | 1,542 | 1,195 | 0,99 | -0,01 | 8,4E-01 |
| Q61029 | Lamina-associated polypeptide 2, isoforms beta/delta/epsilon/g | 18 | 5 | 7 | 1 | 452 | 50,3 | 9,45 | 3073 | 2,95 | 4 | 1 | 0,868 | 0,827 | 1,187 | 1,3 | 0,692 | 0,75 | 1,47 | 1,087 | 1,05 | 0,05 | 8,4E-01 |
| P68433 | Histone H3.1 OS=Mus musculus OX=10090 GN=Hist1h3a PE=1 S | 38 | 9 | 18 | 9 | 136 | 15,4 | 11,12 | 6895 | 13,46 | 5 | 4 | 0,894 | 0,771 | 1,149 | 1,266 | 1,15 | 0,845 | 1,022 | 0,953 | 1,03 | 0,03 | 8,4E-01 |
| P62334 | 26S proteasome regulatory subunit 10B OS=Mus musculus OX=1 | 35 | 13 | 21 | 13 | 389 | 44,1 | 7,49 | 7928 | 25,2 | 7 | 6 | 1,073 | 1,063 | 1,053 | 1,028 | 1,076 | 1,018 | 1,103 | 1,038 | 1,00 | 0,00 | 8,4E-01 |
| Q9CQR4 | Acyl-coenzyme A thioesterase 13 OS=Mus musculus OX=10090 ( | 21 | 3 | 4 | 3 | 140 | 15,2 | 8,82 | 2187 | 0 | 2 | 1 | 0,942 | 1,008 | 1,048 | 0,98 | 1,156 | 1,033 | 0,867 | 0,976 | 0,99 | -0,01 | 8,4E-01 |
| P62082 | 40S ribosomal protein S7 OS=Mus musculus OX=10090 GN=Rps7 | 14 | 4 | 8 | 4 | 194 | 22,1 | 10,1 | 4094 | 2 | 3 | 1 | 1,009 | 0,963 | 1,036 | 1,085 | 0,861 | 1,118 | 1,034 | 1,03 | 1,01 | 0,01 | 8,4E-01 |
| Q9D0I9 | Arginine--tRNA ligase, cytoplasmic OS=Mus musculus OX=10090 | 10 | 6 | 9 | 6 | 660 | 75,6 | 7,55 | 3359 | 9,95 | 4 | 3 | 1,068 | 0,966 | 0,893 | 1,111 | 0,784 | 1,028 | 1,059 | 1,095 | 1,02 | 0,02 | 8,4E-01 |
| Q9Z2W1 | Serine/threonine-protein kinase 25 OS=Mus musculus OX=1009 | 2 | 1 | 1 | 1 | 426 | 48,1 | 6,81 | 513 | | 1 | | 1,017 | 0,78 | 1,301 | 0,668 | 1,088 | 0,804 | 1,224 | 0,797 | 0,96 | -0,04 | 8,4E-01 |
| Q62446 | Peptidyl-prolyl cis-trans isomerase FKBP3 OS=Mus musculus OX= | 12 | 2 | 3 | 2 | 224 | 25,1 | 9,28 | 1753 | | 2 | | 0,994 | 0,857 | 0,968 | 1,005 | 0,943 | 1,038 | 1,044 | 0,847 | 0,99 | -0,01 | 8,4E-01 |
| P80318 | T-complex protein 1 subunit gamma OS=Mus musculus OX=1009 | 36 | 24 | 35 | 24 | 545 | 60,6 | 6,7 | 11532 | 41,08 | 14 | 10 | 0,937 | 1,061 | 1,033 | 0,978 | 1,088 | 0,972 | 0,982 | 0,999 | 0,99 | -0,01 | 8,4E-01 |
| Q8VCI5 | Peroxisomal biogenesis factor 19 OS=Mus musculus OX=10090 ( | 26 | 6 | 10 | 6 | 299 | 32,7 | 4,34 | 2968 | 11,6 | 3 | 4 | 1,013 | 1,013 | 0,877 | 1,195 | 1,122 | 0,901 | 1,052 | 0,955 | 1,02 | 0,02 | 8,4E-01 |
| Q923D4 | Splicing factor 3B subunit 5 OS=Mus musculus OX=10090 GN=Sf | 13 | 1 | 1 | 1 | 86 | 10,1 | 6,35 | 469 | | 1 | | 0,888 | 0,699 | 0,999 | 1,068 | 0,862 | 1,349 | 0,813 | 0,761 | 0,97 | -0,03 | 8,4E-01 |
| Q99J83 | Autophagy protein 5 OS=Mus musculus OX=10090 GN=Atg5 PE= | 5 | 1 | 1 | 1 | 275 | 32,4 | 5,91 | | 2,62 | | 1 | 0,955 | 0,855 | 1,095 | 1,258 | 0,85 | 1,094 | 1,154 | 1,159 | 0,98 | -0,02 | 8,4E-01 |
| Q9DBG9 | Tax1-binding protein 3 OS=Mus musculus OX=10090 GN=Tax1bp | 14 | 1 | 1 | 1 | 124 | 13,7 | 8,48 | 554 | | 1 | | 0,919 | 1,104 | 0,824 | 0,976 | 0,948 | 0,787 | 1,07 | 0,95 | 1,02 | 0,02 | 8,4E-01 |
| Q924D0 | Reticulon-4-interacting protein 1, mitochondrial OS=Mus muscu | 3 | 1 | 2 | 1 | 396 | 43,3 | 9,2 | 1018 | | 1 | | 1,037 | 1,099 | 0,952 | 1,029 | 1,248 | 0,818 | 0,955 | 1,018 | 1,02 | 0,02 | 8,4E-01 |
| Q80XN0 | D-beta-hydroxybutyrate dehydrogenase, mitochondrial OS=Mus | 29 | 9 | 22 | 9 | 343 | 38,3 | 9,01 | 6902 | 26,95 | 5 | 5 | 1,028 | 0,961 | 1,111 | 0,941 | 0,877 | 1,115 | 0,945 | 1,05 | 1,01 | 0,01 | 8,4E-01 |
| Q91WC3 | Long-chain-fatty-acid--CoA ligase 6 OS=Mus musculus OX=1009( | 23 | 13 | 17 | 13 | 697 | 78 | 7,34 | 5368 | 15,94 | 8 | 5 | 0,996 | 1,003 | 0,971 | 0,948 | 0,861 | 1,02 | 1,02 | 1,054 | 0,99 | -0,01 | 8,4E-01 |
| Q9R0K7 | Plasma membrane calcium-transporting ATPase 2 OS=Mus musc | 33 | 39 | 91 | 25 | 1198 | 132,5 | 5,96 | 32161 | 98,49 | 23 | 19 | 0,999 | 0,971 | 1,003 | 0,956 | 0,96 | 1,028 | 0,955 | 1,003 | 1,00 | 0,00 | 8,5E-01 |
| Q8C8R3 | Ankyrin-2 OS=Mus musculus OX=10090 GN=Ank2 PE=1 SV=2 | 38 | 118 | 239 | 1 | 3898 | 426 | 5,17 | 87801 | 254,97 | 78 | 52 | 1,159 | 1,017 | 1,184 | 1,052 | 1,096 | 0,982 | 1,043 | 1,236 | 1,01 | 0,01 | 8,5E-01 |
| Q9Z0G0 | PDZ domain-containing protein GIPC1 OS=Mus musculus OX=10 | 7 | 2 | 2 | 2 | 333 | 36,1 | 5,91 | 522 | 2,38 | 1 | 1 | 0,935 | 0,941 | 0,979 | 1,127 | 0,807 | 1,021 | 0,929 | 1,155 | 1,02 | 0,02 | 8,5E-01 |
| P70297 | Signal transducing adapter molecule 1 OS=Mus musculus OX=10 | 11 | 5 | 6 | 5 | 548 | 59,7 | 4,84 | 1417 | 6,27 | 3 | 3 | 1,021 | 0,946 | 1,07 | 1,035 | 1,058 | 0,955 | 1,064 | 0,963 | 1,01 | 0,01 | 8,5E-01 |
| Q9CX56 | 26S proteasome non-ATPase regulatory subunit 8 OS=Mus musc | 12 | 4 | 6 | 4 | 353 | 39,9 | 9,58 | 2777 | | 4 | | 1,017 | 1,004 | 1,135 | 1,147 | 0,889 | 1,082 | 1,099 | 1,175 | 1,01 | 0,01 | 8,5E-01 |
| Q62420 | Endophilin-A1 OS=Mus musculus OX=10090 GN=Sh3gl2 PE=1 SV | 35 | 14 | 24 | 8 | 352 | 39,9 | 5,39 | 9833 | 17,84 | 10 | 4 | 1,019 | 1,001 | 1,011 | 0,976 | 1,026 | 1,037 | 1,002 | 0,958 | 1,00 | 0,00 | 8,5E-01 |
| Q80TG9 | Leucine-rich repeat and fibronectin type-III domain-containing p | 3 | 2 | 2 | 1 | 788 | 84,9 | 6,74 | 1319 | | 2 | | 0,846 | 0,862 | 0,795 | 0,706 | 0,962 | 0,804 | 0,762 | 0,731 | 0,98 | -0,02 | 8,5E-01 |
| Q924Z6-3 | Isoform 3 of Exportin-6 OS=Mus musculus OX=10090 GN=Xpo6 | 4 | 1 | 1 | 1 | 246 | 28,6 | 8,03 | | 0 | | 1 | 0,991 | 1,056 | 1,184 | 1,077 | 0,724 | 1,404 | 0,981 | 1,329 | 0,97 | -0,03 | 8,5E-01 |
| Q9QVP9 | Protein-tyrosine kinase 2-beta OS=Mus musculus OX=10090 GN | 6 | 6 | 6 | 6 | 1009 | 115,7 | 6,35 | 2843 | | 6 | | 1,068 | 1,081 | 1,005 | 0,915 | 0,911 | 1,077 | 1,031 | 1,009 | 1,01 | 0,01 | 8,5E-01 |
| Q9WVG6 | Histone-arginine methyltransferase CARM1 OS=Mus musculus C | 3 | 2 | 2 | 2 | 608 | 65,8 | 6,77 | 1112 | | 2 | | 0,869 | 0,948 | 0,8 | 1,078 | 0,772 | 0,893 | 0,975 | 1,131 | 0,98 | -0,02 | 8,5E-01 |
| Q99MR8 | Methylcrotonoyl-CoA carboxylase subunit alpha, mitochondrial | 5 | 3 | 3 | 3 | 717 | 79,3 | 7,83 | 1669 | | 3 | | 0,912 | 0,896 | 1,103 | 1,24 | 1,377 | 1,066 | 0,858 | 0,96 | 0,97 | -0,03 | 8,5E-01 |
| P30275 | Creatine kinase U-type, mitochondrial OS=Mus musculus OX=10 | 25 | 10 | 31 | 8 | 418 | 47 | 8,16 | 15075 | 1,99 | 9 | 1 | 1,044 | 1,082 | 0,955 | 1,018 | 0,991 | 0,996 | 1,003 | 1,082 | 1,01 | 0,01 | 8,5E-01 |
| P62849 | 40S ribosomal protein S24 OS=Mus musculus OX=10090 GN=Rp: | 13 | 2 | 3 | 2 | 133 | 15,4 | 10,78 | 1170 | 2,1 | 2 | 1 | 1,058 | 0,978 | 0,897 | 1,103 | 1,041 | 0,997 | 0,877 | 1,072 | 1,01 | 0,01 | 8,5E-01 |
| Q9CRB6 | Tubulin polymerization-promoting protein family member 3 OS= | 35 | 8 | 11 | 8 | 176 | 19 | 9,11 | 2774 | 8,85 | 4 | 4 | 1,058 | 0,957 | 0,907 | 1,069 | 0,919 | 1,017 | 1,131 | 0,971 | 0,99 | -0,01 | 8,5E-01 |
| Q8CAQ8-2 | Isoform 2 of MICOS complex subunit Mic60 OS=Mus musculus C | 50 | 34 | 53 | 34 | 746 | 82,9 | 6,61 | 21438 | 54,92 | 22 | 16 | 1,041 | 1,015 | 1,005 | 0,993 | 1,003 | 0,959 | 1,049 | 1,063 | 1,00 | 0,00 | 8,5E-01 |
| O35127 | Protein C10 OS=Mus musculus OX=10090 GN=Grcc10 PE=1 SV=1 | 22 | 2 | 3 | 2 | 126 | 13,2 | 5,14 | | 7,34 | | 2 | 0,757 | 0,85 | 0,848 | 1,006 | 0,736 | 0,894 | 0,898 | 0,99 | 0,98 | -0,02 | 8,5E-01 |

| | | | | | | | | | | | | | | | | | | | | | | | |
|---|---|---|---|---|---|---|---|---|---|---|---|---|---|---|---|---|---|---|---|---|---|---|---|
| Q60902 | Epidermal growth factor receptor substrate 15-like 1 OS=Mus m | 8 | 6 | 9 | 6 | 907 | 99,2 | 5,02 | 3327 | 4,68 | 4 | 2 | 0,925 | 1,142 | 0,89 | 0,993 | 0,937 | 0,926 | 1,025 | 1,015 | 1,01 | 0,01 | 8,5E-01 |
| P31786 | Acyl-CoA-binding protein OS=Mus musculus OX=10090 GN=Dbi | 54 | 5 | 14 | 5 | 87 | 10 | 8,82 | 787 | 26,45 | 1 | 4 | 0,993 | 1,098 | 1,069 | 0,998 | 1,163 | 1,112 | 0,965 | 0,962 | 0,99 | -0,01 | 8,5E-01 |
| Q02395 | Metal-response element-binding transcription factor 2 OS=Mus | 1 | 1 | 1 | 1 | 593 | 66,9 | 8,68 | 668 | | 1 | | 0,905 | 1,13 | 0,868 | 0,845 | 1,106 | 0,954 | 0,924 | 0,83 | 0,98 | -0,02 | 8,6E-01 |
| Q8BVL9 | Janus kinase and microtubule-interacting protein 1 OS=Mus mu | 1 | 1 | 1 | 1 | 626 | 73,1 | 6,09 | 365 | | 1 | | 0,742 | 1,093 | 0,576 | 0,838 | 0,788 | 0,613 | 1,114 | 0,847 | 0,97 | -0,03 | 8,6E-01 |
| Q61205 | Platelet-activating factor acetylhydrolase IB subunit gamma OS= | 19 | 4 | 5 | 4 | 232 | 25,8 | 6,93 | 1549 | 5,98 | 2 | 2 | 1,043 | 1,256 | 1,097 | 1,341 | 1,295 | 1,174 | 1,133 | 1,072 | 1,01 | 0,01 | 8,6E-01 |
| Q9D5V6 | Synapse-associated protein 1 OS=Mus musculus OX=10090 GN= | 7 | 2 | 4 | 2 | 365 | 41,3 | 4,54 | | 9,81 | | 2 | 0,941 | 1,112 | 0,918 | 1,182 | 0,902 | 0,985 | 1,132 | 1,205 | 0,98 | -0,02 | 8,6E-01 |
| Q91YR7 | Pre-mRNA-processing factor 6 OS=Mus musculus OX=10090 GN | 3 | 2 | 2 | 2 | 941 | 106,7 | 8,09 | 172 | 1,98 | 1 | 1 | 0,864 | 0,924 | 1,214 | 0,817 | 1,049 | 0,852 | 1,013 | 0,826 | 1,02 | 0,02 | 8,6E-01 |
| Q9CQD1 | Ras-related protein Rab-5A OS=Mus musculus OX=10090 GN=Ra | 24 | 4 | 9 | 2 | 215 | 23,6 | 8,15 | 3632 | 4,68 | 4 | 1 | 0,919 | 0,942 | 0,956 | 0,93 | 1,114 | 0,87 | 0,929 | 0,877 | 0,99 | -0,01 | 8,6E-01 |
| P27546-4 | Isoform 4 of Microtubule-associated protein 4 OS=Mus musculu | 14 | 12 | 22 | 4 | 902 | 94,5 | 9,06 | 12339 | 5,91 | 10 | 3 | 1,014 | 0,862 | 0,974 | 0,955 | 0,83 | 1,089 | 0,91 | 0,929 | 1,01 | 0,01 | 8,6E-01 |
| Q99PG2 | Opioid growth factor receptor OS=Mus musculus OX=10090 GN | 15 | 4 | 5 | 4 | 633 | 70,6 | 4,77 | 2206 | 4,51 | 4 | 1 | 1,029 | 1,216 | 0,897 | 1,08 | 1,154 | 0,938 | 1,096 | 1,094 | 0,99 | -0,01 | 8,6E-01 |
| Q8VCN9 | Tubulin-specific chaperone C OS=Mus musculus OX=10090 GN= | 2 | 1 | 1 | 1 | 341 | 38,1 | 5,3 | 599 | | 1 | | 0,999 | 1,214 | 0,925 | 0,71 | 0,829 | 1,017 | 1,163 | 0,932 | 0,98 | -0,02 | 8,6E-01 |
| P30999 | Catenin delta-1 OS=Mus musculus OX=10090 GN=Ctnnd1 PE=1 S | 15 | 13 | 15 | 13 | 938 | 104,9 | 6,87 | 8080 | 2,88 | 12 | 1 | 0,97 | 0,84 | 0,88 | 0,922 | 0,94 | 0,844 | 0,949 | 0,906 | 0,99 | -0,01 | 8,6E-01 |
| Q9DB15 | 39S ribosomal protein L12, mitochondrial OS=Mus musculus OX | 25 | 5 | 11 | 5 | 201 | 21,7 | 9,29 | 5765 | 9,22 | 4 | 1 | 1,006 | 0,974 | 0,904 | 1,025 | 1,032 | 0,933 | 1,031 | 0,941 | 0,99 | -0,01 | 8,6E-01 |
| Q8BJU2 | Tetraspanin-9 OS=Mus musculus OX=10090 GN=Tspan9 PE=1 SV | 6 | 1 | 1 | 1 | 239 | 26,7 | 7,42 | | 2,49 | | 1 | 1,262 | 1,174 | 0,934 | 1,243 | 1,59 | 0,862 | 0,977 | 1,054 | 1,03 | 0,03 | 8,6E-01 |
| Q9ESN9 | C-Jun-amino-terminal kinase-interacting protein 3 OS=Mus mus | 2 | 3 | 3 | 2 | 1337 | 147,5 | 5,45 | 1202 | 2,65 | 2 | 1 | 1,129 | 0,993 | 1,217 | 0,911 | 1,211 | 1,001 | 1,104 | 0,996 | 0,99 | -0,01 | 8,6E-01 |
| Q8C7R4 | Ubiquitin-like modifier-activating enzyme 6 OS=Mus musculus C | 1 | 1 | 1 | 1 | 1053 | 117,9 | 6,11 | 505 | | 1 | | 0,886 | 0,899 | 1,089 | 0,663 | 0,744 | 1,017 | 0,746 | 0,949 | 1,02 | 0,02 | 8,6E-01 |
| P48320 | Glutamate decarboxylase 2 OS=Mus musculus OX=10090 GN=Ga | 7 | 4 | 7 | 3 | 585 | 65,2 | 6,9 | 2025 | 0 | 3 | 1 | 1,022 | 0,937 | 0,926 | 0,932 | 0,858 | 1,101 | 0,955 | 0,943 | 0,99 | -0,01 | 8,6E-01 |
| Q6PDS3 | Sterile alpha and TIR motif-containing protein 1 OS=Mus muscul | 9 | 5 | 5 | 5 | 724 | 79,6 | 6,27 | 1427 | 7,95 | 2 | 3 | 1,044 | 1,217 | 0,982 | 1,171 | 1,263 | 0,931 | 1,113 | 1,171 | 0,99 | -0,01 | 8,6E-01 |
| Q9QXT0 | Protein canopy homolog 2 OS=Mus musculus OX=10090 GN=Cn | 34 | 5 | 6 | 5 | 182 | 20,8 | 5,07 | 2139 | 2,22 | 3 | 2 | 0,736 | 0,906 | 0,941 | 0,833 | 1,074 | 0,836 | 0,802 | 0,764 | 0,98 | -0,02 | 8,6E-01 |
| P97825 | Jupiter microtubule associated homolog 1 OS=Mus musculus OX | 38 | 4 | 6 | 4 | 154 | 16,1 | 5,31 | 1888 | 2,29 | 4 | 1 | 0,874 | 1,087 | 0,898 | 1,019 | 1,116 | 0,899 | 0,983 | 0,93 | 0,99 | -0,01 | 8,6E-01 |
| Q80XK6 | Autophagy-related protein 2 homolog B OS=Mus musculus OX= | 0 | 1 | 1 | 1 | 2075 | 231,3 | 5,88 | 615 | | 1 | | 0,835 | 1,13 | 1,364 | 0,709 | 1,008 | 1,25 | 0,919 | 0,979 | 0,97 | -0,03 | 8,6E-01 |
| Q6P9K9 | Neurexin-3 OS=Mus musculus OX=10090 GN=Nrxn3 PE=1 SV=2 | 3 | 5 | 6 | 3 | 1571 | 173,3 | 6,14 | 3582 | | 5 | | 1,296 | 1,299 | 1,038 | 1,441 | 1,142 | 1,099 | 1,266 | 1,481 | 1,02 | 0,02 | 8,6E-01 |
| Q9Z2D0 | Myotubularin-related protein 9 OS=Mus musculus OX=10090 GN | 10 | 6 | 6 | 6 | 545 | 62,9 | 6,62 | 1567 | 5,67 | 3 | 3 | 0,996 | 0,951 | 0,961 | 0,945 | 0,927 | 1,15 | 0,902 | 0,917 | 0,99 | -0,01 | 8,6E-01 |
| Q3TPE9 | Ankyrin repeat and MYND domain-containing protein 2 OS=Mus | 7 | 2 | 2 | 2 | 440 | 48,7 | 6,67 | 438 | 0 | 1 | 1 | 1,297 | 0,855 | 0,945 | 1,253 | 1,261 | 1,115 | 1,012 | 1,05 | 0,98 | -0,02 | 8,6E-01 |
| Q921M7 | Protein FAM49B OS=Mus musculus OX=10090 GN=Fam49b PE= | 24 | 7 | 11 | 6 | 324 | 36,8 | 6,06 | 6864 | | 7 | | 1,032 | 1,15 | 1,022 | 1,123 | 0,824 | 1,162 | 1,256 | 1,014 | 1,02 | 0,02 | 8,6E-01 |
| Q9DC61 | Mitochondrial-processing peptidase subunit alpha OS=Mus mus | 7 | 4 | 4 | 4 | 524 | 58,2 | 6,83 | 1718 | 2,19 | 3 | 1 | 1,117 | 1,11 | 1,069 | 1,162 | 1,391 | 0,93 | 1,135 | 1,072 | 0,98 | -0,02 | 8,6E-01 |
| P63044 | Vesicle-associated membrane protein 2 OS=Mus musculus OX=1 | 43 | 5 | 16 | 5 | 116 | 12,7 | 8,13 | 6290 | 17,56 | 4 | 2 | 0,903 | 1,142 | 0,995 | 0,919 | 0,986 | 0,865 | 1,177 | 0,991 | 0,99 | -0,01 | 8,6E-01 |
| P62257 | Ubiquitin-conjugating enzyme E2 H OS=Mus musculus OX=1009 | 5 | 1 | 1 | 1 | 183 | 20,6 | 4,67 | | 2,47 | | 1 | 0,889 | 1,001 | 1,023 | 0,867 | 0,782 | 1,116 | 1,007 | 0,932 | 0,99 | -0,01 | 8,7E-01 |
| P00920 | Carbonic anhydrase 2 OS=Mus musculus OX=10090 GN=Ca2 PE= | 21 | 4 | 7 | 4 | 260 | 29 | 7,01 | 808 | 17,66 | 1 | 3 | 1,017 | 0,996 | 1,072 | 1,051 | 1,075 | 1,019 | 1,049 | 1,009 | 1,00 | 0,00 | 8,7E-01 |
| Q9QZB9 | Dynactin subunit 5 OS=Mus musculus OX=10090 GN=Dctn5 PE= | 14 | 3 | 3 | 3 | 182 | 20,1 | 8,02 | 1391 | 0 | 2 | 1 | 0,966 | 0,943 | 0,866 | 0,89 | 1,184 | 0,793 | 0,876 | 0,875 | 0,98 | -0,02 | 8,7E-01 |
| Q6PGH2 | Jupiter microtubule associated homolog 2 OS=Mus musculus OX | 26 | 3 | 3 | 3 | 190 | 20 | 8,62 | 1942 | | 3 | | 1,209 | 1,038 | 1,731 | 1,127 | 2,132 | 1,123 | 1,045 | 1,021 | 0,96 | -0,04 | 8,7E-01 |
| Q8C570 | mRNA export factor OS=Mus musculus OX=10090 GN=Rae1 PE= | 3 | 2 | 2 | 2 | 368 | 40,9 | 7,83 | 1066 | | 2 | | 1,091 | 0,898 | 0,944 | 1,138 | 0,834 | 0,877 | 1,213 | 1,231 | 0,98 | -0,02 | 8,7E-01 |
| Q9D071 | MMS19 nucleotide excision repair protein homolog OS=Mus mu | 1 | 1 | 1 | 1 | 1031 | 113 | 6,21 | | 2,23 | | 1 | 0,883 | 1,092 | 1,078 | 1,072 | 0,791 | 1,071 | 1,202 | 1,132 | 0,98 | -0,02 | 8,7E-01 |
| Q9Z275 | Retinaldehyde-binding protein 1 OS=Mus musculus OX=10090 G | 12 | 4 | 6 | 4 | 317 | 36,4 | 5,06 | 2051 | 4,15 | 3 | 2 | 0,985 | 0,972 | 1,105 | 1,095 | 1,089 | 0,935 | 1,125 | 0,968 | 1,01 | 0,01 | 8,7E-01 |
| Q03517 | Secretogranin-2 OS=Mus musculus OX=10090 GN=Scg2 PE=1 SV | 14 | 8 | 10 | 8 | 617 | 70,6 | 4,75 | 3340 | 6,07 | 5 | 3 | 1,106 | 1,036 | 1,08 | 1,075 | 1,153 | 1,114 | 1,04 | 0,959 | 1,01 | 0,01 | 8,7E-01 |
| Q9CYN2 | Signal peptidase complex subunit 2 OS=Mus musculus OX=1009 | 8 | 2 | 2 | 2 | 226 | 25 | 8,57 | 1255 | | 2 | | 1,031 | 0,712 | 0,965 | 0,985 | 0,777 | 0,88 | 1,024 | 0,951 | 1,02 | 0,02 | 8,7E-01 |
| Q61701 | ELAV-like protein 4 OS=Mus musculus OX=10090 GN=Elavl4 PE= | 10 | 3 | 3 | 1 | 385 | 42,3 | 9,38 | 718 | 2,82 | 1 | 2 | 0,833 | 0,839 | 0,919 | 1,169 | 1,018 | 0,692 | 1,091 | 1,041 | 0,98 | -0,02 | 8,7E-01 |
| Q9D880 | Mitochondrial import inner membrane translocase subunit TIM | 15 | 5 | 11 | 5 | 353 | 39,8 | 8,13 | 4504 | 3,79 | 3 | 2 | 0,879 | 0,911 | 1,158 | 1,032 | 0,777 | 1,026 | 1,057 | 1,057 | 1,02 | 0,02 | 8,7E-01 |
| P32067 | Lupus La protein homolog OS=Mus musculus OX=10090 GN=Ssb | 9 | 4 | 5 | 4 | 415 | 47,7 | 9,77 | 961 | 8,77 | 2 | 2 | 0,892 | 0,947 | 0,869 | 1,144 | 0,96 | 0,926 | 1,063 | 0,95 | 0,99 | -0,01 | 8,7E-01 |
| Q9D1N9 | 39S ribosomal protein L21, mitochondrial OS=Mus musculus OX | 6 | 1 | 1 | 1 | 209 | 23,4 | 9,99 | 488 | | 1 | | 0,969 | 0,963 | 1,107 | 0,999 | 1,398 | 0,952 | 0,848 | 0,927 | 0,98 | -0,02 | 8,7E-01 |
| P63094 | Guanine nucleotide-binding protein G(s) subunit alpha isoforms | 27 | 9 | 18 | 9 | 394 | 45,6 | 5,96 | 5410 | 20,37 | 8 | 1 | 0,98 | 1,014 | 0,995 | 1,071 | 1,125 | 0,981 | 1,003 | 0,978 | 0,99 | -0,01 | 8,7E-01 |
| O35098 | Dihydropyrimidinase-related protein 4 OS=Mus musculus OX=10 | 58 | 25 | 112 | 23 | 572 | 61,9 | 6,98 | 48541 | 85,77 | 18 | 12 | 0,983 | 0,987 | 0,956 | 1,007 | 0,891 | 1,021 | 0,997 | 1,003 | 1,01 | 0,01 | 8,7E-01 |
| Q99LJ0 | CTTNBP2 N-terminal-like protein OS=Mus musculus OX=10090 G | 4 | 2 | 3 | 2 | 638 | 69,8 | 7,71 | 1005 | | 2 | | 0,959 | 1,023 | 0,94 | 1,099 | 0,884 | 0,98 | 1,066 | 1,054 | 1,01 | 0,01 | 8,7E-01 |
| P09528 | Ferritin heavy chain OS=Mus musculus OX=10090 GN=Fth1 PE=1 | 21 | 4 | 5 | 4 | 182 | 21,1 | 5,88 | 3585 | | 4 | | 1,203 | 1,039 | 0,87 | 1,045 | 0,946 | 1,202 | 1,011 | 0,937 | 1,01 | 0,01 | 8,7E-01 |
| Q61578 | NADPH:adrenodoxin oxidoreductase, mitochondrial OS=Mus mu | 4 | 2 | 2 | 2 | 494 | 54,2 | 8,66 | 641 | 1,66 | 1 | 1 | 0,885 | 0,829 | 1,085 | 0,906 | 1,221 | 0,864 | 0,784 | 0,909 | 0,98 | -0,02 | 8,7E-01 |
| Q99L27 | GMP reductase 2 OS=Mus musculus OX=10090 GN=Gmpr2 PE=1 | 6 | 2 | 5 | 1 | 348 | 38 | 7,44 | 2581 | | 2 | | 0,726 | 0,881 | 0,763 | 0,898 | 0,816 | 0,703 | 0,847 | 0,865 | 1,01 | 0,01 | 8,7E-01 |
| A2CG49 | Kalirin OS=Mus musculus OX=10090 GN=Kalrn PE=1 SV=1 | 2 | 6 | 9 | 4 | 2964 | 336,8 | 6,07 | 2770 | 5,78 | 4 | 2 | 0,99 | 1,018 | 0,819 | 1,018 | 0,832 | 0,985 | 0,991 | 0,996 | 1,01 | 0,01 | 8,7E-01 |
| Q64133 | Amine oxidase [flavin-containing] A OS=Mus musculus OX=1009 | 19 | 10 | 12 | 10 | 526 | 59,6 | 7,81 | 5330 | 6,61 | 7 | 3 | 1,125 | 1,103 | 1,063 | 1,016 | 1,065 | 1,082 | 1,109 | 1,032 | 1,00 | 0,00 | 8,7E-01 |
| Q61553 | Fascin OS=Mus musculus OX=10090 GN=Fscn1 PE=1 SV=4 | 47 | 21 | 40 | 21 | 493 | 54,5 | 6,89 | 15443 | 31,15 | 13 | 8 | 0,939 | 0,881 | 1,111 | 0,884 | 0,933 | 1,033 | 0,907 | 0,982 | 0,99 | -0,01 | 8,7E-01 |
| P57759 | Endoplasmic reticulum resident protein 29 OS=Mus musculus O | 21 | 5 | 8 | 5 | 262 | 28,8 | 6,15 | 2183 | 8,54 | 3 | 2 | 0,969 | 0,951 | 0,725 | 1,262 | 0,913 | 1,029 | 0,963 | 0,928 | 1,02 | 0,02 | 8,8E-01 |
| Q9DBN5 | Lon protease homolog 2, peroxisomal OS=Mus musculus OX=10 | 1 | 1 | 2 | 1 | 852 | 94,5 | 7,23 | 712 | | 1 | | 1,089 | 1,359 | 0,959 | 1,246 | 1,293 | 1,063 | 1,075 | 1,155 | 1,01 | 0,01 | 8,8E-01 |
| P35505 | Fumarylacetoacetase OS=Mus musculus OX=10090 GN=Fah PE= | 5 | 2 | 2 | 2 | 419 | 46,1 | 7,18 | | 2,45 | | 2 | 1,171 | 1,443 | 1,815 | 1,528 | 1,759 | 1,41 | 1,421 | 1,256 | 1,02 | 0,02 | 8,8E-01 |
| P80314 | T-complex protein 1 subunit beta OS=Mus musculus OX=10090 | 43 | 23 | 36 | 23 | 535 | 57,4 | 6,4 | 15510 | 26,89 | 17 | 7 | 1,051 | 0,918 | 1,115 | 1,002 | 0,945 | 1,106 | 0,963 | 1,036 | 1,01 | 0,01 | 8,8E-01 |
| Q8BSL7 | ADP-ribosylation factor 2 OS=Mus musculus OX=10090 GN=Arf2 | 47 | 8 | 16 | 1 | 181 | 20,7 | 6,58 | 7450 | 6,44 | 6 | 3 | 1,381 | 0,817 | 0,912 | 1,105 | 1,096 | 1,124 | 0,929 | 0,98 | 1,02 | 0,02 | 8,8E-01 |
| Q99N84 | 28S ribosomal protein S18b, mitochondrial OS=Mus musculus O | 4 | 1 | 1 | 1 | 254 | 28,7 | 8,35 | | 2,65 | | 1 | 1,136 | 1,081 | 1,526 | 1,054 | 1,484 | 1,212 | 1,055 | 0,942 | 1,02 | 0,02 | 8,8E-01 |
| P51855 | Glutathione synthetase OS=Mus musculus OX=10090 GN=Gss PI | 4 | 1 | 2 | 1 | 474 | 52,2 | 5,8 | 1473 | | 1 | | 2,848 | 2,852 | 1,714 | 2,622 | 3,012 | 1,955 | 2,804 | 2,494 | 0,98 | -0,02 | 8,8E-01 |
| Q9CY18 | Sorting nexin-7 OS=Mus musculus OX=10090 GN=Snx7 PE=1 SV= | 8 | 3 | 4 | 3 | 387 | 45 | 5,07 | 1565 | 2,03 | 2 | 1 | 0,857 | 0,859 | 0,883 | 0,871 | 0,815 | 0,809 | 1,001 | 0,874 | 0,99 | -0,01 | 8,8E-01 |
| Q6GYP7 | Ral GTPase-activating protein subunit alpha-1 OS=Mus musculus | 2 | 4 | 5 | 4 | 2035 | 229,2 | 6,1 | 1450 | 2,52 | 3 | 1 | 1,127 | 0,958 | 1,062 | 1,102 | 1,014 | 1,036 | 0,982 | 1,265 | 0,99 | -0,01 | 8,8E-01 |
| Q64727 | Vinculin OS=Mus musculus OX=10090 GN=Vcl PE=1 SV=4 | 12 | 13 | 15 | 13 | 1066 | 116,6 | 6 | 4245 | 14,23 | 7 | 6 | 0,959 | 1,052 | 1,02 | 1,104 | 1,153 | 0,796 | 1,038 | 1,21 | 0,99 | -0,01 | 8,8E-01 |
| Q99KR7 | Peptidyl-prolyl cis-trans isomerase F, mitochondrial OS=Mus mu | 8 | 2 | 2 | 2 | 206 | 21,7 | 9,16 | 516 | 2,94 | 1 | 1 | 0,984 | 0,928 | 1,042 | 1,037 | 0,986 | 1,049 | 0,961 | 1,016 | 0,99 | -0,01 | 8,8E-01 |
| Q6V4S5 | Protein sidekick-2 OS=Mus musculus OX=10090 GN=Sdk2 PE=1 S | 3 | 5 | 7 | 4 | 2176 | 239,8 | 7,36 | 3335 | | 5 | | 1,345 | 1,15 | 1,308 | 1,284 | 1,249 | 1,39 | 1,106 | 1,393 | 0,99 | -0,01 | 8,8E-01 |
| P47809 | Dual specificity mitogen-activated protein kinase kinase 4 OS=M | 13 | 4 | 5 | 4 | 397 | 44,1 | 8,07 | 2627 | 2,3 | 3 | 1 | 0,956 | 1,116 | 1,156 | 1,063 | 0,951 | 1,286 | 0,971 | 1,027 | 1,01 | 0,01 | 8,8E-01 |
| Q9D1P4 | Cysteine and histidine-rich domain-containing protein 1 OS=Mu | 11 | 3 | 4 | 3 | 331 | 37,3 | 7,9 | 1866 | 0 | 2 | 1 | 1,04 | 0,903 | 1,128 | 0,972 | 0,958 | 1,135 | 0,865 | 1,038 | 1,01 | 0,01 | 8,8E-01 |
| Q920Q6 | RNA-binding protein Musashi homolog 2 OS=Mus musculus OX= | 8 | 2 | 3 | 2 | 346 | 36,9 | 8,47 | 1968 | | 2 | | 0,934 | 0,783 | 0,875 | 1,033 | 0,943 | 0,85 | 0,838 | 0,956 | 1,01 | 0,01 | 8,8E-01 |
| Q5DTL9 | Sodium-driven chloride bicarbonate exchanger OS=Mus musculu | 3 | 3 | 3 | 2 | 1118 | 125,7 | 6,51 | 968 | 2,34 | 2 | 1 | 1,35 | 1,353 | 1,293 | 0,967 | 1,284 | 1,031 | 1,311 | 1,267 | 1,01 | 0,01 | 8,8E-01 |

| | | | | | | | | | | | | | | | | | | | | | | | |
|---|---|---|---|---|---|---|---|---|---|---|---|---|---|---|---|---|---|---|---|---|---|---|---|
| Q8K394 | Inactive phospholipase C-like protein 2 OS=Mus musculus OX=1 | 7 | 7 | 9 | 7 | 1128 | 125,7 | 6,92 | 3687 | 4,46 | 5 | 2 | 0,914 | 0,952 | 0,91 | 1,032 | 0,977 | 0,941 | 1,005 | 0,907 | 0,99 | -0,01 | 8,8E-01 |
| Q9D1P0 | 39S ribosomal protein L13, mitochondrial OS=Mus musculus OX | 20 | 3 | 4 | 3 | 178 | 20,7 | 9,35 | 817 | 2,23 | 2 | 1 | 0,901 | 0,817 | 0,952 | 1,058 | 1,357 | 0,908 | 0,62 | 0,942 | 0,97 | -0,03 | 8,8E-01 |
| Q8R127 | Saccharopine dehydrogenase-like oxidoreductase OS=Mus musc | 12 | 5 | 5 | 5 | 429 | 47,1 | 8,6 | 633 | 12,55 | 1 | 4 | 1,121 | 0,999 | 1,094 | 1,084 | 1,04 | 0,997 | 1,069 | 1,227 | 0,99 | -0,01 | 8,8E-01 |
| Q9CZ49 | Kelch-like protein 35 OS=Mus musculus OX=10090 GN=Klhl35 PE | 1 | 1 | 2 | 1 | 574 | 62,3 | 7,42 | | 4,85 | | 1 | 0,814 | 1,683 | 0,726 | 0,988 | 1,951 | 0,613 | 0,786 | 1,086 | 0,95 | -0,05 | 8,8E-01 |
| Q9JJI8 | 60S ribosomal protein L38 OS=Mus musculus OX=10090 GN=Rpl | 14 | 1 | 2 | 1 | 70 | 8,2 | 10,1 | 1329 | | 1 | | 0,899 | 0,974 | 0,972 | 1,004 | 1,004 | 0,906 | 0,82 | 1,166 | 0,99 | -0,01 | 8,8E-01 |
| Q91WK1 | SPRY domain-containing protein 4 OS=Mus musculus OX=10090 | 11 | 2 | 2 | 2 | 207 | 23,3 | 9,45 | 1246 | | 2 | | 0,82 | 1,015 | 1,07 | 0,98 | 1,03 | 1,036 | 0,751 | 1,015 | 1,01 | 0,01 | 8,8E-01 |
| P04627 | Serine/threonine-protein kinase A-Raf OS=Mus musculus OX=10 | 3 | 2 | 2 | 1 | 604 | 67,5 | 9,11 | 704 | 3,6 | 1 | 1 | 0,697 | 0,848 | 1,237 | 0,544 | 0,413 | 0,927 | 0,957 | 0,911 | 1,04 | 0,04 | 8,9E-01 |
| P26049 | Gamma-aminobutyric acid receptor subunit alpha-3 OS=Mus mu | 14 | 5 | 6 | 5 | 492 | 55,4 | 8,94 | 2718 | 5,27 | 3 | 2 | 1,017 | 1,094 | 0,984 | 0,978 | 1,009 | 0,978 | 1,021 | 1,047 | 1,00 | 0,00 | 8,9E-01 |
| Q8BVE3 | V-type proton ATPase subunit H OS=Mus musculus OX=10090 G | 16 | 7 | 12 | 7 | 483 | 55,8 | 6,61 | 4969 | 8,83 | 4 | 3 | 1,054 | 1,005 | 1,058 | 0,951 | 1,149 | 0,987 | 1,03 | 0,933 | 0,99 | -0,01 | 8,9E-01 |
| Q5DU25 | IQ motif and SEC7 domain-containing protein 2 OS=Mus muscul | 2 | 2 | 2 | 1 | 1478 | 161,7 | 8,56 | 285 | 1,99 | 1 | 1 | 0,859 | 0,902 | 0,849 | 0,777 | 1,159 | 0,59 | 0,881 | 0,681 | 1,02 | 0,02 | 8,9E-01 |
| Q9D2N4 | Dystrobrevin alpha OS=Mus musculus OX=10090 GN=Dtna PE=1 | 4 | 2 | 3 | 2 | 746 | 84 | 6,76 | 1898 | | 2 | | 0,946 | 0,815 | 1 | 0,893 | 0,861 | 0,937 | 0,837 | 0,988 | 1,01 | 0,01 | 8,9E-01 |
| Q91X83 | S-adenosylmethionine synthase isoform type-1 OS=Mus muscul | 13 | 5 | 8 | 1 | 396 | 43,5 | 5,82 | 3862 | 4 | 4 | 1 | 0,728 | 0,737 | 0,632 | 1,01 | 0,632 | 0,713 | 0,922 | 0,78 | 1,02 | 0,02 | 8,9E-01 |
| Q9DB77 | Cytochrome b-c1 complex subunit 2, mitochondrial OS=Mus mu | 37 | 15 | 32 | 15 | 453 | 48,2 | 9,25 | 11166 | 42,63 | 12 | 10 | 1,059 | 0,986 | 1,057 | 0,968 | 1,165 | 1,022 | 0,957 | 0,958 | 0,99 | -0,01 | 8,9E-01 |
| Q8BYP3 | Rho-related GTP-binding protein RhoF OS=Mus musculus OX=10 | 5 | 1 | 1 | 1 | 211 | 23,6 | 8,43 | 474 | | 1 | | 0,845 | 0,925 | 1,099 | 0,65 | 0,72 | 1,108 | 0,752 | 0,864 | 1,02 | 0,02 | 8,9E-01 |
| Q3UMG5 | Leucine-rich repeat and calponin homology domain-containing p | 2 | 1 | 1 | 1 | 773 | 84,9 | 6,58 | | 3,3 | | 1 | 1,053 | 1,276 | 1,236 | 1,294 | 1,206 | 1,282 | 1,167 | 1,168 | 1,01 | 0,01 | 8,9E-01 |
| P47754 | F-actin-capping protein subunit alpha-2 OS=Mus musculus OX=1 | 58 | 11 | 30 | 8 | 286 | 32,9 | 5,85 | 7607 | 54,85 | 7 | 7 | 0,983 | 1,075 | 0,918 | 1,125 | 1,019 | 0,98 | 1,044 | 1,088 | 0,99 | -0,01 | 8,9E-01 |
| P26039 | Talin-1 OS=Mus musculus OX=10090 GN=Tln1 PE=1 SV=2 | 5 | 10 | 11 | 7 | 2541 | 269,7 | 6,18 | 3607 | 6,31 | 6 | 4 | 0,997 | 0,986 | 1,077 | 1,128 | 1,206 | 0,972 | 1,05 | 0,996 | 0,99 | -0,01 | 8,9E-01 |
| Q9CY16 | 28S ribosomal protein S28, mitochondrial OS=Mus musculus OX | 6 | 1 | 1 | 1 | 186 | 20,5 | 8,92 | 673 | | 1 | | 1,704 | 1,442 | 1,476 | 1,13 | 1,806 | 1,292 | 1,211 | 1,546 | 0,98 | -0,02 | 8,9E-01 |
| Q9CPQ1 | Cytochrome c oxidase subunit 6C OS=Mus musculus OX=10090 ( | 47 | 8 | 18 | 8 | 76 | 8,5 | 10,14 | 6847 | 18,49 | 4 | 4 | 1,068 | 0,937 | 0,927 | 0,909 | 0,899 | 0,998 | 0,956 | 0,964 | 1,01 | 0,01 | 8,9E-01 |
| Q9R1R2 | Tripartite motif-containing protein 3 OS=Mus musculus OX=100 | 19 | 14 | 19 | 11 | 744 | 80,7 | 7,81 | 4330 | 32,22 | 6 | 9 | 1,04 | 0,987 | 1,122 | 0,982 | 1,048 | 1,028 | 0,953 | 1,13 | 0,99 | -0,01 | 8,9E-01 |
| Q99LB6 | Methionine adenosyltransferase 2 subunit beta OS=Mus muscul | 11 | 2 | 2 | 2 | 334 | 37,4 | 6,95 | 670 | 2,96 | 1 | 1 | 0,888 | 1,119 | 1,099 | 0,89 | 1,204 | 0,753 | 1,059 | 1,045 | 0,98 | -0,02 | 8,9E-01 |
| Q7TQI3 | Ubiquitin thioesterase OTUB1 OS=Mus musculus OX=10090 GN= | 28 | 6 | 8 | 6 | 271 | 31,3 | 4,94 | 5327 | | 6 | | 1,003 | 1,058 | 0,844 | 1,037 | 0,939 | 0,956 | 1,064 | 1,015 | 0,99 | -0,01 | 8,9E-01 |
| Q8BSS9 | Liprin-alpha-2 OS=Mus musculus OX=10090 GN=Ppfia2 PE=1 SV= | 15 | 16 | 20 | 13 | 1257 | 143,1 | 6,11 | 7518 | 20,94 | 10 | 7 | 0,971 | 1,069 | 0,918 | 1,032 | 1,101 | 0,922 | 1,001 | 0,994 | 0,99 | -0,01 | 8,9E-01 |
| Q9CQY1 | Ubiquitin-like protein ATG12 OS=Mus musculus OX=10090 GN=/ | 6 | 1 | 1 | 1 | 141 | 15,2 | 4,97 | 671 | | 1 | | 1,079 | 1,071 | 0,898 | 1,266 | 0,924 | 1,048 | 1,253 | 1,147 | 0,99 | -0,01 | 8,9E-01 |
| Q9CRD2 | ER membrane protein complex subunit 2 OS=Mus musculus OX= | 3 | 1 | 2 | 1 | 297 | 34,9 | 6,81 | 993 | | 1 | | 0,887 | 1,018 | 0,777 | 0,978 | 1,092 | 0,779 | 0,927 | 0,812 | 1,01 | 0,01 | 8,9E-01 |
| Q9QYF1 | Retinol dehydrogenase 11 OS=Mus musculus OX=10090 GN=Rdl | 10 | 3 | 3 | 3 | 316 | 35,1 | 8,91 | 1141 | 2,41 | 2 | 1 | 0,905 | 0,859 | 1,037 | 0,985 | 1,034 | 0,825 | 0,937 | 1,025 | 0,99 | -0,01 | 8,9E-01 |
| Q9JLT4 | Thioredoxin reductase 2, mitochondrial OS=Mus musculus OX=1 | 10 | 4 | 4 | 4 | 524 | 56,6 | 8,46 | 1250 | 3,29 | 3 | 1 | 0,989 | 0,912 | 1,097 | 0,871 | 1,045 | 1,087 | 0,758 | 1,029 | 0,99 | -0,01 | 8,9E-01 |
| Q641P0 | Actin-related protein 3B OS=Mus musculus OX=10090 GN=Actr3 | 13 | 5 | 15 | 3 | 418 | 47,5 | 6,02 | 5684 | 15,36 | 2 | 3 | 1,088 | 1,123 | 1,339 | 1,051 | 1,266 | 1,095 | 1,042 | 1,245 | 0,99 | -0,01 | 8,9E-01 |
| Q3UE31 | Uncharacterized protein KIAA0930 homolog OS=Mus musculus ( | 3 | 1 | 1 | 1 | 404 | 45,9 | 8,13 | 595 | | 1 | | 0,892 | 0,855 | 0,939 | 0,662 | 0,426 | 1,051 | 1,089 | 0,689 | 1,03 | 0,03 | 9,0E-01 |
| Q99PL6 | UBX domain-containing protein 6 OS=Mus musculus OX=10090 ( | 3 | 1 | 1 | 1 | 442 | 49,8 | 8,54 | 520 | | 1 | | 1,037 | 0,984 | 0,783 | 1,217 | 0,751 | 0,863 | 1,241 | 1,088 | 1,02 | 0,02 | 9,0E-01 |
| Q9JJ28 | Protein flightless-1 homolog OS=Mus musculus OX=10090 GN=F | 3 | 4 | 4 | 4 | 1271 | 144,7 | 6,06 | 1455 | 2,66 | 3 | 1 | 1,027 | 0,89 | 0,904 | 0,843 | 0,917 | 0,844 | 0,944 | 0,986 | 0,99 | -0,01 | 9,0E-01 |
| Q9D1L9 | Ragulator complex protein LAMTOR5 OS=Mus musculus OX=100 | 33 | 2 | 4 | 2 | 91 | 9,6 | 4,87 | 1869 | 2,88 | 1 | 1 | 0,848 | 0,883 | 0,784 | 1,013 | 0,927 | 0,901 | 0,757 | 0,909 | 1,01 | 0,01 | 9,0E-01 |
| Q99JR1 | Sideroflexin-1 OS=Mus musculus OX=10090 GN=Sfxn1 PE=1 SV= | 20 | 7 | 13 | 6 | 322 | 35,6 | 9,23 | 6021 | 7,96 | 5 | 3 | 1,054 | 0,961 | 1,026 | 0,975 | 0,908 | 1,08 | 1,022 | 0,983 | 1,01 | 0,01 | 9,0E-01 |
| Q06138 | Calcium-binding protein 39 OS=Mus musculus OX=10090 GN=Ca | 10 | 4 | 5 | 4 | 341 | 39,8 | 6,89 | 2873 | 2,47 | 3 | 1 | 1,029 | 0,948 | 0,885 | 1,013 | 0,63 | 1,061 | 1,166 | 1,086 | 0,98 | -0,02 | 9,0E-01 |
| Q8R3V5-1 | Isoform 1 of Endophilin-B2 OS=Mus musculus OX=10090 GN=Sh | 20 | 8 | 9 | 8 | 395 | 44,1 | 5,99 | 3806 | 7,56 | 6 | 2 | 0,966 | 0,899 | 1,105 | 0,979 | 0,94 | 1,017 | 0,999 | 0,968 | 1,01 | 0,01 | 9,0E-01 |
| P15105 | Glutamine synthetase OS=Mus musculus OX=10090 GN=Glul PE | 46 | 19 | 52 | 19 | 373 | 42,1 | 7,08 | 21041 | 52,96 | 11 | 11 | 0,951 | 0,983 | 1,264 | 0,921 | 1,071 | 0,993 | 0,982 | 1,029 | 1,01 | 0,01 | 9,0E-01 |
| Q8R574 | Phosphoribosyl pyrophosphate synthase-associated protein 2 O | 11 | 5 | 6 | 3 | 369 | 40,9 | 7,17 | 1909 | 4,78 | 3 | 2 | 0,926 | 0,837 | 0,857 | 0,868 | 0,94 | 0,896 | 0,77 | 0,86 | 1,01 | 0,01 | 9,0E-01 |
| Q9JMH9-1 | Isoform 1 of Unconventional myosin-XVIIIa OS=Mus musculus O | 18 | 33 | 47 | 33 | 2035 | 230,8 | 6,16 | 25068 | 17,7 | 27 | 6 | 0,961 | 0,994 | 0,946 | 0,981 | 0,936 | 0,93 | 1,017 | 1,013 | 1,00 | 0,00 | 9,0E-01 |
| Q8CIN4 | Serine/threonine-protein kinase PAK 2 OS=Mus musculus OX=10 | 20 | 8 | 10 | 6 | 524 | 57,9 | 5,77 | 4969 | 2,61 | 7 | 1 | 1,057 | 1,007 | 1,084 | 1,002 | 1,072 | 1,044 | 0,985 | 1,064 | 1,00 | 0,00 | 9,0E-01 |
| P47963 | 60S ribosomal protein L13 OS=Mus musculus OX=10090 GN=Rpl | 37 | 8 | 12 | 8 | 211 | 24,3 | 11,55 | 4259 | 14,24 | 4 | 4 | 1,035 | 0,884 | 1,047 | 1,002 | 1,083 | 0,976 | 0,965 | 0,969 | 0,99 | -0,01 | 9,0E-01 |
| Q3TWW8 | Serine/arginine-rich splicing factor 6 OS=Mus musculus OX=100 | 4 | 2 | 2 | 2 | 339 | 39 | 11,46 | 429 | 0 | 1 | 1 | 1,262 | 0,982 | 1,296 | 1,396 | 1,162 | 1,297 | 1,404 | 1,131 | 0,99 | -0,01 | 9,0E-01 |
| Q8BU14 | Translocation protein SEC62 OS=Mus musculus OX=10090 GN=S | 2 | 1 | 1 | 1 | 398 | 45,6 | 7,31 | 772 | | 1 | | 0,978 | 0,949 | 0,985 | 0,857 | 0,836 | 0,872 | 1,032 | 1,063 | 0,99 | -0,01 | 9,0E-01 |
| Q69ZT9 | TBC1 domain family member 30 OS=Mus musculus OX=10090 G | 1 | 1 | 4 | 1 | 766 | 84,8 | 8,53 | 1472 | | 1 | | 1,047 | 0,799 | 1,01 | 1,083 | 1,131 | 0,988 | 0,826 | 1,042 | 0,99 | -0,01 | 9,0E-01 |
| Q9JMK2 | Casein kinase I isoform epsilon OS=Mus musculus OX=10090 GN | 8 | 3 | 3 | 2 | 416 | 47,3 | 9,66 | 1825 | | 3 | | 1,505 | 1,257 | 1,374 | 1,341 | 0,836 | 1,568 | 1,667 | 1,51 | 0,98 | -0,02 | 9,0E-01 |
| Q6PE01 | U5 small nuclear ribonucleoprotein 40 kDa protein OS=Mus mus | 4 | 1 | 1 | 1 | 358 | 39,3 | 8,1 | 712 | | 1 | | 0,778 | 0,53 | 0,886 | 1,185 | 0,64 | 0,683 | 1,071 | 1,081 | 0,97 | -0,03 | 9,0E-01 |
| P29387 | Guanine nucleotide-binding protein subunit beta-4 OS=Mus mu | 24 | 7 | 18 | 4 | 340 | 37,4 | 6,16 | 7229 | 8,45 | 5 | 2 | 0,92 | 1,117 | 0,838 | 1,017 | 0,881 | 0,919 | 1,065 | 1,068 | 0,99 | -0,01 | 9,0E-01 |
| P84099 | 60S ribosomal protein L19 OS=Mus musculus OX=10090 GN=Rpl | 26 | 6 | 8 | 6 | 196 | 23,5 | 11,47 | 517 | 11,97 | 1 | 5 | 1,015 | 0,874 | 0,989 | 1,054 | 0,871 | 1,025 | 0,947 | 1,059 | 1,01 | 0,01 | 9,0E-01 |
| Q8R5A6 | TBC1 domain family member 22A OS=Mus musculus OX=10090 | 2 | 1 | 1 | 1 | 516 | 59,3 | 6,46 | 292 | | 1 | | 1,13 | 1,112 | 0,946 | 1,173 | 1,244 | 1,011 | 1,317 | 0,851 | 0,99 | -0,01 | 9,0E-01 |
| Q8VBT6 | Apolipoprotein B receptor OS=Mus musculus OX=10090 GN=Apo | 1 | 1 | 3 | 1 | 942 | 102,6 | 4,36 | 690 | | 1 | | 0,944 | 1,219 | 0,786 | 1,066 | 1,055 | 0,779 | 1,096 | 1,024 | 1,02 | 0,02 | 9,0E-01 |
| P25444 | 40S ribosomal protein S2 OS=Mus musculus OX=10090 GN=Rps2 | 38 | 11 | 18 | 11 | 293 | 31,2 | 10,24 | 7595 | 18,56 | 7 | 4 | 0,96 | 0,989 | 1,109 | 1,022 | 0,975 | 1,014 | 1,037 | 1,074 | 1,00 | 0,00 | 9,0E-01 |
| Q91V76 | Ester hydrolase C11orf54 homolog OS=Mus musculus OX=10090 | 6 | 3 | 3 | 3 | 315 | 35 | 6,29 | 1111 | 0 | 2 | 1 | 1,077 | 1,203 | 1,007 | 1,334 | 1,177 | 1,033 | 1,156 | 1,302 | 0,99 | -0,01 | 9,0E-01 |
| O55106 | Striatin OS=Mus musculus OX=10090 GN=Strn PE=1 SV=2 | 11 | 5 | 7 | 5 | 780 | 85,9 | 5,27 | 3700 | | 5 | | 0,959 | 1,135 | 0,912 | 0,87 | 0,878 | 0,929 | 1,03 | 1,004 | 1,01 | 0,01 | 9,0E-01 |
| Q9WVE8 | Protein kinase C and casein kinase substrate in neurons protein | 3 | 2 | 3 | 1 | 486 | 55,8 | 5,2 | 1269 | | 2 | | 0,958 | 0,859 | 0,891 | 1,065 | 0,723 | 1,092 | 0,885 | 1,025 | 1,01 | 0,01 | 9,0E-01 |
| O35621 | Phosphomannomutase 1 OS=Mus musculus OX=10090 GN=Pmm | 16 | 4 | 6 | 3 | 262 | 29,8 | 5,47 | 3012 | 2,1 | 3 | 1 | 0,926 | 0,977 | 0,978 | 0,959 | 0,893 | 0,961 | 1,023 | 0,948 | 1,00 | 0,00 | 9,0E-01 |
| Q6IR34 | G-protein-signaling modulator 1 OS=Mus musculus OX=10090 G | 16 | 10 | 13 | 9 | 673 | 74,3 | 6,23 | 5187 | 12,83 | 6 | 5 | 0,881 | 0,961 | 0,88 | 0,998 | 1,072 | 0,889 | 0,939 | 0,849 | 0,99 | -0,01 | 9,0E-01 |
| Q3URE1 | Malonate--CoA ligase ACSF3, mitochondrial OS=Mus musculus C | 2 | 1 | 1 | 1 | 583 | 65 | 8,02 | 591 | | 1 | | 0,687 | 1,121 | 1,16 | 0,806 | 0,556 | 1,084 | 0,939 | 1,294 | 0,97 | -0,03 | 9,0E-01 |
| Q8BJU0 | Small glutamine-rich tetratricopeptide repeat-containing protein | 19 | 5 | 8 | 5 | 315 | 34,3 | 5,06 | 4212 | 6,2 | 4 | 1 | 0,986 | 0,841 | 1,013 | 1,01 | 0,807 | 1,074 | 0,978 | 0,956 | 1,01 | 0,01 | 9,0E-01 |
| Q9CXY6 | Interleukin enhancer-binding factor 2 OS=Mus musculus OX=100 | 6 | 3 | 3 | 3 | 390 | 43 | 5,26 | 669 | 0 | 1 | 2 | 1,009 | 0,978 | 1,26 | 1,242 | 1,14 | 1,043 | 1,131 | 1,217 | 0,99 | -0,01 | 9,0E-01 |
| Q99LU0 | Charged multivesicular body protein 1b-1 OS=Mus musculus OX | 9 | 2 | 2 | 2 | 199 | 22,1 | 8,1 | | 0 | | 2 | 1,258 | 1,463 | 1,22 | 1,093 | 1,2 | 1,152 | 1,358 | 1,279 | 1,01 | 0,01 | 9,0E-01 |
| P97742 | Carnitine O-palmitoyltransferase 1, liver isoform OS=Mus muscu | 5 | 3 | 3 | 3 | 773 | 88,2 | 8,62 | 965 | | 3 | | 0,828 | 1,119 | 1,005 | 1,202 | 1,246 | 0,902 | 0,958 | 0,992 | 1,01 | 0,01 | 9,0E-01 |
| P80317 | T-complex protein 1 subunit zeta OS=Mus musculus OX=10090 ( | 31 | 19 | 33 | 14 | 531 | 58 | 7,08 | 15255 | 18,25 | 15 | 5 | 0,988 | 0,926 | 1,284 | 0,934 | 1,311 | 0,98 | 0,991 | 0,912 | 0,99 | -0,01 | 9,0E-01 |
| P0C0A3 | Charged multivesicular body protein 6 OS=Mus musculus OX=10 | 18 | 3 | 4 | 3 | 200 | 23,4 | 5,44 | 2439 | | 3 | | 1,126 | 1,307 | 1,032 | 1,034 | 1,382 | 0,98 | 1,139 | 0,938 | 1,01 | 0,01 | 9,0E-01 |
| Q80TB8 | Synaptic vesicle membrane protein VAT-1 homolog-like OS=Mus | 14 | 5 | 6 | 5 | 417 | 45,8 | 5,06 | 2406 | 4,91 | 4 | 2 | 1,159 | 1,061 | 1,052 | 1,14 | 1,26 | 1,16 | 1,096 | 0,933 | 0,99 | -0,01 | 9,0E-01 |
| Q64455 | Receptor-type tyrosine-protein phosphatase eta OS=Mus muscu | 2 | 2 | 2 | 2 | 1238 | 136,7 | 5,57 | 1119 | | 2 | | 0,87 | 0,979 | 0,999 | 1,1 | 1,043 | 1,001 | 0,883 | 0,992 | 1,01 | 0,01 | 9,0E-01 |
| Q8R326 | Paraspeckle component 1 OS=Mus musculus OX=10090 GN=Psp | 8 | 4 | 5 | 3 | 523 | 58,7 | 6,67 | 1501 | 2,57 | 3 | 1 | 0,764 | 0,909 | 1,089 | 1,193 | 1,073 | 0,932 | 0,884 | 1,121 | 0,99 | -0,01 | 9,1E-01 |

| | | | | | | | | | | | | | | | | | | | | | | | |
|---|---|---|---|---|---|---|---|---|---|---|---|---|---|---|---|---|---|---|---|---|---|---|---|
| Q61768 | Kinesin-1 heavy chain OS=Mus musculus OX=10090 GN=Kif5b PE | 23 | 22 | 27 | 14 | 963 | 109,5 | 6,44 | 9623 | 21,79 | 15 | 7 | 1,031 | 1,011 | 1,016 | 1,064 | 1,059 | 0,985 | 1,086 | 1,005 | 1,00 | 0,00 | 9,1E-01 |
| Q9EPL8 | Importin-7 OS=Mus musculus OX=10090 GN=Ipo7 PE=1 SV=2 | 8 | 7 | 11 | 7 | 1038 | 119,4 | 4,82 | 5749 | 2,44 | 7 | 1 | 1,126 | 0,972 | 1,07 | 1,083 | 1,102 | 1,009 | 1,024 | 1,096 | 1,00 | 0,00 | 9,1E-01 |
| Q9CR21 | Acyl carrier protein, mitochondrial OS=Mus musculus OX=10090 | 15 | 2 | 5 | 2 | 156 | 17,4 | 5,21 | | 13,43 | | 2 | 0,952 | 1,097 | 0,834 | 1,086 | 1,001 | 1,021 | 0,919 | 1,062 | 0,99 | -0,01 | 9,1E-01 |
| Q7TNM2 | Tripartite motif-containing protein 46 OS=Mus musculus OX=10 | 5 | 3 | 3 | 3 | 759 | 83,4 | 7,65 | 1686 | | 3 | | 1,051 | 1,165 | 1,305 | 1,232 | 1,456 | 1,008 | 1,149 | 1,085 | 1,01 | 0,01 | 9,1E-01 |
| Q9D0E3 | LysM and putative peptidoglycan-binding domain-containing pr | 17 | 3 | 3 | 3 | 226 | 24,8 | 8,92 | | 8,58 | | 3 | 0,992 | 1,064 | 1,287 | 1,262 | 1,285 | 1,043 | 1,086 | 1,147 | 1,01 | 0,01 | 9,1E-01 |
| Q8C754 | Vacuolar protein sorting-associated protein 52 homolog OS=Mu | 6 | 3 | 3 | 3 | 723 | 82 | 5,9 | 1634 | | 3 | | 0,851 | 0,687 | 0,898 | 0,766 | 0,763 | 0,858 | 0,86 | 0,748 | 0,99 | -0,01 | 9,1E-01 |
| Q78ZA7 | Nucleosome assembly protein 1-like 4 OS=Mus musculus OX=10 | 19 | 6 | 10 | 5 | 375 | 42,7 | 4,67 | 6244 | 2 | 5 | 1 | 1,066 | 1,074 | 0,855 | 1,03 | 0,981 | 0,952 | 0,996 | 1,127 | 0,99 | -0,01 | 9,1E-01 |
| Q9DBP5 | UMP-CMP kinase OS=Mus musculus OX=10090 GN=Cmpk1 PE=1 | 20 | 4 | 8 | 4 | 196 | 22,2 | 5,83 | 2531 | 0 | 4 | 1 | 0,995 | 1,031 | 0,916 | 1,002 | 0,969 | 1,146 | 0,976 | 0,819 | 1,01 | 0,01 | 9,1E-01 |
| P21126 | Ubiquitin-like protein 4A OS=Mus musculus OX=10090 GN=Ubl4 | 19 | 3 | 3 | 3 | 157 | 17,8 | 8,44 | 1620 | | 3 | | 0,884 | 0,968 | 0,851 | 0,757 | 0,896 | 0,702 | 1,034 | 0,788 | 1,01 | 0,01 | 9,1E-01 |
| O08709 | Peroxiredoxin-6 OS=Mus musculus OX=10090 GN=Prdx6 PE=1 SV | 34 | 6 | 13 | 6 | 224 | 24,9 | 6,01 | 5528 | 8,47 | 5 | 2 | 0,888 | 1,151 | 0,897 | 1,073 | 1,014 | 0,918 | 1,054 | 0,989 | 1,01 | 0,01 | 9,1E-01 |
| Q9CQR6 | Serine/threonine-protein phosphatase 6 catalytic subunit OS=M | 8 | 3 | 4 | 3 | 305 | 35,1 | 5,69 | 2530 | | 3 | | 0,916 | 0,811 | 0,892 | 1,104 | 0,625 | 1,161 | 0,884 | 0,992 | 1,02 | 0,02 | 9,1E-01 |
| P70704 | Phospholipid-transporting ATPase IA OS=Mus musculus OX=100 | 7 | 7 | 8 | 7 | 1164 | 131,3 | 6,84 | 5469 | 3,69 | 6 | 1 | 0,925 | 0,903 | 0,962 | 0,916 | 0,658 | 1,095 | 1,005 | 0,994 | 0,99 | -0,01 | 9,1E-01 |
| Q80TY0 | Formin-binding protein 1 OS=Mus musculus OX=10090 GN=Fnb | 4 | 3 | 3 | 3 | 616 | 71,3 | 5,67 | 718 | 1,73 | 2 | 1 | 1,052 | 0,967 | 0,956 | 1,04 | 0,921 | 1,159 | 0,873 | 1,096 | 0,99 | -0,01 | 9,1E-01 |
| Q921E2 | Ras-related protein Rab-31 OS=Mus musculus OX=10090 GN=Ra | 26 | 4 | 4 | 4 | 194 | 21,3 | 7,4 | 2336 | 2,11 | 3 | 1 | 0,955 | 0,997 | 1,14 | 0,957 | 1,011 | 1,018 | 1,041 | 0,957 | 1,01 | 0,01 | 9,1E-01 |
| Q9JLV5 | Cullin-3 OS=Mus musculus OX=10090 GN=Cul3 PE=1 SV=1 | 11 | 7 | 9 | 7 | 768 | 88,9 | 8,46 | 2563 | 12,23 | 4 | 4 | 1,001 | 1,119 | 0,989 | 1,06 | 1,084 | 1,004 | 1,028 | 1,037 | 1,00 | 0,00 | 9,1E-01 |
| Q61753 | D-3-phosphoglycerate dehydrogenase OS=Mus musculus OX=10 | 29 | 15 | 27 | 15 | 533 | 56,5 | 6,54 | 8729 | 44,87 | 8 | 8 | 0,909 | 0,922 | 1,106 | 1,076 | 1,069 | 0,935 | 1,005 | 1,031 | 0,99 | -0,01 | 9,1E-01 |
| Q80UJ7 | Rab3 GTPase-activating protein catalytic subunit OS=Mus musc | 4 | 4 | 5 | 4 | 981 | 110,1 | 5,73 | 2720 | | 4 | | 1,008 | 0,817 | 1,166 | 0,995 | 1,075 | 0,865 | 1,089 | 0,997 | 0,99 | -0,01 | 9,1E-01 |
| P35290 | Ras-related protein Rab-24 OS=Mus musculus OX=10090 GN=Ra | 10 | 2 | 4 | 2 | 203 | 23,1 | 6,23 | 543 | 7,29 | 1 | 1 | 0,953 | 0,838 | 1,063 | 0,934 | 1,021 | 0,881 | 1,04 | 0,875 | 0,99 | -0,01 | 9,1E-01 |
| Q7TSH4 | Centriolar coiled-coil protein of 110 kDa OS=Mus musculus OX= | 1 | 1 | 1 | 1 | 1004 | 111,1 | 8,85 | 364 | | 1 | | 1,105 | 1,218 | 0,84 | 1,184 | 1,02 | 0,873 | 1,091 | 1,307 | 1,01 | 0,01 | 9,1E-01 |
| Q9JJA9 | General receptor for phosphoinositides 1-associated scaffold pr | 2 | 1 | 1 | 1 | 392 | 42,3 | 9,1 | 459 | | 1 | | 0,815 | 0,994 | 0,944 | 0,882 | 1,151 | 0,882 | 0,728 | 0,83 | 1,01 | 0,01 | 9,1E-01 |
| P70460 | Vasodilator-stimulated phosphoprotein OS=Mus musculus OX=1 | 7 | 2 | 2 | 2 | 375 | 39,6 | 8,53 | 924 | | 2 | | 0,987 | 0,89 | 0,994 | 0,981 | 0,933 | 1,025 | 1,058 | 0,859 | 0,99 | -0,01 | 9,1E-01 |
| Q9JM52 | Misshapen-like kinase 1 OS=Mus musculus OX=10090 GN=Mink | 5 | 6 | 7 | 2 | 1308 | 147,2 | 7,44 | 3516 | 4,95 | 4 | 2 | 0,808 | 0,843 | 0,773 | 0,822 | 0,893 | 0,878 | 0,797 | 0,699 | 0,99 | -0,01 | 9,1E-01 |
| Q8CGF6 | WD repeat-containing protein 47 OS=Mus musculus OX=10090 | 22 | 18 | 21 | 18 | 920 | 102,2 | 5,97 | 4897 | 25,03 | 7 | 11 | 1,063 | 0,979 | 1,133 | 0,945 | 1,017 | 1,114 | 0,955 | 1,01 | 1,01 | 0,01 | 9,1E-01 |
| Q8C761 | WD repeat-containing protein 60 OS=Mus musculus OX=10090 | 2 | 1 | 1 | 1 | 999 | 115,3 | 7,24 | 158 | | 1 | | 1,046 | 1,214 | 0,813 | 1,044 | 1,205 | 0,909 | 0,88 | 1,073 | 1,01 | 0,01 | 9,1E-01 |
| Q6ZWY8 | Thymosin beta-10 OS=Mus musculus OX=10090 GN=Tmsb10 PE | 48 | 5 | 6 | 3 | 44 | 5 | 5,36 | 744 | 14,3 | 1 | 4 | 0,746 | 0,704 | 1,218 | 0,821 | 0,771 | 1,32 | 0,675 | 0,635 | 1,03 | 0,03 | 9,2E-01 |
| P97412 | Lysosomal-trafficking regulator OS=Mus musculus OX=10090 GN | 0 | 1 | 1 | 1 | 3788 | 425 | 6,6 | 224 | | 1 | | 1,198 | 0,954 | 0,672 | 0,856 | 0,803 | 0,767 | 0,732 | 1,47 | 0,98 | -0,02 | 9,2E-01 |
| P50096 | Inosine-5'-monophosphate dehydrogenase 1 OS=Mus musculus | 6 | 3 | 3 | 2 | 514 | 55,2 | 6,8 | 754 | 5,32 | 1 | 2 | 1,039 | 0,904 | 0,947 | 1,123 | 1,256 | 0,997 | 0,833 | 0,971 | 0,99 | -0,01 | 9,2E-01 |
| Q9QXW9 | Large neutral amino acids transporter small subunit 2 OS=Mus m | 2 | 1 | 1 | 1 | 531 | 57,8 | 6,67 | | 2,62 | | 1 | 1,209 | 1,31 | 1,257 | 1,15 | 1,025 | 1,261 | 1,271 | 1,336 | 1,01 | 0,01 | 9,2E-01 |
| Q99LY9 | NADH dehydrogenase [ubiquinone] iron-sulfur protein 5 OS=Mu | 38 | 4 | 7 | 4 | 106 | 12,6 | 8,92 | 2882 | 2,41 | 3 | 2 | 0,89 | 0,962 | 1,067 | 1,087 | 1,098 | 0,914 | 1,035 | 0,985 | 0,99 | -0,01 | 9,2E-01 |
| O35945 | Aldehyde dehydrogenase, cytosolic 1 OS=Mus musculus OX=100 | 10 | 5 | 6 | 2 | 501 | 54,6 | 7,71 | 867 | 15,64 | 1 | 4 | 2,644 | 2,162 | 1,916 | 2,397 | 3,404 | 1,737 | 2,139 | 2,01 | 0,98 | -0,02 | 9,2E-01 |
| Q91WK5 | Glycine cleavage system H protein, mitochondrial OS=Mus musc | 6 | 1 | 1 | 1 | 170 | 18,6 | 4,75 | 486 | | 1 | | 1,275 | 1,413 | 1,054 | 1,453 | 1,248 | 1,083 | 1,414 | 1,505 | 0,99 | -0,01 | 9,2E-01 |
| P15532 | Nucleoside diphosphate kinase A OS=Mus musculus OX=10090 | 55 | 9 | 30 | 4 | 152 | 17,2 | 7,37 | 8661 | 41,71 | 7 | 4 | 1,04 | 1,044 | 0,936 | 0,936 | 1,197 | 0,961 | 0,911 | 0,853 | 1,01 | 0,01 | 9,2E-01 |
| Q9DCI9 | 39S ribosomal protein L32, mitochondrial OS=Mus musculus OX | 12 | 2 | 2 | 2 | 187 | 21,7 | 9,7 | 745 | | 2 | | 0,847 | 1,238 | 1,089 | 0,968 | 0,963 | 1,003 | 0,973 | 1,164 | 1,01 | 0,01 | 9,2E-01 |
| P97449 | Aminopeptidase N OS=Mus musculus OX=10090 GN=Anpep PE= | 3 | 2 | 2 | 2 | 966 | 109,6 | 5,9 | 893 | | 2 | | 0,809 | 0,907 | 0,919 | 1,185 | 1,3 | 0,804 | 0,905 | 0,867 | 0,99 | -0,01 | 9,2E-01 |
| Q9CQE1 | Protein NipSnap homolog 3B OS=Mus musculus OX=10090 GN= | 4 | 1 | 1 | 1 | 247 | 28,3 | 9,48 | | 3,46 | | 1 | 0,908 | 0,895 | 1,313 | 0,721 | 1,464 | 0,891 | 0,776 | 0,789 | 0,98 | -0,02 | 9,2E-01 |
| P62315 | Small nuclear ribonucleoprotein Sm D1 OS=Mus musculus OX=1 | 9 | 1 | 1 | 1 | 119 | 13,3 | 11,56 | 489 | | 1 | | 0,787 | 0,919 | 0,872 | 1,122 | 1,013 | 0,839 | 0,904 | 0,912 | 1,01 | 0,01 | 9,2E-01 |
| Q91YK0 | Leucine-rich repeat-containing protein 49 OS=Mus musculus OX | 1 | 1 | 1 | 1 | 686 | 78,8 | 8,13 | 595 | | 1 | | 1,297 | 0,928 | 0,993 | 0,814 | 1,155 | 0,951 | 0,984 | 0,895 | 1,01 | 0,01 | 9,2E-01 |
| Q60737 | Casein kinase II subunit alpha OS=Mus musculus OX=10090 GN= | 26 | 9 | 16 | 9 | 391 | 45,1 | 7,74 | 6932 | 0 | 8 | 1 | 0,97 | 0,921 | 0,967 | 1,026 | 1,181 | 0,9 | 0,928 | 0,844 | 1,01 | 0,01 | 9,2E-01 |
| Q61035 | Histidine--tRNA ligase, cytoplasmic OS=Mus musculus OX=10090 | 13 | 6 | 10 | 4 | 509 | 57,4 | 6 | 6004 | | 6 | | 1,041 | 1,075 | 1,107 | 0,988 | 1,054 | 1,05 | 0,996 | 1,098 | 1,00 | 0,00 | 9,2E-01 |
| Q9ERI6 | Retinol dehydrogenase 14 OS=Mus musculus OX=10090 GN=Rdl | 3 | 1 | 1 | 1 | 334 | 36,3 | 8,18 | | 2,47 | | 1 | 0,778 | 0,903 | 0,955 | 0,929 | 0,966 | 0,83 | 0,998 | 0,743 | 1,01 | 0,01 | 9,2E-01 |
| P09242 | Alkaline phosphatase, tissue-nonspecific isozyme OS=Mus musc | 2 | 1 | 1 | 1 | 524 | 57,5 | 7,01 | | 2,42 | | 1 | 1,202 | 1,118 | 1,113 | 1,034 | 1,016 | 1,245 | 1,088 | 1,141 | 0,99 | -0,01 | 9,3E-01 |
| P03930 | ATP synthase protein 8 OS=Mus musculus OX=10090 GN=Mtatp | 27 | 2 | 4 | 2 | 67 | 7,8 | 9,88 | 2775 | | 2 | | 1,003 | 0,716 | 1,251 | 0,806 | 1,022 | 0,967 | 0,91 | 0,924 | 0,99 | -0,01 | 9,3E-01 |
| P17751 | Triosephosphate isomerase OS=Mus musculus OX=10090 GN=T | 31 | 9 | 26 | 9 | 299 | 32,2 | 5,74 | 11851 | 18,88 | 6 | 4 | 1,021 | 1,013 | 1,104 | 1,077 | 1,19 | 1,095 | 0,958 | 0,994 | 0,99 | -0,01 | 9,3E-01 |
| Q7M759 | Alpha/beta hydrolase domain-containing protein 17B OS=Mus m | 2 | 1 | 1 | 1 | 288 | 32,2 | 6,29 | 481 | | 1 | | 1,228 | 0,925 | 1,304 | 1,063 | 1,37 | 1,052 | 0,99 | 1,062 | 1,01 | 0,01 | 9,3E-01 |
| Q69ZW3 | EH domain-binding protein 1 OS=Mus musculus OX=10090 GN= | 6 | 6 | 6 | 6 | 1231 | 139 | 5,38 | 988 | 7,2 | 2 | 4 | 0,888 | 0,979 | 0,991 | 0,838 | 0,845 | 0,935 | 1,004 | 0,893 | 1,01 | 0,01 | 9,3E-01 |
| Q61548 | Clathrin coat assembly protein AP180 OS=Mus musculus OX=10 | 17 | 16 | 42 | 15 | 901 | 91,8 | 4,88 | 11203 | 60,36 | 8 | 11 | 1,039 | 0,993 | 0,943 | 1,026 | 0,821 | 1,175 | 0,931 | 1,044 | 1,01 | 0,01 | 9,3E-01 |
| Q9DCP2 | Sodium-coupled neutral amino acid transporter 3 OS=Mus musc | 11 | 4 | 10 | 4 | 505 | 55,6 | 7,09 | 4725 | 5,27 | 3 | 1 | 1,059 | 1,022 | 0,897 | 0,986 | 1,044 | 0,929 | 0,929 | 1,044 | 1,00 | 0,00 | 9,3E-01 |
| Q8VEJ9 | Vacuolar protein sorting-associated protein 4A OS=Mus musculu | 8 | 3 | 4 | 3 | 437 | 48,9 | 7,8 | 1266 | 6,51 | 2 | 1 | 1,07 | 1,049 | 0,887 | 0,871 | 1,337 | 0,799 | 0,833 | 0,958 | 0,99 | -0,01 | 9,3E-01 |
| Q6PH08 | ERC protein 2 OS=Mus musculus OX=10090 GN=Erc2 PE=1 SV=2 | 8 | 9 | 13 | 5 | 957 | 110,6 | 7,03 | 7301 | 2,28 | 8 | 1 | 1,071 | 1,001 | 0,901 | 1,115 | 0,99 | 0,902 | 1,172 | 1,051 | 0,99 | -0,01 | 9,3E-01 |
| Q80U56 | Late secretory pathway protein AVL9 homolog OS=Mus musculu | 7 | 4 | 5 | 4 | 649 | 72,1 | 6,14 | 2750 | | 4 | | 1,061 | 0,977 | 1,022 | 1,055 | 1,121 | 0,996 | 0,953 | 1,06 | 1,00 | 0,00 | 9,3E-01 |
| Q99KJ8 | Dynactin subunit 2 OS=Mus musculus OX=10090 GN=Dctn2 PE= | 43 | 14 | 23 | 14 | 402 | 44,1 | 5,26 | 11293 | 12,81 | 11 | 5 | 0,964 | 1,044 | 0,979 | 1,03 | 0,944 | 0,956 | 1,076 | 1,055 | 1,00 | 0,00 | 9,3E-01 |
| Q9D328 | Transmembrane protein 35A OS=Mus musculus OX=10090 GN= | 9 | 1 | 2 | 1 | 167 | 18,5 | 9,99 | 1278 | | 1 | | 0,825 | 0,951 | 0,868 | 0,81 | 0,804 | 0,898 | 0,953 | 0,816 | 1,00 | 0,00 | 9,3E-01 |
| Q9Z1N7 | AT-rich interactive domain-containing protein 3B OS=Mus musc | 2 | 1 | 1 | 1 | 568 | 61 | 7,47 | 344 | | 1 | | 0,844 | 0,861 | 0,902 | 1,021 | 0,622 | 0,908 | 0,98 | 1,08 | 1,01 | 0,01 | 9,3E-01 |
| Q9WTP7 | GTP:AMP phosphotransferase AK3, mitochondrial OS=Mus mus | 35 | 7 | 13 | 7 | 227 | 25,4 | 8,84 | 6018 | 8,39 | 5 | 3 | 0,969 | 1,033 | 0,933 | 1,095 | 1,001 | 0,9 | 1,05 | 1,061 | 1,00 | 0,00 | 9,3E-01 |
| Q9WV96 | Mitochondrial import inner membrane translocase subunit Tim | 19 | 1 | 1 | 1 | 100 | 11,3 | 7,08 | | 4,01 | | 1 | 1,395 | 1,019 | 0,796 | 0,762 | 1,362 | 0,708 | 1,109 | 0,864 | 0,98 | -0,02 | 9,3E-01 |
| P51830 | Adenylate cyclase type 9 OS=Mus musculus OX=10090 GN=Adc | 3 | 4 | 4 | 4 | 1353 | 150,9 | 7,21 | 1118 | 5,93 | 2 | 2 | 0,902 | 1,085 | 0,934 | 0,943 | 0,995 | 0,942 | 0,995 | 0,947 | 1,00 | 0,00 | 9,3E-01 |
| Q9CQU0 | Thioredoxin domain-containing protein 12 OS=Mus musculus OX | 39 | 6 | 8 | 6 | 170 | 19 | 5,26 | 3674 | | 6 | | 0,957 | 1,015 | 1,055 | 1,141 | 1,205 | 0,966 | 0,995 | 1,025 | 0,99 | -0,01 | 9,3E-01 |
| Q9JIG8 | PRA1 family protein 2 OS=Mus musculus OX=10090 GN=Praf2 P | 6 | 1 | 2 | 1 | 178 | 19,5 | 9,6 | 661 | | 1 | | 0,862 | 1,122 | 0,834 | 0,862 | 0,88 | 0,719 | 1,004 | 1,043 | 1,01 | 0,01 | 9,3E-01 |
| P55066 | Neurocan core protein OS=Mus musculus OX=10090 GN=Ncan F | 1 | 1 | 2 | 1 | 1268 | 137,1 | 5,72 | 1043 | | 1 | | 1,013 | 0,964 | 0,97 | 0,859 | 1,014 | 0,947 | 0,921 | 0,937 | 1,00 | 0,00 | 9,4E-01 |
| O08688 | Calpain-5 OS=Mus musculus OX=10090 GN=Capn5 PE=1 SV=1 | 16 | 8 | 9 | 8 | 640 | 72,9 | 7,36 | 4836 | 3,25 | 6 | 2 | 0,998 | 0,892 | 1,071 | 0,887 | 0,855 | 1,072 | 0,927 | 1,016 | 0,99 | -0,01 | 9,4E-01 |
| Q8BUV3 | Gephyrin OS=Mus musculus OX=10090 GN=Gphn PE=1 SV=2 | 16 | 10 | 13 | 10 | 769 | 83,2 | 5,6 | 3964 | 6,68 | 8 | 2 | 0,944 | 0,93 | 1,077 | 0,967 | 0,877 | 1,1 | 0,981 | 0,979 | 1,00 | 0,00 | 9,4E-01 |
| Q99N89 | 39S ribosomal protein L43, mitochondrial OS=Mus musculus OX | 6 | 1 | 2 | 1 | 183 | 20,2 | 9,52 | 948 | | 1 | | 1,141 | 1,031 | 1,086 | 1,278 | 1,309 | 1,079 | 0,982 | 1,137 | 1,01 | 0,01 | 9,4E-01 |
| Q8BH66 | Atlastin-1 OS=Mus musculus OX=10090 GN=Atl1 PE=1 SV=1 | 13 | 6 | 11 | 5 | 558 | 63,3 | 6,49 | 5659 | 3,14 | 5 | 1 | 1,048 | 0,952 | 0,996 | 0,9 | 0,921 | 1,064 | 0,957 | 0,939 | 1,00 | 0,00 | 9,4E-01 |
| Q62407 | Striated muscle-specific serine/threonine-protein kinase OS=Mu | 1 | 2 | 4 | 2 | 3262 | 354,1 | 8,21 | 1445 | | 2 | | 1,062 | 1,068 | 1,025 | 0,978 | 1,201 | 0,904 | 1,122 | 0,879 | 1,01 | 0,01 | 9,4E-01 |
| P14685 | 26S proteasome non-ATPase regulatory subunit 3 OS=Mus musc | 18 | 9 | 10 | 9 | 530 | 60,7 | 8,44 | 4604 | 5,44 | 8 | 2 | 0,975 | 0,944 | 1,118 | 0,807 | 0,876 | 0,949 | 0,944 | 1,051 | 1,01 | 0,01 | 9,4E-01 |

| | | | | | | | | | | | | | | | | | | | | | | | |
|---|---|---|---|---|---|---|---|---|---|---|---|---|---|---|---|---|---|---|---|---|---|---|---|
| Q9D1D4 | Transmembrane emp24 domain-containing protein 10 OS=Mus | 9 | 2 | 2 | 2 | 219 | 24,9 | 6,7 | 558 | 2,76 | 1 | 1 | 1,211 | 1,149 | 1,228 | 1,385 | 1,414 | 1,086 | 1,305 | 1,196 | 0,99 | -0,01 | 9,4E-01 |
| Q9QWR8 | Alpha-N-acetylgalactosaminidase OS=Mus musculus OX=10090 ( | 4 | 2 | 2 | 2 | 415 | 47,2 | 6,44 | 940 | | 2 | | 1,115 | 0,837 | 1,237 | 1,455 | 1,05 | 1,241 | 1,165 | 1,232 | 0,99 | -0,01 | 9,4E-01 |
| Q9R0P3 | S-formylglutathione hydrolase OS=Mus musculus OX=10090 GN | 13 | 3 | 5 | 3 | 282 | 31,3 | 7,12 | 2577 | 4,28 | 2 | 1 | 0,983 | 0,857 | 1,179 | 0,992 | 0,953 | 1,096 | 1,061 | 0,926 | 0,99 | -0,01 | 9,4E-01 |
| Q7TME0 | Phospholipid phosphatase-related protein type 4 OS=Mus musc | 21 | 12 | 14 | 12 | 766 | 83,2 | 8,84 | 6402 | 10,91 | 7 | 5 | 1,035 | 0,967 | 1,023 | 0,954 | 0,887 | 1,043 | 0,976 | 1,059 | 1,00 | 0,00 | 9,4E-01 |
| O09167 | 60S ribosomal protein L21 OS=Mus musculus OX=10090 GN=Rpl | 20 | 4 | 6 | 4 | 160 | 18,6 | 10,49 | 686 | 11,29 | 1 | 3 | 1,021 | 0,932 | 0,962 | 1,006 | 1,002 | 1 | 0,989 | 0,938 | 1,00 | 0,00 | 9,4E-01 |
| Q8R191 | Synaptogyrin-3 OS=Mus musculus OX=10090 GN=Syngr3 PE=1 S | 11 | 3 | 12 | 3 | 229 | 24,5 | 8,18 | 6759 | 2,59 | 2 | 1 | 1,063 | 1,084 | 1,116 | 1,093 | 1,167 | 0,995 | 1,129 | 1,077 | 1,00 | 0,00 | 9,4E-01 |
| Q922D4 | Serine/threonine-protein phosphatase 6 regulatory subunit 3 OS | 5 | 4 | 4 | 4 | 844 | 94,6 | 4,58 | 1093 | 0 | 2 | 2 | 0,962 | 1,094 | 0,688 | 1,256 | 0,586 | 1,09 | 1,268 | 0,998 | 1,01 | 0,01 | 9,4E-01 |
| P68181 | cAMP-dependent protein kinase catalytic subunit beta OS=Mus | 29 | 12 | 23 | 5 | 351 | 40,7 | 8,56 | 5576 | 34,01 | 5 | 8 | 0,987 | 0,842 | 0,958 | 0,965 | 0,843 | 1,054 | 0,898 | 0,94 | 1,00 | 0,00 | 9,4E-01 |
| Q91ZX7 | Prolow-density lipoprotein receptor-related protein 1 OS=Mus r | 15 | 58 | 74 | 57 | 4545 | 504,4 | 5,36 | 32226 | 46,21 | 43 | 18 | 0,97 | 0,927 | 1,015 | 0,97 | 0,959 | 0,952 | 0,98 | 0,985 | 1,00 | 0,00 | 9,4E-01 |
| P05480 | Neuronal proto-oncogene tyrosine-protein kinase Src OS=Mus n | 20 | 10 | 18 | 8 | 541 | 60,6 | 7,84 | 3718 | 21,51 | 4 | 6 | 1,024 | 0,898 | 0,964 | 0,997 | 1,001 | 1,008 | 0,905 | 0,958 | 1,00 | 0,00 | 9,4E-01 |
| Q8R016 | Bleomycin hydrolase OS=Mus musculus OX=10090 GN=Blmh PE | 23 | 10 | 14 | 10 | 455 | 52,5 | 6,48 | 5228 | 11,58 | 7 | 4 | 1,105 | 1,024 | 1,127 | 1,199 | 0,901 | 1,403 | 1,073 | 1,044 | 1,01 | 0,01 | 9,4E-01 |
| Q8R1A4 | Dedicator of cytokinesis protein 7 OS=Mus musculus OX=10090 | 4 | 9 | 11 | 9 | 2130 | 241,3 | 6,71 | 3975 | 4,3 | 7 | 2 | 1,072 | 0,974 | 1,102 | 1,001 | 1,309 | 0,998 | 0,981 | 0,89 | 0,99 | -0,01 | 9,4E-01 |
| Q9CX84 | Regulator of G-protein signaling 19 OS=Mus musculus OX=1009( | 6 | 1 | 1 | 1 | 216 | 24,7 | 6 | 773 | | 1 | | 1,333 | 1,672 | 1,444 | 1,347 | 1,749 | 1,281 | 1,197 | 1,525 | 1,01 | 0,01 | 9,4E-01 |
| Q9D2M8 | Ubiquitin-conjugating enzyme E2 variant 2 OS=Mus musculus O | 15 | 3 | 6 | 2 | 145 | 16,4 | 8,09 | 1951 | 6,81 | 2 | 1 | 0,942 | 0,94 | 1,135 | 1,068 | 0,911 | 1,151 | 0,92 | 1,127 | 0,99 | -0,01 | 9,4E-01 |
| Q91W86 | Vacuolar protein sorting-associated protein 11 homolog OS=Mu | 1 | 1 | 1 | 1 | 941 | 107,7 | 7,01 | 675 | | 1 | | 1,826 | 1,516 | 1,862 | 1,734 | 1,652 | 1,994 | 1,682 | 1,644 | 1,00 | 0,00 | 9,4E-01 |
| Q9R1P3 | Proteasome subunit beta type-2 OS=Mus musculus OX=10090 G | 27 | 6 | 9 | 6 | 201 | 22,9 | 7,02 | 4650 | 2,89 | 5 | 2 | 0,926 | 0,888 | 0,926 | 0,953 | 0,896 | 1,038 | 0,839 | 0,933 | 1,00 | 0,00 | 9,4E-01 |
| O08919 | Numb-like protein OS=Mus musculus OX=10090 GN=Numbl PE= | 10 | 6 | 8 | 6 | 604 | 64,1 | 8,82 | 2666 | 7,27 | 3 | 3 | 0,966 | 0,896 | 0,961 | 0,843 | 1,004 | 0,804 | 0,944 | 0,929 | 1,00 | 0,00 | 9,4E-01 |
| Q9QUJ7 | Long-chain-fatty-acid--CoA ligase 4 OS=Mus musculus OX=1009( | 4 | 3 | 3 | 3 | 711 | 79 | 8,28 | 1037 | 1,89 | 2 | 1 | 0,987 | 1,145 | 0,924 | 1,024 | 1,207 | 0,995 | 0,879 | 1,023 | 0,99 | -0,01 | 9,4E-01 |
| P26040 | Ezrin OS=Mus musculus OX=10090 GN=Ezr PE=1 SV=3 | 13 | 8 | 12 | 3 | 586 | 69,4 | 6,1 | 4789 | 2,76 | 6 | 2 | 1,202 | 1,012 | 0,853 | 0,985 | 1,251 | 0,784 | 0,971 | 1,081 | 0,99 | -0,01 | 9,4E-01 |
| Q01063 | cAMP-specific 3',5'-cyclic phosphodiesterase 4D OS=Mus muscu | 1 | 1 | 1 | 1 | 747 | 84,5 | 4,91 | 738 | | 1 | | 0,74 | 1,019 | 0,894 | 0,581 | 0,726 | 0,702 | 0,776 | 0,999 | 1,01 | 0,01 | 9,5E-01 |
| P23116 | Eukaryotic translation initiation factor 3 subunit A OS=Mus mus | 10 | 13 | 14 | 13 | 1344 | 161,8 | 6,77 | 6283 | 2,09 | 12 | 2 | 1,024 | 1,045 | 1,027 | 1,053 | 0,888 | 0,968 | 1,156 | 1,155 | 1,00 | 0,00 | 9,5E-01 |
| Q4KUS2 | Protein unc-13 homolog A OS=Mus musculus OX=10090 GN=Un | 3 | 5 | 5 | 5 | 1712 | 193,7 | 5,25 | 2497 | 2,22 | 4 | 1 | 1,071 | 1,032 | 1,089 | 1,064 | 1,136 | 1,028 | 1,044 | 1,055 | 1,00 | 0,00 | 9,5E-01 |
| Q3THK7 | GMP synthase [glutamine-hydrolyzing] OS=Mus musculus OX=1( | 15 | 8 | 9 | 8 | 693 | 76,7 | 6,73 | 4048 | 0 | 6 | 2 | 0,942 | 0,882 | 0,892 | 0,901 | 0,939 | 0,853 | 0,924 | 0,895 | 1,00 | 0,00 | 9,5E-01 |
| O88851 | Putative hydrolase RBBP9 OS=Mus musculus OX=10090 GN=Rbb | 17 | 2 | 3 | 2 | 186 | 20,9 | 5,97 | 856 | 4,3 | 1 | 2 | 0,962 | 0,735 | 1,074 | 0,912 | 0,791 | 0,963 | 1,039 | 0,867 | 1,01 | 0,01 | 9,5E-01 |
| Q8BH58 | TIP41-like protein OS=Mus musculus OX=10090 GN=Tiprl PE=1 S | 15 | 3 | 5 | 3 | 271 | 31,2 | 5,63 | 2735 | | 3 | | 1,063 | 0,982 | 0,896 | 1,058 | 1,136 | 0,953 | 1,008 | 0,885 | 1,00 | 0,00 | 9,5E-01 |
| A2AJA9 | Apical junction component 1 homolog OS=Mus musculus OX=1( | 1 | 1 | 1 | 1 | 974 | 107,1 | 9,11 | 692 | | 1 | | 1,24 | 1,032 | 1,406 | 1,367 | 1,232 | 1,355 | 1,586 | 0,914 | 0,99 | -0,01 | 9,5E-01 |
| Q68FH0 | Plakophilin-4 OS=Mus musculus OX=10090 GN=Pkp4 PE=1 SV=1 | 8 | 7 | 8 | 5 | 1190 | 131,5 | 8,94 | 1489 | 8,68 | 4 | 4 | 1,003 | 0,898 | 1,199 | 0,914 | 1,054 | 0,928 | 1,136 | 0,873 | 1,01 | 0,01 | 9,5E-01 |
| Q9JHR7 | Insulin-degrading enzyme OS=Mus musculus OX=10090 GN=Ide | 7 | 8 | 14 | 8 | 1019 | 117,7 | 6,54 | 5180 | 7,38 | 5 | 3 | 1,048 | 1,014 | 1,058 | 0,979 | 0,943 | 1,115 | 1,03 | 1,001 | 1,00 | 0,00 | 9,5E-01 |
| Q791T5 | Mitochondrial carrier homolog 1 OS=Mus musculus OX=10090 G | 5 | 2 | 2 | 2 | 389 | 41,5 | 9,32 | 776 | 2,51 | 1 | 1 | 1,106 | 0,9 | 1,052 | 0,902 | 0,907 | 0,979 | 1,019 | 1,04 | 1,00 | 0,00 | 9,5E-01 |
| Q6P1F6 | Serine/threonine-protein phosphatase 2A 55 kDa regulatory sub | 19 | 9 | 14 | 7 | 447 | 51,7 | 6,2 | 6236 | 6,99 | 6 | 3 | 1,068 | 0,831 | 1,163 | 0,918 | 1,026 | 1,016 | 1,093 | 0,867 | 0,99 | -0,01 | 9,5E-01 |
| Q6PEV3 | WAS/WASL-interacting protein family member 2 OS=Mus muscu | 4 | 1 | 1 | 1 | 440 | 46,3 | 10,99 | 657 | | 1 | | 0,909 | 0,913 | 1,086 | 1,026 | 1,406 | 0,716 | 0,717 | 1,138 | 0,99 | -0,01 | 9,5E-01 |
| Q60996 | Serine/threonine-protein phosphatase 2A 56 kDa regulatory sub | 4 | 2 | 3 | 1 | 524 | 60,8 | 6,64 | 705 | 5,67 | 1 | 1 | 1,021 | 0,995 | 1,206 | 0,904 | 0,893 | 1,095 | 1,147 | 0,97 | 1,01 | 0,01 | 9,5E-01 |
| Q9WUA6 | RAC-gamma serine/threonine-protein kinase OS=Mus musculus | 4 | 2 | 3 | 1 | 479 | 55,7 | 6,02 | 915 | 2,59 | 1 | 1 | 1,006 | 0,917 | 1,006 | 1,036 | 0,839 | 1,105 | 1,025 | 0,981 | 1,00 | 0,00 | 9,5E-01 |
| Q9JJV2 | Profilin-2 OS=Mus musculus OX=10090 GN=Pfn2 PE=1 SV=3 | 44 | 7 | 14 | 7 | 140 | 15 | 6,99 | 2516 | 21,86 | 4 | 6 | 0,871 | 0,935 | 1,192 | 0,922 | 0,857 | 1,262 | 0,868 | 0,962 | 0,99 | -0,01 | 9,5E-01 |
| Q80TZ3 | Putative tyrosine-protein phosphatase auxilin OS=Mus musculu | 5 | 5 | 9 | 3 | 938 | 102,2 | 7,23 | 4823 | | 5 | | 0,977 | 1,097 | 0,983 | 0,983 | 1,097 | 0,994 | 0,995 | 0,964 | 1,00 | 0,00 | 9,5E-01 |
| Q922Q8 | Leucine-rich repeat-containing protein 59 OS=Mus musculus OX | 2 | 1 | 1 | 1 | 307 | 34,9 | 9,52 | 760 | | 1 | | 1,016 | 0,892 | 1,074 | 1,1 | 0,886 | 0,888 | 1,213 | 1,118 | 0,99 | -0,01 | 9,5E-01 |
| Q9JI90 | E3 ubiquitin-protein ligase RNF14 OS=Mus musculus OX=10090 | 8 | 4 | 5 | 4 | 485 | 54,9 | 4,7 | 2563 | 2,15 | 3 | 1 | 1,006 | 1,011 | 1,098 | 1,129 | 1,07 | 1,007 | 1,068 | 1,108 | 1,00 | 0,00 | 9,5E-01 |
| Q9R1T4 | Septin-6 OS=Mus musculus OX=10090 GN=Septin6 PE=1 SV=4 | 29 | 12 | 18 | 5 | 434 | 49,6 | 6,43 | 9789 | 9,96 | 10 | 2 | 0,975 | 0,965 | 1,056 | 0,961 | 0,916 | 0,942 | 1,094 | 1,016 | 1,00 | 0,00 | 9,5E-01 |
| G3XA57 | Rab11 family-interacting protein 2 OS=Mus musculus OX=10090 | 5 | 2 | 2 | 2 | 512 | 58,2 | 9,45 | 1205 | | 2 | | 0,907 | 1,172 | 0,986 | 1,031 | 1,043 | 0,878 | 1,035 | 1,122 | 1,00 | 0,00 | 9,5E-01 |
| Q61625 | Glutamate receptor ionotropic, delta-2 OS=Mus musculus OX=1 | 1 | 1 | 1 | 1 | 1007 | 113 | 6,1 | 315 | | 1 | | 0,95 | 0,871 | 1,158 | 1,312 | 1,119 | 0,9 | 1,117 | 1,183 | 0,99 | -0,01 | 9,5E-01 |
| P10852 | 4F2 cell-surface antigen heavy chain OS=Mus musculus OX=100( | 31 | 17 | 28 | 17 | 526 | 58,3 | 5,91 | 8124 | 41,66 | 11 | 9 | 1,009 | 0,945 | 0,965 | 0,96 | 0,987 | 0,907 | 0,901 | 1,074 | 1,00 | 0,00 | 9,6E-01 |
| Q5SRX1 | TOM1-like protein 2 OS=Mus musculus OX=10090 GN=Tom1l2 P | 11 | 4 | 5 | 4 | 507 | 55,6 | 4,82 | 2421 | | 4 | | 0,874 | 0,97 | 0,853 | 0,968 | 0,882 | 0,883 | 1,027 | 0,884 | 1,00 | 0,00 | 9,6E-01 |
| Q5PRF0 | HEAT repeat-containing protein 5A OS=Mus musculus OX=1009( | 1 | 1 | 1 | 1 | 2038 | 219,8 | 6,76 | | 2,2 | | 1 | 1,231 | 1,265 | 1,011 | 0,901 | 1,321 | 0,862 | 1,236 | 1,02 | 0,99 | -0,01 | 9,6E-01 |
| Q6RI63 | Constitutive coactivator of peroxisome proliferator-activated re | 1 | 1 | 1 | 1 | 786 | 89,3 | 6,52 | 348 | | 1 | | 0,933 | 0,925 | 0,928 | 0,91 | 0,806 | 1,01 | 1,02 | 0,872 | 1,00 | 0,00 | 9,6E-01 |
| Q62132 | Receptor-type tyrosine-protein phosphatase R OS=Mus musculu | 2 | 1 | 2 | 1 | 656 | 74 | 7,81 | | 6,64 | | 1 | 0,855 | 0,788 | 0,9 | 0,918 | 0,842 | 1,024 | 0,814 | 0,767 | 1,00 | 0,00 | 9,6E-01 |
| Q8VDD5 | Myosin-9 OS=Mus musculus OX=10090 GN=Myh9 PE=1 SV=4 | 21 | 40 | 62 | 30 | 1960 | 226,2 | 5,66 | 19618 | 77,55 | 22 | 20 | 0,896 | 0,953 | 0,913 | 1,216 | 1,083 | 0,816 | 1,099 | 0,958 | 1,01 | 0,01 | 9,6E-01 |
| Q9D6U8 | Protein FAM162A OS=Mus musculus OX=10090 GN=Fam162a PI | 8 | 1 | 1 | 1 | 155 | 17,7 | 9,88 | 605 | | 1 | | 0,985 | 0,894 | 1,145 | 1 | 0,968 | 1,056 | 0,977 | 1,035 | 1,00 | 0,00 | 9,6E-01 |
| P62069 | Ubiquitin carboxyl-terminal hydrolase 46 OS=Mus musculus OX= | 2 | 1 | 1 | 1 | 366 | 42,4 | 6,83 | 589 | | 1 | | 1,024 | 0,965 | 1,136 | 0,948 | 1,244 | 0,967 | 0,893 | 0,95 | 1,00 | 0,00 | 9,6E-01 |
| Q5SVL6 | Rap1 GTPase-activating protein 2 OS=Mus musculus OX=10090 ( | 5 | 3 | 3 | 3 | 712 | 78,2 | 6,43 | 1322 | | 3 | | 1,405 | 1,208 | 1,222 | 1,122 | 1,047 | 1,174 | 1,563 | 1,146 | 1,01 | 0,01 | 9,6E-01 |
| P26041 | Moesin OS=Mus musculus OX=10090 GN=Msn PE=1 SV=3 | 10 | 6 | 10 | 1 | 577 | 67,7 | 6,6 | 4251 | 2,76 | 5 | 1 | 1,073 | 1,321 | 0,931 | 1,186 | 1,269 | 0,864 | 1,295 | 1,056 | 1,01 | 0,01 | 9,6E-01 |
| Q9D855 | Cytochrome b-c1 complex subunit 7 OS=Mus musculus OX=100( | 52 | 7 | 16 | 7 | 111 | 13,5 | 9,11 | 5217 | 6,96 | 4 | 3 | 1,071 | 1,041 | 0,877 | 1,095 | 0,943 | 0,993 | 1,059 | 1,077 | 1,00 | 0,00 | 9,6E-01 |
| P07309 | Transthyretin OS=Mus musculus OX=10090 GN=Ttr PE=1 SV=1 | 10 | 1 | 1 | 1 | 147 | 15,8 | 6,16 | 574 | | 1 | | 0,833 | 1,409 | 1,18 | 1,359 | 1,692 | 1,006 | 1,222 | 0,906 | 0,99 | -0,01 | 9,6E-01 |
| Q9D6J6 | NADH dehydrogenase [ubiquinone] flavoprotein 2, mitochondri | 32 | 7 | 11 | 7 | 248 | 27,3 | 7,4 | 5538 | 7,85 | 6 | 2 | 0,943 | 0,971 | 0,883 | 0,994 | 0,924 | 0,945 | 0,995 | 0,921 | 1,00 | 0,00 | 9,6E-01 |
| Q3UN02 | Lysocardiolipin acyltransferase 1 OS=Mus musculus OX=10090 G | 7 | 3 | 3 | 3 | 376 | 44,4 | 8,53 | 1085 | | 3 | | 1,166 | 1,058 | 1,175 | 1,144 | 1,138 | 1,228 | 0,894 | 1,301 | 1,00 | 0,00 | 9,6E-01 |
| P14602 | Heat shock protein beta-1 OS=Mus musculus OX=10090 GN=Hsp | 5 | 1 | 1 | 1 | 209 | 23 | 6,55 | | 3,04 | | 1 | 1,089 | 1,565 | 0,86 | 0,639 | 1,359 | 0,859 | 0,841 | 1,049 | 1,01 | 0,01 | 9,6E-01 |
| Q8CBE3 | WD repeat-containing protein 37 OS=Mus musculus OX=10090 ( | 13 | 6 | 12 | 6 | 496 | 55 | 7,23 | 6032 | 2,39 | 5 | 1 | 1,003 | 0,935 | 1,173 | 1,017 | 0,934 | 1,149 | 0,951 | 1,108 | 1,00 | 0,00 | 9,6E-01 |
| Q8BHJ7 | Gamma-aminobutyric acid receptor subunit alpha-5 OS=Mus mu | 8 | 3 | 4 | 2 | 463 | 52,2 | 9,01 | 786 | 10,75 | 1 | 2 | 0,982 | 0,974 | 0,949 | 1,061 | 0,85 | 0,976 | 0,989 | 1,164 | 1,00 | 0,00 | 9,6E-01 |
| Q9DC28 | Casein kinase I isoform delta OS=Mus musculus OX=10090 GN=( | 5 | 2 | 2 | 1 | 415 | 47,3 | 9,74 | 1540 | | 2 | | 1,113 | 0,922 | 1,056 | 1,18 | 0,881 | 1,191 | 1,092 | 1,091 | 1,00 | 0,00 | 9,6E-01 |
| B2RQC6 | CAD protein OS=Mus musculus OX=10090 GN=Cad PE=1 SV=1 | 5 | 9 | 12 | 9 | 2225 | 243,1 | 6,43 | 3036 | 14,19 | 4 | 5 | 1,045 | 1,012 | 1,107 | 1,028 | 1,232 | 0,997 | 0,95 | 1,025 | 1,00 | 0,00 | 9,7E-01 |
| Q9Z1Q9 | Valine--tRNA ligase OS=Mus musculus OX=10090 GN=Vars PE=1 | 8 | 6 | 6 | 6 | 1263 | 140,1 | 7,77 | 1714 | 2,11 | 5 | 1 | 0,973 | 1,042 | 1,038 | 1,074 | 0,994 | 1,01 | 1,105 | 1,012 | 1,00 | 0,00 | 9,7E-01 |
| Q3U186 | Probable arginine--tRNA ligase, mitochondrial OS=Mus musculu | 4 | 3 | 4 | 3 | 578 | 65,3 | 8,02 | 1473 | 2,06 | 2 | 1 | 0,952 | 1,061 | 1,224 | 1,083 | 1,266 | 1,077 | 0,831 | 1,127 | 1,00 | 0,00 | 9,7E-01 |
| Q6ZQ18 | Protein EFR3 homolog B OS=Mus musculus OX=10090 GN=Efr3b | 2 | 2 | 2 | 2 | 817 | 92,3 | 6,54 | 718 | 2,58 | 1 | 1 | 0,868 | 0,982 | 0,992 | 0,914 | 0,736 | 1,09 | 0,939 | 0,977 | 1,00 | 0,00 | 9,7E-01 |
| Q5DTT2 | PH and SEC7 domain-containing protein 1 OS=Mus musculus OX | 5 | 5 | 5 | 5 | 1024 | 109,6 | 7,05 | 981 | 1,84 | 2 | 3 | 0,984 | 0,907 | 0,943 | 0,906 | 0,864 | 0,897 | 0,962 | 1,024 | 1,00 | 0,00 | 9,7E-01 |
| Q91V09 | WD repeat-containing protein 13 OS=Mus musculus OX=10090 ( | 8 | 3 | 3 | 3 | 485 | 53,6 | 9,14 | 1402 | 1,78 | 2 | 1 | 0,997 | 0,707 | 1,006 | 0,838 | 0,943 | 0,672 | 1,062 | 0,852 | 1,01 | 0,01 | 9,7E-01 |
| Q64475 | Histone H2B type 1-B OS=Mus musculus OX=10090 GN=Hist1h2 | 41 | 7 | 10 | 7 | 126 | 13,9 | 10,32 | 4374 | 4,55 | 4 | 3 | 0,87 | 0,787 | 1,29 | 1,162 | 1,332 | 0,78 | 1,148 | 0,879 | 0,99 | -0,01 | 9,7E-01 |

| | | | | | | | | | | | | | | | | | | | | | | | |
|---|---|---|---|---|---|---|---|---|---|---|---|---|---|---|---|---|---|---|---|---|---|---|---|
| Q8QZY9 | Splicing factor 3B subunit 4 OS=Mus musculus OX=10090 GN=Sf | 3 | 1 | 1 | 1 | 424 | 44,3 | 8,56 | 601 | | 1 | | 0,909 | 1,06 | 1,311 | 1,209 | 1,243 | 0,985 | 1,182 | 1,097 | 1,00 | 0,00 | 9,7E-01 |
| P30677 | Guanine nucleotide-binding protein subunit alpha-14 OS=Mus m | 9 | 3 | 5 | 1 | 355 | 41,5 | 6,3 | 1833 | 5,3 | 2 | 1 | 1,207 | 1,123 | 2,094 | 1,209 | 1,186 | 1,717 | 1,329 | 1,445 | 0,99 | -0,01 | 9,7E-01 |
| Q9CXI5 | Mesencephalic astrocyte-derived neurotrophic factor OS=Mus n | 32 | 5 | 6 | 5 | 179 | 20,4 | 8,07 | 2659 | 2,91 | 4 | 1 | 1,019 | 1,194 | 1,064 | 1,155 | 1,088 | 0,961 | 1,176 | 1,219 | 1,00 | 0,00 | 9,7E-01 |
| Q8BUK6 | Protein Hook homolog 3 OS=Mus musculus OX=10090 GN=Hook | 6 | 4 | 5 | 4 | 718 | 83,2 | 5,19 | 2601 | | 4 | | 0,945 | 1,082 | 1,065 | 1,021 | 1,125 | 0,8 | 1,059 | 1,115 | 1,00 | 0,00 | 9,7E-01 |
| Q80UG2 | Plexin-A4 OS=Mus musculus OX=10090 GN=Plxna4 PE=1 SV=3 | 15 | 25 | 41 | 17 | 1893 | 212,4 | 6,83 | 13306 | 44,78 | 14 | 13 | 0,986 | 0,971 | 1,031 | 0,953 | 1,026 | 0,978 | 0,946 | 0,995 | 1,00 | 0,00 | 9,7E-01 |
| P56371 | Ras-related protein Rab-4A OS=Mus musculus OX=10090 GN=Ra | 11 | 2 | 2 | 2 | 218 | 24,4 | 6,07 | 1084 | | 2 | | 0,972 | 1,064 | 1,067 | 1,209 | 1,003 | 1,262 | 1,079 | 0,954 | 1,00 | 0,00 | 9,7E-01 |
| P26350 | Prothymosin alpha OS=Mus musculus OX=10090 GN=Ptma PE=1 | 13 | 1 | 1 | 1 | 111 | 12,2 | 3,79 | 656 | | 1 | | 0,866 | 0,903 | 0,733 | 1,186 | 0,925 | 1,153 | 0,771 | 0,86 | 0,99 | -0,01 | 9,7E-01 |
| Q8CDN6 | Thioredoxin-like protein 1 OS=Mus musculus OX=10090 GN=Txn | 18 | 4 | 10 | 4 | 289 | 32,2 | 4,96 | 5025 | 4,75 | 3 | 1 | 1,011 | 1,013 | 0,952 | 0,975 | 0,867 | 0,988 | 1,09 | 0,998 | 1,00 | 0,00 | 9,7E-01 |
| Q9DBR3 | Armadillo repeat-containing protein 8 OS=Mus musculus OX=10 | 2 | 1 | 1 | 1 | 673 | 75,3 | 6,73 | 423 | | 1 | | 0,961 | 0,962 | 1,203 | 0,926 | 0,872 | 1,302 | 0,848 | 1,01 | 1,00 | 0,00 | 9,7E-01 |
| Q8K1R3 | Polyribonucleotide nucleotidyltransferase 1, mitochondrial OS=I | 1 | 1 | 1 | 1 | 783 | 85,6 | 8,03 | 706 | | 1 | | 0,875 | 0,849 | 0,856 | 1,179 | 1,159 | 0,767 | 1,274 | 0,528 | 1,01 | 0,01 | 9,7E-01 |
| P33173 | Kinesin-like protein KIF1A OS=Mus musculus OX=10090 GN=Kif1 | 12 | 18 | 22 | 12 | 1695 | 191,6 | 6,2 | 9954 | 18,71 | 14 | 5 | 0,95 | 0,947 | 1,193 | 0,967 | 1,11 | 0,982 | 0,99 | 0,964 | 1,00 | 0,00 | 9,7E-01 |
| P13595 | Neural cell adhesion molecule 1 OS=Mus musculus OX=10090 G | 44 | 52 | 237 | 13 | 1115 | 119,4 | 4,83 | 87966 | 251,89 | 35 | 29 | 0,983 | 0,954 | 1,066 | 0,939 | 1,039 | 1,016 | 0,959 | 0,934 | 1,00 | 0,00 | 9,7E-01 |
| Q925T6 | Glutamate receptor-interacting protein 1 OS=Mus musculus OX= | 2 | 2 | 2 | 2 | 1127 | 122 | 6,55 | 449 | 3,27 | 1 | 1 | 0,83 | 1,076 | 0,968 | 0,906 | 0,873 | 0,972 | 0,867 | 1,057 | 1,00 | 0,00 | 9,7E-01 |
| Q8VDN2 | Sodium/potassium-transporting ATPase subunit alpha-1 OS=Mu | 39 | 38 | 166 | 22 | 1023 | 112,9 | 5,45 | 77633 | 124,33 | 30 | 16 | 0,995 | 0,955 | 0,987 | 0,915 | 0,974 | 1,008 | 0,945 | 0,929 | 1,00 | 0,00 | 9,7E-01 |
| Q6PFX7 | Neuronal tyrosine-phosphorylated phosphoinositide-3-kinase ac | 2 | 2 | 2 | 2 | 833 | 87,6 | 9,48 | 534 | 0 | 1 | 1 | 0,977 | 0,839 | 0,833 | 0,819 | 0,913 | 0,843 | 0,804 | 0,901 | 1,00 | 0,00 | 9,7E-01 |
| Q9JMD0 | BUB3-interacting and GLEBS motif-containing protein ZNF207 O | 3 | 1 | 1 | 1 | 495 | 52,8 | 9,1 | 587 | | 1 | | 0,714 | 1,241 | 0,952 | 0,937 | 0,937 | 0,999 | 0,94 | 0,951 | 1,00 | 0,00 | 9,7E-01 |
| P14069 | Protein S100-A6 OS=Mus musculus OX=10090 GN=S100a6 PE=1 | 9 | 1 | 1 | 1 | 89 | 10 | 5,48 | | 2,03 | | 1 | 0,99 | 2,919 | 0,706 | 0,697 | 1,099 | 1,234 | 1,614 | 1,28 | 1,02 | 0,02 | 9,7E-01 |
| Q9CYI4 | Putative RNA-binding protein Luc7-like 1 OS=Mus musculus OX= | 5 | 2 | 2 | 2 | 371 | 43,9 | 9,88 | 1378 | | 2 | | 0,977 | 0,841 | 1,12 | 1,078 | 0,939 | 0,888 | 1,073 | 1,129 | 1,00 | 0,00 | 9,7E-01 |
| P09411 | Phosphoglycerate kinase 1 OS=Mus musculus OX=10090 GN=Pg | 55 | 22 | 50 | 22 | 417 | 44,5 | 7,9 | 22039 | 27,91 | 17 | 9 | 1,012 | 1,009 | 1,002 | 1,044 | 0,972 | 1,042 | 1,006 | 1,044 | 1,00 | 0,00 | 9,7E-01 |
| Q8C522 | Endonuclease domain-containing 1 protein OS=Mus musculus O | 10 | 4 | 7 | 4 | 501 | 55,2 | 6,16 | 2726 | 2,58 | 3 | 1 | 1,034 | 1,078 | 0,886 | 0,866 | 1,005 | 1,09 | 0,899 | 0,881 | 1,00 | 0,00 | 9,7E-01 |
| Q8HW98 | IgLON family member 5 OS=Mus musculus OX=10090 GN=Iglon5 | 15 | 3 | 3 | 3 | 336 | 36,7 | 7,69 | 230 | 8,72 | 1 | 2 | 0,782 | 0,856 | 0,854 | 0,698 | 0,818 | 0,719 | 0,728 | 0,916 | 1,00 | 0,00 | 9,7E-01 |
| Q99JF8 | PC4 and SFRS1-interacting protein OS=Mus musculus OX=10090 | 9 | 4 | 7 | 3 | 528 | 59,7 | 9,13 | 3341 | 3,39 | 4 | 1 | 0,882 | 0,957 | 0,954 | 1,042 | 0,922 | 0,818 | 1,131 | 0,975 | 1,00 | 0,00 | 9,7E-01 |
| Q3UR70 | Transforming growth factor-beta receptor-associated protein 1 | 1 | 1 | 1 | 1 | 860 | 97,2 | 6,7 | 663 | | 1 | | 1,127 | 1,075 | 0,947 | 1,139 | 0,825 | 1,233 | 1,041 | 1,174 | 1,00 | 0,00 | 9,7E-01 |
| Q3V3R4 | Integrin alpha-1 OS=Mus musculus OX=10090 GN=Itga1 PE=1 SV | 1 | 1 | 1 | 1 | 1179 | 130,7 | 6,2 | | 2,49 | | 1 | 0,923 | 1,523 | 1,149 | 1,549 | 1,943 | 0,846 | 1,133 | 1,262 | 0,99 | -0,01 | 9,7E-01 |
| P68369 | Tubulin alpha-1A chain OS=Mus musculus OX=10090 GN=Tuba1 | 58 | 31 | 314 | 8 | 451 | 50,1 | 5,06 | 121570 | 393,43 | 28 | 16 | 0,986 | 1,025 | 1,036 | 1,008 | 0,903 | 1,102 | 1,008 | 1,036 | 1,00 | 0,00 | 9,7E-01 |
| P21278 | Guanine nucleotide-binding protein subunit alpha-11 OS=Mus m | 39 | 14 | 20 | 8 | 359 | 42 | 5,97 | 6772 | 30,52 | 9 | 6 | 1,07 | 0,996 | 1,114 | 1,064 | 0,991 | 1,15 | 1,094 | 1,015 | 1,00 | 0,00 | 9,7E-01 |
| Q8CIE6 | Coatomer subunit alpha OS=Mus musculus OX=10090 GN=Copa | 12 | 12 | 14 | 12 | 1224 | 138,3 | 7,65 | 4981 | 11,72 | 8 | 4 | 0,962 | 0,916 | 0,98 | 0,981 | 0,902 | 0,964 | 0,952 | 1,025 | 1,00 | 0,00 | 9,7E-01 |
| A2A690 | Protein TANC2 OS=Mus musculus OX=10090 GN=Tanc2 PE=1 SV | 3 | 4 | 6 | 4 | 1994 | 220,1 | 7,97 | 2156 | 5,59 | 2 | 2 | 0,987 | 0,954 | 0,981 | 0,987 | 1,032 | 0,913 | 1,045 | 0,914 | 1,00 | 0,00 | 9,7E-01 |
| Q9QYR6 | Microtubule-associated protein 1A OS=Mus musculus OX=1009( | 18 | 44 | 58 | 43 | 2776 | 300 | 5 | 19039 | 60,67 | 26 | 21 | 1,071 | 1,057 | 1,006 | 1,058 | 0,877 | 0,997 | 1,213 | 1,115 | 1,00 | 0,00 | 9,7E-01 |
| P17439 | Lysosomal acid glucosylceramidase OS=Mus musculus OX=1009( | 3 | 1 | 1 | 1 | 515 | 57,6 | 7,75 | 540 | | 1 | | 1,054 | 0,946 | 0,998 | 0,984 | 1,077 | 0,718 | 1,042 | 1,158 | 1,00 | 0,00 | 9,7E-01 |
| Q3TBD2 | Rho GTPase-activating protein 45 OS=Mus musculus OX=10090 | 1 | 1 | 1 | 1 | 1116 | 122,8 | 5,94 | 294 | | 1 | | 0,949 | 0,861 | 0,962 | 0,933 | 0,695 | 0,819 | 1,028 | 1,177 | 1,00 | 0,00 | 9,8E-01 |
| Q3UVK0 | Endoplasmic reticulum metallopeptidase 1 OS=Mus musculus O | 3 | 2 | 2 | 2 | 898 | 100,1 | 7,49 | 1295 | | 2 | | 1,047 | 0,914 | 1,133 | 0,93 | 1,008 | 1,243 | 0,968 | 0,818 | 1,00 | 0,00 | 9,8E-01 |
| Q5SUC9 | Protein SCO1 homolog, mitochondrial OS=Mus musculus OX=10 | 8 | 2 | 3 | 2 | 284 | 31,6 | 8,47 | 1515 | | 2 | | 1,092 | 0,834 | 1,095 | 0,84 | 1,041 | 1,01 | 0,893 | 0,927 | 1,00 | 0,00 | 9,8E-01 |
| P17426 | AP-2 complex subunit alpha-1 OS=Mus musculus OX=10090 GN= | 25 | 25 | 40 | 18 | 977 | 107,6 | 7,03 | 16002 | 35,87 | 14 | 12 | 1,04 | 0,964 | 1,043 | 1,029 | 1,056 | 1,031 | 0,979 | 1,007 | 1,00 | 0,00 | 9,8E-01 |
| Q922R8 | Protein disulfide-isomerase A6 OS=Mus musculus OX=10090 GN | 24 | 11 | 17 | 11 | 440 | 48,1 | 5,14 | 6583 | 14,07 | 7 | 5 | 1,049 | 0,92 | 1,006 | 0,981 | 1,001 | 0,893 | 1,011 | 1,046 | 1,00 | 0,00 | 9,8E-01 |
| P53994 | Ras-related protein Rab-2A OS=Mus musculus OX=10090 GN=Ra | 37 | 8 | 12 | 7 | 212 | 23,5 | 6,54 | 5596 | 12,81 | 5 | 3 | 0,911 | 0,825 | 0,884 | 0,898 | 0,915 | 0,903 | 0,841 | 0,862 | 1,00 | 0,00 | 9,8E-01 |
| Q9CXW4 | 60S ribosomal protein L11 OS=Mus musculus OX=10090 GN=Rpl | 17 | 3 | 6 | 3 | 178 | 20,2 | 9,6 | 3724 | 3,01 | 2 | 1 | 1,004 | 1,005 | 0,953 | 1,183 | 1,16 | 0,877 | 1,038 | 1,079 | 1,00 | 0,00 | 9,8E-01 |
| Q9DAK9 | 14 kDa phosphohistidine phosphatase OS=Mus musculus OX=10 | 26 | 3 | 3 | 3 | 124 | 14 | 5,53 | 2036 | | 3 | | 0,81 | 0,716 | 0,917 | 0,925 | 0,811 | 1,038 | 0,689 | 0,84 | 1,00 | 0,00 | 9,8E-01 |
| Q99LI8 | Hepatocyte growth factor-regulated tyrosine kinase substrate O | 9 | 7 | 7 | 7 | 775 | 86 | 6,16 | 3270 | 4,89 | 5 | 2 | 0,968 | 1,051 | 1,02 | 0,946 | 1,052 | 0,97 | 1,031 | 0,936 | 1,00 | 0,00 | 9,8E-01 |
| Q8BY87 | Ubiquitin carboxyl-terminal hydrolase 47 OS=Mus musculus OX= | 4 | 5 | 5 | 5 | 1376 | 157,4 | 5,11 | 1825 | 5,35 | 3 | 2 | 1,175 | 1,066 | 1,046 | 1,085 | 1,271 | 1,12 | 1,024 | 0,949 | 1,00 | 0,00 | 9,8E-01 |
| Q9DBL1 | Short/branched chain specific acyl-CoA dehydrogenase, mitocho | 13 | 5 | 5 | 4 | 432 | 47,8 | 7,87 | 1247 | 4,57 | 2 | 3 | 1,156 | 1,02 | 1,013 | 1,087 | 1,191 | 1,137 | 0,958 | 0,983 | 1,00 | 0,00 | 9,8E-01 |
| Q7TPM6 | Fibronectin type III and SPRY domain-containing protein 1 OS=M | 15 | 6 | 7 | 6 | 496 | 55,5 | 6,67 | 3466 | 0 | 5 | 1 | 1,032 | 0,962 | 0,995 | 0,961 | 1,029 | 1,058 | 0,958 | 0,901 | 1,00 | 0,00 | 9,8E-01 |
| Q91ZZ3 | Beta-synuclein OS=Mus musculus OX=10090 GN=Sncb PE=1 SV= | 53 | 7 | 17 | 4 | 133 | 14 | 4,37 | 9046 | 12,13 | 5 | 2 | 0,977 | 1,034 | 1,05 | 1,046 | 0,923 | 1,504 | 0,861 | 0,835 | 1,00 | 0,00 | 9,8E-01 |
| Q6VNB8 | WD repeat and FYVE domain-containing protein 3 OS=Mus mus | 1 | 4 | 5 | 4 | 3508 | 392,1 | 6,8 | 3036 | | 4 | | 1,297 | 1,093 | 1,121 | 1,055 | 1,228 | 1,127 | 1,103 | 1,114 | 1,00 | 0,00 | 9,8E-01 |
| Q61595 | Kinectin OS=Mus musculus OX=10090 GN=Ktn1 PE=1 SV=1 | 6 | 7 | 8 | 7 | 1327 | 152,5 | 5,86 | 2753 | 10,3 | 5 | 2 | 0,997 | 1,009 | 1,062 | 1,169 | 0,99 | 1,026 | 1,076 | 1,14 | 1,00 | 0,00 | 9,8E-01 |
| Q91Z96 | BMP-2-inducible protein kinase OS=Mus musculus OX=10090 GI | 1 | 1 | 2 | 1 | 1138 | 126,1 | 6,8 | 827 | | 1 | | 0,952 | 0,879 | 0,954 | 1,063 | 0,932 | 1,021 | 0,875 | 1,015 | 1,00 | 0,00 | 9,8E-01 |
| P61979-2 | Isoform 2 of Heterogeneous nuclear ribonucleoprotein K OS=Mu | 25 | 12 | 22 | 12 | 464 | 51 | 5,33 | 12357 | 5,18 | 10 | 2 | 0,89 | 1,037 | 1,149 | 1,144 | 1,118 | 1,012 | 1,122 | 0,961 | 1,00 | 0,00 | 9,8E-01 |
| O08992 | Syntenin-1 OS=Mus musculus OX=10090 GN=Sdcbp PE=1 SV=1 | 4 | 1 | 1 | 1 | 299 | 32,4 | 7,15 | 758 | | 1 | | 1,003 | 1,105 | 0,845 | 0,9 | 1,158 | 0,791 | 0,837 | 1,077 | 1,00 | 0,00 | 9,8E-01 |
| Q8VE70 | Programmed cell death protein 10 OS=Mus musculus OX=10090 | 10 | 2 | 3 | 2 | 212 | 24,7 | 8,19 | 1560 | 2,34 | 1 | 1 | 1,041 | 1,022 | 0,916 | 1,114 | 0,964 | 1,048 | 1,04 | 1,037 | 1,00 | 0,00 | 9,8E-01 |
| P20065 | Thymosin beta-4 OS=Mus musculus OX=10090 GN=Tmsb4x PE= | 40 | 5 | 8 | 3 | 50 | 5,7 | 5,06 | | 24,44 | | 5 | 0,92 | 0,669 | 1,398 | 0,73 | 0,694 | 1,525 | 0,787 | 0,733 | 0,99 | -0,01 | 9,8E-01 |
| Q80XU3 | Nuclear ubiquitous casein and cyclin-dependent kinase substrat | 4 | 1 | 1 | 1 | 234 | 26,3 | 5,14 | 842 | | 1 | | 1,058 | 1,054 | 0,891 | 1,151 | 0,776 | 1,028 | 1,15 | 1,209 | 1,00 | 0,00 | 9,8E-01 |
| Q99N93 | 39S ribosomal protein L16, mitochondrial OS=Mus musculus OX | 8 | 2 | 3 | 2 | 251 | 28,8 | 10,27 | 1513 | | 2 | | 1,015 | 0,87 | 0,922 | 1,084 | 0,769 | 1,019 | 1,011 | 1,099 | 1,00 | 0,00 | 9,8E-01 |
| Q7TMM9 | Tubulin beta-2A chain OS=Mus musculus OX=10090 GN=Tubb2a | 69 | 28 | 537 | 2 | 445 | 49,9 | 4,89 | 225036 | 418,83 | 23 | 21 | 0,983 | 0,928 | 1,11 | 0,978 | 1,003 | 1,021 | 0,99 | 0,988 | 1,00 | 0,00 | 9,9E-01 |
| P43406 | Integrin alpha-V OS=Mus musculus OX=10090 GN=Itgav PE=1 SV | 7 | 6 | 6 | 6 | 1044 | 115,3 | 5,63 | 2246 | 3,83 | 4 | 2 | 1,071 | 0,932 | 0,976 | 1,01 | 0,93 | 0,994 | 1,065 | 1,003 | 1,00 | 0,00 | 9,9E-01 |
| Q9DBS8 | Centrosomal protein POC5 OS=Mus musculus OX=10090 GN=Po | 1 | 1 | 2 | 1 | 558 | 60,9 | 8,09 | 908 | | 1 | | 0,916 | 0,844 | 1,142 | 0,878 | 1,111 | 0,809 | 0,983 | 0,87 | 1,00 | 0,00 | 9,9E-01 |
| Q6GQS1 | Calcium-binding mitochondrial carrier protein SCaMC-3 OS=Mus | 1 | 1 | 1 | 1 | 467 | 52,5 | 7,59 | | 0 | | 1 | 0,912 | 0,771 | 0,863 | 0,902 | 0,704 | 0,763 | 1,004 | 0,983 | 1,00 | 0,00 | 9,9E-01 |
| P97797 | Tyrosine-protein phosphatase non-receptor type substrate 1 OS | 12 | 4 | 6 | 4 | 513 | 56,4 | 8,28 | 3127 | 3,14 | 4 | 1 | 0,873 | 1,006 | 0,903 | 1,212 | 1,093 | 1,004 | 0,97 | 0,921 | 1,00 | 0,00 | 9,9E-01 |
| Q00558 | Factor VIII intron 22 protein OS=Mus musculus OX=10090 GN=F | 7 | 2 | 2 | 2 | 380 | 40,5 | 6,89 | 1071 | | 2 | | 0,945 | 0,896 | 0,981 | 0,844 | 0,791 | 1,037 | 0,817 | 1,016 | 1,00 | 0,00 | 9,9E-01 |
| Q8BGA9 | Mitochondrial inner membrane protein OXA1L OS=Mus musculu | 2 | 1 | 1 | 1 | 433 | 48,2 | 9,61 | 479 | | 1 | | 1,031 | 1,391 | 1,096 | 1,201 | 1,236 | 0,875 | 1,384 | 1,233 | 1,00 | 0,00 | 9,9E-01 |
| Q8BL65 | Actin-binding LIM protein 2 OS=Mus musculus OX=10090 GN=Al | 3 | 2 | 3 | 1 | 612 | 68,1 | 8,02 | 561 | 5,12 | 1 | 1 | 0,926 | 0,803 | 1,065 | 0,909 | 0,778 | 0,948 | 1,016 | 0,966 | 1,00 | 0,00 | 9,9E-01 |
| Q9WVQ1 | Membrane-associated guanylate kinase, WW and PDZ domain-c | 3 | 3 | 3 | 3 | 1275 | 140,8 | 6,19 | | 8,62 | | 3 | 0,871 | 0,958 | 1,019 | 1,118 | 1,147 | 0,929 | 0,891 | 0,994 | 1,00 | 0,00 | 9,9E-01 |
| P62631 | Elongation factor 1-alpha 2 OS=Mus musculus OX=10090 GN=Ee | 22 | 10 | 31 | 4 | 463 | 50,4 | 9,03 | 16929 | 14,61 | 8 | 3 | 0,927 | 0,994 | 0,939 | 0,912 | 0,838 | 1,048 | 0,967 | 0,916 | 1,00 | 0,00 | 9,9E-01 |
| P26450 | Phosphatidylinositol 3-kinase regulatory subunit alpha OS=Mus | 2 | 1 | 2 | 1 | 724 | 83,5 | 6,28 | | 2,57 | | 1 | 1,05 | 1,254 | 1,262 | 0,975 | 1,039 | 1,198 | 1,168 | 1,131 | 1,00 | 0,00 | 9,9E-01 |
| Q8VBZ3 | Cleft lip and palate transmembrane protein 1 homolog OS=Mus | 6 | 2 | 2 | 2 | 664 | 75,2 | 6,3 | 219 | 2,53 | 1 | 1 | 0,765 | 0,753 | 1,031 | 0,771 | 0,847 | 0,632 | 0,844 | 0,991 | 1,00 | 0,00 | 9,9E-01 |
| Q8C1A5 | Thimet oligopeptidase OS=Mus musculus OX=10090 GN=Thop1 | 11 | 6 | 7 | 6 | 687 | 78 | 6,06 | 1860 | 9,18 | 3 | 3 | 1,023 | 1,06 | 0,977 | 1,046 | 1,002 | 0,993 | 0,999 | 1,114 | 1,00 | 0,00 | 9,9E-01 |

| | | | | | | | | | | | | | | | | | | | | | | | |
|---|---|---|---|---|---|---|---|---|---|---|---|---|---|---|---|---|---|---|---|---|---|---|---|
| P17897 | Lysozyme C-1 OS=Mus musculus OX=10090 GN=Lyz1 PE=1 SV=1 | 19 | 4 | 8 | 4 | 148 | 16,8 | 9,41 | 3524 | 0 | 4 | 1 | 1,47 | 1,074 | 1,033 | 0,839 | 1,524 | 0,971 | 0,988 | 0,944 | 1,00 | 0,00 | 9,9E-01 |
| Q9D7P6 | Iron-sulfur cluster assembly enzyme ISCU, mitochondrial OS=Mu | 16 | 2 | 2 | 2 | 168 | 18,1 | 9,29 | 696 | 3 | 1 | 1 | 1,025 | 1,113 | 0,943 | 0,971 | 0,903 | 1,027 | 1,035 | 1,09 | 1,00 | 0,00 | 9,9E-01 |
| P47746 | Cannabinoid receptor 1 OS=Mus musculus OX=10090 GN=Cnr1 I | 2 | 1 | 1 | 1 | 473 | 52,8 | 8,27 | 440 | | 1 | | 0,99 | 1,144 | 1,144 | 0,691 | 1,118 | 1,047 | 0,762 | 1,049 | 1,00 | 0,00 | 9,9E-01 |
| O55131 | Septin-7 OS=Mus musculus OX=10090 GN=Septin7 PE=1 SV=1 | 38 | 23 | 38 | 23 | 436 | 50,5 | 8,57 | 12343 | 51,59 | 11 | 15 | 1,002 | 1,009 | 1,056 | 0,969 | 1 | 1,013 | 0,992 | 1,032 | 1,00 | 0,00 | 9,9E-01 |
| Q69ZS7 | HBS1-like protein OS=Mus musculus OX=10090 GN=Hbs1l PE=1 | 7 | 3 | 3 | 3 | 682 | 75,1 | 6,46 | 1263 | 2,83 | 2 | 1 | 0,791 | 0,927 | 0,977 | 0,865 | 0,746 | 0,806 | 1,054 | 0,958 | 1,00 | 0,00 | 9,9E-01 |
| O09053 | Werner syndrome ATP-dependent helicase homolog OS=Mus m | 1 | 1 | 1 | 1 | 1401 | 157,1 | 6,46 | 224 | | 1 | | 0,869 | 0,886 | 0,916 | 0,944 | 0,845 | 1,095 | 0,844 | 0,828 | 1,00 | 0,00 | 9,9E-01 |
| Q8CFH6 | Serine/threonine-protein kinase SIK2 OS=Mus musculus OX=100 | 1 | 1 | 1 | 1 | 931 | 104,1 | 6,13 | 205 | | 1 | | 1,305 | 1,039 | 1,435 | 1,263 | 1,025 | 1,527 | 1,206 | 1,278 | 1,00 | 0,00 | 9,9E-01 |
| Q9WTM5 | RuvB-like 2 OS=Mus musculus OX=10090 GN=Ruvbl2 PE=1 SV=3 | 24 | 11 | 12 | 11 | 463 | 51,1 | 5,64 | 4790 | 7,49 | 8 | 3 | 1,022 | 1,024 | 1,017 | 1,055 | 1,085 | 1,026 | 1,019 | 0,987 | 1,00 | 0,00 | 9,9E-01 |
| Q9CW03 | Structural maintenance of chromosomes protein 3 OS=Mus mus | 1 | 1 | 1 | 1 | 1217 | 141,5 | 7,18 | | 0 | | 1 | 0,545 | 0,668 | 0,792 | 0,885 | 0,728 | 0,802 | 0,795 | 0,569 | 1,00 | 0,00 | 9,9E-01 |
| Q8VBT2 | L-serine dehydratase/L-threonine deaminase OS=Mus musculus | 6 | 1 | 1 | 1 | 327 | 34,6 | 7,12 | 503 | | 1 | | 0,854 | 0,91 | 0,931 | 0,916 | 0,922 | 1,007 | 0,98 | 0,705 | 1,00 | 0,00 | 9,9E-01 |
| P63250 | G protein-activated inward rectifier potassium channel 1 OS=Mu | 2 | 1 | 1 | 1 | 501 | 56,5 | 8,37 | 396 | | 1 | | 0,735 | 0,744 | 0,848 | 0,856 | 1,061 | 0,773 | 0,681 | 0,664 | 1,00 | 0,00 | 9,9E-01 |
| Q8K0Z7 | Translational activator of cytochrome c oxidase 1 OS=Mus musc | 17 | 4 | 4 | 4 | 294 | 32,3 | 8,12 | 1002 | 5,48 | 2 | 2 | 1,047 | 1,23 | 1,189 | 1,4 | 1,174 | 1,298 | 1,222 | 1,169 | 1,00 | 0,00 | 9,9E-01 |
| Q8C5Q4 | G-rich sequence factor 1 OS=Mus musculus OX=10090 GN=Grsf1 | 3 | 1 | 1 | 1 | 479 | 53 | 6,67 | 172 | | 1 | | 0,941 | 0,893 | 0,868 | 1,102 | 0,753 | 1,093 | 0,819 | 1,135 | 1,00 | 0,00 | 9,9E-01 |
| Q8BPN8 | DmX-like protein 2 OS=Mus musculus OX=10090 GN=Dmxl2 PE= | 11 | 28 | 36 | 28 | 3032 | 338 | 6,42 | 16312 | 14,58 | 22 | 6 | 0,969 | 1,018 | 1,002 | 0,946 | 0,952 | 1,016 | 0,938 | 1,028 | 1,00 | 0,00 | 9,9E-01 |
| Q9DC71 | 28S ribosomal protein S15, mitochondrial OS=Mus musculus OX | 8 | 2 | 2 | 2 | 258 | 29,4 | 10,13 | 747 | | 2 | | 0,853 | 1,424 | 0,782 | 0,951 | 0,932 | 0,795 | 1,207 | 1,082 | 1,00 | 0,00 | 9,9E-01 |
| P63248 | cAMP-dependent protein kinase inhibitor alpha OS=Mus muscul | 70 | 3 | 9 | 3 | 76 | 8 | 4,54 | 1074 | 10,04 | 2 | 1 | 1,074 | 1,334 | 0,975 | 1,327 | 1,438 | 1,222 | 0,906 | 1,139 | 1,00 | 0,00 | 9,9E-01 |
| Q6PD28 | Serine/threonine-protein phosphatase 2A 56 kDa regulatory sub | 13 | 5 | 6 | 3 | 497 | 57,3 | 6,84 | 1637 | 7,85 | 3 | 2 | 1,195 | 1,262 | 1,011 | 1,002 | 1,511 | 0,878 | 0,978 | 1,108 | 1,00 | 0,00 | 9,9E-01 |
| P15327 | Bisphosphoglycerate mutase OS=Mus musculus OX=10090 GN=I | 7 | 1 | 2 | 1 | 259 | 30 | 7,06 | 725 | | 1 | | 1,919 | 1,452 | 1,477 | 1,431 | 1,164 | 1,679 | 1,465 | 1,977 | 1,00 | 0,00 | 9,9E-01 |
| Q91VE0 | Long-chain fatty acid transport protein 4 OS=Mus musculus OX= | 11 | 6 | 9 | 6 | 643 | 72,3 | 8,59 | 3969 | 4,53 | 4 | 2 | 1,081 | 1,045 | 1,124 | 1,197 | 1,284 | 0,961 | 1,114 | 1,09 | 1,00 | 0,00 | 9,9E-01 |
| P99026 | Proteasome subunit beta type-4 OS=Mus musculus OX=10090 G | 18 | 5 | 10 | 5 | 264 | 29,1 | 5,64 | 5062 | 5,99 | 5 | 1 | 1,024 | 1,006 | 1,118 | 1,006 | 1,05 | 1,107 | 0,954 | 1,044 | 1,00 | 0,00 | 1,0E+00 |
| Q80ZS3 | 28S ribosomal protein S26, mitochondrial OS=Mus musculus OX | 11 | 2 | 3 | 2 | 200 | 23,4 | 9,96 | 1919 | | 2 | | 1,083 | 1,055 | 0,996 | 1,09 | 0,978 | 0,996 | 1,135 | 1,114 | 1,00 | 0,00 | 1,0E+00 |
| P47856 | Glutamine--fructose-6-phosphate aminotransferase [isomerizing | 5 | 3 | 4 | 3 | 697 | 78,5 | 6,84 | 2044 | | 3 | | 1,221 | 0,881 | 1,179 | 0,911 | 1,033 | 1,108 | 1,072 | 0,981 | 1,00 | 0,00 | 1,0E+00 |
| Q5RJI5 | Serine/threonine-protein kinase BRSK1 OS=Mus musculus OX=1 | 9 | 7 | 7 | 5 | 778 | 85,1 | 9,32 | 1802 | 7,11 | 3 | 4 | 1,05 | 0,919 | 1,093 | 0,92 | 0,997 | 1,038 | 0,992 | 0,954 | 1,00 | 0,00 | 1,0E+00 |
| O35449 | Proline-rich transmembrane protein 1 OS=Mus musculus OX=10 | 5 | 1 | 1 | 1 | 306 | 31,4 | 7,65 | 721 | | 1 | | 0,928 | 0,809 | 1,116 | 1,303 | 1,15 | 0,909 | 1,139 | 0,956 | 1,00 | 0,00 | 1,0E+00 |
| P21619 | Lamin-B2 OS=Mus musculus OX=10090 GN=Lmnb2 PE=1 SV=2 | 5 | 3 | 3 | 3 | 596 | 67,3 | 5,5 | 973 | 2,18 | 2 | 1 | 1,027 | 1,202 | 1,198 | 1,009 | 1,098 | 1,018 | 1,155 | 1,166 | 1,00 | 0,00 | 1,0E+00 |
| Q9WTT4 | V-type proton ATPase subunit G 2 OS=Mus musculus OX=10090 | 43 | 4 | 7 | 4 | 118 | 13,6 | 10,26 | 2975 | 11,61 | 3 | 2 | 0,954 | 1,105 | 1,01 | 0,939 | 0,809 | 1,038 | 1,104 | 1,058 | 1,00 | 0,00 | 1,0E+00 |
| Q8K1Z0 | Ubiquinone biosynthesis protein COQ9, mitochondrial OS=Mus | 6 | 2 | 2 | 2 | 313 | 35,1 | 5,92 | 1397 | | 2 | | 0,931 | 1,218 | 1,021 | 1,178 | 1,23 | 0,966 | 1,026 | 1,125 | 1,00 | 0,00 | 1,0E+00 |
| P49070 | Calcium signal-modulating cyclophilin ligand OS=Mus musculus | 7 | 1 | 1 | 1 | 294 | 32,5 | 7,74 | | 3,69 | | 1 | 0,721 | 0,875 | 0,815 | 1,076 | 1,164 | 0,502 | 1,146 | 0,677 | 1,00 | 0,00 | 1,0E+00 |
| Q60972 | Histone-binding protein RBBP4 OS=Mus musculus OX=10090 GN | 4 | 1 | 1 | 1 | 425 | 47,6 | 4,89 | 220 | | 1 | | 1,122 | 1,157 | 1,552 | 1,177 | 1,513 | 1,028 | 1,011 | 1,455 | 1,00 | 0,00 | 1,0E+00 |
| Q8K2B3 | Succinate dehydrogenase [ubiquinone] flavoprotein subunit, mi | 35 | 21 | 40 | 21 | 664 | 72,5 | 7,37 | 12559 | 44,47 | 10 | 13 | 0,994 | 0,953 | 1,063 | 0,964 | 0,969 | 1,089 | 0,902 | 1,014 | 1,00 | 0,00 | 1,0E+00 |
| P28660 | Nck-associated protein 1 OS=Mus musculus OX=10090 GN=Ncka | 18 | 24 | 39 | 24 | 1128 | 128,7 | 6,62 | 14795 | 21,84 | 19 | 8 | 1,021 | 1,009 | 0,908 | 0,967 | 0,929 | 0,909 | 1,066 | 1,001 | 1,00 | 0,00 | 1,0E+00 |
| Q7TPR4 | Alpha-actinin-1 OS=Mus musculus OX=10090 GN=Actn1 PE=1 SV | 24 | 19 | 22 | 11 | 892 | 103 | 5,38 | 10362 | 14,67 | 15 | 5 | 1,017 | 1,005 | 0,953 | 1,046 | 0,96 | 1,043 | 0,986 | 1,032 | 1,00 | 0,00 | 1,0E+00 |
| Q9CZN7 | Serine hydroxymethyltransferase, mitochondrial OS=Mus muscu | 19 | 8 | 12 | 8 | 504 | 55,7 | 8,47 | 3682 | 11,07 | 4 | 4 | 1,018 | 1,074 | 1,086 | 1,08 | 1,146 | 1,054 | 1,053 | 1,005 | 1,00 | 0,00 | 1,0E+00 |
| Q9EQZ6 | Rap guanine nucleotide exchange factor 4 OS=Mus musculus OX | 6 | 6 | 7 | 6 | 1011 | 115,4 | 6,92 | 4302 | | 6 | | 0,933 | 0,998 | 0,958 | 0,94 | 1,017 | 0,885 | 0,916 | 1,011 | 1,00 | 0,00 | 1,0E+00 |
