## Supplementary material for "SorCS2 modulates neurovascular coupling via glutamatergic and calcium signaling in astrocytes": Expanded View Table 2

| | | | | | | | | | Sample name: | | SorCS2 knockout mouse #1 | SorCS2 knockout mouse #2 | SorCS2 knockout mouse #3 | SorCS2 knockout mouse #4 | Wild type mouse #1 | Wild type mouse #2 | Wild type mouse #3 | Wild type mouse #4 | | | |
|---|---|---|---|---|---|---|---|---|---|---|---|---|---|---|---|---|---|---|---|---|---|
| Accession | Description | Coverage [%] | # Peptides | # PSMs | # Unique Peptides | # AAs | MW [kDa] | calc. pI | Score Sequest HT: Sequest HT | # Peptides (by Search Engine): Sequest HT | Abundance Ratio: (F18, 129N) / (F18, 126) | Abundance Ratio: (F18, 129C) / (F18, 126) | Abundance Ratio: (F18, 130N) / (F18, 126) | Abundance Ratio: (F18, 131) / (F18, 126) | Abundance Ratio: (F18, 127N) / (F18, 126) | Abundance Ratio: (F18, 127C) / (F18, 126) | Abundance Ratio: (F18, 128N) / (F18, 126) | Abundance Ratio: (F18, 128C) / (F18, 126) | Ratio SorCS2 knockout / wild type | Fold change in SorCS2 knockout vs. wild type | *t*-test, *P*-value |
| Q9QYS2 | Metabotropic glutamate receptor 3 OS=Mus musculus OX=10090 G | 19 | 15 | 25 | 14 | 879 | 99,1 | 7,75 | 48,07 | 15 | 1,111 | 1,189 | 1,23 | 1,2 | 0,805 | 0,837 | 0,789 | 0,854 | 1,44 | 0,44 | 1,8E-05 |
| Q6PDC0 | RUN domain-containing protein 3B OS=Mus musculus OX=10090 G | 8 | 2 | 2 | 2 | 408 | 45,1 | 5,17 | 2,85 | 2 | 1,301 | 1,352 | 1,271 | 1,303 | 0,843 | 0,894 | 0,735 | 0,888 | 1,56 | 0,56 | 2,5E-05 |
| Q8CHT0 | Delta-1-pyrroline-5-carboxylate dehydrogenase, mitochondrial OS= | 13 | 7 | 9 | 7 | 562 | 61,8 | 8,24 | 21,47 | 7 | 0,972 | 0,948 | 0,973 | 0,974 | 1,071 | 1,076 | 1,098 | 1,114 | 0,89 | -0,11 | 4,5E-05 |
| Q6P069 | Sorcin OS=Mus musculus OX=10090 GN=Sri PE=1 SV=1 | 20 | 4 | 6 | 4 | 198 | 21,6 | 5,59 | 6,83 | 4 | 1,021 | 1,111 | 1,203 | 1,236 | 0,617 | 0,559 | 0,677 | 0,562 | 1,89 | 0,89 | 7,1E-05 |
| Q59J78 | NADH dehydrogenase [ubiquinone] 1 alpha subcomplex assembly f | 16 | 2 | 3 | 2 | 168 | 19,6 | 8,25 | 5,35 | 2 | 1,183 | 1,203 | 1,163 | 1,141 | 0,986 | 1,007 | 0,942 | 0,941 | 1,21 | 0,21 | 7,2E-05 |
| Q9R257 | Heme-binding protein 1 OS=Mus musculus OX=10090 GN=Hebp1 P | 32 | 5 | 7 | 5 | 190 | 21,1 | 5,26 | 20,61 | 5 | 1,318 | 1,527 | 1,434 | 1,61 | 0,829 | 0,869 | 0,781 | 0,964 | 1,71 | 0,71 | 1,7E-04 |
| Q3TC33 | Coiled-coil domain-containing protein 127 OS=Mus musculus OX=1 | 7 | 2 | 2 | 2 | 260 | 30,5 | 9,13 | 4,8 | 2 | 1,072 | 1,063 | 1,03 | 1,089 | 0,964 | 0,935 | 0,966 | 0,947 | 1,12 | 0,12 | 2,6E-04 |
| P08556 | GTPase NRas OS=Mus musculus OX=10090 GN=Nras PE=1 SV=1 | 44 | 7 | 12 | 1 | 189 | 21,2 | 5,17 | 26,32 | 7 | 0,825 | 0,776 | 0,761 | 0,81 | 0,956 | 1,025 | 0,942 | 0,947 | 0,82 | -0,18 | 3,8E-04 |
| Q8K406 | Leucine-rich repeat LGI family member 3 OS=Mus musculus OX=10 | 11 | 6 | 8 | 6 | 548 | 61,8 | 8,16 | 13,6 | 6 | 1,037 | 1,103 | 1,123 | 1,128 | 0,907 | 0,913 | 0,788 | 0,773 | 1,30 | 0,30 | 1,1E-03 |
| Q8BVI4 | Dihydropteridine reductase OS=Mus musculus OX=10090 GN=Qdpr | 31 | 5 | 8 | 5 | 241 | 25,6 | 7,81 | 23,6 | 5 | 1,101 | 1,123 | 1,097 | 1,075 | 0,988 | 1,031 | 1,031 | 0,997 | 1,09 | 0,09 | 1,1E-03 |
| Q9DB34 | Charged multivesicular body protein 2a OS=Mus musculus OX=100 | 5 | 1 | 1 | 1 | 222 | 25,1 | 5,97 | 0 | 1 | 1,23 | 1,177 | 1,349 | 1,268 | 0,906 | 1,067 | 0,919 | 0,886 | 1,33 | 0,33 | 1,3E-03 |
| Q3V384 | AFG1-like ATPase OS=Mus musculus OX=10090 GN=Afg1l PE=1 SV= | 6 | 2 | 2 | 2 | 480 | 54,3 | 8,03 | 0 | 2 | 0,696 | 0,695 | 0,763 | 0,722 | 1,123 | 1,067 | 1,212 | 1,463 | 0,59 | -0,41 | 1,4E-03 |
| Q6V4S5 | Protein sidekick-2 OS=Mus musculus OX=10090 GN=Sdk2 PE=1 SV= | 2 | 4 | 4 | 4 | 2176 | 239,8 | 7,36 | 10,81 | 4 | 1,089 | 1,188 | 1,149 | 1,068 | 0,963 | 0,926 | 0,973 | 0,885 | 1,20 | 0,20 | 1,5E-03 |
| P36536 | GTP-binding protein SAR1a OS=Mus musculus OX=10090 GN=Sar1a | 10 | 2 | 2 | 2 | 198 | 22,4 | 6,93 | 2,27 | 2 | 0,727 | 0,794 | 0,675 | 0,799 | 1,086 | 0,905 | 1,001 | 0,989 | 0,75 | -0,25 | 2,0E-03 |
| Q9CTY5 | Calcium uptake protein 3, mitochondrial OS=Mus musculus OX=100 | 15 | 7 | 7 | 7 | 523 | 59,8 | 7,15 | 13,21 | 7 | 0,935 | 0,933 | 0,958 | 0,939 | 0,992 | 1,023 | 1,009 | 1,059 | 0,92 | -0,08 | 2,1E-03 |
| Q811D0 | Disks large homolog 1 OS=Mus musculus OX=10090 GN=Dlg1 PE=1 | 21 | 21 | 27 | 15 | 905 | 100,1 | 5,8 | 53,99 | 21 | 1,076 | 1,064 | 1,049 | 1,078 | 0,947 | 0,977 | 1,014 | 1 | 1,08 | 0,08 | 2,2E-03 |
| Q3TMH2 | Secernin-3 OS=Mus musculus OX=10090 GN=Scrn3 PE=1 SV=1 | 9 | 3 | 3 | 3 | 418 | 47,6 | 5,66 | 5,7 | 3 | 1,032 | 1,014 | 1,047 | 1,014 | 0,874 | 0,937 | 0,966 | 0,911 | 1,11 | 0,11 | 2,5E-03 |
| Q9R1V6 | Disintegrin and metalloproteinase domain-containing protein 22 O | 19 | 15 | 22 | 15 | 904 | 99,7 | 7,83 | 42,67 | 15 | 1,203 | 1,109 | 1,219 | 1,044 | 0,945 | 0,955 | 0,87 | 0,904 | 1,25 | 0,25 | 2,6E-03 |
| P23819-4 | Isoform 4 of Glutamate receptor 2 OS=Mus musculus OX=10090 GN | 19 | 20 | 29 | 16 | 921 | 102,8 | 7,72 | 73,75 | 20 | 0,922 | 0,901 | 0,893 | 0,979 | 1,053 | 1,024 | 1,015 | 1,021 | 0,90 | -0,10 | 2,6E-03 |
| P46096 | Synaptotagmin-1 OS=Mus musculus OX=10090 GN=Syt1 PE=1 SV=1 | 31 | 19 | 43 | 13 | 421 | 47,4 | 8,53 | 88,93 | 19 | 0,804 | 0,858 | 0,925 | 0,898 | 0,986 | 1,004 | 1,012 | 1,03 | 0,86 | -0,14 | 2,7E-03 |
| P62761 | Visinin-like protein 1 OS=Mus musculus OX=10090 GN=Vsnl1 PE=1 | 50 | 11 | 28 | 9 | 191 | 22,1 | 5,15 | 54,63 | 11 | 1,075 | 1,206 | 1,101 | 1,1 | 0,925 | 0,98 | 0,925 | 0,994 | 1,17 | 0,17 | 3,0E-03 |
| Q6ZWR6 | Nesprin-1 OS=Mus musculus OX=10090 GN=Syne1 PE=1 SV=2 | 4 | 35 | 39 | 35 | 8799 | 1009,3 | 5,59 | 53,77 | 35 | 0,79 | 0,918 | 0,874 | 0,887 | 0,976 | 1,091 | 1,038 | 1,068 | 0,83 | -0,17 | 3,1E-03 |
| Q921G8 | Gamma-tubulin complex component 2 OS=Mus musculus OX=1009 | 1 | 1 | 1 | 1 | 905 | 103,2 | 6,77 | 0 | 1 | 0,689 | 0,907 | 0,795 | 0,675 | 0,975 | 1,041 | 1,026 | 1,033 | 0,75 | -0,25 | 4,1E-03 |
| Q61941 | NAD(P) transhydrogenase, mitochondrial OS=Mus musculus OX=10 | 10 | 10 | 11 | 10 | 1086 | 113,8 | 7,64 | 21,43 | 10 | 1,818 | 1,183 | 1,242 | 1,875 | 0,555 | 0,724 | 0,631 | 0,78 | 2,27 | 1,27 | 4,1E-03 |
| Q0V8T9 | Contactin-associated protein like 5-1 OS=Mus musculus OX=10090 | 2 | 2 | 2 | 2 | 1304 | 145,6 | 6,28 | 2,93 | 2 | 0,658 | 0,595 | 0,644 | 0,689 | 0,943 | 0,849 | 0,989 | 0,764 | 0,73 | -0,27 | 4,3E-03 |
| Q8CHC4 | Synaptojanin-1 OS=Mus musculus OX=10090 GN=Synj1 PE=1 SV=3 | 21 | 28 | 40 | 28 | 1574 | 172,5 | 6,89 | 100,78 | 28 | 0,909 | 0,937 | 0,951 | 0,947 | 0,978 | 0,987 | 1,025 | 1,004 | 0,94 | -0,06 | 4,3E-03 |
| P54763 | Ephrin type-B receptor 2 OS=Mus musculus OX=10090 GN=Ephb2 P | 4 | 4 | 6 | 4 | 986 | 109,8 | 5,71 | 14,89 | 4 | 0,983 | 0,988 | 0,96 | 0,885 | 1,095 | 1,147 | 1,053 | 1,184 | 0,85 | -0,15 | 4,4E-03 |
| Q9CQZ1 | Heat shock factor-binding protein 1 OS=Mus musculus OX=10090 G | 57 | 3 | 6 | 3 | 76 | 8,6 | 4,31 | 8,67 | 3 | 1,054 | 1,153 | 1,149 | 1,135 | 0,939 | 0,937 | 1,048 | 0,94 | 1,16 | 0,16 | 4,7E-03 |
| Q61017 | Guanine nucleotide-binding protein G(I)/G(S)/G(O) subunit gamma | 14 | 1 | 1 | 1 | 69 | 7,8 | 5,52 | 0 | 1 | 1,041 | 1,132 | 1,227 | 1,258 | 0,986 | 0,905 | 0,795 | 0,782 | 1,34 | 0,34 | 5,0E-03 |
| Q7TME0 | Phospholipid phosphatase-related protein type 4 OS=Mus musculu | 12 | 9 | 10 | 9 | 766 | 83,2 | 8,84 | 17 | 9 | 0,888 | 0,843 | 0,938 | 0,961 | 1,102 | 1,018 | 1,134 | 1,032 | 0,85 | -0,15 | 5,2E-03 |
| Q8R0S2 | IQ motif and SEC7 domain-containing protein 1 OS=Mus musculus | 18 | 16 | 24 | 15 | 961 | 107,9 | 7,14 | 44,75 | 16 | 0,874 | 0,904 | 0,901 | 0,941 | 1,016 | 0,961 | 0,999 | 0,968 | 0,92 | -0,08 | 5,2E-03 |
| Q5DTL9 | Sodium-driven chloride bicarbonate exchanger OS=Mus musculus | 10 | 10 | 14 | 6 | 1118 | 125,7 | 6,51 | 24,55 | 10 | 1,205 | 1,299 | 1,217 | 1,16 | 1,012 | 1,122 | 1,061 | 1,057 | 1,15 | 0,15 | 5,2E-03 |
| Q9D823 | 60S ribosomal protein L37 OS=Mus musculus OX=10090 GN=Rpl37 | 13 | 2 | 2 | 2 | 97 | 11,1 | 11,74 | 3,99 | 2 | 0,877 | 0,858 | 1,029 | 0,852 | 1,097 | 1,057 | 1,083 | 1,124 | 0,83 | -0,17 | 5,6E-03 |
| Q8VCD6 | Receptor expression-enhancing protein 2 OS=Mus musculus OX=10 | 9 | 2 | 2 | 2 | 254 | 28,4 | 9,41 | 2,97 | 2 | 0,737 | 0,809 | 0,642 | 0,658 | 0,884 | 0,935 | 0,855 | 0,879 | 0,80 | -0,20 | 5,7E-03 |
| P36916 | Guanine nucleotide-binding protein-like 1 OS=Mus musculus OX=10 | 13 | 7 | 7 | 7 | 607 | 68,7 | 5,68 | 19,55 | 7 | 0,961 | 0,889 | 0,88 | 0,904 | 0,965 | 1,01 | 1,033 | 1,014 | 0,90 | -0,10 | 5,8E-03 |
| Q9D6E4 | Tetratricopeptide repeat protein 9B OS=Mus musculus OX=10090 G | 12 | 2 | 2 | 2 | 239 | 25,9 | 9,48 | 3,46 | 2 | 1,129 | 1,325 | 1,288 | 1,063 | 0,784 | 1,01 | 0,835 | 0,85 | 1,38 | 0,38 | 5,9E-03 |
| P70188 | Kinesin-associated protein 3 OS=Mus musculus OX=10090 GN=Kifap | 3 | 2 | 3 | 2 | 793 | 91,2 | 5,11 | 2,97 | 2 | 0,338 | 0,735 | 0,556 | 0,895 | 1,091 | 1,182 | 1,176 | 1,098 | 0,56 | -0,44 | 6,1E-03 |
| O35857 | Mitochondrial import inner membrane translocase subunit TIM44 | 25 | 11 | 13 | 11 | 452 | 51,1 | 8,13 | 15,43 | 11 | 0,955 | 0,98 | 0,935 | 0,929 | 1,031 | 1,032 | 1,129 | 1,039 | 0,90 | -0,10 | 6,5E-03 |
| Q9D0A3 | Arpin OS=Mus musculus OX=10090 GN=Arpin PE=1 SV=1 | 5 | 1 | 1 | 1 | 226 | 25,2 | 5,19 | 2,41 | 1 | 1,223 | 1,247 | 1,214 | 1,012 | 1,007 | 0,904 | 0,91 | 0,825 | 1,29 | 0,29 | 7,3E-03 |
| O70305 | Ataxin-2 OS=Mus musculus OX=10090 GN=Atxn2 PE=1 SV=1 | 1 | 1 | 1 | 1 | 1285 | 136,4 | 9,55 | 0 | 1 | 1,424 | 1,159 | 0,83 | 1,333 | 1,801 | 1,832 | 1,716 | 1,611 | 0,68 | -0,32 | 7,5E-03 |
| O55143 | Sarcoplasmic/endoplasmic reticulum calcium ATPase 2 OS=Mus mu | 27 | 30 | 47 | 26 | 1044 | 114,8 | 5,34 | 114,94 | 30 | 0,967 | 0,951 | 0,988 | 0,958 | 1,007 | 1,034 | 0,992 | 1,016 | 0,95 | -0,05 | 8,1E-03 |
| Q8C129 | Leucyl-cystinyl aminopeptidase OS=Mus musculus OX=10090 GN=L | 5 | 5 | 6 | 5 | 1025 | 117,2 | 5,96 | 3,62 | 5 | 1,148 | 1,311 | 1,197 | 1,123 | 0,997 | 1,059 | 1,037 | 1,002 | 1,17 | 0,17 | 8,3E-03 |
| Q9D154 | Leukocyte elastase inhibitor A OS=Mus musculus OX=10090 GN=Se | 13 | 5 | 5 | 5 | 379 | 42,5 | 6,21 | 7,32 | 5 | 1,123 | 1,038 | 1,095 | 0,986 | 0,92 | 0,972 | 0,923 | 0,904 | 1,14 | 0,14 | 8,3E-03 |
| Q8K2C7 | Protein OS-9 OS=Mus musculus OX=10090 GN=Os9 PE=1 SV=2 | 2 | 1 | 1 | 1 | 672 | 76,1 | 4,84 | 0 | 1 | 1,284 | 1,315 | 1,282 | 1,547 | 1,106 | 0,597 | 0,677 | 0,96 | 1,63 | 0,63 | 8,4E-03 |
| Q8BK30 | NADH dehydrogenase [ubiquinone] flavoprotein 3, mitochondrial C | 16 | 2 | 4 | 2 | 104 | 11,8 | 9,35 | 7,53 | 2 | 0,862 | 0,723 | 0,804 | 0,883 | 1,071 | 0,908 | 1,069 | 1,117 | 0,79 | -0,21 | 8,5E-03 |
| P32883-2 | Isoform 2B of GTPase KRas OS=Mus musculus OX=10090 GN=Kras | 48 | 7 | 15 | 3 | 188 | 21,5 | 8,12 | 32,39 | 7 | 1,177 | 1,261 | 1,237 | 1,214 | 1,002 | 1,153 | 1,024 | 1,106 | 1,14 | 0,14 | 8,8E-03 |
| Q68FD5 | Clathrin heavy chain 1 OS=Mus musculus OX=10090 GN=Cltc PE=1 | 32 | 55 | 128 | 55 | 1675 | 191,4 | 5,69 | 366,65 | 55 | 0,93 | 0,935 | 0,964 | 1,01 | 1,039 | 1,014 | 1,05 | 1,038 | 0,93 | -0,07 | 8,9E-03 |
| Q9D051 | Pyruvate dehydrogenase E1 component subunit beta, mitochondri | 35 | 11 | 20 | 11 | 359 | 38,9 | 6,87 | 58,29 | 11 | 0,954 | 0,984 | 0,955 | 0,953 | 1,002 | 1,008 | 1,044 | 0,994 | 0,95 | -0,05 | 9,2E-03 |
| Q9D9E0 | Solute carrier family 22 member 17 OS=Mus musculus OX=10090 G | 2 | 1 | 1 | 1 | 401 | 43,2 | 8,73 | 3,35 | 1 | 1,256 | 1,158 | 1,097 | 1,09 | 1,017 | 0,864 | 0,989 | 0,963 | 1,20 | 0,20 | 9,2E-03 |
| Q9D819 | Inorganic pyrophosphatase OS=Mus musculus OX=10090 GN=Ppa1 | 34 | 8 | 15 | 8 | 289 | 32,6 | 5,6 | 32,6 | 8 | 1,106 | 1,056 | 1,16 | 1,081 | 0,963 | 1,014 | 0,989 | 1,032 | 1,10 | 0,10 | 9,3E-03 |
| Q6X893 | Choline transporter-like protein 1 OS=Mus musculus OX=10090 GN | 3 | 2 | 3 | 2 | 653 | 73 | 8,75 | 9,88 | 2 | 1,208 | 1,214 | 1,164 | 1,111 | 1,006 | 1,048 | 0,809 | 0,969 | 1,23 | 0,23 | 9,3E-03 |
| P11438 | Lysosome-associated membrane glycoprotein 1 OS=Mus musculus | 5 | 2 | 3 | 2 | 406 | 43,8 | 8,4 | 2,54 | 2 | 0,94 | 0,96 | 1,046 | 0,962 | 1,137 | 1,039 | 1,102 | 1,093 | 0,89 | -0,11 | 9,8E-03 |

| | | | | | | | | | | | | | | | | | | | | | |
|---|---|---|---|---|---|---|---|---|---|---|---|---|---|---|---|---|---|---|---|---|---|
| Q61743 | ATP-sensitive inward rectifier potassium channel 11 OS=Mus muscu | 2 | 1 | 1 | 1 | 390 | 43,5 | 8,19 | 0 | 1 | 1,767 | 1,784 | 1,383 | 1,764 | 1,234 | 1,316 | 1,367 | 1,272 | 1,29 | 0,29 | 9,8E-03 |
| Q8CDN6 | Thioredoxin-like protein 1 OS=Mus musculus OX=10090 GN=Txnl1 I | 18 | 4 | 7 | 4 | 289 | 32,2 | 4,96 | 13,45 | 4 | 0,868 | 0,918 | 0,842 | 0,951 | 1,061 | 0,959 | 1,046 | 1,114 | 0,86 | -0,14 | 9,9E-03 |
| P28652 | Calcium/calmodulin-dependent protein kinase type II subunit beta | 36 | 16 | 42 | 11 | 542 | 60,4 | 7,28 | 109,44 | 16 | 0,859 | 0,849 | 0,843 | 0,838 | 1,002 | 0,937 | 0,935 | 0,883 | 0,90 | -0,10 | 9,9E-03 |
| P26443 | Glutamate dehydrogenase 1, mitochondrial OS=Mus musculus OX= | 41 | 24 | 34 | 24 | 558 | 61,3 | 8 | 97,13 | 24 | 0,933 | 0,882 | 0,943 | 0,962 | 1,093 | 0,996 | 0,999 | 1,058 | 0,90 | -0,10 | 1,1E-02 |
| Q8CAQ8-2 | Isoform 2 of MICOS complex subunit Mic60 OS=Mus musculus OX= | 56 | 45 | 73 | 45 | 746 | 82,9 | 6,61 | 162,98 | 45 | 0,851 | 0,911 | 0,918 | 0,928 | 1,003 | 0,949 | 0,97 | 1,003 | 0,92 | -0,08 | 1,1E-02 |
| P98086 | Complement C1q subcomponent subunit A OS=Mus musculus OX= | 5 | 1 | 1 | 1 | 245 | 26 | 9,11 | 3,77 | 1 | 0,858 | 1,174 | 1,155 | 1,107 | 0,838 | 0,79 | 0,819 | 0,766 | 1,34 | 0,34 | 1,1E-02 |
| P80316 | T-complex protein 1 subunit epsilon OS=Mus musculus OX=10090 ( | 23 | 13 | 16 | 13 | 541 | 59,6 | 6,02 | 30,02 | 13 | 1,04 | 0,999 | 1,053 | 1,019 | 0,979 | 0,939 | 0,993 | 0,958 | 1,06 | 0,06 | 1,1E-02 |
| Q9DCA2 | 28S ribosomal protein S11, mitochondrial OS=Mus musculus OX=1( | 14 | 2 | 2 | 2 | 191 | 20,2 | 10,77 | 4,72 | 2 | 1,404 | 1,626 | 1,381 | 1,286 | 0,946 | 1,216 | 1,078 | 1,142 | 1,30 | 0,30 | 1,2E-02 |
| Q93092 | Transaldolase OS=Mus musculus OX=10090 GN=Taldo1 PE=1 SV=2 | 10 | 3 | 4 | 3 | 337 | 37,4 | 7,03 | 7,97 | 3 | 1,194 | 1,098 | 1,011 | 1,185 | 0,941 | 0,92 | 1,015 | 0,879 | 1,20 | 0,20 | 1,2E-02 |
| Q9DBX6 | Cytochrome P450 2S1 OS=Mus musculus OX=10090 GN=Cyp2s1 PE | 3 | 1 | 2 | 1 | 501 | 55,6 | 8,78 | 7,42 | 1 | 0,988 | 1,089 | 1,259 | 0,833 | 1,344 | 1,408 | 1,828 | 1,794 | 0,65 | -0,35 | 1,2E-02 |
| Q5PR73 | GTP-binding protein Di-Ras2 OS=Mus musculus OX=10090 GN=Dira | 38 | 7 | 8 | 6 | 199 | 22,5 | 8,76 | 19,44 | 7 | 0,979 | 0,978 | 1,022 | 0,955 | 1,024 | 1,114 | 1,067 | 1,058 | 0,92 | -0,08 | 1,2E-02 |
| Q9DBT5 | AMP deaminase 2 OS=Mus musculus OX=10090 GN=Ampd2 PE=1 S | 7 | 6 | 6 | 6 | 798 | 92 | 6,23 | 10,36 | 6 | 1,059 | 0,992 | 0,976 | 1,04 | 1,066 | 1,093 | 1,12 | 1,123 | 0,92 | -0,08 | 1,2E-02 |
| Q99LX0 | Protein/nucleic acid deglycase DJ-1 OS=Mus musculus OX=10090 G | 46 | 10 | 17 | 10 | 189 | 20 | 6,77 | 37,85 | 10 | 1,085 | 1,065 | 1,052 | 1,107 | 0,97 | 1,018 | 0,985 | 1,045 | 1,07 | 0,07 | 1,3E-02 |
| Q8BZ98 | Dynamin-3 OS=Mus musculus OX=10090 GN=Dnm3 PE=1 SV=1 | 20 | 18 | 47 | 8 | 863 | 97,1 | 8,35 | 91,55 | 18 | 0,988 | 0,937 | 0,994 | 1,028 | 1,045 | 1,095 | 1,051 | 1,118 | 0,92 | -0,08 | 1,3E-02 |
| Q9JHR7 | Insulin-degrading enzyme OS=Mus musculus OX=10090 GN=Ide PE= | 4 | 3 | 3 | 3 | 1019 | 117,7 | 6,54 | 6,72 | 3 | 1,152 | 1,169 | 1,193 | 1,087 | 1,022 | 0,993 | 1,03 | 1,102 | 1,11 | 0,11 | 1,3E-02 |
| Q6VNB8 | WD repeat and FYVE domain-containing protein 3 OS=Mus muscul | 3 | 9 | 9 | 9 | 3508 | 392,1 | 6,8 | 16,43 | 9 | 0,978 | 0,977 | 0,987 | 0,903 | 0,916 | 0,86 | 0,826 | 0,866 | 1,11 | 0,11 | 1,3E-02 |
| Q9DBP5 | UMP-CMP kinase OS=Mus musculus OX=10090 GN=Cmpk1 PE=1 SV | 23 | 5 | 9 | 5 | 196 | 22,2 | 5,83 | 12,81 | 5 | 1,106 | 1,055 | 1,12 | 1,013 | 0,965 | 0,956 | 0,985 | 1,009 | 1,10 | 0,10 | 1,3E-02 |
| Q9CQN7 | 39S ribosomal protein L41, mitochondrial OS=Mus musculus OX=1( | 7 | 1 | 2 | 1 | 135 | 15,3 | 9,82 | 4,78 | 1 | 1,038 | 0,996 | 1,059 | 1,066 | 1,196 | 1,125 | 1,091 | 1,222 | 0,90 | -0,10 | 1,3E-02 |
| Q3ZT31 | Sorting nexin-25 OS=Mus musculus OX=10090 GN=Snx25 PE=1 SV= | 3 | 2 | 2 | 2 | 840 | 97,1 | 6,29 | 2,79 | 2 | 1,243 | 1,222 | 1,103 | 1,225 | 1,444 | 1,458 | 1,286 | 1,601 | 0,83 | -0,17 | 1,3E-02 |
| Q7TSK2 | Seizure protein 6 OS=Mus musculus OX=10090 GN=Sez6 PE=1 SV=1 | 1 | 1 | 1 | 1 | 991 | 107,4 | 5,36 | 2,24 | 1 | 1,399 | 1,477 | 1,46 | 1,281 | 0,924 | 1,216 | 1,222 | 1,113 | 1,26 | 0,26 | 1,3E-02 |
| P16332 | Methylmalonyl-CoA mutase, mitochondrial OS=Mus musculus OX= | 11 | 8 | 8 | 8 | 748 | 82,8 | 6,89 | 11,96 | 8 | 0,959 | 0,851 | 1,097 | 0,918 | 1,121 | 1,179 | 1,104 | 1,206 | 0,83 | -0,17 | 1,4E-02 |
| Q80TZ3 | Putative tyrosine-protein phosphatase auxilin OS=Mus musculus O | 16 | 14 | 22 | 12 | 938 | 102,2 | 7,23 | 37,96 | 14 | 0,933 | 0,879 | 0,886 | 0,937 | 0,971 | 0,984 | 0,961 | 0,95 | 0,94 | -0,06 | 1,4E-02 |
| P30999 | Catenin delta-1 OS=Mus musculus OX=10090 GN=Ctnnd1 PE=1 SV= | 14 | 14 | 14 | 14 | 938 | 104,9 | 6,87 | 14,19 | 14 | 1,139 | 1,179 | 1,134 | 1,05 | 1,058 | 1,022 | 1,016 | 0,992 | 1,10 | 0,10 | 1,4E-02 |
| Q5QNQ6 | Oxysterol-binding protein 2 OS=Mus musculus OX=10090 GN=Osbp | 4 | 3 | 3 | 2 | 908 | 101,3 | 7,14 | 4,87 | 3 | 0,818 | 0,963 | 0,79 | 0,714 | 1,002 | 0,953 | 1,1 | 1,084 | 0,79 | -0,21 | 1,4E-02 |
| Q9CQ40 | 39S ribosomal protein L49, mitochondrial OS=Mus musculus OX=1( | 13 | 2 | 3 | 2 | 166 | 19,1 | 9,5 | 7,05 | 2 | 1,18 | 1,236 | 1,044 | 1,039 | 0,904 | 0,964 | 0,93 | 0,986 | 1,19 | 0,19 | 1,5E-02 |
| Q8BH95 | Enoyl-CoA hydratase, mitochondrial OS=Mus musculus OX=10090 ( | 23 | 6 | 9 | 6 | 290 | 31,5 | 8,48 | 26,32 | 6 | 0,998 | 0,957 | 0,985 | 0,928 | 1,054 | 1,093 | 1,031 | 1,01 | 0,92 | -0,08 | 1,5E-02 |
| Q9CRB9 | MICOS complex subunit Mic19 OS=Mus musculus OX=10090 GN=C | 32 | 8 | 12 | 8 | 227 | 26,3 | 8,37 | 27,89 | 8 | 0,757 | 0,848 | 0,82 | 0,922 | 1,093 | 0,98 | 0,952 | 0,962 | 0,84 | -0,16 | 1,5E-02 |
| O89026 | Roundabout homolog 1 OS=Mus musculus OX=10090 GN=Robo1 P | 1 | 1 | 1 | 1 | 1612 | 176,3 | 6,18 | 2,09 | 1 | 0,942 | 0,87 | 1,106 | 0,841 | 1,039 | 1,388 | 1,344 | 1,444 | 0,72 | -0,28 | 1,5E-02 |
| P84091 | AP-2 complex subunit mu OS=Mus musculus OX=10090 GN=Ap2m1 | 48 | 19 | 32 | 19 | 435 | 49,6 | 9,54 | 79,15 | 19 | 0,939 | 0,972 | 0,961 | 0,991 | 1,013 | 0,991 | 1,038 | 1,015 | 0,95 | -0,05 | 1,5E-02 |
| O08997 | Copper transport protein ATOX1 OS=Mus musculus OX=10090 GN= | 31 | 2 | 4 | 2 | 68 | 7,3 | 6,51 | 11,31 | 2 | 0,958 | 0,864 | 0,941 | 0,939 | 1,001 | 1,172 | 1,086 | 1,024 | 0,86 | -0,14 | 1,6E-02 |
| Q7TT50 | Serine/threonine-protein kinase MRCK beta OS=Mus musculus OX= | 8 | 15 | 18 | 14 | 1713 | 194,6 | 6,46 | 28,94 | 15 | 1,199 | 1,154 | 1,153 | 1,077 | 0,934 | 1,1 | 0,968 | 1,003 | 1,14 | 0,14 | 1,6E-02 |
| O70252 | Heme oxygenase 2 OS=Mus musculus OX=10090 GN=Hmox2 PE=1 | 6 | 2 | 2 | 2 | 315 | 35,7 | 5,87 | 4,53 | 2 | 0,802 | 0,887 | 0,965 | 0,892 | 1,07 | 0,985 | 0,996 | 1,002 | 0,87 | -0,13 | 1,7E-02 |
| Q91W86 | Vacuolar protein sorting-associated protein 11 homolog OS=Mus m | 5 | 5 | 5 | 5 | 941 | 107,7 | 7,01 | 6,82 | 5 | 0,645 | 0,769 | 0,795 | 0,835 | 1,096 | 0,856 | 1,048 | 0,938 | 0,77 | -0,23 | 1,7E-02 |
| P21440 | Phosphatidylcholine translocator ABCB4 OS=Mus musculus OX=100 | 1 | 1 | 1 | 1 | 1276 | 140,3 | 8,78 | 0 | 1 | 1,239 | 1,245 | 0,903 | 0,961 | 0,812 | 0,783 | 0,765 | 0,536 | 1,50 | 0,50 | 1,7E-02 |
| Q5SXY1-3 | Isoform 4 of Cytospin-B OS=Mus musculus OX=10090 GN=Specc1 | 3 | 3 | 3 | 1 | 987 | 109,8 | 5,86 | 4,24 | 3 | 1,184 | 0,865 | 1,09 | 0,975 | 0,577 | 0,757 | 0,876 | 0,575 | 1,48 | 0,48 | 1,7E-02 |
| Q8K1R7 | Serine/threonine-protein kinase Nek9 OS=Mus musculus OX=1009( | 2 | 1 | 1 | 1 | 984 | 107,1 | 5,63 | 3,62 | 1 | 0,943 | 1,066 | 1,206 | 0,928 | 0,821 | 0,837 | 0,822 | 0,812 | 1,26 | 0,26 | 1,7E-02 |
| Q9R1P1 | Proteasome subunit beta type-3 OS=Mus musculus OX=10090 GN= | 22 | 3 | 4 | 3 | 205 | 22,9 | 6,55 | 15,18 | 3 | 1,428 | 1,313 | 1,191 | 1,38 | 0,96 | 1,208 | 1,091 | 1,112 | 1,22 | 0,22 | 1,8E-02 |
| Q8BGC4 | Prostaglandin reductase-3 OS=Mus musculus OX=10090 GN=Zadh2 | 14 | 5 | 6 | 5 | 377 | 40,5 | 7,42 | 13 | 5 | 1,341 | 1,184 | 1,162 | 1,255 | 1,103 | 1,102 | 1,044 | 1,125 | 1,13 | 0,13 | 1,8E-02 |
| Q8BFS6 | Serine/threonine-protein phosphatase CPPED1 OS=Mus musculus ( | 4 | 1 | 1 | 1 | 312 | 35,2 | 5,34 | 3,5 | 1 | 0,939 | 1,011 | 0,948 | 0,966 | 0,902 | 0,886 | 0,866 | 0,934 | 1,08 | 0,08 | 1,8E-02 |
| Q91XD7 | Protein disulfide isomerase Creld1 OS=Mus musculus OX=10090 GN | 17 | 6 | 8 | 6 | 420 | 45,7 | 5,02 | 12,23 | 6 | 1,037 | 0,985 | 1,026 | 0,995 | 1,099 | 1,045 | 1,187 | 1,12 | 0,91 | -0,09 | 1,9E-02 |
| P29341 | Polyadenylate-binding protein 1 OS=Mus musculus OX=10090 GN= | 15 | 10 | 10 | 10 | 636 | 70,6 | 9,5 | 24,59 | 10 | 1,053 | 0,982 | 1,064 | 1,019 | 0,981 | 0,972 | 0,938 | 0,962 | 1,07 | 0,07 | 1,9E-02 |
| Q9CQ85 | Mitochondrial import inner membrane translocase subunit Tim22 ( | 8 | 1 | 1 | 1 | 194 | 20,1 | 8,22 | 2,33 | 1 | 1,516 | 1,055 | 1,017 | 1,089 | 0,692 | 0,486 | 0,921 | 0,702 | 1,67 | 0,67 | 1,9E-02 |
| Q99L47 | Hsc70-interacting protein OS=Mus musculus OX=10090 GN=St13 PI | 14 | 5 | 9 | 5 | 371 | 41,6 | 5,26 | 18,82 | 5 | 1,03 | 0,974 | 0,954 | 1,005 | 1,063 | 1,052 | 1,026 | 1,067 | 0,94 | -0,06 | 1,9E-02 |
| P62334 | 26S proteasome regulatory subunit 10B OS=Mus musculus OX=100 | 24 | 10 | 14 | 10 | 389 | 44,1 | 7,49 | 23,33 | 10 | 0,887 | 0,907 | 0,997 | 0,968 | 1,041 | 1,026 | 1,057 | 1,172 | 0,88 | -0,13 | 1,9E-02 |
| P47802 | Metaxin-1 OS=Mus musculus OX=10090 GN=Mtx1 PE=1 SV=1 | 27 | 10 | 11 | 10 | 317 | 35,6 | 6,18 | 23,04 | 10 | 0,866 | 0,871 | 0,961 | 0,914 | 1,018 | 1,077 | 1,071 | 0,94 | 0,88 | -0,12 | 1,9E-02 |
| Q9CQC7 | NADH dehydrogenase [ubiquinone] 1 beta subcomplex subunit 4 O | 47 | 6 | 8 | 6 | 129 | 15,1 | 9,89 | 17,19 | 6 | 0,924 | 1,039 | 0,821 | 0,96 | 1,062 | 1,073 | 1,101 | 1,097 | 0,86 | -0,14 | 1,9E-02 |
| Q9Z0R9 | Acyl-CoA 6-desaturase OS=Mus musculus OX=10090 GN=Fads2 PE= | 2 | 1 | 1 | 1 | 444 | 52,4 | 8,82 | 2,28 | 1 | 1,058 | 1,256 | 1,029 | 1,12 | 0,851 | 0,729 | 0,871 | 1,019 | 1,29 | 0,29 | 1,9E-02 |
| P32921 | Tryptophan--tRNA ligase, cytoplasmic OS=Mus musculus OX=1009( | 27 | 11 | 12 | 11 | 481 | 54,3 | 6,89 | 21,45 | 11 | 1,138 | 1,01 | 1,048 | 1,025 | 0,818 | 0,912 | 0,976 | 0,947 | 1,16 | 0,16 | 1,9E-02 |
| Q9WUM5 | Succinate--CoA ligase [ADP/GDP-forming] subunit alpha, mitochon | 21 | 7 | 18 | 7 | 346 | 36,1 | 9,39 | 38,34 | 7 | 0,848 | 0,814 | 0,932 | 0,886 | 1,009 | 0,932 | 1,01 | 0,943 | 0,89 | -0,11 | 2,0E-02 |
| Q00915 | Retinol-binding protein 1 OS=Mus musculus OX=10090 GN=Rbp1 P | 9 | 1 | 2 | 1 | 135 | 15,8 | 5,25 | 2,82 | 1 | 0,901 | 0,826 | 0,625 | 1,053 | 1,184 | 1,186 | 1,165 | 1,056 | 0,74 | -0,26 | 2,0E-02 |
| P62746 | Rho-related GTP-binding protein RhoB OS=Mus musculus OX=1009 | 29 | 6 | 8 | 5 | 196 | 22,1 | 5,24 | 25,75 | 6 | 1,031 | 1,076 | 1,057 | 1,021 | 0,975 | 1,024 | 0,956 | 0,991 | 1,06 | 0,06 | 2,0E-02 |
| Q99JI4 | 26S proteasome non-ATPase regulatory subunit 6 OS=Mus muscul | 23 | 9 | 13 | 9 | 389 | 45,5 | 5,52 | 18,77 | 9 | 0,907 | 0,897 | 1,047 | 0,99 | 1,271 | 1,018 | 1,193 | 1,158 | 0,83 | -0,17 | 2,0E-02 |
| A2A690 | Protein TANC2 OS=Mus musculus OX=10090 GN=Tanc2 PE=1 SV=1 | 6 | 10 | 10 | 10 | 1994 | 220,1 | 7,97 | 19,12 | 10 | 0,926 | 0,919 | 0,923 | 0,942 | 0,96 | 1,021 | 0,95 | 1,011 | 0,94 | -0,06 | 2,0E-02 |
| Q8BGY7 | Protein FAM210A OS=Mus musculus OX=10090 GN=Fam210a PE=1 | 4 | 1 | 1 | 1 | 273 | 31,6 | 9,82 | 2,51 | 1 | 0,643 | 0,74 | 0,879 | 0,781 | 0,896 | 0,914 | 0,964 | 1,095 | 0,79 | -0,21 | 2,1E-02 |
| Q62420 | Endophilin-A1 OS=Mus musculus OX=10090 GN=Sh3gl2 PE=1 SV=2 | 30 | 13 | 19 | 10 | 352 | 39,9 | 5,39 | 52,76 | 13 | 0,969 | 1,026 | 0,985 | 1,075 | 0,924 | 0,852 | 0,908 | 0,962 | 1,11 | 0,11 | 2,1E-02 |
| Q8CEE7 | Retinol dehydrogenase 13 OS=Mus musculus OX=10090 GN=Rdh13 | 2 | 1 | 1 | 1 | 334 | 36,4 | 8,85 | 2,69 | 1 | 1,203 | 1,196 | 1,069 | 1,008 | 0,908 | 0,863 | 0,958 | 1,02 | 1,19 | 0,19 | 2,1E-02 |
| O35075 | Vacuolar protein sorting-associated protein 26C OS=Mus musculus | 10 | 2 | 3 | 2 | 297 | 32,9 | 7,68 | 5,37 | 2 | 0,809 | 0,833 | 0,946 | 1,018 | 1,045 | 1,345 | 1,164 | 1,079 | 0,78 | -0,22 | 2,1E-02 |
| Q8K0Z7 | Translational activator of cytochrome c oxidase 1 OS=Mus musculu | 16 | 4 | 4 | 4 | 294 | 32,3 | 8,12 | 6,08 | 4 | 1,28 | 1,113 | 1,187 | 1,133 | 0,955 | 1,105 | 1,049 | 0,984 | 1,15 | 0,15 | 2,2E-02 |
| Q7TQH0 | Ataxin-2-like protein OS=Mus musculus OX=10090 GN=Atxn2l PE=1 | 2 | 2 | 2 | 2 | 1049 | 110,6 | 8,85 | 4,35 | 2 | 0,911 | 1,027 | 0,849 | 0,898 | 1,071 | 1,013 | 1,086 | 1,027 | 0,88 | -0,12 | 2,2E-02 |
| Q3TUH1 | Phosphatidate cytidylyltransferase, mitochondrial OS=Mus muscul | 3 | 1 | 2 | 1 | 337 | 37,8 | 8,98 | 5,13 | 1 | 1,036 | 1,138 | 1,067 | 1,095 | 1,131 | 1,252 | 1,261 | 1,385 | 0,86 | -0,14 | 2,2E-02 |
| P03995 | Glial fibrillary acidic protein OS=Mus musculus OX=10090 GN=Gfap | 34 | 17 | 22 | 16 | 430 | 49,9 | 5,34 | 54,49 | 17 | 1,096 | 1,184 | 1,022 | 1,036 | 0,927 | 0,898 | 1,011 | 0,959 | 1,14 | 0,14 | 2,2E-02 |
| Q791V5 | Mitochondrial carrier homolog 2 OS=Mus musculus OX=10090 GN= | 19 | 5 | 6 | 5 | 303 | 33,5 | 8,25 | 16,53 | 5 | 0,923 | 0,886 | 0,917 | 0,949 | 1,065 | 0,947 | 1,013 | 0,991 | 0,92 | -0,08 | 2,2E-02 |

| | | | | | | | | | | | | | | | | | | | | |
|---|---|---|---|---|---|---|---|---|---|---|---|---|---|---|---|---|---|---|---|---|
| Q6URW6 | Myosin-14 OS=Mus musculus OX=10090 GN=Myh14 PE=1 SV=1 | 8 | 14 | 16 | 9 | 2000 | 228,4 | 5,55 | 35,82 | 14 | 1,084 | 1,029 | 1,044 | 0,996 | 0,976 | 0,973 | 0,953 | 0,999 | 1,06 | 0,06 | 2,2E-02 |
| P59017 | Bcl-2-like protein 13 OS=Mus musculus OX=10090 GN=Bcl2l13 PE=1 | 2 | 1 | 2 | 1 | 434 | 46,7 | 4,59 | 4,58 | 1 | 0,819 | 0,906 | 0,901 | 0,879 | 0,931 | 1,017 | 1,132 | 0,994 | 0,86 | -0,14 | 2,2E-02 |
| Q9D415 | Disks large-associated protein 1 OS=Mus musculus OX=10090 GN=D | 14 | 11 | 12 | 9 | 992 | 110,3 | 7,09 | 19,71 | 11 | 1,365 | 1,284 | 1,217 | 1,06 | 0,884 | 1,105 | 0,875 | 1,019 | 1,27 | 0,27 | 2,2E-02 |
| O70325 | Phospholipid hydroperoxide glutathione peroxidase OS=Mus muscu | 20 | 5 | 10 | 5 | 197 | 22,2 | 8,46 | 21,07 | 5 | 1,137 | 1,082 | 0,944 | 1,021 | 1,117 | 1,282 | 1,279 | 1,202 | 0,86 | -0,14 | 2,2E-02 |
| P31650 | Sodium- and chloride-dependent GABA transporter 3 OS=Mus mus | 10 | 7 | 9 | 7 | 627 | 69,9 | 6,98 | 23,76 | 7 | 1,208 | 1,085 | 1,174 | 1,098 | 0,959 | 1,105 | 0,956 | 0,983 | 1,14 | 0,14 | 2,3E-02 |
| Q7TT37 | Elongator complex protein 1 OS=Mus musculus OX=10090 GN=Elp1 | 4 | 5 | 5 | 5 | 1333 | 149,5 | 6 | 2,02 | 5 | 0,966 | 1,029 | 0,907 | 0,959 | 0,861 | 0,857 | 0,769 | 0,908 | 1,14 | 0,14 | 2,3E-02 |
| P03921 | NADH-ubiquinone oxidoreductase chain 5 OS=Mus musculus OX=10 | 1 | 1 | 2 | 1 | 607 | 68,4 | 9,03 | 5,13 | 1 | 0,856 | 0,886 | 0,908 | 0,86 | 0,999 | 1,171 | 0,969 | 0,974 | 0,85 | -0,15 | 2,3E-02 |
| Q68FH0 | Plakophilin-4 OS=Mus musculus OX=10090 GN=Pkp4 PE=1 SV=1 | 9 | 11 | 12 | 8 | 1190 | 131,5 | 8,94 | 16,71 | 11 | 0,88 | 0,899 | 0,972 | 0,861 | 1,037 | 0,959 | 0,969 | 1,02 | 0,91 | -0,09 | 2,3E-02 |
| P63038 | 60 kDa heat shock protein, mitochondrial OS=Mus musculus OX=10 | 46 | 27 | 58 | 27 | 573 | 60,9 | 6,18 | 147,09 | 27 | 0,936 | 0,971 | 0,967 | 0,96 | 0,986 | 0,987 | 0,997 | 0,974 | 0,97 | -0,03 | 2,4E-02 |
| Q9JHL1 | Na(+)/H(+) exchange regulatory cofactor NHE-RF2 OS=Mus muscul | 12 | 4 | 4 | 4 | 337 | 37,4 | 7,59 | 4,94 | 4 | 1,421 | 1,208 | 1,353 | 1,046 | 0,889 | 1,075 | 0,946 | 1,011 | 1,28 | 0,28 | 2,4E-02 |
| Q8BH50 | Uncharacterized protein C18orf25 homolog OS=Mus musculus OX= | 9 | 1 | 1 | 1 | 245 | 26,4 | 5,19 | 2,61 | 1 | 1,194 | 0,949 | 1,051 | 1,122 | 0,525 | 0,01 | 0,701 | 0,836 | 2,08 | 1,08 | 2,5E-02 |
| Q922H2 | [Pyruvate dehydrogenase (acetyl-transferring)] kinase isozyme 3, m | 21 | 8 | 11 | 8 | 415 | 47,9 | 8,82 | 21,34 | 8 | 0,961 | 0,994 | 1,039 | 0,983 | 1,125 | 1,1 | 1,014 | 1,081 | 0,92 | -0,08 | 2,5E-02 |
| O08749 | Dihydrolipoyl dehydrogenase, mitochondrial OS=Mus musculus OX | 26 | 15 | 33 | 15 | 509 | 54,2 | 7,9 | 69,35 | 15 | 0,847 | 0,903 | 0,93 | 0,873 | 1,011 | 0,918 | 0,975 | 0,96 | 0,92 | -0,08 | 2,6E-02 |
| Q8K386 | Ras-related protein Rab-15 OS=Mus musculus OX=10090 GN=Rab1 | 16 | 3 | 3 | 2 | 212 | 24,3 | 5,71 | 5,13 | 3 | 1,334 | 1,024 | 1,025 | 0,97 | 0,804 | 0,942 | 0,657 | 0,747 | 1,38 | 0,38 | 2,6E-02 |
| P51150 | Ras-related protein Rab-7a OS=Mus musculus OX=10090 GN=Rab7a | 54 | 12 | 14 | 12 | 207 | 23,5 | 6,7 | 32,22 | 12 | 1,168 | 1,076 | 1,08 | 1,069 | 0,931 | 1,052 | 0,998 | 1,009 | 1,10 | 0,10 | 2,6E-02 |
| Q80VP9 | Aspartate beta-hydroxylase domain-containing protein 2 OS=Mus r | 10 | 3 | 3 | 3 | 343 | 38,7 | 7,05 | 3,37 | 3 | 1,12 | 0,949 | 1,331 | 1,022 | 0,912 | 0,839 | 0,86 | 0,697 | 1,34 | 0,34 | 2,6E-02 |
| P35438 | Glutamate receptor ionotropic, NMDA 1 OS=Mus musculus OX=100 | 10 | 10 | 10 | 10 | 938 | 105,4 | 8,84 | 24,74 | 10 | 0,9 | 0,924 | 0,963 | 0,906 | 0,975 | 0,955 | 0,963 | 0,995 | 0,95 | -0,05 | 2,6E-02 |
| E9Q4S1 | High affinity cAMP-specific and IBMX-insensitive 3',5'-cyclic phosph | 4 | 3 | 3 | 3 | 865 | 96,7 | 6,86 | 2,09 | 3 | 1,365 | 0,967 | 1,108 | 0,945 | 0,816 | 0,525 | 0,829 | 0,8 | 1,48 | 0,48 | 2,6E-02 |
| O55023 | Inositol monophosphatase 1 OS=Mus musculus OX=10090 GN=Imp | 33 | 9 | 13 | 9 | 277 | 30,4 | 5,19 | 34,4 | 9 | 1,056 | 0,958 | 1,016 | 1,071 | 0,933 | 0,926 | 0,983 | 0,917 | 1,09 | 0,09 | 2,7E-02 |
| Q9ERR1 | Nuclear distribution protein nudE-like 1 OS=Mus musculus OX=100 | 23 | 8 | 9 | 6 | 345 | 38,3 | 5,24 | 22,38 | 8 | 0,927 | 0,786 | 0,843 | 0,9 | 0,895 | 1,121 | 1,076 | 1,043 | 0,84 | -0,16 | 2,7E-02 |
| Q9EQQ9 | Protein O-GlcNAcase OS=Mus musculus OX=10090 GN=Oga PE=1 S | 2 | 2 | 2 | 2 | 916 | 103,1 | 4,92 | 5,06 | 2 | 1,143 | 1,312 | 1,21 | 1,068 | 0,978 | 1,047 | 1,068 | 0,98 | 1,16 | 0,16 | 2,7E-02 |
| O88952 | Protein lin-7 homolog C OS=Mus musculus OX=10090 GN=Lin7c PE | 21 | 3 | 6 | 2 | 197 | 21,8 | 8,43 | 11,07 | 3 | 0,828 | 1,021 | 0,813 | 0,922 | 1,008 | 1,075 | 1,018 | 1,09 | 0,86 | -0,14 | 2,7E-02 |
| Q9D7N3 | 28S ribosomal protein S9, mitochondrial OS=Mus musculus OX=100 | 6 | 2 | 3 | 2 | 390 | 44,9 | 8,81 | 0 | 2 | 1,152 | 1,151 | 1,21 | 1,124 | 0,828 | 1,038 | 1,055 | 1,047 | 1,17 | 0,17 | 2,7E-02 |
| Q8R3P0 | Aspartoacylase OS=Mus musculus OX=10090 GN=Aspa PE=1 SV=2 | 15 | 5 | 8 | 5 | 312 | 35,3 | 6,58 | 15,18 | 5 | 1,127 | 1,094 | 1,074 | 1,043 | 0,994 | 1,001 | 1,04 | 0,898 | 1,10 | 0,10 | 2,7E-02 |
| Q8BG51 | Mitochondrial Rho GTPase 1 OS=Mus musculus OX=10090 GN=Rho | 13 | 6 | 6 | 5 | 631 | 72,2 | 6,49 | 7,56 | 6 | 0,837 | 0,793 | 0,845 | 0,909 | 1,085 | 0,881 | 0,966 | 1,034 | 0,85 | -0,15 | 2,7E-02 |
| Q9JI46 | Diphosphoinositol polyphosphate phosphohydrolase 1 OS=Mus mu | 22 | 3 | 3 | 2 | 168 | 19 | 6,34 | 7,55 | 3 | 0,997 | 1,042 | 1,005 | 0,969 | 0,916 | 0,948 | 0,959 | 0,97 | 1,06 | 0,06 | 2,8E-02 |
| Q62448 | Eukaryotic translation initiation factor 4 gamma 2 OS=Mus muscul | 5 | 4 | 5 | 4 | 906 | 102 | 7,14 | 5,21 | 4 | 1,242 | 1,133 | 1,065 | 0,933 | 0,787 | 0,954 | 0,761 | 0,932 | 1,27 | 0,27 | 2,8E-02 |
| Q91WE1 | Sorting nexin-15 OS=Mus musculus OX=10090 GN=Snx15 PE=1 SV= | 3 | 1 | 1 | 1 | 337 | 37,7 | 5,24 | 2,88 | 1 | 1,403 | 1,179 | 1,259 | 1,065 | 0,694 | 1,045 | 1,034 | 0,885 | 1,34 | 0,34 | 2,8E-02 |
| P97445 | Voltage-dependent P/Q-type calcium channel subunit alpha-1A OS | 6 | 13 | 16 | 12 | 2368 | 267,5 | 8,85 | 20,98 | 13 | 0,991 | 0,954 | 0,976 | 1,03 | 1,035 | 1,02 | 1,044 | 1,052 | 0,95 | -0,05 | 2,8E-02 |
| P08249 | Malate dehydrogenase, mitochondrial OS=Mus musculus OX=1009 | 58 | 19 | 68 | 19 | 338 | 35,6 | 8,68 | 164,66 | 19 | 1,02 | 1,034 | 1,033 | 1,008 | 0,955 | 0,992 | 0,996 | 1,006 | 1,04 | 0,04 | 2,8E-02 |
| Q9D4C9 | Clavesin-1 OS=Mus musculus OX=10090 GN=Clvs1 PE=1 SV=1 | 11 | 3 | 3 | 1 | 354 | 40,6 | 6,73 | 10,73 | 3 | 1,179 | 1,467 | 1,13 | 1,25 | 0,894 | 0,771 | 1,173 | 0,892 | 1,35 | 0,35 | 2,9E-02 |
| Q69ZR2 | E3 ubiquitin-protein ligase HECTD1 OS=Mus musculus OX=10090 G | 1 | 3 | 3 | 3 | 2618 | 289,9 | 5,41 | 4,29 | 3 | 1,253 | 1,213 | 1,082 | 1,27 | 0,974 | 1,15 | 0,976 | 1,039 | 1,16 | 0,16 | 2,9E-02 |
| P05532 | Mast/stem cell growth factor receptor Kit OS=Mus musculus OX=10 | 5 | 4 | 4 | 4 | 979 | 109,3 | 7,21 | 2,36 | 4 | 0,918 | 1,003 | 0,811 | 1,041 | 1,155 | 1,36 | 1,221 | 1,025 | 0,79 | -0,21 | 2,9E-02 |
| O08989 | Ras-related protein M-Ras OS=Mus musculus OX=10090 GN=Mras | 24 | 4 | 5 | 4 | 208 | 23,9 | 8,78 | 8,56 | 4 | 1,075 | 1,002 | 1,023 | 1,017 | 1,149 | 1,054 | 1,231 | 1,283 | 0,87 | -0,13 | 2,9E-02 |
| Q8QZT1 | Acetyl-CoA acetyltransferase, mitochondrial OS=Mus musculus OX= | 32 | 13 | 27 | 13 | 424 | 44,8 | 8,51 | 57,02 | 13 | 0,845 | 0,846 | 0,914 | 0,965 | 1,069 | 0,942 | 1,031 | 0,985 | 0,89 | -0,11 | 2,9E-02 |
| Q3UM45 | Protein phosphatase 1 regulatory subunit 7 OS=Mus musculus OX= | 27 | 8 | 9 | 8 | 361 | 41,3 | 4,92 | 23,8 | 8 | 0,995 | 0,991 | 0,991 | 0,986 | 0,868 | 0,93 | 0,932 | 0,977 | 1,07 | 0,07 | 2,9E-02 |
| Q3THS6 | S-adenosylmethionine synthase isoform type-2 OS=Mus musculus | 12 | 4 | 4 | 4 | 395 | 43,7 | 6,48 | 9,41 | 4 | 1,649 | 1,388 | 1,388 | 1,419 | 1,144 | 1,359 | 1,008 | 1,23 | 1,23 | 0,23 | 3,0E-02 |
| P97450 | ATP synthase-coupling factor 6, mitochondrial OS=Mus musculus O | 56 | 11 | 17 | 11 | 108 | 12,5 | 9,36 | 43,95 | 11 | 0,739 | 0,814 | 0,868 | 0,882 | 0,992 | 0,875 | 0,991 | 0,931 | 0,87 | -0,13 | 3,0E-02 |
| O88685 | 26S proteasome regulatory subunit 6A OS=Mus musculus OX=1009 | 15 | 6 | 7 | 6 | 442 | 49,5 | 5,19 | 8,06 | 6 | 1,263 | 1,282 | 1,261 | 1,023 | 0,991 | 0,927 | 1,082 | 1,037 | 1,20 | 0,20 | 3,0E-02 |
| Q9WUA3 | ATP-dependent 6-phosphofructokinase, platelet type OS=Mus mus | 18 | 18 | 28 | 18 | 784 | 85,4 | 7,11 | 54,57 | 18 | 1,102 | 1,067 | 1,097 | 0,999 | 0,953 | 1,042 | 0,951 | 0,938 | 1,10 | 0,10 | 3,0E-02 |
| Q8R5H6 | Wiskott-Aldrich syndrome protein family member 1 OS=Mus muscu | 15 | 7 | 9 | 6 | 559 | 61,5 | 6,37 | 21,85 | 7 | 1,492 | 1,494 | 1,316 | 1,064 | 0,897 | 1,177 | 0,82 | 1,025 | 1,37 | 0,37 | 3,0E-02 |
| P63040 | Complexin-1 OS=Mus musculus OX=10090 GN=Cplx1 PE=1 SV=1 | 40 | 4 | 6 | 2 | 134 | 15,1 | 4,97 | 4,8 | 4 | 0,654 | 0,542 | 0,465 | 0,836 | 1,056 | 0,695 | 1,033 | 1,042 | 0,65 | -0,35 | 3,1E-02 |
| P62702 | 40S ribosomal protein S4, X isoform OS=Mus musculus OX=10090 | 17 | 4 | 4 | 4 | 263 | 29,6 | 10,15 | 4,53 | 4 | 1,221 | 1,233 | 0,952 | 0,882 | 0,799 | 0,902 | 0,7 | 0,774 | 1,35 | 0,35 | 3,2E-02 |
| Q3TDD9 | Protein phosphatase 1 regulatory subunit 21 OS=Mus musculus OX | 15 | 12 | 16 | 12 | 780 | 88,3 | 6,9 | 32,25 | 12 | 1,151 | 1,052 | 1,202 | 1,101 | 0,962 | 1,017 | 1,056 | 1,043 | 1,10 | 0,10 | 3,2E-02 |
| O54988 | STE20-like serine/threonine-protein kinase OS=Mus musculus OX= | 13 | 15 | 17 | 15 | 1233 | 141,4 | 5,14 | 36,91 | 15 | 1,147 | 1,093 | 1,095 | 1,057 | 0,877 | 1,043 | 0,99 | 1,022 | 1,12 | 0,12 | 3,2E-02 |
| P48722 | Heat shock 70 kDa protein 4L OS=Mus musculus OX=10090 GN=Hs | 36 | 28 | 32 | 26 | 838 | 94,3 | 5,74 | 60,15 | 28 | 0,931 | 0,912 | 0,952 | 1,011 | 1,018 | 0,992 | 1,019 | 1,036 | 0,94 | -0,06 | 3,2E-02 |
| Q9JK42 | [Pyruvate dehydrogenase (acetyl-transferring)] kinase isozyme 2, m | 11 | 4 | 4 | 3 | 407 | 46 | 6,61 | 4,19 | 4 | 0,762 | 0,961 | 0,74 | 0,919 | 0,991 | 0,982 | 0,996 | 1,06 | 0,84 | -0,16 | 3,2E-02 |
| Q8C1B7 | Septin-11 OS=Mus musculus OX=10090 GN=Septin11 PE=1 SV=4 | 34 | 14 | 19 | 6 | 431 | 49,7 | 6,68 | 48,3 | 14 | 1,072 | 0,969 | 1,045 | 1,036 | 0,966 | 0,973 | 0,949 | 0,98 | 1,07 | 0,07 | 3,2E-02 |
| P27773 | Protein disulfide-isomerase A3 OS=Mus musculus OX=10090 GN=P | 47 | 22 | 41 | 22 | 505 | 56,6 | 6,21 | 95,3 | 22 | 1,095 | 1,055 | 1,042 | 1,098 | 1,001 | 1,019 | 1,027 | 1,05 | 1,05 | 0,05 | 3,2E-02 |
| Q8BFU3 | RING finger protein 214 OS=Mus musculus OX=10090 GN=Rnf214 P | 14 | 10 | 12 | 9 | 668 | 73,6 | 6,23 | 17,51 | 10 | 1,029 | 1,062 | 0,994 | 0,964 | 0,893 | 0,96 | 0,967 | 0,929 | 1,08 | 0,08 | 3,3E-02 |
| Q61548 | Clathrin coat assembly protein AP180 OS=Mus musculus OX=10090 | 15 | 13 | 28 | 13 | 901 | 91,8 | 4,88 | 46,07 | 13 | 0,859 | 0,885 | 0,966 | 0,99 | 0,999 | 0,99 | 1,122 | 1,106 | 0,88 | -0,12 | 3,3E-02 |
| Q3U7R1 | Extended synaptotagmin-1 OS=Mus musculus OX=10090 GN=Esyt1 | 1 | 1 | 1 | 1 | 1092 | 121,5 | 5,95 | 2,46 | 1 | 2,171 | 2,205 | 1,635 | 1,921 | 1,365 | 1,773 | 1,604 | 1,363 | 1,30 | 0,30 | 3,3E-02 |
| P97807 | Fumarate hydratase, mitochondrial OS=Mus musculus OX=10090 G | 23 | 14 | 27 | 14 | 507 | 54,3 | 9,04 | 60,08 | 14 | 0,906 | 0,954 | 0,951 | 0,972 | 1,026 | 0,96 | 1,003 | 1,017 | 0,94 | -0,06 | 3,3E-02 |
| Q3TLI0 | Trafficking protein particle complex subunit 10 OS=Mus musculus | 6 | 5 | 5 | 5 | 1259 | 141,5 | 5,96 | 8,82 | 5 | 1,186 | 1,085 | 1,099 | 1,085 | 1,14 | 1,413 | 1,265 | 1,315 | 0,87 | -0,13 | 3,4E-02 |
| O35343 | Importin subunit alpha-3 OS=Mus musculus OX=10090 GN=Kpna4 | 9 | 4 | 4 | 2 | 521 | 57,9 | 4,94 | 5,85 | 4 | 0,936 | 0,931 | 1,153 | 1,05 | 1,162 | 1,208 | 1,19 | 1,124 | 0,87 | -0,13 | 3,4E-02 |
| Q80Z38 | SH3 and multiple ankyrin repeat domains protein 2 OS=Mus muscu | 16 | 18 | 19 | 18 | 1476 | 158,9 | 7,18 | 24,89 | 18 | 1,212 | 1,154 | 1,089 | 1,086 | 1,004 | 1,098 | 1,027 | 0,994 | 1,10 | 0,10 | 3,4E-02 |
| P0DP27 | Calmodulin-2 OS=Mus musculus OX=10090 GN=Calm2 PE=1 SV=1 | 52 | 13 | 32 | 7 | 149 | 16,8 | 4,22 | 82,79 | 13 | 0,972 | 1,084 | 1,015 | 1,064 | 0,967 | 0,822 | 0,957 | 0,931 | 1,12 | 0,12 | 3,4E-02 |
| P60879 | Synaptosomal-associated protein 25 OS=Mus musculus OX=10090 | 68 | 20 | 89 | 2 | 206 | 23,3 | 4,77 | 206,75 | 20 | 1,284 | 1,633 | 1,08 | 1,152 | 0,927 | 0,987 | 0,848 | 0,991 | 1,37 | 0,37 | 3,4E-02 |
| Q9D855 | Cytochrome b-c1 complex subunit 7 OS=Mus musculus OX=10090 | 59 | 8 | 15 | 8 | 111 | 13,5 | 9,11 | 31 | 8 | 0,859 | 0,932 | 0,897 | 0,897 | 0,909 | 1,051 | 1,002 | 0,985 | 0,91 | -0,09 | 3,4E-02 |
| P07309 | Transthyretin OS=Mus musculus OX=10090 GN=Ttr PE=1 SV=1 | 4 | 1 | 1 | 1 | 147 | 15,8 | 6,16 | 1,81 | 1 | 0,731 | 0,841 | 0,844 | 0,786 | 0,808 | 1,068 | 1,003 | 1,009 | 0,82 | -0,18 | 3,4E-02 |
| A3KMP2 | Tetratricopeptide repeat protein 38 OS=Mus musculus OX=10090 | 3 | 1 | 1 | 1 | 465 | 52,2 | 6,3 | 0 | 1 | 0,811 | 0,754 | 0,823 | 0,819 | 0,831 | 0,949 | 0,857 | 0,947 | 0,89 | -0,11 | 3,4E-02 |
| Q9R118 | Serine protease HTRA1 OS=Mus musculus OX=10090 GN=Htra1 PE | 5 | 2 | 2 | 2 | 480 | 51,2 | 7,65 | 0 | 2 | 1,031 | 1,554 | 1,388 | 1,128 | 0,69 | 0,924 | 0,935 | 1,028 | 1,43 | 0,43 | 3,4E-02 |

| | | | | | | | | | | | | | | | | | | | | | |
|---|---|---|---|---|---|---|---|---|---|---|---|---|---|---|---|---|---|---|---|---|---|
| P70398 | Probable ubiquitin carboxyl-terminal hydrolase FAF-X OS=Mus mus | 9 | 18 | 21 | 18 | 2559 | 290,5 | 5,87 | 36,46 | 18 | 1,073 | 1,131 | 1,111 | 1,014 | 0,958 | 1,047 | 0,993 | 0,984 | 1,09 | 0,09 | 3,4E-02 |
| Q3UQS2 | Leucine-rich single-pass membrane protein 1 OS=Mus musculus OX | 5 | 2 | 2 | 2 | 128 | 14,2 | 5,48 | 4,33 | 2 | 1,07 | 1,359 | 1,151 | 0,855 | 0,778 | 0,954 | 0,709 | 0,647 | 1,44 | 0,44 | 3,4E-02 |
| P97370 | Sodium/potassium-transporting ATPase subunit beta-3 OS=Mus mu | 12 | 3 | 5 | 3 | 278 | 31,8 | 8,51 | 11,98 | 3 | 1,048 | 1,117 | 1,215 | 1,195 | 1,074 | 1,033 | 0,987 | 0,903 | 1,14 | 0,14 | 3,4E-02 |
| O35474 | EGF-like repeat and discoidin I-like domain-containing protein 3 OS | 9 | 4 | 4 | 4 | 480 | 53,7 | 7,58 | 7,19 | 4 | 1,315 | 1,272 | 1,118 | 1,14 | 0,919 | 1,17 | 0,969 | 0,99 | 1,20 | 0,20 | 3,4E-02 |
| Q80VC9 | Calmodulin-regulated spectrin-associated protein 3 OS=Mus muscu | 11 | 11 | 12 | 11 | 1252 | 135,1 | 8,43 | 12,72 | 11 | 1,303 | 1,024 | 1,197 | 1,163 | 0,992 | 1,026 | 1,036 | 0,995 | 1,16 | 0,16 | 3,5E-02 |
| Q80TQ2 | Ubiquitin carboxyl-terminal hydrolase CYLD OS=Mus musculus OX= | 9 | 7 | 9 | 7 | 952 | 106,5 | 5,63 | 21,83 | 7 | 0,922 | 1,024 | 1,009 | 0,944 | 1,005 | 1,059 | 1,103 | 1,089 | 0,92 | -0,08 | 3,5E-02 |
| Q64436 | Potassium-transporting ATPase alpha chain 1 OS=Mus musculus OX | 6 | 7 | 33 | 2 | 1033 | 113,9 | 5,88 | 69,09 | 7 | 1,005 | 1,115 | 1,122 | 1,121 | 0,896 | 0,863 | 0,994 | 1,049 | 1,15 | 0,15 | 3,5E-02 |
| Q80XI3 | Eukaryotic translation initiation factor 4 gamma 3 OS=Mus musculu | 8 | 10 | 12 | 7 | 1579 | 174,8 | 5,53 | 21,5 | 10 | 0,688 | 0,75 | 0,794 | 0,72 | 0,752 | 0,886 | 0,896 | 0,977 | 0,84 | -0,16 | 3,6E-02 |
| O70435 | Proteasome subunit alpha type-3 OS=Mus musculus OX=10090 GN | 24 | 6 | 7 | 6 | 255 | 28,4 | 5,44 | 12,73 | 6 | 0,927 | 0,914 | 0,894 | 0,935 | 0,937 | 0,995 | 0,943 | 0,967 | 0,96 | -0,04 | 3,6E-02 |
| O55142 | 60S ribosomal protein L35a OS=Mus musculus OX=10090 GN=Rpl3! | 20 | 3 | 3 | 3 | 110 | 12,5 | 10,89 | 1,77 | 3 | 0,555 | 0,798 | 0,904 | 0,882 | 1,185 | 0,967 | 0,998 | 1,003 | 0,76 | -0,24 | 3,6E-02 |
| P46097 | Synaptotagmin-2 OS=Mus musculus OX=10090 GN=Syt2 PE=1 SV=1 | 27 | 11 | 20 | 5 | 422 | 47,2 | 7,99 | 28,44 | 11 | 0,986 | 0,916 | 0,973 | 0,96 | 1,04 | 1,118 | 0,971 | 1,094 | 0,91 | -0,09 | 3,6E-02 |
| Q811S7 | Upstream-binding protein 1 OS=Mus musculus OX=10090 GN=Ubp | 5 | 2 | 2 | 2 | 540 | 60,2 | 6,27 | 2,61 | 2 | 1,279 | 0,967 | 1,12 | 1,002 | 0,955 | 0,653 | 0,855 | 0,879 | 1,31 | 0,31 | 3,6E-02 |
| Q8VDQ8 | NAD-dependent protein deacetylase sirtuin-2 OS=Mus musculus O) | 31 | 10 | 11 | 10 | 389 | 43,2 | 5,35 | 34,34 | 10 | 1,022 | 1,036 | 1,016 | 1,012 | 1,026 | 0,926 | 0,9 | 0,877 | 1,10 | 0,10 | 3,6E-02 |
| Q61102 | ATP-binding cassette sub-family B member 7, mitochondrial OS=Mu | 6 | 4 | 4 | 4 | 752 | 82,5 | 9,32 | 3 | 4 | 1,065 | 1,118 | 1,025 | 1,059 | 1,138 | 1,096 | 1,236 | 1,252 | 0,90 | -0,10 | 3,6E-02 |
| Q9D906 | Ubiquitin-like modifier-activating enzyme ATG7 OS=Mus musculus | 7 | 5 | 6 | 5 | 698 | 77,5 | 6,4 | 13,44 | 5 | 0,905 | 0,895 | 0,84 | 0,983 | 1,055 | 0,993 | 0,953 | 1,011 | 0,90 | -0,10 | 3,6E-02 |
| Q8K310 | Matrin-3 OS=Mus musculus OX=10090 GN=Matr3 PE=1 SV=1 | 5 | 4 | 4 | 4 | 846 | 94,6 | 6,25 | 6,43 | 4 | 0,706 | 0,758 | 0,786 | 0,577 | 0,981 | 1,075 | 0,77 | 0,868 | 0,77 | -0,23 | 3,7E-02 |
| Q9Z1Z0 | General vesicular transport factor p115 OS=Mus musculus OX=100! | 5 | 5 | 6 | 5 | 959 | 106,9 | 4,93 | 17,03 | 5 | 1,219 | 1,111 | 1,159 | 1,082 | 1,264 | 1,236 | 1,184 | 1,273 | 0,92 | -0,08 | 3,7E-02 |
| Q9WTU3 | Sodium channel protein type 8 subunit alpha OS=Mus musculus OX | 3 | 6 | 8 | 4 | 1978 | 225 | 6,25 | 18,63 | 6 | 0,927 | 0,951 | 0,873 | 0,971 | 1,094 | 1,074 | 1,017 | 0,95 | 0,90 | -0,10 | 3,7E-02 |
| O70439 | Syntaxin-7 OS=Mus musculus OX=10090 GN=Stx7 PE=1 SV=3 | 18 | 4 | 5 | 4 | 261 | 29,8 | 5,78 | 15,65 | 4 | 0,848 | 0,648 | 0,882 | 0,984 | 1,267 | 0,97 | 1,059 | 1,068 | 0,77 | -0,23 | 3,8E-02 |
| Q8BNY6 | Neuronal calcium sensor 1 OS=Mus musculus OX=10090 GN=Ncs1 I | 23 | 4 | 5 | 4 | 190 | 21,9 | 4,83 | 9,53 | 4 | 0,938 | 0,784 | 0,889 | 0,992 | 0,986 | 1,168 | 1,094 | 1,003 | 0,85 | -0,15 | 3,8E-02 |
| Q8JZS0 | Protein lin-7 homolog A OS=Mus musculus OX=10090 GN=Lin7a PE | 35 | 7 | 14 | 5 | 233 | 26 | 8,72 | 22,91 | 7 | 1,11 | 1,148 | 1,05 | 1,011 | 1,023 | 0,963 | 0,887 | 0,998 | 1,12 | 0,12 | 3,9E-02 |
| O35405 | Phospholipase D3 OS=Mus musculus OX=10090 GN=Pld3 PE=1 SV= | 7 | 3 | 3 | 3 | 488 | 54,4 | 6,52 | 10,65 | 3 | 0,985 | 0,998 | 0,996 | 0,937 | 0,821 | 0,846 | 0,947 | 0,936 | 1,10 | 0,10 | 3,9E-02 |
| O88665 | Bromodomain-containing protein 7 OS=Mus musculus OX=10090 G | 1 | 1 | 1 | 1 | 651 | 74 | 6,38 | 2,05 | 1 | 0,47 | 0,538 | 0,585 | 0,905 | 0,959 | 0,768 | 1,072 | 0,911 | 0,67 | -0,33 | 3,9E-02 |
| P54823 | Probable ATP-dependent RNA helicase DDX6 OS=Mus musculus OX | 10 | 4 | 4 | 4 | 483 | 54,2 | 8,66 | 6,32 | 4 | 0,922 | 0,771 | 0,82 | 0,898 | 0,997 | 1,131 | 0,987 | 0,907 | 0,85 | -0,15 | 3,9E-02 |
| Q3TY60 | Protein FAM131B OS=Mus musculus OX=10090 GN=Fam131b PE=1 | 20 | 5 | 6 | 5 | 332 | 35,9 | 4,45 | 13,59 | 5 | 0,727 | 0,896 | 0,94 | 0,997 | 1,05 | 0,993 | 1,078 | 1,092 | 0,85 | -0,15 | 3,9E-02 |
| Q9Z0V1 | Potassium voltage-gated channel subfamily D member 3 OS=Mus n | 4 | 2 | 3 | 2 | 655 | 73,4 | 8,25 | 4,62 | 2 | 0,801 | 0,667 | 0,525 | 0,765 | 0,905 | 1,122 | 1,279 | 0,783 | 0,67 | -0,33 | 3,9E-02 |
| Q8CCJ3 | E3 UFM1-protein ligase 1 OS=Mus musculus OX=10090 GN=Ufl1 PE | 1 | 1 | 1 | 1 | 793 | 89,5 | 6,67 | 0 | 1 | 0,918 | 1,27 | 1,455 | 1,23 | 0,774 | 1,105 | 0,674 | 0,797 | 1,45 | 0,45 | 3,9E-02 |
| Q80U63 | Mitofusin-2 OS=Mus musculus OX=10090 GN=Mfn2 PE=1 SV=3 | 19 | 12 | 20 | 12 | 757 | 86,1 | 6,77 | 41,06 | 12 | 1,149 | 1,214 | 1,151 | 1,039 | 0,983 | 1,021 | 1,043 | 1,075 | 1,10 | 0,10 | 4,0E-02 |
| P62858 | 40S ribosomal protein S28 OS=Mus musculus OX=10090 GN=Rps28 | 13 | 1 | 1 | 1 | 69 | 7,8 | 10,7 | 0 | 1 | 0,783 | 0,598 | 0,786 | 0,853 | 0,859 | 0,888 | 1,114 | 0,99 | 0,78 | -0,22 | 4,0E-02 |
| P35290 | Ras-related protein Rab-24 OS=Mus musculus OX=10090 GN=Rab2 | 4 | 1 | 1 | 1 | 203 | 23,1 | 6,23 | 2,13 | 1 | 0,692 | 0,811 | 0,799 | 0,598 | 0,855 | 1,377 | 0,9 | 1,122 | 0,68 | -0,32 | 4,0E-02 |
| Q2WF71 | Leucine-rich repeat and fibronectin type III domain-containing prot | 7 | 3 | 3 | 3 | 766 | 81,9 | 7,59 | 5,04 | 3 | 2,294 | 1,637 | 1,628 | 1,419 | 0,882 | 1,241 | 1,205 | 1,382 | 1,48 | 0,48 | 4,0E-02 |
| Q9CQH3 | NADH dehydrogenase [ubiquinone] 1 beta subcomplex subunit 5, r | 24 | 7 | 11 | 7 | 189 | 21,7 | 9,41 | 18,8 | 7 | 0,78 | 0,822 | 0,778 | 0,916 | 1,111 | 0,841 | 1,006 | 1,012 | 0,83 | -0,17 | 4,0E-02 |
| Q9WV92 | Band 4.1-like protein 3 OS=Mus musculus OX=10090 GN=Epb41l3 F | 44 | 36 | 48 | 6 | 929 | 103,3 | 5,31 | 106,7 | 36 | 1,071 | 1,077 | 0,982 | 1,076 | 0,966 | 0,882 | 0,928 | 1,028 | 1,11 | 0,11 | 4,1E-02 |
| Q8C190 | VPS9 domain-containing protein 1 OS=Mus musculus OX=10090 GN | 1 | 1 | 1 | 1 | 649 | 71,2 | 6,87 | 1,92 | 1 | 0,747 | 0,686 | 0,912 | 1,088 | 1,384 | 1,029 | 1,102 | 1,15 | 0,74 | -0,26 | 4,1E-02 |
| Q56A07 | Sodium channel subunit beta-2 OS=Mus musculus OX=10090 GN=S | 23 | 4 | 4 | 4 | 215 | 24,2 | 6,54 | 2,93 | 4 | 0,763 | 0,889 | 0,928 | 0,896 | 0,953 | 0,938 | 1,017 | 0,988 | 0,89 | -0,11 | 4,1E-02 |
| Q640R3 | Hepatocyte cell adhesion molecule OS=Mus musculus OX=10090 G | 13 | 4 | 4 | 4 | 418 | 46,3 | 9,42 | 7,23 | 4 | 0,965 | 0,975 | 1,003 | 0,935 | 1,012 | 0,993 | 1,066 | 1,099 | 0,93 | -0,07 | 4,1E-02 |
| A2RSJ4 | UHRF1-binding protein 1-like OS=Mus musculus OX=10090 GN=Uh | 4 | 4 | 4 | 4 | 1457 | 161,8 | 6,61 | 8,42 | 4 | 1,041 | 1,141 | 1,042 | 1,077 | 0,943 | 1,046 | 0,977 | 1,003 | 1,08 | 0,08 | 4,1E-02 |
| Q80X73 | Protein pelota homolog OS=Mus musculus OX=10090 GN=Pelo PE= | 9 | 3 | 3 | 3 | 385 | 43,3 | 5,99 | 1,79 | 3 | 1,358 | 1,047 | 1,268 | 0,925 | 0,844 | 0,873 | 0,895 | 0,937 | 1,30 | 0,30 | 4,1E-02 |
| O35129 | Prohibitin-2 OS=Mus musculus OX=10090 GN=Phb2 PE=1 SV=1 | 39 | 11 | 21 | 11 | 299 | 33,3 | 9,83 | 52,17 | 11 | 1,021 | 1,048 | 1,025 | 1,013 | 1,048 | 1,037 | 1,072 | 1,06 | 0,97 | -0,03 | 4,2E-02 |
| P61089 | Ubiquitin-conjugating enzyme E2 N OS=Mus musculus OX=10090 G | 55 | 7 | 8 | 7 | 152 | 17,1 | 6,57 | 15,76 | 7 | 1,385 | 1,315 | 1,161 | 0,979 | 0,882 | 1,059 | 0,898 | 0,981 | 1,27 | 0,27 | 4,2E-02 |
| Q9EPN1 | Neurobeachin OS=Mus musculus OX=10090 GN=Nbea PE=1 SV=1 | 11 | 26 | 31 | 23 | 2936 | 326,5 | 6,2 | 54,56 | 26 | 0,949 | 1,031 | 0,928 | 0,97 | 1,026 | 1,021 | 1,06 | 1,02 | 0,94 | -0,06 | 4,2E-02 |
| P62627 | Dynein light chain roadblock-type 1 OS=Mus musculus OX=10090 G | 23 | 2 | 3 | 2 | 96 | 11 | 7,25 | 9,57 | 2 | 1,005 | 0,989 | 1,04 | 1,073 | 1,031 | 1,232 | 1,253 | 1,145 | 0,88 | -0,12 | 4,2E-02 |
| Q8R164 | Valacyclovir hydrolase OS=Mus musculus OX=10090 GN=Bphl PE=1 | 32 | 8 | 15 | 8 | 291 | 32,8 | 8,94 | 11,98 | 8 | 0,917 | 0,935 | 0,874 | 0,918 | 1,141 | 1,004 | 1,018 | 0,937 | 0,89 | -0,11 | 4,3E-02 |
| P62874 | Guanine nucleotide-binding protein G(I)/G(S)/G(T) subunit beta-1 C | 39 | 11 | 39 | 6 | 340 | 37,4 | 6 | 76,42 | 11 | 0,74 | 0,77 | 0,921 | 0,968 | 1,032 | 0,942 | 1,059 | 0,995 | 0,84 | -0,16 | 4,3E-02 |
| P14211 | Calreticulin OS=Mus musculus OX=10090 GN=Calr PE=1 SV=1 | 33 | 12 | 19 | 12 | 416 | 48 | 4,49 | 42,62 | 12 | 0,806 | 0,837 | 0,94 | 1,013 | 1,057 | 0,959 | 1,073 | 1,063 | 0,87 | -0,13 | 4,3E-02 |
| Q80ZF8 | Adhesion G protein-coupled receptor B3 OS=Mus musculus OX=10( | 3 | 5 | 7 | 4 | 1522 | 171,2 | 7,02 | 14,1 | 5 | 1,335 | 0,991 | 1,146 | 1,198 | 0,921 | 0,988 | 1,038 | 0,956 | 1,20 | 0,20 | 4,4E-02 |
| Q8K1C0 | Protein angel homolog 2 OS=Mus musculus OX=10090 GN=Angel2 | 2 | 1 | 1 | 1 | 544 | 62,4 | 8,21 | 0 | 1 | 0,845 | 0,842 | 0,933 | 0,934 | 1,149 | 1,087 | 1,445 | 0,957 | 0,77 | -0,23 | 4,4E-02 |
| O08585 | Clathrin light chain A OS=Mus musculus OX=10090 GN=Clta PE=1 S\ | 28 | 10 | 16 | 10 | 235 | 25,6 | 4,58 | 33,01 | 10 | 0,74 | 0,8 | 0,826 | 0,975 | 1,039 | 0,879 | 1,019 | 1,057 | 0,84 | -0,16 | 4,4E-02 |
| O54990 | Prominin-1 OS=Mus musculus OX=10090 GN=Prom1 PE=1 SV=1 | 2 | 1 | 1 | 1 | 867 | 97,1 | 6,68 | 0 | 1 | 1,876 | 1,877 | 1,099 | 1,181 | 0,95 | 1,022 | 0,787 | 1,036 | 1,59 | 0,59 | 4,4E-02 |
| Q9Z0Y1 | Dynactin subunit 3 OS=Mus musculus OX=10090 GN=Dctn3 PE=1 S\ | 23 | 4 | 5 | 4 | 186 | 21 | 6,06 | 15,36 | 4 | 0,808 | 0,914 | 0,891 | 0,999 | 1,092 | 0,959 | 1,091 | 0,995 | 0,87 | -0,13 | 4,5E-02 |
| A2A699 | Protein FAM171A2 OS=Mus musculus OX=10090 GN=Fam171a2 PE | 6 | 4 | 5 | 4 | 822 | 87,4 | 8,05 | 7,34 | 4 | 1,269 | 1,23 | 1,131 | 0,972 | 0,822 | 0,994 | 0,953 | 1,027 | 1,21 | 0,21 | 4,5E-02 |
| Q6PIC6 | Sodium/potassium-transporting ATPase subunit alpha-3 OS=Mus m | 47 | 54 | 240 | 34 | 1013 | 111,6 | 5,41 | 557,53 | 54 | 0,864 | 0,91 | 0,939 | 0,985 | 1,023 | 1,003 | 1,026 | 0,953 | 0,92 | -0,08 | 4,5E-02 |
| Q9QY76 | Vesicle-associated membrane protein-associated protein B OS=Mu | 31 | 9 | 9 | 8 | 243 | 26,9 | 7,78 | 14,93 | 9 | 1,202 | 1,087 | 1,214 | 1,035 | 1,036 | 1,057 | 0,995 | 0,944 | 1,13 | 0,13 | 4,6E-02 |
| Q80W04 | Transmembrane and coiled-coil domains protein 2 OS=Mus muscul | 7 | 4 | 4 | 4 | 706 | 77 | 6,84 | 2,09 | 4 | 1,479 | 1,194 | 1,442 | 1,231 | 0,932 | 0,953 | 0,987 | 1,315 | 1,28 | 0,28 | 4,6E-02 |
| Q60715 | Prolyl 4-hydroxylase subunit alpha-1 OS=Mus musculus OX=10090 ( | 4 | 2 | 2 | 2 | 534 | 60,9 | 5,9 | 4,21 | 2 | 1,549 | 1,482 | 1,206 | 1,482 | 1,291 | 0,742 | 1,235 | 0,964 | 1,35 | 0,35 | 4,6E-02 |
| P68037 | Ubiquitin-conjugating enzyme E2 L3 OS=Mus musculus OX=10090 ( | 18 | 4 | 4 | 4 | 154 | 17,9 | 8,51 | 7,86 | 4 | 0,805 | 0,941 | 0,916 | 1,061 | 1,047 | 1,014 | 1,139 | 1,141 | 0,86 | -0,14 | 4,6E-02 |
| Q8CGU1 | Calcium-binding and coiled-coil domain-containing protein 1 OS=M | 10 | 6 | 6 | 6 | 691 | 77,2 | 4,82 | 8,38 | 6 | 1,172 | 0,914 | 1,068 | 1,229 | 1,447 | 1,169 | 1,416 | 1,291 | 0,82 | -0,18 | 4,6E-02 |
| Q62425 | Cytochrome c oxidase subunit NDUFA4 OS=Mus musculus OX=100! | 56 | 7 | 12 | 7 | 82 | 9,3 | 9,52 | 28,48 | 7 | 0,931 | 1,025 | 0,916 | 0,979 | 1,035 | 1,099 | 1,021 | 1,012 | 0,92 | -0,08 | 4,6E-02 |
| P63248 | cAMP-dependent protein kinase inhibitor alpha OS=Mus musculus | 50 | 2 | 2 | 2 | 76 | 8 | 4,54 | 7,27 | 2 | 1,275 | 1,279 | 1,364 | 1,191 | 0,988 | 1,267 | 1,086 | 1,089 | 1,15 | 0,15 | 4,7E-02 |
| Q9Z0X1 | Apoptosis-inducing factor 1, mitochondrial OS=Mus musculus OX=1 | 16 | 10 | 12 | 10 | 612 | 66,7 | 9,17 | 21,5 | 10 | 0,786 | 0,863 | 0,884 | 0,886 | 0,897 | 0,974 | 0,904 | 1,022 | 0,90 | -0,10 | 4,7E-02 |
| Q505F5 | Leucine-rich repeat-containing protein 47 OS=Mus musculus OX=1( | 17 | 8 | 9 | 8 | 581 | 63,6 | 8,1 | 23,25 | 8 | 1,156 | 1,041 | 1,11 | 0,993 | 0,899 | 1,029 | 0,876 | 0,988 | 1,13 | 0,13 | 4,8E-02 |
| Q60932 | Voltage-dependent anion-selective channel protein 1 OS=Mus mus | 65 | 14 | 33 | 13 | 296 | 32,3 | 8,43 | 80,7 | 14 | 0,881 | 0,944 | 1,007 | 1,061 | 1,069 | 1,094 | 1,129 | 1,035 | 0,90 | -0,10 | 4,8E-02 |

| | | | | | | | | | | | | | | | | | | | | | |
|---|---|---|---|---|---|---|---|---|---|---|---|---|---|---|---|---|---|---|---|---|---|
| Q8K010 | 5-oxoprolinase OS=Mus musculus OX=10090 GN=Oplah PE=1 SV=1 | 6 | 7 | 8 | 7 | 1288 | 137,5 | 6,28 | 14,1 | 7 | 1,257 | 1,187 | 1,139 | 1,119 | 1,011 | 1,153 | 0,993 | 1,075 | 1,11 | 0,11 | 4,8E-02 |
| Q8VE95 | UPF0598 protein C8orf82 homolog OS=Mus musculus OX=10090 Pl | 8 | 1 | 1 | 1 | 218 | 24,3 | 8,7 | 3,25 | 1 | 1,575 | 1,513 | 1,429 | 1,215 | 0,82 | 1,346 | 1,102 | 1,143 | 1,30 | 0,30 | 4,9E-02 |
| P62897 | Cytochrome c, somatic OS=Mus musculus OX=10090 GN=Cycs PE=1 | 49 | 6 | 13 | 6 | 105 | 11,6 | 9,58 | 18,37 | 6 | 0,882 | 0,723 | 0,975 | 0,906 | 1,051 | 1,051 | 1,063 | 0,929 | 0,85 | -0,15 | 5,0E-02 |
| P05202 | Aspartate aminotransferase, mitochondrial OS=Mus musculus OX= | 54 | 24 | 49 | 24 | 430 | 47,4 | 9 | 131,36 | 24 | 0,957 | 0,929 | 0,979 | 0,941 | 1,081 | 0,984 | 0,991 | 0,997 | 0,94 | -0,06 | 5,0E-02 |
| Q80XR2 | Calcium-transporting ATPase type 2C member 1 OS=Mus musculus | 4 | 3 | 3 | 3 | 918 | 100,2 | 6,83 | 6,75 | 3 | 1,181 | 1,356 | 1,278 | 1,01 | 1,002 | 1,073 | 1,009 | 0,986 | 1,19 | 0,19 | 5,0E-02 |
| Q8K4K6 | Pantothenate kinase 1 OS=Mus musculus OX=10090 GN=Pank1 PE= | 2 | 1 | 1 | 1 | 548 | 60,1 | 8,03 | 0 | 1 | 0,609 | 0,688 | 0,617 | 0,814 | 0,752 | 1,055 | 0,945 | 0,799 | 0,77 | -0,23 | 5,0E-02 |
| E9Q8I9 | Protein furry homolog OS=Mus musculus OX=10090 GN=Fry PE=1 S | 2 | 6 | 7 | 6 | 3020 | 338,9 | 6,01 | 8,43 | 6 | 1,465 | 1,334 | 1,364 | 1,17 | 0,956 | 1,211 | 1,211 | 1,114 | 1,19 | 0,19 | 5,0E-02 |
| P01831 | Thy-1 membrane glycoprotein OS=Mus musculus OX=10090 GN=Th | 32 | 5 | 22 | 5 | 162 | 18,1 | 8,97 | 55,28 | 5 | 1,026 | 1,036 | 1,043 | 0,995 | 1,075 | 1,113 | 1,038 | 1,06 | 0,96 | -0,04 | 5,0E-02 |
| Q64674 | Spermidine synthase OS=Mus musculus OX=10090 GN=Srm PE=1 S | 5 | 1 | 1 | 1 | 302 | 34 | 5,5 | 2,15 | 1 | 1,662 | 1,227 | 1,695 | 1,379 | 1,194 | 1,124 | 1,335 | 1,077 | 1,26 | 0,26 | 5,0E-02 |
| P60229 | Eukaryotic translation initiation factor 3 subunit E OS=Mus musculu | 4 | 2 | 3 | 2 | 445 | 52,2 | 6,04 | 6,65 | 2 | 0,647 | 0,909 | 0,795 | 0,905 | 0,859 | 1,128 | 1,031 | 1,058 | 0,80 | -0,20 | 5,0E-02 |
| Q6PGB6 | N-alpha-acetyltransferase 50 OS=Mus musculus OX=10090 GN=Naa | 14 | 2 | 2 | 2 | 169 | 19,4 | 8,81 | 2,93 | 2 | 1,36 | 1,262 | 1,237 | 1,21 | 0,978 | 1,002 | 1,098 | 1,27 | 1,17 | 0,17 | 5,1E-02 |
| Q9D6M3 | Mitochondrial glutamate carrier 1 OS=Mus musculus OX=10090 GN | 22 | 7 | 17 | 5 | 323 | 34,6 | 9,09 | 36,79 | 7 | 0,769 | 0,91 | 0,892 | 0,961 | 1,058 | 0,991 | 1,036 | 0,928 | 0,88 | -0,12 | 5,2E-02 |
| Q9CPQ1 | Cytochrome c oxidase subunit 6C OS=Mus musculus OX=10090 GN= | 38 | 7 | 10 | 7 | 76 | 8,5 | 10,14 | 17,14 | 7 | 1,202 | 1,127 | 1,002 | 0,858 | 0,902 | 0,873 | 0,808 | 0,854 | 1,22 | 0,22 | 5,2E-02 |
| Q64105 | Sepiapterin reductase OS=Mus musculus OX=10090 GN=Spr PE=1 S | 30 | 6 | 7 | 6 | 261 | 27,9 | 5,74 | 22,33 | 6 | 1,029 | 1,116 | 1,122 | 0,996 | 0,911 | 1,02 | 0,958 | 0,994 | 1,10 | 0,10 | 5,2E-02 |
| O70311 | Glycylpeptide N-tetradecanoyltransferase 2 OS=Mus musculus OX= | 3 | 2 | 2 | 2 | 529 | 60,4 | 7,8 | 1,65 | 2 | 1,152 | 1,073 | 1,116 | 0,977 | 1,051 | 0,911 | 0,9 | 0,737 | 1,20 | 0,20 | 5,2E-02 |
| Q8C7X2 | ER membrane protein complex subunit 1 OS=Mus musculus OX=10 | 1 | 1 | 1 | 1 | 997 | 111,5 | 7,43 | 3,79 | 1 | 1,877 | 1,698 | 1,215 | 1,16 | 0,811 | 1,33 | 0,774 | 0,93 | 1,55 | 0,55 | 5,3E-02 |
| P35235 | Tyrosine-protein phosphatase non-receptor type 11 OS=Mus musc | 10 | 5 | 5 | 5 | 597 | 68,4 | 7,3 | 10,41 | 5 | 0,936 | 1,07 | 1,119 | 1,098 | 0,868 | 0,976 | 0,969 | 0,947 | 1,12 | 0,12 | 5,3E-02 |
| Q03173 | Protein enabled homolog OS=Mus musculus OX=10090 GN=Enah P | 2 | 1 | 1 | 1 | 802 | 85,8 | 7,72 | 2,69 | 1 | 0,905 | 0,563 | 0,961 | 1,031 | 0,998 | 1,243 | 1,328 | 1,112 | 0,74 | -0,26 | 5,3E-02 |
| Q5M8N4 | Epimerase family protein SDR39U1 OS=Mus musculus OX=10090 G | 3 | 1 | 1 | 1 | 293 | 31,4 | 9,31 | 2,28 | 1 | 0,482 | 0,614 | 0,687 | 0,634 | 0,75 | 1,03 | 0,751 | 0,711 | 0,75 | -0,25 | 5,3E-02 |
| Q6P1B1 | Xaa-Pro aminopeptidase 1 OS=Mus musculus OX=10090 GN=Xpnpe | 8 | 5 | 6 | 5 | 623 | 69,5 | 5,54 | 12,98 | 5 | 1,403 | 1,32 | 1,193 | 1,274 | 1,148 | 1,171 | 1,246 | 1,149 | 1,10 | 0,10 | 5,3E-02 |
| Q80TK0 | AP2-interacting clathrin-endocytosis protein OS=Mus musculus OX= | 6 | 6 | 6 | 6 | 1239 | 134 | 6,61 | 11,07 | 6 | 1,176 | 1,195 | 1,094 | 1,101 | 0,974 | 1,091 | 1,004 | 1,104 | 1,09 | 0,09 | 5,4E-02 |
| P29595 | NEDD8 OS=Mus musculus OX=10090 GN=Nedd8 PE=1 SV=2 | 17 | 1 | 2 | 1 | 81 | 9 | 7,25 | 5,59 | 1 | 0,849 | 0,945 | 0,736 | 0,982 | 0,947 | 1,156 | 1,01 | 1,078 | 0,84 | -0,16 | 5,4E-02 |
| Q7TSQ8 | Pyruvate dehydrogenase phosphatase regulatory subunit, mitocho | 10 | 8 | 10 | 8 | 878 | 99,2 | 6,35 | 14,63 | 8 | 1,09 | 1,075 | 1,117 | 1,125 | 1,011 | 1,017 | 1,102 | 1,049 | 1,05 | 0,05 | 5,4E-02 |
| Q9ERA0 | Alpha-globin transcription factor CP2 OS=Mus musculus OX=10090 | 3 | 2 | 2 | 2 | 502 | 57 | 5,54 | 0 | 2 | 1,133 | 0,907 | 0,774 | 1,179 | 1,051 | 2,348 | 1,632 | 1,656 | 0,60 | -0,40 | 5,4E-02 |
| P56213 | FAD-linked sulfhydryl oxidase ALR OS=Mus musculus OX=10090 GN | 26 | 4 | 5 | 4 | 198 | 22,9 | 7,93 | 7,2 | 4 | 0,802 | 0,921 | 0,946 | 0,99 | 1,159 | 1,37 | 1,001 | 1,024 | 0,80 | -0,20 | 5,4E-02 |
| Q61097 | Kinase suppressor of Ras 1 OS=Mus musculus OX=10090 GN=Ksr1 F | 7 | 5 | 5 | 5 | 873 | 96,7 | 8,54 | 8,11 | 5 | 0,994 | 0,942 | 0,876 | 0,816 | 0,989 | 1,046 | 0,963 | 1,094 | 0,89 | -0,11 | 5,4E-02 |
| Q4PJX1 | Protein odr-4 homolog OS=Mus musculus OX=10090 GN=Odr4 PE= | 3 | 1 | 1 | 1 | 447 | 50 | 6,14 | 0 | 1 | 1,186 | 1,236 | 1,081 | 0,868 | 0,401 | 1,051 | 0,748 | 0,679 | 1,52 | 0,52 | 5,4E-02 |
| P43006 | Excitatory amino acid transporter 2 OS=Mus musculus OX=10090 G | 16 | 9 | 32 | 9 | 572 | 62 | 6,68 | 48,76 | 9 | 0,833 | 0,78 | 0,79 | 0,923 | 1,178 | 0,821 | 1,037 | 1,066 | 0,81 | -0,19 | 5,5E-02 |
| Q8BH66 | Atlastin-1 OS=Mus musculus OX=10090 GN=Atl1 PE=1 SV=1 | 19 | 11 | 13 | 11 | 558 | 63,3 | 6,49 | 17,99 | 11 | 0,994 | 0,992 | 1,034 | 0,99 | 1,093 | 1,058 | 1,06 | 1,003 | 0,95 | -0,05 | 5,5E-02 |
| P54285 | Voltage-dependent L-type calcium channel subunit beta-3 OS=Mus | 20 | 7 | 11 | 5 | 484 | 54,5 | 6,27 | 13,6 | 7 | 1,186 | 1,063 | 1,058 | 1,004 | 0,864 | 1,006 | 0,833 | 1,021 | 1,16 | 0,16 | 5,5E-02 |
| O88643 | Serine/threonine-protein kinase PAK 1 OS=Mus musculus OX=1009 | 29 | 14 | 22 | 8 | 545 | 60,7 | 5,74 | 41,31 | 14 | 1,126 | 1,136 | 0,96 | 1,008 | 0,886 | 0,945 | 0,94 | 0,994 | 1,12 | 0,12 | 5,5E-02 |
| Q8R1Q8 | Cytoplasmic dynein 1 light intermediate chain 1 OS=Mus musculus | 28 | 11 | 14 | 11 | 523 | 56,6 | 6,42 | 34,73 | 11 | 0,929 | 0,925 | 0,948 | 0,969 | 0,974 | 1,061 | 1,064 | 0,957 | 0,93 | -0,07 | 5,5E-02 |
| Q8CHW4 | Translation initiation factor eIF-2B subunit epsilon OS=Mus muscul | 3 | 2 | 2 | 2 | 717 | 80 | 5,07 | 0 | 2 | 0,533 | 0,682 | 0,852 | 0,814 | 1,023 | 0,91 | 0,795 | 1,159 | 0,74 | -0,26 | 5,6E-02 |
| O35098 | Dihydropyrimidinase-related protein 4 OS=Mus musculus OX=1009 | 36 | 14 | 33 | 13 | 572 | 61,9 | 6,98 | 101,54 | 14 | 1,096 | 1,05 | 1,094 | 1,104 | 0,912 | 0,959 | 1,073 | 1,036 | 1,09 | 0,09 | 5,6E-02 |
| Q8R0F8 | Acylpyruvase FAHD1, mitochondrial OS=Mus musculus OX=10090 ( | 3 | 1 | 2 | 1 | 227 | 25,2 | 7,69 | 3,91 | 1 | 0,963 | 0,842 | 1,003 | 1,014 | 1,129 | 1,223 | 1,019 | 1,034 | 0,87 | -0,13 | 5,6E-02 |
| Q9QXE2 | DNA polymerase lambda OS=Mus musculus OX=10090 GN=Poll PE= | 2 | 1 | 1 | 1 | 573 | 62,9 | 8 | 1,8 | 1 | 1,134 | 1,221 | 0,941 | 1,074 | 1,049 | 0,906 | 0,748 | 0,854 | 1,23 | 0,23 | 5,6E-02 |
| A2ALS5 | Rap1 GTPase-activating protein 1 OS=Mus musculus OX=10090 GN= | 12 | 8 | 8 | 7 | 663 | 73,4 | 5,87 | 18,79 | 8 | 1,197 | 1,08 | 1,048 | 0,983 | 1,001 | 0,869 | 0,944 | 0,986 | 1,13 | 0,13 | 5,6E-02 |
| Q9DC70 | NADH dehydrogenase [ubiquinone] iron-sulfur protein 7, mitochon | 18 | 5 | 10 | 5 | 224 | 24,7 | 9,92 | 13,18 | 5 | 0,973 | 0,946 | 0,79 | 0,87 | 1,049 | 0,983 | 0,966 | 1,006 | 0,89 | -0,11 | 5,6E-02 |
| P85094 | Isochorismatase domain-containing protein 2A OS=Mus musculus ( | 5 | 1 | 2 | 1 | 206 | 22,4 | 8,02 | 4,67 | 1 | 1,1 | 0,866 | 0,865 | 0,91 | 0,867 | 0,565 | 0,732 | 0,778 | 1,27 | 0,27 | 5,6E-02 |
| P52196 | Thiosulfate sulfurtransferase OS=Mus musculus OX=10090 GN=Tst | 23 | 5 | 6 | 5 | 297 | 33,4 | 7,85 | 9,33 | 5 | 1,075 | 0,959 | 0,964 | 0,914 | 1,004 | 1,093 | 1,108 | 1,106 | 0,91 | -0,09 | 5,6E-02 |
| Q9WUM4 | Coronin-1C OS=Mus musculus OX=10090 GN=Coro1c PE=1 SV=2 | 26 | 15 | 16 | 13 | 474 | 53,1 | 7,08 | 24,16 | 15 | 0,825 | 0,857 | 0,938 | 1,011 | 1,029 | 0,996 | 1,066 | 0,976 | 0,89 | -0,11 | 5,7E-02 |
| O08532-5 | Isoform 2E of Voltage-dependent calcium channel subunit alpha-2/ | 27 | 26 | 39 | 26 | 1091 | 123,2 | 5,27 | 84,1 | 26 | 0,962 | 0,964 | 0,904 | 0,987 | 1,052 | 0,957 | 1,025 | 1,05 | 0,93 | -0,07 | 5,7E-02 |
| Q9CPV3 | 39S ribosomal protein L42, mitochondrial OS=Mus musculus OX=10 | 9 | 1 | 1 | 1 | 142 | 16,5 | 9,19 | 0 | 1 | 4,583 | 3,142 | 3,445 | 3,44 | 2,708 | 3,148 | 2,692 | 2,908 | 1,28 | 0,28 | 5,7E-02 |
| Q9D0M3 | Cytochrome c1, heme protein, mitochondrial OS=Mus musculus OX | 25 | 6 | 12 | 6 | 325 | 35,3 | 9,16 | 28,08 | 6 | 0,957 | 0,911 | 0,975 | 1,032 | 1,101 | 0,992 | 1,077 | 1,032 | 0,92 | -0,08 | 5,7E-02 |
| O08579 | Emerin OS=Mus musculus OX=10090 GN=Emd PE=1 SV=1 | 10 | 2 | 2 | 2 | 259 | 29,4 | 5,01 | 2,6 | 2 | 1,141 | 1,245 | 1,045 | 1,031 | 0,927 | 0,953 | 1,083 | 0,877 | 1,16 | 0,16 | 5,7E-02 |
| Q80UJ7 | Rab3 GTPase-activating protein catalytic subunit OS=Mus musculus | 17 | 15 | 17 | 15 | 981 | 110,1 | 5,73 | 22,77 | 15 | 1,062 | 1,059 | 0,997 | 1,058 | 1,137 | 1,063 | 1,089 | 1,093 | 0,95 | -0,05 | 5,7E-02 |
| P97355 | Spermine synthase OS=Mus musculus OX=10090 GN=Sms PE=1 SV= | 7 | 3 | 3 | 3 | 366 | 41,3 | 5,06 | 2,08 | 3 | 0,982 | 1,129 | 0,849 | 1,088 | 0,848 | 0,885 | 0,892 | 0,81 | 1,18 | 0,18 | 5,7E-02 |
| Q9CQR6 | Serine/threonine-protein phosphatase 6 catalytic subunit OS=Mus | 13 | 3 | 4 | 3 | 305 | 35,1 | 5,69 | 4,72 | 3 | 1,289 | 1,297 | 1,196 | 0,991 | 0,863 | 1,135 | 0,918 | 0,991 | 1,22 | 0,22 | 5,8E-02 |
| P62823 | Ras-related protein Rab-3C OS=Mus musculus OX=10090 GN=Rab3 | 47 | 12 | 20 | 6 | 227 | 25,9 | 5,24 | 47,09 | 12 | 0,921 | 0,857 | 0,95 | 0,99 | 0,949 | 1,031 | 1,077 | 1,022 | 0,91 | -0,09 | 5,8E-02 |
| Q99KR3 | Endoribonuclease LACTB2 OS=Mus musculus OX=10090 GN=Lactb2 | 4 | 1 | 1 | 1 | 288 | 32,7 | 6,33 | 2,1 | 1 | 1,518 | 1,211 | 1,204 | 1,4 | 0,795 | 1,342 | 0,557 | 0,939 | 1,47 | 0,47 | 5,8E-02 |
| Q64735 | Complement component receptor 1-like protein OS=Mus musculus | 2 | 1 | 1 | 1 | 483 | 53,7 | 6,65 | 0 | 1 | 0,01 | 0,629 | 0,76 | 0,848 | 1,149 | 0,728 | 1,2 | 1,288 | 0,51 | -0,49 | 5,8E-02 |
| Q3TCN2 | Putative phospholipase B-like 2 OS=Mus musculus OX=10090 GN=F | 4 | 2 | 2 | 2 | 594 | 66,2 | 6,13 | 0 | 2 | 0,97 | 1,064 | 0,813 | 1,054 | 1,145 | 1,213 | 1,256 | 1,806 | 0,72 | -0,28 | 5,8E-02 |
| P57746 | V-type proton ATPase subunit D OS=Mus musculus OX=10090 GN=. | 30 | 6 | 8 | 6 | 247 | 28,4 | 9,45 | 19,3 | 6 | 0,949 | 0,925 | 0,914 | 0,994 | 0,956 | 1,01 | 1,008 | 1,026 | 0,95 | -0,05 | 5,8E-02 |
| P56695 | Wolframin OS=Mus musculus OX=10090 GN=Wfs1 PE=1 SV=1 | 5 | 4 | 4 | 4 | 890 | 100,5 | 7,97 | 6,85 | 4 | 1,095 | 1,061 | 1,227 | 1,081 | 0,988 | 0,935 | 1,099 | 0,868 | 1,15 | 0,15 | 5,9E-02 |
| Q6ZQK5 | Arf-GAP with coiled-coil, ANK repeat and PH domain-containing pro | 13 | 11 | 11 | 11 | 770 | 87,2 | 6,68 | 18,75 | 11 | 1,188 | 1,034 | 1,095 | 1,081 | 1,066 | 0,931 | 0,957 | 1,027 | 1,10 | 0,10 | 5,9E-02 |
| Q3UHL1 | CaM kinase-like vesicle-associated protein OS=Mus musculus OX=1 | 21 | 8 | 12 | 8 | 512 | 54,8 | 5,54 | 22,49 | 8 | 0,918 | 0,942 | 0,965 | 0,985 | 0,989 | 1,049 | 0,98 | 0,991 | 0,95 | -0,05 | 5,9E-02 |
| P61294 | Ras-related protein Rab-6B OS=Mus musculus OX=10090 GN=Rab6 | 30 | 8 | 12 | 4 | 208 | 23,4 | 5,53 | 23,31 | 8 | 0,948 | 0,998 | 0,986 | 1,143 | 0,956 | 0,803 | 0,903 | 0,915 | 1,14 | 0,14 | 6,0E-02 |
| Q8BIZ1 | Ankyrin repeat and sterile alpha motif domain-containing protein 1 | 7 | 9 | 13 | 9 | 1259 | 139 | 6,29 | 24,17 | 9 | 1,282 | 1,275 | 1,1 | 1,167 | 1,109 | 1,147 | 1,067 | 1,028 | 1,11 | 0,11 | 6,0E-02 |
| Q99JY0 | Trifunctional enzyme subunit beta, mitochondrial OS=Mus musculu | 9 | 4 | 5 | 4 | 475 | 51,4 | 9,38 | 13,29 | 4 | 0,955 | 0,938 | 1,056 | 0,963 | 1,031 | 1,115 | 1,025 | 1,052 | 0,93 | -0,07 | 6,0E-02 |
| O35609 | Secretory carrier-associated membrane protein 3 OS=Mus musculu | 17 | 4 | 4 | 4 | 349 | 38,4 | 7,64 | 12,35 | 4 | 1,278 | 1,157 | 1,22 | 1,014 | 0,868 | 0,989 | 0,857 | 1,144 | 1,21 | 0,21 | 6,0E-02 |
| Q62348 | Translin OS=Mus musculus OX=10090 GN=Tsn PE=1 SV=1 | 11 | 2 | 2 | 2 | 228 | 26,2 | 6,44 | 9,04 | 2 | 1,005 | 1,105 | 1,001 | 1,149 | 0,878 | 1,02 | 0,875 | 0,996 | 1,13 | 0,13 | 6,0E-02 |
| G3XA57 | Rab11 family-interacting protein 2 OS=Mus musculus OX=10090 GN | 4 | 2 | 2 | 2 | 512 | 58,2 | 9,45 | 5,66 | 2 | 1,033 | 1,081 | 1,038 | 1,07 | 0,943 | 0,977 | 0,95 | 1,067 | 1,07 | 0,07 | 6,1E-02 |

| | | | | | | | | | | | | | | | | | | | |
|---|---|---|---|---|---|---|---|---|---|---|---|---|---|---|---|---|---|---|---|
| Q8BHK2 | Sodium channel subunit beta-3 OS=Mus musculus OX=10090 GN=S | 10 | 2 | 2 | 2 | 215 | 24,8 | 4,74 | 1,76 | 2 | 1,224 | 1,382 | 1,138 | 1,266 | 0,968 | 1,103 | 1,15 | 1,165 | 1,14 | 0,14 | 6,1E-02 |
| Q5SSL4 | Active breakpoint cluster region-related protein OS=Mus musculus | 14 | 13 | 18 | 12 | 859 | 97,6 | 6,58 | 48,41 | 13 | 0,936 | 0,893 | 0,977 | 0,989 | 0,959 | 1,044 | 1,04 | 1,022 | 0,93 | -0,07 | 6,1E-02 |
| Q9D1X0 | Nucleolar protein 3 OS=Mus musculus OX=10090 GN=Nol3 PE=1 SV | 5 | 1 | 1 | 1 | 220 | 24,6 | 4,07 | 1,64 | 1 | 0,404 | 0,449 | 0,619 | 0,666 | 0,687 | 0,816 | 0,723 | 0,614 | 0,75 | -0,25 | 6,1E-02 |
| P70704 | Phospholipid-transporting ATPase IA OS=Mus musculus OX=10090 | 17 | 17 | 20 | 16 | 1164 | 131,3 | 6,84 | 50,39 | 17 | 0,872 | 0,939 | 0,958 | 0,958 | 0,939 | 1,058 | 1,084 | 0,998 | 0,91 | -0,09 | 6,1E-02 |
| P10711 | Transcription elongation factor A protein 1 OS=Mus musculus OX=1 | 9 | 3 | 3 | 3 | 301 | 33,9 | 8,38 | 2,17 | 3 | 0,975 | 1,082 | 0,985 | 1,036 | 0,94 | 0,64 | 0,981 | 0,726 | 1,24 | 0,24 | 6,1E-02 |
| P16546-2 | Isoform 2 of Spectrin alpha chain, non-erythrocytic 1 OS=Mus musc | 51 | 134 | 239 | 1 | 2452 | 282,2 | 5,31 | 615,4 | 134 | 1,183 | 1,004 | 1,097 | 1,114 | 1,055 | 0,947 | 1,011 | 0,988 | 1,10 | 0,10 | 6,1E-02 |
| P99028 | Cytochrome b-c1 complex subunit 6, mitochondrial OS=Mus muscu | 16 | 2 | 4 | 2 | 89 | 10,4 | 4,87 | 4,14 | 2 | 0,7 | 0,743 | 0,919 | 0,994 | 1,116 | 0,981 | 1,006 | 0,967 | 0,82 | -0,18 | 6,2E-02 |
| P16330 | 2',3'-cyclic-nucleotide 3'-phosphodiesterase OS=Mus musculus OX= | 38 | 24 | 47 | 24 | 420 | 47,1 | 8,97 | 101,41 | 24 | 1,021 | 1,023 | 1,03 | 1,117 | 1,06 | 0,884 | 0,924 | 0,848 | 1,13 | 0,13 | 6,2E-02 |
| Q80WM4 | Hyaluronan and proteoglycan link protein 4 OS=Mus musculus OX= | 8 | 3 | 4 | 3 | 400 | 42,8 | 8,85 | 9,09 | 3 | 0,903 | 0,851 | 1,004 | 1,018 | 1,07 | 1,09 | 1,057 | 0,984 | 0,90 | -0,10 | 6,2E-02 |
| Q80VL1 | Tudor and KH domain-containing protein OS=Mus musculus OX=10 | 18 | 8 | 8 | 8 | 560 | 62,1 | 4,94 | 11,78 | 8 | 0,97 | 1,107 | 0,871 | 1,028 | 1,244 | 1,152 | 1,092 | 1,068 | 0,87 | -0,13 | 6,2E-02 |
| Q8K2C6 | NAD-dependent protein deacylase sirtuin-5, mitochondrial OS=Mu | 19 | 5 | 5 | 5 | 310 | 34,1 | 8,53 | 9,07 | 5 | 1,371 | 1,351 | 1,435 | 1,329 | 1,075 | 1,188 | 1,36 | 1,27 | 1,12 | 0,12 | 6,2E-02 |
| Q9EQF5 | Dihydropyrimidinase OS=Mus musculus OX=10090 GN=Dpys PE=1 S | 4 | 2 | 2 | 2 | 519 | 56,7 | 7,2 | 4,6 | 2 | 1,241 | 1,085 | 0,929 | 1,102 | 0,804 | 0,982 | 0,993 | 0,852 | 1,20 | 0,20 | 6,2E-02 |
| Q8C0C7 | Phenylalanine--tRNA ligase alpha subunit OS=Mus musculus OX=10 | 2 | 1 | 1 | 1 | 508 | 57,6 | 8,28 | 0 | 1 | 1,646 | 1,354 | 0,911 | 0,722 | 0,663 | 0,353 | 0,572 | 0,881 | 1,88 | 0,88 | 6,2E-02 |
| Q91YP0 | L-2-hydroxyglutarate dehydrogenase, mitochondrial OS=Mus musc | 13 | 5 | 5 | 5 | 464 | 50,9 | 8,29 | 8,14 | 5 | 1,063 | 1,057 | 1,154 | 1,124 | 1,032 | 0,866 | 1,024 | 1,04 | 1,11 | 0,11 | 6,3E-02 |
| O08992 | Syntenin-1 OS=Mus musculus OX=10090 GN=Sdcbp PE=1 SV=1 | 7 | 2 | 2 | 2 | 299 | 32,4 | 7,15 | 0 | 2 | 1,172 | 0,843 | 1,103 | 1,078 | 0,819 | 0,98 | 0,744 | 0,862 | 1,23 | 0,23 | 6,3E-02 |
| P10126 | Elongation factor 1-alpha 1 OS=Mus musculus OX=10090 GN=Eef1a | 24 | 12 | 23 | 6 | 462 | 50,1 | 9,01 | 54,95 | 12 | 0,771 | 0,831 | 0,934 | 0,978 | 1,072 | 0,926 | 1,117 | 0,981 | 0,86 | -0,14 | 6,3E-02 |
| Q9CZ04 | COP9 signalosome complex subunit 7a OS=Mus musculus OX=1009 | 21 | 6 | 8 | 6 | 275 | 30,2 | 7,87 | 13,31 | 6 | 1,055 | 1,043 | 1,131 | 1,065 | 0,921 | 1,069 | 0,989 | 0,987 | 1,08 | 0,08 | 6,4E-02 |
| P29533 | Vascular cell adhesion protein 1 OS=Mus musculus OX=10090 GN=V | 11 | 6 | 7 | 6 | 739 | 81,3 | 5,3 | 19 | 6 | 1,103 | 0,978 | 1,088 | 1,135 | 0,954 | 0,947 | 1,048 | 0,982 | 1,09 | 0,09 | 6,4E-02 |
| Q9CXT8 | Mitochondrial-processing peptidase subunit beta OS=Mus musculu | 3 | 2 | 3 | 1 | 489 | 54,6 | 6,99 | 6,43 | 2 | 0,878 | 1,064 | 1,152 | 0,862 | 1,061 | 1,367 | 1,235 | 1,161 | 0,82 | -0,18 | 6,4E-02 |
| O55131 | Septin-7 OS=Mus musculus OX=10090 GN=Septin7 PE=1 SV=1 | 36 | 18 | 47 | 17 | 436 | 50,5 | 8,57 | 101,29 | 18 | 1,129 | 1,087 | 1,042 | 1,041 | 0,967 | 1,04 | 0,986 | 1,046 | 1,06 | 0,06 | 6,5E-02 |
| Q9D3D9 | ATP synthase subunit delta, mitochondrial OS=Mus musculus OX=1 | 17 | 3 | 9 | 2 | 168 | 17,6 | 5,08 | 16,59 | 3 | 0,956 | 1,073 | 1,009 | 1,067 | 1,05 | 1,091 | 1,152 | 1,136 | 0,93 | -0,07 | 6,5E-02 |
| P23818 | Glutamate receptor 1 OS=Mus musculus OX=10090 GN=Gria1 PE=1 | 17 | 15 | 19 | 11 | 907 | 101,5 | 7,69 | 48,01 | 15 | 0,908 | 0,936 | 0,928 | 0,985 | 1,046 | 0,93 | 1,072 | 1,025 | 0,92 | -0,08 | 6,5E-02 |
| Q99LC3 | NADH dehydrogenase [ubiquinone] 1 alpha subcomplex subunit 10 | 17 | 6 | 8 | 6 | 355 | 40,6 | 7,78 | 22,88 | 6 | 0,964 | 0,903 | 0,937 | 1,019 | 1,059 | 1,061 | 0,985 | 1,001 | 0,93 | -0,07 | 6,5E-02 |
| P59999 | Actin-related protein 2/3 complex subunit 4 OS=Mus musculus OX= | 13 | 3 | 3 | 3 | 168 | 19,7 | 8,43 | 4,8 | 3 | 1,037 | 0,948 | 1,02 | 1,06 | 0,99 | 0,752 | 0,928 | 0,891 | 1,14 | 0,14 | 6,5E-02 |
| Q9WUC3 | Lymphocyte antigen 6H OS=Mus musculus OX=10090 GN=Ly6h PE= | 12 | 2 | 3 | 2 | 139 | 14,7 | 7,53 | 8,62 | 2 | 0,707 | 0,63 | 0,729 | 1,033 | 1,091 | 0,88 | 1,16 | 0,952 | 0,76 | -0,24 | 6,5E-02 |
| P35564 | Calnexin OS=Mus musculus OX=10090 GN=Canx PE=1 SV=1 | 25 | 15 | 22 | 15 | 591 | 67,2 | 4,64 | 37,29 | 15 | 1,016 | 0,927 | 1,039 | 1,008 | 1,009 | 1,198 | 1,17 | 1,067 | 0,90 | -0,10 | 6,5E-02 |
| P39053-6 | Isoform 6 of Dynamin-1 OS=Mus musculus OX=10090 GN=Dnm1 | 42 | 40 | 96 | 1 | 864 | 97,2 | 7,74 | 194,77 | 40 | 1,069 | 1,015 | 1,025 | 1,125 | 1,006 | 0,931 | 0,853 | 1,025 | 1,11 | 0,11 | 6,5E-02 |
| Q6PEV3 | WAS/WASL-interacting protein family member 2 OS=Mus musculus | 5 | 2 | 2 | 2 | 440 | 46,3 | 10,99 | 5,62 | 2 | 1,159 | 1,268 | 1,042 | 1,202 | 0,934 | 1,118 | 0,988 | 1,067 | 1,14 | 0,14 | 6,6E-02 |
| Q9WUD1 | STIP1 homology and U box-containing protein 1 OS=Mus musculus | 10 | 3 | 4 | 3 | 304 | 34,9 | 6,01 | 6,86 | 3 | 1,129 | 0,939 | 0,993 | 1,022 | 1,167 | 1,132 | 1,153 | 1,055 | 0,91 | -0,09 | 6,6E-02 |
| Q60571 | Corticotropin-releasing factor-binding protein OS=Mus musculus O | 3 | 1 | 1 | 1 | 322 | 36 | 6,6 | 1,69 | 1 | 1,095 | 1,038 | 0,94 | 1,05 | 0,901 | 0,903 | 1,024 | 0,835 | 1,13 | 0,13 | 6,6E-02 |
| P58021 | Transmembrane 9 superfamily member 2 OS=Mus musculus OX=10 | 2 | 1 | 1 | 1 | 662 | 75,3 | 7,43 | 4,12 | 1 | 0,839 | 1,242 | 1,305 | 1,178 | 1,317 | 2,146 | 1,459 | 1,532 | 0,71 | -0,29 | 6,6E-02 |
| Q61037-6 | Isoform E of Tuberin OS=Mus musculus OX=10090 GN=Tsc2 | 7 | 9 | 9 | 9 | 1786 | 198,8 | 7,01 | 14,96 | 9 | 1,087 | 1,255 | 1,075 | 1,177 | 0,958 | 0,94 | 1,082 | 1,092 | 1,13 | 0,13 | 6,6E-02 |
| Q9D0M5 | Dynein light chain 2, cytoplasmic OS=Mus musculus OX=10090 GN= | 64 | 5 | 16 | 2 | 89 | 10,3 | 7,37 | 47,44 | 5 | 1,089 | 1,176 | 1,044 | 1,336 | 1,104 | 1,004 | 0,843 | 0,933 | 1,20 | 0,20 | 6,6E-02 |
| P48320 | Glutamate decarboxylase 2 OS=Mus musculus OX=10090 GN=Gad2 | 14 | 6 | 12 | 5 | 585 | 65,2 | 6,9 | 10,31 | 6 | 0,868 | 0,862 | 1,027 | 0,932 | 1,12 | 1,048 | 1,012 | 0,963 | 0,89 | -0,11 | 6,6E-02 |
| Q8K0G5 | EARP and GARP complex-interacting protein 1 OS=Mus musculus O | 14 | 4 | 5 | 4 | 386 | 43,1 | 5,14 | 10,45 | 4 | 0,741 | 0,875 | 0,977 | 0,937 | 0,996 | 1,017 | 1,054 | 0,959 | 0,88 | -0,12 | 6,6E-02 |
| O08807 | Peroxiredoxin-4 OS=Mus musculus OX=10090 GN=Prdx4 PE=1 SV=1 | 22 | 6 | 6 | 4 | 274 | 31 | 7,15 | 8,77 | 6 | 1,088 | 1,095 | 1,14 | 0,994 | 0,976 | 0,882 | 0,704 | 1,042 | 1,20 | 0,20 | 6,6E-02 |
| P08030 | Adenine phosphoribosyltransferase OS=Mus musculus OX=10090 G | 7 | 1 | 1 | 1 | 180 | 19,7 | 6,79 | 2,8 | 1 | 1,001 | 1,088 | 1,063 | 0,99 | 0,82 | 0,97 | 0,837 | 1,022 | 1,14 | 0,14 | 6,6E-02 |
| Q8BL97 | Serine/arginine-rich splicing factor 7 OS=Mus musculus OX=10090 G | 5 | 1 | 1 | 1 | 267 | 30,8 | 11,9 | 2,49 | 1 | 1,209 | 1,081 | 1,239 | 1,145 | 0,848 | 0,917 | 1,183 | 0,59 | 1,32 | 0,32 | 6,6E-02 |
| P62743 | AP-2 complex subunit sigma OS=Mus musculus OX=10090 GN=Ap2 | 26 | 4 | 4 | 4 | 142 | 17 | 6,18 | 4,87 | 4 | 0,556 | 0,82 | 0,584 | 0,698 | 0,811 | 0,769 | 0,792 | 0,843 | 0,83 | -0,17 | 6,7E-02 |
| O70251 | Elongation factor 1-beta OS=Mus musculus OX=10090 GN=Eef1b PI | 17 | 4 | 5 | 3 | 225 | 24,7 | 4,69 | 11,81 | 4 | 1,119 | 1,022 | 1,001 | 1,023 | 1,12 | 1,065 | 1,149 | 1,114 | 0,94 | -0,06 | 6,7E-02 |
| P97412 | Lysosomal-trafficking regulator OS=Mus musculus OX=10090 GN=L | 1 | 3 | 3 | 3 | 3788 | 425 | 6,6 | 2,57 | 3 | 1,73 | 1,433 | 1,52 | 1,367 | 1,115 | 1,475 | 1,1 | 1,301 | 1,21 | 0,21 | 6,7E-02 |
| P17710 | Hexokinase-1 OS=Mus musculus OX=10090 GN=Hk1 PE=1 SV=3 | 37 | 37 | 98 | 37 | 974 | 108,2 | 6,8 | 208 | 37 | 0,944 | 0,92 | 0,957 | 0,972 | 0,983 | 0,958 | 0,999 | 0,977 | 0,97 | -0,03 | 6,7E-02 |
| Q9CWJ9 | Bifunctional purine biosynthesis protein PURH OS=Mus musculus C | 14 | 7 | 8 | 7 | 592 | 64,2 | 6,76 | 18,15 | 7 | 1,084 | 1,08 | 1,031 | 1,046 | 0,9 | 1,004 | 0,991 | 1,047 | 1,08 | 0,08 | 6,7E-02 |
| P60335 | Poly(rC)-binding protein 1 OS=Mus musculus OX=10090 GN=Pcbp1 | 25 | 6 | 10 | 4 | 356 | 37,5 | 7,09 | 19,59 | 6 | 0,999 | 1,079 | 1,185 | 1,065 | 0,955 | 1,057 | 0,957 | 0,945 | 1,11 | 0,11 | 6,8E-02 |
| A2A9C3 | KICSTOR complex protein SZT2 OS=Mus musculus OX=10090 GN=S | 1 | 1 | 1 | 1 | 3431 | 377,4 | 6,19 | 2,08 | 1 | 2,21 | 1,625 | 1,158 | 1,907 | 0,785 | 1,503 | 0,798 | 1,276 | 1,58 | 0,58 | 6,9E-02 |
| P63158 | High mobility group protein B1 OS=Mus musculus OX=10090 GN=H | 13 | 2 | 2 | 2 | 215 | 24,9 | 5,74 | 6,24 | 2 | 1,231 | 1,278 | 1,208 | 1,317 | 1,211 | 1,189 | 1,219 | 1,117 | 1,06 | 0,06 | 6,9E-02 |
| O88441 | Metaxin-2 OS=Mus musculus OX=10090 GN=Mtx2 PE=1 SV=1 | 10 | 2 | 4 | 2 | 263 | 29,7 | 5,63 | 8,37 | 2 | 0,816 | 0,891 | 1,07 | 1,004 | 1,098 | 1,065 | 1,05 | 1,077 | 0,88 | -0,12 | 6,9E-02 |
| P68404-2 | Isoform Beta-II of Protein kinase C beta type OS=Mus musculus OX= | 29 | 22 | 40 | 16 | 673 | 76,8 | 7,02 | 83,68 | 22 | 0,927 | 0,911 | 0,956 | 1,006 | 0,99 | 1,021 | 1,062 | 0,976 | 0,94 | -0,06 | 7,0E-02 |
| Q8BNE1 | TRPM8 channel-associated factor 1 OS=Mus musculus OX=10090 G | 1 | 1 | 1 | 1 | 924 | 102,7 | 6,96 | 2,43 | 1 | 1,562 | 1,19 | 1,028 | 1,405 | 1,317 | 0,644 | 0,821 | 0,706 | 1,49 | 0,49 | 7,0E-02 |
| Q61885 | Myelin-oligodendrocyte glycoprotein OS=Mus musculus OX=10090 | 26 | 7 | 9 | 7 | 246 | 28,3 | 7,96 | 29,5 | 7 | 1,48 | 1,197 | 1,088 | 1,034 | 1,079 | 1,018 | 0,828 | 0,837 | 1,28 | 0,28 | 7,0E-02 |
| Q4KUS2 | Protein unc-13 homolog A OS=Mus musculus OX=10090 GN=Unc13 | 8 | 14 | 16 | 9 | 1712 | 193,7 | 5,25 | 33,88 | 14 | 1,009 | 0,961 | 0,986 | 1,057 | 1,011 | 1,063 | 1,095 | 1,09 | 0,94 | -0,06 | 7,1E-02 |
| Q7TPM6 | Fibronectin type III and SPRY domain-containing protein 1 OS=Mus | 17 | 7 | 8 | 7 | 496 | 55,5 | 6,67 | 14,75 | 7 | 1,076 | 1,088 | 1,077 | 1,046 | 1,066 | 0,94 | 1,037 | 0,991 | 1,06 | 0,06 | 7,2E-02 |
| O88990 | Alpha-actinin-3 OS=Mus musculus OX=10090 GN=Actn3 PE=2 SV=1 | 10 | 8 | 11 | 1 | 900 | 103 | 5,45 | 16,55 | 8 | 1,014 | 1,082 | 0,934 | 0,886 | 0,862 | 0,758 | 0,9 | 0,912 | 1,14 | 0,14 | 7,3E-02 |
| Q75NR7 | ATP-dependent DNA helicase Q4 OS=Mus musculus OX=10090 GN= | 1 | 1 | 1 | 1 | 1216 | 135 | 8,37 | 2,13 | 1 | 1,01 | 1,303 | 1,059 | 1,03 | 1,059 | 0,878 | 0,901 | 0,84 | 1,20 | 0,20 | 7,3E-02 |
| Q9CQI3 | Glia maturation factor beta OS=Mus musculus OX=10090 GN=Gmfb | 15 | 2 | 3 | 2 | 142 | 16,7 | 5,16 | 8,79 | 2 | 0,946 | 0,994 | 0,964 | 0,955 | 0,874 | 0,925 | 0,904 | 0,966 | 1,05 | 0,05 | 7,4E-02 |
| Q9CS42 | Ribose-phosphate pyrophosphokinase 2 OS=Mus musculus OX=100 | 25 | 6 | 7 | 3 | 318 | 34,8 | 6,61 | 14,92 | 6 | 1,033 | 1,005 | 1,018 | 1,131 | 1,212 | 1,25 | 1,075 | 1,096 | 0,90 | -0,10 | 7,4E-02 |
| O35598 | Disintegrin and metalloproteinase domain-containing protein 10 O | 2 | 1 | 2 | 1 | 749 | 83,9 | 7,94 | 2,91 | 1 | 1,006 | 1,059 | 1,068 | 0,948 | 1,09 | 1,233 | 1,062 | 1,102 | 0,91 | -0,09 | 7,4E-02 |
| P12382 | ATP-dependent 6-phosphofructokinase, liver type OS=Mus muscul | 17 | 14 | 20 | 12 | 780 | 85,3 | 7,17 | 42,38 | 14 | 1,025 | 0,952 | 1,018 | 0,994 | 1,016 | 1,023 | 1,068 | 1,071 | 0,95 | -0,05 | 7,5E-02 |
| Q5SWU9 | Acetyl-CoA carboxylase 1 OS=Mus musculus OX=10090 GN=Acaca P | 5 | 9 | 9 | 9 | 2345 | 265,1 | 6,39 | 12,75 | 9 | 1,045 | 1,055 | 1,093 | 1,05 | 1,019 | 0,979 | 1,039 | 1,046 | 1,04 | 0,04 | 7,5E-02 |
| Q80WT0 | Junctophilin-4 OS=Mus musculus OX=10090 GN=Jph4 PE=2 SV=1 | 3 | 1 | 1 | 1 | 628 | 66 | 6,71 | 2,53 | 1 | 2,315 | 1,637 | 1,598 | 1,14 | 0,998 | 1,358 | 1,133 | 0,994 | 1,49 | 0,49 | 7,5E-02 |
| Q9R1R2 | Tripartite motif-containing protein 3 OS=Mus musculus OX=10090 | 12 | 7 | 7 | 7 | 744 | 80,7 | 7,81 | 13,94 | 7 | 1,09 | 1,015 | 1,031 | 1,097 | 1,018 | 1,043 | 0,948 | 0,958 | 1,07 | 0,07 | 7,5E-02 |
| P21126 | Ubiquitin-like protein 4A OS=Mus musculus OX=10090 GN=Ubl4a P | 19 | 3 | 3 | 3 | 157 | 17,8 | 8,44 | 1,72 | 3 | 1,159 | 0,974 | 0,852 | 0,735 | 0,713 | 0,81 | 0,724 | 0,637 | 1,29 | 0,29 | 7,5E-02 |

| | | | | | | | | | | | | | | | | | | | | | |
|---|---|---|---|---|---|---|---|---|---|---|---|---|---|---|---|---|---|---|---|---|---|
| Q9D2U5 | N-alpha-acetyltransferase 38, NatC auxiliary subunit OS=Mus musc | 7 | 1 | 1 | 1 | 125 | 13,4 | 5,5 | 0 | 1 | 1,332 | 1,243 | 1,408 | 0,886 | 0,684 | 0,766 | 0,836 | 1,196 | 1,40 | 0,40 | 7,5E-02 |
| Q9ERF3 | WD repeat-containing protein 61 OS=Mus musculus OX=10090 GN= | 4 | 1 | 1 | 1 | 305 | 33,8 | 5,36 | 2,78 | 1 | 1,227 | 1,203 | 1,037 | 0,967 | 0,898 | 1,115 | 0,768 | 0,738 | 1,26 | 0,26 | 7,6E-02 |
| Q91ZU6 | Dystonin OS=Mus musculus OX=10090 GN=Dst PE=1 SV=2 | 6 | 39 | 46 | 35 | 7393 | 833,7 | 5,31 | 81,65 | 39 | 1,176 | 1,161 | 1,054 | 0,98 | 0,95 | 0,97 | 1,009 | 1,02 | 1,11 | 0,11 | 7,6E-02 |
| P28741 | Kinesin-like protein KIF3A OS=Mus musculus OX=10090 GN=Kif3a P | 8 | 5 | 7 | 5 | 701 | 80,1 | 6,54 | 8,31 | 5 | 1,363 | 1,337 | 1,134 | 1,189 | 0,997 | 0,931 | 1,024 | 1,272 | 1,19 | 0,19 | 7,6E-02 |
| Q8VHK5 | Membrane protein MLC1 OS=Mus musculus OX=10090 GN=Mlc1 P | 10 | 3 | 3 | 3 | 382 | 41,6 | 7,9 | 5,1 | 3 | 0,896 | 1,196 | 1,005 | 1,077 | 1,146 | 1,277 | 1,194 | 1,153 | 0,88 | -0,12 | 7,7E-02 |
| Q6PHN9 | Ras-related protein Rab-35 OS=Mus musculus OX=10090 GN=Rab3 | 61 | 13 | 17 | 11 | 201 | 23 | 8,29 | 33,29 | 13 | 1,095 | 1,038 | 1,058 | 1,029 | 1,041 | 1,027 | 0,984 | 1,003 | 1,04 | 0,04 | 7,7E-02 |
| Q499X9 | Methionine--tRNA ligase, mitochondrial OS=Mus musculus OX=100 | 2 | 1 | 1 | 1 | 586 | 65,8 | 8,09 | 2,1 | 1 | 1,438 | 1,571 | 1,1 | 1,185 | 0,877 | 1,086 | 1,026 | 1,207 | 1,26 | 0,26 | 7,8E-02 |
| Q6NXK7 | Inactive dipeptidyl peptidase 10 OS=Mus musculus OX=10090 GN= | 26 | 22 | 32 | 20 | 797 | 90,8 | 6,48 | 56,83 | 22 | 0,914 | 0,904 | 0,967 | 0,934 | 0,981 | 0,936 | 0,969 | 0,983 | 0,96 | -0,04 | 7,8E-02 |
| P55096 | ATP-binding cassette sub-family D member 3 OS=Mus musculus OX | 3 | 2 | 2 | 2 | 659 | 75,4 | 9,26 | 2,44 | 2 | 1,03 | 1,031 | 1,047 | 0,995 | 0,84 | 1,079 | 0,8 | 0,824 | 1,16 | 0,16 | 7,8E-02 |
| Q02105 | Complement C1q subcomponent subunit C OS=Mus musculus OX= | 4 | 1 | 1 | 1 | 246 | 26 | 8,54 | 2,33 | 1 | 0,996 | 0,987 | 0,928 | 0,875 | 0,888 | 0,746 | 0,851 | 0,91 | 1,12 | 0,12 | 7,8E-02 |
| A6H630 | Damage-control phosphatase ARMT1 OS=Mus musculus OX=10090 | 8 | 4 | 4 | 4 | 439 | 50,5 | 5,92 | 10,26 | 4 | 0,881 | 0,975 | 0,989 | 1,032 | 1,009 | 1,014 | 1,229 | 1,168 | 0,88 | -0,12 | 7,8E-02 |
| Q5U3K5 | Rab-like protein 6 OS=Mus musculus OX=10090 GN=Rabl6 PE=1 SV= | 8 | 6 | 7 | 6 | 725 | 79,8 | 5,53 | 6,42 | 6 | 0,567 | 0,892 | 0,915 | 1,016 | 1,109 | 0,92 | 1,166 | 1,153 | 0,78 | -0,22 | 7,8E-02 |
| Q99N95 | 39S ribosomal protein L3, mitochondrial OS=Mus musculus OX=100 | 12 | 3 | 3 | 3 | 348 | 39,1 | 9,55 | 5,98 | 3 | 0,884 | 0,726 | 0,943 | 0,921 | 0,934 | 1,272 | 0,973 | 1,059 | 0,82 | -0,18 | 7,8E-02 |
| P22315 | Ferrochelatase, mitochondrial OS=Mus musculus OX=10090 GN=Fe | 8 | 3 | 4 | 3 | 420 | 47,1 | 8,91 | 10,77 | 3 | 0,913 | 0,893 | 1,021 | 1,027 | 1,18 | 1,034 | 1,114 | 0,991 | 0,89 | -0,11 | 7,8E-02 |
| Q0GNC1 | Inverted formin-2 OS=Mus musculus OX=10090 GN=Inf2 PE=1 SV=1 | 1 | 1 | 1 | 1 | 1273 | 138,5 | 5,21 | 2,24 | 1 | 1,214 | 1,209 | 1,184 | 1,163 | 0,88 | 1,059 | 1,148 | 1,143 | 1,13 | 0,13 | 7,8E-02 |
| Q31125 | Zinc transporter SLC39A7 OS=Mus musculus OX=10090 GN=Slc39a7 | 6 | 2 | 2 | 2 | 476 | 50,6 | 6,87 | 0 | 2 | 0,837 | 0,95 | 0,859 | 1,038 | 1,108 | 1,204 | 1,138 | 0,905 | 0,85 | -0,15 | 7,8E-02 |
| Q925H1 | Zinc finger transcription factor Trps1 OS=Mus musculus OX=10090 | 1 | 1 | 1 | 1 | 1281 | 140,9 | 7,59 | 0 | 1 | 2,139 | 1,252 | 1,514 | 2,677 | 1,154 | 1,385 | 1,234 | 1,022 | 1,58 | 0,58 | 7,9E-02 |
| P05201 | Aspartate aminotransferase, cytoplasmic OS=Mus musculus OX=10 | 31 | 12 | 20 | 12 | 413 | 46,2 | 7,14 | 37,15 | 12 | 0,931 | 0,921 | 0,864 | 0,948 | 1,005 | 0,924 | 0,957 | 1,013 | 0,94 | -0,06 | 7,9E-02 |
| P17742 | Peptidyl-prolyl cis-trans isomerase A OS=Mus musculus OX=10090 | 55 | 13 | 40 | 12 | 164 | 18 | 7,9 | 75,35 | 13 | 0,987 | 1,085 | 0,953 | 1,015 | 0,95 | 0,89 | 0,951 | 0,97 | 1,07 | 0,07 | 7,9E-02 |
| Q9CQR4 | Acyl-coenzyme A thioesterase 13 OS=Mus musculus OX=10090 GN= | 21 | 3 | 4 | 3 | 140 | 15,2 | 8,82 | 3,66 | 3 | 0,625 | 0,697 | 0,883 | 0,844 | 1,049 | 0,814 | 0,986 | 0,88 | 0,82 | -0,18 | 7,9E-02 |
| P53811 | Phosphatidylinositol transfer protein beta isoform OS=Mus muscul | 20 | 4 | 4 | 3 | 271 | 31,5 | 6,95 | 5,17 | 4 | 1,026 | 0,68 | 0,935 | 0,893 | 0,883 | 1,691 | 1,158 | 1,373 | 0,69 | -0,31 | 7,9E-02 |
| O55234 | Proteasome subunit beta type-5 OS=Mus musculus OX=10090 GN= | 9 | 3 | 4 | 3 | 264 | 28,5 | 7,02 | 9,77 | 3 | 0,972 | 0,926 | 0,835 | 0,99 | 1,026 | 0,985 | 0,998 | 1,016 | 0,92 | -0,08 | 8,0E-02 |
| P70290 | 55 kDa erythrocyte membrane protein OS=Mus musculus OX=1009 | 18 | 6 | 7 | 6 | 466 | 52,2 | 7,2 | 9,33 | 6 | 1,958 | 2,483 | 1,272 | 1,19 | 0,884 | 1,288 | 1,073 | 0,983 | 1,63 | 0,63 | 8,0E-02 |
| Q9JI91 | Alpha-actinin-2 OS=Mus musculus OX=10090 GN=Actn2 PE=1 SV=2 | 30 | 23 | 32 | 13 | 894 | 103,8 | 5,45 | 62,65 | 23 | 1,117 | 1,067 | 1,036 | 1,082 | 1,137 | 1,134 | 1,294 | 1,115 | 0,92 | -0,08 | 8,0E-02 |
| Q6NS60 | F-box only protein 41 OS=Mus musculus OX=10090 GN=Fbxo41 PE= | 11 | 10 | 12 | 10 | 873 | 94,3 | 8,24 | 24,88 | 10 | 1,083 | 1,341 | 1,079 | 1,089 | 0,933 | 1,072 | 1,017 | 0,974 | 1,15 | 0,15 | 8,0E-02 |
| Q99KB8 | Hydroxyacylglutathione hydrolase, mitochondrial OS=Mus musculu | 28 | 7 | 12 | 7 | 309 | 34,1 | 7,75 | 29,4 | 7 | 1,08 | 1,058 | 1,049 | 1 | 0,957 | 1 | 0,972 | 1,044 | 1,05 | 0,05 | 8,1E-02 |
| Q3UTQ8 | Cyclin-dependent kinase-like 5 OS=Mus musculus OX=10090 GN=C | 9 | 7 | 9 | 6 | 938 | 105,4 | 9,58 | 11,61 | 7 | 0,983 | 0,853 | 1,038 | 1,052 | 1,025 | 1,159 | 1,059 | 1,165 | 0,89 | -0,11 | 8,1E-02 |
| Q8K4R4 | Cytoplasmic phosphatidylinositol transfer protein 1 OS=Mus muscu | 9 | 3 | 3 | 3 | 332 | 38,4 | 6,32 | 4,81 | 3 | 0,01 | 0,727 | 0,597 | 0,843 | 1,008 | 1,005 | 0,855 | 0,894 | 0,58 | -0,42 | 8,1E-02 |
| Q61672 | Equilibrative nucleoside transporter 2 OS=Mus musculus OX=1009 | 2 | 1 | 1 | 1 | 456 | 50,2 | 6,51 | 2,56 | 1 | 0,596 | 1,077 | 0,719 | 0,956 | 0,907 | 1,221 | 1,195 | 1,119 | 0,75 | -0,25 | 8,1E-02 |
| Q9Z1L5 | Voltage-dependent calcium channel subunit alpha-2/delta-3 OS=M | 10 | 9 | 10 | 9 | 1091 | 122,7 | 5,73 | 12,28 | 9 | 1,247 | 1,293 | 1,112 | 1,075 | 1,034 | 1,097 | 1,003 | 1,107 | 1,11 | 0,11 | 8,1E-02 |
| O88737 | Protein bassoon OS=Mus musculus OX=10090 GN=Bsn PE=1 SV=4 | 26 | 77 | 101 | 73 | 3942 | 418,6 | 7,71 | 219,08 | 77 | 0,932 | 0,982 | 0,965 | 0,989 | 0,974 | 1,015 | 0,996 | 1,029 | 0,96 | -0,04 | 8,1E-02 |
| Q99MI1 | ELKS/Rab6-interacting/CAST family member 1 OS=Mus musculus O | 12 | 15 | 16 | 12 | 1120 | 128,3 | 5,87 | 32,93 | 15 | 0,998 | 1,069 | 0,99 | 1,061 | 0,798 | 0,979 | 0,928 | 0,998 | 1,11 | 0,11 | 8,1E-02 |
| Q9D0R2 | Threonine--tRNA ligase, cytoplasmic OS=Mus musculus OX=10090 | 9 | 6 | 6 | 5 | 722 | 83,3 | 7,36 | 14,65 | 6 | 1,205 | 1,046 | 1,109 | 1,025 | 1,03 | 0,986 | 0,988 | 1,028 | 1,09 | 0,09 | 8,1E-02 |
| Q8BKR5 | Protein phosphatase 1 regulatory subunit 37 OS=Mus musculus OX | 1 | 1 | 1 | 1 | 712 | 77,5 | 5,06 | 1,74 | 1 | 0,951 | 0,965 | 1,017 | 0,873 | 0,97 | 1,227 | 1,027 | 1,11 | 0,88 | -0,12 | 8,2E-02 |
| P46735 | Unconventional myosin-Ib OS=Mus musculus OX=10090 GN=Myo1 | 7 | 6 | 6 | 6 | 1107 | 128,5 | 9,26 | 16,07 | 6 | 0,993 | 0,997 | 0,972 | 0,83 | 1,024 | 1,061 | 0,994 | 1,081 | 0,91 | -0,09 | 8,2E-02 |
| Q9JHQ5 | Leucine zipper transcription factor-like protein 1 OS=Mus musculus | 15 | 4 | 4 | 4 | 299 | 34,8 | 5,17 | 8,37 | 4 | 0,956 | 0,981 | 1,072 | 1,054 | 0,901 | 0,846 | 1,017 | 0,922 | 1,10 | 0,10 | 8,3E-02 |
| P46664 | Adenylosuccinate synthetase isozyme 2 OS=Mus musculus OX=100 | 11 | 6 | 6 | 6 | 456 | 50 | 6,38 | 14,91 | 6 | 0,958 | 1,057 | 1,001 | 1,054 | 0,968 | 0,942 | 0,983 | 0,968 | 1,05 | 0,05 | 8,3E-02 |
| Q99KJ8 | Dynactin subunit 2 OS=Mus musculus OX=10090 GN=Dctn2 PE=1 S | 31 | 9 | 11 | 9 | 402 | 44,1 | 5,26 | 33,67 | 9 | 1,229 | 1,115 | 0,999 | 1,101 | 0,907 | 1,095 | 0,959 | 0,97 | 1,13 | 0,13 | 8,3E-02 |
| Q8BI84 | Transport and Golgi organization protein 1 homolog OS=Mus musc | 2 | 3 | 3 | 3 | 1930 | 213,5 | 4,75 | 1,7 | 3 | 1,488 | 1,444 | 1,641 | 1,2 | 0,915 | 1,408 | 0,876 | 1,255 | 1,30 | 0,30 | 8,3E-02 |
| Q9CWK8 | Sorting nexin-2 OS=Mus musculus OX=10090 GN=Snx2 PE=1 SV=2 | 26 | 13 | 16 | 11 | 519 | 58,4 | 5,12 | 36,35 | 13 | 1,084 | 1,167 | 1,09 | 1,036 | 1,002 | 1,085 | 1,016 | 0,984 | 1,07 | 0,07 | 8,3E-02 |
| Q62419 | Endophilin-A2 OS=Mus musculus OX=10090 GN=Sh3gl1 PE=1 SV=1 | 25 | 9 | 9 | 6 | 368 | 41,5 | 5,72 | 12,41 | 9 | 1,09 | 1,021 | 0,948 | 0,971 | 0,789 | 0,993 | 0,917 | 0,895 | 1,12 | 0,12 | 8,3E-02 |
| Q62108 | Disks large homolog 4 OS=Mus musculus OX=10090 GN=Dlg4 PE=1 | 30 | 22 | 28 | 20 | 724 | 80,4 | 5,87 | 85,37 | 22 | 0,917 | 0,914 | 0,872 | 1,008 | 1,078 | 0,93 | 1,051 | 1,01 | 0,91 | -0,09 | 8,3E-02 |
| Q61503 | 5'-nucleotidase OS=Mus musculus OX=10090 GN=Nt5e PE=1 SV=2 | 2 | 1 | 1 | 1 | 576 | 63,8 | 6,64 | 2,34 | 1 | 0,863 | 1,005 | 0,791 | 1,142 | 1,239 | 1,005 | 1,124 | 1,214 | 0,83 | -0,17 | 8,3E-02 |
| Q9Z2D3 | Gasdermin-E OS=Mus musculus OX=10090 GN=Gsdme PE=1 SV=1 | 7 | 3 | 4 | 3 | 512 | 56,6 | 5,39 | 2,85 | 3 | 1,08 | 0,939 | 0,905 | 0,998 | 0,882 | 0,894 | 0,869 | 0,937 | 1,09 | 0,09 | 8,4E-02 |
| Q9WUN2 | Serine/threonine-protein kinase TBK1 OS=Mus musculus OX=1009 | 6 | 4 | 6 | 4 | 729 | 83,4 | 6,87 | 12,15 | 4 | 1,271 | 1,213 | 1,099 | 0,986 | 0,96 | 0,969 | 1,058 | 1,025 | 1,14 | 0,14 | 8,4E-02 |
| Q9CQW2 | ADP-ribosylation factor-like protein 8B OS=Mus musculus OX=1009 | 10 | 3 | 4 | 2 | 186 | 21,5 | 8,43 | 8,08 | 3 | 0,639 | 0,869 | 0,7 | 0,889 | 0,838 | 1,177 | 0,902 | 0,975 | 0,80 | -0,20 | 8,4E-02 |
| Q8VCN9 | Tubulin-specific chaperone C OS=Mus musculus OX=10090 GN=Tbc | 6 | 2 | 3 | 2 | 341 | 38,1 | 5,3 | 6,67 | 2 | 1,456 | 1,182 | 1,32 | 1,288 | 1,172 | 1,203 | 1,182 | 1,215 | 1,10 | 0,10 | 8,4E-02 |
| Q8C437 | PEX5-related protein OS=Mus musculus OX=10090 GN=Pex5l PE=1 | 12 | 6 | 7 | 6 | 567 | 63,1 | 5,33 | 14,68 | 6 | 0,914 | 0,804 | 1,026 | 0,884 | 1,005 | 1,018 | 1,043 | 0,965 | 0,90 | -0,10 | 8,4E-02 |
| Q9D2R6 | Cytochrome c oxidase assembly factor 3 homolog, mitochondrial O | 9 | 1 | 1 | 1 | 108 | 12 | 9,86 | 2,29 | 1 | 0,882 | 0,923 | 1,038 | 0,918 | 1,128 | 1,121 | 0,906 | 1,179 | 0,87 | -0,13 | 8,4E-02 |
| Q9JHG6 | Calcipressin-1 OS=Mus musculus OX=10090 GN=Rcan1 PE=1 SV=2 | 4 | 1 | 1 | 1 | 251 | 28,1 | 5,41 | 2,03 | 1 | 1,452 | 1,28 | 1,693 | 0,927 | 0,93 | 0,937 | 1,205 | 0,732 | 1,41 | 0,41 | 8,5E-02 |
| P20029 | Endoplasmic reticulum chaperone BiP OS=Mus musculus OX=1009 | 38 | 26 | 37 | 24 | 655 | 72,4 | 5,16 | 91,05 | 26 | 1,074 | 1,037 | 1,12 | 0,983 | 0,978 | 0,988 | 0,992 | 1,011 | 1,06 | 0,06 | 8,6E-02 |
| Q9EQ06 | Estradiol 17-beta-dehydrogenase 11 OS=Mus musculus OX=10090 | 5 | 1 | 1 | 1 | 298 | 32,9 | 8,66 | 2,89 | 1 | 0,964 | 0,851 | 0,866 | 0,909 | 1,106 | 1,162 | 0,961 | 0,902 | 0,87 | -0,13 | 8,6E-02 |
| O55102 | Biogenesis of lysosome-related organelles complex 1 subunit 1 OS= | 8 | 1 | 1 | 1 | 125 | 14,3 | 8,15 | 0 | 1 | 1,166 | 1,112 | 0,787 | 1,125 | 0,772 | 0,997 | 0,735 | 0,824 | 1,26 | 0,26 | 8,6E-02 |
| Q7M729 | Sodium channel subunit beta-4 OS=Mus musculus OX=10090 GN=S | 5 | 2 | 3 | 2 | 228 | 25,2 | 8,82 | 7,46 | 2 | 1,001 | 0,684 | 0,866 | 0,908 | 1,075 | 0,916 | 1,365 | 1,051 | 0,78 | -0,22 | 8,6E-02 |
| Q9Z0S6 | Claudin-10 OS=Mus musculus OX=10090 GN=Cldn10 PE=1 SV=2 | 6 | 1 | 1 | 1 | 231 | 24,7 | 7,8 | 3,03 | 1 | 0,913 | 1,153 | 1,535 | 1,176 | 0,964 | 1,03 | 0,864 | 0,764 | 1,32 | 0,32 | 8,6E-02 |
| Q8BHZ0 | Protein FAM49A OS=Mus musculus OX=10090 GN=Fam49a PE=1 S | 16 | 4 | 5 | 2 | 323 | 37,3 | 6,01 | 7,52 | 4 | 1,33 | 1,016 | 0,862 | 0,979 | 0,848 | 0,721 | 0,651 | 0,967 | 1,31 | 0,31 | 8,6E-02 |
| Q9CR00 | 26S proteasome non-ATPase regulatory subunit 9 OS=Mus musculu | 27 | 5 | 6 | 5 | 222 | 24,7 | 6,43 | 11,4 | 5 | 0,902 | 0,963 | 0,844 | 0,875 | 0,972 | 0,95 | 0,943 | 1,122 | 0,90 | -0,10 | 8,6E-02 |
| Q8R1N4 | NudC domain-containing protein 3 OS=Mus musculus OX=10090 G | 8 | 2 | 2 | 2 | 363 | 40,9 | 5,26 | 2,14 | 2 | 1,085 | 1,36 | 1,346 | 1,271 | 1,058 | 1,102 | 1,136 | 1,196 | 1,13 | 0,13 | 8,7E-02 |
| Q5U405 | Transmembrane protease serine 13 OS=Mus musculus OX=10090 | 1 | 1 | 10 | 1 | 543 | 59,8 | 9,01 | 9,37 | 1 | 1,418 | 1,13 | 1,392 | 0,938 | 1,038 | 0,943 | 0,964 | 0,984 | 1,24 | 0,24 | 8,7E-02 |
| Q8R1A4 | Dedicator of cytokinesis protein 7 OS=Mus musculus OX=10090 GN | 1 | 3 | 3 | 3 | 2130 | 241,3 | 6,71 | 4,67 | 3 | 1,864 | 1,663 | 1,501 | 1,322 | 1,099 | 1,547 | 1,251 | 1,233 | 1,24 | 0,24 | 8,7E-02 |
| Q9EPU5 | Tumor necrosis factor receptor superfamily member 21 OS=Mus m | 2 | 1 | 1 | 1 | 655 | 71,9 | 7,46 | 2,45 | 1 | 1,184 | 1,456 | 1,324 | 0,787 | 0,918 | 0,757 | 0,854 | 0,978 | 1,35 | 0,35 | 8,7E-02 |
| Q9Z1Q2 | Phosphatidylserine lipase ABHD16A OS=Mus musculus OX=10090 | 10 | 4 | 7 | 4 | 558 | 63 | 8,25 | 15,2 | 4 | 1,217 | 1,432 | 1,105 | 1,117 | 0,877 | 1,145 | 0,905 | 1,109 | 1,21 | 0,21 | 8,7E-02 |

| | | | | | | | | | | | | | | | | | | | | | |
|---|---|---|---|---|---|---|---|---|---|---|---|---|---|---|---|---|---|---|---|---|---|
| Q8K0U4 | Heat shock 70 kDa protein 12A OS=Mus musculus OX=10090 GN=H | 31 | 19 | 28 | 19 | 675 | 74,8 | 6,77 | 68,75 | 19 | 0,981 | 0,969 | 0,945 | 1,01 | 1,058 | 0,977 | 1,046 | 1,01 | 0,95 | -0,05 | 8,7E-02 |
| Q9QWW1 | Homer protein homolog 2 OS=Mus musculus OX=10090 GN=Home | 10 | 4 | 4 | 3 | 354 | 40,5 | 6,01 | 7,25 | 4 | 0,993 | 0,917 | 0,925 | 1,152 | 1,127 | 1,075 | 1,159 | 1,095 | 0,89 | -0,11 | 8,8E-02 |
| Q9QUJ7 | Long-chain-fatty-acid--CoA ligase 4 OS=Mus musculus OX=10090 GI | 6 | 4 | 4 | 4 | 711 | 79 | 8,28 | 7,13 | 4 | 0,781 | 0,887 | 0,73 | 0,96 | 1,013 | 0,939 | 0,969 | 0,901 | 0,88 | -0,12 | 8,8E-02 |
| Q6PEE3 | Ribonucleoside-diphosphate reductase subunit M2 B OS=Mus musc | 2 | 1 | 1 | 1 | 351 | 40,8 | 5,01 | 2 | 1 | 0,75 | 0,678 | 0,482 | 0,561 | 0,704 | 1,293 | 0,712 | 1,001 | 0,67 | -0,33 | 8,8E-02 |
| Q9DCS9 | NADH dehydrogenase [ubiquinone] 1 beta subcomplex subunit 10 | 28 | 5 | 6 | 5 | 176 | 21 | 8,03 | 12,3 | 5 | 0,8 | 0,817 | 1,108 | 0,968 | 1,099 | 1,182 | 1,093 | 0,989 | 0,85 | -0,15 | 8,8E-02 |
| Q8BP47 | Asparagine--tRNA ligase, cytoplasmic OS=Mus musculus OX=10090 | 17 | 8 | 10 | 8 | 559 | 64,2 | 5,86 | 18 | 8 | 1,126 | 0,944 | 1,042 | 1,085 | 1,043 | 0,872 | 0,915 | 0,933 | 1,12 | 0,12 | 8,8E-02 |
| Q9JKV1 | Proteasomal ubiquitin receptor ADRM1 OS=Mus musculus OX=100 | 2 | 1 | 1 | 1 | 407 | 42 | 5,07 | 2,33 | 1 | 0,973 | 1,099 | 0,871 | 0,932 | 1,04 | 1,021 | 1,133 | 1,165 | 0,89 | -0,11 | 8,9E-02 |
| P50153 | Guanine nucleotide-binding protein G(I)/G(S)/G(O) subunit gamma | 37 | 3 | 3 | 3 | 75 | 8,4 | 7,08 | 3,9 | 3 | 0,823 | 0,795 | 0,855 | 1,061 | 1,002 | 0,948 | 1,086 | 1,043 | 0,87 | -0,13 | 8,9E-02 |
| Q99PU8 | ATP-dependent RNA helicase DHX30 OS=Mus musculus OX=10090 | 1 | 1 | 1 | 1 | 1217 | 136,6 | 8,75 | 2,85 | 1 | 0,461 | 0,599 | 0,574 | 0,69 | 0,823 | 0,651 | 0,886 | 0,617 | 0,78 | -0,22 | 8,9E-02 |
| Q9DC07 | LIM zinc-binding domain-containing Nebulette OS=Mus musculus C | 30 | 8 | 10 | 7 | 270 | 31,1 | 8,31 | 16,8 | 8 | 1,076 | 1,126 | 1,109 | 0,95 | 0,997 | 1,041 | 0,86 | 0,792 | 1,15 | 0,15 | 8,9E-02 |
| Q9CPP6 | NADH dehydrogenase [ubiquinone] 1 alpha subcomplex subunit 5 | 23 | 2 | 3 | 2 | 116 | 13,4 | 8,1 | 5,13 | 2 | 0,661 | 0,792 | 0,784 | 0,967 | 1,138 | 0,748 | 1,08 | 1,209 | 0,77 | -0,23 | 8,9E-02 |
| Q8K2B3 | Succinate dehydrogenase [ubiquinone] flavoprotein subunit, mitoc | 33 | 20 | 39 | 20 | 664 | 72,5 | 7,37 | 109,98 | 20 | 1,023 | 0,975 | 0,97 | 0,935 | 1,044 | 1,077 | 0,999 | 0,997 | 0,95 | -0,05 | 8,9E-02 |
| Q9CPU0 | Lactoylglutathione lyase OS=Mus musculus OX=10090 GN=Glo1 PE= | 9 | 1 | 1 | 1 | 184 | 20,8 | 5,47 | 4,32 | 1 | 1,247 | 0,848 | 0,852 | 1,236 | 1,105 | 1,446 | 1,297 | 1,692 | 0,76 | -0,24 | 9,0E-02 |
| P69566 | Ran-binding protein 9 OS=Mus musculus OX=10090 GN=Ranbp9 PE | 5 | 2 | 2 | 2 | 653 | 71 | 6,84 | 6,09 | 2 | 2,083 | 1,604 | 1,352 | 1,21 | 1,173 | 1,231 | 1,071 | 1,198 | 1,34 | 0,34 | 9,0E-02 |
| P40142 | Transketolase OS=Mus musculus OX=10090 GN=Tkt PE=1 SV=1 | 34 | 19 | 25 | 19 | 623 | 67,6 | 7,5 | 57,8 | 19 | 1,147 | 1,094 | 1,058 | 1,032 | 0,96 | 1,069 | 1,005 | 1,025 | 1,07 | 0,07 | 9,0E-02 |
| Q8BFT9 | Synaptic vesicle 2-related protein OS=Mus musculus OX=10090 GN | 6 | 2 | 2 | 2 | 548 | 60,7 | 5,85 | 7,77 | 2 | 1,23 | 1,442 | 1,277 | 1,309 | 1,083 | 1,266 | 1,227 | 1,197 | 1,10 | 0,10 | 9,0E-02 |
| Q8K0D5 | Elongation factor G, mitochondrial OS=Mus musculus OX=10090 GI | 8 | 5 | 9 | 5 | 751 | 83,5 | 6,92 | 11,82 | 5 | 1,029 | 1,011 | 1,065 | 1,12 | 1,253 | 1,287 | 1,144 | 1,037 | 0,89 | -0,11 | 9,1E-02 |
| Q925I1 | ATPase family AAA domain-containing protein 3 OS=Mus musculus | 26 | 13 | 20 | 13 | 591 | 66,7 | 9,29 | 37,83 | 13 | 0,83 | 0,879 | 0,91 | 0,928 | 1,003 | 0,868 | 0,984 | 1,065 | 0,90 | -0,10 | 9,1E-02 |
| Q8CGM1 | Adhesion G protein-coupled receptor B2 OS=Mus musculus OX=10 | 4 | 4 | 5 | 3 | 1561 | 169,8 | 7,36 | 5,26 | 4 | 1,494 | 1,336 | 0,851 | 1,642 | 1,549 | 2,361 | 1,687 | 1,729 | 0,73 | -0,27 | 9,1E-02 |
| Q8R050 | Eukaryotic peptide chain release factor GTP-binding subunit ERF3A | 16 | 11 | 14 | 4 | 636 | 68,6 | 5,21 | 21,88 | 11 | 1,1 | 1,181 | 1,172 | 0,954 | 0,863 | 1,098 | 0,941 | 0,922 | 1,15 | 0,15 | 9,1E-02 |
| Q9DBL1 | Short/branched chain specific acyl-CoA dehydrogenase, mitochond | 12 | 4 | 4 | 4 | 432 | 47,8 | 7,87 | 11,2 | 4 | 0,956 | 1,039 | 0,994 | 1,059 | 1,122 | 1,176 | 1,027 | 1,052 | 0,92 | -0,08 | 9,2E-02 |
| P41216 | Long-chain-fatty-acid--CoA ligase 1 OS=Mus musculus OX=10090 GI | 13 | 8 | 9 | 7 | 699 | 77,9 | 7,15 | 23,9 | 8 | 0,882 | 0,846 | 0,939 | 0,995 | 1,062 | 0,924 | 1,019 | 1,007 | 0,91 | -0,09 | 9,2E-02 |
| O35226 | 26S proteasome non-ATPase regulatory subunit 4 OS=Mus musculu | 10 | 3 | 3 | 3 | 376 | 40,7 | 4,79 | 5,79 | 3 | 1,317 | 1,147 | 1,078 | 1,117 | 1,108 | 0,972 | 0,987 | 1,087 | 1,12 | 0,12 | 9,2E-02 |
| Q91V57 | N-chimaerin OS=Mus musculus OX=10090 GN=Chn1 PE=1 SV=2 | 3 | 1 | 1 | 1 | 459 | 53,2 | 6,76 | 2,38 | 1 | 1,845 | 1,369 | 1,613 | 1,244 | 1,129 | 1,273 | 1,397 | 0,868 | 1,30 | 0,30 | 9,2E-02 |
| Q9DB77 | Cytochrome b-c1 complex subunit 2, mitochondrial OS=Mus muscu | 39 | 15 | 42 | 15 | 453 | 48,2 | 9,25 | 119,04 | 15 | 0,892 | 0,9 | 0,996 | 0,946 | 1,005 | 0,954 | 1,015 | 0,981 | 0,94 | -0,06 | 9,2E-02 |
| Q6KAR6 | Exocyst complex component 3 OS=Mus musculus OX=10090 GN=E | 11 | 8 | 9 | 8 | 755 | 86,4 | 6,2 | 18,96 | 8 | 0,887 | 0,837 | 0,933 | 1,026 | 1,09 | 1,066 | 1,039 | 0,924 | 0,89 | -0,11 | 9,2E-02 |
| Q8C7K6 | Prenylcysteine oxidase-like OS=Mus musculus OX=10090 GN=Pcyo | 5 | 3 | 3 | 3 | 495 | 54,8 | 7,65 | 4,59 | 3 | 1,181 | 1,082 | 1,309 | 1,042 | 1,035 | 1,111 | 0,944 | 0,956 | 1,14 | 0,14 | 9,3E-02 |
| Q810B7 | SLIT and NTRK-like protein 5 OS=Mus musculus OX=10090 GN=Slitr | 1 | 1 | 1 | 1 | 957 | 107,1 | 6,93 | 2,93 | 1 | 0,803 | 0,726 | 0,854 | 0,883 | 1,018 | 1,022 | 0,97 | 0,783 | 0,86 | -0,14 | 9,3E-02 |
| P39447 | Tight junction protein ZO-1 OS=Mus musculus OX=10090 GN=Tjp1 | 6 | 9 | 9 | 9 | 1745 | 194,6 | 6,64 | 15,74 | 9 | 1,063 | 1,142 | 1,04 | 0,984 | 0,864 | 0,96 | 1,02 | 1,003 | 1,10 | 0,10 | 9,3E-02 |
| B2RUJ5 | Amyloid-beta A4 precursor protein-binding family A member 1 OS= | 8 | 5 | 5 | 5 | 842 | 92,9 | 4,88 | 12,04 | 5 | 0,857 | 0,838 | 0,797 | 0,898 | 0,842 | 1,059 | 1,013 | 0,905 | 0,89 | -0,11 | 9,3E-02 |
| Q8CD19 | LanC-like protein 3 OS=Mus musculus OX=10090 GN=Lancl3 PE=2 S | 3 | 1 | 1 | 1 | 420 | 46,4 | 7,05 | 0 | 1 | 1,01 | 0,891 | 1,012 | 1,176 | 0,816 | 0,81 | 0,901 | 0,988 | 1,16 | 0,16 | 9,3E-02 |
| P26450 | Phosphatidylinositol 3-kinase regulatory subunit alpha OS=Mus mu | 3 | 2 | 2 | 2 | 724 | 83,5 | 6,28 | 0 | 2 | 0,818 | 1,153 | 0,864 | 1,029 | 1,027 | 1,206 | 1,311 | 1,106 | 0,83 | -0,17 | 9,3E-02 |
| P58069 | Ras GTPase-activating protein 2 OS=Mus musculus OX=10090 GN= | 1 | 2 | 2 | 2 | 847 | 96,3 | 7,59 | 0 | 2 | 1,205 | 0,922 | 1,165 | 1,433 | 0,882 | 0,6 | 1,046 | 1,028 | 1,33 | 0,33 | 9,4E-02 |
| P12815 | Programmed cell death protein 6 OS=Mus musculus OX=10090 GN= | 18 | 3 | 3 | 3 | 191 | 21,9 | 5,4 | 8,01 | 3 | 1,169 | 1,295 | 1,352 | 1,283 | 1,192 | 0,96 | 1,275 | 0,988 | 1,15 | 0,15 | 9,4E-02 |
| P62274 | 40S ribosomal protein S29 OS=Mus musculus OX=10090 GN=Rps29 | 13 | 1 | 1 | 1 | 56 | 6,7 | 10,13 | 1,99 | 1 | 0,934 | 0,738 | 1,033 | 0,946 | 1,248 | 1,04 | 0,996 | 1,037 | 0,84 | -0,16 | 9,4E-02 |
| Q61425 | Hydroxyacyl-coenzyme A dehydrogenase, mitochondrial OS=Mus n | 11 | 4 | 4 | 4 | 314 | 34,4 | 8,65 | 7,88 | 4 | 0,712 | 0,867 | 0,995 | 0,912 | 1,06 | 0,89 | 1,039 | 1,082 | 0,86 | -0,14 | 9,4E-02 |
| Q80V42 | Carboxypeptidase M OS=Mus musculus OX=10090 GN=Cpm PE=1 S | 5 | 2 | 2 | 2 | 443 | 50,5 | 7,78 | 0 | 2 | 2,174 | 1,786 | 1,445 | 1,542 | 1,765 | 1,283 | 0,877 | 0,979 | 1,42 | 0,42 | 9,4E-02 |
| Q9CYK1 | Tryptophan--tRNA ligase, mitochondrial OS=Mus musculus OX=100 | 15 | 4 | 4 | 4 | 360 | 40,1 | 8,82 | 2,15 | 4 | 1,109 | 0,819 | 0,985 | 1,064 | 0,903 | 0,772 | 0,751 | 0,931 | 1,18 | 0,18 | 9,5E-02 |
| O08908 | Phosphatidylinositol 3-kinase regulatory subunit beta OS=Mus mus | 2 | 1 | 1 | 1 | 722 | 81,2 | 6,07 | 2,27 | 1 | 1,478 | 1,992 | 0,979 | 1,398 | 0,925 | 1,143 | 1,068 | 1,028 | 1,40 | 0,40 | 9,5E-02 |
| Q8VHI6 | Wiskott-Aldrich syndrome protein family member 3 OS=Mus muscu | 4 | 2 | 3 | 1 | 501 | 55,2 | 6,37 | 6,17 | 2 | 1,655 | 1,366 | 1,015 | 1,546 | 1,964 | 1,889 | 1,392 | 1,843 | 0,79 | -0,21 | 9,6E-02 |
| Q9EQF6 | Dihydropyrimidinase-related protein 5 OS=Mus musculus OX=1009 | 22 | 12 | 20 | 12 | 564 | 61,5 | 7,09 | 38,92 | 12 | 1,027 | 1,196 | 0,981 | 0,963 | 0,829 | 0,932 | 0,968 | 0,951 | 1,13 | 0,13 | 9,6E-02 |
| Q61290 | Voltage-dependent R-type calcium channel subunit alpha-1E OS=M | 6 | 10 | 10 | 10 | 2272 | 257,1 | 8,18 | 16,69 | 10 | 1,226 | 1,183 | 1,16 | 1,073 | 1,002 | 1,155 | 0,943 | 1,092 | 1,11 | 0,11 | 9,6E-02 |
| P50429 | Arylsulfatase B OS=Mus musculus OX=10090 GN=Arsb PE=1 SV=3 | 7 | 3 | 3 | 3 | 534 | 59,6 | 7,24 | 2,24 | 3 | 1,838 | 1,245 | 1,329 | 1,705 | 1,135 | 1,14 | 0,86 | 1,476 | 1,33 | 0,33 | 9,6E-02 |
| P14824 | Annexin A6 OS=Mus musculus OX=10090 GN=Anxa6 PE=1 SV=3 | 21 | 13 | 20 | 13 | 673 | 75,8 | 5,5 | 47 | 13 | 0,973 | 0,902 | 0,958 | 0,944 | 1,022 | 0,926 | 1,037 | 1,031 | 0,94 | -0,06 | 9,6E-02 |
| P62911 | 60S ribosomal protein L32 OS=Mus musculus OX=10090 GN=Rpl32 | 15 | 2 | 2 | 2 | 135 | 15,9 | 11,33 | 4,72 | 2 | 0,894 | 1,082 | 0,949 | 1,018 | 1,023 | 1,054 | 1,112 | 1,205 | 0,90 | -0,10 | 9,6E-02 |
| P67778 | Prohibitin OS=Mus musculus OX=10090 GN=Phb PE=1 SV=1 | 42 | 12 | 18 | 12 | 272 | 29,8 | 5,76 | 40,01 | 12 | 0,93 | 0,941 | 0,942 | 0,876 | 0,982 | 0,932 | 0,95 | 0,982 | 0,96 | -0,04 | 9,7E-02 |
| Q9R1L5 | Microtubule-associated serine/threonine-protein kinase 1 OS=Mus | 5 | 6 | 7 | 6 | 1570 | 170,9 | 8,44 | 5,58 | 6 | 1,231 | 1,049 | 0,944 | 1,011 | 1,034 | 0,807 | 0,772 | 0,943 | 1,19 | 0,19 | 9,7E-02 |
| Q60809 | CCR4-NOT transcription complex subunit 7 OS=Mus musculus OX=1 | 4 | 1 | 1 | 1 | 285 | 32,7 | 4,84 | 0 | 1 | 0,706 | 0,561 | 0,947 | 1,024 | 0,83 | 1,402 | 1,101 | 1,151 | 0,72 | -0,28 | 9,7E-02 |
| P54775 | 26S proteasome regulatory subunit 6B OS=Mus musculus OX=1009 | 15 | 8 | 8 | 8 | 418 | 47,4 | 5,21 | 14,74 | 8 | 0,843 | 0,896 | 0,989 | 0,97 | 1,075 | 0,946 | 1,04 | 0,984 | 0,91 | -0,09 | 9,8E-02 |
| Q9EQC5 | N-terminal kinase-like protein OS=Mus musculus OX=10090 GN=Sc | 5 | 3 | 3 | 3 | 806 | 89,1 | 6,44 | 5,32 | 3 | 1,328 | 1,267 | 1,432 | 1,254 | 1,13 | 1,361 | 1,065 | 1,123 | 1,13 | 0,13 | 9,8E-02 |
| Q9CQ86 | Migration and invasion enhancer 1 OS=Mus musculus OX=10090 G | 10 | 1 | 1 | 1 | 115 | 12,3 | 4,51 | 2,65 | 1 | 1,027 | 0,686 | 0,814 | 0,643 | 0,782 | 1,151 | 1,286 | 1,026 | 0,75 | -0,25 | 9,8E-02 |
| O70133 | ATP-dependent RNA helicase A OS=Mus musculus OX=10090 GN=D | 5 | 6 | 8 | 6 | 1380 | 149,4 | 6,83 | 7,6 | 6 | 1,024 | 1,273 | 1,012 | 0,95 | 1,295 | 1,396 | 1,086 | 1,235 | 0,85 | -0,15 | 9,9E-02 |
| Q99P31 | Hsp70-binding protein 1 OS=Mus musculus OX=10090 GN=Hspbp1 | 14 | 3 | 3 | 3 | 357 | 39,1 | 5,36 | 2,2 | 3 | 1,307 | 1,911 | 1,398 | 1,46 | 1,026 | 1,565 | 0,874 | 1,028 | 1,35 | 0,35 | 9,9E-02 |
| Q80SW1 | S-adenosylhomocysteine hydrolase-like protein 1 OS=Mus musculu | 26 | 14 | 26 | 8 | 530 | 58,9 | 6,89 | 43,98 | 14 | 0,868 | 0,756 | 0,954 | 0,916 | 0,992 | 0,941 | 0,989 | 0,93 | 0,91 | -0,09 | 9,9E-02 |
| Q8K2L8 | Trafficking protein particle complex subunit 12 OS=Mus musculus C | 6 | 4 | 6 | 4 | 797 | 87,6 | 4,81 | 11,81 | 4 | 0,972 | 0,934 | 0,869 | 0,967 | 0,947 | 0,982 | 1,019 | 1,033 | 0,94 | -0,06 | 9,9E-02 |
| P58321 | Ubiquitin carboxyl-terminal hydrolase isozyme L4 OS=Mus musculu | 10 | 3 | 5 | 3 | 233 | 26,4 | 5,1 | 13,93 | 3 | 1,147 | 1,227 | 1,269 | 1,301 | 0,978 | 0,956 | 1,252 | 1,148 | 1,14 | 0,14 | 9,9E-02 |
| P18872 | Guanine nucleotide-binding protein G(o) subunit alpha OS=Mus mu | 44 | 17 | 52 | 16 | 354 | 40,1 | 5,53 | 117,04 | 17 | 0,911 | 0,942 | 0,889 | 1,01 | 1,053 | 0,928 | 1,038 | 1,039 | 0,92 | -0,08 | 9,9E-02 |
| P10605 | Cathepsin B OS=Mus musculus OX=10090 GN=Ctsb PE=1 SV=2 | 13 | 4 | 7 | 4 | 339 | 37,3 | 5,91 | 6,64 | 4 | 0,96 | 0,644 | 0,921 | 0,974 | 1,052 | 0,954 | 1,244 | 1,024 | 0,82 | -0,18 | 9,9E-02 |
| P40237 | CD82 antigen OS=Mus musculus OX=10090 GN=Cd82 PE=1 SV=1 | 9 | 2 | 2 | 2 | 266 | 29,6 | 5,02 | 3,34 | 2 | 0,958 | 0,921 | 1,087 | 0,843 | 0,891 | 0,865 | 0,839 | 0,76 | 1,14 | 0,14 | 9,9E-02 |
| Q8BWT5 | Disco-interacting protein 2 homolog A OS=Mus musculus OX=1009 | 5 | 7 | 10 | 7 | 1523 | 165,1 | 7,61 | 20,93 | 7 | 1,076 | 1,077 | 1,105 | 1,117 | 1,004 | 1,115 | 0,966 | 1,031 | 1,06 | 0,06 | 9,9E-02 |
| Q9CQA3 | Succinate dehydrogenase [ubiquinone] iron-sulfur subunit, mitoch | 34 | 11 | 15 | 11 | 282 | 31,8 | 8,68 | 34,38 | 11 | 1,005 | 1,032 | 0,916 | 0,983 | 1,111 | 1,03 | 1,024 | 1,025 | 0,94 | -0,06 | 1,0E-01 |
| Q99KK7 | Dipeptidyl peptidase 3 OS=Mus musculus OX=10090 GN=Dpp3 PE= | 7 | 5 | 7 | 5 | 738 | 82,8 | 5,38 | 15,14 | 5 | 1,133 | 0,955 | 0,978 | 1,005 | 0,938 | 0,93 | 0,945 | 0,947 | 1,08 | 0,08 | 1,0E-01 |

| | | | | | | | | | | | | | | | | | | | | | |
|---|---|---|---|---|---|---|---|---|---|---|---|---|---|---|---|---|---|---|---|---|---|
| P26638 | Serine--tRNA ligase, cytoplasmic OS=Mus musculus OX=10090 GN= | 19 | 8 | 10 | 8 | 512 | 58,4 | 6,3 | 20,44 | 8 | 0,97 | 0,96 | 1,074 | 1,066 | 0,94 | 0,85 | 0,99 | 0,958 | 1,09 | 0,09 | 1,0E-01 |
| P52795 | Ephrin-B1 OS=Mus musculus OX=10090 GN=Efnb1 PE=1 SV=1 | 5 | 1 | 2 | 1 | 345 | 37,8 | 9,03 | 6,97 | 1 | 1,736 | 1,455 | 1,493 | 1,24 | 0,743 | 1,65 | 0,815 | 0,905 | 1,44 | 0,44 | 1,0E-01 |
| Q8BP40 | Lysophosphatidic acid phosphatase type 6 OS=Mus musculus OX=1 | 13 | 5 | 5 | 5 | 418 | 47,6 | 7,72 | 4,94 | 5 | 0,794 | 0,905 | 0,984 | 0,903 | 0,898 | 1,289 | 0,987 | 1,151 | 0,83 | -0,17 | 1,0E-01 |
| Q70IV5 | Synemin OS=Mus musculus OX=10090 GN=Synm PE=1 SV=2 | 0 | 1 | 1 | 1 | 1561 | 173,1 | 5,12 | 1,62 | 1 | 1,04 | 1,083 | 1,008 | 1,192 | 1,037 | 0,947 | 1,018 | 0,97 | 1,09 | 0,09 | 1,0E-01 |
| Q6DFV7 | Nuclear receptor coactivator 7 OS=Mus musculus OX=10090 GN=N | 1 | 1 | 1 | 1 | 943 | 106,3 | 5,43 | 0 | 1 | 0,863 | 0,766 | 0,991 | 0,69 | 0,884 | 1,256 | 1,043 | 0,939 | 0,80 | -0,20 | 1,0E-01 |
| Q80XU3 | Nuclear ubiquitous casein and cyclin-dependent kinase substrate 1 | 4 | 1 | 1 | 1 | 234 | 26,3 | 5,14 | 2,19 | 1 | 1,532 | 0,978 | 1,115 | 0,857 | 0,891 | 0,853 | 0,755 | 0,822 | 1,35 | 0,35 | 1,0E-01 |
| Q60603 | Potassium voltage-gated channel subfamily H member 1 OS=Mus n | 5 | 4 | 4 | 4 | 989 | 111,2 | 7,47 | 4,58 | 4 | 0,934 | 1,278 | 1,162 | 1,11 | 0,825 | 1,134 | 0,824 | 0,908 | 1,21 | 0,21 | 1,0E-01 |
| Q04735 | Cyclin-dependent kinase 16 OS=Mus musculus OX=10090 GN=Cdk1 | 4 | 2 | 2 | 1 | 496 | 55,9 | 7,87 | 4,57 | 2 | 0,858 | 0,64 | 0,88 | 0,697 | 0,993 | 1,026 | 0,756 | 0,957 | 0,82 | -0,18 | 1,0E-01 |
| Q62261 | Spectrin beta chain, non-erythrocytic 1 OS=Mus musculus OX=1009 | 54 | 120 | 197 | 114 | 2363 | 274,1 | 5,58 | 452,64 | 120 | 0,968 | 0,969 | 0,966 | 0,995 | 1,005 | 0,998 | 1,023 | 0,971 | 0,98 | -0,02 | 1,0E-01 |
| Q9JJC6 | RILP-like protein 1 OS=Mus musculus OX=10090 GN=Rilpl1 PE=1 SV | 5 | 2 | 2 | 2 | 406 | 47,3 | 5,16 | 0 | 2 | 1,243 | 1,397 | 1,532 | 1,03 | 1,152 | 0,869 | 1,212 | 0,892 | 1,26 | 0,26 | 1,0E-01 |
| Q5DU37 | Zinc finger FYVE domain-containing protein 26 OS=Mus musculus C | 0 | 1 | 1 | 1 | 2529 | 282,8 | 6,21 | 0 | 1 | 0,994 | 0,546 | 0,8 | 0,775 | 1,041 | 0,87 | 0,917 | 1,131 | 0,79 | -0,21 | 1,0E-01 |
| Q8R2Y2 | Cell surface glycoprotein MUC18 OS=Mus musculus OX=10090 GN= | 3 | 1 | 1 | 1 | 648 | 71,5 | 5,83 | 4,15 | 1 | 1,172 | 1,163 | 1,125 | 1,017 | 0,792 | 1,049 | 1,039 | 1,04 | 1,14 | 0,14 | 1,0E-01 |
| P62141 | Serine/threonine-protein phosphatase PP1-beta catalytic subunit C | 28 | 8 | 12 | 4 | 327 | 37,2 | 6,19 | 26,03 | 8 | 1,3 | 1,602 | 1,36 | 1,162 | 0,98 | 1,39 | 1,032 | 0,979 | 1,24 | 0,24 | 1,0E-01 |
| Q9CR25 | 2-(3-amino-3-carboxypropyl)histidine synthase subunit 2 OS=Mus r | 2 | 1 | 1 | 1 | 489 | 52,3 | 5,53 | 1,61 | 1 | 1,779 | 1,825 | 1,644 | 1,455 | 0,82 | 1,013 | 1,063 | 1,883 | 1,40 | 0,40 | 1,0E-01 |
| O08539-2 | Isoform 2 of Myc box-dependent-interacting protein 1 OS=Mus mu | 32 | 12 | 21 | 1 | 434 | 48 | 5,15 | 48,21 | 12 | 1,183 | 0,968 | 1,136 | 0,875 | 0,819 | 0,816 | 1,01 | 0,86 | 1,19 | 0,19 | 1,0E-01 |
| Q01768 | Nucleoside diphosphate kinase B OS=Mus musculus OX=10090 GN= | 45 | 6 | 7 | 3 | 152 | 17,4 | 7,5 | 16,73 | 6 | 0,832 | 0,94 | 0,814 | 1,023 | 1,068 | 0,902 | 1,157 | 1,037 | 0,87 | -0,13 | 1,0E-01 |
| P28271 | Cytoplasmic aconitate hydratase OS=Mus musculus OX=10090 GN= | 13 | 11 | 13 | 11 | 889 | 98,1 | 7,5 | 30,31 | 11 | 1,149 | 1,078 | 1,12 | 1,034 | 0,943 | 1,112 | 0,979 | 1,007 | 1,08 | 0,08 | 1,0E-01 |
| Q61036 | Serine/threonine-protein kinase PAK 3 OS=Mus musculus OX=1009 | 18 | 9 | 15 | 2 | 559 | 62,4 | 5,4 | 27,61 | 9 | 1,456 | 1,409 | 1,743 | 1,447 | 0,975 | 1,471 | 1,157 | 1,397 | 1,21 | 0,21 | 1,0E-01 |
| F8VPU2 | FERM, ARHGEF and pleckstrin domain-containing protein 1 OS=Mu | 5 | 4 | 4 | 4 | 1048 | 118,8 | 7,88 | 6,44 | 4 | 2,176 | 1,729 | 1,916 | 1,505 | 1,006 | 1,723 | 1,142 | 1,675 | 1,32 | 0,32 | 1,0E-01 |
| Q80UF9 | Proton channel OTOP3 OS=Mus musculus OX=10090 GN=Otop3 PE | 2 | 1 | 1 | 1 | 577 | 64,7 | 8,06 | 2,36 | 1 | 0,948 | 1,634 | 1,434 | 0,988 | 0,952 | 0,829 | 1,063 | 0,772 | 1,38 | 0,38 | 1,0E-01 |
| Q8VCL2 | Protein SCO2 homolog, mitochondrial OS=Mus musculus OX=1009( | 4 | 1 | 1 | 1 | 255 | 28,9 | 8,29 | 3,13 | 1 | 0,915 | 1,138 | 0,905 | 0,817 | 1,177 | 1,019 | 1,035 | 1,153 | 0,86 | -0,14 | 1,0E-01 |
| B9EKR1 | Receptor-type tyrosine-protein phosphatase zeta OS=Mus musculu | 0 | 1 | 1 | 1 | 2312 | 254,2 | 4,88 | 1,84 | 1 | 1,077 | 0,972 | 0,832 | 1,006 | 0,896 | 0,761 | 0,89 | 0,876 | 1,14 | 0,14 | 1,0E-01 |
| Q01815 | Voltage-dependent L-type calcium channel subunit alpha-1C OS=M | 1 | 2 | 2 | 2 | 2139 | 240 | 6,87 | 2,7 | 2 | 0,733 | 0,94 | 1,168 | 0,93 | 0,903 | 1,263 | 1,304 | 1,321 | 0,79 | -0,21 | 1,0E-01 |
| P17427 | AP-2 complex subunit alpha-2 OS=Mus musculus OX=10090 GN=Ap | 22 | 23 | 41 | 16 | 938 | 104 | 6,93 | 100,85 | 23 | 0,82 | 0,908 | 0,924 | 0,932 | 0,927 | 0,925 | 1,036 | 0,978 | 0,93 | -0,07 | 1,0E-01 |
| O35874 | Neutral amino acid transporter A OS=Mus musculus OX=10090 GN= | 3 | 3 | 3 | 3 | 532 | 56 | 5,87 | 5,07 | 3 | 0,952 | 1,038 | 0,999 | 0,873 | 0,88 | 0,92 | 0,894 | 0,888 | 1,08 | 0,08 | 1,0E-01 |
| Q80XN0 | D-beta-hydroxybutyrate dehydrogenase, mitochondrial OS=Mus m | 30 | 10 | 20 | 10 | 343 | 38,3 | 9,01 | 35,29 | 10 | 0,932 | 0,98 | 1,011 | 0,972 | 0,906 | 0,967 | 0,894 | 0,947 | 1,05 | 0,05 | 1,0E-01 |
| P97792 | Coxsackievirus and adenovirus receptor homolog OS=Mus musculu | 19 | 7 | 8 | 7 | 365 | 39,9 | 6,96 | 10,57 | 7 | 1,032 | 1,014 | 1,066 | 1,269 | 0,966 | 0,993 | 0,94 | 1,014 | 1,12 | 0,12 | 1,0E-01 |
| Q9QYB1 | Chloride intracellular channel protein 4 OS=Mus musculus OX=100 | 25 | 5 | 6 | 5 | 253 | 28,7 | 5,59 | 4,81 | 5 | 0,915 | 0,77 | 0,865 | 0,909 | 1,018 | 1,104 | 0,894 | 0,904 | 0,88 | -0,12 | 1,0E-01 |
| Q0KL02 | Triple functional domain protein OS=Mus musculus OX=10090 GN= | 10 | 29 | 35 | 23 | 3102 | 347,6 | 6,35 | 55,28 | 29 | 1,108 | 1,017 | 1,022 | 1,012 | 0,998 | 0,989 | 0,959 | 1,016 | 1,05 | 0,05 | 1,0E-01 |
| Q9Z1T6-1 | Isoform 2 of 1-phosphatidylinositol 3-phosphate 5-kinase OS=Mus | 2 | 3 | 3 | 3 | 2052 | 231,9 | 6,6 | 0 | 3 | 1,35 | 1,196 | 1,252 | 1,299 | 1,267 | 1,208 | 1,029 | 1,124 | 1,10 | 0,10 | 1,0E-01 |
| Q9WVA3 | Mitotic checkpoint protein BUB3 OS=Mus musculus OX=10090 GN= | 3 | 1 | 1 | 1 | 326 | 36,9 | 6,84 | 0 | 1 | 1,499 | 1,83 | 1,156 | 0,928 | 1,66 | 2,052 | 1,526 | 1,955 | 0,75 | -0,25 | 1,0E-01 |
| Q8BGY9 | High affinity choline transporter 1 OS=Mus musculus OX=10090 GN | 2 | 1 | 1 | 1 | 580 | 63,3 | 5,33 | 0 | 1 | 0,967 | 1,484 | 1,536 | 1,117 | 0,839 | 0,777 | 1,23 | 0,949 | 1,34 | 0,34 | 1,0E-01 |
| P35803-8 | Isoform 8 of Neuronal membrane glycoprotein M6-b OS=Mus musc | 20 | 9 | 15 | 8 | 288 | 31,6 | 5,35 | 28,97 | 9 | 0,693 | 0,834 | 0,654 | 0,918 | 1,022 | 0,753 | 1,126 | 0,96 | 0,80 | -0,20 | 1,0E-01 |
| Q9Z140 | Copine-6 OS=Mus musculus OX=10090 GN=Cpne6 PE=1 SV=1 | 13 | 7 | 8 | 7 | 557 | 61,7 | 5,59 | 22,45 | 7 | 0,867 | 0,759 | 0,95 | 1,046 | 1,059 | 1,033 | 1,091 | 0,955 | 0,88 | -0,12 | 1,0E-01 |
| Q8CBF3 | Ephrin type-B receptor 1 OS=Mus musculus OX=10090 GN=Ephb1 F | 5 | 5 | 5 | 3 | 984 | 109,8 | 6,4 | 6,29 | 5 | 0,829 | 0,742 | 1,035 | 0,935 | 0,993 | 1,109 | 0,935 | 1,076 | 0,86 | -0,14 | 1,0E-01 |
| Q91VH2 | Sorting nexin-9 OS=Mus musculus OX=10090 GN=Snx9 PE=1 SV=1 | 2 | 1 | 1 | 1 | 595 | 66,5 | 5,52 | 2,86 | 1 | 1,068 | 1,25 | 1,058 | 1,286 | 1,221 | 1,255 | 1,445 | 1,334 | 0,89 | -0,11 | 1,0E-01 |
| G5E829 | Plasma membrane calcium-transporting ATPase 1 OS=Mus musculu | 34 | 39 | 77 | 22 | 1220 | 134,7 | 5,91 | 183,88 | 39 | 0,958 | 0,952 | 0,974 | 0,955 | 1,035 | 0,962 | 1,056 | 0,969 | 0,95 | -0,05 | 1,1E-01 |
| Q8VDZ4 | Palmitoyltransferase ZDHHC5 OS=Mus musculus OX=10090 GN=Zd | 2 | 1 | 1 | 1 | 715 | 77,5 | 9,01 | 1,68 | 1 | 4,083 | 3,533 | 2,07 | 2,862 | 2,752 | 2,008 | 2,159 | 2,076 | 1,39 | 0,39 | 1,1E-01 |
| P63143 | Voltage-gated potassium channel subunit beta-1 OS=Mus musculus | 10 | 4 | 5 | 2 | 401 | 44,7 | 9,32 | 9,46 | 4 | 1,101 | 1,401 | 1,109 | 1,321 | 1,242 | 0,839 | 0,97 | 1,021 | 1,21 | 0,21 | 1,1E-01 |
| Q8BGT8 | Phytanoyl-CoA hydroxylase-interacting protein-like OS=Mus muscu | 23 | 7 | 9 | 7 | 375 | 42,3 | 6,35 | 22,8 | 7 | 0,944 | 1,015 | 0,951 | 1,052 | 0,991 | 1,144 | 1,058 | 1,08 | 0,93 | -0,07 | 1,1E-01 |
| Q8JZN5 | Complex I assembly factor ACAD9, mitochondrial OS=Mus musculu | 19 | 11 | 17 | 11 | 625 | 68,7 | 7,46 | 40,35 | 11 | 1,287 | 1,245 | 1,18 | 1,146 | 0,985 | 1,181 | 0,995 | 1,197 | 1,11 | 0,11 | 1,1E-01 |
| Q9Z1A9 | TBC1 domain family member 8 OS=Mus musculus OX=10090 GN=T | 1 | 1 | 1 | 1 | 1134 | 130 | 5,48 | 2,17 | 1 | 1,053 | 0,788 | 1,136 | 1,059 | 0,979 | 1,487 | 1,473 | 1,19 | 0,79 | -0,21 | 1,1E-01 |
| Q9Z110 | Delta-1-pyrroline-5-carboxylate synthase OS=Mus musculus OX=10 | 13 | 9 | 10 | 9 | 795 | 87,2 | 7,55 | 15,6 | 9 | 1,266 | 1,126 | 1,164 | 1,058 | 0,953 | 1,022 | 1,029 | 1,153 | 1,11 | 0,11 | 1,1E-01 |
| Q61792 | LIM and SH3 domain protein 1 OS=Mus musculus OX=10090 GN=La | 40 | 10 | 17 | 9 | 263 | 30 | 7,05 | 26,12 | 10 | 1,026 | 1,094 | 1,054 | 1,115 | 0,918 | 1,033 | 1,025 | 1,044 | 1,07 | 0,07 | 1,1E-01 |
| P63216 | Guanine nucleotide-binding protein G(I)/G(S)/G(O) subunit gamma | 72 | 4 | 10 | 4 | 75 | 8,3 | 7,78 | 15,8 | 4 | 1,181 | 1,156 | 1,196 | 1,109 | 0,959 | 1,113 | 1,043 | 1,162 | 1,09 | 0,09 | 1,1E-01 |
| Q8K2Y9 | Cerebral cavernous malformations protein 2 homolog OS=Mus mus | 5 | 2 | 3 | 2 | 453 | 49,9 | 5,62 | 2,09 | 2 | 0,694 | 0,944 | 0,979 | 0,782 | 1,061 | 0,894 | 1,052 | 0,981 | 0,85 | -0,15 | 1,1E-01 |
| P97798 | Neogenin OS=Mus musculus OX=10090 GN=Neo1 PE=1 SV=1 | 4 | 4 | 4 | 4 | 1493 | 163,1 | 6,47 | 2,64 | 4 | 1,809 | 1,293 | 1,202 | 0,879 | 0,706 | 1,118 | 0,799 | 0,947 | 1,45 | 0,45 | 1,1E-01 |
| Q8C5H8 | NAD kinase 2, mitochondrial OS=Mus musculus OX=10090 GN=Nad | 18 | 8 | 9 | 8 | 452 | 50,8 | 8,07 | 17,28 | 8 | 0,968 | 1,078 | 0,951 | 1,025 | 1,077 | 1,087 | 1,093 | 1,019 | 0,94 | -0,06 | 1,1E-01 |
| P0C7L0 | WAS/WASL-interacting protein family member 3 OS=Mus musculus | 2 | 1 | 2 | 1 | 485 | 49,4 | 10,18 | 3,3 | 1 | 1,197 | 1,349 | 1,267 | 0,908 | 0,814 | 1,116 | 0,912 | 1,004 | 1,23 | 0,23 | 1,1E-01 |
| Q6ZQ82 | Rho GTPase-activating protein 26 OS=Mus musculus OX=10090 GN | 7 | 4 | 5 | 4 | 814 | 92 | 6,68 | 5,61 | 4 | 1,367 | 1,285 | 1,084 | 1,046 | 0,833 | 1,134 | 0,966 | 1,077 | 1,19 | 0,19 | 1,1E-01 |
| P10637-3 | Isoform Tau-B of Microtubule-associated protein tau OS=Mus musc | 53 | 17 | 40 | 2 | 364 | 38,2 | 9,33 | 72,12 | 17 | 1,117 | 1,084 | 1,021 | 1,095 | 0,88 | 1,059 | 0,994 | 1,042 | 1,09 | 0,09 | 1,1E-01 |
| Q9QYJ0 | DnaJ homolog subfamily A member 2 OS=Mus musculus OX=10090 | 23 | 11 | 16 | 11 | 412 | 45,7 | 6,48 | 20,89 | 11 | 0,855 | 0,799 | 0,923 | 1,007 | 1,018 | 0,964 | 1,048 | 0,94 | 0,90 | -0,10 | 1,1E-01 |
| Q8K3W0 | BRISC and BRCA1-A complex member 2 OS=Mus musculus OX=100 | 9 | 3 | 4 | 3 | 383 | 43,5 | 5,94 | 5,65 | 3 | 1,113 | 0,892 | 1,121 | 1,033 | 0,926 | 0,817 | 0,951 | 0,982 | 1,13 | 0,13 | 1,1E-01 |
| Q9JLV5 | Cullin-3 OS=Mus musculus OX=10090 GN=Cul3 PE=1 SV=1 | 19 | 12 | 15 | 12 | 768 | 88,9 | 8,46 | 26,87 | 12 | 0,847 | 0,801 | 0,907 | 1,005 | 0,97 | 0,925 | 1,098 | 0,998 | 0,89 | -0,11 | 1,1E-01 |
| Q8CAK3 | Shiftless antiviral inhibitor of ribosomal frameshifting protein homo | 4 | 1 | 1 | 1 | 290 | 33 | 7,28 | 1,9 | 1 | 0,963 | 1,138 | 0,93 | 0,949 | 0,942 | 0,958 | 0,788 | 0,691 | 1,18 | 0,18 | 1,1E-01 |
| Q08460 | Calcium-activated potassium channel subunit alpha-1 OS=Mus mus | 9 | 9 | 10 | 9 | 1209 | 134,3 | 7,15 | 14,96 | 9 | 0,989 | 1,028 | 1,143 | 0,931 | 1,023 | 1,207 | 1,12 | 1,22 | 0,90 | -0,10 | 1,1E-01 |
| Q9D6F4 | Gamma-aminobutyric acid receptor subunit alpha-4 OS=Mus musc | 7 | 3 | 3 | 3 | 552 | 60,8 | 9,32 | 2,03 | 3 | 1,193 | 1,143 | 1,14 | 0,965 | 1,017 | 1,042 | 0,976 | 1,018 | 1,10 | 0,10 | 1,1E-01 |
| Q9CZN7 | Serine hydroxymethyltransferase, mitochondrial OS=Mus musculus | 6 | 3 | 3 | 3 | 504 | 55,7 | 8,47 | 6,37 | 3 | 1,183 | 1,136 | 1,176 | 0,962 | 0,98 | 0,939 | 0,876 | 1,117 | 1,14 | 0,14 | 1,1E-01 |
| Q9CZM2 | 60S ribosomal protein L15 OS=Mus musculus OX=10090 GN=Rpl15 | 19 | 3 | 3 | 3 | 204 | 24,1 | 11,62 | 8,5 | 3 | 1,082 | 1,045 | 1,133 | 1,021 | 1,072 | 0,997 | 0,988 | 0,984 | 1,06 | 0,06 | 1,1E-01 |
| Q9D0I9 | Arginine--tRNA ligase, cytoplasmic OS=Mus musculus OX=10090 GN | 12 | 9 | 10 | 9 | 660 | 75,6 | 7,55 | 19,67 | 9 | 0,89 | 0,94 | 0,958 | 1,046 | 1,032 | 0,952 | 1,109 | 1,129 | 0,91 | -0,09 | 1,1E-01 |
| P56812 | Programmed cell death protein 5 OS=Mus musculus OX=10090 GN= | 14 | 2 | 2 | 2 | 126 | 14,3 | 5,68 | 1,78 | 2 | 0,916 | 0,99 | 1,184 | 0,981 | 1,075 | 1,305 | 1,136 | 1,127 | 0,88 | -0,12 | 1,1E-01 |
| Q9JKF6 | Nectin-1 OS=Mus musculus OX=10090 GN=Nectin1 PE=1 SV=3 | 9 | 5 | 5 | 5 | 515 | 57 | 6,35 | 19,01 | 5 | 0,792 | 0,891 | 0,856 | 0,834 | 0,972 | 0,823 | 0,91 | 0,999 | 0,91 | -0,09 | 1,1E-01 |

| | | | | | | | | | | | | | | | | | | | | | |
|---|---|---|---|---|---|---|---|---|---|---|---|---|---|---|---|---|---|---|---|---|---|
| Q9QXL2 | Kinesin-like protein KIF21A OS=Mus musculus OX=10090 GN=Kif21a | 13 | 17 | 23 | 17 | 1672 | 186,4 | 6,27 | 57,99 | 17 | 1,096 | 1,032 | 1,1 | 1,078 | 0,964 | 1,062 | 1,032 | 1,048 | 1,05 | 0,05 | 1,1E-01 |
| Q9JL56 | Glycerophosphodiester phosphodiesterase 1 OS=Mus musculus OX | 18 | 6 | 11 | 6 | 331 | 37,6 | 6,9 | 18,52 | 6 | 0,978 | 1,04 | 1,089 | 1,014 | 0,974 | 1,029 | 0,857 | 0,941 | 1,08 | 0,08 | 1,1E-01 |
| Q9Z218 | Dipeptidyl aminopeptidase-like protein 6 OS=Mus musculus OX=10 | 26 | 20 | 34 | 18 | 804 | 91,2 | 6,68 | 85,68 | 20 | 0,91 | 0,95 | 0,898 | 0,974 | 0,979 | 0,939 | 1,055 | 0,982 | 0,94 | -0,06 | 1,1E-01 |
| Q69ZF3 | Non-lysosomal glucosylceramidase OS=Mus musculus OX=10090 G | 8 | 6 | 7 | 6 | 918 | 103,2 | 5,59 | 14,94 | 6 | 1,147 | 0,995 | 1,048 | 0,933 | 0,92 | 0,957 | 0,814 | 0,99 | 1,12 | 0,12 | 1,1E-01 |
| Q6PHU5 | Sortilin OS=Mus musculus OX=10090 GN=Sort1 PE=1 SV=1 | 6 | 4 | 6 | 4 | 825 | 91,1 | 5,88 | 12,58 | 4 | 1,185 | 1,132 | 0,977 | 0,906 | 0,85 | 1,042 | 0,843 | 0,863 | 1,17 | 0,17 | 1,1E-01 |
| G5E870 | E3 ubiquitin-protein ligase TRIP12 OS=Mus musculus OX=10090 GN | 1 | 1 | 1 | 1 | 2025 | 224 | 8,35 | 0 | 1 | 2,15 | 2,28 | 1,194 | 0,89 | 0,525 | 1,316 | 0,858 | 0,97 | 1,78 | 0,78 | 1,1E-01 |
| P09411 | Phosphoglycerate kinase 1 OS=Mus musculus OX=10090 GN=Pgk1 | 54 | 22 | 37 | 22 | 417 | 44,5 | 7,9 | 102,39 | 22 | 1,034 | 0,97 | 1,04 | 1,014 | 0,945 | 0,946 | 1,009 | 0,99 | 1,04 | 0,04 | 1,1E-01 |
| P20917 | Myelin-associated glycoprotein OS=Mus musculus OX=10090 GN=M | 5 | 4 | 8 | 4 | 627 | 69,3 | 5,1 | 15,33 | 4 | 1,153 | 1,13 | 1,122 | 0,948 | 0,988 | 1,07 | 0,943 | 0,928 | 1,11 | 0,11 | 1,1E-01 |
| Q9CZW5 | Mitochondrial import receptor subunit TOM70 OS=Mus musculus O | 33 | 22 | 28 | 22 | 611 | 67,5 | 7,53 | 64,08 | 22 | 0,869 | 0,828 | 0,929 | 0,93 | 1,025 | 0,877 | 1,075 | 0,95 | 0,91 | -0,09 | 1,1E-01 |
| Q9D710 | Thioredoxin-related transmembrane protein 2 OS=Mus musculus O | 18 | 4 | 4 | 4 | 295 | 33,9 | 8,75 | 8,39 | 4 | 1,179 | 1,16 | 1,118 | 1,006 | 1,02 | 1,019 | 0,978 | 1,101 | 1,08 | 0,08 | 1,1E-01 |
| Q8CAY6 | Acetyl-CoA acetyltransferase, cytosolic OS=Mus musculus OX=1009 | 5 | 2 | 2 | 2 | 397 | 41,3 | 7,5 | 0 | 2 | 1,213 | 1,138 | 0,921 | 1,109 | 1,157 | 1,741 | 1,265 | 1,283 | 0,80 | -0,20 | 1,1E-01 |
| Q8BGH7 | CDC42 small effector protein 2 OS=Mus musculus OX=10090 GN=C | 23 | 1 | 3 | 1 | 84 | 9,2 | 8,35 | 5,99 | 1 | 0,876 | 0,673 | 0,803 | 0,72 | 0,983 | 0,767 | 1,035 | 0,85 | 0,85 | -0,15 | 1,1E-01 |
| Q9D4H1 | Exocyst complex component 2 OS=Mus musculus OX=10090 GN=Ex | 8 | 7 | 7 | 7 | 924 | 103,9 | 7,18 | 18,53 | 7 | 1,053 | 0,869 | 0,917 | 0,973 | 1,063 | 1,019 | 1,007 | 1,029 | 0,93 | -0,07 | 1,1E-01 |
| Q61749 | Translation initiation factor eIF-2B subunit delta OS=Mus musculus | 8 | 3 | 3 | 3 | 524 | 57,6 | 9,25 | 2,1 | 3 | 1,538 | 1,486 | 1,365 | 1,454 | 1,537 | 1,268 | 1,082 | 1,188 | 1,15 | 0,15 | 1,1E-01 |
| O35526 | Syntaxin-1A OS=Mus musculus OX=10090 GN=Stx1a PE=1 SV=3 | 36 | 13 | 31 | 12 | 288 | 33 | 5,24 | 66,93 | 13 | 1,076 | 1,235 | 0,967 | 1,021 | 0,94 | 1,003 | 0,908 | 0,99 | 1,12 | 0,12 | 1,1E-01 |
| P12658 | Calbindin OS=Mus musculus OX=10090 GN=Calb1 PE=1 SV=2 | 34 | 10 | 12 | 10 | 261 | 30 | 4,83 | 21,53 | 10 | 1,121 | 1,032 | 0,993 | 1,05 | 0,9 | 1,022 | 0,967 | 1,018 | 1,07 | 0,07 | 1,1E-01 |
| Q505D7 | Optic atrophy 3 protein homolog OS=Mus musculus OX=10090 GN= | 4 | 1 | 1 | 1 | 179 | 20,1 | 8,34 | 2,26 | 1 | 0,624 | 0,727 | 0,732 | 0,938 | 1,227 | 0,622 | 1,113 | 1,241 | 0,72 | -0,28 | 1,1E-01 |
| Q91VW3 | SH3 domain-binding glutamic acid-rich-like protein 3 OS=Mus musc | 29 | 5 | 6 | 5 | 93 | 10,5 | 5,14 | 13,08 | 5 | 1,25 | 1,392 | 1,131 | 1,01 | 0,841 | 1,175 | 0,959 | 1,016 | 1,20 | 0,20 | 1,1E-01 |
| Q9Z0U0 | Xenotropic and polytropic retrovirus receptor 1 OS=Mus musculus | 3 | 2 | 3 | 2 | 695 | 81,7 | 8,24 | 4,41 | 2 | 1,063 | 1,243 | 1,024 | 0,925 | 0,996 | 0,903 | 0,96 | 0,853 | 1,15 | 0,15 | 1,1E-01 |
| Q9D8N0 | Elongation factor 1-gamma OS=Mus musculus OX=10090 GN=Eef1g | 12 | 5 | 5 | 5 | 437 | 50 | 6,74 | 12,84 | 5 | 1,088 | 1,048 | 1,001 | 0,988 | 0,995 | 0,983 | 0,989 | 0,988 | 1,04 | 0,04 | 1,1E-01 |
| Q91W90 | Thioredoxin domain-containing protein 5 OS=Mus musculus OX=10 | 4 | 2 | 2 | 2 | 417 | 46,4 | 5,78 | 1,97 | 2 | 0,804 | 0,943 | 0,872 | 0,956 | 1,17 | 0,882 | 1,046 | 0,99 | 0,87 | -0,13 | 1,1E-01 |
| Q9D6J5 | NADH dehydrogenase [ubiquinone] 1 beta subcomplex subunit 8, r | 24 | 3 | 5 | 3 | 186 | 21,9 | 6,64 | 7,68 | 3 | 0,785 | 1,131 | 0,855 | 1,087 | 1,135 | 1,228 | 1 | 1,251 | 0,84 | -0,16 | 1,1E-01 |
| Q9CQ22 | Ragulator complex protein LAMTOR1 OS=Mus musculus OX=10090 | 16 | 2 | 3 | 2 | 161 | 17,7 | 5,15 | 9,32 | 2 | 0,841 | 0,943 | 0,964 | 0,972 | 0,9 | 0,769 | 0,891 | 0,846 | 1,09 | 0,09 | 1,1E-01 |
| P20108 | Thioredoxin-dependent peroxide reductase, mitochondrial OS=Mu | 32 | 6 | 7 | 6 | 257 | 28,1 | 7,58 | 17,03 | 6 | 0,932 | 0,928 | 0,991 | 1,054 | 1,027 | 1,081 | 1,168 | 0,989 | 0,92 | -0,08 | 1,1E-01 |
| E9Q5F9 | Histone-lysine N-methyltransferase SETD2 OS=Mus musculus OX=1 | 0 | 1 | 1 | 1 | 2537 | 285,5 | 6,3 | 1,69 | 1 | 0,736 | 0,913 | 0,632 | 1,087 | 0,998 | 1,025 | 0,979 | 1,171 | 0,81 | -0,19 | 1,2E-01 |
| P54227 | Stathmin OS=Mus musculus OX=10090 GN=Stmn1 PE=1 SV=2 | 48 | 10 | 19 | 7 | 149 | 17,3 | 5,97 | 47,6 | 10 | 1,139 | 1,114 | 1,071 | 1,106 | 0,978 | 1,027 | 1,114 | 1,072 | 1,06 | 0,06 | 1,2E-01 |
| Q60930 | Voltage-dependent anion-selective channel protein 2 OS=Mus mus | 37 | 9 | 16 | 9 | 295 | 31,7 | 7,49 | 44,64 | 9 | 0,901 | 1,013 | 0,872 | 0,888 | 0,996 | 1,033 | 0,989 | 0,935 | 0,93 | -0,07 | 1,2E-01 |
| P26883 | Peptidyl-prolyl cis-trans isomerase FKBP1A OS=Mus musculus OX=1 | 38 | 3 | 6 | 3 | 108 | 11,9 | 8,16 | 15,6 | 3 | 1,573 | 1,274 | 1,173 | 1,013 | 0,817 | 1,268 | 0,891 | 0,917 | 1,29 | 0,29 | 1,2E-01 |
| Q9EPU0 | Regulator of nonsense transcripts 1 OS=Mus musculus OX=10090 G | 8 | 6 | 6 | 6 | 1124 | 123,9 | 6,61 | 13,98 | 6 | 0,962 | 0,992 | 0,983 | 0,974 | 1,146 | 1,036 | 1,001 | 0,992 | 0,94 | -0,06 | 1,2E-01 |
| P70451 | Tyrosine-protein kinase Fer OS=Mus musculus OX=10090 GN=Fer P | 1 | 1 | 1 | 1 | 823 | 94,5 | 7,05 | 0 | 1 | 1,487 | 1,642 | 1,465 | 1,079 | 1,075 | 1,316 | 1,203 | 1,117 | 1,20 | 0,20 | 1,2E-01 |
| P80314 | T-complex protein 1 subunit beta OS=Mus musculus OX=10090 GN | 28 | 13 | 18 | 13 | 535 | 57,4 | 6,4 | 34,49 | 13 | 1,066 | 1 | 1,069 | 1,031 | 1,022 | 0,993 | 1,023 | 0,993 | 1,03 | 0,03 | 1,2E-01 |
| Q91Z53 | Glyoxylate reductase/hydroxypyruvate reductase OS=Mus musculu | 19 | 6 | 7 | 6 | 328 | 35,3 | 7,65 | 17,14 | 6 | 1,17 | 1,137 | 0,991 | 1,012 | 0,889 | 1,033 | 0,879 | 1,045 | 1,12 | 0,12 | 1,2E-01 |
| Q8BGN8-2 | Isoform 2 of Synaptoporin OS=Mus musculus OX=10090 GN=Synpr | 13 | 4 | 5 | 4 | 285 | 31,3 | 6,33 | 10,16 | 4 | 1,25 | 1,433 | 1,153 | 1,156 | 0,971 | 1,19 | 1,004 | 1,185 | 1,15 | 0,15 | 1,2E-01 |
| Q9CYT6 | Adenylyl cyclase-associated protein 2 OS=Mus musculus OX=10090 | 35 | 15 | 22 | 14 | 476 | 52,8 | 6,43 | 52,1 | 15 | 1,147 | 1,05 | 1,013 | 1,032 | 0,94 | 0,993 | 0,964 | 1,058 | 1,07 | 0,07 | 1,2E-01 |
| Q8BUK6 | Protein Hook homolog 3 OS=Mus musculus OX=10090 GN=Hook3 F | 8 | 4 | 5 | 4 | 718 | 83,2 | 5,19 | 10,28 | 4 | 0,826 | 0,844 | 0,86 | 0,794 | 0,846 | 0,869 | 0,855 | 0,864 | 0,97 | -0,03 | 1,2E-01 |
| P47911 | 60S ribosomal protein L6 OS=Mus musculus OX=10090 GN=Rpl6 PE | 19 | 6 | 9 | 6 | 296 | 33,5 | 10,7 | 23,75 | 6 | 0,897 | 0,924 | 0,885 | 0,953 | 1,075 | 0,912 | 0,979 | 0,965 | 0,93 | -0,07 | 1,2E-01 |
| P03930 | ATP synthase protein 8 OS=Mus musculus OX=10090 GN=Mtatp8 P | 36 | 3 | 4 | 3 | 67 | 7,8 | 9,88 | 6,44 | 3 | 0,934 | 0,986 | 1,026 | 0,9 | 0,889 | 0,887 | 0,859 | 0,957 | 1,07 | 0,07 | 1,2E-01 |
| P60867 | 40S ribosomal protein S20 OS=Mus musculus OX=10090 GN=Rps20 | 19 | 2 | 2 | 2 | 119 | 13,4 | 9,94 | 2,76 | 2 | 0,873 | 0,834 | 1,08 | 0,941 | 0,854 | 1,248 | 1,297 | 1,155 | 0,82 | -0,18 | 1,2E-01 |
| Q8VHH5 | Arf-GAP with GTPase, ANK repeat and PH domain-containing prote | 4 | 4 | 4 | 4 | 910 | 97,9 | 7,75 | 9,39 | 4 | 1,183 | 1,098 | 1,173 | 1,001 | 0,877 | 1,146 | 0,861 | 1 | 1,15 | 0,15 | 1,2E-01 |
| Q8BGN3 | Ectonucleotide pyrophosphatase/phosphodiesterase family memb | 6 | 3 | 4 | 3 | 440 | 50,6 | 7,31 | 8,03 | 3 | 1,185 | 1,023 | 1,11 | 1,228 | 1,122 | 0,922 | 1,054 | 0,998 | 1,11 | 0,11 | 1,2E-01 |
| Q8BVG4 | Dipeptidyl peptidase 9 OS=Mus musculus OX=10090 GN=Dpp9 PE= | 4 | 4 | 4 | 4 | 862 | 97,9 | 6,65 | 2,45 | 4 | 1,183 | 1,159 | 1,107 | 0,976 | 0,955 | 1,067 | 0,994 | 1,03 | 1,09 | 0,09 | 1,2E-01 |
| P29391 | Ferritin light chain 1 OS=Mus musculus OX=10090 GN=Ftl1 PE=1 SV | 20 | 3 | 3 | 3 | 183 | 20,8 | 6 | 8,3 | 3 | 1,31 | 1,18 | 1,14 | 0,952 | 0,952 | 1,091 | 0,979 | 0,976 | 1,15 | 0,15 | 1,2E-01 |
| Q8JZS9 | 39S ribosomal protein L48, mitochondrial OS=Mus musculus OX=10 | 5 | 1 | 1 | 1 | 211 | 23,9 | 9,44 | 0 | 1 | 0,417 | 1,244 | 0,965 | 0,911 | 1,425 | 1,111 | 1,084 | 1,298 | 0,72 | -0,28 | 1,2E-01 |
| Q9WUK2 | Eukaryotic translation initiation factor 4H OS=Mus musculus OX=10 | 19 | 5 | 6 | 5 | 248 | 27,3 | 7,23 | 11,12 | 5 | 0,762 | 0,84 | 0,863 | 0,946 | 0,935 | 0,857 | 1,112 | 0,983 | 0,88 | -0,12 | 1,2E-01 |
| P83510 | Traf2 and NCK-interacting protein kinase OS=Mus musculus OX=10 | 13 | 15 | 16 | 10 | 1323 | 150,3 | 7,27 | 28,55 | 15 | 1,328 | 1,076 | 1,149 | 1,135 | 1,07 | 0,94 | 1,015 | 1,152 | 1,12 | 0,12 | 1,2E-01 |
| O55042 | Alpha-synuclein OS=Mus musculus OX=10090 GN=Snca PE=1 SV=2 | 59 | 10 | 27 | 7 | 140 | 14,5 | 4,77 | 78,33 | 10 | 1,094 | 0,993 | 1,018 | 0,996 | 0,919 | 0,957 | 1,015 | 0,984 | 1,06 | 0,06 | 1,2E-01 |
| Q9DCU6 | 39S ribosomal protein L4, mitochondrial OS=Mus musculus OX=100 | 5 | 1 | 1 | 1 | 294 | 33,1 | 9,82 | 2,38 | 1 | 4,803 | 2,794 | 3,163 | 2,606 | 1,62 | 2,576 | 1,983 | 2,96 | 1,46 | 0,46 | 1,2E-01 |
| P24369 | Peptidyl-prolyl cis-trans isomerase B OS=Mus musculus OX=10090 | 38 | 7 | 7 | 7 | 216 | 23,7 | 9,55 | 15,99 | 7 | 0,995 | 0,923 | 0,99 | 1,015 | 1,033 | 0,973 | 1,057 | 1,09 | 0,94 | -0,06 | 1,2E-01 |
| P11352 | Glutathione peroxidase 1 OS=Mus musculus OX=10090 GN=Gpx1 P | 19 | 3 | 3 | 3 | 201 | 22,3 | 7,21 | 2,83 | 3 | 1,359 | 1,224 | 0,961 | 0,956 | 0,927 | 0,947 | 0,982 | 0,912 | 1,19 | 0,19 | 1,2E-01 |
| Q9CQ26 | STAM-binding protein OS=Mus musculus OX=10090 GN=Stambp PE | 7 | 3 | 3 | 3 | 424 | 48,5 | 6,64 | 3,81 | 3 | 1,078 | 0,881 | 0,947 | 1,441 | 0,805 | 0,939 | 0,769 | 0,889 | 1,28 | 0,28 | 1,2E-01 |
| P46737 | Lys-63-specific deubiquitinase BRCC36 OS=Mus musculus OX=1009 | 14 | 4 | 4 | 4 | 291 | 33,3 | 5,83 | 2,04 | 4 | 1,254 | 1,179 | 1,112 | 1,155 | 1,168 | 1,024 | 1,139 | 1,001 | 1,08 | 0,08 | 1,2E-01 |
| Q71M36 | Chondroitin sulfate proteoglycan 5 OS=Mus musculus OX=10090 G | 6 | 3 | 3 | 3 | 566 | 60,4 | 4,54 | 4,46 | 3 | 1,224 | 0,525 | 0,97 | 0,988 | 1,631 | 1,17 | 1,27 | 1,034 | 0,73 | -0,27 | 1,2E-01 |
| P31324 | cAMP-dependent protein kinase type II-beta regulatory subunit OS | 29 | 12 | 17 | 10 | 416 | 46,1 | 4,98 | 45,62 | 12 | 0,927 | 0,964 | 0,902 | 0,906 | 0,957 | 0,931 | 1,021 | 0,961 | 0,96 | -0,04 | 1,2E-01 |
| P53810 | Phosphatidylinositol transfer protein alpha isoform OS=Mus muscu | 30 | 8 | 9 | 7 | 271 | 31,9 | 6,37 | 22,63 | 8 | 0,722 | 0,793 | 0,746 | 1,03 | 1,076 | 0,807 | 1,102 | 1,006 | 0,82 | -0,18 | 1,2E-01 |
| P51881 | ADP/ATP translocase 2 OS=Mus musculus OX=10090 GN=Slc25a5 P | 34 | 12 | 30 | 6 | 298 | 32,9 | 9,73 | 58,75 | 12 | 1,297 | 1,582 | 1,213 | 0,971 | 0,953 | 1,191 | 0,875 | 1,018 | 1,25 | 0,25 | 1,2E-01 |
| Q9JKK7 | Tropomodulin-2 OS=Mus musculus OX=10090 GN=Tmod2 PE=1 SV= | 26 | 9 | 15 | 8 | 351 | 39,5 | 5,35 | 47,1 | 9 | 1,038 | 1,01 | 1,024 | 1,043 | 1,006 | 1,154 | 1,122 | 1,071 | 0,95 | -0,05 | 1,2E-01 |
| P14576 | Signal recognition particle 54 kDa protein OS=Mus musculus OX=10 | 7 | 3 | 3 | 3 | 504 | 55,7 | 8,75 | 8,29 | 3 | 1,274 | 1,099 | 1,095 | 1,032 | 1,024 | 1,044 | 1,067 | 0,95 | 1,10 | 0,10 | 1,2E-01 |
| Q9R1C6 | Diacylglycerol kinase epsilon OS=Mus musculus OX=10090 GN=Dgk | 5 | 2 | 2 | 2 | 564 | 63,6 | 7,44 | 6,46 | 2 | 1,131 | 1,04 | 1,147 | 0,985 | 0,888 | 0,971 | 1 | 1,064 | 1,10 | 0,10 | 1,2E-01 |
| Q6P1D5 | Seizure 6-like protein OS=Mus musculus OX=10090 GN=Sez6l PE=1 | 1 | 1 | 1 | 1 | 963 | 104,8 | 4,69 | 2,9 | 1 | 1,17 | 1,115 | 1,203 | 1,267 | 1,111 | 1,21 | 0,963 | 1,023 | 1,10 | 0,10 | 1,2E-01 |
| Q8C5Q4 | G-rich sequence factor 1 OS=Mus musculus OX=10090 GN=Grsf1 PI | 4 | 1 | 1 | 1 | 479 | 53 | 6,67 | 3,38 | 1 | 0,918 | 0,974 | 0,903 | 0,892 | 0,733 | 0,772 | 0,926 | 0,896 | 1,11 | 0,11 | 1,2E-01 |
| Q8K394 | Inactive phospholipase C-like protein 2 OS=Mus musculus OX=1009 | 11 | 12 | 14 | 11 | 1128 | 125,7 | 6,92 | 33,8 | 12 | 1,23 | 1,134 | 1,179 | 1,064 | 1,007 | 1,166 | 1,011 | 1,057 | 1,09 | 0,09 | 1,2E-01 |
| Q91Y86 | Mitogen-activated protein kinase 8 OS=Mus musculus OX=10090 G | 5 | 2 | 2 | 2 | 384 | 44,2 | 7,69 | 2,2 | 2 | 0,714 | 0,922 | 0,886 | 0,701 | 0,652 | 0,645 | 0,772 | 0,696 | 1,17 | 0,17 | 1,3E-01 |

| | | | | | | | | | | | | | | | | | | | | | |
|---|---|---|---|---|---|---|---|---|---|---|---|---|---|---|---|---|---|---|---|---|---|
| Q9WTX6 | Cullin-1 OS=Mus musculus OX=10090 GN=Cul1 PE=1 SV=1 | 15 | 9 | 9 | 9 | 776 | 89,6 | 8 | 26,41 | 9 | 0,959 | 0,897 | 0,962 | 1,019 | 0,994 | 1,053 | 0,967 | 1,074 | 0,94 | -0,06 | 1,3E-01 |
| Q8K2H2 | Deubiquitinase OTUD6B OS=Mus musculus OX=10090 GN=Otud6b | 4 | 1 | 1 | 1 | 294 | 33,7 | 5,53 | 2,07 | 1 | 1,709 | 1,178 | 1,488 | 1,535 | 2,1 | 1,74 | 1,394 | 2,212 | 0,79 | -0,21 | 1,3E-01 |
| Q9CRC6 | BLOC-1-related complex subunit 7 OS=Mus musculus OX=10090 GN | 10 | 1 | 1 | 1 | 105 | 11,5 | 6,79 | 1,66 | 1 | 1,023 | 1,391 | 1,143 | 0,836 | 0,787 | 0,926 | 1,002 | 0,757 | 1,27 | 0,27 | 1,3E-01 |
| Q9D7P6 | Iron-sulfur cluster assembly enzyme ISCU, mitochondrial OS=Mus n | 8 | 2 | 2 | 2 | 168 | 18,1 | 9,29 | 4,4 | 2 | 0,845 | 0,973 | 0,825 | 0,809 | 1,099 | 0,831 | 0,949 | 1,14 | 0,86 | -0,14 | 1,3E-01 |
| Q7TSJ2 | Microtubule-associated protein 6 OS=Mus musculus OX=10090 GN | 60 | 48 | 83 | 48 | 906 | 96,4 | 9,5 | 192,87 | 48 | 0,868 | 0,92 | 0,947 | 1,059 | 1,018 | 0,964 | 1,131 | 1,059 | 0,91 | -0,09 | 1,3E-01 |
| Q924D0 | Reticulon-4-interacting protein 1, mitochondrial OS=Mus musculus | 7 | 2 | 3 | 2 | 396 | 43,3 | 9,2 | 9,66 | 2 | 1,138 | 1,107 | 1,064 | 0,972 | 0,83 | 1,049 | 0,937 | 1,029 | 1,11 | 0,11 | 1,3E-01 |
| Q9D6R2 | Isocitrate dehydrogenase [NAD] subunit alpha, mitochondrial OS=M | 34 | 15 | 32 | 15 | 366 | 39,6 | 6,73 | 71,97 | 15 | 0,943 | 0,99 | 1,003 | 0,959 | 1,069 | 0,984 | 0,993 | 1,015 | 0,96 | -0,04 | 1,3E-01 |
| P58252 | Elongation factor 2 OS=Mus musculus OX=10090 GN=Eef2 PE=1 SV | 17 | 14 | 21 | 14 | 858 | 95,3 | 6,83 | 47,11 | 14 | 0,846 | 0,844 | 0,935 | 1 | 0,974 | 0,915 | 1,061 | 1,022 | 0,91 | -0,09 | 1,3E-01 |
| Q8K0D0 | Cyclin-dependent kinase 17 OS=Mus musculus OX=10090 GN=Cdk1 | 7 | 3 | 4 | 2 | 523 | 59,5 | 9,03 | 6,47 | 3 | 1,055 | 2,337 | 1,271 | 1,771 | 2,775 | 2,195 | 2,14 | 1,794 | 0,72 | -0,28 | 1,3E-01 |
| Q9R1T4 | Septin-6 OS=Mus musculus OX=10090 GN=Septin6 PE=1 SV=4 | 31 | 13 | 19 | 6 | 434 | 49,6 | 6,43 | 47,72 | 13 | 1,19 | 1,168 | 1,08 | 0,945 | 0,86 | 1,068 | 0,95 | 1,005 | 1,13 | 0,13 | 1,3E-01 |
| Q9CPX6 | Ubiquitin-like-conjugating enzyme ATG3 OS=Mus musculus OX=100 | 15 | 4 | 4 | 4 | 314 | 35,8 | 4,72 | 6,81 | 4 | 1,283 | 1,171 | 1,025 | 1,026 | 0,879 | 1,058 | 1,023 | 1,023 | 1,13 | 0,13 | 1,3E-01 |
| Q9CXP8 | Guanine nucleotide-binding protein G(I)/G(S)/G(O) subunit gamma | 68 | 3 | 3 | 3 | 68 | 7,2 | 7,85 | 8,95 | 3 | 1,413 | 1,261 | 1,175 | 1,055 | 1,011 | 1,219 | 0,986 | 1,041 | 1,15 | 0,15 | 1,3E-01 |
| P61022 | Calcineurin B homologous protein 1 OS=Mus musculus OX=10090 G | 52 | 8 | 10 | 8 | 195 | 22,4 | 5,1 | 18,42 | 8 | 1,485 | 1,357 | 1,17 | 1,138 | 0,932 | 1,323 | 1,024 | 1,048 | 1,19 | 0,19 | 1,3E-01 |
| Q9Z2W8 | Glutamate receptor 4 OS=Mus musculus OX=10090 GN=Gria4 PE=1 | 7 | 6 | 9 | 2 | 902 | 100,8 | 8,07 | 21,52 | 6 | 0,73 | 0,789 | 0,978 | 1,036 | 1,007 | 1,214 | 1,02 | 0,947 | 0,84 | -0,16 | 1,3E-01 |
| Q9D5V6 | Synapse-associated protein 1 OS=Mus musculus OX=10090 GN=Sya | 8 | 3 | 4 | 3 | 365 | 41,3 | 4,54 | 8,46 | 3 | 1,436 | 1,451 | 1,34 | 1,209 | 1,076 | 1,349 | 1,089 | 1,295 | 1,13 | 0,13 | 1,3E-01 |
| P63001 | Ras-related C3 botulinum toxin substrate 1 OS=Mus musculus OX=1 | 42 | 10 | 19 | 6 | 192 | 21,4 | 8,5 | 38,06 | 10 | 0,849 | 0,945 | 0,905 | 1,047 | 1,019 | 0,972 | 1,035 | 1,029 | 0,92 | -0,08 | 1,3E-01 |
| Q5SNZ0 | Girdin OS=Mus musculus OX=10090 GN=Ccdc88a PE=1 SV=2 | 4 | 7 | 7 | 7 | 1873 | 215,8 | 6,24 | 8,99 | 7 | 1,343 | 1,176 | 1,064 | 1,117 | 0,919 | 1,05 | 1,092 | 1,116 | 1,13 | 0,13 | 1,3E-01 |
| Q3UH66 | Serine/threonine-protein kinase WNK2 OS=Mus musculus OX=1009 | 6 | 10 | 10 | 10 | 2149 | 227,4 | 5,91 | 17,74 | 10 | 0,809 | 0,931 | 0,996 | 1,004 | 0,958 | 1,03 | 1,124 | 1,023 | 0,90 | -0,10 | 1,3E-01 |
| Q9D0Q7 | 39S ribosomal protein L45, mitochondrial OS=Mus musculus OX=10 | 5 | 2 | 2 | 2 | 306 | 35,4 | 9,23 | 4,62 | 2 | 1,01 | 1,262 | 1,07 | 0,888 | 1,006 | 0,949 | 0,809 | 0,829 | 1,18 | 0,18 | 1,3E-01 |
| Q921I1 | Serotransferrin OS=Mus musculus OX=10090 GN=Tf PE=1 SV=1 | 18 | 11 | 11 | 11 | 697 | 76,7 | 7,18 | 22,18 | 11 | 0,974 | 0,947 | 1,076 | 0,994 | 0,978 | 0,923 | 0,889 | 0,962 | 1,06 | 0,06 | 1,3E-01 |
| P56564 | Excitatory amino acid transporter 1 OS=Mus musculus OX=10090 G | 12 | 6 | 31 | 6 | 543 | 59,6 | 8,4 | 56,94 | 6 | 0,923 | 0,997 | 1,098 | 1,117 | 1,104 | 1,138 | 1,202 | 1,066 | 0,92 | -0,08 | 1,3E-01 |
| P97470 | Serine/threonine-protein phosphatase 4 catalytic subunit OS=Mus | 4 | 1 | 2 | 1 | 307 | 35,1 | 5,06 | 3,47 | 1 | 0,931 | 0,782 | 0,885 | 0,912 | 1,036 | 0,993 | 0,989 | 0,853 | 0,91 | -0,09 | 1,3E-01 |
| Q8VDN2 | Sodium/potassium-transporting ATPase subunit alpha-1 OS=Mus m | 47 | 51 | 192 | 31 | 1023 | 112,9 | 5,45 | 422,81 | 51 | 0,868 | 0,945 | 0,931 | 0,964 | 1,004 | 0,936 | 0,954 | 1,001 | 0,95 | -0,05 | 1,3E-01 |
| Q8VD62 | UPF0696 protein C11orf68 homolog OS=Mus musculus OX=10090 G | 14 | 4 | 6 | 4 | 298 | 31,8 | 6,1 | 4,78 | 4 | 1,008 | 1,211 | 1,234 | 1,094 | 1,052 | 1,118 | 0,875 | 0,988 | 1,13 | 0,13 | 1,3E-01 |
| P97393 | Rho GTPase-activating protein 5 OS=Mus musculus OX=10090 GN= | 7 | 8 | 8 | 8 | 1501 | 172 | 6,34 | 10,18 | 8 | 0,951 | 0,914 | 0,964 | 0,999 | 1,077 | 0,965 | 1,024 | 0,977 | 0,95 | -0,05 | 1,3E-01 |
| P54923 | [Protein ADP-ribosylarginine] hydrolase OS=Mus musculus OX=100 | 11 | 2 | 3 | 2 | 362 | 40 | 5,76 | 6,38 | 2 | 2,327 | 1,668 | 1,323 | 1,011 | 0,888 | 1,053 | 1,132 | 1,232 | 1,47 | 0,47 | 1,3E-01 |
| Q7TQ95 | Endoplasmic reticulum junction formation protein lunapark OS=Mu | 20 | 5 | 5 | 5 | 425 | 47,5 | 5,27 | 9,34 | 5 | 1,267 | 1,068 | 1,076 | 1,005 | 0,895 | 1,088 | 0,969 | 0,984 | 1,12 | 0,12 | 1,3E-01 |
| Q91ZP9 | N-terminal EF-hand calcium-binding protein 2 OS=Mus musculus O | 5 | 2 | 3 | 2 | 389 | 43,4 | 5,3 | 6,44 | 2 | 0,884 | 0,82 | 0,85 | 0,912 | 0,937 | 0,868 | 1,092 | 0,928 | 0,91 | -0,09 | 1,3E-01 |
| Q9EPJ9 | ADP-ribosylation factor GTPase-activating protein 1 OS=Mus muscu | 26 | 8 | 10 | 8 | 414 | 45,3 | 5,57 | 21,41 | 8 | 1,105 | 1,044 | 1,023 | 1,031 | 0,973 | 1,055 | 0,951 | 1,018 | 1,05 | 0,05 | 1,3E-01 |
| P14094 | Sodium/potassium-transporting ATPase subunit beta-1 OS=Mus mu | 39 | 14 | 41 | 14 | 304 | 35,2 | 8,65 | 91,19 | 14 | 0,825 | 0,911 | 0,963 | 0,995 | 1,037 | 1,04 | 1,011 | 0,924 | 0,92 | -0,08 | 1,3E-01 |
| Q8C0P5 | Coronin-2A OS=Mus musculus OX=10090 GN=Coro2a PE=2 SV=1 | 5 | 3 | 3 | 2 | 524 | 59,5 | 7,71 | 7,83 | 3 | 0,886 | 0,955 | 0,958 | 0,987 | 0,979 | 1,017 | 0,966 | 0,99 | 0,96 | -0,04 | 1,3E-01 |
| Q80VJ3 | 2'-deoxynucleoside 5'-phosphate N-hydrolase 1 OS=Mus musculus | 8 | 1 | 1 | 1 | 173 | 19 | 5,38 | 2,26 | 1 | 1,733 | 1,792 | 1,716 | 1,385 | 0,01 | 1,545 | 1,264 | 1,325 | 1,60 | 0,60 | 1,3E-01 |
| Q9CQZ6 | NADH dehydrogenase [ubiquinone] 1 beta subcomplex subunit 3 O | 12 | 2 | 3 | 2 | 104 | 11,7 | 9,04 | 7,86 | 2 | 1,604 | 1,499 | 1,312 | 0,987 | 0,846 | 1,224 | 1,089 | 1,15 | 1,25 | 0,25 | 1,3E-01 |
| Q80XP9 | Serine/threonine-protein kinase WNK3 OS=Mus musculus OX=1009 | 0 | 1 | 1 | 1 | 1757 | 193,9 | 6,02 | 1,79 | 1 | 1,062 | 1,239 | 1,166 | 1,027 | 0,894 | 0,913 | 0,984 | 1,163 | 1,14 | 0,14 | 1,3E-01 |
| P60904 | DnaJ homolog subfamily C member 5 OS=Mus musculus OX=10090 | 23 | 6 | 14 | 6 | 198 | 22,1 | 5,07 | 36,7 | 6 | 0,781 | 0,906 | 0,886 | 0,998 | 0,942 | 0,942 | 1,042 | 0,994 | 0,91 | -0,09 | 1,4E-01 |
| Q61035 | Histidine--tRNA ligase, cytoplasmic OS=Mus musculus OX=10090 G | 14 | 6 | 6 | 4 | 509 | 57,4 | 6 | 16,92 | 6 | 1,509 | 1,29 | 1,269 | 1,312 | 1,025 | 1,304 | 1,328 | 1,043 | 1,14 | 0,14 | 1,4E-01 |
| O35188 | Fractalkine OS=Mus musculus OX=10090 GN=Cx3cl1 PE=1 SV=3 | 5 | 1 | 2 | 1 | 395 | 42,1 | 5,54 | 4,24 | 1 | 1,378 | 1,19 | 1,045 | 0,999 | 0,83 | 1,225 | 0,876 | 0,749 | 1,25 | 0,25 | 1,4E-01 |
| B9EJ80 | PDZ domain-containing protein 8 OS=Mus musculus OX=10090 GN= | 1 | 1 | 1 | 1 | 1147 | 127,7 | 6,04 | 2,3 | 1 | 2,205 | 1,221 | 1,123 | 1,075 | 0,676 | 1,209 | 0,887 | 0,855 | 1,55 | 0,55 | 1,4E-01 |
| Q9CZU6 | Citrate synthase, mitochondrial OS=Mus musculus OX=10090 GN=C | 19 | 9 | 22 | 9 | 464 | 51,7 | 8,57 | 48,99 | 9 | 0,766 | 0,931 | 0,941 | 0,932 | 1,012 | 0,887 | 1,025 | 1,013 | 0,91 | -0,09 | 1,4E-01 |
| P58871 | 182 kDa tankyrase-1-binding protein OS=Mus musculus OX=10090 | 7 | 9 | 10 | 9 | 1720 | 181,7 | 4,88 | 18,36 | 9 | 1,1 | 1,187 | 1,104 | 1,029 | 1,005 | 0,947 | 1,064 | 1,091 | 1,08 | 0,08 | 1,4E-01 |
| P70297 | Signal transducing adapter molecule 1 OS=Mus musculus OX=1009 | 6 | 3 | 4 | 3 | 548 | 59,7 | 4,84 | 2,53 | 3 | 0,749 | 0,967 | 0,832 | 1,084 | 0,984 | 0,986 | 1,083 | 1,179 | 0,86 | -0,14 | 1,4E-01 |
| Q00558 | Factor VIII intron 22 protein OS=Mus musculus OX=10090 GN=F8a1 | 7 | 2 | 2 | 2 | 380 | 40,5 | 6,89 | 5,18 | 2 | 0,656 | 0,987 | 0,928 | 0,862 | 0,974 | 1,108 | 1,132 | 0,869 | 0,84 | -0,16 | 1,4E-01 |
| Q8C8U0 | Liprin-beta-1 OS=Mus musculus OX=10090 GN=Ppfibp1 PE=1 SV=3 | 1 | 1 | 1 | 1 | 969 | 108,5 | 5,49 | 1,63 | 1 | 0,847 | 0,863 | 0,794 | 0,848 | 0,878 | 0,86 | 0,988 | 1,247 | 0,84 | -0,16 | 1,4E-01 |
| Q05512 | Serine/threonine-protein kinase MARK2 OS=Mus musculus OX=100 | 17 | 12 | 14 | 10 | 776 | 86,3 | 9,67 | 22,45 | 12 | 1,066 | 1,036 | 1,105 | 1,024 | 0,88 | 1,076 | 0,994 | 0,979 | 1,08 | 0,08 | 1,4E-01 |
| Q9JKC6 | Cell cycle exit and neuronal differentiation protein 1 OS=Mus musc | 36 | 6 | 12 | 6 | 149 | 15 | 8,97 | 20,49 | 6 | 0,969 | 1,028 | 1,154 | 1,142 | 1,059 | 0,759 | 0,999 | 0,936 | 1,14 | 0,14 | 1,4E-01 |
| Q99ME2 | WD repeat-containing protein 6 OS=Mus musculus OX=10090 GN= | 1 | 1 | 1 | 1 | 1125 | 121,8 | 6,98 | 1,91 | 1 | 1,281 | 1,246 | 0,939 | 0,862 | 0,701 | 1 | 0,935 | 0,84 | 1,25 | 0,25 | 1,4E-01 |
| Q3TWN3 | Metal transporter CNNM2 OS=Mus musculus OX=10090 GN=Cnnm | 9 | 6 | 6 | 4 | 875 | 96,6 | 6,21 | 7,01 | 6 | 0,99 | 1,063 | 0,928 | 1,041 | 0,829 | 0,968 | 0,637 | 0,993 | 1,17 | 0,17 | 1,4E-01 |
| Q9CQT1 | Methylthioribose-1-phosphate isomerase OS=Mus musculus OX=10 | 2 | 1 | 1 | 1 | 369 | 39,4 | 5,91 | 1,7 | 1 | 1,273 | 0,997 | 1,113 | 0,87 | 0,994 | 0,915 | 0,884 | 0,828 | 1,17 | 0,17 | 1,4E-01 |
| Q3TCJ1 | BRISC complex subunit Abraxas 2 OS=Mus musculus OX=10090 GN= | 9 | 3 | 3 | 3 | 415 | 46,9 | 6,18 | 2,2 | 3 | 0,6 | 0,858 | 0,946 | 0,943 | 0,854 | 1,125 | 0,975 | 1,087 | 0,83 | -0,17 | 1,4E-01 |
| Q8BYW1 | Rho GTPase-activating protein 25 OS=Mus musculus OX=10090 GN | 6 | 4 | 5 | 3 | 648 | 73,3 | 6,38 | 4,29 | 4 | 0,907 | 1,124 | 0,816 | 0,925 | 0,9 | 0,649 | 0,876 | 0,753 | 1,19 | 0,19 | 1,4E-01 |
| P31938 | Dual specificity mitogen-activated protein kinase kinase 1 OS=Mus | 23 | 9 | 12 | 9 | 393 | 43,4 | 6,7 | 19,02 | 9 | 1,085 | 1,077 | 1,103 | 1,019 | 0,979 | 1,045 | 1,064 | 1,02 | 1,04 | 0,04 | 1,4E-01 |
| P24547 | Inosine-5'-monophosphate dehydrogenase 2 OS=Mus musculus OX | 7 | 2 | 2 | 2 | 514 | 55,8 | 7,28 | 6,83 | 2 | 0,899 | 1,124 | 1,212 | 1,11 | 1,255 | 1,127 | 1,417 | 1,173 | 0,87 | -0,13 | 1,4E-01 |
| P61922 | 4-aminobutyrate aminotransferase, mitochondrial OS=Mus muscul | 40 | 16 | 25 | 16 | 500 | 56,4 | 8,09 | 54,76 | 16 | 0,96 | 0,81 | 1,095 | 0,984 | 1,054 | 1,169 | 1,076 | 1,01 | 0,89 | -0,11 | 1,4E-01 |
| Q8BHS6 | Armadillo repeat-containing X-linked protein 3 OS=Mus musculus C | 6 | 2 | 2 | 2 | 379 | 42,6 | 8,68 | 0 | 2 | 1,024 | 0,754 | 0,605 | 0,926 | 1,279 | 0,98 | 1,195 | 0,812 | 0,78 | -0,22 | 1,4E-01 |
| P97326 | Cadherin-6 OS=Mus musculus OX=10090 GN=Cdh6 PE=1 SV=2 | 5 | 3 | 3 | 3 | 790 | 88,3 | 5 | 2,5 | 3 | 1,571 | 1,56 | 1,233 | 1,064 | 1,077 | 1,277 | 0,999 | 1,134 | 1,21 | 0,21 | 1,4E-01 |
| A2ASS6 | Titin OS=Mus musculus OX=10090 GN=Ttn PE=1 SV=1 | 0 | 2 | 2 | 2 | 35213 | 3904,1 | 6,2 | 0 | 2 | 1,968 | 1,208 | 2,007 | 3,406 | 0,669 | 1,647 | 1,697 | 1,031 | 1,70 | 0,70 | 1,4E-01 |
| P14115 | 60S ribosomal protein L27a OS=Mus musculus OX=10090 GN=Rpl2 | 25 | 4 | 5 | 4 | 148 | 16,6 | 11,12 | 11,59 | 4 | 1,054 | 0,924 | 0,883 | 0,889 | 1,037 | 0,992 | 0,96 | 1,109 | 0,92 | -0,08 | 1,4E-01 |
| Q6WKZ8 | E3 ubiquitin-protein ligase UBR2 OS=Mus musculus OX=10090 GN= | 0 | 1 | 1 | 1 | 1755 | 199 | 6,33 | 0 | 1 | 0,01 | 0,687 | 0,628 | 0,621 | 0,76 | 0,832 | 0,582 | 1,007 | 0,61 | -0,39 | 1,4E-01 |
| Q921M7 | Protein FAM49B OS=Mus musculus OX=10090 GN=Fam49b PE=1 S | 36 | 10 | 12 | 8 | 324 | 36,8 | 6,06 | 24,73 | 10 | 0,936 | 0,907 | 0,958 | 1,028 | 1,065 | 0,971 | 1,096 | 0,974 | 0,93 | -0,07 | 1,4E-01 |
| A2AG50 | MAP7 domain-containing protein 2 OS=Mus musculus OX=10090 G | 2 | 1 | 1 | 1 | 781 | 86 | 8,81 | 0 | 1 | 0,01 | 1,121 | 0,989 | 1,575 | 1,587 | 1,884 | 1,19 | 1,463 | 0,60 | -0,40 | 1,4E-01 |
| Q3UGY8 | Brefeldin A-inhibited guanine nucleotide-exchange protein 3 OS=M | 7 | 11 | 12 | 11 | 2170 | 239,9 | 5,88 | 17,7 | 11 | 1,239 | 1,258 | 1,1 | 1,049 | 1,007 | 1,133 | 1,056 | 1,061 | 1,09 | 0,09 | 1,4E-01 |
| Q62523 | Zyxin OS=Mus musculus OX=10090 GN=Zyx PE=1 SV=2 | 5 | 2 | 2 | 2 | 564 | 60,5 | 6,4 | 2,75 | 2 | 0,375 | 1,006 | 0,953 | 0,95 | 1,18 | 1,265 | 1,112 | 0,878 | 0,74 | -0,26 | 1,4E-01 |

| | | | | | | | | | | | | | | | | | | | | | |
|---|---|---|---|---|---|---|---|---|---|---|---|---|---|---|---|---|---|---|---|---|---|
| Q9QXZ0 | Microtubule-actin cross-linking factor 1 OS=Mus musculus OX=100 | 10 | 64 | 72 | 60 | 7354 | 831,4 | 5,43 | 142,93 | 64 | 1,063 | 1,064 | 1,015 | 1 | 0,994 | 1,006 | 1,019 | 1,007 | 1,03 | 0,03 | 1,4E-01 |
| Q9DAM7 | Transmembrane protein 263 OS=Mus musculus OX=10090 GN=Tm | 62 | 4 | 5 | 4 | 115 | 11,5 | 9,32 | 5,93 | 4 | 1,67 | 1,453 | 1,268 | 1,1 | 0,95 | 1,295 | 1,182 | 1,106 | 1,21 | 0,21 | 1,4E-01 |
| Q9Z0V2 | Potassium voltage-gated channel subfamily D member 2 OS=Mus n | 5 | 2 | 3 | 2 | 630 | 70,5 | 8,07 | 2,9 | 2 | 0,66 | 0,755 | 0,903 | 0,912 | 1,109 | 0,8 | 1,08 | 0,889 | 0,83 | -0,17 | 1,4E-01 |
| P68510 | 14-3-3 protein eta OS=Mus musculus OX=10090 GN=Ywhah PE=1 S | 57 | 17 | 41 | 12 | 246 | 28,2 | 4,89 | 92,61 | 17 | 0,993 | 1,025 | 0,974 | 1,003 | 0,888 | 1,013 | 0,881 | 0,98 | 1,06 | 0,06 | 1,4E-01 |
| P01027 | Complement C3 OS=Mus musculus OX=10090 GN=C3 PE=1 SV=3 | 1 | 2 | 2 | 2 | 1663 | 186,4 | 6,73 | 1,75 | 2 | 0,992 | 1,005 | 0,695 | 0,651 | 0,935 | 1,247 | 0,898 | 1,092 | 0,80 | -0,20 | 1,4E-01 |
| Q61733 | 28S ribosomal protein S31, mitochondrial OS=Mus musculus OX=1 | 6 | 3 | 3 | 3 | 384 | 43,9 | 8,51 | 6,77 | 3 | 0,928 | 0,962 | 0,969 | 0,958 | 0,936 | 1,022 | 0,994 | 1,097 | 0,94 | -0,06 | 1,5E-01 |
| Q5DTT2 | PH and SEC7 domain-containing protein 1 OS=Mus musculus OX=1 | 7 | 5 | 5 | 5 | 1024 | 109,6 | 7,05 | 14,33 | 5 | 1,285 | 1,115 | 1,009 | 1,151 | 1,043 | 1,04 | 0,888 | 1,101 | 1,12 | 0,12 | 1,5E-01 |
| Q9ERB5 | Solute carrier organic anion transporter family member 1C1 OS=Mu | 3 | 2 | 2 | 2 | 715 | 78,3 | 8,81 | 1,75 | 2 | 1,148 | 0,892 | 1,242 | 0,967 | 0,846 | 1,014 | 0,894 | 0,907 | 1,16 | 0,16 | 1,5E-01 |
| O55013 | Trafficking protein particle complex subunit 3 OS=Mus musculus O | 19 | 4 | 4 | 4 | 180 | 20,3 | 4,96 | 9,27 | 4 | 0,905 | 0,881 | 1,034 | 0,971 | 0,951 | 1,105 | 1,038 | 1,011 | 0,92 | -0,08 | 1,5E-01 |
| Q9D358 | Low molecular weight phosphotyrosine protein phosphatase OS=M | 19 | 4 | 4 | 4 | 158 | 18,2 | 6,74 | 5,86 | 4 | 0,758 | 0,857 | 0,729 | 1,062 | 1,067 | 0,891 | 1,045 | 0,972 | 0,86 | -0,14 | 1,5E-01 |
| Q8BG93 | Nucleotide triphosphate diphosphatase NUDT15 OS=Mus musculus | 6 | 1 | 1 | 1 | 170 | 19,6 | 5,27 | 0 | 1 | 1,091 | 0,769 | 0,864 | 0,991 | 0,777 | 0,654 | 0,825 | 0,885 | 1,18 | 0,18 | 1,5E-01 |
| P04370 | Myelin basic protein OS=Mus musculus OX=10090 GN=Mbp PE=1 S | 37 | 9 | 48 | 2 | 250 | 27,2 | 9,58 | 128,23 | 9 | 0,87 | 0,914 | 0,87 | 0,808 | 0,984 | 0,813 | 0,983 | 1,051 | 0,90 | -0,10 | 1,5E-01 |
| Q9CWM4 | Prefoldin subunit 1 OS=Mus musculus OX=10090 GN=Pfdn1 PE=1 S | 24 | 3 | 6 | 3 | 122 | 14,2 | 8,32 | 14,85 | 3 | 1,225 | 1,161 | 1,158 | 1,143 | 0,971 | 1,219 | 0,955 | 1,109 | 1,10 | 0,10 | 1,5E-01 |
| Q8VBY2 | Calcium/calmodulin-dependent protein kinase kinase 1 OS=Mus m | 9 | 3 | 4 | 3 | 505 | 55,8 | 5,99 | 5,22 | 3 | 1,262 | 1,301 | 1,066 | 1,002 | 1,034 | 1,11 | 1,022 | 0,892 | 1,14 | 0,14 | 1,5E-01 |
| A3KGV1 | Outer dense fiber protein 2 OS=Mus musculus OX=10090 GN=Odf2 | 1 | 1 | 1 | 1 | 830 | 95,5 | 7,61 | 2,04 | 1 | 0,686 | 0,849 | 0,73 | 0,973 | 1,029 | 0,8 | 1,112 | 0,92 | 0,84 | -0,16 | 1,5E-01 |
| P50172 | Corticosteroid 11-beta-dehydrogenase isozyme 1 OS=Mus musculu | 9 | 2 | 2 | 2 | 292 | 32,3 | 8,46 | 4,81 | 2 | 1,241 | 1,357 | 1,22 | 1,147 | 0,98 | 1,045 | 1,149 | 1,274 | 1,12 | 0,12 | 1,5E-01 |
| A2A8Z1 | Oxysterol-binding protein-related protein 9 OS=Mus musculus OX= | 1 | 1 | 1 | 1 | 736 | 83,1 | 6,07 | 2,77 | 1 | 1,327 | 1,042 | 1 | 1,107 | 0,634 | 1,104 | 1,044 | 0,833 | 1,24 | 0,24 | 1,5E-01 |
| P23927 | Alpha-crystallin B chain OS=Mus musculus OX=10090 GN=Cryab PE | 23 | 4 | 6 | 4 | 175 | 20,1 | 7,33 | 9,93 | 4 | 0,911 | 0,991 | 0,958 | 1,073 | 0,953 | 0,898 | 0,92 | 0,923 | 1,06 | 0,06 | 1,5E-01 |
| Q99JB2 | Stomatin-like protein 2, mitochondrial OS=Mus musculus OX=1009 | 21 | 6 | 8 | 6 | 353 | 38,4 | 8,87 | 18,81 | 6 | 0,999 | 0,881 | 1,012 | 0,995 | 1,032 | 1,007 | 1,008 | 1,173 | 0,92 | -0,08 | 1,5E-01 |
| P39688 | Tyrosine-protein kinase Fyn OS=Mus musculus OX=10090 GN=Fyn | 7 | 4 | 5 | 3 | 537 | 60,6 | 6,67 | 8,24 | 4 | 1,294 | 1,208 | 1,243 | 1,226 | 1,022 | 1,285 | 1,124 | 1,159 | 1,08 | 0,08 | 1,5E-01 |
| Q8C008 | Double zinc ribbon and ankyrin repeat-containing protein 1 OS=Mu | 3 | 2 | 2 | 2 | 778 | 85 | 8,18 | 1,79 | 2 | 0,76 | 1,065 | 0,967 | 0,734 | 0,581 | 0,816 | 0,753 | 0,747 | 1,22 | 0,22 | 1,5E-01 |
| Q9DC63 | F-box only protein 3 OS=Mus musculus OX=10090 GN=Fbxo3 PE=1 | 2 | 1 | 1 | 1 | 480 | 55,2 | 5,02 | 0 | 1 | 1,543 | 1,184 | 1,015 | 1,103 | 0,741 | 1,288 | 0,831 | 0,872 | 1,30 | 0,30 | 1,5E-01 |
| P60766 | Cell division control protein 42 homolog OS=Mus musculus OX=100 | 27 | 5 | 7 | 1 | 191 | 21,2 | 6,55 | 11,24 | 5 | 1,384 | 0,885 | 1,134 | 1,107 | 0,946 | 0,894 | 1,016 | 0,957 | 1,18 | 0,18 | 1,5E-01 |
| P03911 | NADH-ubiquinone oxidoreductase chain 4 OS=Mus musculus OX=1 | 5 | 2 | 2 | 2 | 459 | 51,8 | 9,38 | 1,63 | 2 | 2,598 | 2,165 | 2,161 | 1,986 | 1,005 | 2,374 | 1,961 | 1,378 | 1,33 | 0,33 | 1,5E-01 |
| Q9D892 | Inosine triphosphate pyrophosphatase OS=Mus musculus OX=1009 | 29 | 4 | 4 | 4 | 198 | 21,9 | 5,87 | 9,11 | 4 | 1,261 | 1,669 | 1,102 | 1,043 | 0,797 | 1,27 | 0,933 | 0,924 | 1,29 | 0,29 | 1,5E-01 |
| Q5SSM3 | Rho GTPase-activating protein 44 OS=Mus musculus OX=10090 GN | 8 | 6 | 8 | 6 | 814 | 88,9 | 6,6 | 15,59 | 6 | 0,974 | 0,91 | 0,965 | 1,07 | 0,987 | 1,041 | 1,081 | 1,071 | 0,94 | -0,06 | 1,5E-01 |
| Q99LC8 | Translation initiation factor eIF-2B subunit alpha OS=Mus musculus | 7 | 2 | 2 | 2 | 305 | 33,8 | 8,32 | 2,37 | 2 | 1,04 | 0,905 | 1,058 | 1,106 | 0,853 | 0,837 | 1,08 | 0,864 | 1,13 | 0,13 | 1,5E-01 |
| Q9QWL7 | Keratin, type I cytoskeletal 17 OS=Mus musculus OX=10090 GN=Krt | 21 | 9 | 12 | 1 | 433 | 48,1 | 5,06 | 25,4 | 9 | 1,974 | 1,852 | 0,907 | 0,811 | 0,906 | 0,907 | 0,951 | 0,732 | 1,59 | 0,59 | 1,5E-01 |
| Q8BWS5 | G protein-regulated inducer of neurite outgrowth 3 OS=Mus muscu | 4 | 2 | 4 | 2 | 763 | 80,4 | 7,02 | 4,43 | 2 | 1,055 | 1,073 | 1,379 | 1,168 | 0,78 | 0,48 | 1,291 | 0,909 | 1,35 | 0,35 | 1,5E-01 |
| Q99LP6 | GrpE protein homolog 1, mitochondrial OS=Mus musculus OX=100 | 31 | 6 | 7 | 6 | 217 | 24,3 | 8,38 | 19,61 | 6 | 0,938 | 0,969 | 1,126 | 0,954 | 1,042 | 1,122 | 1,047 | 1,09 | 0,93 | -0,07 | 1,5E-01 |
| Q8CI32 | BAG family molecular chaperone regulator 5 OS=Mus musculus OX | 7 | 2 | 2 | 2 | 447 | 50,9 | 6,05 | 5,72 | 2 | 0,979 | 1,151 | 1,148 | 1,429 | 0,868 | 1,029 | 1,043 | 1,078 | 1,17 | 0,17 | 1,5E-01 |
| Q6IFX2 | Keratin, type I cytoskeletal 42 OS=Mus musculus OX=10090 GN=Krt | 19 | 8 | 12 | 3 | 452 | 50,1 | 5,16 | 19,53 | 8 | 1,184 | 1,542 | 1,074 | 0,962 | 0,931 | 1,104 | 0,719 | 1,015 | 1,26 | 0,26 | 1,5E-01 |
| Q61206 | Platelet-activating factor acetylhydrolase IB subunit beta OS=Mus r | 8 | 1 | 2 | 1 | 229 | 25,6 | 5,92 | 7,52 | 1 | 1,3 | 1,02 | 1,386 | 1,091 | 0,995 | 0,907 | 1,065 | 1,16 | 1,16 | 0,16 | 1,5E-01 |
| P47738 | Aldehyde dehydrogenase, mitochondrial OS=Mus musculus OX=10 | 22 | 10 | 12 | 9 | 519 | 56,5 | 7,62 | 24,1 | 10 | 0,955 | 0,999 | 0,907 | 0,968 | 1,051 | 0,954 | 0,981 | 1,046 | 0,95 | -0,05 | 1,5E-01 |
| Q61151 | Serine/threonine-protein phosphatase 2A 56 kDa regulatory subun | 17 | 7 | 8 | 5 | 467 | 54,7 | 6,95 | 14,73 | 7 | 0,838 | 1,065 | 0,887 | 0,943 | 0,937 | 1,249 | 1,047 | 1,039 | 0,87 | -0,13 | 1,5E-01 |
| P00920 | Carbonic anhydrase 2 OS=Mus musculus OX=10090 GN=Ca2 PE=1 S | 30 | 5 | 6 | 5 | 260 | 29 | 7,01 | 19,81 | 5 | 0,876 | 0,974 | 0,953 | 0,981 | 0,947 | 0,849 | 0,895 | 0,886 | 1,06 | 0,06 | 1,5E-01 |
| O35239 | Tyrosine-protein phosphatase non-receptor type 9 OS=Mus muscul | 8 | 5 | 7 | 5 | 593 | 67,9 | 8,1 | 9,6 | 5 | 0,01 | 0,626 | 0,859 | 0,866 | 1,021 | 0,8 | 0,911 | 0,994 | 0,63 | -0,37 | 1,5E-01 |
| Q8BYN5 | FSD1-like protein OS=Mus musculus OX=10090 GN=Fsd1l PE=1 SV= | 9 | 4 | 5 | 4 | 507 | 56,9 | 6,7 | 8,07 | 4 | 1,212 | 1,135 | 1,034 | 0,972 | 0,833 | 1,032 | 1,022 | 1 | 1,12 | 0,12 | 1,5E-01 |
| Q80TJ1 | Calcium-dependent secretion activator 1 OS=Mus musculus OX=10 | 31 | 34 | 54 | 1 | 1355 | 153 | 5,74 | 114,12 | 34 | 1,264 | 1,15 | 1,155 | 1,067 | 0,888 | 1,168 | 1,054 | 1,062 | 1,11 | 0,11 | 1,5E-01 |
| P57776 | Elongation factor 1-delta OS=Mus musculus OX=10090 GN=Eef1d P | 17 | 4 | 5 | 3 | 281 | 31,3 | 5,02 | 13,45 | 4 | 0,986 | 1,064 | 1,433 | 1,162 | 0,929 | 1,077 | 0,985 | 0,982 | 1,17 | 0,17 | 1,5E-01 |
| Q9Z307 | Inward rectifier potassium channel 16 OS=Mus musculus OX=1009 | 2 | 1 | 1 | 1 | 419 | 48 | 7,83 | 0 | 1 | 1,005 | 0,832 | 0,832 | 0,883 | 0,913 | 0,93 | 0,995 | 1,069 | 0,91 | -0,09 | 1,5E-01 |
| Q3UFS0 | Protein zyg-11 homolog B OS=Mus musculus OX=10090 GN=Zyg11 | 3 | 2 | 2 | 2 | 744 | 83,9 | 6,87 | 2,85 | 2 | 1,455 | 1,403 | 1,298 | 1,083 | 1,055 | 1,157 | 1,243 | 1,185 | 1,13 | 0,13 | 1,5E-01 |
| P21278 | Guanine nucleotide-binding protein subunit alpha-11 OS=Mus mus | 33 | 10 | 12 | 5 | 359 | 42 | 5,97 | 33,13 | 10 | 1,049 | 1,02 | 0,989 | 1,061 | 1,035 | 1,051 | 1,128 | 1,075 | 0,96 | -0,04 | 1,5E-01 |
| P62827 | GTP-binding nuclear protein Ran OS=Mus musculus OX=10090 GN= | 21 | 5 | 8 | 5 | 216 | 24,4 | 7,49 | 13,98 | 5 | 1,018 | 0,98 | 1,016 | 1,042 | 0,92 | 1,019 | 0,976 | 0,983 | 1,04 | 0,04 | 1,5E-01 |
| Q9D8X2 | Coiled-coil domain-containing protein 124 OS=Mus musculus OX=1 | 6 | 1 | 1 | 1 | 217 | 25,3 | 9,64 | 2,36 | 1 | 1,821 | 1,506 | 0,984 | 0,736 | 0,647 | 0,581 | 1,021 | 1,014 | 1,55 | 0,55 | 1,5E-01 |
| Q921G7 | Electron transfer flavoprotein-ubiquinone oxidoreductase, mitoch | 16 | 9 | 9 | 9 | 616 | 68 | 7,58 | 10,97 | 9 | 1,008 | 0,994 | 1,018 | 1,044 | 0,992 | 1,141 | 1,052 | 1,1 | 0,95 | -0,05 | 1,5E-01 |
| Q8CHY3 | Dymeclin OS=Mus musculus OX=10090 GN=Dym PE=1 SV=1 | 1 | 1 | 1 | 1 | 669 | 75,8 | 5,77 | 0 | 1 | 1,277 | 0,793 | 1,278 | 1,021 | 0,939 | 1,064 | 0,645 | 0,736 | 1,29 | 0,29 | 1,5E-01 |
| P53995 | Anaphase-promoting complex subunit 1 OS=Mus musculus OX=100 | 0 | 1 | 1 | 1 | 1944 | 215,9 | 6,35 | 1,96 | 1 | 1,235 | 1,258 | 1,423 | 1,096 | 1,104 | 1,325 | 0,9 | 0,846 | 1,20 | 0,20 | 1,5E-01 |
| Q8VC42 | Regulator of MON1-CCZ1 complex OS=Mus musculus OX=10090 GN | 3 | 2 | 2 | 2 | 657 | 74,9 | 7,83 | 2,01 | 2 | 1,132 | 0,776 | 0,889 | 0,836 | 1,072 | 0,991 | 1,115 | 1,001 | 0,87 | -0,13 | 1,5E-01 |
| P35283 | Ras-related protein Rab-12 OS=Mus musculus OX=10090 GN=Rab1 | 21 | 5 | 6 | 4 | 243 | 27,3 | 8,41 | 14,53 | 5 | 0,896 | 0,869 | 1,033 | 1,007 | 0,966 | 1,061 | 1,028 | 1,047 | 0,93 | -0,07 | 1,5E-01 |
| Q80W21 | Glutathione S-transferase Mu 7 OS=Mus musculus OX=10090 GN= | 28 | 6 | 8 | 4 | 218 | 25,7 | 6,8 | 23,07 | 6 | 1,376 | 1,099 | 1,136 | 1,094 | 0,938 | 1,149 | 0,93 | 1,112 | 1,14 | 0,14 | 1,5E-01 |
| P35550 | rRNA 2'-O-methyltransferase fibrillarin OS=Mus musculus OX=1009 | 3 | 1 | 1 | 1 | 327 | 34,3 | 10,24 | 0 | 1 | 3,47 | 1,78 | 0,864 | 1,506 | 0,893 | 0,998 | 0,724 | 1,296 | 1,95 | 0,95 | 1,5E-01 |
| P09528 | Ferritin heavy chain OS=Mus musculus OX=10090 GN=Fth1 PE=1 S | 21 | 4 | 4 | 4 | 182 | 21,1 | 5,88 | 5,91 | 4 | 0,65 | 0,811 | 0,865 | 0,759 | 0,927 | 0,721 | 1,01 | 0,928 | 0,86 | -0,14 | 1,5E-01 |
| B0V2N1-3 | Isoform 3 of Receptor-type tyrosine-protein phosphatase S OS=Mu | 9 | 12 | 14 | 10 | 1501 | 168,2 | 6,81 | 28,09 | 12 | 1,04 | 0,996 | 1,043 | 1,053 | 1,038 | 1,046 | 1,071 | 1,091 | 0,97 | -0,03 | 1,5E-01 |
| P09671 | Superoxide dismutase [Mn], mitochondrial OS=Mus musculus OX= | 23 | 6 | 16 | 6 | 222 | 24,6 | 8,62 | 44,93 | 6 | 0,869 | 0,988 | 0,9 | 0,937 | 0,993 | 0,939 | 0,95 | 1,017 | 0,95 | -0,05 | 1,5E-01 |
| Q923X4 | Glutaredoxin-2, mitochondrial OS=Mus musculus OX=10090 GN=Gl | 7 | 1 | 1 | 1 | 156 | 17,3 | 9,95 | 0 | 1 | 1,394 | 1,036 | 1,082 | 1,085 | 1,496 | 1,205 | 1,213 | 1,392 | 0,87 | -0,13 | 1,5E-01 |
| Q9D0S9 | Histidine triad nucleotide-binding protein 2, mitochondrial OS=Mus | 21 | 3 | 3 | 3 | 163 | 17,3 | 9,82 | 11,34 | 3 | 1,149 | 1,085 | 1,191 | 1,04 | 0,87 | 1,191 | 0,91 | 0,981 | 1,13 | 0,13 | 1,6E-01 |
| Q6Y685 | Transforming acidic coiled-coil-containing protein 1 OS=Mus muscu | 5 | 4 | 5 | 3 | 774 | 83,9 | 5,03 | 10,95 | 4 | 1,028 | 1,217 | 1,118 | 1,086 | 0,967 | 1,148 | 0,978 | 0,971 | 1,09 | 0,09 | 1,6E-01 |
| Q99JP0 | Mitogen-activated protein kinase kinase kinase kinase 3 OS=Mus m | 4 | 3 | 3 | 3 | 894 | 101,1 | 7,64 | 5,69 | 3 | 1,416 | 1,164 | 1,422 | 1,386 | 1,101 | 1,265 | 1,208 | 1,315 | 1,10 | 0,10 | 1,6E-01 |
| Q9QUG9 | RAS guanyl-releasing protein 2 OS=Mus musculus OX=10090 GN=R | 6 | 3 | 3 | 3 | 608 | 69,4 | 7,66 | 4,56 | 3 | 1,519 | 1,232 | 1,202 | 1,405 | 1,356 | 1,427 | 1,819 | 1,58 | 0,87 | -0,13 | 1,6E-01 |
| Q7TPR4 | Alpha-actinin-1 OS=Mus musculus OX=10090 GN=Actn1 PE=1 SV=1 | 44 | 38 | 64 | 25 | 892 | 103 | 5,38 | 135,36 | 38 | 0,984 | 0,89 | 0,879 | 0,943 | 1,069 | 0,901 | 0,987 | 1,017 | 0,93 | -0,07 | 1,6E-01 |
| P70296 | Phosphatidylethanolamine-binding protein 1 OS=Mus musculus OX | 29 | 4 | 8 | 4 | 187 | 20,8 | 5,4 | 20,5 | 4 | 1,506 | 1,31 | 1,278 | 1,092 | 0,878 | 1,361 | 1,041 | 1,051 | 1,20 | 0,20 | 1,6E-01 |

| | | | | | | | | | | | | | | | | | | | | | |
|---|---|---|---|---|---|---|---|---|---|---|---|---|---|---|---|---|---|---|---|---|---|
| P52189 | Inward rectifier potassium channel 4 OS=Mus musculus OX=10090 | 3 | 1 | 1 | 1 | 445 | 49,9 | 6,21 | 2,37 | 1 | 1,279 | 1,181 | 1,317 | 1,221 | 0,921 | 1,351 | 1,076 | 1,023 | 1,14 | 0,14 | 1,6E-01 |
| O89116 | Vesicle transport through interaction with t-SNAREs homolog 1A O | 8 | 2 | 2 | 2 | 217 | 25 | 6,4 | 2,19 | 2 | 0,823 | 0,584 | 0,93 | 1,173 | 1,092 | 0,949 | 1,209 | 1,126 | 0,80 | -0,20 | 1,6E-01 |
| Q9ES74 | Serine/threonine-protein kinase Nek7 OS=Mus musculus OX=1009 | 7 | 2 | 2 | 2 | 302 | 34,5 | 8,25 | 2,12 | 2 | 1,023 | 0,811 | 1,058 | 0,929 | 0,815 | 0,73 | 0,959 | 0,846 | 1,14 | 0,14 | 1,6E-01 |
| Q9JM76 | Actin-related protein 2/3 complex subunit 3 OS=Mus musculus OX= | 10 | 2 | 3 | 2 | 178 | 20,5 | 8,59 | 7,09 | 2 | 0,772 | 0,69 | 0,848 | 0,948 | 0,929 | 0,875 | 0,973 | 0,87 | 0,89 | -0,11 | 1,6E-01 |
| Q3TW96 | UDP-N-acetylhexosamine pyrophosphorylase-like protein 1 OS=Mu | 3 | 1 | 1 | 1 | 507 | 56,6 | 5,43 | 3,39 | 1 | 2,871 | 1,788 | 1,874 | 1,415 | 1,062 | 1,88 | 1,368 | 1,349 | 1,40 | 0,40 | 1,6E-01 |
| Q5M8N0 | CB1 cannabinoid receptor-interacting protein 1 OS=Mus musculus | 21 | 3 | 3 | 3 | 164 | 18,6 | 7,96 | 7,4 | 3 | 0,883 | 0,801 | 0,929 | 1,067 | 1,03 | 0,942 | 1,102 | 1,023 | 0,90 | -0,10 | 1,6E-01 |
| Q80Y83 | Dixin OS=Mus musculus OX=10090 GN=Dixdc1 PE=1 SV=1 | 2 | 1 | 1 | 1 | 711 | 80,2 | 6,18 | 2,32 | 1 | 0,655 | 0,96 | 1,151 | 0,806 | 0,81 | 0,417 | 0,568 | 0,835 | 1,36 | 0,36 | 1,6E-01 |
| Q80UP3 | Diacylglycerol kinase zeta OS=Mus musculus OX=10090 GN=Dgkz P | 11 | 8 | 9 | 8 | 929 | 104 | 8,09 | 14,91 | 8 | 0,928 | 0,911 | 1,013 | 0,987 | 0,975 | 1,106 | 1,033 | 0,977 | 0,94 | -0,06 | 1,6E-01 |
| Q9CR21 | Acyl carrier protein, mitochondrial OS=Mus musculus OX=10090 GI | 12 | 2 | 2 | 2 | 156 | 17,4 | 5,21 | 0 | 2 | 0,815 | 1,04 | 0,873 | 0,933 | 1,015 | 1,101 | 0,974 | 0,948 | 0,91 | -0,09 | 1,6E-01 |
| Q9Z2G9 | Oxidoreductase HTATIP2 OS=Mus musculus OX=10090 GN=Htatip2 | 10 | 2 | 2 | 2 | 242 | 26,9 | 8,59 | 0 | 2 | 1,208 | 0,887 | 0,787 | 0,968 | 0,807 | 0,648 | 0,77 | 0,935 | 1,22 | 0,22 | 1,6E-01 |
| P11627 | Neural cell adhesion molecule L1 OS=Mus musculus OX=10090 GN= | 7 | 7 | 8 | 7 | 1260 | 140,9 | 5,97 | 20,37 | 7 | 0,96 | 0,784 | 0,96 | 0,939 | 1,057 | 0,892 | 1,093 | 0,994 | 0,90 | -0,10 | 1,6E-01 |
| Q99LY9 | NADH dehydrogenase [ubiquinone] iron-sulfur protein 5 OS=Mus n | 45 | 4 | 5 | 4 | 106 | 12,6 | 8,92 | 16,84 | 4 | 1,113 | 1,159 | 1,128 | 1,051 | 1,044 | 1,121 | 0,938 | 1,063 | 1,07 | 0,07 | 1,6E-01 |
| Q9EP75 | Leukotriene-B4 omega-hydroxylase 3 OS=Mus musculus OX=10090 | 2 | 1 | 1 | 1 | 524 | 59,8 | 6,87 | 0 | 1 | 1,442 | 1,272 | 1,097 | 1,114 | 0,963 | 1,155 | 1,144 | 1,075 | 1,14 | 0,14 | 1,6E-01 |
| O88544 | COP9 signalosome complex subunit 4 OS=Mus musculus OX=10090 | 32 | 12 | 14 | 12 | 406 | 46,3 | 5,83 | 26,05 | 12 | 0,915 | 0,987 | 0,972 | 0,995 | 0,98 | 0,987 | 1,008 | 1,031 | 0,97 | -0,03 | 1,6E-01 |
| Q9D8U8 | Sorting nexin-5 OS=Mus musculus OX=10090 GN=Snx5 PE=1 SV=1 | 16 | 5 | 5 | 5 | 404 | 46,8 | 6,62 | 11,94 | 5 | 1,303 | 1,51 | 1,229 | 1,102 | 0,949 | 1,265 | 1,15 | 1,091 | 1,15 | 0,15 | 1,6E-01 |
| Q9D172 | Glutamine amidotransferase-like class 1 domain-containing proteir | 51 | 9 | 11 | 9 | 266 | 28,1 | 8,78 | 40,53 | 9 | 1,115 | 0,922 | 1,135 | 1,17 | 1,202 | 1,098 | 1,263 | 1,196 | 0,91 | -0,09 | 1,6E-01 |
| Q6P4T0 | Autophagy-related protein 2 homolog A OS=Mus musculus OX=100 | 1 | 1 | 1 | 1 | 1914 | 210,8 | 6,06 | 2,03 | 1 | 1,273 | 1,182 | 1,181 | 0,983 | 1,052 | 1,154 | 0,766 | 0,995 | 1,16 | 0,16 | 1,6E-01 |
| P60202 | Myelin proteolipid protein OS=Mus musculus OX=10090 GN=Plp1 F | 20 | 5 | 23 | 5 | 277 | 30,1 | 8,35 | 51,67 | 5 | 0,944 | 1,028 | 1,013 | 1,125 | 1,018 | 1,009 | 0,865 | 0,816 | 1,11 | 0,11 | 1,6E-01 |
| Q8BXK8 | Arf-GAP with GTPase, ANK repeat and PH domain-containing prote | 5 | 4 | 4 | 4 | 857 | 94,4 | 7,94 | 10,02 | 4 | 1,149 | 1,067 | 1,211 | 0,978 | 1,013 | 1,09 | 0,964 | 0,966 | 1,09 | 0,09 | 1,6E-01 |
| Q62188 | Dihydropyrimidinase-related protein 3 OS=Mus musculus OX=1009 | 36 | 16 | 40 | 14 | 570 | 61,9 | 6,49 | 106,49 | 16 | 0,913 | 0,999 | 1,056 | 1,141 | 0,944 | 0,891 | 0,972 | 0,974 | 1,09 | 0,09 | 1,6E-01 |
| Q9DBC7 | cAMP-dependent protein kinase type I-alpha regulatory subunit OS | 15 | 7 | 7 | 5 | 381 | 43,2 | 5,35 | 16,5 | 7 | 0,775 | 0,847 | 0,914 | 0,944 | 1,103 | 0,865 | 0,94 | 0,967 | 0,90 | -0,10 | 1,6E-01 |
| O54774 | AP-3 complex subunit delta-1 OS=Mus musculus OX=10090 GN=Ap | 11 | 12 | 16 | 12 | 1199 | 135 | 7,37 | 29,98 | 12 | 0,866 | 0,975 | 1,041 | 1,061 | 0,989 | 1,193 | 1,068 | 1,078 | 0,91 | -0,09 | 1,6E-01 |
| A2AP18-2 | Isoform 2 of 1-phosphatidylinositol 4,5-bisphosphate phosphodiest | 10 | 11 | 11 | 11 | 1238 | 136,2 | 7,2 | 16,33 | 11 | 1,197 | 1,108 | 1,137 | 1,01 | 1,082 | 1,085 | 1,014 | 0,958 | 1,08 | 0,08 | 1,6E-01 |
| Q8BPQ7 | Small G protein signaling modulator 1 OS=Mus musculus OX=1009 | 6 | 6 | 7 | 6 | 1093 | 123,1 | 5,66 | 8,27 | 6 | 1,008 | 0,92 | 0,941 | 0,833 | 1,099 | 1,026 | 0,905 | 1,013 | 0,92 | -0,08 | 1,6E-01 |
| Q9JL26 | Formin-like protein 1 OS=Mus musculus OX=10090 GN=Fmnl1 PE=1 | 5 | 5 | 5 | 5 | 1094 | 122 | 5,82 | 5,09 | 5 | 1,118 | 1,047 | 1,006 | 1,143 | 0,948 | 0,926 | 1,036 | 1,09 | 1,08 | 0,08 | 1,6E-01 |
| Q8R3Q2 | Serine/threonine-protein phosphatase 6 regulatory subunit 2 OS=N | 5 | 3 | 4 | 3 | 923 | 100,4 | 4,82 | 11,75 | 3 | 1,553 | 1,474 | 1,305 | 1,284 | 1,236 | 1,316 | 1,186 | 1,382 | 1,10 | 0,10 | 1,6E-01 |
| Q9D9V3 | Ethylmalonyl-CoA decarboxylase OS=Mus musculus OX=10090 GN= | 7 | 2 | 2 | 2 | 322 | 35,4 | 7,01 | 5,32 | 2 | 1,484 | 1,308 | 1,384 | 1,051 | 1,083 | 1,308 | 1,118 | 1,005 | 1,16 | 0,16 | 1,6E-01 |
| Q9CQC6 | Basic leucine zipper and W2 domain-containing protein 1 OS=Mus i | 10 | 4 | 4 | 4 | 419 | 48 | 5,92 | 7,81 | 4 | 1,125 | 1,049 | 1,119 | 1,053 | 0,896 | 1,142 | 0,924 | 1,011 | 1,09 | 0,09 | 1,7E-01 |
| O88533 | Aromatic-L-amino-acid decarboxylase OS=Mus musculus OX=1009 | 3 | 1 | 2 | 1 | 480 | 53,8 | 6,6 | 4,26 | 1 | 1,149 | 0,846 | 1,132 | 0,761 | 1,16 | 1,24 | 1,497 | 0,945 | 0,80 | -0,20 | 1,7E-01 |
| Q8CFI5 | Probable proline--tRNA ligase, mitochondrial OS=Mus musculus OX | 5 | 3 | 3 | 3 | 475 | 53,5 | 7,84 | 4,31 | 3 | 0,856 | 0,77 | 1,056 | 1,157 | 1,197 | 1,143 | 1,059 | 1,046 | 0,86 | -0,14 | 1,7E-01 |
| Q9Z280 | Phospholipase D1 OS=Mus musculus OX=10090 GN=Pld1 PE=1 SV= | 1 | 1 | 1 | 1 | 1074 | 123,9 | 8,72 | 2,14 | 1 | 0,68 | 1,16 | 1,093 | 0,883 | 0,788 | 0,944 | 0,479 | 0,682 | 1,32 | 0,32 | 1,7E-01 |
| Q8CB59 | Protein FAM161B OS=Mus musculus OX=10090 GN=Fam161b PE=1 | 3 | 1 | 1 | 1 | 589 | 66,9 | 9,54 | 0 | 1 | 2,358 | 1,82 | 1,361 | 1,311 | 0,84 | 1,652 | 1,212 | 1,283 | 1,37 | 0,37 | 1,7E-01 |
| P83940 | Elongin-C OS=Mus musculus OX=10090 GN=Eloc PE=1 SV=1 | 38 | 3 | 3 | 3 | 112 | 12,5 | 4,78 | 10,15 | 3 | 2,055 | 1,421 | 1,44 | 1,292 | 0,977 | 1,519 | 1,027 | 1,335 | 1,28 | 0,28 | 1,7E-01 |
| P84075 | Neuron-specific calcium-binding protein hippocalcin OS=Mus musc | 48 | 9 | 12 | 4 | 193 | 22,4 | 4,97 | 29,05 | 9 | 0,811 | 0,673 | 1,006 | 1,011 | 1,005 | 1,172 | 1,057 | 0,897 | 0,85 | -0,15 | 1,7E-01 |
| P45952 | Medium-chain specific acyl-CoA dehydrogenase, mitochondrial OS= | 23 | 8 | 10 | 8 | 421 | 46,5 | 8,37 | 19,21 | 8 | 1,174 | 1,247 | 1,136 | 1,009 | 0,945 | 1,17 | 0,984 | 1,026 | 1,11 | 0,11 | 1,7E-01 |
| Q9CWE0 | Mitochondrial fission regulator 1-like OS=Mus musculus OX=10090 | 8 | 2 | 3 | 2 | 289 | 31,7 | 6,1 | 2,85 | 2 | 1,176 | 1,458 | 0,859 | 0,948 | 0,886 | 0,861 | 0,908 | 0,937 | 1,24 | 0,24 | 1,7E-01 |
| P24527 | Leukotriene A-4 hydrolase OS=Mus musculus OX=10090 GN=Lta4h | 17 | 9 | 12 | 9 | 611 | 69 | 6,42 | 26,19 | 9 | 1,1 | 1,066 | 1,046 | 1,003 | 0,893 | 1,083 | 0,961 | 0,998 | 1,07 | 0,07 | 1,7E-01 |
| Q99JT2 | Serine/threonine-protein kinase 26 OS=Mus musculus OX=10090 G | 10 | 4 | 4 | 2 | 416 | 46,6 | 5,24 | 8,18 | 4 | 1,327 | 1,183 | 1,335 | 1,06 | 0,843 | 1,083 | 1,02 | 1,269 | 1,16 | 0,16 | 1,7E-01 |
| Q8VD37 | SH3-containing GRB2-like protein 3-interacting protein 1 OS=Mus n | 18 | 11 | 16 | 11 | 806 | 86 | 7,87 | 40,77 | 11 | 0,866 | 0,862 | 0,934 | 0,934 | 0,933 | 0,876 | 1,022 | 1,028 | 0,93 | -0,07 | 1,7E-01 |
| Q9D2G2 | Dihydrolipoyllysine-residue succinyltransferase component of 2-ox | 17 | 6 | 13 | 6 | 454 | 49 | 8,95 | 18,21 | 6 | 0,898 | 0,876 | 0,983 | 0,921 | 1 | 0,9 | 0,973 | 1,039 | 0,94 | -0,06 | 1,7E-01 |
| P70174 | Histamine H1 receptor OS=Mus musculus OX=10090 GN=Hrh1 PE= | 3 | 1 | 1 | 1 | 488 | 55,6 | 9,32 | 2,67 | 1 | 1,294 | 1,23 | 1,494 | 1,134 | 1,115 | 0,736 | 0,814 | 1,411 | 1,26 | 0,26 | 1,7E-01 |
| Q9CWD8 | Iron-sulfur protein NUBPL OS=Mus musculus OX=10090 GN=Nubpl | 13 | 3 | 3 | 3 | 319 | 34,1 | 9,07 | 6,05 | 3 | 0,979 | 1,16 | 0,89 | 0,842 | 0,916 | 0,96 | 0,676 | 0,661 | 1,20 | 0,20 | 1,7E-01 |
| P51174 | Long-chain specific acyl-CoA dehydrogenase, mitochondrial OS=Mu | 14 | 5 | 6 | 5 | 430 | 47,9 | 8,31 | 10,26 | 5 | 0,946 | 0,891 | 0,995 | 1,202 | 0,851 | 0,969 | 0,902 | 0,853 | 1,13 | 0,13 | 1,7E-01 |
| Q6NVG1 | Lysophospholipid acyltransferase LPCAT4 OS=Mus musculus OX=1C | 3 | 1 | 1 | 1 | 524 | 57,1 | 8,75 | 0 | 1 | 1,467 | 1,732 | 1,116 | 0,802 | 0,609 | 1,232 | 0,664 | 1,037 | 1,44 | 0,44 | 1,7E-01 |
| Q8R574 | Phosphoribosyl pyrophosphate synthase-associated protein 2 OS=N | 10 | 4 | 5 | 3 | 369 | 40,9 | 7,17 | 9,57 | 4 | 0,67 | 0,993 | 0,936 | 0,976 | 1,013 | 0,954 | 1,024 | 1,084 | 0,88 | -0,12 | 1,7E-01 |
| O88492 | Perilipin-4 OS=Mus musculus OX=10090 GN=Plin4 PE=1 SV=2 | 5 | 2 | 3 | 2 | 1403 | 139,3 | 8,59 | 7,86 | 2 | 1,52 | 1,528 | 1,154 | 0,989 | 0,824 | 1,25 | 0,905 | 1,159 | 1,25 | 0,25 | 1,7E-01 |
| Q6ZPY7 | Lysine-specific demethylase 3B OS=Mus musculus OX=10090 GN=K | 1 | 1 | 1 | 1 | 1562 | 170,8 | 6,92 | 0 | 1 | 0,839 | 0,809 | 1,047 | 0,786 | 1,069 | 0,857 | 1,14 | 0,955 | 0,87 | -0,13 | 1,7E-01 |
| P45591 | Cofilin-2 OS=Mus musculus OX=10090 GN=Cfl2 PE=1 SV=1 | 21 | 5 | 5 | 3 | 166 | 18,7 | 7,88 | 14,34 | 5 | 0,825 | 0,919 | 1,01 | 1,004 | 1,042 | 1,023 | 1,009 | 0,971 | 0,93 | -0,07 | 1,7E-01 |
| Q9Z2U1 | Proteasome subunit alpha type-5 OS=Mus musculus OX=10090 GN | 22 | 5 | 10 | 5 | 241 | 26,4 | 4,79 | 18,82 | 5 | 0,983 | 1,057 | 0,917 | 0,939 | 0,973 | 0,869 | 0,905 | 0,914 | 1,06 | 0,06 | 1,7E-01 |
| Q9CZT8 | Ras-related protein Rab-3B OS=Mus musculus OX=10090 GN=Rab3 | 32 | 8 | 16 | 3 | 219 | 24,7 | 5,11 | 32,43 | 8 | 0,809 | 0,819 | 0,993 | 1,003 | 0,998 | 1,024 | 1,14 | 0,908 | 0,89 | -0,11 | 1,7E-01 |
| Q61696 | Heat shock 70 kDa protein 1A OS=Mus musculus OX=10090 GN=Hs | 29 | 18 | 29 | 11 | 641 | 70 | 5,72 | 57,34 | 18 | 1,091 | 1,041 | 0,952 | 1,026 | 1,013 | 1,095 | 1,145 | 1,106 | 0,94 | -0,06 | 1,7E-01 |
| Q61490 | CD166 antigen OS=Mus musculus OX=10090 GN=Alcam PE=1 SV=3 | 11 | 9 | 11 | 9 | 583 | 65,1 | 6,15 | 16,14 | 9 | 0,962 | 0,962 | 0,952 | 0,947 | 1,063 | 0,928 | 1,105 | 0,978 | 0,94 | -0,06 | 1,7E-01 |
| P28828 | Receptor-type tyrosine-protein phosphatase mu OS=Mus musculus | 2 | 2 | 2 | 2 | 1452 | 163,5 | 6,68 | 2,01 | 2 | 1,337 | 0,937 | 1,295 | 1,904 | 1,326 | 0,896 | 0,956 | 0,891 | 1,35 | 0,35 | 1,7E-01 |
| Q9ER72 | Cysteine--tRNA ligase, cytoplasmic OS=Mus musculus OX=10090 GI | 4 | 3 | 3 | 3 | 831 | 94,8 | 6,76 | 7,38 | 3 | 1,371 | 1,185 | 1,046 | 1,037 | 0,921 | 1,05 | 0,989 | 1,123 | 1,14 | 0,14 | 1,7E-01 |
| Q8QZS1 | 3-hydroxyisobutyryl-CoA hydrolase, mitochondrial OS=Mus muscul | 23 | 9 | 10 | 9 | 385 | 43 | 8,06 | 19,93 | 9 | 0,996 | 0,969 | 0,982 | 0,93 | 1,013 | 1,082 | 0,995 | 0,965 | 0,96 | -0,04 | 1,7E-01 |
| Q8C878 | NEDD8-activating enzyme E1 catalytic subunit OS=Mus musculus O | 15 | 4 | 4 | 4 | 462 | 51,7 | 5,45 | 5,59 | 4 | 1,276 | 1,219 | 1,101 | 1 | 0,862 | 1,162 | 0,898 | 1,083 | 1,15 | 0,15 | 1,7E-01 |
| Q7TPD2 | Protein FAM185A OS=Mus musculus OX=10090 GN=Fam185a PE=2 | 4 | 1 | 1 | 1 | 378 | 40 | 6,38 | 2,72 | 1 | 0,694 | 0,807 | 1,257 | 1,278 | 1,088 | 1,259 | 1,326 | 1,384 | 0,80 | -0,20 | 1,7E-01 |
| Q91YT0 | NADH dehydrogenase [ubiquinone] flavoprotein 1, mitochondrial C | 33 | 12 | 22 | 12 | 464 | 50,8 | 8,21 | 57,8 | 12 | 0,839 | 0,863 | 0,962 | 0,969 | 1,016 | 0,905 | 0,991 | 0,976 | 0,93 | -0,07 | 1,7E-01 |
| Q61753 | D-3-phosphoglycerate dehydrogenase OS=Mus musculus OX=1009 | 26 | 11 | 14 | 11 | 533 | 56,5 | 6,54 | 34,97 | 11 | 1,193 | 1,089 | 1,1 | 1,021 | 0,953 | 1,165 | 0,916 | 0,954 | 1,10 | 0,10 | 1,7E-01 |
| Q924S8 | Sprouty-related, EVH1 domain-containing protein 1 OS=Mus muscu | 3 | 1 | 1 | 1 | 444 | 50,6 | 6,47 | 3,45 | 1 | 1,252 | 1,277 | 1,005 | 1,516 | 1,074 | 1,064 | 1,225 | 0,952 | 1,17 | 0,17 | 1,7E-01 |
| Q8BHL5-3 | Isoform 3 of Engulfment and cell motility protein 2 OS=Mus muscul | 7 | 4 | 4 | 3 | 798 | 91,2 | 6,57 | 5,77 | 4 | 1,327 | 0,924 | 1,296 | 1,124 | 1,049 | 0,982 | 1,028 | 1,031 | 1,14 | 0,14 | 1,7E-01 |
| O88447 | Kinesin light chain 1 OS=Mus musculus OX=10090 GN=Klc1 PE=1 SV | 23 | 12 | 17 | 8 | 541 | 61,4 | 5,68 | 27,09 | 12 | 1,021 | 1,096 | 0,948 | 0,954 | 0,853 | 0,936 | 0,955 | 0,993 | 1,08 | 0,08 | 1,7E-01 |

| | | | | | | | | | | | | | | | | | | | | | |
|---|---|---|---|---|---|---|---|---|---|---|---|---|---|---|---|---|---|---|---|---|---|
| Q9QYR6 | Microtubule-associated protein 1A OS=Mus musculus OX=10090 G | 20 | 55 | 70 | 54 | 2776 | 300 | 5 | 122,04 | 55 | 0,957 | 0,919 | 1,007 | 0,997 | 1,022 | 0,98 | 1,014 | 1,001 | 0,97 | -0,03 | 1,7E-01 |
| Q8CHU3 | Epsin-2 OS=Mus musculus OX=10090 GN=Epn2 PE=1 SV=1 | 17 | 7 | 7 | 7 | 595 | 63,4 | 8,16 | 18,26 | 7 | 1,14 | 1,05 | 0,96 | 1,045 | 0,908 | 1,003 | 1,024 | 0,984 | 1,07 | 0,07 | 1,7E-01 |
| Q9Z0H8 | CAP-Gly domain-containing linker protein 2 OS=Mus musculus OX= | 24 | 21 | 28 | 15 | 1047 | 115,8 | 6,48 | 57,79 | 21 | 1,167 | 1,099 | 1,125 | 1,013 | 0,909 | 1,093 | 1,053 | 1,032 | 1,08 | 0,08 | 1,7E-01 |
| P20444 | Protein kinase C alpha type OS=Mus musculus OX=10090 GN=Prkca | 23 | 16 | 23 | 9 | 672 | 76,8 | 7,14 | 38,91 | 16 | 0,96 | 0,897 | 0,947 | 1,017 | 1,147 | 0,926 | 1,068 | 1,007 | 0,92 | -0,08 | 1,7E-01 |
| Q91YX5 | Acyl-CoA:lysophosphatidylglycerol acyltransferase 1 OS=Mus musc | 6 | 2 | 3 | 2 | 370 | 43,1 | 8,79 | 2,52 | 2 | 0,918 | 1,018 | 0,856 | 0,92 | 0,971 | 0,947 | 1,066 | 0,989 | 0,93 | -0,07 | 1,7E-01 |
| P60755 | MAM domain-containing glycosylphosphatidylinositol anchor prote | 1 | 1 | 1 | 1 | 949 | 106,6 | 7,02 | 2,32 | 1 | 1,175 | 0,781 | 0,937 | 0,911 | 1,124 | 1,039 | 1,144 | 1,034 | 0,88 | -0,12 | 1,7E-01 |
| Q9CQ06 | 39S ribosomal protein L24, mitochondrial OS=Mus musculus OX=1( | 4 | 1 | 1 | 1 | 216 | 24,9 | 9,5 | 2,4 | 1 | 0,708 | 1,095 | 0,816 | 0,964 | 1,025 | 0,963 | 1,022 | 1,149 | 0,86 | -0,14 | 1,7E-01 |
| Q6P5F7 | Protein tweety homolog 3 OS=Mus musculus OX=10090 GN=Ttyh3 | 4 | 2 | 4 | 2 | 524 | 57,7 | 5,85 | 8,36 | 2 | 1,322 | 1,202 | 1,103 | 1,193 | 1,019 | 1,057 | 1,099 | 1,239 | 1,09 | 0,09 | 1,7E-01 |
| P62754 | 40S ribosomal protein S6 OS=Mus musculus OX=10090 GN=Rps6 PE | 13 | 4 | 5 | 4 | 249 | 28,7 | 10,84 | 12,47 | 4 | 1,301 | 1,294 | 1,215 | 1,055 | 0,968 | 1,286 | 0,971 | 1,06 | 1,14 | 0,14 | 1,7E-01 |
| Q14CH7 | Alanine--tRNA ligase, mitochondrial OS=Mus musculus OX=10090 G | 4 | 2 | 2 | 2 | 980 | 106,7 | 6,18 | 2,79 | 2 | 1,538 | 1,219 | 1,195 | 0,84 | 0,851 | 1,135 | 0,687 | 1,045 | 1,29 | 0,29 | 1,7E-01 |
| Q5SSH7 | Zinc finger ZZ-type and EF-hand domain-containing protein 1 OS=M | 2 | 4 | 4 | 4 | 2924 | 328,1 | 6,11 | 5,87 | 4 | 1,237 | 1,116 | 1,101 | 1,17 | 1,106 | 1,064 | 1,114 | 1,131 | 1,05 | 0,05 | 1,7E-01 |
| Q811I0 | ATP synthase mitochondrial F1 complex assembly factor 1 OS=Mus | 7 | 2 | 2 | 2 | 324 | 36,3 | 8,19 | 0 | 2 | 0,834 | 1,205 | 1,162 | 0,924 | 0,994 | 0,622 | 0,55 | 1,021 | 1,29 | 0,29 | 1,7E-01 |
| Q923S9 | Ras-related protein Rab-30 OS=Mus musculus OX=10090 GN=Rab3( | 10 | 3 | 4 | 1 | 203 | 23 | 4,97 | 11,09 | 3 | 0,986 | 0,854 | 0,92 | 1,07 | 0,964 | 1,187 | 0,977 | 1,168 | 0,89 | -0,11 | 1,7E-01 |
| Q05920 | Pyruvate carboxylase, mitochondrial OS=Mus musculus OX=10090 | 19 | 22 | 29 | 22 | 1178 | 129,6 | 6,71 | 69,46 | 22 | 0,994 | 1,015 | 1,006 | 0,997 | 1,084 | 1,077 | 0,992 | 1,007 | 0,96 | -0,04 | 1,8E-01 |
| P31750 | RAC-alpha serine/threonine-protein kinase OS=Mus musculus OX=: | 5 | 2 | 2 | 2 | 480 | 55,7 | 5,9 | 0 | 2 | 0,601 | 0,844 | 1,199 | 0,991 | 1,201 | 0,991 | 1,096 | 1,17 | 0,82 | -0,18 | 1,8E-01 |
| Q8VHJ5 | Serine/threonine-protein kinase MARK1 OS=Mus musculus OX=100 | 16 | 10 | 11 | 7 | 795 | 88,3 | 9,39 | 17,26 | 10 | 1,14 | 1,016 | 0,988 | 1,046 | 0,991 | 0,915 | 1,049 | 0,971 | 1,07 | 0,07 | 1,8E-01 |
| Q99J77 | Sialic acid synthase OS=Mus musculus OX=10090 GN=Nans PE=1 SV | 7 | 2 | 2 | 2 | 359 | 40 | 7,06 | 5,41 | 2 | 1,099 | 1,422 | 1,168 | 1,06 | 1,08 | 1,088 | 1,017 | 1,056 | 1,12 | 0,12 | 1,8E-01 |
| P00405 | Cytochrome c oxidase subunit 2 OS=Mus musculus OX=10090 GN=I | 18 | 5 | 9 | 5 | 227 | 26 | 4,73 | 21,3 | 5 | 1,31 | 1,349 | 1,216 | 0,938 | 0,937 | 1,123 | 0,955 | 1,141 | 1,16 | 0,16 | 1,8E-01 |
| P62881 | Guanine nucleotide-binding protein subunit beta-5 OS=Mus muscu | 14 | 5 | 6 | 5 | 395 | 43,5 | 6,46 | 16,71 | 5 | 0,67 | 0,829 | 0,881 | 1,1 | 1,209 | 0,788 | 1,097 | 1,193 | 0,81 | -0,19 | 1,8E-01 |
| Q8BL03 | Mitochondrial basic amino acids transporter OS=Mus musculus OX: | 4 | 1 | 1 | 1 | 306 | 32,7 | 8,62 | 2,36 | 1 | 1 | 1,003 | 1,232 | 1,157 | 0,843 | 0,862 | 0,883 | 1,19 | 1,16 | 0,16 | 1,8E-01 |
| Q99NB1 | Acetyl-coenzyme A synthetase 2-like, mitochondrial OS=Mus muscu | 8 | 5 | 5 | 5 | 682 | 74,6 | 6,98 | 6,64 | 5 | 0,726 | 1,19 | 0,992 | 1,103 | 1,08 | 1,18 | 1,15 | 1,255 | 0,86 | -0,14 | 1,8E-01 |
| Q8BMF3 | NADP-dependent malic enzyme, mitochondrial OS=Mus musculus ( | 20 | 11 | 16 | 10 | 604 | 67,1 | 7,83 | 36,42 | 11 | 0,924 | 0,997 | 1,011 | 1,002 | 1,021 | 1,004 | 1,084 | 0,997 | 0,96 | -0,04 | 1,8E-01 |
| Q99N28 | Cell adhesion molecule 3 OS=Mus musculus OX=10090 GN=Cadm3 | 30 | 10 | 19 | 10 | 396 | 42,9 | 5,8 | 40,08 | 10 | 0,946 | 0,948 | 0,97 | 1,009 | 0,997 | 0,952 | 1,036 | 1,046 | 0,96 | -0,04 | 1,8E-01 |
| Q99KI3 | ER membrane protein complex subunit 3 OS=Mus musculus OX=10 | 8 | 2 | 2 | 2 | 261 | 30 | 6,81 | 4,44 | 2 | 0,694 | 1,032 | 0,949 | 1,024 | 1,053 | 1,186 | 1,143 | 0,919 | 0,86 | -0,14 | 1,8E-01 |
| P04925 | Major prion protein OS=Mus musculus OX=10090 GN=Prnp PE=1 S\ | 9 | 2 | 4 | 2 | 254 | 28 | 9,33 | 13,3 | 2 | 0,81 | 0,794 | 1,014 | 0,973 | 1,187 | 1,014 | 1,162 | 0,832 | 0,86 | -0,14 | 1,8E-01 |
| Q9Z1N9 | Protein unc-13 homolog B OS=Mus musculus OX=10090 GN=Unc13 | 4 | 6 | 6 | 1 | 1602 | 181,7 | 5,95 | 6,78 | 6 | 0,718 | 0,879 | 0,937 | 0,818 | 0,77 | 0,57 | 0,885 | 0,579 | 1,20 | 0,20 | 1,8E-01 |
| Q9CPY7 | Cytosol aminopeptidase OS=Mus musculus OX=10090 GN=Lap3 PE: | 25 | 11 | 19 | 11 | 519 | 56,1 | 7,72 | 27,76 | 11 | 1,199 | 1,029 | 1,12 | 1,21 | 1,093 | 1,001 | 1,028 | 1,126 | 1,07 | 0,07 | 1,8E-01 |
| Q9D0M1 | Phosphoribosyl pyrophosphate synthase-associated protein 1 OS=N | 5 | 2 | 3 | 1 | 356 | 39,4 | 7,2 | 7,54 | 2 | 1,166 | 1,382 | 1,006 | 1,033 | 0,898 | 1,141 | 1,056 | 0,746 | 1,19 | 0,19 | 1,8E-01 |
| O35855 | Branched-chain-amino-acid aminotransferase, mitochondrial OS=M | 6 | 2 | 2 | 2 | 393 | 44,1 | 8,29 | 2 | 2 | 1,036 | 1,217 | 1,046 | 1,307 | 0,91 | 1,107 | 1,085 | 1,018 | 1,12 | 0,12 | 1,8E-01 |
| Q68FL4 | Putative adenosylhomocysteinase 3 OS=Mus musculus OX=10090 ( | 18 | 10 | 15 | 4 | 613 | 66,9 | 7,36 | 28,47 | 10 | 0,718 | 0,709 | 1,031 | 0,957 | 1,008 | 0,953 | 1,051 | 0,931 | 0,87 | -0,13 | 1,8E-01 |
| P51855 | Glutathione synthetase OS=Mus musculus OX=10090 GN=Gss PE=1 | 8 | 3 | 3 | 3 | 474 | 52,2 | 5,8 | 2,2 | 3 | 1,943 | 0,924 | 1,237 | 1,027 | 1,001 | 0,971 | 0,899 | 0,847 | 1,38 | 0,38 | 1,8E-01 |
| Q9JHI5 | Isovaleryl-CoA dehydrogenase, mitochondrial OS=Mus musculus O) | 20 | 7 | 8 | 7 | 424 | 46,3 | 8,29 | 16,37 | 7 | 1,094 | 0,831 | 1,048 | 0,894 | 1,14 | 1,059 | 1,016 | 1,062 | 0,90 | -0,10 | 1,8E-01 |
| Q9WV27 | Sodium/potassium-transporting ATPase subunit alpha-4 OS=Mus m | 9 | 11 | 59 | 1 | 1032 | 114,8 | 5,71 | 128,76 | 11 | 1,156 | 1,474 | 1,015 | 1,016 | 0,976 | 1,046 | 0,921 | 1,037 | 1,17 | 0,17 | 1,8E-01 |
| P48542 | G protein-activated inward rectifier potassium channel 2 OS=Mus r | 2 | 1 | 2 | 1 | 425 | 48,6 | 5,38 | 4,85 | 1 | 0,631 | 0,996 | 0,649 | 0,86 | 0,968 | 0,923 | 0,797 | 1,111 | 0,83 | -0,17 | 1,8E-01 |
| Q9D7S9 | Charged multivesicular body protein 5 OS=Mus musculus OX=1009( | 7 | 1 | 1 | 1 | 219 | 24,6 | 4,79 | 2,1 | 1 | 1,891 | 1,378 | 1,383 | 1,137 | 0,855 | 1,448 | 1,281 | 0,897 | 1,29 | 0,29 | 1,8E-01 |
| Q8VEK0 | Cell cycle control protein 50A OS=Mus musculus OX=10090 GN=Tm | 10 | 3 | 4 | 3 | 364 | 41 | 8,37 | 7,07 | 3 | 0,938 | 0,909 | 0,912 | 0,936 | 0,884 | 0,973 | 1,178 | 1,039 | 0,91 | -0,09 | 1,8E-01 |
| P62264 | 40S ribosomal protein S14 OS=Mus musculus OX=10090 GN=Rps14 | 40 | 6 | 8 | 6 | 151 | 16,3 | 10,05 | 17,21 | 6 | 1,281 | 1,271 | 1,128 | 0,927 | 0,955 | 1,075 | 1,047 | 1,004 | 1,13 | 0,13 | 1,8E-01 |
| Q9DC69 | NADH dehydrogenase [ubiquinone] 1 alpha subcomplex subunit 9, | 30 | 11 | 12 | 11 | 377 | 42,5 | 9,74 | 24,79 | 11 | 0,874 | 0,927 | 0,947 | 1,039 | 1,126 | 0,924 | 1,048 | 1,017 | 0,92 | -0,08 | 1,8E-01 |
| Q9JJR8 | Transmembrane protein 9B OS=Mus musculus OX=10090 GN=Tme | 7 | 2 | 2 | 2 | 199 | 22,6 | 8,18 | 1,95 | 2 | 0,383 | 0,747 | 0,717 | 0,919 | 1,078 | 0,639 | 0,923 | 1,029 | 0,75 | -0,25 | 1,8E-01 |
| Q9R0P9 | Ubiquitin carboxyl-terminal hydrolase isozyme L1 OS=Mus musculu | 54 | 12 | 24 | 12 | 223 | 24,8 | 5,24 | 55,23 | 12 | 1,053 | 1,1 | 0,971 | 0,99 | 0,891 | 1,057 | 0,918 | 0,964 | 1,07 | 0,07 | 1,8E-01 |
| Q9QYS9 | Protein quaking OS=Mus musculus OX=10090 GN=Qki PE=1 SV=1 | 4 | 1 | 1 | 1 | 341 | 37,6 | 8,56 | 2,3 | 1 | 0,01 | 0,93 | 0,631 | 1,246 | 1,009 | 1,128 | 1,34 | 1,003 | 0,63 | -0,37 | 1,8E-01 |
| Q8BGQ7 | Alanine--tRNA ligase, cytoplasmic OS=Mus musculus OX=10090 GN | 20 | 18 | 24 | 18 | 968 | 106,8 | 5,67 | 43,44 | 18 | 1,155 | 1,076 | 1,036 | 1,038 | 0,937 | 1,012 | 1,045 | 1,069 | 1,06 | 0,06 | 1,8E-01 |
| Q8C460 | ERI1 exoribonuclease 3 OS=Mus musculus OX=10090 GN=Eri3 PE=1 | 6 | 2 | 2 | 2 | 337 | 37,2 | 8,07 | 2,12 | 2 | 0,346 | 0,661 | 0,752 | 0,846 | 0,824 | 1,004 | 0,773 | 0,748 | 0,78 | -0,22 | 1,8E-01 |
| Q99NE5 | Regulating synaptic membrane exocytosis protein 1 OS=Mus muscu | 13 | 16 | 19 | 6 | 1463 | 163,1 | 9,42 | 33,44 | 16 | 1,007 | 1,005 | 0,904 | 1,035 | 0,962 | 0,907 | 0,946 | 0,948 | 1,05 | 0,05 | 1,8E-01 |
| Q80TL7 | Protein MON2 homolog OS=Mus musculus OX=10090 GN=Mon2 PE | 2 | 4 | 4 | 4 | 1715 | 189 | 6,13 | 9,76 | 4 | 1,387 | 1,193 | 1,206 | 0,981 | 0,898 | 1,192 | 0,922 | 1,096 | 1,16 | 0,16 | 1,8E-01 |
| Q6ZPR4 | Potassium channel subfamily T member 1 OS=Mus musculus OX=1( | 1 | 1 | 1 | 1 | 1224 | 138 | 7,52 | 2,69 | 1 | 1,337 | 1,059 | 1,502 | 0,986 | 1,066 | 1,117 | 0,976 | 0,964 | 1,18 | 0,18 | 1,8E-01 |
| Q810J8 | Zinc finger FYVE domain-containing protein 1 OS=Mus musculus O) | 15 | 9 | 10 | 9 | 777 | 86,9 | 7,34 | 7,44 | 9 | 1,313 | 1,306 | 1,142 | 1,047 | 0,935 | 1,097 | 1,019 | 1,22 | 1,13 | 0,13 | 1,8E-01 |
| O89001 | Carboxypeptidase D OS=Mus musculus OX=10090 GN=Cpd PE=1 SV | 1 | 1 | 1 | 1 | 1377 | 152,3 | 6,18 | 0 | 1 | 0,528 | 0,794 | 1,133 | 0,994 | 0,872 | 1,678 | 1,131 | 1,076 | 0,73 | -0,27 | 1,8E-01 |
| P58044 | Isopentenyl-diphosphate Delta-isomerase 1 OS=Mus musculus OX= | 10 | 2 | 2 | 2 | 227 | 26,3 | 6,16 | 0 | 2 | 1,157 | 1,014 | 1,068 | 1,002 | 0,985 | 0,773 | 0,946 | 1,088 | 1,12 | 0,12 | 1,8E-01 |
| Q922Q9 | Chitinase domain-containing protein 1 OS=Mus musculus OX=1009 | 9 | 3 | 3 | 3 | 393 | 44,9 | 7,97 | 4,16 | 3 | 0,999 | 0,888 | 0,844 | 1,113 | 1,155 | 1,073 | 1,004 | 1,027 | 0,90 | -0,10 | 1,8E-01 |
| Q8VE38 | Oxidoreductase NAD-binding domain-containing protein 1 OS=Mus | 11 | 3 | 3 | 3 | 311 | 34,7 | 8,13 | 6,66 | 3 | 1,243 | 1,267 | 1,137 | 1,129 | 0,952 | 1,153 | 1,14 | 1,161 | 1,08 | 0,08 | 1,8E-01 |
| Q6P1F6 | Serine/threonine-protein phosphatase 2A 55 kDa regulatory subun | 18 | 10 | 12 | 4 | 447 | 51,7 | 6,2 | 24,76 | 10 | 0,933 | 0,946 | 0,99 | 1,077 | 0,896 | 0,732 | 0,964 | 0,968 | 1,11 | 0,11 | 1,8E-01 |
| Q3TXS7 | 26S proteasome non-ATPase regulatory subunit 1 OS=Mus musculu | 13 | 12 | 13 | 12 | 953 | 105,7 | 5,39 | 24,51 | 12 | 1 | 0,934 | 1,036 | 1,006 | 1,016 | 1,004 | 1,044 | 1,067 | 0,96 | -0,04 | 1,8E-01 |
| Q61495 | Desmoglein-1-alpha OS=Mus musculus OX=10090 GN=Dsg1a PE=2 | 2 | 2 | 3 | 2 | 1057 | 114,5 | 4,89 | 4,65 | 2 | 2,619 | 2,096 | 1,582 | 0,913 | 0,858 | 1,546 | 1,059 | 1,367 | 1,49 | 0,49 | 1,8E-01 |
| Q80YA9 | Connector enhancer of kinase suppressor of ras 2 OS=Mus musculu | 6 | 6 | 6 | 6 | 1032 | 117,3 | 6,79 | 9,61 | 6 | 0,745 | 0,969 | 0,967 | 1,066 | 1,16 | 1,083 | 0,978 | 1,004 | 0,89 | -0,11 | 1,8E-01 |
| Q69Z98 | Serine/threonine-protein kinase BRSK2 OS=Mus musculus OX=1009 | 17 | 10 | 11 | 8 | 735 | 81,7 | 8,79 | 21,97 | 10 | 0,952 | 1,004 | 1,005 | 1,141 | 1,019 | 1,153 | 1,158 | 1,085 | 0,93 | -0,07 | 1,8E-01 |
| Q9D1H6 | NADH dehydrogenase [ubiquinone] 1 alpha subcomplex assembly f | 11 | 2 | 2 | 2 | 173 | 20,1 | 9,39 | 4,59 | 2 | 0,837 | 0,92 | 0,869 | 0,922 | 0,887 | 1,049 | 0,976 | 0,896 | 0,93 | -0,07 | 1,8E-01 |
| Q9D4H8 | Cullin-2 OS=Mus musculus OX=10090 GN=Cul2 PE=1 SV=2 | 6 | 5 | 5 | 5 | 745 | 86,8 | 7,01 | 8,95 | 5 | 0,878 | 0,689 | 0,98 | 1,027 | 1,041 | 0,959 | 1,116 | 0,961 | 0,88 | -0,12 | 1,8E-01 |
| Q9EPW0 | Inositol polyphosphate-4-phosphatase type I A OS=Mus musculus ( | 10 | 7 | 9 | 7 | 939 | 105,5 | 7,05 | 22,08 | 7 | 0,903 | 0,948 | 0,998 | 1,027 | 1,068 | 0,984 | 1,012 | 1,008 | 0,95 | -0,05 | 1,8E-01 |
| Q05BC3 | Echinoderm microtubule-associated protein-like 1 OS=Mus muscul | 12 | 8 | 11 | 8 | 814 | 89,6 | 7,02 | 23,46 | 8 | 1,195 | 0,974 | 1,121 | 1,044 | 0,946 | 1,046 | 1,033 | 0,992 | 1,08 | 0,08 | 1,8E-01 |
| Q8BIK4 | Dedicator of cytokinesis protein 9 OS=Mus musculus OX=10090 GN | 1 | 3 | 3 | 3 | 2055 | 235,2 | 7,25 | 7,44 | 3 | 1,255 | 1,165 | 1,115 | 1,006 | 0,977 | 1,194 | 0,923 | 0,968 | 1,12 | 0,12 | 1,8E-01 |
| Q9JJK2 | LanC-like protein 2 OS=Mus musculus OX=10090 GN=Lancl2 PE=1 S | 15 | 6 | 9 | 6 | 450 | 50,7 | 7,28 | 17,39 | 6 | 1,538 | 1,143 | 1,142 | 1 | 0,978 | 1,112 | 0,927 | 1,068 | 1,18 | 0,18 | 1,8E-01 |

| | | | | | | | | | | | | | | | | | | | | | |
|---|---|---|---|---|---|---|---|---|---|---|---|---|---|---|---|---|---|---|---|---|---|
| O08915 | AH receptor-interacting protein OS=Mus musculus OX=10090 GN= | 15 | 4 | 5 | 4 | 330 | 37,6 | 6,4 | 5,32 | 4 | 1,094 | 1,061 | 1,139 | 1,143 | 1,064 | 0,996 | 1,143 | 0,989 | 1,06 | 0,06 | 1,8E-01 |
| A2APX8 | Sodium channel protein type 1 subunit alpha OS=Mus musculus O | 7 | 15 | 20 | 8 | 2009 | 228,7 | 5,87 | 44,48 | 15 | 0,758 | 0,897 | 0,81 | 0,94 | 0,964 | 0,84 | 0,921 | 1,005 | 0,91 | -0,09 | 1,8E-01 |
| Q8K411 | Presequence protease, mitochondrial OS=Mus musculus OX=1009 | 7 | 6 | 7 | 6 | 1036 | 117,3 | 7,2 | 17,69 | 6 | 0,909 | 0,937 | 0,931 | 0,957 | 0,92 | 1,033 | 0,951 | 0,987 | 0,96 | -0,04 | 1,9E-01 |
| P99027 | 60S acidic ribosomal protein P2 OS=Mus musculus OX=10090 GN= | 25 | 3 | 4 | 3 | 115 | 11,6 | 4,54 | 5,47 | 3 | 1,02 | 0,861 | 0,99 | 1,139 | 0,867 | 0,983 | 0,942 | 0,797 | 1,12 | 0,12 | 1,9E-01 |
| Q6KAU4 | Multivesicular body subunit 12B OS=Mus musculus OX=10090 GN= | 6 | 2 | 2 | 2 | 317 | 35,4 | 7,93 | 2,44 | 2 | 0,769 | 1,215 | 0,7 | 0,723 | 1,097 | 0,998 | 1,023 | 1,029 | 0,82 | -0,18 | 1,9E-01 |
| Q61831 | Mitogen-activated protein kinase 10 OS=Mus musculus OX=10090 | 11 | 4 | 4 | 3 | 464 | 52,5 | 6,96 | 12,66 | 4 | 1,174 | 1,143 | 1,07 | 0,913 | 0,88 | 1,025 | 1,02 | 0,973 | 1,10 | 0,10 | 1,9E-01 |
| Q60649 | Caseinolytic peptidase B protein homolog OS=Mus musculus OX=1 | 5 | 3 | 3 | 3 | 677 | 76 | 8,51 | 4,82 | 3 | 0,98 | 0,94 | 1,174 | 1,023 | 1,086 | 1,311 | 1,174 | 1,026 | 0,90 | -0,10 | 1,9E-01 |
| P21812 | Mast cell protease 4 OS=Mus musculus OX=10090 GN=Mcpt4 PE=1 | 5 | 1 | 1 | 1 | 246 | 27,2 | 9,48 | 3,05 | 1 | 2,201 | 1,293 | 1,614 | 1,16 | 0,761 | 1,334 | 1,054 | 1,454 | 1,36 | 0,36 | 1,9E-01 |
| Q3TWI9 | CSC1-like protein 2 OS=Mus musculus OX=10090 GN=Tmem63b PE | 1 | 1 | 1 | 1 | 832 | 94,7 | 7,49 | 0 | 1 | 0,791 | 0,932 | 0,73 | 0,899 | 0,934 | 0,896 | 0,858 | 1,178 | 0,87 | -0,13 | 1,9E-01 |
| P60487 | Pyridoxal phosphate phosphatase OS=Mus musculus OX=10090 GN | 15 | 4 | 4 | 4 | 292 | 31,5 | 5,74 | 9,52 | 4 | 1,244 | 1,027 | 1,073 | 0,937 | 0,795 | 1,029 | 0,907 | 1,036 | 1,14 | 0,14 | 1,9E-01 |
| P14869 | 60S acidic ribosomal protein P0 OS=Mus musculus OX=10090 GN= | 16 | 4 | 5 | 4 | 317 | 34,2 | 6,25 | 3,62 | 4 | 0,879 | 1,023 | 1,025 | 0,83 | 0,872 | 0,906 | 0,855 | 0,796 | 1,10 | 0,10 | 1,9E-01 |
| Q91YP2 | Neurolysin, mitochondrial OS=Mus musculus OX=10090 GN=Nln PE | 8 | 5 | 5 | 5 | 704 | 80,4 | 6,44 | 12,32 | 5 | 0,8 | 0,866 | 0,881 | 0,879 | 0,899 | 1,055 | 0,845 | 0,917 | 0,92 | -0,08 | 1,9E-01 |
| O08599-2 | Isoform 2 of Syntaxin-binding protein 1 OS=Mus musculus OX=100 | 53 | 34 | 86 | 1 | 603 | 68,7 | 6,77 | 194,12 | 34 | 1,247 | 1,308 | 0,961 | 0,915 | 0,907 | 1,072 | 0,883 | 0,925 | 1,17 | 0,17 | 1,9E-01 |
| Q99LG2 | Transportin-2 OS=Mus musculus OX=10090 GN=Tnpo2 PE=1 SV=1 | 5 | 3 | 5 | 3 | 887 | 100,4 | 4,98 | 5,74 | 3 | 1,131 | 1,316 | 1,212 | 1,145 | 0,905 | 1,259 | 1,056 | 1,084 | 1,12 | 0,12 | 1,9E-01 |
| Q8K3H0 | DCC-interacting protein 13-alpha OS=Mus musculus OX=10090 GN | 21 | 11 | 12 | 11 | 707 | 79,3 | 5,41 | 31,97 | 11 | 1,121 | 1,121 | 1,064 | 0,988 | 0,929 | 1,102 | 0,928 | 1,023 | 1,08 | 0,08 | 1,9E-01 |
| P62821 | Ras-related protein Rab-1A OS=Mus musculus OX=10090 GN=Rab1 | 44 | 11 | 23 | 7 | 205 | 22,7 | 6,21 | 65,1 | 11 | 0,986 | 0,901 | 1,013 | 1,037 | 1,048 | 0,989 | 1,096 | 1,025 | 0,95 | -0,05 | 1,9E-01 |
| Q99K46 | Ubiquitin carboxyl-terminal hydrolase 11 OS=Mus musculus OX=10 | 2 | 2 | 2 | 2 | 921 | 105,3 | 4,96 | 4,46 | 2 | 0,658 | 1,176 | 1 | 1,421 | 1,481 | 1,067 | 1,599 | 1,287 | 0,78 | -0,22 | 1,9E-01 |
| Q9Z351-1 | Isoform 12 of Potassium voltage-gated channel subfamily KQT men | 6 | 4 | 5 | 4 | 870 | 95,7 | 9,23 | 13,64 | 4 | 1,02 | 0,97 | 0,851 | 1,079 | 1,054 | 1,123 | 1,014 | 1,047 | 0,92 | -0,08 | 1,9E-01 |
| Q78PY7 | Staphylococcal nuclease domain-containing protein 1 OS=Mus mus | 8 | 7 | 8 | 7 | 910 | 102 | 7,43 | 9,39 | 7 | 1,462 | 1,471 | 1,207 | 0,913 | 0,769 | 1,294 | 0,919 | 1,048 | 1,25 | 0,25 | 1,9E-01 |
| Q80U30 | Protein CLEC16A OS=Mus musculus OX=10090 GN=Clec16a PE=1 S | 2 | 2 | 2 | 2 | 1036 | 116,2 | 6,05 | 2,25 | 2 | 0,849 | 0,844 | 0,959 | 0,885 | 1 | 0,989 | 0,845 | 0,967 | 0,93 | -0,07 | 1,9E-01 |
| Q9D8Y1 | Transmembrane protein 126A OS=Mus musculus OX=10090 GN=Tr | 9 | 2 | 2 | 2 | 196 | 21,5 | 9,41 | 3,97 | 2 | 1,053 | 1,088 | 0,853 | 0,864 | 1,143 | 1,47 | 0,855 | 1,226 | 0,82 | -0,18 | 1,9E-01 |
| P08003 | Protein disulfide-isomerase A4 OS=Mus musculus OX=10090 GN=P | 14 | 10 | 13 | 10 | 638 | 71,9 | 5,31 | 23,75 | 10 | 1,138 | 1,067 | 1,216 | 1,141 | 1,091 | 1,095 | 1,092 | 1,103 | 1,04 | 0,04 | 1,9E-01 |
| P84086 | Complexin-2 OS=Mus musculus OX=10090 GN=Cplx2 PE=1 SV=1 | 36 | 4 | 6 | 2 | 134 | 15,4 | 5,08 | 8,17 | 4 | 0,875 | 0,619 | 0,917 | 0,999 | 0,923 | 0,996 | 1,002 | 0,986 | 0,87 | -0,13 | 1,9E-01 |
| Q6GQT9 | Nodal modulator 1 OS=Mus musculus OX=10090 GN=Nomo1 PE=1 | 9 | 10 | 10 | 10 | 1214 | 133,3 | 6,09 | 17,94 | 10 | 0,854 | 0,889 | 0,935 | 0,876 | 0,947 | 1,106 | 0,851 | 0,977 | 0,92 | -0,08 | 1,9E-01 |
| P34152 | Focal adhesion kinase 1 OS=Mus musculus OX=10090 GN=Ptk2 PE= | 5 | 5 | 6 | 5 | 1052 | 119,2 | 6,62 | 13,75 | 5 | 1,144 | 1,097 | 1,136 | 0,935 | 0,802 | 1,053 | 0,995 | 1,021 | 1,11 | 0,11 | 1,9E-01 |
| Q8BLC3 | Ly6/PLAUR domain-containing protein 1 OS=Mus musculus OX=10 | 9 | 1 | 1 | 1 | 141 | 15,3 | 7,64 | 1,67 | 1 | 0,89 | 1,448 | 1,66 | 1,191 | 1,1 | 0,963 | 1,08 | 1,048 | 1,24 | 0,24 | 1,9E-01 |
| Q5DU25 | IQ motif and SEC7 domain-containing protein 2 OS=Mus musculus | 14 | 17 | 19 | 16 | 1478 | 161,7 | 8,56 | 34,84 | 17 | 1,039 | 1,083 | 1,019 | 1,002 | 1,037 | 1,022 | 0,978 | 0,924 | 1,05 | 0,05 | 1,9E-01 |
| Q9DCL8 | Protein phosphatase inhibitor 2 OS=Mus musculus OX=10090 GN= | 12 | 1 | 1 | 1 | 206 | 23,1 | 4,83 | 3,9 | 1 | 1,249 | 1,017 | 0,853 | 0,851 | 0,78 | 0,881 | 0,964 | 0,686 | 1,20 | 0,20 | 1,9E-01 |
| Q9WTL7 | Acyl-protein thioesterase 2 OS=Mus musculus OX=10090 GN=Lypla | 8 | 1 | 1 | 1 | 231 | 24,8 | 7,23 | 2,97 | 1 | 1,417 | 1,065 | 1,305 | 1,458 | 0,848 | 1,444 | 0,952 | 1,076 | 1,21 | 0,21 | 1,9E-01 |
| Q8JZU2 | Tricarboxylate transport protein, mitochondrial OS=Mus musculus | 15 | 5 | 5 | 5 | 311 | 33,9 | 9,89 | 13,36 | 5 | 0,77 | 0,792 | 0,999 | 1,019 | 1,079 | 0,964 | 1,103 | 0,909 | 0,88 | -0,12 | 1,9E-01 |
| Q9DBC0 | Protein adenylyltransferase SelO, mitochondrial OS=Mus musculus | 4 | 2 | 2 | 2 | 667 | 74,2 | 5,83 | 4,06 | 2 | 1,263 | 1,134 | 1,175 | 1,144 | 1,146 | 0,991 | 1,175 | 1,11 | 1,07 | 0,07 | 1,9E-01 |
| Q7TNM2 | Tripartite motif-containing protein 46 OS=Mus musculus OX=1009 | 4 | 2 | 2 | 2 | 759 | 83,4 | 7,65 | 3,12 | 2 | 1,531 | 1,508 | 1,364 | 1,044 | 0,969 | 1,319 | 1,297 | 1,003 | 1,19 | 0,19 | 1,9E-01 |
| Q8C547 | HEAT repeat-containing protein 5B OS=Mus musculus OX=10090 G | 1 | 3 | 3 | 3 | 2070 | 224,2 | 7,14 | 4,52 | 3 | 1,03 | 1,143 | 1,188 | 1,175 | 1,173 | 1,355 | 1,185 | 1,164 | 0,93 | -0,07 | 1,9E-01 |
| Q8R464 | Cell adhesion molecule 4 OS=Mus musculus OX=10090 GN=Cadm4 | 19 | 5 | 5 | 5 | 388 | 42,7 | 6,3 | 14,47 | 5 | 1,276 | 1,169 | 1,073 | 1,032 | 0,963 | 1,162 | 1,017 | 0,997 | 1,10 | 0,10 | 1,9E-01 |
| Q6PIP5 | NudC domain-containing protein 1 OS=Mus musculus OX=10090 G | 5 | 2 | 2 | 2 | 582 | 66,7 | 5,3 | 2,97 | 2 | 0,957 | 1,061 | 0,972 | 0,924 | 0,968 | 1,079 | 1,055 | 1,034 | 0,95 | -0,05 | 1,9E-01 |
| Q91YN5 | UDP-N-acetylhexosamine pyrophosphorylase OS=Mus musculus O | 3 | 1 | 1 | 1 | 522 | 58,6 | 6,49 | 3,42 | 1 | 1,24 | 0,949 | 1,08 | 1,07 | 0,77 | 1,067 | 1,031 | 0,947 | 1,14 | 0,14 | 1,9E-01 |
| Q9DCR2 | AP-3 complex subunit sigma-1 OS=Mus musculus OX=10090 GN=A | 11 | 2 | 2 | 2 | 193 | 21,7 | 5,39 | 4,98 | 2 | 1,383 | 1,149 | 1,065 | 1,01 | 0,87 | 0,966 | 0,913 | 1,205 | 1,17 | 0,17 | 1,9E-01 |
| P59759 | Myocardin-related transcription factor B OS=Mus musculus OX=10 | 4 | 3 | 3 | 3 | 1080 | 117,5 | 6,16 | 2,49 | 3 | 1,2 | 1,144 | 1,653 | 1,235 | 1,23 | 1,103 | 1,023 | 1,147 | 1,16 | 0,16 | 1,9E-01 |
| Q8CHS8 | Vacuolar protein sorting-associated protein 37A OS=Mus musculus | 6 | 2 | 2 | 2 | 397 | 44,4 | 5,67 | 2,63 | 2 | 0,972 | 0,988 | 0,975 | 1,018 | 1,111 | 1,023 | 1,02 | 0,976 | 0,96 | -0,04 | 1,9E-01 |
| Q8BKX6 | Serine/threonine-protein kinase SMG1 OS=Mus musculus OX=100 | 0 | 1 | 1 | 1 | 3658 | 409,5 | 6,4 | 2,51 | 1 | 1,24 | 0,991 | 1 | 1,35 | 1,009 | 0,981 | 0,91 | 1,106 | 1,14 | 0,14 | 1,9E-01 |
| Q8R5M8- | Isoform 4 of Cell adhesion molecule 1 OS=Mus musculus OX=1009 | 9 | 2 | 3 | 2 | 417 | 45,8 | 5,12 | 10,21 | 2 | 2,224 | 1,575 | 1,162 | 1,12 | 0,94 | 1,366 | 1,097 | 1,094 | 1,35 | 0,35 | 1,9E-01 |
| Q09LZ8 | Cytosolic carboxypeptidase 6 OS=Mus musculus OX=10090 GN=Ag | 6 | 2 | 2 | 2 | 540 | 62,2 | 7,36 | 2,08 | 2 | 1,441 | 1,718 | 1,488 | 1,588 | 1,713 | 1,049 | 1,427 | 0,996 | 1,20 | 0,20 | 1,9E-01 |
| Q9DAS9 | Guanine nucleotide-binding protein G(I)/G(S)/G(O) subunit gamma | 46 | 4 | 4 | 4 | 72 | 8 | 8,97 | 11,23 | 4 | 1,413 | 1,189 | 1,138 | 1,005 | 1,094 | 1,091 | 1,045 | 1,003 | 1,12 | 0,12 | 1,9E-01 |
| Q8K596 | Sodium/calcium exchanger 2 OS=Mus musculus OX=10090 GN=Slc | 18 | 12 | 17 | 12 | 921 | 100,6 | 5,12 | 47,68 | 12 | 0,89 | 0,942 | 0,822 | 1,05 | 1,069 | 0,901 | 1,114 | 1,009 | 0,90 | -0,10 | 1,9E-01 |
| O70579 | Peroxisomal membrane protein PMP34 OS=Mus musculus OX=100 | 4 | 1 | 1 | 1 | 307 | 34,4 | 10,11 | 0 | 1 | 1,081 | 1,214 | 1,096 | 0,776 | 1,069 | 0,668 | 0,847 | 0,857 | 1,21 | 0,21 | 1,9E-01 |
| Q8BP92 | Reticulocalbin-2 OS=Mus musculus OX=10090 GN=Rcn2 PE=1 SV=1 | 26 | 6 | 7 | 6 | 320 | 37,2 | 4,42 | 12,41 | 6 | 0,881 | 1,03 | 0,979 | 0,821 | 1,044 | 1,185 | 0,944 | 0,961 | 0,90 | -0,10 | 1,9E-01 |
| P21107-2 | Isoform 2 of Tropomyosin alpha-3 chain OS=Mus musculus OX=100 | 50 | 19 | 23 | 7 | 248 | 29 | 4,78 | 55,17 | 19 | 1,202 | 1,047 | 1,117 | 0,984 | 0,955 | 1,023 | 1,032 | 1,043 | 1,07 | 0,07 | 1,9E-01 |
| Q9CQQ7 | ATP synthase F(0) complex subunit B1, mitochondrial OS=Mus mus | 38 | 16 | 31 | 16 | 256 | 28,9 | 9,06 | 69,4 | 16 | 0,935 | 0,945 | 0,923 | 0,92 | 1,031 | 0,925 | 0,957 | 0,948 | 0,96 | -0,04 | 1,9E-01 |
| Q7TMC8 | L-fucose kinase OS=Mus musculus OX=10090 GN=Fcsk PE=1 SV=1 | 2 | 2 | 2 | 2 | 1090 | 119,2 | 6,6 | 1,96 | 2 | 1,059 | 1,24 | 0,969 | 0,959 | 0,84 | 0,834 | 0,995 | 1,057 | 1,13 | 0,13 | 1,9E-01 |
| Q9R0H0 | Peroxisomal acyl-coenzyme A oxidase 1 OS=Mus musculus OX=100 | 4 | 2 | 2 | 2 | 661 | 74,6 | 8,48 | 4,87 | 2 | 1,483 | 1,269 | 1,37 | 1,247 | 1,149 | 1,426 | 1,031 | 1,186 | 1,12 | 0,12 | 2,0E-01 |
| Q99NB8 | Ubiquilin-4 OS=Mus musculus OX=10090 GN=Ubqln4 PE=1 SV=1 | 5 | 2 | 2 | 2 | 596 | 63,5 | 5,03 | 2,09 | 2 | 1,685 | 1,406 | 1,484 | 1,173 | 0,894 | 1,605 | 1,21 | 0,846 | 1,26 | 0,26 | 2,0E-01 |
| Q99K70 | Ras-related GTP-binding protein C OS=Mus musculus OX=10090 G | 11 | 3 | 4 | 3 | 398 | 44,1 | 5,1 | 6,84 | 3 | 1,84 | 1,7 | 1,295 | 1,015 | 0,935 | 1,388 | 1,022 | 1,251 | 1,27 | 0,27 | 2,0E-01 |
| Q8BHE3 | Caytaxin OS=Mus musculus OX=10090 GN=Atcay PE=1 SV=1 | 17 | 5 | 6 | 5 | 372 | 42,2 | 4,68 | 11,29 | 5 | 0,617 | 0,888 | 0,833 | 1,13 | 1,134 | 1,045 | 1,157 | 0,86 | 0,83 | -0,17 | 2,0E-01 |
| Q91YA2 | Serine/threonine-protein kinase H1 OS=Mus musculus OX=10090 | 2 | 1 | 1 | 1 | 424 | 48,1 | 9,83 | 2,06 | 1 | 0,808 | 0,984 | 0,683 | 0,752 | 0,968 | 1,076 | 0,838 | 0,844 | 0,87 | -0,13 | 2,0E-01 |
| Q8BG92 | Clavesin-2 OS=Mus musculus OX=10090 GN=Clvs2 PE=1 SV=1 | 16 | 4 | 4 | 2 | 327 | 37,9 | 6,2 | 10,53 | 4 | 0,923 | 0,904 | 1,036 | 1,053 | 0,88 | 0,787 | 0,856 | 1,026 | 1,10 | 0,10 | 2,0E-01 |
| Q8VE88 | Protein FAM114A2 OS=Mus musculus OX=10090 GN=Fam114a2 PE | 9 | 4 | 4 | 4 | 497 | 54 | 4,93 | 4,14 | 4 | 0,93 | 1,232 | 1,225 | 1,101 | 0,976 | 1,049 | 0,904 | 1,086 | 1,12 | 0,12 | 2,0E-01 |
| Q52KF3 | Protein spire homolog 1 OS=Mus musculus OX=10090 GN=Spire1 P | 8 | 4 | 5 | 4 | 598 | 68,4 | 9,01 | 2,08 | 4 | 1,069 | 0,866 | 0,82 | 1,026 | 0,877 | 0,491 | 0,763 | 0,961 | 1,22 | 0,22 | 2,0E-01 |
| Q78JW9 | Ubiquitin domain-containing protein UBFD1 OS=Mus musculus OX= | 12 | 4 | 4 | 4 | 368 | 40,1 | 8,85 | 4,12 | 4 | 1,276 | 1,157 | 1,305 | 1,235 | 0,964 | 0,97 | 1,205 | 1,305 | 1,12 | 0,12 | 2,0E-01 |
| P63318 | Protein kinase C gamma type OS=Mus musculus OX=10090 GN=Prk | 34 | 23 | 42 | 22 | 697 | 78,3 | 7,46 | 80,82 | 23 | 0,931 | 0,961 | 0,956 | 1,028 | 1,013 | 1,031 | 1,012 | 0,965 | 0,96 | -0,04 | 2,0E-01 |
| Q8QZR5 | Alanine aminotransferase 1 OS=Mus musculus OX=10090 GN=Gpt | 7 | 3 | 3 | 3 | 496 | 55,1 | 6,64 | 8,74 | 3 | 1,331 | 1,147 | 1,283 | 0,951 | 0,904 | 1,233 | 0,977 | 0,945 | 1,16 | 0,16 | 2,0E-01 |
| Q9JLI6 | Selenocysteine lyase OS=Mus musculus OX=10090 GN=Scly PE=1 S | 3 | 1 | 1 | 1 | 432 | 47,1 | 6,8 | 2,58 | 1 | 0,922 | 1,364 | 1,046 | 0,761 | 0,823 | 1,045 | 0,596 | 0,705 | 1,29 | 0,29 | 2,0E-01 |
| Q7TNS2 | MICOS complex subunit Mic10 OS=Mus musculus OX=10090 GN=M | 11 | 1 | 1 | 1 | 76 | 8,6 | 8,94 | 3 | 1 | 0,586 | 0,787 | 0,823 | 0,932 | 0,972 | 0,855 | 0,913 | 0,84 | 0,87 | -0,13 | 2,0E-01 |

| | | | | | | | | | | | | | | | | | | | | | |
|---|---|---|---|---|---|---|---|---|---|---|---|---|---|---|---|---|---|---|---|---|---|
| Q9EQZ6 | Rap guanine nucleotide exchange factor 4 OS=Mus musculus OX=1( | 17 | 16 | 18 | 16 | 1011 | 115,4 | 6,92 | 35,24 | 16 | 0,884 | 0,963 | 0,933 | 0,965 | 0,933 | 1,028 | 0,964 | 0,978 | 0,96 | -0,04 | 2,0E-01 |
| Q3V0G7 | GTPase-activating Rap/Ran-GAP domain-like protein 3 OS=Mus mu | 4 | 3 | 3 | 3 | 1038 | 115,4 | 7,17 | 2,3 | 3 | 0,821 | 0,809 | 0,89 | 0,734 | 0,553 | 0,868 | 0,62 | 0,764 | 1,16 | 0,16 | 2,0E-01 |
| Q9CZP7 | Hsp90 co-chaperone Cdc37-like 1 OS=Mus musculus OX=10090 GN | 3 | 1 | 1 | 1 | 335 | 38,4 | 5,39 | 2,74 | 1 | 0,92 | 0,668 | 1,018 | 1,135 | 0,959 | 1,236 | 1,126 | 1,082 | 0,85 | -0,15 | 2,0E-01 |
| Q8VCT3 | Aminopeptidase B OS=Mus musculus OX=10090 GN=Rnpep PE=1 S' | 15 | 8 | 12 | 8 | 650 | 72,4 | 5,35 | 20,7 | 8 | 1,043 | 1,111 | 0,904 | 0,981 | 1,033 | 1,083 | 1,074 | 1,127 | 0,94 | -0,06 | 2,0E-01 |
| Q9CQ75 | NADH dehydrogenase [ubiquinone] 1 alpha subcomplex subunit 2 ( | 39 | 5 | 8 | 5 | 99 | 10,9 | 9,99 | 17,26 | 5 | 0,761 | 0,879 | 0,878 | 0,943 | 1,046 | 0,818 | 0,976 | 0,975 | 0,91 | -0,09 | 2,0E-01 |
| Q8K1R3 | Polyribonucleotide nucleotidyltransferase 1, mitochondrial OS=Mu | 12 | 9 | 9 | 9 | 783 | 85,6 | 8,03 | 10,65 | 9 | 1,095 | 1,089 | 0,924 | 1,005 | 0,933 | 1,007 | 0,929 | 0,986 | 1,07 | 0,07 | 2,0E-01 |
| Q8K0F1 | TBC1 domain family member 23 OS=Mus musculus OX=10090 GN= | 2 | 1 | 1 | 1 | 684 | 76,4 | 5,36 | 2,58 | 1 | 2,242 | 1,301 | 1,613 | 1,333 | 1,279 | 1,486 | 1,283 | 1,108 | 1,26 | 0,26 | 2,0E-01 |
| Q8CGF6 | WD repeat-containing protein 47 OS=Mus musculus OX=10090 GN= | 11 | 9 | 10 | 9 | 920 | 102,2 | 5,97 | 18,63 | 9 | 1,121 | 1,145 | 1,164 | 1,094 | 0,992 | 1,07 | 1,145 | 1,109 | 1,05 | 0,05 | 2,0E-01 |
| B2RSH2 | Guanine nucleotide-binding protein G(i) subunit alpha-1 OS=Mus m | 21 | 7 | 14 | 4 | 354 | 40,3 | 5,97 | 36,91 | 7 | 1,21 | 1,217 | 1,018 | 1,047 | 0,977 | 1,154 | 1,014 | 0,944 | 1,10 | 0,10 | 2,0E-01 |
| Q9CQ92 | Mitochondrial fission 1 protein OS=Mus musculus OX=10090 GN=F | 17 | 3 | 4 | 3 | 152 | 17 | 8,53 | 7,9 | 3 | 0,866 | 0,941 | 1,065 | 0,957 | 0,986 | 0,981 | 1,129 | 1,041 | 0,93 | -0,07 | 2,0E-01 |
| P21107 | Tropomyosin alpha-3 chain OS=Mus musculus OX=10090 GN=Tpm3 | 48 | 17 | 23 | 4 | 285 | 33 | 4,72 | 39,42 | 17 | 0,734 | 0,679 | 0,954 | 1,159 | 1,292 | 0,818 | 1,147 | 1,122 | 0,81 | -0,19 | 2,0E-01 |
| Q9ERL9 | Guanylate cyclase soluble subunit alpha-1 OS=Mus musculus OX=1( | 5 | 3 | 3 | 3 | 691 | 77,5 | 7,17 | 3,56 | 3 | 1,254 | 1,081 | 1,173 | 1,045 | 0,984 | 1,219 | 0,976 | 0,876 | 1,12 | 0,12 | 2,0E-01 |
| Q8VD04 | GRIP1-associated protein 1 OS=Mus musculus OX=10090 GN=Gripa | 18 | 14 | 16 | 14 | 806 | 92,7 | 5,25 | 25,95 | 14 | 0,91 | 0,987 | 1,085 | 0,948 | 0,915 | 0,888 | 0,974 | 0,913 | 1,07 | 0,07 | 2,0E-01 |
| Q8BWM0 | Prostaglandin E synthase 2 OS=Mus musculus OX=10090 GN=Ptges | 11 | 3 | 4 | 3 | 384 | 43,3 | 9 | 4,79 | 3 | 1,168 | 1,093 | 1,271 | 1,184 | 1,226 | 1,295 | 1,194 | 1,243 | 0,95 | -0,05 | 2,0E-01 |
| Q2M3X8 | Phosphatase and actin regulator 1 OS=Mus musculus OX=10090 GN | 17 | 10 | 15 | 10 | 580 | 66,2 | 6,96 | 37,32 | 10 | 1,17 | 1,092 | 1,034 | 1,085 | 1,005 | 1,101 | 1,034 | 1,043 | 1,05 | 0,05 | 2,0E-01 |
| Q8BGA9 | Mitochondrial inner membrane protein OXA1L OS=Mus musculus ( | 5 | 2 | 2 | 2 | 433 | 48,2 | 9,61 | 2,43 | 2 | 0,688 | 1,112 | 1,062 | 1,183 | 1,74 | 0,842 | 1,217 | 1,532 | 0,76 | -0,24 | 2,0E-01 |
| P56375 | Acylphosphatase-2 OS=Mus musculus OX=10090 GN=Acyp2 PE=1 S | 19 | 2 | 5 | 2 | 106 | 11,9 | 9,26 | 12,07 | 2 | 0,679 | 0,778 | 0,889 | 1,122 | 1,04 | 0,877 | 1,142 | 1,036 | 0,85 | -0,15 | 2,0E-01 |
| Q8R105 | Vacuolar protein sorting-associated protein 37C OS=Mus musculus | 16 | 4 | 4 | 4 | 352 | 38,4 | 5,31 | 10,38 | 4 | 1,36 | 1,094 | 1,212 | 1,086 | 1,173 | 1,124 | 0,908 | 1,055 | 1,12 | 0,12 | 2,0E-01 |
| Q8CH77 | Neuron navigator 1 OS=Mus musculus OX=10090 GN=Nav1 PE=1 S' | 2 | 3 | 3 | 3 | 1875 | 202,2 | 8,06 | 0 | 3 | 1,052 | 0,963 | 1,058 | 1,114 | 0,948 | 1,075 | 0,7 | 0,976 | 1,13 | 0,13 | 2,0E-01 |
| Q7TN29 | Stromal membrane-associated protein 2 OS=Mus musculus OX=10( | 6 | 2 | 2 | 2 | 428 | 46,5 | 8,87 | 2,67 | 2 | 1,242 | 1,56 | 1,183 | 0,948 | 0,791 | 1,35 | 0,92 | 0,854 | 1,26 | 0,26 | 2,0E-01 |
| O35435 | Dihydroorotate dehydrogenase (quinone), mitochondrial OS=Mus r | 26 | 8 | 8 | 8 | 395 | 42,7 | 9,55 | 15,11 | 8 | 1,192 | 1,168 | 1,034 | 1,075 | 0,922 | 1,13 | 0,997 | 1,082 | 1,08 | 0,08 | 2,0E-01 |
| Q80W65 | Proprotein convertase subtilisin/kexin type 9 OS=Mus musculus OX | 2 | 1 | 1 | 1 | 694 | 74,8 | 6,96 | 0 | 1 | 2,282 | 1,493 | 1,593 | 0,855 | 0,703 | 1,492 | 0,74 | 1,283 | 1,48 | 0,48 | 2,0E-01 |
| Q9R0N0 | Galactokinase OS=Mus musculus OX=10090 GN=Galk1 PE=1 SV=2 | 9 | 3 | 3 | 3 | 392 | 42,3 | 5,26 | 5,6 | 3 | 1,358 | 1,261 | 1,111 | 1,018 | 0,893 | 1,272 | 0,954 | 0,984 | 1,16 | 0,16 | 2,0E-01 |
| P51863 | V-type proton ATPase subunit d 1 OS=Mus musculus OX=10090 GN | 27 | 9 | 11 | 9 | 351 | 40,3 | 5 | 15,53 | 9 | 0,95 | 0,963 | 0,929 | 0,962 | 0,953 | 1,163 | 1,006 | 0,962 | 0,93 | -0,07 | 2,1E-01 |
| Q9EPE9 | Manganese-transporting ATPase 13A1 OS=Mus musculus OX=1009 | 4 | 4 | 4 | 4 | 1200 | 132,3 | 8,03 | 3,72 | 4 | 1,18 | 1,085 | 1,053 | 0,84 | 0,808 | 1,073 | 0,904 | 0,849 | 1,14 | 0,14 | 2,1E-01 |
| Q8BPU7 | Engulfment and cell motility protein 1 OS=Mus musculus OX=1009( | 3 | 2 | 2 | 1 | 727 | 83,9 | 6,28 | 4,75 | 2 | 0,723 | 0,949 | 0,376 | 1,11 | 1,036 | 0,882 | 0,963 | 1,342 | 0,75 | -0,25 | 2,1E-01 |
| Q69ZU6 | Thrombospondin type-1 domain-containing protein 7A OS=Mus mu | 4 | 5 | 5 | 5 | 1645 | 183,4 | 7,2 | 4,65 | 5 | 0,969 | 0,715 | 1,034 | 0,782 | 1,031 | 0,89 | 1,115 | 0,969 | 0,87 | -0,13 | 2,1E-01 |
| Q9JIY5 | Serine protease HTRA2, mitochondrial OS=Mus musculus OX=1009 | 9 | 3 | 4 | 3 | 458 | 49,3 | 9,6 | 6,49 | 3 | 1,045 | 1,138 | 1,225 | 1,044 | 0,902 | 0,996 | 1,152 | 1,021 | 1,09 | 0,09 | 2,1E-01 |
| Q91YQ5 | Dolichyl-diphosphooligosaccharide--protein glycosyltransferase sul | 11 | 7 | 8 | 7 | 608 | 68,5 | 6,46 | 19,82 | 7 | 1,031 | 0,974 | 1,044 | 1,073 | 0,991 | 1,02 | 0,953 | 1,013 | 1,04 | 0,04 | 2,1E-01 |
| Q9Z1G4 | V-type proton ATPase 116 kDa subunit a isoform 1 OS=Mus muscul | 26 | 24 | 57 | 24 | 839 | 96,4 | 6,76 | 144,92 | 24 | 0,989 | 1,006 | 1,052 | 1,058 | 0,949 | 0,963 | 1,034 | 1,011 | 1,04 | 0,04 | 2,1E-01 |
| Q3UXZ6 | Protein FAM81A OS=Mus musculus OX=10090 GN=Fam81a PE=1 S' | 16 | 6 | 7 | 6 | 364 | 41,7 | 8,75 | 18,62 | 6 | 0,784 | 1,023 | 0,865 | 0,999 | 1,035 | 0,993 | 1,078 | 0,93 | 0,91 | -0,09 | 2,1E-01 |
| O88531 | Palmitoyl-protein thioesterase 1 OS=Mus musculus OX=10090 GN= | 8 | 2 | 2 | 2 | 306 | 34,5 | 8 | 2,05 | 2 | 1,031 | 0,865 | 0,744 | 0,755 | 0,884 | 0,866 | 1,299 | 1,022 | 0,83 | -0,17 | 2,1E-01 |
| P58771 | Tropomyosin alpha-1 chain OS=Mus musculus OX=10090 GN=Tpm1 | 40 | 15 | 20 | 5 | 284 | 32,7 | 4,74 | 41,45 | 15 | 0,77 | 0,784 | 0,855 | 1,058 | 1,205 | 0,785 | 1,06 | 1,033 | 0,85 | -0,15 | 2,1E-01 |
| Q00612 | Glucose-6-phosphate 1-dehydrogenase X OS=Mus musculus OX=10 | 6 | 3 | 3 | 3 | 515 | 59,2 | 6,49 | 5,69 | 3 | 1,028 | 0,903 | 0,934 | 0,967 | 0,961 | 0,919 | 0,907 | 0,832 | 1,06 | 0,06 | 2,1E-01 |
| Q91VI7 | Ribonuclease inhibitor OS=Mus musculus OX=10090 GN=Rnh1 PE=1 | 14 | 6 | 6 | 6 | 456 | 49,8 | 4,78 | 14,01 | 6 | 1,229 | 1,089 | 1,081 | 0,954 | 0,941 | 1,088 | 0,941 | 1,011 | 1,09 | 0,09 | 2,1E-01 |
| Q4VAE3 | Transmembrane protein 65 OS=Mus musculus OX=10090 GN=Tmer | 20 | 6 | 8 | 6 | 234 | 24,9 | 7,78 | 15,41 | 6 | 0,937 | 1,026 | 1,025 | 0,958 | 0,971 | 1,05 | 1,022 | 1,116 | 0,95 | -0,05 | 2,1E-01 |
| Q9CWV1 | DNA helicase MCM8 OS=Mus musculus OX=10090 GN=Mcm8 PE=1 | 1 | 1 | 2 | 1 | 833 | 92,3 | 7,62 | 4,46 | 1 | 0,868 | 0,782 | 1,019 | 0,984 | 1,082 | 0,868 | 1,01 | 1,147 | 0,89 | -0,11 | 2,1E-01 |
| Q68EF4 | Metabotropic glutamate receptor 4 OS=Mus musculus OX=10090 G | 3 | 2 | 2 | 2 | 912 | 101,8 | 8,82 | 6,01 | 2 | 1,06 | 1,276 | 1,226 | 1,146 | 1,261 | 0,882 | 1,053 | 0,992 | 1,12 | 0,12 | 2,1E-01 |
| Q9JK48 | Endophilin-B1 OS=Mus musculus OX=10090 GN=Sh3glb1 PE=1 SV=1 | 22 | 8 | 9 | 8 | 365 | 40,8 | 6,04 | 13,82 | 8 | 1,26 | 1,059 | 0,987 | 0,964 | 0,974 | 0,941 | 1,012 | 0,956 | 1,10 | 0,10 | 2,1E-01 |
| P83882 | 60S ribosomal protein L36a OS=Mus musculus OX=10090 GN=Rpl3( | 15 | 2 | 2 | 2 | 106 | 12,4 | 10,58 | 2,61 | 2 | 1,005 | 0,821 | 0,889 | 0,809 | 0,916 | 1,072 | 0,847 | 1,191 | 0,88 | -0,12 | 2,1E-01 |
| Q8BH82 | N-acyl-phosphatidylethanolamine-hydrolyzing phospholipase D OS | 2 | 1 | 1 | 1 | 396 | 45,8 | 5,94 | 1,72 | 1 | 0,546 | 0,569 | 0,699 | 1,024 | 1,052 | 0,75 | 0,723 | 1,258 | 0,75 | -0,25 | 2,1E-01 |
| P70232 | Neural cell adhesion molecule L1-like protein OS=Mus musculus OX | 2 | 2 | 2 | 2 | 1209 | 135 | 5,57 | 0 | 2 | 0,625 | 0,841 | 1,172 | 0,986 | 1,223 | 0,954 | 1,055 | 1,111 | 0,83 | -0,17 | 2,1E-01 |
| Q9ESJ1 | CDK5 and ABL1 enzyme substrate 1 OS=Mus musculus OX=10090 G | 2 | 1 | 1 | 1 | 568 | 61,4 | 9,23 | 0 | 1 | 0,798 | 1,138 | 1,057 | 1,164 | 1,272 | 1,753 | 0,816 | 1,538 | 0,77 | -0,23 | 2,1E-01 |
| P97315 | Cysteine and glycine-rich protein 1 OS=Mus musculus OX=10090 GI | 41 | 8 | 12 | 8 | 193 | 20,6 | 8,57 | 28,16 | 8 | 1,098 | 1,058 | 1,008 | 1,016 | 1,04 | 1,182 | 1,056 | 1,117 | 0,95 | -0,05 | 2,1E-01 |
| Q99KV1 | DnaJ homolog subfamily B member 11 OS=Mus musculus OX=1009 | 13 | 3 | 3 | 3 | 358 | 40,5 | 6,32 | 11,37 | 3 | 1,221 | 1,159 | 1,21 | 1,139 | 0,98 | 1,279 | 1,021 | 1,06 | 1,09 | 0,09 | 2,1E-01 |
| F6SEU4 | Ras/Rap GTPase-activating protein SynGAP OS=Mus musculus OX=1 | 27 | 29 | 47 | 26 | 1340 | 148,1 | 8,98 | 90,28 | 29 | 1,195 | 1,133 | 1,021 | 1,024 | 0,994 | 1,041 | 1,055 | 1,033 | 1,06 | 0,06 | 2,1E-01 |
| Q9JLT4 | Thioredoxin reductase 2, mitochondrial OS=Mus musculus OX=100! | 8 | 4 | 6 | 4 | 524 | 56,6 | 8,46 | 14,19 | 4 | 1,002 | 1,217 | 1,061 | 1,107 | 1,104 | 0,891 | 1,01 | 1,028 | 1,09 | 0,09 | 2,1E-01 |
| Q8BG32 | 26S proteasome non-ATPase regulatory subunit 11 OS=Mus muscu | 32 | 12 | 15 | 12 | 422 | 47,4 | 6,48 | 31,88 | 12 | 1,099 | 1,128 | 1,09 | 1,037 | 1,023 | 1,069 | 1,03 | 1,092 | 1,03 | 0,03 | 2,1E-01 |
| Q8VI75 | Importin-4 OS=Mus musculus OX=10090 GN=Ipo4 PE=1 SV=1 | 1 | 1 | 1 | 1 | 1082 | 119,2 | 5,03 | 2,12 | 1 | 1,439 | 1,204 | 1,239 | 1,062 | 0,883 | 1,255 | 0,792 | 1,226 | 1,19 | 0,19 | 2,1E-01 |
| Q9CQ02 | COMM domain-containing protein 4 OS=Mus musculus OX=10090 ( | 6 | 1 | 1 | 1 | 199 | 21,8 | 6,67 | 2,89 | 1 | 0,559 | 1,293 | 0,876 | 1,255 | 1,196 | 1,169 | 1,315 | 1,29 | 0,80 | -0,20 | 2,1E-01 |
| Q4U2R1 | E3 ubiquitin-protein ligase HERC2 OS=Mus musculus OX=10090 GN | 1 | 2 | 2 | 2 | 4836 | 527,1 | 6,27 | 6,13 | 2 | 1,742 | 1,138 | 1,625 | 1,168 | 0,713 | 1,519 | 0,984 | 1,176 | 1,29 | 0,29 | 2,1E-01 |
| Q9CRB8 | Mitochondrial fission process protein 1 OS=Mus musculus OX=100! | 19 | 3 | 3 | 3 | 166 | 18,3 | 8,68 | 2,01 | 3 | 2,059 | 1,366 | 1,236 | 0,816 | 0,943 | 1,072 | 0,802 | 1,159 | 1,38 | 0,38 | 2,1E-01 |
| Q8VD65 | Phosphoinositide 3-kinase regulatory subunit 4 OS=Mus musculus ( | 5 | 7 | 8 | 7 | 1358 | 152,5 | 7,12 | 12,84 | 7 | 0,941 | 1,053 | 1,138 | 0,94 | 0,87 | 0,935 | 0,946 | 1,01 | 1,08 | 0,08 | 2,1E-01 |
| Q8BH79 | Anoctamin-10 OS=Mus musculus OX=10090 GN=Ano10 PE=1 SV=1 | 3 | 2 | 2 | 2 | 659 | 76,1 | 6,7 | 4,75 | 2 | 1,019 | 1,051 | 1,095 | 0,969 | 1,072 | 1,103 | 1,049 | 1,07 | 0,96 | -0,04 | 2,1E-01 |
| P13020 | Gelsolin OS=Mus musculus OX=10090 GN=Gsn PE=1 SV=3 | 10 | 8 | 9 | 8 | 780 | 85,9 | 6,18 | 17,38 | 8 | 1,021 | 0,993 | 1,233 | 1,094 | 1,056 | 1,017 | 0,985 | 0,965 | 1,08 | 0,08 | 2,1E-01 |
| O70274 | Protein tyrosine phosphatase type IVA 2 OS=Mus musculus OX=10( | 8 | 1 | 1 | 1 | 167 | 19,1 | 8,37 | 0 | 1 | 1,4 | 1,316 | 0,986 | 1,396 | 0,955 | 1,166 | 1,115 | 1,228 | 1,14 | 0,14 | 2,1E-01 |
| Q9DC61 | Mitochondrial-processing peptidase subunit alpha OS=Mus muscul | 18 | 9 | 12 | 9 | 524 | 58,2 | 6,83 | 25,82 | 9 | 1,239 | 1,109 | 1,153 | 0,965 | 1,029 | 1,108 | 0,949 | 1,014 | 1,09 | 0,09 | 2,1E-01 |
| Q99MR1 | GRB10-interacting GYF protein 1 OS=Mus musculus OX=10090 GN= | 1 | 1 | 1 | 1 | 1044 | 116,2 | 5,39 | 1,74 | 1 | 0,955 | 0,988 | 1,19 | 0,99 | 0,975 | 0,856 | 0,962 | 0,01 | 1,47 | 0,47 | 2,1E-01 |
| Q14C51 | Pentatricopeptide repeat domain-containing protein 3, mitochondr | 3 | 2 | 2 | 2 | 685 | 77,7 | 5,88 | 4,84 | 2 | 1,249 | 1,511 | 0,753 | 1,409 | 1,439 | 2,724 | 1,474 | 1,332 | 0,71 | -0,29 | 2,1E-01 |
| P70303 | CTP synthase 2 OS=Mus musculus OX=10090 GN=Ctps2 PE=1 SV=1 | 6 | 3 | 3 | 3 | 586 | 65,5 | 6,49 | 5 | 3 | 0,541 | 0,847 | 1,126 | 1,26 | 1,367 | 1,035 | 1,349 | 1,043 | 0,79 | -0,21 | 2,2E-01 |
| P68134 | Actin, alpha skeletal muscle OS=Mus musculus OX=10090 GN=Acta | 37 | 16 | 148 | 2 | 377 | 42 | 5,39 | 333,76 | 16 | 0,935 | 0,919 | 0,966 | 1,099 | 1,047 | 0,954 | 1,155 | 1,088 | 0,92 | -0,08 | 2,2E-01 |
| Q6PDG8 | Vacuolar fusion protein MON1 homolog A OS=Mus musculus OX=1( | 2 | 1 | 1 | 1 | 556 | 62,1 | 6,1 | 0 | 1 | 0,719 | 0,92 | 0,965 | 0,727 | 0,512 | 0,776 | 0,877 | 0,576 | 1,22 | 0,22 | 2,2E-01 |

| | | | | | | | | | | | | | | | | | | | | | |
|---|---|---|---|---|---|---|---|---|---|---|---|---|---|---|---|---|---|---|---|---|---|
| Q922F4 | Tubulin beta-6 chain OS=Mus musculus OX=10090 GN=Tubb6 PE=1 | 34 | 14 | 64 | 2 | 447 | 50,1 | 4,89 | 140,84 | 14 | 0,779 | 1,243 | 0,907 | 0,864 | 0,639 | 0,868 | 0,892 | 0,744 | 1,21 | 0,21 | 2,2E-01 |
| Q7TPH6 | E3 ubiquitin-protein ligase MYCBP2 OS=Mus musculus OX=10090 G | 1 | 6 | 7 | 6 | 4749 | 520,9 | 7,09 | 5,13 | 6 | 1,415 | 1,089 | 1,224 | 1,181 | 1,049 | 1,267 | 0,986 | 1,102 | 1,11 | 0,11 | 2,2E-01 |
| Q9Z1W9 | STE20/SPS1-related proline-alanine-rich protein kinase OS=Mus mu | 4 | 2 | 2 | 2 | 556 | 60,3 | 6,29 | 5,34 | 2 | 0,721 | 0,677 | 0,964 | 1,015 | 1,076 | 0,841 | 1,059 | 0,958 | 0,86 | -0,14 | 2,2E-01 |
| P48774 | Glutathione S-transferase Mu 5 OS=Mus musculus OX=10090 GN=( | 33 | 9 | 13 | 8 | 224 | 26,6 | 7,21 | 22,75 | 9 | 1,148 | 1,231 | 1,053 | 1,04 | 0,972 | 1,042 | 1,005 | 1,137 | 1,08 | 0,08 | 2,2E-01 |
| Q9R1P3 | Proteasome subunit beta type-2 OS=Mus musculus OX=10090 GN= | 27 | 8 | 13 | 8 | 201 | 22,9 | 7,02 | 19,77 | 8 | 1,057 | 1,057 | 1,106 | 0,994 | 1,07 | 1,126 | 1,048 | 1,16 | 0,96 | -0,04 | 2,2E-01 |
| Q3UZ39-2 | Isoform 2 of Leucine-rich repeat flightless-interacting protein 1 OS= | 4 | 3 | 3 | 3 | 628 | 71,3 | 6,04 | 1,72 | 3 | 1,013 | 1,009 | 0,623 | 0,909 | 1,045 | 0,954 | 1,197 | 0,955 | 0,86 | -0,14 | 2,2E-01 |
| Q9DB05 | Alpha-soluble NSF attachment protein OS=Mus musculus OX=1009 | 61 | 15 | 28 | 14 | 295 | 33,2 | 5,45 | 65,7 | 15 | 1,005 | 1,01 | 1,096 | 1,062 | 1,048 | 1,131 | 1,068 | 1,081 | 0,96 | -0,04 | 2,2E-01 |
| Q8K2M0 | 39S ribosomal protein L38, mitochondrial OS=Mus musculus OX=1( | 3 | 1 | 2 | 1 | 380 | 45 | 8,1 | 0 | 1 | 0,908 | 1,138 | 0,937 | 1,009 | 1,052 | 1,36 | 1,134 | 0,981 | 0,88 | -0,12 | 2,2E-01 |
| Q61301 | Catenin alpha-2 OS=Mus musculus OX=10090 GN=Ctnna2 PE=1 SV= | 25 | 21 | 28 | 16 | 953 | 105,2 | 5,71 | 72,27 | 21 | 0,827 | 0,838 | 0,932 | 1,018 | 1,001 | 0,875 | 1,054 | 1,011 | 0,92 | -0,08 | 2,2E-01 |
| Q8BR63 | Protein FAM177A1 OS=Mus musculus OX=10090 GN=Fam177a1 PE | 15 | 3 | 3 | 3 | 207 | 23,6 | 4,59 | 6,03 | 3 | 1,311 | 1,179 | 0,816 | 1,024 | 0,821 | 1,124 | 0,818 | 0,852 | 1,20 | 0,20 | 2,2E-01 |
| Q8CCB4 | Vacuolar protein sorting-associated protein 53 homolog OS=Mus m | 8 | 7 | 9 | 7 | 832 | 94,4 | 6,61 | 13,67 | 7 | 1,01 | 1,06 | 1,147 | 1,045 | 1,094 | 1,136 | 1,214 | 1,061 | 0,95 | -0,05 | 2,2E-01 |
| Q9EPR5 | VPS10 domain-containing receptor SorCS2 OS=Mus musculus OX=1 | 1 | 1 | 1 | 1 | 1159 | 128,8 | 7,34 | 2,06 | 1 | 2,546 | 2,024 | 1,864 | 1,14 | 0,803 | 1,774 | 1,409 | 1,605 | 1,35 | 0,35 | 2,2E-01 |
| P24668 | Cation-dependent mannose-6-phosphate receptor OS=Mus muscul | 4 | 2 | 2 | 2 | 278 | 31,2 | 5,39 | 2,27 | 2 | 0,964 | 1,035 | 0,844 | 0,877 | 1,12 | 1,092 | 0,983 | 0,896 | 0,91 | -0,09 | 2,2E-01 |
| Q9WU60 | Attractin OS=Mus musculus OX=10090 GN=Atrn PE=1 SV=3 | 2 | 3 | 3 | 3 | 1428 | 158 | 7,27 | 4,57 | 3 | 1,194 | 1,118 | 1,112 | 0,971 | 1,085 | 1,082 | 0,979 | 0,848 | 1,10 | 0,10 | 2,2E-01 |
| Q80WC1 | Ubinuclein-2 OS=Mus musculus OX=10090 GN=Ubn2 PE=1 SV=2 | 1 | 2 | 4 | 2 | 1314 | 141,7 | 9,17 | 5,1 | 2 | 0,692 | 0,774 | 1,007 | 0,996 | 0,837 | 1,111 | 1,005 | 1,064 | 0,86 | -0,14 | 2,2E-01 |
| Q6GQS1 | Calcium-binding mitochondrial carrier protein SCaMC-3 OS=Mus m | 18 | 8 | 8 | 7 | 467 | 52,5 | 7,59 | 18,24 | 8 | 1,202 | 1,07 | 1,098 | 1,086 | 0,975 | 1,169 | 0,954 | 1,044 | 1,08 | 0,08 | 2,2E-01 |
| Q9CZ42 | ATP-dependent (S)-NAD(P)H-hydrate dehydratase OS=Mus muscul | 13 | 4 | 5 | 4 | 343 | 36,7 | 7,77 | 4,37 | 4 | 1,321 | 1,436 | 1,347 | 1,08 | 1,064 | 1,174 | 1,234 | 1,237 | 1,10 | 0,10 | 2,2E-01 |
| Q78IK4 | MICOS complex subunit Mic27 OS=Mus musculus OX=10090 GN=A | 14 | 3 | 3 | 3 | 265 | 29,2 | 9,31 | 5,84 | 3 | 1,143 | 1,367 | 1,038 | 0,951 | 1,105 | 1,428 | 1,206 | 1,427 | 0,87 | -0,13 | 2,2E-01 |
| P48771 | Cytochrome c oxidase subunit 7A2, mitochondrial OS=Mus muscul | 28 | 2 | 7 | 2 | 83 | 9,3 | 10,27 | 19,37 | 2 | 0,664 | 1,015 | 0,789 | 0,911 | 1,208 | 0,795 | 1,074 | 0,944 | 0,84 | -0,16 | 2,2E-01 |
| P28474 | Alcohol dehydrogenase class-3 OS=Mus musculus OX=10090 GN=A | 11 | 3 | 3 | 3 | 374 | 39,5 | 7,25 | 6,56 | 3 | 1,363 | 0,953 | 1,205 | 1,165 | 0,941 | 1,134 | 1,059 | 1,038 | 1,12 | 0,12 | 2,2E-01 |
| Q9CXW3 | Calcyclin-binding protein OS=Mus musculus OX=10090 GN=Cacybp | 11 | 2 | 2 | 2 | 229 | 26,5 | 7,87 | 5 | 2 | 0,963 | 0,643 | 0,934 | 0,773 | 0,831 | 0,899 | 1,099 | 1,007 | 0,86 | -0,14 | 2,2E-01 |
| Q9WVK4 | EH domain-containing protein 1 OS=Mus musculus OX=10090 GN=I | 18 | 9 | 9 | 7 | 534 | 60,6 | 6,83 | 21,51 | 9 | 1,006 | 0,972 | 1,016 | 0,98 | 0,971 | 1,12 | 1,101 | 0,995 | 0,95 | -0,05 | 2,2E-01 |
| Q8C0I1 | Alkyldihydroxyacetonephosphate synthase, peroxisomal OS=Mus n | 6 | 3 | 3 | 3 | 645 | 71,6 | 7,5 | 4,32 | 3 | 1,983 | 1,981 | 1,165 | 1,302 | 1,066 | 1,454 | 1,398 | 1,223 | 1,25 | 0,25 | 2,2E-01 |
| Q922E4 | Ethanolamine-phosphate cytidylyltransferase OS=Mus musculus O) | 13 | 5 | 6 | 5 | 404 | 45,2 | 6,58 | 6,09 | 5 | 1,008 | 0,878 | 1,111 | 1,075 | 1,136 | 0,973 | 1,182 | 1,167 | 0,91 | -0,09 | 2,2E-01 |
| Q8BHJ5 | F-box-like/WD repeat-containing protein TBL1XR1 OS=Mus muscul | 3 | 1 | 1 | 1 | 514 | 55,6 | 5,63 | 2,37 | 1 | 1,318 | 1,239 | 1,075 | 1,12 | 1,032 | 1,109 | 0,94 | 1,223 | 1,10 | 0,10 | 2,2E-01 |
| P62137 | Serine/threonine-protein phosphatase PP1-alpha catalytic subunit | 24 | 7 | 11 | 3 | 330 | 37,5 | 6,33 | 30,26 | 7 | 0,918 | 0,937 | 1,001 | 0,997 | 1,022 | 0,949 | 0,992 | 1,079 | 0,95 | -0,05 | 2,2E-01 |
| Q9R0N7-4 | Isoform 4 of Synaptotagmin-7 OS=Mus musculus OX=10090 GN=Sy | 21 | 8 | 10 | 8 | 567 | 63,1 | 9,44 | 22 | 8 | 0,797 | 0,898 | 0,932 | 1,099 | 0,973 | 1,039 | 1,108 | 0,988 | 0,91 | -0,09 | 2,2E-01 |
| O55229 | Choline/ethanolamine kinase OS=Mus musculus OX=10090 GN=Chl | 9 | 3 | 4 | 3 | 394 | 45,1 | 5,41 | 9,5 | 3 | 1,552 | 0,974 | 1,216 | 1,257 | 0,946 | 1,186 | 0,827 | 1,213 | 1,20 | 0,20 | 2,2E-01 |
| Q8BTZ7 | Mannose-1-phosphate guanyltransferase beta OS=Mus musculus C | 6 | 2 | 2 | 2 | 360 | 39,9 | 6,74 | 5,99 | 2 | 0,91 | 0,997 | 1,204 | 0,973 | 0,96 | 1,311 | 1,436 | 1,068 | 0,86 | -0,14 | 2,2E-01 |
| B1AR13 | CDGSH iron-sulfur domain-containing protein 3, mitochondrial OS= | 22 | 3 | 3 | 3 | 137 | 15,7 | 9,79 | 7,81 | 3 | 1,095 | 1,008 | 1,016 | 0,923 | 0,999 | 0,974 | 0,878 | 0,952 | 1,06 | 0,06 | 2,2E-01 |
| P97820 | Mitogen-activated protein kinase kinase kinase kinase 4 OS=Mus m | 6 | 7 | 7 | 3 | 1233 | 140,5 | 7,47 | 13,14 | 7 | 1,315 | 1,053 | 0,665 | 0,79 | 0,8 | 0,624 | 0,688 | 0,868 | 1,28 | 0,28 | 2,2E-01 |
| Q9QXW9 | Large neutral amino acids transporter small subunit 2 OS=Mus mus | 2 | 1 | 1 | 1 | 531 | 57,8 | 6,67 | 2,21 | 1 | 0,832 | 1,376 | 0,905 | 0,779 | 0,941 | 1,13 | 1,384 | 1,389 | 0,80 | -0,20 | 2,2E-01 |
| Q8R092 | Uncharacterized protein C1orf43 homolog OS=Mus musculus OX=1 | 16 | 3 | 3 | 3 | 253 | 28,7 | 9,36 | 5,44 | 3 | 1,395 | 1,243 | 1,219 | 1,16 | 0,827 | 1,245 | 1,032 | 1,279 | 1,14 | 0,14 | 2,2E-01 |
| Q99KC8 | von Willebrand factor A domain-containing protein 5A OS=Mus mu | 6 | 4 | 4 | 4 | 793 | 87,1 | 6,58 | 6,19 | 4 | 0,01 | 1,314 | 0,683 | 1,035 | 1,074 | 1,211 | 1,021 | 1,311 | 0,66 | -0,34 | 2,2E-01 |
| Q8VEJ9 | Vacuolar protein sorting-associated protein 4A OS=Mus musculus ( | 12 | 4 | 5 | 3 | 437 | 48,9 | 7,8 | 7,58 | 4 | 1,176 | 1,167 | 1,084 | 1,09 | 0,945 | 1,175 | 1,066 | 1,041 | 1,07 | 0,07 | 2,2E-01 |
| Q9CXJ1 | Probable glutamate--tRNA ligase, mitochondrial OS=Mus musculus | 2 | 1 | 1 | 1 | 523 | 58,3 | 8,75 | 2,31 | 1 | 1,375 | 1,439 | 1,221 | 1,17 | 1,154 | 1,02 | 1,256 | 1,295 | 1,10 | 0,10 | 2,2E-01 |
| Q9D1Q4 | Dolichol-phosphate mannosyltransferase subunit 3 OS=Mus muscu | 13 | 1 | 1 | 1 | 92 | 10,1 | 7,08 | 3,38 | 1 | 2,02 | 1,27 | 1,513 | 1,257 | 1,322 | 1,323 | 1,382 | 0,856 | 1,24 | 0,24 | 2,2E-01 |
| Q99JW2 | Aminoacylase-1 OS=Mus musculus OX=10090 GN=Acy1 PE=1 SV=1 | 4 | 2 | 3 | 2 | 408 | 45,8 | 6,32 | 2,27 | 2 | 0,783 | 1,062 | 1,038 | 1,173 | 0,978 | 0,741 | 0,878 | 0,931 | 1,15 | 0,15 | 2,2E-01 |
| F6W8I0 | YjeF N-terminal domain-containing protein 3 OS=Mus musculus OX | 18 | 4 | 4 | 4 | 251 | 27 | 7,4 | 5,2 | 4 | 1,185 | 1,481 | 1,035 | 1,2 | 1,101 | 1,053 | 1,124 | 1,11 | 1,12 | 0,12 | 2,2E-01 |
| Q9QZQ1 | Afadin OS=Mus musculus OX=10090 GN=Afdn PE=1 SV=3 | 11 | 18 | 21 | 18 | 1820 | 206,4 | 6,32 | 39,66 | 18 | 1,104 | 1,064 | 0,997 | 1,076 | 0,969 | 0,989 | 1,035 | 1,072 | 1,04 | 0,04 | 2,2E-01 |
| Q0VGU4 | Neurosecretory protein VGF OS=Mus musculus OX=10090 GN=Vgf | 12 | 6 | 13 | 6 | 617 | 68,2 | 4,7 | 34,43 | 6 | 1,231 | 1,127 | 1,027 | 1,024 | 0,939 | 1,09 | 1,032 | 1,032 | 1,08 | 0,08 | 2,2E-01 |
| P18760 | Cofilin-1 OS=Mus musculus OX=10090 GN=Cfl1 PE=1 SV=3 | 72 | 13 | 19 | 11 | 166 | 18,5 | 8,09 | 36,53 | 13 | 1,333 | 1,05 | 1,321 | 1,02 | 0,868 | 1,207 | 1,04 | 1,016 | 1,14 | 0,14 | 2,2E-01 |
| Q8K0S0 | Phytanoyl-CoA hydroxylase-interacting protein OS=Mus musculus ( | 20 | 6 | 12 | 6 | 330 | 37,5 | 7,01 | 33,71 | 6 | 0,932 | 0,917 | 0,943 | 0,992 | 1,102 | 0,971 | 0,984 | 0,939 | 0,95 | -0,05 | 2,2E-01 |
| Q8K1N1 | Calcium-independent phospholipase A2-gamma OS=Mus musculus | 9 | 5 | 5 | 5 | 776 | 87,3 | 9,23 | 7,53 | 5 | 1,156 | 1,145 | 1,007 | 1,098 | 0,973 | 1,092 | 1,04 | 1,07 | 1,06 | 0,06 | 2,2E-01 |
| Q9CQ80 | Vacuolar protein-sorting-associated protein 25 OS=Mus musculus ( | 11 | 3 | 4 | 3 | 176 | 20,7 | 6,34 | 14,05 | 3 | 1,107 | 1,159 | 1,062 | 1,053 | 0,977 | 1,116 | 0,939 | 1,085 | 1,06 | 0,06 | 2,3E-01 |
| E9Q6P5 | Tetratricopeptide repeat protein 7B OS=Mus musculus OX=10090 ( | 14 | 11 | 13 | 11 | 843 | 94,1 | 6,9 | 35,53 | 11 | 0,974 | 0,942 | 0,995 | 0,967 | 0,945 | 1,063 | 1,004 | 1,009 | 0,96 | -0,04 | 2,3E-01 |
| Q9CWF2 | Tubulin beta-2B chain OS=Mus musculus OX=10090 GN=Tubb2b PE | 62 | 25 | 161 | 1 | 445 | 49,9 | 4,89 | 336,52 | 25 | 1,689 | 1,706 | 1,129 | 1,189 | 0,747 | 1,646 | 0,782 | 1,131 | 1,33 | 0,33 | 2,3E-01 |
| Q8BLK9 | Ribosomal protein S6 kinase delta-1 OS=Mus musculus OX=10090 ( | 2 | 2 | 2 | 2 | 1056 | 115,6 | 4,89 | 2,1 | 2 | 0,887 | 0,52 | 0,96 | 1,01 | 1,099 | 0,911 | 1,222 | 0,884 | 0,82 | -0,18 | 2,3E-01 |
| Q78ZA7 | Nucleosome assembly protein 1-like 4 OS=Mus musculus OX=1009( | 15 | 4 | 6 | 3 | 375 | 42,7 | 4,67 | 17,51 | 4 | 1,318 | 1,185 | 1,178 | 0,919 | 0,937 | 1,085 | 0,923 | 1,126 | 1,13 | 0,13 | 2,3E-01 |
| Q91Z38 | Tetratricopeptide repeat protein 1 OS=Mus musculus OX=10090 GI | 14 | 3 | 4 | 3 | 292 | 33,2 | 5,01 | 6,03 | 3 | 1,308 | 1,343 | 1,234 | 1,135 | 1,019 | 1,218 | 1,147 | 1,259 | 1,08 | 0,08 | 2,3E-01 |
| P61226 | Ras-related protein Rap-2b OS=Mus musculus OX=10090 GN=Rap2 | 32 | 6 | 11 | 4 | 183 | 20,5 | 4,81 | 10,85 | 6 | 0,469 | 0,643 | 0,662 | 0,822 | 1,026 | 0,451 | 1,076 | 0,904 | 0,75 | -0,25 | 2,3E-01 |
| Q9WV98 | Mitochondrial import inner membrane translocase subunit Tim9 O: | 53 | 4 | 8 | 4 | 89 | 10,3 | 7,21 | 21,53 | 4 | 0,689 | 0,926 | 0,805 | 0,983 | 1,007 | 0,906 | 0,974 | 0,897 | 0,90 | -0,10 | 2,3E-01 |
| P11798 | Calcium/calmodulin-dependent protein kinase type II subunit alpha | 26 | 14 | 41 | 11 | 478 | 54,1 | 7,08 | 94,59 | 14 | 1,032 | 1,029 | 0,925 | 0,985 | 1,041 | 1,098 | 1,004 | 1,007 | 0,96 | -0,04 | 2,3E-01 |
| Q3UMB5 | Guanine nucleotide exchange protein SMCR8 OS=Mus musculus O) | 1 | 1 | 1 | 1 | 935 | 104,9 | 5,4 | 2,09 | 1 | 0,679 | 0,741 | 0,49 | 0,642 | 0,784 | 0,655 | 0,703 | 0,732 | 0,89 | -0,11 | 2,3E-01 |
| Q9CXZ1 | NADH dehydrogenase [ubiquinone] iron-sulfur protein 4, mitochon | 37 | 6 | 7 | 6 | 175 | 19,8 | 9,99 | 12,6 | 6 | 0,876 | 0,932 | 1,112 | 0,821 | 0,972 | 1,156 | 0,988 | 1,032 | 0,90 | -0,10 | 2,3E-01 |
| Q8VI56 | Low-density lipoprotein receptor-related protein 4 OS=Mus muscul | 1 | 1 | 1 | 1 | 1905 | 211,8 | 5,39 | 0 | 1 | 0,912 | 0,918 | 1,48 | 1,094 | 1,021 | 1,715 | 1,521 | 1,238 | 0,80 | -0,20 | 2,3E-01 |
| P62259 | 14-3-3 protein epsilon OS=Mus musculus OX=10090 GN=Ywhae PE | 64 | 20 | 71 | 17 | 255 | 29,2 | 4,74 | 151,72 | 20 | 0,939 | 0,953 | 0,983 | 1,037 | 0,925 | 0,924 | 0,954 | 0,975 | 1,04 | 0,04 | 2,3E-01 |
| P17809 | Solute carrier family 2, facilitated glucose transporter member 1 OS | 9 | 4 | 4 | 4 | 492 | 53,9 | 8,87 | 8,95 | 4 | 0,878 | 0,991 | 0,925 | 0,987 | 0,921 | 0,858 | 0,896 | 0,935 | 1,05 | 0,05 | 2,3E-01 |
| Q9JI19 | Acidic fibroblast growth factor intracellular-binding protein OS=Mu | 8 | 3 | 3 | 3 | 357 | 41,2 | 6,76 | 3,8 | 3 | 2,226 | 1,29 | 1,954 | 0,994 | 0,75 | 1,566 | 0,997 | 1,339 | 1,39 | 0,39 | 2,3E-01 |
| P20352 | Tissue factor OS=Mus musculus OX=10090 GN=F3 PE=1 SV=2 | 6 | 1 | 1 | 1 | 294 | 32,9 | 9,32 | 3,41 | 1 | 1,495 | 1,056 | 1,478 | 1,279 | 1,262 | 0,986 | 1,306 | 1,067 | 1,15 | 0,15 | 2,3E-01 |
| Q9QVP9 | Protein-tyrosine kinase 2-beta OS=Mus musculus OX=10090 GN=Pt | 14 | 12 | 13 | 12 | 1009 | 115,7 | 6,35 | 19,36 | 12 | 1,191 | 1,509 | 1,038 | 1,073 | 0,933 | 1,126 | 1,042 | 1,093 | 1,15 | 0,15 | 2,3E-01 |
| Q921J2 | GTP-binding protein Rheb OS=Mus musculus OX=10090 GN=Rheb F | 14 | 3 | 3 | 3 | 184 | 20,4 | 5,92 | 9,41 | 3 | 0,738 | 0,88 | 0,878 | 1,019 | 1,024 | 0,856 | 1,022 | 0,986 | 0,90 | -0,10 | 2,3E-01 |

| | | | | | | | | | | | | | | | | | | | | | |
|---|---|---|---|---|---|---|---|---|---|---|---|---|---|---|---|---|---|---|---|---|---|
| P63005 | Platelet-activating factor acetylhydrolase IB subunit alpha OS=Mus | 40 | 12 | 20 | 12 | 410 | 46,6 | 7,37 | 36,48 | 12 | 1,106 | 1,092 | 1,064 | 1,055 | 0,985 | 1,122 | 0,996 | 1,036 | 1,04 | 0,04 | 2,3E-01 |
| Q9CR95 | Adaptin ear-binding coat-associated protein 1 OS=Mus musculus O | 15 | 5 | 6 | 5 | 275 | 29,6 | 6,38 | 5,13 | 5 | 0,747 | 0,789 | 0,838 | 1,027 | 1,035 | 0,777 | 1,037 | 1,028 | 0,88 | -0,12 | 2,3E-01 |
| Q9JM14 | 5'(3')-deoxyribonucleotidase, cytosolic type OS=Mus musculus OX= | 16 | 3 | 4 | 3 | 200 | 23,1 | 5,49 | 6,15 | 3 | 1,292 | 1,085 | 1,086 | 1,283 | 0,955 | 1,195 | 1 | 1,155 | 1,10 | 0,10 | 2,3E-01 |
| O54833 | Casein kinase II subunit alpha' OS=Mus musculus OX=10090 GN=Cs | 24 | 7 | 7 | 7 | 350 | 41,2 | 8,56 | 13,52 | 7 | 1,095 | 1,311 | 1,064 | 1,044 | 0,851 | 1,123 | 1,006 | 1,076 | 1,11 | 0,11 | 2,3E-01 |
| P80315 | T-complex protein 1 subunit delta OS=Mus musculus OX=10090 GN | 24 | 12 | 21 | 12 | 539 | 58 | 8,02 | 47,9 | 12 | 0,942 | 0,865 | 1,012 | 0,985 | 0,999 | 0,973 | 1,027 | 0,987 | 0,95 | -0,05 | 2,3E-01 |
| Q9ERZ3 | Muscarinic acetylcholine receptor M3 OS=Mus musculus OX=1009( | 2 | 1 | 1 | 1 | 589 | 66,2 | 9,07 | 0 | 1 | 0,786 | 1,406 | 0,895 | 1,569 | 1,467 | 1,975 | 1,924 | 0,935 | 0,74 | -0,26 | 2,3E-01 |
| Q9JI39 | ATP-binding cassette sub-family B member 10, mitochondrial OS=M | 7 | 4 | 4 | 4 | 715 | 77,1 | 9,6 | 1,76 | 4 | 1,044 | 1,166 | 0,814 | 0,972 | 0,762 | 0,867 | 0,655 | 1,085 | 1,19 | 0,19 | 2,3E-01 |
| Q921S7 | 39S ribosomal protein L37, mitochondrial OS=Mus musculus OX=1( | 17 | 6 | 7 | 6 | 423 | 48,3 | 8,84 | 18,57 | 6 | 1,33 | 1,33 | 1,188 | 1,006 | 0,921 | 1,32 | 0,954 | 1,024 | 1,15 | 0,15 | 2,3E-01 |
| Q8BIJ6 | Isoleucine--tRNA ligase, mitochondrial OS=Mus musculus OX=1009( | 17 | 14 | 20 | 14 | 1012 | 112,7 | 6,81 | 38,09 | 14 | 1,014 | 1,032 | 1,009 | 0,996 | 0,991 | 1,074 | 1,023 | 1,099 | 0,97 | -0,03 | 2,3E-01 |
| Q8BSF4 | Phosphatidylserine decarboxylase proenzyme, mitochondrial OS=M | 2 | 1 | 1 | 1 | 406 | 45,9 | 9,57 | 1,69 | 1 | 0,813 | 0,993 | 1,36 | 1,496 | 1,092 | 1,44 | 1,462 | 1,837 | 0,80 | -0,20 | 2,3E-01 |
| P26043 | Radixin OS=Mus musculus OX=10090 GN=Rdx PE=1 SV=3 | 18 | 13 | 15 | 5 | 583 | 68,5 | 6,2 | 22,76 | 13 | 1,36 | 1,005 | 1,055 | 0,997 | 0,988 | 1,019 | 0,97 | 0,978 | 1,12 | 0,12 | 2,3E-01 |
| P56942 | Pro-MCH OS=Mus musculus OX=10090 GN=Pmch PE=1 SV=2 | 7 | 1 | 1 | 1 | 165 | 18,5 | 7,14 | 2,9 | 1 | 0,718 | 1,021 | 1,028 | 1,097 | 1,024 | 1,421 | 1,399 | 0,874 | 0,82 | -0,18 | 2,3E-01 |
| P08553 | Neurofilament medium polypeptide OS=Mus musculus OX=10090 ( | 34 | 30 | 41 | 26 | 848 | 95,9 | 4,77 | 92,34 | 30 | 1,112 | 1,083 | 1,003 | 0,974 | 1,023 | 1,012 | 0,997 | 0,943 | 1,05 | 0,05 | 2,3E-01 |
| Q64516-1 | Isoform 1 of Glycerol kinase OS=Mus musculus OX=10090 GN=Gk | 31 | 14 | 19 | 14 | 524 | 57,4 | 5,64 | 40,79 | 14 | 1,011 | 0,882 | 0,973 | 0,929 | 1,01 | 1,021 | 0,946 | 0,99 | 0,96 | -0,04 | 2,3E-01 |
| P62843 | 40S ribosomal protein S15 OS=Mus musculus OX=10090 GN=Rps15 | 8 | 1 | 1 | 1 | 145 | 17 | 10,39 | 2,81 | 1 | 1,382 | 0,929 | 1,245 | 1,013 | 0,933 | 0,914 | 0,805 | 1,204 | 1,18 | 0,18 | 2,3E-01 |
| P10493 | Nidogen-1 OS=Mus musculus OX=10090 GN=Nid1 PE=1 SV=2 | 2 | 1 | 1 | 1 | 1245 | 136,5 | 5,44 | 2,86 | 1 | 2,266 | 1,042 | 1,814 | 1,394 | 0,845 | 1,921 | 0,887 | 0,886 | 1,44 | 0,44 | 2,3E-01 |
| Q0PHV7 | Dapper homolog 3 OS=Mus musculus OX=10090 GN=Dact3 PE=1 S\ | 2 | 1 | 2 | 1 | 610 | 63,2 | 10,42 | 0 | 1 | 0,955 | 1,099 | 0,592 | 1,165 | 0,447 | 0,842 | 0,652 | 0,964 | 1,31 | 0,31 | 2,3E-01 |
| Q8CIW6 | Solute carrier family 26 member 6 OS=Mus musculus OX=10090 GN | 4 | 2 | 3 | 2 | 758 | 82,8 | 8,65 | 4,5 | 2 | 0,442 | 0,855 | 0,543 | 0,865 | 0,844 | 0,673 | 1,108 | 0,827 | 0,78 | -0,22 | 2,3E-01 |
| Q3THE2 | Myosin regulatory light chain 12B OS=Mus musculus OX=10090 GN | 41 | 8 | 14 | 8 | 172 | 19,8 | 4,84 | 20,03 | 8 | 0,825 | 0,877 | 0,906 | 0,964 | 1,029 | 0,834 | 1,048 | 0,96 | 0,92 | -0,08 | 2,3E-01 |
| Q8CJ19 | [F-actin]-monooxygenase MICAL3 OS=Mus musculus OX=10090 GN | 13 | 20 | 22 | 19 | 1993 | 223,6 | 5,47 | 29,54 | 20 | 1,042 | 1,118 | 0,937 | 0,99 | 1,054 | 1,057 | 1,099 | 1,09 | 0,95 | -0,05 | 2,3E-01 |
| Q9D7X1 | BTB/POZ domain-containing protein KCTD4 OS=Mus musculus OX= | 9 | 2 | 3 | 2 | 259 | 30 | 7,06 | 5,46 | 2 | 1,989 | 1,384 | 1,559 | 1,153 | 0,99 | 1,541 | 1,168 | 1,27 | 1,22 | 0,22 | 2,3E-01 |
| Q69ZX8 | Actin-binding LIM protein 3 OS=Mus musculus OX=10090 GN=Ablin | 6 | 3 | 3 | 3 | 682 | 77,6 | 8,54 | 0 | 3 | 1,213 | 1,505 | 1,282 | 1,165 | 1,263 | 1,25 | 1,155 | 0,902 | 1,13 | 0,13 | 2,3E-01 |
| Q8C729-2 | Isoform 2 of Protein FAM126B OS=Mus musculus OX=10090 GN=Fa | 8 | 5 | 5 | 5 | 586 | 64,4 | 6,67 | 9,23 | 5 | 0,882 | 0,967 | 1,061 | 1,131 | 1,154 | 1,023 | 1,095 | 1,089 | 0,93 | -0,07 | 2,3E-01 |
| P53798 | Squalene synthase OS=Mus musculus OX=10090 GN=Fdft1 PE=1 SV | 2 | 1 | 1 | 1 | 416 | 48,1 | 6,32 | 0 | 1 | 0,983 | 0,652 | 0,785 | 0,922 | 1,031 | 0,779 | 0,944 | 1,169 | 0,85 | -0,15 | 2,4E-01 |
| P58742 | Aladin OS=Mus musculus OX=10090 GN=Aaas PE=1 SV=1 | 3 | 1 | 1 | 1 | 546 | 59,4 | 6,87 | 0 | 1 | 2,725 | 1,597 | 1,687 | 0,921 | 0,947 | 1,497 | 1,14 | 1,288 | 1,42 | 0,42 | 2,4E-01 |
| P63328 | Serine/threonine-protein phosphatase 2B catalytic subunit alpha is | 30 | 15 | 49 | 9 | 521 | 58,6 | 5,86 | 67,91 | 15 | 1,054 | 1,048 | 1,013 | 1,025 | 0,99 | 0,927 | 1,045 | 1,03 | 1,04 | 0,04 | 2,4E-01 |
| Q8R010 | Aminoacyl tRNA synthase complex-interacting multifunctional prot | 5 | 2 | 2 | 2 | 320 | 35,4 | 7,83 | 4,4 | 2 | 1,132 | 0,969 | 1,22 | 1,063 | 1,043 | 1,058 | 1,022 | 0,958 | 1,07 | 0,07 | 2,4E-01 |
| Q91VZ6 | Stromal membrane-associated protein 1 OS=Mus musculus OX=10( | 11 | 5 | 6 | 5 | 440 | 47,6 | 8,51 | 17,41 | 5 | 0,759 | 0,774 | 0,829 | 1,12 | 1,024 | 0,913 | 1,006 | 1,003 | 0,88 | -0,12 | 2,4E-01 |
| Q9WVJ3 | Carboxypeptidase Q OS=Mus musculus OX=10090 GN=Cpq PE=1 S\ | 4 | 2 | 4 | 2 | 470 | 51,8 | 6,21 | 5,13 | 2 | 1,69 | 1,355 | 1,153 | 0,984 | 0,666 | 1,325 | 0,925 | 1,166 | 1,27 | 0,27 | 2,4E-01 |
| Q4V9Z5 | Seizure 6-like protein 2 OS=Mus musculus OX=10090 GN=Sez6l2 PE | 4 | 3 | 4 | 3 | 910 | 97,4 | 4,87 | 6,45 | 3 | 0,663 | 0,841 | 0,798 | 0,84 | 0,938 | 0,718 | 0,871 | 1,01 | 0,89 | -0,11 | 2,4E-01 |
| Q64331 | Unconventional myosin-VI OS=Mus musculus OX=10090 GN=Myo6 | 5 | 6 | 9 | 6 | 1265 | 146,3 | 8,85 | 19,16 | 6 | 0,968 | 0,876 | 1,082 | 1,04 | 1,09 | 1,107 | 1,105 | 0,963 | 0,93 | -0,07 | 2,4E-01 |
| Q80U95 | Ubiquitin-protein ligase E3C OS=Mus musculus OX=10090 GN=Ube | 5 | 5 | 6 | 5 | 1083 | 123,9 | 6,39 | 5,8 | 5 | 0,964 | 0,947 | 0,854 | 1,153 | 1,034 | 1,192 | 0,973 | 1,139 | 0,90 | -0,10 | 2,4E-01 |
| Q60865 | Caprin-1 OS=Mus musculus OX=10090 GN=Caprin1 PE=1 SV=2 | 5 | 3 | 5 | 3 | 707 | 78,1 | 5,25 | 3,6 | 3 | 1,656 | 1,254 | 1,577 | 1,321 | 1,267 | 1,315 | 1,378 | 1,323 | 1,10 | 0,10 | 2,4E-01 |
| P50580 | Proliferation-associated protein 2G4 OS=Mus musculus OX=10090 ( | 18 | 6 | 6 | 6 | 394 | 43,7 | 6,86 | 16,52 | 6 | 0,952 | 0,941 | 0,859 | 1,058 | 1,056 | 0,921 | 1,086 | 1,033 | 0,93 | -0,07 | 2,4E-01 |
| Q8BLF1 | Neutral cholesterol ester hydrolase 1 OS=Mus musculus OX=10090 | 16 | 6 | 6 | 6 | 408 | 45,7 | 7,05 | 20,27 | 6 | 1,181 | 1,125 | 1,066 | 1,128 | 1,074 | 1,038 | 1,063 | 1,149 | 1,04 | 0,04 | 2,4E-01 |
| P84309 | Adenylate cyclase type 5 OS=Mus musculus OX=10090 GN=Adcy5 F | 6 | 8 | 9 | 8 | 1262 | 139 | 7,06 | 10,84 | 8 | 0,883 | 0,738 | 0,893 | 0,82 | 1,033 | 0,804 | 1,335 | 0,832 | 0,83 | -0,17 | 2,4E-01 |
| Q8K215 | LYR motif-containing protein 4 OS=Mus musculus OX=10090 GN=Ly | 29 | 2 | 3 | 2 | 91 | 10,8 | 10,13 | 8,66 | 2 | 1,107 | 1,024 | 0,882 | 0,86 | 0,937 | 1,12 | 1,066 | 1,142 | 0,91 | -0,09 | 2,4E-01 |
| Q9Z1P6 | NADH dehydrogenase [ubiquinone] 1 alpha subcomplex subunit 7 ( | 54 | 7 | 15 | 7 | 113 | 12,6 | 10,17 | 31,66 | 7 | 0,958 | 1,039 | 1,038 | 0,945 | 0,957 | 0,978 | 0,943 | 0,964 | 1,04 | 0,04 | 2,4E-01 |
| Q8K097 | Protein lifeguard 2 OS=Mus musculus OX=10090 GN=Faim2 PE=2 S\ | 7 | 1 | 1 | 1 | 317 | 35,2 | 6,92 | 2,45 | 1 | 0,9 | 1,051 | 0,753 | 0,741 | 0,932 | 0,938 | 1,105 | 0,917 | 0,89 | -0,11 | 2,4E-01 |
| Q8BGR9 | Ubiquitin-like domain-containing CTD phosphatase 1 OS=Mus musc | 4 | 1 | 1 | 1 | 318 | 36,8 | 6,46 | 0 | 1 | 1,159 | 0,908 | 0,528 | 0,748 | 0,679 | 0,693 | 0,712 | 0,527 | 1,28 | 0,28 | 2,4E-01 |
| Q62418 | Drebrin-like protein OS=Mus musculus OX=10090 GN=Dbnl PE=1 S\ | 16 | 5 | 7 | 5 | 436 | 48,7 | 4,92 | 9,74 | 5 | 0,84 | 0,766 | 0,996 | 0,841 | 0,919 | 0,905 | 0,896 | 1,014 | 0,92 | -0,08 | 2,4E-01 |
| Q9R1P4 | Proteasome subunit alpha type-1 OS=Mus musculus OX=10090 GN | 41 | 11 | 15 | 11 | 263 | 29,5 | 6,46 | 39,19 | 11 | 1,117 | 1,083 | 0,995 | 1,011 | 0,963 | 0,917 | 1,036 | 1,06 | 1,06 | 0,06 | 2,4E-01 |
| Q91YR1 | Twinfilin-1 OS=Mus musculus OX=10090 GN=Twf1 PE=1 SV=2 | 17 | 5 | 6 | 4 | 350 | 40,1 | 6,67 | 12,01 | 5 | 0,927 | 0,986 | 0,955 | 1,009 | 1,069 | 0,997 | 1,041 | 0,942 | 0,96 | -0,04 | 2,4E-01 |
| Q8VD75 | Huntingtin-interacting protein 1 OS=Mus musculus OX=10090 GN= | 8 | 7 | 7 | 7 | 1029 | 115,1 | 5,43 | 15,4 | 7 | 1,084 | 0,991 | 1,038 | 0,901 | 0,857 | 0,967 | 0,843 | 1,034 | 1,08 | 0,08 | 2,4E-01 |
| P13595-2 | Isoform 2 of Neural cell adhesion molecule 1 OS=Mus musculus OX | 36 | 27 | 43 | 2 | 848 | 93,5 | 4,91 | 99,4 | 27 | 1,233 | 0,893 | 1,094 | 0,996 | 0,886 | 0,966 | 0,924 | 1,03 | 1,11 | 0,11 | 2,4E-01 |
| Q9R0N5 | Synaptotagmin-5 OS=Mus musculus OX=10090 GN=Syt5 PE=1 SV=1 | 11 | 5 | 13 | 3 | 386 | 43,1 | 9,6 | 21,45 | 5 | 0,898 | 1,17 | 0,965 | 1,093 | 0,976 | 0,719 | 0,994 | 0,97 | 1,13 | 0,13 | 2,4E-01 |
| Q91YQ3 | Cold shock domain-containing protein C2 OS=Mus musculus OX=1( | 5 | 1 | 2 | 1 | 154 | 16,8 | 7,55 | 3,47 | 1 | 1,1 | 0,721 | 0,944 | 0,988 | 1,008 | 1,016 | 1,037 | 1,13 | 0,90 | -0,10 | 2,4E-01 |
| O35382 | Exocyst complex component 4 OS=Mus musculus OX=10090 GN=Ex | 14 | 12 | 13 | 12 | 975 | 110,5 | 6,49 | 9,15 | 12 | 1,059 | 0,853 | 1,135 | 0,995 | 0,993 | 1,292 | 1,077 | 1,131 | 0,90 | -0,10 | 2,4E-01 |
| Q9DB90 | Protein SMG9 OS=Mus musculus OX=10090 GN=Smg9 PE=1 SV=1 | 2 | 1 | 1 | 1 | 520 | 57,6 | 7,01 | 0 | 1 | 0,448 | 0,676 | 0,673 | 0,654 | 0,896 | 0,762 | 0,497 | 0,834 | 0,82 | -0,18 | 2,4E-01 |
| P70663 | SPARC-like protein 1 OS=Mus musculus OX=10090 GN=Sparcl1 PE= | 10 | 4 | 5 | 4 | 650 | 72,2 | 4,6 | 4,94 | 4 | 1,814 | 1,45 | 1,276 | 0,99 | 0,772 | 1,389 | 0,809 | 1,326 | 1,29 | 0,29 | 2,4E-01 |
| Q9WVL0 | Maleylacetoacetate isomerase OS=Mus musculus OX=10090 GN=G | 9 | 2 | 2 | 2 | 216 | 24,3 | 7,85 | 4,54 | 2 | 1,144 | 0,984 | 1,178 | 1,006 | 1,184 | 1,201 | 1,032 | 1,239 | 0,93 | -0,07 | 2,4E-01 |
| Q9ERN0 | Secretory carrier-associated membrane protein 2 OS=Mus musculu | 4 | 1 | 1 | 1 | 329 | 36,4 | 6,32 | 2,76 | 1 | 1,23 | 1,1 | 0,863 | 0,799 | 1,051 | 1,867 | 0,871 | 1,463 | 0,76 | -0,24 | 2,4E-01 |
| P26516 | 26S proteasome non-ATPase regulatory subunit 7 OS=Mus musculu | 7 | 2 | 5 | 2 | 321 | 36,5 | 6,77 | 3,33 | 2 | 1,195 | 0,903 | 1,165 | 1,136 | 0,81 | 1,204 | 0,865 | 0,954 | 1,15 | 0,15 | 2,4E-01 |
| Q3UQ44 | Ras GTPase-activating-like protein IQGAP2 OS=Mus musculus OX=1 | 4 | 4 | 4 | 4 | 1575 | 180,4 | 5,64 | 3,14 | 4 | 1,943 | 1,292 | 1,03 | 2,709 | 0,923 | 1,335 | 1,167 | 1,511 | 1,41 | 0,41 | 2,4E-01 |
| P0C0A3 | Charged multivesicular body protein 6 OS=Mus musculus OX=1009( | 27 | 5 | 7 | 5 | 200 | 23,4 | 5,44 | 10,08 | 5 | 1,081 | 0,804 | 0,954 | 1,104 | 1,046 | 1,296 | 1,105 | 0,99 | 0,89 | -0,11 | 2,4E-01 |
| Q3UHE1 | Membrane-associated phosphatidylinositol transfer protein 3 OS=N | 6 | 4 | 5 | 4 | 974 | 106,4 | 7,12 | 9,43 | 4 | 0,964 | 0,898 | 0,957 | 0,947 | 0,882 | 1,109 | 0,974 | 1,078 | 0,93 | -0,07 | 2,5E-01 |
| Q8BRH0 | Protein O-mannosyl-transferase TMTC3 OS=Mus musculus OX=100 | 1 | 1 | 1 | 1 | 920 | 104,1 | 8,53 | 2,29 | 1 | 0,772 | 1,039 | 0,873 | 0,919 | 0,975 | 1,028 | 0,904 | 1,015 | 0,92 | -0,08 | 2,5E-01 |
| Q5RJI5 | Serine/threonine-protein kinase BRSK1 OS=Mus musculus OX=1009 | 15 | 10 | 14 | 8 | 778 | 85,1 | 9,32 | 31,32 | 10 | 1,238 | 1,143 | 1,094 | 1,1 | 0,929 | 1,247 | 1,029 | 0,966 | 1,10 | 0,10 | 2,5E-01 |
| Q03249 | Galactose-1-phosphate uridylyltransferase OS=Mus musculus OX=1 | 3 | 1 | 1 | 1 | 379 | 43,2 | 6,77 | 2,32 | 1 | 0,853 | 0,894 | 0,847 | 0,859 | 0,886 | 1,124 | 0,829 | 0,947 | 0,91 | -0,09 | 2,5E-01 |
| Q99N96 | 39S ribosomal protein L1, mitochondrial OS=Mus musculus OX=10( | 21 | 6 | 6 | 6 | 336 | 37,6 | 8,72 | 9,24 | 6 | 1,096 | 1,032 | 0,884 | 0,965 | 1,15 | 0,949 | 1,063 | 1,157 | 0,92 | -0,08 | 2,5E-01 |
| Q91VN4 | MICOS complex subunit Mic25 OS=Mus musculus OX=10090 GN=C | 37 | 9 | 14 | 9 | 273 | 29,8 | 8,41 | 41,38 | 9 | 0,944 | 1,011 | 1,077 | 1,011 | 1,075 | 1,061 | 1,045 | 1,016 | 0,96 | -0,04 | 2,5E-01 |
| P97765 | WW domain-binding protein 2 OS=Mus musculus OX=10090 GN=W | 4 | 2 | 2 | 2 | 261 | 28 | 6,33 | 2,36 | 2 | 1,357 | 1,051 | 0,95 | 1,017 | 0,908 | 1,073 | 0,817 | 1,026 | 1,14 | 0,14 | 2,5E-01 |

| | | | | | | | | | | | | | | | | | | | |
|---|---|---|---|---|---|---|---|---|---|---|---|---|---|---|---|---|---|---|---|
| Q5SRX1 | TOM1-like protein 2 OS=Mus musculus OX=10090 GN=Tom1l2 PE=: | 7 | 2 | 7 | 2 | 507 | 55,6 | 4,82 | 6,78 | 2 | 1,498 | 1,006 | 1,125 | 0,81 | 0,823 | 0,948 | 1,002 | 0,897 | 1,21 | 0,21 | 2,5E-01 |
| Q8VDM4 | 26S proteasome non-ATPase regulatory subunit 2 OS=Mus musculu | 8 | 7 | 10 | 7 | 908 | 100,1 | 5,17 | 24,14 | 7 | 0,919 | 0,913 | 1,017 | 1,085 | 1,096 | 1,053 | 1,016 | 1,006 | 0,94 | -0,06 | 2,5E-01 |
| Q3TDK6 | Protein rogdi homolog OS=Mus musculus OX=10090 GN=Rogdi PE= | 29 | 7 | 7 | 7 | 287 | 32,1 | 8,18 | 23,3 | 7 | 0,964 | 0,886 | 0,947 | 1,109 | 1,043 | 1 | 1,113 | 1,022 | 0,93 | -0,07 | 2,5E-01 |
| Q69ZF7 | Metal transporter CNNM4 OS=Mus musculus OX=10090 GN=Cnnm | 3 | 2 | 2 | 1 | 771 | 86,6 | 6,07 | 4,34 | 2 | 0,855 | 0,979 | 1,049 | 1,145 | 1,002 | 1,211 | 1,092 | 1,104 | 0,91 | -0,09 | 2,5E-01 |
| Q99JI6 | Ras-related protein Rap-1b OS=Mus musculus OX=10090 GN=Rap1 | 37 | 7 | 16 | 2 | 184 | 20,8 | 5,78 | 36,49 | 7 | 1,159 | 0,891 | 1,232 | 1,218 | 1,042 | 0,963 | 1,088 | 0,973 | 1,11 | 0,11 | 2,5E-01 |
| Q9D1D4 | Transmembrane emp24 domain-containing protein 10 OS=Mus mu | 9 | 2 | 3 | 2 | 219 | 24,9 | 6,7 | 4,44 | 2 | 0,797 | 0,941 | 0,827 | 1,155 | 0,969 | 0,725 | 0,8 | 0,714 | 1,16 | 0,16 | 2,5E-01 |
| P63330 | Serine/threonine-protein phosphatase 2A catalytic subunit alpha is | 28 | 6 | 13 | 6 | 309 | 35,6 | 5,54 | 32,72 | 6 | 1,098 | 1,08 | 1,028 | 0,965 | 0,954 | 1,04 | 1,004 | 0,996 | 1,04 | 0,04 | 2,5E-01 |
| Q6PAJ1 | Breakpoint cluster region protein OS=Mus musculus OX=10090 GN= | 4 | 5 | 6 | 4 | 1270 | 143 | 6,92 | 9,15 | 5 | 1,443 | 1,24 | 1,238 | 1,17 | 1,24 | 1,117 | 1,242 | 1,15 | 1,07 | 0,07 | 2,5E-01 |
| Q8R3V5-1 | Isoform 1 of Endophilin-B2 OS=Mus musculus OX=10090 GN=Sh3gl | 27 | 10 | 17 | 10 | 395 | 44,1 | 5,99 | 30,93 | 10 | 0,907 | 0,916 | 0,998 | 0,973 | 0,944 | 0,953 | 1,035 | 1,029 | 0,96 | -0,04 | 2,5E-01 |
| Q91ZJ5 | UTP--glucose-1-phosphate uridylyltransferase OS=Mus musculus O | 30 | 15 | 18 | 15 | 508 | 56,9 | 7,61 | 39,21 | 15 | 1,09 | 1,006 | 1,099 | 0,987 | 0,988 | 1,006 | 0,995 | 1,037 | 1,04 | 0,04 | 2,5E-01 |
| Q923Z3 | Protein MTO1 homolog, mitochondrial OS=Mus musculus OX=1009 | 5 | 3 | 3 | 3 | 669 | 74,3 | 8,92 | 2,6 | 3 | 0,516 | 0,815 | 1,005 | 0,829 | 1,04 | 0,915 | 0,77 | 1,059 | 0,84 | -0,16 | 2,5E-01 |
| Q8R3T5 | Syntaxin-binding protein 6 OS=Mus musculus OX=10090 GN=Stxbp | 14 | 3 | 4 | 3 | 210 | 23,7 | 9,19 | 6,45 | 3 | 1,097 | 1,053 | 1,012 | 1,033 | 1,041 | 0,915 | 0,939 | 1,079 | 1,06 | 0,06 | 2,5E-01 |
| P21279 | Guanine nucleotide-binding protein G(q) subunit alpha OS=Mus mu | 31 | 10 | 11 | 5 | 359 | 42,1 | 5,68 | 29,3 | 10 | 0,797 | 0,922 | 0,8 | 0,97 | 0,904 | 0,952 | 0,98 | 0,897 | 0,93 | -0,07 | 2,5E-01 |
| P47757-4 | Isoform 3 of F-actin-capping protein subunit beta OS=Mus musculu | 47 | 13 | 22 | 13 | 301 | 33,7 | 6,43 | 47,4 | 13 | 1,118 | 1,049 | 1,029 | 1,132 | 1,01 | 1,039 | 1,063 | 1,07 | 1,03 | 0,03 | 2,5E-01 |
| Q99KR7 | Peptidyl-prolyl cis-trans isomerase F, mitochondrial OS=Mus muscu | 21 | 6 | 6 | 4 | 206 | 21,7 | 9,16 | 10,94 | 6 | 1,126 | 1,081 | 1,103 | 0,936 | 0,957 | 1,07 | 0,994 | 0,973 | 1,06 | 0,06 | 2,5E-01 |
| Q8CFV4 | Neuritin OS=Mus musculus OX=10090 GN=Nrn1 PE=1 SV=1 | 23 | 4 | 7 | 4 | 142 | 15,3 | 6,99 | 18,6 | 4 | 0,825 | 0,913 | 0,988 | 0,905 | 0,913 | 0,986 | 0,954 | 0,964 | 0,95 | -0,05 | 2,5E-01 |
| Q6PDI5 | Proteasome adapter and scaffold protein ECM29 OS=Mus musculu | 1 | 2 | 2 | 2 | 1840 | 203,6 | 7,06 | 0 | 2 | 1,551 | 1,268 | 1,366 | 0,883 | 0,813 | 1,32 | 0,772 | 1,174 | 1,24 | 0,24 | 2,5E-01 |
| O70228 | Probable phospholipid-transporting ATPase IIA OS=Mus musculus ( | 5 | 5 | 6 | 3 | 1047 | 118,5 | 7,69 | 12,84 | 5 | 1,147 | 0,907 | 1,047 | 0,988 | 1,055 | 1,104 | 1,105 | 1,088 | 0,94 | -0,06 | 2,5E-01 |
| Q80TA9 | Ectopic P granules protein 5 homolog OS=Mus musculus OX=10090 | 1 | 3 | 3 | 3 | 2572 | 290,6 | 6,37 | 4,58 | 3 | 1,267 | 0,909 | 0,828 | 0,712 | 0,692 | 0,908 | 0,854 | 0,491 | 1,26 | 0,26 | 2,5E-01 |
| Q9EQU5 | Protein SET OS=Mus musculus OX=10090 GN=Set PE=1 SV=1 | 8 | 2 | 2 | 2 | 289 | 33,4 | 4,32 | 3,81 | 2 | 1,406 | 1,289 | 1,506 | 1,479 | 1,339 | 1,436 | 1,859 | 1,698 | 0,90 | -0,10 | 2,5E-01 |
| Q99N87 | 28S ribosomal protein S5, mitochondrial OS=Mus musculus OX=100 | 9 | 4 | 4 | 4 | 432 | 48,2 | 10,14 | 9 | 4 | 0,863 | 0,819 | 0,97 | 1,016 | 1,057 | 0,868 | 0,981 | 1,124 | 0,91 | -0,09 | 2,5E-01 |
| Q80Y56 | Rabenosyn-5 OS=Mus musculus OX=10090 GN=Rbsn PE=1 SV=1 | 6 | 5 | 5 | 5 | 783 | 88,4 | 5,3 | 4,35 | 5 | 0,813 | 0,878 | 1,053 | 0,963 | 0,956 | 0,971 | 1,034 | 1,028 | 0,93 | -0,07 | 2,5E-01 |
| Q99JB8 | Protein kinase C and casein kinase II substrate protein 3 OS=Mus m | 6 | 2 | 2 | 2 | 424 | 48,6 | 6,1 | 5,22 | 2 | 0,877 | 0,866 | 0,974 | 1,071 | 1,031 | 0,95 | 1,039 | 1,033 | 0,93 | -0,07 | 2,5E-01 |
| O08583 | THO complex subunit 4 OS=Mus musculus OX=10090 GN=Alyref PE | 11 | 2 | 2 | 2 | 255 | 26,9 | 11,15 | 5,52 | 2 | 0,782 | 0,737 | 0,708 | 0,89 | 0,672 | 0,944 | 0,89 | 1,114 | 0,86 | -0,14 | 2,5E-01 |
| Q9Z0E0 | Neurochondrin OS=Mus musculus OX=10090 GN=Ncdn PE=1 SV=1 | 21 | 14 | 21 | 14 | 729 | 78,8 | 5,54 | 46,8 | 14 | 1,063 | 1,111 | 1,094 | 1,117 | 0,952 | 1,168 | 1,018 | 1,005 | 1,06 | 0,06 | 2,5E-01 |
| Q62167 | ATP-dependent RNA helicase DDX3X OS=Mus musculus OX=10090 | 12 | 7 | 7 | 3 | 662 | 73,1 | 7,18 | 10,43 | 7 | 1,056 | 1,113 | 1,037 | 0,955 | 1,054 | 1,043 | 1,174 | 1,115 | 0,95 | -0,05 | 2,5E-01 |
| Q7TMY8 | E3 ubiquitin-protein ligase HUWE1 OS=Mus musculus OX=10090 GI | 3 | 10 | 15 | 10 | 4377 | 482,3 | 5,22 | 33,64 | 10 | 1,031 | 1,029 | 1,138 | 1,112 | 1,07 | 1,164 | 1,131 | 1,117 | 0,96 | -0,04 | 2,5E-01 |
| Q9CQA1 | Trafficking protein particle complex subunit 5 OS=Mus musculus O) | 5 | 1 | 1 | 1 | 188 | 20,8 | 9,66 | 1,77 | 1 | 2,162 | 1,641 | 1,113 | 1,606 | 1,488 | 1,528 | 1,155 | 1,153 | 1,23 | 0,23 | 2,5E-01 |
| Q63829 | COMM domain-containing protein 3 OS=Mus musculus OX=10090 ( | 12 | 2 | 2 | 2 | 195 | 22 | 5,59 | 5,59 | 2 | 1,421 | 1,419 | 1,154 | 1,021 | 1,155 | 0,977 | 1,208 | 1,114 | 1,13 | 0,13 | 2,5E-01 |
| P56395 | Cytochrome b5 OS=Mus musculus OX=10090 GN=Cyb5a PE=1 SV=2 | 34 | 3 | 5 | 3 | 134 | 15,2 | 5,07 | 14,58 | 3 | 1,475 | 1,367 | 1,068 | 0,876 | 0,97 | 1,173 | 0,899 | 0,993 | 1,19 | 0,19 | 2,5E-01 |
| Q8R4H2 | Rho guanine nucleotide exchange factor 12 OS=Mus musculus OX= | 8 | 11 | 11 | 11 | 1543 | 172,2 | 5,74 | 20,08 | 11 | 0,985 | 0,995 | 1,011 | 1,092 | 0,972 | 1,133 | 1,095 | 1,099 | 0,95 | -0,05 | 2,5E-01 |
| Q9JKL5 | Calcineurin B homologous protein 3 OS=Mus musculus OX=10090 ( | 18 | 3 | 3 | 3 | 214 | 24,6 | 4,98 | 6,65 | 3 | 1,11 | 0,946 | 1,111 | 1,048 | 1,212 | 1,048 | 1,182 | 1,06 | 0,94 | -0,06 | 2,6E-01 |
| Q9D7M1 | Glucose-induced degradation protein 8 homolog OS=Mus musculus | 3 | 1 | 1 | 1 | 228 | 26,8 | 4,97 | 2,11 | 1 | 0,953 | 0,945 | 1,067 | 0,915 | 1,015 | 0,799 | 0,794 | 0,948 | 1,09 | 0,09 | 2,6E-01 |
| Q8K341 | Alpha-tubulin N-acetyltransferase 1 OS=Mus musculus OX=10090 G | 12 | 4 | 6 | 4 | 421 | 47,1 | 9,98 | 8,69 | 4 | 0,851 | 1,062 | 0,88 | 1,213 | 1,01 | 1,214 | 1,146 | 1,111 | 0,89 | -0,11 | 2,6E-01 |
| Q9EPK7 | Exportin-7 OS=Mus musculus OX=10090 GN=Xpo7 PE=1 SV=3 | 6 | 6 | 7 | 6 | 1087 | 123,7 | 6,38 | 15,68 | 6 | 1,089 | 1,08 | 1,082 | 1,024 | 0,943 | 1,082 | 0,924 | 1,091 | 1,06 | 0,06 | 2,6E-01 |
| P47867 | Secretogranin-3 OS=Mus musculus OX=10090 GN=Scg3 PE=1 SV=1 | 5 | 2 | 2 | 2 | 471 | 53,3 | 4,93 | 5,98 | 2 | 1,156 | 0,881 | 0,844 | 1,033 | 0,989 | 1,205 | 0,983 | 1,23 | 0,89 | -0,11 | 2,6E-01 |
| A2AJA9 | Apical junction component 1 homolog OS=Mus musculus OX=1009 | 8 | 6 | 6 | 6 | 974 | 107,1 | 9,11 | 16,87 | 6 | 1,373 | 1,204 | 0,942 | 1,048 | 0,875 | 1,167 | 0,974 | 0,99 | 1,14 | 0,14 | 2,6E-01 |
| P16125 | L-lactate dehydrogenase B chain OS=Mus musculus OX=10090 GN= | 38 | 14 | 29 | 13 | 334 | 36,5 | 6,05 | 62,38 | 14 | 0,799 | 0,937 | 0,906 | 0,972 | 0,951 | 0,94 | 1,03 | 0,917 | 0,94 | -0,06 | 2,6E-01 |
| Q9JMI0 | Endothelin-converting enzyme-like 1 OS=Mus musculus OX=10090 | 1 | 1 | 1 | 1 | 775 | 87,9 | 7,8 | 1,91 | 1 | 1,398 | 1,061 | 1,011 | 1,103 | 1,12 | 1,096 | 0,928 | 0,917 | 1,13 | 0,13 | 2,6E-01 |
| Q8BV13 | COP9 signalosome complex subunit 7b OS=Mus musculus OX=1009 | 10 | 2 | 3 | 2 | 264 | 29,7 | 6,32 | 7,07 | 2 | 1,403 | 1,305 | 1,314 | 1,116 | 0,955 | 1,359 | 1,138 | 1,174 | 1,11 | 0,11 | 2,6E-01 |
| Q80TL0 | Protein phosphatase 1E OS=Mus musculus OX=10090 GN=Ppm1e P | 10 | 6 | 8 | 6 | 749 | 83,4 | 4,97 | 15,1 | 6 | 1,077 | 1,008 | 1,159 | 0,937 | 0,959 | 1,045 | 0,994 | 0,903 | 1,07 | 0,07 | 2,6E-01 |
| P62270 | 40S ribosomal protein S18 OS=Mus musculus OX=10090 GN=Rps18 | 12 | 2 | 3 | 2 | 152 | 17,7 | 10,99 | 7,59 | 2 | 1,127 | 1,051 | 1,09 | 0,936 | 1,044 | 0,981 | 0,751 | 1,03 | 1,10 | 0,10 | 2,6E-01 |
| Q62393-3 | Isoform 3 of Tumor protein D52 OS=Mus musculus OX=10090 GN=T | 16 | 3 | 5 | 3 | 199 | 21,6 | 5,03 | 6,5 | 3 | 0,594 | 0,824 | 0,819 | 0,929 | 0,901 | 0,778 | 1,109 | 0,873 | 0,86 | -0,14 | 2,6E-01 |
| Q9Z1J3 | Cysteine desulfurase, mitochondrial OS=Mus musculus OX=10090 ( | 21 | 9 | 12 | 9 | 459 | 50,5 | 8,16 | 26,38 | 9 | 1,141 | 1,083 | 1,034 | 0,982 | 1,004 | 1,05 | 1,002 | 1,005 | 1,04 | 0,04 | 2,6E-01 |
| Q7TQD2 | Tubulin polymerization-promoting protein OS=Mus musculus OX=1 | 33 | 9 | 17 | 9 | 218 | 23,6 | 9,42 | 35,08 | 9 | 0,879 | 0,991 | 0,938 | 0,988 | 0,981 | 0,911 | 1,104 | 1,045 | 0,94 | -0,06 | 2,6E-01 |
| P16460 | Argininosuccinate synthase OS=Mus musculus OX=10090 GN=Ass1 | 18 | 8 | 9 | 8 | 412 | 46,6 | 8,22 | 19,04 | 8 | 0,797 | 0,863 | 0,94 | 1,063 | 1,029 | 0,903 | 1,121 | 0,975 | 0,91 | -0,09 | 2,6E-01 |
| Q9JL62 | Glycolipid transfer protein OS=Mus musculus OX=10090 GN=Gltp P | 4 | 1 | 1 | 1 | 209 | 23,7 | 7,39 | 2,04 | 1 | 1,203 | 1,996 | 0,822 | 0,748 | 1,132 | 0,69 | 0,73 | 0,7 | 1,47 | 0,47 | 2,6E-01 |
| A6X919 | Probable C-mannosyltransferase DPY19L1 OS=Mus musculus OX=1 | 2 | 1 | 1 | 1 | 746 | 84,1 | 9,16 | 0 | 1 | 0,653 | 2,549 | 0,591 | 0,709 | 0,524 | 0,363 | 0,551 | 0,68 | 2,13 | 1,13 | 2,6E-01 |
| O08747 | Netrin receptor UNC5C OS=Mus musculus OX=10090 GN=Unc5c PE | 1 | 1 | 1 | 1 | 931 | 103 | 6,01 | 1,72 | 1 | 1,152 | 1,195 | 1,618 | 1,143 | 1,089 | 0,833 | 1,329 | 1,098 | 1,17 | 0,17 | 2,6E-01 |
| Q8CGA4 | Maturin OS=Mus musculus OX=10090 GN=Mturn PE=1 SV=1 | 6 | 1 | 1 | 1 | 131 | 14,8 | 4,17 | 2,53 | 1 | 1,263 | 1,132 | 1,105 | 1,128 | 0,886 | 1,199 | 1,167 | 0,95 | 1,10 | 0,10 | 2,6E-01 |
| P63054 | Calmodulin regulator protein PCP4 OS=Mus musculus OX=10090 GI | 44 | 3 | 7 | 3 | 62 | 6,8 | 6,71 | 13,45 | 3 | 1,036 | 0,857 | 1,075 | 1,084 | 0,841 | 1,024 | 0,998 | 0,816 | 1,10 | 0,10 | 2,6E-01 |
| P62313 | U6 snRNA-associated Sm-like protein LSm6 OS=Mus musculus OX= | 10 | 1 | 1 | 1 | 80 | 9,1 | 9,58 | 1,93 | 1 | 1,135 | 0,96 | 0,679 | 0,999 | 0,877 | 0,878 | 0,728 | 0,78 | 1,16 | 0,16 | 2,6E-01 |
| Q8BFZ3 | Beta-actin-like protein 2 OS=Mus musculus OX=10090 GN=Actbl2 P | 26 | 10 | 82 | 3 | 376 | 42 | 5,49 | 172,35 | 10 | 1,089 | 1,272 | 1,546 | 4,767 | 0,719 | 1,33 | 1,142 | 1,114 | 2,01 | 1,01 | 2,6E-01 |
| O88845 | A-kinase anchor protein 10, mitochondrial OS=Mus musculus OX=1 | 6 | 3 | 3 | 3 | 662 | 73,6 | 6,79 | 2,53 | 3 | 2,292 | 2,309 | 1,939 | 1,987 | 1,669 | 1,574 | 1,843 | 2,402 | 1,14 | 0,14 | 2,6E-01 |
| Q8K0T4 | Katanin p60 ATPase-containing subunit A-like 1 OS=Mus musculus ( | 5 | 2 | 2 | 2 | 488 | 55,1 | 7,09 | 2,7 | 2 | 0,676 | 0,937 | 0,593 | 0,805 | 1,013 | 0,813 | 0,948 | 0,728 | 0,86 | -0,14 | 2,6E-01 |
| P48318 | Glutamate decarboxylase 1 OS=Mus musculus OX=10090 GN=Gad1 | 13 | 6 | 9 | 5 | 593 | 66,6 | 7,17 | 20,89 | 6 | 1,007 | 1,069 | 1,058 | 1,017 | 0,983 | 1,32 | 1,14 | 1,07 | 0,92 | -0,08 | 2,6E-01 |
| Q9Z275 | Retinaldehyde-binding protein 1 OS=Mus musculus OX=10090 GN= | 3 | 1 | 2 | 1 | 317 | 36,4 | 5,06 | 5,09 | 1 | 1,104 | 0,949 | 1,178 | 1,076 | 0,953 | 0,931 | 0,909 | 1,151 | 1,09 | 0,09 | 2,6E-01 |
| P61222 | ATP-binding cassette sub-family E member 1 OS=Mus musculus OX | 5 | 3 | 3 | 3 | 599 | 67,3 | 8,34 | 6,81 | 3 | 1,409 | 0,907 | 0,941 | 0,936 | 0,775 | 0,865 | 0,938 | 0,979 | 1,18 | 0,18 | 2,6E-01 |
| Q6RHR9 | Membrane-associated guanylate kinase, WW and PDZ domain-con | 5 | 7 | 8 | 7 | 1471 | 161,9 | 7,36 | 15 | 7 | 0,797 | 0,876 | 0,873 | 0,833 | 0,855 | 0,869 | 0,949 | 0,852 | 0,96 | -0,04 | 2,6E-01 |
| Q7TSH2 | Phosphorylase b kinase regulatory subunit beta OS=Mus musculus | 2 | 2 | 2 | 2 | 1085 | 123,8 | 6,83 | 0 | 2 | 0,45 | 0,526 | 0,468 | 0,931 | 0,894 | 0,698 | 0,97 | 0,547 | 0,76 | -0,24 | 2,6E-01 |
| P42125 | Enoyl-CoA delta isomerase 1, mitochondrial OS=Mus musculus OX= | 22 | 6 | 8 | 6 | 289 | 32,2 | 8,98 | 9,21 | 6 | 0,972 | 1,139 | 1,09 | 0,941 | 1,061 | 1,105 | 1,055 | 1,223 | 0,93 | -0,07 | 2,6E-01 |
| Q9Z2M6 | Ubiquitin-like protein 3 OS=Mus musculus OX=10090 GN=Ubl3 PE= | 23 | 2 | 2 | 2 | 117 | 13,2 | 6,92 | 2,39 | 2 | 2,322 | 1,416 | 1,057 | 0,966 | 0,873 | 1,186 | 0,992 | 1,141 | 1,37 | 0,37 | 2,6E-01 |

| | | | | | | | | | | | | | | | | | | | | | |
|---|---|---|---|---|---|---|---|---|---|---|---|---|---|---|---|---|---|---|---|---|---|
| Q9QYH6 | Melanoma-associated antigen D1 OS=Mus musculus OX=10090 GN | 2 | 1 | 1 | 1 | 775 | 85,6 | 7,5 | 3,36 | 1 | 0,971 | 0,723 | 0,858 | 0,959 | 0,74 | 0,844 | 0,847 | 0,766 | 1,10 | 0,10 | 2,6E-01 |
| P0C192 | Leucine-rich repeat-containing protein 4B OS=Mus musculus OX=10 | 6 | 4 | 4 | 4 | 709 | 76,1 | 7,24 | 7,67 | 4 | 1,356 | 1,074 | 1,118 | 0,92 | 0,938 | 1,076 | 0,991 | 0,996 | 1,12 | 0,12 | 2,6E-01 |
| Q9ESN9 | C-Jun-amino-terminal kinase-interacting protein 3 OS=Mus muscul | 8 | 8 | 8 | 6 | 1337 | 147,5 | 5,45 | 17,65 | 8 | 0,911 | 0,936 | 0,945 | 0,916 | 0,883 | 1,001 | 1,084 | 0,952 | 0,95 | -0,05 | 2,6E-01 |
| O08553 | Dihydropyrimidinase-related protein 2 OS=Mus musculus OX=1009 | 59 | 24 | 108 | 21 | 572 | 62,2 | 6,38 | 284,93 | 24 | 0,952 | 0,934 | 0,977 | 0,994 | 0,981 | 0,939 | 1,024 | 1,049 | 0,97 | -0,03 | 2,6E-01 |
| Q8BLK3 | Limbic system-associated membrane protein OS=Mus musculus OX | 18 | 6 | 15 | 6 | 341 | 38,1 | 6,65 | 38,96 | 6 | 1,042 | 0,808 | 1,042 | 0,863 | 1,074 | 0,951 | 1,184 | 0,952 | 0,90 | -0,10 | 2,6E-01 |
| Q9CR26 | Vacuolar protein sorting-associated protein VTA1 homolog OS=Mu | 6 | 1 | 1 | 1 | 309 | 33,9 | 6,13 | 4,36 | 1 | 1,443 | 1,292 | 1,186 | 1,027 | 0,887 | 1,377 | 1,049 | 0,947 | 1,16 | 0,16 | 2,7E-01 |
| O35114 | Lysosome membrane protein 2 OS=Mus musculus OX=10090 GN=S | 2 | 1 | 1 | 1 | 478 | 54 | 5,1 | 2,35 | 1 | 0,984 | 1,121 | 1,038 | 0,963 | 0,918 | 1,204 | 1,105 | 1,454 | 0,88 | -0,12 | 2,7E-01 |
| Q9D0W5 | Peptidyl-prolyl cis-trans isomerase-like 1 OS=Mus musculus OX=10 | 15 | 2 | 2 | 2 | 166 | 18,2 | 7,99 | 0 | 2 | 1,986 | 1,275 | 1,331 | 1,57 | 0,969 | 1,731 | 1,122 | 1,204 | 1,23 | 0,23 | 2,7E-01 |
| Q8BYA0 | Tubulin-specific chaperone D OS=Mus musculus OX=10090 GN=Tb | 5 | 5 | 5 | 5 | 1196 | 133,2 | 6,51 | 7,85 | 5 | 1,077 | 1,315 | 1 | 0,953 | 0,951 | 1,138 | 0,825 | 0,922 | 1,13 | 0,13 | 2,7E-01 |
| Q9DBS1 | Transmembrane protein 43 OS=Mus musculus OX=10090 GN=Tme | 6 | 2 | 2 | 2 | 400 | 44,8 | 7,36 | 2,57 | 2 | 1,776 | 1,738 | 1,293 | 1,64 | 1,45 | 1,298 | 1,148 | 1,734 | 1,15 | 0,15 | 2,7E-01 |
| Q8VDP6 | CDP-diacylglycerol--inositol 3-phosphatidyltransferase OS=Mus mu | 10 | 2 | 3 | 2 | 213 | 23,6 | 8,27 | 5,58 | 2 | 1,458 | 1,309 | 1,064 | 0,913 | 0,811 | 1,161 | 0,952 | 1,107 | 1,18 | 0,18 | 2,7E-01 |
| Q8QZY1 | Eukaryotic translation initiation factor 3 subunit L OS=Mus muscul | 7 | 4 | 5 | 4 | 564 | 66,6 | 6,44 | 4,15 | 4 | 0,98 | 0,91 | 1,154 | 1,15 | 1,037 | 0,895 | 0,965 | 0,964 | 1,09 | 0,09 | 2,7E-01 |
| Q8BYM5 | Neuroligin-3 OS=Mus musculus OX=10090 GN=Nlgn3 PE=1 SV=2 | 3 | 2 | 3 | 2 | 825 | 91,1 | 5,8 | 4,43 | 2 | 2,061 | 1,846 | 0,843 | 1,176 | 1,121 | 1,085 | 1,288 | 1,011 | 1,32 | 0,32 | 2,7E-01 |
| Q80U87 | Ubiquitin carboxyl-terminal hydrolase 8 OS=Mus musculus OX=100 | 2 | 2 | 2 | 2 | 1080 | 122,5 | 8,47 | 0 | 2 | 0,849 | 0,962 | 1,147 | 1,113 | 0,933 | 0,92 | 0,862 | 0,993 | 1,10 | 0,10 | 2,7E-01 |
| P0DN34 | NADH dehydrogenase [ubiquinone] 1 beta subcomplex subunit 1 O | 19 | 1 | 1 | 1 | 57 | 7 | 8,21 | 3,02 | 1 | 0,81 | 0,98 | 0,955 | 1,071 | 1,038 | 0,914 | 1,324 | 1,04 | 0,88 | -0,12 | 2,7E-01 |
| O35864 | COP9 signalosome complex subunit 5 OS=Mus musculus OX=10090 | 14 | 5 | 5 | 5 | 334 | 37,5 | 6,54 | 7,16 | 5 | 1,206 | 1,184 | 1,201 | 1,125 | 0,836 | 1,204 | 1,017 | 1,213 | 1,10 | 0,10 | 2,7E-01 |
| Q61214 | Dual specificity tyrosine-phosphorylation-regulated kinase 1A OS=N | 4 | 3 | 4 | 3 | 763 | 85,4 | 8,75 | 8,11 | 3 | 1,083 | 1,03 | 0,979 | 0,969 | 0,942 | 1,239 | 1,074 | 1,134 | 0,93 | -0,07 | 2,7E-01 |
| Q8C3W1 | Uncharacterized protein C1orf198 homolog OS=Mus musculus OX= | 8 | 2 | 2 | 2 | 322 | 35,3 | 5,24 | 4,67 | 2 | 1,088 | 1,132 | 1,156 | 0,933 | 0,858 | 0,956 | 0,958 | 1,151 | 1,10 | 0,10 | 2,7E-01 |
| P70295 | Ancient ubiquitous protein 1 OS=Mus musculus OX=10090 GN=Aup | 2 | 1 | 1 | 1 | 410 | 46,1 | 8,4 | 0 | 1 | 0,964 | 0,824 | 1,044 | 1,239 | 0,937 | 0,974 | 0,73 | 0,93 | 1,14 | 0,14 | 2,7E-01 |
| P97797 | Tyrosine-protein phosphatase non-receptor type substrate 1 OS=M | 25 | 9 | 28 | 9 | 513 | 56,4 | 8,28 | 59,58 | 9 | 0,901 | 0,817 | 1,009 | 1,104 | 1,121 | 0,928 | 1,152 | 1,022 | 0,91 | -0,09 | 2,7E-01 |
| Q920N7 | Synaptotagmin-12 OS=Mus musculus OX=10090 GN=Syt12 PE=1 SV | 15 | 6 | 8 | 6 | 421 | 46,7 | 5,64 | 24,21 | 6 | 1,145 | 1,132 | 1,012 | 1,03 | 0,957 | 1,086 | 1,007 | 1,054 | 1,05 | 0,05 | 2,7E-01 |
| O88696 | ATP-dependent Clp protease proteolytic subunit, mitochondrial OS | 21 | 4 | 8 | 4 | 272 | 29,8 | 7,47 | 20,72 | 4 | 1,259 | 1,207 | 1,193 | 1,021 | 0,966 | 1,223 | 1,011 | 1,107 | 1,09 | 0,09 | 2,7E-01 |
| Q91WL8 | WW domain-containing oxidoreductase OS=Mus musculus OX=100 | 10 | 4 | 4 | 4 | 414 | 46,5 | 7,02 | 3,79 | 4 | 0,541 | 0,837 | 0,723 | 0,84 | 0,884 | 0,673 | 0,923 | 0,903 | 0,87 | -0,13 | 2,7E-01 |
| Q9CQE5 | Regulator of G-protein signaling 10 OS=Mus musculus OX=10090 G | 7 | 1 | 1 | 1 | 181 | 21,1 | 6,81 | 2,06 | 1 | 0,979 | 1,195 | 1,028 | 1,281 | 1,336 | 1,358 | 1,507 | 0,951 | 0,87 | -0,13 | 2,7E-01 |
| Q921F4 | Heterogeneous nuclear ribonucleoprotein L-like OS=Mus musculus | 2 | 1 | 1 | 1 | 591 | 64,1 | 5,85 | 0 | 1 | 1,944 | 1,057 | 1,328 | 0,935 | 0,814 | 1,124 | 0,784 | 1,298 | 1,31 | 0,31 | 2,7E-01 |
| Q9QZX7 | Serine racemase OS=Mus musculus OX=10090 GN=Srr PE=1 SV=1 | 14 | 4 | 7 | 4 | 339 | 36,3 | 6,02 | 14,84 | 4 | 0,991 | 1,061 | 1,047 | 1,036 | 0,937 | 1,013 | 1,057 | 0,986 | 1,04 | 0,04 | 2,7E-01 |
| P62918 | 60S ribosomal protein L8 OS=Mus musculus OX=10090 GN=Rpl8 PE | 12 | 3 | 3 | 3 | 257 | 28 | 11,03 | 7,07 | 3 | 0,982 | 1,055 | 0,935 | 0,865 | 0,967 | 0,858 | 0,765 | 0,95 | 1,08 | 0,08 | 2,7E-01 |
| Q8CE50 | Sorting nexin-30 OS=Mus musculus OX=10090 GN=Snx30 PE=1 SV= | 11 | 5 | 8 | 5 | 437 | 49,5 | 5,35 | 19,72 | 5 | 0,933 | 0,889 | 0,831 | 0,956 | 1,021 | 0,832 | 0,952 | 1,17 | 0,91 | -0,09 | 2,7E-01 |
| Q8BG40 | Katanin p80 WD40 repeat-containing subunit B1 OS=Mus musculus | 7 | 4 | 4 | 4 | 658 | 72,6 | 7,27 | 9 | 4 | 1,247 | 1,249 | 0,934 | 0,942 | 0,832 | 1,051 | 1,047 | 0,942 | 1,13 | 0,13 | 2,7E-01 |
| Q8BX10-2 | Isoform 2 of Serine/threonine-protein phosphatase PGAM5, mitoch | 38 | 8 | 8 | 8 | 287 | 31,8 | 9,04 | 21,22 | 8 | 1,061 | 1,129 | 1,017 | 1,058 | 1,002 | 1,041 | 1,074 | 1,011 | 1,03 | 0,03 | 2,7E-01 |
| O54781 | SRSF protein kinase 2 OS=Mus musculus OX=10090 GN=Srpk2 PE=1 | 8 | 4 | 4 | 4 | 681 | 76,7 | 4,91 | 4,91 | 4 | 0,939 | 1,012 | 0,921 | 0,872 | 0,845 | 0,953 | 0,862 | 0,902 | 1,05 | 0,05 | 2,7E-01 |
| Q921L5 | Conserved oligomeric Golgi complex subunit 2 OS=Mus musculus C | 2 | 1 | 1 | 1 | 731 | 82 | 6,21 | 1,67 | 1 | 1,15 | 1,408 | 0,912 | 0,82 | 0,69 | 1,168 | 0,573 | 0,953 | 1,27 | 0,27 | 2,7E-01 |
| O08529 | Calpain-2 catalytic subunit OS=Mus musculus OX=10090 GN=Capn2 | 13 | 8 | 9 | 8 | 700 | 79,8 | 4,96 | 23,41 | 8 | 0,974 | 1,054 | 1,043 | 0,977 | 1,047 | 1,078 | 1,067 | 0,993 | 0,97 | -0,03 | 2,7E-01 |
| Q99J85 | Neuronal pentraxin receptor OS=Mus musculus OX=10090 GN=Npt | 6 | 3 | 3 | 3 | 493 | 52,3 | 6,13 | 7,54 | 3 | 0,686 | 0,919 | 0,952 | 1,041 | 0,975 | 0,917 | 1,071 | 1,035 | 0,90 | -0,10 | 2,7E-01 |
| Q6R891 | Neurabin-2 OS=Mus musculus OX=10090 GN=Ppp1r9b PE=1 SV=1 | 11 | 9 | 11 | 9 | 817 | 89,5 | 4,92 | 10,49 | 9 | 1,161 | 1,099 | 1,03 | 0,918 | 0,99 | 1,053 | 0,897 | 0,973 | 1,08 | 0,08 | 2,7E-01 |
| Q8BWD8 | Cyclin-dependent kinase 19 OS=Mus musculus OX=10090 GN=Cdk1 | 2 | 1 | 1 | 1 | 501 | 56,5 | 8,53 | 2,16 | 1 | 0,594 | 1,131 | 0,916 | 1,139 | 1,175 | 0,949 | 1,176 | 1,151 | 0,85 | -0,15 | 2,7E-01 |
| Q9D289 | Trafficking protein particle complex subunit 6B OS=Mus musculus C | 15 | 2 | 2 | 2 | 158 | 17,9 | 8,68 | 5,04 | 2 | 1,351 | 1,229 | 1,107 | 0,999 | 0,993 | 1,187 | 0,993 | 1,084 | 1,10 | 0,10 | 2,7E-01 |
| Q5ND34 | WD repeat-containing protein 81 OS=Mus musculus OX=10090 GN= | 1 | 2 | 2 | 2 | 1934 | 211,8 | 5,77 | 4,88 | 2 | 1,172 | 0,971 | 0,936 | 1,001 | 0,968 | 0,903 | 0,976 | 0,968 | 1,07 | 0,07 | 2,7E-01 |
| P60469 | Liprin-alpha-3 OS=Mus musculus OX=10090 GN=Ppfia3 PE=1 SV=2 | 29 | 31 | 43 | 29 | 1194 | 133,3 | 5,64 | 72,1 | 31 | 1,11 | 1,128 | 1,095 | 1,084 | 0,971 | 1,142 | 0,999 | 1,103 | 1,05 | 0,05 | 2,7E-01 |
| Q99MN1 | Lysine--tRNA ligase OS=Mus musculus OX=10090 GN=Kars PE=1 SV | 4 | 2 | 2 | 2 | 595 | 67,8 | 5,94 | 6,05 | 2 | 0,839 | 0,956 | 0,943 | 0,869 | 1,005 | 1,018 | 0,883 | 0,912 | 0,94 | -0,06 | 2,8E-01 |
| O55125 | Protein NipSnap homolog 1 OS=Mus musculus OX=10090 GN=Nips | 20 | 6 | 10 | 6 | 284 | 33,3 | 9,44 | 22,76 | 6 | 1,008 | 1,015 | 0,946 | 0,913 | 0,92 | 1,157 | 1,021 | 1,046 | 0,94 | -0,06 | 2,8E-01 |
| Q9D5V5 | Cullin-5 OS=Mus musculus OX=10090 GN=Cul5 PE=1 SV=3 | 13 | 12 | 13 | 12 | 780 | 90,9 | 7,81 | 33,88 | 12 | 0,98 | 0,982 | 1,065 | 0,995 | 0,899 | 0,923 | 1,051 | 0,962 | 1,05 | 0,05 | 2,8E-01 |
| Q9DCT1 | 1,5-anhydro-D-fructose reductase OS=Mus musculus OX=10090 GN | 8 | 2 | 2 | 2 | 301 | 34,4 | 7,33 | 2,84 | 2 | 1,773 | 1,559 | 1,087 | 0,935 | 1,035 | 0,881 | 0,801 | 1,46 | 1,28 | 0,28 | 2,8E-01 |
| O54983 | Ketimine reductase mu-crystallin OS=Mus musculus OX=10090 GN= | 19 | 5 | 6 | 5 | 313 | 33,5 | 5,67 | 18,43 | 5 | 0,926 | 0,882 | 1,004 | 1,007 | 1,039 | 0,937 | 1,065 | 0,98 | 0,95 | -0,05 | 2,8E-01 |
| O88448 | Kinesin light chain 2 OS=Mus musculus OX=10090 GN=Klc2 PE=1 SV | 25 | 13 | 15 | 7 | 599 | 66,6 | 7,21 | 38,25 | 13 | 0,744 | 0,923 | 0,876 | 0,995 | 0,928 | 0,861 | 0,988 | 1,153 | 0,90 | -0,10 | 2,8E-01 |
| P29758 | Ornithine aminotransferase, mitochondrial OS=Mus musculus OX=1 | 14 | 5 | 6 | 5 | 439 | 48,3 | 6,62 | 11,72 | 5 | 1,185 | 1,111 | 1,046 | 1,025 | 0,979 | 1,135 | 0,984 | 1,024 | 1,06 | 0,06 | 2,8E-01 |
| Q6PGE7 | Sodium-dependent proline transporter OS=Mus musculus OX=1009 | 3 | 1 | 1 | 1 | 637 | 71 | 6,61 | 2,41 | 1 | 0,903 | 1,249 | 1,207 | 1,045 | 0,628 | 0,948 | 1,206 | 0,94 | 1,18 | 0,18 | 2,8E-01 |
| Q8BXR9 | Oxysterol-binding protein-related protein 6 OS=Mus musculus OX= | 5 | 4 | 4 | 3 | 959 | 108,9 | 7,27 | 4,4 | 4 | 0,972 | 1,145 | 1,577 | 1,101 | 1,049 | 0,946 | 0,986 | 1,15 | 1,16 | 0,16 | 2,8E-01 |
| Q9R1J0 | Sterol-4-alpha-carboxylate 3-dehydrogenase, decarboxylating OS=N | 2 | 1 | 1 | 1 | 362 | 40,7 | 7,85 | 1,77 | 1 | 1,44 | 1,249 | 1,031 | 1,004 | 1,037 | 1,104 | 0,957 | 1,108 | 1,12 | 0,12 | 2,8E-01 |
| Q61655 | ATP-dependent RNA helicase DDX19A OS=Mus musculus OX=1009 | 12 | 5 | 5 | 5 | 478 | 53,9 | 6,67 | 7,67 | 5 | 0,958 | 0,902 | 0,999 | 1,016 | 0,956 | 0,96 | 0,874 | 0,931 | 1,04 | 0,04 | 2,8E-01 |
| Q91VT4 | Carbonyl reductase family member 4 OS=Mus musculus OX=10090 | 18 | 4 | 5 | 4 | 236 | 25,4 | 9,69 | 10,89 | 4 | 1,002 | 0,91 | 0,982 | 0,976 | 0,969 | 1,109 | 0,971 | 1,005 | 0,95 | -0,05 | 2,8E-01 |
| Q6NZJ6 | Eukaryotic translation initiation factor 4 gamma 1 OS=Mus muscul | 4 | 5 | 5 | 2 | 1600 | 176 | 5,4 | 8,74 | 5 | 1,106 | 1,029 | 0,986 | 0,997 | 1,059 | 0,995 | 0,965 | 0,875 | 1,06 | 0,06 | 2,8E-01 |
| Q8C996 | Transmembrane protein 163 OS=Mus musculus OX=10090 GN=Tm | 8 | 2 | 2 | 2 | 288 | 31,2 | 7,93 | 5,77 | 2 | 0,893 | 1,116 | 1,095 | 1,039 | 0,844 | 0,984 | 1,02 | 0,99 | 1,08 | 0,08 | 2,8E-01 |
| O88910 | MAGUK p55 subfamily member 3 OS=Mus musculus OX=10090 GN | 13 | 6 | 7 | 6 | 568 | 64,4 | 6,34 | 10,2 | 6 | 0,852 | 0,928 | 0,837 | 1,043 | 0,992 | 0,961 | 0,945 | 0,994 | 0,94 | -0,06 | 2,8E-01 |
| Q9CQX2 | Cytochrome b5 type B OS=Mus musculus OX=10090 GN=Cyb5b PE= | 18 | 3 | 7 | 3 | 146 | 16,3 | 4,89 | 14,93 | 3 | 1,016 | 0,884 | 0,944 | 0,821 | 0,937 | 0,959 | 1,018 | 0,966 | 0,94 | -0,06 | 2,8E-01 |
| O35658 | Complement component 1 Q subcomponent-binding protein, mito | 10 | 2 | 2 | 2 | 278 | 31 | 4,92 | 3,21 | 2 | 1,483 | 1,469 | 1,05 | 0,873 | 0,86 | 1,178 | 0,747 | 1,187 | 1,23 | 0,23 | 2,8E-01 |
| Q8BJI1 | Sodium-dependent neutral amino acid transporter SLC6A17 OS=Mu | 8 | 4 | 10 | 4 | 727 | 81 | 6,23 | 20,69 | 4 | 1,067 | 0,937 | 1,04 | 0,937 | 0,888 | 0,963 | 1,017 | 0,895 | 1,06 | 0,06 | 2,8E-01 |
| Q8BGB7 | Enolase-phosphatase E1 OS=Mus musculus OX=10090 GN=Enoph1 | 5 | 1 | 1 | 1 | 257 | 28,6 | 4,92 | 2,79 | 1 | 1,537 | 1,197 | 1,194 | 1,022 | 0,835 | 1,267 | 0,988 | 1,172 | 1,16 | 0,16 | 2,8E-01 |
| Q9DCJ5 | NADH dehydrogenase [ubiquinone] 1 alpha subcomplex subunit 8 C | 26 | 4 | 7 | 4 | 172 | 20 | 8,46 | 7,93 | 4 | 0,89 | 1,095 | 1,187 | 0,945 | 1,233 | 1,112 | 1,142 | 1,017 | 0,91 | -0,09 | 2,8E-01 |
| Q8K212 | Phosphofurin acidic cluster sorting protein 1 OS=Mus musculus OX= | 17 | 14 | 22 | 14 | 961 | 104,8 | 7,74 | 39,56 | 14 | 1,031 | 1,01 | 0,946 | 0,991 | 0,934 | 0,949 | 1,025 | 0,936 | 1,03 | 0,03 | 2,8E-01 |
| Q9QXX4 | Calcium-binding mitochondrial carrier protein Aralar2 OS=Mus mus | 5 | 3 | 5 | 1 | 676 | 74,4 | 8,6 | 9,59 | 3 | 1,3 | 1,077 | 1,132 | 0,901 | 1,245 | 2,911 | 1,289 | 1,054 | 0,68 | -0,32 | 2,8E-01 |
| P35700 | Peroxiredoxin-1 OS=Mus musculus OX=10090 GN=Prdx1 PE=1 SV=1 | 56 | 11 | 24 | 9 | 199 | 22,2 | 8,12 | 57,81 | 11 | 0,854 | 0,916 | 0,933 | 0,951 | 0,983 | 0,884 | 0,975 | 0,959 | 0,96 | -0,04 | 2,8E-01 |

| | | | | | | | | | | | | | | | | | | | | | |
|---|---|---|---|---|---|---|---|---|---|---|---|---|---|---|---|---|---|---|---|---|---|
| P09055 | Integrin beta-1 OS=Mus musculus OX=10090 GN=Itgb1 PE=1 SV=1 | 10 | 7 | 7 | 7 | 798 | 88,2 | 5,94 | 9,72 | 7 | 1,328 | 1,267 | 1,162 | 0,929 | 0,881 | 1,196 | 0,97 | 1,106 | 1,13 | 0,13 | 2,8E-01 |
| Q8CHG7 | Rap guanine nucleotide exchange factor 2 OS=Mus musculus OX=1( | 10 | 13 | 15 | 13 | 1496 | 166,3 | 6,61 | 25,68 | 13 | 1,082 | 1,057 | 1,048 | 1,025 | 0,939 | 1,081 | 1,029 | 1,013 | 1,04 | 0,04 | 2,8E-01 |
| Q99L43 | Phosphatidate cytidylyltransferase 2 OS=Mus musculus OX=10090 ( | 5 | 2 | 2 | 2 | 444 | 51,3 | 7,05 | 3,84 | 2 | 0,955 | 1,491 | 1,491 | 1,585 | 1,179 | 1,705 | 1,792 | 1,839 | 0,85 | -0,15 | 2,8E-01 |
| Q01279 | Epidermal growth factor receptor OS=Mus musculus OX=10090 GN | 2 | 2 | 2 | 2 | 1210 | 134,8 | 6,86 | 0 | 2 | 2,967 | 2,329 | 1,385 | 0,909 | 0,822 | 1,526 | 1,265 | 1,625 | 1,45 | 0,45 | 2,8E-01 |
| Q8VBW6 | NEDD8-activating enzyme E1 regulatory subunit OS=Mus musculus | 2 | 1 | 1 | 1 | 534 | 60,2 | 5,52 | 2,1 | 1 | 0,429 | 1,069 | 1,052 | 1,114 | 1,352 | 1,285 | 0,782 | 1,229 | 0,79 | -0,21 | 2,8E-01 |
| Q9D853 | EEF1A lysine methyltransferase 2 OS=Mus musculus OX=10090 GN= | 5 | 1 | 1 | 1 | 244 | 26,8 | 5,15 | 2,5 | 1 | 0,01 | 1,376 | 1,278 | 1,2 | 1,292 | 1,049 | 1,928 | 1,349 | 0,69 | -0,31 | 2,8E-01 |
| Q99J39 | Malonyl-CoA decarboxylase, mitochondrial OS=Mus musculus OX=1 | 9 | 3 | 3 | 3 | 492 | 54,7 | 8,94 | 7,78 | 3 | 1,286 | 1,298 | 1,095 | 0,997 | 1,007 | 1,123 | 0,994 | 1,154 | 1,09 | 0,09 | 2,8E-01 |
| Q3TWL2 | Type 1 phosphatidylinositol 4,5-bisphosphate 4-phosphatase OS=M | 4 | 1 | 1 | 1 | 284 | 30 | 8,82 | 3,35 | 1 | 0,859 | 1,082 | 1,182 | 1,213 | 1,002 | 1,163 | 1,293 | 1,515 | 0,87 | -0,13 | 2,8E-01 |
| O89112 | Glutathione S-transferase LANCL1 OS=Mus musculus OX=10090 GN | 15 | 6 | 10 | 6 | 399 | 45,3 | 7,77 | 23,41 | 6 | 0,925 | 1,047 | 1,091 | 1,132 | 1,156 | 1,065 | 1,125 | 1,082 | 0,95 | -0,05 | 2,8E-01 |
| Q6PDS3 | Sterile alpha and TIR motif-containing protein 1 OS=Mus musculus | 4 | 2 | 2 | 2 | 724 | 79,6 | 6,27 | 1,61 | 2 | 1,253 | 1,29 | 0,982 | 0,851 | 0,755 | 1,076 | 0,965 | 0,985 | 1,16 | 0,16 | 2,8E-01 |
| Q9CWG8 | Protein arginine methyltransferase NDUFAF7, mitochondrial OS=M | 10 | 4 | 4 | 4 | 436 | 48,4 | 6,95 | 7,71 | 4 | 1,056 | 1,149 | 1,097 | 0,939 | 1,062 | 1,135 | 1,187 | 1,102 | 0,95 | -0,05 | 2,8E-01 |
| Q922C1 | Uncharacterized protein C19orf44 homolog OS=Mus musculus OX= | 3 | 1 | 1 | 1 | 641 | 69,2 | 5,85 | 0 | 1 | 1,428 | 1,446 | 1,093 | 1,024 | 0,867 | 1,319 | 1,002 | 1,115 | 1,16 | 0,16 | 2,8E-01 |
| Q62083 | PRKCA-binding protein OS=Mus musculus OX=10090 GN=Pick1 PE= | 8 | 3 | 3 | 3 | 416 | 46,6 | 5,52 | 3,87 | 3 | 1,062 | 0,786 | 1,185 | 1,274 | 1,009 | 1,153 | 1,395 | 1,448 | 0,86 | -0,14 | 2,8E-01 |
| P35802 | Neuronal membrane glycoprotein M6-a OS=Mus musculus OX=100 | 19 | 8 | 21 | 7 | 278 | 31,1 | 5,27 | 45,16 | 8 | 0,985 | 1,48 | 0,987 | 1,031 | 0,948 | 1,071 | 1,014 | 0,835 | 1,16 | 0,16 | 2,8E-01 |
| Q9CQS8 | Protein transport protein Sec61 subunit beta OS=Mus musculus OX | 11 | 1 | 1 | 1 | 96 | 10 | 11,56 | 1,89 | 1 | 0,981 | 1,149 | 1,258 | 1,118 | 1,457 | 1,39 | 0,977 | 1,248 | 0,89 | -0,11 | 2,8E-01 |
| O08917 | Flotillin-1 OS=Mus musculus OX=10090 GN=Flot1 PE=1 SV=1 | 47 | 18 | 30 | 18 | 428 | 47,5 | 7,15 | 69,09 | 18 | 1,229 | 1,142 | 1,014 | 0,995 | 0,866 | 1,09 | 1,027 | 1,05 | 1,09 | 0,09 | 2,9E-01 |
| P62983 | Ubiquitin-40S ribosomal protein S27a OS=Mus musculus OX=10090 | 30 | 5 | 37 | 5 | 156 | 17,9 | 9,64 | 38,82 | 5 | 0,997 | 1,051 | 1,083 | 0,995 | 0,961 | 0,918 | 1,055 | 1,018 | 1,04 | 0,04 | 2,9E-01 |
| Q8BRT1 | CLIP-associating protein 2 OS=Mus musculus OX=10090 GN=Clasp2 | 13 | 14 | 17 | 13 | 1286 | 140,7 | 8,63 | 18,46 | 14 | 1,028 | 1,128 | 0,989 | 0,965 | 0,92 | 0,97 | 0,992 | 1,029 | 1,05 | 0,05 | 2,9E-01 |
| Q925T6 | Glutamate receptor-interacting protein 1 OS=Mus musculus OX=10 | 4 | 4 | 4 | 4 | 1127 | 122 | 6,55 | 3,3 | 4 | 1,628 | 1,433 | 1,532 | 1,32 | 1,25 | 1,49 | 1,393 | 1,394 | 1,07 | 0,07 | 2,9E-01 |
| Q63959 | Potassium voltage-gated channel subfamily C member 3 OS=Mus m | 2 | 1 | 1 | 1 | 769 | 82,1 | 7,09 | 2,83 | 1 | 1,381 | 1,23 | 0,864 | 0,926 | 0,851 | 1,046 | 0,906 | 0,988 | 1,16 | 0,16 | 2,9E-01 |
| Q5FWK3 | Rho GTPase-activating protein 1 OS=Mus musculus OX=10090 GN= | 19 | 7 | 10 | 7 | 439 | 50,4 | 6,44 | 23,16 | 7 | 1,22 | 1,095 | 1,028 | 1,101 | 0,942 | 1,138 | 0,993 | 1,091 | 1,07 | 0,07 | 2,9E-01 |
| Q6ZQ29-2 | Isoform 2 of Serine/threonine-protein kinase TAO2 OS=Mus muscu | 13 | 11 | 11 | 10 | 1055 | 119,9 | 7,28 | 17,45 | 11 | 0,901 | 0,9 | 1,086 | 1,192 | 1,055 | 1,088 | 1,172 | 1,12 | 0,92 | -0,08 | 2,9E-01 |
| P10649 | Glutathione S-transferase Mu 1 OS=Mus musculus OX=10090 GN=( | 52 | 14 | 25 | 12 | 218 | 26 | 7,94 | 55,81 | 14 | 0,975 | 1,014 | 0,928 | 0,938 | 1,008 | 0,983 | 0,952 | 1,035 | 0,97 | -0,03 | 2,9E-01 |
| Q80X80 | Phospholipid transfer protein C2CD2L OS=Mus musculus OX=10090 | 6 | 4 | 4 | 4 | 706 | 76,3 | 7,21 | 9,36 | 4 | 1,16 | 0,946 | 0,893 | 1,076 | 0,961 | 1,236 | 1,187 | 1,091 | 0,91 | -0,09 | 2,9E-01 |
| Q91YI0 | Argininosuccinate lyase OS=Mus musculus OX=10090 GN=Asl PE=1 | 16 | 7 | 8 | 7 | 464 | 51,7 | 6,96 | 14,59 | 7 | 1,031 | 0,912 | 0,909 | 1,046 | 1,015 | 0,933 | 1,062 | 1,203 | 0,93 | -0,07 | 2,9E-01 |
| Q5SQX6 | Cytoplasmic FMR1-interacting protein 2 OS=Mus musculus OX=100 | 13 | 17 | 24 | 13 | 1253 | 145,6 | 7,05 | 30,77 | 17 | 1,03 | 1,044 | 1,064 | 1,007 | 1,015 | 1,089 | 1,064 | 1,069 | 0,98 | -0,02 | 2,9E-01 |
| P24529 | Tyrosine 3-monooxygenase OS=Mus musculus OX=10090 GN=Th PI | 11 | 5 | 5 | 5 | 498 | 56 | 6,15 | 6,02 | 5 | 1,127 | 0,647 | 0,785 | 0,82 | 0,883 | 1,08 | 1,511 | 0,788 | 0,79 | -0,21 | 2,9E-01 |
| Q61578 | NADPH:adrenodoxin oxidoreductase, mitochondrial OS=Mus muscu | 4 | 2 | 2 | 2 | 494 | 54,2 | 8,66 | 0 | 2 | 0,7 | 0,896 | 0,783 | 1,082 | 0,898 | 0,965 | 1,271 | 0,894 | 0,86 | -0,14 | 2,9E-01 |
| Q5F2E8 | Serine/threonine-protein kinase TAO1 OS=Mus musculus OX=1009( | 15 | 13 | 16 | 12 | 1001 | 116 | 7,55 | 30,7 | 13 | 1,093 | 1,174 | 1,06 | 1,058 | 0,929 | 1,134 | 1,058 | 1,03 | 1,06 | 0,06 | 2,9E-01 |
| Q62093 | Serine/arginine-rich splicing factor 2 OS=Mus musculus OX=10090 ( | 3 | 1 | 1 | 1 | 221 | 25,5 | 11,85 | 2,19 | 1 | 0,799 | 0,826 | 0,87 | 0,665 | 1,099 | 0,71 | 0,782 | 1,076 | 0,86 | -0,14 | 2,9E-01 |
| Q99JP7 | Glutathione hydrolase 7 OS=Mus musculus OX=10090 GN=Ggt7 PE= | 13 | 7 | 9 | 7 | 662 | 70,2 | 5,06 | 16,41 | 7 | 1,041 | 1,15 | 0,959 | 1,064 | 0,977 | 1,031 | 0,857 | 1,068 | 1,07 | 0,07 | 2,9E-01 |
| P36993 | Protein phosphatase 1B OS=Mus musculus OX=10090 GN=Ppm1b P | 16 | 5 | 6 | 4 | 390 | 42,8 | 5,19 | 12,9 | 5 | 1,03 | 1,085 | 0,888 | 0,876 | 0,885 | 0,956 | 0,934 | 0,83 | 1,08 | 0,08 | 2,9E-01 |
| Q6PEE2 | CBP80/20-dependent translation initiation factor OS=Mus musculu | 3 | 1 | 1 | 1 | 600 | 67,8 | 6,65 | 0 | 1 | 0,01 | 1,907 | 1,664 | 1,907 | 1,639 | 2,361 | 2,01 | 1,732 | 0,71 | -0,29 | 2,9E-01 |
| Q9DCT8 | Cysteine-rich protein 2 OS=Mus musculus OX=10090 GN=Crip2 PE= | 25 | 4 | 5 | 4 | 208 | 22,7 | 8,63 | 14,01 | 4 | 0,99 | 0,965 | 0,992 | 0,986 | 1,007 | 0,978 | 0,981 | 1,068 | 0,97 | -0,03 | 2,9E-01 |
| Q9WUM3 | Coronin-1B OS=Mus musculus OX=10090 GN=Coro1b PE=1 SV=1 | 17 | 6 | 6 | 6 | 484 | 53,9 | 5,78 | 13,92 | 6 | 1,179 | 1,235 | 1,097 | 1,038 | 0,904 | 1,18 | 1,045 | 1,086 | 1,08 | 0,08 | 2,9E-01 |
| O35680 | 28S ribosomal protein S12, mitochondrial OS=Mus musculus OX=1( | 8 | 1 | 1 | 1 | 139 | 15,4 | 10,71 | 0 | 1 | 0,707 | 0,642 | 0,793 | 0,768 | 0,941 | 0,791 | 0,599 | 1,092 | 0,85 | -0,15 | 2,9E-01 |
| P25444 | 40S ribosomal protein S2 OS=Mus musculus OX=10090 GN=Rps2 PI | 18 | 5 | 6 | 5 | 293 | 31,2 | 10,24 | 10,52 | 5 | 0,918 | 1,034 | 0,995 | 1,026 | 1,019 | 0,838 | 0,856 | 1,006 | 1,07 | 0,07 | 2,9E-01 |
| O08919 | Numb-like protein OS=Mus musculus OX=10090 GN=Numbl PE=1 S | 10 | 6 | 9 | 3 | 604 | 64,1 | 8,82 | 18,16 | 6 | 0,789 | 0,662 | 0,93 | 0,911 | 0,875 | 1,07 | 0,893 | 0,831 | 0,90 | -0,10 | 2,9E-01 |
| Q6AXF6 | SID1 transmembrane family member 1 OS=Mus musculus OX=1009 | 1 | 1 | 1 | 1 | 827 | 93,8 | 6,87 | 2,55 | 1 | 1,299 | 1,262 | 1,118 | 1,043 | 1,14 | 0,971 | 1,228 | 0,98 | 1,09 | 0,09 | 2,9E-01 |
| P46935 | E3 ubiquitin-protein ligase NEDD4 OS=Mus musculus OX=10090 GN | 9 | 8 | 8 | 7 | 887 | 102,6 | 5,26 | 18,53 | 8 | 1,186 | 1,191 | 1,071 | 1,082 | 0,944 | 1,135 | 1,108 | 1,094 | 1,06 | 0,06 | 2,9E-01 |
| Q80U40 | RIMS-binding protein 2 OS=Mus musculus OX=10090 GN=Rimbp2 P | 10 | 10 | 10 | 10 | 1072 | 118,3 | 5,35 | 21,86 | 10 | 1,029 | 1,021 | 1,042 | 1,03 | 0,998 | 1,122 | 1,052 | 1,07 | 0,97 | -0,03 | 2,9E-01 |
| A2AKG8 | Focadhesin OS=Mus musculus OX=10090 GN=Focad PE=1 SV=1 | 1 | 1 | 1 | 1 | 1798 | 198,8 | 6,74 | 0 | 1 | 0,888 | 0,53 | 1,168 | 0,539 | 0,663 | 0,732 | 0,424 | 0,528 | 1,33 | 0,33 | 2,9E-01 |
| Q9JIF7 | Coatomer subunit beta OS=Mus musculus OX=10090 GN=Copb1 PE | 8 | 6 | 6 | 6 | 953 | 107 | 6 | 14,07 | 6 | 1,229 | 1,168 | 1,163 | 1,118 | 0,997 | 1,29 | 1,042 | 1,015 | 1,08 | 0,08 | 2,9E-01 |
| Q5SX40 | Myosin-1 OS=Mus musculus OX=10090 GN=Myh1 PE=1 SV=1 | 0 | 1 | 2 | 1 | 1942 | 223,2 | 5,76 | 4,6 | 1 | 1,662 | 1,136 | 1,242 | 0,81 | 0,686 | 1,295 | 0,878 | 0,989 | 1,26 | 0,26 | 2,9E-01 |
| P63321 | Ras-related protein Ral-A OS=Mus musculus OX=10090 GN=Rala PE | 28 | 7 | 12 | 6 | 206 | 23,5 | 7,11 | 30,43 | 7 | 1,132 | 0,909 | 0,943 | 1,019 | 0,96 | 1,08 | 1,125 | 1,13 | 0,93 | -0,07 | 2,9E-01 |
| Q9QZD8 | Mitochondrial dicarboxylate carrier OS=Mus musculus OX=10090 G | 8 | 2 | 2 | 2 | 287 | 31,7 | 9,32 | 5,05 | 2 | 1,166 | 0,599 | 0,879 | 0,677 | 0,54 | 0,464 | 0,886 | 0,705 | 1,28 | 0,28 | 2,9E-01 |
| Q4KML4 | Costars family protein ABRACL OS=Mus musculus OX=10090 GN=Al | 11 | 1 | 1 | 1 | 81 | 9 | 5,8 | 2,3 | 1 | 1,368 | 1,229 | 1,074 | 0,943 | 0,844 | 1,2 | 1,022 | 1,006 | 1,13 | 0,13 | 2,9E-01 |
| Q8CHP8 | Glycerol-3-phosphate phosphatase OS=Mus musculus OX=10090 G | 24 | 6 | 8 | 6 | 321 | 34,5 | 5,35 | 18,29 | 6 | 1,398 | 1,093 | 1,17 | 0,968 | 1,019 | 1,146 | 0,91 | 1,078 | 1,11 | 0,11 | 2,9E-01 |
| Q9CYR0 | Single-stranded DNA-binding protein, mitochondrial OS=Mus musc | 5 | 1 | 1 | 1 | 152 | 17,3 | 9,92 | 2,43 | 1 | 0,818 | 1,101 | 0,928 | 0,994 | 1,243 | 1,232 | 0,942 | 0,917 | 0,89 | -0,11 | 2,9E-01 |
| Q9Z0N1 | Eukaryotic translation initiation factor 2 subunit 3, X-linked OS=Mu | 12 | 4 | 4 | 4 | 472 | 51 | 8,4 | 2,12 | 4 | 0,947 | 0,942 | 1,169 | 1,061 | 0,959 | 1,371 | 1,273 | 1,03 | 0,89 | -0,11 | 2,9E-01 |
| Q8BGH4 | Receptor expression-enhancing protein 1 OS=Mus musculus OX=1( | 6 | 2 | 2 | 2 | 201 | 22,3 | 9,5 | 0 | 2 | 0,722 | 0,968 | 1,223 | 1,154 | 1,156 | 1,661 | 1,06 | 1,038 | 0,83 | -0,17 | 2,9E-01 |
| Q9CR39 | WD repeat domain phosphoinositide-interacting protein 3 OS=Mus | 9 | 2 | 2 | 2 | 344 | 38 | 7,56 | 7,16 | 2 | 1,169 | 1,411 | 1,126 | 1,214 | 1,037 | 1,12 | 1,071 | 1,298 | 1,09 | 0,09 | 2,9E-01 |
| D3YZU1 | SH3 and multiple ankyrin repeat domains protein 1 OS=Mus muscu | 10 | 18 | 21 | 18 | 2167 | 226,2 | 8,34 | 28,22 | 18 | 1,119 | 1,09 | 1,015 | 0,915 | 0,985 | 1,017 | 0,936 | 0,978 | 1,06 | 0,06 | 2,9E-01 |
| Q8VE19 | GATOR complex protein MIOS OS=Mus musculus OX=10090 GN=M | 1 | 1 | 1 | 1 | 875 | 98,3 | 6,71 | 1,91 | 1 | 0,733 | 0,874 | 0,962 | 0,929 | 0,857 | 0,604 | 0,954 | 0,62 | 1,15 | 0,15 | 2,9E-01 |
| Q9CYA0 | Protein disulfide isomerase Creld2 OS=Mus musculus OX=10090 GN | 7 | 2 | 3 | 2 | 350 | 38,2 | 4,58 | 5,57 | 2 | 0,852 | 0,954 | 0,922 | 0,826 | 1,001 | 0,825 | 0,929 | 1,095 | 0,92 | -0,08 | 2,9E-01 |
| P97351 | 40S ribosomal protein S3a OS=Mus musculus OX=10090 GN=Rps3a | 17 | 4 | 4 | 4 | 264 | 29,9 | 9,73 | 10,76 | 4 | 1,278 | 1,096 | 0,98 | 1,024 | 0,977 | 0,949 | 1,035 | 1,085 | 1,08 | 0,08 | 2,9E-01 |
| P14106 | Complement C1q subcomponent subunit B OS=Mus musculus OX=1 | 11 | 2 | 2 | 2 | 253 | 26,7 | 8,15 | 4,78 | 2 | 1,139 | 1,073 | 0,92 | 0,766 | 0,824 | 1,072 | 0,643 | 0,801 | 1,17 | 0,17 | 3,0E-01 |
| P13595 | Neural cell adhesion molecule 1 OS=Mus musculus OX=10090 GN=I | 35 | 35 | 56 | 10 | 1115 | 119,4 | 4,83 | 129,27 | 35 | 0,77 | 0,845 | 0,907 | 1,05 | 1,014 | 0,862 | 1,047 | 0,978 | 0,92 | -0,08 | 3,0E-01 |
| Q8VEM8 | Phosphate carrier protein, mitochondrial OS=Mus musculus OX=10 | 16 | 5 | 13 | 5 | 357 | 39,6 | 9,26 | 31,27 | 5 | 0,945 | 1,135 | 1,02 | 1,005 | 0,936 | 0,978 | 1,013 | 0,982 | 1,05 | 0,05 | 3,0E-01 |
| Q9CXI5 | Mesencephalic astrocyte-derived neurotrophic factor OS=Mus mus | 12 | 2 | 2 | 2 | 179 | 20,4 | 8,07 | 5,78 | 2 | 1,253 | 1,076 | 1,214 | 1,055 | 1,091 | 1,153 | 0,982 | 1,093 | 1,06 | 0,06 | 3,0E-01 |
| Q9QX60 | Deoxyguanosine kinase, mitochondrial OS=Mus musculus OX=1009 | 8 | 2 | 2 | 2 | 277 | 32,3 | 7,66 | 4,91 | 2 | 1,325 | 1,215 | 1,188 | 1,155 | 0,83 | 1,339 | 0,781 | 1,255 | 1,16 | 0,16 | 3,0E-01 |
| Q9DCT2 | NADH dehydrogenase [ubiquinone] iron-sulfur protein 3, mitochon | 31 | 10 | 18 | 10 | 263 | 30,1 | 7,17 | 51,67 | 10 | 1,364 | 1,239 | 1,243 | 1,187 | 1,044 | 1,327 | 1,045 | 1,246 | 1,08 | 0,08 | 3,0E-01 |

| | | | | | | | | | | | | | | | | | | | | | |
|---|---|---|---|---|---|---|---|---|---|---|---|---|---|---|---|---|---|---|---|---|---|
| P33173 | Kinesin-like protein KIF1A OS=Mus musculus OX=10090 GN=Kif1a P | 9 | 15 | 16 | 8 | 1695 | 191,6 | 6,2 | 25,8 | 15 | 1,095 | 1,145 | 0,9 | 1,023 | 0,946 | 1,061 | 0,849 | 0,99 | 1,08 | 0,08 | 3,0E-01 |
| Q9ERD7 | Tubulin beta-3 chain OS=Mus musculus OX=10090 GN=Tubb3 PE=1 | 39 | 20 | 109 | 3 | 450 | 50,4 | 4,93 | 236,02 | 20 | 0,854 | 0,991 | 0,788 | 0,956 | 0,936 | 0,901 | 1,064 | 0,955 | 0,93 | -0,07 | 3,0E-01 |
| Q8K4Q0 | Regulatory-associated protein of mTOR OS=Mus musculus OX=100 | 3 | 3 | 3 | 3 | 1335 | 149,4 | 6,87 | 5,03 | 3 | 1,146 | 0,949 | 1,087 | 1,19 | 0,79 | 0,761 | 1,332 | 0,824 | 1,18 | 0,18 | 3,0E-01 |
| Q8CC21 | Tetratricopeptide repeat protein 19, mitochondrial OS=Mus muscu | 8 | 3 | 3 | 3 | 365 | 41,2 | 6,21 | 5,61 | 3 | 1,02 | 0,941 | 0,977 | 2,798 | 0,764 | 0,993 | 0,888 | 0,998 | 1,57 | 0,57 | 3,0E-01 |
| Q7TQK5 | Coiled-coil domain-containing protein 93 OS=Mus musculus OX=10 | 6 | 3 | 3 | 3 | 629 | 72,6 | 8,29 | 4,45 | 3 | 1,623 | 1,11 | 1,191 | 1,505 | 1,22 | 1,358 | 1,227 | 0,923 | 1,15 | 0,15 | 3,0E-01 |
| Q8R0Y6 | Cytosolic 10-formyltetrahydrofolate dehydrogenase OS=Mus musc | 24 | 21 | 27 | 21 | 902 | 98,6 | 5,91 | 59,09 | 21 | 1,293 | 1,191 | 1,165 | 1,054 | 0,998 | 1,224 | 1,024 | 1,131 | 1,07 | 0,07 | 3,0E-01 |
| Q922U2 | Keratin, type II cytoskeletal 5 OS=Mus musculus OX=10090 GN=Krt | 25 | 20 | 30 | 10 | 580 | 61,7 | 7,75 | 51,51 | 20 | 1,358 | 1,752 | 0,997 | 0,652 | 1,143 | 1,052 | 0,55 | 0,772 | 1,35 | 0,35 | 3,0E-01 |
| P42669 | Transcriptional activator protein Pur-alpha OS=Mus musculus OX=1 | 19 | 6 | 6 | 6 | 321 | 34,9 | 6,44 | 12,25 | 6 | 0,918 | 0,856 | 0,963 | 0,983 | 1,077 | 0,942 | 0,941 | 0,957 | 0,95 | -0,05 | 3,0E-01 |
| Q02357-6 | Isoform 6 of Ankyrin-1 OS=Mus musculus OX=10090 GN=Ank1 | 11 | 17 | 20 | 15 | 1907 | 209,4 | 6,74 | 39,32 | 17 | 1,195 | 1,064 | 1,028 | 1,028 | 0,984 | 1,041 | 0,998 | 1,084 | 1,05 | 0,05 | 3,0E-01 |
| P08414 | Calcium/calmodulin-dependent protein kinase type IV OS=Mus mu | 12 | 5 | 7 | 5 | 469 | 52,6 | 4,93 | 12,52 | 5 | 1,338 | 1,109 | 1,193 | 1,134 | 1,048 | 1,116 | 1,143 | 1,195 | 1,06 | 0,06 | 3,0E-01 |
| Q2NL51 | Glycogen synthase kinase-3 alpha OS=Mus musculus OX=10090 GN | 20 | 7 | 11 | 3 | 490 | 51,6 | 8,81 | 23,09 | 7 | 1,178 | 1,149 | 1,141 | 1,123 | 1,069 | 1,196 | 1,126 | 1,019 | 1,04 | 0,04 | 3,0E-01 |
| Q6PER3 | Microtubule-associated protein RP/EB family member 3 OS=Mus m | 29 | 9 | 14 | 7 | 281 | 31,9 | 5,54 | 38,09 | 9 | 1,133 | 1,127 | 0,955 | 1,038 | 0,948 | 1,097 | 0,923 | 1,023 | 1,07 | 0,07 | 3,0E-01 |
| Q61249 | Immunoglobulin-binding protein 1 OS=Mus musculus OX=10090 GI | 5 | 1 | 1 | 1 | 340 | 38,9 | 6,18 | 0 | 1 | 1,71 | 0,884 | 1,733 | 1,217 | 0,989 | 1,386 | 0,991 | 1,153 | 1,23 | 0,23 | 3,0E-01 |
| Q3TDN2 | FAS-associated factor 2 OS=Mus musculus OX=10090 GN=Faf2 PE= | 7 | 2 | 2 | 2 | 445 | 52,4 | 5,47 | 3,22 | 2 | 1,628 | 1,427 | 1,319 | 1,031 | 0,812 | 1,504 | 0,987 | 1,218 | 1,20 | 0,20 | 3,0E-01 |
| O88844 | Isocitrate dehydrogenase [NADP] cytoplasmic OS=Mus musculus O | 29 | 14 | 18 | 13 | 414 | 46,6 | 7,17 | 46,79 | 14 | 0,971 | 0,941 | 1,009 | 1,006 | 1,008 | 1,043 | 1,001 | 0,973 | 0,98 | -0,02 | 3,0E-01 |
| Q01405 | Protein transport protein Sec23A OS=Mus musculus OX=10090 GN | 18 | 11 | 14 | 11 | 765 | 86,1 | 7,08 | 29,52 | 11 | 1,18 | 1,031 | 1,177 | 0,989 | 0,984 | 1,103 | 0,987 | 1,045 | 1,06 | 0,06 | 3,0E-01 |
| Q8BYY4 | Tetratricopeptide repeat protein 39B OS=Mus musculus OX=10090 | 3 | 1 | 1 | 1 | 617 | 70,2 | 6,64 | 0 | 1 | 0,996 | 0,775 | 1,002 | 1,105 | 1,095 | 1,186 | 0,826 | 1,383 | 0,86 | -0,14 | 3,0E-01 |
| Q8BH27 | Multiple epidermal growth factor-like domains protein 9 OS=Mus n | 4 | 2 | 3 | 2 | 600 | 62,8 | 5,47 | 4,18 | 2 | 0,957 | 0,717 | 1,046 | 0,94 | 1,137 | 0,92 | 1,035 | 0,953 | 0,90 | -0,10 | 3,0E-01 |
| Q9Z2H2 | Regulator of G-protein signaling 6 OS=Mus musculus OX=10090 GN | 34 | 14 | 24 | 10 | 472 | 54,5 | 7,42 | 52,54 | 14 | 1,17 | 1,128 | 1,164 | 1,156 | 1,03 | 1,235 | 0,952 | 1,122 | 1,06 | 0,06 | 3,0E-01 |
| P27546 | Microtubule-associated protein 4 OS=Mus musculus OX=10090 GN | 12 | 10 | 14 | 7 | 1125 | 117,4 | 4,98 | 23,99 | 10 | 1,005 | 0,879 | 0,963 | 0,898 | 0,847 | 0,927 | 0,94 | 0,863 | 1,05 | 0,05 | 3,0E-01 |
| B2RQC6 | CAD protein OS=Mus musculus OX=10090 GN=Cad PE=1 SV=1 | 3 | 5 | 5 | 5 | 2225 | 243,1 | 6,43 | 2,59 | 5 | 1,167 | 0,919 | 1,034 | 0,842 | 0,955 | 0,845 | 0,935 | 0,888 | 1,09 | 0,09 | 3,0E-01 |
| O88712 | C-terminal-binding protein 1 OS=Mus musculus OX=10090 GN=Ctb | 25 | 11 | 15 | 6 | 441 | 47,7 | 6,77 | 27,78 | 11 | 0,951 | 1,102 | 0,995 | 1,011 | 0,978 | 0,981 | 0,996 | 0,956 | 1,04 | 0,04 | 3,0E-01 |
| Q91YD9 | Neural Wiskott-Aldrich syndrome protein OS=Mus musculus OX=10 | 7 | 3 | 3 | 3 | 501 | 54,2 | 7,93 | 1,95 | 3 | 1,706 | 1,258 | 1,396 | 1,047 | 0,907 | 1,33 | 1,195 | 1,229 | 1,16 | 0,16 | 3,0E-01 |
| Q9CR09 | Ubiquitin-fold modifier-conjugating enzyme 1 OS=Mus musculus O | 17 | 3 | 3 | 3 | 167 | 19,5 | 7,4 | 4,87 | 3 | 1,276 | 1,345 | 0,999 | 1,3 | 1,252 | 0,759 | 1,108 | 1,191 | 1,14 | 0,14 | 3,0E-01 |
| Q9QYB5-2 | Isoform 1 of Gamma-adducin OS=Mus musculus OX=10090 GN=Ad | 13 | 8 | 9 | 8 | 674 | 75,4 | 6,34 | 17,44 | 8 | 0,887 | 0,945 | 0,965 | 0,943 | 1,008 | 0,858 | 1,042 | 1,045 | 0,95 | -0,05 | 3,0E-01 |
| P60766-1 | Isoform 1 of Cell division control protein 42 homolog OS=Mus mus | 27 | 5 | 8 | 1 | 191 | 21,3 | 6,04 | 11,56 | 5 | 0,732 | 0,581 | 0,938 | 0,949 | 1,087 | 0,921 | 0,861 | 0,812 | 0,87 | -0,13 | 3,0E-01 |
| Q9R0Y5 | Adenylate kinase isoenzyme 1 OS=Mus musculus OX=10090 GN=Ak | 28 | 5 | 5 | 5 | 194 | 21,5 | 5,81 | 12,8 | 5 | 0,776 | 0,756 | 0,898 | 0,944 | 0,967 | 0,789 | 0,951 | 0,946 | 0,92 | -0,08 | 3,0E-01 |
| Q68EF0 | Rab-3A-interacting protein OS=Mus musculus OX=10090 GN=Rab3i | 2 | 1 | 1 | 1 | 428 | 47,1 | 6,71 | 2,3 | 1 | 1,2 | 1,305 | 1,364 | 1,058 | 1,164 | 0,978 | 1,199 | 1,201 | 1,08 | 0,08 | 3,0E-01 |
| Q9QYB8 | Beta-adducin OS=Mus musculus OX=10090 GN=Add2 PE=1 SV=4 | 25 | 17 | 29 | 16 | 725 | 80,6 | 6,21 | 58,95 | 17 | 0,941 | 0,82 | 0,993 | 0,956 | 0,987 | 0,939 | 1,003 | 0,961 | 0,95 | -0,05 | 3,0E-01 |
| Q6PE13 | Proline-rich transmembrane protein 3 OS=Mus musculus OX=1009 | 8 | 6 | 10 | 6 | 971 | 101,2 | 7,52 | 24,31 | 6 | 1,114 | 0,964 | 0,997 | 1,049 | 0,98 | 1,044 | 0,916 | 0,995 | 1,05 | 0,05 | 3,0E-01 |
| Q8K2T1 | NmrA-like family domain-containing protein 1 OS=Mus musculus O | 15 | 3 | 4 | 3 | 309 | 34,4 | 6,86 | 11,55 | 3 | 1,058 | 0,956 | 0,924 | 1,025 | 1,024 | 0,998 | 0,801 | 0,863 | 1,08 | 0,08 | 3,0E-01 |
| Q9DBS9 | Oxysterol-binding protein-related protein 3 OS=Mus musculus OX= | 5 | 3 | 3 | 2 | 855 | 96,9 | 6,51 | 2,25 | 3 | 0,01 | 1,044 | 0,924 | 0,654 | 0,673 | 0,984 | 1,116 | 0,978 | 0,70 | -0,30 | 3,0E-01 |
| Q8VEK3 | Heterogeneous nuclear ribonucleoprotein U OS=Mus musculus OX | 8 | 6 | 7 | 6 | 800 | 87,9 | 6,24 | 14,88 | 6 | 1,213 | 1,227 | 1,162 | 0,929 | 1,324 | 1,55 | 1,043 | 1,189 | 0,89 | -0,11 | 3,0E-01 |
| Q8C163 | Nuclease EXOG, mitochondrial OS=Mus musculus OX=10090 GN=E | 10 | 2 | 4 | 2 | 368 | 41,4 | 8,12 | 7,82 | 2 | 1,539 | 1,24 | 1,708 | 1,294 | 1,32 | 1,253 | 1,402 | 1,297 | 1,10 | 0,10 | 3,1E-01 |
| Q6PD03 | Serine/threonine-protein phosphatase 2A 56 kDa regulatory subun | 16 | 6 | 8 | 6 | 486 | 56,3 | 7,03 | 13,57 | 6 | 1,003 | 1,093 | 1,016 | 1,072 | 0,88 | 1,075 | 0,921 | 1,065 | 1,06 | 0,06 | 3,1E-01 |
| Q01853 | Transitional endoplasmic reticulum ATPase OS=Mus musculus OX= | 30 | 23 | 33 | 23 | 806 | 89,3 | 5,26 | 79,73 | 23 | 0,888 | 0,925 | 0,936 | 0,988 | 0,958 | 0,894 | 1,04 | 1,018 | 0,96 | -0,04 | 3,1E-01 |
| Q9JJL8 | Serine--tRNA ligase, mitochondrial OS=Mus musculus OX=10090 GI | 6 | 2 | 2 | 2 | 518 | 58,3 | 7,9 | 5,36 | 2 | 0,95 | 0,871 | 0,784 | 0,948 | 1,025 | 0,864 | 0,871 | 1,128 | 0,91 | -0,09 | 3,1E-01 |
| Q8BGD9 | Eukaryotic translation initiation factor 4B OS=Mus musculus OX=10 | 9 | 4 | 8 | 4 | 611 | 68,8 | 5,67 | 9,32 | 4 | 1,451 | 1,178 | 1,176 | 1,142 | 1,067 | 1,141 | 1,163 | 1,224 | 1,08 | 0,08 | 3,1E-01 |
| P14685 | 26S proteasome non-ATPase regulatory subunit 3 OS=Mus musculu | 28 | 14 | 16 | 14 | 530 | 60,7 | 8,44 | 33 | 14 | 0,977 | 0,951 | 0,979 | 1,026 | 1,061 | 0,967 | 1,013 | 1,003 | 0,97 | -0,03 | 3,1E-01 |
| P63137 | Gamma-aminobutyric acid receptor subunit beta-2 OS=Mus muscu | 11 | 5 | 5 | 1 | 512 | 59,2 | 9,31 | 11,55 | 5 | 0,786 | 1,088 | 0,844 | 0,748 | 1,195 | 1,227 | 1,01 | 0,685 | 0,84 | -0,16 | 3,1E-01 |
| Q9EQJ9 | Membrane-associated guanylate kinase, WW and PDZ domain-con | 2 | 3 | 3 | 3 | 1476 | 161,6 | 8,05 | 0 | 3 | 2,334 | 1,948 | 1,218 | 1,446 | 1,52 | 1,644 | 1,412 | 1,161 | 1,21 | 0,21 | 3,1E-01 |
| P60710 | Actin, cytoplasmic 1 OS=Mus musculus OX=10090 GN=Actb PE=1 S | 64 | 25 | 197 | 10 | 375 | 41,7 | 5,48 | 477,23 | 25 | 0,867 | 0,896 | 0,925 | 0,967 | 1,034 | 0,844 | 0,976 | 1,014 | 0,94 | -0,06 | 3,1E-01 |
| O35681 | Synaptotagmin-3 OS=Mus musculus OX=10090 GN=Syt3 PE=1 SV=2 | 8 | 4 | 4 | 4 | 587 | 63,2 | 6,65 | 5,05 | 4 | 0,899 | 0,943 | 0,963 | 1,019 | 0,964 | 0,92 | 1,153 | 1,04 | 0,94 | -0,06 | 3,1E-01 |
| P46460 | Vesicle-fusing ATPase OS=Mus musculus OX=10090 GN=Nsf PE=1 S | 49 | 40 | 92 | 40 | 744 | 82,6 | 6,95 | 230,59 | 40 | 1,019 | 0,996 | 0,978 | 0,997 | 0,999 | 1,024 | 1,018 | 0,997 | 0,99 | -0,01 | 3,1E-01 |
| Q91V92 | ATP-citrate synthase OS=Mus musculus OX=10090 GN=Acly PE=1 S | 27 | 27 | 35 | 27 | 1091 | 119,7 | 7,44 | 65,96 | 27 | 0,942 | 0,954 | 1,036 | 1,023 | 0,999 | 0,971 | 1,127 | 1,043 | 0,96 | -0,04 | 3,1E-01 |
| Q9D3A9 | Protein tweety homolog 1 OS=Mus musculus OX=10090 GN=Ttyh1 | 5 | 2 | 3 | 2 | 450 | 49 | 5,03 | 4,79 | 2 | 1,103 | 0,918 | 0,918 | 1,114 | 1,131 | 0,942 | 1,066 | 1,559 | 0,86 | -0,14 | 3,1E-01 |
| P61028 | Ras-related protein Rab-8B OS=Mus musculus OX=10090 GN=Rab8 | 17 | 5 | 9 | 1 | 207 | 23,6 | 9,07 | 18,43 | 5 | 0,831 | 1,029 | 0,964 | 0,992 | 0,891 | 0,906 | 0,935 | 0,886 | 1,05 | 0,05 | 3,1E-01 |
| Q9QXG4 | Acetyl-coenzyme A synthetase, cytoplasmic OS=Mus musculus OX= | 1 | 1 | 1 | 1 | 701 | 78,8 | 6,64 | 2,11 | 1 | 1,299 | 0,86 | 1,32 | 1,048 | 1,106 | 0,897 | 1,038 | 0,958 | 1,13 | 0,13 | 3,1E-01 |
| Q9Z1D1 | Eukaryotic translation initiation factor 3 subunit G OS=Mus muscul | 4 | 1 | 2 | 1 | 320 | 35,6 | 5,9 | 4,3 | 1 | 0,721 | 0,935 | 1,037 | 0,802 | 0,958 | 0,761 | 1,089 | 1,226 | 0,87 | -0,13 | 3,1E-01 |
| P98203 | Armadillo repeat protein deleted in velo-cardio-facial syndrome ho | 7 | 6 | 7 | 6 | 962 | 105 | 6,64 | 7,48 | 6 | 1,139 | 1,236 | 1,023 | 0,956 | 1,102 | 1,295 | 1,115 | 1,18 | 0,93 | -0,07 | 3,1E-01 |
| P62281 | 40S ribosomal protein S11 OS=Mus musculus OX=10090 GN=Rps11 | 30 | 4 | 4 | 4 | 158 | 18,4 | 10,3 | 11,81 | 4 | 0,988 | 0,921 | 0,999 | 0,904 | 0,959 | 1,03 | 1,03 | 0,941 | 0,96 | -0,04 | 3,1E-01 |
| Q8VIJ6 | Splicing factor, proline- and glutamine-rich OS=Mus musculus OX=1 | 7 | 4 | 5 | 4 | 699 | 75,4 | 9,44 | 10,2 | 4 | 1,259 | 1,21 | 1,284 | 0,871 | 1,437 | 1,694 | 1,052 | 1,198 | 0,86 | -0,14 | 3,1E-01 |
| Q66L42 | Mitogen-activated protein kinase kinase kinase 10 OS=Mus muscul | 2 | 2 | 2 | 2 | 940 | 103,1 | 6,87 | 5,68 | 2 | 0,804 | 1,069 | 0,874 | 0,795 | 0,9 | 1,207 | 0,904 | 0,96 | 0,89 | -0,11 | 3,1E-01 |
| Q8CGK7 | Guanine nucleotide-binding protein G(olf) subunit alpha OS=Mus m | 8 | 3 | 4 | 2 | 381 | 44,3 | 6,65 | 9,7 | 3 | 1,096 | 0,817 | 0,961 | 0,81 | 0,893 | 1,055 | 1,298 | 0,94 | 0,88 | -0,12 | 3,1E-01 |
| Q91Z31 | Polypyrimidine tract-binding protein 2 OS=Mus musculus OX=1009 | 2 | 1 | 1 | 1 | 531 | 57,5 | 8,66 | 0 | 1 | 1,814 | 1,281 | 2,577 | 1,608 | 1,585 | 1,377 | 1,753 | 1,249 | 1,22 | 0,22 | 3,1E-01 |
| P35918 | Vascular endothelial growth factor receptor 2 OS=Mus musculus O | 1 | 1 | 1 | 1 | 1367 | 152,4 | 6,14 | 0 | 1 | 1,5 | 1,355 | 1,248 | 0,976 | 0,527 | 1,497 | 0,775 | 1,205 | 1,27 | 0,27 | 3,1E-01 |
| Q9Z2G6 | Protein sel-1 homolog 1 OS=Mus musculus OX=10090 GN=Sel1l PE | 3 | 2 | 2 | 2 | 790 | 88,3 | 5,57 | 5,65 | 2 | 0,955 | 0,59 | 0,946 | 0,913 | 0,981 | 1,357 | 0,772 | 0,959 | 0,84 | -0,16 | 3,1E-01 |
| Q64737 | Trifunctional purine biosynthetic protein adenosine-3 OS=Mus mus | 9 | 8 | 10 | 8 | 1010 | 107,4 | 6,68 | 19,73 | 8 | 1,203 | 1,094 | 1,371 | 1,125 | 0,98 | 1,06 | 1,251 | 1,129 | 1,08 | 0,08 | 3,1E-01 |
| Q8R5J9 | PRA1 family protein 3 OS=Mus musculus OX=10090 GN=Arl6ip5 PE | 10 | 2 | 3 | 2 | 188 | 21,5 | 9,61 | 7,9 | 2 | 1,019 | 1,121 | 0,867 | 1,005 | 1,015 | 1,089 | 1,065 | 1,085 | 0,94 | -0,06 | 3,1E-01 |
| Q9QYA2 | Mitochondrial import receptor subunit TOM40 homolog OS=Mus m | 21 | 6 | 8 | 6 | 361 | 37,9 | 7,74 | 21,96 | 6 | 1,2 | 1,239 | 1,032 | 0,919 | 0,817 | 1,217 | 0,894 | 0,958 | 1,13 | 0,13 | 3,1E-01 |
| Q6PHZ2-5 | Isoform 4 of Calcium/calmodulin-dependent protein kinase type II | 26 | 14 | 34 | 9 | 512 | 57,7 | 7,15 | 74,63 | 14 | 0,792 | 0,926 | 0,972 | 0,921 | 0,94 | 0,916 | 1,001 | 0,942 | 0,95 | -0,05 | 3,1E-01 |
| P83093 | Stromal interaction molecule 2 OS=Mus musculus OX=10090 GN=S | 6 | 4 | 4 | 4 | 746 | 83,9 | 6,79 | 10,19 | 4 | 1,248 | 0,946 | 0,983 | 0,993 | 0,934 | 1,124 | 0,892 | 0,783 | 1,12 | 0,12 | 3,1E-01 |

| | | | | | | | | | | | | | | | | | | | | | |
|---|---|---|---|---|---|---|---|---|---|---|---|---|---|---|---|---|---|---|---|---|---|
| Q9Z0S1 | 3'(2'),5'-bisphosphate nucleotidase 1 OS=Mus musculus OX=10090 | 35 | 8 | 10 | 8 | 308 | 33,2 | 5,82 | 20,43 | 8 | 1,07 | 1,151 | 1,011 | 1,008 | 0,937 | 1,072 | 0,964 | 1,058 | 1,05 | 0,05 | 3,1E-01 |
| Q6ZQ06 | Centrosomal protein of 162 kDa OS=Mus musculus OX=10090 GN= | 1 | 1 | 2 | 1 | 1403 | 160,8 | 5,53 | 4,71 | 1 | 1,57 | 1,319 | 1,171 | 0,987 | 0,715 | 1,469 | 0,876 | 1,088 | 1,22 | 0,22 | 3,1E-01 |
| A2APV2 | Formin-like protein 2 OS=Mus musculus OX=10090 GN=Fmnl2 PE=1 | 4 | 4 | 4 | 4 | 1086 | 123 | 7,53 | 7,46 | 4 | 1,244 | 1,114 | 1,199 | 1,403 | 1,159 | 1,183 | 1,084 | 1,234 | 1,06 | 0,06 | 3,1E-01 |
| Q99N50 | Synaptotagmin-like protein 2 OS=Mus musculus OX=10090 GN=Syt | 2 | 2 | 2 | 2 | 950 | 106,7 | 6,57 | 2,11 | 2 | 2,228 | 0,803 | 1,656 | 1,438 | 1,727 | 2,381 | 1,522 | 2,024 | 0,80 | -0,20 | 3,1E-01 |
| Q9JLN9 | Serine/threonine-protein kinase mTOR OS=Mus musculus OX=1009 | 3 | 8 | 10 | 8 | 2549 | 288,6 | 7,17 | 26,5 | 8 | 1,352 | 1,317 | 1,106 | 0,993 | 0,869 | 1,327 | 0,988 | 1,016 | 1,14 | 0,14 | 3,2E-01 |
| Q6PEB6 | MOB-like protein phocein OS=Mus musculus OX=10090 GN=Mob4 | 10 | 2 | 2 | 2 | 225 | 26 | 5,78 | 6,95 | 2 | 0,807 | 0,823 | 0,867 | 0,999 | 0,987 | 0,878 | 1,053 | 0,856 | 0,93 | -0,07 | 3,2E-01 |
| Q07417 | Short-chain specific acyl-CoA dehydrogenase, mitochondrial OS=M | 5 | 2 | 2 | 2 | 412 | 44,9 | 8,47 | 5,49 | 2 | 0,904 | 1,032 | 1,076 | 0,708 | 1,187 | 1,073 | 0,866 | 1,057 | 0,89 | -0,11 | 3,2E-01 |
| P26928 | Hepatocyte growth factor-like protein OS=Mus musculus OX=1009 | 2 | 1 | 1 | 1 | 716 | 80,6 | 7,56 | 0 | 1 | 6,541 | 0,981 | 1,725 | 0,817 | 0,835 | 1,031 | 0,902 | 1,358 | 2,44 | 1,44 | 3,2E-01 |
| Q6GYP7 | Ral GTPase-activating protein subunit alpha-1 OS=Mus musculus O | 5 | 7 | 7 | 7 | 2035 | 229,2 | 6,1 | 12,1 | 7 | 1,38 | 1,182 | 1,164 | 1,118 | 0,935 | 1,315 | 1,052 | 1,113 | 1,10 | 0,10 | 3,2E-01 |
| Q61584 | Fragile X mental retardation syndrome-related protein 1 OS=Mus n | 4 | 2 | 2 | 1 | 677 | 76,2 | 6,98 | 5,19 | 2 | 1,249 | 3,131 | 0,853 | 0,978 | 0,891 | 0,787 | 0,719 | 1,398 | 1,64 | 0,64 | 3,2E-01 |
| Q8K448 | ATP-binding cassette sub-family A member 5 OS=Mus musculus OX | 1 | 1 | 1 | 1 | 1642 | 185,8 | 7,2 | 2,59 | 1 | 1,352 | 1,164 | 0,968 | 1,244 | 1,217 | 1,008 | 0,935 | 1,123 | 1,10 | 0,10 | 3,2E-01 |
| P30275 | Creatine kinase U-type, mitochondrial OS=Mus musculus OX=1009 | 30 | 12 | 38 | 9 | 418 | 47 | 8,16 | 116,04 | 12 | 0,94 | 0,867 | 1,015 | 1,092 | 1,122 | 1,071 | 0,987 | 0,989 | 0,94 | -0,06 | 3,2E-01 |
| O88967 | ATP-dependent zinc metalloprotease YME1L1 OS=Mus musculus O | 5 | 3 | 3 | 3 | 715 | 80 | 8,97 | 4,66 | 3 | 1,672 | 1,504 | 1,452 | 0,991 | 1,002 | 1,422 | 1,059 | 1,355 | 1,16 | 0,16 | 3,2E-01 |
| P97952 | Sodium channel subunit beta-1 OS=Mus musculus OX=10090 GN=S | 15 | 4 | 5 | 4 | 218 | 24,6 | 4,83 | 4,4 | 4 | 0,818 | 0,903 | 1,011 | 0,949 | 0,98 | 0,891 | 1,018 | 1,009 | 0,94 | -0,06 | 3,2E-01 |
| Q921M4 | Golgin subfamily A member 2 OS=Mus musculus OX=10090 GN=Go | 1 | 1 | 1 | 1 | 999 | 113,2 | 5 | 0 | 1 | 0,66 | 0,647 | 0,773 | 0,868 | 0,853 | 1,321 | 0,681 | 0,757 | 0,82 | -0,18 | 3,2E-01 |
| Q5PR68 | Centrosomal protein of 112 kDa OS=Mus musculus OX=10090 GN= | 1 | 1 | 1 | 1 | 954 | 112,6 | 6,8 | 2,11 | 1 | 1,055 | 1,45 | 0,941 | 1,279 | 0,883 | 1,098 | 1,22 | 0,924 | 1,15 | 0,15 | 3,2E-01 |
| Q8K207 | Uncharacterized protein C1orf21 homolog OS=Mus musculus OX=1 | 8 | 1 | 1 | 1 | 121 | 13,9 | 5,22 | 2,7 | 1 | 1,166 | 0,503 | 0,932 | 0,69 | 0,972 | 1,349 | 0,789 | 0,988 | 0,80 | -0,20 | 3,2E-01 |
| P63213 | Guanine nucleotide-binding protein G(I)/G(S)/G(O) subunit gamma | 65 | 6 | 9 | 6 | 71 | 7,8 | 7,99 | 23,85 | 6 | 0,987 | 1,017 | 0,967 | 1,063 | 0,943 | 0,993 | 0,961 | 1,02 | 1,03 | 0,03 | 3,2E-01 |
| Q61553 | Fascin OS=Mus musculus OX=10090 GN=Fscn1 PE=1 SV=4 | 28 | 15 | 21 | 15 | 493 | 54,5 | 6,89 | 48,77 | 15 | 1,137 | 0,984 | 1,017 | 1,023 | 1,013 | 0,978 | 1 | 1,02 | 1,04 | 0,04 | 3,2E-01 |
| Q0P5W1 | Vacuolar protein sorting-associated protein 8 homolog OS=Mus mu | 2 | 3 | 3 | 3 | 1427 | 161 | 5,58 | 2,14 | 3 | 1,34 | 1,149 | 1,004 | 1,086 | 1,048 | 0,575 | 1,33 | 0,874 | 1,20 | 0,20 | 3,2E-01 |
| Q8BJS4 | SUN domain-containing protein 2 OS=Mus musculus OX=10090 GN | 5 | 3 | 3 | 3 | 731 | 81,6 | 7,02 | 2,46 | 3 | 1,364 | 0,826 | 0,975 | 0,805 | 0,736 | 1,072 | 0,811 | 0,674 | 1,21 | 0,21 | 3,2E-01 |
| Q80VP0 | Tectonin beta-propeller repeat-containing protein 1 OS=Mus musc | 10 | 9 | 12 | 9 | 1166 | 130,2 | 6,48 | 20,05 | 9 | 1,288 | 1,119 | 1,005 | 1,046 | 1,056 | 1,122 | 0,974 | 1,001 | 1,07 | 0,07 | 3,2E-01 |
| Q99LE6 | ATP-binding cassette sub-family F member 2 OS=Mus musculus OX | 6 | 4 | 5 | 4 | 628 | 71,7 | 7,05 | 6,16 | 4 | 0,958 | 0,983 | 1,045 | 1,096 | 0,933 | 0,905 | 1,009 | 1,042 | 1,05 | 0,05 | 3,2E-01 |
| P39087 | Glutamate receptor ionotropic, kainate 2 OS=Mus musculus OX=10 | 2 | 2 | 3 | 1 | 908 | 102,4 | 7,77 | 1,68 | 2 | 1,484 | 1,189 | 1,402 | 1,051 | 0,949 | 1,255 | 1,115 | 1,271 | 1,12 | 0,12 | 3,2E-01 |
| Q9JKL4 | NADH dehydrogenase [ubiquinone] 1 alpha subcomplex assembly f | 10 | 2 | 2 | 2 | 185 | 20,7 | 8,05 | 5,16 | 2 | 0,895 | 0,959 | 0,91 | 0,89 | 0,85 | 1,151 | 0,931 | 1,007 | 0,93 | -0,07 | 3,2E-01 |
| P19536 | Cytochrome c oxidase subunit 5B, mitochondrial OS=Mus musculus | 43 | 7 | 19 | 7 | 128 | 13,8 | 8,38 | 40,2 | 7 | 1,05 | 0,983 | 0,932 | 0,943 | 1,043 | 1,001 | 1,014 | 0,979 | 0,97 | -0,03 | 3,2E-01 |
| Q9WUB0 | RanBP-type and C3HC4-type zinc finger-containing protein 1 OS=M | 5 | 2 | 2 | 2 | 508 | 57,5 | 6,19 | 2,54 | 2 | 1,17 | 1,317 | 1,216 | 1,202 | 1,261 | 1,162 | 1,155 | 1,148 | 1,04 | 0,04 | 3,2E-01 |
| Q60996 | Serine/threonine-protein phosphatase 2A 56 kDa regulatory subun | 7 | 3 | 5 | 2 | 524 | 60,8 | 6,64 | 3,05 | 3 | 0,922 | 0,785 | 0,91 | 0,874 | 0,797 | 1,134 | 0,996 | 0,901 | 0,91 | -0,09 | 3,2E-01 |
| P29387 | Guanine nucleotide-binding protein subunit beta-4 OS=Mus muscu | 22 | 6 | 16 | 3 | 340 | 37,4 | 6,16 | 31,49 | 6 | 1,436 | 1,434 | 1,136 | 1,163 | 0,971 | 1,385 | 1,153 | 1,149 | 1,11 | 0,11 | 3,2E-01 |
| Q9DCN2 | NADH-cytochrome b5 reductase 3 OS=Mus musculus OX=10090 GN | 24 | 6 | 7 | 6 | 301 | 34,1 | 8,38 | 19,71 | 6 | 1,071 | 1,133 | 1,158 | 1,167 | 1,138 | 0,958 | 1,097 | 1,132 | 1,05 | 0,05 | 3,2E-01 |
| Q6ZQ18 | Protein EFR3 homolog B OS=Mus musculus OX=10090 GN=Efr3b PE | 13 | 10 | 12 | 10 | 817 | 92,3 | 6,54 | 28,85 | 10 | 0,929 | 0,894 | 0,974 | 1,017 | 0,936 | 1,006 | 1,028 | 0,987 | 0,96 | -0,04 | 3,2E-01 |
| P54071 | Isocitrate dehydrogenase [NADP], mitochondrial OS=Mus musculus | 34 | 14 | 22 | 13 | 452 | 50,9 | 8,69 | 45,43 | 14 | 1,146 | 1,151 | 1,062 | 1,019 | 1,079 | 1,097 | 1,002 | 1,032 | 1,04 | 0,04 | 3,2E-01 |
| Q8CI71 | Syndetin OS=Mus musculus OX=10090 GN=Vps50 PE=1 SV=2 | 10 | 9 | 9 | 9 | 964 | 111,1 | 6,07 | 22,59 | 9 | 0,978 | 0,875 | 0,895 | 0,951 | 0,904 | 0,957 | 1,007 | 0,969 | 0,96 | -0,04 | 3,2E-01 |
| Q8CGK3 | Lon protease homolog, mitochondrial OS=Mus musculus OX=10090 | 22 | 17 | 26 | 17 | 949 | 105,8 | 6,57 | 47,39 | 17 | 1,05 | 1,171 | 1,017 | 1,045 | 0,955 | 1,04 | 1,03 | 1,075 | 1,04 | 0,04 | 3,2E-01 |
| P50136 | 2-oxoisovalerate dehydrogenase subunit alpha, mitochondrial OS= | 8 | 3 | 4 | 3 | 442 | 50,3 | 8,06 | 4,07 | 3 | 1,114 | 1,258 | 0,911 | 0,995 | 1,005 | 0,994 | 0,942 | 1,006 | 1,08 | 0,08 | 3,2E-01 |
| Q8C525 | Protein MB21D2 OS=Mus musculus OX=10090 GN=Mb21d2 PE=1 S | 3 | 1 | 1 | 1 | 428 | 48,4 | 7,68 | 2,44 | 1 | 1,625 | 1,451 | 1,072 | 1,018 | 1,019 | 1,214 | 1,086 | 1,185 | 1,15 | 0,15 | 3,2E-01 |
| P62080 | Tetraspanin-5 OS=Mus musculus OX=10090 GN=Tspan5 PE=1 SV=1 | 5 | 1 | 1 | 1 | 268 | 30,3 | 4,78 | 2,37 | 1 | 0,01 | 1,637 | 1,414 | 1,262 | 1,292 | 1,64 | 1,636 | 1,373 | 0,73 | -0,27 | 3,2E-01 |
| Q9CRC9 | Glucosamine-6-phosphate isomerase 2 OS=Mus musculus OX=1009 | 7 | 2 | 3 | 1 | 276 | 31,1 | 6,9 | 2,92 | 2 | 1,199 | 1,044 | 1,175 | 0,983 | 0,945 | 1,197 | 0,923 | 0,986 | 1,09 | 0,09 | 3,2E-01 |
| Q8BHG2 | CXXC motif containing zinc binding protein OS=Mus musculus OX=1 | 8 | 1 | 1 | 1 | 160 | 18 | 5,12 | 2,04 | 1 | 1,46 | 1,607 | 1,484 | 1,76 | 1,174 | 0,988 | 1,649 | 1,693 | 1,15 | 0,15 | 3,2E-01 |
| Q08642 | Protein-arginine deiminase type-2 OS=Mus musculus OX=10090 GN | 1 | 1 | 2 | 1 | 673 | 76,2 | 5,53 | 2,01 | 1 | 1,135 | 1,369 | 0,917 | 0,829 | 0,693 | 1,085 | 1,074 | 0,691 | 1,20 | 0,20 | 3,2E-01 |
| P49586 | Choline-phosphate cytidylyltransferase A OS=Mus musculus OX=10 | 3 | 1 | 1 | 1 | 367 | 41,6 | 7,03 | 2,64 | 1 | 1,236 | 1,389 | 1,267 | 0,869 | 0,886 | 1,292 | 0,973 | 0,998 | 1,15 | 0,15 | 3,2E-01 |
| B1AS29 | Glutamate receptor ionotropic, kainate 3 OS=Mus musculus OX=10 | 4 | 3 | 4 | 2 | 919 | 104 | 7,52 | 5,06 | 3 | 1,286 | 1,284 | 1,039 | 0,923 | 1,029 | 1,189 | 0,844 | 0,974 | 1,12 | 0,12 | 3,2E-01 |
| P09103 | Protein disulfide-isomerase OS=Mus musculus OX=10090 GN=P4hb | 17 | 9 | 11 | 9 | 509 | 57 | 4,88 | 27,65 | 9 | 1,016 | 0,955 | 1,036 | 0,953 | 0,998 | 0,912 | 0,971 | 0,96 | 1,03 | 0,03 | 3,3E-01 |
| Q99L13 | 3-hydroxyisobutyrate dehydrogenase, mitochondrial OS=Mus musc | 20 | 6 | 8 | 6 | 335 | 35,4 | 8,13 | 18,32 | 6 | 0,956 | 0,92 | 0,972 | 0,899 | 0,911 | 1,05 | 0,949 | 0,982 | 0,96 | -0,04 | 3,3E-01 |
| Q7TNC8 | Glycine receptor subunit alpha-2 OS=Mus musculus OX=10090 GN= | 3 | 1 | 1 | 1 | 452 | 52 | 8,9 | 0 | 1 | 0,902 | 0,996 | 0,58 | 1,226 | 1,138 | 0,918 | 1,09 | 1,183 | 0,86 | -0,14 | 3,3E-01 |
| O08795 | Glucosidase 2 subunit beta OS=Mus musculus OX=10090 GN=Prkcs | 12 | 7 | 7 | 7 | 521 | 58,8 | 4,46 | 14,02 | 7 | 0,952 | 0,921 | 1,047 | 1,109 | 1,058 | 0,947 | 1,15 | 1,152 | 0,94 | -0,06 | 3,3E-01 |
| A2AJT4 | Arginine/serine-rich protein PNISR OS=Mus musculus OX=10090 G | 1 | 1 | 1 | 1 | 805 | 92,1 | 10,01 | 2,13 | 1 | 0,651 | 0,703 | 0,944 | 0,955 | 0,995 | 0,682 | 1,085 | 1 | 0,86 | -0,14 | 3,3E-01 |
| Q3UVK0 | Endoplasmic reticulum metallopeptidase 1 OS=Mus musculus OX=1 | 5 | 4 | 5 | 4 | 898 | 100,1 | 7,49 | 14,85 | 4 | 1,187 | 1,081 | 1,098 | 0,979 | 0,915 | 1,107 | 0,916 | 1,108 | 1,07 | 0,07 | 3,3E-01 |
| Q99K28 | ADP-ribosylation factor GTPase-activating protein 2 OS=Mus muscu | 2 | 1 | 1 | 1 | 520 | 56,6 | 8,18 | 2,73 | 1 | 1,161 | 1,338 | 1,007 | 1,312 | 1,071 | 1,197 | 1,077 | 1,123 | 1,08 | 0,08 | 3,3E-01 |
| E9Q557 | Desmoplakin OS=Mus musculus OX=10090 GN=Dsp PE=1 SV=1 | 4 | 13 | 14 | 13 | 2883 | 332,7 | 6,8 | 15,47 | 13 | 1,641 | 1,805 | 1,227 | 0,82 | 1,226 | 1,399 | 0,784 | 0,977 | 1,25 | 0,25 | 3,3E-01 |
| Q60737 | Casein kinase II subunit alpha OS=Mus musculus OX=10090 GN=Csr | 14 | 4 | 7 | 4 | 391 | 45,1 | 7,74 | 13,84 | 4 | 0,999 | 0,992 | 0,972 | 0,979 | 0,966 | 1,011 | 1,173 | 0,993 | 0,95 | -0,05 | 3,3E-01 |
| Q9JLM8 | Serine/threonine-protein kinase DCLK1 OS=Mus musculus OX=1009 | 32 | 21 | 28 | 20 | 756 | 84,1 | 8,87 | 70,39 | 21 | 0,952 | 0,909 | 0,949 | 0,99 | 0,967 | 0,946 | 1,033 | 0,962 | 0,97 | -0,03 | 3,3E-01 |
| Q8R420 | ATP-binding cassette sub-family A member 3 OS=Mus musculus OX | 1 | 1 | 1 | 1 | 1704 | 191,8 | 7,17 | 2,78 | 1 | 0,968 | 0,897 | 0,914 | 0,927 | 0,791 | 0,972 | 1,176 | 1,157 | 0,90 | -0,10 | 3,3E-01 |
| Q3UHJ0 | AP2-associated protein kinase 1 OS=Mus musculus OX=10090 GN=/ | 28 | 20 | 33 | 20 | 959 | 103,3 | 6,7 | 70,93 | 20 | 1,353 | 1,197 | 1,126 | 1,045 | 0,985 | 1,198 | 1,085 | 1,117 | 1,08 | 0,08 | 3,3E-01 |
| Q8VBX6 | Multiple PDZ domain protein OS=Mus musculus OX=10090 GN=Mp | 0 | 1 | 1 | 1 | 2055 | 218,6 | 5,02 | 0 | 1 | 0,623 | 1,162 | 0,948 | 0,756 | 0,945 | 0,01 | 0,659 | 0,849 | 1,42 | 0,42 | 3,3E-01 |
| P19324 | Serpin H1 OS=Mus musculus OX=10090 GN=Serpinh1 PE=1 SV=3 | 11 | 4 | 5 | 4 | 417 | 46,5 | 8,82 | 8,44 | 4 | 1,403 | 1,508 | 1,483 | 0,89 | 0,997 | 1,405 | 1,051 | 1,1 | 1,16 | 0,16 | 3,3E-01 |
| Q9D880 | Mitochondrial import inner membrane translocase subunit TIM50 | 16 | 5 | 7 | 5 | 353 | 39,8 | 8,13 | 8,2 | 5 | 1,068 | 1,011 | 1,148 | 1,045 | 1,108 | 1,226 | 1,05 | 1,091 | 0,95 | -0,05 | 3,3E-01 |
| Q91V17 | E3 ubiquitin-protein ligase ZNRF1 OS=Mus musculus OX=10090 GN | 6 | 1 | 1 | 1 | 227 | 23,8 | 6,67 | 2,2 | 1 | 1,221 | 1,736 | 1,155 | 1,361 | 0,596 | 1,522 | 0,843 | 1,415 | 1,25 | 0,25 | 3,3E-01 |
| O70589 | Peripheral plasma membrane protein CASK OS=Mus musculus OX= | 19 | 17 | 24 | 17 | 926 | 105 | 6,43 | 51,41 | 17 | 1,062 | 1,032 | 1,068 | 0,997 | 0,982 | 1,042 | 1,014 | 1,032 | 1,02 | 0,02 | 3,3E-01 |
| P30416 | Peptidyl-prolyl cis-trans isomerase FKBP4 OS=Mus musculus OX=1 | 19 | 10 | 13 | 10 | 458 | 51,5 | 5,72 | 22,23 | 10 | 0,945 | 0,949 | 0,983 | 0,999 | 1,003 | 0,839 | 0,915 | 0,96 | 1,04 | 0,04 | 3,3E-01 |
| P01942 | Hemoglobin subunit alpha OS=Mus musculus OX=10090 GN=Hba P | 17 | 2 | 4 | 2 | 142 | 15,1 | 8,22 | 7,87 | 2 | 0,99 | 0,858 | 0,567 | 0,567 | 1,264 | 0,712 | 0,473 | 2,708 | 0,58 | -0,42 | 3,3E-01 |
| Q8BKX1 | Brain-specific angiogenesis inhibitor 1-associated protein 2 OS=Mu | 30 | 16 | 19 | 16 | 535 | 59,2 | 9,04 | 44,88 | 16 | 1,056 | 1,067 | 0,974 | 0,989 | 0,986 | 0,981 | 1,003 | 1,012 | 1,03 | 0,03 | 3,3E-01 |

| | | | | | | | | | | | | | | | | | | | | | |
|---|---|---|---|---|---|---|---|---|---|---|---|---|---|---|---|---|---|---|---|---|---|
| P63082 | V-type proton ATPase 16 kDa proteolipid subunit OS=Mus musculu | 12 | 1 | 1 | 1 | 155 | 15,8 | 9,13 | 0 | 1 | 1,27 | 2,118 | 0,868 | 0,74 | 0,705 | 1,051 | 0,876 | 1,004 | 1,37 | 0,37 | 3,3E-01 |
| Q02956 | Protein kinase C zeta type OS=Mus musculus OX=10090 GN=Prkcz I | 3 | 1 | 1 | 1 | 592 | 67,6 | 5,76 | 2,72 | 1 | 0,931 | 0,939 | 0,75 | 0,823 | 0,864 | 0,689 | 0,479 | 0,937 | 1,16 | 0,16 | 3,3E-01 |
| P20152 | Vimentin OS=Mus musculus OX=10090 GN=Vim PE=1 SV=3 | 16 | 7 | 12 | 5 | 466 | 53,7 | 5,12 | 27,37 | 7 | 1,004 | 1,413 | 1,041 | 0,988 | 0,887 | 0,89 | 1,002 | 1,159 | 1,13 | 0,13 | 3,3E-01 |
| Q60751 | Insulin-like growth factor 1 receptor OS=Mus musculus OX=10090 ( | 1 | 1 | 1 | 1 | 1373 | 155,7 | 5,74 | 2,02 | 1 | 0,879 | 0,804 | 0,973 | 0,778 | 0,786 | 0,824 | 0,556 | 0,902 | 1,12 | 0,12 | 3,3E-01 |
| Q9R099 | Transducin beta-like protein 2 OS=Mus musculus OX=10090 GN=Tb | 3 | 1 | 1 | 1 | 442 | 49,6 | 9,04 | 3,23 | 1 | 1,328 | 1,336 | 1,963 | 1,385 | 1,01 | 1,642 | 1,13 | 1,353 | 1,17 | 0,17 | 3,3E-01 |
| Q8R395 | COMM domain-containing protein 5 OS=Mus musculus OX=10090 ( | 5 | 1 | 1 | 1 | 224 | 24,5 | 8,06 | 2,43 | 1 | 0,869 | 0,652 | 0,666 | 1,062 | 1,13 | 0,782 | 0,915 | 0,932 | 0,86 | -0,14 | 3,3E-01 |
| P70414 | Sodium/calcium exchanger 1 OS=Mus musculus OX=10090 GN=Slc8 | 9 | 8 | 9 | 8 | 970 | 108 | 5 | 13,19 | 8 | 0,958 | 0,924 | 0,962 | 0,894 | 0,921 | 0,984 | 0,935 | 1,007 | 0,97 | -0,03 | 3,3E-01 |
| Q3U0M1 | Trafficking protein particle complex subunit 9 OS=Mus musculus O) | 4 | 4 | 4 | 4 | 1148 | 128,2 | 6,47 | 8,78 | 4 | 1,055 | 0,907 | 1,023 | 1,012 | 0,989 | 0,995 | 0,929 | 0,928 | 1,04 | 0,04 | 3,3E-01 |
| Q04447 | Creatine kinase B-type OS=Mus musculus OX=10090 GN=Ckb PE=1 | 47 | 16 | 60 | 14 | 381 | 42,7 | 5,67 | 146,56 | 16 | 1,142 | 1,206 | 1,017 | 1,005 | 0,949 | 1,094 | 1,025 | 1,058 | 1,06 | 0,06 | 3,3E-01 |
| O88597 | Beclin-1 OS=Mus musculus OX=10090 GN=Becn1 PE=1 SV=3 | 10 | 4 | 5 | 4 | 448 | 51,6 | 4,93 | 11 | 4 | 0,735 | 0,799 | 0,78 | 0,951 | 0,804 | 0,882 | 0,905 | 0,894 | 0,94 | -0,06 | 3,3E-01 |
| P23242 | Gap junction alpha-1 protein OS=Mus musculus OX=10090 GN=Gja | 15 | 4 | 4 | 4 | 382 | 43 | 8,76 | 4,67 | 4 | 0,909 | 1,262 | 0,873 | 0,784 | 0,972 | 1,129 | 1,179 | 1,032 | 0,89 | -0,11 | 3,3E-01 |
| Q9D6K8 | FUN14 domain-containing protein 2 OS=Mus musculus OX=10090 ( | 8 | 1 | 1 | 1 | 151 | 16,6 | 9,7 | 2,05 | 1 | 1,83 | 1,897 | 1,155 | 0,997 | 1,232 | 1,262 | 1,392 | 0,947 | 1,22 | 0,22 | 3,3E-01 |
| Q8K1A6 | Coiled-coil and C2 domain-containing protein 1A OS=Mus musculus | 6 | 5 | 5 | 5 | 943 | 103,6 | 7,84 | 7,03 | 5 | 1,392 | 1,138 | 1,136 | 1,037 | 1,099 | 0,711 | 1,327 | 0,931 | 1,16 | 0,16 | 3,3E-01 |
| Q91VR7 | Microtubule-associated proteins 1A/1B light chain 3A OS=Mus mus | 19 | 3 | 3 | 2 | 121 | 14,3 | 8,68 | 7,28 | 3 | 1,424 | 1,285 | 1,223 | 1,059 | 0,957 | 1,32 | 1,027 | 1,213 | 1,10 | 0,10 | 3,3E-01 |
| P08551 | Neurofilament light polypeptide OS=Mus musculus OX=10090 GN= | 27 | 17 | 23 | 14 | 543 | 61,5 | 4,64 | 53,08 | 17 | 0,923 | 0,924 | 0,897 | 0,898 | 1,08 | 0,974 | 0,95 | 0,844 | 0,95 | -0,05 | 3,3E-01 |
| Q8K021 | Secretory carrier-associated membrane protein 1 OS=Mus musculu | 4 | 1 | 2 | 1 | 338 | 38 | 7,71 | 6,35 | 1 | 1,051 | 1,128 | 1,004 | 1,119 | 1,015 | 1,251 | 1,17 | 1,109 | 0,95 | -0,05 | 3,3E-01 |
| P56380 | Bis(5'-nucleosyl)-tetraphosphatase [asymmetrical] OS=Mus muscul | 21 | 3 | 4 | 3 | 147 | 17 | 6,28 | 5,29 | 3 | 1,348 | 1,356 | 1,382 | 0,968 | 1,897 | 2,176 | 1,141 | 1,069 | 0,80 | -0,20 | 3,3E-01 |
| P48455 | Serine/threonine-protein phosphatase 2B catalytic subunit gamma | 11 | 6 | 9 | 2 | 513 | 58,7 | 7,25 | 18,97 | 6 | 1,296 | 0,793 | 0,01 | 0,641 | 0,865 | 12,255 | 1,185 | 0,467 | 0,19 | -0,81 | 3,3E-01 |
| Q9CZV8 | F-box/LRR-repeat protein 20 OS=Mus musculus OX=10090 GN=Fbx | 5 | 2 | 2 | 2 | 436 | 48,4 | 7,49 | 5,46 | 2 | 0,488 | 0,817 | 0,928 | 0,973 | 0,937 | 0,986 | 0,981 | 0,797 | 0,87 | -0,13 | 3,4E-01 |
| P54265 | Myotonin-protein kinase OS=Mus musculus OX=10090 GN=Dmpk P | 2 | 1 | 1 | 1 | 631 | 69,6 | 4,91 | 1,92 | 1 | 1,27 | 1,008 | 0,984 | 0,852 | 0,918 | 4,262 | 1,299 | 1,011 | 0,55 | -0,45 | 3,4E-01 |
| Q5DU31 | Interactor protein for cytohesin exchange factors 1 OS=Mus muscu | 3 | 1 | 4 | 1 | 406 | 45,8 | 6,73 | 9,82 | 1 | 1,325 | 1,508 | 1,317 | 1,26 | 1,361 | 1,39 | 1,213 | 1,044 | 1,08 | 0,08 | 3,4E-01 |
| P34914 | Bifunctional epoxide hydrolase 2 OS=Mus musculus OX=10090 GN= | 6 | 3 | 3 | 3 | 554 | 62,5 | 6,19 | 5,43 | 3 | 1,424 | 1,858 | 1,446 | 1,52 | 1,417 | 1,191 | 1,465 | 1,615 | 1,10 | 0,10 | 3,4E-01 |
| Q50H33 | BTB/POZ domain-containing protein KCTD8 OS=Mus musculus OX= | 2 | 1 | 1 | 1 | 476 | 52,7 | 8,38 | 2,05 | 1 | 1,642 | 1,16 | 0,866 | 0,999 | 1,239 | 1,383 | 1,38 | 1,391 | 0,87 | -0,13 | 3,4E-01 |
| Q03157 | Amyloid-like protein 1 OS=Mus musculus OX=10090 GN=Aplp1 PE= | 5 | 3 | 4 | 3 | 653 | 72,7 | 5,67 | 3,79 | 3 | 1,174 | 1,163 | 0,993 | 1,029 | 0,929 | 1,158 | 0,839 | 1,076 | 1,09 | 0,09 | 3,4E-01 |
| P32848 | Parvalbumin alpha OS=Mus musculus OX=10090 GN=Pvalb PE=1 SV | 25 | 3 | 4 | 3 | 110 | 11,9 | 5,19 | 15,58 | 3 | 0,927 | 1,01 | 1,119 | 0,968 | 0,969 | 0,937 | 0,883 | 1,023 | 1,06 | 0,06 | 3,4E-01 |
| Q02248 | Catenin beta-1 OS=Mus musculus OX=10090 GN=Ctnnb1 PE=1 SV=1 | 23 | 15 | 27 | 13 | 781 | 85,4 | 5,86 | 86,47 | 15 | 1,064 | 1,017 | 1,003 | 0,987 | 1,01 | 1,104 | 1,021 | 1,048 | 0,97 | -0,03 | 3,4E-01 |
| Q80UW2 | F-box only protein 2 OS=Mus musculus OX=10090 GN=Fbxo2 PE=1 | 13 | 3 | 3 | 3 | 297 | 33,7 | 4,28 | 5,92 | 3 | 1,184 | 1,009 | 1,165 | 0,848 | 0,834 | 1,092 | 0,97 | 0,911 | 1,10 | 0,10 | 3,4E-01 |
| Q8R3B1 | 1-phosphatidylinositol 4,5-bisphosphate phosphodiesterase delta-1 | 1 | 1 | 1 | 1 | 756 | 85,8 | 6,21 | 2,27 | 1 | 0,811 | 0,901 | 0,853 | 0,818 | 0,8 | 1,508 | 0,862 | 0,905 | 0,83 | -0,17 | 3,4E-01 |
| Q78J03 | Methionine-R-sulfoxide reductase B2, mitochondrial OS=Mus musc | 6 | 1 | 1 | 1 | 175 | 19,1 | 9,06 | 2,93 | 1 | 0,732 | 0,575 | 1,11 | 1,181 | 1,032 | 0,986 | 1,109 | 1,093 | 0,85 | -0,15 | 3,4E-01 |
| Q9D428 | Golgin subfamily A member 7B OS=Mus musculus OX=10090 GN=G | 13 | 2 | 2 | 2 | 167 | 18,3 | 6,01 | 5,01 | 2 | 1,402 | 1,138 | 1,098 | 1,097 | 1,058 | 1,11 | 1,204 | 1,012 | 1,08 | 0,08 | 3,4E-01 |
| P00493 | Hypoxanthine-guanine phosphoribosyltransferase OS=Mus muscul | 24 | 4 | 5 | 4 | 218 | 24,6 | 6,68 | 6,67 | 4 | 0,882 | 0,85 | 1,252 | 1,083 | 1,071 | 1,098 | 1,233 | 1,086 | 0,91 | -0,09 | 3,4E-01 |
| P42337 | Phosphatidylinositol 4,5-bisphosphate 3-kinase catalytic subunit al | 1 | 1 | 1 | 1 | 1068 | 124,3 | 7,15 | 0 | 1 | 0,648 | 1,129 | 1,407 | 0,904 | 0,818 | 0,821 | 0,959 | 0,8 | 1,20 | 0,20 | 3,4E-01 |
| P38647 | Stress-70 protein, mitochondrial OS=Mus musculus OX=10090 GN= | 34 | 26 | 32 | 25 | 679 | 73,4 | 6,07 | 72,28 | 26 | 1,015 | 0,933 | 0,968 | 0,947 | 0,999 | 0,977 | 0,963 | 1,011 | 0,98 | -0,02 | 3,4E-01 |
| Q922R8 | Protein disulfide-isomerase A6 OS=Mus musculus OX=10090 GN=P( | 14 | 6 | 7 | 6 | 440 | 48,1 | 5,14 | 20,31 | 6 | 1,062 | 1,056 | 1,135 | 1,099 | 1,002 | 1,089 | 1,08 | 1,069 | 1,03 | 0,03 | 3,4E-01 |
| P42208 | Septin-2 OS=Mus musculus OX=10090 GN=Septin2 PE=1 SV=2 | 12 | 4 | 4 | 4 | 361 | 41,5 | 6,55 | 12,15 | 4 | 1,428 | 1,094 | 1,135 | 1,078 | 1,156 | 1,095 | 1,045 | 1,084 | 1,08 | 0,08 | 3,4E-01 |
| Q8C3F2 | Constitutive coactivator of PPAR-gamma-like protein 2 OS=Mus mu | 1 | 1 | 1 | 1 | 1091 | 119,6 | 8,46 | 3,24 | 1 | 1,057 | 0,938 | 1,097 | 0,849 | 0,971 | 2,511 | 0,967 | 1,077 | 0,71 | -0,29 | 3,4E-01 |
| Q80Z24 | Neuronal growth regulator 1 OS=Mus musculus OX=10090 GN=Neg | 23 | 6 | 6 | 6 | 348 | 37,9 | 6,52 | 2,64 | 6 | 1,012 | 0,568 | 0,936 | 0,925 | 1,278 | 0,723 | 1,269 | 0,883 | 0,83 | -0,17 | 3,4E-01 |
| P40240 | CD9 antigen OS=Mus musculus OX=10090 GN=Cd9 PE=1 SV=2 | 3 | 1 | 1 | 1 | 226 | 25,2 | 7,23 | 3,57 | 1 | 1,102 | 1,101 | 1,067 | 0,949 | 0,966 | 1,394 | 1,281 | 1,027 | 0,90 | -0,10 | 3,4E-01 |
| Q6PGF7 | Exocyst complex component 8 OS=Mus musculus OX=10090 GN=Ex | 9 | 6 | 6 | 6 | 716 | 81 | 5,4 | 15,2 | 6 | 1,25 | 1,259 | 1,149 | 1,094 | 1,03 | 1,287 | 1,127 | 0,984 | 1,07 | 0,07 | 3,4E-01 |
| Q9EP89 | Serine beta-lactamase-like protein LACTB, mitochondrial OS=Mus n | 9 | 6 | 7 | 6 | 551 | 60,7 | 8,9 | 11,17 | 6 | 1,02 | 1,001 | 0,932 | 0,953 | 1,075 | 0,952 | 1,084 | 0,964 | 0,96 | -0,04 | 3,4E-01 |
| Q69ZQ1 | Myogenesis-regulating glycosidase OS=Mus musculus OX=10090 GI | 2 | 1 | 1 | 1 | 716 | 81,3 | 8,29 | 2,05 | 1 | 1,093 | 0,809 | 1,152 | 0,908 | 1,184 | 1,01 | 0,935 | 1,324 | 0,89 | -0,11 | 3,4E-01 |
| Q61334 | B-cell receptor-associated protein 29 OS=Mus musculus OX=10090 | 10 | 3 | 3 | 3 | 240 | 27,9 | 9,72 | 2,23 | 3 | 1,508 | 1,121 | 1,365 | 0,848 | 1,093 | 0,977 | 1,04 | 1,118 | 1,15 | 0,15 | 3,4E-01 |
| Q8VHL1 | Histone-lysine N-methyltransferase SETD7 OS=Mus musculus OX=1 | 9 | 2 | 3 | 2 | 366 | 40,5 | 4,65 | 7,92 | 2 | 1,039 | 1,052 | 1,091 | 0,891 | 0,767 | 1,064 | 0,86 | 1,036 | 1,09 | 0,09 | 3,4E-01 |
| Q9QYY8 | Spastin OS=Mus musculus OX=10090 GN=Spast PE=1 SV=3 | 2 | 1 | 1 | 1 | 614 | 66,4 | 9,69 | 2,02 | 1 | 1,19 | 1,039 | 0,908 | 1,149 | 1,033 | 1,004 | 1,504 | 1,301 | 0,89 | -0,11 | 3,4E-01 |
| Q61166 | Microtubule-associated protein RP/EB family member 1 OS=Mus m | 21 | 4 | 4 | 3 | 268 | 30 | 5,22 | 8,43 | 4 | 1,192 | 0,682 | 0,795 | 0,926 | 1,024 | 1,007 | 1,177 | 0,899 | 0,88 | -0,12 | 3,4E-01 |
| Q64487-1 | Isoform L of Receptor-type tyrosine-protein phosphatase delta OS= | 11 | 16 | 19 | 11 | 1501 | 168,8 | 6,71 | 40,56 | 16 | 1,178 | 1,036 | 1,133 | 0,989 | 1,087 | 0,948 | 1,052 | 1,032 | 1,05 | 0,05 | 3,4E-01 |
| P28656 | Nucleosome assembly protein 1-like 1 OS=Mus musculus OX=1009( | 10 | 3 | 4 | 2 | 391 | 45,3 | 4,46 | 9,63 | 3 | 0,968 | 1,13 | 1,165 | 0,978 | 0,758 | 1,259 | 0,895 | 0,819 | 1,14 | 0,14 | 3,4E-01 |
| Q99LT0 | Protein dpy-30 homolog OS=Mus musculus OX=10090 GN=Dpy30 P | 11 | 2 | 2 | 2 | 99 | 11,2 | 4,88 | 5,02 | 2 | 1,039 | 1,13 | 1,051 | 1,116 | 0,954 | 1,036 | 1,008 | 1,146 | 1,05 | 0,05 | 3,4E-01 |
| P60853 | Leucine zipper putative tumor suppressor 1 OS=Mus musculus OX= | 6 | 3 | 3 | 3 | 599 | 67,2 | 7,56 | 2,44 | 3 | 0,929 | 1,169 | 0,866 | 0,865 | 0,86 | 1,021 | 0,811 | 0,757 | 1,11 | 0,11 | 3,4E-01 |
| Q9QUH0 | Glutaredoxin-1 OS=Mus musculus OX=10090 GN=Glrx PE=1 SV=3 | 30 | 2 | 5 | 2 | 107 | 11,9 | 8,37 | 12,37 | 2 | 1,108 | 1,359 | 1,311 | 1,21 | 1,082 | 0,98 | 1,049 | 1,415 | 1,10 | 0,10 | 3,4E-01 |
| Q8BR07 | Protein bicaudal D homolog 1 OS=Mus musculus OX=10090 GN=Bic | 2 | 1 | 1 | 1 | 835 | 95,8 | 5,71 | 2,02 | 1 | 1,69 | 1,512 | 1,626 | 1,018 | 1,449 | 1,187 | 1,256 | 1,285 | 1,13 | 0,13 | 3,4E-01 |
| Q8C1A9 | Protein ABHD18 OS=Mus musculus OX=10090 GN=Abhd18 PE=2 SV | 2 | 1 | 1 | 1 | 464 | 53,1 | 8,79 | 2,55 | 1 | 1,298 | 1,239 | 1,268 | 1,183 | 0,981 | 1,404 | 1,112 | 1,108 | 1,08 | 0,08 | 3,4E-01 |
| P62908 | 40S ribosomal protein S3 OS=Mus musculus OX=10090 GN=Rps3 PE | 45 | 10 | 13 | 10 | 243 | 26,7 | 9,66 | 21,86 | 10 | 1,012 | 1,134 | 1,104 | 0,944 | 0,988 | 0,997 | 0,992 | 1,033 | 1,05 | 0,05 | 3,4E-01 |
| P16388 | Potassium voltage-gated channel subfamily A member 1 OS=Mus n | 8 | 4 | 4 | 2 | 495 | 56,4 | 5,14 | 8,09 | 4 | 0,938 | 0,809 | 0,572 | 1,2 | 0,969 | 1,477 | 1,197 | 0,727 | 0,81 | -0,19 | 3,4E-01 |
| P50516 | V-type proton ATPase catalytic subunit A OS=Mus musculus OX=10 | 44 | 28 | 81 | 28 | 617 | 68,3 | 5,58 | 192,99 | 28 | 0,988 | 0,979 | 0,956 | 0,999 | 0,976 | 0,962 | 1,04 | 1,034 | 0,98 | -0,02 | 3,4E-01 |
| Q3URE1 | Malonate--CoA ligase ACSF3, mitochondrial OS=Mus musculus OX= | 13 | 7 | 7 | 7 | 583 | 65 | 8,02 | 11,67 | 7 | 1,448 | 1,18 | 1,071 | 0,954 | 0,88 | 1,336 | 0,842 | 0,959 | 1,16 | 0,16 | 3,4E-01 |
| O88343 | Electrogenic sodium bicarbonate cotransporter 1 OS=Mus musculus | 3 | 4 | 4 | 4 | 1079 | 121,4 | 6,84 | 7,13 | 4 | 0,846 | 0,81 | 0,929 | 0,972 | 1,086 | 0,749 | 1,086 | 0,997 | 0,91 | -0,09 | 3,4E-01 |
| P09242 | Alkaline phosphatase, tissue-nonspecific isozyme OS=Mus musculu | 2 | 1 | 1 | 1 | 524 | 57,5 | 7,01 | 2,24 | 1 | 0,937 | 0,905 | 0,822 | 0,805 | 0,934 | 0,79 | 1,027 | 0,961 | 0,93 | -0,07 | 3,5E-01 |
| Q6P5F9 | Exportin-1 OS=Mus musculus OX=10090 GN=Xpo1 PE=1 SV=1 | 9 | 8 | 9 | 8 | 1071 | 123 | 6,07 | 17,16 | 8 | 1,166 | 1,156 | 1,204 | 1,007 | 0,958 | 1,135 | 1,093 | 1,107 | 1,06 | 0,06 | 3,5E-01 |
| O54984 | ATPase Asna1 OS=Mus musculus OX=10090 GN=Asna1 PE=1 SV=2 | 19 | 4 | 4 | 4 | 348 | 38,8 | 4,91 | 10,74 | 4 | 1,045 | 1,163 | 1,112 | 1,091 | 0,983 | 1,2 | 0,972 | 1,016 | 1,06 | 0,06 | 3,5E-01 |
| Q64387 | Prepronociceptin OS=Mus musculus OX=10090 GN=Pnoc PE=2 SV= | 6 | 1 | 3 | 1 | 187 | 20,9 | 8,56 | 4,57 | 1 | 0,841 | 0,889 | 0,913 | 1,146 | 0,901 | 1,091 | 1,04 | 1,09 | 0,92 | -0,08 | 3,5E-01 |
| O09172 | Glutamate--cysteine ligase regulatory subunit OS=Mus musculus O) | 15 | 3 | 3 | 3 | 274 | 30,5 | 5,52 | 6,79 | 3 | 1,554 | 1,276 | 1,168 | 1,087 | 0,724 | 1,409 | 0,986 | 1,228 | 1,17 | 0,17 | 3,5E-01 |

| | | | | | | | | | | | | | | | | | | | | | |
|---|---|---|---|---|---|---|---|---|---|---|---|---|---|---|---|---|---|---|---|---|---|
| P70206 | Plexin-A1 OS=Mus musculus OX=10090 GN=Plxna1 PE=1 SV=1 | 15 | 24 | 29 | 17 | 1894 | 211 | 6,9 | 51,56 | 24 | 1,1 | 1,054 | 1,092 | 1,033 | 1,003 | 1,121 | 0,996 | 1,024 | 1,03 | 0,03 | 3,5E-01 |
| Q8K4P8 | E3 ubiquitin-protein ligase HECW1 OS=Mus musculus OX=10090 GN | 2 | 3 | 4 | 2 | 1604 | 179,4 | 5,39 | 6,56 | 3 | 0,743 | 0,942 | 0,726 | 0,792 | 0,782 | 0,459 | 0,556 | 0,927 | 1,18 | 0,18 | 3,5E-01 |
| Q63810 | Calcineurin subunit B type 1 OS=Mus musculus OX=10090 GN=Ppp3 | 44 | 8 | 23 | 8 | 170 | 19,3 | 4,81 | 45,63 | 8 | 1,173 | 1,056 | 0,988 | 1,008 | 1,025 | 0,951 | 1,05 | 1,009 | 1,05 | 0,05 | 3,5E-01 |
| P62301 | 40S ribosomal protein S13 OS=Mus musculus OX=10090 GN=Rps13 | 12 | 2 | 2 | 2 | 151 | 17,2 | 10,54 | 4,2 | 2 | 0,696 | 0,698 | 0,944 | 1,167 | 1,244 | 0,731 | 1,145 | 1,033 | 0,84 | -0,16 | 3,5E-01 |
| Q9QXJ1 | Amyloid-beta A4 precursor protein-binding family B member 1 OS= | 11 | 5 | 5 | 5 | 710 | 77,3 | 5 | 7,99 | 5 | 1,111 | 1,027 | 1,063 | 1,036 | 0,994 | 1,144 | 1,096 | 1,193 | 0,96 | -0,04 | 3,5E-01 |
| Q99KI0 | Aconitate hydratase, mitochondrial OS=Mus musculus OX=10090 G | 51 | 36 | 74 | 36 | 780 | 85,4 | 7,93 | 179,36 | 36 | 0,959 | 0,93 | 0,948 | 1,006 | 1,078 | 0,716 | 0,963 | 0,698 | 1,11 | 0,11 | 3,5E-01 |
| Q8BL65-4 | Isoform 4 of Actin-binding LIM protein 2 OS=Mus musculus OX=100 | 8 | 4 | 5 | 4 | 617 | 68,6 | 8,07 | 12,86 | 4 | 0,984 | 1,029 | 1,116 | 0,981 | 1,07 | 1,095 | 1,001 | 1,104 | 0,96 | -0,04 | 3,5E-01 |
| Q06185 | ATP synthase subunit e, mitochondrial OS=Mus musculus OX=1009 | 56 | 4 | 7 | 4 | 71 | 8,2 | 9,35 | 14,7 | 4 | 0,911 | 1,073 | 0,9 | 0,875 | 0,927 | 1,075 | 0,952 | 1,038 | 0,94 | -0,06 | 3,5E-01 |
| Q8BJU0 | Small glutamine-rich tetratricopeptide repeat-containing protein al | 21 | 6 | 6 | 6 | 315 | 34,3 | 5,06 | 13,41 | 6 | 0,886 | 0,867 | 0,954 | 1,17 | 1,094 | 0,923 | 1,13 | 1,069 | 0,92 | -0,08 | 3,5E-01 |
| Q91YM2 | Rho GTPase-activating protein 35 OS=Mus musculus OX=10090 GN | 9 | 12 | 14 | 12 | 1499 | 170,3 | 6,61 | 21,53 | 12 | 1,169 | 1,022 | 1,034 | 0,994 | 1,014 | 1,043 | 0,978 | 1,016 | 1,04 | 0,04 | 3,5E-01 |
| Q9CQY6 | Ubiquinol-cytochrome-c reductase complex assembly factor 2 OS=I | 18 | 2 | 2 | 2 | 136 | 16,3 | 9,19 | 3,22 | 2 | 1,059 | 1,048 | 0,846 | 0,994 | 0,929 | 1,334 | 1,124 | 0,98 | 0,90 | -0,10 | 3,5E-01 |
| O08582 | GTP-binding protein 1 OS=Mus musculus OX=10090 GN=Gtpbp1 PE | 3 | 2 | 3 | 2 | 668 | 72,3 | 8,29 | 2,22 | 2 | 1,616 | 0,81 | 1,38 | 1,237 | 1,466 | 2,119 | 1,222 | 1,311 | 0,82 | -0,18 | 3,5E-01 |
| Q8BTM8 | Filamin-A OS=Mus musculus OX=10090 GN=Flna PE=1 SV=5 | 3 | 7 | 8 | 7 | 2647 | 281 | 6,04 | 11,41 | 7 | 1,379 | 1,246 | 1,222 | 0,964 | 0,964 | 1,209 | 1,043 | 1,173 | 1,10 | 0,10 | 3,5E-01 |
| A2ADA5 | tRNA pseudouridine synthase-like 1 OS=Mus musculus OX=10090 G | 5 | 1 | 1 | 1 | 291 | 32 | 10,04 | 2,55 | 1 | 0,622 | 0,616 | 0,851 | 0,764 | 0,76 | 0,746 | 0,949 | 0,715 | 0,90 | -0,10 | 3,5E-01 |
| Q99N93 | 39S ribosomal protein L16, mitochondrial OS=Mus musculus OX=10 | 6 | 2 | 2 | 2 | 251 | 28,8 | 10,27 | 4,4 | 2 | 0,952 | 0,784 | 1,087 | 0,891 | 0,961 | 1,001 | 1,037 | 0,98 | 0,93 | -0,07 | 3,5E-01 |
| Q9D3A8 | Carboxyl-terminal PDZ ligand of neuronal nitric oxide synthase prot | 8 | 3 | 3 | 3 | 503 | 55,8 | 6,09 | 3,19 | 3 | 0,985 | 0,659 | 0,865 | 0,803 | 1,043 | 0,795 | 0,935 | 0,885 | 0,91 | -0,09 | 3,5E-01 |
| Q9D061 | Acyl-CoA-binding domain-containing protein 6 OS=Mus musculus C | 17 | 5 | 5 | 5 | 282 | 30,9 | 5,11 | 3,07 | 5 | 0,808 | 0,99 | 1,238 | 1,172 | 0,978 | 1,054 | 0,878 | 0,866 | 1,11 | 0,11 | 3,5E-01 |
| P57759 | Endoplasmic reticulum resident protein 29 OS=Mus musculus OX=1 | 17 | 4 | 8 | 4 | 262 | 28,8 | 6,15 | 12,81 | 4 | 0,975 | 0,826 | 1,098 | 0,976 | 1,008 | 1,091 | 1,136 | 0,931 | 0,93 | -0,07 | 3,5E-01 |
| Q91V76 | Ester hydrolase C11orf54 homolog OS=Mus musculus OX=10090 PE | 6 | 2 | 4 | 2 | 315 | 35 | 6,29 | 11,36 | 2 | 1,231 | 1,061 | 1,028 | 0,997 | 0,954 | 1,033 | 1,051 | 1,048 | 1,06 | 0,06 | 3,5E-01 |
| P43275 | Histone H1.1 OS=Mus musculus OX=10090 GN=Hist1h1a PE=1 SV=2 | 13 | 4 | 6 | 1 | 213 | 21,8 | 10,93 | 14,85 | 4 | 0,888 | 1,298 | 0,87 | 0,84 | 1,094 | 1,256 | 0,73 | 1,883 | 0,79 | -0,21 | 3,5E-01 |
| Q8C1B1 | Calmodulin-regulated spectrin-associated protein 2 OS=Mus muscu | 4 | 4 | 4 | 4 | 1461 | 164,2 | 6,86 | 3,33 | 4 | 1,267 | 1,417 | 1,114 | 1,117 | 1,127 | 1,033 | 1,278 | 1,12 | 1,08 | 0,08 | 3,5E-01 |
| Q9WVG6 | Histone-arginine methyltransferase CARM1 OS=Mus musculus OX= | 3 | 2 | 2 | 2 | 608 | 65,8 | 6,77 | 4,09 | 2 | 0,948 | 0,819 | 0,816 | 0,791 | 0,84 | 1,154 | 0,78 | 0,963 | 0,90 | -0,10 | 3,5E-01 |
| Q9QUM9 | Proteasome subunit alpha type-6 OS=Mus musculus OX=10090 GN | 33 | 7 | 9 | 7 | 246 | 27,4 | 6,76 | 25,49 | 7 | 1,078 | 0,999 | 1,011 | 0,945 | 0,943 | 1,044 | 0,936 | 0,961 | 1,04 | 0,04 | 3,5E-01 |
| Q9Z1R2 | Large proline-rich protein BAG6 OS=Mus musculus OX=10090 GN=E | 9 | 9 | 10 | 9 | 1154 | 121 | 5,71 | 13,74 | 9 | 1,145 | 1,045 | 1,161 | 1,29 | 1,129 | 1,105 | 1,077 | 1,122 | 1,05 | 0,05 | 3,5E-01 |
| P49615 | Cyclin-dependent-like kinase 5 OS=Mus musculus OX=10090 GN=Cc | 32 | 7 | 8 | 6 | 292 | 33,3 | 7,66 | 17,03 | 7 | 1,041 | 1,142 | 1,102 | 1,242 | 1,012 | 1,119 | 1,107 | 1,093 | 1,05 | 0,05 | 3,5E-01 |
| Q6P5U7 | NACHT and WD repeat domain-containing protein 2 OS=Mus musc | 11 | 14 | 14 | 14 | 1742 | 197,3 | 6,15 | 24,04 | 14 | 1,155 | 1,185 | 1,063 | 0,987 | 0,91 | 1,19 | 1,026 | 0,959 | 1,07 | 0,07 | 3,5E-01 |
| Q8R5H1 | Ubiquitin carboxyl-terminal hydrolase 15 OS=Mus musculus OX=10 | 7 | 6 | 6 | 6 | 981 | 112,3 | 5,17 | 12,87 | 6 | 1,142 | 0,973 | 1,178 | 1,183 | 1,074 | 0,997 | 1,073 | 1,111 | 1,05 | 0,05 | 3,5E-01 |
| Q8BK72 | 28S ribosomal protein S27, mitochondrial OS=Mus musculus OX=10 | 9 | 3 | 3 | 3 | 415 | 47,7 | 5,5 | 4,79 | 3 | 1,016 | 1,086 | 1,101 | 0,888 | 0,885 | 0,948 | 0,953 | 1,061 | 1,06 | 0,06 | 3,5E-01 |
| Q8CIB5 | Fermitin family homolog 2 OS=Mus musculus OX=10090 GN=Fermt | 11 | 6 | 7 | 6 | 680 | 77,8 | 6,7 | 9,89 | 6 | 1,009 | 0,971 | 1,172 | 1,051 | 0,988 | 1,037 | 1,031 | 0,956 | 1,05 | 0,05 | 3,5E-01 |
| Q9DCX2 | ATP synthase subunit d, mitochondrial OS=Mus musculus OX=1009 | 60 | 11 | 20 | 11 | 161 | 18,7 | 5,69 | 33,48 | 11 | 0,875 | 0,889 | 0,971 | 1,009 | 1,094 | 0,519 | 1,019 | 0,398 | 1,24 | 0,24 | 3,5E-01 |
| Q6P2B1 | Transportin-3 OS=Mus musculus OX=10090 GN=Tnpo3 PE=1 SV=1 | 10 | 7 | 8 | 7 | 923 | 104,1 | 5,57 | 11,59 | 7 | 0,9 | 1,109 | 1,152 | 1,047 | 0,945 | 1,023 | 1,005 | 1,004 | 1,06 | 0,06 | 3,5E-01 |
| P20357 | Microtubule-associated protein 2 OS=Mus musculus OX=10090 GN | 28 | 41 | 65 | 41 | 1828 | 199 | 4,91 | 130,03 | 41 | 0,98 | 0,956 | 0,983 | 0,972 | 1,005 | 0,948 | 1,034 | 0,981 | 0,98 | -0,02 | 3,6E-01 |
| Q64455 | Receptor-type tyrosine-protein phosphatase eta OS=Mus musculus | 1 | 1 | 1 | 1 | 1238 | 136,7 | 5,57 | 2,14 | 1 | 1,314 | 1,163 | 0,491 | 0,684 | 0,452 | 0,01 | 0,965 | 1,002 | 1,50 | 0,50 | 3,6E-01 |
| O35250 | Exocyst complex component 7 OS=Mus musculus OX=10090 GN=Ex | 13 | 9 | 9 | 9 | 697 | 79,9 | 6,98 | 13,09 | 9 | 0,868 | 1,009 | 1,04 | 0,972 | 0,982 | 1,177 | 1,005 | 0,97 | 0,94 | -0,06 | 3,6E-01 |
| Q99P72 | Reticulon-4 OS=Mus musculus OX=10090 GN=Rtn4 PE=1 SV=2 | 15 | 15 | 18 | 15 | 1162 | 126,5 | 4,54 | 31,78 | 15 | 0,895 | 0,922 | 0,917 | 0,942 | 0,904 | 0,886 | 1,076 | 0,989 | 0,95 | -0,05 | 3,6E-01 |
| Q62283 | Tetraspanin-7 OS=Mus musculus OX=10090 GN=Tspan7 PE=1 SV=2 | 10 | 2 | 3 | 2 | 249 | 27,5 | 7,2 | 2,89 | 2 | 0,613 | 1,195 | 0,78 | 0,86 | 0,848 | 0,549 | 0,783 | 0,715 | 1,19 | 0,19 | 3,6E-01 |
| P63325 | 40S ribosomal protein S10 OS=Mus musculus OX=10090 GN=Rps10 | 5 | 1 | 1 | 1 | 165 | 18,9 | 10,15 | 2,16 | 1 | 1,173 | 1,159 | 0,96 | 0,975 | 1,099 | 1,023 | 0,888 | 0,966 | 1,07 | 0,07 | 3,6E-01 |
| P41105 | 60S ribosomal protein L28 OS=Mus musculus OX=10090 GN=Rpl28 | 9 | 2 | 2 | 2 | 137 | 15,7 | 12,02 | 0 | 2 | 0,96 | 0,838 | 0,845 | 0,767 | 0,945 | 0,628 | 0,717 | 0,806 | 1,10 | 0,10 | 3,6E-01 |
| Q8BTH8 | Casein kinase I isoform gamma-1 OS=Mus musculus OX=10090 GN= | 5 | 2 | 2 | 1 | 459 | 52,7 | 9,01 | 4,76 | 2 | 0,704 | 0,846 | 0,831 | 0,872 | 0,816 | 0,715 | 1,079 | 1,013 | 0,90 | -0,10 | 3,6E-01 |
| B9EJA2 | Cortactin-binding protein 2 OS=Mus musculus OX=10090 GN=Cttnb | 10 | 14 | 17 | 14 | 1648 | 178,7 | 8,19 | 34,72 | 14 | 1,001 | 0,92 | 1,02 | 0,973 | 0,993 | 0,983 | 1,001 | 1,035 | 0,98 | -0,02 | 3,6E-01 |
| Q3U0V1 | Far upstream element-binding protein 2 OS=Mus musculus OX=100 | 2 | 1 | 1 | 1 | 748 | 76,7 | 7,33 | 2,76 | 1 | 1,006 | 1,421 | 1,382 | 0,704 | 0,848 | 1,229 | 0,697 | 0,928 | 1,22 | 0,22 | 3,6E-01 |
| Q6ZQF0 | DNA topoisomerase 2-binding protein 1 OS=Mus musculus OX=100 | 1 | 1 | 1 | 1 | 1515 | 168,8 | 6,67 | 1,78 | 1 | 1,992 | 1,564 | 1,249 | 1,009 | 0,9 | 1,619 | 0,941 | 1,275 | 1,23 | 0,23 | 3,6E-01 |
| Q9WV96 | Mitochondrial import inner membrane translocase subunit Tim10 E | 44 | 3 | 3 | 3 | 100 | 11,3 | 7,08 | 9,04 | 3 | 1,071 | 1,01 | 0,898 | 0,908 | 0,995 | 1,028 | 1,04 | 0,996 | 0,96 | -0,04 | 3,6E-01 |
| Q8VEH5 | EPM2A-interacting protein 1 OS=Mus musculus OX=10090 GN=Epn | 10 | 6 | 6 | 6 | 606 | 70,1 | 5,87 | 9,59 | 6 | 0,993 | 0,976 | 1,113 | 1,08 | 1,062 | 1,024 | 0,932 | 0,969 | 1,04 | 0,04 | 3,6E-01 |
| Q9QXK3 | Coatomer subunit gamma-2 OS=Mus musculus OX=10090 GN=Copg | 3 | 2 | 2 | 2 | 871 | 97,6 | 5,8 | 5,63 | 2 | 1,346 | 0,986 | 0,858 | 0,883 | 0,952 | 1,025 | 0,784 | 0,808 | 1,14 | 0,14 | 3,6E-01 |
| Q60738 | Zinc transporter 1 OS=Mus musculus OX=10090 GN=Slc30a1 PE=1 S | 12 | 5 | 5 | 5 | 503 | 54,7 | 6,62 | 4,6 | 5 | 1,011 | 1,038 | 0,868 | 0,965 | 0,893 | 1,053 | 1,072 | 1,102 | 0,94 | -0,06 | 3,6E-01 |
| P19096 | Fatty acid synthase OS=Mus musculus OX=10090 GN=Fasn PE=1 SV | 15 | 34 | 47 | 34 | 2504 | 272,3 | 6,58 | 96,09 | 34 | 0,976 | 0,986 | 1,014 | 0,955 | 0,955 | 0,918 | 0,996 | 0,979 | 1,02 | 0,02 | 3,6E-01 |
| Q3TVA9 | Coiled-coil domain-containing protein 136 OS=Mus musculus OX=1 | 8 | 7 | 7 | 7 | 1136 | 131,8 | 4,84 | 16,97 | 7 | 1,135 | 1,207 | 1,148 | 0,922 | 0,956 | 1,028 | 1,053 | 1,1 | 1,07 | 0,07 | 3,6E-01 |
| P47199 | Quinone oxidoreductase OS=Mus musculus OX=10090 GN=Cryz PE= | 8 | 2 | 2 | 2 | 331 | 35,2 | 8,07 | 4,72 | 2 | 0,98 | 0,794 | 0,97 | 0,974 | 0,89 | 0,953 | 1,039 | 1,086 | 0,94 | -0,06 | 3,6E-01 |
| Q80Y17 | Lethal(2) giant larvae protein homolog 1 OS=Mus musculus OX=100 | 9 | 7 | 7 | 7 | 1036 | 112,5 | 6,48 | 14,87 | 7 | 0,973 | 0,954 | 1,25 | 1,004 | 0,843 | 1,1 | 0,977 | 0,911 | 1,09 | 0,09 | 3,6E-01 |
| Q3UV17 | Keratin, type II cytoskeletal 2 oral OS=Mus musculus OX=10090 GN | 7 | 6 | 10 | 2 | 594 | 62,8 | 8,43 | 20,54 | 6 | 0,682 | 2,339 | 0,851 | 0,564 | 1,19 | 0,441 | 0,442 | 0,571 | 1,68 | 0,68 | 3,6E-01 |
| E9PVA8 | eIF-2-alpha kinase activator GCN1 OS=Mus musculus OX=10090 GN | 1 | 3 | 3 | 3 | 2671 | 292,8 | 7,36 | 2,32 | 3 | 0,991 | 1,244 | 0,749 | 0,729 | 0,878 | 0,85 | 0,698 | 0,781 | 1,16 | 0,16 | 3,6E-01 |
| P68368 | Tubulin alpha-4A chain OS=Mus musculus OX=10090 GN=Tuba4a P | 54 | 25 | 100 | 5 | 448 | 49,9 | 5,06 | 274,69 | 25 | 0,994 | 0,979 | 0,966 | 0,911 | 0,963 | 1,06 | 1,01 | 0,943 | 0,97 | -0,03 | 3,6E-01 |
| Q9WTT4 | V-type proton ATPase subunit G 2 OS=Mus musculus OX=10090 GN | 47 | 8 | 16 | 8 | 118 | 13,6 | 10,26 | 31,58 | 8 | 0,768 | 0,858 | 0,886 | 1,03 | 0,912 | 0,964 | 0,893 | 1,014 | 0,94 | -0,06 | 3,6E-01 |
| O09131 | Glutathione S-transferase omega-1 OS=Mus musculus OX=10090 G | 11 | 3 | 4 | 3 | 240 | 27,5 | 7,36 | 10,18 | 3 | 0,785 | 0,897 | 0,761 | 0,932 | 0,974 | 0,775 | 0,96 | 0,91 | 0,93 | -0,07 | 3,6E-01 |
| Q11011 | Puromycin-sensitive aminopeptidase OS=Mus musculus OX=10090 | 24 | 22 | 34 | 22 | 920 | 103,3 | 5,9 | 69,58 | 22 | 1,023 | 1,009 | 1,026 | 1,032 | 0,954 | 1,024 | 1,048 | 0,977 | 1,02 | 0,02 | 3,6E-01 |
| Q9D6P8 | Calmodulin-like protein 3 OS=Mus musculus OX=10090 GN=Calml3 | 27 | 7 | 12 | 1 | 149 | 16,7 | 4,31 | 23,2 | 7 | 1,72 | 1,606 | 1,132 | 0,649 | 0,661 | 1,595 | 0,699 | 0,858 | 1,34 | 0,34 | 3,6E-01 |
| A2ADY9 | Protein DDI1 homolog 2 OS=Mus musculus OX=10090 GN=Ddi2 PE= | 6 | 2 | 2 | 2 | 399 | 44,6 | 5,05 | 5,64 | 2 | 0,741 | 0,802 | 0,947 | 0,961 | 0,753 | 0,812 | 0,901 | 0,72 | 1,08 | 0,08 | 3,6E-01 |
| Q8BYL4 | Tyrosine--tRNA ligase, mitochondrial OS=Mus musculus OX=10090 | 8 | 4 | 4 | 4 | 472 | 52,6 | 9,16 | 9,16 | 4 | 0,934 | 1,053 | 1,018 | 0,963 | 1,004 | 0,875 | 0,89 | 1,018 | 1,05 | 0,05 | 3,6E-01 |
| Q3V0K9 | Plastin-1 OS=Mus musculus OX=10090 GN=Pls1 PE=1 SV=1 | 2 | 1 | 1 | 1 | 630 | 70,4 | 5,38 | 2,56 | 1 | 1,004 | 1,028 | 1,169 | 1,015 | 1,013 | 0,981 | 1,123 | 0,774 | 1,08 | 0,08 | 3,6E-01 |
| P38585 | Tubulin--tyrosine ligase OS=Mus musculus OX=10090 GN=Ttl PE=1 | 7 | 2 | 2 | 2 | 377 | 43,1 | 6,46 | 4,29 | 2 | 1,432 | 1,338 | 1,131 | 0,885 | 0,855 | 1,076 | 1,215 | 1,079 | 1,13 | 0,13 | 3,6E-01 |
| P84099 | 60S ribosomal protein L19 OS=Mus musculus OX=10090 GN=Rpl19 | 14 | 3 | 3 | 3 | 196 | 23,5 | 11,47 | 8,36 | 3 | 0,971 | 0,921 | 0,905 | 0,797 | 0,911 | 0,887 | 0,834 | 0,775 | 1,05 | 0,05 | 3,6E-01 |

| | | | | | | | | | | | | | | | | | | | | | |
|---|---|---|---|---|---|---|---|---|---|---|---|---|---|---|---|---|---|---|---|---|---|
| O54784 | Death-associated protein kinase 3 OS=Mus musculus OX=10090 GN | 3 | 1 | 1 | 1 | 448 | 51,4 | 8,7 | 0 | 1 | 0,657 | 1,512 | 1,027 | 1,189 | 0,981 | 2,193 | 1,348 | 1,136 | 0,78 | -0,22 | 3,6E-01 |
| P97411 | Islet cell autoantigen 1 OS=Mus musculus OX=10090 GN=Ica1 PE=1 | 12 | 5 | 5 | 5 | 478 | 54,3 | 5,8 | 9,46 | 5 | 0,669 | 1,016 | 0,995 | 1,016 | 1,058 | 0,912 | 1,086 | 1,007 | 0,91 | -0,09 | 3,6E-01 |
| Q8VE22 | 28S ribosomal protein S23, mitochondrial OS=Mus musculus OX=10 | 21 | 3 | 4 | 3 | 177 | 20,3 | 8,59 | 9,27 | 3 | 1,29 | 1,336 | 1,251 | 0,855 | 0,799 | 1,307 | 1,016 | 1,012 | 1,14 | 0,14 | 3,6E-01 |
| Q80X95 | Ras-related GTP-binding protein A OS=Mus musculus OX=10090 GN | 6 | 2 | 2 | 2 | 313 | 36,5 | 7,72 | 4,71 | 2 | 1,147 | 1,016 | 1,141 | 0,922 | 1,412 | 1,094 | 0,998 | 1,131 | 0,91 | -0,09 | 3,6E-01 |
| Q8BVE3 | V-type proton ATPase subunit H OS=Mus musculus OX=10090 GN=A | 27 | 12 | 26 | 12 | 483 | 55,8 | 6,61 | 47,11 | 12 | 0,943 | 0,942 | 0,934 | 1,031 | 0,97 | 0,972 | 1,041 | 0,979 | 0,97 | -0,03 | 3,6E-01 |
| Q8R071 | Inositol-trisphosphate 3-kinase A OS=Mus musculus OX=10090 GN= | 23 | 10 | 17 | 10 | 459 | 50,9 | 7,72 | 37,09 | 10 | 0,983 | 0,983 | 0,977 | 1,022 | 1,037 | 0,993 | 1,016 | 0,982 | 0,98 | -0,02 | 3,6E-01 |
| Q8BHL8 | Proteasome inhibitor PI31 subunit OS=Mus musculus OX=10090 GN | 5 | 1 | 1 | 1 | 271 | 29,6 | 5,25 | 3,3 | 1 | 1,079 | 0,886 | 0,806 | 0,842 | 0,771 | 0,616 | 0,931 | 0,919 | 1,12 | 0,12 | 3,6E-01 |
| E9PZ19 | Protein turtle homolog B OS=Mus musculus OX=10090 GN=Igsf9b P | 3 | 4 | 4 | 4 | 1328 | 144,9 | 6,7 | 7,81 | 4 | 0,807 | 1,031 | 0,873 | 0,885 | 0,907 | 1,059 | 0,894 | 0,973 | 0,94 | -0,06 | 3,7E-01 |
| Q8K1M6 | Dynamin-1-like protein OS=Mus musculus OX=10090 GN=Dnm1l PE | 30 | 20 | 29 | 20 | 742 | 82,6 | 7,05 | 76,57 | 20 | 1,083 | 1,075 | 1,011 | 0,983 | 0,963 | 1,078 | 1,001 | 0,969 | 1,04 | 0,04 | 3,7E-01 |
| P55088 | Aquaporin-4 OS=Mus musculus OX=10090 GN=Aqp4 PE=1 SV=2 | 6 | 2 | 3 | 2 | 323 | 34,4 | 7,42 | 3,99 | 2 | 0,71 | 0,594 | 0,857 | 0,676 | 0,926 | 0,654 | 0,768 | 0,796 | 0,90 | -0,10 | 3,7E-01 |
| P04104 | Keratin, type II cytoskeletal 1 OS=Mus musculus OX=10090 GN=Krt | 5 | 4 | 11 | 2 | 637 | 65,6 | 8,15 | 31,76 | 4 | 1,219 | 2,066 | 0,826 | 0,462 | 1,113 | 0,825 | 0,475 | 0,715 | 1,46 | 0,46 | 3,7E-01 |
| Q80TB8 | Synaptic vesicle membrane protein VAT-1 homolog-like OS=Mus m | 11 | 5 | 13 | 5 | 417 | 45,8 | 5,06 | 14,94 | 5 | 0,76 | 0,763 | 0,844 | 1,042 | 0,809 | 0,881 | 1,193 | 0,943 | 0,89 | -0,11 | 3,7E-01 |
| Q9ERS2 | NADH dehydrogenase [ubiquinone] 1 alpha subcomplex subunit 13 | 42 | 8 | 17 | 8 | 144 | 16,8 | 9,48 | 15,67 | 8 | 1,413 | 1,176 | 1,09 | 0,976 | 0,939 | 1,24 | 1,009 | 1,024 | 1,11 | 0,11 | 3,7E-01 |
| P59281 | Rho GTPase-activating protein 39 OS=Mus musculus OX=10090 GN | 6 | 6 | 7 | 6 | 1107 | 125,1 | 7,97 | 13,3 | 6 | 0,78 | 0,818 | 0,953 | 1,012 | 1,016 | 0,826 | 0,973 | 1,03 | 0,93 | -0,07 | 3,7E-01 |
| O88545 | COP9 signalosome complex subunit 6 OS=Mus musculus OX=10090 | 14 | 3 | 3 | 3 | 324 | 35,9 | 5,73 | 9,46 | 3 | 0,929 | 0,979 | 0,901 | 0,833 | 0,956 | 0,889 | 0,964 | 0,974 | 0,96 | -0,04 | 3,7E-01 |
| Q922S4 | cGMP-dependent 3',5'-cyclic phosphodiesterase OS=Mus musculus | 21 | 15 | 21 | 15 | 939 | 105,6 | 5,41 | 52,86 | 15 | 0,917 | 0,872 | 1,016 | 1,006 | 1,017 | 0,888 | 1,128 | 1,013 | 0,94 | -0,06 | 3,7E-01 |
| P61164 | Alpha-centractin OS=Mus musculus OX=10090 GN=Actr1a PE=1 SV= | 25 | 8 | 11 | 3 | 376 | 42,6 | 6,64 | 25,39 | 8 | 1,284 | 0,838 | 1,016 | 1,017 | 0,8 | 1,043 | 0,945 | 0,958 | 1,11 | 0,11 | 3,7E-01 |
| Q6PDM2 | Serine/arginine-rich splicing factor 1 OS=Mus musculus OX=10090 ( | 4 | 1 | 1 | 1 | 248 | 27,7 | 10,36 | 1,94 | 1 | 1,161 | 1,005 | 1,205 | 0,974 | 1,056 | 1,007 | 0,787 | 1,131 | 1,09 | 0,09 | 3,7E-01 |
| Q8R2H9 | Phosphoethanolamine/phosphocholine phosphatase OS=Mus musc | 17 | 4 | 4 | 4 | 267 | 29,9 | 7,94 | 8,31 | 4 | 1,176 | 1,073 | 1,011 | 0,934 | 0,937 | 1,084 | 0,963 | 0,974 | 1,06 | 0,06 | 3,7E-01 |
| P17182 | Alpha-enolase OS=Mus musculus OX=10090 GN=Eno1 PE=1 SV=3 | 50 | 24 | 64 | 22 | 434 | 47,1 | 6,8 | 160,01 | 24 | 0,618 | 0,686 | 0,848 | 0,945 | 1,165 | 0,349 | 0,389 | 0,364 | 1,37 | 0,37 | 3,7E-01 |
| Q91YW3 | DnaJ homolog subfamily C member 3 OS=Mus musculus OX=10090 | 9 | 4 | 4 | 4 | 504 | 57,4 | 5,85 | 2,44 | 4 | 0,804 | 0,803 | 1,099 | 1,136 | 1,122 | 1,054 | 0,89 | 1,223 | 0,90 | -0,10 | 3,7E-01 |
| P08730 | Keratin, type I cytoskeletal 13 OS=Mus musculus OX=10090 GN=Krt | 10 | 6 | 13 | 1 | 437 | 47,7 | 4,86 | 24,46 | 6 | 0,802 | 0,818 | 0,641 | 0,584 | 1,762 | 0,577 | 0,641 | 0,949 | 0,72 | -0,28 | 3,7E-01 |
| Q8K3A0 | Iron-sulfur cluster co-chaperone protein HscB OS=Mus musculus O | 15 | 4 | 4 | 4 | 234 | 26,6 | 7,37 | 8,08 | 4 | 1,155 | 1,017 | 1,009 | 0,878 | 0,863 | 1,058 | 0,899 | 0,963 | 1,07 | 0,07 | 3,7E-01 |
| Q91VE0 | Long-chain fatty acid transport protein 4 OS=Mus musculus OX=10 | 12 | 6 | 6 | 6 | 643 | 72,3 | 8,59 | 14,12 | 6 | 1,056 | 0,789 | 1,017 | 0,896 | 0,995 | 1,005 | 0,946 | 1,067 | 0,94 | -0,06 | 3,7E-01 |
| Q8K2K6 | Arf-GAP domain and FG repeat-containing protein 1 OS=Mus musc | 10 | 5 | 5 | 5 | 561 | 58 | 8,63 | 6,09 | 5 | 1,084 | 1,034 | 1,075 | 0,915 | 0,988 | 0,924 | 1,037 | 0,983 | 1,04 | 0,04 | 3,7E-01 |
| D3Z7P3 | Glutaminase kidney isoform, mitochondrial OS=Mus musculus OX= | 27 | 16 | 30 | 16 | 674 | 73,9 | 7,99 | 60,59 | 16 | 1,081 | 1,002 | 0,963 | 0,932 | 0,983 | 1,157 | 0,984 | 1,057 | 0,95 | -0,05 | 3,7E-01 |
| Q9QYF9 | Protein NDRG3 OS=Mus musculus OX=10090 GN=Ndrg3 PE=1 SV=1 | 19 | 7 | 10 | 6 | 375 | 41,5 | 5,25 | 14,56 | 7 | 1,276 | 1,166 | 1,137 | 1,047 | 0,883 | 1,285 | 0,961 | 1,107 | 1,09 | 0,09 | 3,7E-01 |
| Q9WVK8 | Cholesterol 24-hydroxylase OS=Mus musculus OX=10090 GN=Cyp4 | 9 | 5 | 5 | 5 | 500 | 56,8 | 8,82 | 10,31 | 5 | 1,016 | 1,061 | 1,1 | 1,05 | 1,121 | 1,077 | 1,047 | 1,072 | 0,98 | -0,02 | 3,7E-01 |
| Q8C633 | Calcium-binding and spermatid-specific protein 1 OS=Mus musculu | 2 | 1 | 1 | 1 | 391 | 42,3 | 4,18 | 2,42 | 1 | 1,333 | 0,879 | 1,016 | 1,159 | 0,842 | 1,173 | 0,837 | 1,041 | 1,13 | 0,13 | 3,7E-01 |
| Q8K0T0 | Reticulon-1 OS=Mus musculus OX=10090 GN=Rtn1 PE=1 SV=1 | 18 | 14 | 23 | 13 | 780 | 83,5 | 4,58 | 51,03 | 14 | 0,899 | 0,862 | 0,916 | 0,957 | 0,908 | 0,872 | 1,106 | 0,962 | 0,94 | -0,06 | 3,7E-01 |
| Q64669 | NAD(P)H dehydrogenase [quinone] 1 OS=Mus musculus OX=10090 | 5 | 1 | 1 | 1 | 274 | 30,9 | 8,72 | 0 | 1 | 1,315 | 1,241 | 0,834 | 0,969 | 1,005 | 0,787 | 0,873 | 1,155 | 1,14 | 0,14 | 3,7E-01 |
| Q8BNW9 | Kelch repeat and BTB domain-containing protein 11 OS=Mus musc | 14 | 7 | 8 | 7 | 633 | 67,9 | 5,39 | 17,49 | 7 | 0,844 | 0,945 | 0,9 | 0,901 | 0,888 | 0,894 | 0,939 | 0,993 | 0,97 | -0,03 | 3,7E-01 |
| Q9CYW4 | Haloacid dehalogenase-like hydrolase domain-containing protein 3 | 4 | 1 | 1 | 1 | 251 | 28 | 6,8 | 1,67 | 1 | 1,363 | 1,24 | 1,05 | 0,881 | 0,79 | 1,112 | 0,953 | 1,156 | 1,13 | 0,13 | 3,7E-01 |
| P19783 | Cytochrome c oxidase subunit 4 isoform 1, mitochondrial OS=Mus | 60 | 13 | 23 | 13 | 169 | 19,5 | 9,23 | 53,76 | 13 | 0,811 | 0,81 | 0,749 | 0,962 | 1,221 | 0,623 | 0,981 | 1,021 | 0,87 | -0,13 | 3,7E-01 |
| Q3TL44 | NLR family member X1 OS=Mus musculus OX=10090 GN=Nlrx1 PE= | 5 | 4 | 5 | 4 | 975 | 107,8 | 7,37 | 12,17 | 4 | 0,771 | 0,917 | 1,168 | 1,058 | 1,051 | 1,059 | 1,043 | 1,097 | 0,92 | -0,08 | 3,7E-01 |
| Q8K2G4 | Bardet-Biedl syndrome 7 protein homolog OS=Mus musculus OX=1 | 2 | 1 | 1 | 1 | 715 | 80,3 | 5,81 | 2,06 | 1 | 0,862 | 1,013 | 1,098 | 0,994 | 1,112 | 1,42 | 1,083 | 0,845 | 0,89 | -0,11 | 3,7E-01 |
| Q9QXV0 | ProSAAS OS=Mus musculus OX=10090 GN=Pcsk1n PE=1 SV=2 | 28 | 6 | 7 | 6 | 258 | 27,3 | 5,85 | 17,74 | 6 | 0,917 | 0,932 | 0,957 | 1,044 | 0,896 | 1,245 | 1,076 | 0,951 | 0,92 | -0,08 | 3,7E-01 |
| Q9Z1F9 | SUMO-activating enzyme subunit 2 OS=Mus musculus OX=10090 G | 4 | 2 | 2 | 2 | 638 | 70,5 | 5,24 | 4,74 | 2 | 1,103 | 1,059 | 1,333 | 0,788 | 1,242 | 1,17 | 1,255 | 1,075 | 0,90 | -0,10 | 3,7E-01 |
| Q8C726 | BTB/POZ domain-containing protein 9 OS=Mus musculus OX=1009( | 1 | 1 | 1 | 1 | 612 | 69,1 | 5,68 | 1,61 | 1 | 1,318 | 0,894 | 1,084 | 0,952 | 1,193 | 1,119 | 1,217 | 1,098 | 0,92 | -0,08 | 3,7E-01 |
| Q61165 | Sodium/hydrogen exchanger 1 OS=Mus musculus OX=10090 GN=Sl | 6 | 4 | 6 | 4 | 820 | 91,4 | 7,12 | 10,68 | 4 | 1,446 | 1,132 | 1,093 | 1,114 | 0,96 | 1,165 | 1,049 | 1,219 | 1,09 | 0,09 | 3,7E-01 |
| P53986 | Monocarboxylate transporter 1 OS=Mus musculus OX=10090 GN=S | 7 | 2 | 2 | 2 | 493 | 53,2 | 7,47 | 3,14 | 2 | 0,722 | 0,735 | 0,999 | 0,841 | 0,736 | 0,851 | 1,064 | 1,033 | 0,89 | -0,11 | 3,7E-01 |
| Q9CQX8 | 28S ribosomal protein S36, mitochondrial OS=Mus musculus OX=10 | 47 | 4 | 9 | 4 | 102 | 11,1 | 9,99 | 17,25 | 4 | 1,029 | 0,921 | 0,849 | 0,926 | 0,914 | 1,091 | 1,084 | 0,888 | 0,94 | -0,06 | 3,7E-01 |
| Q3THK7 | GMP synthase [glutamine-hydrolyzing] OS=Mus musculus OX=1009 | 6 | 4 | 6 | 4 | 693 | 76,7 | 6,73 | 14,45 | 4 | 0,833 | 0,921 | 0,922 | 0,899 | 0,909 | 0,844 | 0,948 | 1,087 | 0,94 | -0,06 | 3,7E-01 |
| P67871 | Casein kinase II subunit beta OS=Mus musculus OX=10090 GN=Csn | 23 | 4 | 5 | 4 | 215 | 24,9 | 5,55 | 9,49 | 4 | 1,372 | 1,14 | 1,074 | 1,17 | 1,03 | 1,198 | 0,903 | 1,239 | 1,09 | 0,09 | 3,8E-01 |
| Q9D1P4 | Cysteine and histidine-rich domain-containing protein 1 OS=Mus m | 21 | 4 | 4 | 4 | 331 | 37,3 | 7,9 | 2,8 | 4 | 1,247 | 0,885 | 0,852 | 0,997 | 0,713 | 0,7 | 0,935 | 1,12 | 1,15 | 0,15 | 3,8E-01 |
| O54865 | Guanylate cyclase soluble subunit beta-1 OS=Mus musculus OX=10 | 11 | 7 | 9 | 7 | 620 | 70,6 | 5,31 | 22,77 | 7 | 1,142 | 1,002 | 1,067 | 1,069 | 1,033 | 1,243 | 1,083 | 1,124 | 0,95 | -0,05 | 3,8E-01 |
| P62242 | 40S ribosomal protein S8 OS=Mus musculus OX=10090 GN=Rps8 PE | 27 | 5 | 8 | 5 | 208 | 24,2 | 10,32 | 22,79 | 5 | 0,991 | 0,979 | 1,001 | 0,92 | 1,065 | 0,968 | 0,995 | 0,973 | 0,97 | -0,03 | 3,8E-01 |
| Q03265 | ATP synthase subunit alpha, mitochondrial OS=Mus musculus OX=1 | 49 | 30 | 118 | 30 | 553 | 59,7 | 9,19 | 304,53 | 30 | 0,989 | 1,081 | 1,013 | 0,947 | 0,981 | 0,998 | 0,974 | 0,967 | 1,03 | 0,03 | 3,8E-01 |
| Q9D1E8 | 1-acyl-sn-glycerol-3-phosphate acyltransferase epsilon OS=Mus mu | 6 | 2 | 2 | 2 | 365 | 42,2 | 9,38 | 4,66 | 2 | 1,22 | 0,868 | 1,105 | 0,805 | 0,957 | 0,994 | 0,873 | 0,746 | 1,12 | 0,12 | 3,8E-01 |
| O08914 | Fatty-acid amide hydrolase 1 OS=Mus musculus OX=10090 GN=Faa | 8 | 4 | 4 | 4 | 579 | 63,2 | 7,87 | 7,14 | 4 | 1,371 | 1,232 | 1,166 | 0,882 | 0,918 | 1,133 | 1,109 | 1,058 | 1,10 | 0,10 | 3,8E-01 |
| Q99JX3 | Golgi reassembly-stacking protein 2 OS=Mus musculus OX=10090 ( | 4 | 1 | 1 | 1 | 451 | 47 | 4,79 | 3,92 | 1 | 1,35 | 0,901 | 1,094 | 1,085 | 0,938 | 1,033 | 0,982 | 1,101 | 1,09 | 0,09 | 3,8E-01 |
| Q64299 | CCN family member 3 OS=Mus musculus OX=10090 GN=Ccn3 PE=1 | 3 | 1 | 1 | 1 | 354 | 38,9 | 7,64 | 1,76 | 1 | 0,729 | 1,086 | 0,817 | 0,685 | 0,772 | 0,577 | 0,928 | 0,564 | 1,17 | 0,17 | 3,8E-01 |
| Q6P2L7 | Protein CASC4 OS=Mus musculus OX=10090 GN=Casc4 PE=1 SV=1 | 2 | 1 | 1 | 1 | 435 | 49,4 | 5,45 | 1,88 | 1 | 0,884 | 1,244 | 1,003 | 0,81 | 0,798 | 0,904 | 0,589 | 1,104 | 1,16 | 0,16 | 3,8E-01 |
| P21836 | Acetylcholinesterase OS=Mus musculus OX=10090 GN=Ache PE=1 S | 3 | 2 | 3 | 2 | 614 | 68,1 | 6,33 | 4,72 | 2 | 1,116 | 0,57 | 0,851 | 0,932 | 1,014 | 1,35 | 0,978 | 0,761 | 0,85 | -0,15 | 3,8E-01 |
| Q9DBF1 | Alpha-aminoadipic semialdehyde dehydrogenase OS=Mus musculu | 24 | 10 | 12 | 10 | 539 | 58,8 | 7,47 | 16,95 | 10 | 1,055 | 0,942 | 0,981 | 0,904 | 0,964 | 1,13 | 0,946 | 1,044 | 0,95 | -0,05 | 3,8E-01 |
| Q61140 | Breast cancer anti-estrogen resistance protein 1 OS=Mus musculus | 1 | 1 | 2 | 1 | 874 | 94,2 | 5,68 | 0 | 1 | 1,038 | 1,092 | 0,852 | 1,625 | 0,807 | 1,201 | 0,935 | 0,959 | 1,18 | 0,18 | 3,8E-01 |
| Q9CQL6 | 39S ribosomal protein L35, mitochondrial OS=Mus musculus OX=10 | 4 | 1 | 1 | 1 | 188 | 21,5 | 11,47 | 2,34 | 1 | 1,098 | 1,142 | 0,942 | 0,947 | 0,802 | 1,187 | 0,874 | 0,887 | 1,10 | 0,10 | 3,8E-01 |
| O89094 | Caspase-14 OS=Mus musculus OX=10090 GN=Casp14 PE=1 SV=1 | 2 | 1 | 1 | 1 | 257 | 29,4 | 4,86 | 2,08 | 1 | 1,447 | 1,297 | 1,055 | 0,581 | 0,83 | 1,189 | 0,8 | 0,744 | 1,23 | 0,23 | 3,8E-01 |
| Q9CQI6 | Coactosin-like protein OS=Mus musculus OX=10090 GN=Cotl1 PE=1 | 25 | 4 | 4 | 4 | 142 | 15,9 | 5,4 | 10,48 | 4 | 0,666 | 0,861 | 1,002 | 0,957 | 1,013 | 0,811 | 1,121 | 0,92 | 0,90 | -0,10 | 3,8E-01 |
| Q9DB15 | 39S ribosomal protein L12, mitochondrial OS=Mus musculus OX=10 | 24 | 4 | 6 | 4 | 201 | 21,7 | 9,29 | 12,31 | 4 | 1,146 | 1,013 | 1,277 | 1,081 | 1,071 | 1,031 | 1,081 | 1,111 | 1,05 | 0,05 | 3,8E-01 |
| O35143 | ATPase inhibitor, mitochondrial OS=Mus musculus OX=10090 GN=A | 33 | 4 | 7 | 4 | 106 | 12,2 | 9,64 | 10,16 | 4 | 0,754 | 0,841 | 0,978 | 0,991 | 0,992 | 0,837 | 1,021 | 0,981 | 0,93 | -0,07 | 3,8E-01 |
| Q3UFY7 | 7-methylguanosine phosphate-specific 5'-nucleotidase OS=Mus mu | 4 | 1 | 1 | 1 | 300 | 34,4 | 6,24 | 2,09 | 1 | 0,845 | 0,859 | 0,856 | 0,834 | 1,082 | 0,989 | 0,746 | 0,858 | 0,92 | -0,08 | 3,8E-01 |

| | | | | | | | | | | | | | | | | | | | | | |
|---|---|---|---|---|---|---|---|---|---|---|---|---|---|---|---|---|---|---|---|---|---|
| Q9D5R2 | WD repeat-containing protein 20 OS=Mus musculus OX=10090 GN | 2 | 1 | 1 | 1 | 567 | 62,7 | 8,48 | 1,98 | 1 | 1,753 | 1,549 | 1,115 | 0,891 | 0,941 | 1,431 | 0,977 | 1,098 | 1,19 | 0,19 | 3,8E-01 |
| Q6P4S6 | Serine/threonine-protein kinase SIK3 OS=Mus musculus OX=10090 | 4 | 5 | 5 | 5 | 1311 | 145,7 | 6,92 | 7,21 | 5 | 1,045 | 1,172 | 1,042 | 1,025 | 0,917 | 1,182 | 1,009 | 0,898 | 1,07 | 0,07 | 3,8E-01 |
| Q3UHC7 | Disabled homolog 2-interacting protein OS=Mus musculus OX=100 | 5 | 6 | 6 | 3 | 1189 | 131,6 | 8,72 | 10,1 | 6 | 1,054 | 0,968 | 1,13 | 1,051 | 1,127 | 1,08 | 1,008 | 1,17 | 0,96 | -0,04 | 3,8E-01 |
| Q80YD1 | ATP-dependent RNA helicase SUPV3L1, mitochondrial OS=Mus mu | 4 | 2 | 2 | 2 | 779 | 87 | 7,84 | 5,1 | 2 | 1,617 | 1,304 | 1,347 | 1,061 | 0,833 | 1,066 | 1,266 | 1,484 | 1,15 | 0,15 | 3,8E-01 |
| Q61187 | Tumor susceptibility gene 101 protein OS=Mus musculus OX=1009 | 15 | 5 | 6 | 5 | 391 | 44,1 | 6,71 | 11,14 | 5 | 0,874 | 0,995 | 0,835 | 0,987 | 0,894 | 1,008 | 0,976 | 0,993 | 0,95 | -0,05 | 3,8E-01 |
| Q6NZL0 | Protein SOGA3 OS=Mus musculus OX=10090 GN=Soga3 PE=1 SV=2 | 16 | 16 | 23 | 16 | 945 | 103,4 | 6,14 | 41,46 | 16 | 1,17 | 1,109 | 1,045 | 1,051 | 1,028 | 1,118 | 1,027 | 1,065 | 1,03 | 0,03 | 3,8E-01 |
| Q91XF0 | Pyridoxine-5'-phosphate oxidase OS=Mus musculus OX=10090 GN | 23 | 4 | 4 | 4 | 261 | 30,1 | 8,22 | 4,94 | 4 | 0,678 | 0,987 | 0,898 | 0,708 | 0,884 | 0,982 | 0,76 | 0,993 | 0,90 | -0,10 | 3,8E-01 |
| Q91W53 | Golgin subfamily A member 7 OS=Mus musculus OX=10090 GN=Go | 14 | 2 | 2 | 2 | 137 | 15,8 | 7,05 | 4,72 | 2 | 1,336 | 1,043 | 0,846 | 0,992 | 0,812 | 1,021 | 1 | 0,957 | 1,11 | 0,11 | 3,8E-01 |
| P54728 | UV excision repair protein RAD23 homolog B OS=Mus musculus OX | 14 | 6 | 8 | 6 | 416 | 43,5 | 4,83 | 19,49 | 6 | 0,837 | 0,933 | 0,963 | 0,991 | 0,959 | 0,92 | 0,992 | 0,996 | 0,96 | -0,04 | 3,8E-01 |
| Q9CQJ8 | NADH dehydrogenase [ubiquinone] 1 beta subcomplex subunit 9 O | 30 | 4 | 9 | 4 | 179 | 22 | 7,8 | 19,7 | 4 | 1,394 | 1,21 | 1,129 | 1,059 | 1,044 | 1,159 | 1,127 | 1,169 | 1,07 | 0,07 | 3,8E-01 |
| Q3UDP0 | WD repeat-containing protein 41 OS=Mus musculus OX=10090 GN | 6 | 2 | 2 | 2 | 460 | 51,5 | 5,15 | 5,31 | 2 | 1,801 | 1,231 | 1,266 | 1,017 | 0,886 | 1,435 | 1,019 | 1,202 | 1,17 | 0,17 | 3,8E-01 |
| P63080 | Gamma-aminobutyric acid receptor subunit beta-3 OS=Mus muscu | 15 | 6 | 7 | 3 | 473 | 54,1 | 9,1 | 17,07 | 6 | 1,426 | 1,347 | 1,2 | 0,981 | 0,925 | 1,418 | 1,07 | 0,985 | 1,13 | 0,13 | 3,8E-01 |
| Q8K0P3 | MTOR-associated protein MEAK7 OS=Mus musculus OX=10090 GN | 2 | 1 | 2 | 1 | 455 | 50,8 | 6,52 | 3,16 | 1 | 1,12 | 1,128 | 0,804 | 0,982 | 0,982 | 0,868 | 0,874 | 0,996 | 1,08 | 0,08 | 3,8E-01 |
| Q8HW98 | IgLON family member 5 OS=Mus musculus OX=10090 GN=Iglon5 PE | 9 | 2 | 2 | 2 | 336 | 36,7 | 7,69 | 2,74 | 2 | 0,942 | 1,004 | 1,055 | 0,962 | 1,294 | 0,826 | 1,028 | 1,222 | 0,91 | -0,09 | 3,8E-01 |
| P52825 | Carnitine O-palmitoyltransferase 2, mitochondrial OS=Mus muscul | 5 | 3 | 3 | 3 | 658 | 73,9 | 8,37 | 3,97 | 3 | 1,663 | 1,608 | 1,287 | 1,124 | 1,139 | 1,315 | 1,239 | 1,446 | 1,11 | 0,11 | 3,8E-01 |
| P62482 | Voltage-gated potassium channel subunit beta-2 OS=Mus musculu | 14 | 6 | 7 | 4 | 367 | 41 | 9 | 15,23 | 6 | 0,768 | 0,872 | 0,869 | 0,918 | 0,947 | 0,832 | 0,94 | 0,867 | 0,96 | -0,04 | 3,8E-01 |
| Q923T9 | Calcium/calmodulin-dependent protein kinase type II subunit gam | 22 | 11 | 32 | 5 | 529 | 59,6 | 7,58 | 82,45 | 11 | 0,783 | 0,754 | 0,952 | 0,975 | 1,012 | 0,877 | 0,927 | 0,89 | 0,93 | -0,07 | 3,8E-01 |
| Q04899 | Cyclin-dependent kinase 18 OS=Mus musculus OX=10090 GN=Cdk1 | 6 | 2 | 2 | 1 | 451 | 51,8 | 8,51 | 2,39 | 2 | 1,281 | 1,268 | 0,925 | 0,71 | 0,532 | 1,136 | 0,722 | 1,05 | 1,22 | 0,22 | 3,8E-01 |
| Q61609 | Sodium-dependent phosphate transporter 1 OS=Mus musculus OX | 1 | 1 | 1 | 1 | 681 | 74,1 | 6,89 | 1,78 | 1 | 0,01 | 1,067 | 1,244 | 1,04 | 0,88 | 1,574 | 0,862 | 1,278 | 0,73 | -0,27 | 3,8E-01 |
| Q61081 | Hsp90 co-chaperone Cdc37 OS=Mus musculus OX=10090 GN=Cdc3 | 24 | 8 | 13 | 8 | 379 | 44,6 | 5,34 | 20,01 | 8 | 1,039 | 1,066 | 0,932 | 1,014 | 0,923 | 0,981 | 0,972 | 1,036 | 1,04 | 0,04 | 3,8E-01 |
| P35396 | Peroxisome proliferator-activated receptor delta OS=Mus musculu | 2 | 1 | 1 | 1 | 440 | 49,7 | 7,02 | 2,6 | 1 | 1,789 | 1,554 | 1,131 | 0,746 | 0,619 | 1,551 | 0,858 | 1,052 | 1,28 | 0,28 | 3,8E-01 |
| Q9D387 | Lysosome-associated membrane glycoprotein 5 OS=Mus musculus | 7 | 2 | 2 | 2 | 280 | 31,7 | 6,14 | 2,47 | 2 | 0,855 | 0,943 | 1,156 | 0,869 | 0,93 | 1,177 | 1,171 | 0,92 | 0,91 | -0,09 | 3,8E-01 |
| B2RUR8 | OTU domain-containing protein 7B OS=Mus musculus OX=10090 G | 3 | 2 | 2 | 1 | 840 | 91,9 | 6,86 | 0 | 2 | 1,566 | 1,571 | 0,783 | 1,376 | 1,026 | 1,291 | 0,972 | 1,249 | 1,17 | 0,17 | 3,8E-01 |
| Q9CQA6 | Coiled-coil-helix-coiled-coil-helix domain-containing protein 1 OS= | 9 | 1 | 1 | 1 | 118 | 13,6 | 10,15 | 2,75 | 1 | 1,464 | 1,494 | 1,266 | 0,935 | 0,819 | 1,47 | 0,935 | 1,207 | 1,16 | 0,16 | 3,8E-01 |
| Q8C3Q5 | Protein shisa-7 OS=Mus musculus OX=10090 GN=Shisa7 PE=1 SV=3 | 16 | 8 | 11 | 8 | 558 | 58,3 | 9,82 | 22,08 | 8 | 1,165 | 1,119 | 1,194 | 1,099 | 1,007 | 1,201 | 1,098 | 1,102 | 1,04 | 0,04 | 3,9E-01 |
| P48036 | Annexin A5 OS=Mus musculus OX=10090 GN=Anxa5 PE=1 SV=1 | 14 | 5 | 7 | 5 | 319 | 35,7 | 4,96 | 17,3 | 5 | 1,155 | 1,08 | 1,147 | 0,877 | 0,965 | 1,087 | 0,963 | 0,977 | 1,07 | 0,07 | 3,9E-01 |
| Q99N92 | 39S ribosomal protein L27, mitochondrial OS=Mus musculus OX=1 | 5 | 1 | 1 | 1 | 148 | 15,9 | 10,17 | 0 | 1 | 0,573 | 0,885 | 0,964 | 1,097 | 0,994 | 1,049 | 0,914 | 0,991 | 0,89 | -0,11 | 3,9E-01 |
| Q6ZWY8 | Thymosin beta-10 OS=Mus musculus OX=10090 GN=Tmsb10 PE=1 | 45 | 3 | 3 | 1 | 44 | 5 | 5,36 | 6,7 | 3 | 0,986 | 1,108 | 1,184 | 1,104 | 1,166 | 1,053 | 0,582 | 1,068 | 1,13 | 0,13 | 3,9E-01 |
| B2RR83 | 3'-5' RNA helicase YTHDC2 OS=Mus musculus OX=10090 GN=Ythdc | 1 | 1 | 1 | 1 | 1445 | 161 | 8,51 | 1,7 | 1 | 1,543 | 1,252 | 1,244 | 1,322 | 1,553 | 0,959 | 1,226 | 1,076 | 1,11 | 0,11 | 3,9E-01 |
| Q7TMK9-2 | Isoform 2 of Heterogeneous nuclear ribonucleoprotein Q OS=Mus | 10 | 6 | 7 | 6 | 562 | 62,6 | 7,56 | 17,05 | 6 | 1,048 | 0,958 | 1,036 | 0,945 | 1,03 | 0,955 | 0,906 | 0,959 | 1,04 | 0,04 | 3,9E-01 |
| Q8CHX7 | Raftlin-2 OS=Mus musculus OX=10090 GN=Rftn2 PE=1 SV=3 | 6 | 2 | 2 | 2 | 500 | 54,9 | 5,5 | 2,62 | 2 | 0,656 | 1,138 | 1,137 | 1,179 | 0,857 | 1,11 | 0,912 | 0,648 | 1,17 | 0,17 | 3,9E-01 |
| Q3TEA8 | Heterochromatin protein 1-binding protein 3 OS=Mus musculus OX | 9 | 5 | 5 | 5 | 554 | 60,8 | 9,7 | 7,5 | 5 | 0,98 | 1,221 | 0,956 | 0,706 | 1,083 | 1,53 | 0,811 | 1,117 | 0,85 | -0,15 | 3,9E-01 |
| O35887 | Calumenin OS=Mus musculus OX=10090 GN=Calu PE=1 SV=1 | 20 | 4 | 6 | 4 | 315 | 37 | 4,67 | 8,76 | 4 | 1,328 | 1,481 | 1,075 | 1,079 | 1,006 | 1,274 | 0,902 | 1,27 | 1,11 | 0,11 | 3,9E-01 |
| Q9WVA2 | Mitochondrial import inner membrane translocase subunit Tim8 A | 29 | 3 | 3 | 3 | 97 | 11 | 5,16 | 8,72 | 3 | 0,562 | 0,894 | 0,785 | 0,936 | 1,098 | 0,774 | 0,921 | 0,8 | 0,88 | -0,12 | 3,9E-01 |
| P56399 | Ubiquitin carboxyl-terminal hydrolase 5 OS=Mus musculus OX=100 | 22 | 15 | 22 | 15 | 858 | 95,8 | 5,01 | 47,11 | 15 | 1,044 | 0,958 | 1,072 | 1,047 | 0,986 | 0,914 | 1,072 | 0,997 | 1,04 | 0,04 | 3,9E-01 |
| Q69Z26 | Contactin-4 OS=Mus musculus OX=10090 GN=Cntn4 PE=1 SV=2 | 2 | 2 | 3 | 2 | 1026 | 113,4 | 7,36 | 7,89 | 2 | 1,06 | 0,832 | 0,977 | 0,859 | 1,114 | 0,824 | 1,137 | 0,985 | 0,92 | -0,08 | 3,9E-01 |
| Q61423 | Potassium voltage-gated channel subfamily A member 4 OS=Mus n | 3 | 2 | 2 | 1 | 654 | 73,4 | 5,29 | 4,96 | 2 | 0,784 | 1,586 | 1,538 | 1,549 | 1,54 | 0,759 | 1,452 | 0,527 | 1,28 | 0,28 | 3,9E-01 |
| Q9CQD1 | Ras-related protein Rab-5A OS=Mus musculus OX=10090 GN=Rab5 | 27 | 5 | 7 | 2 | 215 | 23,6 | 8,15 | 15,97 | 5 | 1,017 | 1,408 | 1,083 | 1,325 | 1,027 | 1,116 | 1,205 | 1,111 | 1,08 | 0,08 | 3,9E-01 |
| Q8BKC5 | Importin-5 OS=Mus musculus OX=10090 GN=Ipo5 PE=1 SV=3 | 11 | 10 | 15 | 9 | 1097 | 123,5 | 4,93 | 32,47 | 10 | 1,151 | 1,053 | 1,066 | 1,048 | 1 | 1,091 | 1,062 | 1,051 | 1,03 | 0,03 | 3,9E-01 |
| Q9CYR6 | Phosphoacetylglucosamine mutase OS=Mus musculus OX=10090 G | 5 | 3 | 3 | 3 | 542 | 59,4 | 6,2 | 5,12 | 3 | 0,834 | 0,984 | 1,193 | 1,246 | 1,047 | 1,319 | 1,202 | 1,106 | 0,91 | -0,09 | 3,9E-01 |
| Q9D871 | Carcinoembryonic antigen-related cell adhesion molecule 18 OS=M | 2 | 1 | 1 | 1 | 376 | 42,2 | 6,76 | 0 | 1 | 1,087 | 0,988 | 0,906 | 1,34 | 0,817 | 1,126 | 0,935 | 1,018 | 1,11 | 0,11 | 3,9E-01 |
| Q60973 | Histone-binding protein RBBP7 OS=Mus musculus OX=10090 GN=R | 2 | 1 | 1 | 1 | 425 | 47,8 | 5,05 | 0 | 1 | 0,664 | 1,254 | 1,589 | 1,04 | 1,291 | 0,639 | 0,867 | 0,872 | 1,24 | 0,24 | 3,9E-01 |
| P68372 | Tubulin beta-4B chain OS=Mus musculus OX=10090 GN=Tubb4b PE | 53 | 23 | 150 | 1 | 445 | 49,8 | 4,89 | 288,37 | 23 | 1,191 | 0,951 | 0,982 | 0,915 | 1,014 | 1,046 | 1,028 | 1,276 | 0,93 | -0,07 | 3,9E-01 |
| Q8BUR9 | Mitotic-spindle organizing protein 1 OS=Mus musculus OX=10090 | 21 | 1 | 1 | 1 | 78 | 8,1 | 4,78 | 2,16 | 1 | 1,116 | 1,375 | 1,563 | 0,992 | 0,945 | 1,527 | 0,99 | 0,843 | 1,17 | 0,17 | 3,9E-01 |
| O55029 | Coatomer subunit beta' OS=Mus musculus OX=10090 GN=Copb2 P | 3 | 2 | 2 | 2 | 905 | 102,4 | 5,3 | 4,34 | 2 | 1,109 | 1,138 | 1,536 | 1,598 | 1,5 | 1,752 | 1,523 | 1,227 | 0,90 | -0,10 | 3,9E-01 |
| D3YVF0 | A-kinase anchor protein 5 OS=Mus musculus OX=10090 GN=Akap5 | 39 | 16 | 21 | 16 | 745 | 79,4 | 4,75 | 45,86 | 16 | 1,137 | 0,894 | 1,025 | 1,01 | 1,031 | 1,062 | 1,211 | 1,011 | 0,94 | -0,06 | 3,9E-01 |
| Q78IK2 | ATP synthase membrane subunit DAPIT, mitochondrial OS=Mus mu | 17 | 1 | 1 | 1 | 58 | 6,4 | 9,83 | 1,83 | 1 | 1,286 | 0,867 | 1,047 | 0,967 | 1,028 | 1,306 | 0,998 | 1,283 | 0,90 | -0,10 | 3,9E-01 |
| Q8CG76 | Aflatoxin B1 aldehyde reductase member 2 OS=Mus musculus OX= | 9 | 2 | 2 | 2 | 367 | 40,6 | 8,12 | 7,15 | 2 | 0,786 | 0,923 | 0,716 | 0,728 | 0,815 | 0,844 | 0,739 | 1,076 | 0,91 | -0,09 | 3,9E-01 |
| P62900 | 60S ribosomal protein L31 OS=Mus musculus OX=10090 GN=Rpl31 | 26 | 4 | 4 | 4 | 125 | 14,5 | 10,54 | 5,37 | 4 | 0,817 | 0,809 | 0,861 | 0,968 | 1,136 | 0,869 | 0,863 | 0,87 | 0,92 | -0,08 | 3,9E-01 |
| O35633 | Vesicular inhibitory amino acid transporter OS=Mus musculus OX= | 23 | 9 | 11 | 9 | 525 | 57,3 | 6,64 | 30,59 | 9 | 1,186 | 1,125 | 1,032 | 0,981 | 0,905 | 1,094 | 0,993 | 1,093 | 1,06 | 0,06 | 3,9E-01 |
| P47962 | 60S ribosomal protein L5 OS=Mus musculus OX=10090 GN=Rpl5 PE | 14 | 4 | 5 | 4 | 297 | 34,4 | 9,77 | 9,77 | 4 | 1,065 | 1,035 | 0,929 | 0,999 | 1,117 | 0,961 | 1,028 | 1,09 | 0,96 | -0,04 | 3,9E-01 |
| Q9Z1S5 | Neuronal-specific septin-3 OS=Mus musculus OX=10090 GN=Septi | 42 | 16 | 32 | 16 | 350 | 40 | 6,81 | 71,01 | 16 | 0,81 | 0,929 | 0,943 | 1,051 | 0,961 | 0,908 | 1,016 | 1,072 | 0,94 | -0,06 | 3,9E-01 |
| Q3UHD2 | Glucose-fructose oxidoreductase domain-containing protein 1 OS= | 5 | 2 | 2 | 2 | 390 | 43,3 | 5,92 | 4,61 | 2 | 0,753 | 1,523 | 1,029 | 1,264 | 1,505 | 1,241 | 1,435 | 1,088 | 0,87 | -0,13 | 3,9E-01 |
| Q8C1Z7 | Bardet-Biedl syndrome 4 protein homolog OS=Mus musculus OX=1 | 2 | 1 | 1 | 1 | 520 | 58,2 | 8,1 | 2,32 | 1 | 1,243 | 1,059 | 1,881 | 1,617 | 1,367 | 1,354 | 1,185 | 1,189 | 1,14 | 0,14 | 3,9E-01 |
| Q8K4Z0 | Leucine-rich repeat LGI family member 2 OS=Mus musculus OX=10 | 7 | 4 | 4 | 4 | 550 | 63 | 6,54 | 10,44 | 4 | 0,866 | 1,069 | 0,846 | 1,027 | 0,903 | 0,93 | 0,921 | 0,831 | 1,06 | 0,06 | 3,9E-01 |
| Q9JMA1 | Ubiquitin carboxyl-terminal hydrolase 14 OS=Mus musculus OX=10 | 20 | 9 | 13 | 9 | 493 | 56 | 5,24 | 24,39 | 9 | 1,144 | 1,052 | 1,055 | 1,054 | 0,908 | 1,108 | 1,04 | 1,069 | 1,04 | 0,04 | 3,9E-01 |
| Q9Z1M0 | P2X purinoceptor 7 OS=Mus musculus OX=10090 GN=P2rx7 PE=1 S | 2 | 1 | 1 | 1 | 595 | 68,3 | 8,22 | 2,91 | 1 | 0,924 | 1,027 | 1,117 | 0,709 | 0,792 | 0,916 | 0,764 | 0,943 | 1,11 | 0,11 | 3,9E-01 |
| Q3UUI3 | Acyl-coenzyme A thioesterase THEM4 OS=Mus musculus OX=1009 | 17 | 3 | 3 | 3 | 230 | 26 | 9,64 | 2,53 | 3 | 0,979 | 0,946 | 0,718 | 1,108 | 1,113 | 1,092 | 1,268 | 0,767 | 0,88 | -0,12 | 3,9E-01 |
| P02088 | Hemoglobin subunit beta-1 OS=Mus musculus OX=10090 GN=Hbb- | 34 | 4 | 4 | 4 | 147 | 15,8 | 7,65 | 12,7 | 4 | 1,082 | 1,062 | 0,714 | 0,597 | 1,056 | 0,904 | 0,604 | 2,553 | 0,68 | -0,32 | 3,9E-01 |
| Q8C5W0 | Calmin OS=Mus musculus OX=10090 GN=Clmn PE=1 SV=2 | 1 | 1 | 1 | 1 | 1052 | 117,2 | 5,03 | 0 | 1 | 1,495 | 1,332 | 1,133 | 1,136 | 1,176 | 0,965 | 1,305 | 1,232 | 1,09 | 0,09 | 3,9E-01 |
| P10833 | Ras-related protein R-Ras OS=Mus musculus OX=10090 GN=Rras PE | 19 | 3 | 7 | 1 | 218 | 23,7 | 6,79 | 9,64 | 3 | 1,223 | 0,716 | 0,777 | 0,916 | 0,801 | 0,831 | 0,651 | 0,893 | 1,14 | 0,14 | 3,9E-01 |
| Q8CJH3 | Plexin-B1 OS=Mus musculus OX=10090 GN=Plxnb1 PE=1 SV=2 | 2 | 5 | 5 | 5 | 2119 | 231,2 | 5,55 | 12,85 | 5 | 1,002 | 0,808 | 1,118 | 1,126 | 1,091 | 1,048 | 1,13 | 1,065 | 0,94 | -0,06 | 3,9E-01 |

| | | | | | | | | | | | | | | | | | | | | | |
|---|---|---|---|---|---|---|---|---|---|---|---|---|---|---|---|---|---|---|---|---|---|
| Q8VCX5 | Calcium uptake protein 1, mitochondrial OS=Mus musculus OX=10( | 12 | 5 | 5 | 5 | 477 | 54,3 | 8,59 | 9,27 | 5 | 1,054 | 0,819 | 1,151 | 1,197 | 0,739 | 1,111 | 0,938 | 1,01 | 1,11 | 0,11 | 3,9E-01 |
| Q9EQH3 | Vacuolar protein sorting-associated protein 35 OS=Mus musculus C | 18 | 13 | 19 | 13 | 796 | 91,7 | 5,44 | 40,62 | 13 | 0,86 | 0,807 | 1,009 | 1,038 | 1,017 | 0,942 | 0,99 | 0,979 | 0,95 | -0,05 | 3,9E-01 |
| Q9DBE8 | Alpha-1,3/1,6-mannosyltransferase ALG2 OS=Mus musculus OX=10 | 7 | 3 | 3 | 3 | 415 | 47,4 | 7,97 | 6,24 | 3 | 1,143 | 1,109 | 0,973 | 1,018 | 0,882 | 1,141 | 0,901 | 1,05 | 1,07 | 0,07 | 3,9E-01 |
| P28658 | Ataxin-10 OS=Mus musculus OX=10090 GN=Atxn10 PE=1 SV=2 | 5 | 2 | 4 | 2 | 475 | 53,7 | 5,25 | 4,25 | 2 | 1,151 | 0,89 | 0,519 | 0,7 | 1,142 | 0,787 | 1,15 | 0,801 | 0,84 | -0,16 | 3,9E-01 |
| Q8R1Y2 | bMERB domain-containing protein 1 OS=Mus musculus OX=10090 | 6 | 2 | 2 | 2 | 203 | 23,5 | 5,63 | 5,05 | 2 | 1,721 | 1,058 | 0,937 | 1,401 | 0,963 | 1,175 | 1,237 | 1,056 | 1,15 | 0,15 | 4,0E-01 |
| Q07113 | Cation-independent mannose-6-phosphate receptor OS=Mus musc | 1 | 2 | 2 | 2 | 2483 | 273,6 | 5,71 | 4,2 | 2 | 0,95 | 1,106 | 0,956 | 0,86 | 0,91 | 1,241 | 0,948 | 1,109 | 0,92 | -0,08 | 4,0E-01 |
| Q8BGQ1 | Spermatogenesis-defective protein 39 homolog OS=Mus musculus | 10 | 4 | 4 | 4 | 491 | 56,6 | 7,17 | 2,29 | 4 | 1,076 | 0,953 | 0,83 | 0,989 | 0,823 | 0,941 | 0,95 | 0,919 | 1,06 | 0,06 | 4,0E-01 |
| Q8C0J2 | Autophagy-related protein 16-1 OS=Mus musculus OX=10090 GN=/ | 1 | 1 | 1 | 1 | 607 | 68,1 | 6,38 | 2,32 | 1 | 1,138 | 0,924 | 1,093 | 0,942 | 1,143 | 1,012 | 0,993 | 1,236 | 0,93 | -0,07 | 4,0E-01 |
| P70429 | Ena/VASP-like protein OS=Mus musculus OX=10090 GN=Evl PE=1 S | 9 | 3 | 5 | 3 | 414 | 44,3 | 8,85 | 8,19 | 3 | 1,304 | 1,033 | 0,906 | 1,11 | 0,945 | 1,025 | 1,088 | 0,969 | 1,08 | 0,08 | 4,0E-01 |
| P35288 | Ras-related protein Rab-23 OS=Mus musculus OX=10090 GN=Rab2 | 14 | 2 | 2 | 2 | 237 | 26,7 | 6,79 | 3,16 | 2 | 1,305 | 1,36 | 1,35 | 1,162 | 0,937 | 1,557 | 1,069 | 1,092 | 1,11 | 0,11 | 4,0E-01 |
| Q07409 | Contactin-3 OS=Mus musculus OX=10090 GN=Cntn3 PE=1 SV=2 | 3 | 3 | 3 | 3 | 1028 | 113,2 | 6,23 | 4,97 | 3 | 1,155 | 0,995 | 1,08 | 0,852 | 1,021 | 0,809 | 0,976 | 0,983 | 1,08 | 0,08 | 4,0E-01 |
| Q8BHE8 | m-AAA protease-interacting protein 1, mitochondrial OS=Mus mus | 8 | 2 | 2 | 2 | 291 | 33 | 9,14 | 3,09 | 2 | 1,045 | 1,138 | 1,238 | 1,128 | 1,033 | 1,106 | 1,11 | 1,135 | 1,04 | 0,04 | 4,0E-01 |
| Q80XK6 | Autophagy-related protein 2 homolog B OS=Mus musculus OX=10( | 1 | 1 | 1 | 1 | 2075 | 231,3 | 5,88 | 2,06 | 1 | 1,144 | 0,982 | 0,932 | 1,089 | 1,002 | 0,761 | 1,068 | 1,011 | 1,08 | 0,08 | 4,0E-01 |
| P62814 | V-type proton ATPase subunit B, brain isoform OS=Mus musculus C | 37 | 21 | 68 | 21 | 511 | 56,5 | 5,81 | 159 | 21 | 1,038 | 0,975 | 1,037 | 1,072 | 1,025 | 0,929 | 1,01 | 1,041 | 1,03 | 0,03 | 4,0E-01 |
| Q9ES34 | Ubiquitin-protein ligase E3B OS=Mus musculus OX=10090 GN=Ube | 1 | 1 | 1 | 1 | 1070 | 122,7 | 8,32 | 0 | 1 | 1,016 | 0,7 | 0,754 | 0,961 | 0,85 | 0,838 | 0,712 | 0,719 | 1,10 | 0,10 | 4,0E-01 |
| Q9R0X4 | Acyl-coenzyme A thioesterase 9, mitochondrial OS=Mus musculus ( | 19 | 9 | 10 | 9 | 439 | 50,5 | 8,59 | 25,62 | 9 | 1,158 | 1,096 | 1,023 | 0,987 | 1,066 | 1,192 | 1,075 | 1,105 | 0,96 | -0,04 | 4,0E-01 |
| Q9D832 | DnaJ homolog subfamily B member 4 OS=Mus musculus OX=10090 | 30 | 8 | 10 | 7 | 337 | 37,8 | 8,59 | 19,06 | 8 | 0,943 | 0,829 | 0,936 | 0,973 | 0,889 | 0,918 | 1,095 | 0,981 | 0,95 | -0,05 | 4,0E-01 |
| Q8CC35 | Synaptopodin OS=Mus musculus OX=10090 GN=Synpo PE=1 SV=2 | 5 | 4 | 5 | 4 | 929 | 99,5 | 9,42 | 9,8 | 4 | 1,113 | 0,901 | 0,915 | 0,954 | 1,033 | 0,977 | 1,085 | 0,987 | 0,95 | -0,05 | 4,0E-01 |
| Q9Z0K8 | Pantetheinase OS=Mus musculus OX=10090 GN=Vnn1 PE=1 SV=3 | 3 | 1 | 1 | 1 | 512 | 57,1 | 5,94 | 2,07 | 1 | 1,56 | 1,973 | 1,653 | 0,662 | 0,631 | 1,682 | 1,056 | 1,188 | 1,28 | 0,28 | 4,0E-01 |
| Q9EPL4 | Methyltransferase-like protein 9 OS=Mus musculus OX=10090 GN= | 4 | 1 | 1 | 1 | 318 | 36,4 | 6,68 | 2,47 | 1 | 1,109 | 1,267 | 1,6 | 0,796 | 1,287 | 1,472 | 1,07 | 1,74 | 0,86 | -0,14 | 4,0E-01 |
| Q9JME5 | AP-3 complex subunit beta-2 OS=Mus musculus OX=10090 GN=Ap3 | 15 | 17 | 28 | 15 | 1082 | 119,1 | 5,63 | 49,75 | 17 | 1,031 | 1,016 | 1,05 | 1,045 | 0,972 | 1,006 | 1,022 | 1,066 | 1,02 | 0,02 | 4,0E-01 |
| Q80W47 | WD repeat domain phosphoinositide-interacting protein 2 OS=Mus | 4 | 2 | 2 | 2 | 445 | 48,4 | 5,86 | 4,21 | 2 | 1,008 | 1,228 | 1,1 | 0,912 | 0,761 | 1,22 | 0,977 | 0,855 | 1,11 | 0,11 | 4,0E-01 |
| P70265 | 6-phosphofructo-2-kinase/fructose-2,6-bisphosphatase 2 OS=Mus | 4 | 2 | 2 | 2 | 519 | 59,9 | 8,12 | 5,55 | 2 | 1,061 | 1,164 | 1,045 | 1,092 | 1,006 | 1,173 | 1,057 | 0,898 | 1,06 | 0,06 | 4,0E-01 |
| O35449 | Proline-rich transmembrane protein 1 OS=Mus musculus OX=1009( | 4 | 1 | 1 | 1 | 306 | 31,4 | 7,65 | 3,55 | 1 | 0,847 | 1,327 | 0,877 | 0,876 | 0,833 | 0,959 | 0,809 | 0,891 | 1,12 | 0,12 | 4,0E-01 |
| Q61730 | Interleukin-1 receptor accessory protein OS=Mus musculus OX=10( | 2 | 2 | 2 | 2 | 570 | 65,7 | 7,77 | 0 | 2 | 1,562 | 1,23 | 1,397 | 1,012 | 1,281 | 1,129 | 1,142 | 1,205 | 1,09 | 0,09 | 4,0E-01 |
| A2AF47 | Dedicator of cytokinesis protein 11 OS=Mus musculus OX=10090 G | 1 | 1 | 1 | 1 | 2073 | 237,6 | 7,78 | 1,83 | 1 | 1,264 | 1,09 | 1,066 | 1,011 | 1,014 | 1,352 | 0,961 | 1,952 | 0,84 | -0,16 | 4,0E-01 |
| Q91VR5 | ATP-dependent RNA helicase DDX1 OS=Mus musculus OX=10090 G | 5 | 3 | 3 | 3 | 740 | 82,4 | 7,21 | 4,67 | 3 | 1,067 | 1,406 | 1,11 | 1,166 | 1,123 | 1,034 | 1,171 | 1,128 | 1,07 | 0,07 | 4,0E-01 |
| Q91W67 | Ubiquitin-like protein 7 OS=Mus musculus OX=10090 GN=Ubl7 PE= | 2 | 1 | 2 | 1 | 380 | 40,4 | 5,01 | 0 | 1 | 1,075 | 1,217 | 1,049 | 1,126 | 1,013 | 1,109 | 1,113 | 1,074 | 1,04 | 0,04 | 4,0E-01 |
| O88998 | Noelin OS=Mus musculus OX=10090 GN=Olfm1 PE=1 SV=1 | 4 | 2 | 2 | 2 | 485 | 55,4 | 6,95 | 4,25 | 2 | 1,043 | 0,833 | 0,805 | 0,78 | 1,056 | 0,908 | 0,788 | 1,017 | 0,92 | -0,08 | 4,0E-01 |
| Q6WQJ1 | Sn1-specific diacylglycerol lipase alpha OS=Mus musculus OX=1009 | 2 | 2 | 2 | 2 | 1044 | 115,3 | 6,42 | 4,38 | 2 | 1,13 | 1,26 | 1,198 | 1,227 | 1,24 | 1,544 | 1,037 | 1,4 | 0,92 | -0,08 | 4,0E-01 |
| Q8VED9 | Galectin-related protein OS=Mus musculus OX=10090 GN=Lgalsl PE | 26 | 4 | 6 | 4 | 172 | 18,9 | 5,35 | 12,69 | 4 | 0,689 | 0,813 | 1,089 | 1,042 | 0,918 | 1,073 | 1,02 | 0,984 | 0,91 | -0,09 | 4,0E-01 |
| Q3UUG6 | TBC1 domain family member 24 OS=Mus musculus OX=10090 GN= | 16 | 9 | 13 | 9 | 561 | 63,2 | 7,24 | 24,43 | 9 | 1,144 | 1,124 | 1,028 | 1,18 | 0,936 | 1,152 | 1,082 | 1,102 | 1,05 | 0,05 | 4,0E-01 |
| Q9JKY5 | Huntingtin-interacting protein 1-related protein OS=Mus musculus | 11 | 13 | 17 | 13 | 1068 | 119,4 | 6,52 | 33,92 | 13 | 1,098 | 1,227 | 1,028 | 0,984 | 1,008 | 1,071 | 1,08 | 0,963 | 1,05 | 0,05 | 4,0E-01 |
| Q9WUB3 | Glycogen phosphorylase, muscle form OS=Mus musculus OX=1009( | 21 | 16 | 19 | 11 | 842 | 97,2 | 7,11 | 46,18 | 16 | 1,109 | 1,063 | 0,974 | 0,975 | 0,917 | 1,092 | 0,916 | 1,003 | 1,05 | 0,05 | 4,0E-01 |
| Q9CX00 | IST1 homolog OS=Mus musculus OX=10090 GN=Ist1 PE=1 SV=1 | 5 | 2 | 2 | 2 | 362 | 39,4 | 5,44 | 2,34 | 2 | 1,002 | 1,217 | 1,232 | 1,027 | 0,827 | 1,166 | 1,134 | 0,998 | 1,09 | 0,09 | 4,0E-01 |
| P06801 | NADP-dependent malic enzyme OS=Mus musculus OX=10090 GN=I | 13 | 7 | 11 | 7 | 572 | 63,9 | 7,44 | 26,05 | 7 | 0,932 | 0,945 | 1,054 | 0,955 | 0,93 | 1,039 | 1,028 | 1,024 | 0,97 | -0,03 | 4,0E-01 |
| O88738 | Baculoviral IAP repeat-containing protein 6 OS=Mus musculus OX= | 1 | 2 | 2 | 2 | 4882 | 531,8 | 6,07 | 2,71 | 2 | 1,545 | 1,185 | 1,133 | 1,02 | 0,91 | 1,431 | 0,998 | 0,95 | 1,14 | 0,14 | 4,0E-01 |
| Q9CQL4 | 39S ribosomal protein L20, mitochondrial OS=Mus musculus OX=10 | 7 | 1 | 1 | 1 | 149 | 17,6 | 11,11 | 2,38 | 1 | 0,926 | 0,742 | 0,732 | 0,765 | 0,977 | 0,775 | 0,864 | 0,783 | 0,93 | -0,07 | 4,1E-01 |
| Q9D164 | FXYD domain-containing ion transport regulator 6 OS=Mus muscul | 14 | 1 | 1 | 1 | 94 | 10,4 | 5,11 | 2,45 | 1 | 1,307 | 1,507 | 1,158 | 1,165 | 0,995 | 1,357 | 0,956 | 1,343 | 1,10 | 0,10 | 4,1E-01 |
| G5E8K5 | Ankyrin-3 OS=Mus musculus OX=10090 GN=Ank3 PE=1 SV=1 | 16 | 27 | 36 | 22 | 1961 | 213,9 | 6,7 | 72,58 | 27 | 1,053 | 0,943 | 0,967 | 0,95 | 0,969 | 1,041 | 0,968 | 1,06 | 0,97 | -0,03 | 4,1E-01 |
| Q8CAK1 | Putative transferase CAF17 homolog, mitochondrial OS=Mus musc | 13 | 4 | 4 | 4 | 358 | 38,4 | 9,01 | 9,17 | 4 | 1,391 | 1,007 | 0,955 | 0,921 | 0,98 | 1,004 | 0,985 | 0,909 | 1,10 | 0,10 | 4,1E-01 |
| Q9WUP7 | Ubiquitin carboxyl-terminal hydrolase isozyme L5 OS=Mus musculu | 8 | 2 | 3 | 2 | 329 | 37,6 | 5,33 | 4,84 | 2 | 0,809 | 0,9 | 0,584 | 0,773 | 0,848 | 0,899 | 0,828 | 0,752 | 0,92 | -0,08 | 4,1E-01 |
| Q6P5F6 | Zinc transporter ZIP10 OS=Mus musculus OX=10090 GN=Slc39a10 I | 1 | 1 | 1 | 1 | 833 | 94,3 | 6,71 | 2,07 | 1 | 1,172 | 0,81 | 0,801 | 0,78 | 0,901 | 1,526 | 0,774 | 1,038 | 0,84 | -0,16 | 4,1E-01 |
| P59648 | FXYD domain-containing ion transport regulator 7 OS=Mus musculu | 20 | 2 | 7 | 2 | 80 | 8,5 | 7,84 | 10,17 | 2 | 1,388 | 1,012 | 1,003 | 1,014 | 0,948 | 1,227 | 0,939 | 0,861 | 1,11 | 0,11 | 4,1E-01 |
| Q99J08 | SEC14-like protein 2 OS=Mus musculus OX=10090 GN=Sec14l2 PE= | 11 | 5 | 7 | 5 | 403 | 46,3 | 7,11 | 11,15 | 5 | 1,044 | 0,961 | 1,037 | 0,972 | 0,984 | 0,944 | 0,984 | 1,011 | 1,02 | 0,02 | 4,1E-01 |
| Q9CZC8 | Secernin-1 OS=Mus musculus OX=10090 GN=Scrn1 PE=1 SV=1 | 26 | 11 | 12 | 11 | 414 | 46,3 | 4,79 | 23,45 | 11 | 1,158 | 1,035 | 1,204 | 2,099 | 0,959 | 1,212 | 1,141 | 1,282 | 1,20 | 0,20 | 4,1E-01 |
| Q0GA42 | Metal transporter CNNM1 OS=Mus musculus OX=10090 GN=Cnnm | 3 | 3 | 3 | 2 | 951 | 103,9 | 6,51 | 2,36 | 3 | 1,162 | 0,868 | 1,01 | 0,884 | 0,806 | 1,071 | 0,842 | 0,885 | 1,09 | 0,09 | 4,1E-01 |
| Q80T41 | Gamma-aminobutyric acid type B receptor subunit 2 OS=Mus musc | 6 | 6 | 7 | 6 | 940 | 105,6 | 8,72 | 13,45 | 6 | 1,068 | 0,967 | 0,92 | 0,988 | 1,026 | 1,084 | 0,97 | 1,002 | 0,97 | -0,03 | 4,1E-01 |
| Q8C6B2 | Rhotekin OS=Mus musculus OX=10090 GN=Rtkn PE=1 SV=3 | 1 | 1 | 1 | 1 | 564 | 63 | 6,67 | 0 | 1 | 1,133 | 0,992 | 1,103 | 0,979 | 1,099 | 1,208 | 0,663 | 0,747 | 1,13 | 0,13 | 4,1E-01 |
| P08113 | Endoplasmin OS=Mus musculus OX=10090 GN=Hsp90b1 PE=1 SV=2 | 25 | 21 | 29 | 19 | 802 | 92,4 | 4,82 | 58,95 | 21 | 0,985 | 0,97 | 1,018 | 0,984 | 1,025 | 0,913 | 0,962 | 0,968 | 1,02 | 0,02 | 4,1E-01 |
| P51830 | Adenylate cyclase type 9 OS=Mus musculus OX=10090 GN=Adcy9 F | 11 | 14 | 15 | 14 | 1353 | 150,9 | 7,21 | 35,29 | 14 | 1,046 | 1,141 | 0,968 | 1,042 | 1,08 | 1,046 | 1,06 | 1,172 | 0,96 | -0,04 | 4,1E-01 |
| P15209 | BDNF/NT-3 growth factors receptor OS=Mus musculus OX=10090 ( | 14 | 9 | 11 | 6 | 821 | 92,1 | 6,55 | 25,73 | 9 | 1,371 | 0,999 | 1,122 | 0,937 | 0,912 | 1,126 | 0,997 | 1,02 | 1,09 | 0,09 | 4,1E-01 |
| Q8CAL5 | Glypican-5 OS=Mus musculus OX=10090 GN=Gpc5 PE=2 SV=1 | 2 | 1 | 1 | 1 | 572 | 63,8 | 7,3 | 0 | 1 | 0,866 | 1,032 | 0,644 | 0,818 | 0,988 | 1,007 | 0,91 | 0,788 | 0,91 | -0,09 | 4,1E-01 |
| Q06138 | Calcium-binding protein 39 OS=Mus musculus OX=10090 GN=Cab3 | 24 | 9 | 11 | 9 | 341 | 39,8 | 6,89 | 16,98 | 9 | 0,976 | 0,928 | 0,923 | 1,016 | 0,964 | 0,924 | 1,02 | 1,075 | 0,96 | -0,04 | 4,1E-01 |
| Q9WV18 | Gamma-aminobutyric acid type B receptor subunit 1 OS=Mus musc | 8 | 6 | 6 | 6 | 960 | 108,1 | 8,21 | 12 | 6 | 0,924 | 1,022 | 1,113 | 0,972 | 0,905 | 0,893 | 0,984 | 1,054 | 1,05 | 0,05 | 4,1E-01 |
| Q8BMG7 | Rab3 GTPase-activating protein non-catalytic subunit OS=Mus mus | 10 | 12 | 13 | 12 | 1366 | 152,4 | 6,23 | 28,22 | 12 | 0,939 | 0,985 | 1,009 | 1,047 | 1,008 | 1,014 | 0,99 | 1,067 | 0,98 | -0,02 | 4,1E-01 |
| Q8K3F6 | Potassium voltage-gated channel subfamily KQT member 3 OS=Mu | 1 | 1 | 1 | 1 | 873 | 96,8 | 8,79 | 2,42 | 1 | 0,823 | 1,06 | 0,944 | 0,932 | 0,915 | 0,969 | 1,229 | 0,952 | 0,92 | -0,08 | 4,1E-01 |
| Q9Z2D0 | Myotubularin-related protein 9 OS=Mus musculus OX=10090 GN=N | 12 | 6 | 7 | 6 | 545 | 62,9 | 6,62 | 14,16 | 6 | 0,729 | 0,868 | 0,788 | 0,921 | 0,774 | 0,898 | 0,856 | 1,004 | 0,94 | -0,06 | 4,1E-01 |
| Q8BKF1 | DNA-directed RNA polymerase, mitochondrial OS=Mus musculus O | 1 | 1 | 1 | 1 | 1207 | 136,6 | 9,09 | 0 | 1 | 1,773 | 1,317 | 1,137 | 0,919 | 1,191 | 0,742 | 1,429 | 0,957 | 1,19 | 0,19 | 4,1E-01 |
| P00416 | Cytochrome c oxidase subunit 3 OS=Mus musculus OX=10090 GN=r | 5 | 1 | 2 | 1 | 261 | 29,9 | 7,3 | 3,01 | 1 | 0,903 | 1,417 | 0,968 | 0,672 | 0,8 | 0,974 | 0,89 | 0,711 | 1,17 | 0,17 | 4,1E-01 |
| Q9QZS8 | SH2 domain-containing protein 3C OS=Mus musculus OX=10090 GI | 1 | 1 | 1 | 1 | 854 | 94,3 | 7,53 | 0 | 1 | 0,01 | 0,779 | 0,807 | 0,964 | 0,731 | 0,891 | 0,898 | 0,807 | 0,77 | -0,23 | 4,1E-01 |
| Q9CQE8 | RNA transcription, translation and transport factor protein OS=Mus | 4 | 1 | 1 | 1 | 244 | 28,1 | 6,89 | 2,85 | 1 | 1,199 | 1,074 | 1,05 | 1,201 | 1,178 | 0,902 | 1,176 | 0,985 | 1,07 | 0,07 | 4,1E-01 |

| | | | | | | | | | | | | | | | | | | | | | |
|---|---|---|---|---|---|---|---|---|---|---|---|---|---|---|---|---|---|---|---|---|---|
| Q8BH59 | Calcium-binding mitochondrial carrier protein Aralar1 OS=Mus mus | 23 | 15 | 31 | 13 | 677 | 74,5 | 8,25 | 86,94 | 15 | 1,307 | 1,273 | 1,122 | 0,99 | 1 | 1,213 | 1,038 | 1,133 | 1,07 | 0,07 | 4,1E-01 |
| Q99L85 | Elongator complex protein 5 OS=Mus musculus OX=10090 GN=Elp5 | 3 | 1 | 1 | 1 | 300 | 33,5 | 5,49 | 1,86 | 1 | 0,968 | 1,06 | 0,697 | 1,168 | 0,81 | 1,143 | 1,319 | 1,136 | 0,88 | -0,12 | 4,1E-01 |
| P12787 | Cytochrome c oxidase subunit 5A, mitochondrial OS=Mus musculus | 34 | 6 | 18 | 6 | 146 | 16,1 | 6,54 | 37,3 | 6 | 1,06 | 1,112 | 0,983 | 0,952 | 0,983 | 0,975 | 0,978 | 1,034 | 1,03 | 0,03 | 4,1E-01 |
| P61027 | Ras-related protein Rab-10 OS=Mus musculus OX=10090 GN=Rab1( | 30 | 7 | 9 | 5 | 200 | 22,5 | 8,38 | 22,61 | 7 | 0,945 | 1,102 | 1,053 | 0,879 | 0,932 | 0,948 | 0,994 | 0,918 | 1,05 | 0,05 | 4,1E-01 |
| P21995 | Embigin OS=Mus musculus OX=10090 GN=Emb PE=1 SV=2 | 6 | 2 | 2 | 2 | 330 | 37 | 6,02 | 2,48 | 2 | 1,293 | 1,075 | 1,09 | 1,073 | 1,378 | 1,193 | 1,154 | 1,092 | 0,94 | -0,06 | 4,1E-01 |
| Q9R0D8 | WD repeat-containing protein 54 OS=Mus musculus OX=10090 GN= | 7 | 2 | 2 | 2 | 334 | 35,6 | 6,21 | 5,46 | 2 | 1,012 | 1,198 | 1,071 | 1,272 | 1,202 | 0,957 | 1,468 | 1,369 | 0,91 | -0,09 | 4,1E-01 |
| P49446 | Receptor-type tyrosine-protein phosphatase epsilon OS=Mus musc | 4 | 3 | 3 | 2 | 699 | 80,6 | 7,12 | 4,24 | 3 | 0,819 | 0,792 | 0,917 | 0,807 | 0,848 | 0,915 | 0,957 | 0,782 | 0,95 | -0,05 | 4,2E-01 |
| Q64010 | Adapter molecule crk OS=Mus musculus OX=10090 GN=Crk PE=1 S\ | 14 | 4 | 4 | 4 | 304 | 33,8 | 5,55 | 7,68 | 4 | 0,894 | 0,887 | 0,787 | 0,865 | 1,065 | 0,834 | 0,829 | 0,916 | 0,94 | -0,06 | 4,2E-01 |
| Q9QYK7 | RING finger protein 11 OS=Mus musculus OX=10090 GN=Rnf11 PE= | 14 | 2 | 2 | 2 | 154 | 17,4 | 4,78 | 3,2 | 2 | 1,089 | 1,156 | 0,948 | 1,139 | 0,885 | 0,959 | 0,971 | 1,216 | 1,07 | 0,07 | 4,2E-01 |
| Q3U214 | Microtubule-associated serine/threonine-protein kinase 3 OS=Mus | 4 | 4 | 4 | 4 | 1321 | 144,1 | 8,44 | 7,82 | 4 | 1,328 | 1,286 | 1,169 | 0,905 | 0,972 | 1,167 | 1,093 | 1,095 | 1,08 | 0,08 | 4,2E-01 |
| Q60902 | Epidermal growth factor receptor substrate 15-like 1 OS=Mus musc | 19 | 13 | 14 | 13 | 907 | 99,2 | 5,02 | 27,37 | 13 | 1,146 | 1,035 | 1,146 | 0,894 | 0,862 | 1,106 | 0,893 | 1,063 | 1,08 | 0,08 | 4,2E-01 |
| Q8CIP4 | MAP/microtubule affinity-regulating kinase 4 OS=Mus musculus OX | 1 | 1 | 1 | 1 | 752 | 82,6 | 9,67 | 0 | 1 | 1,709 | 1,146 | 0,919 | 1,166 | 1,064 | 0,973 | 1,076 | 1,218 | 1,14 | 0,14 | 4,2E-01 |
| Q8BWY3 | Eukaryotic peptide chain release factor subunit 1 OS=Mus musculu | 5 | 2 | 2 | 2 | 437 | 49 | 5,71 | 5,87 | 2 | 1,097 | 0,67 | 0,88 | 1,103 | 1,064 | 0,99 | 0,985 | 1,079 | 0,91 | -0,09 | 4,2E-01 |
| Q3UPL0 | Protein transport protein Sec31A OS=Mus musculus OX=10090 GN= | 5 | 6 | 7 | 6 | 1230 | 133,5 | 6,76 | 18,27 | 6 | 1,034 | 1,096 | 1,038 | 0,98 | 0,952 | 1,041 | 1,066 | 0,96 | 1,03 | 0,03 | 4,2E-01 |
| P28663 | Beta-soluble NSF attachment protein OS=Mus musculus OX=10090 | 57 | 17 | 54 | 16 | 298 | 33,5 | 5,47 | 103,46 | 17 | 0,997 | 0,993 | 0,96 | 0,964 | 0,955 | 0,952 | 0,998 | 0,959 | 1,01 | 0,01 | 4,2E-01 |
| Q8C145 | Zinc transporter ZIP6 OS=Mus musculus OX=10090 GN=Slc39a6 PE= | 3 | 2 | 2 | 2 | 765 | 86,3 | 6,84 | 2,52 | 2 | 0,729 | 0,859 | 0,985 | 1,027 | 1,024 | 0,644 | 1,249 | 1,219 | 0,87 | -0,13 | 4,2E-01 |
| Q60770 | Syntaxin-binding protein 3 OS=Mus musculus OX=10090 GN=Stxbp | 7 | 4 | 5 | 4 | 592 | 67,9 | 8,02 | 6 | 4 | 1,182 | 1,144 | 1,24 | 1,059 | 1,033 | 1,236 | 1,115 | 1,027 | 1,05 | 0,05 | 4,2E-01 |
| Q3V1M1 | Immunoglobulin superfamily member 10 OS=Mus musculus OX=10 | 0 | 1 | 1 | 1 | 2594 | 285,4 | 9,29 | 0 | 1 | 1,055 | 1,155 | 1,48 | 0,982 | 0,691 | 0,893 | 1,172 | 1,305 | 1,15 | 0,15 | 4,2E-01 |
| Q3USB7 | Inactive phospholipase C-like protein 1 OS=Mus musculus OX=1009 | 11 | 9 | 10 | 8 | 1096 | 122,6 | 5,64 | 26,17 | 9 | 1,197 | 1,194 | 1,128 | 1,129 | 0,975 | 1,245 | 1,087 | 1,136 | 1,05 | 0,05 | 4,2E-01 |
| Q61418 | H(+)/Cl(-) exchange transporter 4 OS=Mus musculus OX=10090 GN= | 1 | 1 | 1 | 1 | 747 | 83,7 | 6,86 | 2,65 | 1 | 1,944 | 1,217 | 0,943 | 1,073 | 0,947 | 1,224 | 1,003 | 1,194 | 1,19 | 0,19 | 4,2E-01 |
| Q8VDQ1 | Prostaglandin reductase 2 OS=Mus musculus OX=10090 GN=Ptgr2 | 3 | 1 | 1 | 1 | 351 | 38 | 5,41 | 0 | 1 | 1,384 | 0,99 | 0,756 | 0,878 | 0,991 | 0,958 | 0,461 | 0,955 | 1,19 | 0,19 | 4,2E-01 |
| Q6P8X1 | Sorting nexin-6 OS=Mus musculus OX=10090 GN=Snx6 PE=1 SV=2 | 13 | 6 | 8 | 6 | 406 | 46,6 | 6,16 | 12,03 | 6 | 0,885 | 0,878 | 0,841 | 0,949 | 0,87 | 0,872 | 0,932 | 1,029 | 0,96 | -0,04 | 4,2E-01 |
| Q6IR34 | G-protein-signaling modulator 1 OS=Mus musculus OX=10090 GN= | 13 | 7 | 7 | 7 | 673 | 74,3 | 6,23 | 9,96 | 7 | 1,04 | 1,061 | 1,002 | 1,082 | 1,057 | 1,053 | 1,153 | 1,032 | 0,97 | -0,03 | 4,2E-01 |
| P18572 | Basigin OS=Mus musculus OX=10090 GN=Bsg PE=1 SV=2 | 21 | 7 | 9 | 7 | 389 | 42,4 | 5,85 | 24,71 | 7 | 0,937 | 0,994 | 0,947 | 0,863 | 0,895 | 1,001 | 0,986 | 0,984 | 0,97 | -0,03 | 4,2E-01 |
| O35459 | Delta(3,5)-Delta(2,4)-dienoyl-CoA isomerase, mitochondrial OS=Mu | 13 | 4 | 4 | 4 | 327 | 36,1 | 7,71 | 5,63 | 4 | 0,9 | 1,004 | 0,979 | 1,084 | 1,066 | 1,198 | 0,869 | 1,106 | 0,94 | -0,06 | 4,2E-01 |
| P61979 | Heterogeneous nuclear ribonucleoprotein K OS=Mus musculus OX= | 27 | 12 | 15 | 12 | 463 | 50,9 | 5,54 | 26,97 | 12 | 0,938 | 0,921 | 0,964 | 0,974 | 1,116 | 1,062 | 0,861 | 0,957 | 0,95 | -0,05 | 4,2E-01 |
| Q9EP53-3 | Isoform 3 of Hamartin OS=Mus musculus OX=10090 GN=Tsc1 | 6 | 5 | 6 | 5 | 1155 | 128,2 | 6,37 | 10,02 | 5 | 1,033 | 1,095 | 1,087 | 0,914 | 0,867 | 1,156 | 0,948 | 0,891 | 1,07 | 0,07 | 4,2E-01 |
| Q80ZI6 | E3 ubiquitin-protein ligase LRSAM1 OS=Mus musculus OX=10090 G | 5 | 3 | 3 | 3 | 727 | 83,9 | 6,14 | 7,25 | 3 | 1,177 | 1,138 | 0,985 | 0,871 | 0,961 | 0,943 | 0,954 | 1,055 | 1,07 | 0,07 | 4,2E-01 |
| Q91XU3 | Phosphatidylinositol 5-phosphate 4-kinase type-2 gamma OS=Mus | 14 | 6 | 7 | 6 | 421 | 47,3 | 6,89 | 12,44 | 6 | 0,991 | 1,179 | 0,943 | 0,921 | 1,128 | 1,092 | 0,929 | 1,149 | 0,94 | -0,06 | 4,2E-01 |
| Q9JJY4 | Probable ATP-dependent RNA helicase DDX20 OS=Mus musculus O | 1 | 1 | 1 | 1 | 825 | 91,7 | 6,74 | 1,91 | 1 | 1,022 | 0,268 | 1,139 | 1,115 | 0,862 | 0,762 | 0,494 | 0,666 | 1,27 | 0,27 | 4,2E-01 |
| Q61410 | cGMP-dependent protein kinase 2 OS=Mus musculus OX=10090 GN | 1 | 1 | 1 | 1 | 762 | 87 | 8,22 | 2,44 | 1 | 1,009 | 0,997 | 1,001 | 1,116 | 0,923 | 2,194 | 0,954 | 1,095 | 0,80 | -0,20 | 4,2E-01 |
| Q61644 | Protein kinase C and casein kinase substrate in neurons protein 1 C | 42 | 17 | 29 | 16 | 441 | 50,5 | 5,24 | 71,96 | 17 | 0,866 | 0,878 | 0,852 | 0,996 | 0,952 | 0,853 | 0,976 | 0,959 | 0,96 | -0,04 | 4,2E-01 |
| P33175 | Kinesin heavy chain isoform 5A OS=Mus musculus OX=10090 GN=K | 12 | 11 | 14 | 7 | 1027 | 116,9 | 5,94 | 21,46 | 11 | 1,328 | 1,393 | 1,162 | 1,182 | 0,804 | 1,184 | 1,429 | 1,168 | 1,10 | 0,10 | 4,3E-01 |
| Q6WVG3 | BTB/POZ domain-containing protein KCTD12 OS=Mus musculus OX | 15 | 4 | 6 | 4 | 327 | 35,9 | 5,81 | 14,51 | 4 | 1,295 | 1,162 | 0,986 | 1,152 | 1,139 | 1,501 | 1,043 | 1,32 | 0,92 | -0,08 | 4,3E-01 |
| P60761 | Neurogranin OS=Mus musculus OX=10090 GN=Nrgn PE=1 SV=1 | 18 | 1 | 4 | 1 | 78 | 7,5 | 7,05 | 2,19 | 1 | 0,499 | 0,799 | 0,935 | 1,066 | 1,042 | 1,022 | 0,802 | 0,89 | 0,88 | -0,12 | 4,3E-01 |
| Q8C1A5 | Thimet oligopeptidase OS=Mus musculus OX=10090 GN=Thop1 PE= | 16 | 9 | 12 | 9 | 687 | 78 | 6,06 | 22,09 | 9 | 0,884 | 0,848 | 0,983 | 0,951 | 0,923 | 0,945 | 0,941 | 0,966 | 0,97 | -0,03 | 4,3E-01 |
| P62631 | Elongation factor 1-alpha 2 OS=Mus musculus OX=10090 GN=Eef1a | 21 | 9 | 14 | 3 | 463 | 50,4 | 9,03 | 33,28 | 9 | 0,711 | 0,927 | 0,858 | 0,997 | 1,018 | 0,806 | 0,994 | 0,938 | 0,93 | -0,07 | 4,3E-01 |
| Q501J6 | Probable ATP-dependent RNA helicase DDX17 OS=Mus musculus O | 4 | 2 | 2 | 1 | 650 | 72,4 | 8,59 | 5,66 | 2 | 0,761 | 1,287 | 1,106 | 0,883 | 1,141 | 1,232 | 1,07 | 1,021 | 0,90 | -0,10 | 4,3E-01 |
| Q3UN02 | Lysocardiolipin acyltransferase 1 OS=Mus musculus OX=10090 GN= | 13 | 4 | 4 | 4 | 376 | 44,4 | 8,53 | 4,16 | 4 | 1,048 | 1,064 | 0,911 | 0,884 | 1,021 | 0,667 | 0,924 | 0,982 | 1,09 | 0,09 | 4,3E-01 |
| Q8BGD8 | Cytochrome c oxidase assembly factor 6 homolog OS=Mus musculu | 49 | 3 | 3 | 3 | 79 | 9,3 | 8,16 | 3,27 | 3 | 1,195 | 1,237 | 1,132 | 0,952 | 0,958 | 1,171 | 1,015 | 1,105 | 1,06 | 0,06 | 4,3E-01 |
| P56546 | C-terminal-binding protein 2 OS=Mus musculus OX=10090 GN=Ctb| | 11 | 6 | 8 | 1 | 445 | 48,9 | 6,95 | 17,2 | 6 | 1,16 | 1,21 | 0,826 | 0,833 | 0,839 | 1,036 | 1,278 | 1,452 | 0,87 | -0,13 | 4,3E-01 |
| Q6ZQI3 | Malectin OS=Mus musculus OX=10090 GN=Mlec PE=1 SV=2 | 13 | 3 | 3 | 3 | 291 | 32,3 | 6,05 | 0 | 3 | 0,01 | 0,826 | 0,731 | 0,797 | 0,791 | 0,986 | 0,719 | 0,587 | 0,77 | -0,23 | 4,3E-01 |
| Q8C8T7 | Protein ELFN1 OS=Mus musculus OX=10090 GN=Elfn1 PE=2 SV=1 | 2 | 2 | 2 | 1 | 828 | 90,8 | 8,18 | 3,88 | 2 | 1,256 | 0,779 | 0,687 | 0,741 | 0,863 | 1,049 | 0,943 | 1,086 | 0,88 | -0,12 | 4,3E-01 |
| Q9JJU8 | SH3 domain-binding glutamic acid-rich-like protein OS=Mus muscu | 22 | 2 | 3 | 2 | 114 | 12,8 | 4,92 | 0 | 2 | 0,944 | 0,855 | 0,899 | 1,005 | 0,937 | 0,922 | 0,972 | 0,994 | 0,97 | -0,03 | 4,3E-01 |
| O88485 | Cytoplasmic dynein 1 intermediate chain 1 OS=Mus musculus OX=1 | 12 | 7 | 8 | 7 | 628 | 70,7 | 5,12 | 15,39 | 7 | 1,161 | 0,987 | 1,068 | 1,105 | 1,002 | 1,121 | 1,048 | 0,991 | 1,04 | 0,04 | 4,3E-01 |
| Q9CY18 | Sorting nexin-7 OS=Mus musculus OX=10090 GN=Snx7 PE=1 SV=1 | 20 | 7 | 8 | 7 | 387 | 45 | 5,07 | 14,87 | 7 | 1,215 | 1,182 | 1,04 | 1,091 | 1,025 | 1,238 | 0,98 | 1,049 | 1,05 | 0,05 | 4,3E-01 |
| Q60864 | Stress-induced-phosphoprotein 1 OS=Mus musculus OX=10090 GN | 38 | 20 | 29 | 20 | 543 | 62,5 | 6,8 | 63,17 | 20 | 0,874 | 0,852 | 0,919 | 1,067 | 1,034 | 0,825 | 1,061 | 1,041 | 0,94 | -0,06 | 4,3E-01 |
| Q8CGE9 | Regulator of G-protein signaling 12 OS=Mus musculus OX=10090 G | 1 | 2 | 3 | 2 | 1381 | 149,5 | 7,59 | 2,02 | 2 | 1,416 | 1,284 | 1,183 | 1,178 | 0,994 | 1,469 | 1,084 | 1,113 | 1,09 | 0,09 | 4,3E-01 |
| Q8BVI5 | Syntaxin-16 OS=Mus musculus OX=10090 GN=Stx16 PE=1 SV=3 | 2 | 1 | 2 | 1 | 326 | 37,1 | 5,86 | 4,06 | 1 | 0,73 | 0,865 | 0,868 | 0,785 | 0,746 | 0,763 | 0,862 | 0,724 | 1,05 | 0,05 | 4,3E-01 |
| Q9CPS8 | Small membrane A-kinase anchor protein OS=Mus musculus OX=1( | 27 | 2 | 3 | 2 | 106 | 11,9 | 4,25 | 6,67 | 2 | 0,794 | 0,724 | 0,945 | 0,906 | 0,906 | 0,837 | 0,98 | 0,85 | 0,94 | -0,06 | 4,3E-01 |
| Q9D1H7 | Golgi to ER traffic protein 4 homolog OS=Mus musculus OX=10090 | 12 | 3 | 4 | 3 | 327 | 36,5 | 5,41 | 5,12 | 3 | 1,499 | 1,267 | 1,165 | 0,996 | 0,956 | 1,436 | 0,887 | 1,103 | 1,12 | 0,12 | 4,3E-01 |
| Q9R269 | Periplakin OS=Mus musculus OX=10090 GN=Ppl PE=1 SV=1 | 1 | 1 | 1 | 1 | 1755 | 203,9 | 5,54 | 0 | 1 | 1,045 | 0,854 | 1,014 | 0,95 | 0,91 | 0,872 | 1,067 | 0,752 | 1,07 | 0,07 | 4,3E-01 |
| Q64343 | ATP-binding cassette sub-family G member 1 OS=Mus musculus OX | 2 | 1 | 1 | 1 | 666 | 74 | 7,06 | 0 | 1 | 0,979 | 0,759 | 1,018 | 1,34 | 1,068 | 0,894 | 0,863 | 0,828 | 1,12 | 0,12 | 4,3E-01 |
| Q61024 | Asparagine synthetase [glutamine-hydrolyzing] OS=Mus musculus ( | 6 | 3 | 3 | 3 | 561 | 64,2 | 6,58 | 2,69 | 3 | 0,962 | 1,082 | 1,22 | 1,299 | 1,017 | 0,931 | 1,231 | 1,054 | 1,08 | 0,08 | 4,3E-01 |
| Q8CGY8 | UDP-N-acetylglucosamine--peptide N-acetylglucosaminyltransferas | 19 | 17 | 22 | 17 | 1046 | 116,9 | 6,7 | 38,09 | 17 | 1,001 | 1,016 | 1,037 | 1,015 | 0,99 | 1,027 | 0,984 | 1,023 | 1,01 | 0,01 | 4,3E-01 |
| P50428 | Arylsulfatase A OS=Mus musculus OX=10090 GN=Arsa PE=1 SV=2 | 2 | 1 | 1 | 1 | 506 | 53,7 | 5,87 | 1,8 | 1 | 0,932 | 1,092 | 1,294 | 1,18 | 1,199 | 1,241 | 1,52 | 0,988 | 0,91 | -0,09 | 4,3E-01 |
| Q9WTR1 | Transient receptor potential cation channel subfamily V member 2 | 6 | 4 | 4 | 4 | 756 | 85,9 | 6,99 | 7,94 | 4 | 1,045 | 1,313 | 0,988 | 0,971 | 0,893 | 1,098 | 0,923 | 1,081 | 1,08 | 0,08 | 4,3E-01 |
| Q8R191 | Synaptogyrin-3 OS=Mus musculus OX=10090 GN=Syngr3 PE=1 SV=: | 12 | 2 | 3 | 2 | 229 | 24,5 | 8,18 | 8,8 | 2 | 1,238 | 1,203 | 0,962 | 0,897 | 0,794 | 1,08 | 1,04 | 1,025 | 1,09 | 0,09 | 4,3E-01 |
| P97772 | Metabotropic glutamate receptor 1 OS=Mus musculus OX=10090 G | 6 | 7 | 9 | 6 | 1199 | 133,1 | 6,86 | 3,65 | 7 | 0,985 | 1,059 | 1,059 | 1,109 | 1,026 | 1,055 | 1,059 | 0,952 | 1,03 | 0,03 | 4,3E-01 |
| Q9JII6 | Aldo-keto reductase family 1 member A1 OS=Mus musculus OX=10 | 22 | 7 | 9 | 7 | 325 | 36,6 | 7,39 | 16,82 | 7 | 0,945 | 1,108 | 1,056 | 0,909 | 0,899 | 0,941 | 0,975 | 1,023 | 1,05 | 0,05 | 4,3E-01 |
| Q61194 | Phosphatidylinositol 4-phosphate 3-kinase C2 domain-containing su | 1 | 1 | 1 | 1 | 1686 | 190,6 | 8,02 | 0 | 1 | 0,901 | 0,742 | 1,135 | 0,813 | 0,726 | 0,821 | 1,01 | 0,642 | 1,12 | 0,12 | 4,3E-01 |
| P55194 | SH3 domain-binding protein 1 OS=Mus musculus OX=10090 GN=Sh | 2 | 1 | 1 | 1 | 680 | 74,1 | 6,6 | 0 | 1 | 2,078 | 2,779 | 3,147 | 1,452 | 2,452 | 2,083 | 1,622 | 1,904 | 1,17 | 0,17 | 4,3E-01 |

| | | | | | | | | | | | | | | | | | | | | |
|---|---|---|---|---|---|---|---|---|---|---|---|---|---|---|---|---|---|---|---|---|
| Q80VQ0 | Aldehyde dehydrogenase family 3 member B1 OS=Mus musculus O | 3 | 1 | 1 | 1 | 468 | 52,3 | 7,55 | 3,43 | 1 | 1,083 | 1,068 | 0,969 | 1,064 | 0,904 | 1,201 | 0,925 | 0,89 | 1,07 | 0,07 | 4,3E-01 |
| Q9QYI5 | DnaJ homolog subfamily B member 2 OS=Mus musculus OX=10090 | 8 | 2 | 2 | 2 | 324 | 35,6 | 5,91 | 4,62 | 2 | 0,881 | 0,973 | 1,12 | 0,908 | 1,018 | 1,505 | 0,911 | 0,947 | 0,89 | -0,11 | 4,3E-01 |
| Q9D1M4 | Eukaryotic translation elongation factor 1 epsilon-1 OS=Mus muscu | 11 | 2 | 2 | 2 | 174 | 19,8 | 8,59 | 5,9 | 2 | 1,006 | 0,961 | 1,105 | 1,216 | 1,119 | 1,071 | 1,219 | 1,098 | 0,95 | -0,05 | 4,3E-01 |
| P42567 | Epidermal growth factor receptor substrate 15 OS=Mus musculus ( | 3 | 2 | 2 | 2 | 897 | 98,4 | 4,6 | 5,08 | 2 | 1,257 | 1,281 | 0,935 | 0,854 | 0,855 | 1,233 | 0,763 | 0,975 | 1,13 | 0,13 | 4,3E-01 |
| Q61315 | Adenomatous polyposis coli protein OS=Mus musculus OX=10090 ( | 1 | 3 | 3 | 3 | 2845 | 310,9 | 7,58 | 2,17 | 3 | 1,396 | 1,536 | 1,08 | 1,23 | 0,812 | 1,286 | 1,236 | 1,374 | 1,11 | 0,11 | 4,3E-01 |
| O88951 | Protein lin-7 homolog B OS=Mus musculus OX=10090 GN=Lin7b PE | 23 | 4 | 6 | 2 | 207 | 22,9 | 8,68 | 11,89 | 4 | 1,102 | 0,884 | 0,883 | 1,25 | 0,993 | 1,135 | 1,167 | 1,153 | 0,93 | -0,07 | 4,3E-01 |
| Q91YD6 | Villin-like protein OS=Mus musculus OX=10090 GN=Vill PE=1 SV=1 | 1 | 1 | 1 | 1 | 859 | 96,4 | 6,35 | 2,19 | 1 | 1,759 | 2,427 | 1,854 | 1,752 | 1,744 | 1,408 | 2,14 | 1,764 | 1,10 | 0,10 | 4,4E-01 |
| Q6P9K8 | Caskin-1 OS=Mus musculus OX=10090 GN=Caskin1 PE=1 SV=2 | 16 | 17 | 22 | 16 | 1431 | 150,4 | 9,17 | 47,92 | 17 | 1,056 | 1,069 | 1,001 | 1,003 | 0,897 | 1,067 | 0,956 | 1,059 | 1,04 | 0,04 | 4,4E-01 |
| P32020 | Non-specific lipid-transfer protein OS=Mus musculus OX=10090 GN | 6 | 3 | 3 | 3 | 547 | 59,1 | 7,44 | 8,05 | 3 | 1,227 | 0,788 | 1,088 | 1,051 | 1,11 | 1,15 | 1,104 | 1,099 | 0,93 | -0,07 | 4,4E-01 |
| Q9CQX5 | Claudin domain-containing protein 1 OS=Mus musculus OX=10090 | 4 | 1 | 2 | 1 | 253 | 28,6 | 6,1 | 4,92 | 1 | 1,104 | 1,231 | 1,104 | 1,125 | 0,983 | 1,106 | 1,049 | 1,226 | 1,05 | 0,05 | 4,4E-01 |
| Q60597 | 2-oxoglutarate dehydrogenase, mitochondrial OS=Mus musculus O | 31 | 26 | 42 | 26 | 1023 | 116,4 | 6,83 | 98,04 | 26 | 1,138 | 0,997 | 1,069 | 1,023 | 1,03 | 1,084 | 1,001 | 0,986 | 1,03 | 0,03 | 4,4E-01 |
| O88746 | Target of Myb protein 1 OS=Mus musculus OX=10090 GN=Tom1 PE | 9 | 4 | 4 | 4 | 492 | 54,3 | 4,94 | 2,7 | 4 | 0,879 | 0,94 | 0,884 | 0,922 | 1,022 | 0,874 | 0,861 | 1,028 | 0,96 | -0,04 | 4,4E-01 |
| Q8VCM4 | Lipoyltransferase 1, mitochondrial OS=Mus musculus OX=10090 GN | 4 | 1 | 1 | 1 | 373 | 42,1 | 8,48 | 0 | 1 | 1,472 | 1,055 | 1,128 | 1,34 | 1,188 | 1,459 | 0,857 | 0,935 | 1,13 | 0,13 | 4,4E-01 |
| P47963 | 60S ribosomal protein L13 OS=Mus musculus OX=10090 GN=Rpl13 | 23 | 5 | 6 | 5 | 211 | 24,3 | 11,55 | 14,88 | 5 | 0,995 | 1,063 | 0,981 | 0,944 | 1,062 | 1,074 | 0,928 | 1,061 | 0,97 | -0,03 | 4,4E-01 |
| Q62421 | Endophilin-A3 OS=Mus musculus OX=10090 GN=Sh3gl3 PE=1 SV=1 | 13 | 5 | 5 | 5 | 347 | 38,9 | 5,12 | 9,92 | 5 | 0,767 | 0,96 | 0,902 | 1,067 | 0,943 | 0,924 | 1,014 | 1,044 | 0,94 | -0,06 | 4,4E-01 |
| Q9JJI8 | 60S ribosomal protein L38 OS=Mus musculus OX=10090 GN=Rpl38 | 14 | 1 | 1 | 1 | 70 | 8,2 | 10,1 | 2,11 | 1 | 1,21 | 1,033 | 1,184 | 1,259 | 1,015 | 0,925 | 0,978 | 1,384 | 1,09 | 0,09 | 4,4E-01 |
| Q9CVB6 | Actin-related protein 2/3 complex subunit 2 OS=Mus musculus OX= | 33 | 11 | 22 | 11 | 300 | 34,3 | 7,36 | 45,67 | 11 | 0,963 | 1,036 | 1,004 | 1,107 | 1,058 | 0,928 | 1,009 | 0,979 | 1,03 | 0,03 | 4,4E-01 |
| Q8CE90 | Dual specificity mitogen-activated protein kinase kinase 7 OS=Mus | 2 | 1 | 2 | 1 | 535 | 59,3 | 9,11 | 0 | 1 | 0,889 | 1,001 | 1,057 | 1,128 | 0,971 | 0,882 | 0,645 | 1,175 | 1,11 | 0,11 | 4,4E-01 |
| P63101 | 14-3-3 protein zeta/delta OS=Mus musculus OX=10090 GN=Ywhaz | 53 | 19 | 54 | 12 | 245 | 27,8 | 4,79 | 150,99 | 19 | 1,04 | 0,909 | 1,018 | 1,144 | 0,985 | 0,924 | 1,028 | 0,998 | 1,04 | 0,04 | 4,4E-01 |
| Q8BJ42 | Disks large-associated protein 2 OS=Mus musculus OX=10090 GN=I | 11 | 10 | 12 | 7 | 1059 | 119 | 6,81 | 23,51 | 10 | 1,102 | 0,998 | 1,045 | 1,052 | 1,002 | 1,262 | 1,011 | 1,138 | 0,95 | -0,05 | 4,4E-01 |
| O88703 | Potassium/sodium hyperpolarization-activated cyclic nucleotide-ga | 12 | 8 | 8 | 7 | 863 | 94,7 | 8,73 | 17,27 | 8 | 0,852 | 1,052 | 1,026 | 1,055 | 1,018 | 1,014 | 1,036 | 1,088 | 0,96 | -0,04 | 4,4E-01 |
| P11881 | Inositol 1,4,5-trisphosphate receptor type 1 OS=Mus musculus OX= | 10 | 26 | 32 | 26 | 2749 | 313 | 6,04 | 52,05 | 26 | 0,982 | 0,996 | 1,003 | 0,957 | 1,072 | 0,925 | 1,034 | 1,016 | 0,97 | -0,03 | 4,4E-01 |
| Q58A65 | C-Jun-amino-terminal kinase-interacting protein 4 OS=Mus muscul | 6 | 7 | 7 | 5 | 1321 | 146,1 | 5,15 | 15,63 | 7 | 0,916 | 0,615 | 0,67 | 0,695 | 0,815 | 0,639 | 0,844 | 0,881 | 0,91 | -0,09 | 4,4E-01 |
| Q8VHW2 | Voltage-dependent calcium channel gamma-8 subunit OS=Mus mu | 7 | 2 | 2 | 2 | 423 | 43,4 | 9,2 | 2,98 | 2 | 1,153 | 0,936 | 0,917 | 0,889 | 0,776 | 1,184 | 0,842 | 0,675 | 1,12 | 0,12 | 4,4E-01 |
| Q9CQJ6 | Density-regulated protein OS=Mus musculus OX=10090 GN=Denr P | 13 | 2 | 2 | 2 | 198 | 22,2 | 5,3 | 5,47 | 2 | 1,761 | 1,364 | 1,35 | 1,393 | 1,474 | 1,14 | 1,088 | 1,625 | 1,10 | 0,10 | 4,4E-01 |
| Q64133 | Amine oxidase [flavin-containing] A OS=Mus musculus OX=10090 G | 17 | 9 | 10 | 9 | 526 | 59,6 | 7,81 | 26,77 | 9 | 1,046 | 0,974 | 0,96 | 0,885 | 0,897 | 1,241 | 1,007 | 0,987 | 0,94 | -0,06 | 4,4E-01 |
| P23506 | Protein-L-isoaspartate(D-aspartate) O-methyltransferase OS=Mus r | 36 | 7 | 8 | 7 | 227 | 24,6 | 7,65 | 16,26 | 7 | 0,951 | 0,854 | 1,061 | 1,098 | 0,995 | 0,86 | 0,938 | 0,964 | 1,06 | 0,06 | 4,4E-01 |
| Q6I6G8 | E3 ubiquitin-protein ligase HECW2 OS=Mus musculus OX=10090 GI | 3 | 4 | 4 | 3 | 1578 | 176,1 | 5,41 | 5,17 | 4 | 1,48 | 1,061 | 1,216 | 1,357 | 1,253 | 1,148 | 1,096 | 1,284 | 1,07 | 0,07 | 4,4E-01 |
| Q6PB44 | Tyrosine-protein phosphatase non-receptor type 23 OS=Mus musc | 6 | 9 | 10 | 9 | 1692 | 185,1 | 6,8 | 21,59 | 9 | 1,185 | 1,23 | 1,257 | 1,113 | 0,96 | 1,249 | 1,075 | 1,246 | 1,06 | 0,06 | 4,4E-01 |
| Q80YN3 | Breast carcinoma-amplified sequence 1 homolog OS=Mus musculu | 10 | 5 | 6 | 5 | 633 | 67,3 | 6,21 | 14,98 | 5 | 0,885 | 0,91 | 0,96 | 1,019 | 1,18 | 1,007 | 0,957 | 0,868 | 0,94 | -0,06 | 4,4E-01 |
| Q8QZV4 | Serine/threonine-protein kinase 32C OS=Mus musculus OX=10090 | 11 | 4 | 6 | 4 | 488 | 55,2 | 6,16 | 2,46 | 4 | 0,77 | 1,098 | 0,972 | 0,847 | 0,931 | 0,98 | 0,908 | 1,184 | 0,92 | -0,08 | 4,4E-01 |
| Q9Z2A0 | 3-phosphoinositide-dependent protein kinase 1 OS=Mus musculus | 11 | 5 | 7 | 5 | 559 | 63,7 | 7,37 | 4,34 | 5 | 1,159 | 0,741 | 0,957 | 0,918 | 0,921 | 1,1 | 1,012 | 1,052 | 0,92 | -0,08 | 4,4E-01 |
| Q3U186 | Probable arginine--tRNA ligase, mitochondrial OS=Mus musculus O | 3 | 2 | 2 | 2 | 578 | 65,3 | 8,02 | 2,59 | 2 | 1,294 | 1,109 | 1,183 | 1,232 | 1,124 | 2,829 | 1,201 | 1,074 | 0,77 | -0,23 | 4,4E-01 |
| Q8BHH2 | Ras-related protein Rab-9B OS=Mus musculus OX=10090 GN=Rab9 | 12 | 2 | 2 | 2 | 201 | 22,7 | 4,93 | 0 | 2 | 1,11 | 1,53 | 1,395 | 1,084 | 0,655 | 1,358 | 1,104 | 1,349 | 1,15 | 0,15 | 4,4E-01 |
| Q9Z0F7 | Gamma-synuclein OS=Mus musculus OX=10090 GN=Sncg PE=1 SV= | 35 | 6 | 6 | 4 | 123 | 13,2 | 4,65 | 12,69 | 6 | 0,931 | 0,854 | 0,978 | 0,812 | 0,906 | 0,873 | 0,952 | 0,996 | 0,96 | -0,04 | 4,4E-01 |
| P56135 | ATP synthase subunit f, mitochondrial OS=Mus musculus OX=1009( | 33 | 3 | 8 | 3 | 88 | 10,3 | 9,95 | 20,57 | 3 | 0,869 | 0,981 | 1,011 | 0,942 | 0,948 | 0,95 | 1,002 | 1,021 | 0,97 | -0,03 | 4,4E-01 |
| P99026 | Proteasome subunit beta type-4 OS=Mus musculus OX=10090 GN= | 6 | 1 | 1 | 1 | 264 | 29,1 | 5,64 | 3,16 | 1 | 1,315 | 0,96 | 1,163 | 0,965 | 0,826 | 1,289 | 0,899 | 0,949 | 1,11 | 0,11 | 4,4E-01 |
| Q8BLY2 | Threonine--tRNA ligase 2, cytoplasmic OS=Mus musculus OX=1009( | 7 | 5 | 5 | 4 | 790 | 91,3 | 7,53 | 8,18 | 5 | 1,122 | 1,112 | 1,15 | 0,825 | 1,01 | 1,043 | 0,949 | 0,944 | 1,07 | 0,07 | 4,4E-01 |
| Q9CZX8 | 40S ribosomal protein S19 OS=Mus musculus OX=10090 GN=Rps19 | 28 | 4 | 4 | 4 | 145 | 16,1 | 10,4 | 12,52 | 4 | 1,107 | 0,988 | 1,169 | 1,225 | 1,219 | 0,924 | 1,034 | 1,051 | 1,06 | 0,06 | 4,4E-01 |
| Q6PE15 | Mycophenolic acid acyl-glucuronide esterase, mitochondrial OS=M | 12 | 4 | 4 | 4 | 297 | 33 | 8,79 | 5,58 | 4 | 1,019 | 0,94 | 1,076 | 0,969 | 0,994 | 1,242 | 0,98 | 1,014 | 0,95 | -0,05 | 4,4E-01 |
| Q9QYF1 | Retinol dehydrogenase 11 OS=Mus musculus OX=10090 GN=Rdh11 | 12 | 3 | 4 | 3 | 316 | 35,1 | 8,91 | 11,53 | 3 | 0,99 | 0,881 | 0,932 | 0,92 | 0,931 | 1,149 | 0,863 | 0,994 | 0,95 | -0,05 | 4,4E-01 |
| Q9WTQ5 | A-kinase anchor protein 12 OS=Mus musculus OX=10090 GN=Akap | 8 | 11 | 14 | 11 | 1684 | 180,6 | 4,44 | 23,34 | 11 | 1,16 | 1,211 | 1,218 | 0,981 | 0,976 | 1,22 | 1,02 | 1,101 | 1,06 | 0,06 | 4,4E-01 |
| Q9CXW2 | 28S ribosomal protein S22, mitochondrial OS=Mus musculus OX=1( | 10 | 3 | 5 | 3 | 359 | 41,2 | 8,56 | 12,11 | 3 | 0,995 | 1,2 | 1,02 | 1,05 | 1,039 | 1,008 | 1,061 | 0,999 | 1,04 | 0,04 | 4,4E-01 |
| Q9ESX5 | H/ACA ribonucleoprotein complex subunit DKC1 OS=Mus musculus | 2 | 1 | 1 | 1 | 509 | 57,4 | 9,28 | 0 | 1 | 1,84 | 1,535 | 1,437 | 1,077 | 1,131 | 1,747 | 1,149 | 1,146 | 1,14 | 0,14 | 4,4E-01 |
| Q9D8B4 | NADH dehydrogenase [ubiquinone] 1 alpha subcomplex subunit 11 | 24 | 2 | 2 | 2 | 141 | 15 | 8,35 | 4,39 | 2 | 1,083 | 1,437 | 1,323 | 1,469 | 1,435 | 1,083 | 1,313 | 1,071 | 1,08 | 0,08 | 4,4E-01 |
| Q9D7G0 | Ribose-phosphate pyrophosphokinase 1 OS=Mus musculus OX=100 | 20 | 5 | 6 | 2 | 318 | 34,8 | 6,98 | 12,01 | 5 | 0,96 | 0,973 | 1,235 | 0,958 | 0,805 | 1,048 | 0,962 | 1,025 | 1,07 | 0,07 | 4,4E-01 |
| Q0KK55 | Kinase non-catalytic C-lobe domain-containing protein 1 OS=Mus n | 1 | 1 | 1 | 1 | 1742 | 191,2 | 6,3 | 0 | 1 | 0,79 | 0,864 | 0,78 | 0,997 | 0,991 | 0,525 | 1,139 | 1,397 | 0,85 | -0,15 | 4,4E-01 |
| P50247 | Adenosylhomocysteinase OS=Mus musculus OX=10090 GN=Ahcy P | 26 | 12 | 18 | 12 | 432 | 47,7 | 6,54 | 32,45 | 12 | 0,933 | 0,921 | 0,958 | 1,039 | 1,009 | 0,926 | 1,042 | 0,992 | 0,97 | -0,03 | 4,4E-01 |
| P17426 | AP-2 complex subunit alpha-1 OS=Mus musculus OX=10090 GN=Ap | 31 | 30 | 62 | 23 | 977 | 107,6 | 7,03 | 134,58 | 30 | 0,897 | 0,935 | 1,01 | 0,989 | 0,978 | 0,944 | 1,03 | 0,981 | 0,97 | -0,03 | 4,4E-01 |
| Q01063 | cAMP-specific 3',5'-cyclic phosphodiesterase 4D OS=Mus musculus | 6 | 4 | 4 | 3 | 747 | 84,5 | 4,91 | 10,67 | 4 | 1,062 | 1,173 | 1,241 | 1,055 | 1,191 | 1,081 | 1,043 | 1,025 | 1,04 | 0,04 | 4,5E-01 |
| Q8VE70 | Programmed cell death protein 10 OS=Mus musculus OX=10090 GI | 17 | 4 | 5 | 4 | 212 | 24,7 | 8,19 | 10,43 | 4 | 0,964 | 1,002 | 1,002 | 0,964 | 0,932 | 1,101 | 0,995 | 1,024 | 0,97 | -0,03 | 4,5E-01 |
| Q8VHQ3 | Protein phosphatase 1 regulatory inhibitor subunit 16B OS=Mus mu | 4 | 2 | 2 | 2 | 568 | 63,5 | 6,58 | 5,83 | 2 | 1,279 | 1,206 | 1,046 | 0,836 | 1,016 | 1,137 | 0,881 | 0,968 | 1,09 | 0,09 | 4,5E-01 |
| Q9D2V7 | Coronin-7 OS=Mus musculus OX=10090 GN=Coro7 PE=1 SV=2 | 3 | 2 | 3 | 2 | 922 | 100,7 | 5,77 | 6,76 | 2 | 1,112 | 1,184 | 1,031 | 0,916 | 0,949 | 1,149 | 0,895 | 0,991 | 1,07 | 0,07 | 4,5E-01 |
| Q8R1S0 | Ubiquinone biosynthesis monooxygenase COQ6, mitochondrial OS= | 5 | 2 | 2 | 2 | 476 | 51,4 | 7,17 | 4,61 | 2 | 0,984 | 0,778 | 0,959 | 0,965 | 1,043 | 1,076 | 0,749 | 1,142 | 0,92 | -0,08 | 4,5E-01 |
| Q9CZ28 | Vacuolar-sorting protein SNF8 OS=Mus musculus OX=10090 GN=Sn | 14 | 3 | 4 | 3 | 258 | 28,9 | 6,65 | 8,31 | 3 | 1,287 | 1,133 | 0,956 | 0,892 | 0,912 | 1,038 | 0,904 | 1,087 | 1,08 | 0,08 | 4,5E-01 |
| Q8BIG7 | Catechol O-methyltransferase domain-containing protein 1 OS=Mu | 8 | 2 | 2 | 2 | 262 | 28,9 | 8,32 | 5,34 | 2 | 0,81 | 0,784 | 0,776 | 0,897 | 0,817 | 0,942 | 0,943 | 0,747 | 0,95 | -0,05 | 4,5E-01 |
| E9Q3L2 | Phosphatidylinositol 4-kinase alpha OS=Mus musculus OX=10090 G | 15 | 27 | 31 | 27 | 2105 | 236,9 | 7,06 | 69,5 | 27 | 0,97 | 1,019 | 0,981 | 0,973 | 0,997 | 1,055 | 0,985 | 0,975 | 0,98 | -0,02 | 4,5E-01 |
| P97427 | Dihydropyrimidinase-related protein 1 OS=Mus musculus OX=1009 | 30 | 13 | 34 | 11 | 572 | 62,1 | 7,12 | 91,46 | 13 | 1,059 | 1,083 | 0,991 | 1,029 | 0,879 | 1,008 | 1,068 | 1,053 | 1,04 | 0,04 | 4,5E-01 |
| Q9D1G5 | Leucine-rich repeat-containing protein 57 OS=Mus musculus OX=1( | 14 | 3 | 4 | 3 | 239 | 26,7 | 8,35 | 6,97 | 3 | 1,08 | 1 | 1,037 | 1,051 | 0,942 | 0,964 | 1,068 | 1,071 | 1,03 | 0,03 | 4,5E-01 |
| Q9CPV4 | Glyoxalase domain-containing protein 4 OS=Mus musculus OX=100 | 27 | 6 | 8 | 6 | 298 | 33,3 | 5,47 | 24,22 | 6 | 1,089 | 1,019 | 1,129 | 1,124 | 1,091 | 1,223 | 1,057 | 1,133 | 0,97 | -0,03 | 4,5E-01 |
| P58281 | Dynamin-like 120 kDa protein, mitochondrial OS=Mus musculus OX | 34 | 30 | 40 | 30 | 960 | 111,3 | 7,55 | 86,44 | 30 | 1,013 | 1,017 | 1,038 | 0,956 | 0,957 | 1,08 | 1,057 | 1,034 | 0,97 | -0,03 | 4,5E-01 |
| Q9ES28 | Rho guanine nucleotide exchange factor 7 OS=Mus musculus OX=1 | 14 | 12 | 12 | 12 | 862 | 97 | 6,8 | 23,2 | 12 | 1,122 | 1,055 | 1,103 | 0,941 | 0,9 | 1,113 | 1,016 | 0,998 | 1,05 | 0,05 | 4,5E-01 |

| | | | | | | | | | | | | | | | | | | | | | |
|---|---|---|---|---|---|---|---|---|---|---|---|---|---|---|---|---|---|---|---|---|---|
| O35621 | Phosphomannomutase 1 OS=Mus musculus OX=10090 GN=Pmm1 | 10 | 2 | 2 | 2 | 262 | 29,8 | 5,47 | 2,75 | 2 | 0,871 | 0,833 | 1,035 | 1,18 | 1,155 | 0,85 | 1,189 | 1,084 | 0,92 | -0,08 | 4,5E-01 |
| Q64525 | Histone H2B type 2-B OS=Mus musculus OX=10090 GN=Hist2h2bb | 62 | 9 | 11 | 9 | 126 | 13,9 | 10,32 | 25,91 | 9 | 0,861 | 1,417 | 0,803 | 0,554 | 1,112 | 1,635 | 0,669 | 1,092 | 0,81 | -0,19 | 4,5E-01 |
| Q8BJY1 | 26S proteasome non-ATPase regulatory subunit 5 OS=Mus musculu | 7 | 3 | 3 | 3 | 504 | 55,9 | 5,21 | 11,3 | 3 | 1,48 | 1,082 | 1,316 | 1,078 | 1,04 | 1,233 | 1,098 | 1,231 | 1,08 | 0,08 | 4,5E-01 |
| Q8VBV7 | COP9 signalosome complex subunit 8 OS=Mus musculus OX=10090 | 24 | 3 | 3 | 3 | 209 | 23,2 | 5,2 | 10,89 | 3 | 1,157 | 1,331 | 1,126 | 0,993 | 0,847 | 1,268 | 0,961 | 1,151 | 1,09 | 0,09 | 4,5E-01 |
| O88587 | Catechol O-methyltransferase OS=Mus musculus OX=10090 GN=Co | 12 | 3 | 4 | 3 | 265 | 29,5 | 5,83 | 4,76 | 3 | 0,933 | 0,977 | 0,954 | 0,97 | 1,007 | 0,934 | 1,054 | 0,938 | 0,97 | -0,03 | 4,5E-01 |
| Q3TYD6 | Serine/threonine-protein kinase LMTK2 OS=Mus musculus OX=100 | 1 | 2 | 2 | 2 | 1471 | 160,4 | 4,51 | 2,06 | 2 | 1,541 | 1,438 | 1,259 | 1,031 | 1,23 | 1,342 | 0,942 | 1,291 | 1,10 | 0,10 | 4,5E-01 |
| Q99M71 | Mammalian ependymin-related protein 1 OS=Mus musculus OX=1( | 21 | 4 | 7 | 4 | 224 | 25,5 | 7,58 | 13,7 | 4 | 1,113 | 1,158 | 1,093 | 1,21 | 1,039 | 1,261 | 0,964 | 1,089 | 1,05 | 0,05 | 4,5E-01 |
| Q924C1 | Exportin-5 OS=Mus musculus OX=10090 GN=Xpo5 PE=1 SV=1 | 2 | 2 | 2 | 2 | 1204 | 136,9 | 5,82 | 0 | 2 | 1,064 | 1,866 | 1,241 | 1,606 | 1,145 | 1,084 | 1,226 | 1,62 | 1,14 | 0,14 | 4,5E-01 |
| A2A5R2 | Brefeldin A-inhibited guanine nucleotide-exchange protein 2 OS=M | 6 | 10 | 10 | 8 | 1792 | 202,1 | 6,55 | 21,04 | 10 | 0,929 | 0,915 | 0,984 | 1,009 | 1,086 | 0,938 | 0,948 | 0,996 | 0,97 | -0,03 | 4,5E-01 |
| Q99J47 | Dehydrogenase/reductase SDR family member 7B OS=Mus muscul | 4 | 1 | 1 | 1 | 323 | 35 | 9,63 | 2,29 | 1 | 0,924 | 1,429 | 0,773 | 1,967 | 0,759 | 1,232 | 1,031 | 1,136 | 1,22 | 0,22 | 4,5E-01 |
| Q9D7V2 | LysM and putative peptidoglycan-binding domain-containing prote | 10 | 2 | 2 | 1 | 215 | 23,7 | 5,71 | 4,33 | 2 | 0,816 | 0,925 | 0,666 | 1,262 | 0,928 | 0,705 | 0,886 | 0,698 | 1,14 | 0,14 | 4,5E-01 |
| Q60803 | TNF receptor-associated factor 3 OS=Mus musculus OX=10090 GN= | 8 | 4 | 4 | 4 | 567 | 64,3 | 8,05 | 5,29 | 4 | 1,233 | 1,172 | 1,006 | 1,06 | 1,061 | 1,166 | 1,06 | 0,975 | 1,05 | 0,05 | 4,5E-01 |
| P98200 | Phospholipid-transporting ATPase IB OS=Mus musculus OX=10090 | 7 | 8 | 10 | 7 | 1148 | 129,3 | 7,33 | 13,28 | 8 | 0,943 | 0,985 | 0,929 | 0,873 | 0,89 | 1,204 | 0,949 | 0,929 | 0,94 | -0,06 | 4,5E-01 |
| Q99KQ4 | Nicotinamide phosphoribosyltransferase OS=Mus musculus OX=10 | 16 | 6 | 6 | 6 | 491 | 55,4 | 7,15 | 14,19 | 6 | 1,156 | 1,115 | 0,941 | 1,079 | 0,848 | 0,956 | 1,054 | 1,167 | 1,07 | 0,07 | 4,5E-01 |
| Q6PHS9 | Voltage-dependent calcium channel subunit alpha-2/delta-2 OS=M | 5 | 6 | 7 | 6 | 1154 | 130,3 | 5,66 | 10,12 | 6 | 1,036 | 0,977 | 1,1 | 1,039 | 0,969 | 1,287 | 1,199 | 0,97 | 0,94 | -0,06 | 4,5E-01 |
| P12367 | cAMP-dependent protein kinase type II-alpha regulatory subunit O | 16 | 8 | 10 | 6 | 401 | 45,4 | 4,89 | 26,75 | 8 | 0,94 | 0,9 | 0,849 | 0,923 | 0,922 | 0,837 | 0,972 | 1,024 | 0,96 | -0,04 | 4,5E-01 |
| Q8K012 | Formin-binding protein 1-like OS=Mus musculus OX=10090 GN=Fnl | 8 | 5 | 6 | 5 | 605 | 69,8 | 6,64 | 9,8 | 5 | 0,876 | 0,959 | 0,957 | 1,162 | 0,983 | 0,981 | 0,861 | 0,911 | 1,06 | 0,06 | 4,5E-01 |
| Q91VM9 | Inorganic pyrophosphatase 2, mitochondrial OS=Mus musculus OX= | 13 | 4 | 7 | 4 | 330 | 38,1 | 6,98 | 11,97 | 4 | 1,067 | 0,991 | 1,15 | 0,981 | 0,837 | 1,179 | 0,952 | 0,959 | 1,07 | 0,07 | 4,5E-01 |
| P14152 | Malate dehydrogenase, cytoplasmic OS=Mus musculus OX=10090 ( | 41 | 14 | 29 | 14 | 334 | 36,5 | 6,58 | 70,07 | 14 | 0,814 | 0,794 | 0,941 | 1,011 | 0,975 | 0,822 | 1,066 | 0,93 | 0,94 | -0,06 | 4,5E-01 |
| Q9R0Q7 | Prostaglandin E synthase 3 OS=Mus musculus OX=10090 GN=Ptges | 29 | 4 | 5 | 4 | 160 | 18,7 | 4,55 | 14,93 | 4 | 0,868 | 0,907 | 1,057 | 1,118 | 0,897 | 0,887 | 0,985 | 0,973 | 1,06 | 0,06 | 4,5E-01 |
| Q9CZ44 | NSFL1 cofactor p47 OS=Mus musculus OX=10090 GN=Nsfl1c PE=1 S | 30 | 10 | 12 | 10 | 370 | 40,7 | 5,15 | 31,15 | 10 | 0,999 | 1,04 | 0,939 | 0,961 | 1,001 | 0,899 | 0,978 | 0,961 | 1,03 | 0,03 | 4,5E-01 |
| P63044 | Vesicle-associated membrane protein 2 OS=Mus musculus OX=100 | 43 | 6 | 18 | 4 | 116 | 12,7 | 8,13 | 43,11 | 6 | 1,302 | 1,317 | 1,272 | 1,08 | 0,874 | 1,433 | 1,131 | 1,126 | 1,09 | 0,09 | 4,5E-01 |
| Q8CE33 | Kelch-like protein 11 OS=Mus musculus OX=10090 GN=Klhl11 PE=1 | 1 | 1 | 1 | 1 | 709 | 80,4 | 6,01 | 0 | 1 | 0,01 | 0,966 | 1,147 | 1,269 | 1,069 | 0,954 | 1,374 | 0,964 | 0,78 | -0,22 | 4,5E-01 |
| Q64521 | Glycerol-3-phosphate dehydrogenase, mitochondrial OS=Mus mus | 34 | 22 | 45 | 22 | 727 | 80,9 | 6,61 | 111,03 | 22 | 1,087 | 1,121 | 1,084 | 1,038 | 1,005 | 1,134 | 1,025 | 1,06 | 1,03 | 0,03 | 4,5E-01 |
| Q7TNG8 | Probable D-lactate dehydrogenase, mitochondrial OS=Mus muscul | 3 | 1 | 1 | 1 | 484 | 51,8 | 6,62 | 2,87 | 1 | 1,31 | 0,992 | 0,996 | 0,771 | 0,884 | 1,058 | 0,799 | 0,933 | 1,11 | 0,11 | 4,5E-01 |
| Q80UG2 | Plexin-A4 OS=Mus musculus OX=10090 GN=Plxna4 PE=1 SV=3 | 14 | 19 | 24 | 12 | 1893 | 212,4 | 6,83 | 44,71 | 19 | 0,959 | 0,933 | 0,947 | 0,972 | 0,937 | 0,922 | 0,98 | 0,92 | 1,01 | 0,01 | 4,5E-01 |
| P26039 | Talin-1 OS=Mus musculus OX=10090 GN=Tln1 PE=1 SV=2 | 5 | 9 | 9 | 6 | 2541 | 269,7 | 6,18 | 9,64 | 9 | 1,294 | 1,131 | 1,164 | 0,992 | 0,889 | 1,244 | 1,098 | 1,043 | 1,07 | 0,07 | 4,5E-01 |
| Q68EF6 | Brain-enriched guanylate kinase-associated protein OS=Mus muscu | 16 | 7 | 8 | 7 | 600 | 65,3 | 5,87 | 17,57 | 7 | 1,076 | 1,214 | 0,96 | 0,961 | 0,842 | 0,998 | 1,069 | 1,049 | 1,06 | 0,06 | 4,5E-01 |
| G3X9J0 | Signal-induced proliferation-associated 1-like protein 3 OS=Mus mu | 2 | 3 | 4 | 2 | 1776 | 194,9 | 8,32 | 2,58 | 3 | 1,351 | 1,314 | 1,084 | 0,564 | 0,811 | 1,211 | 0,791 | 0,839 | 1,18 | 0,18 | 4,6E-01 |
| Q8K3J1 | NADH dehydrogenase [ubiquinone] iron-sulfur protein 8, mitochon | 27 | 5 | 6 | 5 | 212 | 24 | 6,21 | 14,19 | 5 | 1,089 | 0,937 | 0,908 | 0,897 | 0,877 | 1,23 | 0,982 | 1,018 | 0,93 | -0,07 | 4,6E-01 |
| Q9QZE7 | Translin-associated protein X OS=Mus musculus OX=10090 GN=Tsn | 7 | 2 | 2 | 2 | 290 | 32,9 | 6,55 | 2,64 | 2 | 1,33 | 1,058 | 1,058 | 0,983 | 1,008 | 1,23 | 1,097 | 1,536 | 0,91 | -0,09 | 4,6E-01 |
| P18654 | Ribosomal protein S6 kinase alpha-3 OS=Mus musculus OX=10090 | 9 | 6 | 6 | 2 | 740 | 83,6 | 6,89 | 12,23 | 6 | 1,05 | 1,087 | 0,972 | 0,935 | 1,026 | 1,031 | 0,906 | 0,929 | 1,04 | 0,04 | 4,6E-01 |
| Q8BSZ2 | AP-3 complex subunit sigma-2 OS=Mus musculus OX=10090 GN=Ap | 20 | 4 | 4 | 4 | 193 | 22 | 5,22 | 7,04 | 4 | 1,207 | 0,932 | 1,117 | 0,941 | 0,918 | 0,915 | 0,909 | 1,162 | 1,08 | 0,08 | 4,6E-01 |
| Q9CPU2 | NADH dehydrogenase [ubiquinone] 1 beta subcomplex subunit 2, r | 7 | 1 | 1 | 1 | 105 | 12 | 6,52 | 2,21 | 1 | 1,32 | 1,047 | 0,989 | 1,033 | 0,984 | 1,231 | 0,959 | 0,871 | 1,09 | 0,09 | 4,6E-01 |
| Q8VDT9 | 39S ribosomal protein L50, mitochondrial OS=Mus musculus OX=10 | 18 | 2 | 2 | 2 | 159 | 18,2 | 9,33 | 6,32 | 2 | 1,526 | 1,393 | 1,264 | 1,168 | 1,126 | 1,497 | 1,114 | 1,237 | 1,08 | 0,08 | 4,6E-01 |
| Q60960 | Importin subunit alpha-5 OS=Mus musculus OX=10090 GN=Kpna1 I | 6 | 2 | 3 | 1 | 538 | 60,1 | 5,01 | 6,12 | 2 | 1,396 | 1,16 | 1,23 | 1,352 | 1,165 | 1,241 | 1,546 | 1,551 | 0,93 | -0,07 | 4,6E-01 |
| Q8R123 | FAD synthase OS=Mus musculus OX=10090 GN=Flad1 PE=1 SV=1 | 5 | 2 | 2 | 2 | 492 | 54,7 | 6,64 | 4,1 | 2 | 1,384 | 1,015 | 1,141 | 0,948 | 0,949 | 1,092 | 0,969 | 1,139 | 1,08 | 0,08 | 4,6E-01 |
| Q6Q477 | Plasma membrane calcium-transporting ATPase 4 OS=Mus muscul | 22 | 25 | 51 | 12 | 1205 | 133 | 6,13 | 136,54 | 25 | 0,94 | 1,002 | 0,989 | 1,022 | 0,949 | 1,051 | 1,009 | 1,034 | 0,98 | -0,02 | 4,6E-01 |
| O70481 | E3 ubiquitin-protein ligase UBR1 OS=Mus musculus OX=10090 GN= | 2 | 3 | 3 | 3 | 1757 | 200,1 | 6,05 | 2,77 | 3 | 1,402 | 1,427 | 1,213 | 1,162 | 0,841 | 1,541 | 1,193 | 1,126 | 1,11 | 0,11 | 4,6E-01 |
| Q9DB10 | Essential MCU regulator, mitochondrial OS=Mus musculus OX=100 | 9 | 1 | 1 | 1 | 107 | 11,5 | 8,62 | 0 | 1 | 1,061 | 1,543 | 0,928 | 1,313 | 1,124 | 1,119 | 1,184 | 0,959 | 1,10 | 0,10 | 4,6E-01 |
| Q91X97 | Neurocalcin-delta OS=Mus musculus OX=10090 GN=Ncald PE=1 SV | 42 | 8 | 9 | 6 | 193 | 22,2 | 5,35 | 13,82 | 8 | 1,387 | 1,356 | 1,247 | 0,937 | 0,859 | 1,482 | 0,9 | 1,128 | 1,13 | 0,13 | 4,6E-01 |
| Q9WVM3 | Anaphase-promoting complex subunit 7 OS=Mus musculus OX=100 | 6 | 3 | 3 | 3 | 565 | 63 | 5,72 | 2,17 | 3 | 1,4 | 1,201 | 1,175 | 1,255 | 1,35 | 1,078 | 1,031 | 1,28 | 1,06 | 0,06 | 4,6E-01 |
| Q8BHW2 | Protein OSCP1 OS=Mus musculus OX=10090 GN=Oscp1 PE=1 SV=1 | 14 | 4 | 4 | 4 | 379 | 43,2 | 7,2 | 8,23 | 4 | 1,097 | 0,961 | 1,017 | 1,078 | 0,85 | 1,114 | 1,185 | 1,346 | 0,92 | -0,08 | 4,6E-01 |
| Q8BMA6 | Signal recognition particle subunit SRP68 OS=Mus musculus OX=10 | 3 | 2 | 2 | 2 | 625 | 70,5 | 8,57 | 0 | 2 | 0,674 | 0,787 | 1,151 | 0,847 | 1,153 | 0,648 | 0,921 | 1,286 | 0,86 | -0,14 | 4,6E-01 |
| P63168 | Dynein light chain 1, cytoplasmic OS=Mus musculus OX=10090 GN= | 64 | 5 | 10 | 2 | 89 | 10,4 | 7,4 | 22,92 | 5 | 0,899 | 1,052 | 0,932 | 0,936 | 0,79 | 0,966 | 0,408 | 1,145 | 1,15 | 0,15 | 4,6E-01 |
| Q6W8Q3 | Purkinje cell protein 4-like protein 1 OS=Mus musculus OX=10090 ( | 60 | 3 | 4 | 3 | 68 | 7,5 | 5,52 | 6,07 | 3 | 1,246 | 1,026 | 1,26 | 1,111 | 1,119 | 1,223 | 1,055 | 1,019 | 1,05 | 0,05 | 4,6E-01 |
| O70293 | G protein-coupled receptor kinase 6 OS=Mus musculus OX=10090 ( | 7 | 3 | 3 | 3 | 576 | 65,9 | 8,1 | 0 | 3 | 1,515 | 1,115 | 1,044 | 1,06 | 1,046 | 1,195 | 1,027 | 1,094 | 1,09 | 0,09 | 4,6E-01 |
| Q8VE09 | Tetratricopeptide repeat protein 39C OS=Mus musculus OX=10090 | 2 | 1 | 1 | 1 | 580 | 65,4 | 6,87 | 2,72 | 1 | 1,39 | 0,99 | 0,981 | 0,779 | 0,687 | 0,95 | 1,178 | 0,8 | 1,15 | 0,15 | 4,6E-01 |
| Q923D2 | Flavin reductase (NADPH) OS=Mus musculus OX=10090 GN=Blvrb F | 13 | 2 | 2 | 2 | 206 | 22,2 | 7,01 | 2,8 | 2 | 1,542 | 1,686 | 1,054 | 0,949 | 1,329 | 0,938 | 1,222 | 1,114 | 1,14 | 0,14 | 4,6E-01 |
| P23492 | Purine nucleoside phosphorylase OS=Mus musculus OX=10090 GN= | 4 | 1 | 1 | 1 | 289 | 32,3 | 6,16 | 0 | 1 | 0,01 | 1,203 | 0,735 | 0,843 | 0,903 | 0,51 | 0,922 | 1,456 | 0,74 | -0,26 | 4,6E-01 |
| Q99LH9 | SH3 domain-binding protein 5-like OS=Mus musculus OX=10090 GN | 3 | 1 | 1 | 1 | 392 | 43,3 | 6 | 0 | 1 | 0,527 | 1,073 | 1,17 | 0,774 | 0,663 | 0,669 | 0,881 | 0,836 | 1,16 | 0,16 | 4,6E-01 |
| P50518 | V-type proton ATPase subunit E 1 OS=Mus musculus OX=10090 GN | 42 | 13 | 45 | 13 | 226 | 26,1 | 8,43 | 74 | 13 | 1,034 | 1,087 | 1,051 | 0,986 | 0,879 | 1,005 | 1,062 | 1,061 | 1,04 | 0,04 | 4,6E-01 |
| Q69ZS8 | Kazrin OS=Mus musculus OX=10090 GN=Kazn PE=1 SV=2 | 5 | 3 | 3 | 3 | 779 | 86,7 | 6,9 | 2,88 | 3 | 1,276 | 0,932 | 0,966 | 1,058 | 1,061 | 1,034 | 0,911 | 0,959 | 1,07 | 0,07 | 4,6E-01 |
| P47857 | ATP-dependent 6-phosphofructokinase, muscle type OS=Mus musc | 30 | 19 | 40 | 17 | 780 | 85,2 | 8 | 100,77 | 19 | 1,043 | 0,976 | 0,987 | 0,976 | 0,987 | 1,219 | 1,001 | 0,967 | 0,95 | -0,05 | 4,6E-01 |
| Q61627 | Glutamate receptor ionotropic, delta-1 OS=Mus musculus OX=1009 | 4 | 4 | 4 | 4 | 1009 | 112,1 | 6,7 | 7,1 | 4 | 1,337 | 1,171 | 1,104 | 1,052 | 0,984 | 1,301 | 0,996 | 1,081 | 1,07 | 0,07 | 4,6E-01 |
| Q8CFE4 | SCY1-like protein 2 OS=Mus musculus OX=10090 GN=Scyl2 PE=1 SV | 5 | 3 | 4 | 3 | 930 | 103,3 | 8,02 | 8,62 | 3 | 1,518 | 1,082 | 1,394 | 1,584 | 1,39 | 1,398 | 1,237 | 1,157 | 1,08 | 0,08 | 4,6E-01 |
| P13707 | Glycerol-3-phosphate dehydrogenase [NAD(+)], cytoplasmic OS=M | 18 | 6 | 8 | 6 | 349 | 37,5 | 7,17 | 18,6 | 6 | 1,05 | 0,987 | 1,141 | 1,116 | 0,979 | 1,172 | 1,207 | 1,127 | 0,96 | -0,04 | 4,6E-01 |
| Q9CY27 | Very-long-chain enoyl-CoA reductase OS=Mus musculus OX=10090 | 12 | 5 | 7 | 5 | 308 | 36,1 | 9,55 | 5,2 | 5 | 0,98 | 1,046 | 1,104 | 0,859 | 1,022 | 1,149 | 1,06 | 0,963 | 0,95 | -0,05 | 4,6E-01 |
| Q91YM4 | FAST kinase domain-containing protein 4 OS=Mus musculus OX=10 | 3 | 2 | 2 | 2 | 630 | 71,5 | 8,34 | 5,43 | 2 | 1,194 | 0,932 | 1,229 | 1,084 | 1,048 | 1,15 | 1,019 | 0,985 | 1,06 | 0,06 | 4,6E-01 |
| P15208 | Insulin receptor OS=Mus musculus OX=10090 GN=Insr PE=1 SV=2 | 2 | 2 | 2 | 2 | 1372 | 155,5 | 5,95 | 1,63 | 2 | 1,264 | 1,005 | 0,558 | 0,789 | 0,948 | 0,955 | 0,561 | 0,564 | 1,19 | 0,19 | 4,6E-01 |
| Q9Z0P5 | Twinfilin-2 OS=Mus musculus OX=10090 GN=Twf2 PE=1 SV=1 | 19 | 6 | 6 | 5 | 349 | 39,4 | 6,8 | 15,77 | 6 | 0,971 | 0,908 | 0,992 | 1,076 | 0,974 | 0,895 | 0,959 | 0,992 | 1,03 | 0,03 | 4,6E-01 |
| P61971 | Nuclear transport factor 2 OS=Mus musculus OX=10090 GN=Nutf2 | 19 | 2 | 2 | 2 | 127 | 14,5 | 5,38 | 2,57 | 2 | 1,091 | 1,535 | 0,986 | 1,122 | 0,838 | 1,033 | 1,398 | 0,926 | 1,13 | 0,13 | 4,6E-01 |

| | | | | | | | | | | | | | | | | | | | | | |
|---|---|---|---|---|---|---|---|---|---|---|---|---|---|---|---|---|---|---|---|---|---|
| Q9R1Q9 | V-type proton ATPase subunit S1 OS=Mus musculus OX=10090 GN= | 3 | 1 | 1 | 1 | 463 | 51 | 5,92 | 0 | 1 | 1,369 | 1,568 | 0,812 | 1,312 | 1,129 | 1,544 | 0,882 | 0,772 | 1,17 | 0,17 | 4,6E-01 |
| Q80V91 | Probable E3 ubiquitin-protein ligase DTX3 OS=Mus musculus OX=1( | 9 | 2 | 2 | 2 | 347 | 38 | 8,73 | 2,44 | 2 | 1,57 | 1,366 | 1,363 | 1,363 | 1,425 | 1,399 | 1,392 | 1,237 | 1,04 | 0,04 | 4,6E-01 |
| P80318 | T-complex protein 1 subunit gamma OS=Mus musculus OX=10090 ( | 32 | 16 | 22 | 16 | 545 | 60,6 | 6,7 | 52,4 | 16 | 1,075 | 1,07 | 1,13 | 1,011 | 0,922 | 1,148 | 1,027 | 1,026 | 1,04 | 0,04 | 4,6E-01 |
| Q9D2N4-2 | Isoform 2 of Dystrobrevin alpha OS=Mus musculus OX=10090 GN=I | 11 | 8 | 8 | 8 | 688 | 77,7 | 6,8 | 12,98 | 8 | 1,074 | 1,117 | 0,999 | 1,105 | 1,05 | 1,05 | 1,178 | 1,15 | 0,97 | -0,03 | 4,7E-01 |
| Q8CFI0 | E3 ubiquitin-protein ligase NEDD4-like OS=Mus musculus OX=1009( | 5 | 6 | 8 | 5 | 1004 | 115,3 | 6,15 | 20,15 | 6 | 1,107 | 1,141 | 0,894 | 0,981 | 1,09 | 1,086 | 1,013 | 1,127 | 0,96 | -0,04 | 4,7E-01 |
| Q8R001 | Microtubule-associated protein RP/EB family member 2 OS=Mus m | 33 | 8 | 12 | 7 | 326 | 36,9 | 5,38 | 16,63 | 8 | 1,226 | 1,19 | 1,125 | 1,077 | 1,071 | 1,306 | 0,979 | 1,009 | 1,06 | 0,06 | 4,7E-01 |
| Q80V26 | Inositol monophosphatase 3 OS=Mus musculus OX=10090 GN=Imp | 2 | 1 | 1 | 1 | 356 | 38,6 | 6,47 | 0 | 1 | 0,962 | 0,947 | 1,002 | 1,424 | 0,975 | 0,932 | 1,196 | 0,791 | 1,11 | 0,11 | 4,7E-01 |
| Q921Q7 | Ras and Rab interactor 1 OS=Mus musculus OX=10090 GN=Rin1 PE | 4 | 3 | 3 | 3 | 763 | 83 | 8,72 | 4,31 | 3 | 0,9 | 0,87 | 0,822 | 0,833 | 0,772 | 0,894 | 0,877 | 0,761 | 1,04 | 0,04 | 4,7E-01 |
| Q8R016 | Bleomycin hydrolase OS=Mus musculus OX=10090 GN=Blmh PE=1 | 15 | 7 | 13 | 7 | 455 | 52,5 | 6,48 | 17,52 | 7 | 0,97 | 0,966 | 1,149 | 1,073 | 1,027 | 1,163 | 1,082 | 1,051 | 0,96 | -0,04 | 4,7E-01 |
| P39053-3 | Isoform 3 of Dynamin-1 OS=Mus musculus OX=10090 GN=Dnm1 | 45 | 43 | 99 | 2 | 851 | 95,9 | 6,76 | 202,98 | 43 | 0,71 | 0,923 | 0,937 | 0,928 | 0,973 | 0,804 | 1,01 | 0,931 | 0,94 | -0,06 | 4,7E-01 |
| Q8BTZ5 | Ankyrin repeat domain-containing protein 46 OS=Mus musculus OX | 5 | 1 | 1 | 1 | 228 | 25,2 | 5,73 | 0 | 1 | 1,573 | 1,791 | 1,209 | 1,529 | 0,744 | 1,458 | 1,567 | 1,604 | 1,14 | 0,14 | 4,7E-01 |
| Q14B80 | Potassium voltage-gated channel subfamily C member 2 OS=Mus m | 5 | 2 | 2 | 2 | 642 | 70,5 | 7,69 | 1,75 | 2 | 0,993 | 1,158 | 0,954 | 0,97 | 1,038 | 0,845 | 0,887 | 1,077 | 1,06 | 0,06 | 4,7E-01 |
| Q920E5 | Farnesyl pyrophosphate synthase OS=Mus musculus OX=10090 GN | 10 | 4 | 5 | 4 | 353 | 40,6 | 5,66 | 2,25 | 4 | 1,482 | 1,345 | 1,491 | 1,386 | 1,307 | 1,44 | 1,624 | 1,583 | 0,96 | -0,04 | 4,7E-01 |
| Q03717 | Potassium voltage-gated channel subfamily B member 1 OS=Mus m | 2 | 1 | 2 | 1 | 857 | 95,5 | 8,16 | 7,2 | 1 | 1,65 | 1,138 | 1,566 | 1,478 | 1,733 | 1,486 | 1,353 | 1,708 | 0,93 | -0,07 | 4,7E-01 |
| Q0VBK2 | Keratin, type II cytoskeletal 80 OS=Mus musculus OX=10090 GN=Kr | 2 | 1 | 1 | 1 | 452 | 50,6 | 6,27 | 1,8 | 1 | 1,022 | 1,44 | 0,755 | 0,76 | 1,04 | 0,732 | 0,717 | 0,933 | 1,16 | 0,16 | 4,7E-01 |
| Q62465 | Synaptic vesicle membrane protein VAT-1 homolog OS=Mus muscu | 27 | 8 | 12 | 8 | 406 | 43,1 | 6,37 | 25,65 | 8 | 1,186 | 1,111 | 1,141 | 1,048 | 0,962 | 1,18 | 1,169 | 0,967 | 1,05 | 0,05 | 4,7E-01 |
| Q99L45 | Eukaryotic translation initiation factor 2 subunit 2 OS=Mus musculu | 16 | 5 | 5 | 5 | 331 | 38,1 | 5,8 | 6,78 | 5 | 1,017 | 1,012 | 1,068 | 1,013 | 1,086 | 0,97 | 1,07 | 1,078 | 0,98 | -0,02 | 4,7E-01 |
| Q9Z2Y8 | Pyridoxal phosphate homeostasis protein OS=Mus musculus OX=1( | 26 | 7 | 9 | 7 | 274 | 30 | 8,27 | 21,95 | 7 | 1,048 | 0,913 | 1,004 | 1,015 | 1,055 | 0,939 | 1,012 | 1,123 | 0,96 | -0,04 | 4,7E-01 |
| E9Q7G0 | Nuclear mitotic apparatus protein 1 OS=Mus musculus OX=10090 ( | 0 | 1 | 1 | 1 | 2094 | 235,5 | 5,87 | 0 | 1 | 1,482 | 1,227 | 1,496 | 0,993 | 0,971 | 1,628 | 0,782 | 1,146 | 1,15 | 0,15 | 4,7E-01 |
| Q0VE82 | Copine-7 OS=Mus musculus OX=10090 GN=Cpne7 PE=1 SV=1 | 13 | 5 | 5 | 5 | 557 | 61,9 | 5,4 | 10,04 | 5 | 1,235 | 1,107 | 1,335 | 1,091 | 1,17 | 1,256 | 1,02 | 1,084 | 1,05 | 0,05 | 4,7E-01 |
| P35486 | Pyruvate dehydrogenase E1 component subunit alpha, somatic for | 47 | 17 | 37 | 17 | 390 | 43,2 | 8,19 | 70,27 | 17 | 1,046 | 0,964 | 0,996 | 1,041 | 1,014 | 1,006 | 0,967 | 0,992 | 1,02 | 0,02 | 4,7E-01 |
| Q8K183 | Pyridoxal kinase OS=Mus musculus OX=10090 GN=Pdxk PE=1 SV=1 | 22 | 5 | 11 | 5 | 312 | 35 | 6,29 | 22,7 | 5 | 1,051 | 1,317 | 0,84 | 0,969 | 0,933 | 0,891 | 0,979 | 1,047 | 1,08 | 0,08 | 4,7E-01 |
| Q6NWV3 | Intraflagellar transport protein 122 homolog OS=Mus musculus OX | 1 | 1 | 1 | 1 | 1182 | 134,7 | 6,98 | 2,82 | 1 | 1,585 | 1,647 | 0,888 | 0,907 | 1,486 | 1,34 | 1,472 | 1,378 | 0,89 | -0,11 | 4,7E-01 |
| Q64727 | Vinculin OS=Mus musculus OX=10090 GN=Vcl PE=1 SV=4 | 20 | 19 | 21 | 19 | 1066 | 116,6 | 6 | 37,53 | 19 | 1,171 | 1,221 | 1,081 | 0,957 | 0,946 | 1,193 | 0,998 | 1,051 | 1,06 | 0,06 | 4,7E-01 |
| Q8C8R3-2 | Isoform 2 of Ankyrin-2 OS=Mus musculus OX=10090 GN=Ank2 | 29 | 85 | 115 | 4 | 3926 | 428,5 | 5,16 | 246,89 | 85 | 1,35 | 1,279 | 1,106 | 1,054 | 1,04 | 1,266 | 1,042 | 1,168 | 1,06 | 0,06 | 4,7E-01 |
| Q8BG39 | Synaptic vesicle glycoprotein 2B OS=Mus musculus OX=10090 GN= | 14 | 10 | 19 | 9 | 683 | 77,4 | 5,57 | 49,79 | 10 | 1,009 | 1,095 | 0,937 | 0,984 | 0,963 | 0,997 | 0,968 | 0,992 | 1,03 | 0,03 | 4,7E-01 |
| P47934 | Carnitine O-acetyltransferase OS=Mus musculus OX=10090 GN=Cra | 13 | 8 | 9 | 8 | 626 | 70,8 | 8,44 | 7,2 | 8 | 1,247 | 1,051 | 1,251 | 1,053 | 0,932 | 1,201 | 1,038 | 1,172 | 1,06 | 0,06 | 4,7E-01 |
| Q91YH5 | Atlastin-3 OS=Mus musculus OX=10090 GN=Atl3 PE=1 SV=1 | 2 | 1 | 1 | 1 | 541 | 60,5 | 6,1 | 0 | 1 | 1,467 | 0,764 | 0,637 | 0,766 | 0,826 | 0,864 | 0,73 | 0,61 | 1,20 | 0,20 | 4,7E-01 |
| Q9DAW9 | Calponin-3 OS=Mus musculus OX=10090 GN=Cnn3 PE=1 SV=1 | 8 | 2 | 2 | 2 | 330 | 36,4 | 5,72 | 0 | 2 | 1,14 | 1,157 | 1,242 | 0,893 | 1,126 | 1,396 | 1,432 | 0,916 | 0,91 | -0,09 | 4,7E-01 |
| P11531 | Dystrophin OS=Mus musculus OX=10090 GN=Dmd PE=1 SV=3 | 0 | 1 | 1 | 1 | 3678 | 425,6 | 5,94 | 0 | 1 | 1,432 | 1,303 | 1,717 | 1,2 | 0,994 | 1,325 | 1,193 | 1,614 | 1,10 | 0,10 | 4,7E-01 |
| P70236 | Dual specificity mitogen-activated protein kinase kinase 6 OS=Mus | 16 | 4 | 5 | 4 | 334 | 37,4 | 7,39 | 2,03 | 4 | 1,022 | 1,437 | 0,975 | 0,801 | 0,727 | 0,556 | 1,199 | 1,124 | 1,17 | 0,17 | 4,7E-01 |
| Q9EPL9 | Peroxisomal acyl-coenzyme A oxidase 3 OS=Mus musculus OX=100 | 2 | 1 | 1 | 1 | 700 | 78,4 | 7,12 | 2,73 | 1 | 1,236 | 1,511 | 1,179 | 0,849 | 1,06 | 1,049 | 1,102 | 1,143 | 1,10 | 0,10 | 4,7E-01 |
| Q60790 | Ras GTPase-activating protein 3 OS=Mus musculus OX=10090 GN=I | 9 | 8 | 8 | 8 | 834 | 95,9 | 7,34 | 9,47 | 8 | 1,275 | 1,225 | 1,033 | 1,152 | 0,98 | 1,245 | 1,104 | 1,125 | 1,05 | 0,05 | 4,7E-01 |
| Q9D967 | Magnesium-dependent phosphatase 1 OS=Mus musculus OX=1009 | 13 | 2 | 2 | 2 | 164 | 18,6 | 6,8 | 3,13 | 2 | 1,156 | 1,056 | 0,911 | 0,918 | 0,938 | 1,082 | 0,855 | 0,935 | 1,06 | 0,06 | 4,7E-01 |
| Q8VEH3 | ADP-ribosylation factor-like protein 8A OS=Mus musculus OX=1009 | 17 | 4 | 5 | 3 | 186 | 21,4 | 7,77 | 9,78 | 4 | 0,522 | 0,72 | 0,751 | 1,044 | 0,787 | 0,742 | 0,802 | 1,136 | 0,88 | -0,12 | 4,7E-01 |
| Q5H8C4 | Vacuolar protein sorting-associated protein 13A OS=Mus musculus | 6 | 15 | 16 | 14 | 3166 | 359,2 | 6,19 | 31,8 | 15 | 1,166 | 1,065 | 1,173 | 0,999 | 1,03 | 1,11 | 1,007 | 1,105 | 1,04 | 0,04 | 4,7E-01 |
| Q8K2Q9 | Shootin-1 OS=Mus musculus OX=10090 GN=Shtn1 PE=1 SV=1 | 4 | 2 | 2 | 2 | 631 | 71,3 | 5,44 | 4,68 | 2 | 1,158 | 1,209 | 0,982 | 0,946 | 0,946 | 1,4 | 1,154 | 1,141 | 0,93 | -0,07 | 4,7E-01 |
| Q3UYH7 | Beta-adrenergic receptor kinase 2 OS=Mus musculus OX=10090 GN | 7 | 4 | 5 | 2 | 688 | 79,6 | 7,69 | 14,02 | 4 | 0,937 | 0,913 | 0,929 | 1,319 | 1,109 | 1,299 | 0,935 | 1,131 | 0,92 | -0,08 | 4,7E-01 |
| Q9CRY7 | Lysophospholipase D GDPD1 OS=Mus musculus OX=10090 GN=Gdp | 12 | 3 | 3 | 3 | 314 | 35,8 | 8,31 | 2,38 | 3 | 1,08 | 1,168 | 1,093 | 1,063 | 1,041 | 1,256 | 1,177 | 1,091 | 0,96 | -0,04 | 4,7E-01 |
| Q8BL66 | Early endosome antigen 1 OS=Mus musculus OX=10090 GN=Eea1 F | 13 | 16 | 19 | 16 | 1411 | 160,8 | 5,77 | 49,58 | 16 | 1,072 | 0,932 | 0,973 | 0,964 | 0,983 | 1,074 | 0,983 | 1,014 | 0,97 | -0,03 | 4,7E-01 |
| Q9QUR8 | Semaphorin-7A OS=Mus musculus OX=10090 GN=Sema7a PE=1 SV | 7 | 3 | 3 | 3 | 664 | 74,9 | 7,74 | 6,1 | 3 | 1,191 | 1,141 | 1,158 | 0,944 | 1,031 | 1,107 | 0,995 | 1,11 | 1,05 | 0,05 | 4,8E-01 |
| Q9D0E1 | Heterogeneous nuclear ribonucleoprotein M OS=Mus musculus OX | 4 | 3 | 3 | 3 | 729 | 77,6 | 8,63 | 4,63 | 3 | 1,299 | 1,036 | 1,047 | 0,982 | 1,164 | 1,44 | 0,966 | 1,161 | 0,92 | -0,08 | 4,8E-01 |
| P10639 | Thioredoxin OS=Mus musculus OX=10090 GN=Txn PE=1 SV=3 | 31 | 3 | 5 | 3 | 105 | 11,7 | 4,92 | 13,2 | 3 | 0,861 | 0,936 | 1,033 | 0,897 | 0,901 | 0,959 | 0,977 | 1,028 | 0,96 | -0,04 | 4,8E-01 |
| Q3UMR5 | Calcium uniporter protein, mitochondrial OS=Mus musculus OX=10 | 23 | 10 | 15 | 10 | 350 | 39,7 | 8,56 | 20,57 | 10 | 0,979 | 1,025 | 1,021 | 0,898 | 0,931 | 0,928 | 0,989 | 0,974 | 1,03 | 0,03 | 4,8E-01 |
| Q921H9 | Cytochrome c oxidase assembly factor 7 OS=Mus musculus OX=10( | 12 | 3 | 3 | 3 | 231 | 25,6 | 6,29 | 6,72 | 3 | 0,94 | 0,827 | 0,992 | 0,954 | 0,983 | 0,903 | 0,979 | 0,97 | 0,97 | -0,03 | 4,8E-01 |
| P62751 | 60S ribosomal protein L23a OS=Mus musculus OX=10090 GN=Rpl2 | 31 | 4 | 5 | 4 | 156 | 17,7 | 10,45 | 4,76 | 4 | 1,446 | 1,211 | 1,148 | 0,847 | 1,185 | 1,395 | 1,188 | 1,287 | 0,92 | -0,08 | 4,8E-01 |
| Q9Z2H5 | Band 4.1-like protein 1 OS=Mus musculus OX=10090 GN=Epb41l1 F | 30 | 26 | 40 | 24 | 879 | 98,3 | 5,62 | 83,63 | 26 | 1,054 | 1,036 | 1,06 | 1,084 | 1,027 | 1,123 | 1,103 | 1,054 | 0,98 | -0,02 | 4,8E-01 |
| Q9DC71 | 28S ribosomal protein S15, mitochondrial OS=Mus musculus OX=1( | 9 | 2 | 2 | 2 | 258 | 29,4 | 10,13 | 2,22 | 2 | 1,156 | 0,974 | 0,839 | 0,92 | 1,029 | 1,329 | 0,983 | 0,897 | 0,92 | -0,08 | 4,8E-01 |
| P26645 | Myristoylated alanine-rich C-kinase substrate OS=Mus musculus O) | 49 | 10 | 17 | 10 | 309 | 29,6 | 4,34 | 35,71 | 10 | 0,975 | 0,891 | 1,098 | 0,96 | 0,747 | 0,994 | 1,005 | 0,954 | 1,06 | 0,06 | 4,8E-01 |
| Q91WK5 | Glycine cleavage system H protein, mitochondrial OS=Mus musculu | 6 | 1 | 1 | 1 | 170 | 18,6 | 4,75 | 2,55 | 1 | 0,812 | 0,471 | 0,915 | 1,186 | 1,249 | 0,724 | 0,886 | 1,09 | 0,86 | -0,14 | 4,8E-01 |
| Q9R1V4 | Disintegrin and metalloproteinase domain-containing protein 11 O | 12 | 6 | 6 | 6 | 773 | 84,1 | 7,5 | 13,3 | 6 | 1,197 | 0,973 | 1,195 | 0,978 | 0,945 | 1,135 | 0,986 | 1,048 | 1,06 | 0,06 | 4,8E-01 |
| Q00PI9 | Heterogeneous nuclear ribonucleoprotein U-like protein 2 OS=Mus | 4 | 3 | 4 | 3 | 745 | 84,9 | 4,89 | 1,62 | 3 | 0,907 | 1,082 | 0,74 | 1,18 | 1,044 | 0,915 | 0,805 | 0,806 | 1,09 | 0,09 | 4,8E-01 |
| P43276 | Histone H1.5 OS=Mus musculus OX=10090 GN=Hist1h1b PE=1 SV=2 | 13 | 4 | 6 | 2 | 223 | 22,6 | 10,92 | 13,37 | 4 | 0,867 | 1,539 | 0,864 | 0,676 | 1,1 | 1,68 | 0,713 | 1,285 | 0,83 | -0,17 | 4,8E-01 |
| P97352 | Protein S100-A13 OS=Mus musculus OX=10090 GN=S100a13 PE=1 | 42 | 4 | 7 | 4 | 98 | 11,2 | 6,13 | 14,87 | 4 | 0,97 | 0,921 | 1,026 | 0,954 | 0,975 | 0,887 | 0,962 | 0,958 | 1,02 | 0,02 | 4,8E-01 |
| Q8BHN3 | Neutral alpha-glucosidase AB OS=Mus musculus OX=10090 GN=Ga | 17 | 15 | 21 | 15 | 944 | 106,8 | 6,06 | 38,77 | 15 | 1,218 | 1,017 | 1,021 | 0,948 | 0,988 | 0,951 | 0,973 | 1,093 | 1,05 | 0,05 | 4,8E-01 |
| Q62048 | Astrocytic phosphoprotein PEA-15 OS=Mus musculus OX=10090 GN | 28 | 3 | 7 | 3 | 130 | 15 | 5,02 | 16,9 | 3 | 0,926 | 0,867 | 1,126 | 1,171 | 0,957 | 1 | 0,985 | 0,918 | 1,06 | 0,06 | 4,8E-01 |
| O08688 | Calpain-5 OS=Mus musculus OX=10090 GN=Capn5 PE=1 SV=1 | 17 | 10 | 13 | 10 | 640 | 72,9 | 7,36 | 25,63 | 10 | 1,07 | 1,074 | 0,987 | 1,141 | 1,139 | 0,951 | 1,024 | 1,006 | 1,04 | 0,04 | 4,8E-01 |
| P97950 | Ras-related protein Rab-33A OS=Mus musculus OX=10090 GN=Rab | 15 | 3 | 3 | 3 | 237 | 26,5 | 7,88 | 7,9 | 3 | 0,778 | 1,01 | 0,861 | 0,963 | 0,86 | 0,913 | 0,965 | 1,088 | 0,94 | -0,06 | 4,8E-01 |
| Q99LD9 | Translation initiation factor eIF-2B subunit beta OS=Mus musculus ( | 4 | 1 | 1 | 1 | 351 | 38,9 | 6,24 | 3,23 | 1 | 0,87 | 0,74 | 0,978 | 0,898 | 0,92 | 1,049 | 0,975 | 0,772 | 0,94 | -0,06 | 4,8E-01 |
| Q9CZD3 | Glycine--tRNA ligase OS=Mus musculus OX=10090 GN=Gars PE=1 S | 12 | 11 | 16 | 11 | 729 | 81,8 | 6,65 | 25,31 | 11 | 0,928 | 1,024 | 1,037 | 1,065 | 0,972 | 0,936 | 1,033 | 1,004 | 1,03 | 0,03 | 4,8E-01 |
| Q9CQV7 | Mitochondrial import inner membrane translocase subunit TIM14 ( | 28 | 3 | 3 | 3 | 116 | 12,4 | 10,1 | 5,97 | 3 | 0,808 | 1,103 | 1,071 | 0,989 | 1,045 | 1,022 | 1,089 | 1,019 | 0,95 | -0,05 | 4,8E-01 |
| Q8BTX9 | Inactive hydroxysteroid dehydrogenase-like protein 1 OS=Mus mus | 6 | 2 | 2 | 2 | 330 | 36,8 | 8,48 | 2,45 | 2 | 0,924 | 0,821 | 0,599 | 0,901 | 0,999 | 0,697 | 0,889 | 0,959 | 0,92 | -0,08 | 4,8E-01 |

| | | | | | | | | | | | | | | | | | | | | | |
|---|---|---|---|---|---|---|---|---|---|---|---|---|---|---|---|---|---|---|---|---|---|
| Q5NCE8 | Magnesium transporter MRS2 homolog, mitochondrial OS=Mus mu | 2 | 1 | 1 | 1 | 434 | 49,3 | 7,53 | 1,92 | 1 | 0,401 | 0,775 | 1,129 | 0,936 | 0,802 | 0,833 | 0,911 | 1,25 | 0,85 | -0,15 | 4,8E-01 |
| Q99JT1 | Glutamyl-tRNA(Gln) amidotransferase subunit B, mitochondrial OS= | 4 | 2 | 2 | 2 | 557 | 62,1 | 8,54 | 5,09 | 2 | 0,995 | 1,173 | 1,014 | 0,871 | 0,911 | 1,036 | 0,924 | 0,978 | 1,05 | 0,05 | 4,8E-01 |
| Q9JIA1 | Leucine-rich glioma-inactivated protein 1 OS=Mus musculus OX=10 | 13 | 7 | 9 | 7 | 557 | 63,6 | 8,02 | 17,53 | 7 | 0,9 | 0,901 | 0,869 | 1,05 | 1,093 | 0,87 | 0,977 | 0,963 | 0,95 | -0,05 | 4,8E-01 |
| Q9JMH6 | Thioredoxin reductase 1, cytoplasmic OS=Mus musculus OX=10090 | 9 | 6 | 7 | 6 | 613 | 67 | 7,44 | 10,88 | 6 | 1,145 | 0,922 | 1,046 | 0,972 | 0,897 | 1,029 | 0,991 | 1 | 1,04 | 0,04 | 4,8E-01 |
| Q9WV85 | Nucleoside diphosphate kinase 3 OS=Mus musculus OX=10090 GN= | 10 | 1 | 1 | 1 | 169 | 19,1 | 6,68 | 3,93 | 1 | 1,173 | 1,183 | 0,938 | 0,826 | 0,82 | 1,173 | 0,947 | 0,817 | 1,10 | 0,10 | 4,8E-01 |
| Q91VF2 | Histamine N-methyltransferase OS=Mus musculus OX=10090 GN=H | 3 | 1 | 2 | 1 | 295 | 33,6 | 5,06 | 4,36 | 1 | 0,868 | 0,965 | 0,995 | 1,065 | 1,039 | 1,007 | 1,033 | 0,95 | 0,97 | -0,03 | 4,8E-01 |
| P15508 | Spectrin beta chain, erythrocytic OS=Mus musculus OX=10090 GN= | 13 | 23 | 28 | 17 | 2128 | 245,1 | 5,33 | 58,58 | 23 | 1,367 | 1,198 | 1,066 | 0,984 | 1,005 | 1,2 | 1,101 | 1,028 | 1,06 | 0,06 | 4,9E-01 |
| Q3UHK6 | Teneurin-4 OS=Mus musculus OX=10090 GN=Tenm4 PE=1 SV=2 | 2 | 4 | 4 | 2 | 2771 | 308,2 | 6,57 | 9,91 | 4 | 1,081 | 1,024 | 1,077 | 1,033 | 0,969 | 0,949 | 1,489 | 1,184 | 0,92 | -0,08 | 4,9E-01 |
| Q8C561 | G-protein coupled receptor-associated protein LMBRD2 OS=Mus m | 2 | 1 | 1 | 1 | 694 | 81 | 7,34 | 3,18 | 1 | 0,646 | 0,778 | 0,726 | 1,19 | 0,667 | 1,031 | 0,981 | 1,128 | 0,88 | -0,12 | 4,9E-01 |
| Q922R1 | UPF0183 protein C16orf70 homolog OS=Mus musculus OX=10090 I | 5 | 1 | 1 | 1 | 422 | 47,4 | 7,74 | 3,65 | 1 | 1,201 | 1,359 | 1,621 | 1,315 | 1,241 | 1,625 | 1,025 | 1,136 | 1,09 | 0,09 | 4,9E-01 |
| Q8C0M9 | Isoaspartyl peptidase/L-asparaginase OS=Mus musculus OX=10090 | 39 | 10 | 16 | 10 | 326 | 33,9 | 7,65 | 32,97 | 10 | 0,913 | 0,96 | 1,08 | 1,053 | 1,042 | 1,032 | 1,04 | 1,01 | 0,97 | -0,03 | 4,9E-01 |
| P55012 | Solute carrier family 12 member 2 OS=Mus musculus OX=10090 GN | 8 | 8 | 9 | 8 | 1205 | 131 | 7,33 | 10,2 | 8 | 0,982 | 0,885 | 1,07 | 1,138 | 1 | 1,035 | 0,891 | 0,963 | 1,05 | 0,05 | 4,9E-01 |
| Q8C3X4 | Translation factor Guf1, mitochondrial OS=Mus musculus OX=1009 | 3 | 2 | 2 | 2 | 651 | 72,4 | 8,76 | 4,43 | 2 | 1,232 | 1,36 | 0,835 | 1,088 | 0,795 | 1,11 | 1,126 | 1,079 | 1,10 | 0,10 | 4,9E-01 |
| Q99K48 | Non-POU domain-containing octamer-binding protein OS=Mus mu: | 4 | 2 | 2 | 2 | 473 | 54,5 | 8,95 | 3,24 | 2 | 0,987 | 1,02 | 1,111 | 0,767 | 1,249 | 1,291 | 0,784 | 0,975 | 0,90 | -0,10 | 4,9E-01 |
| Q60676 | Serine/threonine-protein phosphatase 5 OS=Mus musculus OX=100 | 21 | 10 | 16 | 10 | 499 | 56,8 | 6,2 | 33,45 | 10 | 0,94 | 0,932 | 0,921 | 1,012 | 1,041 | 0,877 | 0,987 | 1,025 | 0,97 | -0,03 | 4,9E-01 |
| Q9Z2K1 | Keratin, type I cytoskeletal 16 OS=Mus musculus OX=10090 GN=Krt | 14 | 6 | 17 | 2 | 469 | 51,6 | 5,2 | 33,2 | 6 | 0,891 | 1,282 | 0,896 | 0,886 | 1,939 | 1,298 | 0,735 | 0,842 | 0,82 | -0,18 | 4,9E-01 |
| Q9CQF9 | Prenylcysteine oxidase OS=Mus musculus OX=10090 GN=Pcyox1 PE | 8 | 3 | 4 | 3 | 505 | 56,5 | 6,92 | 6,4 | 3 | 1,254 | 1,225 | 1,4 | 1,257 | 1,117 | 1,463 | 1,518 | 1,326 | 0,95 | -0,05 | 4,9E-01 |
| Q8K2C9 | Very-long-chain (3R)-3-hydroxyacyl-CoA dehydratase 3 OS=Mus mu | 2 | 1 | 1 | 1 | 362 | 43,1 | 9,13 | 2,15 | 1 | 1,093 | 1,078 | 1,165 | 1,068 | 0,971 | 0,847 | 1,149 | 1,192 | 1,06 | 0,06 | 4,9E-01 |
| Q9JME7 | Trafficking protein particle complex subunit 2-like protein OS=Mus | 22 | 3 | 3 | 3 | 139 | 16 | 6,77 | 4,55 | 3 | 1,222 | 1,044 | 0,997 | 1,055 | 1,017 | 1,281 | 1,023 | 1,253 | 0,94 | -0,06 | 4,9E-01 |
| Q9D8B7 | Junctional adhesion molecule C OS=Mus musculus OX=10090 GN=J | 9 | 3 | 3 | 3 | 310 | 34,8 | 7,03 | 3,67 | 3 | 0,827 | 1,083 | 0,974 | 0,933 | 0,963 | 0,922 | 0,791 | 0,947 | 1,05 | 0,05 | 4,9E-01 |
| Q8BG05 | Heterogeneous nuclear ribonucleoprotein A3 OS=Mus musculus O) | 13 | 5 | 6 | 5 | 379 | 39,6 | 9,01 | 13,59 | 5 | 1,397 | 1,353 | 1,08 | 1,072 | 1,201 | 1,32 | 1,028 | 1,02 | 1,07 | 0,07 | 4,9E-01 |
| Q99KN9 | Clathrin interactor 1 OS=Mus musculus OX=10090 GN=Clint1 PE=1 | 10 | 6 | 6 | 6 | 631 | 68,5 | 6,25 | 7,41 | 6 | 1,097 | 1,08 | 1,019 | 0,999 | 0,954 | 1,188 | 1,049 | 1,185 | 0,96 | -0,04 | 4,9E-01 |
| Q8C0Z1 | Protein FAM234A OS=Mus musculus OX=10090 GN=Fam234a PE=1 | 3 | 1 | 1 | 1 | 555 | 60,5 | 5,68 | 0 | 1 | 1,608 | 0,859 | 2,037 | 1,023 | 1,375 | 0,801 | 1,312 | 1,156 | 1,19 | 0,19 | 4,9E-01 |
| Q8VCY8 | Phospholipid phosphatase-related protein type 2 OS=Mus musculu | 6 | 1 | 1 | 1 | 343 | 36,9 | 9,5 | 4,65 | 1 | 1,688 | 1,549 | 1,328 | 1,204 | 1,119 | 1,473 | 1,299 | 1,473 | 1,08 | 0,08 | 4,9E-01 |
| Q76MZ3 | Serine/threonine-protein phosphatase 2A 65 kDa regulatory subun | 34 | 19 | 33 | 19 | 589 | 65,3 | 5,11 | 69,84 | 19 | 0,999 | 1,064 | 0,98 | 1,013 | 0,972 | 0,992 | 0,997 | 1,031 | 1,02 | 0,02 | 4,9E-01 |
| Q9WUB4 | Dynactin subunit 6 OS=Mus musculus OX=10090 GN=Dctn6 PE=1 S\ | 9 | 1 | 2 | 1 | 190 | 20,7 | 6,67 | 2,98 | 1 | 0,642 | 1,101 | 0,866 | 1,6 | 1,186 | 1,058 | 1,44 | 1,173 | 0,87 | -0,13 | 4,9E-01 |
| Q99KH8 | Serine/threonine-protein kinase 24 OS=Mus musculus OX=10090 G | 8 | 3 | 3 | 1 | 431 | 47,9 | 5,43 | 10,05 | 3 | 1,595 | 1,487 | 1,197 | 1,197 | 0,671 | 1,555 | 1,078 | 1,497 | 1,14 | 0,14 | 4,9E-01 |
| Q91W52 | Transmembrane protein 19 OS=Mus musculus OX=10090 GN=Tmei | 4 | 1 | 1 | 1 | 336 | 36,3 | 7,72 | 3,11 | 1 | 1,645 | 1,285 | 1,369 | 1,407 | 1,651 | 1,652 | 1,481 | 1,271 | 0,94 | -0,06 | 4,9E-01 |
| Q9JMC3 | DnaJ homolog subfamily A member 4 OS=Mus musculus OX=10090 | 9 | 4 | 4 | 4 | 397 | 44,9 | 7,58 | 2,85 | 4 | 0,696 | 0,749 | 1,099 | 1,069 | 1,127 | 0,965 | 0,882 | 0,982 | 0,91 | -0,09 | 4,9E-01 |
| Q91VD9 | NADH-ubiquinone oxidoreductase 75 kDa subunit, mitochondrial O | 42 | 29 | 54 | 29 | 727 | 79,7 | 5,72 | 110,65 | 29 | 1,053 | 1,033 | 1,032 | 1,016 | 1,037 | 1,103 | 1,009 | 1,047 | 0,99 | -0,01 | 4,9E-01 |
| Q9CPS6 | Histidine triad nucleotide-binding protein 3 OS=Mus musculus OX= | 24 | 3 | 3 | 3 | 165 | 18,8 | 6,33 | 10,91 | 3 | 1,28 | 1,181 | 1,276 | 0,967 | 1,007 | 1,178 | 1,108 | 1,167 | 1,05 | 0,05 | 4,9E-01 |
| Q91WS0 | CDGSH iron-sulfur domain-containing protein 1 OS=Mus musculus ( | 48 | 5 | 10 | 5 | 108 | 12,1 | 9,06 | 34,84 | 5 | 1,237 | 1,251 | 1,082 | 0,991 | 0,995 | 1,253 | 1,016 | 1,043 | 1,06 | 0,06 | 4,9E-01 |
| Q9D1T0 | Leucine-rich repeat and immunoglobulin-like domain-containing nc | 11 | 5 | 5 | 5 | 614 | 69,1 | 8,31 | 15,72 | 5 | 1,632 | 1,144 | 1,268 | 0,971 | 1,062 | 1,276 | 1,134 | 1,112 | 1,09 | 0,09 | 4,9E-01 |
| Q80ZJ1 | Ras-related protein Rap-2a OS=Mus musculus OX=10090 GN=Rap2i | 33 | 4 | 6 | 2 | 183 | 20,6 | 4,82 | 10,81 | 4 | 1,639 | 1,199 | 1,2 | 0,985 | 1,009 | 1,421 | 1 | 1,098 | 1,11 | 0,11 | 4,9E-01 |
| Q9ERI6 | Retinol dehydrogenase 14 OS=Mus musculus OX=10090 GN=Rdh14 | 18 | 5 | 5 | 5 | 334 | 36,3 | 8,18 | 13,82 | 5 | 0,895 | 1,036 | 0,907 | 0,939 | 0,932 | 0,944 | 1,087 | 0,955 | 0,96 | -0,04 | 4,9E-01 |
| Q99LR1 | Lysophosphatidylserine lipase ABHD12 OS=Mus musculus OX=1009 | 6 | 2 | 2 | 2 | 398 | 45,2 | 8,72 | 2,87 | 2 | 1,092 | 1,085 | 1,028 | 0,9 | 1,002 | 1,048 | 1,201 | 1,037 | 0,96 | -0,04 | 4,9E-01 |
| E9PV86 | Multiple C2 and transmembrane domain-containing protein 1 OS=I | 2 | 2 | 2 | 2 | 951 | 106,7 | 8,37 | 1,84 | 2 | 0,968 | 1,251 | 1,194 | 1,174 | 1,074 | 1,031 | 1,128 | 1,156 | 1,05 | 0,05 | 4,9E-01 |
| O35593 | 26S proteasome non-ATPase regulatory subunit 14 OS=Mus muscu | 6 | 2 | 5 | 2 | 310 | 34,6 | 6,52 | 12,39 | 2 | 1,106 | 1,142 | 1,155 | 0,994 | 0,978 | 1,133 | 1,056 | 1,087 | 1,03 | 0,03 | 4,9E-01 |
| Q922Q1 | Mitochondrial amidoxime reducing component 2 OS=Mus musculu | 9 | 3 | 5 | 3 | 338 | 38,2 | 8,68 | 7,66 | 3 | 0,751 | 0,899 | 1,125 | 0,925 | 1,008 | 0,924 | 0,989 | 1,011 | 0,94 | -0,06 | 4,9E-01 |
| P56391 | Cytochrome c oxidase subunit 6B1 OS=Mus musculus OX=10090 GI | 71 | 7 | 9 | 7 | 86 | 10,1 | 8,72 | 26,51 | 7 | 0,914 | 0,973 | 0,978 | 0,945 | 1,035 | 0,949 | 0,937 | 0,966 | 0,98 | -0,02 | 4,9E-01 |
| P21619 | Lamin-B2 OS=Mus musculus OX=10090 GN=Lmnb2 PE=1 SV=2 | 13 | 8 | 8 | 6 | 596 | 67,3 | 5,5 | 14,11 | 8 | 1,318 | 1,31 | 1,204 | 0,846 | 1,269 | 1,656 | 1,029 | 1,225 | 0,90 | -0,10 | 4,9E-01 |
| Q6P8M1 | Putative deoxyribonuclease TATDN1 OS=Mus musculus OX=10090 ( | 4 | 1 | 1 | 1 | 295 | 33,3 | 6,18 | 0 | 1 | 1,296 | 0,987 | 1,102 | 0,782 | 0,986 | 1,416 | 1,023 | 1,165 | 0,91 | -0,09 | 4,9E-01 |
| A2AGT5 | Cytoskeleton-associated protein 5 OS=Mus musculus OX=10090 GN | 12 | 21 | 24 | 21 | 2032 | 225,5 | 7,96 | 36,2 | 21 | 0,962 | 1,065 | 0,993 | 1,054 | 0,953 | 1,085 | 1,109 | 1,05 | 0,97 | -0,03 | 4,9E-01 |
| Q60631 | Growth factor receptor-bound protein 2 OS=Mus musculus OX=100 | 29 | 6 | 9 | 6 | 217 | 25,2 | 6,32 | 17,5 | 6 | 1,089 | 1,031 | 0,986 | 0,924 | 0,984 | 1,015 | 0,971 | 0,951 | 1,03 | 0,03 | 4,9E-01 |
| Q62376 | U1 small nuclear ribonucleoprotein 70 kDa OS=Mus musculus OX=1 | 2 | 1 | 1 | 1 | 448 | 52 | 9,94 | 0 | 1 | 1,256 | 1,107 | 1,072 | 0,789 | 1,074 | 1,1 | 0,794 | 0,902 | 1,09 | 0,09 | 4,9E-01 |
| Q99M87 | DnaJ homolog subfamily A member 3, mitochondrial OS=Mus musc | 14 | 5 | 8 | 5 | 480 | 52,4 | 9,22 | 22,19 | 5 | 1,328 | 1,165 | 1,02 | 0,991 | 0,86 | 1,278 | 0,891 | 1,111 | 1,09 | 0,09 | 4,9E-01 |
| Q9DA03 | Complex III assembly factor LYRM7 OS=Mus musculus OX=10090 G | 18 | 1 | 1 | 1 | 104 | 12 | 9,45 | 6,3 | 1 | 1,35 | 0,893 | 1,067 | 0,847 | 0,807 | 1,261 | 0,826 | 0,797 | 1,13 | 0,13 | 4,9E-01 |
| Q8R4F1 | Netrin-G2 OS=Mus musculus OX=10090 GN=Ntng2 PE=1 SV=2 | 6 | 4 | 4 | 4 | 589 | 66,1 | 6,13 | 1,9 | 4 | 1,184 | 0,844 | 0,88 | 0,927 | 0,985 | 1,11 | 1,023 | 0,96 | 0,94 | -0,06 | 5,0E-01 |
| Q9D8Z2 | TP53-regulated inhibitor of apoptosis 1 OS=Mus musculus OX=1009 | 37 | 3 | 3 | 3 | 76 | 8,8 | 5,48 | 4,28 | 3 | 1,444 | 1,224 | 1,387 | 1,172 | 1,04 | 1,57 | 0,975 | 1,213 | 1,09 | 0,09 | 5,0E-01 |
| P28661 | Septin-4 OS=Mus musculus OX=10090 GN=Septin4 PE=1 SV=1 | 28 | 13 | 20 | 11 | 478 | 54,9 | 5,87 | 21,43 | 13 | 0,851 | 0,938 | 0,954 | 1,056 | 0,993 | 1,165 | 0,862 | 0,996 | 0,95 | -0,05 | 5,0E-01 |
| Q06890 | Clusterin OS=Mus musculus OX=10090 GN=Clu PE=1 SV=1 | 19 | 8 | 8 | 8 | 448 | 51,6 | 5,67 | 19,04 | 8 | 1,412 | 1,104 | 1,19 | 0,948 | 0,994 | 1,214 | 1,086 | 1,049 | 1,07 | 0,07 | 5,0E-01 |
| Q8R1B4 | Eukaryotic translation initiation factor 3 subunit C OS=Mus musculu | 5 | 4 | 5 | 4 | 911 | 105,5 | 5,78 | 8,88 | 4 | 1,392 | 1,089 | 1,05 | 0,991 | 1,081 | 1,026 | 0,979 | 1,155 | 1,07 | 0,07 | 5,0E-01 |
| Q5SXY1 | Cytospin-B OS=Mus musculus OX=10090 GN=Specc1 PE=1 SV=2 | 3 | 3 | 3 | 1 | 1067 | 118 | 6,64 | 6,24 | 3 | 1,46 | 0,926 | 0,883 | 1,224 | 1,244 | 0,556 | 1,182 | 0,913 | 1,15 | 0,15 | 5,0E-01 |
| Q9JLB0-2 | Isoform Alpha of MAGUK p55 subfamily member 6 OS=Mus muscu | 26 | 13 | 15 | 12 | 539 | 60,9 | 6,18 | 35,86 | 13 | 1,031 | 1,042 | 1,046 | 0,976 | 0,952 | 1,07 | 0,974 | 1,011 | 1,02 | 0,02 | 5,0E-01 |
| Q9WU78 | Programmed cell death 6-interacting protein OS=Mus musculus OX | 21 | 18 | 19 | 18 | 869 | 96 | 6,52 | 50,28 | 18 | 0,907 | 0,873 | 0,907 | 0,986 | 0,996 | 0,857 | 0,988 | 0,947 | 0,97 | -0,03 | 5,0E-01 |
| Q99PT1 | Rho GDP-dissociation inhibitor 1 OS=Mus musculus OX=10090 GN= | 43 | 9 | 13 | 9 | 204 | 23,4 | 5,2 | 28,92 | 9 | 0,819 | 0,88 | 1,051 | 1,061 | 1,009 | 0,923 | 1,074 | 1,002 | 0,95 | -0,05 | 5,0E-01 |
| Q3UHD9 | Arf-GAP with GTPase, ANK repeat and PH domain-containing prote | 15 | 16 | 17 | 16 | 1186 | 124,4 | 9,89 | 30,98 | 16 | 1,128 | 1,042 | 1,005 | 1,055 | 1,051 | 1,038 | 1,085 | 0,939 | 1,03 | 0,03 | 5,0E-01 |
| Q61285 | ATP-binding cassette sub-family D member 2 OS=Mus musculus OX | 2 | 1 | 1 | 1 | 741 | 83,4 | 9,09 | 2,46 | 1 | 1,331 | 1,06 | 1,064 | 0,967 | 0,852 | 1,356 | 0,946 | 0,855 | 1,10 | 0,10 | 5,0E-01 |
| A2ASZ8-5 | Isoform 5 of Calcium-binding mitochondrial carrier protein SCaMC- | 12 | 7 | 7 | 6 | 501 | 56 | 8,87 | 11,27 | 7 | 1,045 | 1,346 | 1,03 | 0,878 | 0,867 | 1,165 | 0,957 | 0,975 | 1,08 | 0,08 | 5,0E-01 |
| Q8CAA7 | Glucose 1,6-bisphosphate synthase OS=Mus musculus OX=10090 G | 14 | 8 | 11 | 8 | 621 | 70,2 | 6,49 | 25,32 | 8 | 1,242 | 1,085 | 1,072 | 0,984 | 0,956 | 1,134 | 0,967 | 1,119 | 1,05 | 0,05 | 5,0E-01 |
| Q924M7 | Mannose-6-phosphate isomerase OS=Mus musculus OX=10090 GN | 6 | 2 | 4 | 2 | 423 | 46,5 | 5,95 | 5,67 | 2 | 1,378 | 1,145 | 0,993 | 1,143 | 0,887 | 1,449 | 1,471 | 1,304 | 0,91 | -0,09 | 5,0E-01 |
| Q91WA3 | Histone deacetylase 11 OS=Mus musculus OX=10090 GN=Hdac11 P | 6 | 2 | 2 | 2 | 347 | 39,1 | 7,14 | 0 | 2 | 0,76 | 1,255 | 0,816 | 1,181 | 0,996 | 0,93 | 1,275 | 1,252 | 0,90 | -0,10 | 5,0E-01 |

| | | | | | | | | | | | | | | | | | | | | | |
|---|---|---|---|---|---|---|---|---|---|---|---|---|---|---|---|---|---|---|---|---|---|
| Q91Z61 | GTP-binding protein Di-Ras1 OS=Mus musculus OX=10090 GN=Dira | 25 | 5 | 7 | 4 | 198 | 22,3 | 8,9 | 14,9 | 5 | 0,803 | 1,111 | 0,867 | 0,968 | 0,929 | 1,035 | 0,977 | 1,012 | 0,95 | -0,05 | 5,0E-01 |
| Q99M51 | Cytoplasmic protein NCK1 OS=Mus musculus OX=10090 GN=Nck1 P | 13 | 3 | 4 | 3 | 377 | 42,9 | 6,47 | 2,53 | 3 | 0,93 | 1,158 | 0,915 | 0,821 | 0,859 | 1,345 | 1,082 | 0,913 | 0,91 | -0,09 | 5,0E-01 |
| Q9D1K7 | UPF0687 protein C20orf27 homolog OS=Mus musculus OX=10090 | 11 | 1 | 1 | 1 | 174 | 19,5 | 6,68 | 0 | 1 | 1,845 | 1,081 | 1,084 | 1,014 | 0,954 | 1,494 | 0,998 | 0,886 | 1,16 | 0,16 | 5,0E-01 |
| P11404 | Fatty acid-binding protein, heart OS=Mus musculus OX=10090 GN= | 20 | 3 | 4 | 3 | 133 | 14,8 | 6,57 | 11,09 | 3 | 0,776 | 0,935 | 0,947 | 1,017 | 0,904 | 0,781 | 0,916 | 0,902 | 1,05 | 0,05 | 5,0E-01 |
| O08547 | Vesicle-trafficking protein SEC22b OS=Mus musculus OX=10090 GN | 16 | 3 | 3 | 3 | 215 | 24,7 | 8,51 | 10,7 | 3 | 0,881 | 0,988 | 1,005 | 1,031 | 0,967 | 0,855 | 1,017 | 0,93 | 1,04 | 0,04 | 5,0E-01 |
| Q80UG5 | Septin-9 OS=Mus musculus OX=10090 GN=Septin9 PE=1 SV=1 | 19 | 11 | 16 | 11 | 583 | 65,5 | 8,9 | 27,12 | 11 | 0,859 | 0,854 | 0,851 | 0,932 | 0,843 | 0,867 | 0,912 | 0,973 | 0,97 | -0,03 | 5,0E-01 |
| P63073 | Eukaryotic translation initiation factor 4E OS=Mus musculus OX=10 | 10 | 2 | 2 | 2 | 217 | 25 | 6,15 | 5,92 | 2 | 1,343 | 1,211 | 1,028 | 1,07 | 1,234 | 1,126 | 0,968 | 1,064 | 1,06 | 0,06 | 5,0E-01 |
| O54734 | Dolichyl-diphosphooligosaccharide--protein glycosyltransferase 48 | 12 | 5 | 5 | 5 | 441 | 49 | 5,83 | 10,23 | 5 | 1,171 | 0,962 | 1,301 | 0,995 | 0,99 | 1,16 | 0,929 | 1,081 | 1,06 | 0,06 | 5,0E-01 |
| Q8BFQ8 | Glutamine amidotransferase-like class 1 domain-containing protein | 19 | 3 | 3 | 3 | 220 | 23,3 | 7,05 | 4,7 | 3 | 1,001 | 1,339 | 0,765 | 1,002 | 0,895 | 1,04 | 0,889 | 0,931 | 1,09 | 0,09 | 5,0E-01 |
| Q7TPS5 | C2 domain-containing protein 5 OS=Mus musculus OX=10090 GN=( | 5 | 4 | 5 | 4 | 1016 | 111,6 | 5,78 | 5,1 | 4 | 0,764 | 0,804 | 1,006 | 0,973 | 0,973 | 0,932 | 0,957 | 0,869 | 0,95 | -0,05 | 5,0E-01 |
| Q9DCZ1 | GMP reductase 1 OS=Mus musculus OX=10090 GN=Gmpr PE=1 SV= | 12 | 3 | 3 | 3 | 345 | 37,5 | 7,09 | 4,79 | 3 | 1,177 | 0,747 | 1,002 | 0,782 | 1 | 0,763 | 0,788 | 0,832 | 1,10 | 0,10 | 5,0E-01 |
| P12960 | Contactin-1 OS=Mus musculus OX=10090 GN=Cntn1 PE=1 SV=1 | 25 | 24 | 52 | 24 | 1020 | 113,3 | 6,16 | 108,35 | 24 | 1,103 | 0,833 | 1,03 | 0,956 | 1,098 | 0,977 | 1,069 | 0,967 | 0,95 | -0,05 | 5,0E-01 |
| P67984 | 60S ribosomal protein L22 OS=Mus musculus OX=10090 GN=Rpl22 | 10 | 1 | 2 | 1 | 128 | 14,8 | 9,19 | 6,45 | 1 | 1,103 | 1,125 | 1,111 | 0,953 | 1,103 | 0,903 | 1,016 | 1,095 | 1,04 | 0,04 | 5,0E-01 |
| Q9CY50 | Translocon-associated protein subunit alpha OS=Mus musculus OX= | 7 | 2 | 3 | 2 | 286 | 32 | 4,45 | 0 | 2 | 0,888 | 0,854 | 0,941 | 0,8 | 0,969 | 1,034 | 0,761 | 0,905 | 0,95 | -0,05 | 5,0E-01 |
| O88307 | Sortilin-related receptor OS=Mus musculus OX=10090 GN=Sorl1 PE | 2 | 4 | 4 | 4 | 2215 | 246,9 | 5,54 | 4,22 | 4 | 1,157 | 1,288 | 1,16 | 0,944 | 0,947 | 1,264 | 1,057 | 0,998 | 1,07 | 0,07 | 5,0E-01 |
| Q9DB20 | ATP synthase subunit O, mitochondrial OS=Mus musculus OX=1009 | 36 | 8 | 20 | 8 | 213 | 23,3 | 9,99 | 33,54 | 8 | 1,005 | 0,985 | 0,954 | 0,91 | 1,026 | 0,988 | 0,945 | 0,971 | 0,98 | -0,02 | 5,0E-01 |
| Q920R0 | Alsin OS=Mus musculus OX=10090 GN=Als2 PE=1 SV=3 | 2 | 4 | 4 | 4 | 1651 | 182,5 | 6,3 | 5,15 | 4 | 1,014 | 0,97 | 0,956 | 1,143 | 0,931 | 1,097 | 0,956 | 0,934 | 1,04 | 0,04 | 5,0E-01 |
| Q8BYK5 | Phosphatase and actin regulator 3 OS=Mus musculus OX=10090 GN | 2 | 1 | 1 | 1 | 558 | 62,6 | 8,31 | 2,08 | 1 | 1,297 | 1,167 | 0,845 | 1,416 | 1,12 | 0,762 | 1,164 | 1,222 | 1,11 | 0,11 | 5,0E-01 |
| Q8BGZ4 | Cell division cycle protein 23 homolog OS=Mus musculus OX=10090 | 3 | 1 | 1 | 1 | 597 | 68,5 | 7,18 | 3,97 | 1 | 1,103 | 1,035 | 0,991 | 0,964 | 1,017 | 1,145 | 0,825 | 0,887 | 1,06 | 0,06 | 5,1E-01 |
| Q80U28 | MAP kinase-activating death domain protein OS=Mus musculus OX | 14 | 17 | 20 | 17 | 1577 | 175,1 | 6,04 | 44,84 | 17 | 1,118 | 1,136 | 1,002 | 1,041 | 0,974 | 1,087 | 1,054 | 1,068 | 1,03 | 0,03 | 5,1E-01 |
| P53994 | Ras-related protein Rab-2A OS=Mus musculus OX=10090 GN=Rab2 | 40 | 7 | 13 | 2 | 212 | 23,5 | 6,54 | 41,66 | 7 | 1,051 | 0,985 | 0,883 | 0,96 | 0,895 | 1,23 | 1,035 | 0,949 | 0,94 | -0,06 | 5,1E-01 |
| Q8BLQ9 | Cell adhesion molecule 2 OS=Mus musculus OX=10090 GN=Cadm2 | 23 | 9 | 20 | 9 | 435 | 47,5 | 5,41 | 50,75 | 9 | 0,955 | 0,886 | 0,966 | 1,018 | 0,931 | 0,842 | 0,931 | 1,001 | 1,03 | 0,03 | 5,1E-01 |
| P97300 | Neuroplastin OS=Mus musculus OX=10090 GN=Nptn PE=1 SV=3 | 30 | 13 | 37 | 5 | 397 | 44,3 | 7,74 | 75,48 | 13 | 1,126 | 0,78 | 1,136 | 0,901 | 1,168 | 0,951 | 1,123 | 0,989 | 0,93 | -0,07 | 5,1E-01 |
| Q8K298 | Anillin OS=Mus musculus OX=10090 GN=Anln PE=1 SV=2 | 3 | 3 | 3 | 3 | 1121 | 122,7 | 6,98 | 2,21 | 3 | 1,207 | 1,276 | 0,946 | 0,961 | 0,979 | 1,228 | 0,853 | 1,005 | 1,08 | 0,08 | 5,1E-01 |
| P60334 | Cysteine dioxygenase type 1 OS=Mus musculus OX=10090 GN=Cdo | 4 | 1 | 1 | 1 | 200 | 23 | 6,46 | 1,73 | 1 | 0,715 | 1,069 | 1,033 | 0,899 | 0,909 | 0,724 | 0,814 | 0,99 | 1,08 | 0,08 | 5,1E-01 |
| Q8R127 | Saccharopine dehydrogenase-like oxidoreductase OS=Mus musculu | 17 | 8 | 10 | 8 | 429 | 47,1 | 8,6 | 23,23 | 8 | 1,186 | 1,041 | 1,111 | 0,976 | 0,923 | 1,131 | 1,01 | 1,071 | 1,04 | 0,04 | 5,1E-01 |
| P19253 | 60S ribosomal protein L13a OS=Mus musculus OX=10090 GN=Rpl13 | 15 | 3 | 4 | 3 | 203 | 23,4 | 11,02 | 10,62 | 3 | 0,91 | 1,212 | 1,092 | 1,082 | 1,014 | 1,273 | 1,065 | 1,185 | 0,95 | -0,05 | 5,1E-01 |
| Q9R1V7 | Disintegrin and metalloproteinase domain-containing protein 23 OS | 15 | 10 | 11 | 10 | 829 | 91,5 | 7,68 | 12,04 | 10 | 1,158 | 1,099 | 1,132 | 0,936 | 0,883 | 1,145 | 0,996 | 1,087 | 1,05 | 0,05 | 5,1E-01 |
| Q3UHH2 | Solute carrier family 22 member 23 OS=Mus musculus OX=10090 G | 7 | 4 | 4 | 4 | 689 | 74,3 | 7,83 | 5,85 | 4 | 0,971 | 0,994 | 1,045 | 1,024 | 0,864 | 0,94 | 1,101 | 0,982 | 1,04 | 0,04 | 5,1E-01 |
| O70318 | Band 4.1-like protein 2 OS=Mus musculus OX=10090 GN=Epb41l2 P | 21 | 17 | 20 | 15 | 988 | 109,9 | 5,43 | 41,5 | 17 | 0,987 | 0,876 | 1,044 | 0,99 | 1,009 | 0,963 | 1,007 | 1,024 | 0,97 | -0,03 | 5,1E-01 |
| A6H5Z3 | Exocyst complex component 6B OS=Mus musculus OX=10090 GN= | 2 | 2 | 3 | 2 | 810 | 94,1 | 6,39 | 2,38 | 2 | 0,911 | 1,123 | 0,808 | 0,991 | 1,037 | 0,935 | 1,064 | 1 | 0,95 | -0,05 | 5,1E-01 |
| Q8C1Y8 | Vacuolar fusion protein CCZ1 homolog OS=Mus musculus OX=1009 | 8 | 3 | 3 | 3 | 480 | 55,5 | 6,02 | 3,07 | 3 | 1,204 | 1,708 | 0,794 | 1,398 | 1,255 | 1,061 | 1,626 | 1,926 | 0,87 | -0,13 | 5,1E-01 |
| Q8C7H1 | Methylmalonic aciduria type A homolog, mitochondrial OS=Mus mu | 5 | 2 | 3 | 2 | 415 | 45,9 | 9,32 | 5,73 | 2 | 1,354 | 1,12 | 0,949 | 1,025 | 0,946 | 1,251 | 0,916 | 1,008 | 1,08 | 0,08 | 5,1E-01 |
| O35136 | Neural cell adhesion molecule 2 OS=Mus musculus OX=10090 GN= | 26 | 22 | 30 | 22 | 837 | 93,1 | 6,07 | 52,5 | 22 | 0,949 | 0,91 | 1,002 | 1,007 | 1,024 | 0,924 | 1,015 | 0,996 | 0,98 | -0,02 | 5,1E-01 |
| P07901 | Heat shock protein HSP 90-alpha OS=Mus musculus OX=10090 GN= | 39 | 36 | 67 | 21 | 733 | 84,7 | 5,01 | 159,79 | 36 | 0,984 | 0,972 | 0,98 | 1,075 | 1,002 | 0,898 | 1,009 | 1,001 | 1,03 | 0,03 | 5,1E-01 |
| Q80TE7 | Leucine-rich repeat-containing protein 7 OS=Mus musculus OX=100 | 15 | 18 | 19 | 18 | 1490 | 166,8 | 6,61 | 38,81 | 18 | 1,343 | 1,12 | 1,071 | 0,978 | 0,93 | 1,235 | 1,008 | 1,056 | 1,07 | 0,07 | 5,1E-01 |
| Q9DBG6 | Dolichyl-diphosphooligosaccharide--protein glycosyltransferase sub | 15 | 8 | 8 | 8 | 631 | 69 | 5,81 | 10,66 | 8 | 1,139 | 1,02 | 1,143 | 1,056 | 0,981 | 1,178 | 0,928 | 1,092 | 1,04 | 0,04 | 5,1E-01 |
| Q62443 | Neuronal pentraxin-1 OS=Mus musculus OX=10090 GN=Nptx1 PE= | 3 | 2 | 2 | 2 | 432 | 47,1 | 6,44 | 4,79 | 2 | 0,882 | 0,905 | 0,969 | 1,03 | 0,996 | 0,858 | 0,95 | 0,843 | 1,04 | 0,04 | 5,1E-01 |
| Q9D1C8 | Vacuolar protein sorting-associated protein 28 homolog OS=Mus m | 19 | 4 | 4 | 4 | 221 | 25,4 | 5,54 | 9 | 4 | 0,876 | 0,905 | 0,981 | 0,891 | 0,901 | 1,117 | 0,863 | 0,942 | 0,96 | -0,04 | 5,1E-01 |
| Q9WUT3 | Ribosomal protein S6 kinase alpha-2 OS=Mus musculus OX=10090 | 9 | 6 | 6 | 2 | 733 | 83,1 | 8,5 | 10,5 | 6 | 1,111 | 0,722 | 0,699 | 0,753 | 0,764 | 0,885 | 1,081 | 0,883 | 0,91 | -0,09 | 5,1E-01 |
| Q3U5Q7 | UMP-CMP kinase 2, mitochondrial OS=Mus musculus OX=10090 GN | 13 | 5 | 7 | 5 | 447 | 50 | 7,27 | 13,84 | 5 | 1,087 | 1,163 | 1,074 | 0,954 | 1,003 | 1,253 | 1,019 | 1,223 | 0,95 | -0,05 | 5,1E-01 |
| Q5SP85 | Coiled-coil domain-containing protein 85A OS=Mus musculus OX=1 | 2 | 1 | 1 | 1 | 500 | 54,5 | 8 | 2,05 | 1 | 0,892 | 1,341 | 0,768 | 0,895 | 0,979 | 1,236 | 1,083 | 0,987 | 0,91 | -0,09 | 5,1E-01 |
| P61957 | Small ubiquitin-related modifier 2 OS=Mus musculus OX=10090 GN | 23 | 2 | 4 | 2 | 95 | 10,9 | 5,5 | 11,02 | 2 | 0,882 | 1,146 | 1,01 | 1,306 | 0,974 | 0,997 | 1,087 | 1,023 | 1,06 | 0,06 | 5,1E-01 |
| Q00493 | Carboxypeptidase E OS=Mus musculus OX=10090 GN=Cpe PE=1 SV | 27 | 11 | 14 | 11 | 476 | 53,2 | 5,19 | 33,16 | 11 | 0,975 | 0,967 | 1 | 1,013 | 0,932 | 1,007 | 1,006 | 0,948 | 1,02 | 0,02 | 5,1E-01 |
| Q6P6M7 | O-phosphoseryl-tRNA(Sec) selenium transferase OS=Mus musculus | 1 | 1 | 1 | 1 | 504 | 55,3 | 8,06 | 2,51 | 1 | 0,95 | 1,032 | 1 | 0,778 | 0,933 | 0,975 | 1,019 | 0,999 | 0,96 | -0,04 | 5,1E-01 |
| Q9Z1Q9 | Valine--tRNA ligase OS=Mus musculus OX=10090 GN=Vars PE=1 SV | 4 | 4 | 4 | 4 | 1263 | 140,1 | 7,77 | 2,09 | 4 | 0,891 | 1,045 | 1,017 | 1,12 | 0,99 | 0,67 | 1,055 | 1,066 | 1,08 | 0,08 | 5,1E-01 |
| Q9R0K7 | Plasma membrane calcium-transporting ATPase 2 OS=Mus musculu | 36 | 43 | 91 | 29 | 1198 | 132,5 | 5,96 | 252,13 | 43 | 1,092 | 1,062 | 0,974 | 0,96 | 0,985 | 0,999 | 0,998 | 1,014 | 1,02 | 0,02 | 5,1E-01 |
| P59108 | Copine-2 OS=Mus musculus OX=10090 GN=Cpne2 PE=1 SV=1 | 3 | 1 | 1 | 1 | 548 | 61 | 5,96 | 3,59 | 1 | 1,719 | 1,415 | 1,324 | 1,239 | 1,374 | 1,709 | 1,429 | 1,542 | 0,94 | -0,06 | 5,1E-01 |
| O88935 | Synapsin-1 OS=Mus musculus OX=10090 GN=Syn1 PE=1 SV=2 | 51 | 27 | 79 | 25 | 706 | 74,1 | 9,8 | 241,75 | 27 | 1,067 | 1,095 | 1,009 | 1,034 | 0,961 | 1,066 | 1,043 | 1,051 | 1,02 | 0,02 | 5,1E-01 |
| Q91VC9 | Growth hormone-inducible transmembrane protein OS=Mus muscu | 3 | 1 | 1 | 1 | 346 | 37,3 | 9,8 | 3,27 | 1 | 0,822 | 1,022 | 1,031 | 0,869 | 0,887 | 0,965 | 1,018 | 1,053 | 0,95 | -0,05 | 5,1E-01 |
| O88456 | Calpain small subunit 1 OS=Mus musculus OX=10090 GN=Capns1 P | 5 | 1 | 1 | 1 | 269 | 28,4 | 5,63 | 3,32 | 1 | 0,693 | 0,55 | 0,868 | 0,714 | 0,702 | 0,882 | 0,689 | 0,771 | 0,93 | -0,07 | 5,1E-01 |
| Q8CIQ7 | Dedicator of cytokinesis protein 3 OS=Mus musculus OX=10090 GN | 4 | 8 | 9 | 8 | 2027 | 232,8 | 6,99 | 12,4 | 8 | 0,953 | 0,951 | 0,922 | 0,812 | 0,855 | 1,028 | 0,999 | 0,902 | 0,96 | -0,04 | 5,1E-01 |
| Q8K4G5 | Actin-binding LIM protein 1 OS=Mus musculus OX=10090 GN=Ablin | 8 | 6 | 8 | 5 | 861 | 96,7 | 8,63 | 12,23 | 6 | 1,194 | 1,487 | 1,203 | 1,077 | 0,984 | 1,456 | 0,905 | 1,196 | 1,09 | 0,09 | 5,1E-01 |
| Q8BW75 | Amine oxidase [flavin-containing] B OS=Mus musculus OX=10090 G | 17 | 9 | 15 | 9 | 520 | 58,5 | 8,29 | 36,41 | 9 | 1,222 | 1,102 | 1,099 | 0,935 | 0,953 | 1,117 | 1,029 | 1,069 | 1,05 | 0,05 | 5,1E-01 |
| P15327 | Bisphosphoglycerate mutase OS=Mus musculus OX=10090 GN=Bpg | 4 | 1 | 1 | 1 | 259 | 30 | 7,06 | 0 | 1 | 1,055 | 1,225 | 0,522 | 1,068 | 1,004 | 0,54 | 0,766 | 1,028 | 1,16 | 0,16 | 5,1E-01 |
| Q9D125 | 28S ribosomal protein S25, mitochondrial OS=Mus musculus OX=10 | 9 | 2 | 2 | 2 | 171 | 19,9 | 9,23 | 4,49 | 2 | 1,395 | 1,289 | 1,063 | 0,886 | 1,08 | 1,186 | 1,074 | 0,95 | 1,08 | 0,08 | 5,1E-01 |
| Q3UHK1 | Proton myo-inositol cotransporter OS=Mus musculus OX=10090 GN | 5 | 3 | 3 | 3 | 637 | 69 | 6,86 | 0 | 3 | 1,655 | 1,252 | 0,913 | 1,645 | 1,235 | 1,679 | 0,981 | 0,871 | 1,15 | 0,15 | 5,2E-01 |
| O89084 | cAMP-specific 3',5'-cyclic phosphodiesterase 4A OS=Mus musculus | 7 | 4 | 4 | 3 | 844 | 93,5 | 5,26 | 12,14 | 4 | 1,038 | 0,758 | 0,979 | 0,89 | 0,958 | 0,939 | 0,958 | 0,98 | 0,96 | -0,04 | 5,2E-01 |
| P62852 | 40S ribosomal protein S25 OS=Mus musculus OX=10090 GN=Rps25 | 9 | 1 | 1 | 1 | 125 | 13,7 | 10,11 | 0 | 1 | 0,891 | 1,531 | 1,16 | 1,122 | 0,966 | 1,294 | 0,987 | 1,035 | 1,10 | 0,10 | 5,2E-01 |
| Q64442 | Sorbitol dehydrogenase OS=Mus musculus OX=10090 GN=Sord PE= | 14 | 5 | 5 | 5 | 357 | 38,2 | 7,02 | 13,32 | 5 | 0,914 | 1,038 | 1,078 | 1,09 | 0,956 | 1,05 | 1,087 | 1,21 | 0,96 | -0,04 | 5,2E-01 |
| Q8BIE6 | FERM domain-containing protein 4A OS=Mus musculus OX=10090 | 2 | 2 | 2 | 2 | 1020 | 113,8 | 8,92 | 4,59 | 2 | 1,465 | 0,955 | 1,001 | 0,783 | 0,897 | 0,914 | 0,958 | 1,025 | 1,11 | 0,11 | 5,2E-01 |
| Q9JJA9 | General receptor for phosphoinositides 1-associated scaffold prote | 2 | 1 | 1 | 1 | 392 | 42,3 | 9,1 | 1,91 | 1 | 1,057 | 0,971 | 1,279 | 0,882 | 0,846 | 1,102 | 0,91 | 1,045 | 1,07 | 0,07 | 5,2E-01 |

| | | | | | | | | | | | | | | | | | | | | | |
|---|---|---|---|---|---|---|---|---|---|---|---|---|---|---|---|---|---|---|---|---|---|
| P16283 | Anion exchange protein 3 OS=Mus musculus OX=10090 GN=Slc4a3 | 5 | 4 | 4 | 4 | 1227 | 135,3 | 6,51 | 12,75 | 4 | 1,398 | 1,164 | 1,102 | 0,955 | 0,872 | 1,238 | 1,086 | 1,095 | 1,08 | 0,08 | 5,2E-01 |
| P57780 | Alpha-actinin-4 OS=Mus musculus OX=10090 GN=Actn4 PE=1 SV=1 | 43 | 34 | 50 | 23 | 912 | 104,9 | 5,41 | 117,59 | 34 | 1,031 | 1,053 | 1,021 | 0,995 | 0,973 | 1,022 | 0,992 | 1,054 | 1,01 | 0,01 | 5,2E-01 |
| Q8BRK8 | 5'-AMP-activated protein kinase catalytic subunit alpha-2 OS=Mus | 10 | 4 | 4 | 3 | 552 | 62 | 7,87 | 5,5 | 4 | 1,4 | 1,082 | 1,016 | 0,947 | 0,882 | 1,31 | 0,998 | 0,858 | 1,10 | 0,10 | 5,2E-01 |
| Q9CZS1 | Aldehyde dehydrogenase X, mitochondrial OS=Mus musculus OX=1 | 17 | 8 | 9 | 7 | 519 | 57,5 | 7,02 | 24,5 | 8 | 0,885 | 0,866 | 1,018 | 1,047 | 1,065 | 0,91 | 0,985 | 1,01 | 0,96 | -0,04 | 5,2E-01 |
| Q9D394 | Protein RUFY3 OS=Mus musculus OX=10090 GN=Rufy3 PE=1 SV=1 | 30 | 14 | 18 | 14 | 469 | 53 | 5,49 | 34,15 | 14 | 1,169 | 1,017 | 1,028 | 1,053 | 0,965 | 1,067 | 1,023 | 1,09 | 1,03 | 0,03 | 5,2E-01 |
| Q62234 | Myomesin-1 OS=Mus musculus OX=10090 GN=Myom1 PE=1 SV=2 | 1 | 1 | 1 | 1 | 1667 | 185,3 | 6,16 | 0 | 1 | 0,741 | 1,121 | 1,013 | 0,949 | 0,723 | 1,034 | 1,409 | 1,102 | 0,90 | -0,10 | 5,2E-01 |
| Q8BGU5 | Cyclin-Y OS=Mus musculus OX=10090 GN=Ccny PE=1 SV=1 | 12 | 3 | 5 | 3 | 341 | 39,4 | 7,2 | 16,36 | 3 | 1,262 | 1,105 | 1,192 | 1,044 | 1,115 | 1,136 | 1,077 | 1,138 | 1,03 | 0,03 | 5,2E-01 |
| P26048 | Gamma-aminobutyric acid receptor subunit alpha-2 OS=Mus muscu | 5 | 2 | 3 | 1 | 451 | 51,1 | 9,06 | 7,65 | 2 | 1,263 | 1,314 | 0,763 | 1,116 | 1,326 | 0,993 | 1,079 | 1,538 | 0,90 | -0,10 | 5,2E-01 |
| Q9WUR2 | Enoyl-CoA delta isomerase 2, mitochondrial OS=Mus musculus OX= | 7 | 3 | 4 | 3 | 391 | 43,2 | 8,92 | 2,25 | 3 | 0,756 | 1,248 | 0,955 | 0,984 | 0,895 | 1,146 | 1,174 | 1,054 | 0,92 | -0,08 | 5,2E-01 |
| Q8CI59 | Metalloreductase STEAP3 OS=Mus musculus OX=10090 GN=Steap3 | 2 | 1 | 1 | 1 | 488 | 54,7 | 9,22 | 0 | 1 | 1,671 | 1,179 | 1,27 | 1,409 | 1,05 | 1,349 | 1,486 | 1,258 | 1,08 | 0,08 | 5,2E-01 |
| Q08189 | Protein-glutamine gamma-glutamyltransferase E OS=Mus musculus | 2 | 1 | 1 | 1 | 693 | 77,3 | 6,81 | 2,47 | 1 | 1,482 | 1,857 | 1,288 | 1,742 | 1,153 | 1,843 | 1,328 | 1,51 | 1,09 | 0,09 | 5,2E-01 |
| Q7M750 | Opalin OS=Mus musculus OX=10090 GN=Opalin PE=1 SV=1 | 26 | 2 | 2 | 2 | 143 | 15,8 | 4,88 | 6,1 | 2 | 1,299 | 1,078 | 0,661 | 0,708 | 0,96 | 1,049 | 0,748 | 0,424 | 1,18 | 0,18 | 5,2E-01 |
| Q9Z1Z2 | Serine-threonine kinase receptor-associated protein OS=Mus musc | 32 | 9 | 11 | 9 | 350 | 38,4 | 5,12 | 22,53 | 9 | 1,341 | 1,033 | 1,065 | 1,068 | 0,999 | 1,15 | 0,989 | 1,139 | 1,05 | 0,05 | 5,2E-01 |
| O88543 | COP9 signalosome complex subunit 3 OS=Mus musculus OX=10090 | 7 | 3 | 3 | 3 | 423 | 47,8 | 6,65 | 6,7 | 3 | 0,965 | 0,831 | 1,025 | 1,035 | 0,967 | 0,949 | 1,076 | 1,014 | 0,96 | -0,04 | 5,2E-01 |
| Q9Z0R6 | Intersectin-2 OS=Mus musculus OX=10090 GN=Itsn2 PE=1 SV=2 | 6 | 8 | 9 | 5 | 1659 | 188,8 | 8,13 | 9,65 | 8 | 1,388 | 1,093 | 1,076 | 1,222 | 1,016 | 1,31 | 1,061 | 1,127 | 1,06 | 0,06 | 5,2E-01 |
| Q9D8T7 | SRA stem-loop-interacting RNA-binding protein, mitochondrial OS= | 21 | 2 | 2 | 2 | 112 | 12,6 | 9,82 | 5,62 | 2 | 1,168 | 1,097 | 1,061 | 1,063 | 1,245 | 1,072 | 1,085 | 1,115 | 0,97 | -0,03 | 5,2E-01 |
| Q9Z1G3 | V-type proton ATPase subunit C 1 OS=Mus musculus OX=10090 GN | 30 | 16 | 27 | 16 | 382 | 43,9 | 7,46 | 62,18 | 16 | 0,899 | 0,869 | 0,936 | 0,988 | 0,988 | 0,85 | 0,996 | 0,975 | 0,97 | -0,03 | 5,2E-01 |
| Q8K4X7 | 1-acyl-sn-glycerol-3-phosphate acyltransferase delta OS=Mus musc | 6 | 2 | 3 | 2 | 378 | 43,8 | 8,38 | 4,1 | 2 | 0,785 | 0,867 | 1 | 0,839 | 0,866 | 0,809 | 0,773 | 0,897 | 1,04 | 0,04 | 5,2E-01 |
| Q9R0P6 | Signal peptidase complex catalytic subunit SEC11A OS=Mus muscul | 4 | 1 | 1 | 1 | 179 | 20,6 | 9,33 | 0 | 1 | 0,702 | 0,602 | 0,87 | 0,851 | 0,77 | 1,031 | 0,752 | 0,731 | 0,92 | -0,08 | 5,2E-01 |
| Q99N89 | 39S ribosomal protein L43, mitochondrial OS=Mus musculus OX=10 | 10 | 2 | 2 | 2 | 183 | 20,2 | 9,52 | 4,76 | 2 | 1,234 | 1,085 | 1,131 | 1,032 | 1,233 | 1,099 | 1,179 | 1,115 | 0,97 | -0,03 | 5,2E-01 |
| Q6P542 | ATP-binding cassette sub-family F member 1 OS=Mus musculus OX | 3 | 2 | 2 | 2 | 837 | 94,9 | 6,51 | 2,14 | 2 | 1,238 | 1,334 | 0,856 | 1,063 | 0,829 | 0,787 | 1,022 | 1,384 | 1,12 | 0,12 | 5,2E-01 |
| P14206 | 40S ribosomal protein SA OS=Mus musculus OX=10090 GN=Rpsa PI | 12 | 3 | 4 | 3 | 295 | 32,8 | 4,87 | 9,1 | 3 | 1,094 | 1,062 | 0,967 | 0,902 | 0,955 | 1,018 | 0,917 | 1 | 1,03 | 0,03 | 5,2E-01 |
| Q6A028 | Switch-associated protein 70 OS=Mus musculus OX=10090 GN=Swa | 2 | 1 | 1 | 1 | 585 | 69 | 6,05 | 2,12 | 1 | 0,985 | 0,913 | 1,326 | 1,092 | 0,922 | 0,887 | 1,076 | 1,136 | 1,07 | 0,07 | 5,2E-01 |
| Q99JY9 | Actin-related protein 3 OS=Mus musculus OX=10090 GN=Actr3 PE= | 23 | 8 | 25 | 5 | 418 | 47,3 | 5,88 | 53,36 | 8 | 1,124 | 0,94 | 0,946 | 1,11 | 0,964 | 0,943 | 1,042 | 1,02 | 1,04 | 0,04 | 5,2E-01 |
| Q61016 | Guanine nucleotide-binding protein G(I)/G(S)/G(O) subunit gamma | 57 | 3 | 7 | 3 | 68 | 7,5 | 8,51 | 20,56 | 3 | 0,974 | 0,908 | 0,982 | 0,945 | 1,023 | 0,933 | 1,202 | 0,858 | 0,95 | -0,05 | 5,2E-01 |
| Q8R0I4 | TM2 domain-containing protein 2 OS=Mus musculus OX=10090 GN | 5 | 1 | 1 | 1 | 213 | 22,8 | 5,22 | 2,39 | 1 | 1,105 | 1,27 | 1,51 | 1,119 | 1,101 | 1,405 | 1,043 | 1,117 | 1,07 | 0,07 | 5,2E-01 |
| Q9Z2I9 | Succinate--CoA ligase [ADP-forming] subunit beta, mitochondrial O | 41 | 18 | 34 | 18 | 463 | 50,1 | 7,01 | 87,5 | 18 | 0,996 | 0,984 | 0,992 | 0,914 | 0,907 | 1,109 | 0,984 | 1,011 | 0,97 | -0,03 | 5,2E-01 |
| Q9WV32 | Actin-related protein 2/3 complex subunit 1B OS=Mus musculus OX | 3 | 1 | 1 | 1 | 372 | 41 | 8,35 | 2,5 | 1 | 1,439 | 1,221 | 1,248 | 1,507 | 1,229 | 1,095 | 1,246 | 1,531 | 1,06 | 0,06 | 5,2E-01 |
| Q9D1A2 | Cytosolic non-specific dipeptidase OS=Mus musculus OX=10090 GN | 22 | 9 | 12 | 9 | 475 | 52,7 | 5,66 | 20,6 | 9 | 1,061 | 1,066 | 1,019 | 0,969 | 0,939 | 1,072 | 0,893 | 1,072 | 1,03 | 0,03 | 5,2E-01 |
| Q9D6J6 | NADH dehydrogenase [ubiquinone] flavoprotein 2, mitochondrial C | 44 | 9 | 28 | 9 | 248 | 27,3 | 7,4 | 71,66 | 9 | 0,971 | 0,947 | 1,055 | 0,985 | 1,024 | 1,031 | 0,938 | 1,06 | 0,98 | -0,02 | 5,2E-01 |
| Q505D1 | Serine/threonine-protein phosphatase 6 regulatory ankyrin repeat | 3 | 2 | 2 | 2 | 1053 | 112,8 | 6,27 | 2,46 | 2 | 1,781 | 1,18 | 1,769 | 1,129 | 1,058 | 1,931 | 0,906 | 1,178 | 1,15 | 0,15 | 5,2E-01 |
| P70697 | Uroporphyrinogen decarboxylase OS=Mus musculus OX=10090 GN | 10 | 2 | 2 | 2 | 367 | 40,7 | 6,65 | 2,23 | 2 | 2,517 | 1,725 | 1,621 | 0,816 | 0,939 | 1,884 | 1,402 | 1,377 | 1,19 | 0,19 | 5,2E-01 |
| Q8BVQ5 | Protein phosphatase methylesterase 1 OS=Mus musculus OX=1009 | 11 | 5 | 11 | 5 | 386 | 42,2 | 5,97 | 27,95 | 5 | 0,936 | 1,005 | 0,981 | 1,103 | 0,952 | 0,93 | 0,99 | 1,038 | 1,03 | 0,03 | 5,2E-01 |
| P56376 | Acylphosphatase-1 OS=Mus musculus OX=10090 GN=Acyp1 PE=1 S | 27 | 2 | 2 | 2 | 99 | 11,2 | 9,04 | 5,45 | 2 | 2,163 | 1,343 | 1,512 | 1,097 | 1,01 | 1,787 | 1,241 | 1,32 | 1,14 | 0,14 | 5,2E-01 |
| Q8BMF4 | Dihydrolipoyllysine-residue acetyltransferase component of pyruva | 21 | 11 | 17 | 11 | 642 | 67,9 | 8,57 | 43,2 | 11 | 1,085 | 0,997 | 1,089 | 1,02 | 1,061 | 1,098 | 1,043 | 1,059 | 0,98 | -0,02 | 5,2E-01 |
| Q8BP67 | 60S ribosomal protein L24 OS=Mus musculus OX=10090 GN=Rpl24 | 8 | 1 | 5 | 1 | 157 | 17,8 | 11,25 | 10,97 | 1 | 0,848 | 0,876 | 1,083 | 1,089 | 1,011 | 0,944 | 1,029 | 1,11 | 0,95 | -0,05 | 5,2E-01 |
| P18242 | Cathepsin D OS=Mus musculus OX=10090 GN=Ctsd PE=1 SV=1 | 6 | 2 | 2 | 2 | 410 | 44,9 | 7,15 | 5,12 | 2 | 0,954 | 0,877 | 1,048 | 0,964 | 1,074 | 0,998 | 0,985 | 0,915 | 0,97 | -0,03 | 5,2E-01 |
| P15116 | Cadherin-2 OS=Mus musculus OX=10090 GN=Cdh2 PE=1 SV=2 | 10 | 8 | 11 | 8 | 906 | 99,7 | 4,78 | 14,03 | 8 | 1,172 | 1,1 | 1,051 | 0,888 | 1,032 | 1,24 | 1,077 | 1,067 | 0,95 | -0,05 | 5,3E-01 |
| Q60900 | ELAV-like protein 3 OS=Mus musculus OX=10090 GN=Elavl3 PE=1 S | 5 | 2 | 2 | 1 | 367 | 39,5 | 9,28 | 2,42 | 2 | 1,426 | 1,269 | 1,124 | 0,953 | 1,039 | 1,394 | 0,988 | 0,968 | 1,09 | 0,09 | 5,3E-01 |
| Q91Z67 | SLIT-ROBO Rho GTPase-activating protein 2 OS=Mus musculus OX= | 10 | 10 | 16 | 10 | 1071 | 120,7 | 6,64 | 23,3 | 10 | 1,125 | 1,122 | 1,071 | 1,095 | 1,033 | 1,112 | 1,177 | 1,196 | 0,98 | -0,02 | 5,3E-01 |
| Q9D2P8 | Myelin-associated oligodendrocyte basic protein OS=Mus musculus | 12 | 2 | 2 | 2 | 170 | 19,2 | 11,14 | 2,33 | 2 | 1,051 | 0,784 | 0,955 | 0,83 | 0,818 | 2,485 | 0,769 | 0,717 | 0,76 | -0,24 | 5,3E-01 |
| Q3UYG8 | ADP-ribose glycohydrolase MACROD2 OS=Mus musculus OX=10090 | 7 | 3 | 3 | 3 | 475 | 52,1 | 4,75 | 7,63 | 3 | 0,94 | 0,911 | 0,885 | 0,986 | 0,9 | 1,133 | 0,942 | 0,906 | 0,96 | -0,04 | 5,3E-01 |
| P21803 | Fibroblast growth factor receptor 2 OS=Mus musculus OX=10090 G | 2 | 1 | 1 | 1 | 821 | 91,9 | 5,74 | 0 | 1 | 0,568 | 1,015 | 1,058 | 1,041 | 0,915 | 0,617 | 0,417 | 1,182 | 1,18 | 0,18 | 5,3E-01 |
| Q8CCN5 | Breast carcinoma-amplified sequence 3 homolog OS=Mus musculu | 11 | 9 | 9 | 9 | 928 | 101 | 6,7 | 15,73 | 9 | 1,174 | 0,807 | 1,004 | 0,9 | 0,877 | 0,865 | 0,871 | 1,034 | 1,07 | 0,07 | 5,3E-01 |
| A2AGL3 | Ryanodine receptor 3 OS=Mus musculus OX=10090 GN=Ryr3 PE=1 | 1 | 4 | 4 | 1 | 4863 | 551 | 5,78 | 9,49 | 4 | 0,809 | 1 | 1,097 | 1,03 | 1,006 | 1,065 | 1,114 | 0,944 | 0,95 | -0,05 | 5,3E-01 |
| Q8C079 | Striatin-interacting protein 1 OS=Mus musculus OX=10090 GN=Stri | 4 | 3 | 3 | 1 | 837 | 95,5 | 6,25 | 5,37 | 3 | 0,822 | 0,889 | 0,911 | 0,765 | 0,947 | 0,689 | 0,878 | 1,136 | 0,93 | -0,07 | 5,3E-01 |
| P47753 | F-actin-capping protein subunit alpha-1 OS=Mus musculus OX=100 | 35 | 7 | 7 | 7 | 286 | 32,9 | 5,55 | 15,31 | 7 | 1,08 | 1,106 | 1,167 | 1,146 | 1,075 | 1,189 | 1,037 | 1,097 | 1,02 | 0,02 | 5,3E-01 |
| Q8VE52 | Opioid growth factor receptor-like protein 1 OS=Mus musculus OX= | 4 | 2 | 3 | 2 | 464 | 52,2 | 6,57 | 0 | 2 | 0,837 | 1,198 | 0,839 | 0,901 | 1,055 | 1,154 | 0,872 | 0,974 | 0,93 | -0,07 | 5,3E-01 |
| Q9WTN0 | Geranylgeranyl pyrophosphate synthase OS=Mus musculus OX=100 | 3 | 1 | 1 | 1 | 300 | 34,7 | 6,46 | 2,52 | 1 | 1,112 | 1,251 | 1,183 | 1,089 | 1,045 | 1,193 | 1,089 | 1,173 | 1,03 | 0,03 | 5,3E-01 |
| Q8JZW4 | Copine-5 OS=Mus musculus OX=10090 GN=Cpne5 PE=1 SV=1 | 21 | 10 | 11 | 6 | 593 | 65,6 | 5,77 | 20,63 | 10 | 1,039 | 0,943 | 0,937 | 0,9 | 0,864 | 0,932 | 1,032 | 0,857 | 1,04 | 0,04 | 5,3E-01 |
| Q8BH57 | WD repeat-containing protein 48 OS=Mus musculus OX=10090 GN= | 24 | 14 | 16 | 14 | 676 | 76 | 7,17 | 35,29 | 14 | 0,945 | 1,071 | 0,981 | 0,976 | 0,972 | 1,131 | 0,984 | 1,007 | 0,97 | -0,03 | 5,3E-01 |
| P32233 | Developmentally-regulated GTP-binding protein 1 OS=Mus musculu | 4 | 1 | 1 | 1 | 367 | 40,5 | 8,9 | 0 | 1 | 0,716 | 1,423 | 0,718 | 0,608 | 0,784 | 0,649 | 0,549 | 0,937 | 1,19 | 0,19 | 5,3E-01 |
| P52760 | 2-iminobutanoate/2-iminopropanoate deaminase OS=Mus musculu | 10 | 1 | 1 | 1 | 135 | 14,2 | 8,68 | 2,53 | 1 | 1,036 | 1,119 | 1,15 | 0,97 | 1,063 | 1,134 | 0,911 | 1,001 | 1,04 | 0,04 | 5,3E-01 |
| Q9DC28 | Casein kinase I isoform delta OS=Mus musculus OX=10090 GN=Csn | 9 | 3 | 4 | 1 | 415 | 47,3 | 9,74 | 5,43 | 3 | 1,084 | 1,003 | 1,312 | 1,16 | 1,098 | 0,783 | 1,41 | 0,853 | 1,10 | 0,10 | 5,3E-01 |
| Q80UM3 | N-alpha-acetyltransferase 15, NatA auxiliary subunit OS=Mus musc | 7 | 6 | 7 | 6 | 865 | 100,9 | 7,62 | 15,59 | 6 | 1,169 | 1,138 | 1,091 | 1,276 | 1,196 | 1,182 | 1,193 | 1,208 | 0,98 | -0,02 | 5,3E-01 |
| Q9WTI7 | Unconventional myosin-Ic OS=Mus musculus OX=10090 GN=Myo1 | 1 | 1 | 1 | 1 | 1063 | 121,9 | 9,35 | 2,11 | 1 | 0,75 | 1,26 | 1,177 | 0,861 | 0,954 | 0,422 | 1,052 | 1,095 | 1,15 | 0,15 | 5,3E-01 |
| Q80ZK0 | 28S ribosomal protein S10, mitochondrial OS=Mus musculus OX=10 | 7 | 1 | 1 | 1 | 160 | 18,7 | 8,84 | 0 | 1 | 1,91 | 2,648 | 1,578 | 2,261 | 2,154 | 1,867 | 1,913 | 1,823 | 1,08 | 0,08 | 5,3E-01 |
| Q8BH58 | TIP41-like protein OS=Mus musculus OX=10090 GN=Tiprl PE=1 SV= | 22 | 4 | 5 | 4 | 271 | 31,2 | 5,63 | 6,22 | 4 | 0,979 | 0,988 | 0,815 | 0,999 | 0,852 | 1,101 | 1,114 | 0,922 | 0,95 | -0,05 | 5,3E-01 |
| Q99JW4 | LIM and senescent cell antigen-like-containing domain protein 1 OS | 7 | 2 | 2 | 2 | 325 | 37,2 | 8,05 | 4,01 | 2 | 1,088 | 1,315 | 1,318 | 1,066 | 1,397 | 1,48 | 1,229 | 1,011 | 0,94 | -0,06 | 5,3E-01 |
| Q6NVE8 | WD repeat-containing protein 44 OS=Mus musculus OX=10090 GN= | 18 | 12 | 13 | 12 | 915 | 101,5 | 5,24 | 33,3 | 12 | 1,035 | 1,254 | 1,088 | 1,071 | 1 | 1,128 | 1,04 | 1,126 | 1,04 | 0,04 | 5,3E-01 |
| Q3V3Q7 | Phosphofurin acidic cluster sorting protein 2 OS=Mus musculus OX | 1 | 1 | 1 | 1 | 862 | 94,9 | 6,51 | 0 | 1 | 1,613 | 1,113 | 1,566 | 1,629 | 1,172 | 1,434 | 1,246 | 1,641 | 1,08 | 0,08 | 5,3E-01 |
| P34884 | Macrophage migration inhibitory factor OS=Mus musculus OX=100 | 6 | 1 | 2 | 1 | 115 | 12,5 | 7,34 | 4,71 | 1 | 1,256 | 0,885 | 1,111 | 0,94 | 0,9 | 0,961 | 1,067 | 1,021 | 1,06 | 0,06 | 5,3E-01 |

| | | | | | | | | | | | | | | | | | | | | | |
|---|---|---|---|---|---|---|---|---|---|---|---|---|---|---|---|---|---|---|---|---|---|
| B1AXV0 | DOMON domain-containing protein FRRS1L OS=Mus musculus OX= | 4 | 1 | 1 | 1 | 293 | 32,5 | 5,43 | 0 | 1 | 0,721 | 0,732 | 0,706 | 0,649 | 0,604 | 0,787 | 0,696 | 0,591 | 1,05 | 0,05 | 5,3E-01 |
| Q8CA95-3 | Isoform 3 of cAMP and cAMP-inhibited cGMP 3',5'-cyclic phosphod | 10 | 7 | 10 | 7 | 796 | 90,3 | 6,65 | 15,56 | 7 | 1,177 | 0,895 | 1,048 | 0,82 | 0,869 | 1,072 | 1,281 | 1,025 | 0,93 | -0,07 | 5,3E-01 |
| Q99M04 | Lipoyl synthase, mitochondrial OS=Mus musculus OX=10090 GN=Li | 2 | 1 | 1 | 1 | 373 | 41,9 | 8,88 | 1,91 | 1 | 0,546 | 0,75 | 0,916 | 0,887 | 0,806 | 0,704 | 0,824 | 1,065 | 0,91 | -0,09 | 5,3E-01 |
| Q9D8E6 | 60S ribosomal protein L4 OS=Mus musculus OX=10090 GN=Rpl4 PE | 15 | 6 | 7 | 6 | 419 | 47,1 | 11 | 19,48 | 6 | 1,114 | 1,025 | 1,06 | 0,898 | 1,077 | 1,06 | 0,969 | 1,145 | 0,96 | -0,04 | 5,3E-01 |
| P09470 | Angiotensin-converting enzyme OS=Mus musculus OX=10090 GN=/ | 1 | 1 | 1 | 1 | 1312 | 150,8 | 6,55 | 2,64 | 1 | 2,029 | 1,542 | 1,06 | 1,106 | 0,876 | 0,787 | 1,772 | 1,444 | 1,18 | 0,18 | 5,3E-01 |
| Q3ULD5 | Methylcrotonoyl-CoA carboxylase beta chain, mitochondrial OS=M | 10 | 5 | 6 | 5 | 563 | 61,3 | 8 | 13,5 | 5 | 0,982 | 0,926 | 1,101 | 1,006 | 0,846 | 1,1 | 0,924 | 0,975 | 1,04 | 0,04 | 5,4E-01 |
| P97379 | Ras GTPase-activating protein-binding protein 2 OS=Mus musculus | 5 | 2 | 2 | 2 | 482 | 54,1 | 5,62 | 2,47 | 2 | 1,068 | 0,87 | 1,12 | 0,866 | 1,066 | 0,93 | 0,92 | 1,297 | 0,93 | -0,07 | 5,4E-01 |
| Q8K4F5 | Protein ABHD11 OS=Mus musculus OX=10090 GN=Abhd11 PE=1 SV | 7 | 3 | 4 | 3 | 307 | 33,5 | 9,61 | 7,34 | 3 | 1,351 | 1,167 | 1,116 | 1,006 | 1,129 | 1,082 | 1,118 | 1,12 | 1,04 | 0,04 | 5,4E-01 |
| Q9CQP0 | 39S ribosomal protein L33, mitochondrial OS=Mus musculus OX=1( | 12 | 1 | 1 | 1 | 65 | 7,4 | 10,89 | 2,36 | 1 | 0,994 | 1,487 | 1,316 | 1,44 | 1,431 | 0,859 | 1,151 | 1,352 | 1,09 | 0,09 | 5,4E-01 |
| P35922 | Synaptic functional regulator FMR1 OS=Mus musculus OX=10090 G | 4 | 3 | 3 | 2 | 614 | 68,9 | 7,62 | 7,93 | 3 | 0,871 | 0,823 | 0,93 | 0,911 | 1,171 | 0,837 | 0,896 | 0,847 | 0,94 | -0,06 | 5,4E-01 |
| A2AWA9 | Rab GTPase-activating protein 1 OS=Mus musculus OX=10090 GN= | 12 | 12 | 14 | 10 | 1064 | 120,7 | 5,25 | 19,49 | 12 | 1,079 | 1,149 | 1,142 | 1,07 | 1,04 | 1,294 | 1,107 | 1,15 | 0,97 | -0,03 | 5,4E-01 |
| Q99P47 | Contactin-associated protein-like 4 OS=Mus musculus OX=10090 G | 2 | 3 | 3 | 3 | 1310 | 144,6 | 7,06 | 5,94 | 3 | 1,194 | 1,459 | 1,142 | 1,309 | 1,064 | 1,033 | 1,238 | 1,455 | 1,07 | 0,07 | 5,4E-01 |
| Q61161 | Mitogen-activated protein kinase kinase kinase 2 OS=Mus m | 3 | 2 | 3 | 2 | 821 | 91,2 | 6,46 | 6,42 | 2 | 1,175 | 1,874 | 1,024 | 1,254 | 1,102 | 0,944 | 1,359 | 1,363 | 1,12 | 0,12 | 5,4E-01 |
| Q9Z100 | Probable carboxypeptidase X1 OS=Mus musculus OX=10090 GN=Cp | 1 | 1 | 1 | 1 | 722 | 80,9 | 6,89 | 2,08 | 1 | 0,985 | 1,078 | 1,074 | 1,106 | 1,235 | 0,88 | 1,016 | 0,883 | 1,06 | 0,06 | 5,4E-01 |
| Q80XI4 | Phosphatidylinositol 5-phosphate 4-kinase type-2 beta OS=Mus mu | 21 | 8 | 9 | 4 | 416 | 47,3 | 7,33 | 26,24 | 8 | 1,112 | 1,064 | 1,061 | 1,106 | 0,912 | 1,058 | 1,089 | 1,148 | 1,03 | 0,03 | 5,4E-01 |
| Q9DAK9 | 14 kDa phosphohistidine phosphatase OS=Mus musculus OX=1009( | 19 | 2 | 2 | 2 | 124 | 14 | 5,53 | 4,15 | 2 | 1,042 | 0,693 | 1,101 | 1,076 | 0,912 | 1,24 | 1,029 | 1,038 | 0,93 | -0,07 | 5,4E-01 |
| Q9EPK2 | Protein XRP2 OS=Mus musculus OX=10090 GN=Rp2 PE=1 SV=3 | 2 | 1 | 1 | 1 | 347 | 39,4 | 5,24 | 1,92 | 1 | 0,969 | 0,927 | 1,141 | 0,981 | 0,834 | 1,19 | 0,787 | 0,942 | 1,07 | 0,07 | 5,4E-01 |
| P63300 | Selenoprotein W OS=Mus musculus OX=10090 GN=Selenow PE=1 S | 22 | 3 | 3 | 3 | 88 | 9,7 | 8,72 | 2,04 | 3 | 1,285 | 0,92 | 1,38 | 1,424 | 1,672 | 0,878 | 1,456 | 1,551 | 0,90 | -0,10 | 5,4E-01 |
| P09541 | Myosin light chain 4 OS=Mus musculus OX=10090 GN=Myl4 PE=1 S | 10 | 2 | 2 | 2 | 193 | 21,1 | 5,03 | 3,13 | 2 | 0,915 | 1,01 | 1,12 | 0,767 | 0,989 | 0,826 | 0,911 | 0,871 | 1,06 | 0,06 | 5,4E-01 |
| P47743 | Metabotropic glutamate receptor 8 OS=Mus musculus OX=10090 G | 1 | 1 | 1 | 1 | 908 | 101,8 | 8,15 | 2,67 | 1 | 0,703 | 0,79 | 1,199 | 0,957 | 0,92 | 0,708 | 1,105 | 1,387 | 0,89 | -0,11 | 5,4E-01 |
| Q66JS6 | Eukaryotic translation initiation factor 3 subunit J-B OS=Mus muscu | 13 | 4 | 4 | 4 | 263 | 29,5 | 4,81 | 7,54 | 4 | 1,164 | 1,379 | 1,468 | 1,315 | 1,283 | 1,328 | 1,329 | 1,202 | 1,04 | 0,04 | 5,4E-01 |
| P35505 | Fumarylacetoacetase OS=Mus musculus OX=10090 GN=Fah PE=1 S | 10 | 4 | 4 | 4 | 419 | 46,1 | 7,18 | 6,58 | 4 | 0,916 | 0,913 | 1,138 | 0,938 | 1,055 | 0,825 | 0,933 | 0,904 | 1,05 | 0,05 | 5,4E-01 |
| Q62415 | Apoptosis-stimulating of p53 protein 1 OS=Mus musculus OX=1009 | 3 | 2 | 3 | 2 | 1087 | 119,1 | 6,68 | 2,09 | 2 | 0,58 | 1,095 | 0,743 | 0,899 | 0,478 | 0,777 | 0,958 | 0,719 | 1,13 | 0,13 | 5,4E-01 |
| Q9DBS5 | Kinesin light chain 4 OS=Mus musculus OX=10090 GN=Klc4 PE=1 SV | 9 | 5 | 5 | 1 | 619 | 68,6 | 6,09 | 13,5 | 5 | 0,816 | 0,838 | 0,89 | 0,993 | 0,758 | 1,087 | 0,739 | 1,342 | 0,90 | -0,10 | 5,4E-01 |
| P61087 | Ubiquitin-conjugating enzyme E2 K OS=Mus musculus OX=10090 G | 14 | 2 | 2 | 2 | 200 | 22,4 | 5,44 | 3,33 | 2 | 0,743 | 0,783 | 1,071 | 1,077 | 1,008 | 1,185 | 0,943 | 0,839 | 0,92 | -0,08 | 5,4E-01 |
| Q14BI2 | Metabotropic glutamate receptor 2 OS=Mus musculus OX=10090 G | 8 | 8 | 15 | 7 | 872 | 95,8 | 8,06 | 26,42 | 8 | 1,159 | 1,083 | 1,073 | 1,097 | 1,247 | 1,149 | 1,104 | 1,037 | 0,97 | -0,03 | 5,4E-01 |
| P62996 | Transformer-2 protein homolog beta OS=Mus musculus OX=10090 | 6 | 2 | 2 | 2 | 288 | 33,6 | 11,25 | 1,74 | 2 | 1,131 | 1,33 | 1,086 | 0,993 | 1,213 | 1,303 | 0,978 | 1,322 | 0,94 | -0,06 | 5,4E-01 |
| Q9WTK7 | Serine/threonine-protein kinase STK11 OS=Mus musculus OX=1009 | 6 | 2 | 2 | 2 | 436 | 49,2 | 6,84 | 5,82 | 2 | 0,849 | 0,88 | 1,068 | 0,978 | 0,938 | 1,118 | 0,946 | 0,945 | 0,96 | -0,04 | 5,4E-01 |
| Q9ESN6 | Tripartite motif-containing protein 2 OS=Mus musculus OX=10090 | 16 | 9 | 12 | 9 | 744 | 81,4 | 6,96 | 25,02 | 9 | 1,192 | 1,036 | 1,104 | 1,002 | 0,899 | 1,094 | 1,088 | 1,088 | 1,04 | 0,04 | 5,4E-01 |
| Q9WTL4 | Insulin receptor-related protein OS=Mus musculus OX=10090 GN=I | 1 | 1 | 1 | 1 | 1300 | 144,8 | 6,64 | 1,89 | 1 | 1,432 | 2,143 | 1,075 | 0,649 | 1,624 | 1,366 | 0,784 | 0,451 | 1,25 | 0,25 | 5,4E-01 |
| Q8K1J6 | CCA tRNA nucleotidyltransferase 1, mitochondrial OS=Mus muscul | 9 | 4 | 4 | 4 | 434 | 49,9 | 8,56 | 9,51 | 4 | 0,903 | 0,994 | 1,105 | 1,049 | 1,124 | 1,04 | 1,091 | 0,946 | 0,96 | -0,04 | 5,4E-01 |
| Q920P3 | BMP/retinoic acid-inducible neural-specific protein 1 OS=Mus musc | 8 | 5 | 6 | 5 | 760 | 88,6 | 8,97 | 12,1 | 5 | 0,87 | 0,938 | 1,103 | 1,138 | 1,015 | 1,038 | 1,131 | 1,044 | 0,96 | -0,04 | 5,4E-01 |
| P12849 | cAMP-dependent protein kinase type I-beta regulatory subunit OS= | 11 | 5 | 5 | 3 | 381 | 43,2 | 5,96 | 13,21 | 5 | 0,724 | 0,803 | 1,005 | 0,978 | 1,058 | 0,89 | 0,935 | 0,84 | 0,94 | -0,06 | 5,4E-01 |
| P68181 | cAMP-dependent protein kinase catalytic subunit beta OS=Mus mu | 24 | 7 | 9 | 2 | 351 | 40,7 | 8,56 | 25,33 | 7 | 0,925 | 0,91 | 0,945 | 1,156 | 1,131 | 0,81 | 1,191 | 1,067 | 0,94 | -0,06 | 5,4E-01 |
| Q8CBE3 | WD repeat-containing protein 37 OS=Mus musculus OX=10090 GN= | 20 | 8 | 12 | 8 | 496 | 55 | 7,23 | 30,4 | 8 | 1,116 | 1,143 | 1,082 | 0,983 | 0,934 | 1,159 | 1,011 | 1,068 | 1,04 | 0,04 | 5,4E-01 |
| Q9JKR6 | Hypoxia up-regulated protein 1 OS=Mus musculus OX=10090 GN=H | 12 | 10 | 16 | 10 | 999 | 111,1 | 5,19 | 29,09 | 10 | 0,981 | 0,914 | 0,993 | 1,024 | 1,042 | 0,948 | 1,02 | 0,982 | 0,98 | -0,02 | 5,4E-01 |
| Q64520 | Guanylate kinase OS=Mus musculus OX=10090 GN=Guk1 PE=1 SV= | 22 | 3 | 3 | 3 | 198 | 21,9 | 6,55 | 7,13 | 3 | 1,885 | 0,867 | 1,122 | 0,818 | 1,107 | 1,541 | 1,241 | 1,489 | 0,87 | -0,13 | 5,4E-01 |
| Q9R049 | E3 ubiquitin-protein ligase AMFR OS=Mus musculus OX=10090 GN= | 1 | 1 | 1 | 1 | 643 | 73,1 | 6,46 | 0 | 1 | 0,754 | 1,153 | 0,876 | 1,023 | 0,909 | 0,766 | 1,058 | 0,792 | 1,08 | 0,08 | 5,4E-01 |
| Q06335 | Amyloid-like protein 2 OS=Mus musculus OX=10090 GN=Aplp2 PE= | 6 | 4 | 5 | 3 | 707 | 80,4 | 4,7 | 7,39 | 4 | 1,241 | 0,892 | 1,193 | 0,981 | 0,899 | 0,892 | 1,023 | 1,207 | 1,07 | 0,07 | 5,4E-01 |
| Q9CXF4 | TBC1 domain family member 15 OS=Mus musculus OX=10090 GN= | 6 | 4 | 4 | 4 | 671 | 76,5 | 5,3 | 2,99 | 4 | 1,38 | 0,958 | 1,021 | 1,159 | 1,245 | 1,06 | 1,148 | 1,355 | 0,94 | -0,06 | 5,4E-01 |
| Q99PL6 | UBX domain-containing protein 6 OS=Mus musculus OX=10090 GN | 12 | 4 | 4 | 4 | 442 | 49,8 | 8,54 | 7,27 | 4 | 0,873 | 0,98 | 0,829 | 0,978 | 0,907 | 0,974 | 0,968 | 0,918 | 0,97 | -0,03 | 5,4E-01 |
| Q8BH44 | Coronin-2B OS=Mus musculus OX=10090 GN=Coro2b PE=1 SV=2 | 10 | 4 | 6 | 4 | 480 | 54,9 | 8,27 | 11,85 | 4 | 1,233 | 0,987 | 0,997 | 0,866 | 0,918 | 1,088 | 0,888 | 0,962 | 1,06 | 0,06 | 5,5E-01 |
| Q9JJ69 | Kv channel-interacting protein 2 OS=Mus musculus OX=10090 GN= | 13 | 3 | 3 | 3 | 270 | 30,9 | 5,05 | 5,08 | 3 | 0,947 | 0,925 | 1,019 | 0,923 | 0,732 | 1,094 | 0,997 | 0,754 | 1,07 | 0,07 | 5,5E-01 |
| Q8JZP2 | Synapsin-3 OS=Mus musculus OX=10090 GN=Syn3 PE=1 SV=2 | 13 | 5 | 7 | 5 | 579 | 63,3 | 9,38 | 13,69 | 5 | 1,183 | 0,997 | 0,905 | 1,023 | 0,891 | 1,078 | 0,923 | 1,03 | 1,05 | 0,05 | 5,5E-01 |
| P29699 | Alpha-2-HS-glycoprotein OS=Mus musculus OX=10090 GN=Ahsg PE | 2 | 1 | 1 | 1 | 345 | 37,3 | 6,51 | 2,67 | 1 | 1,344 | 1,175 | 0,763 | 0,557 | 1,455 | 1,544 | 1,026 | 0,556 | 0,84 | -0,16 | 5,5E-01 |
| A3KFX0 | Cytosolic 5'-nucleotidase 1A OS=Mus musculus OX=10090 GN=Nt5c | 3 | 1 | 1 | 1 | 365 | 40,7 | 7,06 | 1,8 | 1 | 0,01 | 1,197 | 1,44 | 1,212 | 1,302 | 0,787 | 1,724 | 1,018 | 0,80 | -0,20 | 5,5E-01 |
| Q3LAC4 | Phosphatidylinositol 3,4,5-trisphosphate-dependent Rac exchanger | 2 | 3 | 3 | 3 | 1598 | 181,6 | 7,34 | 0 | 3 | 1,192 | 0,974 | 0,815 | 0,817 | 1,017 | 0,946 | 0,605 | 0,906 | 1,09 | 0,09 | 5,5E-01 |
| Q9DCH4 | Eukaryotic translation initiation factor 3 subunit F OS=Mus muscul | 2 | 1 | 1 | 1 | 361 | 38 | 5,58 | 0 | 1 | 0,943 | 0,98 | 1,239 | 0,861 | 1,245 | 1,076 | 1,089 | 0,891 | 0,94 | -0,06 | 5,5E-01 |
| Q3B7Z2 | Oxysterol-binding protein 1 OS=Mus musculus OX=10090 GN=Osbp | 13 | 8 | 8 | 7 | 805 | 88,7 | 7,2 | 16,47 | 8 | 1,306 | 1,228 | 1,205 | 0,926 | 0,943 | 1,193 | 1,003 | 1,245 | 1,06 | 0,06 | 5,5E-01 |
| Q99JR6 | Nicotinamide/nicotinic acid mononucleotide adenylyltransferase 3 | 9 | 2 | 2 | 2 | 245 | 27,7 | 8,56 | 4,57 | 2 | 0,54 | 0,924 | 0,845 | 0,818 | 0,883 | 0,901 | 0,764 | 0,807 | 0,93 | -0,07 | 5,5E-01 |
| Q6P9Q6 | FK506-binding protein 15 OS=Mus musculus OX=10090 GN=Fkbp15 | 1 | 1 | 1 | 1 | 1216 | 132,9 | 5,07 | 1,8 | 1 | 1,382 | 1,088 | 1,096 | 0,905 | 0,819 | 1,052 | 1,183 | 1,098 | 1,08 | 0,08 | 5,5E-01 |
| Q5U458 | DnaJ homolog subfamily C member 11 OS=Mus musculus OX=1009 | 9 | 5 | 5 | 5 | 559 | 63,2 | 8,32 | 7,7 | 5 | 0,966 | 0,949 | 1,074 | 0,986 | 0,981 | 0,883 | 1,046 | 0,954 | 1,03 | 0,03 | 5,5E-01 |
| Q99KF1 | Transmembrane emp24 domain-containing protein 9 OS=Mus mus | 7 | 2 | 2 | 2 | 235 | 27,1 | 8,41 | 5,31 | 2 | 0,92 | 1,15 | 1,026 | 1,039 | 1,074 | 0,935 | 1,028 | 0,953 | 1,04 | 0,04 | 5,5E-01 |
| Q8R066 | Complement C1q tumor necrosis factor-related protein 4 OS=Mus | 12 | 3 | 3 | 3 | 326 | 35 | 8,98 | 7,39 | 3 | 0,74 | 0,883 | 1,165 | 1,022 | 1,086 | 0,957 | 0,948 | 1,068 | 0,94 | -0,06 | 5,5E-01 |
| Q8R550 | SH3 domain-containing kinase-binding protein 1 OS=Mus musculus | 1 | 1 | 1 | 1 | 709 | 78,1 | 7,55 | 0 | 1 | 1,575 | 1,598 | 0,893 | 1,561 | 1,675 | 1,325 | 1,607 | 1,495 | 0,92 | -0,08 | 5,5E-01 |
| A2BDX3 | Adenylyltransferase and sulfurtransferase MOCS3 OS=Mus muscul | 9 | 3 | 3 | 3 | 460 | 49,3 | 7,49 | 5,69 | 3 | 0,715 | 0,823 | 0,877 | 1,143 | 0,994 | 0,896 | 0,872 | 1,049 | 0,93 | -0,07 | 5,5E-01 |
| Q9D1I5 | Methylmalonyl-CoA epimerase, mitochondrial OS=Mus musculus O | 7 | 1 | 2 | 1 | 178 | 19 | 9,09 | 7,46 | 1 | 1,344 | 1,079 | 1,318 | 1,242 | 1,12 | 1,212 | 1,254 | 1,229 | 1,03 | 0,03 | 5,5E-01 |
| Q00288 | Paired box protein Pax-8 OS=Mus musculus OX=10090 GN=Pax8 PE | 4 | 1 | 1 | 1 | 457 | 48,8 | 7,84 | 0 | 1 | 0,529 | 1,115 | 1,097 | 0,928 | 1,041 | 0,872 | 1,059 | 1,06 | 0,91 | -0,09 | 5,5E-01 |
| Q01097 | Glutamate receptor ionotropic, NMDA 2B OS=Mus musculus OX=1( | 8 | 10 | 10 | 10 | 1482 | 165,9 | 6,87 | 15,17 | 10 | 0,987 | 1,159 | 0,925 | 0,899 | 0,84 | 1,038 | 1,015 | 0,885 | 1,05 | 0,05 | 5,5E-01 |
| Q91V14 | Solute carrier family 12 member 5 OS=Mus musculus OX=10090 GN | 14 | 15 | 34 | 13 | 1138 | 126,2 | 6,74 | 77,9 | 15 | 1,032 | 1,08 | 1,024 | 0,959 | 0,951 | 1,069 | 0,957 | 1,023 | 1,02 | 0,02 | 5,5E-01 |
| Q6QD59 | Vesicle transport protein SEC20 OS=Mus musculus OX=10090 GN=E | 4 | 1 | 1 | 1 | 228 | 26,2 | 8,75 | 0 | 1 | 1,008 | 0,965 | 0,775 | 0,979 | 0,814 | 0,703 | 0,93 | 1,049 | 1,07 | 0,07 | 5,5E-01 |
| Q99PU5 | Long-chain-fatty-acid--CoA ligase ACSBG1 OS=Mus musculus OX=1( | 14 | 9 | 12 | 9 | 721 | 80,4 | 5,94 | 27,77 | 9 | 1,159 | 1,065 | 1,107 | 0,958 | 0,946 | 1,132 | 1,059 | 1,005 | 1,04 | 0,04 | 5,5E-01 |

| | | | | | | | | | | | | | | | | | | | | | |
|---|---|---|---|---|---|---|---|---|---|---|---|---|---|---|---|---|---|---|---|---|---|
| Q4VC33 | E3 ubiquitin-protein transferase MAEA OS=Mus musculus OX=1009 | 2 | 1 | 1 | 1 | 396 | 45,3 | 8,69 | 0 | 1 | 0,955 | 1,366 | 1,729 | 1,739 | 1,479 | 1,849 | 1,355 | 1,647 | 0,91 | -0,09 | 5,5E-01 |
| O55017 | Voltage-dependent N-type calcium channel subunit alpha-1B OS=M | 5 | 11 | 14 | 10 | 2327 | 261,3 | 8,59 | 13,71 | 11 | 0,912 | 0,973 | 0,944 | 1,006 | 0,892 | 0,941 | 0,998 | 1,159 | 0,96 | -0,04 | 5,5E-01 |
| P10637 | Microtubule-associated protein tau OS=Mus musculus OX=10090 G | 39 | 23 | 47 | 8 | 733 | 76,2 | 6,79 | 82,96 | 23 | 0,897 | 0,894 | 0,977 | 1,106 | 0,961 | 0,947 | 1,041 | 1,072 | 0,96 | -0,04 | 5,5E-01 |
| P27612 | Phospholipase A-2-activating protein OS=Mus musculus OX=10090 | 16 | 11 | 11 | 11 | 794 | 87,2 | 6,14 | 26,61 | 11 | 0,986 | 1,001 | 1,105 | 1,035 | 0,931 | 1,063 | 1,039 | 0,995 | 1,02 | 0,02 | 5,5E-01 |
| P54822 | Adenylosuccinate lyase OS=Mus musculus OX=10090 GN=Adsl PE=1 | 10 | 3 | 3 | 3 | 484 | 54,8 | 7,27 | 9,98 | 3 | 1,268 | 0,999 | 1,145 | 0,988 | 1,008 | 1,267 | 1,254 | 1,101 | 0,95 | -0,05 | 5,5E-01 |
| P54731 | FAS-associated factor 1 OS=Mus musculus OX=10090 GN=Faf1 PE=1 | 5 | 3 | 3 | 3 | 649 | 73,8 | 4,86 | 2,63 | 3 | 1,287 | 1,188 | 1,683 | 1,019 | 1,141 | 1,255 | 1,138 | 1,276 | 1,08 | 0,08 | 5,5E-01 |
| O89114 | DnaJ homolog subfamily B member 5 OS=Mus musculus OX=10090 | 3 | 1 | 1 | 1 | 348 | 39,1 | 9,04 | 0 | 1 | 0,796 | 1,423 | 1,087 | 0,841 | 1,742 | 1,224 | 1,298 | 0,584 | 0,86 | -0,14 | 5,5E-01 |
| Q8BZZ3 | NEDD4-like E3 ubiquitin-protein ligase WWP1 OS=Mus musculus O | 5 | 4 | 4 | 4 | 918 | 104,6 | 6,38 | 5,48 | 4 | 1,158 | 1,121 | 1,103 | 1,07 | 0,919 | 1,174 | 1,029 | 1,169 | 1,04 | 0,04 | 5,5E-01 |
| Q9CRD0 | OCIA domain-containing protein 1 OS=Mus musculus OX=10090 GN | 17 | 3 | 4 | 3 | 247 | 27,6 | 7,81 | 2,27 | 3 | 1,076 | 1,113 | 0,76 | 0,952 | 0,917 | 1,305 | 0,969 | 1,007 | 0,93 | -0,07 | 5,5E-01 |
| Q924N4 | Solute carrier family 12 member 6 OS=Mus musculus OX=10090 GN | 6 | 6 | 9 | 4 | 1150 | 127,4 | 7,03 | 8,52 | 6 | 1,387 | 1,128 | 1,311 | 1,033 | 1,217 | 1,249 | 1,069 | 1,09 | 1,05 | 0,05 | 5,5E-01 |
| Q8R4G0 | Netrin-G1 OS=Mus musculus OX=10090 GN=Ntng1 PE=1 SV=2 | 2 | 1 | 1 | 1 | 539 | 60,5 | 6,24 | 2,43 | 1 | 1,496 | 1,055 | 1,283 | 1,189 | 1,411 | 1,118 | 1,159 | 1,022 | 1,07 | 0,07 | 5,5E-01 |
| P63280 | SUMO-conjugating enzyme UBC9 OS=Mus musculus OX=10090 GN | 8 | 1 | 1 | 1 | 158 | 18 | 8,66 | 2,36 | 1 | 1,645 | 1,611 | 0,783 | 1,312 | 1,096 | 1,065 | 1,639 | 0,909 | 1,14 | 0,14 | 5,5E-01 |
| Q922Q4 | Pyrroline-5-carboxylate reductase 2 OS=Mus musculus OX=10090 G | 35 | 10 | 10 | 10 | 320 | 33,6 | 7,77 | 18,64 | 10 | 1,583 | 1,101 | 1,205 | 0,929 | 1,051 | 1,302 | 1,066 | 1,014 | 1,09 | 0,09 | 5,5E-01 |
| Q6PGN3 | Serine/threonine-protein kinase DCLK2 OS=Mus musculus OX=1009 | 14 | 9 | 10 | 8 | 756 | 82,9 | 8,44 | 17,02 | 9 | 1,114 | 1,131 | 1,162 | 1,044 | 1,214 | 1,193 | 1,141 | 1,026 | 0,97 | -0,03 | 5,5E-01 |
| P62869 | Elongin-B OS=Mus musculus OX=10090 GN=Elob PE=1 SV=1 | 32 | 5 | 8 | 5 | 118 | 13,2 | 5,01 | 19,24 | 5 | 0,966 | 1,014 | 1,065 | 1,102 | 1,019 | 0,925 | 1,052 | 1,046 | 1,03 | 0,03 | 5,5E-01 |
| Q3UTJ2 | Sorbin and SH3 domain-containing protein 2 OS=Mus musculus OX | 11 | 11 | 13 | 11 | 1180 | 132,3 | 8,53 | 31,47 | 11 | 1,227 | 1,013 | 1,049 | 1,055 | 0,94 | 1,106 | 1,03 | 1,111 | 1,04 | 0,04 | 5,5E-01 |
| Q3TTY5 | Keratin, type II cytoskeletal 2 epidermal OS=Mus musculus OX=100 | 11 | 12 | 21 | 8 | 707 | 70,9 | 8,06 | 40,17 | 12 | 1,136 | 1,747 | 0,875 | 0,685 | 1,45 | 0,88 | 0,596 | 0,774 | 1,20 | 0,20 | 5,5E-01 |
| Q3V038 | Tetratricopeptide repeat protein 9A OS=Mus musculus OX=10090 G | 11 | 2 | 2 | 2 | 219 | 24,3 | 8,85 | 2,12 | 2 | 0,01 | 0,969 | 0,846 | 1,122 | 0,722 | 0,744 | 1,048 | 1,105 | 0,81 | -0,19 | 5,5E-01 |
| P70362 | Ubiquitin recognition factor in ER-associated degradation protein 1 | 12 | 3 | 3 | 3 | 307 | 34,5 | 6,7 | 2,53 | 3 | 1,731 | 1,134 | 1,247 | 1,061 | 1,083 | 1,356 | 0,931 | 1,344 | 1,10 | 0,10 | 5,5E-01 |
| P62889 | 60S ribosomal protein L30 OS=Mus musculus OX=10090 GN=Rpl30 | 16 | 3 | 3 | 3 | 115 | 12,8 | 9,63 | 3,43 | 3 | 1,055 | 0,829 | 0,884 | 1,136 | 1,163 | 0,856 | 1,001 | 1,136 | 0,94 | -0,06 | 5,5E-01 |
| Q6NSR8 | Probable aminopeptidase NPEPL1 OS=Mus musculus OX=10090 GN | 2 | 1 | 1 | 1 | 524 | 55,9 | 6,84 | 0 | 1 | 0,01 | 0,696 | 0,961 | 1,558 | 1,029 | 0,721 | 0,792 | 1,641 | 0,77 | -0,23 | 5,5E-01 |
| P62880 | Guanine nucleotide-binding protein G(I)/G(S)/G(T) subunit beta-2 C | 29 | 10 | 33 | 4 | 340 | 37,3 | 6 | 64,97 | 10 | 1,17 | 0,943 | 1,138 | 1,034 | 1 | 1,128 | 1,053 | 0,941 | 1,04 | 0,04 | 5,5E-01 |
| P53395 | Lipoamide acyltransferase component of branched-chain alpha-ket | 12 | 6 | 7 | 6 | 482 | 53,2 | 8,6 | 14,45 | 6 | 1,122 | 1,063 | 1,102 | 1,055 | 1,027 | 1,092 | 1,018 | 1,128 | 1,02 | 0,02 | 5,6E-01 |
| Q9DB41 | Mitochondrial glutamate carrier 2 OS=Mus musculus OX=10090 GN | 24 | 8 | 13 | 6 | 320 | 34,1 | 9,19 | 28,03 | 8 | 1,126 | 0,983 | 1,069 | 1,022 | 1,043 | 1,093 | 0,916 | 1,027 | 1,03 | 0,03 | 5,6E-01 |
| P21300 | Aldo-keto reductase family 1 member B7 OS=Mus musculus OX=10 | 2 | 1 | 1 | 1 | 316 | 36 | 7,25 | 2,08 | 1 | 1,182 | 1,021 | 0,93 | 1,013 | 1,074 | 1,278 | 1,056 | 0,953 | 0,95 | -0,05 | 5,6E-01 |
| Q3URS9 | Coiled-coil domain-containing protein 51 OS=Mus musculus OX=10 | 7 | 2 | 2 | 2 | 406 | 45,1 | 8,09 | 1,78 | 2 | 1,398 | 1,297 | 0,807 | 1,033 | 0,929 | 1,477 | 1,205 | 1,37 | 0,91 | -0,09 | 5,6E-01 |
| Q60854 | Serpin B6 OS=Mus musculus OX=10090 GN=Serpinb6 PE=1 SV=1 | 20 | 7 | 8 | 7 | 378 | 42,6 | 5,74 | 18,67 | 7 | 1,199 | 1,005 | 1,151 | 1,092 | 1,046 | 1,028 | 1,171 | 1,071 | 1,03 | 0,03 | 5,6E-01 |
| Q99J99 | 3-mercaptopyruvate sulfurtransferase OS=Mus musculus OX=1009 | 18 | 4 | 6 | 4 | 297 | 33,1 | 6,47 | 15,61 | 4 | 0,983 | 1,057 | 0,892 | 0,968 | 1,024 | 0,921 | 0,953 | 0,892 | 1,03 | 0,03 | 5,6E-01 |
| P63094 | Guanine nucleotide-binding protein G(s) subunit alpha isoforms sho | 18 | 6 | 8 | 5 | 394 | 45,6 | 5,96 | 18,62 | 6 | 1,009 | 1,009 | 1,061 | 0,932 | 0,966 | 1,201 | 0,939 | 1,067 | 0,96 | -0,04 | 5,6E-01 |
| Q6VNS1 | NT-3 growth factor receptor OS=Mus musculus OX=10090 GN=Ntrk | 5 | 5 | 5 | 2 | 825 | 92,7 | 6,67 | 8,37 | 5 | 1,152 | 1,02 | 1,06 | 1,064 | 1,225 | 0,828 | 0,988 | 1,04 | 1,05 | 0,05 | 5,6E-01 |
| O35465 | Peptidyl-prolyl cis-trans isomerase FKBP8 OS=Mus musculus OX=10 | 6 | 2 | 4 | 2 | 402 | 43,5 | 5,16 | 4,98 | 2 | 1,193 | 0,934 | 0,772 | 0,728 | 0,937 | 0,952 | 1,006 | 0,996 | 0,93 | -0,07 | 5,6E-01 |
| P70404 | Isocitrate dehydrogenase [NAD] subunit gamma 1, mitochondrial C | 25 | 7 | 19 | 7 | 393 | 42,8 | 9,01 | 42,16 | 7 | 1,29 | 1,103 | 1,088 | 0,976 | 1,04 | 1,138 | 1,041 | 1,067 | 1,04 | 0,04 | 5,6E-01 |
| Q60625 | Intercellular adhesion molecule 5 OS=Mus musculus OX=10090 GN | 13 | 10 | 13 | 10 | 917 | 96,9 | 6,32 | 22,61 | 10 | 1,231 | 0,982 | 1,127 | 1,002 | 1,14 | 1,132 | 1,153 | 1,068 | 0,97 | -0,03 | 5,6E-01 |
| Q8R313 | Exocyst complex component 6 OS=Mus musculus OX=10090 GN=Ex | 4 | 3 | 3 | 3 | 802 | 93 | 6,15 | 2,21 | 3 | 1,166 | 1,01 | 1,011 | 0,681 | 0,793 | 1,138 | 0,667 | 0,917 | 1,10 | 0,10 | 5,6E-01 |
| Q9Z2I0 | Mitochondrial proton/calcium exchanger protein OS=Mus musculu | 18 | 12 | 18 | 12 | 738 | 82,9 | 6,52 | 46,36 | 12 | 0,974 | 1,007 | 0,967 | 0,973 | 0,951 | 0,957 | 1,01 | 1,08 | 0,98 | -0,02 | 5,6E-01 |
| Q91YY2 | Beta-1,4-galactosyltransferase 3 OS=Mus musculus OX=10090 GN= | 3 | 1 | 1 | 1 | 395 | 44,1 | 9,25 | 0 | 1 | 1,201 | 1,099 | 0,94 | 0,908 | 0,952 | 1,213 | 0,773 | 0,929 | 1,07 | 0,07 | 5,6E-01 |
| Q9D024 | Coiled-coil domain-containing protein 47 OS=Mus musculus OX=10 | 2 | 1 | 1 | 1 | 483 | 55,8 | 4,84 | 3,54 | 1 | 1,482 | 1,61 | 1,26 | 0,903 | 1,229 | 1,493 | 1,225 | 0,783 | 1,11 | 0,11 | 5,6E-01 |
| Q91YS8 | Calcium/calmodulin-dependent protein kinase type 1 OS=Mus mus | 11 | 3 | 3 | 2 | 374 | 41,6 | 5,35 | 9,62 | 3 | 1,186 | 1,049 | 0,955 | 1,216 | 1,155 | 1,052 | 1,151 | 1,22 | 0,96 | -0,04 | 5,6E-01 |
| Q61335 | B-cell receptor-associated protein 31 OS=Mus musculus OX=10090 | 21 | 4 | 4 | 4 | 245 | 27,9 | 8,7 | 6,03 | 4 | 1,042 | 1,201 | 1,142 | 0,809 | 0,851 | 1,148 | 0,99 | 0,944 | 1,07 | 0,07 | 5,6E-01 |
| P27601 | Guanine nucleotide-binding protein subunit alpha-13 OS=Mus mus | 23 | 8 | 18 | 7 | 377 | 44 | 8,21 | 33,55 | 8 | 1,106 | 1,125 | 1,032 | 1,036 | 1,058 | 1,153 | 1,054 | 1,117 | 0,98 | -0,02 | 5,6E-01 |
| Q9ERT9 | Protein phosphatase 1 regulatory subunit 1A OS=Mus musculus OX | 32 | 3 | 5 | 3 | 171 | 18,7 | 5,25 | 6,98 | 3 | 1,126 | 0,96 | 1,073 | 0,937 | 0,75 | 1,17 | 0,961 | 0,977 | 1,06 | 0,06 | 5,6E-01 |
| Q9WTM5 | RuvB-like 2 OS=Mus musculus OX=10090 GN=Ruvbl2 PE=1 SV=3 | 17 | 8 | 10 | 8 | 463 | 51,1 | 5,64 | 14,07 | 8 | 1,153 | 1,024 | 1,038 | 0,91 | 0,986 | 1,009 | 1,035 | 0,968 | 1,03 | 0,03 | 5,6E-01 |
| Q71RI9 | Kynurenine--oxoglutarate transaminase 3 OS=Mus musculus OX=10 | 8 | 2 | 2 | 2 | 455 | 51,1 | 8,37 | 3,13 | 2 | 1,254 | 1,244 | 1,046 | 0,99 | 0,999 | 1,123 | 1,1 | 1,13 | 1,04 | 0,04 | 5,6E-01 |
| Q9Z0P4 | Paralemmin-1 OS=Mus musculus OX=10090 GN=Palm PE=1 SV=1 | 39 | 13 | 21 | 12 | 383 | 41,6 | 4,84 | 48,09 | 13 | 0,906 | 1,042 | 1,025 | 1,013 | 0,943 | 1,039 | 1,064 | 1,04 | 0,98 | -0,02 | 5,6E-01 |
| Q8BTY2-2 | Isoform 2 of Sodium bicarbonate cotransporter 3 OS=Mus musculu | 3 | 3 | 5 | 2 | 936 | 104,6 | 6,38 | 9,38 | 3 | 0,842 | 0,714 | 1,393 | 1,293 | 1,232 | 0,907 | 0,634 | 0,963 | 1,14 | 0,14 | 5,6E-01 |
| P80317 | T-complex protein 1 subunit zeta OS=Mus musculus OX=10090 GN= | 24 | 14 | 16 | 10 | 531 | 58 | 7,08 | 31,19 | 14 | 0,986 | 0,944 | 1,048 | 0,937 | 0,949 | 1,006 | 1,015 | 1,019 | 0,98 | -0,02 | 5,6E-01 |
| P52623 | Uridine-cytidine kinase 1 OS=Mus musculus OX=10090 GN=Uck1 PE | 9 | 2 | 2 | 2 | 277 | 31 | 7,88 | 5,53 | 2 | 1,587 | 1,452 | 1,035 | 0,985 | 0,979 | 1,39 | 1,034 | 1,224 | 1,09 | 0,09 | 5,6E-01 |
| B2RW38 | Cilia- and flagella-associated protein 58 OS=Mus musculus OX=1009 | 1 | 1 | 1 | 1 | 873 | 103,5 | 8,16 | 0 | 1 | 1,031 | 0,768 | 0,756 | 0,819 | 0,854 | 0,612 | 0,809 | 0,883 | 1,07 | 0,07 | 5,6E-01 |
| Q6IRU2 | Tropomyosin alpha-4 chain OS=Mus musculus OX=10090 GN=Tpm4 | 13 | 4 | 6 | 3 | 248 | 28,5 | 4,68 | 16 | 4 | 1,101 | 1,087 | 1,274 | 1,602 | 1,181 | 1,172 | 1,127 | 1,281 | 1,06 | 0,06 | 5,6E-01 |
| Q60870 | Receptor expression-enhancing protein 5 OS=Mus musculus OX=10 | 4 | 1 | 2 | 1 | 185 | 21 | 8,38 | 4,05 | 1 | 0,921 | 0,975 | 0,97 | 0,853 | 1,029 | 0,815 | 1,189 | 0,897 | 0,95 | -0,05 | 5,6E-01 |
| P14148 | 60S ribosomal protein L7 OS=Mus musculus OX=10090 GN=Rpl7 PE | 29 | 8 | 8 | 8 | 270 | 31,4 | 10,89 | 17,69 | 8 | 1,075 | 0,895 | 0,89 | 0,911 | 1,022 | 1,039 | 0,891 | 0,955 | 0,97 | -0,03 | 5,6E-01 |
| G3X9K3 | Brefeldin A-inhibited guanine nucleotide-exchange protein 1 OS=M | 3 | 5 | 5 | 3 | 1846 | 208,4 | 5,86 | 8,14 | 5 | 1,308 | 1,27 | 1,232 | 0,959 | 0,9 | 1,37 | 0,936 | 1,225 | 1,08 | 0,08 | 5,6E-01 |
| P22933 | Gamma-aminobutyric acid receptor subunit delta OS=Mus musculu | 3 | 1 | 1 | 1 | 449 | 50,5 | 7,85 | 3,45 | 1 | 1,093 | 1,12 | 1,013 | 0,982 | 0,859 | 1,123 | 0,916 | 1,124 | 1,05 | 0,05 | 5,7E-01 |
| P42932 | T-complex protein 1 subunit theta OS=Mus musculus OX=10090 GN | 34 | 18 | 27 | 18 | 548 | 59,5 | 5,62 | 70,46 | 18 | 1,078 | 1,003 | 1,042 | 1,012 | 0,962 | 1,103 | 1,008 | 0,974 | 1,02 | 0,02 | 5,7E-01 |
| Q9QXS6 | Drebrin OS=Mus musculus OX=10090 GN=Dbn1 PE=1 SV=4 | 19 | 11 | 18 | 11 | 706 | 77,2 | 4,49 | 40,01 | 11 | 0,842 | 0,888 | 0,963 | 0,946 | 1,009 | 0,84 | 0,949 | 0,95 | 0,97 | -0,03 | 5,7E-01 |
| P24638 | Lysosomal acid phosphatase OS=Mus musculus OX=10090 GN=Acp | 3 | 1 | 1 | 1 | 423 | 48,5 | 7,02 | 2,3 | 1 | 0,917 | 1,036 | 0,705 | 0,957 | 0,841 | 0,881 | 1,019 | 0,602 | 1,08 | 0,08 | 5,7E-01 |
| Q8K003 | Translation machinery-associated protein 7 OS=Mus musculus OX= | 14 | 1 | 1 | 1 | 64 | 7,1 | 9,99 | 2,12 | 1 | 1,326 | 1,197 | 1,063 | 1,314 | 1,099 | 1,372 | 1,14 | 1,064 | 1,05 | 0,05 | 5,7E-01 |
| P35980 | 60S ribosomal protein L18 OS=Mus musculus OX=10090 GN=Rpl18 | 27 | 6 | 8 | 6 | 188 | 21,6 | 11,78 | 8,45 | 6 | 0,875 | 0,942 | 1,02 | 0,999 | 0,97 | 1,05 | 1,033 | 0,898 | 0,97 | -0,03 | 5,7E-01 |
| P70302 | Stromal interaction molecule 1 OS=Mus musculus OX=10090 GN=S | 9 | 6 | 6 | 6 | 685 | 77,5 | 6,54 | 9,31 | 6 | 1,258 | 0,833 | 0,92 | 1,018 | 0,922 | 0,787 | 1,126 | 0,914 | 1,07 | 0,07 | 5,7E-01 |
| Q7TPB0 | Phospholipid phosphatase-related protein type 3 OS=Mus musculu | 6 | 3 | 4 | 3 | 716 | 76,6 | 6,01 | 6,48 | 3 | 0,764 | 1,057 | 1,137 | 0,766 | 0,797 | 1,004 | 1,206 | 1,028 | 0,92 | -0,08 | 5,7E-01 |
| Q9DCD0 | 6-phosphogluconate dehydrogenase, decarboxylating OS=Mus mus | 8 | 4 | 4 | 4 | 483 | 53,2 | 7,23 | 7,71 | 4 | 1,312 | 1,127 | 1,13 | 0,905 | 0,857 | 1,349 | 0,981 | 0,957 | 1,08 | 0,08 | 5,7E-01 |
| Q6A0A9 | Constitutive coactivator of PPAR-gamma-like protein 1 OS=Mus mu | 5 | 4 | 7 | 4 | 1112 | 121,6 | 8,92 | 15,28 | 4 | 0,795 | 1,032 | 0,895 | 1,373 | 0,988 | 0,886 | 0,99 | 0,919 | 1,08 | 0,08 | 5,7E-01 |

| | | | | | | | | | | | | | | | | | | | | | |
|---|---|---|---|---|---|---|---|---|---|---|---|---|---|---|---|---|---|---|---|---|---|
| P16054 | Protein kinase C epsilon type OS=Mus musculus OX=10090 GN=Prk | 15 | 10 | 12 | 10 | 737 | 83,5 | 7,03 | 24,08 | 10 | 0,934 | 0,988 | 1,022 | 1,07 | 0,999 | 1,078 | 1,03 | 0,991 | 0,98 | -0,02 | 5,7E-01 |
| Q68FL6 | Methionine--tRNA ligase, cytoplasmic OS=Mus musculus OX=10090 | 4 | 4 | 5 | 4 | 902 | 101,4 | 7,14 | 8,93 | 4 | 1,213 | 1,126 | 0,987 | 0,946 | 1,09 | 1,303 | 1,005 | 1,088 | 0,95 | -0,05 | 5,7E-01 |
| Q91XL9 | Oxysterol-binding protein-related protein 1 OS=Mus musculus OX= | 7 | 6 | 9 | 5 | 950 | 107,7 | 6,44 | 20,17 | 6 | 1,036 | 1,045 | 0,999 | 1,076 | 1,04 | 0,969 | 0,967 | 1,096 | 1,02 | 0,02 | 5,7E-01 |
| Q68EF8 | Rap guanine nucleotide exchange factor-like 1 OS=Mus musculus O | 1 | 1 | 1 | 1 | 662 | 73,6 | 6,21 | 0 | 1 | 0,828 | 0,898 | 0,855 | 0,864 | 1,053 | 0,997 | 0,53 | 1,224 | 0,91 | -0,09 | 5,7E-01 |
| Q9JM52 | Misshapen-like kinase 1 OS=Mus musculus OX=10090 GN=Mink1 P | 11 | 14 | 14 | 9 | 1308 | 147,2 | 7,44 | 28,41 | 14 | 0,861 | 0,83 | 1,182 | 1,284 | 1,167 | 1,023 | 1,302 | 0,989 | 0,93 | -0,07 | 5,7E-01 |
| Q9CWX2 | Complex I intermediate-associated protein 30, mitochondrial OS=M | 14 | 5 | 5 | 5 | 328 | 37,8 | 8,6 | 9,97 | 5 | 0,977 | 1,052 | 0,987 | 0,98 | 1,055 | 1,057 | 0,939 | 1,024 | 0,98 | -0,02 | 5,7E-01 |
| P07146 | Anionic trypsin-2 OS=Mus musculus OX=10090 GN=Prss2 PE=1 SV= | 8 | 1 | 23 | 1 | 246 | 26,2 | 4,63 | 48,17 | 1 | 2,186 | 1,273 | 1,446 | 0,819 | 0,72 | 1,902 | 0,939 | 1,242 | 1,19 | 0,19 | 5,7E-01 |
| P61460 | GATOR complex protein DEPDC5 OS=Mus musculus OX=10090 GN= | 1 | 2 | 2 | 2 | 1591 | 180,3 | 6,74 | 4,42 | 2 | 1,231 | 1,032 | 1,116 | 0,961 | 0,919 | 1,044 | 1,009 | 1,178 | 1,05 | 0,05 | 5,7E-01 |
| P42859 | Huntingtin OS=Mus musculus OX=10090 GN=Htt PE=1 SV=2 | 5 | 11 | 11 | 11 | 3119 | 344,5 | 6,29 | 29,78 | 11 | 0,946 | 1,093 | 0,956 | 0,995 | 0,985 | 0,86 | 0,988 | 1,036 | 1,03 | 0,03 | 5,7E-01 |
| Q9CQF4 | Mitochondrial transcription rescue factor 1 OS=Mus musculus OX= | 4 | 1 | 1 | 1 | 240 | 27,8 | 9,36 | 0 | 1 | 0,595 | 1,288 | 0,941 | 0,969 | 0,864 | 1,161 | 1,119 | 1,024 | 0,91 | -0,09 | 5,7E-01 |
| Q6PIE5 | Sodium/potassium-transporting ATPase subunit alpha-2 OS=Mus m | 48 | 53 | 178 | 32 | 1020 | 112,1 | 5,55 | 374,01 | 53 | 1,04 | 0,944 | 1,008 | 0,966 | 1,032 | 0,972 | 0,942 | 0,939 | 1,02 | 0,02 | 5,7E-01 |
| Q9D8S3 | ADP-ribosylation factor GTPase-activating protein 3 OS=Mus muscu | 2 | 1 | 1 | 1 | 523 | 57,4 | 8,47 | 2,74 | 1 | 1,183 | 0,705 | 0,572 | 0,786 | 0,694 | 0,691 | 0,838 | 0,696 | 1,11 | 0,11 | 5,7E-01 |
| Q9WTP7 | GTP:AMP phosphotransferase AK3, mitochondrial OS=Mus musculu | 44 | 9 | 12 | 9 | 227 | 25,4 | 8,84 | 27,34 | 9 | 1,057 | 1,024 | 1,016 | 0,988 | 0,991 | 1,057 | 1,048 | 1,038 | 0,99 | -0,01 | 5,7E-01 |
| Q80YA7 | Dipeptidyl peptidase 8 OS=Mus musculus OX=10090 GN=Dpp8 PE= | 5 | 3 | 3 | 3 | 892 | 102,1 | 5,81 | 7,62 | 3 | 1,092 | 1,071 | 1,103 | 0,991 | 1,006 | 1,111 | 0,883 | 1,113 | 1,04 | 0,04 | 5,7E-01 |
| O54724 | Caveolae-associated protein 1 OS=Mus musculus OX=10090 GN=Ca | 3 | 1 | 1 | 1 | 392 | 43,9 | 5,52 | 2,02 | 1 | 0,532 | 0,947 | 1,189 | 1,022 | 1,04 | 1,066 | 1,042 | 0,89 | 0,91 | -0,09 | 5,7E-01 |
| Q63844 | Mitogen-activated protein kinase 3 OS=Mus musculus OX=10090 G | 30 | 11 | 13 | 7 | 380 | 43 | 6,61 | 15,92 | 11 | 1,141 | 1,003 | 0,985 | 0,944 | 0,877 | 1,173 | 0,904 | 0,926 | 1,05 | 0,05 | 5,7E-01 |
| Q2TPA8 | Hydroxysteroid dehydrogenase-like protein 2 OS=Mus musculus OX | 2 | 1 | 1 | 1 | 490 | 54,2 | 6,74 | 2,94 | 1 | 1,194 | 0,743 | 0,979 | 0,888 | 0,831 | 1,006 | 1,045 | 0,584 | 1,10 | 0,10 | 5,7E-01 |
| Q60875 | Rho guanine nucleotide exchange factor 2 OS=Mus musculus OX=1 | 19 | 17 | 17 | 17 | 985 | 111,9 | 7,25 | 28,26 | 17 | 1,344 | 1,191 | 1,043 | 0,991 | 0,905 | 1,36 | 0,958 | 1,037 | 1,07 | 0,07 | 5,7E-01 |
| P35293 | Ras-related protein Rab-18 OS=Mus musculus OX=10090 GN=Rab1 | 43 | 8 | 11 | 8 | 206 | 23 | 5,36 | 28,23 | 8 | 1,065 | 1,128 | 1,04 | 1,011 | 0,988 | 1,078 | 1,052 | 1,051 | 1,02 | 0,02 | 5,7E-01 |
| Q6NSW3 | A-kinase anchor protein SPHKAP OS=Mus musculus OX=10090 GN= | 1 | 1 | 1 | 1 | 1687 | 185 | 5,06 | 2,21 | 1 | 0,643 | 0,683 | 1,14 | 0,905 | 0,75 | 1,077 | 1,023 | 0,849 | 0,91 | -0,09 | 5,7E-01 |
| P19157 | Glutathione S-transferase P 1 OS=Mus musculus OX=10090 GN=Gst | 31 | 5 | 15 | 5 | 210 | 23,6 | 7,87 | 41,15 | 5 | 1,352 | 1,178 | 1,199 | 0,992 | 0,921 | 1,326 | 1,06 | 1,146 | 1,06 | 0,06 | 5,7E-01 |
| P49722 | Proteasome subunit alpha type-2 OS=Mus musculus OX=10090 GN | 17 | 4 | 5 | 4 | 234 | 25,9 | 7,43 | 8,47 | 4 | 1,076 | 1,202 | 1,037 | 1,001 | 1,119 | 1,296 | 1,046 | 1,033 | 0,96 | -0,04 | 5,7E-01 |
| Q5RJH3 | Cadherin-12 OS=Mus musculus OX=10090 GN=Cdh12 PE=2 SV=1 | 1 | 1 | 1 | 1 | 794 | 88,4 | 4,83 | 2,68 | 1 | 1,186 | 1,274 | 1,071 | 0,96 | 1,105 | 1,314 | 0,767 | 0,988 | 1,08 | 0,08 | 5,7E-01 |
| P50396 | Rab GDP dissociation inhibitor alpha OS=Mus musculus OX=10090 | 38 | 20 | 34 | 17 | 447 | 50,5 | 5,08 | 70,75 | 20 | 0,981 | 1,001 | 0,985 | 1,012 | 0,924 | 1,069 | 1,063 | 1,005 | 0,98 | -0,02 | 5,7E-01 |
| P61963 | DDB1- and CUL4-associated factor 7 OS=Mus musculus OX=10090 | 11 | 3 | 3 | 3 | 342 | 38,9 | 5,52 | 8,75 | 3 | 1,481 | 1,285 | 1,245 | 1,188 | 1,095 | 1,322 | 1,177 | 1,386 | 1,04 | 0,04 | 5,7E-01 |
| Q8JZM0 | Dimethyladenosine transferase 1, mitochondrial OS=Mus musculus | 2 | 1 | 1 | 1 | 345 | 38,9 | 9,47 | 0 | 1 | 0,758 | 0,804 | 1,127 | 0,885 | 0,849 | 0,654 | 0,948 | 0,877 | 1,07 | 0,07 | 5,7E-01 |
| Q9ESW4 | Acylglycerol kinase, mitochondrial OS=Mus musculus OX=10090 GN | 25 | 10 | 11 | 10 | 421 | 46,9 | 8,4 | 22,91 | 10 | 0,899 | 0,971 | 0,935 | 0,941 | 0,952 | 0,962 | 0,921 | 0,952 | 0,99 | -0,01 | 5,7E-01 |
| P06537 | Glucocorticoid receptor OS=Mus musculus OX=10090 GN=Nr3c1 PE | 1 | 1 | 1 | 1 | 783 | 86 | 6,35 | 1,8 | 1 | 1,54 | 1,116 | 1,522 | 1,207 | 0,931 | 1,647 | 0,985 | 1,348 | 1,10 | 0,10 | 5,7E-01 |
| Q3TY65 | Islet cell autoantigen 1-like protein OS=Mus musculus OX=10090 G | 5 | 2 | 2 | 2 | 431 | 48,1 | 4,91 | 6,07 | 2 | 1,192 | 0,958 | 0,96 | 1,106 | 0,763 | 1,118 | 0,999 | 1,099 | 1,06 | 0,06 | 5,7E-01 |
| Q9DC16 | Endoplasmic reticulum-Golgi intermediate compartment protein 1 | 16 | 3 | 3 | 3 | 290 | 32,5 | 7,06 | 9,5 | 3 | 1,131 | 1,237 | 1,009 | 0,976 | 0,934 | 1,271 | 0,909 | 0,996 | 1,06 | 0,06 | 5,7E-01 |
| Q9CX56 | 26S proteasome non-ATPase regulatory subunit 8 OS=Mus musculu | 10 | 4 | 4 | 4 | 353 | 39,9 | 9,58 | 7,27 | 4 | 0,997 | 1,121 | 1,087 | 1,105 | 1,069 | 1,145 | 0,994 | 0,994 | 1,03 | 0,03 | 5,7E-01 |
| P62075 | Mitochondrial import inner membrane translocase subunit Tim13 | 12 | 1 | 3 | 1 | 95 | 10,5 | 8,18 | 4,58 | 1 | 0,605 | 0,992 | 1,809 | 1,333 | 1,221 | 1,2 | 1,897 | 1,158 | 0,87 | -0,13 | 5,7E-01 |
| P70441 | Na(+)/H(+) exchange regulatory cofactor NHE-RF1 OS=Mus musculu | 39 | 12 | 19 | 12 | 355 | 38,6 | 5,9 | 31,79 | 12 | 0,883 | 0,904 | 1,013 | 0,99 | 0,98 | 0,86 | 1,012 | 1,066 | 0,97 | -0,03 | 5,7E-01 |
| P51660 | Peroxisomal multifunctional enzyme type 2 OS=Mus musculus OX= | 21 | 13 | 16 | 13 | 735 | 79,4 | 8,57 | 33,74 | 13 | 1,147 | 1,107 | 1,046 | 1,008 | 1,033 | 1,061 | 1,041 | 1,093 | 1,02 | 0,02 | 5,8E-01 |
| Q03137 | Ephrin type-A receptor 4 OS=Mus musculus OX=10090 GN=Epha4 | 14 | 13 | 17 | 11 | 986 | 109,7 | 6,51 | 27,23 | 13 | 0,916 | 0,94 | 0,958 | 0,937 | 1,035 | 0,934 | 0,903 | 0,949 | 0,98 | -0,02 | 5,8E-01 |
| Q99LF4 | tRNA-splicing ligase RtcB homolog OS=Mus musculus OX=10090 GN | 11 | 5 | 6 | 5 | 505 | 55,2 | 7,23 | 9,12 | 5 | 1,115 | 1,018 | 1,127 | 0,982 | 0,93 | 1,141 | 0,995 | 1,041 | 1,03 | 0,03 | 5,8E-01 |
| Q91ZX7 | Prolow-density lipoprotein receptor-related protein 1 OS=Mus mus | 7 | 28 | 31 | 28 | 4545 | 504,4 | 5,36 | 68,34 | 28 | 1,144 | 0,98 | 0,995 | 0,979 | 1,049 | 1,005 | 0,942 | 0,994 | 1,03 | 0,03 | 5,8E-01 |
| P24270 | Catalase OS=Mus musculus OX=10090 GN=Cat PE=1 SV=4 | 16 | 8 | 8 | 8 | 527 | 59,8 | 7,88 | 15,23 | 8 | 1,222 | 1,1 | 1,019 | 1,04 | 1,143 | 1,129 | 1,026 | 1,229 | 0,97 | -0,03 | 5,8E-01 |
| Q3TYX3 | SET and MYND domain-containing protein 5 OS=Mus musculus OX= | 5 | 1 | 1 | 1 | 416 | 47,1 | 5,21 | 2,6 | 1 | 1,203 | 0,905 | 0,84 | 1,137 | 0,918 | 1,38 | 1,012 | 1,089 | 0,93 | -0,07 | 5,8E-01 |
| P06745 | Glucose-6-phosphate isomerase OS=Mus musculus OX=10090 GN= | 30 | 20 | 38 | 20 | 558 | 62,7 | 8,13 | 79,16 | 20 | 0,842 | 0,929 | 0,996 | 1,004 | 0,964 | 0,938 | 1,002 | 0,961 | 0,98 | -0,02 | 5,8E-01 |
| Q920Q4 | Vacuolar protein sorting-associated protein 16 homolog OS=Mus m | 4 | 3 | 4 | 3 | 839 | 94,9 | 7,01 | 2,18 | 3 | 0,877 | 1,288 | 1,07 | 1,238 | 1,18 | 0,955 | 1,211 | 1,452 | 0,93 | -0,07 | 5,8E-01 |
| Q9D6F9 | Tubulin beta-4A chain OS=Mus musculus OX=10090 GN=Tubb4a PE | 54 | 23 | 139 | 5 | 444 | 49,6 | 4,88 | 270,78 | 23 | 1,051 | 0,971 | 0,98 | 1,108 | 1,002 | 1,045 | 1,093 | 1,058 | 0,98 | -0,02 | 5,8E-01 |
| Q9CQD4 | Charged multivesicular body protein 1b-2 OS=Mus musculus OX=1 | 12 | 3 | 3 | 3 | 199 | 22,1 | 8,1 | 7,77 | 3 | 1,115 | 0,918 | 0,999 | 0,95 | 0,937 | 1,085 | 1,027 | 1,06 | 0,97 | -0,03 | 5,8E-01 |
| Q8VCM5 | Mitochondrial ubiquitin ligase activator of NFKB 1 OS=Mus musculu | 2 | 1 | 1 | 1 | 352 | 39,8 | 7,96 | 0 | 1 | 1,012 | 0,968 | 0,887 | 0,703 | 0,99 | 0,944 | 0,823 | 1 | 0,95 | -0,05 | 5,8E-01 |
| Q921W0 | Charged multivesicular body protein 1a OS=Mus musculus OX=100 | 9 | 2 | 2 | 2 | 196 | 21,6 | 8,06 | 1,88 | 2 | 0,696 | 1,131 | 0,809 | 0,96 | 0,851 | 1,202 | 1,027 | 0,82 | 0,92 | -0,08 | 5,8E-01 |
| Q7M759 | Alpha/beta hydrolase domain-containing protein 17B OS=Mus mus | 10 | 3 | 3 | 3 | 288 | 32,2 | 6,29 | 7,21 | 3 | 0,716 | 0,953 | 0,977 | 1,106 | 1,13 | 0,964 | 0,958 | 0,92 | 0,94 | -0,06 | 5,8E-01 |
| Q68ED2 | Metabotropic glutamate receptor 7 OS=Mus musculus OX=10090 G | 10 | 7 | 9 | 7 | 915 | 102,2 | 7,88 | 17,87 | 7 | 1,205 | 1,385 | 1,33 | 0,953 | 1,048 | 1,336 | 1,064 | 1,152 | 1,06 | 0,06 | 5,8E-01 |
| Q8CHT1 | Ephexin-1 OS=Mus musculus OX=10090 GN=Ngef PE=1 SV=1 | 12 | 10 | 11 | 10 | 710 | 82,1 | 5,99 | 22,01 | 10 | 1,23 | 1,095 | 1,076 | 0,961 | 1,014 | 1,343 | 1,103 | 1,111 | 0,95 | -0,05 | 5,8E-01 |
| Q8C166 | Copine-1 OS=Mus musculus OX=10090 GN=Cpne1 PE=1 SV=1 | 8 | 5 | 5 | 5 | 536 | 58,8 | 5,66 | 10,19 | 5 | 0,947 | 0,928 | 0,908 | 1,117 | 1,039 | 0,942 | 1,007 | 1,036 | 0,97 | -0,03 | 5,8E-01 |
| Q8K2I1 | Protein farnesyltransferase subunit beta OS=Mus musculus OX=100 | 4 | 1 | 1 | 1 | 437 | 48,8 | 5,8 | 2,72 | 1 | 1,197 | 1,047 | 1,113 | 1,097 | 1,102 | 1,322 | 0,997 | 1,212 | 0,96 | -0,04 | 5,8E-01 |
| Q8K0C9 | GDP-mannose 4,6 dehydratase OS=Mus musculus OX=10090 GN=G | 8 | 2 | 2 | 2 | 372 | 42 | 7,03 | 0 | 2 | 1,483 | 0,884 | 1,165 | 1,743 | 1,444 | 1,195 | 1,402 | 1,744 | 0,91 | -0,09 | 5,8E-01 |
| Q3UHB1 | 5'-nucleotidase domain-containing protein 3 OS=Mus musculus OX | 17 | 10 | 11 | 10 | 546 | 63,1 | 8,56 | 22,21 | 10 | 0,899 | 0,933 | 0,955 | 1,128 | 1,044 | 0,912 | 1,064 | 1,039 | 0,96 | -0,04 | 5,8E-01 |
| O35295 | Transcriptional activator protein Pur-beta OS=Mus musculus OX=1 | 9 | 2 | 3 | 2 | 324 | 33,9 | 5,43 | 9,42 | 2 | 1,012 | 1,143 | 0,924 | 0,955 | 1,131 | 0,997 | 1,022 | 1,017 | 0,97 | -0,03 | 5,8E-01 |
| Q99K51 | Plastin-3 OS=Mus musculus OX=10090 GN=Pls3 PE=1 SV=3 | 20 | 12 | 12 | 10 | 630 | 70,7 | 5,62 | 20,5 | 12 | 1,056 | 1,115 | 0,927 | 1,236 | 1,001 | 1,065 | 1,034 | 1,079 | 1,04 | 0,04 | 5,8E-01 |
| P14231 | Sodium/potassium-transporting ATPase subunit beta-2 OS=Mus mu | 38 | 10 | 17 | 10 | 290 | 33,3 | 8,31 | 44,92 | 10 | 1,023 | 0,955 | 0,936 | 0,978 | 1,064 | 0,904 | 1,012 | 1,001 | 0,98 | -0,02 | 5,8E-01 |
| Q5SYD0 | Unconventional myosin-Id OS=Mus musculus OX=10090 GN=Myo1 | 8 | 7 | 7 | 7 | 1006 | 116 | 9,41 | 15,95 | 7 | 0,914 | 1,011 | 0,957 | 1,046 | 1,073 | 1,122 | 0,912 | 0,954 | 0,97 | -0,03 | 5,8E-01 |
| Q9EP72 | ER membrane protein complex subunit 7 OS=Mus musculus OX=10 | 5 | 1 | 2 | 1 | 241 | 26,3 | 9,23 | 2,56 | 1 | 0,01 | 1,385 | 0,896 | 0,865 | 0,919 | 0,91 | 1,036 | 0,958 | 0,83 | -0,17 | 5,8E-01 |
| Q6PAR5 | GTPase-activating protein and VPS9 domain-containing protein 1 O | 6 | 6 | 6 | 6 | 1458 | 162,3 | 5,19 | 16,07 | 6 | 1,35 | 1,27 | 1,134 | 1,001 | 0,849 | 1,378 | 1,02 | 1,19 | 1,07 | 0,07 | 5,8E-01 |
| Q9D1G1 | Ras-related protein Rab-1B OS=Mus musculus OX=10090 GN=Rab1 | 39 | 8 | 15 | 4 | 201 | 22,2 | 5,73 | 43,37 | 8 | 0,789 | 0,972 | 1,099 | 1,064 | 0,998 | 0,956 | 1,034 | 1,115 | 0,96 | -0,04 | 5,8E-01 |
| Q6ZPJ0 | Testis-expressed protein 2 OS=Mus musculus OX=10090 GN=Tex2 | 1 | 2 | 2 | 2 | 1128 | 125,1 | 5,8 | 4,22 | 2 | 1,388 | 1,259 | 1,084 | 1,112 | 1,052 | 1,257 | 1,136 | 1,205 | 1,04 | 0,04 | 5,8E-01 |
| Q07076 | Annexin A7 OS=Mus musculus OX=10090 GN=Anxa7 PE=1 SV=2 | 24 | 11 | 15 | 11 | 463 | 49,9 | 6,18 | 32,01 | 11 | 1,116 | 1,001 | 1,039 | 0,961 | 0,933 | 1,066 | 0,996 | 1,022 | 1,02 | 0,02 | 5,8E-01 |
| Q8K400 | Syntaxin-binding protein 5 OS=Mus musculus OX=10090 GN=Stxbp | 12 | 10 | 12 | 10 | 1152 | 127,6 | 7,21 | 27,41 | 10 | 1,107 | 1,086 | 1,124 | 1,164 | 0,967 | 1,22 | 1,039 | 1,123 | 1,03 | 0,03 | 5,8E-01 |

| | | | | | | | | | | | | | | | | | | | | | |
|---|---|---|---|---|---|---|---|---|---|---|---|---|---|---|---|---|---|---|---|---|---|
| P17156 | Heat shock-related 70 kDa protein 2 OS=Mus musculus OX=10090 ( | 27 | 18 | 48 | 7 | 633 | 69,6 | 5,67 | 91,63 | 18 | 1,043 | 0,98 | 1,029 | 0,972 | 0,903 | 1,079 | 1,064 | 1,087 | 0,97 | -0,03 | 5,8E-01 |
| Q6ZQ38 | Cullin-associated NEDD8-dissociated protein 1 OS=Mus musculus O | 18 | 22 | 28 | 22 | 1230 | 136,2 | 5,78 | 46,67 | 22 | 1,209 | 1,025 | 1,166 | 0,964 | 0,988 | 1,209 | 0,987 | 0,996 | 1,04 | 0,04 | 5,8E-01 |
| Q9QXT0 | Protein canopy homolog 2 OS=Mus musculus OX=10090 GN=Cnpy2 | 25 | 4 | 4 | 4 | 182 | 20,8 | 5,07 | 4,96 | 4 | 1,22 | 1,096 | 0,974 | 0,96 | 0,944 | 1,122 | 0,966 | 1,049 | 1,04 | 0,04 | 5,8E-01 |
| P61329 | Fibroblast growth factor 12 OS=Mus musculus OX=10090 GN=Fgf12 | 4 | 1 | 2 | 1 | 243 | 27,4 | 9,96 | 0 | 1 | 0,01 | 1,228 | 1,586 | 1,54 | 0,754 | 2,222 | 1,521 | 1,002 | 0,79 | -0,21 | 5,8E-01 |
| P62835 | Ras-related protein Rap-1A OS=Mus musculus OX=10090 GN=Rap1 | 37 | 7 | 11 | 2 | 184 | 21 | 6,67 | 20,2 | 7 | 0,925 | 1,217 | 1,149 | 1,185 | 1,085 | 1,346 | 0,799 | 0,936 | 1,07 | 0,07 | 5,9E-01 |
| Q9Z2V5 | Histone deacetylase 6 OS=Mus musculus OX=10090 GN=Hdac6 PE= | 5 | 5 | 5 | 5 | 1149 | 125,7 | 5,78 | 9,79 | 5 | 1,127 | 0,954 | 1,135 | 1,182 | 0,99 | 1,14 | 1,041 | 1,09 | 1,03 | 0,03 | 5,9E-01 |
| Q9Z0U1 | Tight junction protein ZO-2 OS=Mus musculus OX=10090 GN=Tjp2 | 7 | 7 | 7 | 7 | 1167 | 131,2 | 6,79 | 6,83 | 7 | 0,92 | 1,085 | 1,151 | 1,034 | 0,958 | 1,12 | 0,989 | 0,983 | 1,03 | 0,03 | 5,9E-01 |
| Q62095 | ATP-dependent RNA helicase DDX3Y OS=Mus musculus OX=10090 | 11 | 6 | 6 | 2 | 658 | 73,4 | 7,53 | 7,57 | 6 | 0,856 | 1,198 | 1,219 | 1,263 | 1,234 | 1,099 | 1,129 | 1,318 | 0,95 | -0,05 | 5,9E-01 |
| P62071 | Ras-related protein R-Ras2 OS=Mus musculus OX=10090 GN=Rras2 | 47 | 9 | 16 | 7 | 204 | 23,4 | 6,01 | 27,27 | 9 | 1,231 | 1,039 | 1,063 | 1,078 | 0,953 | 1,256 | 1,196 | 1,189 | 0,96 | -0,04 | 5,9E-01 |
| Q2PFD7 | PH and SEC7 domain-containing protein 3 OS=Mus musculus OX=1( | 14 | 14 | 17 | 14 | 1037 | 114,7 | 6,24 | 46,44 | 14 | 0,919 | 1,015 | 0,921 | 1,03 | 0,947 | 0,923 | 0,928 | 1,006 | 1,02 | 0,02 | 5,9E-01 |
| Q9JKX6 | ADP-sugar pyrophosphatase OS=Mus musculus OX=10090 GN=Nud | 19 | 3 | 3 | 3 | 218 | 24 | 5,48 | 6,27 | 3 | 1,189 | 1,127 | 1,161 | 0,907 | 0,97 | 1,099 | 1,045 | 1,106 | 1,04 | 0,04 | 5,9E-01 |
| Q9CQW1 | Synaptobrevin homolog YKT6 OS=Mus musculus OX=10090 GN=Ykt | 12 | 2 | 2 | 2 | 198 | 22,3 | 6,35 | 6,2 | 2 | 0,868 | 0,925 | 0,838 | 0,773 | 0,696 | 1,028 | 0,813 | 0,665 | 1,06 | 0,06 | 5,9E-01 |
| Q8VEH8 | Endoplasmic reticulum lectin 1 OS=Mus musculus OX=10090 GN=Er | 3 | 1 | 1 | 1 | 483 | 54,9 | 6,25 | 2,51 | 1 | 2,587 | 2,791 | 5,177 | 1,737 | 2,23 | 2,843 | 2,02 | 3,371 | 1,17 | 0,17 | 5,9E-01 |
| Q3UKC1 | Tax1-binding protein 1 homolog OS=Mus musculus OX=10090 GN=T | 3 | 2 | 2 | 2 | 814 | 93,6 | 5,33 | 4,07 | 2 | 0,581 | 0,947 | 1,284 | 1,203 | 1,305 | 0,844 | 1,234 | 1,063 | 0,90 | -0,10 | 5,9E-01 |
| Q810B6 | Rabankyrin-5 OS=Mus musculus OX=10090 GN=Ankfy1 PE=1 SV=2 | 2 | 3 | 4 | 3 | 1169 | 128,6 | 5,91 | 7,73 | 3 | 1,229 | 1,078 | 0,836 | 0,981 | 1,023 | 1,26 | 0,982 | 1,094 | 0,95 | -0,05 | 5,9E-01 |
| B1AXP6 | Mitochondrial import receptor subunit TOM5 homolog OS=Mus mu | 45 | 3 | 3 | 3 | 51 | 6 | 9,6 | 6,34 | 3 | 0,85 | 0,9 | 0,891 | 0,878 | 0,992 | 0,66 | 0,866 | 0,843 | 1,05 | 0,05 | 5,9E-01 |
| Q8BGF9 | Solute carrier family 25 member 44 OS=Mus musculus OX=10090 G | 4 | 1 | 2 | 1 | 314 | 35,3 | 9,66 | 2,27 | 1 | 1,055 | 1,138 | 1,038 | 1,128 | 0,923 | 1,435 | 1,152 | 1,098 | 0,95 | -0,05 | 5,9E-01 |
| Q9CQP2 | Trafficking protein particle complex subunit 2 OS=Mus musculus OX | 14 | 2 | 2 | 2 | 140 | 16,4 | 6,52 | 2,56 | 2 | 0,98 | 0,782 | 0,956 | 1 | 1,126 | 1,006 | 0,97 | 0,805 | 0,95 | -0,05 | 5,9E-01 |
| Q922D4 | Serine/threonine-protein phosphatase 6 regulatory subunit 3 OS=M | 4 | 3 | 3 | 3 | 844 | 94,6 | 4,58 | 0 | 3 | 0,881 | 0,681 | 0,915 | 0,799 | 0,719 | 1,291 | 1,023 | 0,612 | 0,90 | -0,10 | 5,9E-01 |
| Q3UVG3 | Protein FAM91A1 OS=Mus musculus OX=10090 GN=Fam91a1 PE=1 | 1 | 1 | 1 | 1 | 837 | 93,4 | 6,35 | 0 | 1 | 0,939 | 0,661 | 0,589 | 0,657 | 0,655 | 0,711 | 0,377 | 0,824 | 1,11 | 0,11 | 5,9E-01 |
| Q7TMS5 | ATP-binding cassette sub-family G member 2 OS=Mus musculus OX | 6 | 4 | 4 | 4 | 657 | 72,9 | 8,54 | 3,14 | 4 | 1,129 | 0,843 | 0,983 | 1,052 | 0,76 | 0,905 | 1,05 | 1,076 | 1,06 | 0,06 | 5,9E-01 |
| Q61137 | Astrotactin-1 OS=Mus musculus OX=10090 GN=Astn1 PE=1 SV=4 | 8 | 8 | 11 | 8 | 1302 | 144,8 | 5,21 | 19,43 | 8 | 1,152 | 0,979 | 1,058 | 0,908 | 0,94 | 1,02 | 0,907 | 1,081 | 1,04 | 0,04 | 5,9E-01 |
| Q9Z2Z6 | Mitochondrial carnitine/acylcarnitine carrier protein OS=Mus musc | 3 | 1 | 1 | 1 | 301 | 33 | 9,11 | 2,49 | 1 | 1,39 | 1,066 | 1,138 | 0,866 | 0,917 | 1,265 | 1,015 | 0,96 | 1,07 | 0,07 | 5,9E-01 |
| Q7TNR6 | Immunoglobulin superfamily member 21 OS=Mus musculus OX=10 | 7 | 3 | 4 | 3 | 468 | 51,9 | 6,95 | 8,79 | 3 | 0,737 | 0,933 | 0,801 | 1,033 | 0,94 | 0,713 | 0,983 | 1,108 | 0,94 | -0,06 | 5,9E-01 |
| P48453 | Serine/threonine-protein phosphatase 2B catalytic subunit beta iso | 22 | 11 | 27 | 5 | 525 | 59,1 | 5,91 | 41,53 | 11 | 0,941 | 1,174 | 0,919 | 0,901 | 0,867 | 1,12 | 0,993 | 1,165 | 0,95 | -0,05 | 5,9E-01 |
| Q02053 | Ubiquitin-like modifier-activating enzyme 1 OS=Mus musculus OX= | 24 | 24 | 36 | 24 | 1058 | 117,7 | 5,66 | 87,65 | 24 | 0,858 | 0,885 | 0,981 | 1,004 | 1,001 | 0,845 | 1,011 | 0,991 | 0,97 | -0,03 | 5,9E-01 |
| Q8K004 | Spermatogenesis-associated protein 2 OS=Mus musculus OX=1009( | 2 | 1 | 1 | 1 | 515 | 57,8 | 8,63 | 0 | 1 | 1,044 | 1,178 | 1,138 | 1,322 | 0,986 | 1,702 | 1,217 | 1,149 | 0,93 | -0,07 | 5,9E-01 |
| P63034 | Cytohesin-2 OS=Mus musculus OX=10090 GN=Cyth2 PE=1 SV=2 | 12 | 3 | 3 | 2 | 400 | 46,6 | 5,63 | 0 | 3 | 0,532 | 1,916 | 0,289 | 0,654 | 0,719 | 1,65 | 1,079 | 0,885 | 0,78 | -0,22 | 5,9E-01 |
| Q9D8B3 | Charged multivesicular body protein 4b OS=Mus musculus OX=100( | 22 | 6 | 6 | 6 | 224 | 24,9 | 4,82 | 13,24 | 6 | 1,099 | 1,114 | 1,065 | 1,187 | 1,011 | 1,153 | 1,031 | 1,163 | 1,02 | 0,02 | 5,9E-01 |
| Q6PF93 | Phosphatidylinositol 3-kinase catalytic subunit type 3 OS=Mus mus | 7 | 5 | 5 | 5 | 887 | 101,4 | 6,73 | 6,63 | 5 | 0,752 | 1,132 | 0,905 | 1,009 | 1,143 | 0,95 | 0,834 | 1,116 | 0,94 | -0,06 | 5,9E-01 |
| Q8R104 | NAD-dependent protein deacetylase sirtuin-3 OS=Mus musculus OX | 7 | 2 | 2 | 2 | 334 | 36,6 | 7,44 | 6,6 | 2 | 1,257 | 1,231 | 0,914 | 1,077 | 1,049 | 0,87 | 1,287 | 1,008 | 1,06 | 0,06 | 5,9E-01 |
| Q9CXW4 | 60S ribosomal protein L11 OS=Mus musculus OX=10090 GN=Rpl11 | 13 | 2 | 3 | 2 | 178 | 20,2 | 9,6 | 9,24 | 2 | 1,071 | 1,124 | 0,968 | 1,053 | 1,082 | 1,037 | 1,042 | 1,148 | 0,98 | -0,02 | 5,9E-01 |
| Q8BWZ3 | N-alpha-acetyltransferase 25, NatB auxiliary subunit OS=Mus musc | 1 | 1 | 1 | 1 | 972 | 111,6 | 6,52 | 0 | 1 | 0,66 | 1,167 | 0,934 | 0,96 | 0,871 | 1,496 | 0,889 | 0,883 | 0,90 | -0,10 | 5,9E-01 |
| Q8R059 | UDP-glucose 4-epimerase OS=Mus musculus OX=10090 GN=Gale P | 3 | 1 | 1 | 1 | 347 | 38,2 | 6,74 | 0 | 1 | 0,768 | 0,828 | 0,699 | 0,959 | 0,996 | 0,587 | 0,778 | 1,228 | 0,91 | -0,09 | 5,9E-01 |
| Q7TNF0 | Double C2-like domain-containing protein alpha OS=Mus musculus | 11 | 4 | 4 | 3 | 405 | 44,6 | 7,12 | 10,7 | 4 | 1,024 | 0,814 | 1,066 | 1,133 | 1,013 | 1,058 | 0,974 | 1,175 | 0,96 | -0,04 | 5,9E-01 |
| Q91V36 | Nuclear receptor-binding protein 2 OS=Mus musculus OX=10090 G | 11 | 6 | 6 | 6 | 499 | 57,3 | 6,35 | 7,98 | 6 | 1,077 | 1,081 | 1,13 | 0,982 | 0,819 | 1,212 | 0,992 | 1,052 | 1,05 | 0,05 | 5,9E-01 |
| Q9CQH7 | Transcription factor BTF3 homolog 4 OS=Mus musculus OX=10090 | 8 | 1 | 1 | 1 | 158 | 17,3 | 6,35 | 0 | 1 | 1,364 | 1,434 | 1,339 | 1,583 | 0,99 | 2,249 | 1,93 | 1,224 | 0,89 | -0,11 | 6,0E-01 |
| P51410 | 60S ribosomal protein L9 OS=Mus musculus OX=10090 GN=Rpl9 PE | 12 | 2 | 2 | 2 | 192 | 21,9 | 9,95 | 0 | 2 | 1,039 | 0,64 | 0,895 | 0,913 | 1,182 | 0,808 | 0,782 | 0,995 | 0,93 | -0,07 | 6,0E-01 |
| Q04690 | Neurofibromin OS=Mus musculus OX=10090 GN=Nf1 PE=1 SV=1 | 4 | 10 | 12 | 10 | 2841 | 319,4 | 7,39 | 12,52 | 10 | 1,162 | 1,038 | 1,101 | 0,98 | 0,975 | 1,097 | 1,025 | 1,077 | 1,03 | 0,03 | 6,0E-01 |
| Q8R361 | Rab11 family-interacting protein 5 OS=Mus musculus OX=10090 GN | 4 | 2 | 3 | 2 | 645 | 69,5 | 9,07 | 6,27 | 2 | 2,128 | 1,464 | 1,345 | 1,1 | 1,171 | 1,269 | 1,029 | 1,912 | 1,12 | 0,12 | 6,0E-01 |
| O54916 | RalBP1-associated Eps domain-containing protein 1 OS=Mus muscu | 11 | 6 | 6 | 6 | 795 | 86,5 | 5,58 | 7,65 | 6 | 0,848 | 0,836 | 1,027 | 0,813 | 0,87 | 0,76 | 0,988 | 0,737 | 1,05 | 0,05 | 6,0E-01 |
| Q9D7N9 | Adipocyte plasma membrane-associated protein OS=Mus musculus | 9 | 4 | 4 | 4 | 415 | 46,4 | 6,32 | 9,3 | 4 | 1,143 | 1,036 | 1,21 | 0,831 | 0,939 | 1,157 | 0,936 | 0,969 | 1,05 | 0,05 | 6,0E-01 |
| Q9WTX5 | S-phase kinase-associated protein 1 OS=Mus musculus OX=10090 ( | 25 | 4 | 4 | 4 | 163 | 18,7 | 4,54 | 10,13 | 4 | 1,211 | 1,298 | 0,889 | 1,005 | 1,063 | 1,131 | 0,989 | 0,999 | 1,05 | 0,05 | 6,0E-01 |
| P59279 | Ras-related protein Rab-2B OS=Mus musculus OX=10090 GN=Rab2 | 34 | 7 | 12 | 2 | 216 | 24,2 | 6,68 | 31,17 | 7 | 0,01 | 1,718 | 1,851 | 1,183 | 0,998 | 1,173 | 0,68 | 0,95 | 1,25 | 0,25 | 6,0E-01 |
| Q9D0L7 | Armadillo repeat-containing protein 10 OS=Mus musculus OX=1009 | 7 | 2 | 2 | 2 | 306 | 33,3 | 7,99 | 1,83 | 2 | 1,687 | 1,047 | 1,628 | 0,936 | 1,067 | 1,327 | 1,16 | 1,292 | 1,09 | 0,09 | 6,0E-01 |
| Q9DBR3 | Armadillo repeat-containing protein 8 OS=Mus musculus OX=1009( | 10 | 5 | 5 | 5 | 673 | 75,3 | 6,73 | 6,95 | 5 | 1,141 | 1,025 | 1,254 | 1,179 | 1,098 | 1,14 | 1,151 | 1,1 | 1,02 | 0,02 | 6,0E-01 |
| Q9Z130 | Heterogeneous nuclear ribonucleoprotein D-like OS=Mus musculus | 11 | 3 | 3 | 2 | 301 | 33,5 | 7,31 | 6,26 | 3 | 1,159 | 1,071 | 1,01 | 0,983 | 0,912 | 1,387 | 1,053 | 1,108 | 0,95 | -0,05 | 6,0E-01 |
| Q5XJY5 | Coatomer subunit delta OS=Mus musculus OX=10090 GN=Arcn1 PE | 14 | 6 | 6 | 6 | 511 | 57,2 | 6,21 | 6,86 | 6 | 1,25 | 1,002 | 0,895 | 0,986 | 1,003 | 0,896 | 1,069 | 0,979 | 1,05 | 0,05 | 6,0E-01 |
| P31648 | Sodium- and chloride-dependent GABA transporter 1 OS=Mus mus | 6 | 2 | 3 | 2 | 599 | 67 | 7,96 | 11,45 | 2 | 0,821 | 1,008 | 0,896 | 0,925 | 0,868 | 0,943 | 1,014 | 0,933 | 0,97 | -0,03 | 6,0E-01 |
| Q8R3Z5 | Voltage-dependent L-type calcium channel subunit beta-1 OS=Mus | 13 | 7 | 12 | 4 | 597 | 65,4 | 6,62 | 18,39 | 7 | 1,234 | 1,126 | 0,905 | 0,88 | 0,98 | 1,299 | 0,981 | 1,139 | 0,94 | -0,06 | 6,0E-01 |
| Q9QYC0 | Alpha-adducin OS=Mus musculus OX=10090 GN=Add1 PE=1 SV=2 | 23 | 16 | 24 | 15 | 735 | 80,6 | 5,9 | 48,8 | 16 | 0,977 | 0,906 | 0,951 | 0,987 | 1,001 | 0,939 | 0,969 | 0,961 | 0,99 | -0,01 | 6,0E-01 |
| Q7TSS2 | Ubiquitin-conjugating enzyme E2 Q1 OS=Mus musculus OX=10090 | 5 | 2 | 2 | 2 | 422 | 46,1 | 5,1 | 3,28 | 2 | 1,245 | 1,105 | 1,107 | 1,026 | 0,867 | 1,258 | 0,892 | 1,216 | 1,06 | 0,06 | 6,0E-01 |
| P58389 | Serine/threonine-protein phosphatase 2A activator OS=Mus muscu | 17 | 4 | 4 | 4 | 323 | 36,7 | 6,39 | 9,92 | 4 | 1,17 | 1,024 | 1,097 | 0,909 | 0,955 | 1,191 | 0,977 | 0,887 | 1,05 | 0,05 | 6,0E-01 |
| Q8K0V2 | DCN1-like protein 3 OS=Mus musculus OX=10090 GN=Dcun1d3 PE= | 4 | 1 | 2 | 1 | 304 | 34,4 | 5,12 | 2,33 | 1 | 1,376 | 1,395 | 0,764 | 0,674 | 0,777 | 1,347 | 0,547 | 0,972 | 1,16 | 0,16 | 6,0E-01 |
| Q8C8R3 | Ankyrin-2 OS=Mus musculus OX=10090 GN=Ank2 PE=1 SV=2 | 28 | 82 | 112 | 1 | 3898 | 426 | 5,17 | 240,74 | 82 | 0,809 | 1,045 | 0,83 | 0,916 | 1,016 | 0,937 | 0,829 | 0,964 | 0,96 | -0,04 | 6,0E-01 |
| P06837 | Neuromodulin OS=Mus musculus OX=10090 GN=Gap43 PE=1 SV=1 | 70 | 19 | 50 | 19 | 227 | 23,6 | 4,73 | 94,68 | 19 | 0,712 | 0,685 | 0,91 | 1,053 | 0,919 | 0,708 | 1,049 | 0,93 | 0,93 | -0,07 | 6,0E-01 |
| Q9R0N3 | Synaptotagmin-11 OS=Mus musculus OX=10090 GN=Syt11 PE=1 SV | 5 | 2 | 2 | 2 | 430 | 48,3 | 9,11 | 2,89 | 2 | 1,094 | 1,221 | 0,639 | 1,192 | 0,95 | 0,885 | 1,188 | 0,769 | 1,09 | 0,09 | 6,0E-01 |
| Q8BHF7 | CDP-diacylglycerol--glycerol-3-phosphate 3-phosphatidyltransferas | 4 | 2 | 3 | 2 | 553 | 62,5 | 8,9 | 2,92 | 2 | 1,028 | 1,167 | 1 | 0,888 | 0,933 | 1,366 | 0,987 | 1,044 | 0,94 | -0,06 | 6,0E-01 |
| P10630 | Eukaryotic initiation factor 4A-II OS=Mus musculus OX=10090 GN=I | 17 | 7 | 9 | 5 | 407 | 46,4 | 5,48 | 18,89 | 7 | 0,943 | 1,087 | 1,015 | 0,973 | 0,951 | 0,99 | 1,046 | 0,946 | 1,02 | 0,02 | 6,0E-01 |
| O08808 | Protein diaphanous homolog 1 OS=Mus musculus OX=10090 GN=D | 7 | 9 | 9 | 9 | 1255 | 139,3 | 5,57 | 18,58 | 9 | 1,046 | 1,009 | 1,067 | 1,121 | 0,999 | 0,999 | 1,115 | 1,051 | 1,02 | 0,02 | 6,0E-01 |
| P68369 | Tubulin alpha-1A chain OS=Mus musculus OX=10090 GN=Tuba1a P | 60 | 26 | 106 | 4 | 451 | 50,1 | 5,06 | 308,3 | 26 | 0,862 | 0,952 | 0,965 | 0,93 | 0,919 | 0,99 | 0,943 | 0,919 | 0,98 | -0,02 | 6,0E-01 |
| Q61771 | Kinesin-like protein KIF3B OS=Mus musculus OX=10090 GN=Kif3b P | 10 | 7 | 7 | 5 | 747 | 85,2 | 7,69 | 16,92 | 7 | 0,854 | 0,912 | 0,982 | 0,987 | 1,008 | 0,758 | 0,915 | 0,921 | 1,04 | 0,04 | 6,0E-01 |

| | | | | | | | | | | | | | | | | | | | | |
|---|---|---|---|---|---|---|---|---|---|---|---|---|---|---|---|---|---|---|---|---|
| Q8BH55 | Threonine synthase-like 1 OS=Mus musculus OX=10090 GN=Thnsl1 | 5 | 3 | 4 | 3 | 747 | 83 | 7,2 | 6,57 | 3 | 1,083 | 1,079 | 0,962 | 0,94 | 0,969 | 1,129 | 0,87 | 0,952 | 1,04 | 0,04 | 6,1E-01 |
| A6H8H5 | Potassium voltage-gated channel subfamily B member 2 OS=Mus m | 1 | 1 | 1 | 1 | 907 | 102,3 | 6,1 | 0 | 1 | 1,556 | 1,314 | 1,12 | 0,799 | 0,902 | 1,273 | 0,912 | 1,283 | 1,10 | 0,10 | 6,1E-01 |
| P05063 | Fructose-bisphosphate aldolase C OS=Mus musculus OX=10090 GN | 42 | 21 | 37 | 15 | 363 | 39,4 | 7,12 | 87,83 | 21 | 1,091 | 1,254 | 0,963 | 0,946 | 0,956 | 1,076 | 0,992 | 1,063 | 1,04 | 0,04 | 6,1E-01 |
| O88569 | Heterogeneous nuclear ribonucleoproteins A2/B1 OS=Mus musculu | 31 | 11 | 16 | 11 | 353 | 37,4 | 8,95 | 40,18 | 11 | 1,21 | 1,094 | 1,017 | 0,814 | 1,245 | 1,344 | 0,861 | 0,988 | 0,93 | -0,07 | 6,1E-01 |
| Q9D6Y7 | Mitochondrial peptide methionine sulfoxide reductase OS=Mus mu | 16 | 3 | 3 | 3 | 233 | 26 | 8,41 | 5,15 | 3 | 1,025 | 1,28 | 0,865 | 0,863 | 0,953 | 1,184 | 0,992 | 1,151 | 0,94 | -0,06 | 6,1E-01 |
| P61750 | ADP-ribosylation factor 4 OS=Mus musculus OX=10090 GN=Arf4 PE | 24 | 4 | 5 | 1 | 180 | 20,4 | 7,14 | 18,05 | 4 | 0,786 | 1,042 | 0,925 | 0,902 | 0,888 | 0,983 | 0,916 | 0,995 | 0,97 | -0,03 | 6,1E-01 |
| Q61207 | Prosaposin OS=Mus musculus OX=10090 GN=Psap PE=1 SV=2 | 11 | 6 | 6 | 6 | 557 | 61,4 | 5,19 | 6,97 | 6 | 0,96 | 1,114 | 0,972 | 1,151 | 1,059 | 1,085 | 1,06 | 1,101 | 0,97 | -0,03 | 6,1E-01 |
| B2RXS4 | Plexin-B2 OS=Mus musculus OX=10090 GN=Plxnb2 PE=1 SV=1 | 6 | 9 | 10 | 9 | 1842 | 206,1 | 5,87 | 16,5 | 9 | 1,186 | 0,949 | 1,053 | 0,982 | 0,95 | 1,188 | 0,925 | 0,928 | 1,04 | 0,04 | 6,1E-01 |
| O55106 | Striatin OS=Mus musculus OX=10090 GN=Strn PE=1 SV=2 | 9 | 6 | 6 | 6 | 780 | 85,9 | 5,27 | 20,08 | 6 | 1,083 | 0,908 | 1,078 | 1,054 | 1,098 | 0,993 | 1,113 | 1,028 | 0,97 | -0,03 | 6,1E-01 |
| Q9CYH2 | Peroxiredoxin-like 2A OS=Mus musculus OX=10090 GN=Prxl2a PE=1 | 15 | 3 | 3 | 3 | 218 | 24,4 | 9,17 | 6,24 | 3 | 0,919 | 0,955 | 1,018 | 0,874 | 0,876 | 0,977 | 0,911 | 0,922 | 1,02 | 0,02 | 6,1E-01 |
| Q99J31 | Oligophrenin-1 OS=Mus musculus OX=10090 GN=Ophn1 PE=1 SV=1 | 2 | 2 | 2 | 2 | 802 | 91,9 | 7,96 | 2 | 2 | 0,927 | 1,036 | 1,155 | 1,073 | 1,012 | 0,892 | 1,155 | 0,975 | 1,04 | 0,04 | 6,1E-01 |
| Q9WUA6 | RAC-gamma serine/threonine-protein kinase OS=Mus musculus OX | 5 | 2 | 2 | 2 | 479 | 55,7 | 6,02 | 4,32 | 2 | 0,804 | 0,873 | 0,857 | 1,081 | 0,961 | 0,798 | 0,87 | 0,834 | 1,04 | 0,04 | 6,1E-01 |
| Q3U0D9 | E3 ubiquitin-protein ligase HACE1 OS=Mus musculus OX=10090 GN | 4 | 2 | 3 | 2 | 909 | 102 | 5,76 | 3,06 | 2 | 1,49 | 1,243 | 0,953 | 1,246 | 1,117 | 1,619 | 1,412 | 1,136 | 0,93 | -0,07 | 6,1E-01 |
| P14131 | 40S ribosomal protein S16 OS=Mus musculus OX=10090 GN=Rps16 | 5 | 1 | 1 | 1 | 146 | 16,4 | 10,21 | 2,3 | 1 | 1,238 | 1,312 | 1,082 | 0,889 | 1,019 | 1,062 | 1,104 | 1,127 | 1,05 | 0,05 | 6,1E-01 |
| Q9CXE7 | Transmembrane emp24 domain-containing protein 5 OS=Mus mus | 8 | 2 | 2 | 2 | 229 | 26,2 | 4,93 | 2,02 | 2 | 1,185 | 1,116 | 1,166 | 1,123 | 1,124 | 1,17 | 1,267 | 1,113 | 0,98 | -0,02 | 6,1E-01 |
| Q0VBD0 | Integrin beta-8 OS=Mus musculus OX=10090 GN=Itgb8 PE=1 SV=1 | 4 | 2 | 2 | 2 | 767 | 84,5 | 7,09 | 6,1 | 2 | 1,273 | 1,046 | 1,149 | 1,067 | 1,045 | 1,318 | 0,999 | 1,422 | 0,95 | -0,05 | 6,1E-01 |
| A2CG49-9 | Isoform 9 of Kalirin OS=Mus musculus OX=10090 GN=Kalrn | 19 | 23 | 27 | 1 | 1654 | 191,1 | 6,07 | 61,37 | 23 | 0,592 | 1,298 | 0,457 | 1,01 | 0,962 | 0,567 | 0,692 | 0,682 | 1,16 | 0,16 | 6,1E-01 |
| Q6PCP5 | Mitochondrial fission factor OS=Mus musculus OX=10090 GN=Mff I | 15 | 3 | 3 | 3 | 291 | 32,9 | 6,83 | 2,73 | 3 | 1,03 | 1,384 | 1,299 | 1,269 | 1,171 | 1,123 | 1,307 | 1,197 | 1,04 | 0,04 | 6,1E-01 |
| P63089 | Pleiotrophin OS=Mus musculus OX=10090 GN=Ptn PE=1 SV=1 | 4 | 1 | 1 | 1 | 168 | 18,9 | 9,6 | 1,93 | 1 | 0,769 | 0,908 | 0,988 | 0,905 | 1,115 | 0,602 | 0,853 | 0,747 | 1,08 | 0,08 | 6,1E-01 |
| Q9R1P0 | Proteasome subunit alpha type-4 OS=Mus musculus OX=10090 GN | 18 | 5 | 7 | 5 | 261 | 29,5 | 7,72 | 20,88 | 5 | 1,24 | 0,976 | 1,123 | 1,035 | 1,01 | 1,125 | 1,054 | 1,051 | 1,03 | 0,03 | 6,1E-01 |
| P97789 | 5'-3' exoribonuclease 1 OS=Mus musculus OX=10090 GN=Xrn1 PE= | 1 | 1 | 1 | 1 | 1719 | 194,2 | 7,5 | 2,56 | 1 | 1,391 | 1,319 | 0,796 | 0,874 | 0,873 | 1,146 | 0,791 | 1,182 | 1,10 | 0,10 | 6,1E-01 |
| O54692 | Centromere/kinetochore protein zw10 homolog OS=Mus musculus | 2 | 2 | 2 | 2 | 779 | 88 | 5,92 | 0 | 2 | 1,466 | 1,122 | 0,946 | 1,201 | 1,234 | 1,23 | 0,81 | 1,144 | 1,07 | 0,07 | 6,1E-01 |
| Q02819 | Nucleobindin-1 OS=Mus musculus OX=10090 GN=Nucb1 PE=1 SV=2 | 10 | 3 | 4 | 3 | 459 | 53,4 | 5,07 | 8,21 | 3 | 2,438 | 2,277 | 0,833 | 1,225 | 1,319 | 1,541 | 1,412 | 1,643 | 1,15 | 0,15 | 6,1E-01 |
| Q69ZH9 | Rho GTPase-activating protein 23 OS=Mus musculus OX=10090 GN | 5 | 6 | 7 | 6 | 1483 | 161,7 | 8,98 | 3,16 | 6 | 1,316 | 1,025 | 0,91 | 1,064 | 0,911 | 1,348 | 1,262 | 1,073 | 0,94 | -0,06 | 6,1E-01 |
| Q8BM85 | TBC domain-containing protein kinase-like protein OS=Mus muscul | 4 | 3 | 3 | 3 | 762 | 86,3 | 6,2 | 6,51 | 3 | 1,306 | 1,048 | 1,153 | 1,071 | 1,074 | 1,259 | 1,243 | 1,157 | 0,97 | -0,03 | 6,1E-01 |
| Q2TA57 | Aspartate beta-hydroxylase domain-containing protein 1 OS=Mus r | 5 | 1 | 1 | 1 | 360 | 38,2 | 8,25 | 2,95 | 1 | 1,245 | 1,402 | 1,191 | 0,975 | 0,96 | 1,232 | 1,273 | 1,745 | 0,92 | -0,08 | 6,1E-01 |
| Q61171 | Peroxiredoxin-2 OS=Mus musculus OX=10090 GN=Prdx2 PE=1 SV=3 | 27 | 5 | 8 | 5 | 198 | 21,8 | 5,41 | 19,08 | 5 | 0,913 | 0,858 | 0,939 | 1,085 | 1,059 | 0,843 | 1,043 | 0,998 | 0,96 | -0,04 | 6,1E-01 |
| Q9CRA9 | FGFR1 oncogene partner 2 homolog OS=Mus musculus OX=10090 ( | 11 | 2 | 3 | 2 | 253 | 29,4 | 5,83 | 8,4 | 2 | 1,04 | 0,835 | 0,93 | 0,973 | 0,995 | 0,872 | 1,176 | 0,907 | 0,96 | -0,04 | 6,1E-01 |
| Q6P4T2 | U5 small nuclear ribonucleoprotein 200 kDa helicase OS=Mus musc | 2 | 5 | 5 | 5 | 2136 | 244,4 | 6,06 | 1,67 | 5 | 1,27 | 1,42 | 0,828 | 0,91 | 1,049 | 1,192 | 0,783 | 1,05 | 1,09 | 0,09 | 6,1E-01 |
| Q6ZWZ2 | Ubiquitin-conjugating enzyme E2 R2 OS=Mus musculus OX=10090 ( | 13 | 3 | 3 | 3 | 238 | 27,1 | 4,42 | 1,82 | 3 | 1,548 | 1,323 | 1,298 | 1,12 | 1,325 | 0,995 | 1,192 | 1,486 | 1,06 | 0,06 | 6,1E-01 |
| P12970 | 60S ribosomal protein L7a OS=Mus musculus OX=10090 GN=Rpl7a | 23 | 7 | 9 | 7 | 266 | 30 | 10,56 | 21,98 | 7 | 1,192 | 1,049 | 1,111 | 0,94 | 1,03 | 1,077 | 1,003 | 1,063 | 1,03 | 0,03 | 6,1E-01 |
| Q9QY42 | Prosaposin receptor GPR37 OS=Mus musculus OX=10090 GN=Gpr3 | 2 | 1 | 1 | 1 | 600 | 66,7 | 8,12 | 1,7 | 1 | 0,716 | 0,866 | 0,843 | 0,841 | 0,549 | 0,869 | 0,876 | 0,793 | 1,06 | 0,06 | 6,1E-01 |
| P40124 | Adenylyl cyclase-associated protein 1 OS=Mus musculus OX=10090 | 25 | 14 | 20 | 13 | 474 | 51,5 | 7,52 | 38,66 | 14 | 1,153 | 1,039 | 1,13 | 1,012 | 0,946 | 1,145 | 1,058 | 1,071 | 1,03 | 0,03 | 6,1E-01 |
| Q99LI8 | Hepatocyte growth factor-regulated tyrosine kinase substrate OS=I | 9 | 7 | 7 | 7 | 775 | 86 | 6,16 | 14,96 | 7 | 0,922 | 0,932 | 0,933 | 1,038 | 0,938 | 0,895 | 0,973 | 0,951 | 1,02 | 0,02 | 6,1E-01 |
| Q9DBS2 | Tumor protein p63-regulated gene 1-like protein OS=Mus musculus | 17 | 4 | 5 | 4 | 266 | 29,8 | 7,37 | 9,6 | 4 | 1,26 | 0,995 | 0,987 | 1,11 | 0,928 | 1,219 | 0,999 | 1,016 | 1,05 | 0,05 | 6,1E-01 |
| Q9DBR7 | Protein phosphatase 1 regulatory subunit 12A OS=Mus musculus O | 7 | 6 | 8 | 6 | 1029 | 114,9 | 5,49 | 16,35 | 6 | 0,903 | 0,913 | 0,959 | 1,104 | 1,011 | 0,852 | 0,939 | 0,956 | 1,03 | 0,03 | 6,1E-01 |
| Q8BH04 | Phosphoenolpyruvate carboxykinase [GTP], mitochondrial OS=Mus | 15 | 8 | 12 | 8 | 640 | 70,5 | 7,28 | 13,21 | 8 | 0,876 | 1,135 | 0,92 | 0,959 | 0,907 | 1,161 | 0,931 | 1,066 | 0,96 | -0,04 | 6,1E-01 |
| O08709 | Peroxiredoxin-6 OS=Mus musculus OX=10090 GN=Prdx6 PE=1 SV=3 | 38 | 7 | 10 | 7 | 224 | 24,9 | 6,01 | 20,28 | 7 | 0,94 | 0,979 | 1,101 | 0,924 | 1,04 | 1,055 | 0,93 | 1,023 | 0,97 | -0,03 | 6,1E-01 |
| Q6ZWU9 | 40S ribosomal protein S27 OS=Mus musculus OX=10090 GN=Rps27 | 38 | 3 | 4 | 3 | 84 | 9,5 | 9,45 | 9,44 | 3 | 1,819 | 1,111 | 1,243 | 1,329 | 1,171 | 1,112 | 1,558 | 1,271 | 1,08 | 0,08 | 6,1E-01 |
| Q9CQU0 | Thioredoxin domain-containing protein 12 OS=Mus musculus OX=1 | 18 | 2 | 3 | 2 | 170 | 19 | 5,26 | 9,46 | 2 | 1,077 | 1,208 | 1,262 | 1,218 | 1,035 | 1,277 | 1,136 | 1,181 | 1,03 | 0,03 | 6,2E-01 |
| Q6ZPU9 | KIF1-binding protein OS=Mus musculus OX=10090 GN=Kif1bp PE=1 | 12 | 4 | 5 | 4 | 617 | 71 | 5,48 | 0 | 4 | 1,175 | 1,962 | 1,144 | 1,3 | 0,921 | 1,283 | 1,265 | 1,608 | 1,10 | 0,10 | 6,2E-01 |
| Q08274 | Dystrophia myotonica WD repeat-containing protein OS=Mus musc | 11 | 5 | 5 | 5 | 665 | 69,8 | 7,24 | 13,31 | 5 | 1,138 | 1,158 | 1,074 | 0,956 | 0,945 | 1,252 | 1,119 | 1,179 | 0,96 | -0,04 | 6,2E-01 |
| Q9DB50 | AP-1 complex subunit sigma-2 OS=Mus musculus OX=10090 GN=Ap | 8 | 1 | 1 | 1 | 160 | 18,9 | 5,48 | 2,41 | 1 | 0,807 | 0,689 | 1,011 | 1,095 | 0,863 | 1,175 | 1,567 | 0,512 | 0,87 | -0,13 | 6,2E-01 |
| Q91Y44 | Bromodomain testis-specific protein OS=Mus musculus OX=10090 ( | 1 | 1 | 2 | 1 | 956 | 107,2 | 8,97 | 5,29 | 1 | 0,764 | 2,184 | 0,874 | 0,316 | 1,585 | 0,449 | 0,345 | 0,72 | 1,34 | 0,34 | 6,2E-01 |
| Q9WV95 | Pleckstrin homology-like domain family A member 3 OS=Mus musc | 8 | 1 | 1 | 1 | 125 | 13,7 | 9,67 | 1,77 | 1 | 1,357 | 0,739 | 0,527 | 0,932 | 0,756 | 1,284 | 0,877 | 1,086 | 0,89 | -0,11 | 6,2E-01 |
| Q8R086 | Sulfite oxidase, mitochondrial OS=Mus musculus OX=10090 GN=Su | 4 | 2 | 2 | 2 | 546 | 60,7 | 6,54 | 4,58 | 2 | 1,193 | 1,323 | 1,037 | 1,017 | 0,856 | 1,394 | 0,935 | 1,091 | 1,07 | 0,07 | 6,2E-01 |
| Q62165 | Dystroglycan OS=Mus musculus OX=10090 GN=Dag1 PE=1 SV=4 | 1 | 1 | 1 | 1 | 893 | 96,8 | 8,44 | 2,28 | 1 | 1,118 | 0,662 | 0,925 | 0,84 | 0,886 | 0,721 | 0,912 | 0,806 | 1,07 | 0,07 | 6,2E-01 |
| Q9Z1S8 | GRB2-associated-binding protein 2 OS=Mus musculus OX=10090 GI | 1 | 1 | 1 | 1 | 665 | 73,2 | 8,31 | 0 | 1 | 0,713 | 0,839 | 0,843 | 1,107 | 0,78 | 0,7 | 1,044 | 0,738 | 1,07 | 0,07 | 6,2E-01 |
| Q80VP1 | Epsin-1 OS=Mus musculus OX=10090 GN=Epn1 PE=1 SV=3 | 11 | 6 | 8 | 6 | 575 | 60,2 | 4,81 | 18,52 | 6 | 0,962 | 1,031 | 0,957 | 1,08 | 1,012 | 0,988 | 0,987 | 0,979 | 1,02 | 0,02 | 6,2E-01 |
| Q99KY4 | Cyclin-G-associated kinase OS=Mus musculus OX=10090 GN=Gak P | 10 | 12 | 15 | 10 | 1305 | 143,6 | 5,73 | 19,4 | 12 | 1,017 | 1,072 | 0,887 | 1,158 | 0,969 | 1,089 | 1,108 | 1,107 | 0,97 | -0,03 | 6,2E-01 |
| P05132 | cAMP-dependent protein kinase catalytic subunit alpha OS=Mus m | 22 | 8 | 9 | 3 | 351 | 40,5 | 8,79 | 14,47 | 8 | 0,869 | 1,023 | 0,926 | 1,015 | 1,011 | 0,864 | 0,978 | 0,869 | 1,03 | 0,03 | 6,2E-01 |
| P24288 | Branched-chain-amino-acid aminotransferase, cytosolic OS=Mus m | 10 | 3 | 3 | 3 | 386 | 42,8 | 5,39 | 4,51 | 3 | 0,914 | 0,887 | 1,181 | 0,899 | 0,955 | 1,157 | 1,058 | 0,901 | 0,95 | -0,05 | 6,2E-01 |
| O88958 | Glucosamine-6-phosphate isomerase 1 OS=Mus musculus OX=1009 | 8 | 2 | 5 | 1 | 289 | 32,5 | 6,6 | 5,73 | 2 | 1,373 | 1,126 | 1,334 | 1,344 | 1,129 | 1,397 | 1,292 | 1,187 | 1,03 | 0,03 | 6,2E-01 |
| P97300-3 | Isoform 3 of Neuroplastin OS=Mus musculus OX=10090 GN=Nptn | 31 | 10 | 31 | 2 | 277 | 30,8 | 7,49 | 60,66 | 10 | 0,821 | 0,933 | 0,997 | 1,182 | 1,02 | 1,049 | 0,981 | 1,045 | 0,96 | -0,04 | 6,2E-01 |
| Q8R0A7 | Uncharacterized protein KIAA0513 OS=Mus musculus OX=10090 GI | 30 | 10 | 14 | 10 | 407 | 46,3 | 5,02 | 37,79 | 10 | 0,876 | 0,878 | 0,925 | 1,109 | 0,975 | 0,932 | 1,01 | 0,992 | 0,97 | -0,03 | 6,2E-01 |
| Q8C1E7 | Transmembrane protein 120A OS=Mus musculus OX=10090 GN=Tr | 5 | 1 | 1 | 1 | 343 | 40,7 | 8,88 | 2,85 | 1 | 1,778 | 2,375 | 2,025 | 1,405 | 1,02 | 2,551 | 1,515 | 1,704 | 1,12 | 0,12 | 6,2E-01 |
| Q9WV80 | Sorting nexin-1 OS=Mus musculus OX=10090 GN=Snx1 PE=1 SV=1 | 14 | 8 | 9 | 6 | 522 | 58,9 | 5,22 | 14,8 | 8 | 0,79 | 0,982 | 0,964 | 0,846 | 0,898 | 0,889 | 0,83 | 0,863 | 1,03 | 0,03 | 6,2E-01 |
| P10852 | 4F2 cell-surface antigen heavy chain OS=Mus musculus OX=10090 ( | 29 | 16 | 31 | 16 | 526 | 58,3 | 5,91 | 66,17 | 16 | 0,986 | 0,935 | 0,979 | 0,965 | 0,962 | 0,984 | 0,979 | 0,966 | 0,99 | -0,01 | 6,2E-01 |
| Q8R3S6 | Exocyst complex component 1 OS=Mus musculus OX=10090 GN=Ex | 9 | 7 | 7 | 7 | 894 | 101,8 | 6,52 | 10,88 | 7 | 1,058 | 1,032 | 0,855 | 1,051 | 1,054 | 0,826 | 0,991 | 0,982 | 1,04 | 0,04 | 6,2E-01 |
| P00688 | Pancreatic alpha-amylase OS=Mus musculus OX=10090 GN=Amy2 I | 2 | 1 | 1 | 1 | 508 | 57,3 | 7,28 | 3,49 | 1 | 0,878 | 1,42 | 1,035 | 1,006 | 0,684 | 1,529 | 0,994 | 0,64 | 1,13 | 0,13 | 6,2E-01 |
| Q9JLB2 | MAGUK p55 subfamily member 5 OS=Mus musculus OX=10090 GN | 4 | 2 | 2 | 2 | 675 | 77,2 | 6,09 | 3,85 | 2 | 0,576 | 1,092 | 1,055 | 1,291 | 1,064 | 1,337 | 0,948 | 1,027 | 0,92 | -0,08 | 6,2E-01 |
| O35453 | Serine protease hepsin OS=Mus musculus OX=10090 GN=Hpn PE=1 | 2 | 1 | 1 | 1 | 436 | 46,8 | 7,43 | 0 | 1 | 1,798 | 0,805 | 1,16 | 0,856 | 1,086 | 1,195 | 0,822 | 1,013 | 1,12 | 0,12 | 6,2E-01 |

| | | | | | | | | | | | | | | | | | | | | | |
|---|---|---|---|---|---|---|---|---|---|---|---|---|---|---|---|---|---|---|---|---|---|
| P97390 | Vacuolar protein sorting-associated protein 45 OS=Mus musculus C | 2 | 1 | 1 | 1 | 570 | 65 | 8,25 | 2,8 | 1 | 1,874 | 1,117 | 1,182 | 1,594 | 0,901 | 0,832 | 1,693 | 1,715 | 1,12 | 0,12 | 6,2E-01 |
| Q80TI0 | Protein Aster-B OS=Mus musculus OX=10090 GN=Gramd1b PE=1 S\ | 3 | 2 | 2 | 2 | 738 | 85,3 | 6,14 | 1,65 | 2 | 1,03 | 1,251 | 0,85 | 1,004 | 0,867 | 1,263 | 0,855 | 0,882 | 1,07 | 0,07 | 6,2E-01 |
| Q8BVL9 | Janus kinase and microtubule-interacting protein 1 OS=Mus muscu | 3 | 2 | 2 | 2 | 626 | 73,1 | 6,09 | 2,72 | 2 | 1,196 | 1,098 | 1,483 | 1,092 | 0,871 | 1,431 | 0,977 | 1,262 | 1,07 | 0,07 | 6,2E-01 |
| Q5GH64 | XK-related protein 7 OS=Mus musculus OX=10090 GN=Xkr7 PE=2 S\ | 7 | 3 | 3 | 3 | 580 | 64,3 | 8,73 | 2,59 | 3 | 1,461 | 0,931 | 0,741 | 0,946 | 0,966 | 0,978 | 1,076 | 0,697 | 1,10 | 0,10 | 6,2E-01 |
| Q80XA6 | RalBP1-associated Eps domain-containing protein 2 OS=Mus muscu | 11 | 4 | 4 | 4 | 521 | 57,7 | 7,53 | 4,36 | 4 | 1,706 | 1,423 | 1,156 | 1,004 | 0,775 | 1,553 | 1,134 | 1,356 | 1,10 | 0,10 | 6,2E-01 |
| P70195 | Proteasome subunit beta type-7 OS=Mus musculus OX=10090 GN= | 20 | 6 | 11 | 6 | 277 | 29,9 | 7,99 | 20,91 | 6 | 0,812 | 0,881 | 0,943 | 1,036 | 0,916 | 0,926 | 0,952 | 0,981 | 0,97 | -0,03 | 6,2E-01 |
| O08586 | Phosphatidylinositol 3,4,5-trisphosphate 3-phosphatase and dual-s | 7 | 3 | 4 | 3 | 403 | 47,1 | 6,37 | 10,05 | 3 | 1,052 | 1,041 | 1,132 | 1,075 | 1,135 | 1,101 | 1,004 | 1,137 | 0,98 | -0,02 | 6,2E-01 |
| A2AMZ4 | NADH dehydrogenase [ubiquinone] 1 alpha subcomplex assembly f | 11 | 1 | 1 | 1 | 74 | 7,8 | 9,55 | 2,13 | 1 | 0,795 | 0,741 | 1,174 | 0,823 | 1,105 | 0,666 | 1,06 | 0,991 | 0,92 | -0,08 | 6,2E-01 |
| P46061 | Ran GTPase-activating protein 1 OS=Mus musculus OX=10090 GN= | 4 | 2 | 2 | 2 | 589 | 63,5 | 4,68 | 2,96 | 2 | 1,137 | 1,231 | 1,075 | 0,841 | 0,796 | 1,18 | 0,845 | 1,186 | 1,07 | 0,07 | 6,2E-01 |
| P63028 | Translationally-controlled tumor protein OS=Mus musculus OX=10( | 24 | 4 | 7 | 4 | 172 | 19,5 | 4,86 | 9,88 | 4 | 0,96 | 0,923 | 0,937 | 0,873 | 0,859 | 1,213 | 0,883 | 0,913 | 0,95 | -0,05 | 6,2E-01 |
| Q8BW96 | Calcium/calmodulin-dependent protein kinase type 1D OS=Mus mu | 15 | 6 | 8 | 5 | 385 | 42,9 | 7,17 | 10,84 | 6 | 1,185 | 1,028 | 1,054 | 1,092 | 1,039 | 1,178 | 1,18 | 1,066 | 0,98 | -0,02 | 6,2E-01 |
| Q8JZN7 | Mitochondrial Rho GTPase 2 OS=Mus musculus OX=10090 GN=Rho | 5 | 2 | 2 | 1 | 620 | 69 | 6 | 2,98 | 2 | 1,573 | 1,415 | 1,756 | 1,215 | 0,865 | 1,933 | 1,058 | 1,553 | 1,10 | 0,10 | 6,3E-01 |
| Q791T5 | Mitochondrial carrier homolog 1 OS=Mus musculus OX=10090 GN= | 5 | 2 | 2 | 2 | 389 | 41,5 | 9,32 | 5,05 | 2 | 1,641 | 1,336 | 1,288 | 1,034 | 1 | 1,63 | 1,125 | 1,163 | 1,08 | 0,08 | 6,3E-01 |
| Q8BK75 | Elongator complex protein 6 OS=Mus musculus OX=10090 GN=Elp6 | 5 | 1 | 1 | 1 | 266 | 29,3 | 6,61 | 2,95 | 1 | 1,32 | 1,59 | 0,652 | 1,287 | 1,153 | 1,412 | 1,594 | 1,153 | 0,91 | -0,09 | 6,3E-01 |
| Q8BGV7 | BTB/POZ domain-containing adapter for CUL3-mediated RhoA degr | 3 | 1 | 1 | 1 | 329 | 36,4 | 6,99 | 0 | 1 | 1,249 | 0,788 | 1,28 | 1,568 | 1,316 | 0,817 | 1,258 | 1,092 | 1,09 | 0,09 | 6,3E-01 |
| Q9QZE5 | Coatomer subunit gamma-1 OS=Mus musculus OX=10090 GN=Cop | 1 | 1 | 1 | 1 | 874 | 97,5 | 5,35 | 2,57 | 1 | 1,621 | 1,218 | 1,005 | 1,163 | 1,212 | 1,304 | 1,128 | 1,076 | 1,06 | 0,06 | 6,3E-01 |
| P70202 | Latexin OS=Mus musculus OX=10090 GN=Lxn PE=1 SV=2 | 16 | 2 | 2 | 2 | 222 | 25,5 | 5,74 | 2,2 | 2 | 1,052 | 1,319 | 0,95 | 1,216 | 1,141 | 1,116 | 1,041 | 1,064 | 1,04 | 0,04 | 6,3E-01 |
| Q9DBJ1 | Phosphoglycerate mutase 1 OS=Mus musculus OX=10090 GN=Pgar | 63 | 14 | 29 | 9 | 254 | 28,8 | 7,18 | 69,54 | 14 | 0,952 | 1,064 | 0,922 | 1,039 | 0,951 | 0,987 | 0,955 | 1,009 | 1,02 | 0,02 | 6,3E-01 |
| O89032 | SH3 and PX domain-containing protein 2A OS=Mus musculus OX=1( | 1 | 1 | 1 | 1 | 1124 | 124,1 | 8,75 | 0 | 1 | 0,01 | 0,596 | 0,713 | 1,183 | 0,837 | 0,476 | 0,598 | 1,176 | 0,81 | -0,19 | 6,3E-01 |
| O35607 | Bone morphogenetic protein receptor type-2 OS=Mus musculus O> | 4 | 3 | 4 | 3 | 1038 | 114,9 | 6,23 | 2,79 | 3 | 1,029 | 1,396 | 1,253 | 1,14 | 1,098 | 1,087 | 1,296 | 1,149 | 1,04 | 0,04 | 6,3E-01 |
| Q9R1Q8 | Transgelin-3 OS=Mus musculus OX=10090 GN=Tagln3 PE=1 SV=1 | 39 | 9 | 14 | 8 | 199 | 22,5 | 7,33 | 32,78 | 9 | 1,011 | 0,943 | 0,951 | 1,073 | 0,985 | 1,011 | 0,984 | 1,073 | 0,98 | -0,02 | 6,3E-01 |
| Q9D071 | MMS19 nucleotide excision repair protein homolog OS=Mus muscu | 3 | 3 | 3 | 3 | 1031 | 113 | 6,21 | 3,49 | 3 | 0,903 | 0,935 | 1,088 | 0,816 | 0,92 | 0,771 | 0,838 | 1,046 | 1,05 | 0,05 | 6,3E-01 |
| Q8VE97 | Serine/arginine-rich splicing factor 4 OS=Mus musculus OX=10090 ( | 2 | 1 | 1 | 1 | 489 | 55,9 | 11,4 | 2,54 | 1 | 1,454 | 1,385 | 1,074 | 0,905 | 1,071 | 1,463 | 0,956 | 0,97 | 1,08 | 0,08 | 6,3E-01 |
| Q149F3 | Eukaryotic peptide chain release factor GTP-binding subunit ERF3B | 13 | 9 | 12 | 2 | 632 | 69,1 | 5,2 | 15,94 | 9 | 0,837 | 1,108 | 1,233 | 1,245 | 1,154 | 0,941 | 1,054 | 1,062 | 1,05 | 0,05 | 6,3E-01 |
| Q9D7B6 | Isobutyryl-CoA dehydrogenase, mitochondrial OS=Mus musculus O | 23 | 9 | 12 | 9 | 413 | 45 | 8,13 | 24,06 | 9 | 1,063 | 0,908 | 1,037 | 0,978 | 1,067 | 0,954 | 0,89 | 0,973 | 1,03 | 0,03 | 6,3E-01 |
| Q68FD9 | UPF0606 protein KIAA1549 OS=Mus musculus OX=10090 GN=Kiaa1 | 3 | 6 | 7 | 6 | 1940 | 209,1 | 6,1 | 7,82 | 6 | 1,167 | 1,025 | 0,96 | 1,03 | 0,964 | 1,093 | 0,978 | 1,04 | 1,03 | 0,03 | 6,3E-01 |
| Q80W54 | CAAX prenyl protease 1 homolog OS=Mus musculus OX=10090 GN= | 2 | 1 | 1 | 1 | 475 | 54,7 | 6,95 | 3,12 | 1 | 0,95 | 0,976 | 0,953 | 0,867 | 1,026 | 1,099 | 1,034 | 0,751 | 0,96 | -0,04 | 6,3E-01 |
| O54991 | Contactin-associated protein 1 OS=Mus musculus OX=10090 GN=C | 10 | 11 | 17 | 11 | 1385 | 156,2 | 7,06 | 32,23 | 11 | 1,039 | 1,08 | 1,115 | 1,113 | 1,102 | 1,121 | 0,992 | 1,064 | 1,02 | 0,02 | 6,3E-01 |
| Q8CDG3 | Deubiquitinating protein VCIP135 OS=Mus musculus OX=10090 GN | 12 | 12 | 13 | 12 | 1220 | 134,4 | 7,17 | 21,02 | 12 | 0,969 | 0,958 | 0,994 | 0,961 | 0,976 | 1,086 | 0,988 | 0,908 | 0,98 | -0,02 | 6,3E-01 |
| Q9D338 | 39S ribosomal protein L19, mitochondrial OS=Mus musculus OX=1( | 11 | 3 | 4 | 3 | 292 | 33,6 | 9,44 | 2,09 | 3 | 1,233 | 0,879 | 0,925 | 0,795 | 0,722 | 0,837 | 0,793 | 1,193 | 1,08 | 0,08 | 6,3E-01 |
| P70452 | Syntaxin-4 OS=Mus musculus OX=10090 GN=Stx4 PE=1 SV=1 | 5 | 2 | 2 | 2 | 298 | 34,1 | 6,14 | 2,22 | 2 | 1,025 | 1,134 | 0,994 | 1,162 | 1,009 | 1,015 | 0,708 | 1,318 | 1,07 | 0,07 | 6,3E-01 |
| O70451 | Monocarboxylate transporter 2 OS=Mus musculus OX=10090 GN=S | 1 | 1 | 1 | 1 | 484 | 52,6 | 9,31 | 1,83 | 1 | 0,616 | 0,87 | 1,161 | 1,066 | 1,1 | 0,768 | 0,942 | 1,216 | 0,92 | -0,08 | 6,3E-01 |
| Q99MK8 | Beta-adrenergic receptor kinase 1 OS=Mus musculus OX=10090 GN | 9 | 6 | 7 | 4 | 689 | 79,6 | 7,28 | 20,98 | 6 | 0,963 | 1,304 | 1,025 | 1,086 | 1,019 | 1,068 | 1,019 | 1,115 | 1,04 | 0,04 | 6,3E-01 |
| Q5EG47 | 5'-AMP-activated protein kinase catalytic subunit alpha-1 OS=Mus | 8 | 3 | 3 | 2 | 559 | 63,9 | 8,12 | 8,7 | 3 | 0,773 | 1,007 | 0,73 | 1,024 | 1,029 | 0,981 | 1,009 | 0,726 | 0,94 | -0,06 | 6,3E-01 |
| Q80Y24 | Prickle-like protein 2 OS=Mus musculus OX=10090 GN=Prickle2 PE= | 15 | 9 | 10 | 9 | 845 | 95,7 | 7,27 | 14,81 | 9 | 1,099 | 1,361 | 1,11 | 1,172 | 0,847 | 1,253 | 1,149 | 1,262 | 1,05 | 0,05 | 6,3E-01 |
| Q8R366 | Immunoglobulin superfamily member 8 OS=Mus musculus OX=100 | 14 | 9 | 15 | 9 | 611 | 65 | 7,99 | 40,81 | 9 | 1,101 | 0,777 | 1,134 | 0,928 | 1,12 | 0,956 | 1,132 | 0,93 | 0,95 | -0,05 | 6,3E-01 |
| P59325 | Eukaryotic translation initiation factor 5 OS=Mus musculus OX=100 | 7 | 3 | 4 | 3 | 429 | 48,9 | 5,52 | 8,15 | 3 | 0,911 | 1,097 | 1,071 | 1,074 | 0,984 | 1,178 | 1,112 | 1,005 | 0,97 | -0,03 | 6,3E-01 |
| P27600 | Guanine nucleotide-binding protein subunit alpha-12 OS=Mus mus | 5 | 2 | 2 | 1 | 379 | 44,1 | 9,83 | 5,16 | 2 | 1,281 | 0,991 | 0,875 | 0,686 | 1,023 | 1,093 | 0,473 | 0,869 | 1,11 | 0,11 | 6,3E-01 |
| Q9Z1K7 | Adenomatous polyposis coli protein 2 OS=Mus musculus OX=1009( | 1 | 2 | 2 | 2 | 2274 | 243 | 8,88 | 2,3 | 2 | 1,127 | 0,888 | 0,561 | 1,122 | 0,884 | 0,944 | 1,112 | 1,045 | 0,93 | -0,07 | 6,3E-01 |
| Q80TE4 | Signal-induced proliferation-associated 1-like protein 2 OS=Mus mu | 2 | 3 | 3 | 2 | 1722 | 189,3 | 6,79 | 6,52 | 3 | 1,865 | 1,238 | 0,885 | 0,927 | 1,079 | 1,187 | 0,907 | 1,262 | 1,11 | 0,11 | 6,3E-01 |
| Q9D0K2 | Succinyl-CoA:3-ketoacid coenzyme A transferase 1, mitochondrial ( | 29 | 14 | 33 | 14 | 520 | 56 | 8,53 | 65,46 | 14 | 0,951 | 1,005 | 1,016 | 0,957 | 0,951 | 0,988 | 1,019 | 1,017 | 0,99 | -0,01 | 6,3E-01 |
| Q80WQ2 | Protein VAC14 homolog OS=Mus musculus OX=10090 GN=Vac14 P | 11 | 7 | 8 | 7 | 782 | 88 | 6,13 | 12,3 | 7 | 1,119 | 0,925 | 1,086 | 1,059 | 1,016 | 1,073 | 1,034 | 0,971 | 1,02 | 0,02 | 6,4E-01 |
| Q9QZM0 | Ubiquilin-2 OS=Mus musculus OX=10090 GN=Ubqln2 PE=1 SV=2 | 8 | 4 | 7 | 4 | 638 | 67,3 | 5,22 | 22,09 | 4 | 1,028 | 1,039 | 1,076 | 1,071 | 0,906 | 1,126 | 1,028 | 1,059 | 1,02 | 0,02 | 6,4E-01 |
| Q9DC51 | Guanine nucleotide-binding protein G(i) subunit alpha OS=Mus mu | 23 | 8 | 8 | 6 | 354 | 40,5 | 5,69 | 23,24 | 8 | 0,986 | 0,935 | 0,983 | 1,104 | 1,003 | 0,856 | 1,071 | 0,963 | 1,03 | 0,03 | 6,4E-01 |
| Q99JT9 | 1,2-dihydroxy-3-keto-5-methylthiopentene dioxygenase OS=Mus m | 5 | 1 | 1 | 1 | 179 | 21,5 | 5,5 | 0 | 1 | 0,402 | 1,188 | 0,904 | 2,257 | 0,994 | 1,063 | 0,861 | 1,046 | 1,20 | 0,20 | 6,4E-01 |
| P62830 | 60S ribosomal protein L23 OS=Mus musculus OX=10090 GN=Rpl23 | 28 | 3 | 3 | 3 | 140 | 14,9 | 10,51 | 1,71 | 3 | 1,588 | 1,114 | 1,799 | 1,228 | 1,275 | 1,69 | 1,25 | 1,112 | 1,08 | 0,08 | 6,4E-01 |
| Q6ZWV3 | 60S ribosomal protein L10 OS=Mus musculus OX=10090 GN=Rpl10 | 15 | 3 | 3 | 3 | 214 | 24,6 | 10,08 | 7,86 | 3 | 0,759 | 0,59 | 1,032 | 1,111 | 1,014 | 1,026 | 0,835 | 0,876 | 0,93 | -0,07 | 6,4E-01 |
| Q80WC7 | Arf-GAP domain and FG repeat-containing protein 2 OS=Mus musc | 3 | 2 | 2 | 2 | 479 | 48,9 | 9,1 | 2,15 | 2 | 1,163 | 1,019 | 0,913 | 1,04 | 1,054 | 0,919 | 1,033 | 1,011 | 1,03 | 0,03 | 6,4E-01 |
| Q9Z1T1 | AP-3 complex subunit beta-1 OS=Mus musculus OX=10090 GN=Ap3 | 3 | 3 | 5 | 1 | 1105 | 122,7 | 5,66 | 8,35 | 3 | 1,057 | 1,214 | 0,863 | 0,918 | 0,88 | 0,846 | 0,993 | 1,131 | 1,05 | 0,05 | 6,4E-01 |
| Q9Z0J4 | Nitric oxide synthase, brain OS=Mus musculus OX=10090 GN=Nos1 | 6 | 8 | 9 | 8 | 1429 | 160,4 | 7,15 | 9,49 | 8 | 0,932 | 1,156 | 0,966 | 0,934 | 0,734 | 1,072 | 1,068 | 0,923 | 1,05 | 0,05 | 6,4E-01 |
| Q925E7 | Serine/threonine-protein phosphatase 2A 55 kDa regulatory subun | 12 | 7 | 7 | 1 | 453 | 51,9 | 6,49 | 12,83 | 7 | 1,139 | 1,216 | 1,046 | 1,556 | 1,062 | 1,359 | 1,138 | 1,143 | 1,05 | 0,05 | 6,4E-01 |
| Q9QWY8 | Arf-GAP with SH3 domain, ANK repeat and PH domain-containing p | 10 | 11 | 12 | 10 | 1147 | 127,3 | 7,64 | 12,36 | 11 | 0,926 | 1,005 | 0,969 | 0,983 | 0,944 | 1,009 | 0,937 | 0,946 | 1,01 | 0,01 | 6,4E-01 |
| O35379 | Multidrug resistance-associated protein 1 OS=Mus musculus OX=1( | 2 | 2 | 2 | 2 | 1528 | 171,1 | 7,36 | 2,04 | 2 | 0,01 | 1,186 | 0,725 | 0,8 | 0,93 | 0,743 | 1,026 | 0,552 | 0,84 | -0,16 | 6,4E-01 |
| P37804 | Transgelin OS=Mus musculus OX=10090 GN=Tagln PE=1 SV=3 | 14 | 3 | 5 | 2 | 201 | 22,6 | 8,81 | 13,56 | 3 | 1,333 | 1,338 | 1,603 | 0,817 | 0,732 | 1,685 | 0,999 | 1,16 | 1,11 | 0,11 | 6,4E-01 |
| Q6NVE9 | Protein phosphatase PTC7 homolog OS=Mus musculus OX=10090 ( | 10 | 2 | 2 | 2 | 310 | 33 | 5,27 | 3,89 | 2 | 1,307 | 1,11 | 1,003 | 1,052 | 0,78 | 1,298 | 1,013 | 1,128 | 1,06 | 0,06 | 6,4E-01 |
| Q8BT60 | Copine-3 OS=Mus musculus OX=10090 GN=Cpne3 PE=1 SV=2 | 2 | 1 | 1 | 1 | 533 | 59,5 | 5,78 | 0 | 1 | 1,263 | 1,689 | 0,652 | 0,82 | 1,31 | 0,787 | 0,754 | 1,044 | 1,14 | 0,14 | 6,4E-01 |
| Q9CWS0 | N(G),N(G)-dimethylarginine dimethylaminohydrolase 1 OS=Mus mu | 32 | 8 | 9 | 8 | 285 | 31,4 | 5,97 | 19,99 | 8 | 0,951 | 1,053 | 1,031 | 0,986 | 0,941 | 1,042 | 1,06 | 1,049 | 0,98 | -0,02 | 6,4E-01 |
| Q68FF6 | ARF GTPase-activating protein GIT1 OS=Mus musculus OX=10090 G | 23 | 14 | 16 | 14 | 770 | 85,2 | 6,93 | 33,87 | 14 | 0,907 | 0,892 | 0,881 | 1,116 | 1,01 | 0,896 | 1,01 | 1,004 | 0,97 | -0,03 | 6,4E-01 |
| P47740 | Aldehyde dehydrogenase family 3 member A2 OS=Mus musculus C | 16 | 7 | 11 | 7 | 484 | 53,9 | 8,35 | 25,16 | 7 | 1,335 | 1,191 | 0,936 | 0,915 | 0,989 | 1,187 | 0,957 | 1,018 | 1,05 | 0,05 | 6,4E-01 |
| P47746 | Cannabinoid receptor 1 OS=Mus musculus OX=10090 GN=Cnr1 PE= | 2 | 1 | 1 | 1 | 473 | 52,8 | 8,27 | 0 | 1 | 2,781 | 2,856 | 1,923 | 1,588 | 1,625 | 2,182 | 2,525 | 2,092 | 1,09 | 0,09 | 6,4E-01 |
| P0DP60 | Ly-6/neurotoxin-like protein 1 OS=Mus musculus OX=10090 GN=Ly | 15 | 1 | 1 | 1 | 116 | 12,8 | 8,19 | 2,72 | 1 | 1,307 | 1,049 | 1,168 | 1,033 | 1,21 | 1,284 | 0,701 | 1,076 | 1,07 | 0,07 | 6,4E-01 |
| Q6QWF9 | Calcium/calmodulin-dependent protein kinase II inhibitor 1 OS=Mu | 36 | 2 | 2 | 2 | 78 | 8,5 | 5,45 | 7,07 | 2 | 0,876 | 0,953 | 0,848 | 0,899 | 0,906 | 0,841 | 1,002 | 0,906 | 0,98 | -0,02 | 6,4E-01 |

| | | | | | | | | | | | | | | | | | | | | | |
|---|---|---|---|---|---|---|---|---|---|---|---|---|---|---|---|---|---|---|---|---|---|
| Q52KR3 | Protein prune homolog 2 OS=Mus musculus OX=10090 GN=Prune2 | 0 | 1 | 1 | 1 | 3084 | 339,3 | 4,46 | 2,4 | 1 | 1,406 | 0,613 | 1,161 | 1,2 | 1,323 | 1,124 | 1,12 | 1,161 | 0,93 | -0,07 | 6,4E-01 |
| Q61234 | Alpha-1-syntrophin OS=Mus musculus OX=10090 GN=Snta1 PE=1 S | 4 | 2 | 2 | 2 | 503 | 53,6 | 6,86 | 3,81 | 2 | 1,04 | 0,93 | 0,931 | 1,107 | 1,035 | 0,874 | 1,112 | 1,131 | 0,97 | -0,03 | 6,4E-01 |
| Q62446 | Peptidyl-prolyl cis-trans isomerase FKBP3 OS=Mus musculus OX=10 | 17 | 3 | 4 | 3 | 224 | 25,1 | 9,28 | 2,53 | 3 | 1,306 | 1,207 | 1,032 | 1,007 | 1,069 | 1,211 | 1,114 | 0,99 | 1,04 | 0,04 | 6,4E-01 |
| Q6ZPE2 | Myotubularin-related protein 5 OS=Mus musculus OX=10090 GN=S | 8 | 13 | 15 | 13 | 1867 | 208,6 | 7,12 | 22,34 | 13 | 1,473 | 1,077 | 1,307 | 1,015 | 0,981 | 1,439 | 0,961 | 1,188 | 1,07 | 0,07 | 6,4E-01 |
| Q9CPT4 | Myeloid-derived growth factor OS=Mus musculus OX=10090 GN=M | 7 | 1 | 2 | 1 | 166 | 18 | 6,79 | 2,79 | 1 | 0,828 | 0,707 | 0,763 | 1,011 | 0,885 | 0,632 | 1,041 | 0,97 | 0,94 | -0,06 | 6,4E-01 |
| Q9D898 | Actin-related protein 2/3 complex subunit 5-like protein OS=Mus m | 24 | 4 | 6 | 4 | 153 | 17 | 6,8 | 13,04 | 4 | 0,96 | 0,942 | 0,961 | 1,042 | 0,863 | 1,072 | 1,027 | 1,046 | 0,97 | -0,03 | 6,4E-01 |
| Q8BYH7 | TBC1 domain family member 17 OS=Mus musculus OX=10090 GN= | 3 | 2 | 2 | 2 | 645 | 72,8 | 5,25 | 0 | 2 | 1,043 | 1,225 | 1,129 | 1,609 | 0,964 | 1,508 | 1,414 | 1,47 | 0,93 | -0,07 | 6,4E-01 |
| Q8BU30 | Isoleucine--tRNA ligase, cytoplasmic OS=Mus musculus OX=10090 ( | 5 | 5 | 6 | 5 | 1262 | 144,2 | 6,55 | 7,38 | 5 | 0,543 | 1,611 | 0,676 | 0,902 | 0,995 | 0,543 | 0,972 | 0,708 | 1,16 | 0,16 | 6,4E-01 |
| Q9D7I5 | Phospholysine phosphohistidine inorganic pyrophosphate phospha | 10 | 2 | 2 | 2 | 270 | 29,1 | 5,1 | 0 | 2 | 0,808 | 1,195 | 1,584 | 1,077 | 0,904 | 0,925 | 1,253 | 1,217 | 1,08 | 0,08 | 6,4E-01 |
| Q9CR46 | Spindle and kinetochore-associated protein 2 OS=Mus musculus OX | 6 | 1 | 1 | 1 | 120 | 13,7 | 5,55 | 2,14 | 1 | 1,528 | 1,462 | 1,283 | 0,813 | 0,803 | 1,493 | 0,997 | 1,35 | 1,10 | 0,10 | 6,4E-01 |
| P60122 | RuvB-like 1 OS=Mus musculus OX=10090 GN=Ruvbl1 PE=1 SV=1 | 23 | 8 | 8 | 8 | 456 | 50,2 | 6,42 | 12,35 | 8 | 1,025 | 1,339 | 1,332 | 1,158 | 1,059 | 1,371 | 1,099 | 1,123 | 1,04 | 0,04 | 6,4E-01 |
| P49813 | Tropomodulin-1 OS=Mus musculus OX=10090 GN=Tmod1 PE=1 SV= | 24 | 7 | 7 | 6 | 359 | 40,4 | 5,1 | 8,6 | 7 | 1,253 | 0,985 | 1,058 | 1,086 | 1,128 | 1,162 | 1,101 | 1,105 | 0,97 | -0,03 | 6,4E-01 |
| E9PUL5 | Proline-rich transmembrane protein 2 OS=Mus musculus OX=1009( | 21 | 6 | 8 | 6 | 346 | 35,9 | 4,63 | 8,57 | 6 | 0,498 | 0,782 | 0,848 | 1,002 | 1,038 | 0,425 | 0,975 | 1,05 | 0,90 | -0,10 | 6,4E-01 |
| Q60829 | Protein phosphatase 1 regulatory subunit 1B OS=Mus musculus OX | 23 | 3 | 7 | 3 | 194 | 21,8 | 4,65 | 2,25 | 3 | 0,455 | 1,247 | 0,808 | 1,134 | 0,871 | 1,349 | 0,823 | 1,019 | 0,90 | -0,10 | 6,4E-01 |
| O89053 | Coronin-1A OS=Mus musculus OX=10090 GN=Coro1a PE=1 SV=5 | 35 | 18 | 24 | 17 | 461 | 51 | 6,48 | 52,96 | 18 | 0,884 | 0,856 | 0,966 | 1,058 | 1,023 | 0,902 | 0,996 | 0,946 | 0,97 | -0,03 | 6,4E-01 |
| P15105 | Glutamine synthetase OS=Mus musculus OX=10090 GN=Glul PE=1 S | 36 | 17 | 39 | 17 | 373 | 42,1 | 7,08 | 106,37 | 17 | 1,023 | 0,936 | 0,996 | 0,986 | 1,047 | 1,037 | 0,908 | 1,021 | 0,98 | -0,02 | 6,4E-01 |
| Q8VDK1 | Deaminated glutathione amidase OS=Mus musculus OX=10090 GN | 23 | 6 | 6 | 6 | 323 | 35,7 | 7,94 | 10,24 | 6 | 1,085 | 0,979 | 0,994 | 0,988 | 0,997 | 0,939 | 1,049 | 0,996 | 1,02 | 0,02 | 6,4E-01 |
| P58404 | Striatin-4 OS=Mus musculus OX=10090 GN=Strn4 PE=1 SV=2 | 6 | 5 | 6 | 4 | 760 | 81,6 | 5,38 | 15,83 | 5 | 1,143 | 1,157 | 0,956 | 0,822 | 0,851 | 0,924 | 1,018 | 1,097 | 1,05 | 0,05 | 6,4E-01 |
| Q8BMK4 | Cytoskeleton-associated protein 4 OS=Mus musculus OX=10090 GN | 13 | 7 | 8 | 7 | 575 | 63,7 | 5,64 | 16,48 | 7 | 1,045 | 1,192 | 1,196 | 1,041 | 1,084 | 1,233 | 0,951 | 1,065 | 1,03 | 0,03 | 6,4E-01 |
| Q8BMD5 | Cytidine and dCMP deaminase domain-containing protein 1 OS=Mu | 4 | 1 | 1 | 1 | 523 | 59 | 7,8 | 0 | 1 | 3,27 | 1,575 | 1,662 | 0,799 | 1,103 | 1,725 | 1,652 | 1,774 | 1,17 | 0,17 | 6,4E-01 |
| Q61735 | Leukocyte surface antigen CD47 OS=Mus musculus OX=10090 GN= | 4 | 1 | 3 | 1 | 303 | 33,1 | 8,63 | 12,05 | 1 | 1,034 | 1,072 | 1,05 | 0,968 | 0,957 | 0,986 | 1,065 | 1,05 | 1,02 | 0,02 | 6,5E-01 |
| P55066 | Neurocan core protein OS=Mus musculus OX=10090 GN=Ncan PE= | 3 | 4 | 4 | 4 | 1268 | 137,1 | 5,72 | 7,35 | 4 | 1,044 | 0,85 | 0,982 | 0,869 | 1,129 | 0,977 | 0,955 | 0,832 | 0,96 | -0,04 | 6,5E-01 |
| Q9CR84 | ATP synthase F(0) complex subunit C1, mitochondrial OS=Mus mus | 5 | 1 | 3 | 1 | 136 | 14,2 | 10,01 | 7,14 | 1 | 0,54 | 1 | 0,851 | 0,915 | 1,005 | 0,736 | 1,078 | 0,745 | 0,93 | -0,07 | 6,5E-01 |
| Q3U1J4 | DNA damage-binding protein 1 OS=Mus musculus OX=10090 GN=D | 16 | 17 | 20 | 17 | 1140 | 126,8 | 5,26 | 40,66 | 17 | 1,008 | 1,033 | 1,033 | 0,998 | 0,954 | 1,09 | 1,027 | 1,06 | 0,99 | -0,01 | 6,5E-01 |
| Q9CQ91 | NADH dehydrogenase [ubiquinone] 1 alpha subcomplex subunit 3 ( | 8 | 1 | 1 | 1 | 84 | 9,3 | 8,47 | 2,36 | 1 | 1,147 | 0,881 | 0,875 | 0,951 | 0,971 | 0,868 | 0,95 | 0,935 | 1,03 | 0,03 | 6,5E-01 |
| O54829 | Regulator of G-protein signaling 7 OS=Mus musculus OX=10090 GN | 25 | 12 | 17 | 8 | 469 | 54,8 | 8,13 | 43,59 | 12 | 1,222 | 1,042 | 1,01 | 0,995 | 0,969 | 1,112 | 0,989 | 1,078 | 1,03 | 0,03 | 6,5E-01 |
| P42225 | Signal transducer and activator of transcription 1 OS=Mus musculu | 6 | 4 | 4 | 4 | 749 | 87,1 | 5,58 | 2,21 | 4 | 0,839 | 1,125 | 1,045 | 0,83 | 0,814 | 1,28 | 0,932 | 1,052 | 0,94 | -0,06 | 6,5E-01 |
| P97478 | 5-demethoxyubiquinone hydroxylase, mitochondrial OS=Mus musc | 4 | 1 | 2 | 1 | 217 | 24 | 7,17 | 3,08 | 1 | 1,158 | 1,06 | 1,048 | 0,971 | 0,974 | 1,191 | 0,749 | 1,121 | 1,05 | 0,05 | 6,5E-01 |
| P62267 | 40S ribosomal protein S23 OS=Mus musculus OX=10090 GN=Rps23 | 13 | 2 | 3 | 2 | 143 | 15,8 | 10,49 | 5,02 | 2 | 0,947 | 0,788 | 1,02 | 0,85 | 0,909 | 0,922 | 0,947 | 0,927 | 0,97 | -0,03 | 6,5E-01 |
| Q9Z127 | Large neutral amino acids transporter small subunit 1 OS=Mus mus | 6 | 3 | 4 | 3 | 512 | 55,8 | 7,9 | 9,82 | 3 | 0,646 | 1,364 | 0,758 | 0,781 | 0,681 | 0,856 | 0,674 | 0,994 | 1,11 | 0,11 | 6,5E-01 |
| Q9CZW4 | Long-chain-fatty-acid--CoA ligase 3 OS=Mus musculus OX=10090 G | 3 | 2 | 3 | 2 | 720 | 80,4 | 8,54 | 2,56 | 2 | 0,955 | 0,795 | 0,847 | 1,085 | 0,978 | 0,8 | 0,915 | 0,845 | 1,04 | 0,04 | 6,5E-01 |
| P52503 | NADH dehydrogenase [ubiquinone] iron-sulfur protein 6, mitochon | 66 | 8 | 12 | 8 | 116 | 13 | 8,65 | 29,34 | 8 | 0,982 | 0,777 | 1,05 | 1,04 | 1,105 | 0,987 | 0,998 | 0,904 | 0,96 | -0,04 | 6,5E-01 |
| Q80WG5 | Volume-regulated anion channel subunit LRRC8A OS=Mus musculu | 7 | 4 | 4 | 4 | 810 | 94,1 | 7,94 | 9,62 | 4 | 1,325 | 1,341 | 1,219 | 1,1 | 0,955 | 1,488 | 1,143 | 1,162 | 1,05 | 0,05 | 6,5E-01 |
| O88342 | WD repeat-containing protein 1 OS=Mus musculus OX=10090 GN=\ | 27 | 14 | 18 | 14 | 606 | 66,4 | 6,6 | 47,18 | 14 | 1,11 | 1,014 | 0,977 | 0,965 | 1,022 | 1,003 | 0,956 | 1,016 | 1,02 | 0,02 | 6,5E-01 |
| Q3UH93 | Plexin-D1 OS=Mus musculus OX=10090 GN=Plxnd1 PE=1 SV=1 | 6 | 10 | 10 | 10 | 1925 | 211,5 | 7,11 | 9,31 | 10 | 1,083 | 1,079 | 0,922 | 0,953 | 0,875 | 1,038 | 0,974 | 1,041 | 1,03 | 0,03 | 6,5E-01 |
| P40336 | Vacuolar protein sorting-associated protein 26A OS=Mus musculus | 8 | 2 | 4 | 2 | 327 | 38,1 | 6,57 | 11,48 | 2 | 1,33 | 1,155 | 0,968 | 1,031 | 0,957 | 1,299 | 1,026 | 0,989 | 1,05 | 0,05 | 6,5E-01 |
| P59823 | Interleukin-1 receptor accessory protein-like 1 OS=Mus musculus C | 7 | 3 | 3 | 3 | 695 | 79,6 | 6,32 | 9,04 | 3 | 1,392 | 1,211 | 1,29 | 1,271 | 0,86 | 1,598 | 1,141 | 1,265 | 1,06 | 0,06 | 6,5E-01 |
| P55821 | Stathmin-2 OS=Mus musculus OX=10090 GN=Stmn2 PE=1 SV=1 | 16 | 4 | 6 | 1 | 179 | 20,8 | 8,32 | 10,86 | 4 | 1,024 | 0,979 | 0,576 | 0,873 | 0,896 | 0,783 | 0,909 | 1,091 | 0,94 | -0,06 | 6,5E-01 |
| Q99KE1 | NAD-dependent malic enzyme, mitochondrial OS=Mus musculus O | 16 | 8 | 8 | 7 | 589 | 65,8 | 7,61 | 16,65 | 8 | 1,282 | 1,099 | 1,057 | 1,013 | 1,04 | 1,18 | 1,152 | 1,212 | 0,97 | -0,03 | 6,5E-01 |
| O35954 | Membrane-associated phosphatidylinositol transfer protein 1 OS=M | 9 | 9 | 10 | 9 | 1243 | 134,9 | 6,06 | 14,93 | 9 | 1,124 | 0,992 | 0,974 | 1,096 | 1,098 | 1,091 | 1,065 | 1,012 | 0,98 | -0,02 | 6,5E-01 |
| P70336 | Rho-associated protein kinase 2 OS=Mus musculus OX=10090 GN=I | 26 | 36 | 49 | 36 | 1388 | 160,5 | 5,99 | 104,89 | 36 | 1,003 | 0,96 | 1,047 | 1,029 | 0,992 | 1,027 | 1,027 | 1,033 | 0,99 | -0,01 | 6,5E-01 |
| O35685 | Nuclear migration protein nudC OS=Mus musculus OX=10090 GN=I | 31 | 10 | 12 | 10 | 332 | 38,3 | 5,26 | 19,54 | 10 | 1,203 | 1,21 | 1,051 | 1,061 | 0,899 | 1,251 | 1,085 | 1,129 | 1,04 | 0,04 | 6,5E-01 |
| Q9CRA7 | ATP synthase subunit s, mitochondrial OS=Mus musculus OX=1009( | 4 | 1 | 1 | 1 | 200 | 23,3 | 8,06 | 2,67 | 1 | 0,929 | 1,098 | 1,039 | 1,078 | 1,032 | 1,042 | 1,187 | 0,991 | 0,97 | -0,03 | 6,5E-01 |
| P20060 | Beta-hexosaminidase subunit beta OS=Mus musculus OX=10090 GI | 10 | 5 | 6 | 5 | 536 | 61,1 | 8,12 | 12,58 | 5 | 1,073 | 1,012 | 0,956 | 0,96 | 1,077 | 0,975 | 1,031 | 0,986 | 0,98 | -0,02 | 6,5E-01 |
| Q9JJV5 | Voltage-dependent calcium channel gamma-3 subunit OS=Mus mu | 6 | 3 | 4 | 1 | 315 | 35,5 | 9,48 | 10,34 | 3 | 0,895 | 0,637 | 0,793 | 0,819 | 0,846 | 0,535 | 1,02 | 0,973 | 0,93 | -0,07 | 6,5E-01 |
| B2RXC1 | Trafficking protein particle complex subunit 11 OS=Mus musculus ( | 5 | 4 | 4 | 4 | 1133 | 128,3 | 7,58 | 7,31 | 4 | 1,508 | 1,009 | 0,916 | 1,015 | 1,255 | 1,204 | 1,056 | 1,198 | 0,94 | -0,06 | 6,5E-01 |
| Q9QUP5 | Hyaluronan and proteoglycan link protein 1 OS=Mus musculus OX= | 7 | 2 | 2 | 2 | 356 | 40,5 | 7,8 | 4,36 | 2 | 1,552 | 1,044 | 0,788 | 0,821 | 1,138 | 1,618 | 1,006 | 0,891 | 0,90 | -0,10 | 6,5E-01 |
| Q9Z1X4 | Interleukin enhancer-binding factor 3 OS=Mus musculus OX=10090 | 3 | 2 | 2 | 2 | 898 | 96 | 8,76 | 3,57 | 2 | 1,221 | 0,688 | 1,457 | 1,035 | 0,945 | 1,043 | 1,348 | 0,66 | 1,10 | 0,10 | 6,6E-01 |
| Q91WJ8 | Far upstream element-binding protein 1 OS=Mus musculus OX=100 | 1 | 1 | 1 | 1 | 651 | 68,5 | 7,93 | 2,04 | 1 | 0,526 | 0,825 | 0,652 | 0,755 | 0,647 | 0,98 | 0,677 | 0,649 | 0,93 | -0,07 | 6,6E-01 |
| Q8BK63 | Casein kinase I isoform alpha OS=Mus musculus OX=10090 GN=Csn | 13 | 4 | 4 | 4 | 337 | 38,9 | 9,57 | 3,33 | 4 | 1,039 | 1,019 | 1,018 | 1,012 | 0,895 | 1,139 | 1 | 1,176 | 0,97 | -0,03 | 6,6E-01 |
| Q9D9M0 | Serine protease 52 OS=Mus musculus OX=10090 GN=Prss52 PE=2 S | 2 | 1 | 3 | 1 | 321 | 35,9 | 8,95 | 6,26 | 1 | 1,281 | 0,819 | 1,084 | 1,006 | 0,991 | 1,027 | 0,831 | 1,775 | 0,91 | -0,09 | 6,6E-01 |
| Q9EPL8 | Importin-7 OS=Mus musculus OX=10090 GN=Ipo7 PE=1 SV=2 | 10 | 8 | 9 | 8 | 1038 | 119,4 | 4,82 | 20 | 8 | 1,066 | 1,033 | 1,045 | 0,986 | 0,964 | 1,103 | 1,044 | 1,085 | 0,98 | -0,02 | 6,6E-01 |
| Q8WTY4 | Anamorsin OS=Mus musculus OX=10090 GN=Ciapin1 PE=1 SV=1 | 13 | 3 | 3 | 3 | 309 | 33,4 | 5,2 | 2,27 | 3 | 1,017 | 0,917 | 1,017 | 0,966 | 1,008 | 1,14 | 0,937 | 0,933 | 0,97 | -0,03 | 6,6E-01 |
| Q7TQG1 | Pleckstrin homology domain-containing family A member 6 OS=Mu | 1 | 1 | 1 | 1 | 1173 | 131,3 | 8,97 | 2,44 | 1 | 1,606 | 2,38 | 1,987 | 1,513 | 1,798 | 2,075 | 1,338 | 1,807 | 1,07 | 0,07 | 6,6E-01 |
| Q9CQY5 | Magnesium transporter protein 1 OS=Mus musculus OX=10090 GN | 6 | 2 | 2 | 2 | 335 | 37,9 | 9,76 | 0 | 2 | 1,063 | 1,711 | 1,039 | 0,99 | 1,045 | 1,046 | 1,332 | 1,034 | 1,08 | 0,08 | 6,6E-01 |
| Q6PA06 | Atlastin-2 OS=Mus musculus OX=10090 GN=Atl2 PE=1 SV=1 | 3 | 2 | 3 | 2 | 583 | 66,2 | 5,43 | 4,97 | 2 | 1,024 | 1,004 | 0,966 | 0,941 | 0,95 | 1,057 | 0,952 | 1,039 | 0,98 | -0,02 | 6,6E-01 |
| Q9QX11 | Cytohesin-1 OS=Mus musculus OX=10090 GN=Cyth1 PE=1 SV=2 | 9 | 2 | 2 | 1 | 398 | 46,2 | 5,63 | 0 | 2 | 1,004 | 1,065 | 1,154 | 0,988 | 0,959 | 1,09 | 1,246 | 0,684 | 1,06 | 0,06 | 6,6E-01 |
| Q60700 | Mitogen-activated protein kinase kinase kinase 12 OS=Mus muscul | 2 | 2 | 2 | 2 | 888 | 96 | 6,24 | 4,99 | 2 | 1,038 | 0,888 | 0,958 | 0,973 | 0,91 | 1,128 | 0,929 | 0,998 | 0,97 | -0,03 | 6,6E-01 |
| P47809 | Dual specificity mitogen-activated protein kinase kinase 4 OS=Mus | 18 | 6 | 7 | 6 | 397 | 44,1 | 8,07 | 16,55 | 6 | 1,092 | 1,13 | 0,968 | 0,892 | 0,852 | 1,165 | 0,935 | 0,97 | 1,04 | 0,04 | 6,6E-01 |
| A2A432 | Cullin-4B OS=Mus musculus OX=10090 GN=Cul4b PE=1 SV=1 | 8 | 6 | 6 | 5 | 970 | 110,6 | 8,37 | 0 | 6 | 1,123 | 1,179 | 1,146 | 0,919 | 0,819 | 1,225 | 1,093 | 1,039 | 1,05 | 0,05 | 6,6E-01 |
| P70670 | Nascent polypeptide-associated complex subunit alpha, muscle-spe | 1 | 1 | 1 | 1 | 2187 | 220,4 | 9,35 | 3,89 | 1 | 1,29 | 1,262 | 0,964 | 1,186 | 0,912 | 1,432 | 0,775 | 1,269 | 1,07 | 0,07 | 6,6E-01 |
| Q9CPR5 | 39S ribosomal protein L15, mitochondrial OS=Mus musculus OX=10 | 16 | 4 | 5 | 4 | 295 | 33,5 | 10,07 | 9,45 | 4 | 1,182 | 0,987 | 1,099 | 1 | 1,052 | 1,287 | 0,98 | 1,097 | 0,97 | -0,03 | 6,6E-01 |

| | | | | | | | | | | | | | | | | | | | | | |
|---|---|---|---|---|---|---|---|---|---|---|---|---|---|---|---|---|---|---|---|---|---|
| Q8BFR5 | Elongation factor Tu, mitochondrial OS=Mus musculus OX=10090 G | 41 | 17 | 30 | 17 | 452 | 49,5 | 7,56 | 73,41 | 17 | 0,94 | 1,019 | 0,959 | 0,978 | 0,932 | 0,965 | 0,94 | 1,013 | 1,01 | 0,01 | 6,6E-01 |
| Q9ET80 | Junctophilin-1 OS=Mus musculus OX=10090 GN=Jph1 PE=1 SV=1 | 3 | 2 | 2 | 2 | 660 | 71,9 | 9,22 | 0 | 2 | 1,417 | 1,081 | 1,461 | 1,192 | 1,491 | 1,095 | 1,193 | 1,136 | 1,05 | 0,05 | 6,6E-01 |
| P28740-1 | Isoform 1 of Kinesin-like protein KIF2A OS=Mus musculus OX=1009 | 18 | 12 | 17 | 12 | 716 | 80,8 | 6,16 | 42,07 | 12 | 1,077 | 0,973 | 1,045 | 1,046 | 1,006 | 1,136 | 1,03 | 1,036 | 0,98 | -0,02 | 6,6E-01 |
| Q9JI75 | Ribosyldihydronicotinamide dehydrogenase [quinone] OS=Mus mu | 9 | 2 | 2 | 2 | 231 | 26,2 | 7,01 | 3,3 | 2 | 0,752 | 0,789 | 1,025 | 0,815 | 0,718 | 0,945 | 0,775 | 0,799 | 1,04 | 0,04 | 6,6E-01 |
| Q6ZPQ6 | Membrane-associated phosphatidylinositol transfer protein 2 OS=M | 1 | 1 | 1 | 1 | 1335 | 147,9 | 7,93 | 0 | 1 | 1,485 | 1,391 | 0,746 | 1,151 | 1,278 | 0,849 | 1,068 | 1,226 | 1,08 | 0,08 | 6,6E-01 |
| Q02257 | Junction plakoglobin OS=Mus musculus OX=10090 GN=Jup PE=1 SV | 10 | 7 | 8 | 5 | 745 | 81,7 | 6,14 | 18,61 | 7 | 1,183 | 1,396 | 1,001 | 0,866 | 1,111 | 1,203 | 0,855 | 1,025 | 1,06 | 0,06 | 6,6E-01 |
| P63017 | Heat shock cognate 71 kDa protein OS=Mus musculus OX=10090 G | 53 | 36 | 98 | 25 | 646 | 70,8 | 5,52 | 243,48 | 36 | 0,956 | 0,94 | 0,996 | 1,002 | 0,969 | 0,966 | 1,008 | 0,984 | 0,99 | -0,01 | 6,6E-01 |
| Q03517 | Secretogranin-2 OS=Mus musculus OX=10090 GN=Scg2 PE=1 SV=1 | 12 | 5 | 6 | 5 | 617 | 70,6 | 4,75 | 9,04 | 5 | 1,333 | 0,719 | 0,818 | 0,843 | 0,78 | 0,922 | 0,93 | 0,819 | 1,08 | 0,08 | 6,6E-01 |
| P51432 | 1-phosphatidylinositol 4,5-bisphosphate phosphodiesterase beta-3 | 1 | 1 | 1 | 1 | 1234 | 139,4 | 5,94 | 0 | 1 | 1,137 | 0,521 | 0,976 | 1,227 | 0,941 | 0,841 | 0,878 | 0,91 | 1,08 | 0,08 | 6,6E-01 |
| Q8VD33 | Small glutamine-rich tetratricopeptide repeat-containing protein be | 13 | 3 | 3 | 3 | 304 | 33,4 | 4,92 | 5,59 | 3 | 1,052 | 0,919 | 0,893 | 0,919 | 0,998 | 0,961 | 0,882 | 0,851 | 1,02 | 0,02 | 6,6E-01 |
| Q3V3R1 | Monofunctional C1-tetrahydrofolate synthase, mitochondrial OS=M | 14 | 11 | 13 | 10 | 977 | 105,7 | 7,02 | 26,6 | 11 | 1,013 | 1,012 | 0,941 | 1,052 | 0,953 | 1,022 | 0,999 | 0,994 | 1,01 | 0,01 | 6,6E-01 |
| P40630 | Transcription factor A, mitochondrial OS=Mus musculus OX=10090 | 28 | 6 | 6 | 6 | 243 | 28 | 9,69 | 16,56 | 6 | 0,91 | 0,907 | 0,989 | 0,993 | 1,013 | 0,875 | 0,972 | 1,013 | 0,98 | -0,02 | 6,6E-01 |
| P61082 | NEDD8-conjugating enzyme Ubc12 OS=Mus musculus OX=10090 G | 22 | 4 | 4 | 4 | 183 | 20,9 | 7,69 | 4,45 | 4 | 1,223 | 1,404 | 1,026 | 0,926 | 1,013 | 1,153 | 1,109 | 1,102 | 1,05 | 0,05 | 6,6E-01 |
| O89051 | Integral membrane protein 2B OS=Mus musculus OX=10090 GN=Itr | 14 | 3 | 3 | 3 | 266 | 30,2 | 5,3 | 4,99 | 3 | 1,126 | 1,279 | 0,962 | 1,21 | 1,172 | 1,242 | 0,849 | 1,113 | 1,05 | 0,05 | 6,6E-01 |
| O35927 | Catenin delta-2 OS=Mus musculus OX=10090 GN=Ctnnd2 PE=1 SV= | 17 | 16 | 20 | 13 | 1247 | 134,9 | 7,65 | 30,67 | 16 | 1,216 | 1,178 | 0,848 | 1,055 | 0,916 | 1,015 | 1,095 | 1,1 | 1,04 | 0,04 | 6,6E-01 |
| Q99KK9 | Probable histidine--tRNA ligase, mitochondrial OS=Mus musculus C | 10 | 5 | 5 | 3 | 505 | 56,9 | 8,32 | 11,85 | 5 | 0,948 | 0,927 | 1,078 | 0,989 | 1,094 | 0,834 | 0,89 | 1,002 | 1,03 | 0,03 | 6,6E-01 |
| Q8BXC6 | COMM domain-containing protein 2 OS=Mus musculus OX=10090 ( | 5 | 1 | 1 | 1 | 199 | 22,8 | 6,62 | 0 | 1 | 0,835 | 0,635 | 0,721 | 0,793 | 0,906 | 0,862 | 0,709 | 0,645 | 0,96 | -0,04 | 6,6E-01 |
| Q8K1L5 | E3 ubiquitin-protein ligase PPP1R11 OS=Mus musculus OX=10090 ( | 11 | 1 | 1 | 1 | 131 | 14,5 | 6,09 | 3,91 | 1 | 1,102 | 1,514 | 1,722 | 1,11 | 1,234 | 1,882 | 1,301 | 1,417 | 0,93 | -0,07 | 6,7E-01 |
| Q08331 | Calretinin OS=Mus musculus OX=10090 GN=Calb2 PE=1 SV=3 | 25 | 6 | 8 | 6 | 271 | 31,4 | 5,02 | 14,42 | 6 | 0,819 | 0,831 | 1,104 | 0,915 | 0,845 | 0,972 | 1,126 | 0,89 | 0,96 | -0,04 | 6,7E-01 |
| Q8C015 | Serine/threonine-protein kinase PAK 5 OS=Mus musculus OX=1009 | 3 | 2 | 3 | 2 | 719 | 80,9 | 8,21 | 7,91 | 2 | 1,222 | 1,216 | 1,059 | 1,055 | 1,065 | 1,151 | 1,018 | 1,204 | 1,03 | 0,03 | 6,7E-01 |
| Q9D113 | DNL-type zinc finger protein OS=Mus musculus OX=10090 GN=Dnlz | 6 | 1 | 1 | 1 | 177 | 19,4 | 9,48 | 2,96 | 1 | 1,302 | 1,314 | 0,819 | 0,517 | 0,777 | 1,016 | 0,875 | 0,919 | 1,10 | 0,10 | 6,7E-01 |
| Q8CGA0 | Protein phosphatase 1F OS=Mus musculus OX=10090 GN=Ppm1f P | 8 | 2 | 2 | 2 | 452 | 49,6 | 5,3 | 4,61 | 2 | 1,031 | 0,961 | 0,715 | 1,152 | 0,875 | 0,894 | 0,901 | 1,013 | 1,05 | 0,05 | 6,7E-01 |
| Q8BMS1 | Trifunctional enzyme subunit alpha, mitochondrial OS=Mus muscul | 19 | 15 | 22 | 15 | 763 | 82,6 | 9,14 | 47,25 | 15 | 1,03 | 1,043 | 1,004 | 0,937 | 0,978 | 1,011 | 0,968 | 1,01 | 1,01 | 0,01 | 6,7E-01 |
| O35099 | Mitogen-activated protein kinase kinase kinase 5 OS=Mus musculu | 3 | 4 | 4 | 4 | 1380 | 154,4 | 5,78 | 9,94 | 4 | 0,999 | 1,141 | 1,093 | 0,989 | 0,932 | 1,166 | 1,239 | 1,026 | 0,97 | -0,03 | 6,7E-01 |
| Q9JKW0 | ADP-ribosylation factor-like protein 6-interacting protein 1 OS=Mus | 5 | 1 | 1 | 1 | 203 | 23,4 | 9,32 | 4,15 | 1 | 0,867 | 0,899 | 0,91 | 1,229 | 1,061 | 1,048 | 1,075 | 0,892 | 0,96 | -0,04 | 6,7E-01 |
| Q8BZA9 | Fructose-2,6-bisphosphatase TIGAR OS=Mus musculus OX=10090 G | 10 | 2 | 3 | 2 | 269 | 29,2 | 8,18 | 6,27 | 2 | 2,215 | 1,264 | 1,351 | 1,15 | 0,88 | 1,798 | 1,277 | 1,467 | 1,10 | 0,10 | 6,7E-01 |
| Q8CEC5 | NF-kappa-B inhibitor-interacting Ras-like protein 1 OS=Mus muscul | 9 | 2 | 2 | 2 | 192 | 21,6 | 7,33 | 1,75 | 2 | 1,081 | 0,782 | 0,886 | 1,03 | 1,128 | 0,923 | 1,01 | 0,876 | 0,96 | -0,04 | 6,7E-01 |
| Q6PAV2 | Probable E3 ubiquitin-protein ligase HERC4 OS=Mus musculus OX=1 | 2 | 2 | 2 | 2 | 1057 | 118,3 | 6,29 | 4,16 | 2 | 1,108 | 1,034 | 1,305 | 1,159 | 1,143 | 1,257 | 1,072 | 0,989 | 1,03 | 0,03 | 6,7E-01 |
| Q8BXR1 | Probable cationic amino acid transporter OS=Mus musculus OX=10 | 4 | 3 | 6 | 3 | 771 | 83,9 | 5,35 | 15,47 | 3 | 1,054 | 1,046 | 1,001 | 1,015 | 0,909 | 0,827 | 1,133 | 1,11 | 1,03 | 0,03 | 6,7E-01 |
| Q8C5T8 | Coiled-coil domain-containing protein 113 OS=Mus musculus OX=1 | 2 | 1 | 1 | 1 | 377 | 44,2 | 8,46 | 1,98 | 1 | 1,021 | 0,985 | 1,038 | 1,145 | 1,025 | 1,13 | 1,047 | 1,061 | 0,98 | -0,02 | 6,7E-01 |
| Q61205 | Platelet-activating factor acetylhydrolase IB subunit gamma OS=Mu | 14 | 3 | 3 | 3 | 232 | 25,8 | 6,93 | 7,96 | 3 | 1,155 | 1,109 | 0,975 | 1,058 | 0,941 | 1,083 | 0,946 | 1,197 | 1,03 | 0,03 | 6,7E-01 |
| Q91YJ2 | Sorting nexin-4 OS=Mus musculus OX=10090 GN=Snx4 PE=1 SV=1 | 14 | 6 | 6 | 6 | 450 | 51,7 | 5,8 | 11,01 | 6 | 1,393 | 1,208 | 1,129 | 1,133 | 1,163 | 1,17 | 1,244 | 1,17 | 1,02 | 0,02 | 6,7E-01 |
| Q91W43 | Glycine dehydrogenase (decarboxylating), mitochondrial OS=Mus r | 6 | 6 | 8 | 6 | 1025 | 113,2 | 7,56 | 19,42 | 6 | 0,998 | 1,116 | 1,19 | 1,022 | 1,05 | 1,07 | 1,042 | 1,083 | 1,02 | 0,02 | 6,7E-01 |
| Q3V1L4 | Cytosolic purine 5'-nucleotidase OS=Mus musculus OX=10090 GN=I | 5 | 2 | 2 | 2 | 560 | 64,8 | 6,21 | 3,23 | 2 | 1,11 | 1,01 | 0,776 | 1,044 | 0,822 | 0,986 | 0,968 | 1,013 | 1,04 | 0,04 | 6,7E-01 |
| Q9JJZ2 | Tubulin alpha-8 chain OS=Mus musculus OX=10090 GN=Tuba8 PE=1 | 41 | 18 | 70 | 4 | 449 | 50 | 5,1 | 177,95 | 18 | 0,661 | 0,831 | 0,742 | 0,727 | 0,713 | 0,619 | 0,961 | 0,813 | 0,95 | -0,05 | 6,7E-01 |
| Q64314 | Hematopoietic progenitor cell antigen CD34 OS=Mus musculus OX= | 10 | 4 | 4 | 4 | 382 | 41 | 5,3 | 7,23 | 4 | 0,898 | 1,214 | 0,944 | 1,006 | 0,973 | 1,085 | 0,867 | 0,989 | 1,04 | 0,04 | 6,7E-01 |
| P27546-4 | Isoform 4 of Microtubule-associated protein 4 OS=Mus musculus O | 8 | 5 | 8 | 2 | 902 | 94,5 | 9,06 | 20,7 | 5 | 0,927 | 0,85 | 1,141 | 0,982 | 0,889 | 0,76 | 1,085 | 0,999 | 1,04 | 0,04 | 6,7E-01 |
| P59644 | Phosphatidylinositol 4,5-bisphosphate 5-phosphatase A OS=Mus m | 3 | 3 | 3 | 3 | 1003 | 107,5 | 9,36 | 6,23 | 3 | 0,752 | 0,806 | 1,045 | 0,983 | 0,848 | 0,843 | 0,997 | 1,055 | 0,96 | -0,04 | 6,7E-01 |
| Q9ERK4 | Exportin-2 OS=Mus musculus OX=10090 GN=Cse1l PE=1 SV=1 | 8 | 8 | 8 | 8 | 971 | 110,4 | 5,77 | 15,54 | 8 | 1,085 | 0,935 | 1,037 | 0,889 | 0,937 | 0,965 | 0,896 | 1,049 | 1,03 | 0,03 | 6,7E-01 |
| Q812E0 | Cytoplasmic polyadenylation element-binding protein 2 OS=Mus m | 2 | 1 | 1 | 1 | 521 | 58,4 | 7,5 | 2,42 | 1 | 1,07 | 1,315 | 1,125 | 1,057 | 1,161 | 1,357 | 0,943 | 1,302 | 0,96 | -0,04 | 6,7E-01 |
| Q8R146 | Acylamino-acid-releasing enzyme OS=Mus musculus OX=10090 GN | 2 | 2 | 2 | 2 | 732 | 81,5 | 5,59 | 4,3 | 2 | 0,715 | 1,016 | 1,114 | 1,149 | 0,993 | 0,914 | 1,05 | 0,844 | 1,05 | 0,05 | 6,7E-01 |
| P12023 | Amyloid-beta A4 protein OS=Mus musculus OX=10090 GN=App PE= | 9 | 7 | 7 | 6 | 770 | 86,7 | 4,79 | 17,08 | 7 | 0,993 | 0,924 | 0,941 | 0,998 | 0,963 | 0,978 | 0,932 | 1,033 | 0,99 | -0,01 | 6,7E-01 |
| Q9JM13 | Rab5 GDP/GTP exchange factor OS=Mus musculus OX=10090 GN=F | 4 | 2 | 2 | 2 | 491 | 56,8 | 6,79 | 6,26 | 2 | 0,978 | 0,992 | 0,996 | 0,993 | 0,866 | 0,994 | 1,079 | 1,12 | 0,98 | -0,02 | 6,7E-01 |
| Q920M5 | Coronin-6 OS=Mus musculus OX=10090 GN=Coro6 PE=1 SV=1 | 7 | 3 | 3 | 2 | 471 | 52,6 | 5,96 | 0 | 3 | 1,3 | 1,72 | 1,444 | 1,607 | 1,349 | 1,373 | 1,989 | 0,943 | 1,07 | 0,07 | 6,7E-01 |
| P46660 | Alpha-internexin OS=Mus musculus OX=10090 GN=Ina PE=1 SV=3 | 34 | 18 | 25 | 14 | 501 | 55,3 | 5,4 | 58,08 | 18 | 0,99 | 1,036 | 0,897 | 0,972 | 1,133 | 0,924 | 0,976 | 0,959 | 0,98 | -0,02 | 6,7E-01 |
| Q8BLE7 | Vesicular glutamate transporter 2 OS=Mus musculus OX=10090 GN | 8 | 4 | 4 | 4 | 582 | 64,5 | 6,68 | 11,55 | 4 | 0,795 | 1,202 | 0,968 | 1,014 | 0,993 | 1,023 | 1,034 | 1,081 | 0,96 | -0,04 | 6,7E-01 |
| Q8C522 | Endonuclease domain-containing 1 protein OS=Mus musculus OX=1 | 10 | 5 | 7 | 5 | 501 | 55,2 | 6,16 | 13,98 | 5 | 1,241 | 1,319 | 1,169 | 1,092 | 1,196 | 1,191 | 1,199 | 1,146 | 1,02 | 0,02 | 6,7E-01 |
| Q8BUV3 | Gephyrin OS=Mus musculus OX=10090 GN=Gphn PE=1 SV=2 | 23 | 15 | 18 | 15 | 769 | 83,2 | 5,6 | 36,19 | 15 | 1,026 | 1,065 | 1,025 | 0,888 | 0,911 | 1,06 | 1,001 | 0,942 | 1,02 | 0,02 | 6,7E-01 |
| Q811P8 | Rho GTPase-activating protein 32 OS=Mus musculus OX=10090 GN | 4 | 7 | 7 | 7 | 2089 | 229,6 | 6,86 | 16,9 | 7 | 1,178 | 1,151 | 1,212 | 0,917 | 0,939 | 1,289 | 1,012 | 1,039 | 1,04 | 0,04 | 6,7E-01 |
| Q03141 | MAP/microtubule affinity-regulating kinase 3 OS=Mus musculus OX | 9 | 5 | 7 | 2 | 753 | 84,3 | 9,51 | 6,07 | 5 | 1,023 | 1,01 | 0,8 | 0,908 | 1,048 | 0,693 | 1,057 | 0,75 | 1,05 | 0,05 | 6,7E-01 |
| Q91YJ5 | Translation initiation factor IF-2, mitochondrial OS=Mus musculus ( | 1 | 1 | 1 | 1 | 727 | 81,2 | 7,24 | 0 | 1 | 1,2 | 0,997 | 0,696 | 1,068 | 0,835 | 1,439 | 0,766 | 1,266 | 0,92 | -0,08 | 6,7E-01 |
| Q80TM6 | R3H domain-containing protein 2 OS=Mus musculus OX=10090 GN= | 4 | 3 | 3 | 3 | 1044 | 114,5 | 8,91 | 1,69 | 3 | 1,432 | 2,198 | 1,124 | 1,143 | 0,97 | 1,858 | 0,987 | 1,499 | 1,11 | 0,11 | 6,7E-01 |
| Q8BY89 | Choline transporter-like protein 2 OS=Mus musculus OX=10090 GN | 1 | 1 | 1 | 1 | 706 | 80,1 | 8,79 | 2,87 | 1 | 0,814 | 1,094 | 1,173 | 1,362 | 1,094 | 1,159 | 1,297 | 0,537 | 1,09 | 0,09 | 6,8E-01 |
| Q3UTH8 | Rho guanine nucleotide exchange factor 9 OS=Mus musculus OX=1 | 7 | 4 | 6 | 4 | 516 | 60,9 | 5,64 | 12,87 | 4 | 1,102 | 1,013 | 0,994 | 0,902 | 1,091 | 0,96 | 0,921 | 0,94 | 1,03 | 0,03 | 6,8E-01 |
| Q8R0H9 | ADP-ribosylation factor-binding protein GGA1 OS=Mus musculus O | 2 | 1 | 1 | 1 | 635 | 69,9 | 5,27 | 2,9 | 1 | 1,562 | 1,18 | 1,136 | 0,942 | 0,923 | 1,389 | 1,116 | 1,108 | 1,06 | 0,06 | 6,8E-01 |
| Q62178 | Semaphorin-4A OS=Mus musculus OX=10090 GN=Sema4a PE=1 SV | 3 | 2 | 2 | 2 | 760 | 83,4 | 7,72 | 1,66 | 2 | 1,311 | 0,569 | 0,785 | 0,684 | 1,19 | 0,747 | 0,832 | 0,914 | 0,91 | -0,09 | 6,8E-01 |
| Q6PHZ8 | Kv channel-interacting protein 4 OS=Mus musculus OX=10090 GN= | 15 | 3 | 3 | 3 | 250 | 28,7 | 5,21 | 9,79 | 3 | 0,85 | 1,151 | 1,26 | 1,127 | 0,989 | 1,137 | 1,291 | 1,159 | 0,96 | -0,04 | 6,8E-01 |
| Q8CIN4 | Serine/threonine-protein kinase PAK 2 OS=Mus musculus OX=1009 | 11 | 4 | 4 | 2 | 524 | 57,9 | 5,77 | 13,43 | 4 | 0,651 | 0,895 | 0,978 | 0,82 | 0,894 | 0,937 | 0,832 | 0,813 | 0,96 | -0,04 | 6,8E-01 |
| Q9QYJ3 | DnaJ homolog subfamily B member 1 OS=Mus musculus OX=10090 | 11 | 4 | 6 | 3 | 340 | 38,1 | 8,63 | 9,56 | 4 | 1,014 | 1,122 | 1,091 | 0,997 | 1,05 | 1,018 | 0,981 | 1,105 | 1,02 | 0,02 | 6,8E-01 |
| Q8VE33 | Ganglioside-induced differentiation-associated protein 1-like 1 OS= | 24 | 7 | 8 | 6 | 370 | 42,3 | 6,6 | 19,07 | 7 | 1 | 1,117 | 0,86 | 0,935 | 1,01 | 0,98 | 0,95 | 1,079 | 0,97 | -0,03 | 6,8E-01 |
| P58802 | TBC1 domain family member 10A OS=Mus musculus OX=10090 GN | 6 | 2 | 3 | 2 | 500 | 56,2 | 7,85 | 2,94 | 2 | 1,302 | 1,423 | 1,453 | 1,428 | 1,085 | 1,789 | 1,423 | 1,575 | 0,95 | -0,05 | 6,8E-01 |
| Q8C7R4 | Ubiquitin-like modifier-activating enzyme 6 OS=Mus musculus OX= | 6 | 5 | 8 | 5 | 1053 | 117,9 | 6,11 | 5,07 | 5 | 1,375 | 1,276 | 0,907 | 1,093 | 0,945 | 1,245 | 1,2 | 1,043 | 1,05 | 0,05 | 6,8E-01 |

| | | | | | | | | | | | | | | | | | | | | | |
|---|---|---|---|---|---|---|---|---|---|---|---|---|---|---|---|---|---|---|---|---|---|
| Q61595 | Kinectin OS=Mus musculus OX=10090 GN=Ktn1 PE=1 SV=1 | 4 | 5 | 5 | 5 | 1327 | 152,5 | 5,86 | 7,05 | 5 | 1,292 | 1,031 | 1,233 | 1,332 | 0,996 | 1,088 | 1,102 | 1,481 | 1,05 | 0,05 | 6,8E-01 |
| Q8CE22 | Cyclin-D-binding Myb-like transcription factor 1 OS=Mus musculus | 1 | 1 | 1 | 1 | 761 | 84,6 | 4,68 | 0 | 1 | 1,66 | 1,787 | 0,646 | 0,937 | 0,999 | 1,603 | 0,943 | 0,924 | 1,13 | 0,13 | 6,8E-01 |
| Q8CIG8 | Protein arginine N-methyltransferase 5 OS=Mus musculus OX=1009 | 1 | 1 | 1 | 1 | 637 | 72,6 | 6,42 | 0 | 1 | 1,671 | 1,156 | 1,488 | 1,636 | 1,626 | 1,918 | 1,387 | 1,331 | 0,95 | -0,05 | 6,8E-01 |
| Q8K354 | Carbonyl reductase [NADPH] 3 OS=Mus musculus OX=10090 GN=Cl | 20 | 5 | 6 | 4 | 277 | 30,9 | 6,57 | 6,18 | 5 | 0,944 | 0,939 | 0,883 | 0,871 | 0,813 | 0,81 | 0,985 | 0,944 | 1,02 | 0,02 | 6,8E-01 |
| Q61330 | Contactin-2 OS=Mus musculus OX=10090 GN=Cntn2 PE=1 SV=2 | 9 | 7 | 7 | 7 | 1040 | 113,1 | 7,62 | 17,69 | 7 | 1,149 | 0,805 | 1,044 | 0,82 | 0,852 | 1,069 | 0,837 | 0,885 | 1,05 | 0,05 | 6,8E-01 |
| Q8BXA5 | Cleft lip and palate transmembrane protein 1-like protein OS=Mus | 2 | 1 | 1 | 1 | 539 | 62,1 | 8,84 | 2,34 | 1 | 0,859 | 1,119 | 0,805 | 1,109 | 1,164 | 0,839 | 0,997 | 1,079 | 0,95 | -0,05 | 6,8E-01 |
| Q9CPW0 | Contactin-associated protein-like 2 OS=Mus musculus OX=10090 G | 9 | 11 | 13 | 11 | 1332 | 148,1 | 6,77 | 31,64 | 11 | 1,228 | 0,864 | 1,073 | 1,02 | 1,292 | 0,98 | 1,108 | 0,987 | 0,96 | -0,04 | 6,8E-01 |
| Q9DAM5 | Mitochondrial thiamine pyrophosphate carrier OS=Mus musculus C | 3 | 1 | 1 | 1 | 318 | 35,6 | 9,2 | 2,51 | 1 | 1,014 | 1,051 | 0,532 | 0,903 | 0,833 | 1,041 | 0,741 | 0,629 | 1,08 | 0,08 | 6,8E-01 |
| P97929 | Breast cancer type 2 susceptibility protein homolog OS=Mus muscu | 0 | 1 | 1 | 1 | 3329 | 370,4 | 6,67 | 0 | 1 | 1,689 | 0,911 | 0,797 | 1,157 | 1,058 | 0,911 | 1,043 | 1,184 | 1,09 | 0,09 | 6,8E-01 |
| Q9DCM0 | Persulfide dioxygenase ETHE1, mitochondrial OS=Mus musculus OX | 17 | 3 | 5 | 3 | 254 | 27,7 | 7,23 | 13,83 | 3 | 1,288 | 1,199 | 0,893 | 1,025 | 1,013 | 1,31 | 0,994 | 0,864 | 1,05 | 0,05 | 6,8E-01 |
| Q9QXB9 | Developmentally-regulated GTP-binding protein 2 OS=Mus musculu | 7 | 2 | 2 | 2 | 364 | 40,7 | 8,88 | 5,62 | 2 | 1,018 | 0,993 | 1,04 | 0,985 | 1,071 | 1,011 | 0,945 | 0,954 | 1,01 | 0,01 | 6,8E-01 |
| Q9R0M6 | Ras-related protein Rab-9A OS=Mus musculus OX=10090 GN=Rab9 | 7 | 1 | 1 | 1 | 201 | 22,9 | 5,66 | 2,33 | 1 | 0,889 | 1,116 | 0,749 | 0,814 | 0,818 | 0,935 | 0,666 | 0,967 | 1,05 | 0,05 | 6,8E-01 |
| Q9EPV8 | Ubiquitin-like protein 5 OS=Mus musculus OX=10090 GN=Ubl5 PE= | 12 | 1 | 1 | 1 | 73 | 8,5 | 8,44 | 1,65 | 1 | 0,708 | 1,067 | 0,931 | 1,208 | 1,316 | 0,925 | 0,98 | 0,937 | 0,94 | -0,06 | 6,8E-01 |
| P17751 | Triosephosphate isomerase OS=Mus musculus OX=10090 GN=Tpi1 | 41 | 12 | 27 | 12 | 299 | 32,2 | 5,74 | 56,63 | 12 | 0,923 | 0,873 | 0,993 | 1,024 | 0,978 | 0,939 | 1,011 | 0,95 | 0,98 | -0,02 | 6,8E-01 |
| Q8R5C5 | Beta-centractin OS=Mus musculus OX=10090 GN=Actr1b PE=1 SV= | 26 | 9 | 12 | 4 | 376 | 42,3 | 6,4 | 24,33 | 9 | 0,918 | 0,853 | 0,822 | 0,995 | 1,007 | 0,85 | 0,922 | 0,896 | 0,98 | -0,02 | 6,8E-01 |
| P63085 | Mitogen-activated protein kinase 1 OS=Mus musculus OX=10090 G | 30 | 12 | 21 | 8 | 358 | 41,2 | 6,98 | 40,77 | 12 | 0,941 | 0,945 | 1,067 | 1,065 | 0,961 | 0,985 | 1,12 | 1,038 | 0,98 | -0,02 | 6,8E-01 |
| Q8BSK8 | Ribosomal protein S6 kinase beta-1 OS=Mus musculus OX=10090 G | 5 | 2 | 2 | 2 | 525 | 59,1 | 6,81 | 8,07 | 2 | 1,267 | 1,16 | 1,224 | 0,988 | 0,98 | 1,404 | 1,045 | 1,01 | 1,05 | 0,05 | 6,8E-01 |
| Q6IFZ6 | Keratin, type II cytoskeletal 1b OS=Mus musculus OX=10090 GN=Kr | 7 | 5 | 9 | 1 | 572 | 61,3 | 8,02 | 25,5 | 5 | 1,834 | 1,049 | 0,627 | 1,142 | 0,623 | 1,239 | 1,03 | 1,261 | 1,12 | 0,12 | 6,8E-01 |
| P62774 | Myotrophin OS=Mus musculus OX=10090 GN=Mtpn PE=1 SV=2 | 38 | 4 | 6 | 4 | 118 | 12,9 | 5,52 | 17,43 | 4 | 0,937 | 0,921 | 0,982 | 0,999 | 0,938 | 0,88 | 1,063 | 1,038 | 0,98 | -0,02 | 6,8E-01 |
| Q3UR70 | Transforming growth factor-beta receptor-associated protein 1 OS= | 2 | 1 | 2 | 1 | 860 | 97,2 | 6,7 | 4,52 | 1 | 1,161 | 0,732 | 1,162 | 0,962 | 1,121 | 1,16 | 0,935 | 0,998 | 0,95 | -0,05 | 6,8E-01 |
| Q9WUQ2 | Prolactin regulatory element-binding protein OS=Mus musculus OX | 11 | 4 | 4 | 4 | 417 | 45,4 | 8,76 | 7,65 | 4 | 1,417 | 0,945 | 1,12 | 1,068 | 1,073 | 1,162 | 1,001 | 1,132 | 1,04 | 0,04 | 6,8E-01 |
| P49443 | Protein phosphatase 1A OS=Mus musculus OX=10090 GN=Ppm1a F | 33 | 11 | 14 | 10 | 382 | 42,4 | 5,36 | 28,19 | 11 | 1,019 | 0,892 | 1,049 | 1,064 | 0,968 | 1,104 | 1,041 | 0,995 | 0,98 | -0,02 | 6,8E-01 |
| O35409 | Glutamate carboxypeptidase 2 OS=Mus musculus OX=10090 GN=Fo | 6 | 3 | 4 | 3 | 752 | 84,5 | 8,1 | 10,7 | 3 | 1,059 | 1,236 | 1,006 | 0,971 | 0,935 | 1,23 | 0,95 | 1,003 | 1,04 | 0,04 | 6,8E-01 |
| Q8C6G8 | WD repeat-containing protein 26 OS=Mus musculus OX=10090 GN= | 6 | 5 | 5 | 5 | 641 | 70,5 | 6,16 | 9,18 | 5 | 0,976 | 1,082 | 1,049 | 0,867 | 0,915 | 1,025 | 0,98 | 0,964 | 1,02 | 0,02 | 6,8E-01 |
| Q60610 | T-lymphoma invasion and metastasis-inducing protein 1 OS=Mus m | 2 | 3 | 3 | 2 | 1591 | 177,4 | 6,67 | 2,6 | 3 | 0,929 | 1,2 | 0,749 | 0,884 | 0,881 | 1,078 | 1,015 | 0,964 | 0,96 | -0,04 | 6,9E-01 |
| O09061 | Proteasome subunit beta type-1 OS=Mus musculus OX=10090 GN= | 30 | 6 | 12 | 6 | 240 | 26,4 | 7,81 | 36,86 | 6 | 1,295 | 1,26 | 1,164 | 1,068 | 0,986 | 1,313 | 1,155 | 1,189 | 1,03 | 0,03 | 6,9E-01 |
| Q8K382 | DENN domain-containing protein 1A OS=Mus musculus OX=10090 | 2 | 2 | 2 | 2 | 1016 | 111,5 | 6,79 | 0 | 2 | 0,99 | 0,455 | 0,641 | 0,761 | 0,784 | 0,572 | 0,756 | 0,971 | 0,92 | -0,08 | 6,9E-01 |
| P97384 | Annexin A11 OS=Mus musculus OX=10090 GN=Anxa11 PE=1 SV=2 | 15 | 6 | 6 | 6 | 503 | 54 | 7,66 | 16,03 | 6 | 1,035 | 1,353 | 0,928 | 1,068 | 0,959 | 1,263 | 1,121 | 1,235 | 0,96 | -0,04 | 6,9E-01 |
| O08756 | 3-hydroxyacyl-CoA dehydrogenase type-2 OS=Mus musculus OX=1( | 20 | 4 | 5 | 4 | 261 | 27,4 | 8,41 | 16,37 | 4 | 1,053 | 0,98 | 1,01 | 0,953 | 0,982 | 0,966 | 1,041 | 1,06 | 0,99 | -0,01 | 6,9E-01 |
| Q8BTG7 | Protein NDRG4 OS=Mus musculus OX=10090 GN=Ndrg4 PE=1 SV=1 | 10 | 3 | 3 | 2 | 352 | 38,5 | 6,32 | 5,25 | 3 | 1,15 | 0,993 | 1,103 | 1,146 | 0,942 | 1,219 | 1,113 | 0,996 | 1,03 | 0,03 | 6,9E-01 |
| Q9JJY3 | Sphingomyelin phosphodiesterase 3 OS=Mus musculus OX=10090 ( | 11 | 5 | 6 | 5 | 655 | 71,2 | 5,88 | 12,45 | 5 | 1,061 | 1,214 | 0,922 | 0,971 | 1,013 | 0,948 | 1,156 | 1,199 | 0,97 | -0,03 | 6,9E-01 |
| Q8C080 | Sorting nexin-16 OS=Mus musculus OX=10090 GN=Snx16 PE=1 SV= | 6 | 2 | 2 | 2 | 344 | 38,8 | 5,21 | 4,58 | 2 | 0,932 | 0,596 | 0,901 | 1,184 | 0,755 | 0,768 | 0,879 | 0,987 | 1,07 | 0,07 | 6,9E-01 |
| Q9QZB9 | Dynactin subunit 5 OS=Mus musculus OX=10090 GN=Dctn5 PE=1 S\ | 15 | 3 | 3 | 3 | 182 | 20,1 | 8,02 | 4,92 | 3 | 1,167 | 1,006 | 1,111 | 0,981 | 0,957 | 1,055 | 0,949 | 1,185 | 1,03 | 0,03 | 6,9E-01 |
| Q9JI78 | Peptide-N(4)-(N-acetyl-beta-glucosaminyl)asparagine amidase OS= | 2 | 1 | 1 | 1 | 651 | 74,2 | 6,89 | 0 | 1 | 0,721 | 1,282 | 0,928 | 0,836 | 0,881 | 1,026 | 1,094 | 0,984 | 0,95 | -0,05 | 6,9E-01 |
| Q9QYI3 | DnaJ homolog subfamily C member 7 OS=Mus musculus OX=10090 | 7 | 3 | 4 | 3 | 494 | 56,4 | 6,49 | 5,91 | 3 | 1,33 | 0,909 | 1,107 | 1,079 | 0,976 | 0,997 | 1,092 | 1,192 | 1,04 | 0,04 | 6,9E-01 |
| O35127 | Protein C10 OS=Mus musculus OX=10090 GN=Grcc10 PE=1 SV=1 | 12 | 1 | 1 | 1 | 126 | 13,2 | 5,14 | 3,3 | 1 | 0,794 | 1,138 | 1,112 | 0,876 | 0,894 | 0,995 | 0,999 | 0,878 | 1,04 | 0,04 | 6,9E-01 |
| Q8R502 | Volume-regulated anion channel subunit LRRC8C OS=Mus musculu | 5 | 3 | 3 | 3 | 803 | 92,3 | 7,81 | 0 | 3 | 1,931 | 1,923 | 1,334 | 1,188 | 1,869 | 1,591 | 1,516 | 1,754 | 0,95 | -0,05 | 6,9E-01 |
| P19258 | Protein Mpv17 OS=Mus musculus OX=10090 GN=Mpv17 PE=1 SV=1 | 6 | 1 | 1 | 1 | 176 | 19,7 | 9,44 | 2,29 | 1 | 0,823 | 0,99 | 0,622 | 0,662 | 0,773 | 0,932 | 0,827 | 0,725 | 0,95 | -0,05 | 6,9E-01 |
| Q9Z2C4 | Myotubularin-related protein 1 OS=Mus musculus OX=10090 GN=N | 8 | 4 | 4 | 4 | 669 | 75,3 | 6,8 | 9,11 | 4 | 1,178 | 1,125 | 0,918 | 1,008 | 0,975 | 0,947 | 1,013 | 1,166 | 1,03 | 0,03 | 6,9E-01 |
| P70699 | Lysosomal alpha-glucosidase OS=Mus musculus OX=10090 GN=Gaa | 1 | 1 | 1 | 1 | 953 | 106,2 | 5,83 | 0 | 1 | 1,201 | 0,992 | 1,031 | 1,17 | 0,942 | 0,559 | 1,055 | 1,502 | 1,08 | 0,08 | 6,9E-01 |
| Q8BSS9 | Liprin-alpha-2 OS=Mus musculus OX=10090 GN=Ppfia2 PE=1 SV=2 | 16 | 19 | 20 | 17 | 1257 | 143,1 | 6,11 | 42,4 | 19 | 1,088 | 1,167 | 1,047 | 1,017 | 0,972 | 1,15 | 1,021 | 1,091 | 1,02 | 0,02 | 6,9E-01 |
| Q8BFU2 | Histone H2A type 3 OS=Mus musculus OX=10090 GN=Hist3h2a PE= | 15 | 3 | 9 | 1 | 130 | 14,1 | 11,05 | 20,87 | 3 | 1,299 | 1,976 | 0,898 | 0,644 | 0,969 | 1,764 | 0,436 | 0,981 | 1,16 | 0,16 | 6,9E-01 |
| Q8VBZ3 | Cleft lip and palate transmembrane protein 1 homolog OS=Mus mu | 7 | 3 | 4 | 3 | 664 | 75,2 | 6,3 | 7,96 | 3 | 1,079 | 1,241 | 1,098 | 0,984 | 1,017 | 1,112 | 0,974 | 1,181 | 1,03 | 0,03 | 6,9E-01 |
| P45376 | Aldo-keto reductase family 1 member B1 OS=Mus musculus OX=10 | 36 | 11 | 13 | 11 | 316 | 35,7 | 7,18 | 29,87 | 11 | 1,016 | 1,043 | 1,084 | 1,044 | 0,972 | 1,108 | 0,988 | 1,061 | 1,01 | 0,01 | 6,9E-01 |
| Q9QZS3 | Protein numb homolog OS=Mus musculus OX=10090 GN=Numb PE | 8 | 5 | 7 | 2 | 653 | 70,8 | 8,65 | 16,69 | 5 | 1,117 | 0,73 | 1,145 | 1,376 | 1,23 | 0,728 | 1,146 | 0,975 | 1,07 | 0,07 | 6,9E-01 |
| Q9R0M0 | Cadherin EGF LAG seven-pass G-type receptor 2 OS=Mus musculus | 1 | 2 | 2 | 2 | 2919 | 316,8 | 5,47 | 1,79 | 2 | 1,186 | 1,067 | 1,003 | 1,086 | 1,02 | 0,878 | 1,309 | 0,968 | 1,04 | 0,04 | 6,9E-01 |
| Q99JG2 | G-protein coupled receptor 37-like 1 OS=Mus musculus OX=10090 | 2 | 1 | 3 | 1 | 481 | 52,7 | 7,36 | 7,26 | 1 | 0,986 | 1,146 | 1,048 | 0,941 | 0,979 | 1,148 | 0,952 | 0,932 | 1,03 | 0,03 | 6,9E-01 |
| O70591 | Prefoldin subunit 2 OS=Mus musculus OX=10090 GN=Pfdn2 PE=1 S | 34 | 5 | 6 | 5 | 154 | 16,5 | 6,58 | 14,21 | 5 | 1,384 | 1,167 | 1,209 | 1,188 | 1,035 | 1,353 | 1,261 | 1,159 | 1,03 | 0,03 | 6,9E-01 |
| Q9JIF0 | Protein arginine N-methyltransferase 1 OS=Mus musculus OX=1009 | 17 | 6 | 7 | 4 | 371 | 42,4 | 5,43 | 19,9 | 6 | 1,259 | 1,212 | 1,186 | 1,183 | 1,026 | 1,346 | 1,115 | 1,234 | 1,03 | 0,03 | 6,9E-01 |
| Q9D8W5 | 26S proteasome non-ATPase regulatory subunit 12 OS=Mus muscu | 20 | 8 | 9 | 8 | 456 | 52,9 | 7,06 | 11,92 | 8 | 0,921 | 0,977 | 1,118 | 1,014 | 1,033 | 1,096 | 0,99 | 0,991 | 0,98 | -0,02 | 6,9E-01 |
| Q8R3F5 | Malonyl-CoA-acyl carrier protein transacylase, mitochondrial OS=M | 8 | 2 | 2 | 2 | 381 | 41,9 | 8,1 | 3,35 | 2 | 1,434 | 1,036 | 1,017 | 1,016 | 0,887 | 1,401 | 0,923 | 1,034 | 1,06 | 0,06 | 6,9E-01 |
| Q91W61 | F-box/LRR-repeat protein 15 OS=Mus musculus OX=10090 GN=Fbx | 4 | 1 | 1 | 1 | 300 | 33,1 | 7,43 | 0 | 1 | 0,899 | 0,714 | 1,151 | 0,916 | 0,803 | 1,389 | 0,971 | 0,791 | 0,93 | -0,07 | 6,9E-01 |
| Q9WV69 | Dematin OS=Mus musculus OX=10090 GN=Dmtn PE=1 SV=1 | 15 | 6 | 6 | 5 | 405 | 45,4 | 8,41 | 9,31 | 6 | 1,021 | 0,923 | 0,915 | 1,072 | 1,061 | 0,835 | 1,024 | 0,904 | 1,03 | 0,03 | 6,9E-01 |
| Q64336 | T-box brain protein 1 OS=Mus musculus OX=10090 GN=Tbr1 PE=1 S | 6 | 3 | 3 | 3 | 681 | 73,9 | 7,33 | 3,45 | 3 | 1,048 | 0,987 | 0,99 | 1,007 | 1,131 | 0,93 | 0,994 | 0,886 | 1,02 | 0,02 | 6,9E-01 |
| Q9Z204 | Heterogeneous nuclear ribonucleoproteins C1/C2 OS=Mus musculu | 8 | 2 | 2 | 2 | 313 | 34,4 | 5,05 | 2,51 | 2 | 1,189 | 1,39 | 1,069 | 0,955 | 0,916 | 1,451 | 1,004 | 0,979 | 1,06 | 0,06 | 7,0E-01 |
| Q61598 | Rab GDP dissociation inhibitor beta OS=Mus musculus OX=10090 G | 38 | 18 | 19 | 15 | 445 | 50,5 | 6,25 | 43,12 | 18 | 0,896 | 0,95 | 1,042 | 1,051 | 0,994 | 0,869 | 1,032 | 0,96 | 1,02 | 0,02 | 7,0E-01 |
| Q8R2R9 | AP-3 complex subunit mu-2 OS=Mus musculus OX=10090 GN=Ap3r | 16 | 6 | 10 | 6 | 418 | 46,9 | 7,56 | 21,75 | 6 | 1,162 | 1,116 | 1,052 | 1,106 | 0,985 | 1,102 | 1,107 | 1,169 | 1,02 | 0,02 | 7,0E-01 |
| Q63912 | Oligodendrocyte-myelin glycoprotein OS=Mus musculus OX=10090 | 9 | 4 | 4 | 4 | 440 | 49,3 | 8,41 | 8,93 | 4 | 1,467 | 0,748 | 0,963 | 0,814 | 1,651 | 0,769 | 1,095 | 0,893 | 0,91 | -0,09 | 7,0E-01 |
| Q99L04 | Dehydrogenase/reductase SDR family member 1 OS=Mus musculus | 27 | 8 | 9 | 8 | 313 | 34 | 8,35 | 11,09 | 8 | 1,119 | 0,958 | 1,086 | 0,998 | 1,087 | 0,97 | 1,014 | 1,017 | 1,02 | 0,02 | 7,0E-01 |
| Q80TV8 | CLIP-associating protein 1 OS=Mus musculus OX=10090 GN=Clasp1 | 5 | 7 | 7 | 6 | 1535 | 169,1 | 9,03 | 12,88 | 7 | 0,898 | 1,149 | 1,14 | 1,072 | 0,989 | 1,204 | 1,058 | 0,856 | 1,04 | 0,04 | 7,0E-01 |
| Q62074 | Protein kinase C iota type OS=Mus musculus OX=10090 GN=Prkci P | 2 | 1 | 1 | 1 | 595 | 68,2 | 5,85 | 0 | 1 | 1,088 | 0,966 | 0,578 | 1,003 | 0,911 | 1,036 | 0,864 | 1,021 | 0,95 | -0,05 | 7,0E-01 |
| O88986 | 2-amino-3-ketobutyrate coenzyme A ligase, mitochondrial OS=Mus | 8 | 2 | 3 | 2 | 416 | 44,9 | 7,33 | 5,42 | 2 | 1,545 | 1,212 | 0,891 | 1,008 | 0,81 | 1,226 | 1,024 | 1,301 | 1,07 | 0,07 | 7,0E-01 |

| | | | | | | | | | | | | | | | | | | | | | |
|---|---|---|---|---|---|---|---|---|---|---|---|---|---|---|---|---|---|---|---|---|---|
| Q99MN9 | Propionyl-CoA carboxylase beta chain, mitochondrial OS=Mus musc | 24 | 11 | 15 | 11 | 541 | 58,4 | 7,66 | 15,11 | 11 | 1,086 | 1,013 | 0,922 | 0,919 | 0,959 | 1,03 | 0,864 | 0,999 | 1,02 | 0,02 | 7,0E-01 |
| Q99LD8 | N(G),N(G)-dimethylarginine dimethylaminohydrolase 2 OS=Mus mu | 3 | 1 | 1 | 1 | 285 | 29,6 | 6,01 | 1,92 | 1 | 0,918 | 0,616 | 1,13 | 1,02 | 1,04 | 1,008 | 0,939 | 0,885 | 0,95 | -0,05 | 7,0E-01 |
| P61202 | COP9 signalosome complex subunit 2 OS=Mus musculus OX=10090 | 16 | 7 | 8 | 7 | 443 | 51,6 | 5,53 | 14,32 | 7 | 0,997 | 0,968 | 0,978 | 0,936 | 0,951 | 0,955 | 1,005 | 0,999 | 0,99 | -0,01 | 7,0E-01 |
| Q3U319 | E3 ubiquitin-protein ligase BRE1B OS=Mus musculus OX=10090 GN | 1 | 1 | 1 | 1 | 1001 | 113,9 | 6,48 | 0 | 1 | 1,656 | 0,767 | 1,385 | 1,011 | 0,814 | 1,526 | 0,944 | 1,13 | 1,09 | 0,09 | 7,0E-01 |
| Q9DC53 | Copine-8 OS=Mus musculus OX=10090 GN=Cpne8 PE=2 SV=3 | 11 | 5 | 6 | 2 | 577 | 64,6 | 5,97 | 6,78 | 5 | 1,465 | 1,023 | 1,048 | 1,096 | 0,85 | 1,042 | 1,18 | 1,326 | 1,05 | 0,05 | 7,0E-01 |
| Q66L44 | Voltage-dependent calcium channel beta subunit-associated regula | 4 | 3 | 3 | 3 | 698 | 74,1 | 6,33 | 7,66 | 3 | 1,142 | 1,085 | 1,028 | 1,052 | 1,177 | 1,03 | 1,018 | 1,006 | 1,02 | 0,02 | 7,0E-01 |
| Q8BL86 | Metallo-beta-lactamase domain-containing protein 2 OS=Mus mus | 13 | 3 | 4 | 3 | 279 | 31,2 | 6,86 | 4,08 | 3 | 1,119 | 1,169 | 0,985 | 0,977 | 0,955 | 1,173 | 0,889 | 1,102 | 1,03 | 0,03 | 7,0E-01 |
| P16546 | Spectrin alpha chain, non-erythrocytic 1 OS=Mus musculus OX=100 | 51 | 135 | 239 | 2 | 2472 | 284,4 | 5,33 | 614,85 | 135 | 0,858 | 0,859 | 1,034 | 0,967 | 1,059 | 0,761 | 1,03 | 0,998 | 0,97 | -0,03 | 7,0E-01 |
| P35278 | Ras-related protein Rab-5C OS=Mus musculus OX=10090 GN=Rab5 | 36 | 7 | 9 | 4 | 216 | 23,4 | 8,41 | 20,06 | 7 | 0,97 | 0,973 | 1,001 | 0,967 | 1,124 | 0,779 | 1,067 | 1,067 | 0,97 | -0,03 | 7,0E-01 |
| P61264 | Syntaxin-1B OS=Mus musculus OX=10090 GN=Stx1b PE=1 SV=1 | 49 | 19 | 51 | 18 | 288 | 33,2 | 5,38 | 126,13 | 19 | 0,983 | 1,064 | 0,986 | 1,009 | 0,915 | 1,069 | 0,996 | 1,003 | 1,01 | 0,01 | 7,0E-01 |
| Q8C167 | Prolyl endopeptidase-like OS=Mus musculus OX=10090 GN=Prepl P | 7 | 4 | 4 | 4 | 725 | 83,1 | 6,65 | 8,3 | 4 | 1,086 | 0,966 | 1,075 | 1,012 | 0,998 | 1,049 | 1,036 | 1,116 | 0,99 | -0,01 | 7,0E-01 |
| O88444 | Adenylate cyclase type 1 OS=Mus musculus OX=10090 GN=Adcy1 P | 2 | 2 | 2 | 2 | 1118 | 123,3 | 8,47 | 5,63 | 2 | 0,513 | 0,993 | 0,727 | 1,009 | 0,928 | 0,835 | 0,703 | 0,991 | 0,94 | -0,06 | 7,0E-01 |
| Q8BSY0 | Aspartyl/asparaginyl beta-hydroxylase OS=Mus musculus OX=1009 | 2 | 1 | 1 | 1 | 741 | 83 | 5,08 | 2,99 | 1 | 1,304 | 1,111 | 0,827 | 0,857 | 0,643 | 1,344 | 0,638 | 1,136 | 1,09 | 0,09 | 7,0E-01 |
| Q8BFY6 | Peflin OS=Mus musculus OX=10090 GN=Pef1 PE=1 SV=1 | 13 | 3 | 4 | 3 | 275 | 29,2 | 6,3 | 9,08 | 3 | 0,869 | 0,937 | 0,794 | 0,828 | 0,882 | 0,863 | 0,95 | 0,802 | 0,98 | -0,02 | 7,0E-01 |
| P61161 | Actin-related protein 2 OS=Mus musculus OX=10090 GN=Actr2 PE= | 20 | 8 | 11 | 8 | 394 | 44,7 | 6,74 | 28,61 | 8 | 1,033 | 1,012 | 1,048 | 1,097 | 1,006 | 0,989 | 1,109 | 1,035 | 1,01 | 0,01 | 7,0E-01 |
| Q8BGR2 | Volume-regulated anion channel subunit LRRC8D OS=Mus musculu | 1 | 1 | 1 | 1 | 859 | 98,1 | 7,44 | 2,27 | 1 | 0,957 | 1,062 | 1,022 | 1,032 | 1,022 | 1,045 | 0,867 | 1,06 | 1,02 | 0,02 | 7,0E-01 |
| O35943 | Frataxin, mitochondrial OS=Mus musculus OX=10090 GN=Fxn PE=1 | 12 | 2 | 2 | 2 | 207 | 22,9 | 8,02 | 6,09 | 2 | 0,805 | 0,841 | 1,106 | 1,009 | 1,036 | 0,828 | 0,905 | 0,858 | 1,04 | 0,04 | 7,1E-01 |
| Q920Q6 | RNA-binding protein Musashi homolog 2 OS=Mus musculus OX=10 | 11 | 3 | 3 | 3 | 346 | 36,9 | 8,47 | 6,29 | 3 | 1,129 | 0,897 | 0,98 | 1,142 | 1,007 | 1,279 | 1,005 | 1,001 | 0,97 | -0,03 | 7,1E-01 |
| A2APY7 | Arginine-hydroxylase NDUFAF5, mitochondrial OS=Mus musculus C | 3 | 1 | 2 | 1 | 343 | 38,4 | 6,1 | 4,98 | 1 | 0,718 | 1,07 | 0,948 | 1,09 | 0,825 | 0,989 | 1,162 | 1,024 | 0,96 | -0,04 | 7,1E-01 |
| Q7TMF3 | NADH dehydrogenase [ubiquinone] 1 alpha subcomplex subunit 12 | 50 | 7 | 9 | 7 | 145 | 17,1 | 9,36 | 24,21 | 7 | 1,114 | 1,071 | 1,046 | 1,034 | 1,001 | 1,276 | 1,027 | 1,064 | 0,98 | -0,02 | 7,1E-01 |
| Q80ZS3 | 28S ribosomal protein S26, mitochondrial OS=Mus musculus OX=10 | 11 | 2 | 2 | 2 | 200 | 23,4 | 9,96 | 4,91 | 2 | 1,275 | 0,872 | 0,99 | 0,787 | 0,925 | 1,146 | 0,931 | 1,114 | 0,95 | -0,05 | 7,1E-01 |
| Q9DCQ7 | Alpha-1-antitrypsin 1-6 OS=Mus musculus OX=10090 GN=Serpina1 | 2 | 1 | 2 | 1 | 411 | 46,3 | 8,59 | 2,2 | 1 | 1,41 | 1,104 | 1,195 | 0,976 | 0,943 | 1,305 | 1,114 | 1,138 | 1,04 | 0,04 | 7,1E-01 |
| P25785 | Metalloproteinase inhibitor 2 OS=Mus musculus OX=10090 GN=Tin | 9 | 1 | 1 | 1 | 220 | 24,3 | 7,49 | 0 | 1 | 1,603 | 1,11 | 1,047 | 0,951 | 1,276 | 1,341 | 0,842 | 0,954 | 1,07 | 0,07 | 7,1E-01 |
| P55937 | Golgin subfamily A member 3 OS=Mus musculus OX=10090 GN=Go | 6 | 6 | 6 | 6 | 1487 | 167,1 | 5,4 | 4,51 | 6 | 0,905 | 1,198 | 0,549 | 1,246 | 1,138 | 0,939 | 0,935 | 1,156 | 0,94 | -0,06 | 7,1E-01 |
| Q9ESP1 | Stromal cell-derived factor 2-like protein 1 OS=Mus musculus OX=1 | 4 | 1 | 1 | 1 | 221 | 23,6 | 7,42 | 0 | 1 | 1,218 | 0,99 | 0,889 | 0,878 | 1,053 | 0,981 | 0,867 | 1,251 | 0,96 | -0,04 | 7,1E-01 |
| Q922J3 | CAP-Gly domain-containing linker protein 1 OS=Mus musculus OX= | 21 | 27 | 27 | 21 | 1391 | 155,7 | 5,24 | 59,91 | 27 | 0,899 | 0,953 | 1,034 | 1,015 | 1,015 | 0,961 | 1,034 | 0,949 | 0,99 | -0,01 | 7,1E-01 |
| Q9CXR1 | Dehydrogenase/reductase SDR family member 7 OS=Mus musculus | 3 | 1 | 2 | 1 | 338 | 38,1 | 8,32 | 4,63 | 1 | 1,302 | 1,216 | 1,009 | 1,119 | 0,982 | 1,349 | 1,098 | 1,058 | 1,04 | 0,04 | 7,1E-01 |
| Q9R1K9 | Centrin-2 OS=Mus musculus OX=10090 GN=Cetn2 PE=1 SV=1 | 19 | 3 | 4 | 3 | 172 | 19,8 | 5 | 8,18 | 3 | 0,875 | 0,885 | 0,923 | 1,004 | 0,932 | 0,844 | 1,052 | 0,94 | 0,98 | -0,02 | 7,1E-01 |
| P49442 | Inositol polyphosphate 1-phosphatase OS=Mus musculus OX=1009 | 9 | 4 | 4 | 4 | 396 | 43,3 | 5,01 | 5,3 | 4 | 1,156 | 1,043 | 1,076 | 1,107 | 0,838 | 1,267 | 1,101 | 1,032 | 1,03 | 0,03 | 7,1E-01 |
| Q8C9H6 | Striatin-interacting proteins 2 OS=Mus musculus OX=10090 GN=Str | 7 | 5 | 5 | 3 | 844 | 96,2 | 5,77 | 3,85 | 5 | 1,33 | 0,844 | 1,173 | 0,807 | 1,297 | 1,326 | 1,05 | 0,766 | 0,94 | -0,06 | 7,1E-01 |
| Q8K1X1 | WD repeat-containing protein 11 OS=Mus musculus OX=10090 GN= | 5 | 5 | 5 | 5 | 1223 | 135,9 | 7,11 | 5,37 | 5 | 0,941 | 0,983 | 0,865 | 0,894 | 0,874 | 1,051 | 0,855 | 0,811 | 1,03 | 0,03 | 7,1E-01 |
| Q9CQE1 | Protein NipSnap homolog 3B OS=Mus musculus OX=10090 GN=Nip | 9 | 2 | 2 | 2 | 247 | 28,3 | 9,48 | 5,19 | 2 | 1 | 0,997 | 0,891 | 0,805 | 0,877 | 0,939 | 0,967 | 0,993 | 0,98 | -0,02 | 7,1E-01 |
| Q9DB73 | NADH-cytochrome b5 reductase 1 OS=Mus musculus OX=10090 GN | 15 | 4 | 5 | 4 | 305 | 34,1 | 8,87 | 10,15 | 4 | 1,037 | 1,005 | 0,928 | 0,92 | 0,902 | 1,102 | 0,941 | 1,028 | 0,98 | -0,02 | 7,1E-01 |
| P50446 | Keratin, type II cytoskeletal 6A OS=Mus musculus OX=10090 GN=Kr | 16 | 12 | 16 | 3 | 553 | 59,3 | 7,94 | 24,72 | 12 | 1,318 | 0,939 | 0,719 | 0,424 | 1,507 | 1,127 | 0,547 | 0,67 | 0,88 | -0,12 | 7,1E-01 |
| Q60634 | Flotillin-2 OS=Mus musculus OX=10090 GN=Flot2 PE=1 SV=2 | 39 | 16 | 28 | 16 | 428 | 47 | 5,2 | 60,97 | 16 | 0,998 | 0,982 | 1,086 | 1,128 | 1,019 | 0,93 | 1,056 | 1,109 | 1,02 | 0,02 | 7,1E-01 |
| Q8R151 | NFX1-type zinc finger-containing protein 1 OS=Mus musculus OX=1 | 2 | 3 | 3 | 3 | 1909 | 218,7 | 7,47 | 4,07 | 3 | 1,061 | 1,114 | 0,925 | 1,072 | 1,048 | 1,033 | 1,192 | 0,992 | 0,98 | -0,02 | 7,1E-01 |
| Q9R111 | Guanine deaminase OS=Mus musculus OX=10090 GN=Gda PE=1 SV | 39 | 14 | 21 | 14 | 454 | 51 | 5,53 | 53,51 | 14 | 0,884 | 0,878 | 0,935 | 1,052 | 0,972 | 0,872 | 1,021 | 0,963 | 0,98 | -0,02 | 7,1E-01 |
| Q6NZM9 | Histone deacetylase 4 OS=Mus musculus OX=10090 GN=Hdac4 PE= | 4 | 3 | 3 | 3 | 1076 | 118,5 | 6,92 | 4,42 | 3 | 0,914 | 0,662 | 0,941 | 0,938 | 0,877 | 1,059 | 0,705 | 0,971 | 0,96 | -0,04 | 7,1E-01 |
| Q3UH68 | LIM and calponin homology domains-containing protein 1 OS=Mus | 1 | 1 | 2 | 1 | 1057 | 118,1 | 5,48 | 5,73 | 1 | 1,466 | 2,053 | 0,652 | 1,322 | 1,743 | 1,602 | 1,135 | 1,501 | 0,92 | -0,08 | 7,1E-01 |
| Q99PL5 | Ribosome-binding protein 1 OS=Mus musculus OX=10090 GN=Rrbp | 1 | 2 | 2 | 2 | 1605 | 172,8 | 9,33 | 2,07 | 2 | 0,761 | 0,952 | 1,175 | 0,998 | 0,95 | 0,771 | 0,948 | 1,057 | 1,04 | 0,04 | 7,1E-01 |
| Q9CZX0 | Elongator complex protein 3 OS=Mus musculus OX=10090 GN=Elp3 | 5 | 3 | 3 | 3 | 547 | 62,3 | 8,9 | 4,92 | 3 | 1,141 | 0,934 | 1,091 | 0,92 | 1,089 | 1,138 | 1,033 | 0,935 | 0,97 | -0,03 | 7,1E-01 |
| Q920I9 | WD repeat-containing protein 7 OS=Mus musculus OX=10090 GN= | 19 | 23 | 30 | 23 | 1489 | 163,3 | 6,9 | 68,82 | 23 | 1,207 | 1,068 | 1,087 | 1,029 | 1,016 | 1,273 | 1,125 | 1,08 | 0,98 | -0,02 | 7,1E-01 |
| P38060 | Hydroxymethylglutaryl-CoA lyase, mitochondrial OS=Mus musculus | 12 | 4 | 7 | 4 | 325 | 34,2 | 8,41 | 14,52 | 4 | 0,913 | 0,887 | 1,044 | 1,084 | 1,061 | 0,933 | 1,04 | 0,981 | 0,98 | -0,02 | 7,1E-01 |
| Q8BQP9 | Regulator of G-protein signaling 7-binding protein OS=Mus muscul | 18 | 3 | 3 | 3 | 257 | 29 | 8,46 | 3,34 | 3 | 1,327 | 1,155 | 1,353 | 1,157 | 1,082 | 1,552 | 1,235 | 1,295 | 0,97 | -0,03 | 7,1E-01 |
| Q9JIG8 | PRA1 family protein 2 OS=Mus musculus OX=10090 GN=Praf2 PE=1 | 12 | 2 | 3 | 2 | 178 | 19,5 | 9,6 | 9,19 | 2 | 0,971 | 1,066 | 0,851 | 0,963 | 1,029 | 0,865 | 1,008 | 0,85 | 1,03 | 0,03 | 7,1E-01 |
| P61458 | Pterin-4-alpha-carbinolamine dehydratase OS=Mus musculus OX=1 | 10 | 1 | 1 | 1 | 104 | 12 | 6,8 | 2,44 | 1 | 0,867 | 1,172 | 1,055 | 1,175 | 0,837 | 1,246 | 0,993 | 1,022 | 1,04 | 0,04 | 7,1E-01 |
| Q8R4N0 | Citramalyl-CoA lyase, mitochondrial OS=Mus musculus OX=10090 C | 24 | 8 | 11 | 8 | 338 | 37,5 | 8,54 | 31,13 | 8 | 1,348 | 1,15 | 0,967 | 0,957 | 1,054 | 1,271 | 1,122 | 1,133 | 0,97 | -0,03 | 7,1E-01 |
| Q64514 | Tripeptidyl-peptidase 2 OS=Mus musculus OX=10090 GN=Tpp2 PE= | 10 | 14 | 17 | 14 | 1262 | 139,8 | 6,58 | 33,05 | 14 | 1,003 | 0,851 | 1,03 | 1,079 | 1,011 | 1,059 | 0,927 | 1,055 | 0,98 | -0,02 | 7,1E-01 |
| P35436 | Glutamate receptor ionotropic, NMDA 2A OS=Mus musculus OX=1 | 3 | 4 | 5 | 4 | 1464 | 165,3 | 7,01 | 6,93 | 4 | 1,13 | 0,984 | 1,03 | 0,981 | 1,099 | 1,101 | 0,943 | 1,06 | 0,98 | -0,02 | 7,1E-01 |
| Q8BG89 | Protein ZNF365 OS=Mus musculus OX=10090 GN=Znf365 PE=1 SV= | 2 | 1 | 1 | 1 | 408 | 46,8 | 9,14 | 2,18 | 1 | 1,061 | 1,293 | 1,112 | 0,997 | 1,02 | 1,03 | 1,073 | 1,22 | 1,03 | 0,03 | 7,2E-01 |
| P70168 | Importin subunit beta-1 OS=Mus musculus OX=10090 GN=Kpnb1 P | 15 | 11 | 14 | 11 | 876 | 97,1 | 4,78 | 27,06 | 11 | 1,062 | 1,053 | 1,087 | 1,068 | 1,032 | 1,117 | 1,068 | 1,082 | 0,99 | -0,01 | 7,2E-01 |
| P28660 | Nck-associated protein 1 OS=Mus musculus OX=10090 GN=Nckap1 | 19 | 25 | 34 | 25 | 1128 | 128,7 | 6,62 | 63,43 | 25 | 1,011 | 1,126 | 1,007 | 1,028 | 1,011 | 1,1 | 1,086 | 1,029 | 0,99 | -0,01 | 7,2E-01 |
| Q8CBY8 | Dynactin subunit 4 OS=Mus musculus OX=10090 GN=Dctn4 PE=1 S | 9 | 3 | 5 | 3 | 467 | 53 | 7,72 | 2,95 | 3 | 1,03 | 1,061 | 1,026 | 1,092 | 1,093 | 1,267 | 0,95 | 1,007 | 0,97 | -0,03 | 7,2E-01 |
| Q6DFV3 | Rho GTPase-activating protein 21 OS=Mus musculus OX=10090 GN | 4 | 7 | 7 | 7 | 1944 | 215,6 | 7,64 | 9,73 | 7 | 1,609 | 1,332 | 1,163 | 1,269 | 1,257 | 1,477 | 1,19 | 1,276 | 1,03 | 0,03 | 7,2E-01 |
| Q9Z1K5 | E3 ubiquitin-protein ligase ARIH1 OS=Mus musculus OX=10090 GN= | 7 | 3 | 3 | 3 | 555 | 64 | 5,08 | 8,7 | 3 | 1,273 | 1,332 | 1,586 | 1,198 | 1,346 | 1,039 | 1,674 | 1,581 | 0,96 | -0,04 | 7,2E-01 |
| Q922G0 | Solute carrier family 25 member 36 OS=Mus musculus OX=10090 G | 9 | 2 | 2 | 2 | 311 | 34,1 | 8,65 | 5,73 | 2 | 1,424 | 1,603 | 1,42 | 1,29 | 1,362 | 1,652 | 1,296 | 1,26 | 1,03 | 0,03 | 7,2E-01 |
| Q8JZW5 | SH2 domain-containing protein 5 OS=Mus musculus OX=10090 GN= | 8 | 3 | 3 | 3 | 429 | 47,4 | 8,47 | 2,38 | 3 | 1,172 | 0,763 | 0,801 | 0,918 | 0,769 | 1,27 | 0,818 | 1,02 | 0,94 | -0,06 | 7,2E-01 |
| Q571H0 | Nucleolar pre-ribosomal-associated protein 1 OS=Mus musculus O | 0 | 1 | 2 | 1 | 2274 | 254,5 | 6,96 | 3,96 | 1 | 0,959 | 0,897 | 1,13 | 1,151 | 1,135 | 1,046 | 1,103 | 0,964 | 0,97 | -0,03 | 7,2E-01 |
| P31230 | Aminoacyl tRNA synthase complex-interacting multifunctional prot | 13 | 3 | 3 | 3 | 310 | 34 | 8,35 | 6,82 | 3 | 1,046 | 0,92 | 1,093 | 1,021 | 1,012 | 1,023 | 0,951 | 1,032 | 1,02 | 0,02 | 7,2E-01 |
| Q9DCG9 | Multifunctional methyltransferase subunit TRM112-like protein OS | 10 | 1 | 1 | 1 | 125 | 14,1 | 5,27 | 0 | 1 | 1,117 | 0,811 | 0,999 | 0,875 | 0,538 | 1,005 | 0,945 | 1,1 | 1,06 | 0,06 | 7,2E-01 |
| Q9Z2I8 | Succinate--CoA ligase [GDP-forming] subunit beta, mitochondrial O | 7 | 3 | 3 | 3 | 433 | 46,8 | 7,02 | 5,32 | 3 | 0,867 | 1,083 | 0,986 | 1,015 | 1,039 | 0,88 | 0,973 | 1,175 | 0,97 | -0,03 | 7,2E-01 |
| Q9DBL7 | Bifunctional coenzyme A synthase OS=Mus musculus OX=10090 GN | 11 | 5 | 7 | 5 | 563 | 62 | 7,11 | 7,72 | 5 | 0,859 | 0,973 | 0,977 | 1,275 | 0,995 | 1,004 | 1,069 | 0,867 | 1,04 | 0,04 | 7,2E-01 |

| | | | | | | | | | | | | | | | | | | | | | |
|---|---|---|---|---|---|---|---|---|---|---|---|---|---|---|---|---|---|---|---|---|---|
| Q3UHD3 | Microtubule-associated tumor suppressor candidate 2 homolog OS | 4 | 5 | 6 | 5 | 1353 | 147,3 | 8,24 | 12,25 | 5 | 1,222 | 1,195 | 0,982 | 1,093 | 1,092 | 1,114 | 1,093 | 1,11 | 1,02 | 0,02 | 7,2E-01 |
| P52480-2 | Isoform M1 of Pyruvate kinase PKM OS=Mus musculus OX=10090 ( | 44 | 25 | 70 | 2 | 531 | 57,9 | 7,14 | 188,32 | 25 | 0,889 | 0,923 | 0,9 | 1,103 | 0,88 | 0,773 | 1,34 | 1,023 | 0,95 | -0,05 | 7,2E-01 |
| Q9QZC2 | Plexin-C1 OS=Mus musculus OX=10090 GN=Plxnc1 PE=1 SV=1 | 3 | 4 | 4 | 4 | 1574 | 176,4 | 7,75 | 4,39 | 4 | 1,133 | 0,86 | 0,903 | 1,027 | 0,852 | 1,155 | 1,058 | 0,992 | 0,97 | -0,03 | 7,2E-01 |
| Q8BGN5 | NIPA-like protein 3 OS=Mus musculus OX=10090 GN=Nipal3 PE=2 S | 2 | 1 | 1 | 1 | 410 | 44,8 | 7,75 | 3,42 | 1 | 0,779 | 1,101 | 1,111 | 1,112 | 1,116 | 0,977 | 1,195 | 0,965 | 0,96 | -0,04 | 7,2E-01 |
| P14733 | Lamin-B1 OS=Mus musculus OX=10090 GN=Lmnb1 PE=1 SV=3 | 16 | 11 | 11 | 9 | 588 | 66,7 | 5,16 | 22,72 | 11 | 1,26 | 1,481 | 1,076 | 0,893 | 1,226 | 1,645 | 0,841 | 1,309 | 0,94 | -0,06 | 7,2E-01 |
| Q9CPR4 | 60S ribosomal protein L17 OS=Mus musculus OX=10090 GN=Rpl17 | 14 | 3 | 3 | 3 | 184 | 21,4 | 10,18 | 5,46 | 3 | 1,103 | 1,084 | 1,152 | 0,983 | 1,084 | 1,197 | 1,048 | 1,066 | 0,98 | -0,02 | 7,2E-01 |
| Q99NE5-7 | Isoform 7 of Regulating synaptic membrane exocytosis protein 1 O | 14 | 13 | 15 | 3 | 1184 | 132,9 | 9,33 | 22,47 | 13 | 1,07 | 0,946 | 1,051 | 1,042 | 1,168 | 1,063 | 0,966 | 0,991 | 0,98 | -0,02 | 7,2E-01 |
| P47754 | F-actin-capping protein subunit alpha-2 OS=Mus musculus OX=100 | 46 | 8 | 11 | 8 | 286 | 32,9 | 5,85 | 19,65 | 8 | 1,288 | 1,033 | 1,037 | 0,924 | 0,96 | 1,157 | 0,999 | 1,035 | 1,03 | 0,03 | 7,2E-01 |
| Q8R554 | OTU domain-containing protein 7A OS=Mus musculus OX=10090 G | 4 | 3 | 3 | 2 | 926 | 100,7 | 8,22 | 0 | 3 | 0,968 | 1,032 | 0,618 | 0,967 | 1,007 | 0,745 | 0,773 | 0,894 | 1,05 | 0,05 | 7,2E-01 |
| Q9EP69 | Phosphatidylinositide phosphatase SAC1 OS=Mus musculus OX=10 | 13 | 8 | 12 | 8 | 587 | 66,9 | 7,3 | 16,44 | 8 | 1,378 | 1,093 | 0,934 | 0,91 | 1,012 | 1,149 | 0,905 | 1,072 | 1,04 | 0,04 | 7,2E-01 |
| Q8VCE6 | 5'(3')-deoxyribonucleotidase, mitochondrial OS=Mus musculus OX= | 4 | 1 | 1 | 1 | 220 | 25,6 | 8,35 | 2 | 1 | 0,901 | 1,058 | 0,896 | 1,075 | 1,057 | 0,929 | 1,146 | 0,908 | 0,97 | -0,03 | 7,2E-01 |
| Q5DTZ6 | RING finger protein 150 OS=Mus musculus OX=10090 GN=Rnf150 F | 3 | 1 | 1 | 1 | 437 | 48 | 5,43 | 0 | 1 | 1,36 | 0,639 | 0,929 | 0,676 | 0,996 | 0,41 | 0,96 | 0,918 | 1,10 | 0,10 | 7,2E-01 |
| Q61768 | Kinesin-1 heavy chain OS=Mus musculus OX=10090 GN=Kif5b PE=1 | 18 | 15 | 19 | 13 | 963 | 109,5 | 6,44 | 33,19 | 15 | 0,952 | 0,779 | 1,087 | 1,069 | 0,992 | 0,927 | 1,062 | 1,019 | 0,97 | -0,03 | 7,2E-01 |
| P62245 | 40S ribosomal protein S15a OS=Mus musculus OX=10090 GN=Rps1 | 5 | 1 | 1 | 1 | 130 | 14,8 | 10,13 | 0 | 1 | 1,224 | 1,371 | 1,305 | 1,053 | 1,37 | 1,07 | 0,989 | 1,82 | 0,94 | -0,06 | 7,2E-01 |
| Q9CX34 | Protein SGT1 homolog OS=Mus musculus OX=10090 GN=Sugt1 PE= | 21 | 6 | 6 | 6 | 336 | 38,1 | 5,45 | 12,8 | 6 | 1,044 | 1,144 | 1,143 | 1,134 | 1,062 | 1,095 | 1,017 | 1,218 | 1,02 | 0,02 | 7,2E-01 |
| Q9Z2Q6 | Septin-5 OS=Mus musculus OX=10090 GN=Septin5 PE=1 SV=2 | 35 | 15 | 28 | 14 | 369 | 42,7 | 6,67 | 49,57 | 15 | 0,813 | 0,832 | 0,974 | 1,051 | 1,01 | 0,845 | 0,95 | 0,963 | 0,97 | -0,03 | 7,3E-01 |
| P08228 | Superoxide dismutase [Cu-Zn] OS=Mus musculus OX=10090 GN=So | 53 | 8 | 16 | 8 | 154 | 15,9 | 6,51 | 42,38 | 8 | 1,027 | 0,931 | 1,077 | 1,154 | 1,041 | 0,928 | 1,068 | 1,068 | 1,02 | 0,02 | 7,3E-01 |
| Q61739 | Integrin alpha-6 OS=Mus musculus OX=10090 GN=Itga6 PE=1 SV=3 | 2 | 2 | 2 | 2 | 1091 | 122,1 | 7,03 | 4,99 | 2 | 0,917 | 0,88 | 1,2 | 1,092 | 1,017 | 0,923 | 1,041 | 0,992 | 1,03 | 0,03 | 7,3E-01 |
| P04370-4 | Isoform 4 of Myelin basic protein OS=Mus musculus OX=10090 GN= | 37 | 8 | 48 | 1 | 195 | 21,5 | 11,24 | 131,33 | 8 | 0,943 | 0,802 | 0,714 | 1,235 | 1,168 | 0,977 | 0,773 | 0,98 | 0,95 | -0,05 | 7,3E-01 |
| Q3TIU4 | 2',5'-phosphodiesterase 12 OS=Mus musculus OX=10090 GN=Pde1 | 4 | 2 | 2 | 2 | 608 | 67,5 | 6,92 | 2,02 | 2 | 1,711 | 1,316 | 1,25 | 1,161 | 1,262 | 1,525 | 1,227 | 1,657 | 0,96 | -0,04 | 7,3E-01 |
| Q9JMH9-1 | Isoform 1 of Unconventional myosin-XVIIIa OS=Mus musculus OX= | 17 | 31 | 39 | 31 | 2035 | 230,8 | 6,16 | 71,25 | 31 | 1,101 | 1,072 | 1,027 | 0,997 | 0,987 | 1,091 | 1,029 | 1,044 | 1,01 | 0,01 | 7,3E-01 |
| Q9Z2W9 | Glutamate receptor 3 OS=Mus musculus OX=10090 GN=Gria3 PE=1 | 19 | 17 | 21 | 13 | 888 | 100,5 | 8,38 | 50,42 | 17 | 0,923 | 1,013 | 0,991 | 0,971 | 1 | 1,035 | 0,878 | 1,048 | 0,98 | -0,02 | 7,3E-01 |
| Q6IME9 | Keratin, type II cytoskeletal 72 OS=Mus musculus OX=10090 GN=Kr | 4 | 2 | 5 | 1 | 520 | 56,7 | 7,4 | 5,49 | 2 | 1,205 | 1,412 | 1,128 | 0,777 | 1,302 | 1,096 | 0,731 | 1,133 | 1,06 | 0,06 | 7,3E-01 |
| Q9DCM2 | Glutathione S-transferase kappa 1 OS=Mus musculus OX=10090 GN | 12 | 2 | 3 | 2 | 226 | 25,7 | 8,88 | 4,1 | 2 | 1,2 | 1,186 | 1,188 | 1,018 | 0,975 | 1,309 | 1,135 | 1,051 | 1,03 | 0,03 | 7,3E-01 |
| A2CG49 | Kalirin OS=Mus musculus OX=10090 GN=Kalrn PE=1 SV=1 | 11 | 25 | 30 | 3 | 2964 | 336,8 | 6,07 | 68,26 | 25 | 0,771 | 0,909 | 0,968 | 0,892 | 0,99 | 0,879 | 0,724 | 1,072 | 0,97 | -0,03 | 7,3E-01 |
| Q8VCM8 | Nicalin OS=Mus musculus OX=10090 GN=Ncln PE=1 SV=2 | 3 | 2 | 2 | 2 | 563 | 62,9 | 6,49 | 4,23 | 2 | 0,721 | 0,942 | 0,934 | 0,789 | 0,868 | 0,861 | 0,7 | 0,858 | 1,03 | 0,03 | 7,3E-01 |
| Q9D6K5 | Synaptojanin-2-binding protein OS=Mus musculus OX=10090 GN=S | 14 | 2 | 3 | 2 | 145 | 15,8 | 6,3 | 10,05 | 2 | 1,137 | 0,96 | 1,13 | 1,017 | 1,245 | 1,212 | 0,951 | 0,966 | 0,97 | -0,03 | 7,3E-01 |
| Q9CYH5 | Glucose-fructose oxidoreductase domain-containing protein 2 OS= | 3 | 1 | 1 | 1 | 385 | 42,2 | 6,25 | 0 | 1 | 0,01 | 1,061 | 1,305 | 1,369 | 0,87 | 1,174 | 1,109 | 1,059 | 0,89 | -0,11 | 7,3E-01 |
| P97813 | Phospholipase D2 OS=Mus musculus OX=10090 GN=Pld2 PE=1 SV= | 4 | 4 | 4 | 4 | 933 | 106,1 | 7,39 | 4,73 | 4 | 1,083 | 1,07 | 0,98 | 0,88 | 0,963 | 1,079 | 1,027 | 1,02 | 0,98 | -0,02 | 7,3E-01 |
| P56818 | Beta-secretase 1 OS=Mus musculus OX=10090 GN=Bace1 PE=1 SV= | 2 | 1 | 1 | 1 | 501 | 55,7 | 5,6 | 3,04 | 1 | 0,896 | 0,946 | 0,757 | 1,135 | 0,887 | 0,938 | 1,105 | 0,936 | 0,97 | -0,03 | 7,3E-01 |
| Q99020 | Heterogeneous nuclear ribonucleoprotein A/B OS=Mus musculus C | 13 | 3 | 3 | 2 | 285 | 30,8 | 7,91 | 3,55 | 3 | 1,026 | 0,99 | 1,004 | 0,71 | 0,942 | 1,221 | 0,804 | 0,929 | 0,96 | -0,04 | 7,3E-01 |
| P43404 | Tyrosine-protein kinase ZAP-70 OS=Mus musculus OX=10090 GN=Z | 1 | 1 | 1 | 1 | 618 | 70,1 | 7,83 | 2,03 | 1 | 1,021 | 0,964 | 1,175 | 0,885 | 0,959 | 0,986 | 0,977 | 1,032 | 1,02 | 0,02 | 7,3E-01 |
| B2RXA1 | PI-PLC X domain-containing protein 2 OS=Mus musculus OX=10090 | 2 | 1 | 1 | 1 | 340 | 38,6 | 8,88 | 1,84 | 1 | 1,02 | 1,482 | 1,215 | 1,373 | 1,289 | 1,057 | 1,613 | 0,856 | 1,06 | 0,06 | 7,3E-01 |
| Q91WQ3 | Tyrosine--tRNA ligase, cytoplasmic OS=Mus musculus OX=10090 G | 32 | 17 | 22 | 17 | 528 | 59,1 | 7,01 | 41,78 | 17 | 0,937 | 0,995 | 0,973 | 1,016 | 0,935 | 1,058 | 0,954 | 1,022 | 0,99 | -0,01 | 7,3E-01 |
| Q80ZJ6 | Protein zer-1 homolog OS=Mus musculus OX=10090 GN=Zer1 PE=1 | 3 | 2 | 2 | 2 | 779 | 89 | 5,94 | 2,14 | 2 | 1,03 | 0,913 | 1,046 | 0,877 | 1,154 | 1,052 | 0,842 | 0,933 | 0,97 | -0,03 | 7,3E-01 |
| Q9WV60 | Glycogen synthase kinase-3 beta OS=Mus musculus OX=10090 GN= | 42 | 12 | 14 | 8 | 420 | 46,7 | 8,78 | 30,59 | 12 | 0,897 | 0,914 | 0,927 | 0,99 | 0,982 | 0,871 | 0,944 | 0,978 | 0,99 | -0,01 | 7,3E-01 |
| P62073 | Mitochondrial import inner membrane translocase subunit Tim10 ( | 32 | 5 | 11 | 5 | 90 | 10,3 | 6,29 | 17,16 | 5 | 0,822 | 1,098 | 0,973 | 1,016 | 1,005 | 0,948 | 0,918 | 0,951 | 1,02 | 0,02 | 7,3E-01 |
| Q148V7-2 | Isoform 2 of RAB11-binding protein RELCH OS=Mus musculus OX=1 | 9 | 10 | 10 | 10 | 1192 | 131,8 | 5,29 | 19,88 | 10 | 0,942 | 0,98 | 0,97 | 0,84 | 0,896 | 0,882 | 0,989 | 1,035 | 0,98 | -0,02 | 7,3E-01 |
| Q5DU14 | Unconventional myosin-XVI OS=Mus musculus OX=10090 GN=Myo | 0 | 1 | 1 | 1 | 1919 | 211,2 | 6,9 | 2,84 | 1 | 0,833 | 0,954 | 0,906 | 0,849 | 0,88 | 0,799 | 0,965 | 0,968 | 0,98 | -0,02 | 7,3E-01 |
| P51163 | Uroporphyrinogen-III synthase OS=Mus musculus OX=10090 GN=U | 9 | 2 | 3 | 2 | 265 | 28,5 | 6,54 | 6,77 | 2 | 1,017 | 1,456 | 1,276 | 1,065 | 1,108 | 1,312 | 1,162 | 1,068 | 1,04 | 0,04 | 7,3E-01 |
| O08810 | 116 kDa U5 small nuclear ribonucleoprotein component OS=Mus m | 2 | 2 | 2 | 2 | 971 | 109,3 | 5 | 4,16 | 2 | 1,415 | 1,202 | 1,12 | 1,205 | 1,359 | 1,389 | 1,085 | 0,926 | 1,04 | 0,04 | 7,3E-01 |
| B7ZMP1 | Xaa-Pro aminopeptidase 3 OS=Mus musculus OX=10090 GN=Xpnpe | 4 | 2 | 2 | 2 | 506 | 56,6 | 7,68 | 2,3 | 2 | 1,245 | 1,189 | 1,262 | 1,234 | 1,091 | 1,216 | 1,294 | 1,434 | 0,98 | -0,02 | 7,3E-01 |
| Q9D5T0 | ATPase family AAA domain-containing protein 1 OS=Mus musculus | 7 | 2 | 2 | 2 | 361 | 40,7 | 6,9 | 2,6 | 2 | 1,019 | 0,433 | 0,728 | 0,766 | 0,809 | 0,83 | 0,659 | 0,829 | 0,94 | -0,06 | 7,3E-01 |
| Q99JN2 | Kelch-like protein 22 OS=Mus musculus OX=10090 GN=Klhl22 PE=1 | 2 | 1 | 1 | 1 | 634 | 71,6 | 5,71 | 0 | 1 | 2,971 | 1,803 | 2,428 | 1,329 | 2,732 | 1,88 | 2,155 | 2,335 | 0,94 | -0,06 | 7,3E-01 |
| Q8BTI9 | Phosphatidylinositol 4,5-bisphosphate 3-kinase catalytic subunit be | 3 | 3 | 3 | 3 | 1064 | 121,6 | 7,09 | 2,02 | 3 | 0,877 | 1,11 | 1,403 | 1,251 | 1,362 | 1,03 | 1,493 | 0,993 | 0,95 | -0,05 | 7,3E-01 |
| P56480 | ATP synthase subunit beta, mitochondrial OS=Mus musculus OX=1( | 48 | 24 | 80 | 24 | 529 | 56,3 | 5,34 | 237,44 | 24 | 1,207 | 1,132 | 1,042 | 0,971 | 0,989 | 1,172 | 1,012 | 1,085 | 1,02 | 0,02 | 7,3E-01 |
| Q920M7 | Synaptotagmin-17 OS=Mus musculus OX=10090 GN=Syt17 PE=1 SV | 12 | 5 | 6 | 5 | 470 | 53,3 | 7,15 | 11,84 | 5 | 0,825 | 0,913 | 0,999 | 0,998 | 0,934 | 0,822 | 1,08 | 0,996 | 0,97 | -0,03 | 7,3E-01 |
| P42232 | Signal transducer and activator of transcription 5B OS=Mus muscul | 2 | 1 | 1 | 1 | 786 | 89,9 | 6,05 | 2,97 | 1 | 0,821 | 1,378 | 1,37 | 1,107 | 1,233 | 0,976 | 1,094 | 1,17 | 1,05 | 0,05 | 7,3E-01 |
| P28028 | Serine/threonine-protein kinase B-raf OS=Mus musculus OX=10090 | 11 | 7 | 8 | 5 | 804 | 88,7 | 7,65 | 14,92 | 7 | 1,292 | 1,281 | 0,966 | 1,188 | 1,013 | 1,29 | 1,152 | 1,138 | 1,03 | 0,03 | 7,4E-01 |
| P57080 | Ubiquitin carboxyl-terminal hydrolase 25 OS=Mus musculus OX=10 | 1 | 1 | 1 | 1 | 1055 | 121,3 | 5,31 | 2,23 | 1 | 0,679 | 1,404 | 0,942 | 0,856 | 0,792 | 0,77 | 1,09 | 0,984 | 1,07 | 0,07 | 7,4E-01 |
| Q8VDJ3 | Vigilin OS=Mus musculus OX=10090 GN=Hdlbp PE=1 SV=1 | 5 | 6 | 6 | 6 | 1268 | 141,7 | 6,87 | 10,86 | 6 | 1,097 | 1,129 | 0,936 | 0,875 | 0,918 | 1,046 | 1 | 0,978 | 1,02 | 0,02 | 7,4E-01 |
| Q8C4X2 | Casein kinase I isoform gamma-3 OS=Mus musculus OX=10090 GN= | 11 | 4 | 4 | 3 | 424 | 48,9 | 9,11 | 6,88 | 4 | 0,84 | 1,113 | 0,925 | 0,951 | 0,93 | 0,964 | 0,949 | 1,08 | 0,98 | -0,02 | 7,4E-01 |
| Q99MR8 | Methylcrotonoyl-CoA carboxylase subunit alpha, mitochondrial OS | 16 | 9 | 9 | 9 | 717 | 79,3 | 7,83 | 20,9 | 9 | 1,279 | 1,176 | 0,981 | 0,941 | 0,953 | 1,234 | 0,946 | 1,095 | 1,04 | 0,04 | 7,4E-01 |
| Q7TN79 | A-kinase anchor protein 7 isoform gamma OS=Mus musculus OX=1 | 7 | 2 | 2 | 2 | 314 | 35,5 | 7,87 | 3 | 2 | 0,965 | 0,753 | 0,863 | 0,915 | 0,726 | 0,912 | 0,835 | 0,931 | 1,03 | 0,03 | 7,4E-01 |
| O35286 | Pre-mRNA-splicing factor ATP-dependent RNA helicase DHX15 OS= | 5 | 4 | 4 | 4 | 795 | 90,9 | 7,46 | 5,34 | 4 | 0,91 | 0,99 | 0,976 | 0,993 | 1,107 | 0,861 | 1,027 | 0,953 | 0,98 | -0,02 | 7,4E-01 |
| Q8JZR0 | Long-chain-fatty-acid--CoA ligase 5 OS=Mus musculus OX=10090 G | 12 | 8 | 9 | 7 | 683 | 76,2 | 7,09 | 18,53 | 8 | 1,265 | 1,214 | 1,251 | 1,123 | 0,971 | 1,401 | 1,107 | 1,237 | 1,03 | 0,03 | 7,4E-01 |
| P83741 | Serine/threonine-protein kinase WNK1 OS=Mus musculus OX=1009 | 3 | 6 | 6 | 6 | 2377 | 250,8 | 6,43 | 10,12 | 6 | 0,99 | 1,186 | 0,999 | 1,023 | 0,926 | 1,022 | 1,064 | 1,102 | 1,02 | 0,02 | 7,4E-01 |
| Q5EBJ4 | Ermin OS=Mus musculus OX=10090 GN=Ermn PE=1 SV=1 | 19 | 3 | 3 | 3 | 281 | 32,1 | 4,59 | 3,38 | 3 | 1,398 | 0,654 | 0,76 | 1,148 | 1,004 | 0,496 | 1,126 | 1,02 | 1,09 | 0,09 | 7,4E-01 |
| Q6PFD5 | Disks large-associated protein 3 OS=Mus musculus OX=10090 GN=I | 10 | 8 | 10 | 7 | 977 | 105,8 | 8,76 | 9,12 | 8 | 0,987 | 0,953 | 0,889 | 0,968 | 0,998 | 0,849 | 0,964 | 0,932 | 1,01 | 0,01 | 7,4E-01 |
| Q9QZQ8 | Core histone macro-H2A.1 OS=Mus musculus OX=10090 GN=H2afy | 14 | 4 | 4 | 4 | 372 | 39,7 | 9,8 | 8,65 | 4 | 1,345 | 1,396 | 1,118 | 0,848 | 1,074 | 1,853 | 0,923 | 1,195 | 0,93 | -0,07 | 7,4E-01 |
| Q9CX84 | Regulator of G-protein signaling 19 OS=Mus musculus OX=10090 G | 5 | 1 | 1 | 1 | 216 | 24,7 | 6 | 2,16 | 1 | 0,814 | 1,055 | 1,036 | 1,463 | 1,174 | 0,764 | 0,99 | 1,203 | 1,06 | 0,06 | 7,4E-01 |
| Q9QZ73 | DCN1-like protein 1 OS=Mus musculus OX=10090 GN=Dcun1d1 PE= | 12 | 3 | 3 | 3 | 259 | 30,1 | 5,34 | 6,7 | 3 | 1,062 | 0,963 | 1,066 | 1,004 | 1,014 | 1,174 | 1,009 | 0,97 | 0,98 | -0,02 | 7,4E-01 |

| | | | | | | | | | | | | | | | | | | | | | |
|---|---|---|---|---|---|---|---|---|---|---|---|---|---|---|---|---|---|---|---|---|---|
| Q9Z239 | Phospholemman OS=Mus musculus OX=10090 GN=Fxyd1 PE=1 SV= | 13 | 1 | 1 | 1 | 92 | 10,3 | 8,54 | 4,16 | 1 | 1,063 | 0,988 | 0,833 | 0,889 | 0,932 | 0,984 | 0,804 | 0,962 | 1,02 | 0,02 | 7,4E-01 |
| Q6A078 | Centrosomal protein of 290 kDa OS=Mus musculus OX=10090 GN= | 0 | 1 | 1 | 1 | 2472 | 288,9 | 6,1 | 1,73 | 1 | 0,944 | 0,605 | 0,915 | 0,98 | 0,899 | 0,755 | 0,755 | 0,901 | 1,04 | 0,04 | 7,4E-01 |
| Q6PCN3 | Tau-tubulin kinase 1 OS=Mus musculus OX=10090 GN=Ttbk1 PE=1 | 4 | 4 | 4 | 4 | 1308 | 141,5 | 5,62 | 0 | 4 | 1,513 | 1,035 | 1,083 | 1,301 | 1,243 | 1,34 | 1,203 | 1,305 | 0,97 | -0,03 | 7,4E-01 |
| Q3TPX4 | Exocyst complex component 5 OS=Mus musculus OX=10090 GN=Ex | 12 | 8 | 10 | 8 | 708 | 81,7 | 6,71 | 13,24 | 8 | 1,121 | 1,046 | 1,137 | 1,107 | 0,928 | 1,235 | 1,092 | 1,064 | 1,02 | 0,02 | 7,4E-01 |
| P48758 | Carbonyl reductase [NADPH] 1 OS=Mus musculus OX=10090 GN=Cl | 30 | 7 | 12 | 6 | 277 | 30,6 | 8,31 | 31,59 | 7 | 0,985 | 1,058 | 0,991 | 1,041 | 0,989 | 1,021 | 1,127 | 0,99 | 0,99 | -0,01 | 7,4E-01 |
| P58854 | Gamma-tubulin complex component 3 OS=Mus musculus OX=1009 | 5 | 4 | 4 | 4 | 905 | 103,4 | 8,32 | 10,37 | 4 | 1,206 | 0,95 | 1,108 | 0,887 | 0,884 | 1,508 | 0,967 | 1,013 | 0,95 | -0,05 | 7,4E-01 |
| Q9QYX7 | Protein piccolo OS=Mus musculus OX=10090 GN=Pclo PE=1 SV=4 | 17 | 70 | 88 | 66 | 5068 | 550,5 | 6,51 | 178,14 | 70 | 1,092 | 1,037 | 1,022 | 1,004 | 0,936 | 1,072 | 1,046 | 1,051 | 1,01 | 0,01 | 7,4E-01 |
| Q9JKK1 | Syntaxin-6 OS=Mus musculus OX=10090 GN=Stx6 PE=1 SV=1 | 7 | 2 | 2 | 2 | 255 | 29 | 4,92 | 5,05 | 2 | 0,744 | 1,129 | 1,051 | 0,848 | 1,009 | 0,627 | 0,874 | 1,077 | 1,05 | 0,05 | 7,4E-01 |
| O88602 | Voltage-dependent calcium channel gamma-2 subunit OS=Mus mu | 9 | 4 | 5 | 2 | 323 | 35,9 | 8,98 | 13,38 | 4 | 0,963 | 1,062 | 0,865 | 0,978 | 1,03 | 1,235 | 0,89 | 0,846 | 0,97 | -0,03 | 7,4E-01 |
| Q9CY64 | Biliverdin reductase A OS=Mus musculus OX=10090 GN=Blvra PE=1 | 13 | 4 | 4 | 4 | 295 | 33,5 | 7,02 | 11,44 | 4 | 0,89 | 1,058 | 1,028 | 0,958 | 0,957 | 0,99 | 0,994 | 0,938 | 1,01 | 0,01 | 7,4E-01 |
| Q8C052 | Microtubule-associated protein 1S OS=Mus musculus OX=10090 GI | 2 | 2 | 4 | 2 | 973 | 102,9 | 7,02 | 6,33 | 2 | 0,933 | 0,856 | 1,132 | 1,025 | 0,983 | 1,111 | 0,933 | 1,016 | 0,98 | -0,02 | 7,4E-01 |
| Q99KP6 | Pre-mRNA-processing factor 19 OS=Mus musculus OX=10090 GN=F | 6 | 3 | 3 | 3 | 504 | 55,2 | 6,61 | 4,66 | 3 | 1,245 | 1,244 | 1,102 | 1,021 | 1,123 | 1,283 | 1,009 | 1,087 | 1,02 | 0,02 | 7,4E-01 |
| Q8K1Z0 | Ubiquinone biosynthesis protein COQ9, mitochondrial OS=Mus mu | 6 | 2 | 4 | 2 | 313 | 35,1 | 5,92 | 7,29 | 2 | 0,947 | 1,159 | 0,859 | 0,736 | 0,924 | 0,934 | 0,806 | 0,908 | 1,04 | 0,04 | 7,4E-01 |
| P12961 | Neuroendocrine protein 7B2 OS=Mus musculus OX=10090 GN=Scg | 6 | 2 | 2 | 2 | 212 | 23,9 | 5,81 | 3,98 | 2 | 0,562 | 1,339 | 1,055 | 1,672 | 1,677 | 0,81 | 1,594 | 0,986 | 0,91 | -0,09 | 7,4E-01 |
| Q9D6S7 | Ribosome-recycling factor, mitochondrial OS=Mus musculus OX=10 | 11 | 2 | 2 | 2 | 262 | 29 | 9,85 | 6,7 | 2 | 1,123 | 1,226 | 1,024 | 1,263 | 0,955 | 1,235 | 1,291 | 1,021 | 1,03 | 0,03 | 7,4E-01 |
| P97930 | Thymidylate kinase OS=Mus musculus OX=10090 GN=Dtymk PE=1 | 9 | 2 | 2 | 2 | 212 | 23,9 | 7,9 | 4,89 | 2 | 0,851 | 1,034 | 0,902 | 0,932 | 1,234 | 0,684 | 1,084 | 0,889 | 0,96 | -0,04 | 7,4E-01 |
| Q9DAU1 | Protein canopy homolog 3 OS=Mus musculus OX=10090 GN=Cnpy3 | 15 | 3 | 3 | 3 | 276 | 30,5 | 5,62 | 4,64 | 3 | 0,834 | 0,898 | 0,864 | 0,843 | 1,164 | 0,842 | 1,08 | 0,544 | 0,95 | -0,05 | 7,4E-01 |
| Q9CZN8 | Glutamyl-tRNA(Gln) amidotransferase subunit A, mitochondrial OS | 8 | 4 | 5 | 4 | 525 | 56,7 | 5,95 | 8,99 | 4 | 1,303 | 1,233 | 1,313 | 0,99 | 1,143 | 1,375 | 1,028 | 1,15 | 1,03 | 0,03 | 7,4E-01 |
| P35979 | 60S ribosomal protein L12 OS=Mus musculus OX=10090 GN=Rpl12 | 39 | 5 | 6 | 5 | 165 | 17,8 | 9,42 | 17,79 | 5 | 0,877 | 0,776 | 1,03 | 0,898 | 0,924 | 1,024 | 0,791 | 0,939 | 0,97 | -0,03 | 7,4E-01 |
| Q9JIS5 | Synaptic vesicle glycoprotein 2A OS=Mus musculus OX=10090 GN= | 16 | 12 | 18 | 11 | 742 | 82,6 | 5,57 | 39,9 | 12 | 0,761 | 0,888 | 0,913 | 1,044 | 0,906 | 0,814 | 0,994 | 0,99 | 0,97 | -0,03 | 7,4E-01 |
| Q8R1B5 | Complexin-3 OS=Mus musculus OX=10090 GN=Cplx3 PE=1 SV=1 | 41 | 8 | 8 | 8 | 158 | 17,6 | 4,89 | 20,51 | 8 | 0,948 | 0,936 | 1,074 | 1,076 | 0,982 | 0,948 | 0,985 | 1,058 | 1,02 | 0,02 | 7,5E-01 |
| Q9CYL5 | Golgi-associated plant pathogenesis-related protein 1 OS=Mus mus | 6 | 1 | 1 | 1 | 154 | 17,1 | 9,51 | 2,33 | 1 | 0,982 | 0,96 | 1,118 | 0,951 | 1,13 | 0,472 | 0,881 | 1,282 | 1,07 | 0,07 | 7,5E-01 |
| Q9D6G9 | CKLF-like MARVEL transmembrane domain-containing protein 5 OS | 12 | 1 | 1 | 1 | 156 | 17,4 | 4,84 | 0 | 1 | 0,862 | 1,726 | 1,052 | 0,615 | 0,77 | 1,346 | 0,965 | 0,806 | 1,09 | 0,09 | 7,5E-01 |
| Q8C170 | Unconventional myosin-IXa OS=Mus musculus OX=10090 GN=Myo | 1 | 2 | 2 | 2 | 2542 | 291,9 | 8,97 | 2,2 | 2 | 3,183 | 2,372 | 1,812 | 1,579 | 2,118 | 2,361 | 2,493 | 1,388 | 1,07 | 0,07 | 7,5E-01 |
| Q5XJV6 | Serine/threonine-protein kinase LMTK3 OS=Mus musculus OX=100 | 1 | 1 | 1 | 1 | 1424 | 150,8 | 4,88 | 0 | 1 | 1,845 | 1,279 | 1,627 | 0,832 | 0,833 | 1,609 | 1,155 | 1,595 | 1,08 | 0,08 | 7,5E-01 |
| Q8CF89 | TGF-beta-activated kinase 1 and MAP3K7-binding protein 1 OS=Mu | 2 | 1 | 1 | 1 | 502 | 54,6 | 5,52 | 0 | 1 | 2,139 | 1,844 | 1,905 | 1,451 | 0,841 | 3,085 | 1,626 | 2,476 | 0,91 | -0,09 | 7,5E-01 |
| Q9CPQ8 | ATP synthase subunit g, mitochondrial OS=Mus musculus OX=1009 | 9 | 1 | 1 | 1 | 103 | 11,4 | 9,74 | 2,01 | 1 | 1,183 | 1,106 | 1,507 | 1,359 | 1,465 | 1,042 | 1,172 | 1,712 | 0,96 | -0,04 | 7,5E-01 |
| Q6P9S0 | Protein MTSS 2 OS=Mus musculus OX=10090 GN=Mtss2 PE=1 SV=1 | 3 | 2 | 2 | 2 | 715 | 76,8 | 6,99 | 4,43 | 2 | 0,709 | 0,736 | 1,005 | 1,02 | 0,967 | 0,695 | 0,872 | 0,799 | 1,04 | 0,04 | 7,5E-01 |
| B9EJ86 | Oxysterol-binding protein-related protein 8 OS=Mus musculus OX= | 6 | 4 | 5 | 4 | 889 | 101,2 | 6,96 | 9,59 | 4 | 1,076 | 0,798 | 1,04 | 0,959 | 0,99 | 0,967 | 1,062 | 0,944 | 0,98 | -0,02 | 7,5E-01 |
| O55222 | Integrin-linked protein kinase OS=Mus musculus OX=10090 GN=Ilk | 2 | 1 | 1 | 1 | 452 | 51,3 | 8,07 | 1,88 | 1 | 1,137 | 1,231 | 0,959 | 1,116 | 1,053 | 1,308 | 1,244 | 0,969 | 0,97 | -0,03 | 7,5E-01 |
| Q8R081 | Heterogeneous nuclear ribonucleoprotein L OS=Mus musculus OX= | 8 | 4 | 5 | 4 | 586 | 63,9 | 8,1 | 10,64 | 4 | 1,491 | 1,035 | 1,224 | 0,88 | 1,05 | 1,623 | 1,096 | 1,114 | 0,95 | -0,05 | 7,5E-01 |
| Q7M6Y3 | Phosphatidylinositol-binding clathrin assembly protein OS=Mus mu | 10 | 6 | 10 | 6 | 660 | 71,5 | 7,91 | 24,04 | 6 | 1,094 | 0,972 | 1,045 | 1,086 | 0,998 | 1,04 | 1,119 | 1,092 | 0,99 | -0,01 | 7,5E-01 |
| Q8VDL4 | ADP-dependent glucokinase OS=Mus musculus OX=10090 GN=Adp | 6 | 2 | 2 | 2 | 496 | 53,9 | 5,62 | 3,35 | 2 | 1,338 | 0,992 | 0,903 | 1,029 | 0,755 | 1,157 | 0,959 | 1,204 | 1,05 | 0,05 | 7,5E-01 |
| Q9JMG1 | Endothelial differentiation-related factor 1 OS=Mus musculus OX=1 | 10 | 1 | 1 | 1 | 148 | 16,4 | 9,99 | 0 | 1 | 2,115 | 1,64 | 1,389 | 0,531 | 1,157 | 1,997 | 0,774 | 1,185 | 1,11 | 0,11 | 7,5E-01 |
| Q3UTZ3 | Microtubule-associated protein 11 OS=Mus musculus OX=10090 GI | 3 | 2 | 2 | 2 | 580 | 62,7 | 8,68 | 2,54 | 2 | 0,989 | 0,837 | 1,25 | 1,239 | 1,095 | 0,759 | 1,357 | 0,883 | 1,05 | 0,05 | 7,5E-01 |
| P47791 | Glutathione reductase, mitochondrial OS=Mus musculus OX=10090 | 9 | 3 | 3 | 3 | 500 | 53,6 | 7,99 | 3,5 | 3 | 0,698 | 1,037 | 0,63 | 0,742 | 0,782 | 0,968 | 0,753 | 0,743 | 0,96 | -0,04 | 7,5E-01 |
| Q80YV4 | Pantothenate kinase 4 OS=Mus musculus OX=10090 GN=Pank4 PE= | 6 | 4 | 5 | 4 | 820 | 91,5 | 6,71 | 2,55 | 4 | 0,713 | 0,96 | 0,597 | 1,009 | 0,721 | 0,815 | 0,777 | 1,15 | 0,95 | -0,05 | 7,5E-01 |
| Q91WK0 | Leucine-rich repeat flightless-interacting protein 2 OS=Mus muscul | 5 | 2 | 2 | 2 | 415 | 47,1 | 5,68 | 2,24 | 2 | 0,887 | 1,199 | 0,979 | 1,055 | 1,175 | 1,424 | 0,885 | 0,838 | 0,95 | -0,05 | 7,5E-01 |
| Q6ZWN5 | 40S ribosomal protein S9 OS=Mus musculus OX=10090 GN=Rps9 PI | 10 | 2 | 2 | 2 | 194 | 22,6 | 10,65 | 5,06 | 2 | 1,572 | 1,056 | 1,197 | 1,007 | 1,028 | 1,223 | 1,113 | 1,282 | 1,04 | 0,04 | 7,5E-01 |
| Q9Z2A5 | Arginyl-tRNA--protein transferase 1 OS=Mus musculus OX=10090 G | 2 | 1 | 1 | 1 | 516 | 59,1 | 8,19 | 2,58 | 1 | 0,589 | 0,949 | 0,655 | 0,798 | 0,811 | 0,821 | 0,743 | 0,727 | 0,96 | -0,04 | 7,5E-01 |
| Q3UVX5 | Metabotropic glutamate receptor 5 OS=Mus musculus OX=10090 G | 10 | 12 | 15 | 11 | 1203 | 131,8 | 7,84 | 13,45 | 12 | 1,042 | 1,02 | 0,898 | 1,046 | 0,992 | 1,018 | 1,121 | 0,943 | 0,98 | -0,02 | 7,5E-01 |
| Q7TMB8 | Cytoplasmic FMR1-interacting protein 1 OS=Mus musculus OX=100 | 6 | 7 | 10 | 3 | 1253 | 145,1 | 6,9 | 18,77 | 7 | 1,031 | 0,96 | 1,01 | 0,967 | 0,96 | 1,071 | 0,986 | 0,99 | 0,99 | -0,01 | 7,5E-01 |
| Q9WV91 | Prostaglandin F2 receptor negative regulator OS=Mus musculus OX | 7 | 6 | 7 | 6 | 879 | 98,7 | 6,61 | 12,91 | 6 | 1,068 | 1,043 | 0,777 | 0,954 | 0,953 | 0,99 | 0,958 | 1,031 | 0,98 | -0,02 | 7,5E-01 |
| P60764 | Ras-related C3 botulinum toxin substrate 3 OS=Mus musculus OX= | 27 | 5 | 10 | 1 | 192 | 21,4 | 8,15 | 21,21 | 5 | 1,375 | 1,302 | 1,089 | 0,981 | 0,992 | 1,381 | 1,084 | 1,127 | 1,04 | 0,04 | 7,5E-01 |
| Q9CXY6 | Interleukin enhancer-binding factor 2 OS=Mus musculus OX=10090 | 7 | 2 | 2 | 2 | 390 | 43 | 5,26 | 0 | 2 | 1,055 | 1,119 | 0,966 | 0,804 | 0,981 | 1,251 | 0,946 | 0,903 | 0,97 | -0,03 | 7,5E-01 |
| P48678 | Prelamin-A/C OS=Mus musculus OX=10090 GN=Lmna PE=1 SV=2 | 20 | 13 | 16 | 11 | 665 | 74,2 | 6,98 | 25,66 | 13 | 1,198 | 1,525 | 1,029 | 0,86 | 0,996 | 1,324 | 0,761 | 1,277 | 1,06 | 0,06 | 7,5E-01 |
| O88704 | Potassium/sodium hyperpolarization-activated cyclic nucleotide-ga | 13 | 9 | 9 | 8 | 910 | 102,4 | 8,37 | 15,74 | 9 | 0,904 | 0,973 | 0,983 | 1 | 1,023 | 1,018 | 0,921 | 0,942 | 0,99 | -0,01 | 7,5E-01 |
| Q8C115 | Pleckstrin homology domain-containing family H member 2 OS=Mu | 0 | 1 | 1 | 1 | 1491 | 167,6 | 7,47 | 2,55 | 1 | 1,009 | 0,781 | 1,007 | 1,037 | 0,876 | 0,87 | 1,064 | 0,926 | 1,03 | 0,03 | 7,5E-01 |
| P14069 | Protein S100-A6 OS=Mus musculus OX=10090 GN=S100a6 PE=1 SV | 17 | 2 | 2 | 2 | 89 | 10 | 5,48 | 5,03 | 2 | 1,435 | 1,096 | 1,166 | 0,858 | 1,045 | 1,224 | 0,932 | 1,176 | 1,04 | 0,04 | 7,5E-01 |
| Q9CZP5 | Mitochondrial chaperone BCS1 OS=Mus musculus OX=10090 GN=B | 8 | 3 | 3 | 3 | 418 | 47,4 | 7,93 | 10,98 | 3 | 0,946 | 1,078 | 0,953 | 1,084 | 0,992 | 1,011 | 1,005 | 1,115 | 0,98 | -0,02 | 7,5E-01 |
| Q8BYM8 | Probable cysteine--tRNA ligase, mitochondrial OS=Mus musculus O | 7 | 4 | 4 | 4 | 551 | 61,2 | 8,18 | 4,88 | 4 | 0,886 | 1,295 | 1,064 | 1,193 | 1,064 | 0,928 | 1,096 | 1,212 | 1,03 | 0,03 | 7,6E-01 |
| O70166 | Stathmin-3 OS=Mus musculus OX=10090 GN=Stmn3 PE=1 SV=1 | 11 | 2 | 2 | 2 | 180 | 20,9 | 7,49 | 5,63 | 2 | 0,867 | 1,085 | 1,058 | 1,314 | 0,962 | 0,909 | 1,215 | 1,089 | 1,04 | 0,04 | 7,6E-01 |
| P98192 | Dihydroxyacetone phosphate acyltransferase OS=Mus musculus OX | 2 | 1 | 1 | 1 | 678 | 76,8 | 8,13 | 0 | 1 | 1,061 | 0,829 | 1,611 | 1,233 | 1,262 | 1,03 | 1,258 | 1,423 | 0,95 | -0,05 | 7,6E-01 |
| P15379 | CD44 antigen OS=Mus musculus OX=10090 GN=Cd44 PE=1 SV=3 | 1 | 1 | 1 | 1 | 778 | 85,6 | 4,96 | 2,06 | 1 | 1,853 | 1,273 | 1,807 | 0,764 | 1,362 | 1,248 | 1,676 | 1,035 | 1,07 | 0,07 | 7,6E-01 |
| Q9D2R8 | 28S ribosomal protein S33, mitochondrial OS=Mus musculus OX=1( | 11 | 1 | 1 | 1 | 106 | 12,5 | 10,26 | 3,63 | 1 | 1,414 | 0,969 | 0,991 | 1,194 | 1,131 | 1,249 | 1,025 | 1,01 | 1,03 | 0,03 | 7,6E-01 |
| Q9D517 | 1-acyl-sn-glycerol-3-phosphate acyltransferase gamma OS=Mus mu | 10 | 4 | 4 | 4 | 376 | 43,3 | 8,51 | 9,46 | 4 | 0,899 | 0,849 | 0,988 | 1,024 | 1,083 | 0,919 | 0,942 | 0,892 | 0,98 | -0,02 | 7,6E-01 |
| Q8K221 | Arfaptin-2 OS=Mus musculus OX=10090 GN=Arfip2 PE=1 SV=2 | 14 | 5 | 8 | 5 | 341 | 37,7 | 5,87 | 16,13 | 5 | 0,933 | 0,942 | 0,999 | 1,022 | 0,926 | 0,991 | 1,063 | 0,963 | 0,99 | -0,01 | 7,6E-01 |
| Q7TQI3 | Ubiquitin thioesterase OTUB1 OS=Mus musculus OX=10090 GN=Ot | 34 | 8 | 11 | 8 | 271 | 31,3 | 4,94 | 30,23 | 8 | 1,072 | 1,083 | 1,055 | 1,07 | 0,972 | 1,19 | 1,014 | 1,042 | 1,01 | 0,01 | 7,6E-01 |
| Q91VR2 | ATP synthase subunit gamma, mitochondrial OS=Mus musculus OX | 32 | 12 | 34 | 12 | 298 | 32,9 | 9,01 | 73,19 | 12 | 0,989 | 0,974 | 1,016 | 0,935 | 1,025 | 0,905 | 1,034 | 0,994 | 0,99 | -0,01 | 7,6E-01 |
| Q9Z2I2 | Peptidyl-prolyl cis-trans isomerase FKBP1B OS=Mus musculus OX=1 | 13 | 1 | 5 | 1 | 108 | 11,8 | 8,47 | 10,35 | 1 | 1,173 | 0,917 | 0,968 | 0,927 | 0,835 | 1,232 | 0,949 | 1,106 | 0,97 | -0,03 | 7,6E-01 |
| Q6PNC0 | DmX-like protein 1 OS=Mus musculus OX=10090 GN=Dmxl1 PE=1 S | 4 | 9 | 10 | 8 | 3013 | 335,8 | 6,42 | 14,35 | 9 | 1,045 | 1,176 | 1,235 | 1,293 | 1,129 | 1,146 | 1,227 | 1,173 | 1,02 | 0,02 | 7,6E-01 |
| Q9EQ80 | NIF3-like protein 1 OS=Mus musculus OX=10090 GN=Nif3l1 PE=1 S\ | 14 | 4 | 7 | 4 | 376 | 41,7 | 6,76 | 14,04 | 4 | 1,155 | 1,192 | 1,102 | 1,095 | 1,014 | 1,256 | 1,084 | 1,118 | 1,02 | 0,02 | 7,6E-01 |

| | | | | | | | | | | | | | | | | | | | | | |
|---|---|---|---|---|---|---|---|---|---|---|---|---|---|---|---|---|---|---|---|---|---|
| Q8R2Q4 | Ribosome-releasing factor 2, mitochondrial OS=Mus musculus OX= | 1 | 1 | 1 | 1 | 779 | 86,1 | 6,43 | 1,66 | 1 | 1,125 | 1,279 | 0,751 | 0,845 | 0,918 | 1,164 | 0,781 | 0,949 | 1,05 | 0,05 | 7,6E-01 |
| Q62172 | RalA-binding protein 1 OS=Mus musculus OX=10090 GN=Ralbp1 PE | 8 | 5 | 6 | 5 | 648 | 75 | 5,92 | 6,9 | 5 | 0,718 | 1,031 | 0,93 | 1 | 0,965 | 0,752 | 0,857 | 0,99 | 1,03 | 0,03 | 7,6E-01 |
| Q9D1E6 | Tubulin-folding cofactor B OS=Mus musculus OX=10090 GN=Tbcb P | 21 | 5 | 5 | 5 | 244 | 27,4 | 5,24 | 9,25 | 5 | 0,872 | 0,944 | 0,909 | 1,008 | 1,019 | 0,843 | 0,998 | 0,936 | 0,98 | -0,02 | 7,6E-01 |
| Q3ULJ0 | Glycerol-3-phosphate dehydrogenase 1-like protein OS=Mus muscu | 21 | 8 | 10 | 8 | 351 | 38,2 | 6,77 | 28,1 | 8 | 0,886 | 0,952 | 1,024 | 1,213 | 1,055 | 0,909 | 1,032 | 0,979 | 1,03 | 0,03 | 7,6E-01 |
| Q8BGR6 | ADP-ribosylation factor-like protein 15 OS=Mus musculus OX=1009 | 25 | 4 | 4 | 4 | 204 | 22,9 | 5,29 | 10,13 | 4 | 1,306 | 1,391 | 1,171 | 1,178 | 1,097 | 1,455 | 1,134 | 1,236 | 1,03 | 0,03 | 7,6E-01 |
| Q8BWR2 | PITH domain-containing protein 1 OS=Mus musculus OX=10090 GN | 6 | 1 | 2 | 1 | 211 | 24,2 | 5,74 | 2,33 | 1 | 0,928 | 0,706 | 1,204 | 0,748 | 0,467 | 1,264 | 1,04 | 1,08 | 0,93 | -0,07 | 7,6E-01 |
| P60603 | Reactive oxygen species modulator 1 OS=Mus musculus OX=10090 | 10 | 1 | 1 | 1 | 79 | 8,2 | 9,33 | 2,21 | 1 | 1,071 | 1,168 | 1,044 | 0,985 | 0,97 | 1,184 | 0,99 | 1,045 | 1,02 | 0,02 | 7,6E-01 |
| Q3UVL4 | Vacuolar protein sorting-associated protein 51 homolog OS=Mus m | 10 | 7 | 7 | 7 | 782 | 86,1 | 6,24 | 20,22 | 7 | 1,103 | 0,97 | 1,253 | 1,126 | 1,11 | 1,15 | 1,189 | 1,083 | 0,98 | -0,02 | 7,6E-01 |
| Q9D7X8 | Gamma-glutamylcyclotransferase OS=Mus musculus OX=10090 GN | 12 | 2 | 3 | 2 | 188 | 21,2 | 5,67 | 8,37 | 2 | 0,749 | 1 | 1,078 | 1,258 | 0,952 | 1,077 | 1,136 | 1,064 | 0,97 | -0,03 | 7,6E-01 |
| A2AQ25 | Sickle tail protein OS=Mus musculus OX=10090 GN=Skt PE=1 SV=1 | 11 | 20 | 21 | 20 | 1946 | 212,9 | 7,77 | 40,59 | 20 | 1,005 | 1,166 | 1,035 | 1,087 | 1,051 | 1,11 | 1,021 | 1,06 | 1,01 | 0,01 | 7,6E-01 |
| Q921H8 | 3-ketoacyl-CoA thiolase A, peroxisomal OS=Mus musculus OX=1009 | 8 | 2 | 2 | 2 | 424 | 43,9 | 8,44 | 7,05 | 2 | 0,908 | 0,901 | 1,017 | 1,183 | 1,126 | 0,672 | 1,109 | 0,943 | 1,04 | 0,04 | 7,6E-01 |
| Q5SV85-2 | Isoform 2 of Synergin gamma OS=Mus musculus OX=10090 GN=Syr | 8 | 6 | 6 | 6 | 1138 | 122 | 5 | 6,56 | 6 | 1,057 | 0,944 | 1,147 | 1,128 | 0,992 | 0,984 | 1,026 | 1,189 | 1,02 | 0,02 | 7,6E-01 |
| Q61543 | Golgi apparatus protein 1 OS=Mus musculus OX=10090 GN=Glg1 Pl | 6 | 5 | 6 | 5 | 1175 | 133,6 | 6,84 | 8,39 | 5 | 0,829 | 1,033 | 0,953 | 1,077 | 0,979 | 1,056 | 0,991 | 0,942 | 0,98 | -0,02 | 7,6E-01 |
| Q80TL1 | Adenylate cyclase type 2 OS=Mus musculus OX=10090 GN=Adcy2 F | 4 | 4 | 5 | 4 | 1090 | 123,2 | 8,31 | 8,22 | 4 | 1,316 | 1,121 | 0,85 | 1,003 | 1,147 | 1,109 | 0,988 | 1,183 | 0,97 | -0,03 | 7,6E-01 |
| Q80TR1 | Adhesion G protein-coupled receptor L1 OS=Mus musculus OX=100 | 5 | 5 | 6 | 5 | 1466 | 161,6 | 6,48 | 11,73 | 5 | 0,994 | 0,939 | 0,963 | 0,95 | 0,977 | 1,139 | 0,843 | 0,966 | 0,98 | -0,02 | 7,6E-01 |
| Q6P1B3 | PILR alpha-associated neural protein OS=Mus musculus OX=10090 | 3 | 1 | 2 | 1 | 278 | 29,7 | 8,72 | 3,93 | 1 | 0,979 | 0,955 | 0,862 | 0,938 | 0,984 | 0,807 | 1,156 | 0,888 | 0,97 | -0,03 | 7,6E-01 |
| P60882 | Multiple epidermal growth factor-like domains protein 8 OS=Mus n | 1 | 2 | 2 | 2 | 2789 | 297,3 | 6,8 | 2,2 | 2 | 1,018 | 1,087 | 0,81 | 0,833 | 1,037 | 1,063 | 0,835 | 0,923 | 0,97 | -0,03 | 7,6E-01 |
| P46471 | 26S proteasome regulatory subunit 7 OS=Mus musculus OX=10090 | 28 | 12 | 13 | 12 | 433 | 48,6 | 5,95 | 19,63 | 12 | 1,281 | 1,137 | 1,101 | 0,967 | 0,968 | 1,231 | 1,005 | 1,167 | 1,03 | 0,03 | 7,6E-01 |
| O35643 | AP-1 complex subunit beta-1 OS=Mus musculus OX=10090 GN=Ap1 | 23 | 22 | 39 | 8 | 943 | 103,9 | 5,17 | 91,57 | 22 | 1,099 | 1,023 | 1,015 | 0,987 | 0,959 | 1,083 | 1,035 | 1,097 | 0,99 | -0,01 | 7,6E-01 |
| P35279 | Ras-related protein Rab-6A OS=Mus musculus OX=10090 GN=Rab6 | 29 | 8 | 11 | 4 | 208 | 23,6 | 5,54 | 20,42 | 8 | 1,277 | 1,194 | 1,073 | 0,875 | 0,978 | 1,329 | 1,215 | 1,047 | 0,97 | -0,03 | 7,6E-01 |
| Q99LI2 | Chloride channel CLIC-like protein 1 OS=Mus musculus OX=10090 G | 6 | 2 | 2 | 2 | 539 | 60,6 | 5,68 | 4,93 | 2 | 1,098 | 1,016 | 1,098 | 1,136 | 1,036 | 1,272 | 0,978 | 0,966 | 1,02 | 0,02 | 7,6E-01 |
| Q61701 | ELAV-like protein 4 OS=Mus musculus OX=10090 GN=Elavl4 PE=1 S | 8 | 3 | 3 | 1 | 385 | 42,3 | 9,38 | 1,97 | 3 | 2,029 | 1,044 | 1,658 | 1,62 | 1,262 | 1,967 | 2,123 | 1,373 | 0,94 | -0,06 | 7,6E-01 |
| Q8BHL3 | TBC1 domain family member 10B OS=Mus musculus OX=10090 GN | 6 | 4 | 5 | 4 | 798 | 87,2 | 8,9 | 15,48 | 4 | 0,774 | 0,957 | 0,997 | 1,065 | 1,017 | 0,938 | 0,992 | 0,929 | 0,98 | -0,02 | 7,6E-01 |
| Q61120 | SHC-transforming protein 3 OS=Mus musculus OX=10090 GN=Shc3 | 9 | 4 | 4 | 4 | 474 | 52,1 | 7,3 | 0 | 4 | 1,357 | 1,303 | 1,446 | 1,379 | 1,129 | 1,659 | 1,436 | 1,403 | 0,97 | -0,03 | 7,6E-01 |
| Q9CQN1 | Heat shock protein 75 kDa, mitochondrial OS=Mus musculus OX=1( | 14 | 9 | 11 | 8 | 706 | 80,2 | 6,68 | 24,65 | 9 | 1,039 | 1,168 | 1,099 | 0,976 | 0,93 | 1,117 | 1,067 | 1,094 | 1,02 | 0,02 | 7,6E-01 |
| P07356 | Annexin A2 OS=Mus musculus OX=10090 GN=Anxa2 PE=1 SV=2 | 13 | 4 | 4 | 4 | 339 | 38,7 | 7,69 | 2,9 | 4 | 1,406 | 1,267 | 0,842 | 0,816 | 0,983 | 1,431 | 1,062 | 1,081 | 0,95 | -0,05 | 7,6E-01 |
| Q8C6M1 | Ubiquitin carboxyl-terminal hydrolase 20 OS=Mus musculus OX=10 | 3 | 2 | 2 | 2 | 916 | 102,1 | 6,16 | 2,08 | 2 | 1,161 | 1,015 | 1,358 | 1,435 | 0,741 | 1,595 | 1,415 | 1,489 | 0,95 | -0,05 | 7,6E-01 |
| P35285 | Ras-related protein Rab-22A OS=Mus musculus OX=10090 GN=Rab | 19 | 3 | 3 | 1 | 194 | 21,8 | 8,15 | 7,67 | 3 | 1,637 | 0,697 | 1,278 | 1,328 | 1,338 | 1,06 | 1,373 | 1,437 | 0,95 | -0,05 | 7,6E-01 |
| O55126 | Protein NipSnap homolog 2 OS=Mus musculus OX=10090 GN=Nipsi | 21 | 6 | 11 | 6 | 281 | 32,9 | 9,26 | 26,37 | 6 | 1,01 | 0,96 | 0,979 | 1,046 | 1,03 | 0,879 | 0,994 | 1,04 | 1,01 | 0,01 | 7,6E-01 |
| P15864 | Histone H1.2 OS=Mus musculus OX=10090 GN=Hist1h1c PE=1 SV=2 | 28 | 9 | 14 | 4 | 212 | 21,3 | 11 | 35,15 | 9 | 1,186 | 1,437 | 0,897 | 0,644 | 1,011 | 1,522 | 0,694 | 1,245 | 0,93 | -0,07 | 7,6E-01 |
| Q9JLZ3 | Methylglutaconyl-CoA hydratase, mitochondrial OS=Mus musculus | 33 | 8 | 13 | 8 | 314 | 33,4 | 9,51 | 34,42 | 8 | 1,111 | 1,067 | 1,129 | 0,974 | 0,951 | 1,173 | 1,003 | 1,08 | 1,02 | 0,02 | 7,6E-01 |
| Q3TFQ1 | SPRY domain-containing protein 7 OS=Mus musculus OX=10090 GN | 5 | 1 | 1 | 1 | 196 | 21,7 | 6,7 | 2,54 | 1 | 1,181 | 1,197 | 1,244 | 1,184 | 0,938 | 1,533 | 0,889 | 1,258 | 1,04 | 0,04 | 7,7E-01 |
| Q61411 | GTPase HRas OS=Mus musculus OX=10090 GN=Hras PE=1 SV=2 | 60 | 9 | 17 | 3 | 189 | 21,3 | 5,31 | 38,98 | 9 | 1,005 | 1,112 | 0,953 | 0,935 | 0,857 | 1,018 | 1,127 | 0,914 | 1,02 | 0,02 | 7,7E-01 |
| Q80X50 | Ubiquitin-associated protein 2-like OS=Mus musculus OX=10090 GI | 1 | 1 | 1 | 1 | 1107 | 116,7 | 7,11 | 3,2 | 1 | 1,19 | 1,072 | 1,201 | 0,982 | 0,894 | 1,365 | 1,024 | 1,021 | 1,03 | 0,03 | 7,7E-01 |
| P35585 | AP-1 complex subunit mu-1 OS=Mus musculus OX=10090 GN=Ap1r | 9 | 3 | 3 | 3 | 423 | 48,5 | 7,3 | 4,42 | 3 | 0,798 | 1,033 | 0,852 | 1,085 | 0,861 | 1,015 | 0,965 | 1,024 | 0,97 | -0,03 | 7,7E-01 |
| Q8R570 | Synaptosomal-associated protein 47 OS=Mus musculus OX=10090 | 14 | 5 | 5 | 5 | 413 | 46,5 | 5,76 | 12,97 | 5 | 1,097 | 1,199 | 1,239 | 1,052 | 1,015 | 1,253 | 1,12 | 1,294 | 0,98 | -0,02 | 7,7E-01 |
| P63087-2 | Isoform 2 of Serine/threonine-protein phosphatase PP1-gamma ca | 21 | 6 | 10 | 2 | 337 | 38,5 | 6,13 | 24,11 | 6 | 1,512 | 1,028 | 0,544 | 1,07 | 1,267 | 0,962 | 0,766 | 0,883 | 1,07 | 0,07 | 7,7E-01 |
| Q922H4 | Mannose-1-phosphate guanyltransferase alpha OS=Mus musculus | 10 | 3 | 3 | 3 | 420 | 46,2 | 7,62 | 4,88 | 3 | 0,784 | 0,769 | 0,891 | 0,755 | 0,812 | 0,719 | 0,898 | 0,829 | 0,98 | -0,02 | 7,7E-01 |
| P22005 | Proenkephalin-A OS=Mus musculus OX=10090 GN=Penk PE=1 SV=2 | 16 | 4 | 4 | 4 | 268 | 31 | 5,68 | 4,19 | 4 | 1,095 | 0,777 | 1,098 | 1,009 | 0,743 | 1,081 | 1,527 | 0,859 | 0,95 | -0,05 | 7,7E-01 |
| P61982 | 14-3-3 protein gamma OS=Mus musculus OX=10090 GN=Ywhag PE | 57 | 17 | 35 | 10 | 247 | 28,3 | 4,89 | 82,96 | 17 | 0,774 | 0,876 | 0,93 | 1,015 | 0,975 | 0,778 | 0,947 | 0,98 | 0,98 | -0,02 | 7,7E-01 |
| Q9JHG2 | Calcipressin-2 OS=Mus musculus OX=10090 GN=Rcan2 PE=1 SV=1 | 8 | 1 | 1 | 1 | 197 | 22 | 6,68 | 2,66 | 1 | 1,392 | 1,447 | 1,153 | 1,122 | 0,895 | 1,513 | 1,123 | 1,387 | 1,04 | 0,04 | 7,7E-01 |
| P11499 | Heat shock protein HSP 90-beta OS=Mus musculus OX=10090 GN=I | 46 | 37 | 70 | 20 | 724 | 83,2 | 5,03 | 178,28 | 37 | 0,948 | 0,956 | 1,037 | 1,016 | 0,996 | 0,92 | 1,017 | 0,987 | 1,01 | 0,01 | 7,7E-01 |
| A2AWP8 | Rho guanine nucleotide exchange factor 10-like protein OS=Mus m | 2 | 2 | 2 | 2 | 1280 | 139,9 | 5,82 | 0 | 2 | 1,719 | 1,594 | 2,61 | 1,664 | 1,3 | 2,676 | 1,659 | 2,442 | 0,94 | -0,06 | 7,7E-01 |
| Q68FM6 | Protein phosphatase 1 regulatory subunit 29 OS=Mus musculus OX | 4 | 3 | 3 | 2 | 823 | 90 | 7,52 | 4,63 | 3 | 1,337 | 2,049 | 2,12 | 1,458 | 1,71 | 1,417 | 1,916 | 1,647 | 1,04 | 0,04 | 7,7E-01 |
| P02802 | Metallothionein-1 OS=Mus musculus OX=10090 GN=Mt1 PE=1 SV= | 20 | 1 | 1 | 1 | 61 | 6 | 7,96 | 2,8 | 1 | 1,308 | 0,73 | 1,391 | 1,165 | 1,268 | 0,93 | 1,307 | 1,299 | 0,96 | -0,04 | 7,7E-01 |
| Q9WV34 | MAGUK p55 subfamily member 2 OS=Mus musculus OX=10090 GN | 36 | 17 | 22 | 16 | 552 | 61,5 | 6,44 | 37,95 | 17 | 0,947 | 0,964 | 1,017 | 0,995 | 0,979 | 0,927 | 0,962 | 1,024 | 1,01 | 0,01 | 7,7E-01 |
| Q8BMJ2 | Leucine--tRNA ligase, cytoplasmic OS=Mus musculus OX=10090 GN | 5 | 4 | 4 | 4 | 1178 | 134,1 | 7,05 | 3,02 | 4 | 1,203 | 1,274 | 1,099 | 0,996 | 0,916 | 1,299 | 1,09 | 1,146 | 1,03 | 0,03 | 7,7E-01 |
| Q9JKF7 | 39S ribosomal protein L39, mitochondrial OS=Mus musculus OX=10 | 10 | 3 | 3 | 3 | 336 | 38,5 | 7,94 | 2,43 | 3 | 0,824 | 1,163 | 1,043 | 0,959 | 0,996 | 0,874 | 0,887 | 1,121 | 1,03 | 0,03 | 7,7E-01 |
| P63042 | Stathmin-4 OS=Mus musculus OX=10090 GN=Stmn4 PE=2 SV=1 | 5 | 1 | 1 | 1 | 189 | 22,1 | 5,99 | 2,27 | 1 | 1,065 | 1,035 | 0,969 | 0,884 | 1,144 | 0,892 | 0,887 | 1,129 | 0,98 | -0,02 | 7,7E-01 |
| Q99NF2 | NMDA receptor synaptonuclear signaling and neuronal migration fa | 8 | 3 | 3 | 3 | 532 | 60,3 | 8,98 | 5,2 | 3 | 1,441 | 1,298 | 1,74 | 0,879 | 1,013 | 1,74 | 1,197 | 1,116 | 1,06 | 0,06 | 7,7E-01 |
| Q8CCT4 | Transcription elongation factor A protein-like 5 OS=Mus musculus ( | 7 | 1 | 1 | 1 | 200 | 22 | 6,2 | 2,77 | 1 | 1,083 | 1,291 | 1,207 | 1,083 | 1,255 | 0,846 | 1,116 | 1,661 | 0,96 | -0,04 | 7,7E-01 |
| Q99K01 | Pyridoxal-dependent decarboxylase domain-containing protein 1 O | 3 | 2 | 2 | 2 | 787 | 87,3 | 5,48 | 5,33 | 2 | 1,199 | 0,913 | 1,077 | 1,209 | 1,143 | 1,135 | 1,022 | 1,004 | 1,02 | 0,02 | 7,7E-01 |
| Q9CPQ3 | Mitochondrial import receptor subunit TOM22 homolog OS=Mus m | 8 | 1 | 1 | 1 | 142 | 15,5 | 4,34 | 2,46 | 1 | 1,025 | 0,767 | 0,673 | 0,677 | 0,63 | 0,746 | 0,915 | 0,729 | 1,04 | 0,04 | 7,7E-01 |
| Q9DCP2 | Sodium-coupled neutral amino acid transporter 3 OS=Mus musculu | 8 | 3 | 3 | 3 | 505 | 55,6 | 7,09 | 7,99 | 3 | 1,329 | 1,136 | 1,061 | 0,921 | 0,972 | 1,017 | 1,137 | 1,201 | 1,03 | 0,03 | 7,7E-01 |
| O08599 | Syntaxin-binding protein 1 OS=Mus musculus OX=10090 GN=Stxbp | 54 | 35 | 84 | 2 | 594 | 67,5 | 6,96 | 191,02 | 35 | 0,938 | 0,954 | 0,621 | 0,708 | 0,864 | 0,547 | 0,924 | 0,745 | 1,05 | 0,05 | 7,7E-01 |
| Q14CH0 | Protein FAM171B OS=Mus musculus OX=10090 GN=Fam171b PE=1 | 9 | 5 | 6 | 5 | 825 | 92 | 8,37 | 10,21 | 5 | 1,352 | 1,188 | 1,109 | 1,096 | 0,967 | 1,38 | 1,156 | 1,118 | 1,03 | 0,03 | 7,7E-01 |
| O08788 | Dynactin subunit 1 OS=Mus musculus OX=10090 GN=Dctn1 PE=1 S' | 30 | 34 | 42 | 34 | 1281 | 141,6 | 5,9 | 99,82 | 34 | 1,06 | 1,028 | 1,008 | 1,009 | 0,96 | 1,07 | 0,99 | 1,051 | 1,01 | 0,01 | 7,8E-01 |
| Q9R0N4 | Synaptotagmin-10 OS=Mus musculus OX=10090 GN=Syt10 PE=1 SV | 2 | 1 | 1 | 1 | 523 | 59 | 7,77 | 0 | 1 | 1,648 | 0,608 | 1,233 | 0,758 | 0,957 | 0,949 | 1,46 | 1,198 | 0,93 | -0,07 | 7,8E-01 |
| Q9Z2U0 | Proteasome subunit alpha type-7 OS=Mus musculus OX=10090 GN | 23 | 5 | 5 | 5 | 248 | 27,8 | 8,46 | 15,26 | 5 | 1,138 | 1,035 | 1,043 | 0,961 | 0,939 | 1,149 | 0,943 | 1,071 | 1,02 | 0,02 | 7,8E-01 |
| Q9R0P3 | S-formylglutathione hydrolase OS=Mus musculus OX=10090 GN=Es | 23 | 5 | 8 | 5 | 282 | 31,3 | 7,12 | 13,16 | 5 | 1,016 | 1,052 | 1,155 | 1,129 | 1,155 | 1,147 | 1,013 | 1,092 | 0,99 | -0,01 | 7,8E-01 |
| Q9D273 | Corrinoid adenosyltransferase OS=Mus musculus OX=10090 GN=M | 19 | 4 | 4 | 4 | 237 | 26,3 | 9,2 | 7,06 | 4 | 1,106 | 1,045 | 1,163 | 1,052 | 1,034 | 1,165 | 1,049 | 1,07 | 1,01 | 0,01 | 7,8E-01 |
| P14873 | Microtubule-associated protein 1B OS=Mus musculus OX=10090 GI | 22 | 47 | 60 | 46 | 2464 | 270,1 | 4,83 | 128,88 | 47 | 0,909 | 0,908 | 1,019 | 1,03 | 0,969 | 0,929 | 1,037 | 0,979 | 0,99 | -0,01 | 7,8E-01 |

| | | | | | | | | | | | | | | | | | | | | | |
|---|---|---|---|---|---|---|---|---|---|---|---|---|---|---|---|---|---|---|---|---|---|
| Q810U3 | Neurofascin OS=Mus musculus OX=10090 GN=Nfasc PE=1 SV=1 | 11 | 12 | 16 | 12 | 1240 | 137,9 | 6,19 | 34,15 | 12 | 1,174 | 0,938 | 1,118 | 0,861 | 1,044 | 1,008 | 0,976 | 0,973 | 1,02 | 0,02 | 7,8E-01 |
| Q9R1T2 | SUMO-activating enzyme subunit 1 OS=Mus musculus OX=10090 G | 3 | 1 | 1 | 1 | 350 | 38,6 | 5,36 | 0 | 1 | 1,603 | 0,651 | 1,282 | 1,362 | 1,527 | 0,618 | 1,285 | 1,136 | 1,07 | 0,07 | 7,8E-01 |
| Q99JR1 | Sideroflexin-1 OS=Mus musculus OX=10090 GN=Sfxn1 PE=1 SV=3 | 22 | 6 | 9 | 6 | 322 | 35,6 | 9,23 | 16,69 | 6 | 1,185 | 1,208 | 1,071 | 0,912 | 0,975 | 1,194 | 1,039 | 1,071 | 1,02 | 0,02 | 7,8E-01 |
| Q01065 | Calcium/calmodulin-dependent 3',5'-cyclic nucleotide phosphodies | 8 | 3 | 3 | 3 | 535 | 61,2 | 5,72 | 5,5 | 3 | 1,18 | 1,281 | 0,945 | 0,836 | 0,972 | 1,049 | 1,059 | 1,038 | 1,03 | 0,03 | 7,8E-01 |
| Q8BJF9 | Charged multivesicular body protein 2b OS=Mus musculus OX=100 | 11 | 3 | 3 | 2 | 213 | 23,9 | 8,78 | 4,61 | 3 | 0,762 | 0,87 | 0,915 | 0,901 | 0,803 | 0,823 | 0,856 | 1,043 | 0,98 | -0,02 | 7,8E-01 |
| Q9QXY6 | EH domain-containing protein 3 OS=Mus musculus OX=10090 GN= | 23 | 13 | 15 | 10 | 535 | 60,8 | 6,46 | 36,98 | 13 | 1,054 | 0,986 | 1,009 | 1,018 | 0,886 | 1,108 | 1,066 | 1,068 | 0,99 | -0,01 | 7,8E-01 |
| O55100 | Synaptogyrin-1 OS=Mus musculus OX=10090 GN=Syngr1 PE=1 SV= | 10 | 2 | 3 | 2 | 234 | 25,6 | 4,65 | 6,36 | 2 | 1,223 | 1,076 | 1,046 | 0,905 | 0,873 | 1,245 | 0,988 | 1,024 | 1,03 | 0,03 | 7,8E-01 |
| Q8BY87 | Ubiquitin carboxyl-terminal hydrolase 47 OS=Mus musculus OX=10 | 3 | 4 | 4 | 4 | 1376 | 157,4 | 5,11 | 4,87 | 4 | 0,944 | 1,003 | 1,01 | 1,129 | 1,024 | 0,85 | 1,11 | 1,023 | 1,02 | 0,02 | 7,8E-01 |
| Q9CPW3 | 39S ribosomal protein L54, mitochondrial OS=Mus musculus OX=10 | 13 | 2 | 2 | 2 | 135 | 15,4 | 10,04 | 5,3 | 2 | 0,894 | 0,93 | 1,195 | 0,928 | 0,972 | 0,979 | 1,019 | 1,063 | 0,98 | -0,02 | 7,8E-01 |
| P23116 | Eukaryotic translation initiation factor 3 subunit A OS=Mus muscul | 6 | 6 | 6 | 6 | 1344 | 161,8 | 6,77 | 11,8 | 6 | 1,203 | 1,026 | 1,243 | 1,045 | 0,985 | 1,226 | 1,024 | 1,187 | 1,02 | 0,02 | 7,8E-01 |
| Q8BJZ4 | 28S ribosomal protein S35, mitochondrial OS=Mus musculus OX=10 | 8 | 2 | 2 | 2 | 320 | 36 | 8,59 | 1,85 | 2 | 1,132 | 1,056 | 1,072 | 1,09 | 0,96 | 1,287 | 1,216 | 0,985 | 0,98 | -0,02 | 7,8E-01 |
| Q8C0T5 | Signal-induced proliferation-associated 1-like protein 1 OS=Mus mu | 10 | 15 | 16 | 14 | 1782 | 196,9 | 8,13 | 27,91 | 15 | 0,99 | 0,979 | 1,041 | 0,971 | 0,896 | 0,988 | 1,044 | 1,109 | 0,99 | -0,01 | 7,8E-01 |
| Q3UHD6 | Sorting nexin-27 OS=Mus musculus OX=10090 GN=Snx27 PE=1 SV= | 10 | 4 | 5 | 4 | 539 | 61 | 6,4 | 13,24 | 4 | 1,02 | 1,131 | 0,857 | 1,077 | 1,01 | 1,022 | 1,022 | 1,105 | 0,98 | -0,02 | 7,8E-01 |
| Q9D7X3 | Dual specificity protein phosphatase 3 OS=Mus musculus OX=1009 | 31 | 4 | 8 | 4 | 185 | 20,5 | 6,54 | 26,58 | 4 | 0,932 | 1,12 | 1,079 | 1,058 | 1,019 | 0,982 | 1,184 | 1,074 | 0,98 | -0,02 | 7,8E-01 |
| Q80TS3 | Adhesion G protein-coupled receptor L3 OS=Mus musculus OX=100 | 3 | 5 | 8 | 5 | 1537 | 171 | 6,71 | 17,13 | 5 | 1,106 | 0,998 | 1,088 | 1,098 | 1,038 | 1,077 | 1,094 | 1,048 | 1,01 | 0,01 | 7,8E-01 |
| P63141 | Potassium voltage-gated channel subfamily A member 2 OS=Mus m | 11 | 5 | 8 | 3 | 499 | 56,7 | 4,86 | 20,13 | 5 | 1,073 | 1,208 | 1,382 | 1,154 | 1,093 | 1,394 | 1,078 | 1,379 | 0,97 | -0,03 | 7,8E-01 |
| Q9D2N9 | Vacuolar protein sorting-associated protein 33A OS=Mus musculus | 5 | 3 | 4 | 3 | 598 | 67,5 | 7,08 | 2,3 | 3 | 1,467 | 1,222 | 0,769 | 1,078 | 1,26 | 0,953 | 1,131 | 1,004 | 1,04 | 0,04 | 7,8E-01 |
| Q8N7N5 | DDB1- and CUL4-associated factor 8 OS=Mus musculus OX=10090 | 4 | 2 | 2 | 2 | 591 | 66 | 5,87 | 2,26 | 2 | 0,493 | 1,423 | 1,142 | 1,714 | 1,312 | 0,931 | 1,324 | 1,543 | 0,93 | -0,07 | 7,8E-01 |
| P70255-4 | Isoform 4 of Nuclear factor 1 C-type OS=Mus musculus OX=10090 | 6 | 2 | 2 | 2 | 397 | 44,1 | 7,97 | 3,93 | 2 | 0,65 | 0,945 | 0,888 | 0,636 | 0,654 | 0,696 | 0,839 | 1,074 | 0,96 | -0,04 | 7,8E-01 |
| Q3UZV7 | UPF0577 protein KIAA1324-like homolog OS=Mus musculus OX=10 | 2 | 2 | 2 | 2 | 1028 | 113,6 | 6,14 | 2,23 | 2 | 0,594 | 0,44 | 0,713 | 1,188 | 0,922 | 0,886 | 0,644 | 0,687 | 0,94 | -0,06 | 7,8E-01 |
| Q8BHG1 | Nardilysin OS=Mus musculus OX=10090 GN=Nrdc PE=1 SV=1 | 3 | 3 | 4 | 3 | 1161 | 132,8 | 4,87 | 9,77 | 3 | 1,015 | 1,007 | 1,283 | 1,138 | 0,956 | 1,123 | 1,148 | 1,125 | 1,02 | 0,02 | 7,8E-01 |
| Q62132 | Receptor-type tyrosine-protein phosphatase R OS=Mus musculus C | 6 | 3 | 4 | 3 | 656 | 74 | 7,81 | 2,52 | 3 | 1,111 | 1,243 | 1,438 | 1,023 | 0,916 | 1,253 | 1,402 | 1,414 | 0,97 | -0,03 | 7,8E-01 |
| Q9CR61 | NADH dehydrogenase [ubiquinone] 1 beta subcomplex subunit 7 O | 43 | 5 | 8 | 5 | 137 | 16,3 | 8,18 | 14,31 | 5 | 1,287 | 1,152 | 1,061 | 0,977 | 0,929 | 1,268 | 1,08 | 1,089 | 1,03 | 0,03 | 7,8E-01 |
| P68040 | Receptor of activated protein C kinase 1 OS=Mus musculus OX=100 | 10 | 3 | 4 | 3 | 317 | 35,1 | 7,69 | 7,32 | 3 | 1,135 | 0,909 | 0,893 | 0,915 | 0,714 | 1,106 | 0,891 | 1,022 | 1,03 | 0,03 | 7,8E-01 |
| Q9Z0G0 | PDZ domain-containing protein GIPC1 OS=Mus musculus OX=1009 | 12 | 3 | 4 | 3 | 333 | 36,1 | 5,91 | 5,55 | 3 | 1,277 | 1,184 | 1,192 | 1,003 | 1,204 | 1,298 | 0,917 | 1,122 | 1,03 | 0,03 | 7,8E-01 |
| P60843 | Eukaryotic initiation factor 4A-I OS=Mus musculus OX=10090 GN=E | 21 | 8 | 10 | 6 | 406 | 46,1 | 5,48 | 28,22 | 8 | 1,119 | 1,115 | 1,061 | 1,1 | 1,087 | 1,251 | 1,053 | 1,06 | 0,99 | -0,01 | 7,8E-01 |
| Q91WD5 | NADH dehydrogenase [ubiquinone] iron-sulfur protein 2, mitochon | 30 | 12 | 24 | 12 | 463 | 52,6 | 6,99 | 49,5 | 12 | 1,076 | 1,099 | 0,981 | 1,021 | 1,002 | 1,137 | 1,015 | 1,07 | 0,99 | -0,01 | 7,8E-01 |
| P05064 | Fructose-bisphosphate aldolase A OS=Mus musculus OX=10090 GN | 53 | 27 | 70 | 21 | 364 | 39,3 | 8,09 | 193,47 | 27 | 0,95 | 0,96 | 0,974 | 1,037 | 0,922 | 0,963 | 1,029 | 0,973 | 1,01 | 0,01 | 7,8E-01 |
| Q8K0X8 | Fasciculation and elongation protein zeta-1 OS=Mus musculus OX= | 5 | 2 | 3 | 2 | 392 | 45,2 | 4,36 | 4,34 | 2 | 1,113 | 0,858 | 0,953 | 1,03 | 1,005 | 1,065 | 1,011 | 0,942 | 0,98 | -0,02 | 7,8E-01 |
| Q9QYR9 | Acyl-coenzyme A thioesterase 2, mitochondrial OS=Mus musculus | 7 | 3 | 4 | 1 | 453 | 49,6 | 7,36 | 8,03 | 3 | 0,707 | 1,054 | 1,187 | 1,025 | 0,941 | 0,725 | 1,081 | 1,075 | 1,04 | 0,04 | 7,8E-01 |
| Q6PAK3 | Protein arginine N-methyltransferase 8 OS=Mus musculus OX=1009 | 10 | 4 | 6 | 2 | 394 | 45,2 | 6,93 | 19,9 | 4 | 1,303 | 1,041 | 1,121 | 0,973 | 1,103 | 1,092 | 1,164 | 0,987 | 1,02 | 0,02 | 7,8E-01 |
| Q9CPW2 | Ferredoxin-2, mitochondrial OS=Mus musculus OX=10090 GN=Fdx2 | 13 | 2 | 2 | 2 | 174 | 18,8 | 6,04 | 2,51 | 2 | 0,72 | 1,016 | 1,004 | 0,961 | 0,784 | 0,823 | 1,021 | 0,97 | 1,03 | 0,03 | 7,8E-01 |
| Q9QYG0 | Protein NDRG2 OS=Mus musculus OX=10090 GN=Ndrg2 PE=1 SV=1 | 29 | 8 | 19 | 7 | 371 | 40,8 | 5,4 | 36,93 | 8 | 0,975 | 0,92 | 1,117 | 1,093 | 1,164 | 1,011 | 0,973 | 1,029 | 0,98 | -0,02 | 7,8E-01 |
| Q03059 | Choline O-acetyltransferase OS=Mus musculus OX=10090 GN=Chat | 2 | 1 | 1 | 1 | 641 | 71,8 | 7,74 | 2,55 | 1 | 1,861 | 1,435 | 1,25 | 1,232 | 1,269 | 1,336 | 1,885 | 1,021 | 1,05 | 0,05 | 7,8E-01 |
| Q6NXK8 | Acid-sensing ion channel 1 OS=Mus musculus OX=10090 GN=Asic1 | 2 | 1 | 1 | 1 | 526 | 59,6 | 5,58 | 1,98 | 1 | 0,964 | 0,729 | 1,52 | 1,176 | 1,003 | 1,085 | 1,407 | 0,629 | 1,06 | 0,06 | 7,8E-01 |
| Q78JE5 | F-box only protein 22 OS=Mus musculus OX=10090 GN=Fbxo22 PE | 5 | 2 | 3 | 2 | 402 | 44,2 | 7,61 | 4,03 | 2 | 0,706 | 0,65 | 0,759 | 1,142 | 0,802 | 0,736 | 0,818 | 0,772 | 1,04 | 0,04 | 7,8E-01 |
| Q8CJG1 | Protein argonaute-1 OS=Mus musculus OX=10090 GN=Ago1 PE=1 S | 5 | 3 | 3 | 2 | 857 | 97,2 | 9,16 | 0 | 3 | 1,07 | 1,26 | 0,962 | 1,331 | 1,279 | 1,317 | 0,829 | 1,37 | 0,96 | -0,04 | 7,9E-01 |
| Q91WG2 | Rab GTPase-binding effector protein 2 OS=Mus musculus OX=1009 | 4 | 2 | 2 | 2 | 554 | 62,1 | 4,89 | 5,15 | 2 | 0,945 | 0,612 | 0,744 | 0,836 | 0,728 | 1,083 | 0,653 | 0,513 | 1,05 | 0,05 | 7,9E-01 |
| Q6ZWV7 | 60S ribosomal protein L35 OS=Mus musculus OX=10090 GN=Rpl35 | 8 | 1 | 1 | 1 | 123 | 14,5 | 11,05 | 3,12 | 1 | 1,063 | 1,643 | 1,54 | 1,608 | 1,645 | 1,621 | 1,497 | 1,273 | 0,97 | -0,03 | 7,9E-01 |
| P61205 | ADP-ribosylation factor 3 OS=Mus musculus OX=10090 GN=Arf3 PE | 46 | 8 | 12 | 5 | 181 | 20,6 | 7,43 | 37,01 | 8 | 0,985 | 1,104 | 0,928 | 0,972 | 0,848 | 1,103 | 0,977 | 0,988 | 1,02 | 0,02 | 7,9E-01 |
| Q9D0F3 | Protein ERGIC-53 OS=Mus musculus OX=10090 GN=Lman1 PE=1 SV | 5 | 2 | 3 | 2 | 517 | 57,8 | 6,34 | 5,66 | 2 | 1,05 | 1,312 | 1,007 | 0,936 | 1,05 | 1,013 | 0,76 | 1,322 | 1,04 | 0,04 | 7,9E-01 |
| Q3TDQ1 | Dolichyl-diphosphooligosaccharide--protein glycosyltransferase sub | 3 | 2 | 2 | 2 | 823 | 93,2 | 8,95 | 5,8 | 2 | 1,084 | 1,373 | 1,381 | 1,018 | 0,975 | 1,269 | 1,013 | 1,436 | 1,03 | 0,03 | 7,9E-01 |
| Q8R1V4 | Transmembrane emp24 domain-containing protein 4 OS=Mus mus | 5 | 1 | 1 | 1 | 227 | 26 | 8,18 | 2,54 | 1 | 1,153 | 1,289 | 1,723 | 1,157 | 1,533 | 1,604 | 1,023 | 1,372 | 0,96 | -0,04 | 7,9E-01 |
| P32037 | Solute carrier family 2, facilitated glucose transporter member 3 O | 10 | 4 | 5 | 4 | 493 | 53,4 | 4,98 | 12,03 | 4 | 0,726 | 1,034 | 0,884 | 1,102 | 0,98 | 0,743 | 1,005 | 0,903 | 1,03 | 0,03 | 7,9E-01 |
| Q6IRU5-2 | Isoform 2 of Clathrin light chain B OS=Mus musculus OX=10090 GN | 31 | 10 | 14 | 10 | 211 | 23,2 | 4,68 | 34,86 | 10 | 0,869 | 0,873 | 1,033 | 1,026 | 1,04 | 0,739 | 0,944 | 0,988 | 1,02 | 0,02 | 7,9E-01 |
| Q5SRY7 | F-box/WD repeat-containing protein 11 OS=Mus musculus OX=100 | 2 | 1 | 1 | 1 | 542 | 62 | 7,12 | 2,19 | 1 | 1,063 | 1,235 | 1,453 | 1,171 | 0,934 | 1,232 | 1,384 | 1,232 | 1,03 | 0,03 | 7,9E-01 |
| Q60692 | Proteasome subunit beta type-6 OS=Mus musculus OX=10090 GN= | 20 | 5 | 5 | 5 | 238 | 25,4 | 5,11 | 14,48 | 5 | 1,27 | 1,129 | 1,015 | 0,891 | 1,045 | 1,101 | 1,004 | 1,062 | 1,02 | 0,02 | 7,9E-01 |
| Q925H0 | Acid-sensing ion channel 2 OS=Mus musculus OX=10090 GN=Asic2 | 5 | 3 | 3 | 3 | 512 | 57,7 | 5,19 | 4,72 | 3 | 0,973 | 0,845 | 1,05 | 0,94 | 1,142 | 0,744 | 0,968 | 1,061 | 0,97 | -0,03 | 7,9E-01 |
| Q05186 | Reticulocalbin-1 OS=Mus musculus OX=10090 GN=Rcn1 PE=1 SV=1 | 12 | 3 | 4 | 3 | 325 | 38,1 | 4,84 | 9,91 | 3 | 1,149 | 0,824 | 1,396 | 1,306 | 1,032 | 0,978 | 1,393 | 1,098 | 1,04 | 0,04 | 7,9E-01 |
| P17183 | Gamma-enolase OS=Mus musculus OX=10090 GN=Eno2 PE=1 SV=2 | 20 | 9 | 22 | 7 | 434 | 47,3 | 5,11 | 58,76 | 9 | 1,038 | 1,059 | 0,968 | 0,949 | 0,95 | 1,068 | 1,022 | 1,014 | 0,99 | -0,01 | 7,9E-01 |
| Q6P5E8 | Diacylglycerol kinase theta OS=Mus musculus OX=10090 GN=Dgkq | 7 | 5 | 5 | 5 | 934 | 102,2 | 7,52 | 8,21 | 5 | 1,168 | 0,879 | 1,169 | 1,091 | 0,973 | 1,247 | 1,066 | 1,12 | 0,98 | -0,02 | 7,9E-01 |
| Q8BM13 | Noelin-2 OS=Mus musculus OX=10090 GN=Olfm2 PE=1 SV=2 | 5 | 2 | 2 | 2 | 448 | 50,7 | 8,51 | 4,5 | 2 | 0,946 | 0,807 | 0,811 | 1,256 | 1,152 | 0,993 | 0,821 | 0,993 | 0,96 | -0,04 | 7,9E-01 |
| Q9CRC8 | Leucine-rich repeat-containing protein 40 OS=Mus musculus OX=10 | 5 | 3 | 3 | 3 | 602 | 68 | 6,89 | 7,72 | 3 | 1,244 | 1,414 | 1,197 | 1,169 | 1,096 | 1,411 | 1,228 | 1,194 | 1,02 | 0,02 | 7,9E-01 |
| Q8K377 | Leucine-rich repeat transmembrane neuronal protein 1 OS=Mus m | 4 | 2 | 2 | 2 | 522 | 58,7 | 7,11 | 2,29 | 2 | 1,355 | 1,827 | 1,532 | 1,391 | 1,632 | 1,446 | 1,565 | 1,321 | 1,02 | 0,02 | 7,9E-01 |
| Q9DBH5 | Vesicular integral-membrane protein VIP36 OS=Mus musculus OX= | 3 | 1 | 1 | 1 | 358 | 40,4 | 6,95 | 2,37 | 1 | 1,221 | 0,909 | 1,046 | 1,054 | 1,185 | 1,165 | 0,81 | 0,948 | 1,03 | 0,03 | 7,9E-01 |
| Q9CWP6 | Motile sperm domain-containing protein 2 OS=Mus musculus OX=1 | 2 | 1 | 1 | 1 | 518 | 59,8 | 5,67 | 0 | 1 | 0,549 | 0,864 | 1,005 | 1,083 | 0,893 | 0,832 | 0,795 | 0,849 | 1,04 | 0,04 | 7,9E-01 |
| A2ARS0 | Ankyrin repeat domain-containing protein 63 OS=Mus musculus O | 2 | 1 | 1 | 1 | 390 | 41 | 10,83 | 2,77 | 1 | 1,326 | 0,926 | 1,002 | 1,002 | 1,083 | 1,064 | 1,401 | 0,865 | 0,96 | -0,04 | 7,9E-01 |
| P97434 | Myosin phosphatase Rho-interacting protein OS=Mus musculus OX | 3 | 3 | 3 | 3 | 1024 | 116,3 | 6,21 | 7,67 | 3 | 1,034 | 0,721 | 0,978 | 0,924 | 1,042 | 0,703 | 1,146 | 0,896 | 0,97 | -0,03 | 7,9E-01 |
| Q9D404 | 3-oxoacyl-[acyl-carrier-protein] synthase, mitochondrial OS=Mus m | 14 | 4 | 5 | 4 | 459 | 48,6 | 7,06 | 5,56 | 4 | 1,578 | 1,294 | 1,4 | 1,011 | 1,067 | 1,545 | 1,344 | 1,151 | 1,03 | 0,03 | 7,9E-01 |
| P62862 | 40S ribosomal protein S30 OS=Mus musculus OX=10090 GN=Fau P | 19 | 1 | 1 | 1 | 59 | 6,6 | 12,15 | 2,11 | 1 | 1,191 | 0,749 | 0,881 | 0,876 | 1,157 | 0,686 | 0,991 | 0,699 | 1,05 | 0,05 | 7,9E-01 |
| Q921M3 | Splicing factor 3B subunit 3 OS=Mus musculus OX=10090 GN=Sf3b | 2 | 2 | 2 | 2 | 1217 | 135,5 | 5,26 | 1,7 | 2 | 1,25 | 1,251 | 1,151 | 0,883 | 0,977 | 1,319 | 0,986 | 1,123 | 1,03 | 0,03 | 7,9E-01 |
| Q99J83 | Autophagy protein 5 OS=Mus musculus OX=10090 GN=Atg5 PE=1 S | 5 | 1 | 2 | 1 | 275 | 32,4 | 5,91 | 2,1 | 1 | 1,724 | 0,901 | 1,069 | 0,831 | 1,029 | 0,874 | 0,922 | 1,436 | 1,06 | 0,06 | 7,9E-01 |

| | | | | | | | | | | | | | | | | | | | | | |
|---|---|---|---|---|---|---|---|---|---|---|---|---|---|---|---|---|---|---|---|---|---|
| Q62245 | Son of sevenless homolog 1 OS=Mus musculus OX=10090 GN=Sos1 | 2 | 3 | 3 | 3 | 1319 | 150,8 | 6,89 | 4,47 | 3 | 0,92 | 0,962 | 0,831 | 0,91 | 0,89 | 1,003 | 0,922 | 0,854 | 0,99 | -0,01 | 7,9E-01 |
| Q9D0R8 | Protein LSM12 homolog OS=Mus musculus OX=10090 GN=Lsm12 P | 7 | 1 | 1 | 1 | 195 | 21,7 | 7,74 | 3,84 | 1 | 1,282 | 0,97 | 1,08 | 1,038 | 0,906 | 1,24 | 1,023 | 1,095 | 1,02 | 0,02 | 7,9E-01 |
| Q9JIX8 | Apoptotic chromatin condensation inducer in the nucleus OS=Mus | 1 | 1 | 1 | 1 | 1338 | 150,6 | 5,91 | 0 | 1 | 1,454 | 1,087 | 1,227 | 0,938 | 1,287 | 1,128 | 1,451 | 1 | 0,97 | -0,03 | 7,9E-01 |
| Q69ZW3 | EH domain-binding protein 1 OS=Mus musculus OX=10090 GN=Ehb | 4 | 4 | 4 | 4 | 1231 | 139 | 5,38 | 9,75 | 4 | 1,049 | 1,196 | 0,98 | 1,052 | 0,877 | 0,971 | 1,125 | 1,209 | 1,02 | 0,02 | 8,0E-01 |
| P62069 | Ubiquitin carboxyl-terminal hydrolase 46 OS=Mus musculus OX=10 | 2 | 1 | 1 | 1 | 366 | 42,4 | 6,83 | 0 | 1 | 1,112 | 0,701 | 1,09 | 0,731 | 1,104 | 0,588 | 0,801 | 1,361 | 0,94 | -0,06 | 8,0E-01 |
| A2ALU4 | Protein Shroom2 OS=Mus musculus OX=10090 GN=Shroom2 PE=1 | 2 | 3 | 3 | 3 | 1481 | 164,6 | 6,7 | 2,42 | 3 | 0,982 | 0,918 | 0,935 | 0,878 | 0,94 | 0,812 | 1,061 | 0,96 | 0,98 | -0,02 | 8,0E-01 |
| Q8K4Z3 | NAD(P)H-hydrate epimerase OS=Mus musculus OX=10090 GN=Nax | 20 | 5 | 6 | 5 | 282 | 31 | 7,69 | 17,01 | 5 | 0,95 | 0,915 | 1,164 | 0,891 | 0,979 | 0,898 | 0,963 | 1,173 | 0,98 | -0,02 | 8,0E-01 |
| Q8C8N2 | Protein SCAI OS=Mus musculus OX=10090 GN=Scai PE=1 SV=2 | 12 | 6 | 8 | 6 | 606 | 70,2 | 8,6 | 4,68 | 6 | 1,132 | 1,338 | 1,266 | 1,179 | 1,094 | 1,458 | 1,09 | 1,167 | 1,02 | 0,02 | 8,0E-01 |
| Q9CQ21 | Malignant T-cell-amplified sequence 2 OS=Mus musculus OX=1009 | 17 | 2 | 2 | 2 | 181 | 20,4 | 9,11 | 5,07 | 2 | 0,962 | 0,625 | 0,628 | 1,001 | 0,573 | 1,077 | 0,843 | 0,88 | 0,95 | -0,05 | 8,0E-01 |
| O70310 | Glycylpeptide N-tetradecanoyltransferase 1 OS=Mus musculus OX= | 8 | 3 | 3 | 3 | 496 | 56,9 | 8 | 6,87 | 3 | 1,388 | 0,987 | 1,024 | 1,024 | 1,09 | 1,043 | 1,119 | 1,068 | 1,02 | 0,02 | 8,0E-01 |
| Q8C7M3 | E3 ubiquitin-protein ligase TRIM9 OS=Mus musculus OX=10090 GN | 7 | 5 | 5 | 5 | 817 | 90,8 | 7,24 | 8,26 | 5 | 1,205 | 1,106 | 1,228 | 0,967 | 0,946 | 1,297 | 0,892 | 1,244 | 1,03 | 0,03 | 8,0E-01 |
| Q8BGD5 | Carnitine O-palmitoyltransferase 1, brain isoform OS=Mus musculu | 5 | 4 | 5 | 4 | 798 | 90 | 7,49 | 5,04 | 4 | 1,125 | 1,154 | 1,047 | 1,104 | 0,931 | 1,292 | 0,965 | 1,149 | 1,02 | 0,02 | 8,0E-01 |
| Q8BWF0 | Succinate-semialdehyde dehydrogenase, mitochondrial OS=Mus m | 32 | 15 | 20 | 15 | 523 | 55,9 | 8,25 | 41,38 | 15 | 1,236 | 1,075 | 1,093 | 0,964 | 1,014 | 1,273 | 1,054 | 1,112 | 0,98 | -0,02 | 8,0E-01 |
| Q3UH60 | Disco-interacting protein 2 homolog B OS=Mus musculus OX=1009 | 8 | 9 | 11 | 9 | 1574 | 171 | 8,09 | 14,15 | 9 | 1,333 | 0,903 | 1,151 | 0,829 | 0,953 | 1,091 | 0,975 | 1,069 | 1,03 | 0,03 | 8,0E-01 |
| P11983 | T-complex protein 1 subunit alpha OS=Mus musculus OX=10090 GN | 23 | 12 | 18 | 12 | 556 | 60,4 | 6,16 | 40,45 | 12 | 1,084 | 1,13 | 1,056 | 0,985 | 0,965 | 1,135 | 1,059 | 1,047 | 1,01 | 0,01 | 8,0E-01 |
| Q91VC3 | Eukaryotic initiation factor 4A-III OS=Mus musculus OX=10090 GN= | 5 | 2 | 2 | 2 | 411 | 46,8 | 6,73 | 2,12 | 2 | 1,023 | 1,25 | 0,906 | 0,85 | 0,814 | 1,462 | 0,776 | 1,173 | 0,95 | -0,05 | 8,0E-01 |
| Q9CR35 | Chymotrypsinogen B OS=Mus musculus OX=10090 GN=Ctrb1 PE=1 | 3 | 1 | 1 | 1 | 263 | 27,8 | 5,07 | 2,4 | 1 | 1,821 | 1,161 | 1,217 | 0,669 | 0,862 | 1,603 | 0,916 | 1,179 | 1,07 | 0,07 | 8,0E-01 |
| Q99LS3 | Phosphoserine phosphatase OS=Mus musculus OX=10090 GN=Psph | 8 | 2 | 2 | 2 | 225 | 25,1 | 6,14 | 2,74 | 2 | 1,207 | 0,844 | 1,11 | 1,199 | 1,044 | 1,067 | 1,01 | 1,364 | 0,97 | -0,03 | 8,0E-01 |
| Q9R0N8 | Synaptotagmin-6 OS=Mus musculus OX=10090 GN=Syt6 PE=1 SV=2 | 5 | 2 | 2 | 2 | 511 | 57,2 | 8,19 | 1,83 | 2 | 0,566 | 0,952 | 1,022 | 0,803 | 0,863 | 0,633 | 0,865 | 0,859 | 1,04 | 0,04 | 8,0E-01 |
| Q9D8Z6 | Autophagy-related protein 101 OS=Mus musculus OX=10090 GN=A | 5 | 1 | 1 | 1 | 218 | 25 | 6,15 | 2,09 | 1 | 1,106 | 1,358 | 1,305 | 1,364 | 1,263 | 1,795 | 0,59 | 1,216 | 1,06 | 0,06 | 8,0E-01 |
| Q91V41 | Ras-related protein Rab-14 OS=Mus musculus OX=10090 GN=Rab1 | 49 | 11 | 18 | 9 | 215 | 23,9 | 6,21 | 45,74 | 11 | 0,844 | 0,971 | 1,008 | 1,177 | 0,951 | 0,972 | 1,023 | 0,98 | 1,02 | 0,02 | 8,0E-01 |
| P10922 | Histone H1.0 OS=Mus musculus OX=10090 GN=H1f0 PE=2 SV=4 | 16 | 3 | 5 | 3 | 194 | 20,8 | 10,9 | 14,53 | 3 | 1,044 | 1,501 | 0,926 | 0,576 | 0,982 | 1,496 | 0,719 | 1,113 | 0,94 | -0,06 | 8,0E-01 |
| P42227 | Signal transducer and activator of transcription 3 OS=Mus musculu | 3 | 2 | 2 | 2 | 770 | 88 | 6,3 | 2,7 | 2 | 0,818 | 1,119 | 0,877 | 1,024 | 0,856 | 1,045 | 1,06 | 0,964 | 0,98 | -0,02 | 8,0E-01 |
| Q64337 | Sequestosome-1 OS=Mus musculus OX=10090 GN=Sqstm1 PE=1 SV | 11 | 3 | 3 | 3 | 442 | 48,1 | 5,21 | 9,28 | 3 | 1,481 | 1,262 | 1,144 | 1,112 | 1,026 | 1,642 | 1,019 | 1,134 | 1,04 | 0,04 | 8,0E-01 |
| Q8CGC7 | Bifunctional glutamate/proline--tRNA ligase OS=Mus musculus OX= | 9 | 12 | 13 | 12 | 1512 | 170 | 7,66 | 24,17 | 12 | 1,098 | 1,015 | 1,03 | 0,952 | 0,97 | 1,031 | 1,031 | 1,028 | 1,01 | 0,01 | 8,0E-01 |
| Q9EQK5 | Major vault protein OS=Mus musculus OX=10090 GN=Mvp PE=1 SV | 2 | 1 | 1 | 1 | 861 | 95,9 | 5,59 | 0 | 1 | 1,254 | 0,713 | 0,672 | 0,718 | 0,662 | 0,509 | 0,972 | 1,021 | 1,06 | 0,06 | 8,0E-01 |
| P19246 | Neurofilament heavy polypeptide OS=Mus musculus OX=10090 GN | 20 | 22 | 26 | 19 | 1090 | 116,9 | 5,81 | 54,74 | 22 | 1,042 | 0,979 | 0,983 | 0,957 | 1,126 | 1,006 | 0,9 | 0,865 | 1,02 | 0,02 | 8,0E-01 |
| Q9Z0R4 | Intersectin-1 OS=Mus musculus OX=10090 GN=Itsn1 PE=1 SV=2 | 20 | 30 | 35 | 27 | 1714 | 194,2 | 7,91 | 54,27 | 30 | 1,096 | 1,069 | 1,042 | 1,022 | 1,003 | 1,057 | 1,076 | 1,069 | 1,01 | 0,01 | 8,0E-01 |
| P14602 | Heat shock protein beta-1 OS=Mus musculus OX=10090 GN=Hspb1 | 5 | 1 | 1 | 1 | 209 | 23 | 6,55 | 2,89 | 1 | 2,32 | 2,048 | 1,252 | 0,959 | 1,334 | 2,208 | 1,305 | 1,326 | 1,07 | 0,07 | 8,0E-01 |
| Q7TSE6 | Serine/threonine-protein kinase 38-like OS=Mus musculus OX=100 | 8 | 3 | 3 | 3 | 464 | 53,7 | 6,96 | 3,23 | 3 | 0,696 | 1,198 | 1,006 | 1,005 | 0,98 | 0,918 | 1,032 | 1,089 | 0,97 | -0,03 | 8,0E-01 |
| Q3UBX0 | Transmembrane protein 109 OS=Mus musculus OX=10090 GN=Tm | 5 | 1 | 1 | 1 | 243 | 26,3 | 9,89 | 1,64 | 1 | 1,133 | 1,286 | 1,38 | 0,909 | 1,036 | 1,348 | 0,796 | 1,353 | 1,04 | 0,04 | 8,0E-01 |
| Q9D9H8 | UPF0565 protein C2orf69 homolog OS=Mus musculus OX=10090 PI | 5 | 2 | 2 | 2 | 365 | 41,7 | 7,87 | 2,37 | 2 | 0,933 | 1,145 | 1,063 | 1,147 | 1,057 | 0,911 | 1,097 | 1,149 | 1,02 | 0,02 | 8,1E-01 |
| Q920P5 | Adenylate kinase isoenzyme 5 OS=Mus musculus OX=10090 GN=Ak | 6 | 3 | 3 | 3 | 562 | 63,3 | 5,29 | 4,53 | 3 | 1,11 | 1 | 0,986 | 1,083 | 0,981 | 1,185 | 0,898 | 1,045 | 1,02 | 0,02 | 8,1E-01 |
| P61080 | Ubiquitin-conjugating enzyme E2 D1 OS=Mus musculus OX=10090 | 12 | 2 | 2 | 2 | 147 | 16,6 | 7,42 | 1,62 | 2 | 0,693 | 0,899 | 1,122 | 0,967 | 1,18 | 0,732 | 0,869 | 0,76 | 1,04 | 0,04 | 8,1E-01 |
| Q4VAA2 | Protein CDV3 OS=Mus musculus OX=10090 GN=Cdv3 PE=1 SV=2 | 16 | 3 | 3 | 3 | 281 | 29,7 | 6,1 | 7,48 | 3 | 1,02 | 0,868 | 1,038 | 1,053 | 0,872 | 1,049 | 1,111 | 1,015 | 0,98 | -0,02 | 8,1E-01 |
| Q60771 | Claudin-11 OS=Mus musculus OX=10090 GN=Cldn11 PE=1 SV=1 | 11 | 2 | 3 | 2 | 207 | 22,1 | 7,91 | 8,1 | 2 | 0,941 | 0,988 | 0,885 | 1,186 | 1,154 | 0,999 | 0,9 | 1,033 | 0,98 | -0,02 | 8,1E-01 |
| P97467 | Peptidyl-glycine alpha-amidating monooxygenase OS=Mus musculu | 3 | 3 | 3 | 3 | 979 | 108,9 | 6,55 | 4,05 | 3 | 0,884 | 0,923 | 0,945 | 1,093 | 0,984 | 1,087 | 0,893 | 0,944 | 0,98 | -0,02 | 8,1E-01 |
| A6H6A9-3 | Isoform 3 of Rab GTPase-activating protein 1-like OS=Mus musculu | 16 | 5 | 7 | 3 | 370 | 42,7 | 7,33 | 5,83 | 5 | 1,228 | 1,294 | 1,072 | 1,055 | 1,121 | 1,228 | 1,036 | 1,357 | 0,98 | -0,02 | 8,1E-01 |
| Q9CZ13 | Cytochrome b-c1 complex subunit 1, mitochondrial OS=Mus muscu | 27 | 11 | 27 | 10 | 480 | 52,8 | 6,21 | 69,62 | 11 | 1,102 | 1,02 | 1,031 | 0,958 | 0,996 | 1,085 | 0,972 | 1,019 | 1,01 | 0,01 | 8,1E-01 |
| Q9D7N6 | 39S ribosomal protein L30, mitochondrial OS=Mus musculus OX=10 | 12 | 2 | 2 | 2 | 160 | 18,3 | 10,15 | 2,98 | 2 | 0,766 | 0,919 | 0,97 | 0,83 | 0,935 | 0,794 | 0,851 | 0,85 | 1,02 | 0,02 | 8,1E-01 |
| P97447 | Four and a half LIM domains protein 1 OS=Mus musculus OX=1009 | 16 | 4 | 4 | 4 | 280 | 31,9 | 8,37 | 8,74 | 4 | 1,17 | 0,994 | 1,135 | 1,163 | 1,009 | 1,132 | 1,215 | 1,043 | 1,01 | 0,01 | 8,1E-01 |
| Q4ACU6 | SH3 and multiple ankyrin repeat domains protein 3 OS=Mus muscu | 12 | 15 | 19 | 15 | 1730 | 185,3 | 8,88 | 32,38 | 15 | 1,054 | 1,007 | 0,969 | 0,917 | 0,928 | 1,099 | 0,994 | 0,973 | 0,99 | -0,01 | 8,1E-01 |
| P47708 | Rabphilin-3A OS=Mus musculus OX=10090 GN=Rph3a PE=1 SV=2 | 29 | 16 | 21 | 15 | 681 | 75,4 | 8,27 | 46,11 | 16 | 1,066 | 0,969 | 0,966 | 0,977 | 0,997 | 1,005 | 0,996 | 0,953 | 1,01 | 0,01 | 8,1E-01 |
| P70207 | Plexin-A2 OS=Mus musculus OX=10090 GN=Plxna2 PE=1 SV=2 | 8 | 14 | 19 | 6 | 1894 | 211,4 | 6,54 | 33,55 | 14 | 1,061 | 0,885 | 1,021 | 1,001 | 1,075 | 0,884 | 0,883 | 1,06 | 1,02 | 0,02 | 8,1E-01 |
| Q5RKR3 | Immunoglobulin superfamily containing leucine-rich repeat protein | 3 | 2 | 2 | 2 | 745 | 79,7 | 5,49 | 5,01 | 2 | 1,045 | 0,893 | 1,046 | 1,096 | 1,026 | 1,228 | 0,878 | 1,033 | 0,98 | -0,02 | 8,1E-01 |
| A2RT62 | F-box/LRR-repeat protein 16 OS=Mus musculus OX=10090 GN=Fbx | 13 | 6 | 7 | 6 | 479 | 51,8 | 6,6 | 17,63 | 6 | 0,97 | 0,971 | 1,067 | 1,006 | 0,955 | 1,075 | 1,032 | 0,987 | 0,99 | -0,01 | 8,1E-01 |
| Q9JHD1 | Histone acetyltransferase KAT2B OS=Mus musculus OX=10090 GN= | 1 | 1 | 1 | 1 | 813 | 91,7 | 9,14 | 0 | 1 | 1,459 | 1,51 | 1,066 | 0,469 | 0,556 | 1,814 | 0,768 | 0,999 | 1,09 | 0,09 | 8,1E-01 |
| O55033 | Cytoplasmic protein NCK2 OS=Mus musculus OX=10090 GN=Nck2 F | 10 | 3 | 3 | 3 | 380 | 42,9 | 6,95 | 2,82 | 3 | 0,853 | 0,993 | 0,882 | 1,001 | 0,885 | 1,149 | 0,856 | 0,916 | 0,98 | -0,02 | 8,1E-01 |
| P84096 | Rho-related GTP-binding protein RhoG OS=Mus musculus OX=1009 | 26 | 4 | 5 | 3 | 191 | 21,3 | 8,12 | 15,72 | 4 | 0,966 | 0,98 | 1,044 | 1,033 | 1,02 | 1,095 | 0,944 | 0,92 | 1,01 | 0,01 | 8,1E-01 |
| Q8BU31 | Ras-related protein Rap-2c OS=Mus musculus OX=10090 GN=Rap2 | 25 | 3 | 5 | 1 | 183 | 20,7 | 4,94 | 6,87 | 3 | 0,825 | 0,87 | 1,049 | 0,897 | 0,853 | 1,023 | 0,791 | 0,905 | 1,02 | 0,02 | 8,1E-01 |
| Q60805 | Tyrosine-protein kinase Mer OS=Mus musculus OX=10090 GN=Mer | 1 | 1 | 1 | 1 | 994 | 110,1 | 5,5 | 2,07 | 1 | 0,988 | 0,759 | 0,87 | 0,951 | 0,706 | 0,87 | 0,976 | 1,116 | 0,97 | -0,03 | 8,1E-01 |
| P21460 | Cystatin-C OS=Mus musculus OX=10090 GN=Cst3 PE=1 SV=2 | 12 | 2 | 3 | 2 | 140 | 15,5 | 9 | 3,35 | 2 | 1,103 | 1,012 | 0,449 | 0,715 | 0,784 | 0,665 | 0,831 | 0,846 | 1,05 | 0,05 | 8,1E-01 |
| Q3UQ84 | Threonine--tRNA ligase, mitochondrial OS=Mus musculus OX=1009 | 9 | 7 | 8 | 7 | 723 | 81,6 | 7,87 | 11,01 | 7 | 0,964 | 1,052 | 1,088 | 0,964 | 0,943 | 1,115 | 0,872 | 1,074 | 1,02 | 0,02 | 8,1E-01 |
| Q3UHD1 | Adhesion G protein-coupled receptor B1 OS=Mus musculus OX=10 | 4 | 6 | 6 | 6 | 1582 | 173,2 | 7,56 | 13,82 | 6 | 1,295 | 1,007 | 1,006 | 0,934 | 1,09 | 1,133 | 1 | 1,103 | 0,98 | -0,02 | 8,1E-01 |
| Q8BHC1 | Ras-related protein Rab-39B OS=Mus musculus OX=10090 GN=Rab | 22 | 5 | 6 | 3 | 213 | 24,6 | 7,83 | 17,14 | 5 | 1,183 | 1,066 | 1,046 | 1,151 | 0,95 | 1,208 | 1,155 | 1,201 | 0,98 | -0,02 | 8,1E-01 |
| P59764 | Dedicator of cytokinesis protein 4 OS=Mus musculus OX=10090 GN | 2 | 4 | 4 | 4 | 1978 | 226,4 | 7,65 | 4,03 | 4 | 0,927 | 0,825 | 1,012 | 1,02 | 1,172 | 0,757 | 1,031 | 0,921 | 0,98 | -0,02 | 8,1E-01 |
| Q9DCS2 | Methyltransferase-like 26 OS=Mus musculus OX=10090 GN=Mettl2 | 5 | 1 | 1 | 1 | 204 | 22,7 | 6,52 | 2,16 | 1 | 1,098 | 0,878 | 1,151 | 1,16 | 1,19 | 1,252 | 0,868 | 1,082 | 0,98 | -0,02 | 8,1E-01 |
| O88983 | Syntaxin-8 OS=Mus musculus OX=10090 GN=Stx8 PE=1 SV=1 | 4 | 1 | 1 | 1 | 236 | 26,9 | 5,01 | 2,31 | 1 | 1,164 | 0,93 | 0,812 | 0,985 | 1,008 | 1,017 | 1,22 | 0,762 | 0,97 | -0,03 | 8,2E-01 |
| P62082 | 40S ribosomal protein S7 OS=Mus musculus OX=10090 GN=Rps7 PI | 4 | 1 | 1 | 1 | 194 | 22,1 | 10,1 | 2,21 | 1 | 0,653 | 1,26 | 1,152 | 1,034 | 0,902 | 0,843 | 1,102 | 1,107 | 1,04 | 0,04 | 8,2E-01 |
| Q61699 | Heat shock protein 105 kDa OS=Mus musculus OX=10090 GN=Hsph | 32 | 23 | 31 | 20 | 858 | 96,3 | 5,53 | 56,26 | 23 | 0,918 | 0,924 | 1,007 | 0,993 | 0,976 | 0,922 | 0,991 | 0,98 | 0,99 | -0,01 | 8,2E-01 |
| Q8R4E6 | Purine-rich element-binding protein gamma OS=Mus musculus OX= | 3 | 1 | 1 | 1 | 350 | 39,9 | 9,51 | 1,73 | 1 | 1,386 | 1,154 | 1,117 | 0,963 | 0,903 | 1,294 | 1,229 | 1,074 | 1,03 | 0,03 | 8,2E-01 |
| P58774 | Tropomyosin beta chain OS=Mus musculus OX=10090 GN=Tpm2 PI | 26 | 11 | 14 | 1 | 284 | 32,8 | 4,7 | 26,23 | 11 | 1,714 | 1,161 | 1,163 | 1,296 | 1,174 | 1,735 | 1,197 | 1,406 | 0,97 | -0,03 | 8,2E-01 |

| | | | | | | | | | | | | | | | | | | | | | |
|---|---|---|---|---|---|---|---|---|---|---|---|---|---|---|---|---|---|---|---|---|---|
| Q149L6 | DnaJ homolog subfamily B member 14 OS=Mus musculus OX=1009 | 3 | 1 | 1 | 1 | 379 | 42,3 | 8,59 | 2 | 1 | 0,87 | 1,138 | 0,768 | 1,197 | 0,95 | 0,962 | 0,852 | 1,098 | 1,03 | 0,03 | 8,2E-01 |
| Q8CH72 | E3 ubiquitin-protein ligase TRIM32 OS=Mus musculus OX=10090 GI | 14 | 8 | 8 | 8 | 655 | 72 | 6,9 | 9,72 | 8 | 0,951 | 1,019 | 0,998 | 1,005 | 1,016 | 1,012 | 1,006 | 0,911 | 1,01 | 0,01 | 8,2E-01 |
| Q91XD6 | Vacuolar protein-sorting-associated protein 36 OS=Mus musculus ( | 6 | 2 | 2 | 2 | 386 | 43,7 | 7,15 | 2,81 | 2 | 1,304 | 1,652 | 1,488 | 0,98 | 1,004 | 1,699 | 1,299 | 1,226 | 1,04 | 0,04 | 8,2E-01 |
| Q3UDE2 | Tubulin--tyrosine ligase-like protein 12 OS=Mus musculus OX=1009 | 5 | 3 | 4 | 3 | 639 | 74 | 5,63 | 6,83 | 3 | 0,963 | 1,199 | 1,173 | 1,285 | 1,074 | 0,958 | 1,124 | 1,36 | 1,02 | 0,02 | 8,2E-01 |
| O70172 | Phosphatidylinositol 5-phosphate 4-kinase type-2 alpha OS=Mus m | 19 | 7 | 7 | 3 | 405 | 46,1 | 6,99 | 14,4 | 7 | 1,007 | 0,972 | 1,194 | 1,16 | 1,071 | 1,105 | 1,024 | 1,078 | 1,01 | 0,01 | 8,2E-01 |
| Q9D773 | 39S ribosomal protein L2, mitochondrial OS=Mus musculus OX=10( | 9 | 3 | 3 | 3 | 306 | 33,3 | 11,06 | 4,19 | 3 | 0,934 | 0,978 | 1,063 | 0,982 | 1,045 | 0,971 | 0,98 | 0,991 | 0,99 | -0,01 | 8,2E-01 |
| O08759 | Ubiquitin-protein ligase E3A OS=Mus musculus OX=10090 GN=Ube | 6 | 4 | 5 | 4 | 870 | 99,8 | 5,08 | 6,54 | 4 | 0,684 | 0,778 | 0,945 | 1,095 | 1,024 | 0,696 | 1,017 | 0,878 | 0,97 | -0,03 | 8,2E-01 |
| Q91V12 | Cytosolic acyl coenzyme A thioester hydrolase OS=Mus musculus O | 17 | 5 | 8 | 5 | 381 | 42,5 | 8,68 | 23,23 | 5 | 0,912 | 0,973 | 0,964 | 1,003 | 0,956 | 0,895 | 1,076 | 0,965 | 0,99 | -0,01 | 8,2E-01 |
| Q01730 | Ras suppressor protein 1 OS=Mus musculus OX=10090 GN=Rsu1 PE | 6 | 1 | 1 | 1 | 277 | 31,5 | 8,88 | 0 | 1 | 1,218 | 1,171 | 1,286 | 0,921 | 1,033 | 1,323 | 1,002 | 1,136 | 1,02 | 0,02 | 8,2E-01 |
| Q60931 | Voltage-dependent anion-selective channel protein 3 OS=Mus mus | 34 | 9 | 12 | 8 | 283 | 30,7 | 8,79 | 42,59 | 9 | 0,903 | 1,078 | 0,932 | 0,978 | 0,983 | 1,011 | 1,036 | 0,906 | 0,99 | -0,01 | 8,2E-01 |
| Q6ZQA6 | Immunoglobulin superfamily member 3 OS=Mus musculus OX=100 | 1 | 1 | 1 | 1 | 1194 | 134,6 | 6,05 | 1,9 | 1 | 0,968 | 1,041 | 0,685 | 0,981 | 1,012 | 0,983 | 0,719 | 0,863 | 1,03 | 0,03 | 8,2E-01 |
| P07724 | Serum albumin OS=Mus musculus OX=10090 GN=Alb PE=1 SV=3 | 24 | 13 | 63 | 13 | 608 | 68,6 | 6,07 | 144,19 | 13 | 1,052 | 0,921 | 0,978 | 0,815 | 0,685 | 1,253 | 0,857 | 1,099 | 0,97 | -0,03 | 8,2E-01 |
| Q5SYH2 | Transmembrane protein 199 OS=Mus musculus OX=10090 GN=Tm( | 5 | 1 | 1 | 1 | 208 | 23,1 | 9,1 | 2,17 | 1 | 1,91 | 1,255 | 1,138 | 1,448 | 1,425 | 1,756 | 1,27 | 1,092 | 1,04 | 0,04 | 8,2E-01 |
| P09405 | Nucleolin OS=Mus musculus OX=10090 GN=Ncl PE=1 SV=2 | 7 | 4 | 4 | 4 | 707 | 76,7 | 4,75 | 6,75 | 4 | 1,413 | 1,25 | 1,103 | 0,809 | 0,992 | 1,341 | 0,981 | 1,117 | 1,03 | 0,03 | 8,2E-01 |
| Q8BLR2 | Copine-4 OS=Mus musculus OX=10090 GN=Cpne4 PE=1 SV=1 | 15 | 7 | 10 | 7 | 557 | 62,4 | 6,33 | 17,66 | 7 | 1,119 | 1,011 | 1,037 | 1,124 | 1,033 | 1,102 | 1,189 | 1,013 | 0,99 | -0,01 | 8,2E-01 |
| A2ARP1 | Inositol hexakisphosphate and diphosphoinositol-pentakisphospha | 1 | 1 | 1 | 1 | 1436 | 159,8 | 5,39 | 2,26 | 1 | 1,015 | 0,979 | 1,16 | 1,46 | 1,181 | 1,437 | 0,759 | 1,07 | 1,04 | 0,04 | 8,2E-01 |
| P35762 | CD81 antigen OS=Mus musculus OX=10090 GN=Cd81 PE=1 SV=2 | 8 | 1 | 5 | 1 | 236 | 25,8 | 5,83 | 8,28 | 1 | 1,097 | 1,43 | 0,83 | 1,089 | 0,939 | 1,033 | 0,824 | 1,474 | 1,04 | 0,04 | 8,2E-01 |
| Q8VDI9 | Alpha-1,2-mannosyltransferase ALG9 OS=Mus musculus OX=10090 | 2 | 1 | 1 | 1 | 611 | 69,5 | 8,94 | 0 | 1 | 1,245 | 1,076 | 1,48 | 0,912 | 1,095 | 1,378 | 0,874 | 1,215 | 1,03 | 0,03 | 8,2E-01 |
| Q6PDY2 | 2-aminoethanethiol dioxygenase OS=Mus musculus OX=10090 GN= | 32 | 6 | 6 | 6 | 256 | 28,4 | 5,97 | 11,8 | 6 | 0,639 | 1,026 | 1,03 | 1,074 | 1,014 | 0,754 | 1,053 | 1,065 | 0,97 | -0,03 | 8,2E-01 |
| Q63850 | Nuclear pore glycoprotein p62 OS=Mus musculus OX=10090 GN=N | 3 | 1 | 1 | 1 | 526 | 53,2 | 5,31 | 2,11 | 1 | 1,315 | 1,264 | 1,357 | 1,503 | 1,246 | 1,147 | 1,188 | 1,724 | 1,03 | 0,03 | 8,2E-01 |
| Q3U2A8 | Valine--tRNA ligase, mitochondrial OS=Mus musculus OX=10090 GI | 1 | 1 | 1 | 1 | 1060 | 118,4 | 7,14 | 2,74 | 1 | 0,942 | 1,622 | 1,055 | 0,768 | 1,142 | 0,819 | 1,088 | 1,151 | 1,04 | 0,04 | 8,2E-01 |
| Q9CR51 | V-type proton ATPase subunit G 1 OS=Mus musculus OX=10090 GN | 24 | 2 | 2 | 2 | 118 | 13,7 | 7,97 | 4,65 | 2 | 1,181 | 0,931 | 1,305 | 1,836 | 1,101 | 0,744 | 1,496 | 1,652 | 1,05 | 0,05 | 8,2E-01 |
| Q80TY0 | Formin-binding protein 1 OS=Mus musculus OX=10090 GN=Fnbp1 I | 10 | 6 | 7 | 6 | 616 | 71,3 | 5,67 | 8,96 | 6 | 0,957 | 0,866 | 0,962 | 1,081 | 0,944 | 1,065 | 1,017 | 0,894 | 0,99 | -0,01 | 8,2E-01 |
| Q3U2P1 | Protein transport protein Sec24A OS=Mus musculus OX=10090 GN= | 1 | 1 | 1 | 1 | 1090 | 118,7 | 7,83 | 2,54 | 1 | 0,637 | 0,936 | 1,054 | 0,969 | 0,892 | 1,003 | 0,907 | 0,69 | 1,03 | 0,03 | 8,2E-01 |
| Q9JHW4 | Selenocysteine-specific elongation factor OS=Mus musculus OX=10 | 11 | 5 | 5 | 5 | 583 | 63,5 | 8,29 | 4,45 | 5 | 1,04 | 1,171 | 0,968 | 1,278 | 1,121 | 1,286 | 0,917 | 1,232 | 0,98 | -0,02 | 8,2E-01 |
| Q7TPD3 | Roundabout homolog 2 OS=Mus musculus OX=10090 GN=Robo2 P | 4 | 4 | 4 | 4 | 1470 | 161,1 | 6,33 | 5,48 | 4 | 1,244 | 0,696 | 1,077 | 1,543 | 1,29 | 1,009 | 1,1 | 0,986 | 1,04 | 0,04 | 8,3E-01 |
| Q80ZW2 | Protein THEM6 OS=Mus musculus OX=10090 GN=Them6 PE=1 SV= | 33 | 8 | 11 | 8 | 207 | 23,8 | 9,11 | 24,28 | 8 | 1,227 | 1,147 | 1,028 | 0,907 | 0,973 | 1,119 | 1,021 | 1,123 | 1,02 | 0,02 | 8,3E-01 |
| P60840 | Alpha-endosulfine OS=Mus musculus OX=10090 GN=Ensa PE=1 SV= | 64 | 7 | 12 | 6 | 121 | 13,3 | 7,24 | 26,71 | 7 | 0,966 | 1,046 | 0,917 | 0,981 | 0,919 | 0,968 | 0,972 | 1,093 | 0,99 | -0,01 | 8,3E-01 |
| Q9D0L4 | Uncharacterized aarF domain-containing protein kinase 1 OS=Mus | 9 | 3 | 4 | 3 | 525 | 59,7 | 8,05 | 4,92 | 3 | 1,993 | 1,386 | 1,135 | 1,073 | 0,775 | 1,95 | 1,605 | 1,556 | 0,95 | -0,05 | 8,3E-01 |
| Q9R0P5 | Destrin OS=Mus musculus OX=10090 GN=Dstn PE=1 SV=3 | 28 | 6 | 8 | 5 | 165 | 18,5 | 7,97 | 13,66 | 6 | 1,12 | 1,1 | 1,232 | 0,98 | 1,08 | 1,223 | 1,026 | 1,04 | 1,01 | 0,01 | 8,3E-01 |
| P63037 | DnaJ homolog subfamily A member 1 OS=Mus musculus OX=10090 | 12 | 5 | 7 | 5 | 397 | 44,8 | 7,08 | 8,93 | 5 | 1,202 | 1,124 | 1,095 | 1,02 | 0,97 | 1,322 | 1,081 | 1,144 | 0,98 | -0,02 | 8,3E-01 |
| Q9CY16 | 28S ribosomal protein S28, mitochondrial OS=Mus musculus OX=1( | 11 | 2 | 4 | 2 | 186 | 20,5 | 8,92 | 7,37 | 2 | 1,236 | 0,99 | 1,039 | 0,91 | 1,016 | 1,127 | 0,953 | 1,007 | 1,02 | 0,02 | 8,3E-01 |
| P09240 | Cholecystokinin OS=Mus musculus OX=10090 GN=Cck PE=1 SV=3 | 9 | 1 | 2 | 1 | 115 | 12,8 | 9,35 | 4,44 | 1 | 1,369 | 0,825 | 0,826 | 1,078 | 1,176 | 0,892 | 1,161 | 1,003 | 0,97 | -0,03 | 8,3E-01 |
| P22892 | AP-1 complex subunit gamma-1 OS=Mus musculus OX=10090 GN=/ | 11 | 10 | 10 | 10 | 822 | 91,3 | 6,8 | 16,57 | 10 | 0,885 | 0,891 | 1,064 | 1,053 | 0,998 | 0,959 | 0,979 | 1,003 | 0,99 | -0,01 | 8,3E-01 |
| Q9Z0L0 | Trophoblast glycoprotein OS=Mus musculus OX=10090 GN=Tpbg P | 12 | 4 | 4 | 4 | 426 | 46,4 | 6,83 | 8,73 | 4 | 1,077 | 1,014 | 1,103 | 0,955 | 0,907 | 1,188 | 0,982 | 1,01 | 1,02 | 0,02 | 8,3E-01 |
| Q9ERR7 | Selenoprotein F OS=Mus musculus OX=10090 GN=Selenof PE=1 SV= | 7 | 1 | 1 | 1 | 162 | 17,8 | 5,35 | 2,5 | 1 | 1,094 | 1,113 | 0,616 | 0,958 | 0,929 | 0,99 | 0,735 | 1,008 | 1,03 | 0,03 | 8,3E-01 |
| Q5IRJ6 | Zinc transporter 9 OS=Mus musculus OX=10090 GN=Slc30a9 PE=1 S | 4 | 3 | 3 | 3 | 567 | 62,8 | 8,69 | 4,82 | 3 | 0,817 | 0,883 | 1,128 | 0,958 | 1,107 | 0,882 | 0,98 | 0,894 | 0,98 | -0,02 | 8,3E-01 |
| D3YZG8 | Probable bifunctional methylenetetrahydrofolate dehydrogenase/( | 4 | 1 | 1 | 1 | 338 | 36,4 | 9,39 | 0 | 1 | 1,93 | 1,281 | 1,489 | 0,642 | 0,809 | 1,993 | 0,874 | 1,321 | 1,07 | 0,07 | 8,3E-01 |
| Q9JLC8 | Sacsin OS=Mus musculus OX=10090 GN=Sacs PE=1 SV=2 | 2 | 7 | 8 | 7 | 4582 | 520,4 | 7,11 | 10,69 | 7 | 1,17 | 0,984 | 1,064 | 1,07 | 0,979 | 1,092 | 1,084 | 1,184 | 0,99 | -0,01 | 8,3E-01 |
| Q91V51 | Probable tubulin polyglutamylase TTLL1 OS=Mus musculus OX=100 | 2 | 1 | 1 | 1 | 423 | 49,1 | 8,72 | 1,6 | 1 | 1,129 | 1,069 | 1,157 | 1,06 | 1,022 | 1,355 | 0,975 | 0,978 | 1,02 | 0,02 | 8,3E-01 |
| Q8C031 | Leucine-rich repeat-containing protein 4C OS=Mus musculus OX=1( | 4 | 2 | 2 | 2 | 640 | 71,9 | 7,15 | 3,29 | 2 | 1,627 | 0,808 | 0,932 | 1,094 | 1,162 | 0,69 | 0,949 | 1,444 | 1,05 | 0,05 | 8,3E-01 |
| P37040 | NADPH--cytochrome P450 reductase OS=Mus musculus OX=10090 | 16 | 11 | 17 | 11 | 678 | 77 | 5,53 | 16,32 | 11 | 1,018 | 0,927 | 0,969 | 0,998 | 1,027 | 0,953 | 0,979 | 0,927 | 1,01 | 0,01 | 8,3E-01 |
| Q8JZK9 | Hydroxymethylglutaryl-CoA synthase, cytoplasmic OS=Mus muscul | 6 | 3 | 3 | 3 | 520 | 57,5 | 5,99 | 4,37 | 3 | 1,612 | 1,626 | 1,152 | 1,136 | 1,49 | 1,554 | 1,117 | 1,21 | 1,03 | 0,03 | 8,3E-01 |
| Q99LB2 | Dehydrogenase/reductase SDR family member 4 OS=Mus musculus | 3 | 1 | 1 | 1 | 279 | 29,9 | 9,38 | 2,67 | 1 | 1,173 | 0,996 | 1,155 | 0,928 | 0,988 | 1,257 | 0,97 | 1,117 | 0,98 | -0,02 | 8,3E-01 |
| Q3TIR1 | Trafficking protein particle complex subunit 13 OS=Mus musculus ( | 4 | 2 | 2 | 2 | 417 | 46,4 | 5,49 | 1,74 | 2 | 1,076 | 1,265 | 1,17 | 1,116 | 1,272 | 1,162 | 1,142 | 1,1 | 0,99 | -0,01 | 8,3E-01 |
| A2A6Q5 | Cell division cycle protein 27 homolog OS=Mus musculus OX=1009( | 2 | 1 | 1 | 1 | 825 | 91,8 | 7,02 | 2,87 | 1 | 1,83 | 1,518 | 1,002 | 1,091 | 1,116 | 1,621 | 1,263 | 1,244 | 1,04 | 0,04 | 8,3E-01 |
| Q62351 | Transferrin receptor protein 1 OS=Mus musculus OX=10090 GN=Tfi | 11 | 7 | 10 | 7 | 763 | 85,7 | 6,57 | 21,96 | 7 | 0,941 | 1,022 | 1,049 | 1,113 | 1,077 | 0,979 | 0,977 | 1,053 | 1,01 | 0,01 | 8,3E-01 |
| P84084 | ADP-ribosylation factor 5 OS=Mus musculus OX=10090 GN=Arf5 PE | 42 | 7 | 8 | 3 | 180 | 20,5 | 6,79 | 19,65 | 7 | 1,15 | 1,173 | 0,806 | 1,167 | 1,067 | 1,294 | 1,028 | 1,006 | 0,98 | -0,02 | 8,3E-01 |
| Q91YE6 | Importin-9 OS=Mus musculus OX=10090 GN=Ipo9 PE=1 SV=3 | 2 | 2 | 2 | 2 | 1041 | 116 | 4,81 | 3,28 | 2 | 1,164 | 0,941 | 1,241 | 1,006 | 0,941 | 1,237 | 1,018 | 1,073 | 1,02 | 0,02 | 8,3E-01 |
| Q9D8W7 | OCIA domain-containing protein 2 OS=Mus musculus OX=10090 GI | 17 | 3 | 3 | 3 | 154 | 16,9 | 9,41 | 5,05 | 3 | 1,006 | 1,209 | 1,051 | 0,897 | 0,99 | 1,337 | 0,956 | 0,979 | 0,98 | -0,02 | 8,3E-01 |
| P26049 | Gamma-aminobutyric acid receptor subunit alpha-3 OS=Mus musc | 10 | 4 | 5 | 3 | 492 | 55,4 | 8,94 | 13,38 | 4 | 0,856 | 1,336 | 0,914 | 0,953 | 1,011 | 1,074 | 1,1 | 0,974 | 0,98 | -0,02 | 8,3E-01 |
| Q61233 | Plastin-2 OS=Mus musculus OX=10090 GN=Lcp1 PE=1 SV=4 | 5 | 3 | 4 | 1 | 627 | 70,1 | 5,33 | 7,23 | 3 | 0,911 | 1,138 | 0,998 | 1,118 | 1,232 | 0,773 | 1,141 | 1,12 | 0,98 | -0,02 | 8,3E-01 |
| Q60605 | Myosin light polypeptide 6 OS=Mus musculus OX=10090 GN=Myl6 | 36 | 6 | 7 | 4 | 151 | 16,9 | 4,65 | 14,26 | 6 | 0,838 | 0,941 | 0,937 | 0,797 | 0,909 | 0,758 | 0,935 | 0,864 | 1,01 | 0,01 | 8,3E-01 |
| Q9ESK9 | RB1-inducible coiled-coil protein 1 OS=Mus musculus OX=10090 GI | 7 | 9 | 9 | 9 | 1588 | 182,2 | 5,49 | 14,58 | 9 | 0,787 | 1,074 | 0,864 | 0,972 | 0,844 | 0,907 | 0,985 | 0,9 | 1,02 | 0,02 | 8,3E-01 |
| A3KGB4 | TBC1 domain family member 8B OS=Mus musculus OX=10090 GN= | 1 | 1 | 1 | 1 | 1114 | 127,8 | 5,49 | 1,77 | 1 | 0,818 | 1,057 | 1,326 | 1,003 | 1,035 | 1,124 | 1,031 | 1,109 | 0,98 | -0,02 | 8,3E-01 |
| Q99J27 | Acetyl-coenzyme A transporter 1 OS=Mus musculus OX=10090 GN= | 1 | 1 | 1 | 1 | 550 | 61 | 8,16 | 0 | 1 | 2,072 | 1,632 | 1,329 | 0,713 | 0,773 | 2,413 | 0,948 | 1,2 | 1,08 | 0,08 | 8,3E-01 |
| P80560 | Receptor-type tyrosine-protein phosphatase N2 OS=Mus musculus | 5 | 5 | 5 | 4 | 1001 | 111,4 | 5,95 | 4,49 | 5 | 0,703 | 0,865 | 0,601 | 0,785 | 0,782 | 0,437 | 0,972 | 0,877 | 0,96 | -0,04 | 8,3E-01 |
| P50571 | Gamma-aminobutyric acid receptor subunit beta-1 OS=Mus muscu | 15 | 6 | 6 | 2 | 474 | 54,1 | 8,78 | 12,62 | 6 | 0,871 | 1,374 | 0,895 | 0,96 | 0,963 | 0,927 | 0,977 | 1,123 | 1,03 | 0,03 | 8,3E-01 |
| Q9QY81 | Nuclear pore membrane glycoprotein 210 OS=Mus musculus OX=1( | 1 | 1 | 1 | 1 | 1886 | 204 | 6,65 | 2,59 | 1 | 1,514 | 0,995 | 1,164 | 1,354 | 0,899 | 1,489 | 1,246 | 1,248 | 1,03 | 0,03 | 8,3E-01 |
| Q9QYI4 | DnaJ homolog subfamily B member 12 OS=Mus musculus OX=1009 | 3 | 1 | 1 | 1 | 376 | 42 | 8,51 | 2,14 | 1 | 1,011 | 1,362 | 1,075 | 1,114 | 1,216 | 1,481 | 0,962 | 1,025 | 0,97 | -0,03 | 8,3E-01 |
| P62311 | U6 snRNA-associated Sm-like protein LSm3 OS=Mus musculus OX=: | 12 | 1 | 1 | 1 | 102 | 11,8 | 4,7 | 2,41 | 1 | 1,134 | 1,162 | 0,973 | 1,017 | 0,903 | 1,25 | 1,05 | 1,159 | 0,98 | -0,02 | 8,4E-01 |
| Q80WJ7 | Protein LYRIC OS=Mus musculus OX=10090 GN=Mtdh PE=1 SV=1 | 8 | 4 | 4 | 4 | 579 | 63,8 | 9,33 | 6 | 4 | 1,491 | 1,333 | 1,153 | 1,091 | 1,401 | 0,959 | 1,088 | 1,486 | 1,03 | 0,03 | 8,4E-01 |

| | | | | | | | | | | | | | | | | | | | | | |
|---|---|---|---|---|---|---|---|---|---|---|---|---|---|---|---|---|---|---|---|---|---|
| P01864 | Ig gamma-2A chain C region secreted form OS=Mus musculus OX=1 | 7 | 2 | 3 | 2 | 335 | 36,6 | 8,22 | 8,56 | 2 | 8,414 | 1,065 | 1,098 | 0,926 | 0,972 | 6,191 | 1,177 | 1,222 | 1,20 | 0,20 | 8,4E-01 |
| Q9D2R0 | Acetoacetyl-CoA synthetase OS=Mus musculus OX=10090 GN=Aacs | 5 | 3 | 3 | 3 | 672 | 75,2 | 6,71 | 2,67 | 3 | 0,906 | 1,833 | 1,592 | 1,572 | 1,087 | 1,974 | 1,695 | 1,386 | 0,96 | -0,04 | 8,4E-01 |
| Q640M6 | Glycerophosphodiester phosphodiesterase domain-containing prot | 3 | 2 | 2 | 2 | 607 | 68,8 | 8 | 4,06 | 2 | 1,245 | 1,081 | 1,127 | 0,987 | 0,87 | 1,499 | 0,947 | 0,992 | 1,03 | 0,03 | 8,4E-01 |
| Q8R5L3 | Vam6/Vps39-like protein OS=Mus musculus OX=10090 GN=Vps39 I | 6 | 4 | 4 | 4 | 886 | 101,6 | 6,99 | 5,62 | 4 | 1,222 | 1,226 | 1,263 | 1,096 | 1,171 | 1,345 | 1,211 | 1,131 | 0,99 | -0,01 | 8,4E-01 |
| P46638 | Ras-related protein Rab-11B OS=Mus musculus OX=10090 GN=Rab | 40 | 9 | 12 | 9 | 218 | 24,5 | 5,94 | 28,89 | 9 | 0,956 | 0,95 | 0,987 | 1,037 | 1,004 | 0,894 | 1,053 | 1,013 | 0,99 | -0,01 | 8,4E-01 |
| Q3TZZ7 | Extended synaptotagmin-2 OS=Mus musculus OX=10090 GN=Esyt2 | 4 | 3 | 3 | 3 | 845 | 94,1 | 7,75 | 3,45 | 3 | 1,113 | 1,168 | 0,777 | 1,197 | 1,213 | 0,864 | 1,054 | 1,02 | 1,03 | 0,03 | 8,4E-01 |
| Q9CZR3 | Mitochondrial import receptor subunit TOM40B OS=Mus musculus | 6 | 2 | 2 | 2 | 308 | 34 | 7,27 | 2 | 2 | 0,711 | 2,232 | 1,612 | 1,32 | 0,685 | 1,944 | 1,356 | 1,537 | 1,06 | 0,06 | 8,4E-01 |
| Q9DCW4 | Electron transfer flavoprotein subunit beta OS=Mus musculus OX=1 | 42 | 8 | 11 | 8 | 255 | 27,6 | 8,1 | 28,61 | 8 | 1,132 | 1,066 | 0,983 | 0,966 | 1,002 | 1,16 | 1,019 | 1,012 | 0,99 | -0,01 | 8,4E-01 |
| P63030 | Mitochondrial pyruvate carrier 1 OS=Mus musculus OX=10090 GN= | 7 | 1 | 1 | 1 | 109 | 12,4 | 9,61 | 1,65 | 1 | 0,975 | 1,553 | 0,863 | 0,997 | 0,965 | 1,041 | 1,061 | 1,183 | 1,03 | 0,03 | 8,4E-01 |
| Q9DCS3 | Enoyl-[acyl-carrier-protein] reductase, mitochondrial OS=Mus musc | 7 | 2 | 5 | 2 | 373 | 40,3 | 9,07 | 11,62 | 2 | 0,99 | 1,002 | 0,796 | 1,129 | 0,939 | 0,893 | 0,994 | 1,027 | 1,02 | 0,02 | 8,4E-01 |
| P43406 | Integrin alpha-V OS=Mus musculus OX=10090 GN=Itgav PE=1 SV=2 | 11 | 11 | 12 | 11 | 1044 | 115,3 | 5,63 | 32,54 | 11 | 1,07 | 0,872 | 0,978 | 1,01 | 1,014 | 0,974 | 0,964 | 1,015 | 0,99 | -0,01 | 8,4E-01 |
| P62315 | Small nuclear ribonucleoprotein Sm D1 OS=Mus musculus OX=1009 | 20 | 2 | 2 | 2 | 119 | 13,3 | 11,56 | 6,98 | 2 | 0,932 | 1,111 | 1,051 | 0,887 | 0,9 | 1,244 | 0,898 | 0,85 | 1,02 | 0,02 | 8,4E-01 |
| P16858 | Glyceraldehyde-3-phosphate dehydrogenase OS=Mus musculus OX | 44 | 17 | 59 | 17 | 333 | 35,8 | 8,25 | 148,52 | 17 | 0,916 | 0,986 | 0,981 | 1,022 | 0,975 | 0,936 | 1,011 | 0,96 | 1,01 | 0,01 | 8,4E-01 |
| Q9DBX3 | Sushi domain-containing protein 2 OS=Mus musculus OX=10090 GI | 4 | 2 | 3 | 2 | 820 | 90,6 | 6,62 | 5,98 | 2 | 1,068 | 0,945 | 1,052 | 0,893 | 0,817 | 1,148 | 0,891 | 1,03 | 1,02 | 0,02 | 8,4E-01 |
| Q6P5E4 | UDP-glucose:glycoprotein glucosyltransferase 1 OS=Mus musculus | 10 | 13 | 17 | 13 | 1551 | 176,3 | 5,62 | 29,15 | 13 | 1,248 | 1,13 | 1,044 | 0,883 | 0,996 | 1,177 | 1,049 | 1,009 | 1,02 | 0,02 | 8,4E-01 |
| Q8C050 | Ribosomal protein S6 kinase alpha-5 OS=Mus musculus OX=10090 | 1 | 1 | 1 | 1 | 863 | 96,5 | 7,34 | 2,11 | 1 | 0,897 | 0,757 | 1,201 | 1,245 | 0,904 | 0,95 | 0,986 | 1,15 | 1,03 | 0,03 | 8,4E-01 |
| Q9EQ20 | Methylmalonate-semialdehyde dehydrogenase [acylating], mitocho | 24 | 11 | 15 | 11 | 535 | 57,9 | 8,07 | 36,55 | 11 | 1,036 | 1,09 | 1,025 | 0,98 | 0,958 | 1,099 | 1,017 | 1,091 | 0,99 | -0,01 | 8,4E-01 |
| P07744 | Keratin, type II cytoskeletal 4 OS=Mus musculus OX=10090 GN=Krt4 | 7 | 6 | 10 | 2 | 525 | 56,2 | 8,15 | 17,44 | 6 | 0,968 | 1,369 | 0,941 | 0,831 | 1,539 | 0,861 | 0,82 | 0,701 | 1,05 | 0,05 | 8,4E-01 |
| Q9Z2S7 | TSC22 domain family protein 3 OS=Mus musculus OX=10090 GN=T | 12 | 2 | 3 | 1 | 137 | 15,2 | 4,53 | 4,56 | 2 | 0,677 | 1,058 | 1,046 | 1,14 | 1,178 | 0,781 | 0,928 | 1,152 | 0,97 | -0,03 | 8,4E-01 |
| Q91ZA3 | Propionyl-CoA carboxylase alpha chain, mitochondrial OS=Mus mu | 24 | 17 | 26 | 17 | 724 | 79,9 | 7,25 | 45,87 | 17 | 1,107 | 1,075 | 1,064 | 1,008 | 0,977 | 1,133 | 1,042 | 1,139 | 0,99 | -0,01 | 8,4E-01 |
| Q8K0C4 | Lanosterol 14-alpha demethylase OS=Mus musculus OX=10090 GN | 2 | 1 | 1 | 1 | 503 | 56,7 | 8,41 | 2,22 | 1 | 1,296 | 1,118 | 1,165 | 1,135 | 0,776 | 1,237 | 1,068 | 1,501 | 1,03 | 0,03 | 8,4E-01 |
| O35381 | Acidic leucine-rich nuclear phosphoprotein 32 family member A OS | 10 | 3 | 3 | 3 | 247 | 28,5 | 4,07 | 3,84 | 3 | 1,143 | 0,884 | 1,069 | 1,179 | 0,967 | 1,189 | 0,974 | 1,075 | 1,02 | 0,02 | 8,4E-01 |
| Q3UU96 | Serine/threonine-protein kinase MRCK alpha OS=Mus musculus OX | 5 | 7 | 7 | 7 | 1719 | 195,4 | 6,51 | 12,49 | 7 | 1,061 | 1,066 | 1,232 | 1,173 | 1,062 | 1,139 | 1,148 | 1,228 | 0,99 | -0,01 | 8,4E-01 |
| Q3UJU9 | Regulator of microtubule dynamics protein 3 OS=Mus musculus OX | 14 | 7 | 8 | 7 | 470 | 52 | 5,21 | 13,69 | 7 | 1,51 | 1,149 | 1,183 | 1,224 | 1,142 | 1,458 | 1,187 | 1,372 | 0,98 | -0,02 | 8,4E-01 |
| Q5HZI2 | C2 calcium-dependent domain-containing protein 4C OS=Mus musc | 14 | 4 | 4 | 4 | 419 | 44,6 | 9,73 | 9,26 | 4 | 0,945 | 1,178 | 0,787 | 0,813 | 0,811 | 1,324 | 0,71 | 1,013 | 0,97 | -0,03 | 8,4E-01 |
| Q8CI78 | Required for meiotic nuclear division protein 1 homolog OS=Mus m | 10 | 4 | 4 | 4 | 450 | 51,8 | 7,99 | 9,5 | 4 | 1,193 | 1,062 | 1,232 | 0,98 | 1,104 | 1,246 | 0,975 | 1,075 | 1,02 | 0,02 | 8,4E-01 |
| P49025 | Citron Rho-interacting kinase OS=Mus musculus OX=10090 GN=Cit | 8 | 16 | 16 | 16 | 2055 | 235,2 | 6,54 | 19,03 | 16 | 1,038 | 0,994 | 0,968 | 0,994 | 1,06 | 0,943 | 1,027 | 0,988 | 0,99 | -0,01 | 8,4E-01 |
| P62192 | 26S proteasome regulatory subunit 4 OS=Mus musculus OX=10090 | 27 | 10 | 12 | 9 | 440 | 49,2 | 6,21 | 22 | 10 | 1,363 | 1,071 | 1,074 | 0,971 | 0,92 | 1,404 | 0,962 | 1,079 | 1,03 | 0,03 | 8,4E-01 |
| Q8K245 | UV radiation resistance-associated protein OS=Mus musculus OX=1 | 6 | 4 | 5 | 4 | 698 | 77,5 | 7,97 | 2,23 | 4 | 1,38 | 0,919 | 0,755 | 0,815 | 0,953 | 0,891 | 0,957 | 1,197 | 0,97 | -0,03 | 8,4E-01 |
| O88487 | Cytoplasmic dynein 1 intermediate chain 2 OS=Mus musculus OX=1 | 14 | 7 | 9 | 7 | 612 | 68,4 | 5,29 | 18,99 | 7 | 0,842 | 0,945 | 0,94 | 1,033 | 0,978 | 0,779 | 0,972 | 1,093 | 0,98 | -0,02 | 8,4E-01 |
| Q9WV55 | Vesicle-associated membrane protein-associated protein A OS=Mu | 28 | 7 | 8 | 6 | 249 | 27,8 | 8,4 | 17,62 | 7 | 1,173 | 1,034 | 1,161 | 1,128 | 1,062 | 1,152 | 1,134 | 1,181 | 0,99 | -0,01 | 8,4E-01 |
| Q91W50 | Cold shock domain-containing protein E1 OS=Mus musculus OX=10 | 16 | 10 | 11 | 10 | 798 | 88,7 | 6,37 | 21,37 | 10 | 1,053 | 1,006 | 1,097 | 1,034 | 1,034 | 1,105 | 1,065 | 1,009 | 0,99 | -0,01 | 8,5E-01 |
| D3YZP9 | Coiled-coil domain-containing protein 6 OS=Mus musculus OX=100 | 2 | 1 | 1 | 1 | 469 | 52,9 | 7,34 | 1,65 | 1 | 1,032 | 1,249 | 1,15 | 1,079 | 0,801 | 1,404 | 0,874 | 1,302 | 1,03 | 0,03 | 8,5E-01 |
| Q9CQV8 | 14-3-3 protein beta/alpha OS=Mus musculus OX=10090 GN=Ywhab | 47 | 14 | 32 | 4 | 246 | 28,1 | 4,83 | 71,28 | 14 | 0,902 | 1,005 | 0,973 | 1,032 | 0,954 | 0,883 | 0,973 | 1,064 | 1,01 | 0,01 | 8,5E-01 |
| Q8BJH1 | Zinc finger C2HC domain-containing protein 1A OS=Mus musculus | 8 | 4 | 6 | 4 | 324 | 35,1 | 9,88 | 7,37 | 4 | 1,093 | 0,88 | 0,948 | 1,048 | 0,954 | 1,062 | 0,947 | 0,961 | 1,01 | 0,01 | 8,5E-01 |
| Q9ES00 | Ubiquitin conjugation factor E4 B OS=Mus musculus OX=10090 GN | 3 | 3 | 3 | 3 | 1173 | 133,2 | 6,07 | 2 | 3 | 1,273 | 1,209 | 1,181 | 1,064 | 1,14 | 1,314 | 1,073 | 1,257 | 0,99 | -0,01 | 8,5E-01 |
| Q9JHW2 | Omega-amidase NIT2 OS=Mus musculus OX=10090 GN=Nit2 PE=1 S | 20 | 4 | 4 | 4 | 276 | 30,5 | 6,9 | 6,1 | 4 | 1,079 | 1,362 | 1,452 | 1,233 | 1,175 | 1,587 | 1,185 | 1,065 | 1,02 | 0,02 | 8,5E-01 |
| Q9WUR9 | Adenylate kinase 4, mitochondrial OS=Mus musculus OX=10090 GN | 21 | 4 | 9 | 4 | 223 | 25 | 7,53 | 15,1 | 4 | 1,006 | 1,039 | 0,937 | 1,008 | 1,012 | 0,903 | 1,093 | 0,944 | 1,01 | 0,01 | 8,5E-01 |
| Q8VDU5 | SNF-related serine/threonine-protein kinase OS=Mus musculus OX | 11 | 7 | 8 | 7 | 748 | 81,9 | 7,49 | 7,35 | 7 | 0,996 | 0,868 | 0,948 | 1,24 | 0,844 | 1,251 | 0,97 | 0,889 | 1,02 | 0,02 | 8,5E-01 |
| P97494 | Glutamate--cysteine ligase catalytic subunit OS=Mus musculus OX= | 5 | 2 | 2 | 2 | 637 | 72,5 | 5,9 | 5,76 | 2 | 1,066 | 0,88 | 0,925 | 1,045 | 0,999 | 0,995 | 0,981 | 0,9 | 1,01 | 0,01 | 8,5E-01 |
| Q99LB6 | Methionine adenosyltransferase 2 subunit beta OS=Mus musculus | 3 | 1 | 1 | 1 | 334 | 37,4 | 6,95 | 2,87 | 1 | 1,158 | 0,876 | 0,863 | 0,852 | 0,942 | 1,077 | 0,822 | 0,981 | 0,98 | -0,02 | 8,5E-01 |
| P41731 | CD63 antigen OS=Mus musculus OX=10090 GN=Cd63 PE=1 SV=2 | 5 | 1 | 1 | 1 | 238 | 25,7 | 6,98 | 2,28 | 1 | 1,426 | 1,603 | 1,378 | 1,232 | 1,194 | 1,818 | 1,277 | 1,216 | 1,02 | 0,02 | 8,5E-01 |
| Q6P3A8 | 2-oxoisovalerate dehydrogenase subunit beta, mitochondrial OS=M | 6 | 2 | 2 | 2 | 390 | 42,9 | 6,7 | 5,24 | 2 | 1,094 | 1,171 | 1,171 | 0,855 | 0,888 | 1,19 | 0,988 | 1,143 | 1,02 | 0,02 | 8,5E-01 |
| Q8BXZ1 | Protein disulfide-isomerase TMX3 OS=Mus musculus OX=10090 GN | 17 | 7 | 8 | 7 | 456 | 51,8 | 5,16 | 18,29 | 7 | 0,888 | 0,996 | 0,969 | 1,051 | 1 | 0,914 | 1,035 | 0,989 | 0,99 | -0,01 | 8,5E-01 |
| Q9DCZ4 | MICOS complex subunit Mic26 OS=Mus musculus OX=10090 GN=A | 16 | 3 | 6 | 3 | 198 | 22,6 | 9,25 | 20,01 | 3 | 1,12 | 1,012 | 1,042 | 1,028 | 0,982 | 1,182 | 1,064 | 1,014 | 0,99 | -0,01 | 8,5E-01 |
| P21661 | Neuroendocrine convertase 2 OS=Mus musculus OX=10090 GN=Pc | 9 | 5 | 5 | 5 | 637 | 70,7 | 6,42 | 8,68 | 5 | 1,388 | 0,85 | 1,011 | 1,022 | 0,901 | 1,122 | 1,114 | 1,241 | 0,98 | -0,02 | 8,5E-01 |
| P27661 | Histone H2AX OS=Mus musculus OX=10090 GN=H2afx PE=1 SV=2 | 17 | 4 | 10 | 2 | 143 | 15,1 | 10,74 | 20,21 | 4 | 1,16 | 1,25 | 0,835 | 0,617 | 1,192 | 1,334 | 0,605 | 0,905 | 0,96 | -0,04 | 8,5E-01 |
| Q641P0 | Actin-related protein 3B OS=Mus musculus OX=10090 GN=Actr3b P | 22 | 9 | 18 | 6 | 418 | 47,5 | 6,02 | 23,21 | 9 | 0,993 | 1,216 | 0,997 | 1,053 | 1,084 | 1,084 | 1,013 | 1,034 | 1,01 | 0,01 | 8,5E-01 |
| Q9D8Y0 | EF-hand domain-containing protein D2 OS=Mus musculus OX=1009 | 33 | 10 | 16 | 10 | 240 | 26,8 | 5,06 | 25 | 10 | 0,872 | 0,955 | 1,032 | 0,891 | 0,86 | 0,906 | 0,997 | 0,95 | 1,01 | 0,01 | 8,5E-01 |
| Q9CQM9 | Glutaredoxin-3 OS=Mus musculus OX=10090 GN=Glrx3 PE=1 SV=1 | 17 | 5 | 8 | 5 | 337 | 37,8 | 5,59 | 13,27 | 5 | 1,022 | 1,075 | 1,161 | 1,178 | 0,979 | 1,128 | 1,159 | 1,127 | 1,01 | 0,01 | 8,5E-01 |
| P53996 | Cellular nucleic acid-binding protein OS=Mus musculus OX=10090 | 11 | 2 | 2 | 2 | 178 | 19,6 | 7,58 | 4,02 | 2 | 0,853 | 0,976 | 0,664 | 0,955 | 0,668 | 1,012 | 0,784 | 0,903 | 1,02 | 0,02 | 8,5E-01 |
| Q922D8 | C-1-tetrahydrofolate synthase, cytoplasmic OS=Mus musculus OX= | 18 | 15 | 19 | 14 | 935 | 101,1 | 7,14 | 34,43 | 15 | 1,124 | 1,067 | 1,052 | 1,065 | 1,013 | 1,184 | 1,015 | 1,132 | 0,99 | -0,01 | 8,5E-01 |
| P09925 | Surfeit locus protein 1 OS=Mus musculus OX=10090 GN=Surf1 PE=1 | 5 | 1 | 1 | 1 | 306 | 34,8 | 9,72 | 0 | 1 | 1,099 | 1,092 | 1,12 | 0,922 | 1,108 | 0,954 | 0,642 | 1,4 | 1,03 | 0,03 | 8,5E-01 |
| Q8CCJ4 | APC membrane recruitment protein 2 OS=Mus musculus OX=1009 | 7 | 4 | 4 | 4 | 672 | 69,9 | 6,64 | 6,26 | 4 | 1,108 | 0,96 | 0,93 | 0,957 | 1,059 | 0,853 | 0,928 | 1,064 | 1,01 | 0,01 | 8,5E-01 |
| P10107 | Annexin A1 OS=Mus musculus OX=10090 GN=Anxa1 PE=1 SV=2 | 2 | 1 | 1 | 1 | 346 | 38,7 | 7,37 | 2,01 | 1 | 1,233 | 1,067 | 0,761 | 0,639 | 1,069 | 0,771 | 0,808 | 1,183 | 0,97 | -0,03 | 8,5E-01 |
| Q64433 | 10 kDa heat shock protein, mitochondrial OS=Mus musculus OX=10 | 69 | 8 | 11 | 8 | 102 | 11 | 8,35 | 28,1 | 8 | 1,111 | 1,195 | 1,036 | 0,968 | 0,952 | 1,219 | 1,039 | 1,043 | 1,01 | 0,01 | 8,5E-01 |
| P02535 | Keratin, type I cytoskeletal 10 OS=Mus musculus OX=10090 GN=Krt | 14 | 10 | 20 | 6 | 570 | 57,7 | 5,11 | 43,06 | 10 | 1,303 | 1,226 | 0,774 | 0,819 | 1,438 | 1,004 | 0,619 | 0,894 | 1,04 | 0,04 | 8,5E-01 |
| Q9EPL2 | Calsyntenin-1 OS=Mus musculus OX=10090 GN=Clstn1 PE=1 SV=1 | 4 | 3 | 4 | 3 | 979 | 108,8 | 4,92 | 5,67 | 3 | 1,024 | 1,173 | 1,187 | 1,104 | 1,165 | 1,101 | 1,256 | 0,902 | 1,01 | 0,01 | 8,5E-01 |
| Q8BR92 | Paralemmin-2 OS=Mus musculus OX=10090 GN=Palm2 PE=1 SV=1 | 15 | 7 | 10 | 6 | 376 | 42,1 | 5,15 | 24,52 | 7 | 0,977 | 1,096 | 0,954 | 1,003 | 0,956 | 1,022 | 1,031 | 0,994 | 1,01 | 0,01 | 8,6E-01 |
| Q9CR16 | Peptidyl-prolyl cis-trans isomerase D OS=Mus musculus OX=10090 | 28 | 11 | 13 | 9 | 370 | 40,7 | 7,43 | 36,63 | 11 | 1,053 | 0,945 | 1,036 | 1,119 | 0,959 | 0,984 | 1,133 | 1,037 | 1,01 | 0,01 | 8,6E-01 |
| O35344 | Importin subunit alpha-4 OS=Mus musculus OX=10090 GN=Kpna3 I | 7 | 3 | 3 | 1 | 521 | 57,7 | 4,94 | 6,49 | 3 | 1,318 | 1,104 | 0,908 | 0,908 | 0,912 | 1,104 | 0,999 | 1,139 | 1,02 | 0,02 | 8,6E-01 |
| Q9JHS4 | ATP-dependent Clp protease ATP-binding subunit clpX-like, mitoch | 4 | 2 | 2 | 2 | 634 | 69,2 | 7,71 | 2,23 | 2 | 2,09 | 1,224 | 1,28 | 2,06 | 1,533 | 1,782 | 1,411 | 1,736 | 1,03 | 0,03 | 8,6E-01 |

| | | | | | | | | | | | | | | | | | | | | | |
|---|---|---|---|---|---|---|---|---|---|---|---|---|---|---|---|---|---|---|---|---|---|
| Q9WVI9 | C-Jun-amino-terminal kinase-interacting protein 1 OS=Mus muscul | 2 | 1 | 1 | 1 | 707 | 77,2 | 4,96 | 0 | 1 | 1,334 | 1,482 | 1,616 | 1,46 | 1,674 | 2,024 | 0,95 | 1,046 | 1,03 | 0,03 | 8,6E-01 |
| Q99JF8 | PC4 and SFRS1-interacting protein OS=Mus musculus OX=10090 GN | 2 | 1 | 1 | 1 | 528 | 59,7 | 9,13 | 1,91 | 1 | 1,085 | 1,055 | 1,03 | 0,653 | 0,915 | 0,812 | 0,797 | 1,196 | 1,03 | 0,03 | 8,6E-01 |
| Q6ZQ08 | CCR4-NOT transcription complex subunit 1 OS=Mus musculus OX=1 | 2 | 4 | 4 | 4 | 2375 | 266,6 | 7,11 | 5,66 | 4 | 1,043 | 1,072 | 0,854 | 0,888 | 0,825 | 1,038 | 0,957 | 1,097 | 0,98 | -0,02 | 8,6E-01 |
| P97372 | Proteasome activator complex subunit 2 OS=Mus musculus OX=10 | 11 | 2 | 2 | 2 | 239 | 27 | 5,76 | 0 | 2 | 1,103 | 1,24 | 1,45 | 1,039 | 1,401 | 1,049 | 1,329 | 0,947 | 1,02 | 0,02 | 8,6E-01 |
| B5TVM2 | Immunoglobulin-like domain-containing receptor 2 OS=Mus muscu | 6 | 3 | 3 | 3 | 661 | 73,2 | 7,68 | 5,07 | 3 | 1,331 | 1,167 | 1,134 | 1,301 | 1,359 | 1,243 | 1,229 | 1,041 | 1,01 | 0,01 | 8,6E-01 |
| B1AZP2-2 | Isoform 2 of Disks large-associated protein 4 OS=Mus musculus OX | 11 | 10 | 11 | 9 | 989 | 107,5 | 7,3 | 19,3 | 10 | 1,176 | 1,076 | 1,099 | 1,152 | 1,164 | 1,145 | 1,018 | 1,211 | 0,99 | -0,01 | 8,6E-01 |
| Q3UHB8 | Coiled-coil domain-containing protein 177 OS=Mus musculus OX=1 | 6 | 4 | 4 | 4 | 706 | 79,8 | 10,8 | 7,1 | 4 | 0,971 | 0,975 | 1,129 | 1,372 | 1,045 | 0,836 | 1,108 | 1,353 | 1,02 | 0,02 | 8,6E-01 |
| Q8JZZ7 | Adhesion G protein-coupled receptor L2 OS=Mus musculus OX=10 | 3 | 3 | 3 | 3 | 1487 | 166,5 | 6,71 | 7,01 | 3 | 0,843 | 0,859 | 0,78 | 0,94 | 0,78 | 0,633 | 0,94 | 1,003 | 1,02 | 0,02 | 8,6E-01 |
| Q9D1L0 | Coiled-coil-helix-coiled-coil-helix domain-containing protein 2 OS=M | 31 | 3 | 4 | 3 | 153 | 15,7 | 9,66 | 8,17 | 3 | 1,259 | 1,191 | 1,068 | 1,042 | 1,05 | 1,218 | 1,107 | 1,235 | 0,99 | -0,01 | 8,6E-01 |
| Q9D1N9 | 39S ribosomal protein L21, mitochondrial OS=Mus musculus OX=1 | 10 | 2 | 2 | 2 | 209 | 23,4 | 9,99 | 4,04 | 2 | 0,913 | 1,176 | 1,048 | 0,87 | 0,839 | 1,17 | 0,851 | 1,068 | 1,02 | 0,02 | 8,6E-01 |
| Q8BZJ7 | DCN1-like protein 2 OS=Mus musculus OX=10090 GN=Dcun1d2 PE= | 10 | 2 | 2 | 2 | 259 | 30 | 5,59 | 3,23 | 2 | 1,528 | 1,072 | 1,204 | 1,052 | 1,136 | 1,523 | 1,061 | 1,246 | 0,98 | -0,02 | 8,6E-01 |
| Q921W4 | Quinone oxidoreductase-like protein 1 OS=Mus musculus OX=1009 | 9 | 3 | 3 | 3 | 348 | 38,7 | 5,97 | 8,08 | 3 | 1,064 | 0,865 | 1,039 | 1,078 | 0,963 | 1,022 | 1,035 | 0,988 | 1,01 | 0,01 | 8,6E-01 |
| E9Q735 | Ubiquitin conjugation factor E4 A OS=Mus musculus OX=10090 GN | 1 | 1 | 1 | 1 | 1028 | 118,1 | 5,33 | 0 | 1 | 0,988 | 1,047 | 1,185 | 0,928 | 1,491 | 1,176 | 0,698 | 0,914 | 0,97 | -0,03 | 8,6E-01 |
| Q3TXX4 | Vesicular glutamate transporter 1 OS=Mus musculus OX=10090 GN | 11 | 6 | 11 | 6 | 560 | 61,6 | 7,34 | 11,41 | 6 | 1,001 | 1,03 | 0,896 | 1,073 | 0,966 | 0,976 | 1,062 | 1,028 | 0,99 | -0,01 | 8,6E-01 |
| Q8BYI9 | Tenascin-R OS=Mus musculus OX=10090 GN=Tnr PE=1 SV=2 | 9 | 11 | 11 | 11 | 1358 | 149,5 | 4,94 | 26,05 | 11 | 1,015 | 0,862 | 1,018 | 1,007 | 1,093 | 0,825 | 1,025 | 1,009 | 0,99 | -0,01 | 8,6E-01 |
| Q3V3V9-2 | Isoform 2 of Capping protein, Arp2/3 and myosin-I linker protein 2 | 11 | 5 | 5 | 5 | 738 | 79,7 | 6,77 | 11,73 | 5 | 1,366 | 1,138 | 0,986 | 1,057 | 0,82 | 1,07 | 1,213 | 1,343 | 1,02 | 0,02 | 8,6E-01 |
| P61079 | Ubiquitin-conjugating enzyme E2 D3 OS=Mus musculus OX=10090 | 7 | 1 | 2 | 1 | 147 | 16,7 | 7,8 | 2,29 | 1 | 0,87 | 0,752 | 1,09 | 1,16 | 0,839 | 0,997 | 1,268 | 0,867 | 0,98 | -0,02 | 8,6E-01 |
| P24549 | Retinal dehydrogenase 1 OS=Mus musculus OX=10090 GN=Aldh1a1 | 17 | 7 | 7 | 6 | 501 | 54,4 | 7,8 | 14,49 | 7 | 1,256 | 0,987 | 1,153 | 0,928 | 0,95 | 1,069 | 1,163 | 1,21 | 0,98 | -0,02 | 8,6E-01 |
| Q9JHK4 | Geranylgeranyl transferase type-2 subunit alpha OS=Mus musculus | 5 | 3 | 5 | 3 | 567 | 64,9 | 5,8 | 4,62 | 3 | 0,967 | 0,669 | 0,911 | 0,861 | 0,937 | 0,835 | 0,885 | 0,802 | 0,99 | -0,01 | 8,6E-01 |
| Q91WM2 | Haloacid dehalogenase-like hydrolase domain-containing 5 OS=Mu | 2 | 1 | 1 | 1 | 419 | 46,3 | 7,88 | 1,78 | 1 | 1,18 | 1,211 | 1,38 | 1,167 | 1,153 | 1,504 | 0,947 | 1,244 | 1,02 | 0,02 | 8,6E-01 |
| P80313 | T-complex protein 1 subunit eta OS=Mus musculus OX=10090 GN= | 21 | 12 | 17 | 12 | 544 | 59,6 | 7,84 | 28,91 | 12 | 1,043 | 0,992 | 1,17 | 1,125 | 1,017 | 1,145 | 1,048 | 1,158 | 0,99 | -0,01 | 8,6E-01 |
| Q9JIW9 | Ras-related protein Ral-B OS=Mus musculus OX=10090 GN=Ralb PE | 13 | 3 | 7 | 2 | 206 | 23,3 | 6,62 | 9,8 | 3 | 0,665 | 0,764 | 0,562 | 0,757 | 0,568 | 0,677 | 0,704 | 0,755 | 1,02 | 0,02 | 8,6E-01 |
| P54754 | Ephrin type-B receptor 3 OS=Mus musculus OX=10090 GN=Ephb3 F | 7 | 6 | 6 | 4 | 993 | 109,6 | 6,28 | 8,27 | 6 | 1,327 | 1,259 | 0,963 | 0,792 | 0,767 | 1,417 | 0,955 | 1,07 | 1,03 | 0,03 | 8,6E-01 |
| Q62407 | Striated muscle-specific serine/threonine-protein kinase OS=Mus m | 1 | 2 | 2 | 2 | 3262 | 354,1 | 8,21 | 3,74 | 2 | 0,857 | 1,286 | 0,825 | 1,071 | 0,832 | 1,395 | 0,869 | 1,062 | 0,97 | -0,03 | 8,6E-01 |
| Q8BRU6 | Synaptic vesicular amine transporter OS=Mus musculus OX=10090 | 2 | 1 | 1 | 1 | 517 | 55,7 | 6,23 | 0 | 1 | 1,478 | 0,532 | 0,947 | 0,82 | 1,023 | 0,812 | 1,176 | 0,916 | 0,96 | -0,04 | 8,7E-01 |
| Q62277 | Synaptophysin OS=Mus musculus OX=10090 GN=Syp PE=1 SV=2 | 15 | 5 | 6 | 5 | 314 | 34 | 4,94 | 15,08 | 5 | 0,658 | 1,125 | 0,814 | 0,891 | 0,938 | 0,823 | 0,977 | 0,824 | 0,98 | -0,02 | 8,7E-01 |
| Q8R307 | Vacuolar protein sorting-associated protein 18 homolog OS=Mus m | 2 | 2 | 2 | 2 | 973 | 110,1 | 6,09 | 2,07 | 2 | 0,994 | 1,177 | 1,221 | 0,999 | 1,041 | 1,325 | 0,963 | 0,99 | 1,02 | 0,02 | 8,7E-01 |
| Q9CR98 | Protein FAM136A OS=Mus musculus OX=10090 GN=Fam136a PE=1 | 30 | 4 | 6 | 4 | 138 | 15,7 | 7,61 | 18,28 | 4 | 1,027 | 0,959 | 1,071 | 0,908 | 0,963 | 0,95 | 1,083 | 1,002 | 0,99 | -0,01 | 8,7E-01 |
| Q9CXJ4 | ATP-binding cassette sub-family B member 8, mitochondrial OS=M | 6 | 4 | 4 | 4 | 717 | 78 | 9,07 | 10,61 | 4 | 1,044 | 1,219 | 1,16 | 1,001 | 0,947 | 1,258 | 1,046 | 1,115 | 1,01 | 0,01 | 8,7E-01 |
| P61759 | Prefoldin subunit 3 OS=Mus musculus OX=10090 GN=Vbp1 PE=1 S\ | 24 | 5 | 6 | 5 | 196 | 22,4 | 6,28 | 10,81 | 5 | 0,875 | 0,813 | 0,879 | 1,006 | 0,882 | 0,808 | 0,961 | 0,96 | 0,99 | -0,01 | 8,7E-01 |
| P68433 | Histone H3.1 OS=Mus musculus OX=10090 GN=Hist1h3a PE=1 SV=2 | 26 | 5 | 9 | 5 | 136 | 15,4 | 11,12 | 15,26 | 5 | 0,963 | 1,626 | 0,877 | 0,511 | 1,039 | 1,663 | 0,509 | 0,996 | 0,95 | -0,05 | 8,7E-01 |
| Q9D1R9 | 60S ribosomal protein L34 OS=Mus musculus OX=10090 GN=Rpl34 | 8 | 2 | 2 | 2 | 117 | 13,3 | 11,47 | 7,65 | 2 | 1,728 | 1,219 | 1,243 | 0,849 | 0,832 | 1,8 | 1,041 | 1,175 | 1,04 | 0,04 | 8,7E-01 |
| Q9CPW4 | Actin-related protein 2/3 complex subunit 5 OS=Mus musculus OX= | 34 | 4 | 11 | 4 | 151 | 16,3 | 5,67 | 27,97 | 4 | 1,252 | 1,242 | 1,065 | 0,975 | 0,941 | 1,222 | 1,101 | 1,205 | 1,01 | 0,01 | 8,7E-01 |
| Q6ZQ58 | La-related protein 1 OS=Mus musculus OX=10090 GN=Larp1 PE=1 S | 3 | 3 | 3 | 3 | 1072 | 121,1 | 8,79 | 7,5 | 3 | 1,233 | 1,191 | 1,162 | 0,931 | 0,936 | 1,306 | 1,18 | 1,166 | 0,98 | -0,02 | 8,7E-01 |
| P56371 | Ras-related protein Rab-4A OS=Mus musculus OX=10090 GN=Rab4 | 27 | 5 | 6 | 3 | 218 | 24,4 | 6,07 | 18,35 | 5 | 1,196 | 1,254 | 0,875 | 1,119 | 1,106 | 1,128 | 0,997 | 1,151 | 1,01 | 0,01 | 8,7E-01 |
| O55026 | Ectonucleoside triphosphate diphosphohydrolase 2 OS=Mus muscu | 4 | 2 | 2 | 2 | 495 | 54,3 | 8,37 | 2,67 | 2 | 1,291 | 0,75 | 1,109 | 1,171 | 1,104 | 0,919 | 0,918 | 1,504 | 0,97 | -0,03 | 8,7E-01 |
| Q8R317 | Ubiquilin-1 OS=Mus musculus OX=10090 GN=Ubqln1 PE=1 SV=1 | 10 | 4 | 5 | 4 | 582 | 61,9 | 4,94 | 6,5 | 4 | 1,377 | 1,046 | 1,259 | 0,866 | 0,825 | 1,197 | 1,064 | 1,353 | 1,02 | 0,02 | 8,7E-01 |
| Q99LD4 | COP9 signalosome complex subunit 1 OS=Mus musculus OX=10090 | 10 | 5 | 6 | 5 | 471 | 53,4 | 6,84 | 13,84 | 5 | 0,866 | 0,859 | 1,02 | 1,062 | 0,989 | 0,888 | 0,963 | 1,007 | 0,99 | -0,01 | 8,7E-01 |
| G5E897 | Protein O-glucosyltransferase 3 OS=Mus musculus OX=10090 GN=F | 2 | 1 | 1 | 1 | 503 | 57,7 | 7,74 | 0 | 1 | 0,955 | 1,284 | 0,895 | 0,765 | 1,044 | 1,19 | 0,543 | 1,255 | 0,97 | -0,03 | 8,7E-01 |
| Q9CZY3 | Ubiquitin-conjugating enzyme E2 variant 1 OS=Mus musculus OX=1 | 42 | 5 | 5 | 2 | 147 | 16,3 | 7,96 | 10,11 | 5 | 1 | 1,098 | 0,725 | 0,926 | 1,024 | 0,818 | 0,935 | 0,911 | 1,02 | 0,02 | 8,7E-01 |
| P62484 | Abl interactor 2 OS=Mus musculus OX=10090 GN=Abi2 PE=1 SV=1 | 23 | 9 | 12 | 5 | 446 | 49,4 | 6,01 | 20,25 | 9 | 1,037 | 0,88 | 1,027 | 1,015 | 1,002 | 0,846 | 0,947 | 1,118 | 1,01 | 0,01 | 8,7E-01 |
| Q8BMD8 | Calcium-binding mitochondrial carrier protein SCaMC-1 OS=Mus m | 2 | 1 | 1 | 1 | 475 | 52,9 | 7,43 | 1,62 | 1 | 1,107 | 0,994 | 0,876 | 0,964 | 1,033 | 1,256 | 0,821 | 0,903 | 0,98 | -0,02 | 8,7E-01 |
| Q99K85 | Phosphoserine aminotransferase OS=Mus musculus OX=10090 GN= | 20 | 8 | 12 | 8 | 370 | 40,4 | 8,03 | 30,36 | 8 | 1,007 | 1,016 | 1,051 | 1,071 | 1,072 | 1,023 | 1,009 | 1,055 | 1,00 | 0,00 | 8,7E-01 |
| P24472 | Glutathione S-transferase A4 OS=Mus musculus OX=10090 GN=Gst | 9 | 3 | 4 | 3 | 222 | 25,5 | 7,39 | 6,95 | 3 | 0,965 | 0,9 | 0,96 | 0,921 | 0,883 | 0,997 | 0,921 | 0,926 | 1,01 | 0,01 | 8,7E-01 |
| Q8R516 | E3 ubiquitin-protein ligase MIB2 OS=Mus musculus OX=10090 GN= | 1 | 1 | 1 | 1 | 973 | 105,9 | 8,09 | 0 | 1 | 1,087 | 1,12 | 1,006 | 0,935 | 0,743 | 1,314 | 0,95 | 1,057 | 1,02 | 0,02 | 8,7E-01 |
| Q9WU28 | Prefoldin subunit 5 OS=Mus musculus OX=10090 GN=Pfdn5 PE=1 S | 30 | 4 | 5 | 4 | 154 | 17,3 | 6,33 | 9,42 | 4 | 1,069 | 0,904 | 1,107 | 0,931 | 1 | 1,036 | 1,018 | 0,991 | 0,99 | -0,01 | 8,7E-01 |
| Q7TNG5 | Echinoderm microtubule-associated protein-like 2 OS=Mus muscul | 11 | 6 | 6 | 6 | 649 | 70,7 | 6,28 | 10,8 | 6 | 0,705 | 0,947 | 1,186 | 1,008 | 0,968 | 1,049 | 0,949 | 0,948 | 0,98 | -0,02 | 8,7E-01 |
| Q91VK4 | Integral membrane protein 2C OS=Mus musculus OX=10090 GN=Itr | 8 | 2 | 2 | 2 | 269 | 30,5 | 8,59 | 2,29 | 2 | 1,136 | 0,968 | 0,906 | 1,038 | 0,771 | 1,265 | 0,978 | 0,959 | 1,02 | 0,02 | 8,7E-01 |
| Q8BXN7 | Protein phosphatase 1K, mitochondrial OS=Mus musculus OX=1009 | 7 | 2 | 2 | 2 | 372 | 40,9 | 6,39 | 0 | 2 | 1,419 | 1,326 | 1,018 | 1,228 | 1,056 | 1,357 | 1,252 | 1,402 | 0,99 | -0,01 | 8,7E-01 |
| Q6PDL0 | Cytoplasmic dynein 1 light intermediate chain 2 OS=Mus musculus | 20 | 9 | 10 | 9 | 492 | 54,2 | 6,28 | 23,95 | 9 | 0,988 | 0,892 | 0,903 | 0,982 | 0,968 | 0,851 | 0,952 | 0,969 | 1,01 | 0,01 | 8,7E-01 |
| Q3UPH1 | Protein PRRC1 OS=Mus musculus OX=10090 GN=Prrc1 PE=1 SV=1 | 3 | 1 | 1 | 1 | 443 | 46,3 | 5,95 | 2,09 | 1 | 0,624 | 1,069 | 0,746 | 0,618 | 0,793 | 0,98 | 0,773 | 0,406 | 1,04 | 0,04 | 8,8E-01 |
| Q9ES07 | Solute carrier family 15 member 2 OS=Mus musculus OX=10090 GN | 2 | 2 | 2 | 2 | 729 | 81,6 | 7,34 | 2,26 | 2 | 0,902 | 1,158 | 0,956 | 1,165 | 1,162 | 1,024 | 0,948 | 1,101 | 0,99 | -0,01 | 8,8E-01 |
| Q9ES97 | Reticulon-3 OS=Mus musculus OX=10090 GN=Rtn3 PE=1 SV=2 | 19 | 18 | 21 | 17 | 964 | 103,8 | 4,92 | 40,08 | 18 | 0,882 | 0,961 | 0,958 | 1,003 | 0,961 | 0,87 | 1,007 | 0,992 | 0,99 | -0,01 | 8,8E-01 |
| Q1RLL3 | Copine-9 OS=Mus musculus OX=10090 GN=Cpne9 PE=1 SV=1 | 10 | 4 | 5 | 1 | 553 | 61,8 | 5,4 | 9,62 | 4 | 0,937 | 1,232 | 0,763 | 0,785 | 0,829 | 0,98 | 0,976 | 0,857 | 1,02 | 0,02 | 8,8E-01 |
| Q8CBW3 | Abl interactor 1 OS=Mus musculus OX=10090 GN=Abi1 PE=1 SV=3 | 27 | 12 | 16 | 8 | 481 | 52,3 | 7,64 | 36,36 | 12 | 1,029 | 0,944 | 1,013 | 1,039 | 0,928 | 0,999 | 1,048 | 1,028 | 1,01 | 0,01 | 8,8E-01 |
| Q8K093 | Thyrotropin-releasing hormone-degrading ectoenzyme OS=Mus mu | 1 | 1 | 1 | 1 | 1025 | 117,4 | 7,06 | 2,15 | 1 | 1,219 | 0,291 | 0,75 | 0,621 | 1,099 | 0,653 | 0,646 | 0,628 | 0,95 | -0,05 | 8,8E-01 |
| P70280 | Vesicle-associated membrane protein 7 OS=Mus musculus OX=100 | 11 | 2 | 2 | 2 | 220 | 25 | 8,6 | 2,6 | 2 | 1,091 | 1,402 | 0,968 | 0,956 | 0,936 | 1,681 | 1,056 | 0,881 | 0,97 | -0,03 | 8,8E-01 |
| O35071 | Kinesin-like protein KIF1C OS=Mus musculus OX=10090 GN=Kif1c P | 2 | 3 | 3 | 1 | 1100 | 122,4 | 7,03 | 2,55 | 3 | 1,24 | 0,999 | 2,037 | 1,797 | 1,913 | 1,16 | 1,213 | 1,594 | 1,03 | 0,03 | 8,8E-01 |
| Q9ERB0 | Synaptosomal-associated protein 29 OS=Mus musculus OX=10090 | 21 | 5 | 5 | 5 | 260 | 29,6 | 5,38 | 2,79 | 5 | 1,072 | 1,066 | 0,99 | 1,043 | 1,073 | 0,952 | 0,986 | 1,131 | 1,01 | 0,01 | 8,8E-01 |
| Q8R1F6 | Protein HID1 OS=Mus musculus OX=10090 GN=Hid1 PE=1 SV=1 | 3 | 2 | 2 | 2 | 788 | 88,7 | 5,94 | 2,67 | 2 | 1,073 | 1,421 | 0,9 | 1,084 | 0,96 | 1,16 | 1,116 | 1,165 | 1,02 | 0,02 | 8,8E-01 |
| Q6ZWX6 | Eukaryotic translation initiation factor 2 subunit 1 OS=Mus muscul | 17 | 6 | 7 | 6 | 315 | 36,1 | 5,08 | 17 | 6 | 0,855 | 0,929 | 1,005 | 0,959 | 0,877 | 1,013 | 0,904 | 0,983 | 0,99 | -0,01 | 8,8E-01 |
| P53026 | 60S ribosomal protein L10a OS=Mus musculus OX=10090 GN=Rpl1 | 24 | 5 | 8 | 5 | 217 | 24,9 | 9,98 | 13,15 | 5 | 0,939 | 1,082 | 0,964 | 0,925 | 1,085 | 0,931 | 0,946 | 0,914 | 1,01 | 0,01 | 8,8E-01 |

| | | | | | | | | | | | | | | | | | | | | | |
|---|---|---|---|---|---|---|---|---|---|---|---|---|---|---|---|---|---|---|---|---|---|
| Q66GT5 | Phosphatidylglycerophosphatase and protein-tyrosine phosphatase | 8 | 2 | 2 | 2 | 193 | 21,9 | 9,72 | 4,57 | 2 | 1,061 | 0,985 | 1,335 | 1,036 | 1,182 | 1,33 | 1,077 | 0,904 | 0,98 | -0,02 | 8,8E-01 |
| Q9CQ54 | NADH dehydrogenase [ubiquinone] 1 subunit C2 OS=Mus musculus | 42 | 8 | 14 | 8 | 120 | 14,2 | 9,2 | 30,94 | 8 | 1,041 | 1,124 | 0,96 | 1,022 | 1,037 | 0,985 | 1,036 | 1,065 | 1,01 | 0,01 | 8,8E-01 |
| B1AY13 | Ubiquitin carboxyl-terminal hydrolase 24 OS=Mus musculus OX=10 | 3 | 6 | 6 | 6 | 2617 | 293,8 | 6,19 | 7,5 | 6 | 1,17 | 0,9 | 1,042 | 1,269 | 0,969 | 1,087 | 1,035 | 1,365 | 0,98 | -0,02 | 8,8E-01 |
| Q8JZR6 | Electroneutral sodium bicarbonate exchanger 1 OS=Mus musculus | 4 | 4 | 4 | 1 | 1089 | 122,3 | 6,6 | 1,68 | 4 | 1,491 | 0,468 | 1,126 | 0,891 | 0,996 | 1,272 | 1,114 | 0,747 | 0,96 | -0,04 | 8,8E-01 |
| Q9JM63 | ATP-sensitive inward rectifier potassium channel 10 OS=Mus muscu | 2 | 1 | 1 | 1 | 379 | 42,4 | 8,29 | 0 | 1 | 1,122 | 0,739 | 0,744 | 0,925 | 0,738 | 1,022 | 0,836 | 1,006 | 0,98 | -0,02 | 8,8E-01 |
| Q9CR62 | Mitochondrial 2-oxoglutarate/malate carrier protein OS=Mus musc | 18 | 5 | 10 | 5 | 314 | 34,1 | 9,94 | 18,34 | 5 | 1,118 | 1,391 | 1,051 | 1,116 | 1,183 | 1,151 | 1,211 | 1,08 | 1,01 | 0,01 | 8,8E-01 |
| Q61656 | Probable ATP-dependent RNA helicase DDX5 OS=Mus musculus OX | 5 | 3 | 3 | 2 | 614 | 69,2 | 8,92 | 6,99 | 3 | 1,08 | 1,294 | 0,924 | 0,948 | 0,874 | 1,274 | 1,074 | 0,947 | 1,02 | 0,02 | 8,8E-01 |
| Q9CZ57 | 5-methylcytosine rRNA methyltransferase NSUN4 OS=Mus musculu | 3 | 1 | 1 | 1 | 381 | 42,8 | 8,25 | 2,06 | 1 | 1,24 | 1,552 | 1,542 | 1,088 | 1,194 | 1,472 | 1,323 | 1,519 | 0,98 | -0,02 | 8,8E-01 |
| Q9Z1X2 | Phosphatidylserine synthase 2 OS=Mus musculus OX=10090 GN=Pt | 3 | 1 | 1 | 1 | 473 | 55 | 7,09 | 2,46 | 1 | 0,981 | 1,149 | 1,173 | 0,906 | 1,066 | 1,43 | 0,969 | 0,834 | 0,98 | -0,02 | 8,8E-01 |
| P26040 | Ezrin OS=Mus musculus OX=10090 GN=Ezr PE=1 SV=3 | 23 | 15 | 18 | 8 | 586 | 69,4 | 6,1 | 27,91 | 15 | 0,945 | 0,926 | 0,896 | 0,845 | 0,997 | 0,768 | 0,836 | 0,974 | 1,01 | 0,01 | 8,8E-01 |
| Q61481 | Calcium/calmodulin-dependent 3',5'-cyclic nucleotide phosphodies | 19 | 9 | 10 | 9 | 545 | 62,3 | 5,91 | 14,74 | 9 | 0,792 | 1,033 | 1,023 | 0,975 | 0,876 | 1,015 | 0,979 | 0,993 | 0,99 | -0,01 | 8,8E-01 |
| Q80YF9 | Rho GTPase-activating protein 33 OS=Mus musculus OX=10090 GN | 3 | 2 | 2 | 2 | 1305 | 139,7 | 9,19 | 2,57 | 2 | 1,21 | 1,082 | 1,047 | 0,914 | 0,947 | 1,244 | 0,827 | 1,164 | 1,02 | 0,02 | 8,8E-01 |
| Q61616 | D(1A) dopamine receptor OS=Mus musculus OX=10090 GN=Drd1 P | 3 | 1 | 1 | 1 | 446 | 49,6 | 6,89 | 0 | 1 | 1,698 | 0,602 | 1,286 | 1,071 | 1,182 | 1,12 | 1,308 | 1,191 | 0,97 | -0,03 | 8,8E-01 |
| P22723 | Gamma-aminobutyric acid receptor subunit gamma-2 OS=Mus mu | 4 | 2 | 2 | 2 | 474 | 55,1 | 8,66 | 6,2 | 2 | 0,853 | 0,811 | 0,903 | 0,977 | 0,913 | 0,824 | 0,916 | 0,865 | 1,01 | 0,01 | 8,8E-01 |
| Q922Q8 | Leucine-rich repeat-containing protein 59 OS=Mus musculus OX=1 | 7 | 2 | 2 | 2 | 307 | 34,9 | 9,52 | 5,4 | 2 | 1,097 | 0,989 | 1,135 | 1,139 | 1,042 | 1,102 | 1,102 | 1,139 | 0,99 | -0,01 | 8,8E-01 |
| Q9DB60 | Prostamide/prostaglandin F synthase OS=Mus musculus OX=10090 | 8 | 2 | 2 | 2 | 201 | 21,7 | 6,74 | 2,07 | 2 | 0,756 | 0,555 | 0,691 | 0,861 | 0,718 | 0,777 | 0,741 | 0,669 | 0,99 | -0,01 | 8,8E-01 |
| Q80V03 | Uncharacterized aarF domain-containing protein kinase 5 OS=Mus | 2 | 1 | 1 | 1 | 582 | 66,6 | 8,76 | 0 | 1 | 0,565 | 1,348 | 1,158 | 0,992 | 1,04 | 1,237 | 1,03 | 0,869 | 0,97 | -0,03 | 8,8E-01 |
| Q922B2 | Aspartate--tRNA ligase, cytoplasmic OS=Mus musculus OX=10090 G | 32 | 15 | 17 | 15 | 501 | 57,1 | 6,49 | 27,08 | 15 | 1,141 | 0,98 | 1 | 0,932 | 0,995 | 1,098 | 0,961 | 1,032 | 0,99 | -0,01 | 8,8E-01 |
| Q8BVU5 | ADP-ribose pyrophosphatase, mitochondrial OS=Mus musculus OX | 4 | 2 | 2 | 2 | 350 | 38,6 | 6,76 | 2,19 | 2 | 0,964 | 1,15 | 1,612 | 0,883 | 0,779 | 1,62 | 1,072 | 1,286 | 0,97 | -0,03 | 8,8E-01 |
| Q9QWR8 | Alpha-N-acetylgalactosaminidase OS=Mus musculus OX=10090 GN | 7 | 2 | 2 | 2 | 415 | 47,2 | 6,44 | 2,59 | 2 | 0,885 | 1,047 | 0,999 | 0,948 | 1,018 | 1,02 | 0,933 | 0,934 | 0,99 | -0,01 | 8,8E-01 |
| Q9EQP2 | EH domain-containing protein 4 OS=Mus musculus OX=10090 GN= | 9 | 4 | 5 | 3 | 541 | 61,4 | 6,76 | 7,84 | 4 | 1,166 | 1,212 | 1,207 | 0,964 | 1,14 | 1,282 | 1,173 | 1,004 | 0,99 | -0,01 | 8,8E-01 |
| Q8BKZ9 | Pyruvate dehydrogenase protein X component, mitochondrial OS= | 27 | 10 | 15 | 10 | 501 | 54 | 7,75 | 37,4 | 10 | 1,143 | 1,084 | 1,101 | 0,948 | 0,967 | 1,169 | 1,057 | 1,047 | 1,01 | 0,01 | 8,8E-01 |
| Q8JZQ9 | Eukaryotic translation initiation factor 3 subunit B OS=Mus musculu | 6 | 4 | 4 | 4 | 803 | 91,3 | 5,02 | 11,74 | 4 | 1,097 | 0,846 | 0,953 | 0,91 | 0,844 | 1,126 | 0,812 | 0,97 | 1,01 | 0,01 | 8,8E-01 |
| Q62318 | Transcription intermediary factor 1-beta OS=Mus musculus OX=10 | 5 | 3 | 3 | 3 | 834 | 88,8 | 5,77 | 2,14 | 3 | 2,061 | 1,057 | 1,243 | 1,27 | 0,967 | 1,999 | 1,194 | 1,28 | 1,04 | 0,04 | 8,8E-01 |
| Q9Z2C9 | Myotubularin-related protein 7 OS=Mus musculus OX=10090 GN= | 5 | 3 | 4 | 3 | 660 | 75,6 | 6,43 | 4,77 | 3 | 0,678 | 1,047 | 1,073 | 1,049 | 1,299 | 0,711 | 0,905 | 1,026 | 0,98 | -0,02 | 8,8E-01 |
| Q99KX1 | Myeloid leukemia factor 2 OS=Mus musculus OX=10090 GN=Mlf2 F | 4 | 1 | 1 | 1 | 247 | 28 | 6,98 | 3,31 | 1 | 1 | 0,84 | 0,963 | 0,959 | 1,06 | 0,78 | 0,882 | 1,089 | 0,99 | -0,01 | 8,9E-01 |
| O88851 | Putative hydrolase RBBP9 OS=Mus musculus OX=10090 GN=Rbbp9 | 16 | 3 | 3 | 3 | 186 | 20,9 | 5,97 | 7,72 | 3 | 0,785 | 0,718 | 1,051 | 0,965 | 0,841 | 0,936 | 0,877 | 0,816 | 1,01 | 0,01 | 8,9E-01 |
| P62317 | Small nuclear ribonucleoprotein Sm D2 OS=Mus musculus OX=1009 | 8 | 1 | 1 | 1 | 118 | 13,5 | 9,91 | 0 | 1 | 1,106 | 1,039 | 0,932 | 0,742 | 0,848 | 1,349 | 0,757 | 0,956 | 0,98 | -0,02 | 8,9E-01 |
| Q9JHU9 | Inositol-3-phosphate synthase 1 OS=Mus musculus OX=10090 GN= | 8 | 5 | 5 | 5 | 557 | 60,9 | 6,42 | 10,87 | 5 | 0,906 | 0,975 | 1,05 | 0,926 | 0,944 | 0,941 | 0,982 | 0,97 | 1,01 | 0,01 | 8,9E-01 |
| Q9CQS4 | Solute carrier family 25 member 46 OS=Mus musculus OX=10090 G | 11 | 5 | 6 | 5 | 418 | 46,2 | 7,64 | 12,51 | 5 | 0,917 | 1,104 | 1,022 | 0,988 | 0,973 | 1,069 | 0,965 | 0,997 | 1,01 | 0,01 | 8,9E-01 |
| Q8CIE6 | Coatomer subunit alpha OS=Mus musculus OX=10090 GN=Copa PE | 6 | 6 | 8 | 6 | 1224 | 138,3 | 7,65 | 13,17 | 6 | 1,171 | 0,986 | 1,066 | 0,915 | 0,977 | 1,194 | 1,001 | 0,918 | 1,01 | 0,01 | 8,9E-01 |
| P50544 | Very long-chain specific acyl-CoA dehydrogenase, mitochondrial OS | 15 | 8 | 8 | 8 | 656 | 70,8 | 8,75 | 11,16 | 8 | 1,151 | 0,999 | 1,196 | 1,008 | 0,952 | 1,203 | 1,112 | 1,044 | 1,01 | 0,01 | 8,9E-01 |
| Q9WVE8 | Protein kinase C and casein kinase substrate in neurons protein 2 C | 17 | 9 | 12 | 8 | 486 | 55,8 | 5,2 | 30,86 | 9 | 0,912 | 1,013 | 0,979 | 0,975 | 0,924 | 0,989 | 0,979 | 1,003 | 1,00 | 0,00 | 8,9E-01 |
| P97441 | Zinc transporter 3 OS=Mus musculus OX=10090 GN=Slc30a3 PE=1 S | 17 | 4 | 5 | 4 | 388 | 41,8 | 6,38 | 12,34 | 4 | 1,279 | 1,254 | 1,147 | 0,997 | 0,954 | 1,321 | 1,094 | 1,247 | 1,01 | 0,01 | 8,9E-01 |
| Q9D0I4 | Syntaxin-17 OS=Mus musculus OX=10090 GN=Stx17 PE=1 SV=1 | 7 | 2 | 2 | 2 | 301 | 33,2 | 6,74 | 5,42 | 2 | 0,99 | 1,075 | 1,023 | 0,785 | 0,946 | 0,904 | 0,906 | 1,073 | 1,01 | 0,01 | 8,9E-01 |
| Q9QWI6 | SRC kinase signaling inhibitor 1 OS=Mus musculus OX=10090 GN=S | 34 | 38 | 46 | 38 | 1250 | 134,8 | 9,32 | 80,73 | 38 | 0,849 | 0,997 | 0,943 | 1,009 | 0,938 | 0,865 | 0,99 | 0,978 | 1,01 | 0,01 | 8,9E-01 |
| Q68FE6 | Rho family-interacting cell polarization regulator 1 OS=Mus muscul | 1 | 1 | 1 | 1 | 1223 | 132,3 | 5,88 | 2,27 | 1 | 0,745 | 0,97 | 0,824 | 0,999 | 0,689 | 1,017 | 0,901 | 0,878 | 1,02 | 0,02 | 8,9E-01 |
| Q8BRN9 | Coiled-coil and C2 domain-containing protein 1B OS=Mus musculus | 2 | 2 | 2 | 2 | 848 | 93 | 5,41 | 3,8 | 2 | 1,238 | 0,905 | 1,177 | 0,693 | 0,745 | 1,17 | 1,069 | 0,938 | 1,02 | 0,02 | 8,9E-01 |
| Q6A4J8 | Ubiquitin carboxyl-terminal hydrolase 7 OS=Mus musculus OX=100 | 7 | 5 | 5 | 5 | 1103 | 128,4 | 5,5 | 5,39 | 5 | 1,013 | 1,186 | 0,863 | 1,149 | 1,034 | 1,153 | 1,076 | 0,995 | 0,99 | -0,01 | 8,9E-01 |
| Q80U23 | Syntaphilin OS=Mus musculus OX=10090 GN=Snph PE=1 SV=3 | 15 | 8 | 9 | 8 | 495 | 53,7 | 5,9 | 18,23 | 8 | 1,032 | 1,03 | 1,058 | 1,05 | 0,984 | 1,088 | 0,969 | 1,108 | 1,01 | 0,01 | 8,9E-01 |
| Q9JLJ2 | 4-trimethylaminobutyraldehyde dehydrogenase OS=Mus musculus | 15 | 6 | 7 | 6 | 494 | 53,5 | 6,98 | 19,06 | 6 | 1,026 | 0,974 | 1 | 0,863 | 0,845 | 1,021 | 0,963 | 1,003 | 1,01 | 0,01 | 8,9E-01 |
| Q91XM9 | Disks large homolog 2 OS=Mus musculus OX=10090 GN=Dlg2 PE=1 | 31 | 22 | 29 | 18 | 852 | 94,8 | 6,24 | 70,38 | 22 | 0,951 | 0,935 | 0,953 | 1,053 | 1,01 | 0,925 | 0,995 | 0,981 | 1,00 | 0,00 | 8,9E-01 |
| Q9D020 | Cytosolic 5'-nucleotidase 3A OS=Mus musculus OX=10090 GN=Nt5 | 7 | 2 | 3 | 2 | 331 | 37,2 | 6,65 | 5,24 | 2 | 1,129 | 1,279 | 1,058 | 1,011 | 0,94 | 1,315 | 1,007 | 1,156 | 1,01 | 0,01 | 8,9E-01 |
| P48962 | ADP/ATP translocase 1 OS=Mus musculus OX=10090 GN=Slc25a4 P | 39 | 15 | 36 | 9 | 298 | 32,9 | 9,72 | 76,93 | 15 | 0,929 | 1,121 | 0,992 | 0,986 | 1,037 | 1,085 | 0,953 | 0,921 | 1,01 | 0,01 | 8,9E-01 |
| Q8CI43 | Myosin light chain 6B OS=Mus musculus OX=10090 GN=Myl6b PE= | 20 | 4 | 4 | 2 | 207 | 22,7 | 5,53 | 5,39 | 4 | 1,084 | 0,956 | 1,176 | 1,45 | 1,144 | 1,613 | 1,073 | 0,94 | 0,98 | -0,02 | 8,9E-01 |
| P08752 | Guanine nucleotide-binding protein G(i) subunit alpha-2 OS=Mus m | 35 | 10 | 14 | 7 | 355 | 40,5 | 5,45 | 35,62 | 10 | 0,927 | 0,87 | 0,974 | 0,81 | 0,819 | 0,943 | 0,844 | 0,947 | 1,01 | 0,01 | 8,9E-01 |
| Q8BH74 | Nuclear pore complex protein Nup107 OS=Mus musculus OX=1009 | 2 | 2 | 3 | 2 | 926 | 106,7 | 5,43 | 2,34 | 2 | 1,428 | 2,435 | 1,038 | 0,995 | 1,056 | 1,731 | 1,215 | 1,679 | 1,04 | 0,04 | 8,9E-01 |
| Q5DTY9 | BTB/POZ domain-containing protein KCTD16 OS=Mus musculus OX | 11 | 7 | 10 | 7 | 427 | 48,9 | 8,19 | 7,54 | 7 | 1,047 | 1,067 | 1,021 | 1,086 | 0,985 | 1,109 | 0,971 | 1,131 | 1,01 | 0,01 | 8,9E-01 |
| O70433 | Four and a half LIM domains protein 2 OS=Mus musculus OX=1009 | 10 | 2 | 2 | 2 | 279 | 32,1 | 7,3 | 3,19 | 2 | 1,207 | 0,972 | 1,124 | 1,262 | 0,767 | 1,027 | 1,389 | 1,294 | 1,02 | 0,02 | 8,9E-01 |
| P16045 | Galectin-1 OS=Mus musculus OX=10090 GN=Lgals1 PE=1 SV=3 | 11 | 1 | 1 | 1 | 135 | 14,9 | 5,49 | 2,78 | 1 | 1,512 | 0,863 | 1,272 | 1,19 | 1,526 | 1,194 | 0,942 | 1,07 | 1,02 | 0,02 | 8,9E-01 |
| P55302 | Alpha-2-macroglobulin receptor-associated protein OS=Mus muscu | 9 | 3 | 3 | 3 | 360 | 42,2 | 7,87 | 4,52 | 3 | 0,927 | 0,951 | 0,869 | 1,115 | 0,994 | 0,912 | 0,966 | 1,023 | 0,99 | -0,01 | 8,9E-01 |
| Q9DCB8 | Iron-sulfur cluster assembly 2 homolog, mitochondrial OS=Mus mu | 30 | 4 | 6 | 4 | 154 | 16,7 | 5,6 | 12,6 | 4 | 1,435 | 1,307 | 0,978 | 1,134 | 1,146 | 1,321 | 1,217 | 1,107 | 1,01 | 0,01 | 8,9E-01 |
| Q9ER88 | 28S ribosomal protein S29, mitochondrial OS=Mus musculus OX=1 | 9 | 3 | 3 | 3 | 391 | 44,7 | 8,94 | 4,53 | 3 | 0,964 | 0,908 | 1,194 | 1,023 | 1,079 | 1,143 | 0,957 | 0,954 | 0,99 | -0,01 | 8,9E-01 |
| Q8BK08 | Transmembrane protein 11, mitochondrial OS=Mus musculus OX=1 | 13 | 2 | 2 | 2 | 190 | 21,3 | 7,36 | 5,79 | 2 | 1,204 | 1,055 | 0,924 | 1,17 | 1,021 | 1,066 | 1,026 | 1,291 | 0,99 | -0,01 | 8,9E-01 |
| Q9WVJ2 | 26S proteasome non-ATPase regulatory subunit 13 OS=Mus muscu | 18 | 7 | 8 | 7 | 376 | 42,8 | 5,71 | 12,78 | 7 | 0,968 | 0,981 | 1,053 | 0,955 | 0,975 | 1,026 | 1,043 | 0,89 | 1,01 | 0,01 | 8,9E-01 |
| Q62433 | Protein NDRG1 OS=Mus musculus OX=10090 GN=Ndrg1 PE=1 SV=1 | 11 | 3 | 3 | 3 | 394 | 43 | 6,1 | 5,07 | 3 | 1,355 | 1,005 | 1,163 | 0,858 | 0,907 | 1,234 | 1,077 | 1,092 | 1,02 | 0,02 | 8,9E-01 |
| Q9CQ00 | Distal membrane-arm assembly complex protein 1 OS=Mus muscul | 13 | 1 | 1 | 1 | 111 | 11,3 | 9,39 | 0 | 1 | 1,316 | 1,503 | 1,356 | 1,046 | 1,501 | 1,589 | 1,162 | 1,059 | 0,98 | -0,02 | 8,9E-01 |
| O54946 | DnaJ homolog subfamily B member 6 OS=Mus musculus OX=10090 | 5 | 2 | 2 | 2 | 365 | 39,8 | 9,36 | 5,24 | 2 | 1,054 | 1,302 | 1,122 | 1,164 | 1,263 | 1,014 | 1,125 | 1,286 | 0,99 | -0,01 | 8,9E-01 |
| Q9CXI0 | 2-methoxy-6-polyprenyl-1,4-benzoquinol methylase, mitochondria | 12 | 3 | 3 | 3 | 327 | 37,3 | 7,49 | 2,46 | 3 | 0,918 | 0,971 | 0,836 | 0,719 | 0,824 | 0,989 | 0,779 | 0,812 | 1,01 | 0,01 | 8,9E-01 |
| Q8BFZ9 | Erlin-2 OS=Mus musculus OX=10090 GN=Erlin2 PE=1 SV=1 | 21 | 7 | 8 | 7 | 340 | 37,8 | 5,5 | 22,72 | 7 | 1,071 | 1,032 | 1,056 | 1,003 | 1,044 | 1,144 | 0,984 | 1,011 | 0,99 | -0,01 | 8,9E-01 |
| Q9JKD3 | Secretory carrier-associated membrane protein 5 OS=Mus musculu | 4 | 1 | 7 | 1 | 235 | 26,1 | 8,54 | 13,61 | 1 | 0,794 | 0,975 | 0,962 | 0,971 | 0,979 | 0,828 | 0,966 | 0,96 | 0,99 | -0,01 | 9,0E-01 |
| Q3UYC0 | Protein phosphatase 1H OS=Mus musculus OX=10090 GN=Ppm1h F | 29 | 11 | 13 | 11 | 513 | 56,3 | 6,73 | 22,64 | 11 | 1,126 | 0,994 | 1,035 | 0,977 | 0,957 | 1,191 | 0,975 | 1,043 | 0,99 | -0,01 | 9,0E-01 |

| | | | | | | | | | | | | | | | | | | | | | |
|---|---|---|---|---|---|---|---|---|---|---|---|---|---|---|---|---|---|---|---|---|---|
| Q80U56 | Late secretory pathway protein AVL9 homolog OS=Mus musculus C | 9 | 5 | 6 | 5 | 649 | 72,1 | 6,14 | 12,16 | 5 | 1,171 | 1,138 | 1,019 | 0,944 | 1,034 | 1,365 | 1,017 | 0,916 | 0,99 | -0,01 | 9,0E-01 |
| Q8CDA1 | Phosphatidylinositide phosphatase SAC2 OS=Mus musculus OX=10( | 1 | 1 | 1 | 1 | 1132 | 127,5 | 7,11 | 2,39 | 1 | 1,552 | 1,094 | 0,758 | 0,849 | 1,182 | 1,269 | 0,772 | 1,143 | 0,97 | -0,03 | 9,0E-01 |
| Q9WVQ1 | Membrane-associated guanylate kinase, WW and PDZ domain-con | 9 | 10 | 10 | 10 | 1275 | 140,8 | 6,19 | 20,4 | 10 | 0,904 | 0,921 | 0,928 | 1,077 | 0,929 | 0,931 | 1 | 0,994 | 0,99 | -0,01 | 9,0E-01 |
| Q9WUA2 | Phenylalanine--tRNA ligase beta subunit OS=Mus musculus OX=100 | 2 | 1 | 1 | 1 | 589 | 65,7 | 7,12 | 3,35 | 1 | 0,91 | 0,996 | 1,118 | 1,063 | 1,193 | 0,959 | 1 | 0,973 | 0,99 | -0,01 | 9,0E-01 |
| P27671 | Ras-specific guanine nucleotide-releasing factor 1 OS=Mus musculu | 6 | 6 | 6 | 6 | 1262 | 144 | 7,17 | 8,8 | 6 | 1,077 | 0,974 | 0,884 | 1,143 | 1,038 | 1,025 | 1,05 | 0,996 | 0,99 | -0,01 | 9,0E-01 |
| P62962 | Profilin-1 OS=Mus musculus OX=10090 GN=Pfn1 PE=1 SV=2 | 36 | 5 | 10 | 5 | 140 | 14,9 | 8,28 | 25,51 | 5 | 0,881 | 0,818 | 1,088 | 1,047 | 0,906 | 0,947 | 1,023 | 0,995 | 0,99 | -0,01 | 9,0E-01 |
| P61961 | Ubiquitin-fold modifier 1 OS=Mus musculus OX=10090 GN=Ufm1 P | 18 | 1 | 1 | 1 | 85 | 9,1 | 9,31 | 0 | 1 | 0,757 | 0,913 | 0,702 | 0,804 | 0,787 | 1,054 | 0,688 | 0,589 | 1,02 | 0,02 | 9,0E-01 |
| P70372 | ELAV-like protein 1 OS=Mus musculus OX=10090 GN=Elavl1 PE=1 S | 9 | 3 | 3 | 2 | 326 | 36,1 | 9,04 | 4,44 | 3 | 1,299 | 1,037 | 0,98 | 0,98 | 1,159 | 1,133 | 1,026 | 0,929 | 1,01 | 0,01 | 9,0E-01 |
| Q9QUI0 | Transforming protein RhoA OS=Mus musculus OX=10090 GN=Rhoa | 40 | 8 | 12 | 7 | 193 | 21,8 | 6,1 | 29,58 | 8 | 0,814 | 1,065 | 0,996 | 0,9 | 0,903 | 0,99 | 0,983 | 0,93 | 0,99 | -0,01 | 9,0E-01 |
| Q8CCM6-2 | Isoform 2 of Mitochondrial import inner membrane translocase sub | 3 | 1 | 1 | 1 | 244 | 27,9 | 9,95 | 1,7 | 1 | 0,732 | 0,702 | 0,78 | 0,846 | 0,872 | 0,53 | 0,839 | 0,775 | 1,01 | 0,01 | 9,0E-01 |
| Q9JJV2 | Profilin-2 OS=Mus musculus OX=10090 GN=Pfn2 PE=1 SV=3 | 42 | 5 | 8 | 5 | 140 | 15 | 6,99 | 11,24 | 5 | 0,974 | 1,055 | 1,092 | 0,991 | 0,914 | 1,016 | 1,248 | 0,975 | 0,99 | -0,01 | 9,0E-01 |
| P62500 | TSC22 domain family protein 1 OS=Mus musculus OX=10090 GN=T: | 2 | 2 | 2 | 1 | 1077 | 109,7 | 5,83 | 6,43 | 2 | 0,982 | 0,949 | 0,901 | 1,168 | 1,031 | 0,944 | 1,006 | 1,052 | 0,99 | -0,01 | 9,0E-01 |
| P70392 | Ras-specific guanine nucleotide-releasing factor 2 OS=Mus musculu | 9 | 12 | 14 | 12 | 1189 | 135,6 | 7,85 | 23,35 | 12 | 0,933 | 0,991 | 1,038 | 1,057 | 1,113 | 0,978 | 0,942 | 1,01 | 0,99 | -0,01 | 9,0E-01 |
| Q810U4-3 | Isoform 3 of Neuronal cell adhesion molecule OS=Mus musculus O) | 10 | 11 | 12 | 8 | 1260 | 139 | 5,92 | 29,02 | 11 | 1,092 | 0,911 | 1,134 | 1,145 | 1,044 | 1,009 | 1,206 | 1,059 | 0,99 | -0,01 | 9,0E-01 |
| Q99104 | Unconventional myosin-Va OS=Mus musculus OX=10090 GN=Myo5 | 36 | 68 | 99 | 68 | 1853 | 215,4 | 8,63 | 174,52 | 68 | 1,09 | 1,054 | 1,017 | 0,997 | 0,983 | 1,096 | 1,043 | 1,052 | 1,00 | 0,00 | 9,0E-01 |
| Q9CYG7 | Mitochondrial import receptor subunit TOM34 OS=Mus musculus C | 10 | 3 | 4 | 3 | 309 | 34,3 | 9,14 | 5,08 | 3 | 0,86 | 1,06 | 1,062 | 1,049 | 1,021 | 0,948 | 1,034 | 1,001 | 1,01 | 0,01 | 9,0E-01 |
| Q60598 | Src substrate cortactin OS=Mus musculus OX=10090 GN=Cttn PE=1 | 20 | 11 | 13 | 11 | 546 | 61,2 | 5,4 | 28,1 | 11 | 0,887 | 1,099 | 0,936 | 1,008 | 0,996 | 0,885 | 1,032 | 1,047 | 0,99 | -0,01 | 9,0E-01 |
| Q9CQF8 | Ribosomal protein 63, mitochondrial OS=Mus musculus OX=10090 | 27 | 3 | 3 | 3 | 102 | 11,9 | 10,15 | 4,64 | 3 | 0,933 | 0,887 | 1,026 | 0,816 | 1,015 | 0,896 | 0,865 | 0,857 | 1,01 | 0,01 | 9,0E-01 |
| Q6NVF0 | Inositol polyphosphate 5-phosphatase OCRL OS=Mus musculus OX= | 6 | 5 | 6 | 5 | 900 | 104,2 | 6,27 | 10,06 | 5 | 1,129 | 1,089 | 1,048 | 0,973 | 1,064 | 1,17 | 1,01 | 1,02 | 0,99 | -0,01 | 9,0E-01 |
| P63046 | Sulfotransferase 4A1 OS=Mus musculus OX=10090 GN=Sult4a1 PE= | 3 | 1 | 1 | 1 | 284 | 33 | 5,53 | 2,72 | 1 | 0,847 | 0,9 | 1,024 | 1,084 | 1,132 | 0,842 | 0,982 | 0,94 | 0,99 | -0,01 | 9,0E-01 |
| O35963 | Ras-related protein Rab-33B OS=Mus musculus OX=10090 GN=Rab | 23 | 4 | 5 | 3 | 229 | 25,8 | 7,69 | 8,8 | 4 | 1,223 | 1,019 | 1,017 | 1,235 | 0,918 | 1,232 | 1,12 | 1,178 | 1,01 | 0,01 | 9,0E-01 |
| Q3TY86 | Apoptosis-inducing factor 3 OS=Mus musculus OX=10090 GN=Aifm | 3 | 2 | 2 | 2 | 605 | 66,8 | 9,04 | 3,9 | 2 | 0,979 | 0,854 | 0,62 | 0,608 | 0,57 | 0,708 | 0,923 | 0,923 | 0,98 | -0,02 | 9,0E-01 |
| P61804 | Dolichyl-diphosphooligosaccharide--protein glycosyltransferase sut | 11 | 1 | 1 | 1 | 113 | 12,5 | 7,08 | 0 | 1 | 1,558 | 1,213 | 1,648 | 1,161 | 1,059 | 1,885 | 1,316 | 1,211 | 1,02 | 0,02 | 9,0E-01 |
| P68254 | 14-3-3 protein theta OS=Mus musculus OX=10090 GN=Ywhaq PE=1 | 51 | 16 | 33 | 9 | 245 | 27,8 | 4,78 | 64,75 | 16 | 0,901 | 0,788 | 1,013 | 1,087 | 0,923 | 0,958 | 0,947 | 0,928 | 1,01 | 0,01 | 9,0E-01 |
| Q8BYR5 | Calcium-dependent secretion activator 2 OS=Mus musculus OX=10 | 14 | 16 | 17 | 10 | 1297 | 147,7 | 6,14 | 34,86 | 16 | 0,935 | 1,049 | 1,151 | 1,103 | 0,923 | 1,126 | 1,017 | 1,138 | 1,01 | 0,01 | 9,1E-01 |
| O08672 | Kinesin-like protein KIFC2 OS=Mus musculus OX=10090 GN=Kifc2 P | 2 | 1 | 1 | 1 | 792 | 85,6 | 6,16 | 2,05 | 1 | 0,574 | 1,12 | 1,156 | 1,125 | 1,182 | 1,113 | 0,882 | 0,877 | 0,98 | -0,02 | 9,1E-01 |
| Q8BFP9 | [Pyruvate dehydrogenase (acetyl-transferring)] kinase isozyme 1, m | 10 | 4 | 4 | 3 | 434 | 49 | 8,19 | 7,12 | 4 | 1,057 | 1,02 | 1,007 | 1,144 | 0,928 | 1,13 | 1,179 | 0,957 | 1,01 | 0,01 | 9,1E-01 |
| P18052 | Receptor-type tyrosine-protein phosphatase alpha OS=Mus muscul | 11 | 10 | 10 | 9 | 829 | 93,6 | 7,05 | 16,77 | 10 | 1,24 | 1,002 | 1,052 | 1,004 | 1,043 | 1,226 | 1,026 | 1,039 | 0,99 | -0,01 | 9,1E-01 |
| A2AN08 | E3 ubiquitin-protein ligase UBR4 OS=Mus musculus OX=10090 GN= | 4 | 19 | 21 | 19 | 5180 | 571,9 | 6,06 | 28,74 | 19 | 1,145 | 1,152 | 1,046 | 1,001 | 1,018 | 1,117 | 1,093 | 1,095 | 1,00 | 0,00 | 9,1E-01 |
| Q91WC3 | Long-chain-fatty-acid--CoA ligase 6 OS=Mus musculus OX=10090 G | 26 | 16 | 18 | 16 | 697 | 78 | 7,34 | 45,4 | 16 | 1,009 | 0,964 | 1,07 | 1,012 | 1,038 | 1,069 | 0,953 | 1,011 | 1,00 | 0,00 | 9,1E-01 |
| Q9CQU5 | ZW10 interactor OS=Mus musculus OX=10090 GN=Zwint PE=1 SV=: | 5 | 1 | 1 | 1 | 252 | 28,7 | 8,43 | 3,06 | 1 | 1,294 | 1,212 | 1,136 | 1,399 | 1,119 | 1,083 | 1,226 | 1,686 | 0,99 | -0,01 | 9,1E-01 |
| Q61686 | Chromobox protein homolog 5 OS=Mus musculus OX=10090 GN=C | 7 | 1 | 1 | 1 | 191 | 22,2 | 5,86 | 0 | 1 | 1,358 | 1,091 | 0,525 | 0,756 | 1,146 | 0,568 | 0,954 | 1,171 | 0,97 | -0,03 | 9,1E-01 |
| Q60673 | Receptor-type tyrosine-protein phosphatase-like N OS=Mus muscu | 5 | 3 | 3 | 2 | 979 | 106 | 7,09 | 4,66 | 3 | 1,215 | 1,561 | 1,186 | 0,884 | 0,875 | 1,693 | 0,892 | 1,274 | 1,02 | 0,02 | 9,1E-01 |
| O08739 | AMP deaminase 3 OS=Mus musculus OX=10090 GN=Ampd3 PE=1 S | 4 | 3 | 3 | 3 | 766 | 88,6 | 7,33 | 4,85 | 3 | 1,169 | 1,34 | 1,119 | 1,251 | 1,038 | 1,499 | 1,164 | 1,229 | 0,99 | -0,01 | 9,1E-01 |
| P08226 | Apolipoprotein E OS=Mus musculus OX=10090 GN=Apoe PE=1 SV=: | 19 | 5 | 6 | 5 | 311 | 35,8 | 5,68 | 15,71 | 5 | 0,902 | 0,931 | 0,968 | 1,015 | 0,986 | 0,884 | 1,044 | 0,922 | 0,99 | -0,01 | 9,1E-01 |
| Q9JHU4 | Cytoplasmic dynein 1 heavy chain 1 OS=Mus musculus OX=10090 G | 27 | 128 | 182 | 128 | 4644 | 531,7 | 6,42 | 386,78 | 128 | 0,999 | 1 | 1,043 | 1,044 | 1,014 | 0,989 | 1,042 | 1,033 | 1,00 | 0,00 | 9,1E-01 |
| Q9CQB5 | CDGSH iron-sulfur domain-containing protein 2 OS=Mus musculus ( | 24 | 3 | 6 | 3 | 135 | 15,2 | 9,51 | 23,5 | 3 | 0,727 | 1,115 | 0,849 | 1,174 | 0,948 | 0,789 | 1,423 | 0,79 | 0,98 | -0,02 | 9,1E-01 |
| Q9DCL9 | Multifunctional protein ADE2 OS=Mus musculus OX=10090 GN=Pai | 17 | 6 | 6 | 6 | 425 | 47 | 7,23 | 10,61 | 6 | 0,83 | 0,879 | 1,08 | 0,995 | 0,976 | 0,981 | 0,922 | 0,932 | 0,99 | -0,01 | 9,1E-01 |
| B1AZA5 | Transmembrane protein 245 OS=Mus musculus OX=10090 GN=Tm | 1 | 1 | 1 | 1 | 876 | 97,3 | 8,75 | 2,04 | 1 | 1,011 | 0,859 | 1,087 | 0,784 | 1,069 | 0,988 | 0,806 | 0,835 | 1,01 | 0,01 | 9,1E-01 |
| P56382 | ATP synthase subunit epsilon, mitochondrial OS=Mus musculus OX | 31 | 2 | 5 | 2 | 52 | 5,8 | 10,01 | 12,07 | 2 | 0,877 | 1,088 | 0,865 | 0,843 | 1,066 | 0,869 | 0,796 | 0,905 | 1,01 | 0,01 | 9,1E-01 |
| Q9QUR6 | Prolyl endopeptidase OS=Mus musculus OX=10090 GN=Prep PE=1 S | 14 | 9 | 11 | 9 | 710 | 80,7 | 5,73 | 9,94 | 9 | 0,868 | 0,906 | 1,052 | 1,14 | 0,999 | 0,878 | 1,085 | 0,969 | 1,01 | 0,01 | 9,1E-01 |
| Q810U4-5 | Isoform 5 of Neuronal cell adhesion molecule OS=Mus musculus O) | 17 | 4 | 4 | 1 | 260 | 28,9 | 8,43 | 8,74 | 4 | 1,873 | 1,467 | 1,481 | 0,964 | 0,952 | 1,513 | 1,447 | 1,758 | 1,02 | 0,02 | 9,1E-01 |
| P70175 | Disks large homolog 3 OS=Mus musculus OX=10090 GN=Dlg3 PE=1 | 31 | 24 | 28 | 23 | 849 | 93,4 | 6,79 | 52,63 | 24 | 1,114 | 1,011 | 1,047 | 0,997 | 0,928 | 1,095 | 1,098 | 1,07 | 0,99 | -0,01 | 9,1E-01 |
| Q9D0F9 | Phosphoglucomutase-1 OS=Mus musculus OX=10090 GN=Pgm1 PE | 31 | 16 | 20 | 16 | 562 | 61,4 | 6,57 | 42,92 | 16 | 0,982 | 0,958 | 0,987 | 1 | 0,975 | 0,919 | 1,03 | 1,015 | 1,00 | 0,00 | 9,1E-01 |
| Q924L1 | LETM1 domain-containing protein 1 OS=Mus musculus OX=10090 ( | 9 | 3 | 4 | 3 | 360 | 41,7 | 10,54 | 5,14 | 3 | 1,509 | 1,338 | 1,301 | 1,339 | 1,272 | 1,334 | 1,295 | 1,55 | 1,01 | 0,01 | 9,1E-01 |
| Q9ERG2 | Striatin-3 OS=Mus musculus OX=10090 GN=Strn3 PE=1 SV=1 | 14 | 8 | 8 | 7 | 796 | 87,1 | 5,29 | 19,78 | 8 | 1,053 | 0,97 | 0,992 | 0,977 | 0,963 | 1,013 | 1,069 | 0,931 | 1,00 | 0,00 | 9,1E-01 |
| Q8BX70 | Vacuolar protein sorting-associated protein 13C OS=Mus musculus | 7 | 21 | 22 | 20 | 3748 | 419,8 | 6,81 | 28,05 | 21 | 1,033 | 1,057 | 0,943 | 1,042 | 0,959 | 1,054 | 1,04 | 1,037 | 1,00 | 0,00 | 9,1E-01 |
| Q9D486 | C-Maf-inducing protein OS=Mus musculus OX=10090 GN=Cmip PE= | 1 | 1 | 1 | 1 | 773 | 86,2 | 6,81 | 2,36 | 1 | 1,127 | 0,754 | 1,163 | 1,126 | 1,211 | 1,327 | 0,923 | 0,78 | 0,98 | -0,02 | 9,1E-01 |
| Q8C0L0 | Thioredoxin-related transmembrane protein 4 OS=Mus musculus C | 7 | 2 | 3 | 2 | 335 | 37,1 | 4,37 | 5,86 | 2 | 0,931 | 1,231 | 0,825 | 0,831 | 0,875 | 0,996 | 0,95 | 0,953 | 1,01 | 0,01 | 9,1E-01 |
| O09114 | Prostaglandin-H2 D-isomerase OS=Mus musculus OX=10090 GN=Pt | 5 | 1 | 1 | 1 | 189 | 21,1 | 8,25 | 0 | 1 | 0,651 | 0,935 | 0,922 | 0,73 | 0,917 | 0,746 | 0,723 | 0,815 | 1,01 | 0,01 | 9,1E-01 |
| Q8R035 | Peptidyl-tRNA hydrolase ICT1, mitochondrial OS=Mus musculus OX | 12 | 2 | 2 | 2 | 206 | 23,5 | 10,18 | 5,5 | 2 | 0,933 | 1,161 | 0,996 | 1,096 | 1,066 | 1,238 | 0,891 | 1,03 | 0,99 | -0,01 | 9,2E-01 |
| Q9Z2Y3 | Homer protein homolog 1 OS=Mus musculus OX=10090 GN=Home | 45 | 14 | 16 | 13 | 366 | 41,4 | 5,53 | 36,65 | 14 | 1,013 | 0,933 | 1,072 | 1,074 | 1,087 | 0,912 | 1,097 | 1,02 | 0,99 | -0,01 | 9,2E-01 |
| P97371 | Proteasome activator complex subunit 1 OS=Mus musculus OX=10( | 20 | 4 | 4 | 4 | 249 | 28,7 | 5,97 | 4,91 | 4 | 0,787 | 0,976 | 1,09 | 1,157 | 1,129 | 1,038 | 0,823 | 0,974 | 1,01 | 0,01 | 9,2E-01 |
| Q80YA3 | Phospholipase DDHD1 OS=Mus musculus OX=10090 GN=Ddhd1 PE | 1 | 1 | 1 | 1 | 547 | 61,8 | 6,61 | 2,02 | 1 | 0,665 | 1,203 | 0,603 | 1,055 | 0,979 | 0,797 | 0,97 | 0,848 | 0,98 | -0,02 | 9,2E-01 |
| Q9CQZ5 | NADH dehydrogenase [ubiquinone] 1 alpha subcomplex subunit 6 ( | 39 | 5 | 8 | 5 | 131 | 15,3 | 10,11 | 22,69 | 5 | 1,047 | 1,196 | 1,039 | 0,911 | 0,975 | 1,162 | 1,058 | 1,029 | 0,99 | -0,01 | 9,2E-01 |
| P27659 | 60S ribosomal protein L3 OS=Mus musculus OX=10090 GN=Rpl3 PE | 11 | 5 | 5 | 5 | 403 | 46,1 | 10,21 | 11,8 | 5 | 1,043 | 1,11 | 1,202 | 0,987 | 1,141 | 1,032 | 1,051 | 1,142 | 0,99 | -0,01 | 9,2E-01 |
| E9Q401 | Ryanodine receptor 2 OS=Mus musculus OX=10090 GN=Ryr2 PE=1 | 12 | 45 | 47 | 42 | 4966 | 564,5 | 6,09 | 77,71 | 45 | 1,03 | 1,015 | 1,014 | 1,051 | 0,976 | 1,018 | 1,041 | 1,066 | 1,00 | 0,00 | 9,2E-01 |
| Q9QXG2 | Rab proteins geranylgeranyltransferase component A 1 OS=Mus m | 7 | 5 | 5 | 5 | 665 | 73,9 | 4,68 | 12,55 | 5 | 0,951 | 1,022 | 1,017 | 1,132 | 1,033 | 1,285 | 0,954 | 0,89 | 0,99 | -0,01 | 9,2E-01 |
| Q9QZ06 | Toll-interacting protein OS=Mus musculus OX=10090 GN=Tollip PE= | 10 | 3 | 4 | 3 | 274 | 30,3 | 5,17 | 4,61 | 3 | 0,797 | 0,854 | 1,02 | 0,972 | 0,887 | 0,831 | 1,006 | 0,946 | 0,99 | -0,01 | 9,2E-01 |
| Q9D0J4 | ADP-ribosylation factor-like protein 2 OS=Mus musculus OX=10090 | 11 | 2 | 2 | 2 | 184 | 20,9 | 5,96 | 5,08 | 2 | 1,176 | 1,12 | 1,161 | 0,941 | 0,902 | 1,213 | 0,872 | 1,356 | 1,01 | 0,01 | 9,2E-01 |
| Q3TIR3 | Synembryn-A OS=Mus musculus OX=10090 GN=Ric8a PE=1 SV=2 | 10 | 5 | 6 | 5 | 530 | 59,8 | 5,68 | 6,47 | 5 | 1,29 | 1,338 | 1,043 | 1,093 | 0,968 | 1,776 | 1,023 | 1,082 | 0,98 | -0,02 | 9,2E-01 |
| O55003 | BCL2/adenovirus E1B 19 kDa protein-interacting protein 3 OS=Mus | 6 | 1 | 2 | 1 | 187 | 21 | 7,14 | 4,8 | 1 | 0,632 | 0,953 | 0,964 | 0,938 | 0,889 | 0,87 | 0,806 | 0,958 | 0,99 | -0,01 | 9,2E-01 |

| | | | | | | | | | | | | | | | | | | | | | |
|---|---|---|---|---|---|---|---|---|---|---|---|---|---|---|---|---|---|---|---|---|---|
| Q9Z1B3 | 1-phosphatidylinositol 4,5-bisphosphate phosphodiesterase beta-1 | 20 | 23 | 31 | 23 | 1216 | 138,3 | 6,13 | 81,12 | 23 | 1,005 | 0,936 | 0,979 | 1,039 | 0,983 | 0,969 | 1,014 | 1,003 | 1,00 | 0,00 | 9,2E-01 |
| Q6PFQ7 | Ras GTPase-activating protein 4 OS=Mus musculus OX=10090 GN=R | 1 | 1 | 1 | 1 | 802 | 90 | 8 | 2,23 | 1 | 1,186 | 0,933 | 1,023 | 0,818 | 1,005 | 1,1 | 0,879 | 0,938 | 1,01 | 0,01 | 9,2E-01 |
| Q3UEZ8 | Sodium/bile acid cotransporter 4 OS=Mus musculus OX=10090 GN= | 3 | 1 | 1 | 1 | 437 | 46,6 | 5,01 | 2,11 | 1 | 1,012 | 0,761 | 1,059 | 0,889 | 0,773 | 0,842 | 1,277 | 0,883 | 0,99 | -0,01 | 9,2E-01 |
| Q99PJ0 | Neurotrimin OS=Mus musculus OX=10090 GN=Ntm PE=1 SV=2 | 17 | 5 | 14 | 5 | 344 | 38 | 7,81 | 32,1 | 5 | 1,001 | 0,926 | 1,029 | 1,023 | 1,023 | 0,879 | 1,015 | 1,082 | 0,99 | -0,01 | 9,2E-01 |
| P63276 | 40S ribosomal protein S17 OS=Mus musculus OX=10090 GN=Rps17 | 9 | 1 | 1 | 1 | 135 | 15,5 | 9,85 | 2,73 | 1 | 0,838 | 0,819 | 0,948 | 1,038 | 0,8 | 0,793 | 1,015 | 1,071 | 0,99 | -0,01 | 9,2E-01 |
| D3Z4S3 | Putative peptidyl-tRNA hydrolase PTRHD1 OS=Mus musculus OX=1 | 9 | 1 | 1 | 1 | 140 | 16 | 9,32 | 2,51 | 1 | 1,316 | 0,987 | 1 | 1,055 | 1,259 | 0,997 | 0,935 | 1,212 | 0,99 | -0,01 | 9,2E-01 |
| P97819 | 85/88 kDa calcium-independent phospholipase A2 OS=Mus muscul | 1 | 1 | 1 | 1 | 807 | 89,5 | 7,24 | 2,07 | 1 | 1,738 | 1,022 | 0,686 | 0,7 | 1,188 | 0,726 | 0,899 | 1,223 | 1,03 | 0,03 | 9,2E-01 |
| P28481-4 | Isoform 4 of Collagen alpha-1(II) chain OS=Mus musculus OX=1009 | 1 | 1 | 1 | 1 | 1458 | 138,9 | 7,09 | 1,62 | 1 | 1,248 | 1,305 | 1,009 | 1,456 | 1,131 | 0,923 | 1,246 | 1,802 | 0,98 | -0,02 | 9,2E-01 |
| Q8K2Y7 | 39S ribosomal protein L47, mitochondrial OS=Mus musculus OX=10 | 3 | 1 | 1 | 1 | 252 | 29,7 | 10,21 | 1,95 | 1 | 1,187 | 0,831 | 1,044 | 1,019 | 0,948 | 0,876 | 0,879 | 1,326 | 1,01 | 0,01 | 9,2E-01 |
| Q6ZPJ3 | (E3-independent) E2 ubiquitin-conjugating enzyme UBE2O OS=Mus | 13 | 15 | 15 | 15 | 1288 | 140,7 | 5,06 | 31,32 | 15 | 1,173 | 1,072 | 1,133 | 1,059 | 1,01 | 1,24 | 1,081 | 1,084 | 1,00 | 0,00 | 9,2E-01 |
| Q80YD6 | Secretion-regulating guanine nucleotide exchange factor OS=Mus r | 5 | 2 | 2 | 2 | 464 | 49,2 | 6,19 | 5,4 | 2 | 0,955 | 1,397 | 1,244 | 0,957 | 0,833 | 1,373 | 1,109 | 1,176 | 1,01 | 0,01 | 9,2E-01 |
| Q8VBT9 | Tether containing UBX domain for GLUT4 OS=Mus musculus OX=10 | 11 | 5 | 5 | 5 | 550 | 59,8 | 6,96 | 7,66 | 5 | 1,232 | 0,972 | 1,06 | 1,073 | 1,066 | 1,074 | 1,123 | 1,052 | 1,01 | 0,01 | 9,3E-01 |
| P58242 | Acid sphingomyelinase-like phosphodiesterase 3b OS=Mus musculu | 6 | 2 | 2 | 2 | 456 | 51,6 | 6,43 | 2,28 | 2 | 1,257 | 1,19 | 1,128 | 1,007 | 0,931 | 1,416 | 0,808 | 1,364 | 1,01 | 0,01 | 9,3E-01 |
| Q9QZ88 | Vacuolar protein sorting-associated protein 29 OS=Mus musculus O | 41 | 7 | 7 | 7 | 182 | 20,5 | 6,79 | 13,32 | 7 | 0,936 | 0,968 | 0,989 | 0,936 | 0,98 | 0,969 | 0,995 | 0,895 | 1,00 | 0,00 | 9,3E-01 |
| Q9ESJ4 | NCK-interacting protein with SH3 domain OS=Mus musculus OX=10 | 13 | 8 | 12 | 8 | 714 | 78,5 | 6,05 | 19,81 | 8 | 1,078 | 1,216 | 1,067 | 1,003 | 1,022 | 1,099 | 1,2 | 1,019 | 1,01 | 0,01 | 9,3E-01 |
| Q812G0 | Alpha-1,3-mannosyl-glycoprotein 4-beta-N-acetylglucosaminyltran | 1 | 1 | 1 | 1 | 535 | 61,4 | 7,62 | 0 | 1 | 0,916 | 0,985 | 0,902 | 0,91 | 1,061 | 0,971 | 0,855 | 0,847 | 0,99 | -0,01 | 9,3E-01 |
| O35678 | Monoglyceride lipase OS=Mus musculus OX=10090 GN=Mgll PE=1 S | 21 | 6 | 7 | 6 | 303 | 33,4 | 7,15 | 14,55 | 6 | 1,053 | 1,053 | 1,069 | 1,169 | 1,283 | 0,985 | 1,099 | 0,946 | 1,01 | 0,01 | 9,3E-01 |
| B9EHT4 | CAP-Gly domain-containing linker protein 3 OS=Mus musculus OX= | 5 | 2 | 2 | 2 | 547 | 59,5 | 7,94 | 4,13 | 2 | 1,221 | 1,194 | 0,941 | 1,324 | 1,098 | 1,497 | 0,95 | 1,08 | 1,01 | 0,01 | 9,3E-01 |
| P62806 | Histone H4 OS=Mus musculus OX=10090 GN=Hist1h4a PE=1 SV=2 | 22 | 3 | 9 | 3 | 103 | 11,4 | 11,36 | 19,84 | 3 | 1,36 | 1,42 | 1,043 | 0,745 | 1,057 | 1,593 | 0,867 | 1,135 | 0,98 | -0,02 | 9,3E-01 |
| Q9CPT3 | N-acylneuraminate-9-phosphatase OS=Mus musculus OX=10090 GN | 4 | 1 | 1 | 1 | 248 | 27,8 | 6,07 | 3,08 | 1 | 1,569 | 0,969 | 0,801 | 0,948 | 0,811 | 1,387 | 1,268 | 0,731 | 1,02 | 0,02 | 9,3E-01 |
| Q99N94 | 39S ribosomal protein L9, mitochondrial OS=Mus musculus OX=100 | 5 | 1 | 1 | 1 | 265 | 30,2 | 10,08 | 0 | 1 | 0,891 | 1,082 | 2,284 | 0,96 | 1,235 | 1,469 | 1,433 | 0,947 | 1,03 | 0,03 | 9,3E-01 |
| S4R1M9 | Oxysterol-binding protein-related protein 10 OS=Mus musculus OX | 6 | 4 | 5 | 4 | 766 | 83,8 | 8,28 | 7,31 | 4 | 1,283 | 1,309 | 1,277 | 1,075 | 1,045 | 1,544 | 1,069 | 1,238 | 1,01 | 0,01 | 9,3E-01 |
| Q6PB66 | Leucine-rich PPR motif-containing protein, mitochondrial OS=Mus r | 22 | 29 | 33 | 29 | 1392 | 156,5 | 6,83 | 67,46 | 29 | 1,107 | 1,026 | 1,13 | 1,053 | 1,047 | 1,074 | 1,08 | 1,105 | 1,00 | 0,00 | 9,3E-01 |
| Q9CRB6 | Tubulin polymerization-promoting protein family member 3 OS=Mu | 18 | 3 | 4 | 3 | 176 | 19 | 9,11 | 13,59 | 3 | 0,914 | 0,96 | 1,064 | 1,136 | 1,057 | 0,863 | 1,104 | 1,023 | 1,01 | 0,01 | 9,3E-01 |
| O35551 | Rab GTPase-binding effector protein 1 OS=Mus musculus OX=1009 | 17 | 12 | 14 | 12 | 862 | 99,5 | 5,01 | 29,62 | 12 | 0,966 | 1 | 0,921 | 1,114 | 0,827 | 1,063 | 1,007 | 1,078 | 1,01 | 0,01 | 9,3E-01 |
| Q91WG7 | Diacylglycerol kinase gamma OS=Mus musculus OX=10090 GN=Dgk | 3 | 3 | 3 | 3 | 788 | 88,5 | 6,76 | 5,32 | 3 | 1,047 | 0,962 | 0,978 | 1,084 | 0,894 | 1,012 | 1,092 | 1,093 | 1,00 | 0,00 | 9,3E-01 |
| Q9QZB7 | Actin-related protein 10 OS=Mus musculus OX=10090 GN=Actr10 P | 7 | 4 | 4 | 4 | 417 | 46,2 | 7,61 | 10,32 | 4 | 1,131 | 0,861 | 1,044 | 1,072 | 1,034 | 1,114 | 0,957 | 1,027 | 0,99 | -0,01 | 9,3E-01 |
| Q64523 | Histone H2A type 2-C OS=Mus musculus OX=10090 GN=Hist2h2ac I | 16 | 3 | 9 | 1 | 129 | 14 | 10,9 | 19,88 | 3 | 1,182 | 1,809 | 1,041 | 0,722 | 1,111 | 1,836 | 0,806 | 1,115 | 0,98 | -0,02 | 9,3E-01 |
| Q9CZR8 | Elongation factor Ts, mitochondrial OS=Mus musculus OX=10090 G | 18 | 4 | 4 | 4 | 324 | 35,3 | 7,06 | 8,53 | 4 | 1,085 | 0,755 | 0,865 | 1,106 | 1,085 | 0,805 | 0,961 | 0,923 | 1,01 | 0,01 | 9,3E-01 |
| Q8R2Y0 | Monoacylglycerol lipase ABHD6 OS=Mus musculus OX=10090 GN=A | 10 | 3 | 4 | 3 | 336 | 38,2 | 8,47 | 5,46 | 3 | 0,973 | 1,094 | 1,113 | 1,178 | 1,099 | 1,073 | 1,113 | 1,057 | 1,00 | 0,00 | 9,3E-01 |
| P55264 | Adenosine kinase OS=Mus musculus OX=10090 GN=Adk PE=1 SV=2 | 14 | 4 | 5 | 4 | 361 | 40,1 | 6,21 | 6,13 | 4 | 1,368 | 1,222 | 1,31 | 1,311 | 1,293 | 1,473 | 1,335 | 1,137 | 0,99 | -0,01 | 9,3E-01 |
| Q7TSF4 | Leucine-rich repeat-containing protein 75A OS=Mus musculus OX=1 | 8 | 2 | 2 | 2 | 339 | 37,6 | 8,76 | 0 | 2 | 0,01 | 2,049 | 3,281 | 1,816 | 2,178 | 3,103 | 1,496 | 0,682 | 0,96 | -0,04 | 9,3E-01 |
| Q4QQM4 | Tumor protein p53-inducible protein 11 OS=Mus musculus OX=100 | 8 | 1 | 1 | 1 | 189 | 20,9 | 9,41 | 4,61 | 1 | 1,123 | 0,911 | 0,838 | 0,761 | 0,734 | 1,126 | 0,918 | 0,895 | 0,99 | -0,01 | 9,3E-01 |
| Q64373 | Bcl-2-like protein 1 OS=Mus musculus OX=10090 GN=Bcl2l1 PE=1 S | 4 | 1 | 2 | 1 | 233 | 26,1 | 4,94 | 4,21 | 1 | 0,451 | 1,241 | 0,961 | 1,106 | 1,123 | 0,929 | 0,928 | 0,844 | 0,98 | -0,02 | 9,3E-01 |
| P39053-4 | Isoform 4 of Dynamin-1 OS=Mus musculus OX=10090 GN=Dnm1 | 43 | 42 | 99 | 1 | 864 | 97,4 | 7,17 | 201,45 | 42 | 0,843 | 1,091 | 1,031 | 1,023 | 1,011 | 1,058 | 0,935 | 1,005 | 0,99 | -0,01 | 9,3E-01 |
| Q9EQI8 | 39S ribosomal protein L46, mitochondrial OS=Mus musculus OX=10 | 16 | 4 | 5 | 4 | 283 | 32,1 | 7,4 | 5,03 | 4 | 1,183 | 0,924 | 1,054 | 1,021 | 1,056 | 0,981 | 1,016 | 1,152 | 0,99 | -0,01 | 9,3E-01 |
| Q8JZQ2 | AFG3-like protein 2 OS=Mus musculus OX=10090 GN=Afg3l2 PE=1 S | 21 | 18 | 27 | 18 | 802 | 89,5 | 8,6 | 57,61 | 18 | 0,974 | 1,015 | 1,008 | 0,993 | 0,98 | 1,046 | 0,948 | 1,008 | 1,00 | 0,00 | 9,3E-01 |
| Q9WTR5 | Cadherin-13 OS=Mus musculus OX=10090 GN=Cdh13 PE=1 SV=2 | 14 | 7 | 8 | 7 | 714 | 78,1 | 5,12 | 16,08 | 7 | 0,925 | 1,017 | 1,142 | 1,216 | 1,044 | 1,205 | 0,985 | 1,094 | 0,99 | -0,01 | 9,3E-01 |
| P61255 | 60S ribosomal protein L26 OS=Mus musculus OX=10090 GN=Rpl26 | 12 | 2 | 2 | 2 | 145 | 17,2 | 10,55 | 2,4 | 2 | 1,37 | 1,562 | 1,286 | 1,172 | 1,187 | 1,343 | 1,449 | 1,447 | 0,99 | -0,01 | 9,3E-01 |
| Q3UV70 | [Pyruvate dehydrogenase [acetyl-transferring]]-phosphatase 1, mit | 9 | 3 | 3 | 3 | 538 | 61,1 | 6,67 | 5,01 | 3 | 0,948 | 1,165 | 0,747 | 1,439 | 1,322 | 0,721 | 1,341 | 0,988 | 0,98 | -0,02 | 9,3E-01 |
| Q9D1Q6 | Endoplasmic reticulum resident protein 44 OS=Mus musculus OX=1 | 15 | 5 | 6 | 5 | 406 | 46,8 | 5,27 | 10,39 | 5 | 1,163 | 1,132 | 0,93 | 0,961 | 1,037 | 1,186 | 0,979 | 1,01 | 0,99 | -0,01 | 9,3E-01 |
| Q9CR68 | Cytochrome b-c1 complex subunit Rieske, mitochondrial OS=Mus m | 27 | 7 | 15 | 7 | 274 | 29,3 | 8,7 | 24,66 | 7 | 0,905 | 0,957 | 0,933 | 0,803 | 0,895 | 0,919 | 0,889 | 0,907 | 1,00 | 0,00 | 9,3E-01 |
| Q9JL04 | Formin-2 OS=Mus musculus OX=10090 GN=Fmn2 PE=1 SV=2 | 3 | 4 | 4 | 4 | 1578 | 167,3 | 5,48 | 9,63 | 4 | 1,05 | 1,084 | 1,184 | 1,233 | 1,151 | 1,171 | 1,124 | 1,089 | 1,00 | 0,00 | 9,3E-01 |
| O54950 | 5'-AMP-activated protein kinase subunit gamma-1 OS=Mus muscul | 6 | 2 | 2 | 2 | 330 | 37,5 | 7,14 | 3,2 | 2 | 1,48 | 1,11 | 1,427 | 1,108 | 1,162 | 1,623 | 1,04 | 1,356 | 0,99 | -0,01 | 9,3E-01 |
| Q8C2E7 | WASH complex subunit 5 OS=Mus musculus OX=10090 GN=Washc5 | 5 | 5 | 5 | 5 | 1159 | 134 | 7,12 | 4,36 | 5 | 0,941 | 1,336 | 1,128 | 1,116 | 1,088 | 1,034 | 1,034 | 1,328 | 1,01 | 0,01 | 9,3E-01 |
| Q9WU79 | Proline dehydrogenase 1, mitochondrial OS=Mus musculus OX=100 | 11 | 5 | 5 | 5 | 599 | 68 | 8,24 | 8,3 | 5 | 1,448 | 0,966 | 0,955 | 0,948 | 0,895 | 1,367 | 0,905 | 1,093 | 1,01 | 0,01 | 9,3E-01 |
| O70493 | Sorting nexin-12 OS=Mus musculus OX=10090 GN=Snx12 PE=1 SV= | 21 | 4 | 4 | 4 | 165 | 19,1 | 7,34 | 10,16 | 4 | 1,015 | 1,005 | 0,987 | 1,172 | 1,008 | 0,987 | 1,089 | 1,078 | 1,00 | 0,00 | 9,3E-01 |
| Q924T7 | E3 ubiquitin-protein ligase RNF31 OS=Mus musculus OX=10090 GN | 2 | 2 | 2 | 2 | 1066 | 119,2 | 6,52 | 2,54 | 2 | 1,046 | 1,257 | 0,783 | 0,897 | 1,084 | 0,876 | 1,024 | 1,037 | 0,99 | -0,01 | 9,4E-01 |
| Q91V61 | Sideroflexin-3 OS=Mus musculus OX=10090 GN=Sfxn3 PE=1 SV=1 | 25 | 7 | 10 | 7 | 321 | 35,4 | 9,51 | 34,48 | 7 | 1,104 | 1,125 | 0,924 | 0,988 | 1,007 | 1,053 | 0,98 | 1,083 | 1,00 | 0,00 | 9,4E-01 |
| Q64511 | DNA topoisomerase 2-beta OS=Mus musculus OX=10090 GN=Top2 | 1 | 2 | 2 | 2 | 1612 | 181,8 | 8,29 | 3,65 | 2 | 1,119 | 1,178 | 0,986 | 0,835 | 1,071 | 1,065 | 0,993 | 0,962 | 1,01 | 0,01 | 9,4E-01 |
| Q6PGL7 | WASH complex subunit 2 OS=Mus musculus OX=10090 GN=Washc2 | 3 | 3 | 3 | 3 | 1334 | 145,2 | 4,77 | 4,39 | 3 | 1,333 | 1,045 | 1,093 | 1,088 | 1,161 | 1,228 | 0,999 | 1,144 | 1,01 | 0,01 | 9,4E-01 |
| Q80TJ1-2 | Isoform 2 of Calcium-dependent secretion activator 1 OS=Mus mus | 31 | 34 | 53 | 1 | 1361 | 153,9 | 5,86 | 109,96 | 34 | 1,271 | 1,146 | 0,994 | 1,012 | 0,804 | 1,453 | 0,884 | 1,34 | 0,99 | -0,01 | 9,4E-01 |
| Q9QXS1 | Plectin OS=Mus musculus OX=10090 GN=Plec PE=1 SV=3 | 27 | 122 | 151 | 122 | 4691 | 533,9 | 5,96 | 292,77 | 122 | 1,06 | 1,004 | 1,061 | 1,012 | 1,099 | 0,941 | 1,051 | 1,034 | 1,00 | 0,00 | 9,4E-01 |
| Q9DB72 | BTB/POZ domain-containing protein 17 OS=Mus musculus OX=1009 | 4 | 2 | 2 | 2 | 478 | 52,6 | 9,42 | 4,45 | 2 | 1,139 | 0,556 | 1,016 | 0,884 | 1,045 | 0,787 | 0,797 | 1,013 | 0,99 | -0,01 | 9,4E-01 |
| P06151 | L-lactate dehydrogenase A chain OS=Mus musculus OX=10090 GN= | 37 | 17 | 29 | 16 | 332 | 36,5 | 7,74 | 59,99 | 17 | 0,873 | 0,945 | 1,018 | 1,003 | 0,928 | 0,915 | 1,036 | 0,946 | 1,00 | 0,00 | 9,4E-01 |
| Q8R1I1 | Cytochrome b-c1 complex subunit 9 OS=Mus musculus OX=10090 G | 27 | 1 | 2 | 1 | 64 | 7,4 | 9,19 | 6,17 | 1 | 0,98 | 1,104 | 0,926 | 0,878 | 0,955 | 1,158 | 0,878 | 0,87 | 1,01 | 0,01 | 9,4E-01 |
| Q8BGH2 | Sorting and assembly machinery component 50 homolog OS=Mus m | 21 | 8 | 11 | 8 | 469 | 51,8 | 6,8 | 19,31 | 8 | 1,085 | 1,126 | 1,024 | 0,907 | 0,956 | 1,115 | 1,022 | 1,068 | 1,00 | 0,00 | 9,4E-01 |
| P49070 | Calcium signal-modulating cyclophilin ligand OS=Mus musculus OX= | 14 | 3 | 3 | 3 | 294 | 32,5 | 7,74 | 4,69 | 3 | 0,888 | 0,791 | 0,76 | 0,873 | 0,938 | 0,775 | 0,754 | 0,862 | 0,99 | -0,01 | 9,4E-01 |
| Q91XV3 | Brain acid soluble protein 1 OS=Mus musculus OX=10090 GN=Basp | 90 | 20 | 105 | 20 | 226 | 22,1 | 4,51 | 252,88 | 20 | 0,672 | 0,723 | 0,934 | 1,145 | 1,035 | 0,623 | 0,961 | 0,9 | 0,99 | -0,01 | 9,4E-01 |
| Q8BPM0 | Disheveled-associated activator of morphogenesis 1 OS=Mus musc | 11 | 11 | 13 | 11 | 1077 | 123,3 | 7,46 | 29,02 | 11 | 1,064 | 1,079 | 1,018 | 1,058 | 1,002 | 1,096 | 1,043 | 1,086 | 1,00 | 0,00 | 9,4E-01 |
| Q8C754 | Vacuolar protein sorting-associated protein 52 homolog OS=Mus m | 11 | 7 | 8 | 7 | 723 | 82 | 5,9 | 15,74 | 7 | 1,167 | 1,115 | 1,012 | 0,969 | 1,041 | 1,12 | 0,97 | 1,151 | 1,00 | 0,00 | 9,4E-01 |
| Q80Y98 | Phospholipase DDHD2 OS=Mus musculus OX=10090 GN=Ddhd2 PE | 1 | 1 | 1 | 1 | 699 | 79,5 | 5,31 | 2,09 | 1 | 0,996 | 1,04 | 1,176 | 0,938 | 0,888 | 1,018 | 1,144 | 1,077 | 1,01 | 0,01 | 9,4E-01 |

| | | | | | | | | | | | | | | | | | | | | | |
|---|---|---|---|---|---|---|---|---|---|---|---|---|---|---|---|---|---|---|---|---|---|
| Q9D924 | Iron-sulfur cluster assembly 1 homolog, mitochondrial OS=Mus mu | 32 | 3 | 5 | 3 | 129 | 14,2 | 9,07 | 11,89 | 3 | 1,12 | 1,275 | 0,961 | 1,02 | 0,865 | 1,239 | 1,286 | 1,023 | 0,99 | -0,01 | 9,4E-01 |
| Q8CC88 | von Willebrand factor A domain-containing protein 8 OS=Mus musc | 5 | 7 | 7 | 7 | 1905 | 213,3 | 6,6 | 10,63 | 7 | 1,127 | 1,141 | 1,043 | 1,017 | 1,027 | 1,185 | 0,953 | 1,144 | 1,00 | 0,00 | 9,4E-01 |
| Q8BWG8 | Beta-arrestin-1 OS=Mus musculus OX=10090 GN=Arrb1 PE=1 SV=1 | 14 | 5 | 5 | 5 | 418 | 46,9 | 6,28 | 6,22 | 5 | 0,971 | 0,882 | 1,051 | 0,988 | 0,989 | 0,71 | 0,973 | 1,188 | 1,01 | 0,01 | 9,4E-01 |
| Q6P9K9 | Neurexin-3 OS=Mus musculus OX=10090 GN=Nrxn3 PE=1 SV=2 | 7 | 8 | 10 | 5 | 1571 | 173,3 | 6,14 | 18,76 | 8 | 1,356 | 1,114 | 1,214 | 1,05 | 1,108 | 1,212 | 1,218 | 1,218 | 1,00 | 0,00 | 9,4E-01 |
| Q05816 | Fatty acid-binding protein 5 OS=Mus musculus OX=10090 GN=Fabp | 66 | 7 | 12 | 7 | 135 | 15,1 | 6,54 | 29,93 | 7 | 1,004 | 0,929 | 1,01 | 1,14 | 0,98 | 1,038 | 1,071 | 0,979 | 1,00 | 0,00 | 9,4E-01 |
| Q91VT1 | E3 SUMO-protein ligase NSE2 OS=Mus musculus OX=10090 GN=Ns | 4 | 1 | 1 | 1 | 247 | 28,2 | 8 | 0 | 1 | 0,684 | 0,417 | 0,435 | 0,904 | 0,683 | 0,551 | 0,608 | 0,634 | 0,99 | -0,01 | 9,4E-01 |
| Q8BUL6 | Pleckstrin homology domain-containing family A member 1 OS=Mu | 5 | 2 | 2 | 2 | 383 | 43,3 | 9,22 | 1,85 | 2 | 0,886 | 1,15 | 0,841 | 0,767 | 0,76 | 1,014 | 0,944 | 0,896 | 1,01 | 0,01 | 9,4E-01 |
| Q8BIP0 | Aspartate--tRNA ligase, mitochondrial OS=Mus musculus OX=1009( | 11 | 5 | 5 | 5 | 653 | 74,1 | 6,98 | 0 | 5 | 1,523 | 1,157 | 1,232 | 1,186 | 1,213 | 1,36 | 1,124 | 1,37 | 1,01 | 0,01 | 9,4E-01 |
| Q9D1K2 | V-type proton ATPase subunit F OS=Mus musculus OX=10090 GN=/ | 35 | 4 | 6 | 4 | 119 | 13,4 | 5,82 | 17,69 | 4 | 1,04 | 1,073 | 0,957 | 0,98 | 0,963 | 1 | 1,045 | 1,052 | 1,00 | 0,00 | 9,4E-01 |
| Q9D0E3 | LysM and putative peptidoglycan-binding domain-containing prote | 25 | 6 | 6 | 5 | 226 | 24,8 | 8,92 | 10,74 | 6 | 0,899 | 1,003 | 1,068 | 1,182 | 0,983 | 0,939 | 1,172 | 1,035 | 1,01 | 0,01 | 9,4E-01 |
| Q8BN59 | La-related protein 6 OS=Mus musculus OX=10090 GN=Larp6 PE=1 S | 2 | 1 | 1 | 1 | 492 | 54,8 | 7,9 | 2,45 | 1 | 0,01 | 1,008 | 0,435 | 0,841 | 0,512 | 0,623 | 0,828 | 0,402 | 0,97 | -0,03 | 9,4E-01 |
| Q69ZK0 | Phosphatidylinositol 3,4,5-trisphosphate-dependent Rac exchanger | 3 | 4 | 5 | 4 | 1650 | 184,8 | 6,29 | 5,15 | 4 | 1,55 | 1,249 | 1,873 | 1,009 | 1,306 | 1,718 | 1,281 | 1,313 | 1,01 | 0,01 | 9,4E-01 |
| Q8BWT1 | 3-ketoacyl-CoA thiolase, mitochondrial OS=Mus musculus OX=1009 | 37 | 10 | 15 | 10 | 397 | 41,8 | 8,09 | 34,87 | 10 | 1,038 | 0,959 | 1,068 | 1,027 | 0,965 | 1,065 | 0,989 | 1,063 | 1,00 | 0,00 | 9,4E-01 |
| P28867 | Protein kinase C delta type OS=Mus musculus OX=10090 GN=Prkcd | 13 | 8 | 9 | 7 | 674 | 77,5 | 7,39 | 14,91 | 8 | 1,197 | 0,998 | 0,875 | 0,942 | 1,018 | 1,074 | 0,911 | 0,987 | 1,01 | 0,01 | 9,5E-01 |
| Q9CR30 | Josephin-2 OS=Mus musculus OX=10090 GN=Josd2 PE=1 SV=1 | 6 | 1 | 1 | 1 | 188 | 20,8 | 6,54 | 2,89 | 1 | 1,612 | 1,7 | 1,559 | 1,214 | 1,305 | 1,635 | 1,675 | 1,431 | 1,01 | 0,01 | 9,5E-01 |
| Q80TL4 | PHD finger protein 24 OS=Mus musculus OX=10090 GN=Phf24 PE=: | 40 | 15 | 25 | 15 | 400 | 45,2 | 5,77 | 62,31 | 15 | 0,912 | 0,958 | 0,977 | 1,099 | 0,971 | 0,937 | 1,075 | 0,977 | 1,00 | 0,00 | 9,5E-01 |
| Q3URD3 | Sarcolemmal membrane-associated protein OS=Mus musculus OX= | 9 | 7 | 7 | 7 | 845 | 96,9 | 5,33 | 8,7 | 7 | 1,311 | 1,124 | 1,086 | 1,144 | 1,02 | 1,258 | 1,122 | 1,244 | 1,00 | 0,00 | 9,5E-01 |
| P28184 | Metallothionein-3 OS=Mus musculus OX=10090 GN=Mt3 PE=1 SV= | 18 | 1 | 1 | 1 | 68 | 7 | 7,47 | 2,98 | 1 | 0,913 | 0,452 | 1,612 | 1,056 | 1,242 | 0,845 | 1,072 | 0,944 | 0,98 | -0,02 | 9,5E-01 |
| Q6A065 | Centrosomal protein of 170 kDa OS=Mus musculus OX=10090 GN= | 5 | 8 | 8 | 8 | 1588 | 174,9 | 7,17 | 14,33 | 8 | 0,917 | 1,096 | 1,056 | 1,237 | 1,12 | 0,99 | 1,192 | 1,026 | 0,99 | -0,01 | 9,5E-01 |
| Q9D424 | Calcium-binding tyrosine phosphorylation-regulated protein OS=M | 2 | 1 | 1 | 1 | 453 | 48,3 | 4,54 | 0 | 1 | 0,721 | 1,432 | 0,662 | 0,425 | 1,818 | 0,875 | 0,308 | 0,351 | 0,97 | -0,03 | 9,5E-01 |
| Q5SVL6 | Rap1 GTPase-activating protein 2 OS=Mus musculus OX=10090 GN | 13 | 8 | 10 | 7 | 712 | 78,2 | 6,43 | 18 | 8 | 0,925 | 0,915 | 0,986 | 1,09 | 1,018 | 0,936 | 1,037 | 0,938 | 1,00 | 0,00 | 9,5E-01 |
| O35215 | D-dopachrome decarboxylase OS=Mus musculus OX=10090 GN=Dc | 24 | 3 | 4 | 3 | 118 | 13,1 | 6,54 | 9,65 | 3 | 1,014 | 1,101 | 1,043 | 0,942 | 0,996 | 1,126 | 0,987 | 0,978 | 1,00 | 0,00 | 9,5E-01 |
| P48428 | Tubulin-specific chaperone A OS=Mus musculus OX=10090 GN=Tbc | 31 | 4 | 4 | 4 | 108 | 12,8 | 5,27 | 5,76 | 4 | 0,86 | 1,112 | 0,836 | 0,999 | 0,906 | 1,06 | 0,997 | 0,865 | 0,99 | -0,01 | 9,5E-01 |
| Q9CWZ7 | Gamma-soluble NSF attachment protein OS=Mus musculus OX=10( | 33 | 12 | 19 | 12 | 312 | 34,7 | 5,41 | 48,56 | 12 | 0,896 | 0,941 | 0,991 | 1,017 | 0,976 | 0,926 | 0,986 | 0,965 | 1,00 | 0,00 | 9,5E-01 |
| P62077 | Mitochondrial import inner membrane translocase subunit Tim8 B | 8 | 1 | 1 | 1 | 83 | 9,3 | 5,12 | 2,59 | 1 | 1,044 | 0,986 | 0,944 | 0,943 | 1,032 | 1,001 | 0,929 | 0,946 | 1,00 | 0,00 | 9,5E-01 |
| Q9EST1 | Gasdermin-A OS=Mus musculus OX=10090 GN=Gsdma PE=2 SV=1 | 2 | 1 | 1 | 1 | 446 | 49,6 | 5,57 | 1,89 | 1 | 1,584 | 1,534 | 1,05 | 1,081 | 1,151 | 1,248 | 1,356 | 1,538 | 0,99 | -0,01 | 9,5E-01 |
| Q7TMM9 | Tubulin beta-2A chain OS=Mus musculus OX=10090 GN=Tubb2a PE | 61 | 25 | 163 | 1 | 445 | 49,9 | 4,89 | 338,83 | 25 | 0,78 | 0,908 | 0,816 | 1,051 | 0,865 | 0,733 | 1,004 | 0,931 | 1,01 | 0,01 | 9,5E-01 |
| Q9QZ08 | N-acetyl-D-glucosamine kinase OS=Mus musculus OX=10090 GN=N | 6 | 2 | 3 | 2 | 343 | 37,2 | 5,73 | 5,27 | 2 | 1,229 | 0,952 | 1,161 | 1,229 | 1,196 | 1,112 | 1,115 | 1,13 | 1,00 | 0,00 | 9,5E-01 |
| P97449 | Aminopeptidase N OS=Mus musculus OX=10090 GN=Anpep PE=1 S | 3 | 3 | 3 | 3 | 966 | 109,6 | 5,9 | 5,02 | 3 | 0,85 | 1,474 | 0,907 | 0,617 | 0,842 | 1,197 | 0,711 | 1,155 | 0,99 | -0,01 | 9,5E-01 |
| Q9EQZ7 | Regulating synaptic membrane exocytosis protein 2 OS=Mus muscu | 2 | 3 | 3 | 3 | 1530 | 172,8 | 9,25 | 3,23 | 3 | 0,774 | 0,898 | 0,969 | 1,154 | 1,046 | 0,988 | 0,869 | 0,915 | 0,99 | -0,01 | 9,5E-01 |
| P60879-2 | Isoform 2 of Synaptosomal-associated protein 25 OS=Mus musculu | 62 | 18 | 80 | 1 | 206 | 23,3 | 4,86 | 190,51 | 18 | 0,862 | 1,151 | 0,659 | 0,901 | 0,912 | 0,846 | 0,945 | 0,843 | 1,01 | 0,01 | 9,5E-01 |
| Q8CHH9 | Septin-8 OS=Mus musculus OX=10090 GN=Septin8 PE=1 SV=4 | 25 | 11 | 13 | 9 | 429 | 49,8 | 6 | 30,47 | 11 | 0,855 | 1,005 | 0,94 | 0,954 | 0,866 | 0,986 | 0,911 | 0,98 | 1,00 | 0,00 | 9,5E-01 |
| Q3TC72 | Fumarylacetoacetate hydrolase domain-containing protein 2A OS= | 6 | 1 | 2 | 1 | 313 | 34,7 | 8,16 | 8,46 | 1 | 0,777 | 0,772 | 0,288 | 0,868 | 0,966 | 0,467 | 0,808 | 0,51 | 0,98 | -0,02 | 9,5E-01 |
| Q9DCD6 | Gamma-aminobutyric acid receptor-associated protein OS=Mus mu | 19 | 2 | 2 | 2 | 117 | 13,9 | 8,79 | 3,67 | 2 | 0,978 | 0,98 | 0,95 | 1,075 | 0,932 | 1,018 | 0,996 | 1,028 | 1,00 | 0,00 | 9,5E-01 |
| Q8VDD5 | Myosin-9 OS=Mus musculus OX=10090 GN=Myh9 PE=1 SV=4 | 21 | 42 | 53 | 35 | 1960 | 226,2 | 5,66 | 103,31 | 42 | 1,007 | 1,187 | 0,997 | 0,947 | 0,978 | 1,124 | 0,994 | 1,058 | 1,00 | 0,00 | 9,5E-01 |
| A2AAE1 | Transmembrane protein KIAA1109 OS=Mus musculus OX=10090 GI | 1 | 3 | 3 | 3 | 5005 | 555 | 6,61 | 4,62 | 3 | 0,693 | 1,013 | 1,348 | 1,035 | 0,65 | 1,764 | 0,616 | 1,136 | 0,98 | -0,02 | 9,5E-01 |
| Q9CRD2 | ER membrane protein complex subunit 2 OS=Mus musculus OX=10 | 8 | 2 | 2 | 2 | 297 | 34,9 | 6,81 | 5,78 | 2 | 1,167 | 1,075 | 0,761 | 0,883 | 0,841 | 1,399 | 0,922 | 0,678 | 1,01 | 0,01 | 9,5E-01 |
| Q60668 | Heterogeneous nuclear ribonucleoprotein D0 OS=Mus musculus O) | 9 | 3 | 3 | 2 | 355 | 38,3 | 7,81 | 3,17 | 3 | 1,136 | 1,028 | 1,036 | 0,855 | 0,933 | 1,233 | 0,798 | 1,063 | 1,01 | 0,01 | 9,5E-01 |
| Q8BGX3 | Leucine-rich repeat and transmembrane domain-containing protei | 4 | 1 | 1 | 1 | 370 | 41 | 6,4 | 2,8 | 1 | 1,304 | 1,277 | 1,225 | 0,907 | 1,182 | 1,008 | 1,157 | 1,337 | 1,01 | 0,01 | 9,5E-01 |
| Q7TQF7 | Amphiphysin OS=Mus musculus OX=10090 GN=Amph PE=1 SV=1 | 30 | 22 | 42 | 20 | 686 | 75 | 4,63 | 111,92 | 22 | 0,984 | 0,982 | 0,979 | 1,018 | 0,916 | 1,013 | 1,033 | 1,008 | 1,00 | 0,00 | 9,5E-01 |
| O35066 | Kinesin-like protein KIF3C OS=Mus musculus OX=10090 GN=Kif3c P | 5 | 4 | 4 | 2 | 796 | 89,9 | 8,06 | 9,08 | 4 | 0,92 | 1,087 | 0,895 | 1,135 | 1,088 | 0,951 | 1,053 | 0,927 | 1,00 | 0,00 | 9,5E-01 |
| P97492 | Regulator of G-protein signaling 14 OS=Mus musculus OX=10090 G | 6 | 3 | 3 | 3 | 547 | 59,8 | 7,43 | 3,13 | 3 | 0,687 | 0,839 | 1,064 | 0,889 | 0,672 | 0,773 | 0,84 | 1,161 | 1,01 | 0,01 | 9,5E-01 |
| P13439 | Uridine 5'-monophosphate synthase OS=Mus musculus OX=10090 | 2 | 1 | 1 | 1 | 481 | 52,3 | 6,61 | 2,82 | 1 | 0,899 | 1,048 | 1,194 | 0,902 | 0,829 | 1,266 | 1,039 | 0,938 | 0,99 | -0,01 | 9,5E-01 |
| Q812A2 | SLIT-ROBO Rho GTPase-activating protein 3 OS=Mus musculus OX= | 15 | 18 | 20 | 18 | 1099 | 124,3 | 6,67 | 37,36 | 18 | 1,064 | 1,022 | 0,962 | 0,994 | 1,022 | 1,035 | 0,957 | 1,035 | 1,00 | 0,00 | 9,5E-01 |
| Q80Y55 | BSD domain-containing protein 1 OS=Mus musculus OX=10090 GN= | 7 | 2 | 3 | 2 | 427 | 46,9 | 4,4 | 4,84 | 2 | 0,948 | 1,057 | 1,327 | 1,214 | 1,186 | 1,038 | 1,17 | 1,13 | 1,00 | 0,00 | 9,5E-01 |
| P70208 | Plexin-A3 OS=Mus musculus OX=10090 GN=Plxna3 PE=1 SV=2 | 3 | 6 | 9 | 1 | 1872 | 207,8 | 7,09 | 17,23 | 6 | 2,01 | 1,278 | 1,877 | 1,351 | 1,65 | 1,629 | 1,558 | 1,634 | 1,01 | 0,01 | 9,5E-01 |
| Q8BK64 | Activator of 90 kDa heat shock protein ATPase homolog 1 OS=Mus | 14 | 5 | 5 | 5 | 338 | 38,1 | 5,53 | 13,45 | 5 | 1,175 | 1,034 | 0,927 | 0,9 | 0,862 | 1,111 | 1,024 | 1,059 | 1,00 | 0,00 | 9,5E-01 |
| Q9DB75 | Cell death-inducing p53-target protein 1 OS=Mus musculus OX=10( | 4 | 1 | 1 | 1 | 208 | 21,8 | 5,66 | 2,12 | 1 | 0,999 | 0,98 | 0,828 | 0,869 | 0,871 | 0,802 | 0,978 | 1,01 | 1,00 | 0,00 | 9,5E-01 |
| Q9CQ60 | 6-phosphogluconolactonase OS=Mus musculus OX=10090 GN=Pgls | 19 | 4 | 4 | 4 | 257 | 27,2 | 5,85 | 5,12 | 4 | 1,074 | 1,248 | 1,048 | 1,093 | 0,995 | 1,312 | 1,117 | 1,019 | 1,00 | 0,00 | 9,6E-01 |
| Q6PH08 | ERC protein 2 OS=Mus musculus OX=10090 GN=Erc2 PE=1 SV=2 | 24 | 26 | 32 | 15 | 957 | 110,6 | 7,03 | 71,28 | 26 | 0,965 | 1,048 | 0,991 | 0,996 | 0,962 | 1,085 | 0,965 | 0,98 | 1,00 | 0,00 | 9,6E-01 |
| P63242 | Eukaryotic translation initiation factor 5A-1 OS=Mus musculus OX= | 36 | 5 | 6 | 5 | 154 | 16,8 | 5,24 | 5,14 | 5 | 1,156 | 1,046 | 1,037 | 1,216 | 1,052 | 0,989 | 1,236 | 1,195 | 1,00 | 0,00 | 9,6E-01 |
| P61358 | 60S ribosomal protein L27 OS=Mus musculus OX=10090 GN=Rpl27 | 7 | 1 | 1 | 1 | 136 | 15,8 | 10,56 | 0 | 1 | 0,95 | 0,952 | 1,134 | 0,715 | 0,919 | 0,722 | 0,991 | 1,092 | 1,01 | 0,01 | 9,6E-01 |
| O35454 | Chloride transport protein 6 OS=Mus musculus OX=10090 GN=Clcn | 3 | 2 | 3 | 2 | 870 | 96,9 | 7,02 | 0 | 2 | 1,775 | 1,035 | 1,068 | 1,216 | 1,526 | 1,208 | 1,133 | 1,182 | 1,01 | 0,01 | 9,6E-01 |
| Q6NS52-2 | Isoform 2 of Diacylglycerol kinase beta OS=Mus musculus OX=1009 | 12 | 8 | 8 | 8 | 795 | 89,4 | 7,93 | 13,08 | 8 | 1,338 | 0,995 | 1,064 | 0,847 | 0,952 | 1,274 | 0,973 | 1,074 | 0,99 | -0,01 | 9,6E-01 |
| Q61390 | T-complex protein 1 subunit zeta-2 OS=Mus musculus OX=10090 G | 11 | 5 | 6 | 1 | 531 | 58,1 | 7,36 | 17,78 | 5 | 1,028 | 0,952 | 1,117 | 1,143 | 0,882 | 0,969 | 1,172 | 1,197 | 1,00 | 0,00 | 9,6E-01 |
| Q69ZK9 | Neuroligin-2 OS=Mus musculus OX=10090 GN=Nlgn2 PE=1 SV=2 | 6 | 3 | 4 | 3 | 836 | 90,9 | 6,18 | 7,35 | 3 | 1,728 | 0,78 | 0,945 | 1,152 | 1,014 | 1,378 | 1,06 | 1,203 | 0,99 | -0,01 | 9,6E-01 |
| P63011 | Ras-related protein Rab-3A OS=Mus musculus OX=10090 GN=Rab3 | 55 | 14 | 47 | 8 | 220 | 25 | 5,03 | 90,29 | 14 | 1,035 | 1,025 | 1,025 | 1,015 | 1,039 | 1,099 | 1,004 | 0,951 | 1,00 | 0,00 | 9,6E-01 |
| Q61191 | Host cell factor 1 OS=Mus musculus OX=10090 GN=Hcfc1 PE=1 SV= | 2 | 4 | 4 | 4 | 2045 | 210,3 | 7,18 | 2,22 | 4 | 1,305 | 1,047 | 1,298 | 1,27 | 1,146 | 1,424 | 1,071 | 1,301 | 1,00 | 0,00 | 9,6E-01 |
| Q8R5A6 | TBC1 domain family member 22A OS=Mus musculus OX=10090 GN | 2 | 1 | 1 | 1 | 516 | 59,3 | 6,46 | 2,01 | 1 | 2,668 | 1,103 | 1,39 | 0,986 | 0,837 | 2,55 | 1,204 | 1,673 | 0,98 | -0,02 | 9,6E-01 |
| Q9CR57 | 60S ribosomal protein L14 OS=Mus musculus OX=10090 GN=Rpl14 | 4 | 1 | 2 | 1 | 217 | 23,5 | 11,02 | 1,93 | 1 | 1,428 | 0,862 | 1 | 1,118 | 1,235 | 1,094 | 0,93 | 1,179 | 0,99 | -0,01 | 9,6E-01 |
| Q8C419 | Probable G-protein coupled receptor 158 OS=Mus musculus OX=10 | 11 | 11 | 12 | 11 | 1200 | 134,3 | 8,09 | 24,53 | 11 | 1,116 | 0,883 | 1,01 | 0,991 | 1,056 | 0,978 | 0,985 | 0,992 | 1,00 | 0,00 | 9,6E-01 |
| P99029 | Peroxiredoxin-5, mitochondrial OS=Mus musculus OX=10090 GN=P | 28 | 5 | 9 | 5 | 210 | 21,9 | 8,85 | 30,37 | 5 | 0,945 | 0,965 | 1,011 | 1,045 | 0,958 | 0,948 | 1,028 | 1,039 | 1,00 | 0,00 | 9,6E-01 |

| | | | | | | | | | | | | | | | | | | | | | |
|---|---|---|---|---|---|---|---|---|---|---|---|---|---|---|---|---|---|---|---|---|---|
| Q9Z2W0 | Aspartyl aminopeptidase OS=Mus musculus OX=10090 GN=Dnpep | 9 | 3 | 6 | 3 | 473 | 52,2 | 7,25 | 5,39 | 3 | 1,105 | 1,122 | 0,718 | 0,934 | 1,074 | 0,839 | 0,844 | 1,097 | 1,01 | 0,01 | 9,6E-01 |
| O88741 | Ganglioside-induced differentiation-associated protein 1 OS=Mus m | 25 | 10 | 14 | 9 | 358 | 41,3 | 8,37 | 32,33 | 10 | 1,008 | 1,16 | 0,915 | 0,985 | 1,03 | 0,927 | 1,083 | 1,041 | 1,00 | 0,00 | 9,6E-01 |
| P15532 | Nucleoside diphosphate kinase A OS=Mus musculus OX=10090 GN= | 44 | 6 | 6 | 3 | 152 | 17,2 | 7,37 | 11,53 | 6 | 1,062 | 1,155 | 1,191 | 1,177 | 1,064 | 1,206 | 1,142 | 1,182 | 1,00 | 0,00 | 9,6E-01 |
| Q61316 | Heat shock 70 kDa protein 4 OS=Mus musculus OX=10090 GN=Hsp | 48 | 35 | 48 | 32 | 841 | 94,1 | 5,24 | 111,22 | 35 | 1,069 | 0,922 | 1,022 | 0,997 | 0,977 | 0,985 | 1,039 | 1,002 | 1,00 | 0,00 | 9,6E-01 |
| Q9CT10 | Ran-binding protein 3 OS=Mus musculus OX=10090 GN=Ranbp3 PE | 2 | 1 | 1 | 1 | 491 | 52,5 | 5,12 | 2,1 | 1 | 1,753 | 1,372 | 0,751 | 1,045 | 0,888 | 1,531 | 1,69 | 0,747 | 1,01 | 0,01 | 9,6E-01 |
| P28738 | Kinesin heavy chain isoform 5C OS=Mus musculus OX=10090 GN=K | 29 | 26 | 40 | 22 | 956 | 109,2 | 6,19 | 82,28 | 26 | 1,004 | 1,066 | 1,019 | 1,035 | 0,927 | 1,132 | 1,03 | 1,044 | 1,00 | 0,00 | 9,6E-01 |
| Q7TSC1 | Protein PRRC2A OS=Mus musculus OX=10090 GN=Prrc2a PE=1 SV= | 1 | 2 | 2 | 2 | 2158 | 229,1 | 9,39 | 2,03 | 2 | 1,448 | 1,487 | 0,926 | 0,35 | 1,241 | 1,32 | 0,739 | 0,971 | 0,99 | -0,01 | 9,6E-01 |
| Q5SUR0 | Phosphoribosylformylglycinamidine synthase OS=Mus musculus OX | 2 | 2 | 2 | 2 | 1337 | 144,5 | 5,67 | 3,19 | 2 | 1,034 | 1,328 | 1,013 | 0,993 | 1,073 | 0,992 | 1,149 | 1,171 | 1,00 | 0,00 | 9,6E-01 |
| Q9D2M8 | Ubiquitin-conjugating enzyme E2 variant 2 OS=Mus musculus OX=1 | 33 | 4 | 4 | 1 | 145 | 16,4 | 8,09 | 10,65 | 4 | 0,967 | 1,143 | 0,958 | 1,043 | 1,121 | 0,986 | 0,942 | 1,051 | 1,00 | 0,00 | 9,6E-01 |
| Q64444 | Carbonic anhydrase 4 OS=Mus musculus OX=10090 GN=Ca4 PE=1 S | 17 | 5 | 5 | 5 | 305 | 34,3 | 8,21 | 8,17 | 5 | 1,166 | 1,033 | 0,776 | 0,812 | 0,771 | 1,222 | 0,774 | 1,047 | 0,99 | -0,01 | 9,6E-01 |
| Q9CQV6 | Microtubule-associated proteins 1A/1B light chain 3B OS=Mus mus | 17 | 2 | 2 | 1 | 125 | 14,6 | 8,43 | 5,68 | 2 | 0,958 | 1,064 | 0,98 | 1,062 | 1,063 | 0,971 | 1,06 | 0,977 | 1,00 | 0,00 | 9,6E-01 |
| B1AWN6 | Sodium channel protein type 2 subunit alpha OS=Mus musculus OX | 7 | 15 | 22 | 8 | 2006 | 227,8 | 5,67 | 61,66 | 15 | 0,948 | 1,08 | 1,12 | 1,114 | 0,988 | 1,101 | 1,084 | 1,098 | 1,00 | 0,00 | 9,6E-01 |
| Q9CS84 | Neurexin-1 OS=Mus musculus OX=10090 GN=Nrxn1 PE=1 SV=3 | 11 | 13 | 15 | 11 | 1514 | 166,1 | 6,09 | 30,44 | 13 | 1,223 | 1,091 | 1,048 | 0,98 | 1,029 | 1,193 | 1,025 | 1,083 | 1,00 | 0,00 | 9,6E-01 |
| Q9QZB0 | Regulator of G-protein signaling 17 OS=Mus musculus OX=10090 G | 4 | 1 | 1 | 1 | 210 | 24,3 | 5,71 | 0 | 1 | 1,047 | 1,02 | 1,141 | 0,947 | 0,961 | 1,014 | 1,184 | 0,984 | 1,00 | 0,00 | 9,6E-01 |
| O55022 | Membrane-associated progesterone receptor component 1 OS=Mu | 31 | 6 | 9 | 5 | 195 | 21,7 | 4,7 | 15,63 | 6 | 1,181 | 1,02 | 1,13 | 0,885 | 1,049 | 1,127 | 1,009 | 1,044 | 1,00 | 0,00 | 9,6E-01 |
| Q923U0 | TOM1-like protein 1 OS=Mus musculus OX=10090 GN=Tom1l1 PE= | 2 | 1 | 1 | 1 | 474 | 52,7 | 5,36 | 3,23 | 1 | 0,835 | 0,939 | 1,321 | 1,729 | 1,327 | 0,787 | 1,098 | 1,661 | 0,99 | -0,01 | 9,7E-01 |
| Q8R180 | ERO1-like protein alpha OS=Mus musculus OX=10090 GN=Ero1a PE | 7 | 3 | 4 | 3 | 464 | 54,1 | 6,54 | 9,48 | 3 | 1,192 | 0,969 | 0,798 | 0,943 | 0,917 | 1,163 | 0,918 | 0,886 | 1,00 | 0,00 | 9,7E-01 |
| Q80UM7 | Mannosyl-oligosaccharide glucosidase OS=Mus musculus OX=1009 | 5 | 3 | 3 | 3 | 834 | 91,8 | 9 | 7,71 | 3 | 1,502 | 1,146 | 1,575 | 1,331 | 1,306 | 1,668 | 1,352 | 1,204 | 1,00 | 0,00 | 9,7E-01 |
| Q80ZD8 | Amphoterin-induced protein 1 OS=Mus musculus OX=10090 GN=A | 11 | 4 | 4 | 4 | 492 | 55,3 | 6,86 | 5,23 | 4 | 0,941 | 1,121 | 0,857 | 0,703 | 0,654 | 0,846 | 1,06 | 1,085 | 0,99 | -0,01 | 9,7E-01 |
| Q8CFJ9 | GATOR complex protein WDR24 OS=Mus musculus OX=10090 GN= | 1 | 1 | 1 | 1 | 790 | 88,1 | 6,46 | 0 | 1 | 0,931 | 0,47 | 1,174 | 0,72 | 0,597 | 0,905 | 1,052 | 0,71 | 1,01 | 0,01 | 9,7E-01 |
| Q8VIM9 | Immunity-related GTPase family Q protein OS=Mus musculus OX=1 | 4 | 2 | 2 | 2 | 583 | 59,3 | 4,83 | 2,98 | 2 | 1,073 | 1,182 | 1,244 | 1,093 | 1,028 | 1,201 | 1,135 | 1,218 | 1,00 | 0,00 | 9,7E-01 |
| Q8BPN8 | DmX-like protein 2 OS=Mus musculus OX=10090 GN=Dmxl2 PE=1 S | 20 | 54 | 72 | 53 | 3032 | 338 | 6,42 | 182,87 | 54 | 1,006 | 0,997 | 1,024 | 1,036 | 1,009 | 1,057 | 1,011 | 0,983 | 1,00 | 0,00 | 9,7E-01 |
| Q8VCW8 | Medium-chain acyl-CoA ligase ACSF2, mitochondrial OS=Mus musc | 11 | 7 | 8 | 7 | 615 | 67,9 | 8,18 | 15,85 | 7 | 0,971 | 1,056 | 1,123 | 0,952 | 0,966 | 1,175 | 0,899 | 1,05 | 1,00 | 0,00 | 9,7E-01 |
| Q3UMB9 | WASH complex subunit 4 OS=Mus musculus OX=10090 GN=Washc | 1 | 1 | 1 | 1 | 1173 | 136,3 | 7,37 | 0 | 1 | 0,994 | 1,014 | 0,822 | 1,065 | 0,791 | 1,315 | 0,576 | 1,183 | 1,01 | 0,01 | 9,7E-01 |
| Q60780 | Growth arrest-specific protein 7 OS=Mus musculus OX=10090 GN= | 10 | 4 | 5 | 4 | 421 | 48,1 | 7,83 | 10,4 | 4 | 1,168 | 0,936 | 1,123 | 1,181 | 1,103 | 1,017 | 1,237 | 1,063 | 1,00 | 0,00 | 9,7E-01 |
| O08539 | Myc box-dependent-interacting protein 1 OS=Mus musculus OX=1 | 31 | 17 | 28 | 6 | 588 | 64,4 | 5,03 | 59,99 | 17 | 0,941 | 0,958 | 1,169 | 1,249 | 1,103 | 1,051 | 1,035 | 1,141 | 1,00 | 0,00 | 9,7E-01 |
| A2AJI0 | MAP7 domain-containing protein 1 OS=Mus musculus OX=10090 G | 4 | 3 | 3 | 3 | 846 | 93,2 | 10,15 | 2,07 | 3 | 0,974 | 1,116 | 0,716 | 0,814 | 0,831 | 1,103 | 0,861 | 0,807 | 1,00 | 0,00 | 9,7E-01 |
| Q9WTX8 | Mitotic spindle assembly checkpoint protein MAD1 OS=Mus muscu | 1 | 1 | 2 | 1 | 717 | 83,5 | 5,66 | 0 | 1 | 1,459 | 1,419 | 1,783 | 0,635 | 0,76 | 2,594 | 0,737 | 1,124 | 1,02 | 0,02 | 9,7E-01 |
| Q80TM9 | Nischarin OS=Mus musculus OX=10090 GN=Nisch PE=1 SV=2 | 5 | 7 | 8 | 7 | 1593 | 174,9 | 5,16 | 5,71 | 7 | 0,796 | 0,86 | 0,874 | 0,995 | 0,861 | 0,894 | 0,915 | 0,862 | 1,00 | 0,00 | 9,7E-01 |
| Q8K1S4 | Netrin receptor UNC5A OS=Mus musculus OX=10090 GN=Unc5a PE | 3 | 2 | 2 | 2 | 898 | 98,8 | 6,92 | 2,09 | 2 | 0,815 | 1,351 | 0,644 | 0,289 | 0,874 | 0,756 | 0,823 | 0,682 | 0,99 | -0,01 | 9,7E-01 |
| Q8K3C3 | Protein LZIC OS=Mus musculus OX=10090 GN=Lzic PE=1 SV=1 | 13 | 2 | 2 | 2 | 190 | 21,5 | 4,96 | 2,34 | 2 | 0,525 | 1,011 | 0,837 | 0,882 | 0,755 | 1,035 | 0,752 | 0,733 | 0,99 | -0,01 | 9,7E-01 |
| Q9D379 | Epoxide hydrolase 1 OS=Mus musculus OX=10090 GN=Ephx1 PE=1 | 5 | 2 | 3 | 2 | 455 | 52,5 | 8,35 | 5,62 | 2 | 1,429 | 1,411 | 1,156 | 1,057 | 1,152 | 1,489 | 1,192 | 1,201 | 1,00 | 0,00 | 9,7E-01 |
| Q9D1B9 | 39S ribosomal protein L28, mitochondrial OS=Mus musculus OX=1 | 5 | 1 | 1 | 1 | 257 | 30,2 | 9,29 | 2,44 | 1 | 0,709 | 1,003 | 0,938 | 1,164 | 1,174 | 0,907 | 0,859 | 0,855 | 1,01 | 0,01 | 9,7E-01 |
| Q64332 | Synapsin-2 OS=Mus musculus OX=10090 GN=Syn2 PE=1 SV=2 | 27 | 13 | 27 | 11 | 586 | 63,3 | 8,43 | 83,68 | 13 | 1,05 | 1,121 | 0,889 | 1,113 | 0,954 | 1,084 | 1,154 | 0,992 | 1,00 | 0,00 | 9,7E-01 |
| Q9QXA5 | U6 snRNA-associated Sm-like protein LSm4 OS=Mus musculus OX= | 9 | 1 | 1 | 1 | 137 | 15,1 | 10,05 | 3,36 | 1 | 1,083 | 1,091 | 1,283 | 0,883 | 0,784 | 1,495 | 0,994 | 1,093 | 0,99 | -0,01 | 9,7E-01 |
| Q9CYZ2 | Tumor protein D54 OS=Mus musculus OX=10090 GN=Tpd52l2 PE=1 | 20 | 4 | 4 | 4 | 220 | 24 | 6,15 | 5,56 | 4 | 0,832 | 1,213 | 1,026 | 0,867 | 0,828 | 1,332 | 0,897 | 0,903 | 0,99 | -0,01 | 9,7E-01 |
| Q9Z2B2 | Brain mitochondrial carrier protein 1 OS=Mus musculus OX=10090 | 6 | 2 | 2 | 2 | 325 | 36,3 | 9,69 | 2,96 | 2 | 1,208 | 1,162 | 0,996 | 0,916 | 0,945 | 0,989 | 1 | 1,331 | 1,00 | 0,00 | 9,7E-01 |
| P26231 | Catenin alpha-1 OS=Mus musculus OX=10090 GN=Ctnna1 PE=1 SV= | 11 | 9 | 9 | 4 | 906 | 100 | 6,23 | 18,81 | 9 | 0,996 | 1,138 | 1,025 | 0,987 | 0,946 | 1,163 | 1,065 | 0,981 | 1,00 | 0,00 | 9,7E-01 |
| Q99LC5 | Electron transfer flavoprotein subunit alpha, mitochondrial OS=Mu | 29 | 8 | 10 | 8 | 333 | 35 | 8,38 | 26,22 | 8 | 1,036 | 0,992 | 1,118 | 0,983 | 1,02 | 1,156 | 0,98 | 0,965 | 1,00 | 0,00 | 9,7E-01 |
| Q9CQ69 | Cytochrome b-c1 complex subunit 8 OS=Mus musculus OX=10090 G | 29 | 3 | 4 | 3 | 82 | 9,8 | 10,26 | 7,17 | 3 | 1,243 | 1,117 | 1,112 | 1,123 | 1,098 | 1,391 | 0,97 | 1,15 | 1,00 | 0,00 | 9,7E-01 |
| O70566 | Protein diaphanous homolog 2 OS=Mus musculus OX=10090 GN=D | 4 | 3 | 3 | 3 | 1098 | 124,8 | 6,92 | 0 | 3 | 0,985 | 1,292 | 0,642 | 0,724 | 0,645 | 1,187 | 0,653 | 1,127 | 1,01 | 0,01 | 9,7E-01 |
| Q9DCB4 | cAMP-regulated phosphoprotein 21 OS=Mus musculus OX=10090 G | 3 | 2 | 2 | 2 | 807 | 88,5 | 7,39 | 5,07 | 2 | 1,3 | 0,832 | 0,99 | 1,18 | 0,907 | 1,227 | 1,181 | 1,006 | 1,00 | 0,00 | 9,7E-01 |
| Q9DD18 | D-aminoacyl-tRNA deacylase 1 OS=Mus musculus OX=10090 GN=D | 13 | 2 | 2 | 2 | 209 | 23,4 | 7,87 | 3,71 | 2 | 1,181 | 1,202 | 0,98 | 1,261 | 1,13 | 1,087 | 1,348 | 1,072 | 1,00 | 0,00 | 9,7E-01 |
| Q91ZZ3 | Beta-synuclein OS=Mus musculus OX=10090 GN=Sncb PE=1 SV=1 | 54 | 8 | 18 | 3 | 133 | 14 | 4,37 | 52,24 | 8 | 0,972 | 1,003 | 1,027 | 1,109 | 0,982 | 1,029 | 1,039 | 1,066 | 1,00 | 0,00 | 9,7E-01 |
| Q6A152 | Cytochrome P450 4X1 OS=Mus musculus OX=10090 GN=Cyp4x1 PE | 2 | 1 | 1 | 1 | 507 | 58,5 | 7,21 | 2,44 | 1 | 0,702 | 0,444 | 0,902 | 0,858 | 0,823 | 0,903 | 0,701 | 0,459 | 1,01 | 0,01 | 9,7E-01 |
| O70161 | Phosphatidylinositol 4-phosphate 5-kinase type-1 gamma OS=Mus | 22 | 10 | 12 | 10 | 661 | 72,4 | 5,59 | 35,97 | 10 | 1,268 | 1,019 | 1,044 | 0,985 | 0,93 | 1,214 | 1,069 | 1,115 | 1,00 | 0,00 | 9,7E-01 |
| Q3U487 | E3 ubiquitin-protein ligase HECTD3 OS=Mus musculus OX=10090 G | 4 | 3 | 3 | 3 | 861 | 97,3 | 5,47 | 7,54 | 3 | 1,362 | 1,004 | 0,992 | 1,19 | 1,042 | 1,044 | 1,082 | 1,364 | 1,00 | 0,00 | 9,7E-01 |
| B1AXH1 | NHS-like protein 2 OS=Mus musculus OX=10090 GN=Nhsl2 PE=1 SV | 1 | 1 | 1 | 1 | 1219 | 132,3 | 8,48 | 2,01 | 1 | 1,018 | 1,063 | 0,887 | 1,386 | 1,026 | 1,091 | 1,383 | 0,833 | 1,00 | 0,00 | 9,7E-01 |
| Q9DBN4 | Putative monooxygenase p33MONOX OS=Mus musculus OX=1009 | 4 | 1 | 1 | 1 | 303 | 32,7 | 9,42 | 0 | 1 | 1,258 | 1,236 | 1,335 | 1,13 | 1,221 | 1,062 | 1,406 | 1,259 | 1,00 | 0,00 | 9,7E-01 |
| P70349 | Histidine triad nucleotide-binding protein 1 OS=Mus musculus OX= | 29 | 3 | 4 | 3 | 126 | 13,8 | 6,87 | 6,93 | 3 | 1,015 | 0,934 | 1,107 | 1,067 | 1,083 | 1,04 | 1,099 | 0,893 | 1,00 | 0,00 | 9,7E-01 |
| O70443 | Guanine nucleotide-binding protein G(z) subunit alpha OS=Mus mu | 28 | 11 | 14 | 10 | 355 | 40,8 | 7,61 | 38,83 | 11 | 1,129 | 1,072 | 1,204 | 1,037 | 1,031 | 1,237 | 1,138 | 1,044 | 1,00 | 0,00 | 9,7E-01 |
| P62812 | Gamma-aminobutyric acid receptor subunit alpha-1 OS=Mus musc | 20 | 8 | 10 | 7 | 455 | 51,7 | 9,31 | 20,64 | 8 | 1,092 | 1,048 | 1,084 | 0,943 | 1,106 | 1,097 | 1,027 | 0,93 | 1,00 | 0,00 | 9,7E-01 |
| Q9D7A8 | Armadillo repeat-containing protein 1 OS=Mus musculus OX=1009 | 16 | 6 | 7 | 6 | 282 | 31,2 | 5,57 | 15,46 | 6 | 1,065 | 1,122 | 1,065 | 1,201 | 1,084 | 1,037 | 1,175 | 1,163 | 1,00 | 0,00 | 9,8E-01 |
| Q80TN7 | Neuron navigator 3 OS=Mus musculus OX=10090 GN=Nav3 PE=1 S | 0 | 1 | 1 | 1 | 2359 | 252,1 | 8,76 | 2,57 | 1 | 0,571 | 0,949 | 0,742 | 1,074 | 0,958 | 0,651 | 0,817 | 0,927 | 0,99 | -0,01 | 9,8E-01 |
| P53612 | Geranylgeranyl transferase type-2 subunit beta OS=Mus musculus | 13 | 4 | 5 | 4 | 339 | 37,8 | 5,16 | 7,42 | 4 | 1,238 | 1,085 | 1,091 | 0,971 | 1,155 | 1,005 | 1,036 | 1,18 | 1,00 | 0,00 | 9,8E-01 |
| Q7SIG6 | Arf-GAP with SH3 domain, ANK repeat and PH domain-containing p | 15 | 10 | 11 | 9 | 958 | 106,7 | 6,65 | 21,02 | 10 | 1,033 | 1,119 | 1,121 | 0,947 | 1,15 | 1,114 | 0,834 | 1,133 | 1,00 | 0,00 | 9,8E-01 |
| Q8BTG3 | T-complex protein 11-like protein 1 OS=Mus musculus OX=10090 G | 4 | 2 | 2 | 2 | 509 | 56,3 | 5,5 | 2,56 | 2 | 1,184 | 1,081 | 1,052 | 1,029 | 0,845 | 1,375 | 0,939 | 1,203 | 1,00 | 0,00 | 9,8E-01 |
| Q3UHX2 | 28 kDa heat- and acid-stable phosphoprotein OS=Mus musculus OX | 28 | 4 | 5 | 4 | 181 | 20,6 | 7,39 | 7,27 | 4 | 0,745 | 0,891 | 0,93 | 0,963 | 0,907 | 0,989 | 0,818 | 0,807 | 1,00 | 0,00 | 9,8E-01 |
| P21956 | Lactadherin OS=Mus musculus OX=10090 GN=Mfge8 PE=1 SV=3 | 5 | 2 | 2 | 2 | 463 | 51,2 | 6,52 | 2,46 | 2 | 0,909 | 1,313 | 1,194 | 0,934 | 1,188 | 1,174 | 0,964 | 1,01 | 1,00 | 0,00 | 9,8E-01 |
| Q4KMM3 | Oxidation resistance protein 1 OS=Mus musculus OX=10090 GN=O | 34 | 29 | 37 | 29 | 866 | 95,9 | 5,33 | 101,33 | 29 | 0,964 | 1,064 | 0,972 | 1,057 | 0,967 | 1,004 | 1,026 | 1,056 | 1,00 | 0,00 | 9,8E-01 |
| Q8BNU0 | Armadillo repeat-containing protein 6 OS=Mus musculus OX=1009 | 14 | 5 | 5 | 5 | 468 | 50,7 | 6 | 10,66 | 5 | 1,261 | 1,095 | 0,964 | 0,935 | 0,804 | 1,255 | 0,93 | 1,283 | 1,00 | 0,00 | 9,8E-01 |
| Q8CJG0 | Protein argonaute-2 OS=Mus musculus OX=10090 GN=Ago2 PE=1 S | 4 | 3 | 3 | 2 | 860 | 97,2 | 9,19 | 0 | 3 | 1,84 | 1,325 | 1,803 | 1,4 | 1,334 | 1,917 | 1,275 | 1,817 | 1,00 | 0,00 | 9,8E-01 |

| | | | | | | | | | | | | | | | | | | | | | |
|---|---|---|---|---|---|---|---|---|---|---|---|---|---|---|---|---|---|---|---|---|---|
| Q60575 | Kinesin-like protein KIF1B OS=Mus musculus OX=10090 GN=Kif1b P | 5 | 8 | 8 | 1 | 1816 | 204 | 5,62 | 8,44 | 8 | 0,738 | 0,729 | 0,94 | 0,728 | 0,903 | 0,567 | 1,122 | 0,56 | 0,99 | -0,01 | 9,8E-01 |
| Q61239 | Protein farnesyltransferase/geranylgeranyltransferase type-1 subu | 7 | 3 | 3 | 3 | 377 | 44 | 4,93 | 6,83 | 3 | 1,073 | 0,771 | 1,114 | 1,003 | 1,122 | 0,759 | 0,997 | 1,096 | 1,00 | 0,00 | 9,8E-01 |
| O70400 | PDZ and LIM domain protein 1 OS=Mus musculus OX=10090 GN=Pd | 3 | 1 | 1 | 1 | 327 | 35,8 | 6,84 | 1,82 | 1 | 1,117 | 1,504 | 1,009 | 1,115 | 1,049 | 2,019 | 0,811 | 0,9 | 0,99 | -0,01 | 9,8E-01 |
| Q0VBF8 | Protein stum homolog OS=Mus musculus OX=10090 GN=Stum PE=1 | 21 | 2 | 3 | 2 | 141 | 15 | 7,21 | 7,29 | 2 | 0,979 | 1,082 | 0,818 | 0,93 | 1,021 | 0,822 | 0,963 | 0,995 | 1,00 | 0,00 | 9,8E-01 |
| Q99JY8 | Phospholipid phosphatase 3 OS=Mus musculus OX=10090 GN=Plpp | 14 | 4 | 4 | 4 | 312 | 35,2 | 9,07 | 10,76 | 4 | 1,057 | 1,28 | 1,002 | 0,948 | 1,032 | 1,176 | 1,024 | 1,064 | 1,00 | 0,00 | 9,8E-01 |
| Q5DQR4 | Syntaxin-binding protein 5-like OS=Mus musculus OX=10090 GN=St | 12 | 11 | 14 | 11 | 1185 | 131,8 | 6,86 | 34,72 | 11 | 1,057 | 1,052 | 1,012 | 1,008 | 0,987 | 1,162 | 0,995 | 0,98 | 1,00 | 0,00 | 9,8E-01 |
| Q9DBG3 | AP-2 complex subunit beta OS=Mus musculus OX=10090 GN=Ap2b | 31 | 30 | 59 | 16 | 937 | 104,5 | 5,38 | 133,74 | 30 | 1,038 | 1,017 | 1,039 | 1,049 | 0,965 | 1,098 | 1,041 | 1,036 | 1,00 | 0,00 | 9,8E-01 |
| Q8BGB8 | Ubiquinone biosynthesis protein COQ4 homolog, mitochondrial OS | 4 | 1 | 1 | 1 | 266 | 30,1 | 9,26 | 2,13 | 1 | 1,108 | 1,135 | 1,072 | 0,793 | 0,965 | 1,161 | 0,868 | 1,103 | 1,00 | 0,00 | 9,8E-01 |
| P62196 | 26S proteasome regulatory subunit 8 OS=Mus musculus OX=10090 | 27 | 9 | 13 | 8 | 406 | 45,6 | 7,55 | 27,56 | 9 | 1,379 | 1,071 | 1,183 | 1,124 | 1,029 | 1,355 | 1,161 | 1,202 | 1,00 | 0,00 | 9,8E-01 |
| Q91W89 | Alpha-mannosidase 2C1 OS=Mus musculus OX=10090 GN=Man2c1 | 4 | 3 | 3 | 3 | 1039 | 115,6 | 6,51 | 0 | 3 | 2,143 | 1,556 | 1,421 | 1,255 | 1,203 | 1,906 | 1,153 | 2,146 | 0,99 | -0,01 | 9,8E-01 |
| Q9ER00 | Syntaxin-12 OS=Mus musculus OX=10090 GN=Stx12 PE=1 SV=1 | 11 | 3 | 3 | 3 | 274 | 31,2 | 5,44 | 8,74 | 3 | 0,813 | 0,957 | 0,903 | 0,866 | 0,834 | 0,781 | 1,007 | 0,923 | 1,00 | 0,00 | 9,8E-01 |
| Q8VDC0 | Probable leucine--tRNA ligase, mitochondrial OS=Mus musculus OX | 5 | 4 | 4 | 4 | 902 | 101,4 | 8,19 | 5,79 | 4 | 1,018 | 0,74 | 0,94 | 0,765 | 0,946 | 0,771 | 0,904 | 0,834 | 1,00 | 0,00 | 9,8E-01 |
| Q91V09 | WD repeat-containing protein 13 OS=Mus musculus OX=10090 GN= | 20 | 8 | 11 | 8 | 485 | 53,6 | 9,14 | 19,16 | 8 | 0,815 | 0,973 | 1,06 | 0,987 | 0,986 | 0,872 | 1,014 | 0,969 | 1,00 | 0,00 | 9,8E-01 |
| Q91VR8 | Protein BRICK1 OS=Mus musculus OX=10090 GN=Brk1 PE=1 SV=1 | 40 | 4 | 4 | 4 | 75 | 8,8 | 5,45 | 4,27 | 4 | 1,17 | 1,053 | 0,933 | 0,86 | 0,877 | 1,134 | 0,879 | 1,136 | 1,00 | 0,00 | 9,8E-01 |
| Q925N0 | Sideroflexin-5 OS=Mus musculus OX=10090 GN=Sfxn5 PE=1 SV=2 | 19 | 6 | 11 | 6 | 342 | 37,3 | 9,44 | 15,73 | 6 | 1,128 | 1,03 | 1,029 | 0,956 | 1,03 | 1,118 | 0,939 | 1,051 | 1,00 | 0,00 | 9,8E-01 |
| Q8CI94 | Glycogen phosphorylase, brain form OS=Mus musculus OX=10090 ( | 28 | 23 | 28 | 18 | 843 | 96,7 | 6,73 | 66,36 | 23 | 0,99 | 1,031 | 0,989 | 0,907 | 0,95 | 1,018 | 0,955 | 0,997 | 1,00 | 0,00 | 9,8E-01 |
| P27048 | Small nuclear ribonucleoprotein-associated protein B OS=Mus mus | 3 | 1 | 1 | 1 | 231 | 23,6 | 10,9 | 2,08 | 1 | 1,432 | 0,915 | 0,917 | 0,925 | 1,305 | 1,149 | 0,739 | 1,013 | 1,00 | 0,00 | 9,8E-01 |
| A2ALK8 | Tyrosine-protein phosphatase non-receptor type 3 OS=Mus muscul | 1 | 1 | 1 | 1 | 913 | 103,8 | 7,2 | 2,1 | 1 | 0,932 | 1,093 | 1,096 | 0,969 | 0,998 | 1,07 | 0,875 | 1,154 | 1,00 | 0,00 | 9,8E-01 |
| Q9CQ62 | 2,4-dienoyl-CoA reductase, mitochondrial OS=Mus musculus OX=10 | 20 | 6 | 6 | 6 | 335 | 36,2 | 8,95 | 11,29 | 6 | 1,068 | 0,875 | 1,121 | 0,937 | 1,012 | 0,928 | 0,988 | 1,067 | 1,00 | 0,00 | 9,8E-01 |
| O35495 | Cyclin-dependent kinase 14 OS=Mus musculus OX=10090 GN=Cdk1 | 3 | 2 | 3 | 2 | 469 | 53 | 8,92 | 4,5 | 2 | 0,763 | 0,749 | 0,908 | 0,913 | 0,884 | 0,844 | 0,851 | 0,749 | 1,00 | 0,00 | 9,8E-01 |
| Q9D0G0 | 28S ribosomal protein S30, mitochondrial OS=Mus musculus OX=10 | 14 | 5 | 5 | 5 | 442 | 49,9 | 9,38 | 7,15 | 5 | 1,032 | 0,804 | 0,886 | 0,757 | 0,799 | 1,12 | 0,697 | 0,853 | 1,00 | 0,00 | 9,8E-01 |
| Q9JIG7 | Coiled-coil domain-containing protein 22 OS=Mus musculus OX=10 | 12 | 6 | 7 | 6 | 627 | 70,8 | 6,01 | 7,49 | 6 | 0,932 | 1,014 | 0,878 | 0,912 | 0,941 | 0,839 | 0,954 | 0,998 | 1,00 | 0,00 | 9,8E-01 |
| P20065 | Thymosin beta-4 OS=Mus musculus OX=10090 GN=Tmsb4x PE=1 SV | 40 | 3 | 3 | 1 | 50 | 5,7 | 5,06 | 4,19 | 3 | 0,849 | 0,92 | 0,993 | 1,042 | 0,923 | 0,814 | 1,051 | 1,01 | 1,00 | 0,00 | 9,8E-01 |
| P43274 | Histone H1.4 OS=Mus musculus OX=10090 GN=Hist1h1e PE=1 SV=2 | 23 | 8 | 13 | 2 | 219 | 22 | 11,11 | 34,84 | 8 | 1,178 | 1,387 | 1,075 | 0,802 | 0,992 | 1,43 | 0,812 | 1,192 | 1,00 | 0,00 | 9,8E-01 |
| Q3UMY5 | Echinoderm microtubule-associated protein-like 4 OS=Mus muscul | 5 | 5 | 5 | 5 | 988 | 110 | 6,54 | 7,62 | 5 | 1,234 | 1,221 | 1,131 | 1,282 | 1,17 | 1,305 | 1,162 | 1,227 | 1,00 | 0,00 | 9,8E-01 |
| Q9Z1N5 | Spliceosome RNA helicase Ddx39b OS=Mus musculus OX=10090 GN | 3 | 1 | 1 | 1 | 428 | 49 | 5,67 | 0 | 1 | 1,159 | 0,925 | 1,2 | 0,975 | 0,827 | 1,715 | 0,876 | 0,821 | 1,00 | 0,00 | 9,8E-01 |
| Q80U49 | Centrosomal protein of 170 kDa protein B OS=Mus musculus OX=10 | 4 | 5 | 5 | 5 | 1574 | 170,7 | 6,87 | 8,76 | 5 | 1,114 | 1,029 | 0,872 | 0,886 | 1,122 | 0,731 | 1,027 | 1,03 | 1,00 | 0,00 | 9,8E-01 |
| Q9CY73 | 39S ribosomal protein L44, mitochondrial OS=Mus musculus OX=10 | 2 | 1 | 1 | 1 | 333 | 37,5 | 8,51 | 2,33 | 1 | 1,076 | 1,199 | 1,178 | 0,922 | 1,139 | 1,269 | 1,012 | 0,963 | 1,00 | 0,00 | 9,8E-01 |
| Q80UK0 | SEC14 domain and spectrin repeat-containing protein 1 OS=Mus m | 2 | 1 | 1 | 1 | 696 | 79,3 | 5,1 | 3,07 | 1 | 1,157 | 1,215 | 0,903 | 0,73 | 0,795 | 1,286 | 0,998 | 0,939 | 1,00 | 0,00 | 9,8E-01 |
| Q62417 | Sorbin and SH3 domain-containing protein 1 OS=Mus musculus OX | 7 | 7 | 7 | 7 | 1290 | 143 | 8,25 | 7,63 | 7 | 1,294 | 0,942 | 0,852 | 0,918 | 0,913 | 1,089 | 1,02 | 0,975 | 1,00 | 0,00 | 9,8E-01 |
| Q8BGS2 | BolA-like protein 2 OS=Mus musculus OX=10090 GN=Bola2 PE=1 SV | 10 | 1 | 1 | 1 | 86 | 10,2 | 6,16 | 2,49 | 1 | 1,437 | 1,265 | 0,952 | 1,066 | 1,142 | 1,322 | 1,086 | 1,18 | 1,00 | 0,00 | 9,8E-01 |
| Q8BRJ3 | RELT-like protein 2 OS=Mus musculus OX=10090 GN=Rell2 PE=1 SV | 7 | 1 | 1 | 1 | 303 | 32,3 | 6,18 | 2,29 | 1 | 0,848 | 0,704 | 0,696 | 0,752 | 0,849 | 0,875 | 0,683 | 0,587 | 1,00 | 0,00 | 9,9E-01 |
| Q9DCC8 | Mitochondrial import receptor subunit TOM20 homolog OS=Mus m | 19 | 3 | 5 | 3 | 145 | 16,3 | 8,6 | 8,62 | 3 | 0,74 | 0,95 | 0,829 | 0,79 | 0,998 | 0,692 | 0,839 | 0,774 | 1,00 | 0,00 | 9,9E-01 |
| Q8BKY8 | Transcription termination factor 2, mitochondrial OS=Mus musculu | 2 | 1 | 1 | 1 | 385 | 43,4 | 8,94 | 2,34 | 1 | 1,048 | 0,982 | 1,1 | 0,792 | 0,764 | 1,235 | 0,941 | 0,973 | 1,00 | 0,00 | 9,9E-01 |
| Q8K2Q0 | COMM domain-containing protein 9 OS=Mus musculus OX=10090 ( | 12 | 2 | 3 | 2 | 198 | 21,8 | 5,82 | 6,24 | 2 | 0,999 | 1,088 | 1,068 | 1,099 | 1,149 | 1,083 | 1,006 | 1,019 | 1,00 | 0,00 | 9,9E-01 |
| O70503 | Very-long-chain 3-oxoacyl-CoA reductase OS=Mus musculus OX=10 | 8 | 2 | 2 | 2 | 312 | 34,7 | 9,52 | 3,98 | 2 | 1,133 | 0,999 | 1,143 | 0,939 | 0,947 | 1,021 | 1,014 | 1,238 | 1,00 | 0,00 | 9,9E-01 |
| P70698 | CTP synthase 1 OS=Mus musculus OX=10090 GN=Ctps1 PE=1 SV=2 | 5 | 2 | 2 | 2 | 591 | 66,6 | 6,58 | 2,12 | 2 | 1,087 | 1,004 | 1,169 | 0,847 | 0,891 | 1,421 | 0,948 | 0,836 | 1,00 | 0,00 | 9,9E-01 |
| Q9Z268 | RasGAP-activating-like protein 1 OS=Mus musculus OX=10090 GN= | 16 | 12 | 16 | 12 | 799 | 89,3 | 6,37 | 38,91 | 12 | 0,984 | 0,991 | 0,949 | 1,098 | 0,958 | 0,973 | 1,077 | 1,011 | 1,00 | 0,00 | 9,9E-01 |
| O54901 | OX-2 membrane glycoprotein OS=Mus musculus OX=10090 GN=Cd | 14 | 5 | 10 | 5 | 278 | 31,2 | 8,79 | 27 | 5 | 0,815 | 0,911 | 0,941 | 1,046 | 0,986 | 0,771 | 0,982 | 0,979 | 1,00 | 0,00 | 9,9E-01 |
| Q9JMK2 | Casein kinase I isoform epsilon OS=Mus musculus OX=10090 GN=C | 9 | 3 | 3 | 1 | 416 | 47,3 | 9,66 | 1,87 | 3 | 0,526 | 0,627 | 0,799 | 0,969 | 0,64 | 0,594 | 1,04 | 0,637 | 1,00 | 0,00 | 9,9E-01 |
| O88384 | Vesicle transport through interaction with t-SNAREs homolog 1B O | 14 | 3 | 3 | 3 | 232 | 26,7 | 8,79 | 5,85 | 3 | 1,228 | 0,951 | 1,072 | 0,99 | 1,06 | 1,181 | 0,994 | 1,001 | 1,00 | 0,00 | 9,9E-01 |
| P62331 | ADP-ribosylation factor 6 OS=Mus musculus OX=10090 GN=Arf6 PE | 15 | 2 | 3 | 2 | 175 | 20,1 | 8,95 | 4,18 | 2 | 1,009 | 0,908 | 0,932 | 1,093 | 1 | 0,96 | 1,03 | 0,955 | 1,00 | 0,00 | 9,9E-01 |
| Q3UHG7 | DENN domain-containing protein 11 OS=Mus musculus OX=10090 ( | 5 | 2 | 2 | 2 | 455 | 51,4 | 5,25 | 0 | 2 | 0,604 | 0,511 | 0,786 | 1,221 | 0,909 | 0,642 | 0,782 | 0,8 | 1,00 | 0,00 | 9,9E-01 |
| Q3UUJ4 | STE20-related kinase adapter protein alpha OS=Mus musculus OX= | 7 | 2 | 2 | 2 | 431 | 48,2 | 6,43 | 2,47 | 2 | 0,519 | 1,223 | 1,112 | 0,923 | 0,774 | 0,952 | 0,946 | 1,094 | 1,00 | 0,00 | 9,9E-01 |
| Q8C078 | Calcium/calmodulin-dependent protein kinase kinase 2 OS=Mus mu | 13 | 5 | 7 | 5 | 588 | 64,6 | 5,9 | 2,91 | 5 | 0,913 | 1,059 | 0,813 | 0,8 | 0,916 | 0,857 | 0,896 | 0,92 | 1,00 | 0,00 | 9,9E-01 |
| A2AHC3 | Calmodulin-regulated spectrin-associated protein 1 OS=Mus muscu | 5 | 7 | 7 | 7 | 1581 | 175,8 | 6,95 | 5,07 | 7 | 1,16 | 0,813 | 0,871 | 0,978 | 0,973 | 0,987 | 0,962 | 0,905 | 1,00 | 0,00 | 9,9E-01 |
| P45878 | Peptidyl-prolyl cis-trans isomerase FKBP2 OS=Mus musculus OX=10 | 28 | 4 | 5 | 4 | 140 | 15,3 | 8,88 | 11,67 | 4 | 1,144 | 0,991 | 1,093 | 0,938 | 0,993 | 1,183 | 0,994 | 0,992 | 1,00 | 0,00 | 9,9E-01 |
| Q8BZW8 | NHL repeat-containing protein 2 OS=Mus musculus OX=10090 GN= | 3 | 2 | 2 | 2 | 725 | 78,4 | 5,54 | 3,29 | 2 | 0,972 | 0,859 | 1,21 | 0,897 | 0,512 | 1,66 | 0,841 | 0,94 | 1,00 | 0,00 | 9,9E-01 |
| Q9WTS4 | Teneurin-1 OS=Mus musculus OX=10090 GN=Tenm1 PE=1 SV=1 | 1 | 3 | 4 | 3 | 2731 | 305,6 | 6,37 | 5,92 | 3 | 1,368 | 0,641 | 0,904 | 1,039 | 0,85 | 0,899 | 1,067 | 1,127 | 1,00 | 0,00 | 9,9E-01 |
| O55091 | Protein IMPACT OS=Mus musculus OX=10090 GN=Impact PE=1 SV= | 15 | 4 | 5 | 4 | 318 | 36,3 | 5,05 | 7,76 | 4 | 1,292 | 0,979 | 0,944 | 1,235 | 1,064 | 1,052 | 1,13 | 1,199 | 1,00 | 0,00 | 9,9E-01 |
| O88522 | NF-kappa-B essential modulator OS=Mus musculus OX=10090 GN= | 8 | 3 | 3 | 3 | 412 | 47,9 | 5,85 | 3,94 | 3 | 1,203 | 1,787 | 1,187 | 1,114 | 1,247 | 1,794 | 1 | 1,238 | 1,00 | 0,00 | 9,9E-01 |
| Q9JKB3 | Y-box-binding protein 3 OS=Mus musculus OX=10090 GN=Ybx3 PE= | 8 | 2 | 2 | 2 | 361 | 38,8 | 9,69 | 2,34 | 2 | 1,001 | 1,499 | 1,178 | 1,441 | 0,982 | 1,459 | 1,163 | 1,524 | 1,00 | 0,00 | 9,9E-01 |
| P62717 | 60S ribosomal protein L18a OS=Mus musculus OX=10090 GN=Rpl18 | 31 | 5 | 7 | 5 | 176 | 20,7 | 10,71 | 14,22 | 5 | 0,933 | 0,903 | 1,008 | 0,973 | 1,019 | 0,882 | 0,917 | 0,997 | 1,00 | 0,00 | 9,9E-01 |
| Q9QZI9 | Serine incorporator 3 OS=Mus musculus OX=10090 GN=Serinc3 PE= | 3 | 1 | 1 | 1 | 472 | 52,6 | 7,23 | 0 | 1 | 0,758 | 0,856 | 0,944 | 1,087 | 0,753 | 0,96 | 1,081 | 0,846 | 1,00 | 0,00 | 9,9E-01 |
| Q3UJD6 | Ubiquitin carboxyl-terminal hydrolase 19 OS=Mus musculus OX=10 | 2 | 2 | 2 | 2 | 1360 | 150,5 | 6,38 | 0 | 2 | 0,978 | 0,813 | 0,559 | 0,623 | 0,696 | 0,731 | 0,645 | 0,906 | 1,00 | 0,00 | 9,9E-01 |
| Q71LX4 | Talin-2 OS=Mus musculus OX=10090 GN=Tln2 PE=1 SV=3 | 18 | 35 | 46 | 32 | 2375 | 253,5 | 5,8 | 77,71 | 35 | 1,111 | 1,014 | 1,076 | 1,051 | 0,952 | 1,248 | 1,024 | 1,031 | 1,00 | 0,00 | 9,9E-01 |
| Q91WG4 | Elongator complex protein 2 OS=Mus musculus OX=10090 GN=Elp2 | 3 | 2 | 2 | 2 | 831 | 93 | 5,82 | 1,89 | 2 | 0,764 | 1,353 | 1,218 | 1,379 | 1,251 | 0,979 | 1,267 | 1,224 | 1,00 | 0,00 | 9,9E-01 |
| Q61879 | Myosin-10 OS=Mus musculus OX=10090 GN=Myh10 PE=1 SV=2 | 28 | 62 | 99 | 55 | 1976 | 228,9 | 5,54 | 201,57 | 62 | 0,998 | 1,023 | 1,004 | 0,963 | 0,939 | 1,031 | 1,023 | 0,996 | 1,00 | 0,00 | 9,9E-01 |
| O09111 | NADH dehydrogenase [ubiquinone] 1 beta subcomplex subunit 11, | 19 | 2 | 3 | 2 | 151 | 17,4 | 5,22 | 7,43 | 2 | 0,991 | 1,17 | 1,01 | 0,818 | 0,877 | 1,182 | 0,926 | 1,008 | 1,00 | 0,00 | 9,9E-01 |
| P59016 | Vacuolar protein sorting-associated protein 33B OS=Mus musculus | 11 | 7 | 7 | 7 | 617 | 70,5 | 6,86 | 15,75 | 7 | 0,893 | 0,989 | 1,076 | 1,11 | 0,994 | 1,05 | 1,015 | 1,011 | 1,00 | 0,00 | 9,9E-01 |
| Q61527 | Receptor tyrosine-protein kinase erbB-4 OS=Mus musculus OX=100 | 2 | 2 | 2 | 2 | 1308 | 146,8 | 6,38 | 0 | 2 | 0,989 | 1,443 | 0,955 | 1,242 | 0,967 | 1,068 | 0,985 | 1,602 | 1,00 | 0,00 | 9,9E-01 |
| Q3UNH4 | G protein-regulated inducer of neurite outgrowth 1 OS=Mus muscu | 34 | 31 | 39 | 31 | 932 | 95,4 | 7,93 | 72,22 | 31 | 0,856 | 0,954 | 1,016 | 1,073 | 0,967 | 0,897 | 1,024 | 1,009 | 1,00 | 0,00 | 9,9E-01 |

| | | | | | | | | | | | | | | | | | | | | | |
|---|---|---|---|---|---|---|---|---|---|---|---|---|---|---|---|---|---|---|---|---|---|
| Q11136 | Xaa-Pro dipeptidase OS=Mus musculus OX=10090 GN=Pepd PE=1 S | 10 | 4 | 4 | 4 | 493 | 55 | 5,78 | 11,43 | 4 | 0,873 | 0,993 | 1,006 | 0,983 | 0,943 | 0,855 | 0,973 | 1,082 | 1,00 | 0,00 | 9,9E-01 |
| P28667 | MARCKS-related protein OS=Mus musculus OX=10090 GN=Marcksl | 36 | 4 | 6 | 4 | 200 | 20,2 | 4,61 | 12,59 | 4 | 0,521 | 0,921 | 1,016 | 1,292 | 1,001 | 0,752 | 1,101 | 0,891 | 1,00 | 0,00 | 9,9E-01 |
| Q8R0S4 | Voltage-dependent L-type calcium channel subunit beta-4 OS=Mus | 13 | 6 | 11 | 3 | 519 | 57,9 | 9,28 | 11,43 | 6 | 1,179 | 0,963 | 1,018 | 1,223 | 1,002 | 1,219 | 1,126 | 1,038 | 1,00 | 0,00 | 1,0E+00 |
| Q61220 | Protein kinase C-binding protein NELL2 OS=Mus musculus OX=1009 | 1 | 1 | 1 | 1 | 819 | 91,4 | 5,82 | 2,52 | 1 | 0,656 | 0,986 | 0,915 | 1,038 | 1,234 | 0,558 | 0,858 | 0,941 | 1,00 | 0,00 | 1,0E+00 |
| Q8VHQ9 | Acyl-coenzyme A thioesterase 11 OS=Mus musculus OX=10090 GN= | 5 | 2 | 2 | 2 | 594 | 67,3 | 6,8 | 4,3 | 2 | 1,882 | 1,643 | 1,547 | 1,506 | 1,537 | 2,091 | 1,202 | 1,753 | 1,00 | 0,00 | 1,0E+00 |
| Q6PHQ8 | N-alpha-acetyltransferase 35, NatC auxiliary subunit OS=Mus musc | 4 | 2 | 2 | 2 | 725 | 83,3 | 7,3 | 1,8 | 2 | 1,011 | 1,064 | 0,563 | 0,997 | 0,88 | 1,08 | 0,861 | 0,817 | 1,00 | 0,00 | 1,0E+00 |
| Q9JJ28 | Protein flightless-1 homolog OS=Mus musculus OX=10090 GN=Flii P | 3 | 3 | 3 | 3 | 1271 | 144,7 | 6,06 | 5,85 | 3 | 1,021 | 1,025 | 1,132 | 0,963 | 1,143 | 1,047 | 0,965 | 0,985 | 1,00 | 0,00 | 1,0E+00 |
| P55258 | Ras-related protein Rab-8A OS=Mus musculus OX=10090 GN=Rab8 | 23 | 6 | 10 | 2 | 207 | 23,7 | 9,07 | 21,19 | 6 | 1,06 | 0,987 | 1,104 | 1,329 | 1,216 | 0,912 | 1,077 | 1,277 | 1,00 | 0,00 | 1,0E+00 |
| Q99K67 | Alpha-aminoadipic semialdehyde synthase, mitochondrial OS=Mus | 2 | 2 | 2 | 2 | 926 | 102,9 | 6,87 | 2,26 | 2 | 1,305 | 1,398 | 0,793 | 1,278 | 1,254 | 1,156 | 0,958 | 1,409 | 1,00 | 0,00 | 1,0E+00 |
| Q8C0E2 | Vacuolar protein sorting-associated protein 26B OS=Mus musculus | 31 | 9 | 14 | 9 | 336 | 39,1 | 7,37 | 22,09 | 9 | 1,114 | 1,127 | 0,976 | 1,177 | 1,114 | 0,986 | 1,157 | 1,138 | 1,00 | 0,00 | 1,0E+00 |
| Q5FWH7 | Zinc transporter ZIP12 OS=Mus musculus OX=10090 GN=Slc39a12 P | 1 | 1 | 1 | 1 | 689 | 76,2 | 5,77 | 1,82 | 1 | 1,518 | 1,43 | 2,576 | 1,419 | 1,462 | 2,179 | 1,462 | 1,835 | 1,00 | 0,00 | 1,0E+00 |
| Q6P9R2 | Serine/threonine-protein kinase OSR1 OS=Mus musculus OX=1009( | 9 | 4 | 4 | 4 | 527 | 58,2 | 6,43 | 5,17 | 4 | 1,051 | 1,17 | 0,97 | 0,998 | 0,925 | 1,264 | 1,269 | 0,729 | 1,00 | 0,00 | 1,0E+00 |
| Q9CZ30 | Obg-like ATPase 1 OS=Mus musculus OX=10090 GN=Ola1 PE=1 SV= | 20 | 8 | 9 | 8 | 396 | 44,7 | 7,81 | 14,68 | 8 | 0,818 | 0,883 | 0,979 | 1,066 | 0,963 | 0,799 | 0,975 | 1,008 | 1,00 | 0,00 | 1,0E+00 |
| P62748 | Hippocalcin-like protein 1 OS=Mus musculus OX=10090 GN=Hpcal1 | 44 | 8 | 9 | 4 | 193 | 22,3 | 5,5 | 22,8 | 8 | 1,17 | 0,782 | 0,748 | 0,968 | 0,794 | 1,205 | 0,959 | 0,712 | 1,00 | 0,00 | 1,0E+00 |
| Q3TES0 | IQ motif and SEC7 domain-containing protein 3 OS=Mus musculus ( | 6 | 5 | 5 | 4 | 1195 | 129 | 6,19 | 8,35 | 5 | 1,114 | 1,322 | 1,266 | 1,01 | 0,961 | 1,621 | 1,111 | 1,021 | 1,00 | 0,00 | 1,0E+00 |
| Q6P9N1 | Hyccin OS=Mus musculus OX=10090 GN=Fam126a PE=1 SV=3 | 2 | 1 | 1 | 1 | 521 | 57,3 | 7,88 | 0 | 1 | 0,01 | 1,29 | 0,94 | 0,997 | 0,675 | 1,112 | 0,436 | 1,015 | 1,00 | 0,00 | 1,0E+00 |
| O08576 | RUN domain-containing protein 3A OS=Mus musculus OX=10090 G | 6 | 3 | 3 | 3 | 446 | 50 | 5,5 | 9,15 | 3 | 1,231 | 1,193 | 1,018 | 0,975 | 0,859 | 1,332 | 0,994 | 1,232 | 1,00 | 0,00 | 1,0E+00 |
| P05480 | Neuronal proto-oncogene tyrosine-protein kinase Src OS=Mus mus | 21 | 10 | 13 | 9 | 541 | 60,6 | 7,84 | 32,69 | 10 | 1,04 | 1,014 | 1,041 | 1,02 | 0,959 | 1,089 | 1,044 | 1,023 | 1,00 | 0,00 | 1,0E+00 |
| Q8BGZ1 | Hippocalcin-like protein 4 OS=Mus musculus OX=10090 GN=Hpcal4 | 27 | 5 | 10 | 3 | 191 | 22,2 | 4,89 | 24,76 | 5 | 1,007 | 0,929 | 1,036 | 1,038 | 0,918 | 1,102 | 1,001 | 0,989 | 1,00 | 0,00 | 1,0E+00 |
| Q9R0Q6 | Actin-related protein 2/3 complex subunit 1A OS=Mus musculus OX | 26 | 9 | 14 | 9 | 370 | 41,6 | 8,18 | 36,09 | 9 | 0,968 | 0,986 | 0,99 | 0,974 | 1,019 | 0,933 | 0,987 | 0,979 | 1,00 | 0,00 | 1,0E+00 |
