## Supplementary material for "SorCS2 modulates neurovascular coupling via glutamatergic and calcium signaling in astrocytes": Expanded View Table 3

© 2000-2022 QIAGEN. All rights reserved.

| **Ingenuity Canonical Pathways** | **-log(*P*-value)** | ***P*-value** | **Z-score** | **Molecules** |
|---|---|---|---|---|
| Synaptogenesis Signaling Pathway | 6,73E00 | 1,9E-07 | -1,000 | AP2M1,Calm1 (includes others),CAMK2B,CPLX1,CTNND1,EPHB2,GRIA2,GRIN1,GRM3,KRAS,MRAS,NRAS,SNAP25,SYT1,SYT2,WASF1 |
| Synaptic Long Term Potentiation | 6,03E00 | 9,3E-07 | -0,632 | Calm1 (includes others),CAMK2B,GRIA2,GRIN1,GRM3,KRAS,MRAS,NRAS,PDIA3,PPP1R7 |
| Oxidative Phosphorylation | 5,64E00 | 2,3E-06 | -3,000 | ATP5PF,CYCS,MT-ND5,NDUFA4,NDUFB4,NDUFB5,NDUFV3,UQCRB,VPS9D1 |
| Sirtuin Signaling Pathway | 4,94E00 | 1,1E-05 | 2,121 | ATG7,ATP5PF,GLUD1,GOT2,MT-ND5,NDUFA4,NDUFB4,NDUFB5,NDUFV3,SIRT2,TIMM22,TIMM44,VDAC1 |
| TCA Cycle II (Eukaryotic) | 4,08E00 | 8,3E-05 | -1,000 | DLD,FH,MDH2,SUCLG1 |
| EIF2 Signaling | 3,95E00 | 0,00011 | -1,342 | EIF4G2,EIF4G3,KRAS,MRAS,NRAS,PABPC1,RPL35A,RPS28,RPS4Y1,WARS1 |
| Opioid Signaling Pathway | 3,81E00 | 0,00015 | -0,905 | AP2M1,CACNA1A,Calm1 (includes others),CAMK2B,CLTA,CLTC,GNB1,GRIN1,KRAS,MRAS,NRAS |
| Calcium Signaling | 3,38E00 | 0,00042 | -1,633 | ATP2A2,ATP2C1,CACNA1A,Calm1 (includes others),CALR,CAMK2B,GRIA2,GRIN1,MYH14 |
| Superpathway of Methionine Degradation | 3,21E00 | 0,00062 | -1,000 | DLD,GOT2,MAT2A,MMUT |
| Apoptosis Signaling | 3,14E00 | 0,00072 | -0,447 | AIFM1,CYCS,ELP1,KRAS,MRAS,NRAS |
| Glioma Invasiveness Signaling | 3,03E00 | 0,00093 | -0,447 | KRAS,MRAS,NRAS,RHOB,RHOT1 |
| Chemokine Signaling | 2,85E00 | 0,00141 | -0,447 | Calm1 (includes others),CAMK2B,KRAS,MRAS,NRAS |
| IL-8 Signaling | 2,8E00 | 0,00158 | -0,378 | ELP1,GNB1,KRAS,MRAS,NRAS,PLD3,RHOB,RHOT1 |
| Thrombin Signaling | 2,78E00 | 0,00166 | -0,378 | CAMK2B,GNB1,KRAS,MRAS,NRAS,PDIA3,RHOB,RHOT1 |
| Reelin Signaling in Neurons | 2,71E00 | 0,00195 | -1,633 | CAMK2B,GRIN1,NDEL1,PDK2,PDK3,WASF1 |
| Dopamine-DARPP32 Feedback in cAMP Signaling | 2,56E00 | 0,00275 | -0,447 | ATP2A2,CACNA1A,Calm1 (includes others),GRIN1,KCNJ11,PDIA3,PPP1R7 |
| Estrogen Receptor Signaling | 2,47E00 | 0,00339 | -0,816 | CACNA1A,GNB1,KRAS,MRAS,MT-ND5,NDUFA4,NDUFB4,NDUFB5,NDUFV3,NRAS,PDIA3 |
| GNRH Signaling | 2,44E00 | 0,00363 | -1,342 | CACNA1A,Calm1 (includes others),CAMK2B,GNB1,KRAS,MRAS,NRAS |
| Senescence Pathway | 2,42E00 | 0,0038 | -1,667 | CACNA1A,Calm1 (includes others),DLD,KRAS,MRAS,NRAS,PDHB,PDK2,PDK3 |
| Synaptic Long Term Depression | 2,38E00 | 0,00417 | -0,378 | CACNA1A,GRIA2,GRM3,KRAS,MRAS,NRAS,PDIA3 |
| Necroptosis Signaling Pathway | 2,24E00 | 0,00575 | -1,633 | CAMK2B,CYLD,GLUD1,TIMM22,TIMM44,VDAC1 |
| GM-CSF Signaling | 2,23E00 | 0,00589 | -1,000 | CAMK2B,KRAS,MRAS,NRAS |
| CDK5 Signaling | 2,18E00 | 0,00661 | -1,342 | CACNA1A,KRAS,MRAS,NRAS,PPP1R7 |
| Cholecystokinin/Gastrin-mediated Signaling | 2,11E00 | 0,00776 | -0,447 | KRAS,MRAS,NRAS,RHOB,RHOT1 |
| CXCR4 Signaling | 2,08E00 | 0,00832 | -0,816 | GNB1,KRAS,MRAS,NRAS,RHOB,RHOT1 |
| Angiopoietin Signaling | 2,08E00 | 0,00832 | -1,000 | ELP1,KRAS,MRAS,NRAS |
| Antiproliferative Role of Somatostatin Receptor 2 | 2,08E00 | 0,00832 | -1,000 | GNB1,KRAS,MRAS,NRAS |
| Glioma Signaling | 2,03E00 | 0,00933 | -0,447 | Calm1 (includes others),CAMK2B,KRAS,MRAS,NRAS |
| G Beta Gamma Signaling | 1,96E00 | 0,01096 | -1,342 | CACNA1A,GNB1,KRAS,MRAS,NRAS |
| P2Y Purigenic Receptor Signaling Pathway | 1,94E00 | 0,01148 | -0,447 | GNB1,KRAS,MRAS,NRAS,PDIA3 |
| PDGF Signaling | 1,92E00 | 0,01202 | -1,000 | KRAS,MRAS,NRAS,SYNJ1 |
| Ceramide Signaling | 1,85E00 | 0,01413 | -1,000 | CYCS,KRAS,MRAS,NRAS |
| Gαi Signaling | 1,82E00 | 0,01514 | -0,447 | GNB1,GRM3,KRAS,MRAS,NRAS |
| Insulin Receptor Signaling | 1,82E00 | 0,01514 | 0,447 | KRAS,MRAS,NRAS,PPP1R7,SYNJ1 |
| Ephrin Receptor Signaling | 1,74E00 | 0,0182 | -1,342 | EPHB2,GNB1,GRIN1,KRAS,MRAS,NRAS |
| Endocannabinoid Neuronal Synapse Pathway | 1,72E00 | 0,01905 | -1,342 | CACNA1A,GNB1,GRIA2,GRIN1,PDIA3 |
| Signaling by Rho Family GTPases | 1,67E00 | 0,02138 | 0,816 | GFAP,GNB1,MRAS,RHOB,RHOT1,SEPTIN11,WASF1 |
| CREB Signaling in Neurons | 1,63E00 | 0,02344 | -0,905 | ADGRB3,CACNA1A,Calm1 (includes others),CAMK2B,GNB1,GRIA2,GRIN1,GRM3,KRAS,MRAS,NRAS,PDIA3 |
| HMGB1 Signaling | 1,53E00 | 0,02951 | -0,447 | KRAS,MRAS,NRAS,RHOB,RHOT1 |
| Endothelin-1 Signaling | 1,31E00 | 0,04898 | 0,447 | KRAS,MRAS,NRAS,PDIA3,PLD3 |
