## Supplementary material for "SorCS2 modulates neurovascular coupling via glutamatergic and calcium signaling in astrocytes": Expanded View figure 1

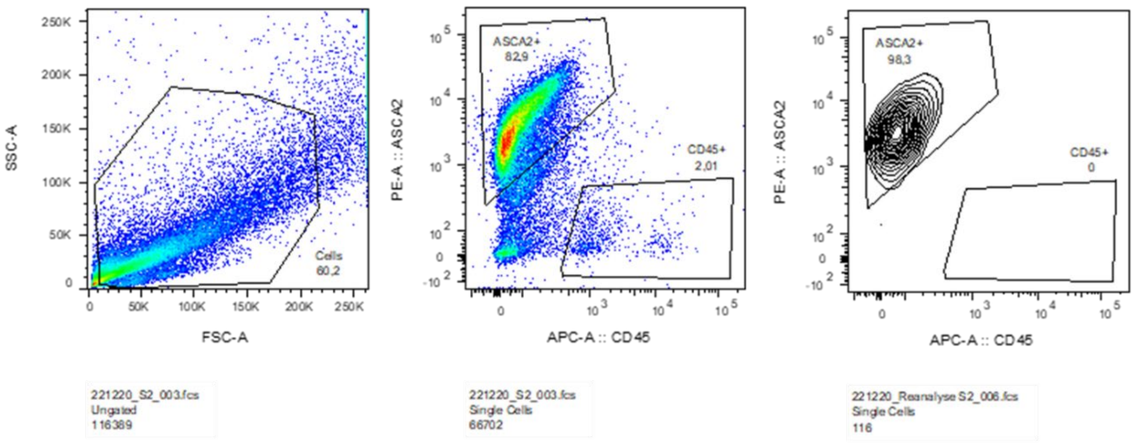


**Expanded View figure 1. Representative results of flow cytometric isolation of astrocytes from 2-month-old mouse brain.** Left panel shows ungated forward (FSC) and side (SSC) scatter plot of isolated cells. Middle panel identifies populations of ASCA2 positive and CD45 positive cells. Right panel reanalyzes the data to determine a fraction of sorted astrocytes.
